# Epstein Barr virus genomes reveal population structure and type 1 association with endemic Burkitt lymphoma

**DOI:** 10.1101/689216

**Authors:** Yasin Kaymaz, Cliff I. Oduor, Ozkan Aydemir, Micah A. Luftig, Juliana A. Otieno, John Michael Ong’echa, Jeffrey A. Bailey, Ann M. Moormann

## Abstract

Endemic Burkitt lymphoma (eBL), the most prevalent pediatric cancer in sub-Saharan Africa, is associated with malaria and Epstein Barr virus (EBV). In order to better understand the role of EBV in eBL, we improved viral DNA enrichment methods and generated a total of 98 new EBV genomes from both eBL cases (N=58) and healthy controls (N=40) residing in the same geographic region in Kenya. Comparing cases and controls, we found that EBV type 1 was significantly associated with eBL with 74.5% of patients (41/55) versus 47.5% of healthy children (19/40) carrying type 1 (OR=3.24, 95% CI=1.36 - 7.71, *P=0.007*). Controlling for EBV type, we also performed a genome-wide association study identifying 6 nonsynonymous variants in the genes EBNA1, EBNA2, BcLF1, and BARF1 that were enriched in eBL patients. Additionally, we observed that viruses isolated from plasma of eBL patients were identical to their tumor counterpart consistent with circulating viral DNA originating from the tumor. We also detected three intertypic recombinants carrying type 1 EBNA2 and type 2 EBNA3 regions as well as one novel genome with a 20 kb deletion resulting in the loss of multiple lytic and virion genes. Comparing EBV types, genes show differential variation rates as type 1 appears to be more divergent. Besides, type 2 demonstrates novel substructures. Overall, our findings address the complexities of EBV population structure and provide new insight into viral variation, which has the potential to influence eBL oncogenesis.

**Key Points:**

- EBV type 1 is more prevalent in eBL patients compared to the geographically matched healthy control group.
- Genome-wide association analysis between cases and controls identifies 6 eBL-associated nonsynonymous variants in EBNA1, EBNA2, BcLF1, and BARF1 genes.
- Analysis of population structure reveals that EBV type 2 exists as two genomic sub groups.

## Introduction

EBV infects more than 90% of the world’s population and typically persists as a chronic asymptomatic infection.^1^ While most individuals endure a lifelong infection with minimal effect, EBV is associated with ~1% of all human malignancies worldwide. EBV was first isolated from an endemic Burkitt lymphoma (eBL) tumor which is the most prevalent pediatric cancer in sub-Saharan Africa.^2^ Repeated *Plasmodium falciparum* infections during childhood appear to drive this increased incidence.^3^ Malaria causes polyclonal B-cell expansion and increased expression of activation-induced cytidine deaminase (AID) dependent DNA damage leading to the hallmark translocation of the *MYC* gene under control of the constitutively active immunoglobulin enhancer.^4–6^ How EBV potentiates eBL is incompletely understood, however, the clonal presence of this virus in almost every eBL tumor suggests a necessary role.

EBV strains are categorized into two types based on the high degree of divergence in the *EBNA2* and *EBNA3* genes.^7–9^ This long standing evolutionary division is also present in orthologous primate viruses,^10^ yet remains unexplained. While EBV type 1 has been extensively studied,^11, 12^ because it causes acute infectious mononucleosis and other diseases in the developed world, type 2 virus studies have not kept pace since infected individuals are less frequent and found primarily in sub-Saharan Africa. While several recent studies have reported both types of EBV circulating in western countries,^13, 14^ the African context provides a better opportunity to examine viral variation because type 1 and type 2 are found in both eBL patients as well as healthy individuals.^8, 15, 16^ Viral variation has been shown to impact differential transformation and growth, and capacity to block apoptosis or immune recognition.^7, 17, 18^ However, studies focusing on only certain genomic regions/proteins potentially miss disease associations of other loci.^19, 20^ Although new studies have been conducted,^21, 22^ genome-wide examinations in case-control studies are few and often lack typing the virus.

To address this shortfall, whole genome sequencing of EBV is now attainable from tumor, blood, or saliva using targeted viral DNA capture methods.^23–28^ However, studying EBV from the blood of healthy individuals remains challenging due to low viral abundance relative to human DNA (1-10 EBV copy/ng blood DNA). In addition, EBV’s GC-rich genome is inefficiently amplified using conventional library preparation methods. Here, we present improved methods for EBV genome enrichment that allow us to sequence virus directly from eBL patients and healthy children. Leveraging these samples, we sought to define the viral population structure and characterize viral subtypes collected from children hailing from the same region of western Kenya. Additionally, we performed the first genome wide association study to identify viral variants that correlate with eBL pathogenesis.

## Materials and Methods

### Ethical approval and sample collection

For this study, we recruited children between 2009 and 2012 with suspected eBL, between 2-14 years of age, undergoing initial diagnosis at Jaramogi Oginga Odinga Teaching and Referral Hospital (JOOTRH; Kisumu), which is a regional referral hospital for pediatric cancer in western Kenya.^29^ We obtained written informed consent from children’s parents or legal guardians to enroll them in this study. Ethical approval was obtained from the Institutional Review Board at the University of Massachusetts Medical School and the Scientific and Ethical Review Unit at the Kenya Medical Research Institute. For this study, primary tumor biopsies were collected using fine needle aspirates (FNA) and transferred into RNAlater at the bedside, prior to induction of chemotherapy. In addition, peripheral blood samples were collected and fractionated by centrifugation prior to freezing into plasma and cell pellets. All samples were stored at −80°C prior to nucleic acid extraction.

### Improved enrichment of GC-rich EBV in low abundance samples

We used Allprep DNA/RNA/Protein mini kit (Qiagen) for DNA isolations from FNAs and QIAamp DNA Kit for blood and plasma. We developed an improved multi-step amplification and enrichment process for the GC-rich EBV genome, particularly in samples with low viral copies. We used EBV-specific whole genome amplification (sWGA) to provide sufficient material and targeted enrichment with hybridization probes after the library preparation. For this, we designed 3’-protected oligos following the instructions from Leichty et al.^30^ (detailed in Supplemental Methods). For low viral load samples, we added a multiplex long-range PCR amplification (mlrPCR) step, comprising two sets of non-overlapping EBV-specific primers^31^ tiling across the genome. We improved the amplification yield for low copy EBV input (**Supplemental Table 1**) by optimizing buffers and reaction conditions (**Supplemental Figure 1A and 1B**).

### Sequencing library preparation and hybrid capture enrichment

Illumina sequencing library preparation steps consisted of DNA shearing, blunt-end repair (Quick Blunting kit, NEB), 3’-adenylation (Klenow Fragment 3’ to 5’ exo-, NEB), and ligation of indexed sequencing adaptors (Quick Ligation kit, NEB). We PCR amplified libraries to a final concentration with 10 cycles using KAPA HiFi HotStart ReadyMix and quantified them using bioanalyzer. We then pooled sample libraries balancing them according to their EBV content and proceeded to target enrichment hybridization using custom EBV-specific biotinylated RNA probes (MyBaits, Arbor Biosciences). We sequenced the libraries using Illumina sequencing instruments with various read lengths ranging from 75bp to 150bp.

### Sequence preprocessing and de novo genome assembly

We checked the sequence quality using FastQC (v0.10.1) after trimming residual adapter and low quality bases (<20) using cutadapt (v1.7.1)^32^ and prinseq (v0.20.4),^33^ respectively. After removing reads that mapped to the human genome (hg38), we de novo assembled the remaining reads into contigs with VelvetOptimiser (v2.2.5)^34^ using a kmer search ranging from 21 to 149 to maximize N50. We then ordered and oriented the contigs guided by the reference using ABACAS, extended with read support using IMAGE,^35^ and merged the overlapping contigs to form larger scaffolds (using in-house scripts). By aligning reads back to scaffolds, we assessed contig quality requiring support from ≥5 unique reads. We created a final genome by demarcating repetitive and missing regions due to low coverage with sequential ambiguous “N” nucleotides. We excluded minor variants (<5% of reads) in final assemblies. Deposited genomes can be accessed from GenBank (accession #) and raw reads can be downloaded from SRA (SRA accession #).

### Diversity and variant association analysis

We used Mafft (v7.215)^36^ for multiple sequence alignment (msa) of genomes, and constructed phylogenetic neighbor-joining trees with Jukes-Cantor substitution model using MEGA (v6.0).^37^ We determined variant sites relative to consensus using snp-sites (v2.3.2)^38^ then projected variant loci on EBV type 1 reference. For principal coordinate analysis (PCoA), we used dartR (v1.0.5).^39^ We calculated dN/dS rates per gene using SNAP (v2.1.1) after excluding frameshift insertions and ambiguous bases.^40^ For variant association analysis, we used ‘v-assoc’ function from PSEQ/PLINK. To control for multiple testing, we calculated empirical p-values with one million permutations (pseq proj v-assoc --phenotype eBL --fix-null --perm 1000000) with EBV type stratification which permutes within types (--strata EBVtype).

## Results

### Study participant characteristics

The objective of this study was to examine EBV genetic variation in a region of western Kenya with a high incidence of eBL^29^ and determine if any variants are associated with eBL pathogenesis. We leveraged specimens from eBL patients and healthy children residing in the same geographic area (**Figure 1A**).^29^ We sequenced the virus isolated from 58 eBL cases and 40 healthy Kenyan children, as controls. Patients aging between 1 and 13 years were predominantly male (74%), consistent with the sex ratio of eBL (**Table 1**).^29^ Healthy controls had similar levels of malaria exposure based on previous epidemiologic studies.^41^ Control samples ranged in age from 1 to 6 years. This difference in age was necessary due to the finding that younger, healthy yet malaria-exposed children have higher average viral loads compared to older children who have developed immune control over this chronic viral infection.^42^

**Figure 1.**
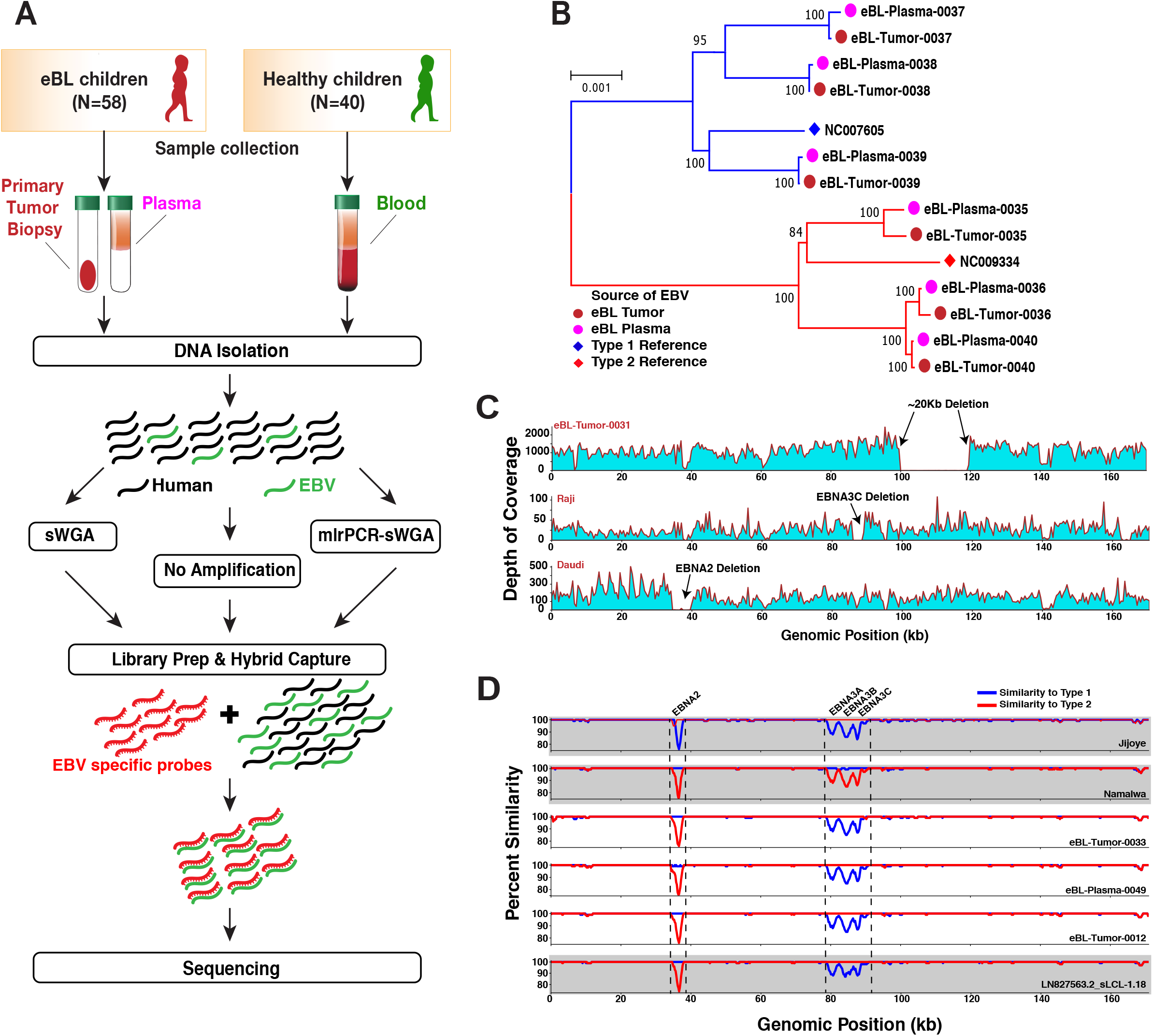
EBV genome sequencing from tumors and primary clinical samples. **A)** Overview of sample collection and methods for sequencing virus from Kenyan children diagnosed with eBL and healthy children as controls. Hybrid capture was universally performed along with additional amplification and enrichment steps to overcome low amounts of virus and input DNA. mlrPCR-sWGA; multiplexed long range PCR - specific whole genome amplification. **B)** Comparison of virus from paired tumor (brown circles) and plasma samples (pink circles) at diagnosis shows viral DNA circulating in the peripheral blood represents the virus in the tumor. The neighbor-Joining tree is scaled (0.001 substitutions per site) and includes standard reference genomes for type 1 (NC007605, blue diamond) and type 2 (NC009334, red diamond). **C)** The depth of coverage showing an absence of reads from approximately 100 kb to 120 kb is indicative of a large deletion in the virus from an eBL tumor (top panel). In the middle and lower panels, although we did not detect any in our tumor or control viruses, we had the power to detect deletions previously described in tumor lines including EBNA3C deletion in Raji and ENBA2 deletion in Daudi strains. **D)** Three intertypic viruses were detected by scanning across the genomes for percent identity in 1kb windows to both type 1 and type 2 references (NC_007605, NC_009334, respectively). Top two graphs (grey) represent controls, Jijoye and Namalwa, followed by 3 intertypic viruses from this study and one publicly available intertypic virus (LN827563.2_sLCL-1.18 in grey).

**Table 1.** Characteristics of children included in EBV sequencing analysis.

|  |  | eBL Patients<br>(N=58) | Healthy Controls<br>(N=40) |
| --- | --- | --- | --- |
| Age at collection,<br>N (%) | <6 (yrs) | 16 (27.6) | 39 (97.5) |
|  | 7 - 13 (yrs) | 42 (72.4) | 1 (2.5) |
| Sex, N (%) | Female/Male | 15/43 (25.9/74.1) | 20/20 (50.0/50.0) |
| Obtained<br>Specimen, N (%) | Tumor biopsy | 41 (41.8) | - |
|  | Blood | - | 40 (100.0) |
|  | Plasma | 14 (14.2) | - |
|  | New cultured eBL | 3 (3.0) | - |

### Sequencing and assembly quality

EBV is a large GC-rich double stranded DNA virus with 172 kb genome of which ~20% is repetitive sequence. For the majority of eBL patients, we prepared sequencing libraries directly from tumor DNA followed by hybrid capture enrichment. For low copy viral samples, such as eBL plasma and healthy control blood, we designed and implemented additional viral whole genome amplification and enrichment prior to library preparation and sequencing (**Figure 1A; Supplementary Figure 1**). We generated a study set of 114 genomes including replicates from cell lines and primary clinical samples, representing 98 cases and controls. In addition, we sequenced 20 technical replicates for quality control purposes such as estimation of re-sequencing error or sWGA bias, and sensitivity of detection of mixed infections. The baseline re-sequencing error rate was limited to ~1.1×10^-5^ bases when our assemblies are compared with high-quality known strain genomes^43^ (**Supplemental Table 2**). The mean error rate was ~2.1×10^-5^ bases for sWGA with GenomiPhi, while it is ~1.1×10^-4^ bases when we used more sensitive mlrPCR-sWGA (Methods). We obtained an average of ~5 million reads, resulting in an average 9,688 depth of coverage across assemblies (**Supplemental Table 3**). De novo sequence assembly created large scaffolds covering non-repetitive regions, except three isolates with low coverage, yielded a median of 137,887bp genomes (ranging 47,534bp - 146,920bp). We determined the types of each isolate by calculating the nucleotide distance to both reference types in addition to read mapping rates against type-specific regions. Despite our ability to experimentally detect mixed types at levels as low as 10% (**Supplemental Figure 2A**), we found no evidence of mixed infections in our cases and controls. Also, to ensure that our sample inclusion was unbiased when selecting healthy individuals with high enough viremias to sequence, we compared the viral loads and found no significant difference between type 1 and 2 (*P=0.126*, **Supplemental Figure 2B**).

### Equivalence of tumor and plasma viral DNA in eBL cases

The viral genomes from eBL cases included virus reconstructed from plasma and tumor samples. We confirmed that viral DNA in the plasma was representative of the virus in the tumor cells by sequencing plasma-tumor pairs from 6 eBL patients (**Figure 1B**). Accounting for the sequencing errors, the pairs appeared to be identical. Besides these plasma-tumor pairs, we further confirmed identical EBV types with additional pairs from 8 separate patients using type-specific PCRs. Overall, these findings demonstrate that viral DNA isolated from plasma represents the tumor virus.

### Structural variation and intertypic recombinants

First, we looked for large deletions within our viral genomes, but did not detect any of the previously described deletions in EBNAs, even though we were able to detect, as positive controls, EBNA3C deletion in Raji and the EBNA2 deletion in Daudi cell lines. However, in one sample we did detect a novel 20kb deletion, spanning from 100 kb to 120 kb in the genome (**Figure 1C**), which contains lytic phase genes *BBRF1/2, BBLF1/3, BGLF1/2/3/4/5*, and *BDLF2/3/4*. Interestingly, none of the latent genes were affected by this deletion.

Next, we interrogated our isolates by comparing the pairwise similarities of each genome against EBV type 1 and type 2 references. By traversing through the genome with a window, we were able to delineate regions that were more similar to one type over the other (**Figure 1D**). As expected, Jijoye, a type 2 strain, displayed less similarity against type 1 reference around its *EBNA2* and *EBNA3* genes, the most divergent region between types, while Namalwa as a type 1 strain shows the same pattern of dissimilarity against type 2 reference around the same regions. Interestingly, we found three patient-derived genomes, eBL-Tumor-0012, eBL-Tumor-0033, and eBL-Plasma-0049, with mixed similarity trends. Similar to a previously detected recombinant strain (LN827563.2_sLCL-1.18),^43^ all of the intertypic isolates carried type 1 *EBNA2* and type 2 *EBNA3* genes. Although not significant (*P=0.268*), these new intertypic hybrids were all isolated from eBL patients while we did not detect any in healthy controls.

### Genomic population structure is driven by type differences with distinct substructure in type 2 viruses

Our samples present a unique opportunity to study population structure of EBV types and their co-evolution within a geographically defined region. As expected, the major bifurcation within the phylogenetic tree based on the entire genome occurs between type 1 and type 2 viruses (**Figure 2A**). Viruses from eBL patients as well as healthy controls appeared to be intermixed almost randomly within the type 1 branch. Interestingly, within type 2 genomes 8 eBL-associated isolates formed a sub-cluster. The hybrid genomes clustered with type 2s, which is consistent with type 2 EBNA3s representing a greater amount of sequence than type 1 EBNA2 region.

**Figure 2.**
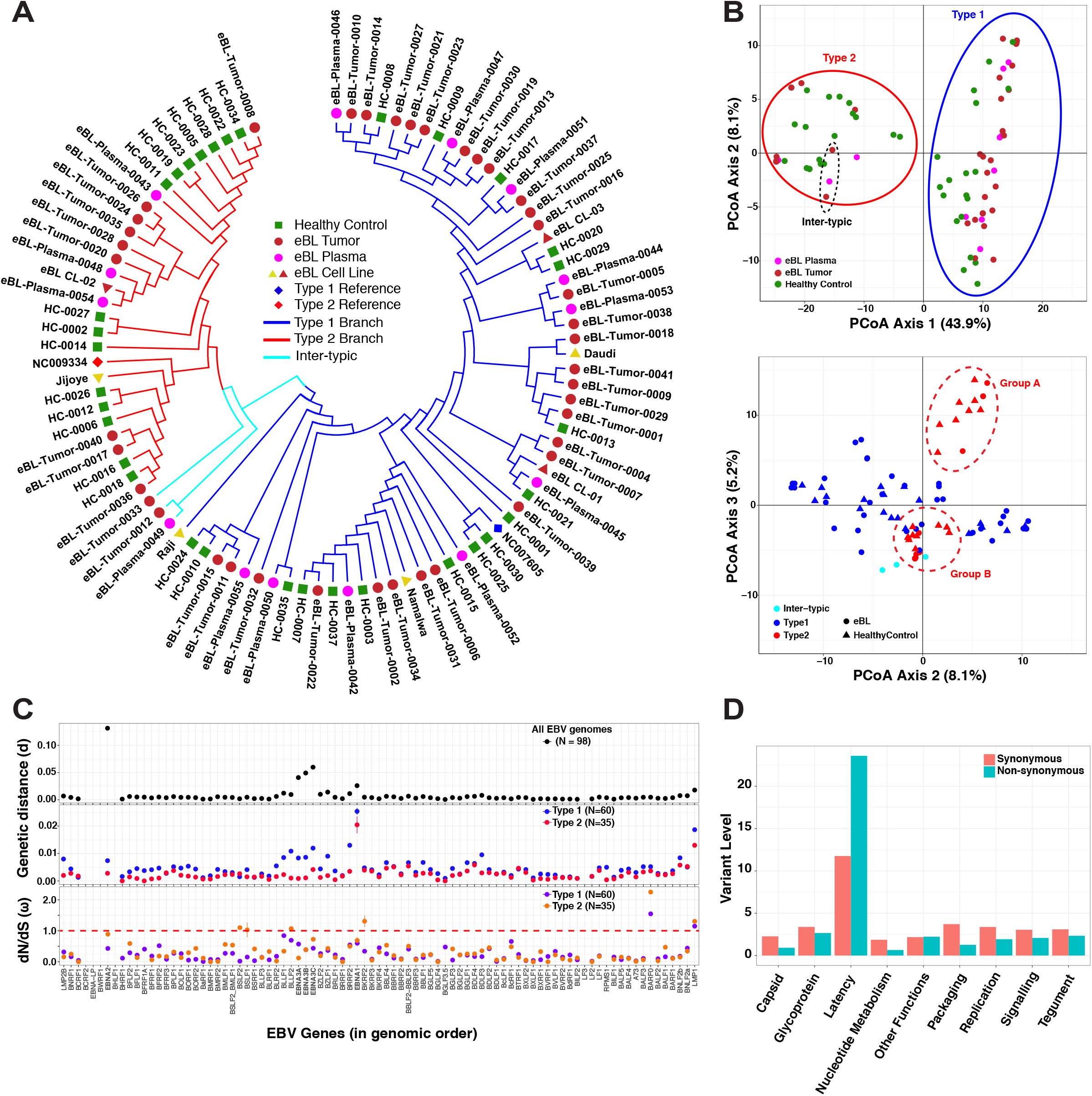
Diversity analysis of EBV genomes and coding genes in Kenyan population. **A)** Phylogenetic tree of the Western Kenya EBV genomes demonstrating the major type 1 and type 2 demarcation (blue and red branches, respectively). Pairwise distance calculations were based on Jukes-Cantor nucleotide substitution model, and the tree was constructed with the simple Neighbor-Joining method. Genomes are colored based on sample type: healthy children blood (green squares), eBL tumors (brown circles), plasma of eBL children (pink circles), and new and previous cell lines (brown and yellow triangles, respectively). Low coverage genomes are excluded. **B)** Principal coordinates analysis plots of nucleotide variations among whole genome sequences with first and second axes (upper plot, colored by sample type), and second and third axes (lower plot, colored by EBV subtype and shapes represent case and control). **C)** Genetic distance metrics of each EBV gene calculated based on Kimura-2-parameter method averaged across all genomes (upper panel) or type 1 / type 2 separately (middle panel). Lower panel shows nonsynonymous to synonymous change (dN/dS) ratios of viral protein coding genes averaged across all pairwise comparisons with in each group separately. Error bars represent standard error of mean. (Three intertypic genomes are excluded). **D)** Average synonymous and non-synonymous variants in genes are summarized as functional categories of genes. Variant level represents the number of variants per gene normalized by gene length in kb.

We further explored viral population structure with principal coordinate analysis (PCoA) of variation across the genome. While the first three components cumulatively explain 57.2% of the total variance, the first component, which solely accounted for 43.9% of the variance, separates genomes based on type 1 and type 2 (**Figure 2B, upper plot**). Similar to the phylogenetic tree, intertypic genomes positioned more closely to type 2s. Interestingly, the second and predominantly third components separate type 2 viruses into two distinct clusters, group A and B (**Figure 2B, lower plot**). These clusters were reflected, although not as distinctly, in the structure of the tree as well. The PCoA loading values, which accounts for 37.1% of the variance between the type 2 groups, are predominantly driven by correlated variation spanning 70kb upstream of EBNA3C (**Supplemental Figure 3A and B**). Together these findings suggest that there are two EBV type 2 strains circulating within this population. We also examined viral variation from the perspective of LMP1. Interestingly, the vast majority of viruses were grouped into Alaskan and Mediterranean strains (**Supplemental Figure 4**). For all available LMP1 type 2 sequences, group A and group B correlated with Mediterranean and Alaskan, respectively.

### EBV type 2 has less diversity compared with type 1

We further explored the pattern and nature of genomic variation across the genome comparing and contrasting EBV type 1 and type 2. Examining the pairwise divergence of coding genes for all viral genomes, we found that the divergence was the highest in the type-specific *EBNA* genes (*EBNA2* and *EBNA3s*), in particular, with *EBNA2* showing the greatest divergence (d=0.1313 ± 2.3×10^-3^) (**Figure 2C, upper panel**). Investigating each type separately, the diversity within types was low for *EBNA2* and *EBNA3Cs*, consistent with type 1 and 2 being separated by many fixed differences (**Figure 2C, middle panel**). In both types, intra-type divergence was greatest for *EBNA1* and *LMP1*. Most remarkable was the fact that type 2 generally showed lower levels of divergence across the genome (0.0047 ± 3.7×10^-3^ and 0.0025 ± 2.7×10^-3^ for type 1 and type 2, respectively). Overall, these measures suggest that EBV gene evolutionary rates differ by types.

To explore signatures of evolutionary selection, we examined the dN/dS ratios within coding sequences (**Figure 2C, lower panel**). Overall most genes showed signals of purifying selection, as indicated by ω < 1.0, except *LMP1, BARF0*, and *BKRF2* (only type 2). Interestingly, with dN/dS measures, *EBNA2, BSLF1, BSLF2*, and *BLLF2* genes had relatively higher rates in type 2 compared to type 1 suggestive of differential evolutionary pressure. Overall, the magnitude of average nonsynonymous and synonymous changes per gene, normalized by gene length, reflect the high-level diversity accumulated in certain genes (**Supplemental Figure 5**). Latency-associated genes generally have the highest non-synonymous variant rates, but they also have the highest synonymous rates consistent with longstanding divergence (**Figure 2D**). Other functional categories, including lytic genes, have relatively low levels of nonsynonymous mutations suggesting stronger purifying selection.

### Global context of Kenyan viruses

To more broadly contextualize our viral population from western Kenya, we examined the phylogeny of the Kenyan viruses along with other publicly available genomes from across the world (**Supplemental Table 4**). Among all isolates, the most polymorphic genomic regions appeared to be around *EBNA2* and *EBNA3* genes (**Supplemental Figure 6A**). Phylogenetic tree shows that the major types, type 1 and type 2, are the main demarcation point regardless of the source or geographic location. The three intertypic genomes from our sample set neatly cluster with the previously isolated intertypic hybrid, sLCL-1.18 (**Supplemental Figure 6B**). Type 1 genomes from our study were split into two groups, with one forming a sub-branch only with Kenyan type 1, including Mutu, Daudi, and several Kenyan LCLs. The second group interspersed with other African (Ghana, Nigeria, North Africa) and non-African isolates. In addition, a few of our genomes from healthy carriers clustered with a group of mainly Australian isolates, however; none of them clustered with South Asian group. Our Kenyan EBV type 2s generally intermixed with other type 2 genomes.

### Viral Genomic Variants and Associations with eBL

After excluding the intertypic hybrids, we compared type frequencies of EBV genomes isolated from eBL patients and healthy controls. We observed a significant difference in frequencies with 74.5% of eBLs carrying type 1 while only 25.5% carried type 2 infections. In contrast, 47.5% vs. 52.5% of type 1 and type 2, respectively were found in healthy controls. EBV type 1 was associated with eBL (OR=3.24, 95% CI=1.36 - 7.71, *P = 0.007*, Fisher’s exact) (**Figure 3A**), independent of age and gender (all *P>0.05*, **Supplemental Figure 7**). We then expanded the association analysis to all 2198 non-synonymous single nucleotide variations across the entire genome (**Figure 3B**). We did an initial association test for each nonsynonymous variant and detected 133 significant associations (**Supplemental Table 5 & Methods**). The vast majority of these variants were located within the type1-type2 region given the highly correlated nature of this region. We then stratified by type to detect variation independent of viral type. This yielded 6 variants solely associated with eBL (**Table 2, Supplemental Table 5**). Variant 37668T>C represents a serine residue change to a proline at the C-terminus of *EBNA2* (S485P) which is carried by 24/54 eBL cases; while this variant was present in only 2/36 healthy controls. Two variants in *EBNA1* at 95773A>T and 95778T>G (N38Y and H39Q, respectively) were both observed in 3/57 eBL isolates while their corresponding frequencies were 11/36 and 12/37 among healthy controls.

**Figure 3.**
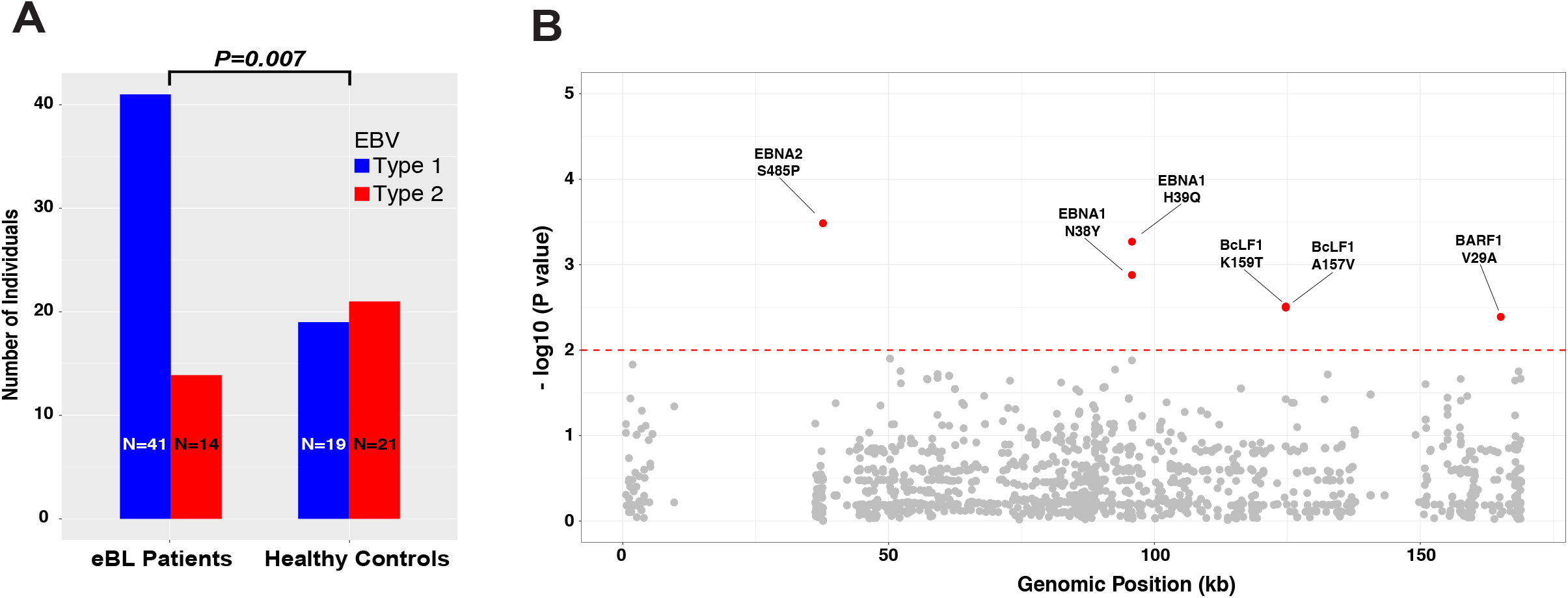
Significant associations of EBV type 1 genomes and single nucleotide variants with eBL. **A)** The frequency of type 1 and type 2 genomes identified from eBL patients and healthy control children (excluding the three intertypic hybrid genomes is significantly different (*P=0.007*, Fisher’s exact). **B)** Manhattan plot for genome-wide associations of non-synonymous single nucleotide variants tested for frequency differences between cases and controls controlling for type specific variants. The significance of each locus association is represented with an empirical p-value (negative log10 scale) that was calculated by 1 million permutations with random label swapping. Permutations were stratified for EBV genome type and adjusted for the missing genotypes due to lack of coverage. All significant variants associated with eBL cases are shown in red (*P < 0.01*). Nucleotide positions are according to type 1 reference genome.

**Table 2.** Single nucleotide variants associated with eBL. Single nucleotide variant association test results with P < 0.01 after type stratification. Table summarizes the statistically significant single nucleotide variant associations and their effects in the coding regions. Reference is the genotype based on the consensus of all genomes in the sequencing set and variant position denotes the projection to type 1 reference genome (NC_007605). The association test has been performed for every variant position comparing the frequency of reference and alternative (minor allele) bases among eBL patient and healthy control children (Fisher’s exact test). Empirical p values were based on one million permutations. *Genomes with missing data (Ns, lack of coverage) were excluded. Ref: reference allele, Alt: alternative/variant allele, AA: amino acid, P: p-value, OR: odds ratio.

| Gene | Position | Ref | Alt | AA<br>Change | eBLs |  | Healthy Controls |  | P | OR |
| --- | --- | --- | --- | --- | --- | --- | --- | --- | --- | --- |
|  |  |  |  |  | Genotypes* | Alt<br>Count | Genotypes* | Alt<br>Count |  |  |
| EBNA2 | 37668 | T | C | S485P | 54 | 24 | 36 | 2 | 0.000328 | 0.1 |
| EBNA1 | 95773 | A | T | N38Y | 57 | 3 | 36 | 11 | 0.001322 | 6.67213 |
| EBNA1 | 95778 | T | G | H39Q | 57 | 3 | 37 | 12 | 0.000538 | 7.16129 |
| BcLF1 | 124703 | T | G | K159T | 56 | 1 | 34 | 7 | 0.003178 | 12.7377 |
| BcLF1 | 124709 | G | A | A157V | 56 | 1 | 34 | 7 | 0.003092 | 12.7377 |
| BARF1 | 165131 | T | C | V29A | 57 | 36 | 36 | 10 | 0.004082 | 0.349462 |

Nucleotide variants in non-coding and promoter regions can affect regulation of viral gene expression and activity within host cell. *BZLF1* is a regulator gene of lytic reactivation and classified based on its promoter as prototype Zp-P (B95-8) and Zp-V3 (M81 strain).^44^ We determined variants at seven positions in the upstream promoter region of *BZLF1* (**Supplemental Table 6**). Interestingly, all of the Kenyan viruses carried C at positions both −525 and −274 (as in Zp-P) regardless of promoter type. We also found that −532 and −524 are variable in our isolates while these two are not variant in both promoter types. Our results show that only 12.5% (5/40) type 1 promoter sequences fully resembled Zp-V3 in eBL group as opposed to 22% (2/9) healthy genomes, while all of the type 2 genomes, without exception, carried Zp-V3 type promoter regardless of disease status.

## Discussion

In this study, we investigated genomic diversity of EBV by sampling virus from children in western Kenya where eBL incidence is high.^41^ Our improved methods allowed us to sequence asymptomatically infected healthy controls with relatively low peripheral blood viral loads, and thereby examine the virus in the population at large.^42^ We performed the first association study comparing viral genomes from eBL patients and geographically matched controls, without the need for viral propagation in LCLs; thus showing that type 1 EBV, as well as potentially several non-type specific variants, are associated with eBL. Furthermore, as the first study that characterized significant numbers of EBV type 2, we were able to compare and contrast both types and explore the viral population, thus discovering novel differences including population substructure in EBV type 2.

Our sequencing data demonstrated that EBV from plasma is representative of the tumor virus in eBL patients. This is consistent with the premise that peripheral EBV DNA originates from apoptotic tumor cells given that cell-free EBV DNA in eBL patients are mostly unprotected against DNase^45^, as opposed to being encapsidated during lytic reactivation, and that plasma EBV levels are associated with tumor burden and stage.^46^ These findings support the use of plasma viremia as a surrogate biomarker and the development of plasma-based prognostic tests with predictive models that could be used during clinical trials.^46^ The lack of mixed infections observed in our healthy controls could be due to the limit of detection in blood compared to virus isolated from saliva.^14^ Further studies are needed to understand the coevolution and dynamics of both EBV types.

In addition, we detected three intertypic recombinant EBV genomes solely found within our eBL patients; similar to those previously described in other cancers.^47^ It is unclear whether the intertypic genomes represent a common event with subsequent mutation and recombination or multiple independent events. If the latter is true, it supports more frequent mixed-type infections given that both parents have to be present in the same cell.^48–50^ It is interesting that all four intertypics observed to date carry the same type *EBNA2/EBNA3* combinations with the type 2 genes being so closely related (**Supplemental Figure 8**). Thus, if multiple events have generated these viruses, it suggests that certain strains may have a greater proclivity to recombine. Further studies will be needed to better define the intertypic population, their origins and their association with disease.

Importantly, we were able to explore EBV population genetics and compare and contrast type 1 and type 2 because of their co-prevalence in Africa. As well described, the major differentiation in terms of genetic variability was the variation correlated with type 1 and type 2 viruses. These viral types showed distinct population characteristics with type 1 harboring greater diversity especially in functionally important latent genes. Combined with the observed nucleotide diversity, latency genes appear to have long standing divergence that has accumulated significant synonymous changes (as opposed to recent sweeps on nonsynonymous changes that would erase synonymous variants). Global phylogenetic analysis emphasizes this diversity by providing two main subgroups for type 1 genomes in our sequencing set. One group represents core local Kenyan viruses while the second group is a mixture of viruses from across the globe, with the exception of South Asian viruses that group apart. While previously sequenced type 2 viruses intermingle with western Kenya isolates, the majority of these originated from East Africa with only a few from West Africa. Interestingly, intermingling is also true for type 2 as we observed two distinct groups. This is more apparent in PCA where type 2 virus forms 2 clusters. Examination via PCA, the loading values are determined by a broad stretch of the genome from the end of *EBNA3C* to *LMP1*, where Mediterranean and Alaskan designations correlate. It remains to be determined whether this substructure might be due to the introduction of previously geographically isolated viruses or distinct evolutionary trajectories within the population. Further study is needed with broader samplings to understand its significance but our findings suggest that there may be significant epistasis potentially including *LMP1*.

By sequencing virus directly from healthy controls, we were able to address the question of relative tumorigenicity between type 1 and 2. We tested the long-standing hypothesis that type 1 virus is more strongly associated with eBL, in contrast to type 2. Our work was able to more definitely answer this question as we were not reliant on LCLs from healthy controls where type 1 bias in transformation might explain the lack of previous associations. We earlier demonstrated, by mutational profiling of EBV positive and negative eBL tumors, that the virus, especially type 1, might mitigate the necessity of certain driver mutations in the host genome.^16^ In addition, our genome-wide results controlling for viral type substantiates investigations of non-type associated variation that could also impart oncogenic risk, as we found suggestive trends for several nonsynonymous variants as well. Only a small subset of type 1 viruses from eBL patients carried *BZLF1* promoter variant, which leads to a gain of function,^44^ while all type 2 viruses carried this variant suggesting this promoter might be beneficial for type 2 but makes it unlikely to be a driver of oncogenesis.

Overall, this population-based study provides the groundwork to unravel the complexities of EBV genome structure and insight into viral variation that influences oncogenesis. Genomic and mutational analysis of BL tumors identified key differences based on viral content suggesting new avenues for the development of prognostic molecular biomarkers and the potential for antiviral therapeutic interventions.

## Acknowledgements

This work was supported by the US National Institutes of Health, National Cancer Institute R01 CA134051, R01 CA189806 (A.M.M., J.A.B, C.I.O, Y.K.) and The Thrasher Research Fund 02833-7 (A.M.M.), UMCCTS Pilot Project Program U1 LTR000161-04 (Y.K., J.A.B., and A.M.M.), Turkish Ministry of National Education Graduate Study Abroad Program (Y.K.). We would like to thank the Kenyan children and their families who participated in this study. Patrick Marsh for helping with EBV genotyping assays, Mercedeh Movassagh for sharing genotyping primers. This publication was approved by the Director of KEMRI.

## Authorship Contributions

Contribution: Y.K., C.I.O., and O.A. designed and performed experiments; Y.K. and C.I.O analyzed and interpreted results; Y.K. made the figures; Y.K., J.A.B. and A.M.M. designed the research and wrote the paper, C.I.O, J.A.O., J.M.O., and A.M.M. organized clinical sample acquisition.

## References

1. Young LS, Rickinson AB. Epstein-Barr virus: 40 years on. Nat. Rev. Cancer. 2004;4(10):757–768.

2. Crawford DH. Biology and disease associations of Epstein-Barr virus. Philos. Trans. R. Soc. Lond. B Biol. Sci. 2001;356(1408):461–473.

3. Moormann AM, Bailey JA. Malaria—how this parasitic infection aids and abets EBV-associated Burkitt lymphomagenesis. Curr. Opin. Virol. 2016;

4. Torgbor C, Awuah P, Deitsch K, et al. A multifactorial role for P. falciparum malaria in endemic Burkitt’s lymphoma pathogenesis. PLoS Pathog. 2014;10(5):e1004170.

5. Simone O, Bejarano MT, Pierce SK, et al. TLRs innate immunereceptors and Plasmodium falciparum erythrocyte membrane protein 1 (PfEMP1) CIDR1α-driven human polyclonal B-cell activation. Acta Trop. 2011;119(2-3): 144–150.

6. Robbiani DF, Deroubaix S, Feldhahn N, et al. Plasmodium Infection Promotes Genomic Instability and AID-Dependent B Cell Lymphoma. Cell. 2015;162(4):727–737.

7. Cohen JI, Wang F, Mannick J, Kieff E. Epstein-Barr virus nuclear protein 2 is a key determinant of lymphocyte transformation. Proc. Natl. Acad. Sci. U. S. A. 1989;86(23):9558–9562.

8. Rowe M, Young LS, Cadwallader K, et al. Distinction between Epstein-Barr virus type A (EBNA 2A) and type B (EBNA 2B) isolates extends to the EBNA 3 family of nuclear proteins. J. Virol. 1989;63(3):1031–1039.

9. Dambaugh T, Hennessy K, Chamnankit L, Kieff E. U2 region of Epstein-Barr virus DNA may encode Epstein-Barr nuclear antigen 2. Proc. Natl. Acad. Sci. U. S. A. 1984;81(23):7632–7636.

10. Cho YG, Gordadze AV, Ling PD, Wang F. Evolution of two types of rhesus lymphocryptovirus similar to type 1 and type 2 Epstein-Barr virus. J. Virol. 1999;73(11):9206–9212.

11. Zimber U, Adldinger HK, Lenoir GM, et al. Geographical prevalence of two types of Epstein-Barr virus. Virology. 1986;154(1):56–66.

12. Apolloni A, Sculley TB. Detection of A-Type and B-Type Epstein-Bart Virus in Throat Washings and Lymphocytes. Virology. 1994;202(2):978–981.

13. Sixbey JW, Shirley P, Chesney PJ, Buntin DM, Resnick L. Detection of a second widespread strain of Epstein-Barr virus. Lancet. 1989;2(8666):761–765.

14. Correia S, Palser A, Elgueta Karstegl C, et al. Natural variation of Epstein-Barr virus genes, proteins and pri-miRNA (revised). J. Virol. 2017;

15. Young LS, Yao QY, Rooney CM, et al. New type B isolates of Epstein-Barr virus from Burkitt’s lymphoma and from normal individuals in endemic areas. J. Gen. Virol. 1987;68 (Pt 11):2853–2862.

16. Kaymaz Y, Oduor CI, Yu H, et al. Comprehensive Transcriptome and Mutational Profiling of Endemic Burkitt Lymphoma Reveals EBV Type-Specific Differences. Mol. Cancer Res. 2017;15(5):563–576.

17. Lucchesi W, Brady G, Dittrich-Breiholz O, et al. Differential gene regulation by Epstein-Barr virus type 1 and type 2 EBNA2. J. Virol. 2008;82(15):7456–7466.

18. Kaye KM, Izumi KM, Kieff E. Epstein-Barr virus latent membrane protein 1 is essential for B-lymphocyte growth transformation. Proc. Natl. Acad. Sci. U. S. A. 1993;90(19):9150–9154.

19. Wohlford EM, Asito AS, Chelimo K, et al. Identification of a novel variant of LMP-1 of EBV in patients with endemic Burkitt lymphoma in western Kenya. Infect. Agent. Cancer. 2013;8(1):34.

20. Chang CM, Yu KJ, Mbulaiteye SM, Hildesheim A, Bhatia K. The extent of genetic diversity of Epstein-Barr virus and its geographic and disease patterns: a need for reappraisal. Virus Res. 2009;143(2):209–221.

21. Chiara M, Manzari C, Lionetti C, et al. Geographic Population Structure in Epstein-Barr Virus Revealed by Comparative Genomics. Genome Biol. Evol. 2016;8(11):3284–3291.

22. Zhou L, Chen J-N, Qiu X-M, et al. Comparative analysis of 22 Epstein–Barr virus genomes from diseased and healthy individuals. J. Gen. Virol. 2017;98(1):96–107.

23. Depledge DP, Palser AL, Watson SJ, et al. Specific capture and whole-genome sequencing of viruses from clinical samples. PLoS One. 2011;6(11):e27805.

24. Kwok H, Wu CW, Palser AL, et al. Genomic diversity of Epstein-Barr virus genomes isolated from primary nasopharyngeal carcinoma biopsy samples. J. Virol. 2014;88(18):10662–10672.

25. Liu Y, Yang W, Pan Y, et al. Genome-wide analysis of Epstein-Barr virus (EBV) isolated from EBV-associated gastric carcinoma (EBVaGC). Oncotarget. 2016;7(4):4903–4914.

26. Wang S, Xiong H, Yan S, Wu N, Lu Z. Identification and Characterization of Epstein-Barr Virus Genomes in Lung Carcinoma Biopsy Samples by Next-Generation Sequencing Technology. Sci. Rep. 2016;6:26156.

27. Lei H, Li T, Li B, et al. Epstein-Barr virus from Burkitt Lymphoma biopsies from Africa and South America share novel LMP-1 promoter and gene variations. Sci. Rep. 2015;5:16706.

28. Parras-Moltó M, López-Bueno A. Methods for Enrichment and Sequencing of Oral Viral Assemblages: Saliva, Oral Mucosa, and Dental Plaque Viromes. Methods Mol. Biol. 2018;1838:143–161.

29. Buckle G, Maranda L, Skiles J, et al. Factors influencing survival among Kenyan children diagnosed with endemic Burkitt lymphoma between 2003 and 2011: A historical cohort study. Int. J. Cancer. 2016;139(6):1231–1240.

30. Leichty AR, Brisson D. Selective whole genome amplification for resequencing target microbial species from complex natural samples. Genetics. 2014;198(2):473–481.

31. Kwok H, Tong AHY, Lin CH, et al. Genomic sequencing and comparative analysis of Epstein-Barr virus genome isolated from primary nasopharyngeal carcinoma biopsy. PLoS One. 2012;7(5):e36939.

32. Martin M. Cutadapt removes adapter sequences from high-throughput sequencing reads. EMBnet.journal. 2011;17(1):10–12.

33. Schmieder R, Edwards R. Quality control and preprocessing of metagenomic datasets. Bioinformatics. 2011;27(6):863–864.

34. Consortium VB, Others. Velvetoptimiser. Available: bioinformatics. net. au/software. velvetoptimiser. shtml. Accessed. 2012;22.:

35. Swain MT, Tsai IJ, Assefa SA, et al. A post-assembly genome-improvement toolkit (PAGIT) to obtain annotated genomes from contigs. Nat. Protoc. 2012;7(7):1260–1284.

36. Katoh K, Standley DM. MAFFT multiple sequence alignment software version 7: improvements in performance and usability. Mol. Biol. Evol. 2013;30(4):772–780.

37. Kumar S, Stecher G, Tamura K. MEGA7: Molecular Evolutionary Genetics Analysis Version 7.0 for Bigger Datasets. Mol. Biol. Evol. 2016;33(7):1870–1874.

38. Page AJ, Taylor B, Delaney AJ, et al. SNP-sites: rapid efficient extraction of SNPs from multi-FASTA alignments. Microb Genom. 2016;2(4):e000056.

39. Gruber B, Unmack PJ, Berry OF, Georges A. dartr: An r package to facilitate analysis of SNP data generated from reduced representation genome sequencing. Mol. Ecol. Resour. 2018;18(3):691–699.

40. Ganeshan S, Dickover RE, Korber BT, Bryson YJ, Wolinsky SM. Human immunodeficiency virus type 1 genetic evolution in children with different rates of development of disease. J. Virol. 1997;71(1):663–677.

41. Rainey JJ, Mwanda WO, Wairiumu P, et al. Spatial distribution of Burkitt’s lymphoma in Kenya and association with malaria risk. Trop. Med. Int. Health. 2007;12(8):936–943.

42. Moormann AM, Chelimo K, Sumba OP, et al. Exposure to holoendemic malaria results in elevated Epstein-Barr virus loads in children. J. Infect. Dis. 2005;191(8):1233–1238.

43. Palser AL, Grayson NE, White RE, et al. Genome diversity of Epstein-Barr virus from multiple tumor types and normal infection. J. Virol. 2015;89(10):5222–5237.

44. Bristol JA, Djavadian R, Albright ER, et al. A cancer-associated Epstein-Barr virus BZLF1 promoter variant enhances lytic infection. PLoS Pathog. 2018;14(7):e1007179.

45. Mulama DH, Bailey JA, Foley J, et al. Sickle cell trait is not associated with endemic Burkitt lymphoma: An ethnicity and malaria endemicity-matched case--control study suggests factors controlling EBV may serve as a predictive biomarker for this pediatric cancer. International Journal of Cancer. 2014;134(3):645–653.

46. Westmoreland KD, Montgomery ND, Stanley CC, et al. Plasma Epstein-Barr virus DNA for pediatric Burkitt lymphoma diagnosis, prognosis and response assessment in Malawi. Int. J. Cancer. 2017;

47. Cho S-G, Lee W-K. Analysis of Genetic Polymorphisms of Epstein-Barr Virus Isolates from Cancer Patients and Healthy Carriers. J. Microbiol. Biotechnol. 2000;10(5):620–627.

48. Burrows JM, Khanna R, Sculley TB, et al. Identification of a naturally occurring recombinant Epstein-Barr virus isolate from New Guinea that encodes both type 1 and type 2 nuclear antigen sequences. J. Virol. 1996;70(7):4829–4833.

49. Yao QY, Tierney RJ, Croom-Carter D, et al. Isolation of intertypic recombinants of Epstein-Barr virus from T-cell-immunocompromised individuals. J. Virol. 1996;70(8):4895–4903.

50. Skare J, Farley J, Strominger JL, et al. Transformation by Epstein-Barr virus requires DNA sequences in the region of BamHI fragments Y and H. J. Virol. 1985;55(2):286–297.

