## Supplemental Methods, Supp. Figure/Table legends for "Epstein Barr virus genomes reveal population structure and type 1 association with endemic Burkitt lymphoma"

#### Cell cultures and controlled mixtures

BL cultured cell lines, Namalwa, Daudi, Raji, and Jijoye were grown in a complete growth medium, RPMI 1640 (Life Technologies), with 2mM L-glutamine adjusted to contain 1.5g/L sodium bicarbonate, 4.5g/L glucose, 10mM HEPES, 1.0mM sodium pyruvate, and 7.5% fetal bovine serum. We used Jijoye and Daudi as representative genomes of type 1 and type 2 strains. For mixing experiments, we created relative ratios of Jijoye:Daudi of 10:90, 25:75, 75:25, 90:10 in addition to sequencing each strain individually.

#### Sequence design for RNA baits

Capture bait sequences were designed using in-house scripts to target both type 1 and type 2. In addition to type 1 and type 2 references, we also designed against other available complete genomes including Mutu I, Akata, GD1 and GD2 to ensure the capture of divergent regions. Specifically, the design consisted of overlapping 120nt probes tiling every 30 bases (4x overlapping tiling) across the genomic sequences with increased number of probes for regions with elevated GC content (>65%). Additional probes were added based on the sequential analysis of additional genomes, when current probes were greater than 5% divergent or there was a gap in coverage for a specific region (**Supplemental Table 7**).

#### Multiplexed long range PCR (mlrPCR)

To increase viral DNA content in low abundant specimens, we applied an initial amplification with long range PCR using a strategy consisting of two multiplexed sets of primers which combined tiled the viral genome as published by Kwok et al.[1] To this, we added EBV type 2 specific primers. Following the initial multiplex long range (mlr) PCRs, we mixed two independent reactions and then performed specific whole genome amplification (sWGA) using phi29 polymerase with EBV-specific oligos. Overall DNA quality and quantities were assessed with NanoDrop and Picogreen and purified with 2x XP-Ampure magnetic beads. We prepared two reaction solutions with separate primer pools (2uL of 10uM each), using 2.5uL 10X long range PCR buffer $Mg^{2+}$ (Qiagen), 1.25uL dNTP (10mM each), 0.15uL long range PCR enzyme mix (Qiagen), 5uL 5X Q-solution (Qiagen) to which we added 10ng of input DNA in 14 uL. The reaction conditions involved initial denaturation at 95°C for 3 min followed by 20 cycles (95°C for 30 sec, gradient annealing 58°C to 49°C for 15 sec each, extension at 72°C for 7 min) and a final extension at 68°C for 10 min. We then mixed two independent reactions, denatured at 95°C for 3 min, and then added sWGA reaction buffer that contains 7uL 10X phi29 reaction buffer (NEB), 3uL dNTP mix (final concentrations: 30mM dGTP and dCTP; 10mM dATP and dTTP), 7uL EBV specific protected oligo Mix (10uM each), 2uL phi29 polymerase (10U/uL, NEB), 0.7uL BSA (0.1ug/uL), and 0.3uL H2O. For samples with higher viral loads that did not require PCR amplification prior to sWGA, we denatured DNA using the same conditions but replaced reaction buffer with TE or TE-Q-solution mix. We incubated the sWGA at 30°C for 16h followed by incubation at 65°C for 15 min to stop the reaction. Instead of random hexamers for MDA (multiple strand displacement amplification) reaction, we used EBV specific hexamers with 3'-end modification to protect against phi29 exonuclease activity. For WGA with Genomiphi v2 kit, we followed the manufacturer's instructions modified by adding extra 2x dGTP and dCTP. For hybrid capture, we followed MyBait protocol as manufacturer's recommendations. Following incubation at 65°C for 72 hours, we purified hybridization products with streptavidin beads and using Kapa HiFi to amplify the captured library. We quantified viral content with bi-plex qPCR using primers for viral BALF5 and human beta-actin gene.[2] For validation of EBV subtypes, we used primers spanning EBNA3C gene producing 153bp and 246bp products for type 1 and type 2, respectively.

We improved the amplification yield by adding extra 2x dGTP/dCTP to the amplification buffers, especially for low EBV inputs (10 EBV copy/uL) (**Supplemental Table 1**). We also tested the effect of Q-solution (Qiagen) on sWGA yield and found that EBV yields were almost doubled (**Supplemental Figure 1A and 1B**). In addition, we found that prolonged sWGA incubation time (16 hours) improved amplification yield compared with relatively shorter time (8 hours). Combining the above methods allowed for adequate input for hybrid capture even from low viral load healthy controls. We performed sequencing using Illumina MiSeq, HiSeq 2000, and NextSeq 500 platforms with 1x75bp, 2x100bp, and 2x150bp, respectively.

| | | | | |
|---|---|---|---|---|
| **Primer sequences used in multiplexed EBV qPCR for viral load measurements.** | | | | |
| BALF5; | Fwd: CGGAAGCCCTCTGGACTTC | | b-actin; | Fwd: TCACCCACACTGTGCCCATCTACGA |
| | Rev: CCCTGTTTATCCGATGGAATG | | | Rev: CAGCGGAACCGCTCATTGCCAATGG |
| **Primer sequences used in determining genomic subtype of EBV (Type I, Type 2)** | | | | |
| EBNA3C; | Fwd: AGAAGGGGAGCGTGTGTTG | | | |
| | Rev: GGCTCGTTTTTGACGTCGG | | | |
| **EBV specific 3'-protected oligos for MDA reactions using Phi29 polymerase** | | | | |
| | GCCGC**O**G | | | |
| | CCGCC**E**C | | | |
| | GGTCT**O**G | | | |
| | GCGGG**O**C | | | |
| | CGCCA**O**C | | | |
| | CCGCC**F**C | | | |
| | GTGGC**O**G | | | |
| | GGGCC**E**T | | | |
| | CGGGG**Z**C | | | |
| | GTCCG**E**G | | | |
| **Two pools of primer sequences used in multiplexed long-range PCR.** | | | | |
| **Pool1** | **Primer sequence** | | **Pool2** | **Primer sequence** |
| Fwd1-1 | TTCTGGTGATGCTTGTGCTC | | Fwd2-1 | CTGTTTATGAGACGCCAGCA |
| Rev1-1 | TGCTGGCGTCTCATAAACAG | | Rev2-1 | TTTTCGCTGCTTGTCCTTTT |
| Fwd1-2 | AAAAGGACAAGCAGCGAAAA | | Fwd2-2 | TTATGGTTCAGTGCGTCGAG |
| Rev1-2 | GTGCAGGAGGCTGTTTCTTC | | Rev2-2 | GAACTGAGGAGGGCATGAAG |
| Fwd1-3 | ATGCCTACATTCTATCTTGCGTTAC | | Fwd2-3 | AGGGATGCCTGGACACAAGA |
| Rev1-3 | TTACTGGATGGAGGGGCGAGGTCTT | | Rev2-3 | AACATGGACTGGGAGTGGAG |
| Fwd1-4 | CTAGAGGTCCGCGAGATTTG | | Fwd2-4 | GCAGGCAGTACGAGATGTCA |
| Rev1-4 | AGAAGGCAAGCGAAAATTGA | | Rev2-4 | TCCCTTCACATCCCAGAGAC |

| | | | |
|---|---|---|---|
| Fwd1-5 | CGACATTGACAGCCTTCTCA | Fwd2-5 | TGCTCCTGATGTTTCTGAGGTGGA |
| Rev1-5 | AAACACGAATGCCAAGAACC | Rev2-5 | AGGTAACTTCTTTGAGCCTCCCGA |
| Fwd1-6 | TTGCTCCATCTGTCAGCAAC | Fwd2-6 | GGTGACCACTGAGGGAGTGT |
| Rev1-6 | CACAAGCCTCCTCTCAGGAC | Rev2-6 | ATTTCAGGACTACCTGCGCGACTT |
| Fwd1-7 | GGACATCTCTGGCTCGAAAG | Fwd2-7 | TCAGGAGGTCGTCAAAATCC |
| Rev1-7 | AGGAGGAGAACCCGAGGATA | Rev2-7 | TTTCACATCCGACTCATTCCCTGC |
| Rev1-7-t2 | AGGAGGAGAACCCGAGGATC | Fwd2-8 | CCAGTCGCCGTTACTCATCT |
| Fwd1-8 | TCCAGGCTGTTGGAGAACACTTCA | Rev2-8 | ACCTTTCATCCGAACTCCTCAGGT |
| Rev1-8 | ATCACAGTCACCCCCAGAAG | Fwd2-9 | GCCTCTATGTCGCTCTGACC |
| Fwd1-9 | CAGACGGTGGCGTATATGAG | Rev2-9 | CGGAGGCGTGGTTAAATAAA |
| Rev1-9 | CAAAGAGCCCCGTAAAGATG | Fwd2-10 | CTCGCGTGTTAGGAAGGAAG |
| Fwd1-10 | GCGAGCCATAAAGCAGTTTC | Rev2-10 | AGGCAAAGCTGGTCAAAGAA |
| Rev1-10 | TCTCCCGAACTAGCAGCATT | Fw2-11-t2 | ATACATAGGAGCCTCACGAA |
| Fwd1-11 | GCCTTCTTTGACCAGCTTTG | Fwd2-11 | GGTGAAACGCGAGAAGAAAG |
| Rev1-11 | GACGGGTTCTACTGGCATGT | Rev2-11 | TTTAGCAGTTCCTCCGCACT |
| Rev1-11-t2 | GACGGGTTCTACTGGCATGG | Fwd2-12 | CCCACCACGTCTTCAACTTT |
| Fwd1-12 | AGTGCGGAGGAACTGCTAAA | Rev2-12 | CCATACCAGGTGCCTTTTGT |
| Rev1-12 | TGCAGAGGATGAGACCAGTG | Fwd2-12 | ACTCCCGGCTGTAAATTCCT |
| Fwd1-13 | TCCAAGGTGACCCCTGTTAG | Rev2-12 | TGGCCAGAAATACACCAACA |
| Rev1-13 | TGATGCAGAGTCGCCTAATG | Fwd2-13 | ACAGACCATCTACGCCAACC |
| Fwd1-14 | CCCATGTTGTCACGTCACTC | Rev2-13 | CCACCACAAGAAGGTGTCCT |
| Rev1-14 | CACCGTGTTGGAGACCTTTT | Fwd2-14 | GATGTTGCTGGGGCTAATGT |
| Fwd1-15 | TACGGGGCACTTAACCTGAC | Rev2-14 | AGAGAGGGAGTTTCGCTTCC |
| Rev1-15 | TGACGGAGCTGTATCACGAG | Fwd2-15 | CGTTGGAAGTTGTTGGGACT |
| Fwd1-16 | GGCACCATAGCATGTCACAC | Rev2-15 | CATTTTACCAGGGACGAGGA |
| Rev1-16 | AGTCCCAACAACTTCCAACG | Fwd2-16 | GGTCTCAACGTGTCCTGGTT |
| Fwd1-17 | CCCGTTCACCAAAACAGTCT | Rev2-16 | GTGAAGGTATGTGCCGGTCT |
| Rev1-17 | AACCAGGACACGTTGAGACC | Fwd2-17 | CCTGAGAACGCTCCAGGTAG |
| Fwd1-18 | ACCTCCCATAGCAACACCAG | Rev2-17 | CCTGGTGAGAAGTTGGTGGT |
| Rev1-18 | CCCGTGCGATGAGTTTATTT | Fwd2-18 | TTTGGGATGCATCACTTTGA |
| Fwd1-19 | CCAGACATACCCCAAACCAC | Rev2-18 | CCTCAAAGGTGTGGTCGTTT |
| Rev1-19 | CTCCAGAGGGCAGACGTTAG | Fwd2-19 | TCGTGGCTCGTACAGACGATTGTT |
| Fwd1-20 | GCCCGTTGGGTTACATTAAGGTGT | Rev2-19 | ACCTGGTACATTGTGCCCATCAGA |

| | | | | |
|---|---|---|---|---|
| Rev1-20 | CATGCAGTGGTGTCAGACAGGAAA | | Fwd2-20 | CCCACACCTTCACTCCTTGT |
| Fwd1-21 | CTTTGGGTTCCATTGTGTGCCCTT | | Rev2-20 | CAGAGCCAGGCACATCTACA |
| Rev1-21 | TTTGCGCCTTCTCCTGGTTTATGC | | Fwd2-21 | TGGAAGAAGGCGTAGAGCAT |
| Fwd1-22 | ACGCCATACCCAAGTGAGTC | | Rev2-21 | GCAAGGCTGACTCACCTGTTTGA |
| Rev1-23 | TCAAGAACCTGACGGAGCTT | | Fwd2-22 | AGGTTGCACACCACATCAAA |
| Fwd1-24 | ACGCCGAGTCATCTCTCATTTGGA | | Rev2-22 | GACTCGCTCACCCAAGAAAG |
| Rev1-24 | CGTGACTACCCCCACGTACT | | Fwd2-23 | CACGGGGTTTATGTTTCTGG |
| Fwd1-25 | GTGCAGAGCCTTGACATTGA | | Rev2-23 | CCCCCTCCACTTTTTCCA |
| Rev1-25 | TGAACACCACCACGATGACT | | EBNA2_Fw_Extra | TGGGAATGGTGTTAACTTTC |
| EBNA2_Fw_Extra | TGGGAATGGTGTTAACTTTC | | EBNA2_Rev_Extra | ATGTGTTGTGTGTGGTTTTG |
| EBNA2_Rev_Extra | ATGTGTTGTGTGTGGTTTTG | | | |

### References

1. Kwok H, Tong AHY, Lin CH, et al. Genomic sequencing and comparative analysis of Epstein-Barr virus genome isolated from primary nasopharyngeal carcinoma biopsy. *PLoS One*. 2012;7(5):e36939.
2. Moormann AM, Chelimo K, Sumba OP, et al. Exposure to holoendemic malaria results in elevated Epstein-Barr virus loads in children. *J. Infect. Dis.* 2005;191(8):1233–1238.

### Supplemental Figure Legends

**Supplemental Figure 1. Optimization of reaction solution and conditions for mlrPCR-sWGA.**

**A)** Optimization results of various dNTP concentrations in preamp reaction measured as EBV copy increase normalized by overall DNA increase. **B)** Incubation buffer, time and temperature optimization for better EBV copy increase.

**Supplemental Figure 2. Controls for putative mixed infections and sampling bias against EBV types.**

**A)** The sensitivity of EBV genome typing approach measured by accurate type assignments of in-lab mixtures with predefined ratios. Each mixture of Daudi (type 1) and Jijoye (type 2) with varying ratios was prepared in replicates. Following the genome assembly, type of the major strain was determined as judged by the distance to both reference viral genomes. **B)** Comparison of viral load levels of individuals carrying different EBV types (*P=0.126*, t-test). The non-significant difference suggests unbiased sampling among either type regardless of viral loads.

**Supplemental Figure 3. PCoA of only type 2 EBV genomes.**

**A)** First and second axises of PCoA using only type 2 genomes showing the separation of two groups. **B)** PCoA loading absolute values of Axis 1 in are plotted throughout the genome averaged across 1 kb window.

**Supplemental Figure 4. Phylogenetic analysis and main categories of LMP1 proteins in sequenced set.**

Tree of LMP1 proteins constructed by using the maximum likelihood method based on the Jukes-Cantor model. The shape of sample labels represents the group of individuals (square, healthy control; circle, eBL patients; and rectangle, cultured isolates) while color represents LMP1 strain classifications, Alaskan-Raji, Med (Mediterranean), China 1, NC (North Carolina). +/- signs of the Med strains denote the presence or absence of a 30bp deletion in the CTAR region.

**Supplemental Figure 5. Nucleotide variant rates per gene.**

Distribution of average synonymous and non-synonymous variants in viral genes normalized by the gene length. Genes appear in order of their genomic positions.

**Supplemental Figure 6. Global phylogenetic analysis of more than 220 isolates and divergent genomic regions.**

**A)** Representation of reference EBV genome in an open circular form as light blue tiles denote coding genes (and EBNAs highlighted as orange). Track showing accumulated variant positions reflected as polymorphic hotspots throughout the genome detected by multiple sequence alignments of our and all other existing assemblies (total 225 genomes). Bar heights represent the frequency of variant positions within 100 bp window outside of the repetitive regions (grey). **B)** Neighbor-Joining phylogenetic tree of the whole genomes demonstrating global diversity. Isolate names are colored based on geographic origin (African, brown; South Asian, red; Australian, blue; European, lime green-yellow; North America, light green; South American, black).

**Supplemental Figure 7. Covariate distributions of patients and individuals included in the study.**

The distribution of age, gender, and genomic type of EBV isolated from **A)** children diagnosed with eBL and **B)** healthy children. P values have been determined with Fisher's exact test.

**Supplemental Figure 8. Phylogenetic trees of various genes**

Intertypic genomes are highlighted.

### Supplemental Table Legends

**Supplemental Table 1.** Optimization of mlrPCR-sWGA and whole genome amplification reactions.

**Supplemental Table 2.** Estimated sequencing error rates based on replicates and controls.

**Supplemental Table 3.** Sample attributes, preprocessing information, and their sequencing data statistics.

**Supplemental Table 4**. EBV genomes previously published and used for comparison.

**Supplemental Table 5.** Single nucleotide variants with significance cut off P > 0.01.

**Supplemental Table 6.** Nucleotide variant compositions of each isolate in their BZLF1 gene upstream promoter region. Nucleotide positions are in upstream relative to TSS of BZLF1 gene. -532 and -524 are variant position within our isolates while these two are not in both promoter types.

**Supplemental Table 7.** RNA bait sequences used in EBV specific hybrid capture enrichment reactions
