## Supplemental Figures 1-8 for "Epstein Barr virus genomes reveal population structure and type 1 association with endemic Burkitt lymphoma"

### Supplemental Figure 1

A

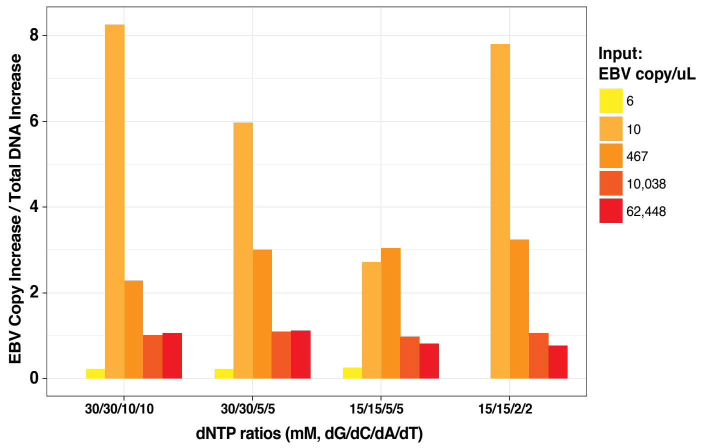


B

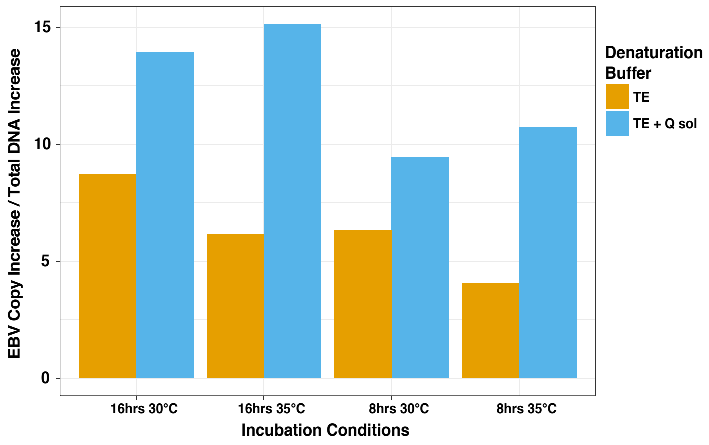

### Supplemental Figure 2

A

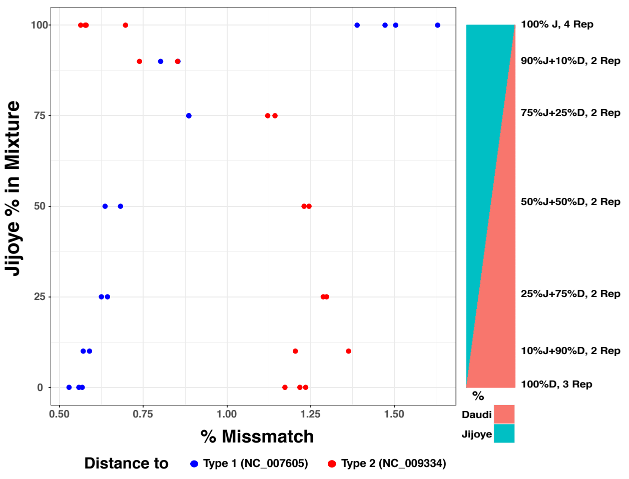


B

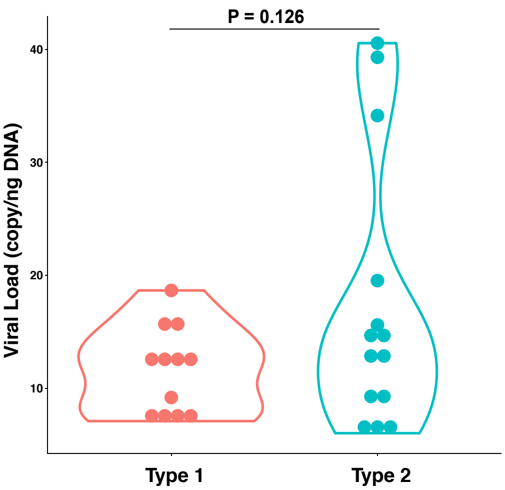

### Supplemental Figure 3

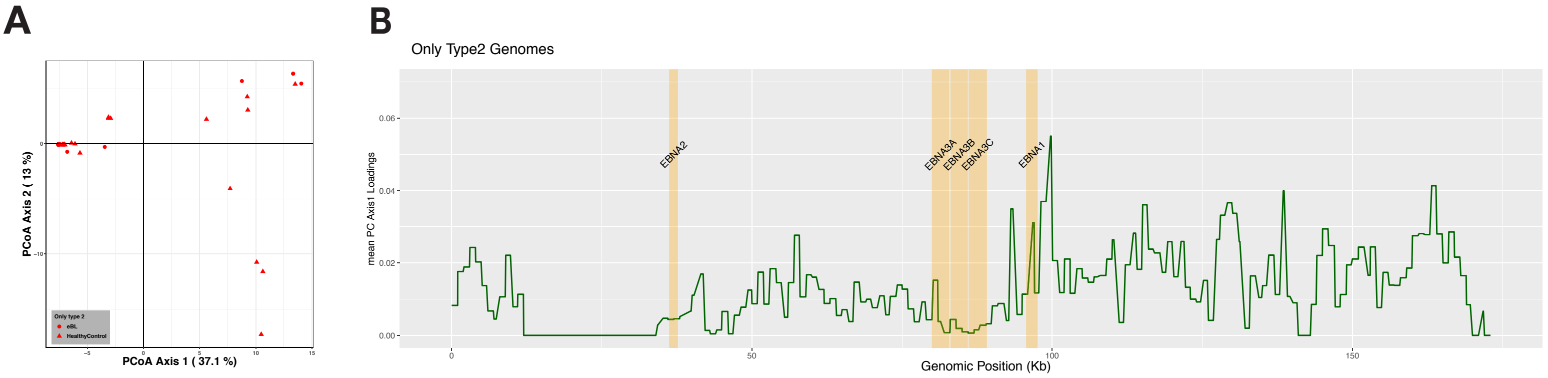

### Supplemental Figure 4

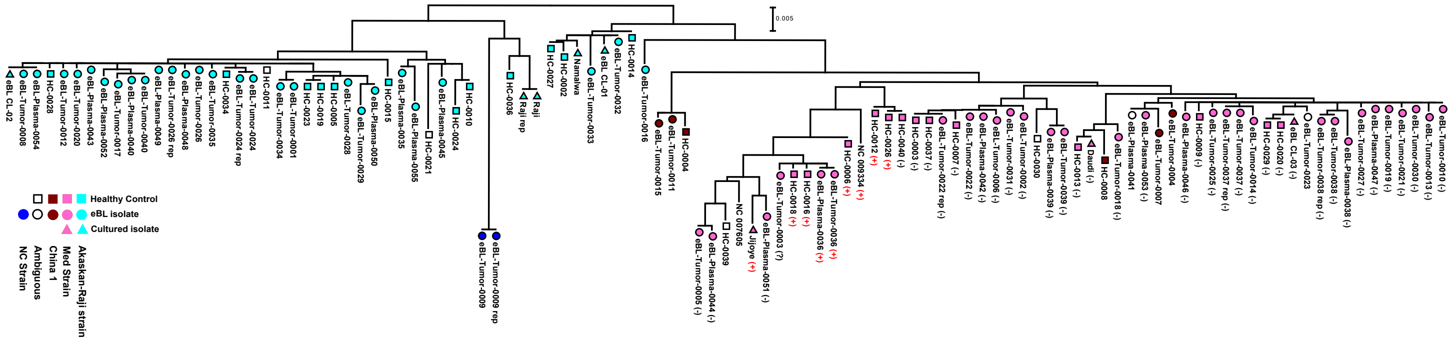

### Supplemental Figure 5

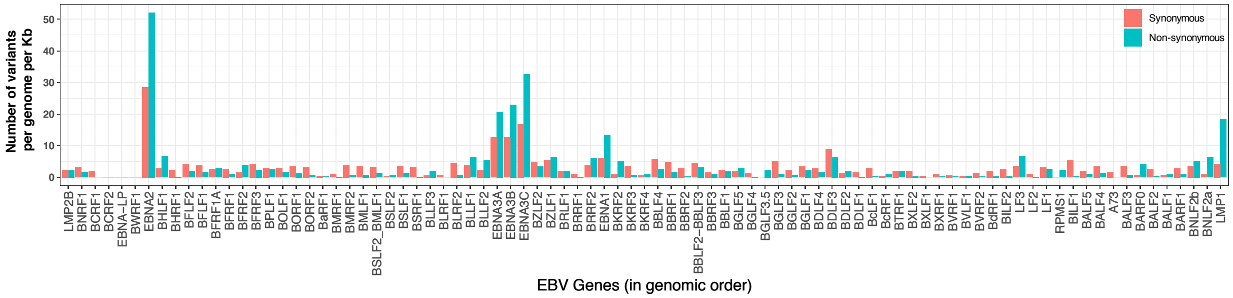

### Supplemental Figure 6

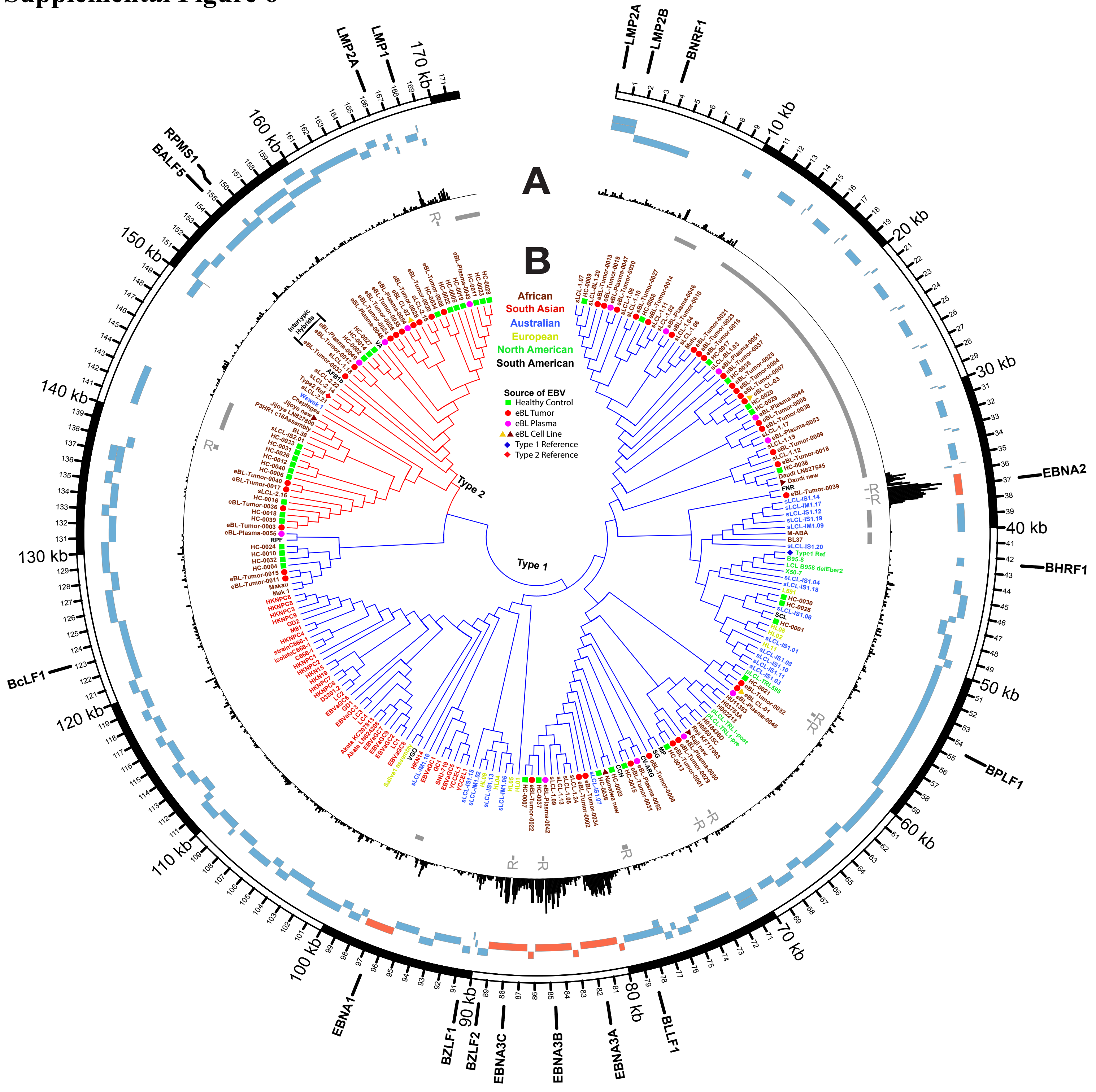

### Supplemental Figure 7

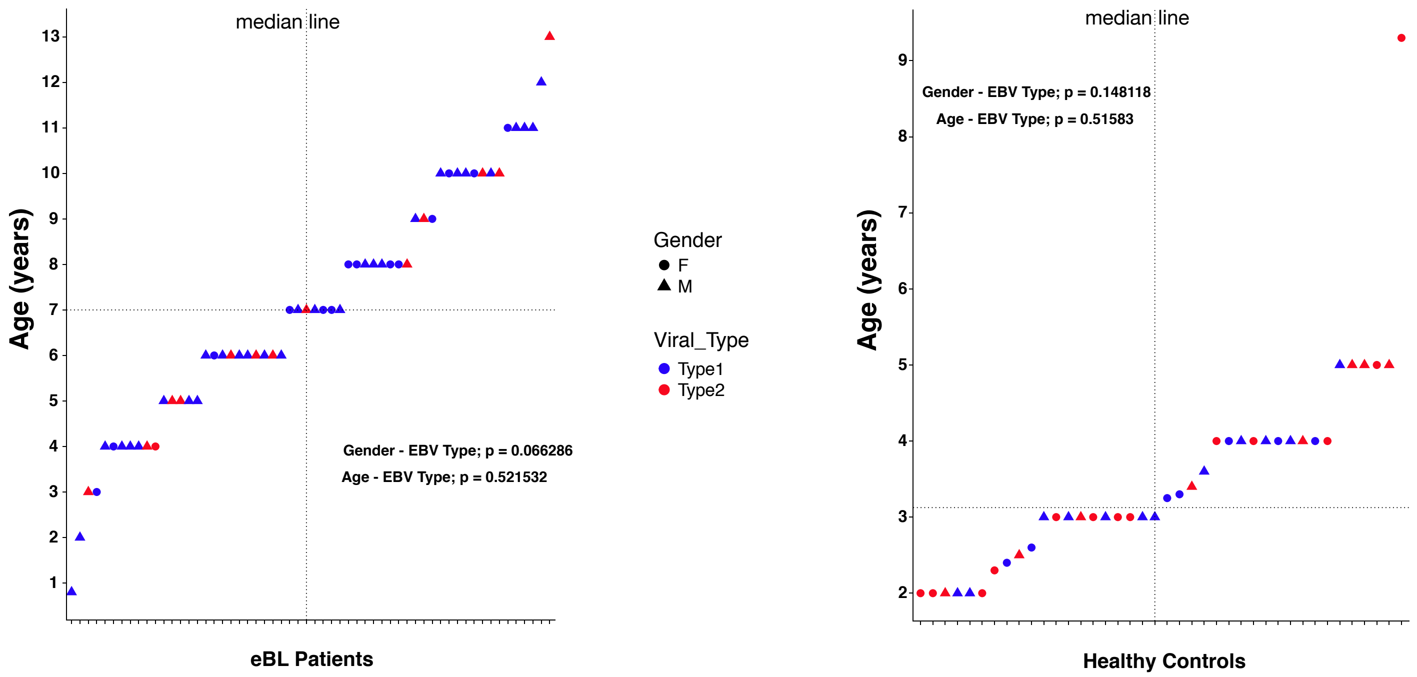

### Supplemental Figure 8

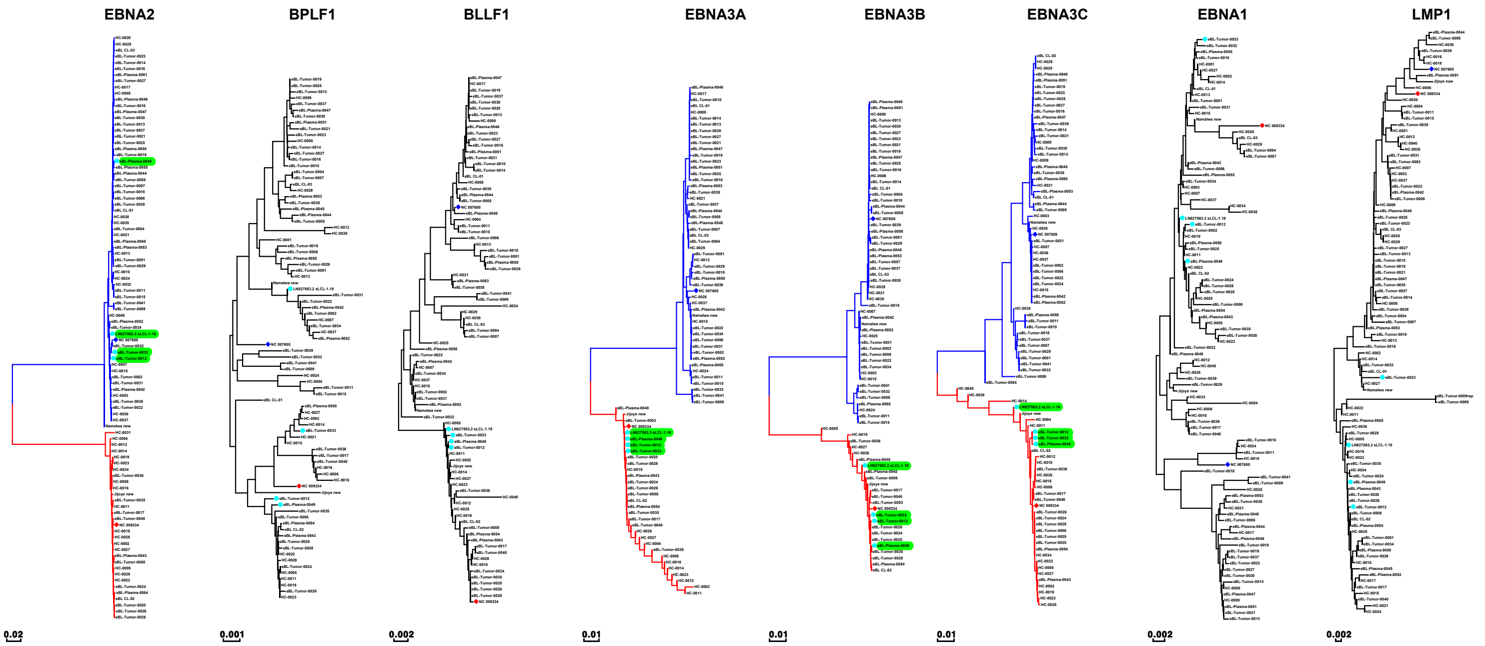
