## Supplemental Tables 1-7 for "Epstein Barr virus genomes reveal population structure and type 1 association with endemic Burkitt lymphoma"

**Supplemental Table 1. Optimization of mlrPCR-sWGA and whole genome amplification reactions.**

| | EBV Input copies for sWGA* | Denaturation Buffer | Average EBV Input (genome copy/uL) | Average Human Input (B-actin/uL) | dNTP composition (G/C/T/A ratio) | Whole Genome Amplifciation Incubation | EBV DNA Output (copy/ng) | Human DNA Output (B-actin/ng ) | EBV Enrichment (EBVpost / EBVpre) / (DNApost / DNApre) | Human Enrichment (Humanpost / Humanpre) / (DNApost / DNApre) | EBV Enrichment / Human Enrichment |
|---|---|---|---|---|---|---|---|---|---|---|---|
| Input EBV copies, dNTP Composition | 10,000 | TE | 62,448 | 3,456 | 30/30/10/10 | 16 hours at 30C | 4,573 | 7 | 1.06 | 0.03 | 35.33 |
| | 1,000 | TE | 10,038 | 13,911 | 30/30/10/10 | 16 hours at 30C | 505 | 59 | 1.01 | 0.08 | 12.63 |
| | 100 | TE | 467 | 10,883 | 30/30/10/10 | 16 hours at 30C | 55 | 70 | 2.28 | 0.12 | 19.00 |
| | 10 | TE | 10 | 11,326 | 30/30/10/10 | 16 hours at 30C | 4 | 68 | 8.26 | 0.12 | 68.83 |
| | 1 | TE | 6 | 19,483 | 30/30/10/10 | 16 hours at 30C | 0 | 55 | 0.21 | 0.06 | 3.50 |
| | 0 | TE | | 11,551 | 30/30/10/10 | 16 hours at 30C | | 76 | | 0.23 | 0.00 |
| | 10,000 | TE | 62,448 | 3,456 | 30/30/5/5 | 16 hours at 30C | 4,847 | 2 | 1.12 | 0.01 | 112.00 |
| | 1,000 | TE | 10,038 | 13,911 | 30/30/5/5 | 16 hours at 30C | 544 | 75 | 1.09 | 0.11 | 9.91 |
| | 100 | TE | 467 | 10,883 | 30/30/5/5 | 16 hours at 30C | 72 | 65 | 3 | 0.12 | 25.00 |
| | 10 | TE | 10 | 11,326 | 30/30/5/5 | 16 hours at 30C | 3 | 65 | 5.97 | 0.11 | 54.27 |
| | 1 | TE | 6 | 19,483 | 30/30/5/5 | 16 hours at 30C | 0 | 81 | 0.22 | 0.09 | 2.44 |
| | 0 | TE | | 11,551 | 30/30/5/5 | 16 hours at 30C | | 71 | | 0.21 | 0.00 |
| | 10,000 | TE | 62,448 | 3,456 | 15/15/5/5 | 16 hours at 30C | 3,483 | 16 | 0.81 | 0.07 | 11.57 |
| | 1,000 | TE | 10,038 | 13,911 | 15/15/5/5 | 16 hours at 30C | 488 | 84 | 0.97 | 0.12 | 8.08 |
| | 100 | TE | 467 | 10,883 | 15/15/5/5 | 16 hours at 30C | 73 | 109 | 3.04 | 0.19 | 16.00 |
| | 10 | TE | 10 | 11,326 | 15/15/5/5 | 16 hours at 30C | 1 | 80 | 2.71 | 0.14 | 19.36 |
| | 1 | TE | 6 | 19,483 | 15/15/5/5 | 16 hours at 30C | 0 | 103 | 0.25 | 0.11 | 2.27 |
| | 0 | TE | | 11,551 | 15/15/5/5 | 16 hours at 30C | | 118 | | 0.35 | 0.00 |
| | 10,000 | TE | 62,448 | 3,456 | 15/15/2/2 | 16 hours at 30C | 3,324 | 12 | 0.77 | 0.05 | 15.40 |
| | 1,000 | TE | 10,038 | 13,911 | 15/15/2/2 | 16 hours at 30C | 533 | 77 | 1.06 | 0.11 | 9.64 |
| | 100 | TE | 467 | 10,883 | 15/15/2/2 | 16 hours at 30C | 78 | 105 | 3.24 | 0.19 | 17.05 |
| | 10 | TE | 10 | 11,326 | 15/15/2/2 | 16 hours at 30C | 4 | 105 | 7.8 | 0.18 | 43.33 |
| | 1 | TE | 6 | 19,483 | 15/15/2/2 | 16 hours at 30C | 0 | 110 | 0 | 0.12 | 0.00 |
| | 0 | TE | | 11,551 | 15/15/2/2 | 16 hours at 30C | | 108 | | 0.32 | 0.00 |
| Denaturation Buffer, Incubation time and temperature | 10 | TE | 10 | 11,326 | 30/30/5/5 | 8 hours at 30C | 2 | 40 | 6.31 | 0.14 | 45.07 |
| | 10 | TE | 10 | 11,326 | 30/30/5/5 | 16 hours at 30C | 2 | 40 | 8.74 | 0.14 | 62.43 |
| | 10 | TE | 10 | 11,326 | 30/30/5/5 | 8 hours at 35C | 1 | 22 | 4.05 | 0.08 | 50.63 |
| | 10 | TE | 10 | 11,326 | 30/30/5/5 | 16 hours at 35C | 2 | 22 | 6.14 | 0.08 | 76.75 |
| | 10 | TE+Q sol | 10 | 11,326 | 30/30/5/5 | 8 hours at 30C | 2 | 41 | 9.44 | 0.14 | 67.43 |
| | 10 | TE+Q sol | 10 | 11,326 | 30/30/5/5 | 16 hours at 30C | 3 | 60 | 13.94 | 0.21 | 66.38 |
| | 10 | TE+Q sol | 10 | 11,326 | 30/30/5/5 | 8 hours at 35C | 3 | 28 | 10.72 | 0.1 | 107.20 |
| | 10 | TE+Q sol | 10 | 11,326 | 30/30/5/5 | 16 hours at 35C | 4 | 49 | 15.13 | 0.17 | 89.00 |

* total EBV genomes 20 uL of input for sWGA

**Supplemental Table 2. Estimated sequencing error rates based on replicates and controls.**

| Prior Amplification | Control Experimental Assembly | Reference Assembly | Number of Substitution Errors | Number of Correct Bases | Error Rate (per base) | Mean Error Rate (per base) |
|---|---|---|---|---|---|---|
| No amplification | Jijoye | Jiyoye Assembly* | 1 | 134,118 | 7.46E-06 | 1.13E-05 |
| No amplification | Daudi | Daudi Assembly* | 0 | 132,780 | 0.00E+00 | |
| No amplification | Raji_Rep1 | Raji Assembly* | 0 | 130,560 | 0.00E+00 | |
| No amplification | Raji_Rep2 | Raji Assembly* | 5 | 132,736 | 3.77E-05 | |
| GenomiPhi-WGA | Raji_wga | Raji Assembly* | 2 | 132,836 | 1.51E-05 | 2.13E-05 |
| GenomiPhi-WGA | eBLtumor-01_EBV_type1_wga | eBLtumor-01_EBV_type1# | 1 | 140,069 | 7.14E-06 | |
| GenomiPhi-WGA | eBLtumor-02_EBV_type2_wga | eBLtumor-02_EBV_type2# | 4 | 142,932 | 2.80E-05 | |
| GenomiPhi-WGA | eBLtumor-03_EBV_type2_wga | eBLtumor-03_EBV_type2# | 1 | 142,154 | 7.03E-06 | |
| GenomiPhi-WGA | eBLtumor-04_EBV_type1_wga | eBLtumor-04_EBV_type1# | 5 | 141,643 | 3.53E-05 | |
| GenomiPhi-WGA | eBLtumor-05_EBV_type1_wga | eBLtumor-05_EBV_type1# | 5 | 141,643 | 3.53E-05 | |
| mlrPCR-sWGA | Raji_Rep1_mlrPCR-sWGA | Raji Assembly* | 6 | 98,507 | 6.09E-05 | 1.09E-04 |
| mlrPCR-sWGA | Raji_Rep2_mlrPCR-sWGA | Raji Assembly* | 11 | 130,538 | 8.43E-05 | |
| mlrPCR-sWGA | Jijoye_Rep1_mlrPCR-sWGA | Jiyoye Assembly* | 13 | 132,045 | 9.84E-05 | |
| mlrPCR-sWGA | Daudi_mlrPCR-sWGA | Daudi Assembly* | 25 | 130,160 | 1.92E-04 | |

*Reference assemblies from Palser et al. 2015 (Jijoye: LN827800, Daudi: LN827545, Raji: KF717093). #The isolates were sequenced without any amplification. Preprocess denotes whether sample DNA was amplified prior to sequencing library preparation. The number of substitutions were determined by pairwise whole genome alignments of control and reference assemblies. Error rates were referred to the average mismatches to reference assemblies after normalizing to total covered genomic regions. GenomiPhi-WGA; whole genome amplification using EBV specific protected hexamers, mlrPCR-sWGA; pre-amplification with PCR primer pools followed by sWGA.

**Supplemental Table 3. Sample attributes, preprocessing information, and their sequencing data statistics.**

| SampleID | Sample Source | Amplification | Capture Method | Viral Type | Illumina Paired-end Read Length | Total Sequence reads | Assembled Genome Size (bases) | Average Depth of Coverage over Assembly |
|---|---|---|---|---|---|---|---|---|
| eBL_CL-01 | New BL_CellLineDNA | No amplification | EBV specific MyBait oligos | Type1 | 300 | 27728 | 137836 | 60.3 |
| eBL_CL-02 | New BL_CellLineDNA | No amplification | EBV specific MyBait oligos | Type2 | 300 | 37293 | 140684 | 79.5 |
| eBL_CL-03 | New BL_CellLineDNA | No amplification | EBV specific MyBait oligos | Type1 | 300 | 130447 | 142246 | 275.1 |
| Daudi | BL_CellLineDNA | No amplification | EBV specific MyBait oligos | Type1 | 300 | 114210 | 138211 | 247.9 |
| Jijoye | BL_CellLineDNA | No amplification | EBV specific MyBait oligos | Type2 | 300 | 15094 | 138424 | 32.7 |
| Namalwa | BL_CellLineDNA | No amplification | EBV specific MyBait oligos | Type1 | 300 | 29866 | 131405 | 68.2 |
| Raji | BL_CellLineDNA | No amplification | EBV specific MyBait oligos | Type1 | 300 | 16399 | 135972 | 36.2 |
| HC-0001 | BloodDNA_fromHealthyKid | mlrPCR-sWGA | EBV specific MyBait oligos | Type1 | 300 | 1989031 | 92740 | 6434.2 |
| HC-0002 | BloodDNA_fromHealthyKid | mlrPCR-sWGA | EBV specific MyBait oligos | Type2 | 300 | 2573344 | 135457 | 5699.2 |
| HC-0003 | BloodDNA_fromHealthyKid | mlrPCR-sWGA | EBV specific MyBait oligos | Type1 | 300 | 3256865 | 101378 | 9637.8 |
| HC-0004 | BloodDNA_fromHealthyKid | mlrPCR-sWGA | EBV specific MyBait oligos | Type2 | 300 | 6191128 | 130528 | 14229.4 |
| HC-0005 | BloodDNA_fromHealthyKid | mlrPCR-sWGA | EBV specific MyBait oligos | Type2 | 300 | 2230574 | 130100 | 5143.5 |
| HC-0006 | BloodDNA_fromHealthyKid | mlrPCR-sWGA | EBV specific MyBait oligos | Type2 | 300 | 14005054 | 113876 | 36895.5 |
| HC-0007 | BloodDNA_fromHealthyKid | mlrPCR-sWGA | EBV specific MyBait oligos | Type1 | 300 | 13554411 | 125967 | 32280.9 |
| HC-0008 | BloodDNA_fromHealthyKid | mlrPCR-sWGA | EBV specific MyBait oligos | Type1 | 300 | 4038485 | 112717 | 10748.6 |
| HC-0009 | BloodDNA_fromHealthyKid | mlrPCR-sWGA | EBV specific MyBait oligos | Type1 | 300 | 5264476 | 132675 | 11903.8 |
| HC-0010 | BloodDNA_fromHealthyKid | mlrPCR-sWGA | EBV specific MyBait oligos | Type1 | 300 | 5080095 | 91925 | 16579 |
| HC-0011 | BloodDNA_fromHealthyKid | mlrPCR-sWGA | EBV specific MyBait oligos | Type2 | 300 | 2361855 | 114610 | 6182.3 |
| HC-0012 | BloodDNA_fromHealthyKid | mlrPCR-sWGA | EBV specific MyBait oligos | Type2 | 300 | 4609922 | 114898 | 12036.6 |
| HC-0013 | BloodDNA_fromHealthyKid | mlrPCR-sWGA | EBV specific MyBait oligos | Type1 | 300 | 399047 | 104101 | 1150 |
| HC-0014 | BloodDNA_fromHealthyKid | mlrPCR-sWGA | EBV specific MyBait oligos | Type2 | 300 | 1373086 | 123167 | 3344.4 |
| HC-0015 | BloodDNA_fromHealthyKid | mlrPCR-sWGA | EBV specific MyBait oligos | Type1 | 300 | 3517550 | 128859 | 8189.3 |
| HC-0016 | BloodDNA_fromHealthyKid | mlrPCR-sWGA | EBV specific MyBait oligos | Type2 | 300 | 5312268 | 131404 | 12128.1 |
| HC-0017 | BloodDNA_fromHealthyKid | mlrPCR-sWGA | EBV specific MyBait oligos | Type1 | 300 | 1194255 | 89724 | 3993.1 |
| HC-0018 | BloodDNA_fromHealthyKid | mlrPCR-sWGA | EBV specific MyBait oligos | Type2 | 300 | 3443411 | 85526 | 12078.5 |
| HC-0019 | BloodDNA_fromHealthyKid | mlrPCR-sWGA | EBV specific MyBait oligos | Type2 | 300 | 1401820 | 137320 | 3062.5 |
| HC-0020 | BloodDNA_fromHealthyKid | mlrPCR-sWGA | EBV specific MyBait oligos | Type1 | 300 | 3297486 | 97564 | 10139.5 |
| HC-0021 | BloodDNA_fromHealthyKid | mlrPCR-sWGA | EBV specific MyBait oligos | Type1 | 300 | 1691945 | 110989 | 4573.3 |
| HC-0022 | BloodDNA_fromHealthyKid | mlrPCR-sWGA | EBV specific MyBait oligos | Type2 | 300 | 986623 | 120580 | 2454.7 |
| HC-0023 | BloodDNA_fromHealthyKid | mlrPCR-sWGA | EBV specific MyBait oligos | Type2 | 300 | 149295 | 132533 | 337.9 |
| HC-0024 | BloodDNA_fromHealthyKid | mlrPCR-sWGA | EBV specific MyBait oligos | Type1 | 300 | 1248198 | 128373 | 2917 |
| HC-0025 | BloodDNA_fromHealthyKid | mlrPCR-sWGA | EBV specific MyBait oligos | Type1 | 300 | 1179837 | 82870 | 4271.2 |
| HC-0026 | BloodDNA_fromHealthyKid | mlrPCR-sWGA | EBV specific MyBait oligos | Type2 | 300 | 555072 | 137682 | 1209.5 |
| HC-0027 | BloodDNA_fromHealthyKid | mlrPCR-sWGA | EBV specific MyBait oligos | Type2 | 300 | 457153 | 136835 | 1002.3 |
| HC-0028 | BloodDNA_fromHealthyKid | mlrPCR-sWGA | EBV specific MyBait oligos | Type2 | 300 | 1910804 | 129668 | 4420.8 |
| HC-0029 | BloodDNA_fromHealthyKid | mlrPCR-sWGA | EBV specific MyBait oligos | Type1 | 300 | 8132282 | 138631 | 17598.4 |
| HC-0030 | BloodDNA_fromHealthyKid | mlrPCR-sWGA | EBV specific MyBait oligos | Type1 | 150 | 514390 | 89660 | 860.6 |
| HC-0031 | BloodDNA_fromHealthyKid | mlrPCR-sWGA | EBV specific MyBait oligos | Type2 | 150 | 896398 | 15578 | 8631.4 |

| | | | | | | | | | |
|---|---|---|---|---|---|---|---|---|---|
| HC-0032 | BloodDNA_fromHealthyKid | mlrPCR-sWGA | EBV specific MyBait oligos | Type1 | 150 | 1230192 | 59837 | 3083.9 | |
| HC-0033 | BloodDNA_fromHealthyKid | mlrPCR-sWGA | EBV specific MyBait oligos | Type2 | 150 | 331520 | 16212 | 3067.4 | |
| HC-0034 | BloodDNA_fromHealthyKid | mlrPCR-sWGA | EBV specific MyBait oligos | Type2 | 150 | 11241210 | 97685 | 17261.4 | |
| HC-0035 | BloodDNA_fromHealthyKid | mlrPCR-sWGA | EBV specific MyBait oligos | Type1 | 150 | 24810 | 77012 | 48.3 | |
| HC-0036 | BloodDNA_fromHealthyKid | mlrPCR-sWGA | EBV specific MyBait oligos | Type1 | 150 | 3822180 | 57343 | 9998.2 | |
| HC-0037 | BloodDNA_fromHealthyKid | mlrPCR-sWGA | EBV specific MyBait oligos | Type1 | 150 | 41852 | 108638 | 57.8 | |
| HC-0038 | BloodDNA_fromHealthyKid | mlrPCR-sWGA | EBV specific MyBait oligos | Type2 | 150 | 107808 | 3440 | 4700.9 | |
| HC-0039 | BloodDNA_fromHealthyKid | mlrPCR-sWGA | EBV specific MyBait oligos | Type2 | 150 | 1028620 | 47534 | 3246 | |
| HC-0040 | BloodDNA_fromHealthyKid | mlrPCR-sWGA | EBV specific MyBait oligos | Type2 | 150 | 9429614 | 111687 | 12664.3 | |
| eBL-Tumor-0001 | eBL_primaryTumorDNA | GenomiPhi-WGA | EBV specific MyBait oligos | Type1 | 200 & 300 | 4222227 | 142384 | 7413.5 | |
| eBL-Tumor-0002 | eBL_primaryTumorDNA | GenomiPhi-WGA | EBV specific MyBait oligos | Type1 | 200 & 300 | 6479721 | 142052 | 11403.8 | |
| eBL-Tumor-0003 | eBL_primaryTumorDNA | GenomiPhi-WGA | EBV specific MyBait oligos | Type2 | 200 & 300 | 8625903 | 60909 | 35404.9 | |
| eBL-Tumor-0004 | eBL_primaryTumorDNA | mlrPCR-sWGA | EBV specific MyBait oligos | Type1 | 300 | 1123872 | 137537 | 2451.4 | |
| eBL-Tumor-0005 | eBL_primaryTumorDNA | GenomiPhi-WGA | EBV specific MyBait oligos | Type1 | 200 & 300 | 4648123 | 142405 | 8160 | |
| eBL-Tumor-0006 | eBL_primaryTumorDNA | GenomiPhi-WGA | EBV specific MyBait oligos | Type1 | 200 & 300 | 6448397 | 143731 | 11216.1 | |
| eBL-Tumor-0007 | eBL_primaryTumorDNA | GenomiPhi-WGA | EBV specific MyBait oligos | Type1 | 200 & 300 | 5448856 | 142866 | 9534.9 | |
| eBL-Tumor-0008 | eBL_primaryTumorDNA | mlrPCR-sWGA | EBV specific MyBait oligos | Type2 | 300 | 10438253 | 141842 | 22077.2 | |
| eBL-Tumor-0009 | eBL_primaryTumorDNA | mlrPCR-sWGA | EBV specific MyBait oligos | Type1 | 300 | 926150 | 136703 | 2032.5 | |
| eBL-Tumor-0010 | eBL_primaryTumorDNA | mlrPCR-sWGA | EBV specific MyBait oligos | Type1 | 300 | 21509972 | 135211 | 47725.3 | |
| eBL-Tumor-0011 | eBL_primaryTumorDNA | GenomiPhi-WGA | EBV specific MyBait oligos | Type1 | 200 & 300 | 14949732 | 144063 | 25943 | |
| eBL-Tumor-0012 | eBL_primaryTumorDNA | GenomiPhi-WGA | EBV specific MyBait oligos | Inter-typic | 200 & 300 | 3673227 | 141985 | 6467.6 | |
| eBL-Tumor-0013 | eBL_primaryTumorDNA | GenomiPhi-WGA | EBV specific MyBait oligos | Type1 | 200 & 300 | 7861933 | 144125 | 13637.4 | |
| eBL-Tumor-0014 | eBL_primaryTumorDNA | GenomiPhi-WGA | EBV specific MyBait oligos | Type1 | 200 & 300 | 8701944 | 143606 | 15149 | |
| eBL-Tumor-0015 | eBL_primaryTumorDNA | GenomiPhi-WGA | EBV specific MyBait oligos | Type1 | 200 & 300 | 5825147 | 144044 | 10110 | |
| eBL-Tumor-0016 | eBL_primaryTumorDNA | GenomiPhi-WGA | EBV specific MyBait oligos | Type1 | 200 & 300 | 1161216 | 139661 | 2078.6 | |
| eBL-Tumor-0017 | eBL_primaryTumorDNA | No amplification | EBV specific MyBait oligos | Type2 | 200 | 10699217 | 145418 | 14715.1 | |
| eBL-Tumor-0018 | eBL_primaryTumorDNA | No amplification | EBV specific MyBait oligos | Type1 | 200 | 11258545 | 143781 | 15660.7 | |
| eBL-Tumor-0019 | eBL_primaryTumorDNA | No amplification | EBV specific MyBait oligos | Type1 | 200 | 16192472 | 143175 | 22619.1 | |
| eBL-Tumor-0020 | eBL_primaryTumorDNA | No amplification | EBV specific MyBait oligos | Type2 | 200 | 14633495 | 146920 | 19920.4 | |
| eBL-Tumor-0021 | eBL_primaryTumorDNA | No amplification | EBV specific MyBait oligos | Type1 | 200 | 14430586 | 143942 | 20050.6 | |
| eBL-Tumor-0022 | eBL_primaryTumorDNA | No amplification | EBV specific MyBait oligos | Type1 | 200 | 7687200 | 143947 | 10680.6 | |
| eBL-Tumor-0022 | eBL_primaryTumorDNA | GenomiPhi-WGA | EBV specific MyBait oligos | Type1 | 200 | 7248781 | 142454 | 10177 | Repeated for error estimation |
| eBL-Tumor-0023 | eBL_primaryTumorDNA | No amplification | EBV specific MyBait oligos | Type1 | 200 | 4481944 | 140842 | 6364.5 | |
| eBL-Tumor-0024 | eBL_primaryTumorDNA | No amplification | EBV specific MyBait oligos | Type2 | 200 | 2585251 | 143612 | 3600.3 | |
| eBL-Tumor-0024 | eBL_primaryTumorDNA | GenomiPhi-WGA | EBV specific MyBait oligos | Type2 | 200 | 4883458 | 142754 | 6841.8 | Repeated for error estimation |
| eBL-Tumor-0025 | eBL_primaryTumorDNA | No amplification | EBV specific MyBait oligos | Type1 | 200 | 9775107 | 143175 | 13654.8 | |
| eBL-Tumor-0026 | eBL_primaryTumorDNA | No amplification | EBV specific MyBait oligos | Type2 | 200 | 4070074 | 142949 | 5694.4 | |
| eBL-Tumor-0026 | eBL_primaryTumorDNA | GenomiPhi-WGA | EBV specific MyBait oligos | Type2 | 200 | 6415050 | 142375 | 9011.5 | Repeated for error estimation |
| eBL-Tumor-0027 | eBL_primaryTumorDNA | No amplification | EBV specific MyBait oligos | Type1 | 200 | 2473450 | 142313 | 3476.1 | |
| eBL-Tumor-0028 | eBL_primaryTumorDNA | No amplification | EBV specific MyBait oligos | Type2 | 200 | 2785996 | 142423 | 3912.3 | |
| eBL-Tumor-0029 | eBL_primaryTumorDNA | No amplification | EBV specific MyBait oligos | Type1 | 200 | 7602074 | 142738 | 10651.8 | |
| eBL-Tumor-0030 | eBL_primaryTumorDNA | No amplification | EBV specific MyBait oligos | Type1 | 200 | 6079669 | 145541 | 8354.6 | |

| | | | | | | | | | |
|---|---|---|---|---|---|---|---|---|---|
| eBL-Tumor-0031 | eBL_primaryTumorDNA | No amplification | EBV specific MyBait oligos | Type1 | 200 | 3972963 | 126974 | 6257.9 | |
| eBL-Tumor-0032 | eBL_primaryTumorDNA | No amplification | EBV specific MyBait oligos | Type1 | 200 | 7118274 | 144616 | 9844.4 | |
| eBL-Tumor-0033 | eBL_primaryTumorDNA | No amplification | EBV specific MyBait oligos | Inter-typic | 200 | 18395334 | 146197 | 25165.1 | |
| eBL-Tumor-0034 | eBL_primaryTumorDNA | No amplification | EBV specific MyBait oligos | Type1 | 200 | 5679398 | 144524 | 7859.5 | |
| eBL-Tumor-0035 | eBL_primaryTumorDNA | GenomiPhi-WGA | EBV specific MyBait oligos | Type2 | 200 & 300 | 6133367 | 142874 | 10732.1 | |
| eBL-Tumor-0036 | eBL_primaryTumorDNA | mlrPCR-sWGA | EBV specific MyBait oligos | Type2 | 300 | 1873678 | 137887 | 4076.6 | |
| eBL-Tumor-0037 | eBL_primaryTumorDNA | No amplification | EBV specific MyBait oligos | Type1 | 200 | 4781768 | 143452 | 6666.7 | |
| eBL-Tumor-0037 | eBL_primaryTumorDNA | GenomiPhi-WGA | EBV specific MyBait oligos | Type1 | 200 | 4097748 | 141698 | 5783.8 | Repeated for error estimation |
| eBL-Tumor-0038 | eBL_primaryTumorDNA | No amplification | EBV specific MyBait oligos | Type1 | 200 | 1652556 | 141363 | 2338 | |
| eBL-Tumor-0038 | eBL_primaryTumorDNA | GenomiPhi-WGA | EBV specific MyBait oligos | Type1 | 200 & 300 | 14668183 | 144437 | 25388.5 | Repeated for error estimation |
| eBL-Tumor-0039 | eBL_primaryTumorDNA | No amplification | EBV specific MyBait oligos | Type1 | 200 | 3484138 | 145453 | 4790.7 | |
| eBL-Tumor-0040 | eBL_primaryTumorDNA | No amplification | EBV specific MyBait oligos | Type2 | 200 | 3758609 | 145880 | 5153 | |
| eBL-Tumor-0041 | eBL_primaryTumorDNA | No amplification | EBV specific MyBait oligos | Type1 | 200 | 6158191 | 145025 | 8492.6 | |
| eBL-Plasma-0035 | PlasmaDNA_fromBLpatient | mlrPCR-sWGA | EBV specific MyBait oligos | Type2 | 300 | 385303 | 131409 | 879.6 | |
| eBL-Plasma-0036 | PlasmaDNA_fromBLpatient | mlrPCR-sWGA | EBV specific MyBait oligos | Type2 | 300 | 727755 | 139446 | 1565.7 | |
| eBL-Plasma-0037 | PlasmaDNA_fromBLpatient | mlrPCR-sWGA | EBV specific MyBait oligos | Type1 | 300 | 52615528 | 125217 | 126058.4 | |
| eBL-Plasma-0038 | PlasmaDNA_fromBLpatient | mlrPCR-sWGA | EBV specific MyBait oligos | Type1 | 300 | 2982299 | 135347 | 6610.3 | |
| eBL-Plasma-0039 | PlasmaDNA_fromBLpatient | mlrPCR-sWGA | EBV specific MyBait oligos | Type1 | 300 | 2834853 | 135720 | 6266.3 | |
| eBL-Plasma-0040 | PlasmaDNA_fromBLpatient | mlrPCR-sWGA | EBV specific MyBait oligos | Type2 | 300 | 336725 | 138187 | 731 | |
| eBL-Plasma-0041 | PlasmaDNA_fromBLpatient | mlrPCR-sWGA | EBV specific MyBait oligos | Type1 | 300 | 1055728 | 112128 | 2824.6 | |
| eBL-Plasma-0042 | PlasmaDNA_fromBLpatient | mlrPCR-sWGA | EBV specific MyBait oligos | Type1 | 300 | 1226736 | 138256 | 2661.9 | |
| eBL-Plasma-0043 | PlasmaDNA_fromBLpatient | mlrPCR-sWGA | EBV specific MyBait oligos | Type2 | 300 | 643566 | 137788 | 1401.2 | |
| eBL-Plasma-0044 | PlasmaDNA_fromBLpatient | mlrPCR-sWGA | EBV specific MyBait oligos | Type1 | 300 | 992336 | 137048 | 2172.2 | |
| eBL-Plasma-0045 | PlasmaDNA_fromBLpatient | mlrPCR-sWGA | EBV specific MyBait oligos | Type1 | 300 | 9600176 | 138091 | 20856.2 | |
| eBL-Plasma-0046 | PlasmaDNA_fromBLpatient | mlrPCR-sWGA | EBV specific MyBait oligos | Type1 | 300 | 7416764 | 112949 | 19699.4 | |
| eBL-Plasma-0047 | PlasmaDNA_fromBLpatient | No amplification | EBV specific MyBait oligos | Type1 | 300 | 2209552 | 141331 | 4690.2 | |
| eBL-Plasma-0048 | PlasmaDNA_fromBLpatient | No amplification | EBV specific MyBait oligos | Type2 | 300 | 117683 | 125628 | 281 | |
| eBL-Plasma-0049 | PlasmaDNA_fromBLpatient | No amplification | EBV specific MyBait oligos | Inter-typic | 300 | 352588 | 136504 | 774.9 | |
| eBL-Plasma-0050 | PlasmaDNA_fromBLpatient | No amplification | EBV specific MyBait oligos | Type1 | 300 | 528192 | 138955 | 1140.4 | |
| eBL-Plasma-0051 | PlasmaDNA_fromBLpatient | No amplification | EBV specific MyBait oligos | Type1 | 300 | 6039246 | 142313 | 12730.9 | |
| eBL-Plasma-0052 | PlasmaDNA_fromBLpatient | mlrPCR-sWGA | EBV specific MyBait oligos | Type1 | 300 | 1682856 | 138645 | 3641.4 | |
| eBL-Plasma-0053 | PlasmaDNA_fromBLpatient | mlrPCR-sWGA | EBV specific MyBait oligos | Type1 | 300 | 9207274 | 135925 | 20321.4 | |
| eBL-Plasma-0054 | PlasmaDNA_fromBLpatient | No amplification | EBV specific MyBait oligos | Type2 | 300 | 1063382 | 142801 | 2234 | |
| eBL-Plasma-0055 | PlasmaDNA_fromBLpatient | No amplification | EBV specific MyBait oligos | Type1 | 300 | 1514583 | 142029 | 3199.2 | |

**Supplemental Table 4. EBV genomes previously published and used for comparison.**

| Sample | NCBI sequence ID |
|---|---|
| LC1 | gi\|1034220039\|gb\|KT823506.1\| Human herpesvirus 4 isolate LC1, partial genome |
| LC2 | gi\|1034220119\|gb\|KT823507.1\| Human herpesvirus 4 isolate LC2, partial genome |
| LC3 | gi\|1034220201\|gb\|KT823508.1\| Human herpesvirus 4 isolate LC3, partial genome |
| LC4 | gi\|1034220278\|gb\|KT823509.1\| Human herpesvirus 4 isolate LC4, partial genome |
| Type2_Ref | gi\|139424470\|ref\|NC_009334.1\| Human herpesvirus 4, complete genome |
| GD2 | gi\|343177302\|gb\|HQ020558.1\| Human herpesvirus 4 strain GD2, complete genome |
| Akata | gi\|428161017\|gb\|KC207813.1\| Human herpesvirus 4 strain Akata, complete genome |
| Mutu | gi\|428161102\|gb\|KC207814.1\| Human herpesvirus 4 strain Mutu, complete genome |
| strainC666-1 | gi\|500154724\|gb\|KC617875.1\| Human herpesvirus 4 strain C666-1, partial genome |
| M81 | gi\|557368242\|gb\|KF373730.1\| Human herpesvirus 4 strain M81, complete genome |
| HKNPC1 | gi\|582051068\|gb\|JQ009376.2\| Human herpesvirus 4 strain HKNPC1, genome |
| HKNPC2 | gi\|663082492\|gb\|KF992564.1\| Human herpesvirus 4 isolate HKNPC2, partial genome |
| HKNPC3 | gi\|663082571\|gb\|KF992565.1\| Human herpesvirus 4 isolate HKNPC3, partial genome |
| HKNPC4 | gi\|663082648\|gb\|KF992566.1\| Human herpesvirus 4 isolate HKNPC4, partial genome |
| HKNPC5 | gi\|663082725\|gb\|KF992567.1\| Human herpesvirus 4 isolate HKNPC5, partial genome |
| HKNPC6 | gi\|663082802\|gb\|KF992568.1\| Human herpesvirus 4 isolate HKNPC6, partial genome |
| HKNPC7 | gi\|663082879\|gb\|KF992569.1\| Human herpesvirus 4 isolate HKNPC7, partial genome |
| HKNPC8 | gi\|663082958\|gb\|KF992570.1\| Human herpesvirus 4 isolate HKNPC8, partial genome |
| HKNPC9 | gi\|663083037\|gb\|KF992571.1\| Human herpesvirus 4 isolate HKNPC9, partial genome |
| isolateC666-1 | gi\|663083114\|gb\|KJ411974.1\| Human herpesvirus 4 isolate C666-1, partial genome |
| Raji | gi\|681961025\|gb\|KF717093.1\| Human herpesvirus 4 strain Raji, complete genome |
| Saliva1_assembly | gi\|764007273\|emb\|LN824142.1\| Human herpesvirus 4 genome assembly Saliva1_assembly, segment : I |
| Mak_1 | gi\|764007353\|emb\|LN824203.1\| Human herpesvirus 4 genome assembly Mak_1, segment : I |
| HL05 | gi\|764007436\|emb\|LN824204.1\| Human herpesvirus 4 genome assembly HL05, segment : I |

| | |
|---|---|
| sLCL-1.12 | gi\|764007519\|emb\|LN824205.1\| Human herpesvirus 4 genome assembly sLCL-1.12, segment : I |
| pLCL-TRL1-post | gi\|764007602\|emb\|LN824206.1\| Human herpesvirus 4 genome assembly pLCL-TRL1-post |
| pLCL-TRL1-pre | gi\|764007685\|emb\|LN824207.1\| Human herpesvirus 4 genome assembly pLCL-TRL1-pre, segment : I |
| Akata_Rep | gi\|764007768\|emb\|LN824208.1\| Human herpesvirus 4 genome assembly Akata, segment : I |
| HKN14 | gi\|764007851\|emb\|LN824209.1\| Human herpesvirus 4 genome assembly HKN14, segment : I |
| HKN19 | gi\|764007935\|emb\|LN824224.1\| Human herpesvirus 4 genome assembly HKN19, segment : I |
| HL08 | gi\|764008018\|emb\|LN824225.1\| Human herpesvirus 4 genome assembly HL08, segment : I |
| HL01 | gi\|764008102\|emb\|LN824226.1\| Human herpesvirus 4 genome assembly HL01, segment : I |
| HL09 | gi\|764008186\|emb\|LN827522.1\| Human herpesvirus 4 genome assembly HL09, segment : I |
| L591 | gi\|764008270\|emb\|LN827523.1\| Human herpesvirus 4 genome assembly L591, segment : I |
| HL11 | gi\|764008352\|emb\|LN827524.1\| Human herpesvirus 4 genome assembly HL11, segment : I |
| C666-1 | gi\|764008436\|emb\|LN827525.1\| Human herpesvirus 4 genome assembly C666-1, segment : I |
| BL37 | gi\|764008516\|emb\|LN827526.1\| Human herpesvirus 4 genome assembly BL37, segment : I |
| M-ABA | gi\|764008600\|emb\|LN827527.1\| Human herpesvirus 4 genome assembly M-ABA, segment : I |
| Wewak_1 | gi\|764008683\|emb\|LN827544.1\| Human herpesvirus 4 genome assembly Wewak_1, segment : I |
| Daudi | gi\|764008767\|emb\|LN827545.1\| Human herpesvirus 4 genome assembly Daudi, segment : I |
| HL02 | gi\|764008849\|emb\|LN827546.1\| Human herpesvirus 4 genome assembly HL02, segment : I |
| HKN15 | gi\|764008933\|emb\|LN827547.1\| Human herpesvirus 4 genome assembly HKN15, segment : I |
| D3201.2 | gi\|764009101\|emb\|LN827549.1\| Human herpesvirus 4 genome assembly D3201.2, segment : I |
| sLCL-1.11 | gi\|764009185\|emb\|LN827550.1\| Human herpesvirus 4 genome assembly sLCL-1.11, segment : I |
| Makau | gi\|764009268\|emb\|LN827551.1\| Human herpesvirus 4 genome assembly Makau, segment : I |
| sLCL-1.08 | gi\|764009351\|emb\|LN827552.1\| Human herpesvirus 4 genome assembly sLCL-1.08, segment : I |
| sLCL-IS1.08 | gi\|764009435\|emb\|LN827553.1\| Human herpesvirus 4 genome assembly sLCL-IS1.08, segment : I |
| AFB1b | gi\|764009519\|emb\|LN827554.1\| Human herpesvirus 4 genome assembly AFB1b, segment : I |
| X50-7 | gi\|764009602\|emb\|LN827555.1\| Human herpesvirus 4 genome assembly X50-7, segment : I |
| Cheptages | gi\|764009684\|emb\|LN827556.1\| Human herpesvirus 4 genome assembly Cheptages, segment : I |
| sLCL-1.02 | gi\|764009848\|emb\|LN827558.1\| Human herpesvirus 4 genome assembly sLCL-1.02, segment : I |

| | |
|---|---|
| pLCL-TRL595 | gi\|764009931\|emb\|LN827559.1\| Human herpesvirus 4 genome assembly pLCL-TRL595, segment : I |
| sLCL-2.14 | gi\|764010014\|emb\|LN827560.1\| Human herpesvirus 4 genome assembly sLCL-2.14, segment : I |
| YCCEL1 | gi\|764010098\|emb\|LN827561.1\| Human herpesvirus 4 genome assembly YCCEL1, segment : I |
| sLCL-1.19 | gi\|764010182\|emb\|LN827562.1\| Human herpesvirus 4 genome assembly sLCL-1.19, segment : I |
| HL04 | gi\|764010348\|emb\|LN827564.1\| Human herpesvirus 4 genome assembly HL04, segment : I |
| sLCL-1.07 | gi\|764010420\|emb\|LN827565.1\| Human herpesvirus 4 genome assembly sLCL-1.07, segment : I |
| sLCL-1.06 | gi\|764010504\|emb\|LN827566.1\| Human herpesvirus 4 genome assembly sLCL-1.06, segment : I |
| sLCL-IM1.09 | gi\|764010588\|emb\|LN827567.1\| Human herpesvirus 4 genome assembly sLCL-IM1.09, segment : I |
| sLCL-1.24 | gi\|764010672\|emb\|LN827568.1\| Human herpesvirus 4 genome assembly sLCL-1.24, segment : I |
| sLCL-IS1.11 | gi\|764010755\|emb\|LN827569.1\| Human herpesvirus 4 genome assembly sLCL-IS1.11, segment : I |
| sLCL-IS1.01 | gi\|764010839\|emb\|LN827570.1\| Human herpesvirus 4 genome assembly sLCL-IS1.01, segment : I |
| sLCL-BL1.20 | gi\|764010923\|emb\|LN827571.1\| Human herpesvirus 4 genome assembly sLCL-BL1.20, segment : I |
| sLCL-IS1.18 | gi\|764011007\|emb\|LN827572.1\| Human herpesvirus 4 genome assembly sLCL-IS1.18, segment : I |
| sLCL-1.10 | gi\|764011091\|emb\|LN827573.1\| Human herpesvirus 4 genome assembly sLCL-1.10, segment : I |
| sLCL-1.09 | gi\|764011175\|emb\|LN827574.1\| Human herpesvirus 4 genome assembly sLCL-1.09, segment : I |
| sLCL-IS1.14 | gi\|764011258\|emb\|LN827575.1\| Human herpesvirus 4 genome assembly sLCL-IS1.14, segment : I |
| sLCL-IS1.20 | gi\|764011341\|emb\|LN827576.1\| Human herpesvirus 4 genome assembly sLCL-IS1.20, segment : I |
| sLCL-1.17 | gi\|764011424\|emb\|LN827577.1\| Human herpesvirus 4 genome assembly sLCL-1.17, segment : I |
| sLCL-IS1.13 | gi\|764011508\|emb\|LN827578.1\| Human herpesvirus 4 genome assembly sLCL-IS1.13, chromosome : I |
| sLCL-1.13 | gi\|764011592\|emb\|LN827579.1\| Human herpesvirus 4 genome assembly sLCL-1.13, segment : I |
| sLCL-2.16 | gi\|764011675\|emb\|LN827580.1\| Human herpesvirus 4 genome assembly sLCL-2.16, segment : I |
| sLCL-1.05 | gi\|764011759\|emb\|LN827581.1\| Human herpesvirus 4 genome assembly sLCL-1.05, segment : I |
| sLCL-BL1.03 | gi\|764011842\|emb\|LN827582.1\| Human herpesvirus 4 genome assembly sLCL-BL1.03, segment : I |
| sLCL-IM1.17 | gi\|764011926\|emb\|LN827583.1\| Human herpesvirus 4 genome assembly sLCL-IM1.17, segment : I |
| sLCL-IS1.06 | gi\|764012009\|emb\|LN827584.1\| Human herpesvirus 4 genome assembly sLCL-IS1.06, segment : I |
| sLCL-1.04 | gi\|764012093\|emb\|LN827585.1\| Human herpesvirus 4 genome assembly sLCL-1.04, segment : I |
| sLCL-IS1.15 | gi\|764012177\|emb\|LN827586.1\| Human herpesvirus 4 genome assembly sLCL-IS1.15, segment : I |

| | |
|---|---|
| sLCL-2.21 | gi\|764012261\|emb\|LN827587.1\| Human herpesvirus 4 genome assembly sLCL-2.21, segment : I |
| sLCL-IS1.19 | gi\|764012345\|emb\|LN827588.1\| Human herpesvirus 4 genome assembly sLCL-IS1.19, segment : I |
| sLCL-IS2.01 | gi\|764012428\|emb\|LN827589.1\| Human herpesvirus 4 genome assembly sLCL-IS2.01, segment : I |
| sLCL-IM1.05 | gi\|764012512\|emb\|LN827590.1\| Human herpesvirus 4 genome assembly sLCL-IM1.05, segment : I |
| sLCL-2.15 | gi\|764012596\|emb\|LN827591.1\| Human herpesvirus 4 genome assembly sLCL-2.15, segment : I |
| sLCL-IS1.10 | gi\|764012680\|emb\|LN827592.1\| Human herpesvirus 4 genome assembly sLCL-IS1.10, segment : I |
| sLCL-IS1.12 | gi\|764012764\|emb\|LN827593.1\| Human herpesvirus 4 genome assembly sLCL-IS1.12, segment : I |
| sLCL-IS1.07 | gi\|764012847\|emb\|LN827594.1\| Human herpesvirus 4 genome assembly sLCL-IS1.07, segment : I |
| sLCL-IS1.03 | gi\|764012930\|emb\|LN827595.1\| Human herpesvirus 4 genome assembly sLCL-IS1.03, segment : I |
| sLCL-IM1.02 | gi\|764013014\|emb\|LN827596.1\| Human herpesvirus 4 genome assembly sLCL-IM1.02, segment : I |
| sLCL-IS1.04 | gi\|764013098\|emb\|LN827597.1\| Human herpesvirus 4 genome assembly sLCL-IS1.04, segment : I |
| LCL_B958_delEber2 | gi\|765364814\|emb\|LN827739.1\| Human herpesvirus 4 genome assembly LCL_B958_delEber2, segment : I |
| sLCL-IM1.16 | gi\|766540480\|emb\|LN827799.1\| Human herpesvirus 4 genome assembly sLCL-IM1.16, segment : I |
| Jijoye | gi\|766540564\|emb\|LN827800.1\| Human herpesvirus 4 genome assembly Jijoye, segment : I |
| CCH | gi\|787749988\|gb\|KP968257.1\| Human herpesvirus 4 strain CCH, partial genome |
| MP | gi\|787750086\|gb\|KP968258.1\| Human herpesvirus 4 strain MP, partial genome |
| SCL | gi\|787750199\|gb\|KP968259.1\| Human herpesvirus 4 strain SCL, partial genome |
| VGO | gi\|787750302\|gb\|KP968260.1\| Human herpesvirus 4 strain VGO, partial genome |
| HU11393 | gi\|787750415\|gb\|KP968261.1\| Human herpesvirus 4 strain HU11393, partial genome |
| H018436D | gi\|787750520\|gb\|KP968262.1\| Human herpesvirus 4 strain H018436D, partial genome |
| H058015C | gi\|787750634\|gb\|KP968263.1\| Human herpesvirus 4 strain H058015C, partial genome |
| H002213 | gi\|787750783\|gb\|KP968264.1\| Human herpesvirus 4 strain H002213, partial genome |
| P3HR1_c16Assembly | gi\|802130350\|emb\|LN827548.2\| Human herpesvirus 4 genome assembly P3HR1_c16 Assembly, segment : I |
| BL36 | gi\|802130370\|emb\|LN827557.2\| Human herpesvirus 4 genome assembly BL36, segment : I |
| sLCL-1.18 | gi\|802130391\|emb\|LN827563.2\| Human herpesvirus 4 genome assembly sLCL-1.18, segment : I |
| sLCL-2.22 | gi\|809279182\|emb\|LN831023.1\| Human herpesvirus 4 genome assembly sLCL-2.22, segment : I |
| GC1 | gi\|810056545\|gb\|KP735248.1\| Human herpesvirus 4 strain GC1, complete genome |

| | |
|---|---|
| Type1_Ref | gi\|82503188\|ref\|NC_007605.1\| Human herpesvirus 4 complete wild type genome |
| GD1 | gi\|84519641\|gb\|AY961628.3\| Human herpesvirus 4 strain GD1, complete genome |
| B95-8 | gi\|94734074\|emb\|V01555.2\| Epstein-Barr virus (EBV) genome, strain B95-8 |
| H03753A | gi\|957760599\|gb\|KR063342.1\| Human herpesvirus 4 strain H03753A, partial genome |
| CV-ARG | gi\|957760676\|gb\|KR063343.1\| Human herpesvirus 4 strain CV-ARG, partial genome |
| RPF | gi\|957760753\|gb\|KR063344.1\| Human herpesvirus 4 strain RPF, partial genome |
| FNR | gi\|957760829\|gb\|KR063345.1\| Human herpesvirus 4 strain FNR, partial genome |
| VA | gi\|957760989\|gb\|KT001102.1\| Human herpesvirus 4 strain VA, partial genome |
| SG | gi\|957761062\|gb\|KT001103.1\| Human herpesvirus 4 strain SG, partial genome |
| EBVaGC3 | gi\|961527962\|gb\|KT254013.1\| Human herpesvirus 4 isolate EBVaGC3, partial genome |
| EBVaGC1 | gi\|971835372\|gb\|KT273942.1\| Human herpesvirus 4 isolate EBVaGC1, partial genome |
| EBVaGC2 | gi\|971835443\|gb\|KT273943.1\| Human herpesvirus 4 isolate EBVaGC2, partial genome |
| EBVaGC5 | gi\|971835583\|gb\|KT273945.1\| Human herpesvirus 4 isolate EBVaGC5, partial genome |
| EBVaGC6 | gi\|971835652\|gb\|KT273946.1\| Human herpesvirus 4 isolate EBVaGC6, partial genome |
| EBVaGC7 | gi\|971835723\|gb\|KT273947.1\| Human herpesvirus 4 isolate EBVaGC7, partial genome |
| EBVaGC8 | gi\|971835791\|gb\|KT273948.1\| Human herpesvirus 4 isolate EBVaGC8, partial genome |
| EBVaGC9 | gi\|971835862\|gb\|KT273949.1\| Human herpesvirus 4 isolate EBVaGC9, partial genome |
| SNU-719 | gi\|995891043\|dbj\|AP015015.1\| Human herpesvirus 4 DNA, complete genome, strain: SNU-719 |
| YCCEL1 | gi\|995891127\|dbj\|AP015016.1\| Human herpesvirus 4 DNA, complete genome, strain: YCCEL1 |

**Supplemental Table 5. Full list of single nucleotide variant association results with significance cut off P > 0.01.**

**EBV genomic variant positions significantly associated with eBL cases in comparison to healthy control individuals. Test With EBV type stratification:**

| chrom | position | REF | ALT | MINA | MINU | OBSA | OBSU | P | OR | VarAAposition | VarType | RefCodon | VarCodon | RefAA | VarAA | Gene |
|---|---|---|---|---|---|---|---|---|---|---|---|---|---|---|---|---|
| NC_007605 | 37668 | T | C | 2 | 24 | 36 | 54 | 0.000328 | 0.1 | 485 | nonsynon | TCC | CCC | S | P | BYRF1 |
| NC_007605 | 95773 | A | T | 11 | 3 | 36 | 57 | 0.001322 | 6.67213 | 38 | nonsynon | AAC | TAC | N | Y | BKRF1 |
| NC_007605 | 95778 | T | G | 12 | 3 | 37 | 57 | 0.000538999 | 7.16129 | 39 | nonsynon | CAT | CAG | H | Q | BKRF1 |
| NC_007605 | 124703 | T | G | 7 | 1 | 34 | 56 | 0.003178 | 12.7377 | 159 | nonsynon | AAG | ACG | K | T | BcLF1 |
| NC_007605 | 124709 | G | A | 7 | 1 | 34 | 56 | 0.003092 | 12.7377 | 157 | nonsynon | GCT | GTT | A | V | BcLF1 |
| NC_007605 | 165131 | T | C | 10 | 36 | 36 | 57 | 0.004082 | 0.349462 | 29 | nonsynon | GTC | GCC | V | A | BARF1 |

**EBV genomic variant positions significantly associated with eBL cases in comparison to healthy control individuals. Test Without EBV type stratification:**

| chrom | position | REF | ALT | MINA | MINU | OBSA | OBSU | P | OR | VarAAposition | VarType | RefCodon | VarCodon | RefAA | VarAA | Gene |
|---|---|---|---|---|---|---|---|---|---|---|---|---|---|---|---|---|
| NC_007605 | 1868 | C | T | 2 | 17 | 35 | 56 | 0.00516199 | 0.16436 | 45 | nonsynon | CCA | TCA | P | S | BNRF1 |
| NC_007605 | 36226 | T | A | 0 | 10 | 34 | 55 | 0.00878399 | 0.0698644 | 4 | nonsynon | TTC | TAC | F | Y | BYRF1 |
| NC_007605 | 36627 | A | C | 18 | 12 | 37 | 53 | 0.00956499 | 2.51786 | 138 | nonsynon | AAT | CAT | N | H | BYRF1 |
| NC_007605 | 36629 | T | A | 18 | 12 | 37 | 53 | 0.00950499 | 2.51786 | 138 | nonsynon | AAT | AAA | N | K | BYRF1 |
| NC_007605 | 36633 | A | C | 18 | 12 | 37 | 53 | 0.00945799 | 2.51786 | 140 | nonsynon | ATT | CTT | I | L | BYRF1 |
| NC_007605 | 36636 | G | C | 18 | 12 | 37 | 53 | 0.00955999 | 2.51786 | 141 | nonsynon | GTA | CTA | V | L | BYRF1 |
| NC_007605 | 36642 | C | A | 18 | 12 | 37 | 53 | 0.00948099 | 2.51786 | 143 | nonsynon | CAA | AAA | Q | K | BYRF1 |
| NC_007605 | 36644 | A | C | 18 | 12 | 37 | 53 | 0.00951899 | 2.51786 | 143 | nonsynon | CAA | CAC | Q | H | BYRF1 |
| NC_007605 | 36645 | T | G | 18 | 12 | 37 | 53 | 0.00955999 | 2.51786 | 144 | nonsynon | TCA | GCA | S | A | BYRF1 |
| NC_007605 | 36669 | T | A | 18 | 12 | 37 | 53 | 0.00953299 | 2.51786 | 152 | nonsynon | TTG | ATG | L | M | BYRF1 |
| NC_007605 | 36670 | T | G | 18 | 12 | 37 | 53 | 0.00952199 | 2.51786 | 152 | nonsynon | TTG | TGG | L | W | BYRF1 |
| NC_007605 | 36880 | C | A | 18 | 12 | 36 | 53 | 0.00876299 | 2.61111 | 222 | nonsynon | ACT | AAT | T | N | BYRF1 |
| NC_007605 | 36883 | T | A | 18 | 13 | 36 | 54 | 0.00979399 | 2.4359 | 223 | nonsynon | CTG | CAG | L | Q | BYRF1 |
| NC_007605 | 36888 | C | G | 18 | 13 | 36 | 54 | 0.00977199 | 2.4359 | 225 | nonsynon | CCC | GCC | P | A | BYRF1 |
| NC_007605 | 36894 | C | A | 18 | 13 | 36 | 54 | 0.00984699 | 2.4359 | 227 | nonsynon | CCA | ACA | P | T | BYRF1 |
| NC_007605 | 36898 | T | C | 18 | 13 | 36 | 54 | 0.00982999 | 2.4359 | 228 | nonsynon | CTA | CCA | L | P | BYRF1 |
| NC_007605 | 36901 | T | C | 18 | 13 | 36 | 54 | 0.00984299 | 2.4359 | 229 | nonsynon | CTC | CCC | L | P | BYRF1 |

| | | | | | | | | | | | | | | | | |
|---|---|---|---|---|---|---|---|---|---|---|---|---|---|---|---|---|
| NC_007605 | 36907 | T | C | 18 | 13 | 36 | 54 | 0.00975699 | 2.4359 | 231 | nonsynon | GTA | GCA | V | A | BYRF1 |
| NC_007605 | 36910 | T | C | 18 | 13 | 36 | 54 | 0.00980199 | 2.4359 | 232 | nonsynon | CTA | CCA | L | P | BYRF1 |
| NC_007605 | 37074 | C | T | 17 | 12 | 35 | 54 | 0.00825199 | 2.56604 | 287 | nonsynon | CCT | TCT | P | S | BYRF1 |
| NC_007605 | 37086 | C | T | 17 | 12 | 35 | 54 | 0.00820199 | 2.56604 | 291 | nonsynon | CCC | TCC | P | S | BYRF1 |
| NC_007605 | 37124 | G | C | 17 | 12 | 35 | 54 | 0.00829799 | 2.56604 | 303 | nonsynon | CAG | CAC | Q | H | BYRF1 |
| NC_007605 | 37206 | C | A | 17 | 13 | 34 | 55 | 0.00883099 | 2.48718 | 331 | nonsynon | CCG | ACG | P | T | BYRF1 |
| NC_007605 | 37210 | A | C | 17 | 13 | 34 | 55 | 0.00882999 | 2.48718 | 332 | nonsynon | CAA | CCA | Q | P | BYRF1 |
| NC_007605 | 37212 | C | A | 17 | 13 | 34 | 55 | 0.00899699 | 2.48718 | 333 | nonsynon | CCC | ACC | P | T | BYRF1 |
| NC_007605 | 37245 | A | G | 17 | 12 | 34 | 53 | 0.00769299 | 2.61111 | 344 | nonsynon | AGC | GGC | S | G | BYRF1 |
| NC_007605 | 37249 | G | A | 17 | 12 | 34 | 53 | 0.00769599 | 2.61111 | 345 | nonsynon | AGG | AAG | R | K | BYRF1 |
| NC_007605 | 37462 | A | C | 18 | 12 | 37 | 53 | 0.00932399 | 2.51786 | 416 | nonsynon | CAC | CCC | H | P | BYRF1 |
| NC_007605 | 37479 | G | A | 18 | 12 | 37 | 54 | 0.00935899 | 2.57143 | 422 | nonsynon | GTT | ATT | V | I | BYRF1 |
| NC_007605 | 37515 | C | G | 18 | 12 | 37 | 54 | 0.00935399 | 2.57143 | 434 | nonsynon | CCA | GCA | P | A | BYRF1 |
| NC_007605 | 37516 | C | A | 18 | 12 | 37 | 54 | 0.00928499 | 2.57143 | 434 | nonsynon | CCA | CAA | P | Q | BYRF1 |
| NC_007605 | 37521 | G | C | 18 | 12 | 37 | 54 | 0.00942099 | 2.57143 | 436 | nonsynon | GCT | CCT | A | P | BYRF1 |
| NC_007605 | 37560 | A | T | 18 | 12 | 37 | 53 | 0.00942999 | 2.51786 | 449 | nonsynon | ATA | TTA | I | L | BYRF1 |
| NC_007605 | 37668 | T | C | 2 | 24 | 36 | 54 | 6.30E-05 | 0.1 | 485 | nonsynon | TCC | CCC | S | P | BYRF1 |
| NC_007605 | 57209 | C | G | 5 | 23 | 38 | 57 | 0.004096 | 0.278628 | 677 | nonsynon | GAG | GAC | E | D | BPLF1 |
| NC_007605 | 57333 | G | A | 5 | 23 | 37 | 57 | 0.004228 | 0.286704 | 636 | nonsynon | CCA | CTA | P | L | BPLF1 |
| NC_007605 | 59221 | C | T | 4 | 0 | 36 | 58 | 0.00979199 | 15.3066 | 7 | nonsynon | GGG | AGG | G | R | BPLF1 |
| NC_007605 | 59230 | C | T | 4 | 21 | 37 | 58 | 0.00581599 | 0.258503 | 4 | nonsynon | GGC | AGC | G | S | BPLF1 |
| NC_007605 | 64132 | G | A | 10 | 33 | 34 | 57 | 0.00713199 | 0.423197 | 5 | nonsynon | AGT | AAT | S | N | BORF2 |
| NC_007605 | 72849 | G | T | 3 | 0 | 16 | 53 | 0.00531599 | 25.2712 | 582 | nonsynon | CCT | CAT | P | H | BSLF1 |
| NC_007605 | 73309 | C | T | 3 | 0 | 18 | 55 | 0.00638299 | 23.0896 | 429 | nonsynon | GCA | ACA | A | T | BSLF1 |
| NC_007605 | 85291 | C | T | 8 | 14 | 12 | 54 | 0.00807599 | 3.35714 | 717 | nonsynon | CCT | TCT | P | S | EBNA3B |
| NC_007605 | 85877 | C | A | 0 | 13 | 32 | 57 | 0.002436 | 0.0582831 | 912 | nonsynon | GCT | GAT | A | D | EBNA3B |
| NC_007605 | 86101 | C | G | 18 | 16 | 30 | 55 | 0.00623499 | 2.51786 | 7 | nonsynon | CAG | GAG | Q | E | EBNA3C |
| NC_007605 | 86114 | G | T | 18 | 16 | 29 | 55 | 0.003552 | 2.64375 | 11 | nonsynon | AGA | ATA | R | I | EBNA3C |
| NC_007605 | 86132 | A | C | 18 | 16 | 30 | 55 | 0.00619199 | 2.51786 | 17 | nonsynon | GAG | GCG | E | A | EBNA3C |
| NC_007605 | 86153 | C | A | 18 | 15 | 29 | 54 | 0.003453 | 2.79 | 24 | nonsynon | ACT | AAT | T | N | EBNA3C |

| | | | | | | | | | | | | | | | | |
|---|---|---|---|---|---|---|---|---|---|---|---|---|---|---|---|---|
| NC_007605 | 86167 | G | A | 18 | 15 | 30 | 54 | 0.003837 | 2.65714 | 29 | nonsynon | GAT | AAT | D | N | EBNA3C |
| NC_007605 | 86168 | A | T | 18 | 15 | 30 | 54 | 0.003826 | 2.65714 | 29 | nonsynon | GAT | GTT | D | V | EBNA3C |
| NC_007605 | 86191 | C | T | 18 | 15 | 30 | 54 | 0.003793 | 2.65714 | 37 | nonsynon | CCA | TCA | P | S | EBNA3C |
| NC_007605 | 86194 | T | A | 18 | 15 | 30 | 54 | 0.003827 | 2.65714 | 38 | nonsynon | TCC | ACC | S | T | EBNA3C |
| NC_007605 | 86197 | A | G | 18 | 15 | 30 | 54 | 0.003815 | 2.65714 | 39 | nonsynon | AGT | GGT | S | G | EBNA3C |
| NC_007605 | 86222 | C | A | 18 | 15 | 29 | 54 | 0.00341 | 2.79 | 47 | nonsynon | CCA | CAA | P | Q | EBNA3C |
| NC_007605 | 86224 | G | A | 18 | 15 | 29 | 54 | 0.003435 | 2.79 | 48 | nonsynon | GAA | AAA | E | K | EBNA3C |
| NC_007605 | 86226 | A | C | 18 | 15 | 29 | 54 | 0.003494 | 2.79 | 48 | nonsynon | GAA | GAC | E | D | EBNA3C |
| NC_007605 | 86230 | T | C | 18 | 15 | 29 | 54 | 0.003445 | 2.79 | 50 | nonsynon | TCA | CCA | S | P | EBNA3C |
| NC_007605 | 86251 | C | T | 18 | 15 | 29 | 54 | 0.003432 | 2.79 | 57 | nonsynon | CCT | TCT | P | S | EBNA3C |
| NC_007605 | 86254 | T | C | 18 | 15 | 29 | 54 | 0.003442 | 2.79 | 58 | nonsynon | TGG | CGG | W | R | EBNA3C |
| NC_007605 | 86265 | C | A | 18 | 16 | 29 | 55 | 0.003616 | 2.64375 | 61 | nonsynon | AGC | AGA | S | R | EBNA3C |
| NC_007605 | 86689 | A | G | 18 | 16 | 30 | 57 | 0.003895 | 2.625 | 178 | nonsynon | CAA | CGA | Q | R | EBNA3C |
| NC_007605 | 86697 | C | T | 17 | 16 | 29 | 57 | 0.00698299 | 2.53963 | 181 | nonsynon | CCC | TCC | P | S | EBNA3C |
| NC_007605 | 86839 | C | T | 17 | 17 | 29 | 57 | 0.00815199 | 2.36585 | 228 | nonsynon | ACA | ATA | T | I | EBNA3C |
| NC_007605 | 86848 | A | C | 18 | 17 | 30 | 57 | 0.00832999 | 2.44538 | 231 | nonsynon | CAT | CCT | H | P | EBNA3C |
| NC_007605 | 86938 | A | T | 18 | 17 | 31 | 57 | 0.00909499 | 2.33422 | 261 | nonsynon | TAT | TTT | Y | F | EBNA3C |
| NC_007605 | 86973 | A | G | 19 | 16 | 31 | 57 | 0.002155 | 2.7064 | 273 | nonsynon | ATC | GTC | I | V | EBNA3C |
| NC_007605 | 87052 | A | G | 18 | 17 | 30 | 56 | 0.00823799 | 2.39496 | 299 | nonsynon | AAT | AGT | N | S | EBNA3C |
| NC_007605 | 87054 | C | T | 18 | 17 | 30 | 56 | 0.00826899 | 2.39496 | 300 | nonsynon | CCA | TCA | P | S | EBNA3C |
| NC_007605 | 87179 | C | A | 18 | 17 | 31 | 56 | 0.00963899 | 2.2861 | 341 | nonsynon | AAC | AAA | N | K | EBNA3C |
| NC_007605 | 87187 | G | T | 18 | 17 | 31 | 56 | 0.00962999 | 2.2861 | 344 | nonsynon | CGA | CTA | R | L | EBNA3C |
| NC_007605 | 87522 | G | A | 17 | 15 | 30 | 54 | 0.00775599 | 2.45116 | 456 | nonsynon | GAA | AAA | E | K | EBNA3C |
| NC_007605 | 87528 | T | C | 17 | 15 | 29 | 55 | 0.00677399 | 2.62602 | 458 | nonsynon | TCA | CCA | S | P | EBNA3C |
| NC_007605 | 87535 | T | A | 17 | 15 | 29 | 55 | 0.00673599 | 2.62602 | 460 | nonsynon | GTG | GAG | V | E | EBNA3C |
| NC_007605 | 87537 | A | C | 17 | 15 | 29 | 55 | 0.00676999 | 2.62602 | 461 | nonsynon | ACC | CCC | T | P | EBNA3C |
| NC_007605 | 87555 | C | A | 18 | 15 | 29 | 55 | 0.001715 | 2.85 | 467 | nonsynon | CCG | ACG | P | T | EBNA3C |
| NC_007605 | 87565 | T | C | 18 | 16 | 30 | 55 | 0.00635399 | 2.51786 | 470 | nonsynon | GTG | GCG | V | A | EBNA3C |
| NC_007605 | 87568 | A | G | 18 | 15 | 30 | 55 | 0.003646 | 2.71429 | 471 | nonsynon | GAG | GGG | E | G | EBNA3C |
| NC_007605 | 87571 | T | G | 18 | 15 | 30 | 55 | 0.003663 | 2.71429 | 472 | nonsynon | ATG | AGG | M | R | EBNA3C |

| | | | | | | | | | | | | | | | | |
|---|---|---|---|---|---|---|---|---|---|---|---|---|---|---|---|---|
| NC_007605 | 87573 | C | T | 18 | 15 | 29 | 55 | 0.001682 | 2.85 | 473 | nonsynon | CCA | TCA | P | S | EBNA3C |
| NC_007605 | 87577 | T | C | 17 | 15 | 28 | 55 | 0.003011 | 2.76068 | 474 | nonsynon | ATG | ACG | M | T | EBNA3C |
| NC_007605 | 87591 | C | G | 16 | 15 | 28 | 55 | 0.00658399 | 2.53333 | 479 | nonsynon | CAA | GAA | Q | E | EBNA3C |
| NC_007605 | 87609 | A | G | 16 | 15 | 28 | 55 | 0.00660899 | 2.53333 | 485 | nonsynon | AAA | GAA | K | E | EBNA3C |
| NC_007605 | 87612 | C | G | 16 | 15 | 28 | 55 | 0.00655499 | 2.53333 | 486 | nonsynon | CCG | GCG | P | A | EBNA3C |
| NC_007605 | 87618 | C | T | 16 | 15 | 28 | 55 | 0.00656199 | 2.53333 | 488 | nonsynon | CCA | TCA | P | S | EBNA3C |
| NC_007605 | 87621 | G | T | 16 | 15 | 28 | 55 | 0.00655099 | 2.53333 | 489 | nonsynon | GTC | TTC | V | F | EBNA3C |
| NC_007605 | 87630 | A | C | 16 | 15 | 28 | 55 | 0.00660599 | 2.53333 | 492 | nonsynon | ACG | CCG | T | P | EBNA3C |
| NC_007605 | 87684 | G | A | 16 | 15 | 27 | 55 | 0.00532199 | 2.66667 | 510 | nonsynon | GTC | ATC | V | I | EBNA3C |
| NC_007605 | 87719 | T | G | 16 | 15 | 27 | 55 | 0.00524299 | 2.66667 | 521 | nonsynon | GAT | GAG | D | E | EBNA3C |
| NC_007605 | 87720 | T | G | 16 | 15 | 27 | 55 | 0.00537199 | 2.66667 | 522 | nonsynon | TCA | GCA | S | A | EBNA3C |
| NC_007605 | 87721 | C | A | 16 | 15 | 27 | 55 | 0.00534799 | 2.66667 | 522 | nonsynon | TCA | TAA | S | Z | EBNA3C |
| NC_007605 | 87723 | T | C | 16 | 15 | 27 | 55 | 0.00531499 | 2.66667 | 523 | nonsynon | TCG | CCG | S | P | EBNA3C |
| NC_007605 | 87732 | T | A | 16 | 15 | 27 | 55 | 0.00535199 | 2.66667 | 526 | nonsynon | TCA | ACA | S | T | EBNA3C |
| NC_007605 | 87742 | A | G | 16 | 15 | 27 | 55 | 0.00533999 | 2.66667 | 529 | nonsynon | AAT | AGT | N | S | EBNA3C |
| NC_007605 | 87760 | A | C | 16 | 15 | 27 | 55 | 0.00533499 | 2.66667 | 535 | nonsynon | CAT | CCT | H | P | EBNA3C |
| NC_007605 | 87804 | G | A | 16 | 15 | 27 | 54 | 0.00544099 | 2.61053 | 550 | nonsynon | GCG | ACG | A | T | EBNA3C |
| NC_007605 | 87805 | C | T | 16 | 15 | 27 | 54 | 0.00546799 | 2.61053 | 550 | nonsynon | GCG | GTG | A | V | EBNA3C |
| NC_007605 | 87813 | A | C | 16 | 15 | 27 | 54 | 0.00554399 | 2.61053 | 553 | nonsynon | ACC | CCC | T | P | EBNA3C |
| NC_007605 | 87831 | A | G | 16 | 15 | 27 | 54 | 0.00547099 | 2.61053 | 559 | nonsynon | ACT | GCT | T | A | EBNA3C |
| NC_007605 | 88162 | T | A | 16 | 17 | 27 | 58 | 0.00675599 | 2.45201 | 669 | nonsynon | CTG | CAG | L | Q | EBNA3C |
| NC_007605 | 88167 | T | C | 16 | 17 | 27 | 58 | 0.00679799 | 2.45201 | 671 | nonsynon | TCC | CCC | S | P | EBNA3C |
| NC_007605 | 88458 | C | G | 6 | 31 | 28 | 56 | 0.003648 | 0.313548 | 768 | nonsynon | CCC | GCC | P | A | EBNA3C |
| NC_007605 | 88501 | A | C | 16 | 16 | 27 | 56 | 0.00610599 | 2.52632 | 782 | nonsynon | CAC | CCC | H | P | EBNA3C |
| NC_007605 | 88554 | T | C | 16 | 17 | 26 | 58 | 0.00561299 | 2.58824 | 800 | nonsynon | TCT | CCT | S | P | EBNA3C |
| NC_007605 | 88566 | T | A | 16 | 17 | 27 | 58 | 0.00676599 | 2.45201 | 804 | nonsynon | TGT | AGT | C | S | EBNA3C |
| NC_007605 | 88632 | G | A | 18 | 17 | 30 | 58 | 0.00455 | 2.4958 | 826 | nonsynon | GAT | AAT | D | N | EBNA3C |
| NC_007605 | 88634 | T | A | 18 | 17 | 30 | 58 | 0.004556 | 2.4958 | 826 | nonsynon | GAT | GAA | D | E | EBNA3C |
| NC_007605 | 88638 | A | T | 18 | 17 | 30 | 58 | 0.00452 | 2.4958 | 828 | nonsynon | AGG | TGG | R | W | EBNA3C |
| NC_007605 | 88641 | G | T | 18 | 17 | 30 | 58 | 0.004551 | 2.4958 | 829 | nonsynon | GCA | TCA | A | S | EBNA3C |

| | | | | | | | | | | | | | | | | |
|---|---|---|---|---|---|---|---|---|---|---|---|---|---|---|---|---|
| NC_007605 | 88644 | C | G | 18 | 16 | 30 | 58 | 0.003963 | 2.67857 | 830 | nonsynon | CCA | GCA | P | A | EBNA3C |
| NC_007605 | 88718 | A | T | 18 | 16 | 30 | 57 | 0.003835 | 2.625 | 854 | nonsynon | CAA | CAT | Q | H | EBNA3C |
| NC_007605 | 89555 | G | T | 24 | 19 | 39 | 57 | 0.00537299 | 2.22222 | 92 | nonsynon | CAA | AAA | Q | K | BZLF2 |
| NC_007605 | 89603 | C | T | 23 | 19 | 38 | 56 | 0.00907499 | 2.12413 | 76 | nonsynon | GTA | ATA | V | I | BZLF2 |
| NC_007605 | 90216 | T | A | 22 | 17 | 39 | 57 | 0.00883799 | 2.2416 | 176 | nonsynon | GAA | GAT | E | D | BZLF1 |
| NC_007605 | 90380 | T | A | 25 | 21 | 38 | 58 | 0.004861 | 2.21755 | 163 | nonsynon | CAA | CTA | Q | L | BZLF1 |
| NC_007605 | 90413 | A | G | 25 | 21 | 38 | 58 | 0.004878 | 2.21755 | 152 | nonsynon | GTG | GCG | V | A | BZLF1 |
| NC_007605 | 90431 | A | G | 25 | 21 | 38 | 58 | 0.004837 | 2.21755 | 146 | nonsynon | GTT | GCT | V | A | BZLF1 |
| NC_007605 | 90642 | A | G | 26 | 21 | 39 | 58 | 0.002836 | 2.2619 | 76 | nonsynon | TCT | CCT | S | P | BZLF1 |
| NC_007605 | 90666 | T | C | 26 | 21 | 39 | 58 | 0.002844 | 2.2619 | 68 | nonsynon | ACC | GCC | T | A | BZLF1 |
| NC_007605 | 92591 | G | A | 3 | 0 | 22 | 57 | 0.00976799 | 19.3133 | 102 | nonsynon | GCG | GTG | A | V | BRLF1 |
| NC_007605 | 95773 | A | T | 11 | 3 | 36 | 57 | 0.000754999 | 6.67213 | 38 | nonsynon | AAC | TAC | N | Y | BKRF1 |
| NC_007605 | 95778 | T | G | 12 | 3 | 37 | 57 | 0.000331 | 7.16129 | 39 | nonsynon | CAT | CAG | H | Q | BKRF1 |
| NC_007605 | 100861 | C | T | 3 | 0 | 15 | 55 | 0.004079 | 28.1273 | 371 | nonsynon | GGG | AGG | G | R | BBLF4 |
| NC_007605 | 100863 | T | C | 3 | 0 | 15 | 55 | 0.004091 | 28.1273 | 370 | nonsynon | CAG | CGG | Q | R | BBLF4 |
| NC_007605 | 100864 | G | A | 3 | 0 | 15 | 55 | 0.004087 | 28.1273 | 370 | nonsynon | CAG | TAG | Q | Z | BBLF4 |
| NC_007605 | 100870 | C | T | 3 | 0 | 15 | 55 | 0.004037 | 28.1273 | 368 | nonsynon | GCC | ACC | A | T | BBLF4 |
| NC_007605 | 100872 | G | A | 3 | 0 | 15 | 55 | 0.004072 | 28.1273 | 367 | nonsynon | ACT | ATT | T | I | BBLF4 |
| NC_007605 | 101364 | T | C | 5 | 0 | 20 | 55 | 0.000414 | 34.2394 | 203 | nonsynon | GAA | GGA | E | G | BBLF4 |
| NC_007605 | 101371 | C | G | 5 | 0 | 20 | 55 | 0.000448 | 34.2394 | 201 | nonsynon | GTG | CTG | V | L | BBLF4 |
| NC_007605 | 124703 | T | G | 7 | 1 | 34 | 56 | 0.002196 | 12.7377 | 159 | nonsynon | AAG | ACG | K | T | BcLF1 |
| NC_007605 | 124709 | G | A | 7 | 1 | 34 | 56 | 0.002193 | 12.7377 | 157 | nonsynon | GCT | GTT | A | V | BcLF1 |
| NC_007605 | 158849 | C | A | 4 | 0 | 34 | 58 | 0.00844099 | 16.2558 | 6 | nonsynon | GTG | TTG | V | L | BALF4 |
| NC_007605 | 165131 | T | C | 10 | 36 | 36 | 57 | 0.000943999 | 0.349462 | 29 | nonsynon | GTC | GCC | V | A | BARF1 |
| NC_007605 | 70812 | G | C | 4 | 0 | 30 | 56 | 0.00643099 | 17.9204 | 361 | nonsynon | GAC | GAG | D | E | BSLF2_BMLF1 |

**Supplemental Table 6. Nucleotide variant compositions of each isolate in their BZLF1 gene upstream promoter region. Nucleotide positions are in upstream relative to TSS of BZLF1 gene. -532 and -524 are variant position within our isolates while these two are not in both promoter types.**

| | | | -532 | -524 | | -525 | -460 | -365 | -274 | -141 | -106 | -100 | Zp promoter type |
|---|---|---|---|---|---|---|---|---|---|---|---|---|---|
| NC_007605 | Type1 Ref | | G | A | | C | A | G | C | A | A | T | Zp-P |
| M81 strain | | | G | A | | A | G | A | A | G | G | G | Zp-V3 |
| eBL-Plasma-0049 | Inter-typic | eBL | A | G | | C | G | A | C | G | G | G | Zp-V3 |
| eBL-Tumor-0033 | Inter-typic | eBL | A | G | | C | G | A | C | A | A | T | Zp-P Chimeric with -141A |
| eBL-Tumor-0012 | Inter-typic | eBL | A | G | | C | G | A | C | G | G | G | Zp-V3 |
| eBL-Tumor-0001 | Type1 | eBL | G | A | | C | A | G | C | A | A | T | Zp-P |
| eBL-Tumor-0018 | Type1 | eBL | A | G | | C | G | A | C | G | G | G | Zp-V3 |
| eBL-Tumor-0002 | Type1 | eBL | G | A | | C | A | G | C | A | A | T | Zp-P |
| eBL-Plasma-0042 | Type1 | eBL | G | A | | C | A | G | C | A | A | T | Zp-P |
| eBL-Plasma-0044 | Type1 | eBL | G | A | | C | A | G | C | A | G | G | Zp-P Chimeric with -141A |
| eBL-Plasma-0045 | Type1 | eBL | G | A | | C | A | G | C | A | A | T | Zp-P |
| eBL-Plasma-0046 | Type1 | eBL | N | N | | N | N | N | N | N | N | N | NA |
| eBL-Plasma-0047 | Type1 | eBL | G | A | | C | A | G | C | A | A | T | Zp-P |
| eBL-Plasma-0050 | Type1 | eBL | G | A | | C | A | G | C | A | A | T | Zp-P |
| eBL-Plasma-0051 | Type1 | eBL | G | A | | C | A | G | C | A | A | T | Zp-P |
| eBL-Plasma-0052 | Type1 | eBL | G | A | | C | A | G | C | A | A | T | Zp-P |
| eBL-Plasma-0053 | Type1 | eBL | G | A | | C | A | G | C | A | A | T | Zp-P |
| eBL-Plasma-0055 | Type1 | eBL | A | G | | C | G | A | C | G | G | G | Zp-V3 |
| eBL-Tumor-0019 | Type1 | eBL | G | A | | C | A | G | C | A | A | T | Zp-P |
| eBL-Tumor-0004 | Type1 | eBL | G | A | | C | A | G | C | A | A | T | Zp-P |
| eBL-Tumor-0005 | Type1 | eBL | G | A | | C | A | G | C | A | A | T | Zp-P |
| eBL-Tumor-0021 | Type1 | eBL | G | A | | C | A | G | C | A | A | T | Zp-P |
| eBL-Tumor-0006 | Type1 | eBL | G | A | | C | A | G | C | A | A | T | Zp-P |
| eBL-Tumor-0007 | Type1 | eBL | G | A | | C | A | G | C | A | A | T | Zp-P |
| eBL-Tumor-0022 | Type1 | eBL | G | A | | C | A | G | C | A | A | T | Zp-P |
| eBL-Tumor-0023 | Type1 | eBL | G | A | | C | A | G | C | A | A | T | Zp-P |
| eBL-Tumor-0025 | Type1 | eBL | A | G | | C | G | A | C | G | G | G | Zp-V3 |
| eBL-Tumor-0027 | Type1 | eBL | G | A | | C | A | G | C | A | A | T | Zp-P |
| eBL-Tumor-0037 | Type1 | eBL | G | A | | C | A | G | C | A | A | T | Zp-P |
| eBL-Tumor-0029 | Type1 | eBL | G | A | | C | A | G | C | A | A | T | Zp-P |

**Supplemental Table 6. Nucleotide variant compositions of each isolate in their BZLF1 gene upstream promoter region. Nucleotide positions are in upstream relative to TSS of BZLF1 gene. -532 and -524 are variant position within our isolates while these two are not in both promoter types.**

| | | | -532 | -524 | | -525 | -460 | -365 | -274 | -141 | -106 | -100 | Zp promoter type |
|---|---|---|---|---|---|---|---|---|---|---|---|---|---|
| NC_007605 | Type1 Ref | | G | A | | C | A | G | C | A | A | T | Zp-P |
| M81 strain | | | G | A | | A | G | A | A | G | G | G | Zp-V3 |
| eBL-Tumor-0038 | Type1 | eBL | G | A | | C | A | G | C | A | A | T | Zp-P |
| eBL-Tumor-0039 | Type1 | eBL | G | A | | C | A | G | C | A | A | T | Zp-P |
| eBL-Tumor-0041 | Type1 | eBL | G | A | | C | A | G | C | A | A | T | Zp-P |
| eBL-Tumor-0030 | Type1 | eBL | G | A | | C | A | G | C | A | A | T | Zp-P |
| eBL-Tumor-0009 | Type1 | eBL | G | A | | C | A | G | C | A | A | T | Zp-P |
| eBL-Tumor-0031 | Type1 | eBL | G | A | | C | A | G | C | A | A | T | Zp-P |
| eBL-Tumor-0032 | Type1 | eBL | A | G | | C | G | A | C | A | A | T | Zp-P Chimeric with -141A |
| eBL-Tumor-0010 | Type1 | eBL | G | A | | C | A | G | C | A | A | T | Zp-P |
| eBL-Tumor-0011 | Type1 | eBL | A | G | | C | G | A | C | G | G | G | Zp-V3 |
| eBL-Tumor-0013 | Type1 | eBL | G | A | | C | A | G | C | A | A | T | Zp-P |
| eBL-Tumor-0034 | Type1 | eBL | G | A | | C | A | G | C | A | A | T | Zp-P |
| eBL-Tumor-0014 | Type1 | eBL | G | A | | C | A | G | C | A | A | T | Zp-P |
| eBL-Tumor-0015 | Type1 | eBL | A | G | | C | G | A | C | G | G | G | Zp-V3 |
| eBL-Tumor-0016 | Type1 | eBL | G | A | | C | A | G | C | A | A | T | Zp-P |
| eBL_CL-01 | Type1 | eBL | G | A | | C | A | G | C | A | A | T | Zp-P |
| eBL_CL-03 | Type1 | eBL | G | A | | C | A | G | C | A | A | T | Zp-P |
| HC-0030 | Type1 | HealthyControl | N | N | | N | N | N | N | N | N | N | NA |
| HC-0032 | Type1 | HealthyControl | N | N | | N | G | N | N | N | N | N | NA |
| HC-0035 | Type1 | HealthyControl | A | G | | C | N | N | C | G | G | G | Most likely Zp-V3 |
| HC-0036 | Type1 | HealthyControl | N | N | | N | N | N | N | N | N | N | NA |
| HC-0037 | Type1 | HealthyControl | G | A | | C | A | G | C | A | A | T | Zp-P |
| HC-0001 | Type1 | HealthyControl | N | N | | N | N | N | N | N | N | N | NA |
| HC-0003 | Type1 | HealthyControl | G | A | | C | A | G | C | A | A | T | Zp-P |
| HC-0007 | Type1 | HealthyControl | N | N | | N | N | G | N | A | A | T | Most likely Zp-P |
| HC-0008 | Type1 | HealthyControl | N | N | | N | N | N | C | N | N | N | NA |
| HC-0009 | Type1 | HealthyControl | G | A | | C | A | G | C | A | A | T | Zp-P |
| HC-0010 | Type1 | HealthyControl | N | N | | N | N | N | N | N | N | N | NA |
| HC-0013 | Type1 | HealthyControl | N | N | | N | N | N | N | N | N | N | NA |

**Supplemental Table 6. Nucleotide variant compositions of each isolate in their BZLF1 gene upstream promoter region. Nucleotide positions are in upstream relative to TSS of BZLF1 gene. -532 and -524 are variant position within our isolates while these two are not in both promoter types.**

| | | | -532 | -524 | | -525 | -460 | -365 | -274 | -141 | -106 | -100 | Zp promoter type |
|---|---|---|---|---|---|---|---|---|---|---|---|---|---|
| NC_007605 | Type1 Ref | | G | A | | C | A | G | C | A | A | T | Zp-P |
| M81 strain | | | G | A | | A | G | A | A | G | G | G | Zp-V3 |
| HC-0015 | Type1 | HealthyControl | A | G | | C | G | A | C | G | G | G | Zp-V3 |
| HC-0017 | Type1 | HealthyControl | N | N | | N | N | N | N | N | N | N | NA |
| HC-0020 | Type1 | HealthyControl | N | N | | N | N | N | N | N | N | N | NA |
| HC-0021 | Type1 | HealthyControl | N | N | | N | A | G | C | A | A | T | Most likely Zp-P |
| HC-0024 | Type1 | HealthyControl | N | N | | N | G | N | N | N | N | N | NA |
| HC-0025 | Type1 | HealthyControl | N | N | | N | A | N | C | A | A | T | Most likely Zp-P |
| HC-0029 | Type1 | HealthyControl | G | A | | C | A | G | C | A | A | T | Zp-P |
| eBL-Tumor-0017 | Type2 | eBL | A | G | | C | G | A | C | G | G | G | Zp-V3 |
| eBL-Plasma-0043 | Type2 | eBL | A | G | | C | G | A | C | G | G | G | Zp-V3 |
| eBL-Plasma-0048 | Type2 | eBL | N | N | | N | N | N | N | N | G | N | NA |
| eBL-Plasma-0054 | Type2 | eBL | A | G | | C | G | A | C | G | G | G | Zp-V3 |
| eBL-Tumor-0003 | Type2 | eBL | N | N | | N | N | N | N | N | N | N | NA |
| eBL-Tumor-0035 | Type2 | eBL | A | G | | C | G | A | C | G | G | G | Zp-V3 |
| eBL-Tumor-0020 | Type2 | eBL | A | G | | C | G | A | C | G | G | G | Zp-V3 |
| eBL-Tumor-0008 | Type2 | eBL | A | G | | C | G | A | C | G | G | G | Zp-V3 |
| eBL-Tumor-0036 | Type2 | eBL | A | G | | C | G | A | C | G | G | G | Zp-V3 |
| eBL-Tumor-0024 | Type2 | eBL | A | G | | C | G | A | C | G | G | G | Zp-V3 |
| eBL-Tumor-0026 | Type2 | eBL | A | G | | C | G | A | C | G | G | G | Zp-V3 |
| eBL-Tumor-0028 | Type2 | eBL | A | G | | C | G | A | C | G | G | G | Zp-V3 |
| eBL-Tumor-0040 | Type2 | eBL | A | G | | C | G | A | C | G | G | G | Zp-V3 |
| eBL_CL-02 | Type2 | eBL | A | G | | C | G | A | C | G | G | G | Zp-V3 |
| HC-0031 | Type2 | HealthyControl | N | N | | N | N | N | N | N | N | N | NA |
| HC-0033 | Type2 | HealthyControl | N | N | | N | N | N | N | N | N | N | NA |
| HC-0034 | Type2 | HealthyControl | N | N | | N | N | N | N | G | G | G | Most likely Zp-V3 |
| HC-0038 | Type2 | HealthyControl | N | N | | N | N | N | N | N | N | N | NA |
| HC-0002 | Type2 | HealthyControl | A | G | | C | G | A | C | G | G | G | Zp-V3 |
| HC-0004 | Type2 | HealthyControl | A | G | | C | G | A | C | G | G | G | Zp-V3 |
| HC-0005 | Type2 | HealthyControl | A | G | | C | G | A | C | G | G | G | Zp-V3 |

**Supplemental Table 6. Nucleotide variant compositions of each isolate in their BZLF1 gene upstream promoter region. Nucleotide positions are in upstream relative to TSS of BZLF1 gene. -532 and -524 are variant position within our isolates while these two are not in both promoter types.**

| | | | -532 | -524 | | -525 | -460 | -365 | -274 | -141 | -106 | -100 | Zp promoter type |
|---|---|---|---|---|---|---|---|---|---|---|---|---|---|
| NC_007605 | Type1 Ref | | G | A | | C | A | G | C | A | A | T | Zp-P |
| M81 strain | | | G | A | | A | G | A | A | G | G | G | Zp-V3 |
| HC-0006 | Type2 | HealthyControl | N | N | | N | N | N | N | N | N | N | NA |
| HC-0011 | Type2 | HealthyControl | N | N | | N | N | N | N | N | N | N | NA |
| HC-0012 | Type2 | HealthyControl | N | N | | N | N | N | N | N | N | N | NA |
| HC-0014 | Type2 | HealthyControl | N | N | | N | N | N | N | G | G | G | Most likely Zp-V3 |
| HC-0016 | Type2 | HealthyControl | A | G | | C | G | A | C | G | G | G | Zp-V3 |
| HC-0039 | Type2 | HealthyControl | N | N | | N | N | N | N | N | N | N | NA |
| HC-0018 | Type2 | HealthyControl | N | N | | N | N | N | N | N | N | N | NA |
| HC-0019 | Type2 | HealthyControl | A | G | | C | G | A | C | G | G | G | Zp-V3 |
| HC-0022 | Type2 | HealthyControl | A | G | | C | G | A | C | G | G | G | Zp-V3 |
| HC-0023 | Type2 | HealthyControl | A | G | | C | G | N | C | N | G | G | Most likely Zp-V3 |
| HC-0040 | Type2 | HealthyControl | A | G | | C | G | A | C | G | G | G | Zp-V3 |
| HC-0026 | Type2 | HealthyControl | A | G | | C | G | A | C | G | G | G | Zp-V3 |
| HC-0027 | Type2 | HealthyControl | A | G | | C | G | A | C | G | G | G | Zp-V3 |
| HC-0028 | Type2 | HealthyControl | A | G | | C | G | A | C | G | G | G | Zp-V3 |

**Supplemental Table 7. RNA bait sequences used in EBV specific hybrid capture enrichment reactions**

| Bait Name | ID | Bait Sequence | Ratio in sythesis |
|---|---|---|---|
| wt-0-120 | 1 | AGAATTCGTCTTGCTCTATTCACCCTTACTTTTCTTCTTGCCCGTTCTCTTTCTTAGTATGAATCCAGTATGCCTGCCTGTAATTGTTGCGCCCTACCTCTTTTGGCTGGCGGCTATTGC | 4 |
| wt-120-240 | 2 | CGCCTCGTGTTTCACGGCCTCAGTTAGTACCGTTGTGACCGCCACCGGCTTGGCCCTCTCACTTCTACTCTTGGCAGCAGTGGCCAGCTCATATGCCGCTGCACAAAGGAAACTGCTGAC | 4 |
| wt-240-360 | 3 | ACCGGTGACAGTGCTTACTGCGGTTGTCACTTGTGAGTACACACGCACCATTTACAATGCATGATGTTCGTGAGATTGATCTGTCTCTAACAGTTCACTTCCTCTGCTTTTCTCCTCAGT | 4 |
| wt-360-480 | 4 | CTTTGCAATTTGCCTAACATGGAGGATTGAGGACCCACCTTTTAATTCTCTTCTGTTTGCATTGCTGGCCGCAGCTGGCGGACTACAAGGCATTTACGGTTAGTGTGCCTCTGTTATGAA | 4 |
| wt-480-600 | 5 | ATGCAGGTTTGACTTCATATGTATGCCTTGGCATGACGTCAACTTTACTTTTATTTCAGTTCTGGTGATGCTTGTGCTCCTGATACTAGCGTACAGAAGGAGATGGCGCCGTTTGACTGT | 4 |
| wt-600-720 | 6 | TTGTGGCGGCATCATGTTTTTGGCATGTGTACTTGTCCTCATCGTCGACGCTGTTTTGCAGCTGAGTCCCCTCCTTGGAGCTGTAACTGTGGTTTCCATGACGCTGCTGCTACTGGCTTT | 4 |
| wt-720-840 | 7 | CGTCCTCTGGCTCTCTTCGCCAGGGGGCCTAGGTACTCTTGGTGCAGCCCTTTTAACATTGGCAGCAGGTAAGCCACACGTGTGACATTGCTTGCCTTTTTGCCACATGTTTTCTGGACA | 4 |
| wt-840-960 | 8 | CAGGACTAACCATGCCATCTCTGATTATAGCTCTGGCACTGCTAGCGTCACTGATTTTGGGCACACTTAACTTGACTACAATGTTCCTTCTCATGCTCCTATGGACACTTGGTAAGTTTT | 4 |
| wt-960-1080 | 9 | CCCTTCCTTTAACTCATTACTTGTTCTTTTGTAATCGCAGCTCTAACTTGGCATCTCTTTTACAGTGGTTCTCCTGATTTGCTCTTCGTGCTCTTCATGTCCACTGAGCAAGATCCTTCT | 4 |
| wt-1080-1200 | 10 | GGCACGACTGTTCCTATATGCTCTCGCACTCTTGTTGCTAGCCTCCGCGCTAATCGCTGGTGGCAGTATTTTGCAAACAAACTTCAAGAGTTTAAGCAGCACTGAATTTATACCCAGTGA | 4 |
| wt-1200-1320 | 11 | GTATCTATTTGTTACTCCTGTTTAGTTGAAGAAAACAAGCTATTGGATTGTAACACACATTTTACGCTTTGTTCCTTAGATTTGTTCTGCATGTTATTACTGATTGTCGCTGGCATACTC | 4 |
| wt-1320-1440 | 12 | TTCATTCTTGCTATCCTGACCGAATGGGGCAGTGGAAATAGAACATACGGTCCAGTTTTTATGTGCCTCGGTGGCCTGCTCACCATGGTAGCCGGCGCTGTGTGGCTGACGGTGATGTCT | 4 |
| wt-1440-1560 | 13 | AACACGCTTTTGTCTGCCTGGATTCTTACAGCAGGATTCCTGATTTTCCTCATTGGTAAGTGTGACACCAACAGGTGTTGCCTTGTTATGTCACCGTTCTGACACATGACTTACATGGGT | 4 |
| wt-1560-1680 | 14 | TTGGCTTTTGTAGGCTTTGCCCTCTTTGGGGTCATTAGATGCTGCCGCTACTGCTGCTACTACTGCCTTACACTGGAAAGTGAGGAGCGCCCACCGACCCCATATCGCAACACTGTATAA | 4 |
| wt-1680-1800 | 15 | AGGTAAGTATTATTAAATTTTAGAGACACTATCACGTGTAACTTGACGTGCAAGGATGGAAGAGAGGGGCAGGGAAACGCAAATGCCGGTTGCCCGGTATGGGGGCCCGTTTATTATGGT | 4 |
| wt-1800-1920 | 16 | AAGGCTCTTCGGGCAAGATGGAGAGGCAAACATACAGGAGGAAAGGCTATATGAGCTACTCTCTGACCCACGCTCCGCGCTCGGCCTAGACCCGGGGCCCCTGATTGCTGAGAACCTGCT | 4 |
| wt-1920-2040 | 17 | GCTAGTGGCGCTGCGTGGCACCAACAACGATCCCAGGCCTCAGCGTCAGGAGAGGGCCAGAGAACTGGCCCTCGTTGGCATTCTACTAGGAAACGGCGAGCAGGGTGAACACTTGGGCAC | 4 |
| wt-2040-2160 | 18 | GGAGAGTGCCCTGGAGGCCTCAGGCAACAACTATGTGTATGCCTACGGACCAGACTGGATGGCAAGGCCTTCCACATGGTCCGCGGAAATCCAGCAATTCCTGCGACTCCTGGGCGCCAC | 4 |
| wt-2160-2280 | 19 | GTACGTGCTTCGCGTGGAGATGGGCAGGCAGTTTGGCTTCGAGGTGCATAGAAGCCGGCCCTCCTTCCGTCAGTTCCAGGCCATCAATCACCTTGTCCTGTTTGACAACGCCCTTCGCAA | 4 |
| wt-2280-2400 | 20 | GTACGATTCCGGCCAGGTGGCGGCGGGCTTCCAGAGGGCCCTTCTGGTGGCCGGGCCAGAGACCGCTGACACGAGGCCGGACCTCCGCAAGCTGAATGAGTGGGTGTTTGGTGGCAGGGC | 8 |
| wt-2400-2520 | 21 | TGCTGGTGGCAGACAGCTGGCCGACGAGCTAAAGATCGTGTCCGCGCTGCGAGACACTTACTCGGGCCACTTGGTCCTTCAGCCCACGGAGACCCTTGACACATGGAAGGTGTTGAGCAG | 4 |
| wt-2520-2640 | 22 | GGACACACGAACCGCTCATAGTTTGGAGCACGGATTCATTCATGCCGCGGGGACCATCCAGGCCAACTGCCCACAGCTGTTTATGAGACGCCAGCACCCCGGCCTCTTTCCCTTCGTTAA | 4 |
| wt-2640-2760 | 23 | TGCAATAGCATCATCGCTGGGCTGGTACTACCAGACCGCCACCGGCCCCGGAGCAGATGCCAGGGCGGCGGCCCGGCGCCAACAGGCCTTTCAGACCAGGGCGGCGGCTGAATGCCATGC | 8 |
| wt-2760-2880 | 24 | CAAAAGCGGGGTGCCGGTCGTGGCCGGCTTCTACAGGACCATCAACGCCACGCTCAAGGGAGGAGAGGGCCTACAGCCCACTATGTTTAACGGGGAGCTGGGGGCCATCAAGCACCAGGC | 4 |
| wt-2880-3000 | 25 | ACTTGACACTGTGAGGTATGACTACGGCCACTATCTCATAATGTTGGGGCCATTCCAGCCATGGAGCGGACTGACGGCCCCTCCGTGCCCCTACGCCGAAAGTTCATGGGCACAGGCGGC | 4 |
| wt-3000-3120 | 26 | CGTGCAGACGGCCCTCGAGCTGTTCTCGGCCCTGTACCCGGCCCCGTGCATCTCGGGCTACGCGCGCCCCCCGGGCCCCAGTGCTGTGATCGAGCATCTGGGGTCCCTAGTTCCAAAGGG | 8 |
| wt-3120-3240 | 27 | GGGTCTGCTGTTGTTTCTGTCTCACCTACCGGATGATGTTAAGGACGGGCTCGGAGAAATGGGGCCGGCCAGGGCCACGGGACCTGGAATGCAGCAGTTTGTCAGCAGCTACTTCCTCAA | 4 |
| wt-3240-3360 | 28 | CCCCGCCTGTTCCAACGTCTTCATTACAGTGAGGCAGCGAGGGGAGAAGATCAACGGCCGTACCGTCCTCCAAGCGCTCGGACGCGCATGCGATATGGCAGGCTGCCAGCACTATGTGCT | 4 |
| wt-3360-3480 | 29 | GGGCTCCACGGTTCCCCTCGGTGGACTCAACTTTGTCAACGACCTGGCGTCCCCGGTTTCCACCGCCGAGATGATGGATGATTTCTCTCCCTTCTTCACCGTGGAGTTTCCCCCGATTCA | 4 |
| wt-3480-3600 | 30 | AGAGGAGGGCGCAAGTTCTCCGGTACCCTTAGATGTGGACGAGAGCATGGACATCTCTCCGTCTTACGAGTTGCCCTGGCTCTCGCTGGAGTCATGCCTCACAAGCATCCTGTCACACCC | 4 |
| wt-3600-3720 | 31 | CACCGTGGGAAGCAAGGAGCACTTGGTCAGGCACACGGACAGGGTCAGCGGAGGACGCGTGGCACAGCAGCCCGGGGTAGGTCCCCTGGACCTGCCGCTGGCGGACTACGCCTTCGTTGC | 8 |
| wt-3720-3840 | 32 | CCACAGTCAGGTCTGGACCAGGCCCGGTGGGGCTCCTCCCTTGCCCTATCGTACCTGGGATCGAATGACAGAGAAGCTGCTTGTCTCCGCAAAACCCGGCGGAGAGAACGTTAAGGTTTC | 4 |
| wt-3840-3960 | 33 | AGGTACCGTGATTACATTGGGAGAACAGGGGTACAAAGTGTCGTTGGATCTGAGGGAGGGAACCAGGCTGGCAATGGCTGAGGCGCTGCTGAACGCAGCATGTGCCCCAATCTTGGATCC | 4 |
| wt-4080-4200 | 35 | CTTTGGCTCGGCCTCCTGCCCCGAGACCGGCTCGTCCGCCTCCAACTTCATGACCGTGGTGGCCTCTGTCTCCGCCCCAGGGGAATTCTCGGGTCCTCTGATCACGCCAGTGCTTCAGAA | 4 |
| wt-4200-4320 | 36 | GACGGGCAGTCTCCTGATTGCGGTGCGTTGCGGGGATGGCAAGATCCAGGGAGGGTCGCTGTTTGAGCAGCTCTTTAGCGACGTGGCCACGACCCCACGGGCACCCGAGGCGTTGTCTCT | 4 |
| wt-4320-4440 | 37 | GAAGAATCTCTTCCGGGCAGTCCAGCAGCTGGTCAAGAGCGGCATCGTGCTGTCAGGGCATGACATCAGCGACGGGGGCCTGGTGACCTGCCTGGTGGAGATGGCCCTGGCCGGGCAGCG | 8 |
| wt-4560-4680 | 39 | CTCCATGAACATGTACCCGGCAGTCCTCGGTCGAGTGGGCGAGCAAGGTCCAGATCAAATGTTTGAGGTGCAGCACGGCCCAGAGACGGTGTTGCGCCAGTCGCTGCGCCTGCTGCTGGG | 4 |
| wt-4680-4800 | 40 | AACCTGGTCATCCTTTGCCAGCGAGCAGTACGAGTGCCTGCGACCAGATCGGATTAACCGGTCCATGCACGTGTCCGACTACGGCTATAACGAAGCACTGGCAGTCTCCCCGTTGACAGG | 4 |
| wt-4920-5040 | 42 | ATGCCTGTGCCGACGGGTGTTCTTTCGCGAGGTTAGGGACAACACGTTCCTCGACAAGTACGTGGGTCTGGCCATCGGAGGAGTTCATGGGGCCAGGGACTCTGCCCTGGCAGGCCGTGC | 4 |
| wt-5040-5160 | 43 | CACCGTGGCGCTGATTAATCGTTTCCCCGCCCTGCGTGACGCTATTCTAAAGTTCCTCAACAGGCCAGATACGTTCTCGGTGGCCTTGGGGGAGCTGGGGGTGCAAGTTTTGGCTGGCCT | 4 |
| wt-5160-5280 | 44 | GGGGGCCGTGGGGTCAACAGATAATCCACCCGCCCCTGGCGTGGAAGTTAATGTCCAGAGATCACCTCTGATTCTGGCCCCCAACGCCTCTGGCATGTTTGAGTCCCGCTGGCTGAACAT | 4 |
| wt-5280-5400 | 45 | TAGCATCCCGGCGACCACCAGCTCTGTCATGCTGCGTGGCCTCCGGGGCTGCGTCCTGCCTTGTTGGGTGCAAGGCTCGTGCCTGGGCCTGCAATTTACTAACCTCGGGATGCCATATGT | 4 |
| wt-5400-5520 | 46 | TTTGCAGAATGCCCACCAGATCGCCTGCCACTTCCACAGCAATGGCACGGATGCCTGGCGCTTTGCTATGAATTATCCAAGAAACCCCACGGAGCAGGGCAACATTGCAGGGCTCTGTTC | 4 |
| wt-5520-5640 | 47 | ACGCGATGGTCGTCATCTGGCTCTCCTGTGTGACCCCTCACTTTGTACAGACTTTTGGCAATGGGAGCACATTCCCCCCGCCTTTGGGCACCCCACGGGGTGCTCCCCCTGGACACTTAT | 4 |
| wt-5640-5760 | 48 | GTTTCAAGCAGCTCACCTATGGTCACTCAGGCACGGTCGCCCCTCCGAGTGACCAGTCACCTTCCAGACTATGCATACACTGAATTTAGCCTGATATTGTCCCCCTAGCCCCGGGCCCAG | 4 |
| wt-5760-5880 | 49 | CCCTCCTCAGAAAACTCTGCATGGAGAAGCTGGACGTGAACCTCCCCCCCAGACCTGTGTGCTGTATTTACAAACACTACAATAAACCCAATGTGCAAATGTGGTTTGTATGGCTACTTT | 4 |
| wt-5880-6000 | 50 | GTGTTCCTAAAAAATGCAACAATAGAAGTGGAAACCCTCAGTCACGGGACATTAACCTCAACCACAAAATGGGGGTTGGAGAAAGTAACCACATATACTGGAGATGATTCATGGGCTGGG | 4 |
| wt-6000-6120 | 51 | GGTTCCCGGACAATACACCCATCTGGAGTTCAACCTAATTACATGGTAGATAAATTAAGAGTCCCTCCTCACCACTCGAAACTATGGCAGACATTCTATAAGATAACGAGGAGAGATGAG | 4 |

| | | | |
|---|---|---|---|
| wt-6120-6240 | 52 | GTGAGGGCAGAGGACATTGGGCAGGTGTGGGCCACGGGGCAGCTGGCCATATCCCCCGCACTACAGAAGTGTAAGCAAAGTGAAGGGCTCGGAAGGCAGGCGGGGCCTAGCAATGTCACA | 4 |
| wt-6240-6360 | 53 | GCTAAATGCCCACCAGGGCACACACTCAAGCGGGGTCTCGGAGCTCCTAGGTCAGACCACGAAAGGTCAGCCTGCAAGGTGGATGGCGTGTTTTCTGAGGTTATCCCCGCTACGTGCAGT | 4 |
| wt-6360-6480 | 54 | GCTGGGTGATAGAGACCCTAGAATGTGTCGAAATGACCAAGCGTCCCCGCAGCGGGGCTCCCAACACGGGTTCCCAGAGAGGGTAAAAGAGGGGGCCATAAAGCCCAGGGTGTAAAACAC | 4 |
| wt-6480-6600 | 55 | CGACCGCGCCACCAGATGGCACACGTGGGGGAAATGAGGGTTAGCATAGGCAACCCCCGCCTACACACCAACTATAGCAAACCCCGCCCCGTCACGGTGACGTAGTCTGTCTTGAGGAGA | 4 |
| wt-6600-6720 | 56 | TGTAGACTTGTAGACACTGCAAAACCTCAGGACCTACGCTGCCCTAGAGGTTTTGCTAGGGAGGAGACGTGTGTGGCTGTAGCCACCCGTCCCGGGTACAAGTCCCGGGTGGTGAGGACG | 4 |
| wt-6720-6840 | 57 | GTGTCTGTGGTTGTCTTCCCAGACTCTGCTTTCTGCCGTCTTCGGTCAAGTACCAGCTGGTGGTCCGCATGTTTTGATCCAAACTTTTGTTTTAGGATTTATGCATCCATTATCCCGCAG | 4 |
| wt-6840-6960 | 58 | TTCCACCTAAACGGGGCTTAACGTTGCATCCCAGAAGATGCACGCTTAACCCCGCCTACAACCGTGACGTAGCTGTTTACCAGCATGTATAGAGTTACGGTTCGCTACATCAAACAGGAC | 4 |
| wt-6960-7080 | 59 | AGCCGTTGCCCTAGTGGTTTCGGACACACCGCCAACGCTCAGTGCGGTGCTACCGACCCGAGGTCAAGTCCCGGGGGAGGAGAAGAGAGGCTTCCCGCCTAGAGCATTTGCAAGTCAGGA | 4 |
| wt-7080-7200 | 60 | TTCTCTAATCCCTCTGGGAGAAGGGTATTCGGCTTGTCCGCTATTTTTTTGTGGCTAGTTTTGCACCCACAACATGTAAGGGCCCGCTACCCCTACAACACAAAACAAACTATCTCCCCT | 4 |
| wt-7200-7320 | 61 | AACCATCCTTTTGCCAATCAATTCTGTGACAGGGTTTCCTGGACACCCAGTCTTAGTTCAGGTAGACACCCAGTTATGCAGTGCCACCAATTCCAACCATTTTTAAACCTCCTGGAATTC | 4 |
| wt-7320-7440 | 62 | TATCATTAAACGGCATGCAGGAAAAGGACAAGCAGCGAAAATTCACGCCCCCTTGGGAGGTGGCGGCATATGCAAAGGATAGCACTCCCACTCTACTACTGGGTATCATATGCTGACTGT | 4 |
| wt-7440-7560 | 63 | ATATGCATGAGGATAGCATATGCTACCCGGATACAGATTAGGATAGCATATACTACCCAGATATAGATTAGGATAGCATATGCTACCCAGATATAGATTAGGATAGCCTATGCTACCCAG | 4 |
| wt-7560-7680 | 64 | ATATAAATTAGGATAGCATATACTACCCAGATATAGATTAGGATAGCATATGCTACCCAGATATAGATTAGGATAGCCTATGCTACCCAGATATAGATTAGGATAGCATATGCTACCCAG | 4 |
| wt-7680-7800 | 65 | ATATAGATTAGGATAGCATATGCTATCCAGATATTTGGGTAGTATATGCTACCCAGATATAAATTAGGATAGCATATACTACCCTAATCTCTATTAGGATAGCATATGCTACCCGGATAC | 4 |
| wt-7800-7920 | 66 | AGATTAGGATAGCATATACTACCCAGATATAGATTAGGATAGCATATGCTACCCAGATATAGATTAGGATAGCCTATGCTACCCAGATATAAATTAGGATAGCATATACTACCCAGATAT | 4 |
| wt-7920-8040 | 67 | AGATTAGGATAGCATATGCTACCCAGATATAGATTAGGATAGCCTATGCTACCCAGATATAGATTAGGATAGCATATGCTATCCAGATATTTGGGTAGTATATGCTACCCATGGCAACAT | 4 |
| wt-8040-8160 | 68 | TAGCCCACCGTGCTCTCAGCGACCTCGTGAATATGAGGACCAACAACCCTGTGCTTGGCGCTCAGGCGCAAGTGTGTGTAATTTGTCCTCCAGATCGCAGCAATCGCGCCCCTATCTTGG | 4 |
| wt-8160-8280 | 69 | CCCGCCCACCTACTTATGCAGGTATTCCCCGGGGTGCCATTAGTGGTTTTGTGGGCAAGTGGTTTGACCGCAGTGGTTAGCGGGGTTACAATCAGCCAAGTTATTACACCCTTATTTTAC | 4 |
| wt-8280-8400 | 70 | AGTCCAAAACCGCAGGGCGGCGTGTGGGGGCTGACGCGTGCCCCCACTCCACAATTTCAAAAAAAAGAGTGGCCACTTGTCTTTGTTTATGGGCCCCATTGGCGTGGAGCCCCGTTTAAT | 4 |
| wt-8400-8520 | 71 | TTTCGGGGGTGTTAGAGACAACCAGTGGAGTCCGCTGCTGTCGGCGTCCACTCTCTTTCCCCTTGTTACAAATAGAGTGTAACAACATGGTTCACCTGTCTTGGTCCCTGCCTGGGACAC | 4 |
| wt-8520-8640 | 72 | ATCTTAATAACCCCAGTATCATATTGCACTAGGATTATGTGTTGCCCATAGCCATAAATTCGTGTGAGATGGACATCCAGTCTTTACGGCTTGTCCCCACCCCATGGATTTCTATTGTTA | 4 |
| wt-8640-8760 | 73 | AAGATATTCAGAATGTTTCATTCCTACACTAGTATTTATTGCCCAAGGGGTTTGTGAGGGTTATATTGGTGTCATAGCACAATGCCACCACTGAACCCCCCGTCCAAATTTTATTCTGGG | 4 |
| wt-8760-8880 | 74 | GGCGTCACCTGAAACCTTGTTTTCGAGCACCTCACATACACCTTACTGTTCACAACTCAGCAGTTATTCTATTAGCTAAACGAAGGAGAATGAAGAAGCAGGCGAAGATTCAGGAGAGTT | 4 |
| wt-8880-9000 | 75 | CACTGCCCGCTCCTTGATCTTCAGCCACTGCCCTTGTGACTAAAATGGTTCACTACCCTCGTGGAATCCTGACCCCATGTAAATAAAACCGTGACAGCTCATGGGGTGGGAGATATCGCT | 4 |
| wt-9000-9120 | 76 | GTTCCTTAGGACCCTTTTACTAACCCTAATTCGATAGCATATGCTTCCCGTTGGGTAACATATGCTATTGAATTAGGGTTAGTCTGGATAGTATATACTACTACCCGGGAAGCATATGCT | 4 |
| wt-9120-9240 | 77 | ACCCGTTTAGGGTTAACAAGGGGGCCTTATAAACACTATTGCTAATGCCCTCTTGAGGGTCCGCTTATCGGTAGCTACACAGGCCCCTCTGATTGACGTTGGTGTAGCCTCCCGTAGTCT | 4 |
| wt-9240-9360 | 78 | TCCTGGGCCCCTGGGAGGTACATGTCCCCCAGCATTGGTGTAAGAGCTTCAGCCAAGAGTTACACATAAAGGCAATGTTGTGTTGCAGTCCACAGACTGCAAAGTCTGCTCCAGGATGAA | 4 |
| wt-9360-9480 | 79 | AGCCACTCAGTGTTGGCAAATGTGCACATCCATTTATAAGGATGTCAACTACAGTCAGAGAACCCCTTTGTGTTTGGTCCCCCCCCGTGTCACATGTGGAACAGGGCCCAGTTGGCAAGT | 4 |
| wt-9480-9600 | 80 | TGTACCAACCAACTGAAGGGATTACATGCACTGCCCCGCGGGAAATACGTCCTACCCAGGAACCCGAAACAGTGTTTCCCAGAAGCTGTAAAAATAGAACGCCCTGGAACTGCCCCACTG | 4 |
| wt-9600-9720 | 81 | TGCAATGCAGCTTTTAGCCATGCCATGCTCTATAAATCACTTCCCTATCTCAGGTAGGCCTGCACACCTTAGGTATGGAGCGAAGGTTAGTGGTCACTCTGCAGTGCCTGGTGCTGCTTT | 4 |
| wt-9720-9840 | 82 | ACCTGGCACCTGAGTGTGGAGGTACAGACCAATGTGACAATTTTCCCCAAATGTTGAGGGACCTAAGAGATGCCTTCAGTCGTGTTAAAACCTTTTTCCAGACAAAGGACGAGGTAGATA | 4 |
| wt-9840-9960 | 83 | ACCTTTTGCTCAAGGAGTCTCTGCTAGAGGACTTTAAGGGCTACCTTGGATGCCAGGCCCTGTCAGAAATGATCCAATTCTACCTGGAGGAAGTCATGCCACAGGCTGAAAACCAGGACC | 4 |
| wt-9960-10080 | 84 | CTGAAGCCAAAGACCATGTCAATTCTTTGGGTGAAAATCTAAAGACCCTACGGCTCCGCCTGCGCAGGTGCCACAGGTTCCTGCCGTGTGAGAACAAGAGTAAAGCTGTGGAACAGATAA | 4 |
| wt-10200-10320 | 85 | TGGAAGCAGGAGATGGGTGCATTTCACCCCAACCCCCCCTTTCGACTGTCATTTACAATAAAATGAAACCTTTTATTCTTGATTGCCTCTTGTGTTCTTGCCGCCCAGGTACCTTCCTGT | 4 |
| wt-10320-10440 | 86 | GTTCTCCCCACGGGAAAAAGAATAGCTTCTGCAGAAGGCCATTGACGCAAGTTTTGCCCGTGGGGATTACCCGACCCAGCCACTTACAGCACATTTTGTTCTAGGTCCATCTTAGGAGCC | 4 |
| wt-10440-10560 | 87 | CGGGCCAGCATTCTATCAGCTTAACGGGAAGAGAAGTGGGGAGGGCACTCGCCCACTAACCTTAACACCTGCAGCCTACAAAAGTACACTAGCTGTTTGCTCTATTCGCCACTAGAGACC | 4 |
| wt-10560-10680 | 88 | GCCAAGATGCGAAACTACAGGCCCGGGCCCAGGCCTTGCAGGGCAGACGGTTAGGCTGACAAGGGGACAAGTGTGGCAGGTGGGCGGGAAGGGGCACAAGAATGCCGGCGAAACTGGACC | 4 |
| wt-10680-10800 | 89 | ACGGTCCACCCCGCCCTCAAGCGTCCGGGAGCCGGGCGGCTCGGCTAAGGAGGGCGGCCTTGCGAACAATTATTAGTAGCTACCAACAAGGGCCCCCAGATGCCCCCCACCAGTCACCCG | 8 |
| wt-10800-10920 | 90 | GCCGTGTCCACTCACATATTCCACTCTTATTTTTAAATTAATGTGTCCCAATTAGAAACCCAAGCGCAGAAATTAGTTGAGAGGCTAGTGTTTTAAACATGCACCCTAGGCCAGCCAGAG | 4 |
| wt-10920-11040 | 91 | ATAATGTCACAAGATTATCAAGTTGGTGTAAACACGCCGTGGGAAAAAATTTATGGTTCAGTGCGTCGAGTGCTATCTTTGGAACAGTAGAAAATTGAACCTTGTTGGCGGGAGAAGGAA | 4 |
| wt-11040-11160 | 92 | TAACGCCTTATCTGGGAGGAGCGACGGATTATAGCCAATAAGAGAGCTCAAGACGCAGGGCTCGCAAAGTATAGTGGCCCCGTGGGACCTTAGAGGTGGAGCAACGTCTAAAGTGGTAAT | 4 |
| wt-11160-11280 | 93 | AACACCAGGCGGGGCTGGGCAAAGGGGTCCTACGGGCGGGATTAATTACGCCTTGCTTACGCAAGCTCAGTTAATTCGCCCACGACTTGAAAAATGTAGCCCTTAACCAATTGGCGGCCC | 4 |
| wt-11280-11400 | 94 | CTAAGGGGGGGACTAAGGTCCCACTACAAAAACTCTGTGTTCTGCTGCAAATTTTAGATCAGATGGCATAGAGACAAGGACACCGAAGACCCCCAGAGCCCTCATCGCAGGGTTCTTACC | 4 |
| wt-11400-11520 | 95 | ATGCGGCCATGTAGGCCCACTTAACACTACAAGACCTACGCCTCTCCATTCATCATGTAACCCACAAATCATCTAAACCGTAAGTCTAAGGGCCTCCTGAGGTTTTCTCAGGAGGCCCTA | 4 |
| wt-11520-11640 | 96 | ATGTATAATTAATCATGCATTTGATTTTAAAAAAGTAGGTTACACTCATTTTAGGCCAGACTTTATTTGCAGATTAATAATTTATGTGATTCTCCTTCCCTCTAGGACTGAAGAAACAGC | 4 |
| wt-11640-11760 | 97 | CTCCTGCACGTGAGCATGTATCTGAAATAATTATTATGTCATAAGTGTAATGATTAGAAAGTCATAAACCCACTTCCCTTTACATGAATCTGGGCACTGAATTTTGGGGTACTTCTAAAG | 4 |
| wt-11760-11880 | 98 | ACTAACGTGTTCGATTTCGGGGTCACTTCCCCTTTTATAAGTGTGTGAACAGTGATTTCAGTAAAACCTAAGAGATATTTGGTGTCACTTCCGCATTTTAAGTTTCAGAAAATTTTAAAA | 4 |
| wt-11880-12000 | 99 | TTAAAATTGAAATTTCTCTCAAAATAATTCCAATGAAAACTTCAAAGAATCTTATGTATGTAATTCTTTTGCCCCAAACTGGGCTTCAGATGCCTTCTATTGCACTCTCACAAAAACATT | 4 |
| wt-12000-12120 | 100 | CTGGACACATGTGCCAGACGCCTGGGCCTCTAAGGCCCTCGGGTCCCCCTGGACCCCGGCCTCAGCAACCCTGCTGCTCCCCTCCTGCCACCCCAGCCTCCCCCCCTCCCCGTCCCCCTT | 8 |
| wt-12120-12240 | 101 | CGCTCCTGTTCCTCCCCCGGTCCCCAGTAGGGCCGCCTGCCCCCCTGCACCCAGTACCTGCCCCTCTTGGCCACGCACCCCGGGCCAGGCCACCTTAGACCCGGCCAAGCCCCATCCCTG | 8 |
| wt-12240-12360 | 102 | AAGACCCAGCGGCCATTCTCTCTGGTAACGAGCAGAGAAGAAGTAGAGGCCCGCGGCCATTGGGCCCAGATTGAGAGACCAGTCCAGGGGCCCGAGGTTGGAGCCAGCGGGCACCCGAGG | 4 |

| | | | |
|---|---|---|---|
| wt-12360-12480 | 103 | TCCCAGCACCCGGTCCCTCCGGGGGGCAGAGACAGGCAGGGCCCCCCGGCAGCTGGCCCCGAGGAGGCGCCCGGAGTGGGGCCGGTCGGCTGGGCTGGCCGAGCCCGGGTCTGGGAGGTC | 12 |
| wt-12480-12600 | 104 | TGGGGTGGCGAGCCTGCTGTCTCAGGAGGGGCCTGGCTCCGCCGGGTGGCCCTGGGGTAAGTCTGGGAGGCAGAGGGTCGGCCTAGGCCCGGGGAAGTGGAGGGGGATCGCCCGGGTCTC | 8 |
| wt-12600-12720 | 105 | TGTTGGCAGAGTCCGGGCGATCCTCTGAGACCCTCCGGGCCCGGACGGTCGCCCTCAGCCCCCCAGACAGACCCCAGGGTCTCCAGGCAGGGTCCGGCATCTTCAGGGGCAGCAGGCTCA | 8 |
| wt-12720-12840 | 106 | CCACCACAGGCCCCCCAGACCCGGGTCTCGGCCAGCCGAGCCGACCGGCCCCGCGCCTGGCGCCTCCTCGGGGCCAGCCGCCGGGGTTGGTTCTGCCCCTCTCTCTGTCCTTCAGAGGAA | 8 |
| wt-12840-12960 | 107 | CCAGGGACCTCGGGCACCCCAGAGCCCCTCGGGCCCGCCTCCAGGCGCCCTCCTGGTCTCCGCTCCCCTCTGAGCCCCGTTAAACCCAAAGAATGTCTGAGGGGAGCCACCCTCGGGGCC | 8 |
| wt-12960-13080 | 108 | CAGGCCCCAGAGTCCAGAGGTCAGGGGCACCTCAGGGTGCCTCCCCGGGTCCCAGGCCAGCCGGAGGGACCCCGGCAGCCCGGGCGGCCCCAGAGGCCGGTTCCTCGCCCCTTCCCCGGG | 12 |
| wt-13080-13200 | 109 | CTTCAGAGCCCAGGATGTCCCCCAGAAGGGACCCTAGGCGTCCCCTCTCCTCCCCTCCAGGCCCGAGCCTCTCCCTCGCGGAGAGGGGCCTCTTTGGGCCCTCAAGTCCAGCCCCACCGA | 8 |
| wt-13200-13320 | 110 | GACCCGAGTGGCCCGGATCCCCCCACCGGCCCTTCTCTCTGTCCCCCTGCTCCTCTCCAACCTTCGCTCCACCCTAGACCCCAGCTTCTGGCCTCCCCGGGTCCACCAGGCCAGCCGGAG | 8 |
| wt-13320-13440 | 111 | GGACCCCGGCAGCCCGGGCGAGTCGCCTTCCCTCTCCCCTGGCCTCTCCTTCCCGCCTCCCACCCGAGCCCCCTCAGCTTGCCTCCCCACCGGGTCCATCAGGCCGGCCGGAGGGACCCC | 12 |
| wt-13440-13560 | 112 | GGCGGCCCGGTGTCAGTCCCCCCTGCAGCCGCCCAGTCTCTGCCTCCAGGCAAGGGCGCCAGCTTTTCTCCCCCCAGCCTGAGGCCCAGTCTCCTGTGCACTGTCTGTAAAGTCCAGCCT | 8 |
| wt-13560-13680 | 113 | CCCACGCCCGTCCACGGCTCCCGGGCCCAGCCTCGTCCACCCCTCCCCACGGTGGACAGGCCCTCTGTCCACCCGGGCCATCCCCGCCCCCCTGTGTCCACCCCAGTCCCGTCCAGGGGG | 12 |
| wt-13680-13800 | 114 | GACTTTATGTGACCCTTGGGCCTGGCTCCCCATAGACTCCCATGTAAGCCTGCCTCGAGTAGGTGCCTCCAGAGCCCCTTTTGCCCCCCTGGCGGCCCAGCCCGACCCCCGGGCGCCCCC | 8 |
| wt-13800-13920 | 115 | AAACTTTGTCCAGATGTCCAGGGGTCCCCGAGGGTGAGGCCCAGCCCCCTCCCGCCCCTGTCCACTGCCCCGGTCCCCCCAGAAGCCCCCAAAAGTAGAGGCTCAGGCCATGCGCGCCCT | 8 |
| wt-13920-14040 | 116 | GTCACCAGGCCTGCCAAAGAGCCAGATCTAAGGCCGGGAGAGGCAGCCCCAAAGCGGGTGCAGTAACAGGTAATCTCTGGTAGTGATTTGGACCCGAAATCTGACACTTTAGAGCTCTGG | 4 |
| wt-14040-14160 | 117 | AGGACTTTAAAACTCTAAAAATCAAAACTTTAGAGGCGAATGGGCGCCATTTTGTCCCCACGCGCGCATAATGGCGGACCTAGGCCTAAAACCCCCAGGAAGCGGGTCTATGGTTGGCTG | 4 |
| wt-14160-14280 | 118 | CGCTGCTGCTATCTTTAGAGGGGAAAAGAGGAATAAGCCCCCAGACAGGGGAGTGGGCTTGTTTGTGACTTCACCAAAGGTCAGGGCCCAAGGGGGTTCGCGTTGCTAGGCCACCTTCTC | 4 |
| wt-14280-14400 | 119 | AGTCCAGCGCGTTTACGTAAGCCAGACAGCAGCCAATTGTCAGTTCTAGGGAGGGGGACCACTGCCCCTGGTATAAAGTGGTCCTGCAGCTATTTCTGGTCGCATCAGAGCGCCAGGAGT | 4 |
| wt-14400-14520 | 120 | CCACACAAATGTAAGAGGGGGTCTTCTACCTCTCCCTAGCCCTCCGCCCCCTCCAAGGACTCGGGCCCAGTTTCTAACTTTTCCCCTTCCCTCCCTCGTCTTGCCCTGCGCCCGGGGCCA | 4 |
| wt-14520-14640 | 121 | CCTTCATCACCGTCGCTGACTCCGCCATCCAAGCCTAGGGGAGACCGAAGTGAAGGCCCTGGACCAACCCGGCCCGGGCCCCCCGGTATCGGGCCAGAGGTAAGTGGACTTTAATTTTTT | 4 |
| wt-14640-14760 | 122 | CTGCTAAGCCCAACACTCCACCACACCCAGGCACACACTACACACACCCACCCGTCTCAGGGTCCCCTCGGACAGCTCCTAAGAAGGCACCGGTCGCCCAGTCCTACCAGAGGGGGCCAA | 4 |
| wt-14760-14880 | 123 | GAACCCAGACGAGTCCGTAGAAGGGTCCTCGTCCAGCAAGAAGAGGAGGTGGTAAGCGGTTCACCTTCAGGGGTAAGTAACCTGACCTCTCCAGGGCTCACATAAAGGGAGGCTTAGTAT | 4 |
| wt-14880-15000 | 124 | ACATGCTTCTTGCTTTTCACAGGAACCTGGGGGCTAGTCTGGGTGGGATTAGGCTGCCTCAAGTTGCATCAGCCAGGGCTTCATGCCCTCCTCAGTTCCCTAGTCCCCGGGCTTCAGGCC | 4 |
| wt-15120-15240 | 126 | CTGGACCCCGGCCTCAGCAACCCTGCTGCTCCCCTCCTGCCACCCCAGCCTCCCCCCCTCCCCGTCCCCCTTCGCTCCTGATCCTCCCCCGGTCCCCAGTAGGGCCGCCTGCCCCCCTGC | 12 |
| wt-15240-15360 | 127 | ACCCAGTACCTGCCCCTCTTGGCCACGCACCCCGGGCCAGGCCACCTTAGACCCGGCCAAGCCCCATCCCTGAAGACCCAGCGGCCATTCTCTCTGGTAACGAGCAGAGAAGAAGTAGAG | 4 |
| wt-15360-15480 | 128 | GCCCGCGGCCATTGGGCCCAGATTGAGAGACCAGTCCAGGGGCCCGAGGTTGGAGCCAGCGGGCACCCGAGGTCCCAGCACCCGGTCCCTCCGGGGGGCAGAGACAGGCAGGGCCCCCCG | 8 |
| wt-15480-15600 | 129 | GCAGCTGGCCCCGAGGAGGCGCCCGGAGTGGGGCCGGTCGGCTGGGCTGGCCGAGCCCGGGTCTGGGAGGTCTGGGGTGGCGAGCCTGCTGTCTCAGGAGGGGCCTGGCTCCGCCGGGTG | 12 |
| wt-15600-15720 | 130 | GCCCTGGGGTAAGTCTGGGAGGCAGAGGGTCGGCCTAGGCCCGGGGAAGTGGAGGGGGATCGCCCGGGTCTCTGTTGGCAGAGTCCGGGCGATCCTCTGAGACCCTCCGGGCCCGGACGG | 8 |
| wt-15720-15840 | 131 | TCGCCCTCAGCCCCCCAGACAGACCCCAGGGTCTCCAGGCAGGGTCCGGCATCTTCAGGGGCAGCAGGCTCACCACCACAGGCCCCCCAGACCCGGGTCTCGGCCAGCCGAGCCGACCGG | 8 |
| wt-15840-15960 | 132 | CCCCGCGCCTGGCGCCTCCTCGGGGCCAGCCGCCGGGGTTGGTTCTGCCCCTCTCTCTGTCCTTCAGAGGAACCAGGGACCTCGGGCACCCCAGAGCCCCTCGGGCCCGCCTCCAGGCGC | 12 |
| wt-15960-16080 | 133 | CCTCCTGGTCTCCGCTCCCCTCTGAGCCCCGTTAAACCCAAAGAATGTCTGAGGGGAGCCACCCTCGGGGCCCAGGCCCCAGAGTCCAGAGGTCAGGGGCACCTCAGGGTGCCTCCCCGG | 8 |
| wt-16080-16200 | 134 | GTCCCAGGCCAGCCGGAGGGACCCCGGCAGCCCGGGCGGCCCCAGAGGCCGGTTCCTCGCCCCTTCCCCGGGCTTCAGAGCCCAGGATGTCCCCCAGAAGGGACCCTAGGCGTCCCCTCT | 8 |
| wt-16200-16320 | 135 | CCTCCCCTCCAGGCCCGAGCCTCTCCCTCGCGGAGAGGGGCCTCTTTGGGCCCTCAAGTCCAGCCCCACCGAGACCCGAGTGGCCCGGATCCCCCCACCGGCCCTTCTCTCTGTCCCCCT | 8 |
| wt-16320-16440 | 136 | GCTCCTCTCCAACCTTCGCTCCACCCTAGACCCCAGCTTCTGGCCTCCCCGGGTCCACCAGGCCAGCCGGAGGGACCCCGGCAGCCCGGGCGAGTCGCCTTCCCTCTCCCCTGGCCTCTC | 8 |
| wt-16440-16560 | 137 | CTTCCCGCCTCCCACCCGAGCCCCCTCAGCTTGCCTCCCCACCGGGTCCATCAGGCCGGCCGGAGGGACCCCGGCGGCCCGGTGTCAGTCCCCCCTGCAGCCGCCCAGTCTCTGCCTCCA | 12 |
| wt-16560-16680 | 138 | GGCAAGGGCGCCAGCTTTTCTCCCCCCAGCCTGAGGCCCAGTCTCCTGTGCACTGTCTGTAAAGTCCAGCCTCCCACGCCCGTCCACGGCTCCCGGGCCCAGCCTCGTCCACCCCTCCCC | 8 |
| wt-16680-16800 | 139 | ACGGTGGACAGGCCCTCTGTCCACCCGGGCCATCCCCGCCCCCCTGTGTCCACCCCAGTCCCGTCCAGGGGGGACTTTATGTGACCCTTGGGCCTGGCTCCCCATAGACTCCCATGTAAG | 8 |
| wt-16800-16920 | 140 | CCTGCCTCGAGTAGGTGCCTCCAGAGCCCCTTTTGCCCCCCTGGCGGCCCAGCCCGACCCCCGGGCGCCCCCAAACTTTGTCCAGATGTCCAGGGGTCCCCGAGGGTGAGGCCCAGCCCC | 8 |
| wt-16920-17040 | 141 | CTCCCGCCCCTGTCCACTGCCCCGGTCCCCCCAGAAGCCCCCAAAAGTAGAGGCTCAGGCCATGCGCGCCCTGTCACCAGGCCTGCCAAAGAGCCAGATCTAAGGCCGGGAGAGGCAGCC | 8 |
| wt-17040-17160 | 142 | CCAAAGCGGGTGCAGTAACAGGTAATCTCTGGTAGTGATTTGGACCCGAAATCTGACACTTTAGAGCTCTGGAGGACTTTAAAACTCTAAAAATCAAAACTTTAGAGGCGAATGGGCGCC | 4 |
| wt-17160-17280 | 143 | ATTTTGTCCCCACGCGCGCATAATGGCGGACCTAGGCCTAAAACCCCCAGGAAGCGGGTCTATGGTTGGCTGCGCTGCTGCTATCTTTAGAGGGGAAAAGAGGAATAAGCCCCCAGACAG | 4 |
| wt-17280-17400 | 144 | GGGAGTGGGCTTGTTTGTGACTTCACCAAAGGTCAGGGCCCAAGGGGGTTCGCGTTGCTAGGCCACCTTCTCAGTCCAGCGCGTTTACGTAAGCCAGACAGCAGCCAATTGTCAGTTCTA | 4 |
| wt-17400-17520 | 145 | GGGAGGGGGACCACTGCCCCTGGTATAAAGTGGTCCTGCAGCTATTTCTGGTCGCATCAGAGCGCCAGGAGTCCACACAAATGTAAGAGGGGGTCTTCTACCTCTCCCTAGCCCTCCGCC | 4 |
| wt-17520-17640 | 146 | CCCTCCAAGGACTCGGGCCCAGTTTCTAACTTTTCCCCTTCCCTCCCTCGTCTTGCCCTGCGCCCGGGGCCACCTTCATCACCGTCGCTGACTCCGCCATCCAAGCCTAGGGGAGACCGA | 4 |
| wt-17640-17760 | 147 | AGTGAAGGCCCTGGACCAACCCGGCCCGGGCCCCCCGGTATCGGGCCAGAGGTAAGTGGACTTTAATTTTTTCTGCTAAGCCCAACACTCCACCACACCCAGGCACACACTACACACACC | 4 |
| wt-17760-17880 | 148 | CACCCGTCTCAGGGTCCCCTCGGACAGCTCCTAAGAAGGCACCGGTCGCCCAGTCCTACCAGAGGGGGCCAAGAACCCAGACGAGTCCGTAGAAGGGTCCTCGTCCAGCAAGAAGAGGAG | 4 |
| wt-17880-18000 | 149 | GTGGTAAGCGGTTCACCTTCAGGGGTAAGTAACCTGACCTCTCCAGGGCTCACATAAAGGGAGGCTTAGTATACATGCTTCTTGCTTTTCACAGGAACCTGGGGGCTAGTCTGGGTGGGA | 4 |
| wt-18000-18120 | 150 | TTAGGCTGCCTCAAGTTGCATCAGCCAGGGCTTCATGCCCTCCTCAGTTCCCTAGTCCCCGGGCTTCAGGCCCCCTCCGTCCCCGTCCTCCAGAGACCCGGGCTTCAGGCCCTGCCTCTC | 8 |
| wt-18240-18360 | 152 | CCTCCCCCCCTCCCCGTCCCCCTTCGCTCCTGATCCTCCCCCGGTCCCCAGTAGGGCCGCCTGCCCCCCTGCACCCAGTACCTGCCCCTCTTGGCCACGCACCCCGGGCCAGGCCACCTT | 8 |
| wt-18360-18480 | 153 | AGACCCGGCCAAGCCCCATCCCTGAAGACCCAGCGGCCATTCTCTCTGGTAACGAGCAGAGAAGAAGTAGAGGCCCGCGGCCATTGGGCCCAGATTGAGAGACCAGTCCAGGGGCCCGAG | 4 |
| wt-18480-18600 | 154 | GTTGGAGCCAGCGGGCACCCGAGGTCCCAGCACCCGGTCCCTCCGGGGGGCAGAGACAGGCAGGGCCCCCCGGCAGCTGGCCCCGAGGAGGCGCCCGGAGTGGGGCCGGTCGGCTGGGCT | 12 |
| wt-18600-18720 | 155 | GGCCGAGCCCGGGTCTGGGAGGTCTGGGGTGGCGAGCCTGCTGTCTCAGGAGGGGCCTGGCTCCGCCGGGTGGCCCTGGGGTAAGTCTGGGAGGCAGAGGGTCGGCCTAGGCCCGGGGAA | 8 |

| | | | |
|---|---|---|---|
| wt-18720-18840 | 156 | GTGGAGGGGGATCGCCCGGGTCTCTGTTGGCAGAGTCCGGGCGATCCTCTGAGACCCTCCGGGCCCGGACGGTCGCCCTCAGCCCCCCAGACAGACCCCAGGGTCTCCAGGCAGGGTCCG | 8 |
| wt-18840-18960 | 157 | GCATCTTCAGGGGCAGCAGGCTCACCACCACAGGCCCCCCAGACCCGGGTCTCGGCCAGCCGAGCCGACCGGCCCCGCGCCTGGCGCCTCCTCGGGGCCAGCCGCCGGGGTTGGTTCTGC | 12 |
| wt-18960-19080 | 158 | CCCTCTCTCTGTCCTTCAGAGGAACCAGGGACCTCGGGCACCCCAGAGCCCCTCGGGCCCGCCTCCAGGCGCCCTCCTGGTCTCCGCTCCCCTCTGAGCCCCGTTAAACCCAAAGAATGT | 8 |
| wt-19080-19200 | 159 | CTGAGGGGAGCCACCCTCGGGGCCCAGGCCCCAGAGTCCAGAGGTCAGGGGCACCTCAGGGTGCCTCCCCGGGTCCCAGGCCAGCCGGAGGGACCCCGGCAGCCCGGGCGGCCCCAGAGG | 12 |
| wt-19200-19320 | 160 | CCGGTTCCTCGCCCCTTCCCCGGGCTTCAGAGCCCAGGATGTCCCCCAGAAGGGACCCTAGGCGTCCCCTCTCCTCCCCTCCAGGCCCGAGCCTCTCCCTCGCGGAGAGGGGCCTCTTTG | 8 |
| wt-19320-19440 | 161 | GGCCCTCAAGTCCAGCCCCACCGAGACCCGAGTGGCCCGGATCCCCCCACCGGCCCTTCTCTCTGTCCCCCTGCTCCTCTCCAACCTTCGCTCCACCCTAGACCCCAGCTTCTGGCCTCC | 8 |
| wt-19440-19560 | 162 | CCGGGTCCACCAGGCCAGCCGGAGGGACCCCGGCAGCCCGGGCGAGTCGCCTTCCCTCTCCCCTGGCCTCTCCTTCCCGCCTCCCACCCGAGCCCCCTCAGCTTGCCTCCCCACCGGGTC | 12 |
| wt-19560-19680 | 163 | CATCAGGCCGGCCGGAGGGACCCCGGCGGCCCGGTGTCAGTCCCCCCTGCAGCCGCCCAGTCTCTGCCTCCAGGCAAGGGCGCCAGCTTTTCTCCCCCCAGCCTGAGGCCCAGTCTCCTG | 8 |
| wt-19680-19800 | 164 | TGCACTGTCTGTAAAGTCCAGCCTCCCACGCCCGTCCACGGCTCCCGGGCCCAGCCTCGTCCACCCCTCCCCACGGTGGACAGGCCCTCTGTCCACCCGGGCCATCCCCGCCCCCCTGTG | 8 |
| wt-19920-20040 | 166 | CCAGCCCGACCCCCGGGCGCCCCCAAACTTTGTCCAGATGTCCAGGGGTCCCCGAGGGTGAGGCCCAGCCCCCTCCCGCCCCTGTCCACTGCCCCGGTCCCCCCAGAAGCCCCCAAAAGT | 8 |
| wt-20040-20160 | 167 | AGAGGCTCAGGCCATGCGCGCCCTGTCACCAGGCCTGCCAAAGAGCCAGATCTAAGGCCGGGAGAGGCAGCCCCAAAGCGGGTGCAGTAACAGGTAATCTCTGGTAGTGATTTGGACCCG | 4 |
| wt-20160-20280 | 168 | AAATCTGACACTTTAGAGCTCTGGAGGACTTTAAAACTCTAAAAATCAAAACTTTAGAGGCGAATGGGCGCCATTTTGTCCCCACGCGCGCATAATGGCGGACCTAGGCCTAAAACCCCC | 4 |
| wt-20280-20400 | 169 | AGGAAGCGGGTCTATGGTTGGCTGCGCTGCTGCTATCTTTAGAGGGGAAAAGAGGAATAAGCCCCCAGACAGGGGAGTGGGCTTGTTTGTGACTTCACCAAAGGTCAGGGCCCAAGGGGG | 4 |
| wt-20400-20520 | 170 | TTCGCGTTGCTAGGCCACCTTCTCAGTCCAGCGCGTTTACGTAAGCCAGACAGCAGCCAATTGTCAGTTCTAGGGAGGGGGACCACTGCCCCTGGTATAAAGTGGTCCTGCAGCTATTTC | 4 |
| wt-20520-20640 | 171 | TGGTCGCATCAGAGCGCCAGGAGTCCACACAAATGTAAGAGGGGGTCTTCTACCTCTCCCTAGCCCTCCGCCCCCTCCAAGGACTCGGGCCCAGTTTCTAACTTTTCCCCTTCCCTCCCT | 4 |
| wt-20640-20760 | 172 | CGTCTTGCCCTGCGCCCGGGGCCACCTTCATCACCGTCGCTGACTCCGCCATCCAAGCCTAGGGGAGACCGAAGTGAAGGCCCTGGACCAACCCGGCCCGGGCCCCCCGGTATCGGGCCA | 8 |
| wt-20760-20880 | 173 | GAGGTAAGTGGACTTTAATTTTTTCTGCTAAGCCCAACACTCCACCACACCCAGGCACACACTACACACACCCACCCGTCTCAGGGTCCCCTCGGACAGCTCCTAAGAAGGCACCGGTCG | 4 |
| wt-20880-21000 | 174 | CCCAGTCCTACCAGAGGGGGCCAAGAACCCAGACGAGTCCGTAGAAGGGTCCTCGTCCAGCAAGAAGAGGAGGTGGTAAGCGGTTCACCTTCAGGGGTAAGTAACCTGACCTCTCCAGGG | 4 |
| wt-21000-21120 | 175 | CTCACATAAAGGGAGGCTTAGTATACATGCTTCTTGCTTTTCACAGGAACCTGGGGGCTAGTCTGGGTGGGATTAGGCTGCCTCAAGTTGCATCAGCCAGGGCTTCATGCCCTCCTCAGT | 4 |
| wt-21120-21240 | 176 | TCCCTAGTCCCCGGGCTTCAGGCCCCCTCCGTCCCCGTCCTCCAGAGACCCGGGCTTCAGGCCCTGCCTCTCCTGTTACCCTTTTAGAACCACAGCCTGGACACATGTGCCAGACGCCTT | 4 |
| wt-21240-21360 | 177 | GGCCTCTAAGGCCCTCGGGTCCCCCTGGACCCCGGCCTCAGCAACCCTGCTGCTCCCCTCCTGCCACCCCAGCCTCCCCCCCTCCCCGTCCCCCTTCGCTCCTGATCCTCCCCCGGTCCC | 8 |
| wt-21360-21480 | 178 | CAGTAGGGCCGCCTGCCCCCCTGCACCCAGTACCTGCCCCTCTTGGCCACGCACCCCGGGCCAGGCCACCTTAGACCCGGCCAAGCCCCATCCCTGAAGACCCAGCGGCCATTCTCTCTG | 8 |
| wt-21480-21600 | 179 | GTAACGAGCAGAGAAGAAGTAGAGGCCCGCGGCCATTGGGCCCAGATTGAGAGACCAGTCCAGGGGCCCGAGGTTGGAGCCAGCGGGCACCCGAGGTCCCAGCACCCGGTCCCTCCGGGG | 8 |
| wt-21600-21720 | 180 | GGCAGAGACAGGCAGGGCCCCCCGGCAGCTGGCCCCGAGGAGGCGCCCGGAGTGGGGCCGGTCGGCTGGGCTGGCCGAGCCCGGGTCTGGGAGGTCTGGGGTGGCGAGCCTGCTGTCTCA | 12 |
| wt-21720-21840 | 181 | GGAGGGGCCTGGCTCCGCCGGGTGGCCCTGGGGTAAGTCTGGGAGGCAGAGGGTCGGCCTAGGCCCGGGGAAGTGGAGGGGGATCGCCCGGGTCTCTGTTGGCAGAGTCCGGGCGATCCT | 8 |
| wt-21840-21960 | 182 | CTGAGACCCTCCGGGCCCGGACGGTCGCCCTCAGCCCCCCAGACAGACCCCAGGGTCTCCAGGCAGGGTCCGGCATCTTCAGGGGCAGCAGGCTCACCACCACAGGCCCCCCAGACCCGG | 8 |
| wt-21960-22080 | 183 | GTCTCGGCCAGCCGAGCCGACCGGCCCCGCGCCTGGCGCCTCCTCGGGGCCAGCCGCCGGGGTTGGTTCTGCCCCTCTCTCTGTCCTTCAGAGGAACCAGGGACCTCGGGCACCCCAGAG | 8 |
| wt-22080-22200 | 184 | CCCCTCGGGCCCGCCTCCAGGCGCCCTCCTGGTCTCCGCTCCCCTCTGAGCCCCGTTAAACCCAAAGAATGTCTGAGGGGAGCCACCCTCGGGGCCCAGGCCCCAGAGTCCAGAGGTCAG | 8 |
| wt-22200-22320 | 185 | GGGCACCTCAGGGTGCCTCCCCGGGTCCCAGGCCAGCCGGAGGGACCCCGGCAGCCCGGGCGGCCCCAGAGGCCGGTTCCTCGCCCCTTCCCCGGGCTTCAGAGCCCAGGATGTCCCCCA | 12 |
| wt-22320-22440 | 186 | GAAGGGACCCTAGGCGTCCCCTCTCCTCCCCTCCAGGCCCGAGCCTCTCCCTCGCGGAGAGGGGCCTCTTTGGGCCCTCAAGTCCAGCCCCACCGAGACCCGAGTGGCCCGGATCCCCCC | 8 |
| wt-22440-22560 | 187 | ACCGGCCCTTCTCTCTGTCCCCCTGCTCCTCTCCAACCTTCGCTCCACCCTAGACCCCAGCTTCTGGCCTCCCCGGGTCCACCAGGCCAGCCGGAGGGACCCCGGCAGCCCGGGCGAGTC | 8 |
| wt-22560-22680 | 188 | GCCTTCCCTCTCCCCTGGCCTCTCCTTCCCGCCTCCCACCCGAGCCCCCTCAGCTTGCCTCCCCACCGGGTCCATCAGGCCGGCCGGAGGGACCCCGGCGGCCCGGTGTCAGTCCCCCCT | 12 |
| wt-22680-22800 | 189 | GCAGCCGCCCAGTCTCTGCCTCCAGGCAAGGGCGCCAGCTTTTCTCCCCCCAGCCTGAGGCCCAGTCTCCTGTGCACTGTCTGTAAAGTCCAGCCTCCCACGCCCGTCCACGGCTCCCGG | 8 |
| wt-22800-22920 | 190 | GCCCAGCCTCGTCCACCCCTCCCCACGGTGGACAGGCCCTCTGTCCACCCGGGCCATCCCCGCCCCCCTGTGTCCACCCCAGTCCCGTCCAGGGGGGACTTTATGTGACCCTTGGGCCTG | 8 |
| wt-22920-23040 | 191 | GCTCCCCATAGACTCCCATGTAAGCCTGCCTCGAGTAGGTGCCTCCAGAGCCCCTTTTGCCCCCCTGGCGGCCCAGCCCGACCCCCGGGCGCCCCCAAACTTTGTCCAGATGTCCAGGGG | 8 |
| wt-23040-23160 | 192 | TCCCCGAGGGTGAGGCCCAGCCCCCTCCCGCCCCTGTCCACTGCCCCGGTCCCCCCAGAAGCCCCCAAAAGTAGAGGCTCAGGCCATGCGCGCCCTGTCACCAGGCCTGCCAAAGAGCCA | 8 |
| wt-23160-23280 | 193 | GATCTAAGGCCGGGAGAGGCAGCCCCAAAGCGGGTGCAGTAACAGGTAATCTCTGGTAGTGATTTGGACCCGAAATCTGACACTTTAGAGCTCTGGAGGACTTTAAAACTCTAAAAATCA | 4 |
| wt-23280-23400 | 194 | AAACTTTAGAGGCGAATGGGCGCCATTTTGTCCCCACGCGCGCATAATGGCGGACCTAGGCCTAAAACCCCCAGGAAGCGGGTCTATGGTTGGCTGCGCTGCTGCTATCTTTAGAGGGGA | 4 |
| wt-23400-23520 | 195 | AAAGAGGAATAAGCCCCCAGACAGGGGAGTGGGCTTGTTTGTGACTTCACCAAAGGTCAGGGCCCAAGGGGGTTCGCGTTGCTAGGCCACCTTCTCAGTCCAGCGCGTTTACGTAAGCCA | 4 |
| wt-23520-23640 | 196 | GACAGCAGCCAATTGTCAGTTCTAGGGAGGGGGACCACTGCCCCTGGTATAAAGTGGTCCTGCAGCTATTTCTGGTCGCATCAGAGCGCCAGGAGTCCACACAAATGTAAGAGGGGGTCT | 4 |
| wt-23760-23880 | 198 | CCATCCAAGCCTAGGGGAGACCGAAGTGAAGGCCCTGGACCAACCCGGCCCGGGCCCCCCGGTATCGGGCCAGAGGTAAGTGGACTTTAATTTTTTCTGCTAAGCCCAACACTCCACCAC | 4 |
| wt-23880-24000 | 199 | ACCCAGGCACACACTACACACACCCACCCGTCTCAGGGTCCCCTCGGACAGCTCCTAAGAAGGCACCGGTCGCCCAGTCCTACCAGAGGGGGCCAAGAACCCAGACGAGTCCGTAGAAGG | 4 |
| wt-24000-24120 | 200 | GTCCTCGTCCAGCAAGAAGAGGAGGTGGTAAGCGGTTCACCTTCAGGGGTAAGTAACCTGACCTCTCCAGGGCTCACATAAAGGGAGGCTTAGTATACATGCTTCTTGCTTTTCACAGGA | 4 |
| wt-24120-24240 | 201 | ACCTGGGGGCTAGTCTGGGTGGGATTAGGCTGCCTCAAGTTGCATCAGCCAGGGCTTCATGCCCTCCTCAGTTCCCTAGTCCCCGGGCTTCAGGCCCCCTCCGTCCCCGTCCTCCAGAGA | 4 |
| wt-24360-24480 | 203 | GCTGCTCCCCTCCTGCCACCCCAGCCTCCCCCCCTCCCCGTCCCCCTTCGCTCCTGATCCTCCCCCGGTCCCCAGTAGGGCCGCCTGCCCCCCTGCACCCAGTACCTGCCCCTCTTGGCC | 8 |
| wt-24480-24600 | 204 | ACGCACCCCGGGCCAGGCCACCTTAGACCCGGCCAAGCCCCATCCCTGAAGACCCAGCGGCCATTCTCTCTGGTAACGAGCAGAGAAGAAGTAGAGGCCCGCGGCCATTGGGCCCAGATT | 4 |
| wt-24600-24720 | 205 | GAGAGACCAGTCCAGGGGCCCGAGGTTGGAGCCAGCGGGCACCCGAGGTCCCAGCACCCGGTCCCTCCGGGGGGCAGAGACAGGCAGGGCCCCCCGGCAGCTGGCCCCGAGGAGGCGCCC | 12 |
| wt-24720-24840 | 206 | GGAGTGGGGCCGGTCGGCTGGGCTGGCCGAGCCCGGGTCTGGGAGGTCTGGGGTGGCGAGCCTGCTGTCTCAGGAGGGGCCTGGCTCCGCCGGGTGGCCCTGGGGTAAGTCTGGGAGGCA | 8 |
| wt-24840-24960 | 207 | GAGGGTCGGCCTAGGCCCGGGGAAGTGGAGGGGGATCGCCCGGGTCTCTGTTGGCAGAGTCCGGGCGATCCTCTGAGACCCTCCGGGCCCGGACGGTCGCCCTCAGCCCCCCAGACAGAC | 8 |
| wt-24960-25080 | 208 | CCCAGGGTCTCCAGGCAGGGTCCGGCATCTTCAGGGGCAGCAGGCTCACCACCACAGGCCCCCCAGACCCGGGTCTCGGCCAGCCGAGCCGACCGGCCCCGCGCCTGGCGCCTCCTCGGG | 12 |
| wt-25080-25200 | 209 | GCCAGCCGCCGGGGTTGGTTCTGCCCCTCTCTCTGTCCTTCAGAGGAACCAGGGACCTCGGGCACCCCAGAGCCCCTCGGGCCCGCCTCCAGGCGCCCTCCTGGTCTCCGCTCCCCTCTG | 8 |

| | | | |
|---|---|---|---|
| wt-25200-25320 | 210 | AGCCCCGTTAAACCCAAAGAATGTCTGAGGGGAGCCACCCTCGGGGCCCAGGCCCCAGAGTCCAGAGGTCAGGGGCACCTCAGGGTGCCTCCCCGGGTCCCAGGCCAGCCGGAGGGACCC | 8 |
| wt-25320-25440 | 211 | CGGCAGCCCGGGCGGCCCCAGAGGCCGGTTCCTCGCCCCTTCCCCGGGCTTCAGAGCCCAGGATGTCCCCCAGAAGGGACCCTAGGCGTCCCCTCTCCTCCCCTCCAGGCCCGAGCCTCT | 8 |
| wt-25440-25560 | 212 | CCCTCGCGGAGAGGGGCCTCTTTGGGCCCTCAAGTCCAGCCCCACCGAGACCCGAGTGGCCCGGATCCCCCCACCGGCCCTTCTCTCTGTCCCCCTGCTCCTCTCCAACCTTCGCTCCAC | 8 |
| wt-25560-25680 | 213 | CCTAGACCCCAGCTTCTGGCCTCCCCGGGTCCACCAGGCCAGCCGGAGGGACCCCGGCAGCCCGGGCGAGTCGCCTTCCCTCTCCCCTGGCCTCTCCTTCCCGCCTCCCACCCGAGCCCC | 12 |
| wt-25680-25800 | 214 | CTCAGCTTGCCTCCCCACCGGGTCCATCAGGCCGGCCGGAGGGACCCCGGCGGCCCGGTGTCAGTCCCCCCTGCAGCCGCCCAGTCTCTGCCTCCAGGCAAGGGCGCCAGCTTTTCTCCC | 8 |
| wt-25800-25920 | 215 | CCCAGCCTGAGGCCCAGTCTCCTGTGCACTGTCTGTAAAGTCCAGCCTCCCACGCCCGTCCACGGCTCCCGGGCCCAGCCTCGTCCACCCCTCCCCACGGTGGACAGGCCCTCTGTCCAC | 8 |
| wt-25920-26040 | 216 | CCGGGCCATCCCCGCCCCCCTGTGTCCACCCCAGTCCCGTCCAGGGGGGACTTTATGTGACCCTTGGGCCTGGCTCCCCATAGACTCCCATGTAAGCCTGCCTCGAGTAGGTGCCTCCAG | 8 |
| wt-26040-26160 | 217 | AGCCCCTTTTGCCCCCCTGGCGGCCCAGCCCGACCCCCGGGCGCCCCCAAACTTTGTCCAGATGTCCAGGGGTCCCCGAGGGTGAGGCCCAGCCCCCTCCCGCCCCTGTCCACTGCCCCG | 8 |
| wt-26160-26280 | 218 | GTCCCCCCAGAAGCCCCCAAAAGTAGAGGCTCAGGCCATGCGCGCCCTGTCACCAGGCCTGCCAAAGAGCCAGATCTAAGGCCGGGAGAGGCAGCCCCAAAGCGGGTGCAGTAACAGGTA | 4 |
| wt-26280-26400 | 219 | ATCTCTGGTAGTGATTTGGACCCGAAATCTGACACTTTAGAGCTCTGGAGGACTTTAAAACTCTAAAAATCAAAACTTTAGAGGCGAATGGGCGCCATTTTGTCCCCACGCGCGCATAAT | 4 |
| wt-26400-26520 | 220 | GGCGGACCTAGGCCTAAAACCCCCAGGAAGCGGGTCTATGGTTGGCTGCGCTGCTGCTATCTTTAGAGGGGAAAAGAGGAATAAGCCCCCAGACAGGGGAGTGGGCTTGTTTGTGACTTC | 4 |
| wt-26520-26640 | 221 | ACCAAAGGTCAGGGCCCAAGGGGGTTCGCGTTGCTAGGCCACCTTCTCAGTCCAGCGCGTTTACGTAAGCCAGACAGCAGCCAATTGTCAGTTCTAGGGAGGGGGACCACTGCCCCTGGT | 4 |
| wt-26640-26760 | 222 | ATAAAGTGGTCCTGCAGCTATTTCTGGTCGCATCAGAGCGCCAGGAGTCCACACAAATGTAAGAGGGGGTCTTCTACCTCTCCCTAGCCCTCCGCCCCCTCCAAGGACTCGGGCCCAGTT | 4 |
| wt-26760-26880 | 223 | TCTAACTTTTCCCCTTCCCTCCCTCGTCTTGCCCTGCGCCCGGGGCCACCTTCATCACCGTCGCTGACTCCGCCATCCAAGCCTAGGGGAGACCGAAGTGAAGGCCCTGGACCAACCCGG | 4 |
| wt-26880-27000 | 224 | CCCGGGCCCCCCGGTATCGGGCCAGAGGTAAGTGGACTTTAATTTTTTCTGCTAAGCCCAACACTCCACCACACCCAGGCACACACTACACACACCCACCCGTCTCAGGGTCCCCTCGGA | 4 |
| wt-27000-27120 | 225 | CAGCTCCTAAGAAGGCACCGGTCGCCCAGTCCTACCAGAGGGGGCCAAGAACCCAGACGAGTCCGTAGAAGGGTCCTCGTCCAGCAAGAAGAGGAGGTGGTAAGCGGTTCACCTTCAGGG | 4 |
| wt-27120-27240 | 226 | GTAAGTAACCTGACCTCTCCAGGGCTCACATAAAGGGAGGCTTAGTATACATGCTTCTTGCTTTTCACAGGAACCTGGGGGCTAGTCTGGGTGGGATTAGGCTGCCTCAAGTTGCATCAG | 4 |
| wt-27360-27480 | 228 | CTGGACACATGTGCCAGACGCCTTGGCCTCTAAGGCCCTCGGGTCCCCCTGGACCCCGGCCTCAGCAACCCTGCTGCTCCCCTCCTGCCACCCCAGCCTCCCCCCCTCCCCGTCCCCCTT | 8 |
| wt-27480-27600 | 229 | CGCTCCTGATCCTCCCCCGGTCCCCAGTAGGGCCGCCTGCCCCCCTGCACCCAGTACCTGCCCCTCTTGGCCACGCACCCCGGGCCAGGCCACCTTAGACCCGGCCAAGCCCCATCCCTG | 8 |
| wt-27600-27720 | 230 | AAGACCCAGCGGCCATTCTCTCTGGTAACGAGCAGAGAAGAAGTAGAGGCCCGCGGCCATTGGGCCCAGATTGAGAGACCAGTCCAGGGGCCCGAGGTTGGAGCCAGCGGGCACCCGAGG | 4 |
| wt-27720-27840 | 231 | TCCCAGCACCCGGTCCCTCCGGGGGGCAGAGACAGGCAGGGCCCCCCGGCAGCTGGCCCCGAGGAGGCGCCCGGAGTGGGGCCGGTCGGCTGGGCTGGCCGAGCCCGGGTCTGGGAGGTC | 12 |
| wt-27840-27960 | 232 | TGGGGTGGCGAGCCTGCTGTCTCAGGAGGGGCCTGGCTCCGCCGGGTGGCCCTGGGGTAAGTCTGGGAGGCAGAGGGTCGGCCTAGGCCCGGGGAAGTGGAGGGGGATCGCCCGGGTCTC | 8 |
| wt-27960-28080 | 233 | TGTTGGCAGAGTCCGGGCGATCCTCTGAGACCCTCCGGGCCCGGACGGTCGCCCTCAGCCCCCCAGACAGACCCCAGGGTCTCCAGGCAGGGTCCGGCATCTTCAGGGGCAGCAGGCTCA | 8 |
| wt-28080-28200 | 234 | CCACCACAGGCCCCCCAGACCCGGGTCTCGGCCAGCCGAGCCGACCGGCCCCGCGCCTGGCGCCTCCTCGGGGCCAGCCGCCGGGGTTGGTTCTGCCCCTCTCTCTGTCCTTCAGAGGAA | 8 |
| wt-28200-28320 | 235 | CCAGGGACCTCGGGCACCCCAGAGCCCCTCGGGCCCGCCTCCAGGCGCCCTCCTGGTCTCCGCTCCCCTCTGAGCCCCGTTAAACCCAAAGAATGTCTGAGGGGAGCCACCCTCGGGGCC | 8 |
| wt-28320-28440 | 236 | CAGGCCCCAGAGTCCAGAGGTCAGGGGCACCTCAGGGTGCCTCCCCGGGTCCCAGGCCAGCCGGAGGGACCCCGGCAGCCCGGGCGGCCCCAGAGGCCGGTTCCTCGCCCCTTCCCCGGG | 12 |
| wt-28440-28560 | 237 | CTTCAGAGCCCAGGATGTCCCCCAGAAGGGACCCTAGGCGTCCCCTCTCCTCCCCTCCAGGCCCGAGCCTCTCCCTCGCGGAGAGGGGCCTCTTTGGGCCCTCAAGTCCAGCCCCACCGA | 8 |
| wt-28560-28680 | 238 | GACCCGAGTGGCCCGGATCCCCCCACCGGCCCTTCTCTCTGTCCCCCTGCTCCTCTCCAACCTTCGCTCCACCCTAGACCCCAGCTTCTGGCCTCCCCGGGTCCACCAGGCCAGCCGGAG | 8 |
| wt-28680-28800 | 239 | GGACCCCGGCAGCCCGGGCGAGTCGCCTTCCCTCTCCCCTGGCCTCTCCTTCCCGCCTCCCACCCGAGCCCCCTCAGCTTGCCTCCCCACCGGGTCCATCAGGCCGGCCGGAGGGACCCC | 12 |
| wt-28800-28920 | 240 | GGCGGCCCGGTGTCAGTCCCCCCTGCAGCCGCCCAGTCTCTGCCTCCAGGCAAGGGCGCCAGCTTTTCTCCCCCCAGCCTGAGGCCCAGTCTCCTGTGCACTGTCTGTAAAGTCCAGCCT | 8 |
| wt-28920-29040 | 241 | CCCACGCCCGTCCACGGCTCCCGGGCCCAGCCTCGTCCACCCCTCCCCACGGTGGACAGGCCCTCTGTCCACCCGGGCCATCCCCGCCCCCCTGTGTCCACCCCAGTCCCGTCCAGGGGG | 12 |
| wt-29040-29160 | 242 | GACTTTATGTGACCCTTGGGCCTGGCTCCCCATAGACTCCCATGTAAGCCTGCCTCGAGTAGGTGCCTCCAGAGCCCCTTTTGCCCCCCTGGCGGCCCAGCCCGACCCCCGGGCGCCCCC | 8 |
| wt-29160-29280 | 243 | AAACTTTGTCCAGATGTCCAGGGGTCCCCGAGGGTGAGGCCCAGCCCCCTCCCGCCCCTGTCCACTGCCCCGGTCCCCCCAGAAGCCCCCAAAAGTAGAGGCTCAGGCCATGCGCGCCCT | 8 |
| wt-29280-29400 | 244 | GTCACCAGGCCTGCCAAAGAGCCAGATCTAAGGCCGGGAGAGGCAGCCCCAAAGCGGGTGCAGTAACAGGTAATCTCTGGTAGTGATTTGGACCCGAAATCTGACACTTTAGAGCTCTGG | 4 |
| wt-29400-29520 | 245 | AGGACTTTAAAACTCTAAAAATCAAAACTTTAGAGGCGAATGGGCGCCATTTTGTCCCCACGCGCGCATAATGGCGGACCTAGGCCTAAAACCCCCAGGAAGCGGGTCTATGGTTGGCTG | 4 |
| wt-29520-29640 | 246 | CGCTGCTGCTATCTTTAGAGGGGAAAAGAGGAATAAGCCCCCAGACAGGGGAGTGGGCTTGTTTGTGACTTCACCAAAGGTCAGGGCCCAAGGGGGTTCGCGTTGCTAGGCCACCTTCTC | 4 |
| wt-29640-29760 | 247 | AGTCCAGCGCGTTTACGTAAGCCAGACAGCAGCCAATTGTCAGTTCTAGGGAGGGGGACCACTGCCCCTGGTATAAAGTGGTCCTGCAGCTATTTCTGGTCGCATCAGAGCGCCAGGAGT | 4 |
| wt-29760-29880 | 248 | CCACACAAATGTAAGAGGGGGTCTTCTACCTCTCCCTAGCCCTCCGCCCCCTCCAAGGACTCGGGCCCAGTTTCTAACTTTTCCCCTTCCCTCCCTCGTCTTGCCCTGCGCCCGGGGCCA | 4 |
| wt-29880-30000 | 249 | CCTTCATCACCGTCGCTGACTCCGCCATCCAAGCCTAGGGGAGACCGAAGTGAAGGCCCTGGACCAACCCGGCCCGGGCCCCCCGGTATCGGGCCAGAGGTAAGTGGACTTTAATTTTTT | 4 |
| wt-30000-30120 | 250 | CTGCTAAGCCCAACACTCCACCACACCCAGGCACACACTACACACACCCACCCGTCTCAGGGTCCCCTCGGACAGCTCCTAAGAAGGCACCGGTCGCCCAGTCCTACCAGAGGGGGCCAA | 4 |
| wt-30120-30240 | 251 | GAACCCAGACGAGTCCGTAGAAGGGTCCTCGTCCAGCAAGAAGAGGAGGTGGTAAGCGGTTCACCTTCAGGGGTAAGTAACCTGACCTCTCCAGGGCTCACATAAAGGGAGGCTTAGTAT | 4 |
| wt-30240-30360 | 252 | ACATGCTTCTTGCTTTTCACAGGAACCTGGGGGCTAGTCTGGGTGGGATTAGGCTGCCTCAAGTTGCATCAGCCAGGGCTTCATGCCCTCCTCAGTTCCCTAGTCCCCGGGCTTCAGGCC | 4 |
| wt-30480-30600 | 254 | CTGGACCCCGGCCTCAGCAACCCTGCTGCTCCCCTCCTGCCACCCCAGCCTCCCCCCCTCCCCGTCCCCCTTCGCTCCTGATCCTCCCCCGGTCCCCAGTAGGGCCGCCTGCCCCCCTGC | 12 |
| wt-30600-30720 | 255 | ACCCAGTACCTGCCCCTCTTGGCCACGCACCCCGGGCCAGGCCACCTTAGACCCGGCCAAGCCCCATCCCTGAAGACCCAGCGGCCATTCTCTCTGGTAACGAGCAGAGAAGAAGTAGAG | 4 |
| wt-30720-30840 | 256 | GCCCGCGGCCATTGGGCCCAGATTGAGAGACCAGTCCAGGGGCCCGAGGTTGGAGCCAGCGGGCACCCGAGGTCCCAGCACCCGGTCCCTCCGGGGGGCAGAGACAGGCAGGGCCCCCCG | 8 |
| wt-30840-30960 | 257 | GCAGCTGGCCCCGAGGAGGCGCCCGGAGTGGGGCCGGTCGGCTGGGCTGGCCGAGCCCGGGTCTGGGAGGTCTGGGGTGGCGAGCCTGCTGTCTCAGGAGGGGCCTGGCTCCGCCGGGTG | 12 |
| wt-30960-31080 | 258 | GCCCTGGGGTAAGTCTGGGAGGCAGAGGGTCGGCCTAGGCCCGGGGAAGTGGAGGGGGATCGCCCGGGTCTCTGTTGGCAGAGTCCGGGCGATCCTCTGAGACCCTCCGGGCCCGGACGG | 8 |
| wt-31080-31200 | 259 | TCGCCCTCAGCCCCCCAGACAGACCCCAGGGTCTCCAGGCAGGGTCCGGCATCTTCAGGGGCAGCAGGCTCACCACCACAGGCCCCCCAGACCCGGGTCTCGGCCAGCCGAGCCGACCGG | 8 |
| wt-31200-31320 | 260 | CCCCGCGCCTGGCGCCTCCTCGGGGCCAGCCGCCGGGGTTGGTTCTGCCCCTCTCTCTGTCCTTCAGAGGAACCAGGGACCTCGGGCACCCCAGAGCCCCTCGGGCCCGCCTCCAGGCGC | 12 |
| wt-31320-31440 | 261 | CCTCCTGGTCTCCGCTCCCCTCTGAGCCCCGTTAAACCCAAAGAATGTCTGAGGGGAGCCACCCTCGGGGCCCAGGCCCCAGAGTCCAGAGGTCAGGGGCACCTCAGGGTGCCTCCCCGG | 8 |
| wt-31440-31560 | 262 | GTCCCAGGCCAGCCGGAGGGACCCCGGCAGCCCGGGCGGCCCCAGAGGCCGGTTCCTCGCCCCTTCCCCGGGCTTCAGAGCCCAGGATGTCCCCCAGAAGGGACCCTAGGCGTCCCCTCT | 8 |

| | | | |
|---|---|---|---|
| wt-31560-31680 | 263 | CCTCCCCTCCAGGCCCGAGCCTCTCCCTCGCGGAGAGGGGCCTCTTTGGGCCCTCAAGTCCAGCCCCACCGAGACCCGAGTGGCCCGGATCCCCCCACCGGCCCTTCTCTCTGTCCCCCT | 8 |
| wt-31680-31800 | 264 | GCTCCTCTCCAACCTTCGCTCCACCCTAGACCCCAGCTTCTGGCCTCCCCGGGTCCACCAGGCCAGCCGGAGGGACCCCGGCAGCCCGGGCGAGTCGCCTTCCCTCTCCCCTGGCCTCTC | 8 |
| wt-31800-31920 | 265 | CTTCCCGCCTCCCACCCGAGCCCCCTCAGCTTGCCTCCCCACCGGGTCCATCAGGCCGGCCGGAGGGACCCCGGCGGCCCGGTGTCAGTCCCCCCTGCAGCCGCCCAGTCTCTGCCTCCA | 12 |
| wt-31920-32040 | 266 | GGCAAGGGCGCCAGCTTTTCTCCCCCCAGCCTGAGGCCCAGTCTCCTGTGCACTGTCTGTAAAGTCCAGCCTCCCACGCCCGTCCACGGCTCCCGGGCCCAGCCTCGTCCACCCCTCCCC | 8 |
| wt-32040-32160 | 267 | ACGGTGGACAGGCCCTCTGTCCACCCGGGCCATCCCCGCCCCCCTGTGTCCACCCCAGTCCCGTCCAGGGGGGACTTTATGTGACCCTTGGGCCTGGCTCCCCATAGACTCCCATGTAAG | 8 |
| wt-32160-32280 | 268 | CCTGCCTCGAGTAGGTGCCTCCAGAGCCCCTTTTGCCCCCCTGGCGGCCCAGCCCGACCCCCGGGCGCCCCCAAACTTTGTCCAGATGTCCAGGGGTCCCCGAGGGTGAGGCCCAGCCCC | 8 |
| wt-32280-32400 | 269 | CTCCCGCCCCTGTCCACTGCCCCGGTCCCCCCAGAAGCCCCCAAAAGTAGAGGCTCAGGCCATGCGCGCCCTGTCACCAGGCCTGCCAAAGAGCCAGATCTAAGGCCGGGAGAGGCAGCC | 8 |
| wt-32400-32520 | 270 | CCAAAGCGGGTGCAGTAACAGGTAATCTCTGGTAGTGATTTGGACCCGAAATCTGACACTTTAGAGCTCTGGAGGACTTTAAAACTCTAAAAATCAAAACTTTAGAGGCGAATGGGCGCC | 4 |
| wt-32520-32640 | 271 | ATTTTGTCCCCACGCGCGCATAATGGCGGACCTAGGCCTAAAACCCCCAGGAAGCGGGTCTATGGTTGGCTGCGCTGCTGCTATCTTTAGAGGGGAAAAGAGGAATAAGCCCCCAGACAG | 4 |
| wt-32640-32760 | 272 | GGGAGTGGGCTTGTTTGTGACTTCACCAAAGGTCAGGGCCCAAGGGGGTTCGCGTTGCTAGGCCACCTTCTCAGTCCAGCGCGTTTACGTAAGCCAGACAGCAGCCAATTGTCAGTTCTA | 4 |
| wt-32760-32880 | 273 | GGGAGGGGGACCACTGCCCCTGGTATAAAGTGGTCCTGCAGCTATTTCTGGTCGCATCAGAGCGCCAGGAGTCCACACAAATGTAAGAGGGGGTCTTCTACCTCTCCCTAGCCCTCCGCC | 4 |
| wt-32880-33000 | 274 | CCCTCCAAGGACTCGGGCCCAGTTTCTAACTTTTCCCCTTCCCTCCCTCGTCTTGCCCTGCGCCCGGGGCCACCTTCATCACCGTCGCTGACTCCGCCATCCAAGCCTAGGGGAGACCGA | 4 |
| wt-33000-33120 | 275 | AGTGAAGGCCCTGGACCAACCCGGCCCGGGCCCCCCGGTATCGGGCCAGAGGTAAGTGGACTTTAATTTTTTCTGCTAAGCCCAACACTCCACCACACCCAGGCACACACTACACACACC | 4 |
| wt-33120-33240 | 276 | CACCCGTCTCAGGGTCCCCTCGGACAGCTCCTAAGAAGGCACCGGTCGCCCAGTCCTACCAGAGGGGGCCAAGAACCCAGACGAGTCCGTAGAAGGGTCCTCGTCCAGCAAGAAGAGGAG | 4 |
| wt-33240-33360 | 277 | GTGGTAAGCGGTTCACCTTCAGGGGTAAGTAACCTGACCTCTCCAGGGCTCACATAAAGGGAGGCTTAGTATACATGCTTCTTGCTTTTCACAGGAACCTGGGGGCTAGTCTGGGTGGGA | 4 |
| wt-33360-33480 | 278 | TTAGGCTGCCTCAAGTTGCATCAGCCAGGGCTTCATGCCCTCCTCAGTTCCCTAGTCCCCGGGCTTCAGGCCCCCTCCGTCCCCGTCCTCCAGAGACCCGGGCTTCAGGCCCTGCCTCTC | 8 |
| wt-33600-33720 | 280 | CCTCCCCCCCTCCCCGTCCCCCTTCGCTCCTGATCCTCCCCCGGTCCCCAGTAGGGCCGCCTGCCCCCCTGCACCCAGTACCTGCCCCTCTTGGCCACGCACCCCGGGCCAGGCCACCTT | 8 |
| wt-33720-33840 | 281 | AGACCCGGCCAAGCCCCATCCCTGAAGACCCAGCGGCCATTCTCTCTGGTAACGAGCAGAGAAGAAGTAGAGGCCCGCGGCCATTGGGCCCAGATTGAGAGACCAGTCCAGGGGCCCGAG | 4 |
| wt-33840-33960 | 282 | GTTGGAGCCAGCGGGCACCCGAGGTCCCAGCACCCGGTCCCTCCGGGGGGCAGAGACAGGCAGGGCCCCCCGGCAGCTGGCCCCGAGGAGGCGCCCGGAGTGGGGCCGGTCGGCTGGGCT | 12 |
| wt-33960-34080 | 283 | GGCCGAGCCCGGGTCTGGGAGGTCTGGGGTGGCGAGCCTGCTGTCTCAGGAGGGGCCTGGCTCCGCCGGGTGGCCCTGGGGTAAGTCTGGGAGGCAGAGGGTCGGCCTAGGCCCGGGGAA | 8 |
| wt-34080-34200 | 284 | GTGGAGGGGGATCGCCCGGGTCTCTGTTGGCAGAGTCCGGGCGATCCTCTGAGACCCTCCGGGCCCGGACGGTCGCCCTCAGCCCCCCAGACAGACCCCAGGGTCTCCAGGCAGGGTCCG | 8 |
| wt-34200-34320 | 285 | GCATCTTCAGGGGCAGCAGGCTCACCACCACAGGCCCCCCAGACCCGGGTCTCGGCCAGCCGAGCCGACCGGCCCCGCGCCTGGCGCCTCCTCGGGGCCAGCCGCCGGGGTTGGTTCTGC | 12 |
| wt-34320-34440 | 286 | CCCTCTCTCTGTCCTTCAGAGGAACCAGGGACCTCGGGCACCCCAGAGCCCCTCGGGCCCGCCTCCAGGCGCCCTCCTGGTCTCCGCTCCCCTCTGAGCCCCGTTAAACCCAAAGAATGT | 8 |
| wt-34440-34560 | 287 | CTGAGGGGAGCCACCCTCGGGGCCCAGGCCCCAGAGTCCAGAGGTCAGGGGCACCTCAGGGTGCCTCCCCGGGTCCCAGGCCAGCCGGAGGGACCCCGGCAGCCCGGGCGGCCCCAGAGG | 12 |
| wt-34560-34680 | 288 | CCGGTTCCTCGCCCCTTCCCCGGGCTTCAGAGCCCAGGATGTCCCCCAGAAGGGACCCTAGGCGTCCCCTCTCCTCCCCTCCAGGCCCGAGCCTCTCCCTCGCGGAGAGGGGCCTCTTTG | 8 |
| wt-34680-34800 | 289 | GGCCCTCAAGTCCAGCCCCACCGAGACCCGAGTGGCCCGGATCCCCCCACCGGCCCTTCTCTCTGTCCCCCTGCTCCTCTCCAACCTTCGCTCCACCCTAGACCCCAGCTTCTGGCCTCC | 8 |
| wt-34800-34920 | 290 | CCGGGTCCACCAGGCCAGCCGGAGGGACCCCGGCAGCCCGGGCGAGTCGCCTTCCCTCTCCCCTGGCCTCTCCTTCCCGCCTCCCACCCGAGCCCCCTCAGCTTGCCTCCCCACCGGGTC | 12 |
| wt-34920-35040 | 291 | CATCAGGCCGGCCGGAGGGACCCCGGCGGCCCGGTGTCAGTCCCCCCTGCAGCCGCCCAGTCTCTGCCTCCAGGCAAGGGCGCCAGCTTTTCTCCCCCCAGCCTGAGGCCCAGTCTCCTG | 8 |
| wt-35040-35160 | 292 | TGCACTGTCTGTAAAGTCCAGCCTCCCACGCCCGTCCACGGCTCCCGGGCCCAGCCTCGTCCACCCCTCCCCACGGTGGACAGGCCCTCTGTCCACCCGGGCCATCCCCGCCCCCCTGTG | 8 |
| wt-35280-35400 | 294 | CCAGCCCGACCCCCGGGCGCCCCCAAACTTTGTCCAGATGTCCAGGGGTCCCCGAGGGTGAGGCCCAGCCCCCTCTCGCCCAAGCTGCTTTGATTCCTGGGATATTTTTGGGAATGGTGT | 4 |
| wt-35400-35520 | 295 | TAACTTTCTCCCCTTGTATTTGCTATTCAATCAACCTGATTCCCCCTGCTCATACCTCCACTTACAACCAAGCCACTACGGCCACGTCCCCGGCCTCCCGCTCGGGTAAGTGCTTTTTCA | 4 |
| wt-35520-35640 | 296 | TTTTTAGCCCCAGCCCCTCCTCTATAAGTTCTAGGCAAACCTCCAATCACCAGCCACCTTCCAATGTAGTCTCTTAGAGAGTGGCTGCTACGCATTAGAGACCACTTTGAGCCACCCACA | 4 |
| wt-35640-35760 | 297 | GTAACCACCCAGCGCCAATCTGTCTACATAGAAGAAGAAGAGGATGAAGACTAAGTCACAGGCTTAGCCAGGTGATTTGTGAATTTCAGTTTATTTACTTTCTTCCAATCAAGCTTTCCC | 4 |
| wt-35760-35880 | 298 | AGCCTCCGCTTGTTAGGTCCTAGTTATGGGTTTTCCATGGGGGACTTAGTATCCGTTCTATTAGATTAACGTGCAAGACGCTAAACTTAACCAAGGTCAGCCAAGGGACGCGTGTTATCC | 4 |
| wt-35880-36000 | 299 | CAGGCTGCCCACCCTGAGGATTTCCCCCCAAAATCCTCCTACCCTCTCTTTATGCCATGTGTGTTGTTGGCTTGTGTTAGTGCTATGTAATGCGTTGCCGCCAGGTGGCAGCCTGTTTAT | 4 |
| wt-36000-36120 | 300 | AGATGTGCAGTACCCCTTAATGTTAGGTCTGCTTTAGGGCTGCCAGGTGGCGCAATCTAGGATTAATTCACCTGTATCCCTTTCCCTCCACCCGCAGTAACCCAGCACTGGCGTGTGACG | 4 |
| wt-36120-36240 | 301 | TGGTGTAAAGTTTTGCCTGAACCTGTGGTTGGGCAGGTACATGCCAACAACCTTCTAAGCACCCGCGCTTGTGTTTTGCTTTATCTGCCGCCATCATGCCTACATTCTATCTTGCGTTAC | 4 |
| wt-36240-36360 | 302 | ATGGGGGACAAACATATCATCTAATTGTTGACACGGATAGTCTTGGAAACCCGTCACTCTCAGTAATTCCCTCGAATCCCTACCAGGAACAACTGTCAGACACTCCATTAATTCCACTAA | 4 |
| wt-36600-36720 | 303 | CTGCTATGCGAATGCTTTGGATGGCTAATTATATTGTAAGACAATCACGGGGTGACCGGGGCCTTATTTTGCCACAAGGCCCACAAACAGCCCCTCAGGCCAGGTTGGTCCAGCCACATG | 4 |
| wt-36720-36840 | 304 | TCCCCCCTCTACGCCCGACAGCACCCACCATTTTGTCACCTCTGTCACAACCGAGGCTTACCCCTCCACAACCACTCATGATGCCACCAAGGCCTACCCCTCCTACCCCTCTGCCACCTG | 4 |
| wt-36840-36960 | 305 | CAACACTAACGGTGCCACCAAGGCCTACCCGTCCTACCACTCTGCCACCCACACCACTACTCACGGTACTACAAAGGCCTACCGAACTTCAACCCACACCATCACCACCACGCATGCATC | 4 |
| wt-36960-37080 | 306 | TCCCTGTCTTGCATGTGCCAGACCAATCAATGCACCCTCTTACTCATCAAAGCACCCCAAATGATCCAGATAGTCCAGAACCACGGTCCCCGACTGTATTTTATAACATTCCACCTATGC | 4 |
| wt-37080-37200 | 307 | CATTACCCCCCTCACAATTGCCACCACCAGCAGCACCAGCACAGCCACCTCCAGGGGTCATCAACGACCAACAATTACATCATCTACCCTCGGGGCCACCATGGTGGCCACCCATCTGCG | 4 |
| wt-37320-37440 | 308 | GACCTTGGAGACCAGAGCCAAACACCTCCAGTCCTAGCATGCCTGAACTAAGTCCAGTCCTCGGTCTTCATCAGGGACAAGGGGCTGGGGACTCACCAACTCCTGGCCCATCCAATGCCG | 4 |
| wt-37440-37560 | 309 | CCCCCGTTTGTAGAAATTCACACACGGCAACCCCTAACGTTTCACCAATACATGAACCGGAGTCCCATAATAGCCCAGAGGCTCCCATTCTCTTCCCCGATGATTGGTATCCTCCATCTA | 4 |
| wt-37560-37680 | 310 | TAGACCCCGCAGACTTAGACGAAAGTTGGGATTACATTTTTGAGACAACAGAATCTCCTAGCTCAGATGAAGATTATGTGGAGGGACCCAGTAAAAGACCTCGCCCCTCCATCCAGTAAA | 4 |
| wt-37680-37800 | 311 | AACCCTTGCCCTCTCCAGCAACCAATGTATCCCAAATAAATGTTACTTCTTTTGCTCTTAACCATTGACACGCCTGTCATTCTATCAATTAAACAAGGGAAAAAGGTTTAGCTATTCCAC | 4 |
| wt-37800-37920 | 312 | CAACACGACCCCAAGGAAGGCTTGCCAAAATTGGTGCCTTGCTCTCAGCACTTTGCCAGCAACTTATAGCATGGTAGGCAGCTCAACTCGGCCCGTCTTACTGCCCAGCCTACTCTCCAC | 4 |
| wt-37920-38040 | 313 | TCCCAGTCCATGTTCGCACTCCTATGCATTTCCTGCCCTCCCACTTTTACCCCAGTCCCAACCCAAAACCACACACAACACATAGAATTGTTAGTTTAAACAGTTTATTGATAGGTGGCT | 4 |
| wt-38040-38160 | 314 | GCTTTTAGCCTAATTGTGTATTGCTCTCGTTGCCAAAACCTGTTGTAAGGGCCGGCACCCGCAACATGGGGAAAACATAACCGCCGCCATCCCATGGGGAGGGTAGAGGCGGTTGACATG | 4 |
| wt-38160-38280 | 315 | TAGGTGAGTAGTGTAAGAAGCATGGCGAAGTAGACAGGTTACTTTTAGAGTGTAGTGTACAGGGCCGGGCGCAACAGTGCCACCAACCCGGGGTCTGAGCATTCCATGGGCAGCAGGGAC | 4 |

| | | | |
|---|---|---|---|
| wt-38280-38400 | 316 | ACTGCACTACCGCCAGGTCCTGGGGCAGCCGGGGTTCCTGGCGCTCCGGGGGCAGCCGGGCGGCCGCCGGTGGGTCCGCTGGGCCGCTGCCCCGCTCCGGGTGGGGGTGGCCCCGCTGG | 12 |
| wt-38400-38520 | 317 | GCACCGCTGCGCCGCCGCCAGGTCCTGGGGCAGCCGGGGTTCCTGGCGCTCCGGGGGCAGCCGGGCGGCCGCCGGTGGGTCCGCTGGGCCGCTGCCCCGCTCCGGGTGGGGGTGGCCCC | 12 |
| wt-38520-38640 | 318 | GCTGGGCACCGCTGCGCCGCCGCCAGGTCCTGGGGCAGCCGGGGTTCCTGGCGCTCCGGGGGCAGCCGGGCGGCCGCCGGTGGGTCCGCTGGGCCGCTGCCCCGCTCCGGGTGGGGGTG | 12 |
| wt-38640-38760 | 319 | GCCCCGCTGGGCACCGCTGCGCCGCCGCCAGGTCCTGGGGCAGCCGGGGTTCCTGGCGCTCCGGGGGCAGCCGGGCGGCCGCCGGTGGGTCCGCTGGGCCGCTGCCCCGCTCCGGGTGGG | 12 |
| wt-38760-38880 | 320 | GGGTGGCCCCGCTGGGCACCGCTGCGCCGCCGCCAGGTCCTGGGGCAGCCGGGGTTCCTGGCGCTCCGGGGGCAGCCGGGCGGCCGCCGGTGGGTCCGCTGGGCCGCTGCCCCGCTCCGG | 12 |
| wt-38880-39000 | 321 | GTGGGGGTGGCCCCGCTGGGCACCGCTGCGCCGCCGCCAGGTCCTGGGGCAGCCGGGGTTCCTGGCGCTCCGGGGGCAGCCGGGCGGCCGCCGGTGGGTCCGCTGGGCCGCTGCCCCGC | 12 |
| wt-39000-39120 | 322 | TCCGGGTGGGGGTGGCCCCGCTGGGCACCGCTGCGCCGCCGCCAGGTCCTGGGGCAGCCGGGGTTCCTGGCGCTCCGGGGGCAGCCGGGCGGCCGCCGGTGGGTCCGCTGGGCCGCTGC | 12 |
| wt-39120-39240 | 323 | CCCGCTCCGGGTGGGGGTGGCCCCGCTGGGCACCGCTGCGCCGCCGCCAGGTCCTGGGGCAGCCGGGGTTCCTGGCGCTCCGGGGGCAGCCGGGCGGCCGCCGGTGGGTCCGCTGGGCC | 12 |
| wt-39240-39360 | 324 | GCTGCCCCGCTCCGGGTGGGGGTGGCCCCGCTGGGCACCGCTGCGCCGCCGCCAGGTCCTGGGGCAGCCGGGGTTCCTGGCGCTCCGGGGGCAGCCGGGCGGCCGCCGGTGGGTCCGCT | 12 |
| wt-39360-39480 | 325 | GGGCCGCTGCCCCGCTCCGGGTGGGGGTGGCCCCGCTGGGCACCGCTGCGCCGCCGCCAGGTCCTGGGGCAGCCGGGGTTCCTGGCGCTCCGGGGGCAGCCGGGCGGCCGCCGGTGGGT | 12 |
| wt-39480-39600 | 326 | CCGCTGGGCCGCTGCCCCGCTCCGGGTGGGGGTGGCCCCGCTGGGCACCGCTGCGCCGCCGCCAGGTCCTGGGGCAGCCGGGGTTCCTGGCGCTCCGGGGGCAGCCGGGCGGCCGCCGG | 16 |
| wt-39600-39720 | 327 | TGGGTCCGCTGGGCCGCTGCCCCGCTCCGGGTGGGGGTGGCCCCGCTGGGCACCGCTGCGCCGCCGCCAGGTCCTGGGGCAGCCGGGGTTCCTGGCGCTCCGGGGGCAGCCGGGCGGCC | 12 |
| wt-39720-39840 | 328 | GCCGGTGGGTCCGCTGGGCCGCTGCCCCGCTCCGGGTGGGGGTGGCCCCGCTGGGCACCGCTGCGCCGCCGCCAGGTCCTGGGGCAGCCGGGGTTCCTGGCGCTCCACTGCACCTGGAA | 12 |
| wt-39840-39960 | 329 | TGCAGGGTGGGGGCGTGGTCCCCTGGACCCCAGCCCCGCCGATCCCTCCCCCAGGGCGTACCCGGCTTGCCTGGTTCTGGGGCTCCTCTGGGGGTCGCTGCATCCGCCGGTAGGGTTCGA | 8 |
| wt-39960-40080 | 330 | ATGGGCGTGGTCCGCTTGCTCTGCTGGCCCGGTACGCCTGGATTGCCGGCTGGGGGCTGGGGTCCCGGGACGCCCCCTCCCTGCTCCCACCCGGTTCCCTCCCCCAGGGCGTGCCCCGCT | 12 |
| wt-40080-40200 | 331 | TGCCTGGTCCTGGAGCTCATCCGGGGATGCTGCATCCGCTAGTCCGACCTGGGTGGGTGCGGTCCGCTGGCCCCACCCTGGGGGTAGCCGCCGGGTCTGCTGGTCCGGTGCACCTGGAAG | 8 |
| wt-40200-40320 | 332 | GCAGGGGGGGGGGCAGTGAGGGAGGGGCGTGGTCCTGGGACCCCGCGCCGACTGGCAGGGGGTCCCCATGGCACAGGCCTAGGGGTCCAGGGGGCAGCCGCGGCCCAGCGCGCCCCGTTC | 12 |
| wt-40320-40440 | 333 | ACGGGGGAGGACCGCGGCCGAGCCACCAGGGGCCCGGCGGGGGTGGGGGGTGCGCTCCCAGGCCGGACCCTGGTGCCAGGCAGGGACCCCGCGCCACCCGCTTCATGGGGGGGGAGGCCG | 12 |
| wt-40440-40560 | 334 | CCGCAAGGACGCCGGGCCGGCTGGGAGGTGTGCACCCCCCGAGCGTCTGGACGACGCTGGCGAGCCGGGCCGGCTCGCCTTCTTTTATCCTCTTTTTGGGGTCTCTGTGTAATACTTTAA | 4 |
| wt-40560-40680 | 335 | GGTTTGCTCAGGAGTGGGGGCTTCTTATTGGTTAATTCAGGTGTGTCATTTTAGCCCGTTGGGTTTCATTAAGGTGTGTCACCAGGTGGGTGGTACCTGGAGGTTATTCTATTGGGATAA | 4 |
| wt-40680-40800 | 336 | CGAGAGGAGGAGGGGCTAGAGGTCCGCGAGATTTGGGGTAGGCGGAGCCTCAGGAGGGTCCCCTCCATAGGGTTGAACCAGGAGGGGGAGGATTGGGCTCCGCCCCGATATACCTAGTGG | 4 |
| wt-40800-40920 | 337 | GTGGAGCCTAGAGGTAGGTATCCATAGGGTTCCATTATCCTGGAGGTATCCTAAGCTCCGCCCCTATATACCAGGTGGGTGGAGCTAGGTAGGATTCAGCTAGGTTCCTACTGGGGTACC | 4 |
| wt-40920-41040 | 338 | CCCCTACCCTACCTTAAGGTGCGCCACCCTTCCTCCTTCCGTTTTAATGGTAGAATAACCTATAGGTTATTAACCTAGTGGTGGAATAGGGTATTGCAGCTGGGTATATACCTATAGGTA | 4 |
| wt-41160-41280 | 340 | GTGCCTTACTGACTTGTCACCTTTGCACATTTGGTCAGCTGACCGATGCTCGCCACTTCCTGGGTCATGACCTGGCCTGTGCCTTGTCCCGTGGACAATGTCCCTCCAGCGTGGTGGCTG | 4 |
| wt-41280-41400 | 341 | CCTTTGGGATGCATCACTTTGAGCCACTAAGCCCCCGTTGCTCGCCTTGCCTGCCTCACCATGACACACTAAGCCCCTGCTAATCCATGAGCCCCGCCTTTAGGAAGCACCACGTCCCGG | 4 |
| wt-41400-41520 | 342 | GGACGGAAGGGGACTTGGGGTGATTTTCTATGTGGGGGTGGAAATATGAGCAAGAATAAGGACGGCTCCTTATTAACCTGATCAGCCCCGGAGTTGCCTGTTTCATCACTAACCCCGGGC | 4 |
| wt-41520-41640 | 343 | CTGAAGAGGTTGACAAGAAGGGTCAAGGTTTCGTCTGTGTGTTGAAGGGCAGGGGCTGTTGGGTGCATCTGGAACGGCTTACCTCGGGTAACTGTTTGCCATTAAAAGGTTGGGGATTAG | 4 |
| wt-41640-41760 | 344 | GTTTAGCCCCTTTAGCTGCCATTTCGAACCGGGGTGTGCAGATGCAGGTCTCCGGGTGGGCAGGCAGTACGAGATGTCACGTTGTGTTGTCTTTCCTCCCACCCCTGTCCTGGCTGTGGC | 4 |
| wt-41760-41880 | 345 | AAATGCGACCCTCATAGAGTTGTGTTTCAGGTCTGTGTCCTGTTTTGCGGTGGGTTATTTCTTCCCTCAGTGTTTGCCAGCTTATTTCCCCAGTTTTCACGTACTGGGGCCTGTGGACAC | 4 |
| wt-41880-42000 | 346 | CTGAGGGAGCGGCCGTTGGTGGGTATGTGTTGGAATTGCTCCCACCCTCAATTTTCGCTTGCCTTCTTCCCTTGTTAACCTGATAGCATAGCCTCTAGGTTTCCTTGTAGGTCTGTTTGG | 4 |
| wt-42000-42120 | 347 | GTTTGTTGGTTCACGTGGTGCTAACTTGAATTTTTTGGTTTTCTAGTTCCCTCTTAATTACATTTGTGCCAGATCTTGTAGAGCAAGATGGCCTATTCAACAAGGGAGATACTGTTAGCC | 4 |
| wt-42120-42240 | 348 | CTGTGTATACGGGACAGTCGTGTGCATGGAAATGGTACCCTGCATCCTGTGTTGGAGCTAGCAGCAAGAGAAACACCTCTCCGCCTTTCGCCAGAGGACACTGTAGTTCTGCGTTATCAT | 4 |
| wt-42240-42360 | 349 | GTGTTGCTTGAGGAGATAATTGAACGAAATTCAGAGACATTTACAGAAACTTGGAACAGATTTATAACACACACCGAACATGTGGATCTGGATTTTAACTCAGTATTTTTAGAGATATTT | 4 |
| wt-42360-42480 | 350 | CACCGTGGAGACCCAAGCCTTGGGCGCGCGTTGGCCTGGATGGCCTGGTGCATGCATGCCTGCAGGACATTGTGTTGTAACCAGTCTACTCCTTACTATGTTGTGGACCTGTCAGTTCGT | 4 |
| wt-42480-42600 | 351 | GGGATGTTAGAAGCCAGCGAAGGCCTGGATGGTTGGATTCATCAACAGGGCGGCTGGTCTACATTAATTGAAGACAACATTCCTGGATCCAGAAGGTTTAGCTGGACTTTGTTTCTTGCT | 4 |
| wt-42600-42720 | 352 | GGACTGACTTTGAGTCTGTTAGTTATATGTAGTTATTTATTTATCTCCAGAGGAAGACACTAATCTATACATTTTCTCAGCACTTTATATGAATCAGGGTCATTGGGCCTGCGGGGAACT | 4 |
| wt-42720-42840 | 353 | GAGCCAGTAGGATATTAGGCAAGGGTGACACAGTGCCCATGCATTATAATTTAACCAAACAGTGGTCGTGAGTTTTAGGCCGGCCATGGGGGCTTACAAGAATAACATGCCAATGACCCG | 4 |
| wt-42840-42960 | 354 | GCCCCCACTTTTAAATTCTGTTGCAGCAGATAGCTGATACCCAATGTTATCTTTTGCGGCAGAAATTGAAAGTGCTGGCCATATCTACAATTGGGTGTCCTAGGTGGGATATACGCCTGT | 4 |
| wt-42960-43080 | 355 | GGTGTTCTAACGGGAAGTGTGTAAGCACACACGTAATTTGCAAGCGGTGCTTCACGCTCTTCGTTAAAATAACACAAGGACAAGATACTAAAGAAATAACTGAGGTGAGTGTGGGAAGAT | 4 |
| wt-43080-43200 | 356 | GGGAATACTATGTGTTATGTTAACGGGTGAGAGCCTATACTGCAGCCCAGACTCGGGGGGAGGAGGAAATGGTAAGAGTTATACTCTACTTATCTTTTTTGACACTACATTTAACTGTTA | 4 |
| wt-43200-43320 | 357 | TGTAACAATGTTTGCTTATTTTCATGTTCAATAAACGCTATGTTAATGATGAAGAACCTGTGTTCTTTGGAAGTGGGCCCAATGGGGTAGTAGGTTTTGGGAGGGTGCCGTGCTAGATAT | 4 |
| wt-43320-43440 | 358 | TTCAACTGCCACAGACCCCATTTTGTCCCACCTGTTACCACATTCTAGGTCCTGCATCCAGTGGGCCAGGTGTCTCACCATGGCTCTTTCTAGGTGGATACCACAGTCCAGGCCCCCAAG | 4 |
| wt-43440-43560 | 359 | GCTACCGTGCTAATTACCTCCTCATGTCCACCCCCACCCTGTGTTACTGTCGCCTGATTATCCTGGCTTAGCAGCCTCCAAGTTTTACAAGACGTCCCATTGCCCTGCCCTTGGTCCAAG | 4 |
| wt-43560-43680 | 360 | TCTCGCCGGTTTTCAGCAGCCTGTTGTAGCCTGCCCCCAAGTTTCGCAGGTTTCCCCCATGCTTCCACCCGTTAACCCAATAGCATGACAGCCAATCCAACACGAGGCAAGTTTTAAGAG | 4 |
| wt-43680-43800 | 361 | TTAAAAGCAACTACTGTTTATTTTCCAAAATGAGCTGGGTATAGTTGATGATCTGTAGGCGCAGCTCATCCCCACATTCCAGGTCCTTGATGGCCTCGTAGATGGCATCTTCGTCGACAT | 4 |
| wt-43800-43920 | 362 | TGACAGCCTTCTCATATACCGTGTCTCTGGGGCTGACCTTTATACAGAAGGCGTCCCCTACTAGGTCCACGGCCAGCTCGTAGGTGGGGCCTATGTTTTCACATAACAGTTTCAAGCAGG | 4 |
| wt-43920-44040 | 363 | TCTCTGGGATGTGAAGGGAGGTGCCCTGGAGCAGGAGATGCATGATTAGGCGCCCTTTTCCATTTGTGCTGAAGATGGGGCAGATGGTGCCACAAAAGTGTCCGGTGACCAGGTAAGCGT | 4 |
| wt-44040-44160 | 364 | AGAGAAGGCTGGGTTGGGAAAGTCCAGCCTTTACTGCACTGGGAGAGCTGCTGAGCAGAGACACATAGAAGGTCTTGTTGGGTATTATCTTGTGGACATTGTTGAAGAAGGAGAGCTGGG | 4 |
| wt-44160-44280 | 365 | TGGAGCTAAACTCCTGAGGCACATGAACCTGGGACCTATTGATGCAGATCTCGCAGTGAGACCCCAGAGTCAGGCTGTGGCCGAAGGGAGACAGGCGAAGGCAGCGCCCGGGGGAGAGAG | 4 |
| wt-44280-44400 | 366 | TGCACAGTGACAGTGGGAGAAACACGGCCTCTGAGACATGTATGGGGGTGTTCATCTCACGCAGAAAATCTTTGCCCAGCTCAAAGTTGGCAGAGATTCCCCTGAAGAAGTCCCGTAGTG | 4 |
| wt-44400-44520 | 367 | AAAAATGGGATCTGTCTACACCATGTCTGGTGTGCCGGGAACATATTGATCGGGCCACACTGCCAACCCTTTCCATTCTTCCCAGCTCTGAGCGAGATTTTCCACACCTGGACACCGACT | 4 |

| | | | |
|---|---|---|---|
| wt-44520-44640 | 368 | TCACGCTATGCGCCGAGGCCTTTGAGGCCGTGTAGTTTCTGTGGTGCGGATGCATTAGGCGGCGCAATGCGGGATCTGCCGGTCGCTGTTGGCGTGCATTCACGGCATCTGGGGTGACCG | 4 |
| wt-44640-44760 | 369 | GGGCCATCGGGTTTACTTTTCACACGTAGACCTGGGAAGTTTGATAGGACTGTACCAGGTCAAGGCCGTGGATGCGCAGGACCACGTCCAGTTCCTTAGTGACATCCACGAGGATTGTTT | 4 |
| wt-44760-44880 | 370 | TGCCCACTCTGGCCACTTGTGTGGATTTAAATATGTACACAAGCGTAATTAACGAGTCACAGACCCCCTGTTCCAGATTCTGACCGGCTGCAAGCGCTGCCTTAAAGGCCTGGAAGCTGG | 4 |
| wt-44880-45000 | 371 | GTGGGTAAATCTGACCAAACAGCACGCTCGGATTCGTGATGCTGTGGTTGATGGCACACAGGGGGTCGCAGAACAGGTGCTTGTGGAAGTCTTGCGGGGTGCACATCTGCAGCCAGGCCC | 4 |
| wt-45000-45120 | 372 | TTAGCCTGGGGCATGGCACATCCAGCAGCGTGTTTTGGGTCTTGATGAGGAACACGATCCTGTCTAGGATTTTGATGTTGTTGCCGAACGAGTCAAGAATCAGGCTCTTGAAGCGGTCAA | 4 |
| wt-45120-45240 | 373 | GGGTGTCCTTGGCGTCCGGGTGGGCCCCGAGGCTCTCGCAGAGTGGGCAGATGGTCCGTGAGGCATTCTTGTGCCTTAGTCCAAACATGGGGGCCAGGAGGCAGGGGCCCTGCGAATGGT | 8 |
| wt-45240-45360 | 374 | CGCCAGCCTCCGGTCTGGTGATGGCCAGGGCCAACTCCGCCAGCTCATCGCCGCTGTATTCCGCGTTTAAACCGATAGCATGGTGGCCTGGCCCCCCGAGCAGGTCCGTCCCCTGCCACG | 8 |
| wt-45360-45480 | 375 | TACCTAATAGTAGTCCACAGTAGTCGGCCTTGGTTGTAATTTCAGGAGAGAGTCCTCCCTTTTCGGCCCTGAGAAATGGATGCTGAACTCGGTTTCTGGTAGGCAGGTGGCAGCACAGGG | 4 |
| wt-45480-45600 | 376 | CGGTGTACAGGCCCCTGCCGACGTCCCCTGGGACATCCTGGGAATCTTTGCAGGTTCTGGGTCCAGGGAGGGTAAGAAAAGTGGGGGTGGTTCTGGGCCACATGGACTTGAAGCAGAAGT | 4 |
| wt-45600-45720 | 377 | TGGCCGGGGACTGGCCGGTGAGGATGGATTTCAGAAACTCCAATTTGTAGTAGCCGAGGTTGGCATTTCTAATCATGTCAGAAGAGGACACAGGGAGGAAGCACCGGCAAATGTAAAAGT | 4 |
| wt-45720-45840 | 378 | GAAGCTGGATGTCAATGGCAAGAATCCTGGAGGGCATGAAGAGGGAATCCAACCCCCCGGCCATGGGGAAGTATTTTATCAGGATGTGTAAAAAGTCCATGCCTGTGATGAGGCTAGAGA | 4 |
| wt-45840-45960 | 379 | TCCAGGCTCGTGGGGCATTTAGACAGTAGTAGCAGAGCAGGGCATAGTCCTCAAAGAAGGCCACGGGGGCATCTGAGTGATTGACCAGGGTGTCGAGCAGATCACAAACTCGGCAGGTGC | 4 |
| wt-45960-46080 | 380 | TGGCTGGAGAGAGGGACTCGTAGGTGTGGACGAGTGGTGGGTAGGCTATGCCTTCTTCCGCGTTGGCTGGAAGATAGGAGTGGGCCATCAAAAGGCCGACTGCCTCGAACTGGCTTTTCA | 4 |
| wt-46080-46200 | 381 | GATTGTCCACGGTCCAGGGCACAAAGTCCTCCATCTTTGGAGTTCTGCCCGCGATCTGTGCCACCTCTGTTACGCCACTCCTCGTGAGGGGGCAGCTGGACAGTCTTTTTCCGGTCAGGG | 4 |
| wt-46200-46320 | 382 | GGTTTGGCTCGTTTGCGCTCGTGACTTTGTGAGCCATGACACATCTGGGTGGCAAGGTGAGGTCTTCTGGGTTTTTAATACCGGGGTCGGCACCAGTTTCTGGGACACCGCCACAAGGAC | 4 |
| wt-46320-46440 | 383 | AAGGTGGGCTAGCAAGTTCTCGAGTCTACGAAGACTCCGGGGGCAGTCTTTTGAGTTTCTCGCCTATGATCCACCCCAATCTCGCCCCCCTAATTGCGCCATCTGCCTACGCGAGGCTGA | 4 |
| wt-46440-46560 | 384 | ACCTCCTGAATCACTGCATCTTTCTTGAGGCGTTTAAAGAAGAGAATAGTGGCCAGGGCCTCGGTGGGGTCCAGCGTGAGGTCTTATTTTTGAAAAGGGATATTATAAAACAGGTCATTG | 4 |
| wt-46560-46680 | 385 | CTCGGATTGTGGCAGCCGATAGCACCCTAGATCTAGTGAATCATGGCGAGCCCGGAAGAGAGGCTCCTAGACGAGCTCAATAACGTAATTGTGTCATTTCTGTGTGACTCTGGGTCTCTG | 4 |
| wt-46680-46800 | 386 | GAAGTGGAGAGATGCTCCGGGGCGCATGTGTTCTCCAGGGGCAGCTCCCAACCCCTCTGCACCGTGAAGCTGCGCCACGGACAGATTTACCACCTGGAGTTTGTCTACAAGTTCCTGGCC | 4 |
| wt-46800-46920 | 387 | TTTAAGCTGAAGAACTGCAACTACCCCTCCTCGCCCGTGTTTGTGATATCCAACAACGGCCTGGCCACCACCCTGAGGTGCTTTTTGCACGAGCCGTCGGGTCTCAGATCGGGCCAGAGC | 4 |
| wt-46920-47040 | 388 | GGCCCTTGCCTGGGTCTCTCAACGGATGTTGACCTACCAAAGAACTCCATCATTATGCTGGGCCAGGATGACTTCATTAAGTTCAAAAGCCCCCTGGTCTTCCCTGCTGAGCTTGATCTC | 4 |
| wt-47040-47160 | 389 | CTGAAATCTATGGTGGTCTGCCGGGCCTACATCACGGAACACCGGACGACGATGCAGTTTCTGGTGTTTCAGGCCGCCAACGCCCAGAAGGCCTCGCGGGTCATGGATATGATTAGTGAT | 4 |
| wt-47160-47280 | 390 | ATGTCTCAGCAACTGTCTCGGTCTGGTCAAGTCGAGGATACGGGCGCCAGAGTCACAGGTGGAGGAGGTCCCAGGCCTGGCGTCACGCACTCGGGGTGTCTTGGGGACTCACACGTTAGG | 4 |
| wt-47280-47400 | 391 | GGGCGCGGTGGTTGGGACTTGGATAACTTTTCAGAAGCTGAGACCGAAGACGAGGCGAGTTACGCTCCTTGGAGGGACAAAGACTCGTGGTCGGAATCCGAGGCGGCGCCGTGGAAGAAG | 4 |
| wt-47400-47520 | 392 | GAACTCGTGAGGCACCCCATCCGCAGGCACCGGACACGCGAGACTCGCCGTATGCGCGGGAGCCATTCACGGGTGGAACACGTGCCCCCCGAGACCCGGGAGACGGTGGTGGGGGGAGCA | 8 |
| wt-47520-47640 | 393 | TGGCGTTATTCTTGGCGCGCCACACCTTATCTGGCACGGGTGCTGGCTGTCACGGCCGTGGCCCTGCTCCTGATGTTTCTGAGGTGGACCTGACGTTGCAGGCCCTTGGGGAGCGGGGGT | 4 |
| wt-47640-47760 | 394 | TCTCCAGGCTCCTGGATCTGGGGCTGGCCTGCCTGGATCTGAGCTATGTGGAAATGAGGGAATTTGTGGTTTGGGGCAGGCCCCCAGCTTCTGAGGCGGCTGTGGCCTCTACGCCAGGCT | 4 |
| wt-47760-47880 | 395 | CGCTTTTCCGAAGCCACTCGTCCGCCTACTGGTTGTCGGAGGTGGAGAGGCCCGGGGGCCTTGTCCGCTGGGCCAGGTCACAGACCAGCCCCTCATCCCTGACCCTCGCGCCCCATCTTG | 8 |
| wt-47880-48000 | 396 | GCCCGTCCCTCTTGTCCCTTTCAGTGGTCACCGGTGGTGGGTGTGGAGCCGTGGCCTTTTGCAACGCCTTTTTCCTAGCTTATTTTTTGGTTGTGCGGTCTGTTTTCCCCGCGTTTTCCG | 4 |
| wt-48000-48120 | 397 | ATAGAATAGCTGCCTGGATCTGCGACCGGTCCCCTTTCTGCGAAAACACCCGGGCCGTGGCCAGGGGTTACCGAGGCCTCGTGAAGAGGTTCTTGGCATTCGTGTTTGAGCGTAGTAGCT | 4 |
| wt-48120-48240 | 398 | ATGACCCCCCCTTGTTGAGGCAAAACTCTAGGCCTGTGGAGCGCTGCTTTGCCATCAAGAATTATGTCCCGGGCCTGGACTCACAAAGCTGTGTGACGGTCCCGAGCTTCTCCCGCTGGG | 4 |
| wt-48240-48360 | 399 | CCCAGTCTCACGCCAGCGAGCTCGATCCCCGGGAGATTCGCGACAGAGTTACACCAGCGACTGCACCTTCGTTCGTGGCTGATCATGCCTCGGCTCTATTGGCCTCCCTCCAGAAGAAGG | 4 |
| wt-48360-48480 | 400 | CCTCCGACACCCCCTGTGGGAATCCCATTCAGTGGATGTGGTACCGCCTGTTGGTAAACTCGTGCCTGAGGAGTGCCCACTGTCTTCTGCCTATACCTGCCGTCTCTGAGGGGGGGAGAA | 4 |
| wt-48480-48600 | 401 | AGACGGGCGGGGGCGTAGGGGAGGAGCTCGTGGGGGCCGGGGGGCCCTGCCTGAGCCGGGATGTTTTCGTGGCGATCGTAAGCCGCAATGTTCTCTCGTGTCTGCTGAACGTGCCTGCCG | 8 |
| wt-48600-48720 | 402 | CGGGTCCCCGGGCCTACAAGTGTTTCAGATCCCACGCCTCCAGACCGGTGTCTGGCCCGGATTACCCTCCCTTGGCCGTGTTTTGCATGGACTGCGGTTACTGCTTGAACTTTGGAAAGC | 4 |
| wt-48720-48840 | 403 | AGACAGGTGTAGGAGGCAGGCTCAATTCCTTTAGACCCACTCTCCAGTTTTATCCCCGTGACCAGAAGGAGAAGCATGTGCTGACCTGCCATGCCAGCGGCCGTGTGTACTGCTCCAACT | 4 |
| wt-48840-48960 | 404 | GCGGCTCTGCGGCGGTGGGCTGCCAGAGGCTGGCTGAGCCACCGAGCGCCCGCTCGGGCTGGCGGCCCCGAATCCGGGCAGTGCTGCCGCACAACGCGGCCTACGAGCTCGACCGTGGCT | 12 |
| wt-48960-49080 | 405 | CCCGCCTCTTGGATGCCATCATCCCCTGCTTGGGACCCGACCGCACTTGCATGCGGCCGGTGGTCCTGCGGGGGGTGACGGTCAGGCAGCTCCTGTATTTAACTTTGCGGACAGAGGCCA | 4 |
| wt-49080-49200 | 406 | GAGCCGTTTGCTCCATCTGTCAGCAACGCCAAGCTCCAGAGGACGCCCGCGACGAGCCTCACCTGTTCTCCTCCTGTTTAGAGGTAGAATTGCCACCTGGTGAGCGGTGTGCGGGCTGCC | 4 |
| wt-49200-49320 | 407 | GTCTCTATCAGACGCGTTATGGCACGCCGGCTGCCCAAGCCCACCCTCCAGGGGAGGCTGGAGGCGGATTTTCCAGACAGTCCCCTGCTTCCTAAATTTCAAGAGCTGAACCAGAATAAT | 4 |
| wt-49320-49440 | 408 | CTCCCCAATGATGTTTTTCGGGAGGCTCAAAGAAGTTACCTGGTATTTCTGACATCCCAGTTCTGCTACGAAGAGTACGTGCAGAGGACTTTTGGGGTGCCTCGGCGCCAACGCGCCATA | 4 |
| wt-49440-49560 | 409 | GACAAGAGGCAGAGAGCCAGTGTGGCTGGGGCTGGTGCTCATGCACACCTTGGCGGGTCATCCGCCACCCCCGTCCAGCAGGCTCAGGCCGCCGCATCCGCTGGGACCGGGGCCTTGGCA | 8 |
| wt-49560-49680 | 410 | TCATCAGCGCCGTCCACGGCCGTAGCCCAGTCCGCGACCCCCTCTGTTTCTTCATCTATTAGCAGCCTCCGGGCCGCGACTTCGGGGGCGACTGCCGCCGCCTCCGCCGCCGCAGCCGTC | 8 |
| wt-49680-49800 | 411 | GATACCGGGTCAGGTGGCGGGGGACAACCCCACGACACCGCCCCACGCGGGGCACGTAAGAAACAGTAGAGGGCACGAAACATGGTGTATGCACTTTATTAATAAACAATTACAGATACA | 4 |
| wt-49800-49920 | 412 | AAAACTTGAGTCTCTCGAGGTCTGCGATGAGGCGGTGGGTGGAACGCTCCAGCTTGCGGCGAAGCTGGCTCACGAAGCGAGACAGTACTCGGCTAGCCTGACTAAGGGTGAGGCTATAAC | 4 |
| wt-49920-50040 | 413 | GCAGGTCCTGTTCCGGGGCGGCGGTGGATAGAGAGGAGGGGGATCCGGAGGGGACCACTAGGTCGCCGGAGGTCGACCCTCCTGTCACCACCTCCCTGATAATGTCTTCAATAGACAGAA | 4 |
| wt-50040-50160 | 414 | ATTGGGTGACCACTGAGGGAGTGTTCCACAGTAATGTTGTCTGGTCGCTAGATGGCGCGGGTGAGGCCACGCTTTGCGAAAACGAAAGTGCTTGAAAAGGCGCGGGATAGCGTGCGCTAC | 4 |
| wt-50160-50280 | 415 | CGGATGGCGGGTAATACATGCTATCCTTACATTTTGGCATTTTGGGCAGCTGGGAGGCGGCGGATGGGGGTGCTTCTTTTCGCACGGTGTATGTTTGGGGACCCGCATGCCGGTACTGGG | 4 |
| wt-50280-50400 | 416 | ATAGGCGCACCTCGGGCCGCGCGCCAGGCTCCGAGCCGGAATGCATTGGGGGCAATGGGATTGCGGGGGATTGTTGCTGTCTGCTCCTGACAGGGAGAGACACGCGCGGCGGAGATGCAG | 8 |
| wt-50400-50520 | 417 | CCGACGGCGGGGCCGCGGTGGGCTGCCCCCGAGGACGGGCGCCGGCCGCCAGCGCCCCCGTGGCCTTTGGCACGGGCCTGGCACCCACCGCTTTAATTGTGGGGGTGGGCAGGGCAGCTG | 12 |
| wt-50520-50640 | 418 | CATCTTGGGGCCTTTGTGCTTGCGTTTTTTGGGGGCGCGGTGCCAATGCACCAACTGGGGTGTGCGCCGGGGCGGCCAAGCCGGACCCCAGGGCGGGTGCCTGGGGGATGGGAAAGCCGG | 8 |

| | | | |
|---|---|---|---|
| wt-50640-50760 | 419 | ACGGCGCTTCTCCCGGGTCGAACGCTGGAGTAGCGGAGGCTGCTGCGCCGGCGGCCACCACGGGCGCACGGGGTCGCAGCCCGACGGCCGTGGGAGGCGGGTGGCGGAGGGCCGAATCT | 12 |
| wt-50760-50880 | 420 | CCGCGGCTTCTTCCCGGCCCCCCTGCTGTTTCTTCTCCCGTTGCATGATAGAATGGCCATAGGGTGGGTCCTGAGAGGAGGCTTGTGTGTCCTGGGGCTGGAGCCCAAAAGTCGTTAAAG | 4 |
| wt-50880-51000 | 421 | ATGCCGCTGATGGTGTGGGAGCTATGCCTCCCGTCGACTGGCCGGGCTTGTAGGGGGCTGAGGGTGGATAACTGGGCTTCTGTGAAGGCACCAACCCTGGAATCTGGATGGTATGTTTCT | 4 |
| wt-51000-51120 | 422 | TCTGTGACCCCGAGGCAGTCGATGGTGTAGAGTGTGGAGACAATGTGTAGACGATGGGCCCTTGTTCAGAAGCCCAGGGACTTGAGGGGGGCTGTTGTGGTGCTGGTTGGGGAAGGAGCT | 4 |
| wt-51120-51240 | 423 | CCAGGGAATCTTTGGGCCATGGCCTTGGGGAGCTTCCCGGAGACCGGTCTGGGCTCTCGGAAGCCCTCGTTTCGGCCCCGAAATAGGGCCTTGCCATCAATCGGGGGCCTGGGAGAGTGA | 4 |
| wt-51240-51360 | 424 | TGGGGGCGGCCAATCCCGGGGTAACTGTCACGTCCCGGGGGGAGGAGGTAGGAGACAGCCAGTCCCTGGGCCTGCCAGGGGCCACCTTCTCTAAGAGGGGGCTCTGTGGGCTGGGAGGGC | 8 |
| wt-51360-51480 | 425 | CAGAGGCCTCAGATTCAGCAGTAGTGCTCCCCTTTTCCCCCTGGTCCGTCTCCCCTCCTCCCAACTGCTGGAGCCGGTCGGAGGAGGCCGGGGTGTTATCTGCTGACTGAAACCCGTCCC | 4 |
| wt-51480-51600 | 426 | CGCTGACCAGTCCGTGCCCCACCCTTGGGGGGAAACCGGAGAACAGCTCCTGGACGTTGCGTGGATTCGGGGGAAGCTGGTATCCAACCGGCAGTGGAGGATCTTCGTGCTCGTAGAAGG | 4 |
| wt-51600-51720 | 427 | AGGGGTTGAGTACATCGGTCGGCCATCGTGAGGCCCCGGCCGCGTTAAAGTAGAACTGCACGTCCGGCAGATTGTGCCGATAGGTGAAACACTTCCAGATGATGTTTTTTCTGTTGGCCA | 4 |
| wt-51720-51840 | 428 | GGATGGCCACGGTGGGGGGCCTGGCCTCCTTAGGTTTGGCGGCCCTGGCCTCGGTGAGAAGCTCGCGTAGCCACACGGCCTGGCGTGCAAAGATGGACATCTCTGGCTCGAAAGACTCGG | 8 |
| wt-51840-51960 | 429 | AGTAGCCGTCCAGGTCCTGCAGAAAATTCAGCGAGATGGTCTCCACCAGGGACCGGAAGGGCTCAGAGTGCCCGTCGCAGTAGAGGAGGGGAGCAACGACCCTGACCTGTCCCAGGGTCT | 4 |
| wt-51960-52080 | 430 | TCAGGTTAAACAGATATTGAGAGGAGACAAAGAGAGTTAGGGGCCGACCGAGGAAGGCCGCCGCCACGGCCGCCTCAAAAACGGAGACGGGGATGGTGTCACCGGAGCCCCTCTTAGGAC | 4 |
| wt-52080-52200 | 431 | CGGTAATGGGAGTGCCATAAGGCATAAGATTTCTCAGGGCCCGGCCGGTAACGGTGCCGTAGGAAGACGGGGTTTCGCGGGGGACCTCGAGTCCCTCCGCCCTGGGGAGCTCTTCTCCGC | 4 |
| wt-52320-52440 | 433 | GGAGCAGCATGCAGGCTCGGGCGCGGGCCGTACTTTGGTTGTGGACCAGGCCCAAGAACTCGGGGTGCGGCCAGAGGGCGGCTCGGGTATCCATCTCCTCCCAGGCGTCCTGGAAGAAGA | 8 |
| wt-52440-52560 | 434 | TGAAGCCGGTGGGTGGACCGGCGATGCGGTGGCGGGTGAGGCGGCGCGCGTCTTCCCCGTCGTTGCTGCCGCGGGTGGTTGAGGGCATGCCCCCCCTCCCGGAGGCTGGACTCCTGACCA | 8 |
| wt-52560-52680 | 435 | GCCTGTAGGTGAGGACCGAGTCCGACAGGAGGTCTCCCAAACCCCCATCTCTCGCTAGAGCCGAGACCAGGCCGAGTCCTGCGTAGAACGATGGGGCGCCCAGGAAGGCGGCAGCGTAGG | 8 |
| wt-52680-52800 | 436 | CCGGATGTGTGCCGACCAGCAGCGCCATCATCTCCCGTTGTTCCAATAGAATAACTTCCCGGTCTGTGGCCGGGGCTGGATAAGGGGGGTGATTCCTAGAGGCGATGAGACTGGCGTGCG | 4 |
| wt-52800-52920 | 437 | CTAAAAGTGTCATGGCCACAATGGGGTTGTCTGCCAGGTCTTCCATCAGGGCTTTGGGCGCAGAGACGTATTCCCGAAGCAGCTCCCCGGCGTTGGACTCCACGTCGGGCCAGGTGTCCC | 4 |
| wt-52920-53040 | 438 | AGTAGGAGTCGGCGGCGGCGGCGCTGAGGCGGGCGGAAGCTACACTGGCCAGGGTTCTTCTCCTCCTCTCTTGGTCATCCTGCGGGGGACCAATAGCTTGGGGGCGTCCGGCTGGGGTCA | 8 |
| wt-53040-53160 | 439 | GGGAAAAGGCCTCTGGGTTCTCCAGCACGGTGGGCATGACATATTCCAGAAAGTTGTGGTAGACGGGGATGTAGTTGAGCGGCTCCTGGGTGTCTGCGGAGACGTAGGCCGGGTTAAGGG | 4 |
| wt-53160-53280 | 440 | GGTCGCAGGGAGACTCTGTTTCCAGCCAGAGGGTGCCGGCGTATTTCGCCGGCCCTGCCGCCGCCAGAAATTGTGCCCGCCGGGTCGGGGCTCCATTGCCCCATCCAGTTGGTGGTGCCG | 8 |
| wt-53280-53400 | 441 | AAATCGTGATGAGGAGGGGCAGGTTGTTGGTCAAGGGATGCTTAACGAAAACGGTAGGCTGGGCGGTCTCGTAAAAAGCCAGGAAACTCTGCTTGGCCGAGGCATAGCGCAGCAGCTTGT | 4 |
| wt-53400-53520 | 442 | CCTTGAGGAGGGCATACTGGGAGCCAGCCGAGGCCCCAAGCGCCAGGCCCCTGGCAGCCTCCACCACGATCTTGAGCTGGCGCGGGTCGGTGTGGCCCCTGGCCTGGGTGACCAGATCCT | 8 |
| wt-53520-53640 | 443 | GCAGCGTTCCCTGCAGCTGGGACTCTTCCTGGGCCTCCTGGATGATGGCCTCCAGTCGGGAGAGGCGCCTTTTCCAGTCTGCGACGGTCTCCTTGCCCCCCGCGACCCGCTTGGGGTCCA | 8 |
| wt-53640-53760 | 444 | ACGTGGCCAGAGCCAACCTCAGCTCCTCCATGCCATCCATGGAGTTCTGGGCCATGCCCTCGACTTCCAGGAGCCGTGTTAGCTCATGAATTTCACCGTCAGCCGCAGCGGCTAGGTTCA | 4 |
| wt-53760-53880 | 445 | GCCAGGCACCCGCGCCCCCAGCTAAGGCCAGGGCTCCTTCGGAAAGACCCCGCACGGCCTCGCAGATGCCCCGGATCCACTTGGCGGCTGCCAGGGATTTCCGGTAGGGCCATGAGCCGT | 8 |
| wt-53880-54000 | 446 | TCCCGGCCGCTGCCCGGGCCAGGGCGGCCTCGAGGGGAGCCTGGACAGGGGCTTTGGGCGGGGAGGGAAGCAGGCTCCGGAGTTCATCGTCGGGGGCTTCGTCGCGTGACCTGGAGAGGA | 8 |
| wt-54000-54120 | 447 | CGGCCTCCAGAGCCGTGTGAAAGCCCCGCCGAGTGCTTGCCGCCATCTCGTGGGCCTTCGCCATCAGGGTCTGGCTCTCCCGGACCTGCTCTTCCAGCGCCCGGACCTCGGCCGCCTCGG | 8 |
| wt-54120-54240 | 448 | CCTCGGTCAGCAGCTCCGAGAAGAAGTCCCCGTGGCCTGGAGGAGATCGTCCCGCTCTCGCCTTGTCAGCAGCTGGGGCTTCTTAGGCCAGAGCGCCGAGTCCGAGGCCAGCCTGGGCG | 8 |
| wt-54480-54600 | 451 | GCGCGAGGGCCGAACGGATGGCATCGGCCGCCGTCCTGGCGCGAAAGAGGGCCCCGGCCGCTTCCTCCGCTCCTCGCCCTCCTCCTCCTTCTTTGGCGGTAGCCGCGGGGGTGGCGGGCC | 12 |
| wt-54600-54720 | 452 | AAGCGTCCAGTCTGGCCAGAGGGCCGGTCTCGATATCCGTGAACCAGCCGGGTTCCGCGGCCTCCATTCTCTCCGCCGCACCACCATCGTCCACGAGCAGGGATCGCAGTCTCTCCCTCC | 8 |
| wt-54720-54840 | 453 | TCACCCTCGTTATTCCCAATAGCATAGCGGCAAGGATCTGTGTGAGGGAGTCCAAGATGTCCGTGTTTCTGGCTACTGCCGCCGCTGCTGCCGCGGCTGAGTCCGTATTGTCTGGCAGCA | 4 |
| wt-54840-54960 | 454 | GGGAGGCCAGCAGGGTGTTCCAGTCATCGGGCGAAGTGGGAGCGGGCTCTGGGCGTGCCCCCAGCGCCTTCCTAATTCTGGCCCAGGCCTCATTCGCCTCTCGCGCTCGCCGCTCCTGCC | 8 |
| wt-54960-55080 | 455 | TCTCCTTGTCTTCCTGTTCTCGGAGCTTCTCCTTTTCCTTGCGCCCGGTCTCCATAAGCTGCCGCAGCTTCTTCTCATACTGTCGCTTGAGCTCTTTGTTGGGGGCAGTGTCCAGAAAGG | 4 |
| wt-55080-55200 | 456 | CCTCGAGCTGTTCCTCGGTGGCGGGCTTAAAGCCTTCGGCCTCCAGGCGCCAGGCCTGCACCTCCTTCTGTCTGAGCTGATCGTTGTTGTTATTCTTCAATTTCTGCAGGTAACTTAGGA | 4 |
| wt-55200-55320 | 457 | AGCGTTTCTTGAGCTTCCCTGGGATGAGCGTTTGGGAGAGCTGATTCTGCAGCCCAGAGAGTAGTCTCAGGGCATCCTCTGGAGCCTGACCTGTGATCGTCGCATCATAGACCGCCAGTA | 4 |
| wt-55320-55440 | 458 | GACCTGGGAGCAGATTCACCGCCGCGGCCGTCTCCTTTAAGGTGCTGTGAGTAGCAAAATTCTGCAAGGCCACTAGGCGCGCTGGCTCCAGCGTCAGCCGGTTGCCCATCTCGAATGTGT | 4 |
| wt-55440-55560 | 459 | GCAGGGCCTCTGAGACCATGGGGTCCAGGATGCGGTCAATGCCATCCTGCACCTCAGGGTCAAGGACCGGCAAGTCACGATAGAGGTGGTCTATGCTCTCCTCGAAGGAGGCAATGTAGT | 4 |
| wt-55560-55680 | 460 | TATCGATGGTGTAGAAGGTGATGGATTTCAGGATGTTCATCAGGTACTTTTTGGAGCGAACAATCTGCTGTATAGTGTCACGTAGGCGGATGTACGTGGGGTTCTTTGCGGCCCCGACTA | 4 |
| wt-55800-55920 | 462 | GAGAGATGGGGGCGCCTGGGGCCGAGGCCACACCGGCCAGGCCCAACATTGCCTCGATGTCGTCCAGGATGGTGCGGAGGCGCTTTTCGTTTTCTCTGGTGGTCTCGAGCTCCTTCTGTT | 4 |
| wt-55920-56040 | 463 | TTTTCGCGACTGTCTCAAACTCTGGAAGGGGGGCAATGCTGGGGTCGTCCTCCTCAACTCGCTCCAGGGGCCAGGGGATACCGCTCATATCACTAAGGGCGGTGCCCAGGTAGAGGAGCT | 4 |
| wt-56040-56160 | 464 | CGCGATAGTCCCATTCAATGGACGTGTACCGGATGTTTAGGAGAGGCAGGGAGGCGATGATCTGGCATGTGTGCCGCAGGTGTGTCAGGAGGTCGTCAAAATCCATCACTGTTGGGAGGC | 4 |
| wt-56160-56280 | 465 | TTGGGTCCTCAAGGTAGGAGAGATAATCGGAGGCCGCCGAGGCCACCTTGTCCCTGATGTCCGCCGTACACCTGCGCACGTGCAGGGCCGCATTCTTGGACCGGACGGCCACGTTGTGGA | 4 |
| wt-56280-56400 | 466 | CAAAGGGGGGCACTGAGGCGGCGGGAGGGGCCCCATACTCTATCGCTGTCAACAGCGCCAAAAAGCGGACGTCCTCCTCATCTACCCCAGCCTGTTGTCTGGCCACGGCCGTTCGGGCGG | 8 |
| wt-56400-56520 | 467 | CCTCCGCCAGGGATAGGAGGCGCTTCCAGCTTTCGTCGTCCAGGACCAAGGGGACGTCCACGTGCGGGCCCCTGTAGATGGAATTATCCTCGGGTTCTCCTCCTCCTTCCCCCGCCTCCT | 4 |
| wt-56520-56640 | 468 | GATCTCCGCCCGAGAGCAGGTCGGTCAGGCGTCTGCGGGCCGCCTCCAGGTCAAATTTTCCATCGTCGCTCTCGGCCAGCTGGGGAATTTCAGCCAGCATCTTAGCACCGGCATCTACAC | 4 |
| wt-56640-56760 | 469 | GGACCGCGTCCTTCGTGGCCAGGGACGGCAGGCAGGCCTCCAGCTTTGCGGCCAGGTGCTTATGGAACTCTCCCGCTCTTCCCTTGTTTTCTGATAGCATGTTTGCGAGGTTTTGGATGT | 4 |
| wt-56760-56880 | 470 | TAAGTTCGGAAGTGAGCAGTTGCTCCAGGTCCAGCGTGGGGACCTGCAGATGTCCCGACCAGTCCTTTAAGAATTCCAGCAGATTTAGCACAGACGATCGGTCCCTACTCCTTATTAGCC | 4 |
| wt-56880-57000 | 471 | CCTGCTCGAGGACCACTGTCACAAGAAGATAGTCTATCATGCTCAAGGCATCTGCCTCTGGCACTTCCCGGTTAGAGGCCGGGTCGTAGACGATGGCCTGTTCCTGGTAGGTATGTCCGG | 4 |
| wt-57000-57120 | 472 | CTATTCTCGCAATGTTGCTCTCGAGGGGCACAAAGTCCATCTCAGGAGTCTCTATGTCAAAGGTGGTCTGATAGTATTGGCTCCTGGCGGTGTCCAGTGTGATGGGGGACGTGGGGGCAC | 4 |
| wt-57120-57240 | 473 | TGGATCCCGATTCCAGGCTGTTGGAGAACACTTCATCTTCGAACATGTCTTCATCCTCTGTGGTGGGGATATCGGAGGCTAAGTCGCTCTCCGCTTCTTCAGAGTCGGACATGGATAGGA | 4 |

| | | | |
|---|---|---|---|
| wt-57240-57360 | 474 | AAGGCTCCTCTAGGTCAGACAGGTAGCGGACGAGGCCAGAACCCCCAGATGCATCATCCCCAAAGGAGGGCTGCTGCCCGAAGGGAGGTGATGGGGATATCTCCGTTCCAGCCCTGTCAG | 4 |
| wt-57360-57480 | 475 | CGGCCGGGGGATGGTTTTTTTCTGGTTCGAGTGTCGTGGCTGATGGTGGGAGCTGCTGAGCAGGAGGAGGAGCCGGGGTAGCTGATGGCAGGGGCTGCTGCTGAGGAGGAAGAGGAGAAG | 4 |
| wt-57480-57600 | 476 | GAGCCCGGGCGGCTGATGGCGGGGGCTGCTGCTGAGGAAGAAGTGGAGAAGGAGCCGGGGCGGCTGATGGCGTGGGCTGCTGCTGAGGAAGAAGTGGAGAAGGAGCCGGGGCGGCTGATT | 8 |
| wt-57600-57720 | 477 | GCGGGGGCTGCTGCTGAGTTGGAGGAGGAGAAAGAGTCGTGGTGGTGGGGGCTGCTGCTGCAGTCGGGGAAGGGGATGGGGTGGTCAGAGGGATTTTTGGGTTCGAGGGAGCTGCCTGTG | 4 |
| wt-57720-57840 | 478 | GCAGAGGGATGGGTATTTGCAAAGGGAGGCGAGGAGATGGAGTGACTGAAGGAGCGATAGTTGAGACTGGCGCGGGGTGGGGTGTCGGGGAGGCGGGTGGTGATTGGTGAGGGATGGGGA | 4 |
| wt-57840-57960 | 479 | TTACTGGAGGGGGAAGGCGAGCTGCTGAAGGGGGGCGATGGGGCGGAACGTGGGTGCGTGGCAGCTGATCATCCTCTGTGTCAGTGGTGGAGGACAGAGGGAGGCGGCGGCCGGAGGTGG | 8 |
| wt-57960-58080 | 480 | GCTTCTTGTGGGGGCTATCTTTGCCCAATCCCTTTTTCCTCTTGGGAGTCTGAGGCGCTGCGCCGCTCGACGCCCTTGGTGGCGTGGAGGGAGCGGGGACCCCGGGGGTGTGACCTAGGC | 8 |
| wt-58080-58200 | 481 | CGGGGATGGGGATGAAGAGGGGAGGGCTGGAGGCCGGGGCCGCGGAGGCCGGGGCCGCGGAGGCCGGGGCCGCGGAGGCCGGGGCCGCGGAGGCCGGGGCCGCGGAGGCCGGGGCCGCAG | 16 |
| wt-58200-58320 | 482 | AGGCCGGGGCCGCAGAGGCCGGGGCCGCAGAGGCCGGAGACGACGGCGGGGAGTTGGTCTTTGCAGGACTATACCTGGCGGCAGGGAATGAGTCGGATGTGAAAGATCGAGAGGGCAGTG | 8 |
| wt-58320-58440 | 483 | GCCTGAGGTTATACGGTATTATTCGCCGTTCAAACGGTAGCATGACGGGAGGGCTGCTATCAGCACCGGGCGTCCCCGCCGCCTCCCCATCACTGGACACAAGCTCGGGCCCCACCAGGT | 4 |
| wt-58440-58560 | 484 | CAAAGCCGCTGCCGTTGGCCTCATAAAAGTCATACACGCCATAGTGTTCCAGCATAAAGATGCGGGGGTCCTCTGTCTCAAAGGCCTCGGGTAGAAAATAGAGATGCACGCAAGTGTACT | 4 |
| wt-58560-58680 | 485 | GGGCCCCTGGTGCCCCCACGTACTGCAGGATGTCGTGCGCATAGGTGCTGACTCTGACATGGGCGGGGGTGCCCGGGGCCGCATCCTTCTGGCAGTGGGGGTCAAACAAGTAGAAGGAGC | 8 |
| wt-58680-58800 | 486 | CATCTGTCTCGATGATGATGGCCCCCGCGTAGATGTCGCAGATGTAGAGGATGAACTGGGCCACCCCGTTGTAACTGCCGTGCAGGACCTCGGCCAGGGACTGAACAACTGCCGAGTTTG | 4 |
| wt-58800-58920 | 487 | CGATCTGGGCAGGGAATAGGACGAGGCCAAAGATCTCCGCCGAGCGGTATATGTGCACGCGCCCACCGCCCCTCAGGACCACGGAGCTGGGCACGTCCGTCAACTGGGCCATCTCGTGCC | 8 |
| wt-58920-59040 | 488 | CCTTGAGGATGCCGCTCTGGCGCATGAGGGCATCCAGCCGCGCCCCCTCGTCCAGGACCTCGTCCAGCTCAGGGCGGGAGGTCAGGGGGCGGCCGGCCAGGAAGCTCTTGACCAGGTAGA | 8 |
| wt-59040-59160 | 489 | GGACGCAGTTGCTGACGCACTGGATGCCGGCAAAGCGGCCAAACTTGCAGTGGGCCTGGTTGCACGAGGCCGTGCCTAGGATGCGGAGGGCCGAGCCTCCACTCCCGCCCCCGGGGGCAT | 8 |
| wt-59160-59280 | 490 | TCACATCCATGGTCCTGATTCCGCGCACGGGGCCGGTTCCCCGGGTGCGCTGGCTTTGCCCCCAGTCGCCGTTACTCATCTTCGGCGGTGGGGCGGGGAGGACGCCTTGTCGCCCCCCTT | 8 |
| wt-59280-59400 | 491 | CTGGTCCGGGGTCTTACGCGGCTGGCGGCGGCAGCCGCCGAGAGATAAGGGGGGTACGTGTGTGCCTCCGCCTCTCCTCTGTCTGGGCCGCCGCCGCCGCTTGCCCGCCTTGAAGGAGAG | 8 |
| wt-59400-59520 | 492 | GGGGTAGTCCGCGGACTGCGTCTGCGGGGGCAGGAGGTCTCAACCTTCTGGGCTCGGGCCGCGGTGTCGATATCCGATGGCCTTTCCCTGTCTTCCTCGTATGCTCCTTCTCCTCCTCCT | 4 |
| wt-59520-59640 | 493 | CCCGGCACGCCCCTGAGATCTGCCTCCCCTCCCTCTCCCTCGTCCTGGTCGGAAAAGTCTGAGGAGGAGAAGGAGAATGGGGAGGAGTCCAAAACGGCACGCCACCTGCCGTGGGGCGGT | 8 |
| wt-59640-59760 | 494 | GGTGACAGGTCCCGGCTGGCCCGGCGCTTGCTCGCGTTCCTGCCGTTACCCAGGAGAATGGCCGCGAGTTTTTTGGCGGGGAGGATGCGGAATGGCGGGGGCGTTTGTCCCACGGGTGAG | 8 |
| wt-59760-59880 | 495 | GGGGAATCGTCGGTTAGGGCCGGCACGAGGTGGTGGGTCTGGACCCGGGCCGTGCGAGCAAAGGCGGCGAGAACCGAGGGGCTTCTGGGGGTGACTGTGATCTGTTCCGGATTTAGGTCC | 8 |
| wt-59880-60000 | 496 | ATGGCGGGTGTGTATGTTTTAATAGGGGTGGTCTCTGGCGCGGCAGGATGATGGTCGAGGACGTCCACCAGGGCCTTGCAGATGCTCTTGCCTAGATACAGGATGTCGTCCATGCTGAGG | 4 |
| wt-60000-60120 | 497 | GGAGGTGGGGTGTCTGCTCCCCCCTGCGGAAGCCGCCTGGGTGCGGGGATGAAGACAGGTGGTGGGCGGGCGTCTCGCCGGACTATGGCCTCGGCACGCTCGGCGTCGATGGCGGGTGGC | 8 |
| wt-60120-60240 | 498 | TGGAACAGGCGGGCGAATGTGTAATCCCGGAACCGGTAGGCGACGCTGCGCCTGAGGGCGCCCGTCAGGCTGTATCCCAGCTCCAGGGCGTGCTCCACCCGCTCGTTGAGCTCCTCGAGA | 8 |
| wt-60240-60360 | 499 | TCCGGACGCAGGGGCTCGCTGGTGTGGGCCCAGAGGGGGTGATCCGCGATGCCCCGGCTCTCCCTGAGGGCCGGCACCAGGAGGCGCCTTCTGAGGGTGGCCGTGTCGGCCGTGGCCAGG | 8 |
| wt-60480-60600 | 501 | ACCACCGCCTCCTCCTCCTCTTCCGTCGCCAGAGGGATCTGCACCCTCCCGGTCTCTGCGTCGTACAGGAGCGGGCGGGAGCACAGCCTCCAAGCTGCCCCCGTCAAGCGCGAGATGTCC | 8 |
| wt-60600-60720 | 502 | TCCGAGAGGGTCTCACCCGAGACCAGAAAGCGGCGGGTGGCCAGGCCCAACGACTCCGCCGTCGTGCTGTATCTCAGGGTGAAGAGGAGTGAAAAGAGGGAGGTGGGCCAGGCAAGCGGT | 4 |
| wt-60720-60840 | 503 | GGTGCTTCCGCCGCCCGCTCTGAAGCTGAGATAGTCTCGGAGATGATGCCTGAGACCTCTCGGACGGCGTCCATGATCCTAAGGACTGCGTCGTGGGACGACAGCCCCCAGGGGCCCCCG | 8 |
| wt-60840-60960 | 504 | CCCTCTTCGTCTTCTGCACCCTCGGCTCCTGCGTCCCCGGCCTTGCCTTCCCCCTCTAAGTTGAGGGGGCGCAGTCCGACCGCCTGGGGGGACTCCCCAGGCATCGGAGGGGCCCCGTCA | 8 |
| wt-60960-61080 | 505 | TAGATCTCCCAGACGGTGGCGTATATGAGCTCGAGAGGACGGCGGGCCCGGGTCAGCTCGGGGGAAGGGAGGGCCAGGTCGCTGCCGAAGGAGACCAGCCAGCGCAGGGCGGCCAGAGAG | 8 |
| wt-61080-61200 | 506 | CGGGTTTTGGGCAGCTCGTTGGAGAGGACCCGGCGAAGGGCGGGCCAGATTTGGAACTCGATGAAGGCGGCCGGGAAGAAGGGGCTGTGGACATAGGCCGGATCCGCGCGCGCCGTTTGG | 8 |
| wt-61200-61320 | 507 | CCGGCCCTCAGGGACCGGCAGTATGCCTCGACGTCTGTCCGCGGGGCCGCCGCCACCGCTGCCGTCCACTGCCTTCTTCCCTGCTCGCCGGGGAGTAGGGGGGGCTTACAGGGGAGGGCC | 8 |
| wt-61320-61440 | 508 | GGAGCCGGGGCCGGGGCCTGCCACAGGCGGCTGTAGCGGACCCATAGCAGAGACCTGAGGAGTTCGGATGAAAGGTCCCCCGCCACCTGCTCATACTCGGCCGCGGGAGGGGGGACGATG | 8 |
| wt-61440-61560 | 509 | AAGATGCGCAGAGGGGTTACGGCGTCCCAAGGGTCCGCCGCCGCCCCCACACCCACAGCCGTCGCGGCGGGGGCGGCGGCGGGCGTAGAGGGGCCGCTGGTGCGCCGGGCTCGTCTGTCC | 12 |
| wt-61560-61680 | 510 | ACGGCCTCGGCCTCCGCCCGCAGGTAGGCCGCCCGGGCCACACGGGCGAAGCGGCTCGTGGGGCTCGCGGTGGGCAGCAGTCGGAAAAAGTGCAGGGCAAAGCCCGATAGACTCTCTAGG | 8 |
| wt-61680-61800 | 511 | AGGGCGGCGGTGGCCTCGAGCCACCTCCACCGCGAGCGGGACACCCGGGGCACAGAGGCCAGCATCATGGCGTAGTCCCCCGCCACGGTCTCGTTGAGCCCGGCCGAGAGCAGAACCGTG | 8 |
| wt-61800-61920 | 512 | GCCACCTGCTCGATGGCGGCTGGAGAGAAGGATGCCCGGCTCCCCGCCGCCTCCTGCACACGAGCGGCCAGGGCCTCCATCTCTGCCGCCATCCCGGCCAGGAAGGCCTCGATGACCGAG | 8 |
| wt-62040-62160 | 514 | GCGACGGGGGGCGTTTGTCCCCCGCTGGCCTCCGCGTACACGCCCAGGATCTCTACCGCCCGCCGCCGGGCCAGGGGCAGCGAGGCCACCACGCTGGAAAGTGACTCGGGGCGGTGAAAG | 8 |
| wt-62160-62280 | 515 | AGACCACCACCGCTTTCTTCACCCTCTCCCCCGCCGGCCCCGCCCCCACTGTGCTCCACCAGCTCCACGGCCATGGCCTTGATGTCCGCGGCCGTGGGCTGACCCTGCCCTGCAGCCGCC | 8 |
| wt-62280-62400 | 516 | CAGGGGTAGCGGTTGGTCTCCGCGTATACGGTGACCAGCCATCTCCCCAGCGTCGTTTTCGCCGCGTTAAAAGCGTAGAATGACAGCCCCTCCCGCGGGAAGGCGTCCCACCGGGCCAGA | 4 |
| wt-62400-62520 | 517 | TAAGTGTCGGCCACCAGCTCTTCCACGAAGGCAAAGGTGGCCGTTGGGCCAGAGACCGCGAGCACCTCCCCGCTGCCCTCTTCGATGATGCGCCGGTACGCGGCCGCCAGGGCCCGGGTC | 8 |
| wt-62520-62640 | 518 | TCTGCGATGAGCCGAGAGCCGTCCAGCGGATCGTCGGTGGCCGGAGAGGCTGTCGTGGGGGGCAGTGAGGATGCCAGCACGTCCAGGGCCGCCTCCAGATGGCCGAGGCCGAAGCTGCGC | 8 |
| wt-62640-62760 | 519 | CTGGAAAAGGAGGCCGCCCGGAGTAGGTAGTAGGCGTGGTGGCGGAGGACCGCCGCCGGGTAAGCGTGGCCGCTCATGAGGGTGAGAGTATTTAAAAAATCGCGCACCAGCACCGGCTGG | 4 |
| wt-62760-62880 | 520 | GCCAAATCCCCCAGTCCAAAGATCCCCAGCTCCAGAGGCATCAGCGCGCGCAGGCGGGCAGCGGGGTCGTCCCCAGACAGCAGCAACTGACGCGTCACGCGGGCGAGCCCCCCGTCCACC | 8 |
| wt-62880-63000 | 521 | TCTGCCAGGGGTGGCTGGGCGTCTGCCCCTCCGCTACCGCCGCTGCTGTCACTCTCCATAGCGGACGCCATGAAGGTCCAGGGGTCCGTCGATCGCCGCCGTCTGCAACGCCGAATCGCG | 8 |
| wt-63000-63120 | 522 | GGGCTGCTGCCCCCTCCGGCCCGGCGTCTAAATATTTCCCGGGGGTCCGAATTCACGCGGGACGTTCGTGGGCTGGTTGAGGAACACGCGCAGGCCTCCTCGCTGAGTGCGGCGGCCGTC | 8 |
| wt-63120-63240 | 523 | TGGCGCGCAGGGCTGCTGGCCCCGGGGGAGGTGGCGGTCGCCGGGGGTGGCAGTGGAGGGGGGAGCTTCAGCTGGTCTGGGTGGCGGCCGCCAGTCTTTGGGGACTTTCTGATACACGCC | 8 |
| wt-63240-63360 | 524 | AGCTCCTTCAACAACGCCGAGGCCACTGGAACGCCCCTTTTCCAATTCAAGCAGAGTGACCCGTTCTCGGGCGTCGACGCGGTATTCACTCCTCTCTCCCTGTTTATCCTAATGAATCAC | 4 |
| wt-63360-63480 | 525 | GGCCGGGGTGTAGCCGCCCGGGTCGAGGCAGGTGGGGGCCTGACGCGGATGGCCAACCTGCTGTACGACAGCCCCGCAACCCTGGCTGACCTGGTCCCGGACTTTGGGCGGCTGGTGGCC | 8 |
| wt-63480-63600 | 526 | GACCGCCGCTTCCACAACTTCATCACCCCTGTGGGCCCCCTGGTGGAGAATATAAAGAGCACCTATCTGAATAAAATCACCACGGTGGTCCACGGGCCTGTGGTCAGCAAGGCCATCCCT | 4 |

| | | | |
|---|---|---|---|
| wt-63600-63720 | 527 | CGCAGCACCGTCAAGGTGACGGTGCCCCAGGAGGCCTTTGTGGATCTGGACGCGTGGCTCTCCGGCGGCGCCGGGGGTGGCGGTGGAGTATGCTTCGTCGGGGGGCTGGGCCTGCAGCCG | 8 |
| wt-63720-63840 | 528 | TGCCCCGCCGATGCGCGCCTCTATGTCGCTCTGACCTATGAGGAAGCCGGGCCGCGGTTTACGTTTTTCCAGTCGTCCCGCGGCCACTGTCAGATCATGAATATCTTAAGAATTTATTAC | 4 |
| wt-63840-63960 | 529 | TCACCATCCATCATGCACCGCTATGCTGTGGTCCAGCCCCTACATATAGAGGAGCTAACCTTCGGGGCGGTTGCCTGTCTGGGGACATTTAGTGCTACTGACGGTTGGAGGAGGTCTGCC | 4 |
| wt-63960-64080 | 530 | TTCAATTACCGTGGCTCTAGCCTCCCCGTGGTGGAGATTGACAGCTTTTATTCCAACGTCTCTGACTGGGAGGTGATTCTCTAGACTTAACGGGAGGAAACAGGAGGAGGAGGGGGACAA | 4 |
| wt-64080-64200 | 531 | GAGCACAAAAGTGGTTCAGTGGACACCCACCACACAGCATGGCAACGACCAGTCATGTCGAGCATGAGCTCCTCTCCAAATTGATTGATGAGTTAAAGGTCAAGGCCAACTCAGACCCCG | 4 |
| wt-64200-64320 | 532 | AGGCTGATGTCCTGGCCGGGCGCCTGCTCCACCGCCTTAAGGCCGAGTCAGTTACACACACAGTAGCCGAATATCTGGAGGTCTTCTCTGACAAATTCTACGATGAGGAATTCTTCCAGA | 4 |
| wt-64320-64440 | 533 | TGCACCGGGATGAGCTGGAGACCCGAGTCTCTGCTTTCGCGCAGAGCCCGGCCTACGAGCGCATCGTCTCCAGCGGCTACCTGTCGGCCCTGCGCTACTATGACACCTATCTGTATGTGG | 4 |
| wt-64440-64560 | 534 | GGCGCAGCGGGAAGCAGGAGAGTGTGCAGCACTTTTACATGCGGTTAGCCGGCTTCTGTGCCTCAACCACCTGCCTCTACGCGGGTCTCAGGGCAGCCCTGCAGCGGGCCAGGCCGGAGA | 8 |
| wt-64560-64680 | 535 | TTGAGAGTGACATGGAGGTGTTTGATTACTACTTTGAGCACCTAACCTCCCAGACGGTGTGCTGCTCCACGCCCTTTATGCGCTTTGCCGGGGTGGAAAACTCCACTCTGGCCAGCTGCA | 4 |
| wt-64680-64800 | 536 | TCCTCACCACCCCCGACCTCAGCTCCGAGTGGGACGTGACCCAGGCCCTCTATAGGCACCTGGGGCGCTACCTCTTTCAGCGAGCCGGGGTGGGTGTAGGGGTGACGGGGGCTGGCCAGG | 8 |
| wt-64800-64920 | 537 | ATGGGAAACACATCAGCCTCCTGATGAGGATGATCAACAGCCACGTGGAGTACCACAACTATGGCTGCAAGAGGCCGGTCAGTGTGGCGGCCTACATGGAGCCCTGGCACAGCCAGATTT | 4 |
| wt-64920-65040 | 538 | TCAAGTTTTTGGAAACGAAGCTGCCGGAGAACCACGAGAGGTGCCCGGGCATCTTTACGGGGCTCTTTGTCCCCGAGCTCTTCTTCAAGCTTTTTAGGGACACGCCCTGGTCGGACTGGT | 4 |
| wt-65040-65160 | 539 | ACCTGTTTGACCCCAAGGACGCCGGGGACCTGGAGAGGCTCTACGGGGAGGAGTTTGAGCGCGAGTACTATCGGCTGGTGACAGCGGGCAAGTTTTGTGGGCGGGTCTCCATCAAGTCCC | 4 |
| wt-65160-65280 | 540 | TGATGTTCTCTATCGTCAACTGCGCCGTCAAGGCCGGCAGCCCCTTCATCCTTTTGAAGGAGGCCTGCAACGCCCACTTTTGGCGCGACCTGCAGGGCGAGGCCATGAACGCCGCCAACC | 4 |
| wt-65280-65400 | 541 | TGTGCGCCGAGGTGCTGCAGCCCTCGAGGAAGTCTGTGGCCACCTGCAATCTGGCCAACATCTGCCTCCCGCGCTGCCTGGTGAATGCGCCTCTGGCGGTGCGGGCACAGCGGGCCGACA | 8 |
| wt-65400-65520 | 542 | CGCAGGGGGATGAACTCCTGCTGGCCCTCCCTCGACTCTCAGTCACCCTACCTGGAGAGGGGGCAGTCGGTGATGGATTCTCGCTAGCCCGCCTCAGAGATGCCACCCAGTGTGCCACCT | 4 |
| wt-65520-65640 | 543 | TTGTGGTGGCCTGCTCCATTCTTCAGGGATCCCCCACTTATGATTCCAGGGATATGGCCTCCATGGGCCTCGGGGTGCAGGGCCTGGCCGATGTCTTTGCGGACCTGGGCTGGCAGTACA | 4 |
| wt-65640-65760 | 544 | CTGACCCTCCCTCTCGCTCGTTAAACAAGGAAATATTCGAACATATGTACTTTACGGCCCTCTGCACCAGTAGTCTGATTGGACTTCACACCAGGAAGATTTTTCCGGGTTTCAAACAGA | 4 |
| wt-65760-65880 | 545 | GCAAGTATGCCGGGGGGTGGTTTCACTGGCACGATTGGGCAGGAACAGACCTTTCTATTCCCAGGGAAATTTGGTCTCGCCTCTCTGAACGCATTGTGAGGGATGGGCTTTTCAATTCAC | 4 |
| wt-65880-66000 | 546 | AGTTTATCGCCCTGATGCCCACCTCAGGCTGTGCCCAGGTGACGGGCTGTTCGGACGCCTTCTACCCCTTCTATGCCAATGCGTCCACCAAGGTCACCAACAAGGAGGAGGCCCTTAGGC | 4 |
| wt-66000-66120 | 547 | CAAACCGGTCTTTTTGGCGTCATGTGCGTCTGGATGACAGGGAAGCTTTGAATCTTGTCGGGGGCCGTGTCTCCTGCCTCCCGGAGGCTCTGCGGCAGCGCTACCTGCGTTTCCAAACGG | 4 |
| wt-66120-66240 | 548 | CCTTTGATTACAACCAGGAGGACCTGATTCAGATGTCCCGGGACAGGGCCCCCTTTGTGGACCAGAGCCAATCTCACAGCCTGTTTTTGCGTGAGGAAGATGCCGCGCGGGCCAGCACGC | 4 |
| wt-66240-66360 | 549 | TAGCCAACCTACTGGTGCGCAGCTACGAGCTGGGCCTGAAGACTATCATGTACTATTGTCGCATTGAGAAGGCCGCCGATCTGGGGGTGATGGAGTGTAAGGCCAGCGCGGCTCTGTCGG | 4 |
| wt-66360-66480 | 550 | TGCCGCGGGAGGAACAGAATGAGCGGAGTCCCGCTGAGCAGATGCCGCCTCGTCCCATGGAACCGGCGCAGGTTGCGGGGCCGGTTGACATCATGAGCAAGGGCCCAGGGGAGGGACCAG | 8 |
| wt-66480-66600 | 551 | GTGGGTGGTGTGTGCCCGGGGGATTGGAAGTGTGCTATAAGTACCGTCAGCTCTTCTCAGAGGATGATCTGTTGGAGACTGACGGTTTTACTGAACGAGCCTGTGAATCTTGCCAATAAA | 4 |
| wt-66600-66720 | 552 | CGTTTATTGCCATGTCCAAGTTGTTGTACGTGCGTGATCATGAGGGCTTTGCCTGCCTAACGGTCGAAACCCACCGCAACCGCTGGTTCGCGGCTCACATTGTCCTCACCAAGGACTGCG | 4 |
| wt-66720-66840 | 553 | GGTGTCTCAAGCTACTCAATGAGAGGGACTTGGAGTTTTACAAGTTCCTCTTTACGTTCCTGGCCATGGCCGAGAAGCTTGTGAACTTTAACATTGATGAACTGGTCACCAGCTTCGAGA | 4 |
| wt-66840-66960 | 554 | GCCACGACATTGATCACTACTACACCGAGCAGAAGGCCATGGAGAACGTCCACGGGGAGACTTATGCTAACATTTTAAACATGCTCTTTGATGGGGACAGGGCGGCGATGAACGCCTACG | 4 |
| wt-66960-67080 | 555 | CAGAGGCCATCATGGCCGACGAGGCCCTGCAAGCCAAGATTTCCTGGCTCCGTGACAAGGTGGCGGCCGCCGTCACCCTGCCGGAGAAGATTCTTGTGTTCCTGCTGATTGAAGGCATCT | 4 |
| wt-67080-67200 | 556 | TCTTCATTAGCTCCTTCTACAGCATAGCCCTGCTGCGGGTCCGGGGCCTAATGCCTGGCATCTGCCTGGCCAATAACTACATAAGTAGGGATGAGCTGCTCCACACCCGCGCTGCCTCCC | 4 |
| wt-67200-67320 | 557 | TGTTATACAATAGCATGACAGCCAAGGCTGACCGACCAAGGGCCACCTGGATCCAGGAGCTGTTTCGCACTGCGGTGGAGGTAGAGACTGCCTTCATCGAGGCTCGTGGAGAGGGGGTTA | 4 |
| wt-67320-67440 | 558 | CCTTGGTGGATGTGCGAGCCATAAAGCAGTTTCTGGAGGCCACGGCCGATCGCATCCTGGGTGACATTGGTCAGGCTCCCTTGTATGGCACACCACCCCCCAAGGACTGCCCGCTCACCT | 4 |
| wt-67440-67560 | 559 | ACATGACTAGCATCAAGCAAACTAATTTCTTTGAGCAAGAGAGTTCCGATTACACCATGCTGGTGGTAGATGACCTTTGAGTCAGGGTGGCTACTTGCTCAGGTTTCTGGGCATAAATTC | 4 |
| wt-67560-67680 | 560 | TCCTGCCTGCCTCTGCTCTGGTACGTTGGCTTCTGCTGCTGCTTGTGATCATGGAAACCACTCAGACTCTCCGCTTTAAGACCAAGGCCCTAGCCGTCCTGTCCAAGTGCTATGACCATG | 4 |
| wt-67680-67800 | 561 | CCCAGACTCATCTCAAGGGAGGAGTGCTGCAGGTAAACCTTCTGTCTGTAAACTATGGAGGCCCCCGGCTGGCCGCCGTGGCCAACGCAGGCACGGCCGGGCTAATCAGCTTCGAGGTCT | 4 |
| wt-67800-67920 | 562 | CCCCTGACGCTGTGGCCGAGTGGCAGAATCACCAGAGCCCAGAGGAGGCCCCGGCCGCCGTGTCATTTAGAAACCTTGCCTACGGGCGCACCTGTGTCCTGGGCAAGGAGCTGTTTGGCT | 4 |
| wt-67920-68040 | 563 | CGGCTGTGGAGCAGGCTTCCCTGCAATTTTACAAGCGGCCACAAGGGGGTTCCCGGCCTGAATTTGTTAAGCTCACTATGGAATATGATGATAAGGTGTCCAAGAGCCACCACACCTGCG | 4 |
| wt-68040-68160 | 564 | CCCTGATGCCCTATATGCCCCCGGCCAGCGACAGGCTGAGGAACGAGCAGATGATTGGGCAGGTGCTGTTGATGCCCAAGACGGCTTCCTCGTTGCAGAAGTGGGCACGCCAGCAAGGCT | 4 |
| wt-68160-68280 | 565 | CAGGCGGCGTTAAGGTGACACTCAATCCGGATCTCTACGTCACCACGTATACTTCTGGGGAGGCCTGCCTCACCCTAGACTACAAGCCTCTGAGTGTGGGGCCATACGAGGCCTTCACTG | 4 |
| wt-68280-68400 | 566 | GCCCTGTGGCCAAGGCTCAGGACGTGGGGGCCGTTGAGGCCCACGTTGTCTGCTCGGTAGCAGCGGACTCGCTGGCGGCGGCGCTTAGCCTCTGCCGCATTCCGGCCGTTAGCGTGCCAA | 8 |
| wt-68400-68520 | 567 | TCTTGAGGTTTTACAGGTCTGGCATCATAGCTGTGGTGGCCGGCCTGCTGACGTCAGCGGGGGACCTGCCGTTGGATCTTAGTGTTATTTTATTTAACCACGCCTCCGAAGAGGCGGCCG | 4 |
| wt-68520-68640 | 568 | CCAGTACGGCCTCTGAGCCAGAAGATAAAAGTCCCCGGGTGCAACCACTGGGCACAGGACTCCAACAACGCCCCAGACATACGGTCAGTCCATCTCCTTCACCTCCGCCACCTCCTAGGA | 4 |
| wt-68640-68760 | 569 | CCCCTACTTGGGAGAGTCCGGCAAGGCCAGAGACACCCTCGCCTGCCATTCCCAGCCACTCCAGCAACACCGCACTGGAGAGGCCTCTGGCTGTTCAGCTCGCGAGGAAAAGGACATCGT | 4 |
| wt-68760-68880 | 570 | CGGAGGCCAGGCAGAAGCAGAAGCACCCCAAGAAAGTGAAGCAGGCCTTTAACCCCCTCATTTAACACCATGTTCTCGTGCAAGCAGCACCTGTCCCTGGGGGCCTGTGTCTTCTGTCTC | 4 |
| wt-68880-69000 | 571 | GGCCTCCTGGCCAGCACCCCCTTCATTTGGTGCTTTGTCTTTGCCAACCTGCTCTCTCTGGAGATCTTCTCACCGTGGCAGACACACGTGTACAGGCTTGGATTCCCGACGGCATGCCTA | 4 |
| wt-69000-69120 | 572 | ATGGCCGTCCTCTGGACGCTGGTACCCGCCAAGCACGCGGTGAGGGCCGTCACTCCAGCCATCATGCTGAATATTGCCAGCGCCTTGATCTTCTTCTCCCTCAGAGTCTACTCGACCAGC | 4 |
| wt-69120-69240 | 573 | ACGTGGGTTTCTGCCCCCTGTCTCTTTCTGGCCAACCTGCCTCTCTTATGCCTGTGGCCCCGGCTGGCCATCGAGATTGTTTACATCTGCCCGGCTATACACCAAAGGTTCTTTGAACTT | 4 |
| wt-69240-69360 | 574 | GGGTTGCTCTTGGCCTGCACCATCTTTGCCCTGTCCGTGGTCTCCAGGGCCCTGGAGGTGTCGGCTGTCTTCATGTCTCCATTTTTCATCTTTCTGGCTTTGGGCTCTGGAAGCCTGGCC | 4 |
| wt-69360-69480 | 575 | GGTGCTCGGCGTAACCAGATTTACACCTCGGGTCTCGAGCGGAGACGCAGCATTTTCTGCGCCCGGGGAGATCATTCGGTGGCATCCCTGAAGGAGACCCTCCATAAATGCCCGTGGGAT | 4 |
| wt-69480-69600 | 576 | CTGCTGGCCATCTCTGCCTTGACCGTTCTTGTCGTCTGTGTGATGATTGTGTTGCATGTGCACGCAGAGGTGTTCTTTGGACTCTCTAGATACCTGCCCCTCTTTCTCTGTGGGGCGATG | 4 |
| wt-69600-69720 | 577 | GCCTCCGGGGGGCTGTACCTGGGCCATTCCAGCATCATTGCATGTGTCATGGCCACCCTCTGCACCCTGACATCTGTTGTGGTATATTTCCTCCATGAAACCCTTGGACCCCTGGGCAAG | 4 |

| | | | |
|---|---|---|---|
| wt-69720-69840 | 578 | ACCGTGCTGTTTATCTCAATCTTTGTCTATTACTTTAGCGGGGTAGCGGCCCTGAGCGCAGCTATGCGCTACAAGCTTAAGAAGTTTGTGAACGGACCCCTGGTCCATCTCCGTGTGGTA | 4 |
| wt-69840-69960 | 579 | TACATGTGCTGTTTTGTCTTTACTTTTTGTGAATATCTGTTGGTGACATTCATTAAATCCTAACGACCGGAGTCCTGTCTCTTTGTGTTCTTGGGGGACTTGAGTTAGCTGTCTTTCCTC | 4 |
| wt-69960-70080 | 580 | TTATTACATTGGGCTAACGGGAGGAAATGAACCCAGGGGTGGCAGTGGATGGGGTCATTTATGGGCAAAACTCACAGGACATGTTTGGGGAGTTAGCATTGGCGTCGGGAAACACAGCTC | 4 |
| wt-70080-70200 | 581 | TGGCAGTTATAACCGCACCAGCTAACAGGACATGTTTGGGGGAGTTGGCATTGGCGTCAGGAGACACGGCTCTGTCAGTTATCACCGTACCATGAGTGCCATGTGTGTCCAGTGCCTAAT | 4 |
| wt-70200-70320 | 582 | CACCGTTCCTCATTTTGTGTGCCTCCTCAAATGTTCCAGAAGTCGGCCACAGGGGAGGTGGCTGAATTAGGGCCTTTTCCCTCATTCCCCCATGAGACCCACGTGGCAGGCCTAGGGGCT | 4 |
| wt-70320-70440 | 583 | ACATTCGCCTCCCACGTTTCCCTTCGCGTGAGGCATCCGATATGACTGAATTTTCGCAGTCTCTTTTCCCTCTTCCCTTGTTATTCCCATAGAATTACAGTGAGGTTACACAGGTGGAGA | 4 |
| wt-70440-70560 | 584 | TTCAGTTTAACCATTTATTGATTTAATCCAGGAACAAAAAACAGTCCTAGTGACCCAGTGCCCGGAGAGAGAATGGCCCTGACAAGTCGGCTGCATGATGCACTTCGGCAGTCACGTGTG | 4 |
| wt-70560-70680 | 585 | TGAGTCTCCACGGCCTCTGTCAAAAGGGAGCTTAGCGTGCCAGGGTTGTAATTCTTGATGTAGTGGCCCAGGAATTCAACTTCATCGTGTCTCCGTCTGCAGTTGGCGTTAATGTAGGCT | 4 |
| wt-70680-70800 | 586 | GGGGCTACTGCCGCATATGCTGCCAAGAGACAGAGGGGCTGCTTCACATATGAGCTGCTCAGGGTCTCCACCACCTTGTTTTGACGGGCCGTGGCACAGGTGATGTAGAAGAGTTGCTTC | 4 |
| wt-70800-70920 | 587 | ACAAAGTTGTAGTCTCGCGTGTTAGGAAGGAAGCAGGGTGCCAGCTCTTTGAGCTTGGTCAGGATCACCTTGCTAAGACTCATGGCGCAGGCCAGGAGGATGTCTTCCGCGGGAGCTAGG | 4 |
| wt-70920-71040 | 588 | GGCAGGTCGCCGTGGTAGGTGATCTCCTGGAGCCAAAAGATGGTCTCTTCTAGCATGGCCACCAGGGTGCAGAGCCCCGCGTTCTGGATCGCCTGCATGCGTGCATCCAGCCATGTGTCC | 4 |
| wt-71040-71160 | 589 | TTGTTGGTTGACTTGGTGAAAAACTCACGTAGTGTCTTGTAGCTCCTGCGCAGCTCGTGTCTGGGTTGCACTTTCTGCCAGGCTCCAATCTCTGGATGGGCGGCCACCGCCAGCATCGAC | 4 |
| wt-71160-71280 | 590 | TGTAGGAACGGGTCTTGGATGGGCTCTAGGGTCAGAGAGGCCAGGGGGCTGGGCAAGGTGACAAATGTAATCTTGGAGACAGGCTTAACCAGACTCATGTCAAACCACGGTTTGTTCGGC | 4 |
| wt-71280-71400 | 591 | AGGGGCCTCTGGCTGCGTTCTTGCCTCGCCTGCTTCCTTGTGCTCCTGCCGGCCCCTCGAGATTCTGACCGGGGACCTCTGGTTGCTCTGTTGCTTCGGGGAGCTCTTGGAGACCTCGGT | 4 |
| wt-71400-71520 | 592 | GCTCTAGGCACCCTGGGGGCCCTTGGGGCTCTGGGCGCTCTTGCTCCCGGGGGCAGGTGTCGGCGCTTGCCATAACTTTCATCGGTGCAGCCATGGACCTCTCCGCGTCGCCTTTTGTGG | 8 |
| wt-71520-71640 | 593 | CCTCTGGTGTAAGAGGAGTTGCCAGTCTCCTCCTTCTCGTCCTCGTCCCTGCACAGGGGTGAGCGATGCAATGTGACTGTCTTGTCCTGTAGGTCCCACTTCTTTCTGGGAATCACAAAC | 4 |
| wt-71640-71760 | 594 | GATGCCGAGGTAGGGGTTATGACCACGCTGGAGGGCCGTGCAGGTATGGCGTGGGCCGGAGTTGGATCTTCATCCTCCTCCTCTGAGGATGAAATCTCTCCATCTGTGGAGTGTTCTTCG | 4 |
| wt-71760-71880 | 595 | CTGCCCTCCATAGGGTCCAGATCGCAGTCTGTGTTGGTGTCTGAGACCGCTTCGAGTTCCAGAATGTGGCTCTCTGCAGAGGGGAGACAAAAGGTGGAGACTGCCTTGAGCACCTCTGTC | 4 |
| wt-71880-72000 | 596 | TCAGGCACCGGATGCCCCCGGCTCCACGGCCCCGGCCACTGGCCGGTGTAGCTTCTTACCTGCGGGATCCTCGTTGGAGGAAATGCTGCTAGTTCGGGAGAGTCTCTGAGAAGGAACCAT | 4 |
| wt-72000-72120 | 597 | CTTGTCTGTCTCTACGACGGGCTAGCTGGGATGTAGTGCTGTCTTGACTGGCCTCAGCCCTATTTATGATTCTGGAGGCGGGCACGCTGATGGAGAAATGGGCGGTCGGTTGATTGGCCC | 4 |
| wt-72120-72240 | 598 | CACAGCGACCGGCGAAGCACTGACTCATGAAGGTGACCGTGATGGCCTGTGATGTGTAGTAGAGTACCAGAAACACCCTCACATTCTTGGAGCTGGCCCTGTGGGTATGCCTCAGGCACG | 4 |
| wt-72240-72360 | 599 | CAAAGTTCCTGCCCCGGGCATGGCACACCTGAACTAAGTTTGGCCCGGTTTGCTCAAACGTGACATGGAGAAACTGGGGGAATTTGTCTTCTGGCACAGCTGTTGCCAGGGTGCTCATGA | 4 |
| wt-72360-72480 | 600 | GCGAGGGCCAGATGCAGGAGCTGACCCAGGCGACGAGATCCAGGCCCAGATGTCCCTCTATCATGGCGCAGACATTCTCCACGGTGGGGGGCAGGGTCTCGCGGGTCCTCTGGATTAGAT | 4 |
| wt-72480-72600 | 601 | AGTCACGCCCATCATCCGCGATGTGGTAGCAGAAGGTTTTGGGGGCCGGCCAGCCCACGTGCAGTGAGTGATGTAAGAGGTTTTGAATGTTGAGGGCATTCTTAACATAGCTGTGCTTGT | 4 |
| wt-72600-72720 | 602 | CTTCCTCTTCCGGATGACAGACAAAGAGGCGCAGCTGCCGGCTAAGACCACCGCCCCTGTCCACCTTGTAGGTATGCGGCAGCCGGATGCACCGCCCGGCGTGATACACGCCGCTGTCAA | 4 |
| wt-72720-72840 | 603 | AAAGCGGGGCCCCAATCTCTTTGATCTTGTGACGCATGCGGCGCAGGCAGGCCGTTAGGCCCATGAGCTTCTGCAGCACAGACACAAACCCTTGTACTGCGCTTGTTCCCACAATAGCAT | 4 |
| wt-72840-72960 | 604 | GGCCTCTAGGTAGGGGGGTGATGACGCGAAAGCCCAGTTTTCCCGTGCATATGCAAAAGGGGAGCACATCTTCCATATTATCCGGGTCGGCGGGTGGACAAGCTGATTTGAAAAAATAGA | 4 |
| wt-73080-73200 | 606 | CCAGGTCCAAGACCAGGTTGCAGACCGGAAGCCGGGGATTAAAGTATTCATGCCGGGAGACAAAGAGCTGCTCGCTCAGGCTGTTCTGTGAATAGTACACTGGGGTGTAGGAGAGGGCCC | 4 |
| wt-73200-73320 | 607 | TGGTGAGACACGTGTCTGGGAGGCGGCAGTTGGTCGGGGTGGAGACGACCTCCGCCAGGTGGGATGAGAAGGGGTCAGCGGCTGTCATTACAAAGTAGTGCCTGTCTGCAAAATGGCAGA | 4 |
| wt-73320-73440 | 608 | GGAAGACCGGTAGCCGCTGCACCCTTCGAAGGACGGTGGGTGGGAGGAATTGTTCCTTGGGATTCCACTGGCCCCGGCAGGTGGCCTGGCCGGCCAAGCATAGAAACCCTTGAAGCGTGG | 4 |
| wt-73440-73560 | 609 | GGGGGTATGTGGGACCCTCATCCGCGTGCCAGCGCGCGAGCTCCACCAGCTCCCGGGCCACGTCCACACTGAGCCCGGCCCAGGCCCGCATGAGTCCGTCATCGGGGTCGGGGTCCCACG | 8 |
| wt-73560-73680 | 610 | TGTATGGGGCCGGGGGCTCCATGCGGATTTTCAGCTGCTGGACACGCACATGCTCAGCCAGGTAAGTCTCCCGGGTGAAGTAGGTGCGCATGTGCTCCGCAAAGCCCCTGTCCAGGAGCG | 4 |
| wt-73680-73800 | 611 | AGGGGAGCACGACGCCCCCCGAAGGCAGACACCCAATTTCTCCCATGCTCGTTAACTGAGAGTATCGCTTAAAGGTTCCCTCGTTGAAGCACTGTGCGTGGGCCAAATAGACGTAGCGCA | 4 |
| wt-73800-73920 | 612 | CGAGATCGGCCGAGGCCAGGGGAAGGCGCCCCCTGTAGGCGTCTATCGTCCTTGCCACAGCGCGGATCTCTCGCGAGTCCCGCCGCAGTTTCTCGTGTGCAAAGTGGGCAAAAGCCTCGG | 8 |
| wt-74040-74160 | 614 | TCGTACAAGGAATGCCCCAAAAATAGTAAGCAGCCGTGACTAGCACGAACTGGGCCTCGTGGGAGCCAAAGGTGCTAATGAACCACCTGGCCGAGATGTTAACGCGGTAGATGCGGCGCA | 4 |
| wt-74160-74280 | 615 | GACAGCCCACGATCTTGGGACGCAGCCACGCCACGCGGCCTCTGGCATCCCCCTGTGGCTGTTTCTTAGCGCTCAGTGTGAGCAGTTCCACGAGGGGCGTGAGCGAGCGCAGGGCCCCCG | 8 |
| wt-74280-74400 | 616 | CGCGATCTAGGTAGGTGGATAGACGGTCCGCGGTGAGCGGCGTGAGGCCGCGCAGGAAGGGGAAGGCCTCCTCCGCCGGCAGGTGCAGCGTCAGAACCAGGCCGCAGCGGCTCTGTGAGG | 8 |
| wt-74400-74520 | 617 | TCAGCCGCTTCTTGGGCAGGTGAAGCTGCAGTTCCACGAGAGAACCCGCCACGTGGTGGAGGGGCGAGGCGTTGTGGCACAAACAAAACAGGCGGAAGCCCTCGTCAGGCCGCGAGAGGA | 4 |
| wt-74520-74640 | 618 | TGGCATCGAGGATGGCCTCCGCAATGTCAGTGTTTGAGGCCACAAGGGCCTTGATGACGACGGGGGCGGACATTATTTAAGACCGGGAGGCCCCAACGGCGGGCTAAACAGAACGATGGC | 4 |
| wt-74640-74760 | 619 | CTTCTATCTCCCAGACTGGTCGTGCTGCGGGCTCTGGCTCTTTGGCCGGCCCAGGAATAGATACAGCCAGCTCCCTGAGGAGCCGGAGACCTTTGAGTGCCCGGACCGCTGGCGAGCCGA | 4 |
| wt-74760-74880 | 620 | GATAGATCTGGGCCTGCCCCCTGGTGTGCAGGTGGGAGATTTGCTAAGAAATGAGCAGACGATGGGCTCACTGAGACAGGTTTATTTGCTCGCTGTTCAAGCCAATAGCATCACGGATCA | 4 |
| wt-74880-75000 | 621 | CCTGAAGCGCTTTGACGCCGTCCGCGTCCCTGAGAGCTGTCGTGGGGTGGTGGAGGCCCAGGTGGCCAAGCTTGAGGCCGTGCGCTCAGTCATCTGGAATACCATGATCTCTCTGGCTGT | 4 |
| wt-75000-75120 | 622 | AAGCGGCATCGAGATGGACGAGAATGGGCTCAAGGCCCTGCTGGACAAACAGGCTGGCGACAGCCTGGCCCTGATGGAGATGGAGAAGGTGGCCACGGCGCTCAAGATGGACGAGACCGG | 4 |
| wt-75120-75240 | 623 | TGCCTGGGCGCAAGAGATCTCGGCCGTTGTCTCATCGGTGACCGCCCCCTCAGCCTCGGCCCCTTTCATCAACTCCGCCTTTGAGCCCGAGGTGCCCACCCCCGTCCTTGCACCGCCTCC | 8 |
| wt-75240-75360 | 624 | CGTGGTGCGGCAGCCGGAGCACTCTGGGCCCACGGAGCTCGCGTTAACGTAGCAACCAGACTCCACACCAAATAAACATTTTATTGGTAAAACAAGGGATATGAAGGTGTCATTGACCCG | 4 |
| wt-75480-75600 | 626 | CAAGGACCCCCGCGTCACTGTGGGGGCTGTAGAAGGAGGTGAGGTGGTGCTTGTGAAGGTAAACAAGCTGACAGAAGCGCCGGTACTTGTTAAGGAACACGGTCTGGTCACTAAAGTTGG | 4 |
| wt-75720-75840 | 628 | CGGTGAGTGTCACTGAGACGGTCTGATGGGGGAAGAGGGCCAGGTCCTTTGGCAAAGAGACGTCCAGGCCCACGTCCCCGGGGTACTGGGGGTGGTTGATGGGACCCTTGTCCTCCTCCA | 4 |
| wt-75840-75960 | 629 | TCTGGGGGGTGGCATATCTGAAGGCAGCCAGGTGGATTTTGAGCTCCGATGGACGCAGCGTGGAGTTGTAGCGCCGCTGATTCTGGAGGATTAGCCGGAGTTCCCCCGTGTAGCCGGGAT | 4 |
| wt-75960-76080 | 630 | CGATGATGCCAACATGAGACGTGACCGGACGGGAGGTGCTGCCCCACAGCATGAGCCCATGACCCTCGGGTGGGCGGGCATAGAGGCCTAGGTCCACAGTTGTGGTCTTCATCGGCGCA | 4 |
| wt-76080-76200 | 631 | GCAGGATGGTGGTCTTGTTGACCAAGGTGAGCCGCCCTACACTAGCCTGCTGGAGCAACAGCTTGTCATTCTGGAAGGCGTAGCGTATGTGTGGACAGGCCTCCATGGTGATGATCTAAC | 4 |
| wt-76200-76320 | 632 | AGACAGGGACGGCGGCGCTATATATAAGAGCCCAAGACCCGGCTCTCTTTACTGCGAAATGGGGAAGGTCCTAAGAAAGCCGTTTGCAAAGGCTGTGCCACTGCTCTTCCTCGCCGCCAC | 4 |

| | | | |
|---|---|---|---|
| wt-76320-76440 | 633 | CTGGCTTCTGACCGGGGTGCTGCCGGCCGGCGCTTCCAGTCCCACAAACGCGGCGGCGGCTTCCCTGACTGAAGCCCAGGACCAGTTCTACTCCTACACATGTAATGCGGACACATTCTC | 4 |
| wt-76440-76560 | 634 | GCCTTCTTTGACCAGCTTTGCCTCCATCTGGGCACTTCTGACGCTTGTCTTAGTCATTATAGCCTCAGCCATCTACCTGATGTACGTCTGCTTTAACAAGTTTGTGAACACGCTGCTGAC | 4 |
| wt-76560-76680 | 635 | GGATTAGATGGGGATATTTAAAAGGGGCAGCAATCTCGGCTGTTTGTACTTCTTCTCTGCTCGTTAAACCAATAGCATGTCAGCTCCACGCAAAGTCAGATTGCCTTCTGTTAAGGCTGT | 4 |
| wt-76680-76800 | 636 | TGACATGAGCATGGAAGACATGGCCGCCCGCCTGGCTCGCCTGGAGTCTGAGAATAAGGCTCTGAAGCAACAGGTCCTCAGAGGGGGTGCCTGTGCCTCGTCTACCTCTGTTCCTTCTGC | 4 |
| wt-76800-76920 | 637 | TCCAGTGCCTCCGCCTGAGCCGCTTACAGCTCGACAGCGAGAGGTAATGATTACGCAGGCCACGGGCCGTTTGGCGTCTCAGGCTATGAAGAAGATTGAAGACAAGGTTCGGAAATCTGT | 4 |
| wt-76920-77040 | 638 | TGACGGTGTAACTACCCGCAATGAAATGGAAAATATATTGCAAAATCTGACCCTCCGCATTCAAGTATCTATGTTGGGTGCAAAAGGCCAACCCAGCCCTGGTGAGGGAACACGACCACG | 4 |
| wt-77040-77160 | 639 | AGAATCAAACGACCCCAACGCCACCCGACGTGCCCGCTCCCGCTCCCGGGGACGTGAAGCAAAGAAAGTGCAAATTTCTGATTAATAAATTTTTATTGACTTTATACATAGGTCTCGGCG | 4 |
| wt-77160-77280 | 640 | TCATCATATGGTGGGGTGGTGTAGGTATGGGATGTAGACAAGTTACGCCTAAAGGCGCAGTCCGCCATGACCAGCAGCAGCAGAAGGGTCAGCACAGCCAGAGAGGCCCACTGCAGTACT | 4 |
| wt-77280-77400 | 641 | AGCATGGAGAGGTTTGAGAATCTGGGCTGGGACGTTGGCGGGACTGGCACGGTGGCTTGGGCTGTGGTAACCGGTGGGCTCGTAAAAGTCCAGCGGGGCCGCAGTTTGCTAGAAGTGCTG | 4 |
| wt-77400-77520 | 642 | GGAGGTAGATAGGTGGTCGCATTGTATCTCGGTCTTGGCGTAGTTGAATCACCGCCGTAATCTGTGGTGGGCTCTGTACTTGTCCGGGCTCCATGTCCTGTGGTGTGCTTTCCACCGGTG | 4 |
| wt-77520-77640 | 643 | GTAGAATTGGCCTTTCCACCTGTTGAGGTGACCGTGGGAACCGCCGTCTTTTGGCCACTGGGGGCCTGGGGCGACGTTGCATTTTGGGGGGGCGTGCCTTTGGTGACATTAACCTCCCCC | 4 |
| wt-77640-77760 | 644 | GGTTTTGTGGATGTGGAACTGTTTCCAGGGCCTGACGCTTGGCTGGTGGTGCCTGGGCGGGGTGCTGGCGAACTGGTGGACACATGATGTGTGCTGATAGAGGCTGGTGTCACCTGTGTT | 4 |
| wt-77760-77880 | 645 | ATATTTTCACCACCTGTTGGGTGAGCGGAGGTTAGTAAAGGCATATGTGACGTTGAATTGTCACTGGTGGAGGGGCTGAGTGTCTCTGGGTTTGAACTGGGTCTCAGTGACATGGAAGAG | 4 |
| wt-77880-78000 | 646 | GTTGAACTTGAAGTTATGTTATGTTGGCCTGTGGTAACAGCACTGGTTGCATTTTTTGGTTGGCTGGTAACTACTGGGGTGGGACTTGTTCCTCCTAAGGTGTGGTTGGTGGCATTTGCC | 4 |
| wt-78000-78120 | 647 | TGTGGACTTGTTTCTCCCACAGTAGGGCCGGTGGCATTTGGGGTTGGGGTAGTCACTGCTGAGGTGGGGCTTGTTTTTCCCAAGGTGGGGCTGGTGGCATTTGGGGTTGGGGTAGTCACT | 4 |
| wt-78120-78240 | 648 | GCTGAGGTGGGGCTTGTTTTTCCCAAGGTGGGGCTGGTGGCATTTGGGGTTGGGGTAGTCACTGCTGAGGTAGGACTTGTTTTTCCCAAGGTGGGGCTGGTGGCATTTGGGGTTGGGGTA | 4 |
| wt-78360-78480 | 650 | TTACTTTCTGTGCCGTTGTCCCATGGAGATGGACTTGGTGTCACCGGTGATGCGCCTGACGTTGTGCCGGCTGGTGTTGGGCTGGTGACATCCGCGGTGGATACAGTGGGGCCTGTGCTT | 4 |
| wt-78480-78600 | 651 | GCAGGTGCGGTGAGGTTGGTAGGCACGTGAGTAGAGCTGGGTAGACCTGTCGTTGTATTGGGATCAGCAAATCCAGTTGTATTCAAGGTAGGGGAGGTGGTGGTGCTCTCGGGTGCCTTG | 4 |
| wt-78600-78720 | 652 | GAGAATATAACCTTGTGGGTTGTTGTGGTGGCATTGGTAGCCGTTCGTGTGATAATGAGTGTCTTGGGGGCCGTGCCAAGACCCGAGACAGTAATGTCAAATGTCCGATTGCTCGCAAAT | 4 |
| wt-78720-78840 | 653 | GCACCAGAAATATTTTCACAACCCGAAGGTGTCCCCGAGGTGAGAGTCCATTTGCACTTAAAGTCAGTTTCAGTGTTGTTTGGCCAGGCCCAAAAGGCAGTCACTGTAACATTTGGCGAG | 4 |
| wt-78840-78960 | 654 | TTTGCGTCCTCAGAAGTGACCATTGGCACTGAATAGGTAGCATTGTCACCCACATATGTGATGTCTGTGGTGTTTGTCGGCATGTCCTGTGAAGCTGGAATCTCATCAGAGAACACAATG | 4 |
| wt-78960-79080 | 655 | TTGGACTGAATGCAGTAATCTCCCCCGCTCGCCTTCGGTCCATTCCCAGAGTAAAACACGTACAGGATACTGTTATTGCCAAGAAATCGTGACACTGGACGTGGTGTCAGACGCAGGCTG | 4 |
| wt-79080-79200 | 656 | TATGCATACCCTGTACCAGGTATTGGGGTGGCCACGGGACTCGTTGATGTGAGAATTCCGCCGCTGGGAACATGGCTCTCGTATCCACTGCAGGTGATGTTAAATTTGTTGTCTCCGGGC | 4 |
| wt-79200-79320 | 657 | AGAACTTGTGAAATTTCGCCATCCTCCATAATACACTCAATATCTATCTCATTACCGAGCATTTCTGTTTTTACGCTGAAATTCGAGTCTTGAGCTGACGTTGGCAAACTTAAGGGTAGC | 4 |
| wt-79320-79440 | 658 | GTGACATCCAGCCCCTGTGCCCTCACTACTGCCGTTATATTGGTAGAATTACAGTTATCCCACTTTATGTATGGCACTGTTTCTGGTATCAGGTACACGGGGTTTTGCATTTCTGCATGG | 4 |
| wt-79440-79560 | 659 | TGGCACCACATGGTTCCAAACACATCTTGAAAGTAGACATCTACAGATTCCAGGCTTACTTGTTGCTCCTCTCCGGTGGTGACGTTAATTGGAAGCTTCTTAGACCGCATAGTTAGAGCC | 4 |
| wt-79560-79680 | 660 | AATTCTCCTGCACCAAGGAGCTCCAGTAGAAAGAGATTGGTGGCATTTTCTGAGCCACCAAATGCACCTCGAGGTTGGTAGACAGCCTTCGTATGGGGTGTCAGCTGGCCAAAGTCAAGA | 4 |
| wt-79680-79800 | 661 | TCAAGTTGATGCTTTTTGCCCCCGACATCGAAATTGATAGTTACATTGACATCTGCCGTGCAAACATTGCATGTGGGGTAAAATGGGAATTCCGGAATCTCAACATTGAAAAAACCAGGA | 4 |
| wt-79800-79920 | 662 | TCTTCACCCGTGAGATGGATCAGGCTCTGGATGGTGTACTGACACACAAGCAAGGCTGCCTCCATTGTCTCGGCACCGATTTCTAGGCAGCATCCTCTTTAATAGGTACAAGGGGGGTGC | 4 |
| wt-79920-80040 | 663 | GGTGTTGGTGAGTCACACTTTTGTTGCAGACAAAATGGACAAGGACAGGCCGGGTCCCCCGGCCCTGGATGACAACATGGAAGAAGAAGTCCCATCTACCTCGGTTGTGCAGGAACAGGT | 4 |
| wt-80040-80160 | 664 | ATCGGCGGGAGATTGGGAAAATGTCCTCATAGAGTTATCAGATAGCAGCTCAGAAAAGGAAGCAGAAGATGCCCACCTGGAGCCGGCCCAGAAGGGTACGAAGAGAAAGCGGGTCGATCA | 4 |
| wt-80160-80280 | 665 | TGATGCCGGTGGGTCAGCTCCAGCACGACCCATGCTCCCACCCCAGCCGGATCTCCCTGGGCGAGAAGCCATTCTCCGCAGGTTTCCACTAGATCTAAGAACACTTCTTCAAGCGATTGG | 4 |
| wt-80280-80400 | 666 | AGCCGCGGCTACGGTGAGCATCCCTATGGCCTAAGTGTGTGATGTGTGTTTTTACCCATCACACAACAACAAGGTAAGTAATTTGTTGCCGTTGGTTTCAGCGCATCGACACACGAGCCA | 4 |
| wt-80400-80520 | 667 | TAGACCAGTTTTTCGGATCCCAGATTTCAAATACCGAGATGTACATAATGTATGCCATGGCCATTCGACAGGCCATTAGAGATCGTCGGAGAAATCCAGCTTCTCGTAGAGATCAGGCCA | 4 |
| wt-80520-80640 | 668 | AATGGAGACTGCAAACCCTGGCCGCCGGATGGCCTATGGGTTACCAGGCATACAGCAGCTGGATGTACAGCTACACCGATCACCAGACGACTCCCACATTCGTACATCTCCAGGCGACAC | 4 |
| wt-80640-80760 | 669 | TTGGGTGCACAGGTGGCCGTAGGTGTCACGTGACCTTTTCCGCCGGCACCTTTAAGCTGCCGCGATGTACCCCCGGGGATCGCCAGTGGTTGTATGTTCAGAGCTCCGTGGGTAACATTG | 4 |
| wt-80760-80880 | 670 | TACAGAGCTGTAATCCCCGCTACAGTATTTTCTTTGACTATATGGCTATACACAGGAGCCTCACGAAAATCTGGGAGGAAGTTTTAACACCTGACCAGCGTGTTTCATTTATGGAATTCC | 4 |
| wt-80880-81000 | 671 | TAGGATTTTTGCAGAGAACGGATTTGTCCTATATCAAGAGCTTTGTCAGCGATGCCCTGGGCACCACTAGTATCCAAACACCGTGGATCGATGACAATCCTAGCACGGAGACGGCACAGG | 4 |
| wt-81000-81120 | 672 | CTTGGAATGCCGGCTTTCTCCGGGGTCGTGCGTATGGGATAGACTTGCTTAGAACTGAAGGGGAACATGTCGAAGGTGCTACCGGTGAAACGCGAGAAGAAAGTGAGGACACGGAGAGCG | 4 |
| wt-81120-81240 | 673 | ATGGAGATGATGAAGATCTTCCTTGTATAGTGTCCAGAGGTGGACCTAAGGTCAAACGACCCCCTATATTTATAAGACGTCTGCACAGGTTGCTGCTGATGAGAGCGGGCAAACGAACAG | 4 |
| wt-81240-81360 | 674 | AACAGGGCAAGGAGGTACTGGAAAAGGCCCGTGGGAGCACTTATGGCACACCTAGGCCGCCTGTTCCGAAACCAAGACCAGAGGTCCCACAAAGCGACGAGACAGCTACCAGTCACGGGT | 4 |
| wt-81360-81480 | 675 | CGGCGCAAGTCCCAGAACCCCCAACCATTCACCTAGCAGCTCAGGGAATGGCATACCCATTACATGAACAACACGGCATGGCCCCGTGTCCGGTAGCACAGGCCCCACCTACGCCCTTGC | 4 |
| wt-81600-81720 | 677 | CTGCGGGGCAGGCCTTTGCACCCGTTAGACCACAACACATGCCAGTAGAACCCGTCCCTGTCCCGACAGTGGCACTTGAGCGACCAGTTTACCCCAAGCCAGTTCGTCCGGCACCTCCTA | 4 |
| wt-81720-81840 | 678 | AGATTGCTATGCAGGGCCCCGGGGAAACTTCTGGCATTAGACGCGCGCGGGAGCGTTGGAGGCCCGCACCTTGGACGCCAAATCCACCCCGTTCTCCCAGTCAGATGTCCGTGCGTGACC | 4 |
| wt-81840-81960 | 679 | GTCTGGCTCGTTTGCGTGCTGAGGCACAGGTCAAACAGGCTAGTGTTGAGGTGCAGCCCCCCCAGTTGACCCAAGTATCCCCTCAGCAACCAATGGAGGGGCCGTTGGTACCAGAGCAGC | 4 |
| wt-81960-82080 | 680 | AGATGTTCCCTGGTGCCCCCTTTAGCCAGGTTGCTGATGTGGTCCGGGCACCTGGGGTACCGGCGATGCAGCCACAGTACTTTGACCTCCCCTTAATTCAACCCATTAGCCAGGGGGCAC | 4 |
| wt-82080-82200 | 681 | CCGTGGCCCCGTTGAGGGCTAGTATGGGCCCGGTACCTCCGGTACCGGCAACACAGCCACAGTATTTTGACATCCCCTTAACTGAACCCATTAACCAGGGGGCATCCGCGGCCCATTTTC | 4 |
| wt-82200-82320 | 682 | TCCCTCAGCAACCGATGGAGGGGCCGTTGGTACCTGAGCAGTGGATGTTCCCAGGTGCCGCCCTGAGCCAGAGTGTTAGGCCAGGGGTAGCGCAGTCACAATATTTTGACCTCCCCTTAA | 4 |
| wt-82320-82440 | 683 | CTCAACCCATTAACCATGGGGCACCCGCAGCCCATTTCCTCCATCAGCCACCAATGGAGGGGCCGTGGGTACCCGAGCAGTGGATGTTCCAAGGTGCCCCCCCTAGCCAAGGCACTGACG | 4 |
| wt-82440-82560 | 684 | TGGTCCAACATCAGCTGGATGCTTTGGGGTATACACTCCATGGTCTTAACCATCCCGGGGTTCCCGTGTCTCCTGCCGTTAACCAATATCATCTCAGCCAGGCTGCCTTTGGGTTACCTA | 4 |
| wt-82560-82680 | 685 | TTGATGAGGATGAGAGTGGCGAGGGGTCCGATACCTCCGAGCCGTGTGAAGCTCTTGATTTGTCAATCCATGGCAGGCCCTGCCCTCAGGCCCCCGAGTGGCCTGTTCAAGAGGAGGGTG | 4 |

| | | | |
|---|---|---|---|
| wt-82680-82800 | 686 | GCCAGGATGCCACCGAGGTTCTTGATTTGTCAATCCATGGCAGGCCCCGCCCTCGGACCCCCGAGTGGCCTGTTCAAGGGGAAGGTGGCCAAAATGTCACAGGCCCTGAAACTAGAAGGG | 4 |
| wt-82800-82920 | 687 | TGGTGGTGTCAGCTGTTGTTCACATGTGTCAGGATGACGAGTTTCCGGATCTACAAGATCCTCCAGATGAGGCCTAAGCAAAGGTGTAGAAGTGTGTCCCCCTCCATTCCACCCACTGAT | 4 |
| wt-82920-83040 | 688 | AATACGCCCGACAATAAAGTTGATGATATTGAATTCCACACCTGCTTGTGTTTGTGATTTTATTTCATATTCCATGAGAGAGACCTCGCATATTTGCAGAAGGGTCACTGAAACATCTTA | 4 |
| wt-83040-83160 | 689 | TCTTAAAACAGTTACACCTGAATAATGAAGAAAGCGTGGCTCAGCAGAGCACAGCAAGCCGATGCCGGGGGGGCATCTGGCTCCGAGGACCCACCAGATTATGGAGATCAAGGTAATGTG | 4 |
| wt-83160-83280 | 690 | ACACAGGTGGGATCTGAGCCTATTTCACCTGAGATTGGCCCCTTTGAACTCTCTGCGGCCAGTGAGGATGATCCTCAATCTGGGCCAGTGGAAGAGAATTTAGATGCCGCTGCAAGAGAG | 4 |
| wt-83280-83400 | 691 | GAAGAGGAACCTCATGAGCAGGAGCACAATGGTGGTGACGATCCCTTGGATGTCCATACTCGCCAGCCTAGATTTGTGGATGTGAACCCAACGCAGGCTCCAGTGATCCAACTAGTCCAC | 4 |
| wt-83400-83520 | 692 | GCTGTCTATGATTCCATGTTGGTAAGAGGCACCTAGAACATTTCCAGATGTTTCGCTTGGATTTTTTGGCCAGTCTTAATTGATTGTCATTGGTTTCAGCAATCGGACCTCCGGCCCCTA | 4 |
| wt-83520-83640 | 693 | GGCAGTTTATTCCTTGAGCAAAACCTGAACATCGAAGAATTTATATGGATGTGCATGACCGTGCGTCACAGATGTCAGGCCATCAGAAAAAAACCATTACCAATTGTTAAGCAGAGGCGT | 4 |
| wt-83640-83760 | 694 | TGGAAGCTCCTGTCATCTTGCAGATCCTGGCGTATGGGTTACCGCACGCATAACCTCAAAGTAAACAGTTTTGAGTCAGGGGGGGACAATGTCCACCCGGTCCTTGTGACTGCTACGCTA | 4 |
| wt-83760-83880 | 695 | GGATGTGATGAGGGCACGCGGCATGCAACAACGTACAGTGCTGGCATTGTACAGATACCACGAATATCAGACCAAAACCAAAAGATCGAAACAGCCTTCCTGATGGCACGTCGTGCTAGG | 4 |
| wt-83880-84000 | 696 | TCACTTTCGGCAGAAAGATATACTTTGTTCTTTGATTTAGTATCCTCCGGAAACACCCTGTATGCTATATGGATTGGGCTGGGCACGAAAAACCGAGTTTCATTTATTGAGTTTGTAGGA | 4 |
| wt-84000-84120 | 697 | TGGTTATGCAAGAAGGACCACACTCATATACGCGAATGGTTCCGCCAGTGCACCGGGAGACCCAAAGCAGCCAAGCCATGGTTAAGAGCGCATCCTGTCGCCATTCCTTATGATGATCCG | 4 |
| wt-84120-84240 | 698 | TTAACAAACGAGGAGATTGATCTGGCCTATGCCCGCGGGCAGGCCATGAATATTGAGGCTCCTAGACTGCCAGATGATCCTATAATTGTTGAGGATGACGACGAAAGTGAGGAAATTGAA | 4 |
| wt-84240-84360 | 699 | GCTGAAAGCGACGAGGAGGAAGACAAGAGTGGAATGGAATCTCTTAAAAATATACCGCAAACACTGCCGTACAATCCAACAGTATACGGCAGGCCCGCGGTGTTTGACCGAAAGTCAGAT | 4 |
| wt-84360-84480 | 700 | GCAAAATCAACCAAAAAATGCAGGGCCATAGTAACTGACTTTAGTGTAATCAAGGCCATTGAAGAGGAACACAGAAAGAAGAAGGCAGCCAGAACAGAGCAGCCAAGAGCCACGCCTGAA | 4 |
| wt-84480-84600 | 701 | TCCCAGGCCCCCACAGTGGTCCTCCAGCGACCACCCACGCAGCAAGAGCCTGGCCCCGTCGGCCCACTGAGTGTCCAGGCTCGGCTGGAGCCATGGCAACCTTTGCCTGGGCCCCAAGTG | 8 |
| wt-84600-84720 | 702 | ACAGCAGTTCTACTTCACGAAGAATCCATGCAGGGTGTCCAAGTACATGGTTCGATGCTAGACCTTCTTGAAAAAGACGATGAAGTCATGGAGCAGAGGGTTATGGCAACCCTACTGCCA | 4 |
| wt-84720-84840 | 703 | CCAGTACCACAACAGCCCCGGGCTGGCAGAAGAGGCCCTTGTGTCTTCACCGGTGACCTAGGCATAGAGAGTGATGAGCCCGCTTCGACAGAGCCGGTTCATGATCAGCTACTGCCTGCC | 4 |
| wt-84840-84960 | 704 | CCAGGACCTGACCCTCTTGAAATCCAACCACTAACGTCCCCCACCACGTCTCAACTTAGCAGTTCGGCACCAAGCTGCGCACAAACTCCATGGCCGGTGGTTCAGCCAAGTCAGACTCCA | 4 |
| wt-84960-85080 | 705 | GATGACCCAACGAAACAGTCCCGGCCACCGGAAACAGCTGCCCCACGCCAGTGGCCAATGCCCCTGCGACCTATCCCTATGCGCCCCTTGCGGATGCAGCCAATCCCATTTAATCATCCA | 4 |
| wt-85080-85200 | 706 | GTGGGACCCACTCCCCATCAGACACCTCAAGTGGAAATAACACCATATAAGCCCACTTGGGCTCAGATAGGGCACATTCCATATCAGCCTACACCAACGGGTCCTGCTACCATGCTGTTG | 4 |
| wt-85200-85320 | 707 | CGCCAGTGGGCACCCGCCACCATGCAGACACCACCGAGAGCGCCCACTCCCATGTCACCACCTGAGGTGCCACCCGTTCCCCGGCAGAGGCCTCGGGGGGCGCCCACTCCCACGCCACCT | 8 |
| wt-85320-85440 | 708 | CCTCAGGTGCCGCCCGTTCCCCGGCAGAGGCCTCGGGGGGCGCCCACTCCCACGCCACCTCCTCAGGTGCTGCCCACTCCCATGCAGCTGGCACTAAGGGCTCCTGCTGGTCAGCAGGGG | 8 |
| wt-85440-85560 | 709 | CCGACAAAGCAAATTTTGCGCCAATTGTTAACGGGGGGCGTCAAGAAAGGGAGACCATCACTTAAGTTACAGGCCGCCCTTGAGCGTCAAGCCGCTGCGGGCTGGCAGCCTTCACCAGGG | 4 |
| wt-85560-85680 | 710 | TCCGGCACCAGTGACAAGATTGTGCAGGCGCCTATTTTCTACCCACCCGTTTTGCAGCCCATACAGGTTATGGGGCAAGGGGGTTCCCCAACGGCCATGGCCGCCTCAGCGGTGACACAG | 4 |
| wt-85680-85800 | 711 | GCACCCACGGAATATACCAGGGAAAGGAGGGGAGTGGGGCCTATGCCTCCCACCGATATTCCGCCGTCTAAACGAGCGAAGATCGAGGCCTATACAGAGCCCGAGATGCCGCACGGGGGG | 4 |
| wt-85800-85920 | 712 | GCCTCGCACTCTCCCGTCGTTATCTTGGAGAATGTCGGCCAGGGGCAACAGCAGACTCTGGAGTGCGGAGGAACTGCTAAACAGGAAAGGGACATGTTGGGGCTGGGGGACATTGCAGTT | 4 |
| wt-85920-86040 | 713 | TCTTCCCCTTCCTCTTCTGAAACATCGAACGATGAGTGATTTCGCCCATGTAACAAGAACTGGGATGAACCCTGGGGCAACAGACTGCGGGGAGGAGGGGGGCAGTGATAAGTCATGACA | 4 |
| wt-86040-86160 | 714 | ATTTTAGATGAGGTAGAAATTTTGCATATTTTCAGACCCACCATGGAATCATTTGAAGGACAGGGGGACTCTAGACAGTCACCCGACAATGAGCGGGGAGATAATGTACAGACTACCGGC | 4 |
| wt-86160-86280 | 715 | GAGCATGATCAGGACCCTGGGCCGGGGCCTCCATCCAGTGGGGCTTCTGAGAGATTGGTACCAGAAGAGTCATACTCAAGAGATCAGCAACCTTGGGGGCAAAGCAGGGGTGATGAAAAC | 4 |
| wt-86280-86400 | 716 | AGAGGCTGGATGCAGCGCATCAGGCGAAGGCGGAGAAGACGGGCTGCCTTGTCCGGCCATCTTTTAGACACGGAAGACAATGTGCCGCCATGGTTGCCTCCACACGACATCACACCATAT | 4 |
| wt-86400-86520 | 717 | ACCGCAAGGAATATCAGGGATGCTGCCTGCCGGGCTGTCAAGGTGAGTATGCCTCTAACTGGGTTCATGGGGGCCATCTAAGGCCCACGTGTGACCCATGTTTCCATTAATTTTAGCAAT | 4 |
| wt-86520-86640 | 718 | CGCACCTGCAAGCGCTATCAAACCTGATACTCGATAGTGGGTTAGACACACAACACATCTTGTGCTTCGTGATGGCAGCCAGGCAGCGTCTTCAGGACATTCGACGTGGACCCTTGGTTG | 4 |
| wt-86640-86760 | 719 | CGGAGGGCGGTGTGGGTTGGCGACATTGGCTTCTAACATCTCCCAGCCAATCCTGGCCCATGGGATATCGTACAGCAACACTACGCACATTAACTCCCGTGCCTAACAGGGTTGGGGCTG | 4 |
| wt-86760-86880 | 720 | ACAGCATCATGTTAACTGCCACATTTGGATGCCAAAATGCGGCACGAACTCTAAACACCTTCTCTGCCACCGTGTGGACACCACCCCATGCTGGACCAAGAGAGCAAGAAAGATACGCTC | 4 |
| wt-86880-87000 | 721 | GGGAAGCCGAGGTACGCTTCCTTCGTGGTAAATGGCAGAGGCGGTACCGAAGAATCTATGATTTGATAGAACTGTGTGGCTCTCTGCACCACATCTGGCAAAACTTGCTCCAGACCGAGG | 4 |
| wt-87000-87120 | 722 | AGAACCTTTTAGATTTCGTGCGTTTCATGGGTGTCATGTCCAGCTGCAATAATCCAGCTGTGAATTACTGGTTTCACAAGACAATCGGAAACTTTAAGCCATATTACCCGTGGAATGCAC | 4 |
| wt-87120-87240 | 723 | CACCTAATGAAAATCCATATCACGCGCGGAGAGGCATAAAAGAACACGTAATCCAGAACGCATTTCGAAAGGCACAAATACAGGGTTTATCAATGTTAGCAACGGGAGGTGAACCCAGAG | 4 |
| wt-87240-87360 | 724 | GTGATGCTACTAGTGAAACGAGCAGTGATGAGGACACCGGTAGACAGGGTTCGGACGTGGAGCTAGAGTCCTCGGACGATGAGCTGCCATATATCGATCCCAATATGGAGCCGGTTCAGC | 4 |
| wt-87360-87480 | 725 | AGAGGCCCGTCATGTTTGTGAGCCGTGTGCCTGCAAAGAAACCGAGGAAACTGCCTTGGCCCACGCCCAAGACGCACCCAGTGAAGCGCACAAATGTTAAGACCTCTGATAGATCTGATA | 4 |
| wt-87480-87600 | 726 | AGGCAGAAGCACAAAGCACCCCTGAAAGGCCGGGCCCTTCCGAACAATCATCAGTGACCGTGGAGCCCGCCCACCCGACCCCGGTGGAGATGCCAATGGTGATTCTCCATCAACCACCTC | 4 |
| wt-87600-87720 | 727 | CAGTGCCCAAACCGGTTCCAGTCAAGCCTACGCCACCGCCTTCCCGTAGGAGAAGGGGAGCGTGTGTTGTGTACGACGATGATGTCATAGAGGTGATTGATGTTGAAACCACCGAAGATT | 4 |
| wt-87720-87840 | 728 | CATCGTCAGTGTCACAGCCAAATAAGCCACATCGGAAACATCAAGACGGCTTTCAACGTTCAGGCCGACGTCAAAAACGAGCCGCGCCTCCCACCGTGAGTCCTTCGGATACTGGGCCTC | 4 |
| wt-87840-87960 | 729 | CTGCCGTGGGGCCTCCTGCCGCGGGGCCTCCTGCCGCGGGGCCTCCTGCCGCGGGGCCTCCTGCCGCGGGGCCTCCTGCCGCGGGGCCTCCTGCCGCGGGGCCTCGCATACTGGCGCCTC | 12 |
| wt-87960-88080 | 730 | TTTCCGCTGGGCCTCCTGCCGCGGGGCCTCACATAGTGACGCCTCCTTCCGCCCGGCCTCGTATAATGGCGCCTCCCGTCGTACGTATGTTTATGAGGGAGCGACAGCTCCCCCAGTCCA | 4 |
| wt-88080-88200 | 731 | CCGGCCGTAAACCTCAGTGCTTCTGGGAAATGCGGGCTGGTCGTGAAATTACACAAATGCAACAAGAACCAAGTTCACACCTGCAGTCCGCCACTCAGCCTACAACGCCTCGCCCATCAT | 4 |
| wt-88200-88320 | 732 | GGGCCCCATCAGTCTGCGCCCTCTCGGTGATGGATGCTGGTAAGGCCCAGCCCATAGAAAGTTCACACTTGAGTTCCATGTCGCCCACACAGCCGATATCGCACGAAGAACAACCCCGGT | 4 |
| wt-88320-88440 | 733 | ATGAGGATCCTGACGCTCCTCTGGATTTAAGTTTACATCCAGACGTTGCTGCTCAACCAGCTCCCCAGGCTCCATACCAGGGATACCAGGAGCCGCCGGCCCCCCAGGCTCCATACCAGG | 4 |
| wt-88440-88560 | 734 | GATACCAGGAGCCGCCGCCCCCCCAGGCTCCATACCAGGGATACCAGGAGCCGCCGGCCCACGGGCTCCAATCATCTTCATATCCAGGATATGCGGGTCCCTGGACCCCAAGGTCTCAAC | 4 |
| wt-88560-88680 | 735 | ATCCATGTTATAGGCACCCCTGGGCACCATGGTCTCAAGATCCTGTGCATGGGCACACCCAGGGTCCATGGGATCCCAGGGCACCACATCTCCCACCTCAGTGGGATGGATCTGCAGGAC | 4 |
| wt-88680-88800 | 736 | ATGGCCAGGATCAGGTCTCCCAGTTCCCACATCTGCAATCGGAGACAGGCCCACCACGTCTTCAACTTTCATTGGTGCCACTGGTCTCATCCTCTGCACCATCATGGTCATCTCCCCAGC | 4 |

| | | | |
|---|---|---|---|
| wt-88800-88920 | 737 | CCCGAGCCCCCATACGCCCCATTCCAACAAGATTCCCCCCTCCCCCTATGCCGTTACAAGATAGCATGGCCGTGGGGTGTGACTCATCAGGTACAGCATGCCCAAGCATGCCCTTTGCCA | 4 |
| wt-88920-89040 | 738 | GTGATTACAGTCAAGGTGCATTTACCCCACTGGACATTAATGCCACCACGCCAAAAAGGCCTCGAGTAGAAGAAAGTTCTCACGGACCTGCCCGGTGTTCCCAAGCTACTGCTGAAGCAC | 4 |
| wt-89040-89160 | 739 | AGGAGATTCTCAGTGACAATTCTGAGATCTCCGTGTTCCCAAAGGACGCGAAGCAGACTGACTACGATGCATCCACTGAAAGTGAGCTAGATTAAGGGGATCCAAGGTGACCCCTGTTAG | 4 |
| wt-89160-89280 | 740 | CTATTTGATCTTTGACTGACACATAAACATGGTTTAAGGAATGAACACTCATGGTGTGAGACTGGAACTGTACTAAATTTGCTGACATATGTACAATGAGAGCCAAAAATTTGATAAACC | 4 |
| wt-89280-89400 | 741 | TTAAAAGTCCCCCCATCTAATGATGTCCAGTTCCCTTCTCCCACCCTGTACACCCCGACCCAAAGGGACTCAATGGCATTCAGATTTCTAGTTACCACAGGTAGAATATCGGGCGTTGGC | 4 |
| wt-89400-89520 | 742 | CCATAAAAATAAGTGCATGGATATAGCTCTGCACAGGCTTGGAAACACCCATTCCAGGTGTGCTTCTTTTTGGTGAAATAAAAACAGCATCCTTTATATGAAAATGTGTATTCTCTGGTG | 4 |
| wt-89520-89640 | 743 | TTGCAGTATGTACAGTTAGCTTTGGTATAGTTTTGGGGTACCTGAAATGTGTGCAGGGTGGGTGTCCAATGTGGCAGTTTTACCTCTTTGTCCCCATACTCCTGCTCGGCCGTCTTGTTA | 4 |
| wt-89640-89760 | 744 | AAGTTAACCGGCGGTGGAGGATCCACCGGCCAGACCTCTACATTTGGTTTGGGTACCCAGGTGATGGCCGCGGCTGCCACCCGCCCTCCTCCTCTTACCCTGGGTGGCAAAAAGTATGCC | 4 |
| wt-89760-89880 | 745 | AGGAGTAGAACAATAACAAGTGCGATGGCGGTAAACAATGGCACCCTCACCTGCTTAAATGAAACCATGGCAACCACTTCAAAGAGAGCCGACAGGAAGATATTTATTAATATTCCATTA | 4 |
| wt-89880-90000 | 746 | GTAAACGAGGCGTGAAGCAGGCGTGGTTTCAATAACGGGAGTTAGAAATTTAAGAGATCCTCGTGTAAAACATCTGGTGTCCGGGGGATAATGGAGTCAACATCCAGGCTTGGGCACATC | 4 |
| wt-90000-90120 | 747 | TGCTTCAACAGGAGGCGCAGCCTGTCATTTTCAGATGATTTGGCAGCAGCCACCTGCGGACAAAAATCAGGCGTTTAGATGGGGCATTTATGTTTGGGACGCTAGCCGCCTGGGCATTCG | 4 |
| wt-90120-90240 | 748 | TGTTAGTATATACTGACCTCACGGTAGTGCTGCAGCAGTTGCTTAAACTTGGCCCGGCATTTTCTGGAAGCCACCCGATTCTTGTATCGCTTTATTTCTAGTTCAGAATCGCATTCCTCC | 4 |
| wt-90240-90360 | 749 | AGCTGCGAGCAAGGGAATGCGTTACTACAAGTGGTGCCTAGTCAGTTGAAACAAGCCCCACCATCCGCTGCCGCCCCTCCATGAGCCCCACCGTCCGCTGCCGCCCCTCCTTGAGCCCCT | 4 |
| wt-90360-90480 | 750 | CCTTACCGATTCTGGCTGTTGTGGTTTCCGTGTGCGTCGTGCCGGGGCAGCCACTGGTGCAGGCTGTGGAACACCAATGTCTGCTAGCTGTTGTCCTTGGTTAGCCCCGGGGCAAGCAAA | 4 |
| wt-90480-90600 | 751 | CACCACTGCTGCTGCTGTTTGAACAGTAGAATTGTCTCCAGGTTGAGGTGCTTCTCCCCCGGCTTGGTTAGTCTGTTGATTCTGGGTTATGTCGGAGACTGGGAACAGCTGAGGTGCTGC | 4 |
| wt-90600-90720 | 752 | ATAAGCTTGATAAGCATTCTCAGGAGCAGGCTGAGGGGCAGAAAACCACGACCCAGTCGGAGCGGTTGAAACATGATAGGCAGTTAGCTGGCCTTGTGGCAGAGGCTCTGGCAGCACCGG | 4 |
| wt-90720-90840 | 753 | CCACAGCACACAAGGCAAAGGAGCTTGCGATGGCCCTCCCAGGTCCTGATAGACTCTGGTAGCTTGGTCAAAAGCTTGTACAAAAGGCACCTGGTATGGGTCAGGTGTAAATTTTACATC | 4 |
| wt-90840-90960 | 754 | TTCAGAAGTCGAGTTTGGGTCCATCATCTTCAGCAAAGATAGCAAAGGTGGCCGGCAAGGTGCAATGTTTAGTGAGTTACCTGTCTAACATCTCCCCTTTAAAGCCAAGGCACCAGCCTC | 4 |
| wt-90960-91080 | 755 | CTCTGTGATGTCATGGTTTGGGACGTGCTAAATTTAGGTGTGTCTATGAGGTACATTAGCAATGCCTGTGGCTCATGCATAGTTTCTAAAAGAGGAGGAGGCAGTTTTCAGAAGTGTCTA | 4 |
| wt-91080-91200 | 756 | AAATAAGCTGGTGTCAAAAATAGACAGCCCAGTTGAAATATGCATGGCATGCAGCAGACATTCATCATTTAGAAATGTATCCAAGATTTCATTAAGTTCGGGGGTCAGGGGGGAGTCCAG | 4 |
| wt-91200-91320 | 757 | ATTCAAATCCTCTGTCATGGACTCTAGTGTTGTGGTCAGTTCGTCCAAATGGCCACGAGGGGGCGGGTGGCTCAGGTCCATCTGTCCACATATGGCTGCTTCCTCCTTCTGGGGAATAAC | 4 |
| wt-91320-91440 | 758 | AGTGTCAGCCATCTCCCTTAGGGCCTTCACGGCCTGACTGGTTTCTTCATCAGGGTCCTCCAACAGATGACTTGCCTCGGGGGTTACTGCGGGGGCCGGGTCAAGTGGCTGGGGCACCGG | 4 |
| wt-91440-91560 | 759 | GGCTGGCGTTAGGGATCCGACCGGTTCATGGACAGGTCCTGTGGGGGTGGGAGCCAAAGAGGCAGGCAGGGGCCGGTTGGCCCACGGGGATCCGGGTGGATGGAAGGGCCTGATCCTCTT | 8 |
| wt-91560-91680 | 760 | TGGCTGACACACCTCTCGCCCCTCGAACACGTCAGATATGGCACTGCCCGCTTCCGGCTTTGGCAGGAACATACCTTCCCGGCTATCCCTGAGGCCCTTCTTCCTTTTAACGGGAGGAAG | 4 |
| wt-91680-91800 | 761 | AAAGGTGGGCTTTGAGGGGTGGGGGAATATGGGTCTCTCATCGCTCTCTTGGTGGACCGCTGCTATCCAAGGCTGTTCAGGTTCCGCCGCGTTGGAAGGACATGGAGTTTGACCACGGTT | 4 |
| wt-91800-91920 | 762 | GGGCCTGGATGTCCGGCGCGACTTTGGGGCCCGCAGGCGCGGGGCCTCGGCCCTGGCCTCTTCCCGCTCGCTCTGCTCGGTGTCACTGTTGCCCGAGTCACTGCTGCTGGAACTGCTGTC | 8 |
| wt-91920-92040 | 763 | ACCGCAGTCGGCGCTTTGGGCACCGGGCTTCAGGGGCATGGTCGGGCTCGGGAGACTTTCGAGTTCATCTGTAAAAGCATGAAACTGTCCGGACTCCGAGTAGCGGGCCTCGGTGTGAGA | 4 |
| wt-92040-92160 | 764 | GGCACCCCCATCATTCCCCATGAGCTCCTCGTCCATCCTGTCGGCTCCGGACACGAGGATAGGAGTTTCCACTGCCTTGGACTTGGTTGACAGCAGGCACGCGGGAAGCACGCCGCTCAC | 4 |
| wt-92160-92280 | 765 | GTAGCTCCTCTGTCCGGCGTGGCTGGAGTAGGAGGCCCGGGGCAGTGTCTTAATCAGAGCCCTGACATCCTTAACATCGTCCGTCAGATGGCCTGTCTTGGACGAGACCATAGTCTGGAA | 4 |
| wt-92280-92400 | 766 | CATCTCCTCGAGGACGGGATAGGTGAACACCCACTTGCAAAAGGCCTTGAACTTGGAGCTTAGGAGGCCTTCCTTCTCCATCCTGTTCAGGTGTTCCACTACCTGCTTGCCGGAGGCCAT | 4 |
| wt-92400-92520 | 767 | GATGGCCGCGCGGTCCACGCCCAGCACCTTGCTGTAGGTGTAGGCCCGCACCCGACTGTGTTTTAGGAGCTTGTACATAGCGGTGCCTATGGTGGCAGGAATCATCACCCGGTTGCTGGG | 4 |
| wt-92520-92640 | 768 | GGCCTGGATGAAGAATCTGTCAGTGACCACTATCAGGTGGTCTAACACGTAGCGCATCACTATAGGGCACGCGATGGAACATGCGTCGTTGCCGGCATTCTCAGCCCGTCTTCTTACCCT | 4 |
| wt-92640-92760 | 769 | GTTGTTTCGGAGAATGGCCCAAAAATTGCAGATGTTGAGCGTGGCCATTAGCCCGCCCCATTCTCGCCCGTGGGCCTTGGCCTCATTTATAAATGCCTTGCATATTTTGTAGGATCTCAG | 4 |
| wt-92760-92880 | 770 | AGTAATCTCCACACTCCCGGCTGTAAATTCCTTGTTGAGGACGTTGCAGTAGTCAGAGACCAGAGAGCCCAGCTGCTTTTTGATTTCAGGAGTTAGCCTCAGAAAGTCTTCCAAGCCATC | 4 |
| wt-92880-93000 | 771 | CTTTTTAGGCCTCATGGCTAGTAGTAACAGAGGAAATGCCCGACCATTAAAATCTTTCCTCCATGAGCTTTACCTGAAACACTATCCCGAAGTGGGGGATGTGGTGCATCTACTGAACAC | 4 |
| wt-93000-93120 | 772 | CATCGGGGTCGACTGCGACCTCCCACCTAGCCACCCACTCCTGACAGCCCAGAGGGGGCTGTTCCTGGCAAGAGTCTTGCAGGCTGTACAGCAGCACAAGCTGCTGGAAGACACCATCGT | 4 |
| wt-93120-93240 | 773 | CCCCAAGATCTTAAAGAAGCTGGCTTATTTCTTAGAGCTGCTAAGCTACTACTCCCCCAAGGATGAACAGCGTGACATCGCCGAGGTTCTTGACCACCTCAAGACGAATCGGGACCTGGG | 4 |
| wt-93240-93360 | 774 | GCTGGACGACAGACTCTGGGCCCTGATTAGGAAACTGCGCCAAGACAGACACCATGCCTCTGTAAATGTCCTCATGCCAGGAAGCGACTACACAGCCGTGTCGCTGCAGTACTACGACGG | 4 |
| wt-93360-93480 | 775 | CATCTCCATAGGTATGAGGAAGGTAATCGCGGATGTCTGCCGCAGTGGCTATGCCTCCATGCCCTCCATGACGGCCACGCACAACCTCTCCCACCAGCTCTTGATGGCGTCCGGGCCCAG | 4 |
| wt-93480-93600 | 776 | TGAGGAACCGTGCGCCTGGCGCGGGTTCTTTAACCAGGTCCTCCTCTGGACTGTGGCCCTCTGCAAGTTTCGCAGATGCATTTACTATAACTACATTCAGGGATCTATAGCCACCATCTC | 4 |
| wt-93600-93720 | 777 | CCAGCTTCTGCACCTCGAGATCAAGGCCCTCTGCAGCTGGATAATATCCCAGGATGGCATGCGCCTCTTTCAACACAGCAGGCCTCTCCTCACCCTCTGGGAGAGCGTGGCCGCAAATCA | 4 |
| wt-93720-93840 | 778 | GGAGGTCACGGATGCCATTACCCTGCCTGACTGCGCTGAATACATAGACCTACTAAAGCACACAAAACATGTCTTAGAAAACTGTTCTGCCATGCAATACAAATAAATTTCTCTTACCTG | 4 |
| wt-93840-93960 | 779 | CGTCTGTTTGTGTAGTGAGGTGTTGTGTCCTGTATGGTATTCTACTTTAAAAAGGCCGGCTGACATGGATTACTGGTCTTTTATGAGCCATTGGCATGGGCGGGACAATCGCAATATAAA | 4 |
| wt-93960-94080 | 780 | ACCCTGACCATCACATGGGGCATTAGGCGACTCTGCATCAGCATCGCTTAAGTATGAGTGGGCAGCAGAGAGGCTCGGTTATTTTGGTTCCTGAACATCTGGCTGGGGCATTAACTAAGC | 4 |
| wt-94080-94200 | 781 | TTATGAGCGATTTTATCACAGGACAAGATGTCACTCTTTCTGGAGGAAATATTGCAGTCAAAATTCGCGATGCTATAAACCAGACCCCCGGGGGTGGTGATGTAGCTATACTTTCTTCCC | 4 |
| wt-94200-94320 | 782 | TGTTTGCTTTATGGAATGCCCTCCCAACATCTGGTAGACAATCCTCCAGGGACGATTTAATCCCAGCCGCCGTGCAGGCCTTAACCACGGCCCACAACTTATGTCTGGGTGTTATTCCAG | 4 |
| wt-94320-94440 | 783 | GTGAGACCTCACACAAGGACACACCCGAGTCATTGCTCCGGGCTATCGTGACGGGTCTCCAAAAATTGTGGGTGGATTCGTGCGGATGTCCAGAGTGCCTACAATGTCTTAAGGGATTGA | 4 |
| wt-94440-94560 | 784 | AGGCAATTAAGCCCGGCCTTTATGAAATCCCTAGGATAATACCACACACTAAGCAGTGTAGTCCTGTCAATCTCCTGAACATGTTGGTCCACAAGCTTGTGGCTTTACGTGGTCATGTGC | 4 |
| wt-94560-94680 | 785 | AGCTTGCATACGACGCCCGTGTCCTGACGCCTGACTTTCACGAAATCCCTGACCTCGATGACTCCGATGCTGTTTTCGCACGCACCTTATTGGCAGCCTTATTTCACCTCAATATGTTCT | 4 |
| wt-94680-94800 | 786 | TTATTCTCAAAGATTACATAACACAAGACTCCATGAGCTTGAAGCAGGCCCTCAGTGGTCATTGGATGTCTGCCACGGGCAACCCCCTGCCTGCAGCACCGGAAACCCTGCGAGACTACT | 4 |
| wt-94800-94920 | 787 | TGGAAGCTTTCCGAAATTCGGATAATCACTTTTATCTCCCGACGACAGGGCCTTTAAACACCTTCCAATTTCCCGAAGAGCTTCTGGGGCGCGTTGTTGTTATTGATTCCTCTTTGTGTG | 4 |

| | | | |
|---|---|---|---|
| wt-94920-95040 | 788 | CCGCCAGTCACGTTCAGGACGTTATCACCCATGGTGTTGGGGCGGGTGTTCCTCGTCCTCGGTTTTCGGCCCTGCCTCCGGCCCCATCCCGCGAGCCCCAGCAGACATGCTCTCAGTTAA | 4 |
| wt-95280-95400 | 791 | CCTCACAGGCCGCACCCAGCTTTTCTTCCGTTGCCCCAGTAGCATCTCTGTCTGGTGACCTTGAAGAGGAAGAGGAGGGGTCCCGAGAATCCCCATCCCTACCGTCCAGCAAAAAGGGGG | 4 |
| wt-95400-95520 | 792 | ACGAGGAATTTGAGGCCTGGCTTGAGGCTCAGGACGCAAATCTTGAGGATGTTCAGCGGGAGTTTTCCGGGCTGCGAGTAATTGGTGATGAGGACGAGGATGGTTCGGAGGATGGGGAAT | 4 |
| wt-95520-95640 | 793 | TTTCAGACCTGGATCTGTCTGACAGCGACCATGAAGGGGATGAGGGTGGGGGGGCTGTTGGAGGGGGCAGGAGTCTGCACTCCCTGTATTCACTGAGCGTCGTCTAATAAAGATGTCTAT | 4 |
| wt-95640-95760 | 794 | TGATCTCTTTTAGTGTGAATCATGTCTGACGAGGGGCCAGGTACAGGACCTGGAAATGGCCTAGGAGAGAAGGGAGACACATCTGGACCAGAAGGCTCCGGCGGCAGTGGACCTCAAAGA | 4 |
| wt-95760-95880 | 795 | AGAGGGGGTGATAACCATGGACGAGGACGGGGAAGAGGACGAGGACGAGGAGGCGGAAGACCAGGAGCCCCGGGCGGCTCAGGATCAGGGCCAAGACATAGAGATGGTGTCCGGAGACCC | 4 |
| wt-95880-96000 | 796 | CAAAAACGTCCAAGTTGCATTGGCTGCAAAGGGACCCACGGTGGAACAGGAGCAGGAGCAGGAGCGGGAGGGGCAGGAGCAGGAGGGGCAGGAGCAGGAGGAGGGGCAGGAGCAGGAGGA | 4 |
| wt-96000-96120 | 797 | GGGGCAGGAGGGGCAGGAGGGGCAGGAGGGGCAGGAGCAGGAGGAGGGGCAGGAGCAGGAGGAGGGGCAGGAGGGGCAGGAGGGGCAGGAGCAGGAGGAGGGGCAGGAGCAGGAGGAGGG | 8 |
| wt-96120-96240 | 798 | GCAGGAGGGGCAGGAGCAGGAGGAGGGGCAGGAGGGGCAGGAGGGGCAGGAGCAGGAGGAGGGGCAGGAGCAGGAGGAGGGGCAGGAGGGGCAGGAGCAGGAGGAGGGGCAGGAGGGGCA | 8 |
| wt-96240-96360 | 799 | GGAGGGGCAGGAGCAGGAGGAGGGGCAGGAGCAGGAGGGGCAGGAGGGGCAGGAGGGGCAGGAGCAGGAGGGGCAGGAGCAGGAGGAGGGGCAGGAGGGGCAGGAGGGGCAGGAGCAGGA | 8 |
| wt-96480-96600 | 800 | GGGGCAGGAGGGGCAGGAGCAGGAGGAGGGGCAGGAGGGGCAGGAGCAGGAGGGGCAGGAGGGGCAGGAGCAGGAGGGGCAGGAGGGGCAGGAGCAGGAGGGGCAGGAGGGGCAGGAGCA | 8 |
| wt-96600-96720 | 801 | GGAGGAGGGGCAGGAGCAGGAGGGGCAGGAGCAGGAGGTGGAGGCCGGGGTCGAGGAGGCAGTGGAGGCCGGGGTCGAGGAGGTAGTGGAGGCCGGGGTCGAGGAGGTAGTGGAGGCCGC | 8 |
| wt-96720-96840 | 802 | CGGGGTAGAGGACGTGAAAGAGCCAGGGGGGGAAGTCGTGAAAGAGCCAGGGGGAGAGGTCGTGGACGTGGAGAAAAGAGGCCCAGGAGTCCCAGTAGTCAGTCATCATCATCCGGGTCT | 4 |
| wt-96840-96960 | 803 | CCACCGCGCAGGCCCCCTCCAGGTAGAAGGCCATTTTTCCACCCTGTAGGGGAAGCCGATTATTTTGAATACCACCAAGAAGGTGGCCCAGATGGTGAGCCTGACGTGCCCCCGGGAGCG | 4 |
| wt-96960-97080 | 804 | ATAGAGCAGGGCCCCGCAGATGACCCAGGAGAAGGCCCAAGCACTGGACCCCGGGGTCAGGGTGATGGAGGCAGGCGCAAAAAAGGAGGGTGGTTTGGAAAGCATCGTGGTCAAGGAGGT | 4 |
| wt-97080-97200 | 805 | TCCAACCCGAAATTTGAGAACATTGCAGAAGGTTTAAGAGCTCTCCTGGCTAGGAGTCACGTAGAAAGGACTACCGACGAAGGAACTTGGGTCGCCGGTGTGTTCGTATATGGAGGTAGT | 4 |
| wt-97200-97320 | 806 | AAGACCTCCCTTTACAACCTAAGGCGAGGAACTGCCCTTGCTATTCCACAATGTCGTCTTACACCATTGAGTCGTCTCCCCTTTGGAATGGCCCCTGGACCCGGCCCACAACCTGGCCCG | 4 |
| wt-97320-97440 | 807 | CTAAGGGAGTCCATTGTCTGTTATTTCATGGTCTTTTTACAAACTCATATATTTGCTGAGGTTTTGAAGGATGCGATTAAGGACCTTGTTATGACAAAGCCCGCTCCTACCTGCAATATC | 4 |
| wt-97440-97560 | 808 | AGGGTGACTGTGTGCAGCTTTGACGATGGAGTAGATTTGCCTCCCTGGTTTCCACCTATGGTGGAAGGGGCTGCCGCGGAGGGTGATGACGGAGATGACGGAGATGAAGGAGGTGATGGA | 4 |
| wt-97560-97680 | 809 | GATGAGGGTGAGGAAGGGCAGGAGTGATGTAACTTGTTAGGAGACGCCCTCAATCGTATTAAAAGCCGTGTATTCCCCCGCACTAAAGAATAAATCCCCAGTAGACATCATGCGTGCTGT | 4 |
| wt-97680-97800 | 810 | TGGTGTATTTCTGGCCATCTGTCTTGTCACCATTTTCGTCCTCCCAACATGGGGCAATTGGGCATACCCATGTTGTCACGTCACTCAGCTCCGCGCTCAACACCTTCTCGCGTTGGAAAA | 4 |
| wt-97800-97920 | 811 | CATTAGCGACATTTACCTGGTGAGCAATCAGACATGCGACGGCTTTAGCCTGGCCTCCTTAAATTCACCTAAGAATGGGAGCAACCAGCTGGTCATCAGCCGCTGCGCAAACGGACTCAA | 4 |
| wt-97920-98040 | 812 | CGTGGTCTCCTTCTTTATCTCCATCCTGAAGCGAAGCAGCTCCGCCCTCACGGGCCATCTCCGTGAGTTGTTAACCACCCTGGAGACTCTTTACGGTTCATTCTCAGTGGAAGACCTGTT | 4 |
| wt-98040-98160 | 813 | TGGTGCCAACTTAAACAGATACGCATGGCATCGCGGGGGCTAGACCTCTGGCTGGATGAGCACGTGTGGAAGAGGAAACAGGAGATTGGTGTGAAAGGAGAAAATCTGCTTCTCCCCGAC | 4 |
| wt-98160-98280 | 814 | TTATGGCTAGATTTCCTACAACTCAGCCCCATCTTCCAGCGCAAGCTTGCTGCCGTTATTGCCTGTGTCCGACGCCTGCGGACTCAGGCCACCGTCTACCCAGAGGAGGACATGTGCATG | 4 |
| wt-98280-98400 | 815 | GCCTGGGCCCGCTTTTGCGACCCCTCTGATATTAAGGTGGTTATTTTGGGCCAGGACCCCTATCACGGGGGTCAAGCAAACGGCCTGGCATTCAGCGTCGCATACGGCTTTCCAGTTCCC | 4 |
| wt-98400-98520 | 816 | CCCAGCCTGAGGAACATCTACGCGGAGCTGCACCGGAGCCTGCCGGAGTTTTCTCCCCCAGATCACGGCTGTCTAGACGCGTGGGCCTCCCAGGGGGTGTTGCTACTCAACACCATCCTG | 4 |
| wt-98520-98640 | 817 | ACCGTGCAAAAGGGCAAGCCCGGCTCGCACGCAGACATTGGCTGGGCGTGGTTTACTGACCACGTAATTTCATTGCTCTCTGAGCGGTTAAAAGCGTGCGTGTTTATGCTGTGGGGTGCG | 4 |
| wt-98640-98760 | 818 | AAGGCGGGAGACAAAGCTTCACTAATCAACTCCAAGAAGCATCTGGTTCTGACCTCTCAGCATCCCTCTCCCCTGGCCCAGAACAGCACCCGAAAGAGTGCCCAGCAGAAGTTCCTGGGC | 4 |
| wt-98760-98880 | 819 | AACAACCACTTTGTCCTCGCTAACAACTTTTTGCGTGAGAAGGGGCTCGGTGAGATAGATTGGAGGCTGTAGAGGGGTCATCACTATGGCCATGTTTCTGAAGTCGCGTGGGGTCCGGTC | 4 |
| wt-98880-99000 | 820 | TTGCAGGGACCGGCGCCTCTTGTCGGACGAGGAGGAAGAGACTTCACAGAGCAGCAGCTACACTCTGGGGTCTCAGGCCTCCCAGTCTATCCAGGAGGAGGACGTGAGTGACACTGATGA | 4 |
| wt-99000-99120 | 821 | GTCTGACTACTCAGATGAAGACGAGGAGATTGATTTGGAGGAAGAGTACCCCAGTGACGAAGACCCATCTGAGGGCAGTGATAGCGACCCCTCGTGGCATCCTTCAGATTCAGACGAGTC | 4 |
| wt-99120-99240 | 822 | TGACTACAGCGAGAGCGACGAGGATGAAGCAACCCCCGGCTCTCAGGCCTCACGATCTTCAAGAGTCTCGCCATCTACCCAACAGTCTTCAGGTCTGACACCCACGCCTTCGTTCTCCCG | 4 |
| wt-99240-99360 | 823 | ACCACGCACCCGGGCACCTCCGAGGCCGCCGGCTCCCGCGCCGGTCAGGGGACGGGCCTCAGCACCTCCCAGGCCACCAGCCCCAGTTCAGCAATCCACCAAAGACAAGGGTCCCCATAG | 8 |
| wt-99360-99480 | 824 | ACCTACGCGACCTGTACTTAGAGGCCCAGCTCCACGCCGCCCCCCTCCACCTTCAAGTCCCAATACATACAATAAACACATGATGGAAACCACCCCCCCCATTAAGGGCAATAACAACTA | 4 |
| wt-99480-99600 | 825 | CAATTGGCCATGGCTGTAAATAAAATGTCATAACCTGGAGTCTGCATGTCTGTTGTTTTATTCAGTAAACCAGTAGTGCGCGTGAGTTCTTTAGGGCATCCACGATGTAGCCGCTCGCGG | 4 |
| wt-99600-99720 | 826 | GGTTCCCCTCCCCAGTGATCATCTCGGATAGGGGATTCCTGTCCATGACCACGCAATTAGAGTGCCGGGCCCGGGACAGCGCCACATACACATGGCCGGGTTTGATGTTTCTGTGGCTGC | 4 |
| wt-99720-99840 | 827 | CGAAGCAGATGGCGACTTTGTTTAGGGACAGACCCTGGGCCTTGGCTATGGTCATGGCCAGCTTTGAGCTAATGCCATAGTCACGGATGCTGCAGAGGTTCAGGGACTTGTCCTCTATCG | 4 |
| wt-99840-99960 | 828 | TCTCATACAGTTTGTTAGTATTGTGTTCCAGGCAGCACACGAAGCCTGCCTCATCCTTGACCATTAGCCTGGGCATGCGTGAACTGCCAGCGTCCTGAGGCTGCTGCTTTCCTCGGATGC | 4 |
| wt-99960-100080 | 829 | CAAAGAAGACGTTGAGATGCGTGTAGCCCAGAAGCGTGTAGTTCTCGGTGGTGGAGGCGTAGTCCAGGAGGCCGTGAAGGAGAGGCTCGTCTGAGGTGAACTCTATGTTGTCGCGAATCA | 4 |
| wt-100080-100200 | 830 | GCATGTTGTTGGTAAATGTGCAGAAGGGGAGGTCCCTGAACTCCCTTCCGCCATAGCGGACGGCCACATCCAGGCATTGCCTGAAATAGGCCCTGAGGTCATTATATATGTTTAACAGGG | 4 |
| wt-100200-100320 | 831 | AGCAGAGGGGGGCAGAATTTGCGGCCGGGGGAGCCAGTACTCGGGCATAGAAGACAGCGGCGGGGCTCCGCTCCCCATCCCAGGCAACCTCCAGCGGCAGTTCGCCCAGCTCCATCCCAG | 8 |
| wt-100320-100440 | 832 | CAGTCACCTCCGGATCCCACGTACGCCCGGGCAGGGGCACAGCACCAAGCTCCGCCACGTATTCCCCGTTTTCACAGAGAGAATGTCCTCCGTGGCTAAAAGCGTAGATGCCTCCGTAGA | 4 |
| wt-100440-100560 | 833 | TGAGTCGGGCCAGGAAGCTGTAGACATACTCGGGCTGCTCATGCCCGTGGGCCTCCACGAAGCTGTCCGCCTCGAGCGTGTCCATAAAGTCCCCGAAGGTGCCGGTATAGCCACAGATGG | 4 |
| wt-100560-100680 | 834 | ACTTTTTGGTCTTGCAGTTGACCGACACCGAGCTGTGCTTGACGTAGGTGACATTGTAGGTGACCTTGACCCGTTCTTCGTCCTGCTCGGTGCCCACCGGGACCATGTCTTGGTCGGCGA | 4 |
| wt-100680-100800 | 835 | ACTGCGAGTAGTTACCGAGGCGTGCATAATTCTTTTGGAGCCAGGTGTGGGCCGTGAGGCCCGGAAGCCCGACCAGGGTCTTGTACTGGGCCAGGGGATCGAGGAAGACCTCGCACTCCA | 4 |
| wt-100800-100920 | 836 | CCGGGCAGGTAAACATGGTCACCCCGCCCCCATCTCCCCCGGTTCCCCGCGCGGCACGCCCCTGCCCGGCAGTCTTGAGCGTGGCATGGAGGGTGGTGAGGAAGGTCTTGACCTCGGCGT | 8 |
| wt-100920-101040 | 837 | GGGAGAGGAAGAGCCGGGTCCAGCCCACGTACTGCGCGGGGTCCATTATGGCCGCCCTGGGGACGACGAAGCGGTCGACGTAGGCCAGGATGTCCGGCGAGAGCTCGAGGCCGTACTCGA | 8 |
| wt-101040-101160 | 838 | GGGTCTTCATGAGGTGTCCAAACTGGACGTCGGTGCAGCGCTTGTTGTTGATGAAGAGGGCCCAGTTGCGGGCCACGTCCACGTAGGTCGCGGCCCTGGGGTTGCCCACCAGGAAGGTGA | 4 |
| wt-101160-101280 | 839 | GGATGTTGTCGCACTCGCGAATCTTGTTTACCTGGGTCTCGTGGCTAAAGGAGGACTGAAAGGCGTCTGTCTGGGTGGGAGAGCCCACGCAGACGATGCAGGGAATGCGGCCCCGGCGGT | 4 |
| wt-101280-101400 | 840 | AGAGTGGGGTACGCAGCCAGGCGTTGAAGAACCAGTAGCAAAAGACCACGGCTGTTAGAATGTGCACGGAAAGCGTTCCAGCTTCGTCCACCACGATCACATTGGTGGTCCATAGCTGCC | 4 |

| | | | |
|---|---|---|---|
| wt-101400-101520 | 841 | CCTGGTGCATGTCTCTCAGGACCTCAAAGGCGGGGCCAGAGACTCCCGAGTAGAGCCCCCTGGGCTTGGTTCGCCTGAACTCGGCGGCAATGTCGGAGAGTACCGGCCAGTATTTGGCCA | 4 |
| wt-101520-101640 | 842 | GGTCCCGCCGCTGGAGTTCCTCTAGGGCGGCGTCCGTAGAGCGACCATGACTGCTGACCCGCTGCGTCATATTTATGTGGCGGCTCTTGAACCCAAAGGCGCTATAGACGGTTGGGCAGT | 4 |
| wt-101640-101760 | 843 | AGGCTCGGAGTGTCTGGGAGAGGTTCTGTGCGGCCACGGTTGTGGCTCCCGTGACCAGGCAGTCCATCGTGTGGTGGAGGCAGCTAACGCTGGTGCTCTTGCCAGCCCCCGCCGTTCCCG | 8 |
| wt-101760-101880 | 844 | TAATTACATAGGCTGAAAAGGGCAGGAAGGGGGGCTCCGAGAGCTCCGGGTCAAACTCGGGGGAGAACGTCTCCATATCCGGGAGTTGTTGGACGCGGCGCCTAGCCAGGGTCCCTATCC | 4 |
| wt-101880-102000 | 845 | TCCTGACTATACGCCTCACGGAGGCGTCTGAGGTCATGTTCAACATGAACGTGGACGAGAGCGCCTCTGGCGCCCTCGGCTCCTCGGCCATTCCTGTTCACCCCACGCCGGCCTCGGTCC | 4 |
| wt-102000-102120 | 846 | GACTTTTTGAGATCCTGCAGGGAAAGTACGCCTACGTCCAGGGACAGACCATCTACGCCAACCTCCGCAACCCCGGAGTCTTCTCGAGGCAGGTGTTTACCCATTTGTTTAAACGAGCCA | 4 |
| wt-102120-102240 | 847 | TCTCTCATTGCACGTACGATGACGTGCTACATGACTGGAACAAGTTCGAGGCCTGCATCCAGAAGCGATGGCCGAGCGATGACTCGTGTGCGAGCCGGTTTCGTGAGTCCACCTTCGAGT | 4 |
| wt-102240-102360 | 848 | CGTGGTCCACGACCATGAAGCTGACCGTGCGTGACCTGCTGACCACCAACATCTACCGAGTGCTACACAGCCGCTCCGTGCTCTCCTATGAGCGTTATGTGGACTGGATCTGCGCCACCG | 4 |
| wt-102360-102480 | 849 | GCATGGTGCCCGCCGTTAAGAAGCCCATAACCCAAGAGCTCCACTCCAAGATAAAGAGCCTGAGGGACAGGTGCGTCTGTCGGGAATTGGGGCACGAGAGGACCATCAGGAGTATCGGGA | 4 |
| wt-102480-102600 | 850 | CGGAATTATATGAGGCAACGAAGGAAATAATAGAGTCGCTCAACTCCACGTTCATCCCCCAGTTTACGGAGGTGACCATCGAGTACCTTCCGAGGAGCGACGAGTATGTGGCCTACTACT | 4 |
| wt-102600-102720 | 851 | GTGGCCGCCGCATCAGGCTGCATGTGCTCTTCCCCCCGGCCATCTTTGCCGGAACGGTGACCTTCGACAGCCCGGTGCAGCGCCTCTACCAGAACATTTTCATGTGCTACCGCACGCTGG | 4 |
| wt-102720-102840 | 852 | AGCATGCCAAGATCTGCCAGCTCCTGAACACGGCCCCTCTCAAGGCCATCGTGGGCCACGGGGGGCGAGACATGTACAAGGACATCCTGGCCCATCTGGAGCAGAACTCACAGCGCAAGG | 4 |
| wt-102840-102960 | 853 | ACCCCAAGAAGGAGCTGCTGAACCTGCTGGTCAAGCTCTCGGAGAACAAGACCATCAGCGGGGTCACGGACGTGGTGGAGGAGTTCATAACGGATGCCTCCAACAACCTGGTGGACCGCA | 4 |
| wt-102960-103080 | 854 | ACCGTCTATTTGGCCAGCCCGGGGAGACAGCTGCACAGGGCCTAAAGAAAAAGGTCTCCAACACGGTGGTCAAGTGTCTGACTGATCAGATAAACGAGCAATTTGACCAGATTAATGGCC | 4 |
| wt-103080-103200 | 855 | TAGAGAAGGAGAGGGAGCTCTATCTAAAGAAGATCCGCTCCATGGAGTCTCAGCTGCAGGCCTCCCTGGGTCCCGGCGGCAACAACCCAGCGGCGTCAGCCCCCGCCGCAGTTGCGGCAG | 4 |
| wt-103200-103320 | 856 | AAGCCGCGTCTGTAGATATACTGACGGGCAGCACCGCCTCCGCAATCGAAAAGCTGTTCAACTCCCCGTCCGCCAGCCTGGGTGCCAGGGTGTCTGGTCACAATGAAAGCATCCTAAACA | 4 |
| wt-103320-103440 | 857 | GTTTCGTTTCTCAATACATCCCCCCTTCGCGGGAAATGACTAAGGATCTGACTGAACTTTGGGAAAGCGAGCTGTTTAACACCTTCAAGTTAACACCCGTGGTTGATAATCAGGGGCAGC | 4 |
| wt-103440-103560 | 858 | GTCTCTACGTCAGATACTCGTCAGACACGATCTCTATATTATTGGGCCCCTTCACCTATCTGGTGGCAGAGCTTTCACCGGTGGAACTCGTGACAGATGTCTACGCCACCCTAGGCATCG | 4 |
| wt-103560-103680 | 859 | TGGAGATCATCGACGAGCTCTACCGGAGCAGTCGCCTGGCCATCTACATCGAGGACCTCGGTCGAAAATACTGCCCCGCGAGCGCGACCGGGGGAGATCATGGCATCCGGCAAGCACCAT | 4 |
| wt-103680-103800 | 860 | CAGCCCGGGGGGACACGGAGCCTGACCATGCAAAAAGTAAGCCTGCGCGTGACCCCCCGCCTGGTGCTGGAAGTTAACCGCCATAACGCCATCTGCGTGGCCACCAACGTCCCTGAGTTC | 4 |
| wt-103800-103920 | 861 | TACAATGCCAGGGGGGACCTTAACATCCGAGACCTCCGGGCCCACGTCAAGGCCCGGATGATCTCGTCCCAGTTTTGCGGCTACGTCCTCGTGAGTCTGCTGGACTCCGAGGACCAGGTC | 4 |
| wt-103920-104040 | 862 | GACCACCTCAACATATTCCCCCACGTGTTCTCCGAGAGGATGATCCTGTACAAACCCAACAATGTGAACCTTATGGAGATGTGCGCCCTGCTCTCGATGATTGAGAATGCCAAGAGCCCC | 4 |
| wt-104040-104160 | 863 | TCCATAGGCCTCTGCCGGGAGGTGCTGGGTCGCCTGACCCTCTTGCACTCCAAGTGCAACAATCTGGACTCTCTGTTTCTGTACAATGGGGCCAGGACGCTGCTGTCCACCCTGGTCAAG | 4 |
| wt-104160-104280 | 864 | TACCACGACCTGGAGGAGGGGGCTGCCACCCCCGGGCCGTGGAATGAGGGCCTGAGTCTCTTTAAGCTGCACAAGGAGCTGAAGCGCGCCCCATCCGAAGCCCGGGACCTCATGCAGAGC | 8 |
| wt-104280-104400 | 865 | CTCTTTCTGACCTCGGGGAAGATGGGGTGCCTGGCCAGGTCACCCAAGGATTACTGCGCGGATCTAAACAAGGAGGAAGATGCCAACTCGGGCTTCACATTTAACCTGTTTTATCAAGAT | 4 |
| wt-104400-104520 | 866 | TCTTTATTGACCAAGCATTTCCAGTGCCAGACCGTCCTCCAGACCTTGAGACGCAAGTGCCTCGGGAGTGACACGGTCTCAAAAATAATTCCCTAGAATAAACTGAGAACAGTCATCAGT | 4 |
| wt-104520-104640 | 867 | AAATCTGTCTCTCGCGTGATTTCCATAGGAATGGTGTAGCCGGGGTGGAGGGCCGATATCACATCAAGCAGAAAGGCCATAATCTCTCGAAAGTAGGCGGTGGGGCTGAGACCATGCTCA | 4 |
| wt-104640-104760 | 868 | GTGGCCGTCTGGCAGGGGGCCGGGCGCGCTCCGTCCTTGTCCAGGAGACACACGTGGCTTCCAGAGAGGCGCAGCCCAGCCCTCCGCAGCCGCTGAAGCCAGGCTCGCGGAAGAGCCCAA | 8 |
| wt-104760-104880 | 869 | AACCTGTTTCGGCGCCGCCCGGGGGCCAGTCTCCGGGTCAGGTCGCGGACCAGGGTCAACAGGTGGTCGTGGGATGGCGGGGCCTTGTCTGCCTCGGGTCTCGCCGCTAGTTGGTCCAGG | 8 |
| wt-104880-105000 | 870 | GTCCAGGAGAAGGCTTCGTGCCAAACCAAAAAGGGCCCCGAGTGCTCCCTACATCCACCCACGTAAAGATCCCCCTGAAAGATGGCCATCAGTAGGCACCCGGGCCCGCGTCGAGCCTTC | 4 |
| wt-105000-105120 | 871 | ACCCGAATGTGTCTGCGGGCCACGGTGGCCTCTCCACCCATCACATCCCGGTCGAGCCGGCTGGCATCCTCCGAGTCTTTCACGCCTTGCAGGAAAGCCTAGGAGATACAGCAACAGAAA | 4 |
| wt-105120-105240 | 872 | GCTATTAGCCGGTGGTTCCCCCACCATCATTCTTCCTGTTAACGGGAAGAATAAGAGTTGGGCAAACCCCGGGGGCCGCGCTCTCCCACCCAGCCCCGCTTCTCACCTGTGCTAGTGGCT | 4 |
| wt-105240-105360 | 873 | CCTCTGAAGGATGGGCGGAGGTTGGTGCCACAAAGCCCAGGATGAACTCGTCTGCATAAGCCCAGGTCAGTCCTAGGTCAGCGGCCGCGTGTAGGAGAACCCGGGTGACGGCGGTGTAGA | 4 |
| wt-105360-105480 | 874 | GGCCCCCGAGTGCCCGTCGCGTGTCTGAGGTGCCATAGCGGTGAAGGGCCCGCAGCCAGGTTTGCGCGTCCCGCGCCTGCCCTCCGCCATCAGGCGTTCCCACGGGGGCGCCCCTGGCAG | 8 |
| wt-105480-105600 | 875 | AGAGGTGGCAGCGGGCCAATTCGTAGAGCCACCAAGTGGCATCAGCCTCAAGGATGGCTGTGGCCTCCGCGCGCCCGACCACCGTCGTCTCGTCCTCCCCCCCTCCCTCGCCGCCTTCCC | 8 |
| wt-105600-105720 | 876 | GCGTGCAAACGTGGCGAGGGTTAATCTCCTTTCGGGTCGGGGGCCAGATTTGTTGTAGGAGCAGCGAGCCGCGTCGTTGCCCTGACCGCGCGTCGAGGCCCAGGAGGGCGTCTGCCAGGG | 8 |
| wt-105720-105840 | 877 | GCGTCCCAGAGACTCCCAGGTTCAGGTCCAGTAGCAGGAGACCCTCGCTGTGTGGCGCCCGGTGCCAGAAGGCCGGCCTCGCCCGTCCCACATAATGGATGGGCAGGAAGGGAAAGCCCG | 8 |
| wt-105840-105960 | 878 | GGACATAGGGCTGGAAATCTGAGCCCCCTGGGCAGAGTTCGGGGTCCAGGAGGTAGAAGATGGGCTTGGTGCCTCTGTGGTTGGCGTAGCAGGAGGCATAGATACTGCGGAGGAAGGCGT | 4 |
| wt-105960-106080 | 879 | AGAGCCCGCCCCCGGCCATACTCCAAGAGTTGACAAGCCAGGACTCGAATCCCCCAGCCGGCTCAAGAATTTTCAGGCTGACGCGGTGCCGTCGGGCGTCCCCACCACGGCCGGTGGCCC | 8 |
| wt-106080-106200 | 880 | CGTCGGACGACACCAGATCTACTTCATAAGTGACCGGTCGCAGGATGTCCCTAAAGGGGACGGGAGAGGGGTCGTCGGGAGTCTCGGTGGAATAGGTGAAAACATCCCCACGCGGTGTCC | 4 |
| wt-106200-106320 | 881 | TGATGTATACGTCCAACTGTCCGGGAGACTCAGAGTGCCTCTGAGCATGGGGGCATGTCTGTTCCCCCTCCATCTCGGACCCGAAGCCATCAACAGGTGGGGGTTGTTGGTCCCGCCCAT | 4 |
| wt-106320-106440 | 882 | CATCCCCCGAGCAGCTTTGGCAGACCACCTGTGCTGGAAAGAGAGGCTGGAAGATGAGGCCCTGCTCATCCTCCACCCTGGCGGCGGACAAGAGTCTGCGGTCTCGGGTTCTAAATGAAA | 4 |
| wt-106440-106560 | 883 | GGTCAAATAGGTCCTTCTCGGCGGCATCGGCGAGCATAGCAATGAGCCCCCCGCTGCGCCTGAGCTCCCGCTCCCATCGCAAAAAGTTGAGTTCGGTAGTCGAGGGCGCGTTGACCACGG | 4 |
| wt-106560-106680 | 884 | GGGGCTCCAGGGAGCCTCCAAGCGGCGGCTGGCAGGCCTGCACCACGATCAGAGTCTCAACGTCCTCCCTTTTGATGGGCACGATGCCCACGACCCAAATCGCCCACCACCGCCCTGCGG | 8 |
| wt-106680-106800 | 885 | TCTGGGTAACATTATAAAAGGTAACCGAGCTGACGCGGGCCCTGACGCTCTCCGCGGGTGTTTCCATCATTGTTTGAGATCTGAGGAGGACTGGACCCTTTAAAACATCCGGTCACGCCC | 4 |
| wt-106800-106920 | 886 | TTTGCAAATTATTTAAAAGGTGAATGCTCAACTGAGACCATCGCAATCATGAAGTCCTCCAAGAATGACACGTTCGTCTATAGAACGTGGGTCAAAACGCTTGTTGTGTACTTTGTGATG | 4 |
| wt-106920-107040 | 887 | TTTGTCATGTCGGCGGTGGTCCCCATCACCGCCATGTTCCCCAACCTGGGGTACCCCTGCTACTTTAACGCACTGGTTGATTACGGGGCACTTAACCTGACCAATTACAACCTGGCCCAC | 4 |
| wt-107040-107160 | 888 | CACCTGACCCCCACGCTCTATCTGGAGCCGCCGGAGATGTTTGTCTACATCACACTGGTCTTTATCGCGGACTGCGTGGCTTTCATCTACTACGCCTGCGGCGAGGTGGCGCTAATCAAG | 4 |
| wt-107160-107280 | 889 | GCCCGAAAAAAGGTCTCGGGTCTTACAGACCTCTCGGCCTGGGTCTCGGCAGTGGGCTCCCCAACCGTGCTGTTTTTGGCCATCCTCAAGCTCTGGTCCATACAGGTCTTCATCCAGGTC | 4 |
| wt-107280-107400 | 890 | CTTTCCTACAAGCACGTCTTTCTCTCGGCCTTTGTGTACTTTTTGCACTTTCTGGCCTCAGTTCTACACGCCTGCGCATGTGTGACCCGCTTCTCCCCGGTCTGGGTGGTCAAGGCCCAG | 4 |
| wt-107400-107520 | 891 | GACAACTCTATTCCCCAGGACACCTTCTTGTGGTGGGTGGTCTTCTACCTGAAGCCCGTAGTTACAAACCTGTACCTGGGGTGCCTTGCCCTGGAGACGCTGGTCTTCTCGCTCAGCGTG | 4 |

| | | | | |
|---|---|---|---|---|
| wt-107520-107640 | 892 | TTCCTGGCCCTGGGCAACAGCTTTTACTTTATGGTGGGGACATGGTGCTGGGAGCCGTGAACCTCTTCCTCATCCTGCCCATCTTCTGGTACATTCTGACGGAGGTGTGGCTGGCCTCC | | 4 |
| wt-107640-107760 | 893 | TTCCTGCGGCACAACTTTGGCTTCTACTGCGGCATGTTCATCGCCTCCATCATCCTGATCCTGCCCTTGGTCAGGTACGAGGCCGTCTTTGTCTCCGCCAAGCTGCACACCACTGTGGCC | | 4 |
| wt-107760-107880 | 894 | ATCAATGTGGCCATCATACCTATCCTGTGCTCGGTGGCCATGCTCATCAGGATATGCCGGATTTTCAAAAGCATGCGCCAGGGCACTGACTATGTCCCTGTCTCGGAGACGGTGGAACTG | | 4 |
| wt-107880-108000 | 895 | GAGCTAGAGTCAGAGCCGAGGCCTAGGCCCTCGCGCACGCCATCACCCGGGCGCAACCGCCGCCGCTCTTCTACGTCCTCATCTTCCTCCAGGTCAACCAGGAGACAGAGGCCCGTCTCT | | 8 |
| wt-108000-108120 | 896 | ACCCAAGCCCTCGTCTCCTCCGTTTTACCGATGACGACGGACAGCGAGGAGGAGATCTTCCCCTAATGCAATAAAAACTTAAAACACTGAGGTTACTTTCCCGTCATTCTTTCGGGGGAA | | 4 |
| wt-108120-108240 | 897 | CGAGGGGAGGCGGGAATTGGGTTAAGATAGGGGCGAAGGGTGGGGGTGGGTGCAAGAATTGGGGCTGGGAATGGAGAGGGGAGTGGGCTAGGTGCCGACACCGGGGTGCCAAGATAATGG | | 4 |
| wt-108240-108360 | 898 | ATTGAGTAAGCATGGGGCTCTGATCGGGTCCGCCGGGTTCTCAGGGGTGTAGTGGGTGGGCATTGCATATTTTTGCCGCGGTGCTGTTGGGCCTTGGACTCGGGGTGATCATCCGTACCA | | 4 |
| wt-108360-108480 | 899 | TCACCCGCACCCGCACCCCAGTCCACAGCCACCGGCCAAGGTCCTGGGCCTCCCACCACCGTTATGCCTCCCCCTTTACCCATTAATTACAAGAGATGTTAGTTTGGTTTTTTATTTGGC | | 4 |
| wt-108480-108600 | 900 | AAAAACAGCAATTCATCATTTTCAGAGTCCTCATCATATTCGAGCCCCTCGTTGGTTTCCCCGCAGGCCCTCCCTTCTTCGGCCGCTATTAGCTTAGTAGTCTCCAGGTTAAACTCCTCA | | 4 |
| wt-108600-108720 | 901 | TAGTCATTATACAGGTTGATTATTCCCCCGTCCACGTCGCCTATGGAGTTGACTCGTCGTCGGCAAAGAGACCAGAGGGCACCCATGGCGCGGTGTCAAAAGTATTGTCTGCGTACGCTT | | 4 |
| wt-108840-108960 | 903 | GCGTGCAGGTCGCCGGGTCTAGAGGCGACCGTTTTCGAAAGAAGGCCGTCACAATGTTCACCCGGGGTGAGCAGTCTCTCCCGGGCTTGCCACCCCCACTGTGGCGGACGTAGTCTCCAA | | 4 |
| wt-108960-109080 | 904 | CAATTTTGTATTGGAGGAGCACCTGGTAGAAGTAGTTGTGCCGTGGATTGATGAAGATGTTGACTGGGACCCGGTCTTTAATACCAATGCGCCCCGCATTTTCGCTTGGGTCCGTCATTA | | 4 |
| wt-109080-109200 | 905 | CGTAGAGCATAGACTCCACCCCCCTGTTGGCAGCTAGGCTGTCTGCCACCAGGTCATGACCGGGGCCCAGTTTGCGCTTACGGACATCTTTAAGATTCCAGGCCTCATCCTGCGTCAACA | | 4 |
| wt-109200-109320 | 906 | GATAGTCACCCTCCGAGGGCAACCGCCCATCCGGGACGTACTCCACGGTAGGACGAGCTATAGAATTGATAAATCTGATAAATGACCTCTTGCATGGCCTCTTGTAAAGCGCAGTGTAGG | | 4 |
| wt-109320-109440 | 907 | ATGGGTAGATGGGGTCAAATTCTGACTTGGAAAAGAGGTACTTGAAGCGGCACTTAATCTCATAAATGCAGCTCCGGTCGGTGAACAGTATAAAGTCTCCCTGTGACTCCACATTGACGC | | 4 |
| wt-109440-109560 | 908 | AAAGATCCAGAGACACCCCAAAAATGCCATCCGTGGGACTAATCATAAAGCCAAATTGACGGTTGGCGGATGCGTCCCCGCAGATGAGCTTACAGACAATGTCCTTGACCGTGTCCTCAC | | 4 |
| wt-109560-109680 | 909 | ACCGCAGGCCAAAGGCCACAGGTCCCCCAAAGTAGTGATTTGTGGAGATGGGAGCTGGCTCAAACACCTTGGTGGGTCCATTCTTAATGGTGGAGAGCAGCTTGGAAGAGGAAATTATGC | | 4 |
| wt-109680-109800 | 910 | CATTTCGCAATATGTCCCACATCAGGTTCTCAGACTGCCCCCTGGTCATGGACTCCACGTACGAGCAGAGAACAGTCCTCTGCTCGTCGGTGGCCTCCTGTAGCCCCCAGTAAATGGATT | | 4 |
| wt-109800-109920 | 911 | TCAGGGAGGGACCGTCCTTGCTGTCATTCTCTTGGACTAACGAGGAGACAAAGTCACAGAAGCCAGTTTCACCAGAGAACTCTTGTATTTGTTTACAGAGGCAATAGAGATAGACAAAGC | | 4 |
| wt-109920-110040 | 912 | GCATGGCCGGCATCTGAGGTGGACGGTCAAGGTTACGGACAAAGGCCTCAGTCTCCGGACTGCGGAGGAAGCGGGCAAACGTGTAGGAGGTCATCTCCTCCATGGGATCCTCGAGCTCAT | | 4 |
| wt-110040-110160 | 913 | CCACGTCGGCCATCTGGACCAAAGAAGTCGTCTGCCAAGAGTTCAGCTACCAGACCTGGAAGATGAGGGTGCTCAAACCGTGGGCGACAGTTGAAGAAGTAGCTCTCCTTGAACCTCTTT | | 4 |
| wt-110160-110280 | 914 | TTAAGGCTCCGGCACCACTGCAAGAATTGACTCATATGCTCCGCCGTGACATCCACGCACGGACTCTCGCCACACGAGGTCAGGCCCATGTCTAAGTTCAGGTTCCACATCTGCGACAGC | | 4 |
| wt-110280-110400 | 915 | ACCTCCAACAGCACCACCTTTGGGGCTGCAAATTGCAAAAAGTAGAGCGGGTCGGATCGGTCAAATCCCATGTCAGGGTTGGGGTAGGGGATTTTGTGGGTGGAGTCAGCGAGGTGCATG | | 4 |
| wt-110400-110520 | 916 | ATACCATAGAGCAGCGAGTAGCCGAGCGACTGCAGATCCAGGCGAAGGGCCGTCTGCGCCCCCACGGGGCCACACGCCGAGGGGTCAGGGATGTGCCCAGCCCCCCTCAAGATGTAGCAC | | 8 |
| wt-110520-110640 | 917 | TTGCTCAAAAGGCAGAGGGGCTTATAGGTGTCCTTGGCTATAGAAAATGGTTCCCTCTGGCAATAGAGGCGATAGAGCTGCCGGCCCTTAGAAGACTTTAGCCGCACATCCAGCATCTTG | | 4 |
| wt-110640-110760 | 918 | TTGCGGTCGTGGAGGGAAGCAGTCCCATAATCAGTCAGGACCAGCCTACCCATGCCCCACATGGTGTCTGTGAAATCCACCAGGATGTTGCTGGGGCTAATGTCCGAATGGAAGAGGCCG | | 4 |
| wt-110760-110880 | 919 | CAGTGCCGATTCAGAAAGTAAACGGCATCTTTGAGGCCCTGAAAGCCCCGCACCAGGGGCTCAATACTACCATCATGCCAGTGGCCATAATCCTGGAGACTGCATCTGAACTGGGGCATA | | 4 |
| wt-110880-111000 | 920 | AACAGGGCGTGGCAGGACGTGCAGGCCGACAGGTAGTCCACCAGGGCCTTGTCCTGCCCATCCTCGGCCGTGGCCTTCCCAATCTGAATCATGTCACACACCATGAGCTCGTGATACAGC | | 4 |
| wt-111000-111120 | 921 | TCCGTCACAGAGTCATAGAGTTTGACCGTGGCATTATCTGCATGTGCATACACGGCCCCGTAGCTCCCCCGCCCCAGCAGATACTCGCAGGTAATGGGGAGGTGATCACAGCGCGTCATG | | 4 |
| wt-111120-111240 | 922 | TTCTCCGGCAGCTTTACATAGAGGGTCTCCGTCATGTCATCAATGTTGGTCACCTTCAGGTGTTTGTGCTGAAAGGTGAAGTAATCAATGACAGTCACCTTCCCCAAAAAGGCCTGGGTC | | 4 |
| wt-111240-111360 | 923 | TCTCGAGGGGGTTCTGGGGAGACACTCAACTCGCCACTGCTGGAGGAGTTCGTCGGGCTCAACTCCGCAGCCATATTCACATCCATGTTCCTCAAATGGCTCGAGGGCCTGTCGCAGCTC | | 4 |
| wt-111360-111480 | 924 | GTCTCTGGCCTCAAGCTCCTGCTCACGGAGCTCCTCCACCCGCTCTAGCTGCTTGTAGTTGATTTTTGGAAATTGAGTCTTGGTCGCGGTGACCACCCTCTGATAGGTAGAAATTAGCTG | | 4 |
| wt-111480-111600 | 925 | TTTGGACTCAAACGTCTCCCTTGCGTGGCGCAGGGACTCTAAGGCACCCCGAGCAGATGTAAACTGTGTTTCAAACAGAGCGTGGTCCCTCCCAAATCTGTCACGTGCGCTCACAGCCGC | | 4 |
| wt-111600-111720 | 926 | TCTCTTTTCTACCGAGGCTCTTAGTTGCTGGGCCACCAGATCTCGCTTAGAACTACTCATCTTCATAAGTCACCATGTCCGCAACTATGGAGCCCAGATCATACGTGGGGTAGAGTACGG | | 4 |
| wt-111720-111840 | 927 | TAGTTCCAGTGGAGGCTTCCCGGTAATTTCCCACAGCGTCCACCATATATCTTTCTGCCTCTCCCGTTAGAATTAGGCAAGGATCATACGTGTCCACCGGCCTTTTATACTGAGCGTTTA | | 4 |
| wt-111840-111960 | 928 | GGTTTTGTTTATGTAGCAAGCACAAAAGGCACACACGAGTGATGCAAAAGGGTTCCTGAGGCAGCAGGCAGAGCTGTTTTGCCATTTTATTCAGGCGGCTAACGTCAAAGGGAGGAGCTA | | 4 |
| wt-111960-112080 | 929 | TATCCTCACCCTTCCAGTCACGCACGTCCAAGTACAGGGCATACACACACCTGGTGAGGTGTGCCAGGAATGCCTCTATGTTGGCACATGGTGTATAAACCGCAGTGGGTAGCAGAATAG | | 4 |
| wt-112080-112200 | 930 | GGCCCCTTTTGCCCCGTGCCGCAGCGTAAACGCAGTGACGCTCTTCGCAGTGGGACCTGGGGCCGTAGAAGAGGGCCCACATCCAAGGGAGTGGGTCTTCAGGCACCAGGGAGGTCCAGG | | 8 |
| wt-112200-112320 | 931 | TTTGGGAGTGGGCCAATATTTGCAAGGCCTGACCTATAACCTCATCTTTGTTCCAGGCCAGCGCAATTCGCATAAGGTCCCCATCAAACACCTCAAAACACAGACCCATGCCCATTTCAG | | 4 |
| wt-112320-112440 | 932 | GCTGAGAGGGCTCCATCCGGCTCGACCAACCTTGTCCACCAAACTGCCATTCTTCTGGTAAACGGGGGTTGAGGGGCAAGAGCTCCAAAGCCAGGCTCGAGAAGTCATAGTCATCCTCGG | | 4 |
| wt-112440-112560 | 933 | CCACACGGCCGGAGCTCCGGGCCTCGTGCCAGGGCCTGTTGTCCTGGGGGAGGATATTGGACACGAGCAGGAAGCTCTTGAGTGGCGTCTCCACCAGCTTAAATTGCTCGGGCGTGTCCT | | 4 |
| wt-112560-112680 | 934 | GGCAGGCCTCCAGTGCCAGTTCCAGACACTGCCCATACCTGCGGGCGAGCATCGGGTCATCGGGCATATCGGCCTTGACCGCGTTGAACATGCTGTATGCCTCGCAGCGCGGCCGTCTGA | | 4 |
| wt-112680-112800 | 935 | CCGAGAACCTAAGAAACGCCCTTCAGCAGGACAGCACCACGCAAGGCTGCCTGGGTGCCGAGACCCCGAGTATTATGTACACAGGGGCCAAGTCAGACAGGTGGGCTCACCCTCTGGTGG | | 4 |
| wt-112800-112920 | 936 | GCACAATTCACGCCAGTAATTTATATTGCCCAATGCTTCGAGCATACTGCCGCCACTATGGCCCCAGGCCCGTGTTTGTAGCTTCTGATGAATCATTACCCATGTTCGGTGCGAGCCCTG | | 4 |
| wt-112920-113040 | 937 | CCCTTCACACCCCAGTCCAGGTCCAGATGTGCCTACTACCAGAGCTACGCGACACGTTACAGCGCCTGCTGCCACCACCCAATCTTGAAGACTCCGAGGCCTTGACGGAATTCAAGACCA | | 4 |
| wt-113040-113160 | 938 | GCGTGTCCTCTGCCCGTGCCATCCTTGAGGACCCCAACTTTTTGGAGATGAGAGAGTTTGTCACCAGCCTGGCCAGCTTCCTGAGTGGTCAGTACAAGCACAAGCCCGCCCGCCTAGAAG | | 4 |
| wt-113160-113280 | 939 | CATTCCAGAAACAAGTAGTGTTACATTCTTTTTATTTTCTGATCTCAATCAAATCTTTAGAGATTACAGACACCATGTTTGACATCTTTCAAAGTGCTTTCGGATTGGAAGAAATGACGC | | 4 |
| wt-113280-113400 | 940 | TGGAGAAGCTGCACATTTTTAAGCAAAAAGCCAGCGTGTTTCTTATCCCCAGGCGCCACGGCAAGACCTGGATAGTCGTGGCCATCATCAGCCTCATCCTCTCGAATCTCTCCAACGTGC | | 4 |
| wt-113400-113520 | 941 | AAATAGGCTACGTGGCTCACCAGAAACATGTCGCGTCCGCCGTTTTCACTGAAATTATTGACACCTTGACCAAGAGCTTCGACTCCAAGCGTGTAGAGGTCAACAAGGAGACCAGCACCA | | 4 |
| wt-113520-113640 | 942 | TCACGTTTAGGCACAGTGGGAAAATCTCCAGCACCGTAATGTGTGCCACCTGCTTCAATAAGAATGTAAGACCTGACGTTTCAGTACTTGGCAATTGTAGAGCATAGCCCGGCTGTAAAG | | 4 |
| wt-113640-113760 | 943 | GTCAGAAAATCGCAGCAGGGTCCAAGGTTGTGCTGTACATGGGACCTCTTTCCCATTAGCAAGAACCCCCTGCAGGACACGTGACATGTCCGGGTGCATTTTGGGTGGGTTAAATCTCAG | | 4 |

| | | | |
|---|---|---|---|
| wt-113760-113880 | 944 | TCCCACCACAAAGGGGGCATCCTCCGGTTTGAACATCAGACCCAACAAAGCCCGATGCCCAGTTATGGGTACGTAGTCGTTGTTCAGGGCCGTGCATGGCAGCAGACAAGGACAGGTGCC | 4 |
| wt-113880-114000 | 945 | AGATGTGCCTGGGCTATCGTCCTCCGTCCAGCCACGCAGGATGTTCACGTGGGCCCCGGCACCATAGCATGTCACACATTCCCCGTTATCACATCTGGTTAGCAGGTTGATAAAATGGGT | 4 |
| wt-114000-114120 | 946 | CAGTGATGGAAAGGTTGGCATATTGGGGCAGCACATCAGCATGTCCATGTTAACGAAAAACATGTACAGGGCCCCTTCTGCATACCAGGCACCACCCCGTCCCAGTGGGATGATCTCCGA | 4 |
| wt-114120-114240 | 947 | GGGTGTGATATCTTGCAGTTCTTCTACTGTTTTAACGGCGGTTGAGGTGGTAAAGACGTGGGCCGTGGTCAGATCTGTGCAGGTGACTACAGGGTTACCCCTAATCTCCACAGGCACCGC | 4 |
| wt-114240-114360 | 948 | CTCACCCACTGCATCTGAGAATACCCCAAAGTACATGAGAGTCAGGCTGTGTGGCCCCTGGACTGCCTTAGTGAAGAGAACCTCGGGCCTGGCCACGGTGGCTAGGGTTCCATTGATGTA | 4 |
| wt-114360-114480 | 949 | GACGGTCACATAGGTGGGCTTCTTCTTGGGCTTCAGCACAATGAGGGTAACATTCATGTAGGTTTTAGGAGGTCCGGCTATCTGAGGCACGTACACAGCTGACACGGCGGTTGTGGCCGT | 4 |
| wt-114480-114600 | 950 | ATAGACTTTCATCTGGGGCGTAGAGGCATCGCTCAGCACCCAGAGGCACTCCTTGTTGAGGAACTTGCGAAGCTGTTCCCGGCTACTGTTCGCGGCGGATGCCATGACGTGCCAGAATAT | 4 |
| wt-114600-114720 | 951 | ATCCCCTCTCCTCGGGGGTGAGTGCCAATTGGCCTTTAATAACAAAGCCCCCAGGCAGCACCAAAAATGCCTGCCCGTCCGATGTGGTGGCCAGGTGGACGCAGTGCCCGTCAGTTCCAA | 4 |
| wt-114720-114840 | 952 | GGGCTACTAGCTGGGAAGCAGCCCCAACCAGCCCACCCGGGGGCCTGGAGTCGATCACCTTACCCCAGGCCGAGGCCCCTTCCTCATACAGCGGGTGGCTATCTATCCATAGGCAGGCAT | 4 |
| wt-114840-114960 | 953 | CCGGCGTCTTTGGTGCATTGGAGATAGTAGCTTTCACCCAACAACTTTCCCAACTAACCCGTGTCTGGACAGTGAAGAACGCTTCCCTGATCAGGTCTGAATTTTTATAGATACGGGAGT | 4 |
| wt-114960-115080 | 954 | AGGAGGTGGGAATAACAACTGGGATTTCTTGTTGTGCTGTCCAGGCCTGCATGGCCAGTTTTTCCCTGAAGCTAGCAGAAATTCTGAGGGCCACTGAAATGAGGAAGCGAAACTCCCTCT | 4 |
| wt-115080-115200 | 955 | CTGGAGCTCCCAAAATTGAAACCTCAGCAAGATCTGTTGCTGGGGAGGCATGGGTGACAGCTGTCATCCTGTGCAGTCTGCCCTGGGCACTCAGCTCTGGATATGTGACAACATAGAGAG | 4 |
| wt-115200-115320 | 956 | CGTGGGGGCTAAAAATATGAGCAATTCCCCTGACCAGGGCCCTGGACTCACGAATGGCCCGACGGGTCTTAGAGAAAGAAACAGGCACCCTCGAGAGTGCCCCCGACCCGACCCCCACAG | 4 |
| wt-115320-115440 | 957 | TGCCGCCAGTCCCTGCTCGGCCTCCGCCGCCTTCCCCACCGGCGCTGCCCCGGATGTTGCTGGGGTTCTCGAGGGCTGGGTGGTGCTTGGACACAGAGGTCTCAGCAGCCGCCTTGGTCT | 8 |
| wt-115440-115560 | 958 | CGGCCCCGGCCCTAAGTCTGAGCCCCAGGCAAAGGGCCGGACTCCCAGCGTGGCCCAACCTCTGCTCCCCTCTATTCTCCTCTTGCGTTATCTCCAATAGAATTTGCTTGAGGTCATACG | 4 |
| wt-115680-115800 | 960 | CCTCTCTGATAAGATGCCCGTCGGTCAGAGCCCTTTTGGCCCCCTCAAAGAGAGACAGGTAATAAATCTGTAGCTCCCCAACCAGCCCTCCTTCATCGTAAAATCGAAGGGCGGCCACGT | 4 |
| wt-115800-115920 | 961 | GGAAGGGGTTGTAGAGCTCTGGAAGGCCCTCATCGCAGTACACTGGCACACTGGTAAACGTGCCCCGATGGCTAGGCCGTCCGGGCAGCATGCCCCGAGCAGCAAACACGCGGCAGACCC | 4 |
| wt-115920-116040 | 962 | TCGTGAGACCCGTCCGGTCACTGAAGAGAGTCTGGCACCAGGCCCCCTCGCAGTTTGGCACGCGATTGGGGCAAAGCTCTGCCATAACCGTGTCGGGAACAAATAGGTGCACGAGGAGGG | 4 |
| wt-116040-116160 | 963 | GGGTCCCGAGGCCACTCAACACTTGGTTGTCAATGTGGACATCCATAGCTCTCTCATGCGTTTGGCTACAGCATCATAGCGCTTGTTTCTGGTGGATTTAAATAACAGGGCCCCGTAGAC | 4 |
| wt-116160-116280 | 964 | AGTCTTTTGTGAGTAAATAGAGATGATGACATGGATGTAGAGACTGAGGACCACATCCACCACCTTCTCGGAGGAGGCCCCCCTAAACAGCATCAGGCAGCAAGGGAACACAAAGGAAAC | 4 |
| wt-116280-116400 | 965 | CAGGGCCGGGATGTGAGGCCTCAGCGCCCCCTCCTGATCAAAGAGGGCCTCGCTGACCCCGGAGATGACATTCTCATTCAGAAAGTAGTGATAGAGGTGATTGACCACAGTCTTAACCAG | 4 |
| wt-116400-116520 | 966 | GCCCTGGACTTGTTCAGGCTCCCACTTGTCCCGCTGGTCCTGTGTGTCTTGTCGGATCTCGGTCCAGGGCCTCAGCGCCGGCTGGAAATGCGGCCCCATGTAGTTGCCTGTAAGGGCGCA | 4 |
| wt-116520-116640 | 967 | CACCACTCCCTCATGGGTCTCAATCAGGGTGCACTCGCTGGATCCATCACATACGTGGTACTCGCCACAGCCCCAGCAGGCAAACACGGAGGCCATGCTCTCAGGTAACGGGAGATGGAA | 4 |
| wt-116640-116760 | 968 | CTCCAGCTTACTATACGAGCACAGGTGGCGAGGATTGGGCTCATCCGTGCCCCCCTCCCCCCGCGGGAGGCTCAATCGGCCTTGGTCTGACATTCCACCCCGGCCAGGTCCAGGAGGGTG | 8 |
| wt-116760-116880 | 969 | CAAATATTCTCCAGGCGCTGCACCTCAGAGACCTCCTGCTCAAAGAGACCTCCCACCGCCACGTAGACGCGGGCCACCGTCCGGGGAAGGTCAGTGGGGTCCCAGCTCAGCAATTCTCCA | 4 |
| wt-116880-117000 | 970 | AATTCTCTCTCCCCAATAGTGCCTCGCTTCTTATCCTGTCTTTCAGAGCATCCGGGGGCAGACATTTCACCTCTTGTTTGTGGACGAGGCTAACTTTATCAAGAAGGAGGCCCTGCCGGC | 4 |
| wt-117000-117120 | 971 | GATCCTGGGCTTTATGCTTCAGAAGGATGCCAAGATTATCTTCATCTCGTCTGTGAACTCGGCTGACCAGGCCACCAGCTTTCTTTATAAGCTGAAGGATGCTCAGGAGCGGCTGCTGAA | 4 |
| wt-117120-117240 | 972 | CGTGGTAAGTTATGTGTGTCAGGAGCATCGGCAAGATTTTGACATGCAGGACAGCATGGTCTCATGCCCCTGCTTTCGCCTGCACATCCCGTCCTACATCACCATGGACAGCAACATCCG | 4 |
| wt-117240-117360 | 973 | AGCAACCACCAACCTCTTTCTGGACGGGGCCTTTAGCACCGAGCTGATGGGTGACACCTCCTCGCTGAGCCAGGGTAGCCTGAGCCGCACTGTGCGTGACGATGCCATCAACCAGCTGGA | 4 |
| wt-117360-117480 | 974 | GCTCTGCCGGGTTGACACCCTCAACCCCCGAGTAGCCGGACGCCTAGCCTCCTCCCTCTACGTGTACGTTGATCCGGCCTATACCAACAACACATCCGCATCAGGCACCGGAATCGCCGC | 4 |
| wt-117480-117600 | 975 | CGTGACTCACGACAGGGCGGACCCTAACAGGGTCATCGTCCTGGGCCTGGAACACTTCTTCCTCAAGGACCTAACAGGGGACGCTGCCCTCCAGATCGCCACCTGCGTCGTGGCCCTCGT | 4 |
| wt-117600-117720 | 976 | CTCCTCGATCGTCACCCTGCACCCCCACTTGGAGGAGGTGAAGGTAGCCGTGGAGGGCAACAGCAGTCAGGACTCTGCGGTGGCCATTGCCTCAATCATTGGGGAATCCTGCCCCCTCCC | 4 |
| wt-117720-117840 | 977 | CTGCGCCTTCGTGCACACCAAGGACAAGACGTCCAGCCTGCAGTGGCCCATGTACCTCCTGACTAATGAGAAGTCCAAGGCCTTTGAGAGGCTCATCTACGCAGTGAACACGGCCAGCCT | 4 |
| wt-117840-117960 | 978 | TTCTGCCAGTCAGGTCACCGTCTCCAACACCATCCAGCTCTCCTTCGATCCGGTCCTCTATCTCATCTCCCAGATCAGGGCCATCAAGCCCATCCCTCTCCGCGACGGTACCTACACCTA | 4 |
| wt-117960-118080 | 979 | CACCGGCAAGCAGCGCAACCTCTCTGACGACGTGCTGGTTGCGCTAGTCATGGCTCATTTTCTCGCAACAACACAGAAGCACACGTTCAAGAAAGTTCATTAAACTTTATTGACTACACC | 4 |
| wt-118080-118200 | 980 | AGTCCCTTGTAAAGCGACGGGTCTCGCGTGACGGCATTCGTGAGCAGGGCTTCGTCCAGGGGCTTGTTCTTGGCGGACATCATTAGCCCAGCCGCAAATATCAGAATTAGCATCAGAAAA | 4 |
| wt-118200-118320 | 981 | GTGAGCCCCACAAACACCAGTGTCCAGAGAGGAAGACCGTAAGATAAAGATGGCTGCCTCTCATCTGGAACGGTGGGAAGCTCAGCAGTTGTTTTTGTGGCATTGGACGTCCCTTTGGAG | 4 |
| wt-118320-118440 | 982 | GACAGCGTGGGGGCCAAGGTGGTAGCGTTGGTAATACGGGTAGTAGCACTGGTGGTGGAGGAGGACCTGGTGGTGACATTGCTAGTCACACCCGTGGAGGTTCCTGTTCCGGCCTCGGTG | 4 |
| wt-118440-118560 | 983 | GCAGTGATGTTCTGTGCAGTAACCTTAGTGGTGACATTGATGGTGGATGCGTTGGAAGTTGTTGGGACTGGTGTGACAGTTGTCCCAGTGAATGTCACCGTGGTTGTGTTGGTGCTCAGA | 4 |
| wt-118560-118680 | 984 | ATAGCAGTTGTGGTTATAGGGGCGCTAGTCGTGGTCAAGGTCGTAGACTGGTTTGTGCTAGGACCCGATGCCGACGGTGATGGTGTAGTCACAGCCGTTGTGCCTGTCACGTTCCCCGCC | 4 |
| wt-118680-118800 | 985 | GAGGCCGTCGAACTGCCACTAGATGTCCAAATAAGGCTTGTCTCACAGATGAGTATCATGGCCATAACAGCGCCTGCCTTGTCTCTGGCGTGTGCCATCGCGTCTGGACGCAGAAGGCCT | 4 |
| wt-118800-118920 | 986 | CCCGGCCTCTTTTATAGCTAGTCTCCACACCCAATACTCTACTGAACCATCACATACATGACCTCCTCGAGGTATGCAGGGAATGAGCGGTCCGTGAGCCGGTCAACACGACATTGCTTC | 4 |
| wt-118920-119040 | 987 | CGTTTCATGCCTCCAGCTGCCCCTGACCAGTTAGGACCCTTGACGGATGTCTTTAACGGCGCGGTGCAGTTGGTCACCAATGACGGCCTAAAGGCCAACACATCCTTGAAGCAGGGCGTA | 4 |
| wt-119040-119160 | 988 | GGAATGGTACCAAACTCGGGGCCCACCCCATCAAAGACATAATATGTCTCATAGTGGCAGTGATGATGCATCACCACCACAGCACTCGCCAGGACCCTCTGCATATCTTGTACAAGGCGC | 4 |
| wt-119160-119280 | 989 | CTTTCAACTCGGCCACTGGCTCTGGTGACGTTAAATGTCCTGTTCCTATTAGTCACAGCCTGTAGATTTGGGCACCCAGACTCAAAAAGTGCAGCTACATGAAGGGCAGCCGCCTCAAAT | 4 |
| wt-119280-119400 | 990 | CCACCATGACCCCCATGGCTGTCCGTGTTGTTGGGGTAATAAGTCACATTGTTAATGACCACGGCCGGGATAAGGGTGTAAACCTTGCAGAATGGATTGGTCGGACACCCATAAGACAGG | 4 |
| wt-119400-119520 | 991 | GGCACCCCAAAATCACGCCCCTTACCCCGAAGCACCTTGGCCCCCACCGGCATAAAGCTGGGCAAAAAGAGTGGGTTAAAACCAAAGGCGAGTAGGGCCAGGAACGCCAAATAGCAGCAG | 4 |
| wt-119520-119640 | 992 | TAATAGATGAAAACAAAGCTCAGCATGAAACAGCGTGGAGGCTCAGCTAGGGTCTCTGCCTCTCCATCATAGACATCTTCCTTGAATCTCATTCTCTCACCGCATACCTCGCTCTTCATC | 4 |
| wt-119640-119760 | 993 | CAGGAGGGGGCCATGGCTGCCATTCTACCAGTTAACGAGGAGAGAGAGAGTAGGTCCGCGGAAATTGGTGCCCCTCTCTGCCCTCCTGACGAGGCCATGGTGTCATCCATCTCCGCAGTC | 4 |
| wt-119760-119880 | 994 | CGTTCTTCAGCTTTGGCATTGGTCCGGGTCCGGGTGGTCTGATTTTGATTCTGATCCTGGGTATTGGTCTTGGTCTCTCCTCCCCCATTGGCATGGATTGGCATAGGTGGGTGTGGCTCA | 4 |
| wt-119880-120000 | 995 | GGCTCAGGTTCCGGCCCTGGGACGGCAGCAGCCGCCGGGACGGTGAAGTCGTGGAAGGTAGAGGCCCGTCCCTCCCGAGGTCGTGGGGCCGGAGCCTTATAAAAGACTTCCACCCTCTCC | 8 |

| | | | |
|---|---|---|---|
| wt-120000-120120 | 996 | CCGCTGGCCAAGACACGCCGCTCGTGGACCACGCCATCTTCCTCCCGGCTGATTGTGTGGCTGACGGTGCCGTGTTCCACCGCCACTTGTTCATCGACCATGGTACCCCCTTTATCTTAA | 4 |
| wt-120120-120240 | 997 | CCAGCAAGTGGCCGTCAGGGTCTCTTGAGAGTATGCCGCTGTGGCCAAGCGAGGCCCCAAATTAAATAGTGATGCCAAAGACTGTAGGTAGGTCATCATCACACGCATGCGTGATAAATC | 4 |
| wt-120240-120360 | 998 | ATCCGCCACTGACAGGTCATCCAGGTCTATCCGGGCTATCTCATCCGGCACCATTTCCTGGAAGAGATTCAAGAGGTCGTGATGCTCATGCCGGATAAGGCCTCGGACCAGGCGCATACT | 4 |
| wt-120360-120480 | 999 | GGCCCTGGGCAGCAGGGTCACCATGATGCAAAAGTAGAGACTCAGATTGTCCAGCAGGGCCAAGCCAAGGGGCCCTGGCACCTCCGGGAGGGCCAACTCGTAGTGGTGCCCCAGGTATGA | 4 |
| wt-120480-120600 | 1000 | AACAGAGCCAAGATGCATGTGTACATCGAGCATGTCTGCGTTCCCGGGAGCCTGCATGACAACCCGGGAGTACACGTTAAACAGGAGAATCTTCTGCAGCACCTCCTCTGCTATGGGCGT | 4 |
| wt-120600-120720 | 1001 | AGGCAGCACCATGGGGAAAACAATGTCCACATCATTGGACTCTAACTTCACGGTGGCATGCTCTCGTCCAAATACCGGGGGCATAACACTGAGGCTCCCGGTCCCATGCCACTGGAAAAA | 4 |
| wt-120720-120840 | 1002 | GGGCTGGTACTTGTTCTTAATGGCGTAGGTCTGACCTGGAACAATCTTGGTGAGTATCAAACTGTCCACGCTAACCTCATCCAGCACGGCCAGGGTGCAATCAGACAGGTAGTTGTACAT | 4 |
| wt-120840-120960 | 1003 | GGACACGTAGTCCGGGACCGTCTCTAGAGAGTACACCTGACCCAAGCCCAATCCCTGCACATTCTGCGTCCCGTGAGTGGAAGCCAGGGGTAAGATGCAGCCAATCCTCTGTTGCATCTT | 4 |
| wt-120960-121080 | 1004 | GGCAATCTCATCGGTATACAGACGAGAGGAGAGAGACACTACCACTTTCAAATCCATCTTTATTGACAATTATCAAAAAACCACCTTATTTCCAAACTTTAATATTCTTCGTACCGGCGC | 4 |
| wt-121080-121200 | 1005 | CACCTCTTCAATTATATAGTGTCCGTAATGGATGGGGGCGTGGGTCTGTTTGACAGACATAAACTCATCGATGAGTGCCCGGGAGGAGGCTGAGAGTGCGGGGAATGCCTCCTGCAGAAA | 4 |
| wt-121200-121320 | 1006 | GCTGCAGGGCTGCTCCAGAAACACGTCAGTGCCAGCAATCACTACAAACTGCACCTCTGTGTTGCTGGTGGCTGGGTGCCCTCCAAGTCGCTGGCTGTACTCGTTGACCATGTTGTAGAG | 4 |
| wt-121320-121440 | 1007 | TCCCCTGTTGTTGCGCAGAAGCTCCTCCTTGTTGAAAAATGCCCGGCAGGGGCTGTAGAGGCCCGGGACGGCCGTCTGGCGATAGGAGGAGTTGTACATGATGTCACCCAGAGAACCCAG | 4 |
| wt-121440-121560 | 1008 | CTGAGATGCCCAGGGATTCACAGTGCTCCGGTATTCATAGGCGGCATCCGGGCGAGAATGGTCATAGATGAGCCCCTCGGCAACCTCCTGATTGTAGTTTTCACAGGAGACCACACAGGC | 4 |
| wt-121560-121680 | 1009 | GGCCCGTCCCCTTGGAGAGTTGGACTTTTGAAAATAAGCCACGTCTGCCGTGACCGGTGTTACGATAATCTCACAGGTGGCCTGCTGGCCGTGGCAGAGTCCTGGAGCTCCATTAACATT | 4 |
| wt-121680-121800 | 1010 | AGTCATACCTGCCAGGTATGTCCTGGGGTCCCGAAGCAGCGTCCCATTGCGCTGAGCGCCCACCTTGGCCTTGATGTAGTCATTGACTTGCTGGTTGCCAAAGGCCTCGGCCGGAAAGAC | 4 |
| wt-121800-121920 | 1011 | GCTAAAGAAGTCTTGGGTGTGGATACCCATGTCAGTAGTGATGGCCGCCACCCTGGCCGGAGTCATGGTCGAGCTATAACTAAGCCCGGTGTCGATGGAGGCCATCTCGTGATGCACCTC | 4 |
| wt-121920-122040 | 1012 | AAAGGTTACCGCGTCCACCCTGGCCTCCCGGCGGCTAACATTTGGGGTCCCAATGAACATGGATGTTGAGGCCCTGGAGCTAAACAATATGTTTTCAGAGAGGATCTCATCGGTCCTGAC | 4 |
| wt-122040-122160 | 1013 | CACGGTCATGGCCACCCCTGGGTGGATCTTGAGCTTGGCCTGGGCAATATAGGCCATGGGGGACATCTTGATGTGCATGGCGGTCATTCCACTGATTGAAACGAGGGAAGGAAGACATTC | 4 |
| wt-122160-122280 | 1014 | GGCCGCGTATTTGCCCATGGGCGAGCGGTGCCACTCCCGGTACTCTGCAAAGAGCTGCTCTGGCCGGTTGAAGGCTTCCACGGCCCGCTGCTGAGGATTGCGCATAACAAAGGTGGCAAC | 4 |
| wt-122280-122400 | 1015 | ATCCTGGTGCATGGTGGCAGCCACTCGCGGGTCCCCGTAAAACATATGGAAAGGAATGGCGTGAAAGAGACACTGGGTGACGGCCCGGGTCCTCTCGGAGAAGGCAAAGGCCACCAGCCC | 4 |
| wt-122400-122520 | 1016 | GTTCACCAAAACAGTCTGCTCTGTCCGCTTGTCGGCGGGATTCGGGGCCAGCTGCTGCGTAACGTCATTGTCCACCGACACACGCACGGCACGGGTGAAAGTGGGGCAGGTCATGAATGA | 4 |
| wt-122520-122640 | 1017 | GGCGCTGAGGTCCCTGATCATGCCCACGGTGGGGCGGAGGTCGGAGATCTCCAGCAGATCCCTGAGCGTCCCATTCTCCAAATTGTCGAGGATGTCCTCGTCCCTGGTAAAATGGTGGCT | 4 |
| wt-122640-122760 | 1018 | GAAGGCTGGCCCGTTGTAGGCCAGGGTCTGGGCCACGTGCTGAAAGTCCACCCCGAGGCCGCACATGTGGGCATTGGTGCAGGTTGGGAGGAAAACGTAGTAAAAGATCTTTTCCAGCAC | 4 |
| wt-122760-122880 | 1019 | ATCCGCATGCCCCTCATCTACATAAGGGCCTAGGTGCAGACGGAAATCGTGGTCGTGGTCTCCGTTAACCCGGTAGCCGTACAAGGCCACAAATTGGGCAGCCATCTCATCCATGTTTCC | 4 |
| wt-122880-123000 | 1020 | AACCCTCTCAATAAACTGGGGCGCGGCCAGGGTGTCAGCGTAAACCTCATTTCCGATAATAATCTGGGGGGCCCGGTCACTAACGGTGAGAAGATGGGTGAAAATGTCTGTGTAGGCCAC | 4 |
| wt-123000-123120 | 1021 | CGGGGGGAGCAGGTTAGGGTCCAGGAGAGCGCAGACATACTGACCCACGCTCTCATCCCCCACAACATCTGACCCGGCCAGGCGCATCAGGGCCTGCTCTAGGGCTATAAGTTCCCCATA | 4 |
| wt-123120-123240 | 1022 | GATTTTTCTATACATGGAATAGGCCTCCTTGGAGATGGCGTTATTTCCCAGGTGGCGGCAGATGAACTTGATCATGGAAAAGCTGTTCACAAAGGCAAGCCTCCCTGACCGTTCCCAGTA | 4 |
| wt-123240-123360 | 1023 | GGTGTTGATGCACAGGGACACCAAAGGCACGTTCATGACAAACTTTTCCTCAAACCCGTGGATCATAGCCTCGACTACGTAGAAGAAGGCTGGATAGGCAGTGTCATAGGCAGTATCCTG | 4 |
| wt-123360-123480 | 1024 | CACAGTCTCAATAACGGCCTGATCCACCACGTGGGCCAGAGATGTGGCGGTCTCAAACTGCTGCCCCCGGGCCTCTTGGAATGCAGCTGGGGCCAGGGGAGTCGGCAGGTTACCCACCAT | 4 |
| wt-123480-123600 | 1025 | TAGCCGGTGCACAGCCCTGTGCCTGGCCCTCTCCCCGGCATCCCTGCCAATGTAAATATCATAAAGGGGGTGCAGCTCCAGCCGCAGCAGGTCATAATTGGACGGGTGGAGGAAGTCTTC | 4 |
| wt-123600-123720 | 1026 | GGTGGGCAGCCCGCACTTGAGAGCTATATCTGTCACGGGGGCTGCATACTTGTTATCATAGAACTCGTCCACAATAACAAGCACATTCATGTGATTGGGCCTCCTGTGTTGCAGGGAGTA | 4 |
| wt-123720-123840 | 1027 | GGTCTCGCGCCTGTCTCGCGGGGCCGGGGCCGCGTTGAGGCTGTTTAGGGTATGGGCGGGTGTGTGGAGTCGGGGGTGACAGAGAACCTTGAGAGCATTCTGTAGGTTAAACGCGAGGAG | 4 |
| wt-123840-123960 | 1028 | AAGGTTATTCTTGTTTACGATCCATGCCTCCACCGGTAGCTGCTGTGTGGGGTTGTCCAGCATTTTGATGGCGGCGGAGGTCGTGTACTTGGGATTGGGCATAAACAGGCCCACTGGGAA | 4 |
| wt-123960-124080 | 1029 | ATAGTAGCTGTACTGCATTCTTCTGTTGAGGGGGTATGGGGACTGAGTGTCATTGTACATCTTTTGCAGGCTTTCCACGGCCACCGCGTGGTTGCCCAGCTTGATGACGGCGGCTGAGAT | 4 |
| wt-124080-124200 | 1030 | CGGCACCCGGGGCTGATCCTCGACCCCTGCGGCCACAGCCGGCAGGTCAGACTTGGTGCTTCCGGCTTTTTCCGGTGAGTCCACGATCCTAGCCATGAAATGCTCAAACGTACGCATCAC | 4 |
| wt-124200-124320 | 1031 | GCGCCCGTAGCTCACGGCAGTGACCAGGTTCTCCCCCCGTACCACAAAAGAAGCATAGCTCGAGGGCCCCATAATCTGGTTGTCGGCCTCCTCACCCAGGAAGGTCAAGAGCTGGCGCAG | 4 |
| wt-124320-124440 | 1032 | AACGTTGTCGGTGACAATAAACACCCCCCCCACTGGCTCTCCCCCCTTGGCGGTCGTGTAGGTACTGACCCCCTTGAGCACGCTCTCCCCGGACACGGCCGCTACCATCTCAGAGAGACG | 4 |
| wt-124440-124560 | 1033 | GCTTCGCACGTACTGAGAAAACCCGGAGCCCATGTTCTCGGCCCGGTCCAGGAAGAAGGAGTGCTCCAGCAGATGCCTCTTGAACATGGCAATGAGGTCAGACTTGACAGTCTTGGAGAA | 4 |
| wt-124560-124680 | 1034 | CCCCCTCTCAGTGAAGGTGGGATCCGCCAGGGTCTGCAGGATAAACATGGGAGGGGCATGGCGAAGCTTCACACTCAGGACGGTGTTAATGAGGCCCCTCTCCAGGGCATCGACCCCAAA | 4 |
| wt-124680-124800 | 1035 | CTGTAGGGCCGAGGCCACGGTCTTGACAGCCCCCACGTACTCTGCGTACTCGACCGGGGTCTCGGGGATACTATGCAGGATCTCCAGATCCAGCATGGACAGTTCCATTTCCGTACTAAT | 4 |
| wt-124800-124920 | 1036 | GTGGTGTTTGTGGCAATTTTTGACCACAATGAATGTCCGCTGCTTGCTGGGTCTCCTTCCGTCCCCGTGAGCAATGGTGGGGACGGAGATTCGAAATTGAATCTTGCCATCCGTCATACG | 4 |
| wt-124920-125040 | 1037 | ACTCAGGTCTTTGAATTCCGTGTTCACACAGGACACGGCCAGTGCCGTCTCCAGGAAGCGAACATATTGGATGGCGTTCGTGTAGACCCCGAGTAGCACCTCAAACTTGATGCCCGCCTC | 4 |
| wt-125040-125160 | 1038 | TCTGGCATCCTTGCCCACCAGCAGGTCAAAGCTATGAAACAACCCCTCAGCCGCTGACTGCCGCAGGTTCGAGAGCAGGTCGGCATCCACCGTCAGATAGGGGAAGGGTCTGTTTTCCAC | 4 |
| wt-125160-125280 | 1039 | ACCCTCATTTGAGGCCATGACACAAGGTAAGAGGGAGATGGGGGGAGGTCTCGAGGGCTTCTCTTCACAGCTGGGTCTCTTTTACGCCCTGGCCTGCAACCGCAGCCCACCCACACTTCC | 4 |
| wt-125280-125400 | 1040 | CGAGGATGCTACCCTTCTAATCAAATGGTTGGACACGGCCCTGGGCAGGGAGGCCACCTTTTACGCGTGTCGGGCTATGCGTCGGCTTCTACTCGGCGTTATCCGAATGAATGACTGCCA | 4 |
| wt-125400-125520 | 1041 | GGAGCTGCCACCCGGTTTAATAATTCTGAGTCCGGGCACCGTCCCTGGCCCCCTTGGAGTCCAGAGTCTGGAGCATACAGACTGCGAAATATGGTCCTCTGCCCACCCTGACCACGCTGC | 4 |
| wt-125520-125640 | 1042 | CCACCTCCCGGTGCCCAGGGTCATCACATACACCGACTGCCCGGGTTCCATAAACACGAGCTCAATGTTTCGCCTTATCATCCGCTACTTGTCTCATCACCAATTTGAGCGCTGCTTCGA | 4 |
| wt-125640-125760 | 1043 | GCAGTTCTGCCGCGTGGTCCCGCGTCGCTTCCTAGGGACCTGTAAGCGAAACTCTGCAAAGATGCTGGCTCATCTGAATCAGGTTACCAGGATCCCCCCCTGTCCGCCCTTCAGCGGGCG | 4 |
| wt-125760-125880 | 1044 | GGAGGCCAGACTCAAGTTCCACTTCTTCTCCTGGAGCACATTCATGCTGTCATGGCCAAACAATGCCACACTCCGGGAGATCAGGACGAGGGCCGCCACCAACCTCACCCACCACCCACA | 4 |
| wt-125880-126000 | 1045 | TCTAGTGGATACTCTGTACCACGCCTCTCCGCAGACCCCATTTCTGACACGCAGCGGTGCTCTATACCGCTTCGTCACCTGTTGCAACTGCACCCTGCCCAATATCTCCATCCAGCAGTG | 4 |
| wt-126000-126120 | 1046 | CAAGGCCGGGGACAGACCGGGGGACCTGGAGATCATTCTACAGAGTAACGGCGGAGGGAGGCCCGCGAGCTTCCAGTTCCCCTCCTCCCCAACTGGCTCCCTATTGCGATGCATAGTTGC | 4 |

| | | | |
|---|---|---|---|
| wt-126120-126240 | 1047 | TGCGTCCCTGCTGCCGGAGGTGTCCGTGGGGCACCAGGAGCTGTCTCCGCTGCGGTCCAGAAGCCAGGGAGGGCAGACGGATGTCAGGTCGGGCCCGGACCCGGCCCGGAGACTGGTGGC | 8 |
| wt-126240-126360 | 1048 | CCTCCTGCGAAGGGAAGATGGGGCACCTAAAGACCCCCCTCTGGGACCGTTTGGACACCCCCGGGGGCCCGGCCCGGCCAAGAGCGAAGACGAGGAGTCTGAGCGTCGAGACGCCCCTCC | 8 |
| wt-126360-126480 | 1049 | ACCCCCGCTCGATTCCAGCTTCCAAGCTTCCCGGTTGGTGCCCGTGGGGCCTGGGTTTCGCCTGCTCGTGTTCAACACCAATCGGGTGATCAACACTAAATTGGTGTGCTCAGAGCCCCT | 4 |
| wt-126480-126600 | 1050 | GGTGAAGATGCGAGTTTGCAATGTCCCCCGCCTCATCAACAACTTTGTAGCCCGCAAGTACGTGGTGAAAGAGACGGCGTTCACCGTCAGTCTATTCTTTACGGACGGGGTGGGGGCCAA | 4 |
| wt-126600-126720 | 1051 | CCTAGCCATCAATGTCAATATCAGTGGCACCTATCTGAGCTTCCTATTGGCCATGACGTCACTGCGGTGCTTCCTGCCTGTGGAGGCTATTTATCCCGCGGCCGTGTCAAACTGGAACTC | 4 |
| wt-126720-126840 | 1052 | GACTCTAGATCTCCATGGGCTGGAAAATCAGAGCCTAGTCAGAGAGAACCGAAGCGGGGTCTTTTGGACTACCAACTTTCCCTCGGTGGTGTCCTGCCGGGACGGTCTCAACGTGTCCTG | 4 |
| wt-126840-126960 | 1053 | GTTTAAGGCCGCAACTGCCACCATATCTCGAGTGCACGGGCAGACATTGGAGCAGCACCTGATCCGTGAAATCACCCCCATCGTGACGCATCGAGAGGCAAAAATCTCCCGGATTAAAAA | 4 |
| wt-126960-127080 | 1054 | CCGGCTCTTTACCCTGCTAGAGCTACGCAATCGGAGTCAGATTCAAGTGCTGCACAAGCGTTTCCTGGAAGGCCTGCTAGACTGCGCCTCCCTCCTGCGCCTGGATCCCAGCTGTATCAA | 4 |
| wt-127080-127200 | 1055 | CCGAATCGCCTCCGAGGGCCTGTTTGATTTCTCCAAGAGAAGCATCGCCCACTCCAAAAACCGACACGAGTGCGCGCTTCTGGGTCACAGACATTCGGCGAACGTGACAAAGCTGGTGGT | 4 |
| wt-127200-127320 | 1056 | AAACGAGCGCAAGACCCGCCTGGACATACTGGGCCGTAACGCTAACTTTTTAACGAGGTGTAAGCATCAGGTTAATCTAAGACAGTCACCTATTTTCCTGACCCTCCTGAGGCACATCCG | 4 |
| wt-127320-127440 | 1057 | CCGACGTCTGGGCCTGGGCCGTGCTTCCGTAAAACGAGAGATTACCCTTCTCCTGGCCCACCTGCGCAAAAAGACAGCCCCCATCCACTGCCGTGATGCTCAAGTGTAAGCAGCCCGGGG | 4 |
| wt-127440-127560 | 1058 | CCCGCTTCATTCACGGGGCCGTGCACCTGCCATCGGGACAGATTGTCTTCCACACCATCCACAGCCCCACTCTTGCCTCGGCGCTGGGACTGCCTGGGGAAAATGTACCCATCCCGGCCC | 4 |
| wt-127560-127680 | 1059 | TCTTCCGTGCCTCGGGCCTCAACGTCCGTGAGAGCCTACCCATGACCAACATGAGGGCACCGATCATCTCGCTGGCTCGCCTCATCCTGGCCCCCAACCCCTATATCCTAGAGGGACAGC | 4 |
| wt-127680-127800 | 1060 | TGACGGTGGGCATGACACAGGACAACGGCATTCCCGTGCTTTTTGCCAGGCCTGTCATTGAGGTAAAAAGCGGGCCTGAGTCCAACATTAAAGCCTCCTCGCAACTTATGATAGCAGAAG | 4 |
| wt-127800-127920 | 1061 | ACTCCTGCCTGAATCAGATCGCCCCCTTTTCCGCATCAGAGCACCCCGCCTTCTCCATGGTTGAGTCCGTAAAACGAGTCCGGGTCGATGAGGGAGCAAACACCCGGCGCACCATCCGGG | 4 |
| wt-127920-128040 | 1062 | ATATTCTGGAGATCCCCGTGACTGTGCTCTCATCCCTGCAACTGTCTCCCACCAAGTCCATCCTGAAAAAGGCACCGGAGCCCCCACCTCCGGAGCCCCAAGCCACCTTCGATGCCACCC | 4 |
| wt-128040-128160 | 1063 | CCTATGCCCGCATCTTTTACGACATCGGGCGACAGGTGCCCAAGCTGGGCAATGCCCCCGCCGCGCAGGTCAGCAACGTGCTCATCGCCAACCGCTCCCACAACTCTCTAAGGCTGGTGC | 4 |
| wt-128160-128280 | 1064 | CCAATCCGGACTTGCTGCCTCTCCAGCATTTGTACCTCAAGCACGTAGTGCTAAAGAGTCTGAATCTGGAGAATATAGTGCAGGACTTTGAGGCCATCTTCACCTCCCCGTCTGATACCA | 4 |
| wt-128280-128400 | 1065 | TCAGTGAGGCTGAAACCAAGGCCTTTGAGAAGCTGGTGGAGCAAGCCAAAAACACCGTAGAGAACATAGTCTTTTGCCTCAACAGCATCTGTTCCACCTCTACACTCCCAGATGTCGTCC | 4 |
| wt-128400-128520 | 1066 | CCGATGTCAATAACCCAAACATTAGCCTGGCTCTAGAGAAGTATTTTCTCATGTTCCCTCCCTCAGGCACCATTATGAGAAATGTCAGATTCGCCACCCCCATCGTCCGGCTCTTGTGCC | 4 |
| wt-128520-128640 | 1067 | AAGGGGCTGAGCTTGGCACCATGGCACAGTTTCTAGGAAAGTACATCAAGGTCAAGAAGGAAACTGGAATGTACACACTGGTCAAGCTTTATTACCTGCTGCGCATCTAAAGGAAAAACA | 4 |
| wt-128640-128760 | 1068 | TAACAATCTTGTGAACCAGAAAGATACCCAGAGCAAAAGCAATAAAGTACAGGATTATTGCCAAAACAACGTGTGCTCTTTCTTCATACAGGCCCGCAATTTCCATGACAGTCCCGTTGG | 4 |
| wt-128760-128880 | 1069 | TGGTCAGCAGCAGATAGTGAACGTGGAGGTTGTCAAAATCAAAGTAGTTGGAGCTCAAGATGGAGTTTTGGACTTCCTGGGAGGTGATGTAGGTTGTAGTTTCCAGGCCTTCCTTTTCAT | 4 |
| wt-128880-129000 | 1070 | CATAACTGAGCAGGGCAAAGCCACAAAAAATGCAGGATTTCTGCGTCCTGGTAAAATTCTGGATCTTTGGAATCTGGCGGGGCTCCCCAGCCACAGCACCCTGCGAACATTTATTCATTA | 4 |
| wt-129000-129120 | 1071 | TAACGGGGGAGAGAAAGAGAGAGCTGCTGAGATAGGTGGTGCTGGCCTCGTATAGCGCCGAGCCTCGGACCTCACGGTCACTAGAGATTATGAATGTCACATTGATGAGCGGGATAATCA | 4 |
| wt-129120-129240 | 1072 | TCAGAACTTTGTCGAGCCTGTCCACGCATTTGTAGGCGGGGAGATGCCACGCATCCCTGTCTTCTCGCTCCAAAGAGAGCCGCCCAAGAAACCCATCCACAGCATTTGAAACGGCCGCCT | 4 |
| wt-129240-129360 | 1073 | GGTCCAGCGTTGCCTCCTGGGGGGCCATGCTCAGCAGCTTGTCTCGTGTGAGGTCAAATCGTAGGCTGAGGTAGCACGGTGAGAAGAGCCCGCTCTCTGTCCCCAGGGCTAGCCCCCGCA | 4 |
| wt-129360-129480 | 1074 | AAACCTCCCCAATCTCTAGGGCCGAGCACAGGGCGGTGGACAGCAGTTGGTATAGGGCAAGGTTGGGCCCCTGGGTAGTCACGTTCAGCCGCAACTCGCGTAGCACCACGTGGCTGCCGA | 4 |
| wt-129480-129600 | 1075 | TAAACAGGGTCTCTCTCATCACGGTATGCAGGGGCTGGAAAAGGGGGTGGCGGTTGTAGGCCGAGAGAAGCACAGATGTGGCGCCTCCAATGAGGCCACTGTAAACCCCGGCCTTGGGGT | 4 |
| wt-129600-129720 | 1076 | AGCCGACGGTGGCTAACCTCAGCGCGTACTCCTGTTTCTCAGTAGTCAGGTGACCCAGCTCCTCCATCTTGACCGTGGCCATCAGCATGGCGGCCAAGCGCTCCAGCCCGTAGGATTGCA | 4 |
| wt-129720-129840 | 1077 | TGCCCTTGACAGTGGCCCCATAACATATGCCGATGATGTCTTTCAGGACAGTCAGCTCAAAGAAGCTCTTGGCCAACCAGCGGAGGTCCACGCAGCCATTGCCAGTCTCACCCACAGCAT | 4 |
| wt-129840-129960 | 1078 | GACCCACCTTAAAGAAGGCCACAGAAACCTCAAACATGGTAGTCAGCGTTTCCGTGTCCAGTTCCGGCTCCCGGCAGCCTCCCTTCATCTCCAGCAGGACCAGTTTCTGGAGAACGTAGC | 4 |
| wt-129960-130080 | 1079 | GAGCGTAGCTGGCGGCTGTCATGGTGACGGCTCGGGAAAACATATCCTTCAGGTTGGGTACAAAGTAGTTGTGAAAGTTGGCATAATGCACAAAGGTTGTAACAATCACCAGGGAATAGT | 4 |
| wt-130080-130200 | 1080 | CCCCGCTTTGGGCACTGGTTAAGGATGGGTAACTAAAAGGCCCCCTCAGATCCGGCAGGTCCTTCGTCTTGCCAAAGACCAGGCTCAACACATGCTCATCTCCCTTCTCGGTCACTCGCT | 4 |
| wt-130200-130320 | 1081 | TGTAGGTGCCCATCAGAAATTTAGAAGTCATGGCCCCCGTGTACTGAAACTTGTCCCCGTTGATGGACAGGGCCACATAAGACAAGTGACAGCGCAGCTGATAAAAGACATAGCTGTGTG | 4 |
| wt-130320-130440 | 1082 | GCCGCGTGTTGGGCAGCATGGTGCCAATATAGTAGAAGAGCTGCTTCTCAAGGGGGGCACTAAGCATGCAGGCAGGGGAATTCAGGCCGCTAATGACTCCGGGATGGACCTTAGATGCAT | 4 |
| wt-130440-130560 | 1083 | CCACTTGCATGGATCCTTCAGAGACAGCAGGGATATCGACAGGCTCGGCCAGCGCAATACCAAGGGTACCAGACGTCTTGTAAATTAACTTGTAGCGGTTAAGCATAGACGCCAAATCTT | 4 |
| wt-130560-130680 | 1084 | CGGTGACATTTGCCTCTCTCCACAGCGCCTCTGGGCTAAGGCCTGGGACCTTTGCCATCAGTTCGGTCCATGGGATGGTGTAATGCGAAGCATGCCCCTCTATGTCCAGGTGCAGCTTAA | 4 |
| wt-130680-130800 | 1085 | CCTCGCTGAGGCTGGCAGCCCCCACCTCCCATAGCAACACCAGGCAAAAAACACAGAGCAACTGCATCCTAGTCCCGATTTCCCCTCTCAAAATCAGAGATCACCTTGCTCAGACCAGCC | 4 |
| wt-130800-130920 | 1086 | CAATCGAAAAACTGAGATCGTATTGCCGGATTCTTCAATGCCTGCATGTAAATCTCCGTCCAGCATCCAGGTAAATCGTCCTGAAACTCTGAGAGGTCCACAAGCACAAACTGAAGGTAG | 4 |
| wt-130920-131040 | 1087 | GCTAGCGTTCGGGTGAACGCAAGACAAACTTCCAACAACACCGCGTCGGCTCGGAAAGGCTGTATGACTTCCTTAAGTACACTAAAGATGCTGTTCTTATACAGCTTCTCGGCCACACCA | 4 |
| wt-131040-131160 | 1088 | CTTCGAATTATGGGGGTGTGGCTTTGATGACATACTGTCGTGATTGTTGTTAGACCGGCACATACCTTCACAATGTCCTCGGGGGCAAAATACTGTGTTAGGAGCCAGGCACAGTAAACG | 4 |
| wt-131160-131280 | 1089 | GCGTGATATGCATCGTTGACACTCTTCAGGTAGCCAGCATCCAGTCCTGACTCATGTTTCCTCCCTCGCTTCTTCAGGCGGCGCATGTTCTCCTCCACGTTTAACTTCATCCAGACTATG | 4 |
| wt-131280-131400 | 1090 | GTGTCCCCCGGGTCTGCGGTAAACGTGGCCAAAACTTGAATAAAGTCACTATAGGAGAGAAGCTGGCTCCGGAGCAGCATTAGAGGGAAAACCACGGAGGCCGACAGCAAATGGCGATCA | 4 |
| wt-131400-131520 | 1091 | TGCAAAATCCAACAATCCAGGGGCGCGACTGACCTGGCACCAGACTCGGTAACCAGCAAGCTCCGCTTCCTAGAGGCCAAGACTCTGAAAGGGGTGGTAAATTTCATCTGGCATGCTAAA | 4 |
| wt-131520-131640 | 1092 | ACCTCAGCCGACGTGTCTTCCCTTCCATGCCTCGCCCGAGTCACATTCTTGTGCATGGCCTTAATGGCATTTTCATACACATGAGTCCAGTACCGCATCGGTTCAGGGACTACAATGGTC | 4 |
| wt-131640-131760 | 1093 | AGGTCCCCAAAGACAGCCTTCAAATGATTCAGCATAGTAGTCTTTCCCACACCAGGGGCACCTTCCAAAAATAGGGAACAGGCAGGTTTGATTACTGGTACATGATTTGTTAGGTGGGTC | 4 |
| wt-131760-131880 | 1094 | ACAATTGGAACCCGCATGCTCTCCTTCCTCTGAGCCTTGGCCTGGCGGGTGTCTTGGGCATCATCCAGATTCAGAACATTCATCACACTCCCACTTAGCCGCTTCAGCTGGGCAGCATGC | 4 |
| wt-131880-132000 | 1095 | TTGGATAACTTACTAAACTCGCGCCCATGGGCGGCCAGGTGTTCGAAGAGACCAGAAGGCTTACCCTTGCCACCATTCTTTTGTTTTAACGCGGAATGAGAAGAGGGCCTGCGGAAATTA | 4 |
| wt-132000-132120 | 1096 | GACTCATCCTCAGACTCACAGTCAGATTTGTCATCGAGCCCAAGGCCGGCCAGGCCCTCCTCAAAGCCTTTCTGGTACATGAAGCTCCGGCTCGTGGAGTCCGCACCTCCTTCTGTGCAC | 4 |
| wt-132120-132240 | 1097 | GAAGTTTTGCGGAACCAGGAGAAGGGGTCTGGCGTCTTGCTGGGGCCACACTCCCGGCTACGGGGCTTCGGGGTAGGGGCAGTAGGCTTTTGGTGTGCGGGTGCTGGTGGCTGGGCTCCC | 8 |

| | | | |
|---|---|---|---|
| wt-132240-132360 | 1098 | CTGGGCAGGGTAAAGGGGCACGATGTGTGCCGGCTACCCGGAGAGTTTCCAGTATTAGATGTCACGGCAGCCTGGGTCCGGCACGGCACCCTCTCCCCAGACAGTCCGGTCGGAGCCATC | 4 |
| wt-132360-132480 | 1099 | AAGGGGGGCCAGTGGGTGGGCACCTGGTAGAGGCCGTCGTCATCTTCCTCACCTGCCCCTGAGTCACTACCGGTTGGGGTAAGAACTGAGGGGGCAAAGTCATCAATCTCAGCGTAAAAG | 4 |
| wt-132480-132600 | 1100 | TTTTCGTGTCTTTCGTTTTCAGGGGACTCATCCTCCTGACATTTTCGCCAGCCGCCGGGCGGGCCGGCCTCCTTTCCTGGAAATCCAGCCATGGATCCCACCCGGGGTCTGTGTGCCCTC | 4 |
| wt-132600-132720 | 1101 | TCCACACACGACCTGGCAAAATTTCACAGTCTTCCCCCGGCTAGAAAGGCGGCAGGTAAGCGAGCGCACCTTCGGTGTTACTCCAAGCTGCTCTCTCTTAAGAGCTGGGAGCAACTGGCC | 4 |
| wt-132720-132840 | 1102 | TCTTTTTTGTCTCTGCCCCCGGGACCCACGTTTACAGACTTTAGACTATTTTTCGAAGTCACCCTGGGTCGGAGAATCGCAGATTGCGTTGTGGTAGCTCTGCAGCCTTATCCCCGGTGT | 4 |
| wt-132840-132960 | 1103 | TATATTGTAGAATTTAAGACGGCCATGAGCAACACGGCCAACCCGCAAAGCGTTACTCGCAAGGCACAGAGGCTAGAGGGCACCGCCCAGTTGTGTGACTGTGCCAATTTTCTTCGCACG | 4 |
| wt-132960-133080 | 1104 | TCCTGCCCCCCCGTGCTGGGCAGTCAGGGCCTGGAAGTCTTGGCGGCGTTGGTATTTAAAAACCAGCGATCCCTGAGAACGCTCCAGGTAGAGTTTCCAGCCCTGGGCCAAAAGACCCTC | 4 |
| wt-133080-133200 | 1105 | CCCACCTCCACCACCGGCCTGCTAAACCTCCTCTCCCGCTGGCAGGATGGCGCTCTCCGGGCACGTCTTGATAGACCCCGCCCGACTGCCCAGGGACACAGGCCCCGAACTCATGTGGGC | 8 |
| wt-133200-133320 | 1106 | CCCAAGCCTTCGCAACTCACTGCGCGTGTCCCCCGAAGCGCTCGAGCTGGCAGAGCGGGAGGCCGAAAGGGCCAGGTCGGAGCGGTGGGACAGGTGTGCCCAGGTGCTCAAAAATAGGCT | 8 |
| wt-133440-133560 | 1108 | CCCCTCCACGGGAGGGCGCTCTGCGCCAGCCCCGCCCTCCCCATCCCCAGCCCAGCCGTTCACTCGGCTCACCGGGAACGCCCAGTATGCAGTCTCAATCTCTCCCACGGACCCCCCTCT | 8 |
| wt-133560-133680 | 1109 | GATGGTGGCCGGCAGCCTGGCTCAAACGCTGCTTGGTAATCTGTACGGGAACATCAACCAGTGGGTACCGTCCTTCGGACCCTGGTACAGGACCATGTCGGCTAATGCCATGCAGCGGCG | 4 |
| wt-133680-133800 | 1110 | CGTGTTCCCTAAGCAGCTGAGGGGCAACCTGAACTTTACCAACTCCGTCTCCCTAAAGCTGATGACAGAAGTGGTGGCGGTGCTTGAGGGCACCACCCAGGACTTTTTCTCAGACGTCAG | 4 |
| wt-133800-133920 | 1111 | GCACCTGCCAGACCTCCAGGCTGCCCTGATCCTCTCGGTGGCCTACCTGCTACTCCAGGGGGGCTCCTCACACCAGCAGCGCCCCCTCCCTGCCTCACGGGAAGAGCTGCTGGAGCTGGG | 8 |
| wt-133920-134040 | 1112 | CCCGGAGAGCCTAGAGAAAATCATCGCCGACCTCAAGGCCAAGTCACCCGGCGGAAATTTTATGATTTTAACAAGCGGAAACAAGGAAGCGCGCCAGTCAATAGCCCCTCTCAACCGACA | 4 |
| wt-134040-134160 | 1113 | GGCGGCATATCCACCCGGCACATTCGCGGACAATAAGATTTACAACCTGTTTGTGGGAGCGGGACTACTGCCCACGACGGCCGCGCTGAACGTGCCCGGGGCGGCGGGTCGGGACCGGGA | 8 |
| wt-134160-134280 | 1114 | CCTGGTGTACCGGATCGCCAACCAGATCTTTGGGGAGGATGTGCCCCCCTTCTCATCTCACCAGTGGAACCTGCGCGTAGGTTTAGCCGCACTCGAGGCCCTGATGCTCGTCTACACGCT | 4 |
| wt-134280-134400 | 1115 | CTGCGAGACCGCCAACCTGGCCGAGGCGGCCACCCGGCGTCTACACCTATCGTCCCTGCTCCCCCAGGCAATGCAGCGGCGCAAGCCTGCCATGGCGTCAGCTGGTATGCCGGGCGCCTA | 8 |
| wt-134400-134520 | 1116 | TCCAGTCCAGACGCTTTTCCGCCACGGGGAGCTCTTCCGCTTCATCTGGGCCCACTACGTGAGGCCCACGGTGGCGGCAGACCCCCAGGCCTCCATCAGCTCTCTTTTCCCCGGGCTGGT | 8 |
| wt-134520-134640 | 1117 | TTTGCTGGCCCTGGAGCTGAAGTTGATGGATGGGCAGGCTCCCTCCCATTATGCCATAAACCTGACCGGACAAAAGTTTGACACCCTCTTTGAGATTATCAACCAGAAGCTTTTATTTCA | 4 |
| wt-134640-134760 | 1118 | CGACCCGGCTGCCATGCTGGCGGCGCGCACACAGCTGCGTCTAGCCTTCGAGGACGGCGTCGGTGTTGCCCTGGGGCGCCCCTCGCCCATGCTTGCGGCGCGGGAGATCCTGGAGCGTCA | 8 |
| wt-134760-134880 | 1119 | GTTCTCAGCCTCGGATGACTACGACCGGCTGTACTTCCTGACGCTGGGCTACCTGGCCTCCCCGGTGGCCCCAAGCTGAGCCAGTTCCTCGCACTGGAGTGGGTCATTGGCAAAAAGGTA | 4 |
| wt-134880-135000 | 1120 | AATAAACTCATCGCACGGGGGTTTTGCCTCCTTCTCGTCTCTTGTTTCGGGTAGGGGAGTAAGGCCGTGCCAGGCCGCCATGCTCAGGGCCACGGCGTGCCAGAGGCCCTCGTAGTCGTG | 4 |
| wt-135120-135240 | 1122 | AAGGCAGTAGAGGACACCCACCACCAGGCGGTTAGGGAGGACACTGCCAAGGTTGAAGAGCAGATTTCCGTCAGCCAGGGTGACCTGGCTCAGGTCCGGCGCCCTGCGCAGTCCAAGCTG | 4 |
| wt-135240-135360 | 1123 | CGCCCACACACATGCACAGACGGCCCCTGTGACATCAGGCCGGTCATGCAAAAACAGACAAAGAGACCGTGAGCGGTTACCGGGGCGCAGGGCCTCTGCCGGGAAGCCCACCCGGGCCAG | 8 |
| wt-135360-135480 | 1124 | GGCCCGGTAAAGCAGGTACCAGTATTCATCCGGCACCTTGCGTGCCAACACACGATTCGTGCGGTTTCCAGTATTTATCACGGCTTCCCGCCACAGGTAAAAGTTAACACTTAGGGTCAG | 4 |
| wt-135480-135600 | 1125 | CAGCTTGGTCAGGGATAGGTGCAAAAACCTGAGCTCGTCCTCGCGCAGAGCGCAAAGCGGCCAGTTCTTTAGCATCTTCAGGAGGAGCCCGTGAATCCCAGGTGTCATTCGCGCGTCATC | 4 |
| wt-135600-135720 | 1126 | CCCGCGCACCCCCAGTCCCATTAACATAGCGGGCACAATGGTGCAGGCACCGTCTGTATACGTCTGCGGCTTCGTGGAGCGCCCGGACGCCCCACCCAAGGACGCCTGCCTTCACCTGGA | 8 |
| wt-135720-135840 | 1127 | TCCCCTCACCGTCAAGAGCCAGCTCCCTCTGAAGAAGCCCTTGCCACTCACGGTGGAACACCTGCCGGATGCTCCGGTCGGCTCAGTCTTTGGCCTTTACCAGAGCCGAGCGGGTCTCTT | 4 |
| wt-135840-135960 | 1128 | TAGCGCAGCCTCGATTACCTCTGGGGACTTCCTGTCCCTGCTGGACTCAATTTACCACGATTGCGATATTGCACAGAGTCAGCGCCTGCCCCTCCCTCGAGAACCCAAGGTGGAGGCTCT | 4 |
| wt-135960-136080 | 1129 | GCACGCCTGGCTCCCCTCACTGTCACTGGCCTCCCTCCACCCAGACATACCCCAAACCACCGCAGATGGAGGCAAGCTGTCCTTCTTTGACCACGTGTCTATCTGTGCCCTGGGTCGTCG | 4 |
| wt-136080-136200 | 1130 | GCGCGGCACCACGGCAGTCTACGGTACAGACCTTGCGTGGGTCCTGAAGCACTTTAGTGACCTGGAACCGTCTATCGCCGCCCAGATTGAGAATGACGCCAATGCCGCAAAGCGTGAATC | 4 |
| wt-136200-136320 | 1131 | CGGATGCCCGGAAGACCACCCTCTGCCCCTCACGAAGCTCATAGCTAAGGCAATCGATGCTGGATTTCTGAGAAACCGCGTGGAGACTCTGAGGCAGGACAGGGGTGTGGCCAATATCCC | 4 |
| wt-136320-136440 | 1132 | AGCCGAGTCGTATTTAAAGGCCAGCGACGCCCCGGACCTACAAAAGCCGGACAAGGCACTTCAGAGCCCACCACCGGCCTCCACAGACCCAGCCACCATGCTATCAGGTAACGCAGGAGA | 4 |
| wt-136440-136560 | 1133 | AGGAGCAACAGCCTGCGGAGGTTCGGCCGCCGCGGGCCAGGACCTCATCAGCGTCCCCCGCAACACCTTTATGACACTGCTTCAGACCAACCTGGACAACAAACCGCCGAGGCAGACCCC | 4 |
| wt-136560-136680 | 1134 | GCTACCCTACGCGGCCCCGCTGCCCCCCTTTTCCCACCAGGCAATAGCCACCGCGCCTTCCTACGGTCCTGGGGCCGGAGCGGTCGCCCCGGCCGGCGGCTACTTTACCTCCCCAGGAGG | 8 |
| wt-136680-136800 | 1135 | TTACTACGCCGGGCCCGCGGGCGGGGACCCGGGTGCCTTCTTGGCGATGGACGCTCACACCTACCACCCCCACCCACACCCCCCTCCGGCCTACTTTGGCTTGCCGGGCCTCTTTGGCCC | 8 |
| wt-136800-136920 | 1136 | CCCTCCACCCGTGCCTCCTTACTACGGATCCCACTTGCGGGCAGACTACGTCCCCGCTCCCTCGCGATCCAACAAGCGGAAAAGAGACCCCGAGGAGGATGAAGAAGGCGGGGGGCTATT | 4 |
| wt-136920-137040 | 1137 | CCCGGGGGAGGACGCCACCCTCTACCGCAAGGACATAGCGGGCCTCTCCAAGAGTGTGAATGAGTTACAGCACACGCTACAGGCCCTGCGCCGGGAGACGCTGTCCTACGGCCACACCGG | 8 |
| wt-137040-137160 | 1138 | AGTCGGATACTGCCCCCAGCAGGGCCCCTGCTACACCCACTCGGGGCCTTACGGATTTCAGCCTCATCAAAGCTACGAAGTGCCCAGATACGTCCCTCATCCGCCCCCACCACCAACTTC | 4 |
| wt-137160-137280 | 1139 | TCACCAGGCAGCTCAGGCGCAGCCTCCACCCCCGGGCACACAGGCCCCCGAAGCCCACTGTGTGGCCGAGTCCACGATCCCTGAGGCGGGAGCAGCCGGGAACTCTGGACCCCGGGAGGA | 8 |
| wt-137400-137520 | 1141 | AGCCCACCTCAAGTCCATCTTTTGCGAGGAATTGCTGAATAAACGCGTGGCTTGAAAGTAAACTTTATTGCGTGTTAGTACCTGTCCATTCACAGGGGTATCCAGCCCTTGCGCCGCCTC | 4 |
| wt-137520-137640 | 1142 | CCCCAGCCCGCCAGCCACCCCAGACAGGAGATGATAATGATGAGGAGCACCGGAGCCACCACAGCACAAGTGATTAGGAGCAGGGCCCAGTGCACCCAGGTGGTCTTAGGGCGCCAGGGA | 4 |
| wt-137640-137760 | 1143 | TCGATTGGAAAAGGGCCCAGGGTCACTGGCTTATGCGTGGGACGTTTAGAAACAGGCCGCCTATGGGGCCTGTGACTGGTGCTTGTGGTGTGGGAGACTAATGTGGTGGGGGCTATGGTA | 4 |
| wt-137760-137880 | 1144 | GTGGCTGGGATAACAGTAAGATGCATACGCTGAGTGAGGGTCCGGTTGGCATGGTATTGGTCGTCTTCTTCCCCTGCAGAGTAATTGCAGTGGACCCCGGAGGCCACACTGCAATTTCTC | 4 |
| wt-137880-138000 | 1145 | AGTGTCACATTGCACGTGTAGTAACCTGCATGCGCAAGGGTCACATTGGGGATTATCAGAGAGACGGAGGTGTTGGAGTCATTTACCCATTCTAGGGTAAGGCTATAATTGTAACCCCCG | 4 |
| wt-138000-138120 | 1146 | TTAGTTATATGAGTTCCGTTGTTGGAAGTAGCTACGGCCAAGGGCAGTTGTCCATCCCCGGGAGTGTATCCCCGGCCCAACTCGATCCGAGAGACCGACTCATTGCTAGGAACGCTGCAG | 4 |
| wt-138120-138240 | 1147 | GTGAGATTCACTCTAGCACCTGCATGGGCGGTGACATTTTCAAATTTAACCAGATCTGAGAAAAATGCACAAACAGACCCCACACAGCAGCACAATAGAAGCACTAAATGAGTCATTCCT | 4 |
| wt-138240-138360 | 1148 | AAACTGTCAGTTTTAAAACTCCCTGCTTCTCAGGCCTAAATATGTGGTGGGGTGTGCTTAGGATCACTTTCATATTCTGCAACAACAGCCATACCCGGAAGAGGAGCTGCCGGTTGCCAT | 4 |
| wt-138360-138480 | 1149 | TTTTCAAGCTGCTAAACCACGAGTGGCAGCAGGCCTAAGAAGCTCCTCAGCAACATGGAGACCTCGAAGGGAAACTGGCAGGAGCAGGGAGTCACGTAGGCACTAGCCTCTTCATGTGAG | 4 |
| wt-138480-138600 | 1150 | GTAAGAGATCGCTAAAAATGGGATCAGGGTATGTAAACCGAGTTTTGCGGGGGATGGTGAGCCAGACACGGCGGGTGGGGGAAGGAGCTGACACGATTGAGTAGAAAGGGCCAAAAATAC | 4 |
| wt-138600-138720 | 1151 | ACCAGCTATAAGGAATTGCTCAGGCCAAAGTTGTTCCTCAGGTGGCTTTAGGCCTAATGTAGGCAATTGCGTGCCTAGAACATTGCTAATGTGCCCTGGGTTTCCTGCCTTCATGCAAAT | 4 |

| | | | |
|---|---|---|---|
| wt-138720-138840 | 1152 | ATTCTACCTCCCCCGGCCTGGTGCAAAATGTCTGCCTCAGAATACTAACAGCTAATCCAAGCTAACATTCTATCAGTAAACGGGCAGAAAACTGATAAGGACCGCGGAGTTTGGCCCTCC | 4 |
| wt-138840-138960 | 1153 | GCGGTGTCCGGTGGTCCTCACACGTGCCCTCCCCCCGGGCCGATGGCTGAGGCCCGGAATATGCAAGTGCATCTTTCTAACCAGTAGGGGCCTCCACCTAGGTGCTTTGTTAATCTTTAG | 4 |
| wt-138960-139080 | 1154 | TGGGAACTAGTGGGAGTGCTGTGCCTCGGGTACCCCTATCCTATAGGTCCTACCGGAGCTCCTTGTCTTGATAATCCCTGTAAACACACACCACCTAAGAACAAGGCATTGTTAACCTTT | 4 |
| wt-139080-139200 | 1155 | GGTGGAACCTAGTGTTAGTGTTGTGCTGTAAATAAGTGTCCAGCGCACCACTAGTCACCAGGTGTCACCGGAGGCTACTTGCCTCAGTGCCACTTTTACCTTCTCAAATCTATACGGGGG | 4 |
| wt-139200-139320 | 1156 | GGGGGGGGGCTCTGTAACATTTGGTGGGACCTGATGCTGCTGGTGTGCTGTAAATAAGTGCCTAGCACATCACGTAGGCACCAGGTGTCACCAGGGCTACTTGCCTCGGCATCTCCTCAC | 4 |
| wt-139320-139440 | 1157 | CGGAGAAGGGGTTAACAAACCCGTGGGGGGTCTTAGTGGAAGTGACGTGCTGTGAATACAGGTCCATAGCACCGCTATCCACTATGTCTCGCCCGGGCTATATGTCGCCTTACCTCCCCT | 4 |
| wt-139440-139560 | 1158 | ATATAGTCACGACCCCACCGAACCAGGCATGATGTAGAATAAAATTTTATGCATCATCTTCTAATCTGTGCCGCTTGGAGGGAAACATGACCACCTGAAGTCTGTTAACCAGGTCAGTGG | 4 |
| wt-139560-139680 | 1159 | TTTTGTTTCCTTGATAGAGACACAAGGACTGCCAGCCCCATTGGGGAGGGGGGGTGGGTACGGGAGAGTTTGGGCTCGTTTAAACAAAGTCTCATCTGATGCTCTGTGGCACCTCAAGGT | 4 |
| wt-139680-139800 | 1160 | GAATATAGCTGCCCATCGACGTATCGCTGGAAACCGGTGGGCCGCTGTTCACCTAAAGTGACGCAAGGTCTGTCAGCCGCCAGGGTCCGTTTACCAGGCTTTCAGGTGTGGAATTTAGAT | 4 |
| wt-139800-139920 | 1161 | AGAGTGGGTGTGTGCTCTTGTTTAATTACACCAAGATCACCACCCTCTATCCATATCCCACAATTGATAAACCTCCGCATGTCCAACCACCACGTTGAACAGGATGTGGCACCCTAAGAG | 4 |
| wt-139920-140040 | 1162 | GACGCAGGCATACAAGGTTATTACCCAGTCCTTGTATGCCTGGTGTCCCCTTAGTGGGACGCAGGCCTAGGTAGCATCATTTACACTAAAAGCAGTGACCTTGTTGGTACTTTAAGGTTG | 4 |
| wt-140040-140160 | 1163 | GTCCAATCCATAGGCTTTTTTTGTGAAAACCCGGGGATCGGACTAGCCTTAGAGTAACTCAAGGCCAAGCATTTCACACCTGCAAATGCACCATGTAACCACAGATCTAAACTGAAAGTT | 4 |
| wt-140160-140280 | 1164 | GCAGCTTTAGATGGCAAGGAAACTTGGGTTTCAGGCATAGAAAGCCTGGCTCACTATAGCAGCCCATGTTTGTTCCAGGGTGGGGGAAAGGCACGTGCCCTTAGAAAACTTAGCTGCAAA | 4 |
| wt-140280-140400 | 1165 | AATTCTATTGTGTTGGGAGAGCCTCTATATCTAAAGGCCTTTCCTCACAATACAAATGTTACTAACGTCTGCCCTCTGGAGACCTGCTATGTGGCTAGACGTATGGCCTACCCAAGACGT | 4 |
| wt-140400-140520 | 1166 | TGGGGGTCTCGGGTAGGCCATGATTCTTCCAGGCATAGGTTACAACCAGTCACTGCTATCAAGCCTACTCAGTTCCCAACGCAGCACATACCCCCCGCCTCTCCTGCCATGAGGACTTAT | 4 |
| wt-140520-140640 | 1167 | GGCAGTGTTTACTGTTCTGCTTTTACTCTTGGACCAGGCTGTCATTCTATCAGAATAACAGGGGAAGCAAGGCCCCCTGCTTCAGCGGGACACGTGTTTCTAGAATCTCGGAGCCAATAA | 4 |
| wt-140640-140760 | 1168 | CTACCTGCCCCTCTAATCTGTATGCTGCATGAAAAACCACATACACGTGATGTAAGTTTAGCCAGTTTATTGTTACACCAATGCCCCGAAAGTCTCCCCCTGTCCCTTTGGGTCTCAGGA | 4 |
| wt-140760-140880 | 1169 | CCCAGCCCTGGAGCTCGGGGGCGGCCGGGTGGCCCACCGGGTCCGCTGGGTCCGCTGCCCCGCTCCGGCGGGGGGTGGCCGGCTGCAGCCGGGTCCGGGGTTCCGGCCCTGGAGCTCGGG | 12 |
| wt-140880-141000 | 1170 | GGGCGGCCGGGTGGCCCACCGGGTCCGCTGGGTCCGCTGCCCCGCTCCGGCGGGGGGTGGCCGGCTGCAGCCGGGTCCGGGGTTCCGGCCCTGGAGCTCGGGGGGCGGCCGGGTGGCCCA | 12 |
| wt-141000-141120 | 1171 | CCGGGTCCGCTGGGTCCGCTGCCCCGCTCCGGCGGGGGGTGGCCGGCTGCAGCCGGGTCCGGGGTTCCGGCCCTGGAGCTCGGGGGGCGGCCGGGTGGCCCACCGGGTCCGCTGGGTCCG | 12 |
| wt-141120-141240 | 1172 | CTGCCCCGCTCCGGCGGGGGGTGGCCGGCTGCAGCCGGGTCCGGGGTTCCGGCCCTGGAGCTCGGGGGGCGGCCGGGTGGCCCACCGGGTCCGCTGGGTCCGCTGCCCCGCTCCGGCGGG | 12 |
| wt-141240-141360 | 1173 | GGGTGGCCGGCTGCAGCCGGGTCCGGGGTTCCGGCCCTGGAGCTCGGGGGGCGGCCGGGTGGCCCACCGGGTCCGCTGGGTCCGCTGCCCCGCTCCGGCGGGGGGTGGCCGGCTGCAGCC | 12 |
| wt-141360-141480 | 1174 | GGGTCCGGGGTTCCGGCCCTGGAGCTCGGGGGGCGGCCGGGTGGCCCACCGGGTCCGCTGGGTCCGCTGCCCCGCTCCGGCGGGGGGTGGCCGGCTGCAGCCGGGTCCGGGGTTCCGGCC | 12 |
| wt-141480-141600 | 1175 | CTGGAGCTCGGGGGGCGGCCGGGTGGCCCACCGGGTCCGCTGGGTCCGCTGCCCCGCTCCGGCGGGGGGTGGCCGGCTGCAGCCGGGTCCGGGGTTCCGGCCCTGGAGCTCGGGGGGCGG | 12 |
| wt-141600-141720 | 1176 | CCGGGTGGCCCACCGGGTCCGCTGGGTCCGCTGCCCCGCTCCGGCGGGGGGTGGCCGGCTGCAGCCGGGTCCGGGGTTCCGGCCCTGGAGCTCGGGGGGCGGCCGGGTGGCCCACCGGGT | 12 |
| wt-141720-141840 | 1177 | CCGCTGGGTCCGCTGCCCCGCTCCGGCGGGGGGTGGCCGGCTGCAGCCGGGTCCGGGGTTCCGGCCCTGGAGCTCGGGGGGCGGCCGGGTGGCCCACCGGGTCCGCTGGGTCCGCTGCCC | 12 |
| wt-141840-141960 | 1178 | CGCTCCGGCGGGGGGTGGCCGGCTGCAGCCGGGTCCGGGGTTCCGGCCCTGGAGCTCGGGGGGCGGCCGGGTGGCCCACCGGGTCCGCTGGGTCCGCTGCCCCGCTCCGGCGGGGGGTGG | 12 |
| wt-141960-142080 | 1179 | CCGGCTGCAGCCGGGTCCGGGGTTCCGGCCCTGGAGCTCGGGGGGCGGCCGGGTGGCCCACCGGGTCCGCTGGGTCCGCTGCCCCGCTCCGGCGGGGGGTGGCCGGCTGCAGCCGGGTCC | 12 |
| wt-142080-142200 | 1180 | GGGGTTCCGGCCCTGGAGCTCGGGGGGCGGCCGGGTGGCCCACCGGGTCCGCTGGGTCCGCTGCCCCGCTCCGGCGGGGGGTGGCCGGCTGCAGCCGGGTCCGGGGTTCCGGCCCTGGAG | 12 |
| wt-142200-142320 | 1181 | CTCGGGGGGCGGCCGGGTGGCCCACCGGGTCCGCTGGGTCCGCTGCCCCGCTCCGGCGGGGGGTGGCCGGCTGCAGCCGGGTCCGGGGTTCCGGCCCTGGAGCTCGGGGGGCGGCCGGGT | 12 |
| wt-142320-142440 | 1182 | GGCCCACCGGGTCCGCTGGGTCCGCTGCCCCGCTCCGGCGGGGGGTGGCCGGCTGCAGCCGGGTCCGGGGTTCCGGCCCTGGAGCTCGGGGGGCGGCCGGGTGGCCCACCGGGTCCGCTG | 12 |
| wt-142440-142560 | 1183 | GGTCCGCTGCCCCGCTCCGGCGGGGGGTGGCCGGCTGCAGCCGGGTCCGGGGTTCCGGCCCTGGAGCTCGGGGGGCGGCCGGGTGGCCCACCGGGTCCGCTGGGTCCGCTGCCCCGCTCC | 12 |
| wt-142560-142680 | 1184 | GGCGGGGGGTGGCCGGCTGCAGCCGGGTCCGGGGTTCCGGCCCTGGAGCTCGGGGGGCGGCCGGGTGGCCCACCGGGTCCGCTGGGTCCGCTGCCCCGCTCCGGCGGGGGGTGGCCGGCT | 12 |
| wt-142680-142800 | 1185 | GCAGCCGGGTCCGGGGTTCCGGCCCTGGAGCTCGGGGGGCGGCCGGGTGGCCCACCGGGTCCGCTGGGTCCGCTGCCCCGCTCCGGCGGGGGGTGGCCGGCTGCAGCCGGGTCCGGGGTT | 12 |
| wt-142800-142920 | 1186 | CCGGCCCTGGAGCTCGGGGGGCGGCCGGGTGGCCCACCGGGTCCGCTGGGTCCGCTGCCCCGCTCCGGCGGGGGGTGGCCGGCTGCAGCCGGGTCCGGGGTTCCGGCCCTGGAGCTCGGG | 12 |
| wt-142920-143040 | 1187 | GGGCGGCCGGGTGGCCCACCGGGTCCGCTGGGTCCGCTGCCCCGCTCCGGCGGGGGGTGGCCGGCTGCAGCCGGGTCCGGGGTTCCGGCCCTGGAGCTCGGGGGGCGGCCGGGTGGCCCA | 12 |
| wt-143040-143160 | 1188 | CCGGGTCCGCTGGGTCCGCTGCCCCGCTCCGGCGGGGGGTGGCCGGCTGCAGCCGGGTCCGGGGTTCCGGCCCTGGAGCTCGGGGGGCGGCCGGGTGGCCCACCGGGTCCGCTGGGTCCG | 12 |
| wt-143160-143280 | 1189 | CTGCCCCGCTCCGGCGGGGGGTGGCCGGCTGCAGCCGGGTCCGGGGTTCCGGCCCTGGAGCTCGGGGGGCGGCCGGGTGGCCCACCGGGTCCGCTGGGTCCGCTGCCCCGCTCCGGCGGG | 12 |
| wt-143280-143400 | 1190 | GATGGGGGTGCGCTCCCAGGCCGGACCCTGGTGCCAGGCAGGGACCCCGCGCCACCCGCTTCATGGGGGGGGAGGCCGCCGCAAGGACGCCGGGCCGGCTGGGAGGTGTGCACCCCCCGA | 12 |
| wt-143400-143520 | 1191 | GCGTCTGGACGACGCTGGCGAGCCGGGCCAGCTCGCCTTCTTTTATCCTCTTTTTGGGGTCTCTGTGCAATACCTTAAGGTTTGCTCAGGAGTGGGGGGCTTCTCATTGGTTAATTCAGG | 4 |
| wt-143520-143640 | 1192 | TGTGTGATTTTAGCCCGTTGGGTTACATTAAGGTGTGTAACCAGGTGGGTGGTACCTGGAGGTCATTCTATTGGGATAACGAGAGGAGGAGGGGCTAGAGGCCCGCGAGATTTGGGGTAG | 4 |
| wt-143640-143760 | 1193 | GCGGAGCCTCAGGAGGGTCCCCTCCATAGGGTTGAACCAGGAGGGGGAGGATCGGGCTCCGCCCCGATATACCTAGTGGGTGGAGCCTAGAGGTAGGTATCCATAGGGTTCCATTATCCT | 4 |
| wt-143760-143880 | 1194 | GGAGGTATCCTAAGCTCCGCCCCTATATACCAGGTGGGTGGAGCTAGGTAGGATTCAGCTAGGTTCCTACTGGGGTACCCCCCTACCCTACCTTAAGGTGCGCCACCCTTCCTCCTTCCG | 4 |
| wt-143880-144000 | 1195 | TTTTAATGGTAGAATAACCTATAGGTTATTAACCTAGTGGTGGAATAGGGTATTGCAGCTGGGTATATACCTATAGGTATATAGAACCTAGAGGAAGGGAACCCTATAGTGTAATCCCTC | 4 |
| wt-144000-144120 | 1196 | CCCCCCCTACCCCCCCCTCCCTTACGGTTGCCTGAGCCCATCCCCCACCCCAGCACCCCGGGGTGACGTGGCACCCCGCGTGCCTTACTGACTTGTCACCTTTGCACATTTGGTCAGCTG | 8 |
| wt-144120-144240 | 1197 | ACCGATGCTCGCCACTTCCTGGGTCATGACCTGGCCTGTGCCTTGTCCCATGGACAATGTCCCTCCAGCGTGGTGGCTGCCTTTGGGATGCATCACTTTGAGCCACTAAGCCCCCGTTGC | 4 |
| wt-144240-144360 | 1198 | TCGCCTTGCCTGCCTCACCATGACACACTAAGCCCCTGCTAATCCATGAGCCCCGCCTTTAGGAAGCACCACGTCCCGGGGACGGAAGCTGGATTTTGGCCAGTCTTCAATTTTGGGGAG | 4 |
| wt-144360-144480 | 1199 | TGGTTTTGTGTGAGCCGGAAGTTGGCAATGGGGTGAGGGTGGCGCTGGTTAAGCTGACGACCTCCCAAGGTCTCTCACCCTGGGTACACAGGTGGGGCGGCAGCCTCTAACTTTGGCTGT | 4 |
| wt-144480-144600 | 1200 | GGCCTCTATTTCCTCCCTTTCCTAGCCAGGGCCATGTGTTCCTGCATGTCTACTTGCCTCCTGTGGTGGCAGAGCTTGGCCCTGGGCCCAACCCCCGCCTTGGGAGCCTGTAGGGGCCAA | 4 |
| wt-144600-144720 | 1201 | CACCCTTGGTTTGTTTGTGTTCCTGTTTGCTGGCAACTTACTGGCAGCCGAGCAGATTCTAATGGGCGCCCGCCTTCTTTCTCTCTTGTTTTATTAATAGAATCTCAGCCAGGACCTATA | 4 |
| wt-144720-144840 | 1202 | CCTGAGACTTCAAAGTCTGGTCCTGGGTTCTGAGACCCCCAAGATTTGTCATGCACACCTGCACACCTGTTGGTATTGGGTTTCTATTCTTGAGTGTGAAAGTTTGTAAAAAAATTCATA | 4 |

| | | | |
|---|---|---|---|
| wt-144840-144960 | 1203 | AAATGTCACTAATTCCTCTTACCTGTTTAGGGTATTGTGCAATTCTTCAGCCTGCCTATTTTCAATTTGCCTAAGGTGGCAATTTAAGATGTGGTTAATTAACCATTTTCCTGTCTGACA | 4 |
| wt-144960-145080 | 1204 | CCACTGCATGGGCAACCGGGTTCCATGGCACATTTAGAGATAAACATAGATGTCTTGTCTTGCTCATGTGCAGAGGAGGGGGTGTTGGTGTGCAATATAGTTTCTGGATTCCAAATTGAG | 4 |
| wt-145080-145200 | 1205 | TTGGGGGTGCTATTTTCACTATGGAATTAAATTACTGACATTAGACAGTGGACACCGGGCTATATGTGGGGATGTCTGTGGCTTGTCATTTCCTCTTAGAAGGTAATCCCCCATCTTAAC | 4 |
| wt-145200-145320 | 1206 | TTCCCTTTAAATTGTGATGCAAGCCCTGGGTTATTTATAGAATGATTATCTAGGTTTGATAGTCTGAAGGCTGGGCAGAGAATGTTTGTAATTTTTATTCACCTTCTTTACCCCCCACGA | 4 |
| wt-145320-145440 | 1207 | GTATCCAGTTCTAGAAGATCTCCTGATATCCCGGGCTGCCATTATTCCCTTGAGTGTTATAGCTTCCTCTTAACTTAAGCAAGAGCTCCAGGATGTTAGCTTTTTTGGTGGGGCTGGTTG | 4 |
| wt-145440-145560 | 1208 | TCAGGAAGAGGTTCCAGTGTTGTCCTTTATTTTTAGATGTTAGCTTTGTGTTAGGTTAGTATGGGCTGGGTATTCACTAGTGAAGGCAACTAACACAGTTAGACGTGCTAGTTGTGCCCA | 4 |
| wt-145560-145680 | 1209 | CTGGTGTTTATCCGGTCCCAAATGTCACCACAGAACACAGGGGGCTGGATTTGGCAGCAGCACTTGTGCTTTTGTTGATTTTTACCCGTGTATCAGAGTGGGGGATGCTAGCCAATTTAG | 4 |
| wt-145680-145800 | 1210 | CTTCCCCTCCCCTTAACAGGGGGTCTCGCGGGGTGCCAATTGTCGCCTGCCTTCCCCCGCTTCCCCTTGTTAACTTATAGCATGATAGGTAGGTCACCTAACGTGGAAGCCTGGTGGGTG | 4 |
| wt-145800-145920 | 1211 | ATCCTTCCTCGGTAGGGAGCGCTTAGGGCTGTTGAGCTCAACAGCCCCACCTGGGTAAAATGTATGTTCTAAAGAGTTACCCAATTATAACAAAACTGTTGTAGGGTAACGAAGACCTGA | 4 |
| wt-145920-146040 | 1212 | TGGAAGTGGTATTGTTGCCGTTGAAAGACGGGTGTCCTGGCTCAAGTTCGCACTTCCTATACAGTGTTAAAGCCTTGTATCGGAAGTTTGGGCTTCGTCCCAGTGTACTCGATAATGTCG | 4 |
| wt-146040-146160 | 1213 | ACTGCTGCGAAAGGTTTGGACCGTCTTCCAGTAGGTGTTGGGGGTCCCAAATCACGAGGTTAGGCAGGTGCACTTGGCTCTTTAGGAGGGACCCTTAAGCCAGACAATGTAGTGCCCCTT | 4 |
| wt-146160-146280 | 1214 | TTTTTTGCAAATTGGCCTTATTATTAATTTCTTGTTAACACTAATTCTGTTCTATGACCCTGTGTTTTTCAGATGCCGTTGAACGTGTCACTGAGCTGAATTTGGACGCAGCTACTTGAC | 4 |
| wt-146280-146400 | 1215 | CTTTGCCCCCGTGCCTCCAGCGCTGATAAGTGCTGCGTCCACTTTGTGTTACAGGTGGGCCAAACCTCCAGAATATCAATTGGTGGGGCCTTGGTGGGCTGCATAAGGCAGTAGGTTTGA | 4 |
| wt-146400-146520 | 1216 | GGTGACCTACTTGGACCATGTGGATCCAGTGTCCTGATCCTGGACCTTGACTATGAAACAATTCTAAAAAAATGCATCATAGTCCAGTGTCCAGGGACAGTGCACTCGGAAGTCTCATCA | 4 |
| wt-146520-146640 | 1217 | TCTCCGTTTGTGTGTTTAGTGTGGCCAGTACGGCCACCCCTGTGCCACGCCCTGGCATGCTGCTGACATCTGGCCGCCAATTTCAGCGGGCCCTTTTCCCCCTTGTTCACCCCATAGCAA | 4 |
| wt-146640-146760 | 1218 | GAAGGGTAGGTTACATGGGTATTTTCCCATCAGCACCTGACTGGCCGGTGCAATTAGAGGAGAGGGCAACAACGCAAGGCTGTTGTTTTATTTGGGTTACAAGAGCTGCGGCGGTCGATG | 4 |
| wt-146760-146880 | 1219 | GGTTCACTGATTACGGTTTCCTAGATTGTACAGATGAACTAGAACTGTCACAATCTATGGGGTCGTAGACAGTGTGCTTACCAGACTTCCATGGAAGATGTGAATTTGCTGCTAGCTATA | 4 |
| wt-146880-147000 | 1220 | TGGGTGGTGCTATGGGCTCCCTAGGGACTCATGTAGTGGGGCTTTGTGATAGCTAATGAATGTGGCAGCTGTTGTTTGTACTGGACCCTGAATTGGAAACAGTAACTTGGATTCTGTAAC | 4 |
| wt-147000-147120 | 1221 | ACTTCATGGGTCCCGTAGTGACAACTATGCTGAATATCTTGAATATGGGAGGAGGGGGGCTTTGGGTTCCATTGTGTGCCCTTTCCTGGCCAACGTGAGGGTCCTAGTGTTATAGGGCGT | 4 |
| wt-147120-147240 | 1222 | GGCAGTTTTCTTGAGGGCTAATAACCCGGGTGAGGCGGTTGTCACAGGTGCTAGACCCTGGAGTTGAACCAGTACCACTCGGTTACAAAGTCATGGTCTAGTAGTTGTGACCCTGCAAAG | 4 |
| wt-147240-147360 | 1223 | CTACGTGGGGATGAGCAGCCAGGGACTTTGGTTGGCAAGCAGACAGGCGGCGCATTGGAACCCCAGAGGAGTGTCCCGGGGCCACCTCTTTGGTTCTGTACATATTTTGTTATTGTACAT | 4 |
| wt-147360-147480 | 1224 | AACCATGGAGTTGGCTGTGGTGCACTCCATCTGGTAAGGGGGCTGGTGCGGACGCCTGTGTTTAGTCTATGCCAATGTTTACCTGCCTTGGGTTACTATTCCAAACGACCACACCTTTGA | 4 |
| wt-147480-147600 | 1225 | GGACACCTGGAGCCCTGATCATTCTCGGCTTTTACTGCCACCTGGCTTCTGTTGGGTCAGACAGTTTGGTGCGCTAGTTGTGTGCTTAGCAGCAACGCACACCAGGCTGACTGCCTTAGC | 4 |
| wt-147600-147720 | 1226 | AGTGTGGCCCTTTATTGTGGCATCCTAAGGAGGGATTCTGGAGTGCCTTTCGCGTGAAGCATGCCCTGAGACGTACTCGAGTTAGGACTTAATCGCTCCTGTGCCGCTGGATGAGGGAGC | 4 |
| wt-147720-147840 | 1227 | GCCAATTTGTACATCCTAGCTCTGGCCATAGAGTTAGCCCACCCTTGTGTCTCCCTTTGGCCTTTGCGGTGCCAATTTCCGGTGGTTTCCCTTTTCCGCCCGTTTATCCAATAGCATGTA | 4 |
| wt-147840-147960 | 1228 | AGAGAGGTTGCCTAGATTTGGCAACTTTGAGGGAACGTTCCGTGTAGCTGGTGACCTAACACCCGCCCATCACCACCGGACAGATTCTGAACTTGTCCTGTGGTGTTTGGTGTGGTTTTG | 4 |
| wt-147960-148080 | 1229 | GGGTACGCAGGAGTACGTTGGAATGCTTTGGAGCCGAGAGGGATGGGCCCGCTTGTGCGCTTATGTGTTACACGGTGCCAATAACCGGCCCGGTGCGGCTGCCCCGTGACCCGTGGGCCT | 4 |
| wt-148080-148200 | 1230 | TACCTTCCTGGCCATCGGGGGACCCTGGTGCTAGGGTCCCTTGTGTTGCTTTCTGCCATAGGGGGGAAAGCATCGCCTTCAGAATTGGCTGCTCCGTTGGAACATTTGAGGCCTACTGTA | 4 |
| wt-148200-148320 | 1231 | TCCGTGTCCTGACAACATTCCCCGCAAACATGACATGGGTTAATTTAAACATGTTTTGTTTGCTTGGGAATGCTCTTAGGGCCTGGAAGCTTGTCATTGGATTCATCGTTTCCTGAACTA | 4 |
| wt-148320-148440 | 1232 | CAGGCGTAGGGCCTATTGTAGCAGGCATGTCTTCATTCCTGCGTACCGAATGGCATGAAGGCACAGCCTGTTACCATTGGCACCTTTTTTCCATGTAAACCTCCGTGATCCTGGGTCCTT | 4 |
| wt-148440-148560 | 1233 | TGGAGACTCAAGTGTGAATTTGTTTTGGTGTTCGGCGCCAGGGTATCTCGACGTTGGAATGTCAACTCAACTTGGGCACCTCGATAACCGGCTCGTGGCTCGTACAGACGATTGTTTGGC | 4 |
| wt-148560-148680 | 1234 | TCTGTAACTTGCCAGGGACGGCTGACGATGTGTTTAGTCTGCCACTTGCATCCGGCGCTTTGGTTACTCGGGAGACTAATGGGGGGTGTGGTATGGCACAGGCTGGGGGTGAGTCTGGGG | 4 |
| wt-148680-148800 | 1235 | ATGTCCCTGGGCGTTGCTGCAGCCCATTCGCCCTCTGGGGATGAGATGTTCAGGGGTGGCCGGTACCCTACGCTGCCGATTTACATAATATAAATTGTAAATGCTGCAGTAGTAGGGATC | 4 |
| wt-148800-148920 | 1236 | TGGACGCGCGACCTGCTACTCTTCGGAAACGCCAACCCAGGAGCGTCGCCTCTGGCCCCATACTCCCGCCATGCGACTGCTCGCCCCCTCCCAGGCCTCCCTGGTGAGCCCTTGCCGCTC | 8 |
| wt-148920-149040 | 1237 | CCCGCATTCCTGCTTTCGGCGCCCCTGCGGATCCCGATGACAGCAGGCCTTTCCTTCCCCCGTTAATGAAAAGAATGACAGTGAGGTTGTGACAGAAGGACAGCTTTATTCAGTTTACAG | 4 |
| wt-149040-149160 | 1238 | AGTGCCCTCGGAGGCTACGATATTCCCGTTAAATGTCTTGTTGATTCTCTCAAAGGTGGGGAGGGAGGAGCTCTCCACAACAATGTTCCCTGGCAGCGTGAGCGCGCAGCCCTGCCGTTG | 4 |
| wt-149160-149280 | 1239 | GATGTATCTTCTCATGATGGTGCTGATAGAGGGGTCTCCGGCGTAGATGAAAAAGGCCTGGGCCATGCTCTGGCCGGTCACGATCGTTATGGGGTTGTTGGAAATGTTCCGGACCGTCAG | 4 |
| wt-149280-149400 | 1240 | CTTGAGGGTCTGGCCCGGCTTCCACTCCTGTGGGTAGACGTAGAAGACCGGGTTGGAGGAGTGGGACACGACAACGGCCGTAATCTTGGAGCTCAGGGGGGCCTCGTAGGTGTTGTTGTA | 4 |
| wt-149400-149520 | 1241 | TTCCAGCTCCGTGATGAAATTAGGAGGAATAATCACAGGGGAGCCAAAGTAGCGGATGTCTGTGGATTCCCCGTCCCAGCGCCAGTGGCTCTTAGGGTAGGGGTTGTAACGGAAGGCAAT | 4 |
| wt-149520-149640 | 1242 | AATCACATCATCCAATAGGGTCATGCCCACCTTGACGTTCAGCGGGCCCTCTCGTTTCAGGTCCGGCGTGTCCACGGAGACTCGGACGTAGCCCTTACCGCGGCGTATGGCGTTTACCGG | 4 |
| wt-149640-149760 | 1243 | ACACACCTTCCCCGGGAATGTGTGAATACGGGCGTATGACTTTAGAAATGGGGGCGTGTGCTGCGCCAGCAGGTAAGGCAGGCACTCGTCCTGGCTGGTGACGGGAGAGCCACTGAGGAA | 4 |
| wt-149760-149880 | 1244 | GATCTGGGGCTCGCTGGTGTTTAGCTTGTCCCCGCTCTGGGTGCAGGAGCGTGTCAGCTGAATGTCGCTCTGCCCGGGCAGAATCTGCAGGTAGAGGTAGGGGTTCTTGACCAATCTGAT | 4 |
| wt-149880-150000 | 1245 | GGGCACAATGTACCAGGTAAACTTCCCTTTCTCTATGAACAGGCTGCGCGGATTCAGGACGCTTAGCACGATGTCCTGGTCAGAGTGCATAACGAAGAAGGGCTTGAGGAATACCTCGTT | 4 |
| wt-150000-150120 | 1246 | GTCTTCCGCTCCAAAGAACAAAAACGCGACCGTAAAGTAGCGGCTGCCGTAGGTGGTCGTGTTGAAGGAGAAAGAAGGTAACTTGAAGCTGAGTATCTGGCCCACCGAGGGGCAGGGAGG | 4 |
| wt-150120-150240 | 1247 | CAGCTCTTGGCACTGCGCGTCCAGCTGCAATACCTGCTTGTTGGTGACGCGGACGTATGAGGGGAAGATCTCGTACTTCCACACGCCTCTCATGAACGACGTGTCTGGTTTTTCAGTGGG | 4 |
| wt-150240-150360 | 1248 | CCGCAGGCGGCGGAGGCTGTTCCTGAACGACGAGCGCCGGGACGCTAGTGCTGCATGGGCTCCTCCGGGGTAAGCTTCGGCCATGGCCGGAGCTCGTCGACGGGCAAGGTGAGAGTCGGG | 8 |
| wt-150360-150480 | 1249 | GGGCGGGCGACGGTGCGGCCCCAATACAACTCTCCGCTCGTTAGCTGGTAGAATATCCGCCCGGCGTCTAGGTTGTCACTTCGCTCGGCCGGCCAGAAGAGCGCAAGTCCAAGTCTGGTG | 4 |
| wt-150480-150600 | 1250 | CTGGGGCCGATGTGCAGCGGTTTGTGCCCGCAGTTGTAGACTGTCATTTTTATGGGCGAGTGGGCGGTCCACACGCGCGGGCGCAGCACCCATTGGTCGCACGCCGCCTCCTGGAATGTA | 4 |
| wt-150600-150720 | 1251 | AACCCCCAGAGAGAGGGCGTGCCGCCCTGGAGATGGCCCTGTGCCATCACATGTATTTCCTCCTTGGGTGGAACAACGGCGTCGTGCTCCGGGTGGAGGGGGAATAGCGTCCAGGCATCT | 4 |
| wt-150720-150840 | 1252 | TTCAGGGTCACGAGACCGGGGTCCATGCTCAGAGAACAGCCCTCCCGGGCGGTGGGCGGCCCGGGCTCCAGCAGAACGTCGCAGACCCAGCCCTCCTCGGCCCTGTCCACCTGTATGTCC | 8 |
| wt-150840-150960 | 1253 | AGGTGCACGGACCCGGAGGCTGCGTCTCGTGACATGGCCAGGCCTGGTGCCAGCCGACCACGTCCCGTGTCCCAGCCGAGGCCGCGCCAGAGCAGAGCCCGGGACTGACTCAGGGCCACA | 8 |

| | | | |
|---|---|---|---|
| wt-150960-151080 | 1254 | TCCCCTCGGCCCGCGGACGCCGCCTCGCCAGCCCCCGGGCCTTCATGGGCCCGCTTTCTACCTCTCTCCGGCACCCCAGCCTGGTCAGCCGCAGAGGAAGCATGACCTTGGGGTGGGACG | 8 |
| wt-151080-151200 | 1255 | GGGCAGGCGTGATCCTGGGCGCAATCTTTGCCGATCCCCACACCTTCACTCCTTGTTAGGTTGATAGAATGTCGGTACCACGCCACGGGGGGCGGGCCCGCATAGGGAAAAGCCAGGGAG | 4 |
| wt-151200-151320 | 1256 | AGCGATGTGGGCGAGGATGGGCTCAGGCGGCCCCAGACACGCAATTTGCCCCCCTGGGCGGCCGCAGCCTGCCCCTCGGCGGCCCGTGCCCCAGCTCCGTCACGGGGGGCGCATAGGAGG | 8 |
| wt-151320-151440 | 1257 | GGTATATCTAGGATAGCCGCACCTACACAAATGAGACACAGACACAGGTCGTGAGGATTTAGGCAACGCAGGCTTGTCTTTATAGTTACAAACATGGGAGCGTGCACCTGGAAGATGCAG | 4 |
| wt-151440-151560 | 1258 | CTGGGGTAGATCTTTACATCTTTACAGGGCGCAGCGGCCGCCAGACACTGAAGGGCAGAGTTCACGGCGGGCACCTCCCAGAGGGAGCCCACCAGCCCGTACCTGGCCACGGCCAGGGCC | 8 |
| wt-151560-151680 | 1259 | TCGTAGGCCGAGACGGGCAGCCGGAGCTTGTGGTACTGTCCCTCCGGCAGGTGGAGTGGGACACAGTTAGAGAACATTAGTCCTCTGGTCCCTATCTCCACCCGCCAGGCCTGTGTGTCA | 4 |
| wt-151680-151800 | 1260 | GTTTGCAGGGCCATCCTCGCACTCAGGTGGACTGGCTAGGCACCCTTCTGAAGTATCTGGCGGTGACTGTCACCTGGTTCTTGAGAGAGTCCATAAAATGGCTGAAGCTCCAGGCGTATA | 4 |
| wt-151800-151920 | 1261 | GTATAATGAGCAACAGGGCCAAACAGGCGGCGGGGCCTGGGTAGTAGCGGGCAACGAGAGACTCTGTGCAATCAAACCCCAGGCTCCCGGCCTCACCCAGGAAGAGCAGCGGCAGGGACA | 4 |
| wt-151920-152040 | 1262 | GCATAAACCAGGAGAAGGCGCAAATGAGTCCGGTGAAGGTGACGTTGCATATCAGGCGCGGCTTCCTTCCGAATTTTGTGCGCAAAAGTTTCCAGATGATGATAACTGTGAGGAGGACGA | 4 |
| wt-152040-152160 | 1263 | TCAGGACTGCCGCCAGTAGGTAGCAGCCGGCTTTCAGTCCTTGGACGGCCGTGTGCATGCCTTTGGTGGGGCCTTCCCTGCACATGTTGGGGCCTCTGTTGAGATTGGCGTCGGGGCCCA | 4 |
| wt-152160-152280 | 1264 | TGGTAATGAGGAGGATGATAATCAGCAGGAGTACCAGACAAAACACGCCCATCAGGTACAGGCACACATTTCTGTGGGAGGTTCTCTTGGGCGTTCGGCTGAACAATGCTAGGGTCTTCT | 4 |
| wt-152280-152400 | 1265 | CCAACGCCATACCCAAGTGAGTCCATACGGAGCACATCAGGCCCAAGAACATCATGTTCTGGGTCAAAAGGCAGAGACCGGTAGACGAGAACTCCTGAATCATTTTTCCCAGCACCCAGA | 4 |
| wt-152400-152520 | 1266 | GCAGCAGTTCTATGAGAAGAGCTATCAGCCAGACATCCATTCGGTGAACCAATTTTCTTACAAAGATGATAAACAAGATGCCAGCCAGTGTTAGCAGAATCAGCAGGACGAGCAGCAGGC | 4 |
| wt-152520-152640 | 1267 | TTGTCATGCCGCTGAGGAAGGCGCTGTAGGATTTAGTGCACGCATCTTCCGTTGCATTGACGGAAGTCATGTTGGCCACCAGGGTCCCCACGGTGGACCCGGGGGCCATGGTGGAGAGCA | 4 |
| wt-152640-152760 | 1268 | TCTTGCTGGTCAGAGCCAGACTGGGTGGTGTCTGCAGCAAAAGAGGAACTTGCCCAGGCAGTCAGTTATTTTGCATGCCACCTCCCTGCCTGGTGGACTTCCAGACTATTTTCTGCATTC | 4 |
| wt-152760-152880 | 1269 | GCCCTTGCGTGTCCATTGTTGCAAGGAGCGATTTGGAGAAAATAAACTGTGAGTTTCACAGATCCACGGGCCACGCTCCCCTGGGGGCTTCATGATCCCACCGCCTTTCCCGATGATGAT | 4 |
| wt-152880-153000 | 1270 | GACAACCGCGGCTGTCTGAAGCGGCTGACGAAATCGGTTGAGATTCTGATGAGAGGCTTGGGGGGGTCTTTGCCCTCAAGGCGAGGCTCCTTCTCCTAGGAATGCCGAGCCCCCTGCACT | 4 |
| wt-153000-153120 | 1271 | AGCTTCGCTCCACTGGGGATCTTTGCCAGCCTTCATACTAGATTCAGCGATCCCCCGGTTGGGAATCTTCGCCAGCCCCCCGTCCTGCTATCCCGCTCGTCGCCGCGCCTCCCATGCTAA | 4 |
| wt-153120-153240 | 1272 | GGGCCCCCTTCCTTTCCCTTGACTTTGGGGATATTCGGAGTCTGCTCTCGCCGCTCTCTTCTCTCGTTTAAACGAGAGAATAGTAGTAGGGTCCAGTCTCAGGCCCCCTCACTTTGGGTC | 4 |
| wt-153240-153360 | 1273 | TTAGAATGGTGGCCGGGCTGTAAAATTCTGGAGGACGGAGAGGGCGGCCCCGGAGTTGTTATCAAAGAGGCACTGGAGGATGTTGGCCGCTCCTTGGAGCAGCTTGTCGAAATAATGATC | 4 |
| wt-153360-153480 | 1274 | CACGGCCACGGGAACGCCGTGCCGCTCGGCGTAGGCCGGGTCCTCGGCCATCTCCGTCTTTCTCGCCCCCTTCACTCCCCCCTTGGGCTCCACAAAGACGTACTGGATGCGGTCGTGGAT | 8 |
| wt-153480-153600 | 1275 | CTGGGGCAGTTCCTCGTTGCGCTCGACGAACTTCTGGTAGACGGCCAGGTGAGGCATCTGGGTGCTCTTGTAGGCTGAGAGCTTGCGGCTGAGCTCCGTTGAAAAGCAGAGCTCCCCCAT | 4 |
| wt-153600-153720 | 1276 | GGGGACCCTGCCTTCACGGAGGTCTGTGTAGGCCTGGTTTAGGATGTCAATGACGGGCAAAAAGCCCACAGGTAGCCCTTGTGTAAATGACTCTTGGAAGGGCCGGTGGGAGAGGAGGCT | 4 |
| wt-153720-153840 | 1277 | GGCCGCCTCCTTTACCCGGGCATCCGCCAGCACCAGGTCGAGCACGCGCCGGCAGCGTGTCTGCACAAACTTGCAGGCCGTCTTCCGGACGAGCTCCACCCCCTTCATCAGGGTCTTGCC | 8 |
| wt-153840-153960 | 1278 | GTCCGTCAGCACCCCCACATATCTCTTCTTTGTAATCAGCATCAGGCAGGAGAAGGTCTTCTCGGCCTCCAGGGAGATGGGGGCCACAAACAGGCTCCGGGTGGTGTGGGCGGCCAGGGC | 4 |
| wt-154080-154200 | 1280 | GGAGGGGGCCAGGGCCTGCAGGTTGGCGGGGCTCAGGGCCTCCACGAAGGCCTTGGCCCGCTCCAACATCGTGCGGCCCTGCAGCGTCACCGTCTCGGCGATGGAGAGGCAGGGAAAGAG | 8 |
| wt-154200-154320 | 1281 | GCCGTTGGCCACCCCGGTGAAGCCGTAGACGGCGTTGCACGTGCACTTGATGGCCAGCTGCTGCTTGTCGAGGATGGTCCTTTGGCGCGGATCCTCGCAGGCCGCCAGCAGCTTCTTGAT | 4 |
| wt-154320-154440 | 1282 | GGCCTTGCGCTTGGCCAGCCAGGAGGTCAACAGACTAGCCAAGAAGGACTCGTGCACGTGCTTCTTTACAAAGTGGTAGACGCCCCCCGTGAGCCTGAAGGACTCATAGTCTTCTCCCGG | 4 |
| wt-154440-154560 | 1283 | GCGCAGGCCGGCTAGCCTGTGCTCTTCTCCCGGCGTTATCATGGTAGAATAACAGAGATTATGAGCCTGAATGATGCTCGGGTAGAGGCTGGCAAAGTCCACCACCAGAACCGGGGAGTT | 4 |
| wt-154560-154680 | 1284 | GTAGAATCCGGACAGGGGCTGGATGACGGTGGCCCCCTGGTAGCCGTCCCGGTCAGAGGCCGAGGGCATGGGCAGGATAAAGTTTTCCTTTTGGGCGGCCGCCAGGAGGCAGGAGAACAC | 4 |
| wt-154680-154800 | 1285 | GCGGATCTGCTGCCCATCGTCCAGCACCCGCCTGCAGGGGATGTGAGCGATCTTGGCAATCTCTGCCACCTCCACGTGGATCACGAAATGGTTTAGCAGATCCATGACCAGGGCCGAGTC | 4 |
| wt-154800-154920 | 1286 | CTGCACGCAGTACATGCCGAGCCGCCTGCGCCCCTCGGGGCCCGCTGCAAAGAGGCGAGGAATCTCCTTGTAATGCACATCCTCCTTCTTGGCCCCCAGTAGGTGCCTGGCTACTGTGTC | 4 |
| wt-154920-155040 | 1287 | CAGCTTGTAGTCTGAGAGGCTGAGCTTGTCCCGGCACACGGCGTACATGTCGATGGGGATGAGGCCGGTGATGCGGACCTTGGTGTTGGCCCGCAAGAAGCCCTTGCCCGCATCATGGGG | 4 |
| wt-155040-155160 | 1288 | TCGCCTGACCTCGCAGACGCCCCCAGCCCTAATTTTGCCCAGAGAGGCTGGGTTGATGCTGTAGATGTGCCTGGCTCTGTCCAGAATGTAGGGCCAGTCAAAGTTGGCCACGTTGTAGCC | 4 |
| wt-155160-155280 | 1289 | GGTCACAATCTCCACGCTGAGGTCTCTGATGAGCTGGAAGAAGGCGTAGAGCATGTCCAGCTCCGATGGGAACTCGTAGACCTCAACCCCCTCTATGTCTTCGCAGGTGCCCAGCGTCAG | 4 |
| wt-155400-155520 | 1291 | ATCGAAGGCCAGGGCCTGGTAGGAGGGCCAGGAGCTGTCTTCACGCCGGACCGAGAGGTCGCCCACCTCACAGTCGTACTCGAGCTCGGCGTACGAGTCCCGGTGCTGGAGGCGGGGGAT | 8 |
| wt-155520-155640 | 1292 | GGCGCGGCGGCAGCTGTACCAGCCAAAGGTGACAAAGTCATTGTCCAGGACAAAGCGGCGCGTGGCATCCACGTTGGCCTCAAAGATCCGACACCCGTGCTTGTCTTGCAGCCACGTGGC | 4 |
| wt-155640-155760 | 1293 | CACGTGACACACACTGTTGGGATGGGAGAGGGTGATCTTGTGGTAGTCGCCGGCATGGTTGCCGTAGCCCATAATGGAACGGCGCGTGACCTTCTCCACCGAGACCCGGCAGGGGGTCCT | 4 |
| wt-155880-156000 | 1295 | ATCGGCCTTGCGCCGGCCCAGTAGCTTGATGACGATGCCACATGGCACCACATACCCCTGTTTATCCGATGGAATGACGGCGCATTTCTCGTGCGTGTACACCGTCTCGAGTATGTCGTA | 4 |
| wt-156000-156120 | 1296 | GACATGGAAGTCCAGAGGGCTTCCGTGGGTGTCTGCCTCCGGCCTTGCCGTGCCCTCTTGGGCACGCTGGCGCCACCACATGCCCTTTCCATCCTCGTCACCCCCCACCACCGTCAGGGA | 8 |
| wt-156120-156240 | 1297 | GTCTTGGTAGAAGCACAGGGGGGGCTGAGGCCCCCGCACATCCACCACCCCTGCGGCGCCTGGTGTCTGGAAACACTTGGGAATGAGACGCAGGTACTCCTTGTCAGGCTTTTTCAGAAG | 4 |
| wt-156240-156360 | 1298 | GCCTTTATTAGGTCTTAGGAAAGGGTTATAGAAGAGTCCCCCAGACATGGTTAAAACTCAGTCTCTGCCTCCCCAAGCAGTGCGGCGGCGGTCTCTGGATCGTGATAGCGTCTTCTGCGT | 4 |
| wt-156360-156480 | 1299 | AGGCCTGGAAAACGGTCCCTGGCTGCCTGCAATGCTCTGCTGGCCACTGAGGGTCCGGCCGCCCTCTGAGCTGCTCTCTTTTGCTCCTGGTTTTGCTCATGCAGCGCTAACATGATGGCT | 4 |
| wt-156480-156600 | 1300 | TGTAATTCTGTCTTACTAATGGGATTAATGCCTGGACCCTCACCAGAGGCATGTTGCTGAGCGAGCTCGTCGATCCCGGGGTAGAGCATCTGCACCGGCTGCTGCGACATCTGGCGCGTG | 4 |
| wt-156600-156720 | 1301 | CGCCTCGTGAGGGAAATAACCAGGATCACCACCCCCGCCACCAGGACCAGAATGAGCATGCCGCCGAAGGGGTTTTTGAAGAAGGAGATGAAACCAGAGACCAGGCTGCTAAACAAACCC | 4 |
| wt-156720-156840 | 1302 | CCCACCGTGCTGACTAGGTTGGTGATGGACTGACCCACGCTACCCAGACTGTCCATAAGTTCCCCCAGGCCGTCCACGAATTGATTTCTTCCGTTTGACACTGCATTGTCCAAATCCTTC | 4 |
| wt-156840-156960 | 1303 | CGCAGGCCGGCGATGTTTTGCGCCTGGAAGTTGTACTCCCGGAAGATGCCCTCCAGGTCAAAGACGTTGGAGGCACGCTGTTCGTCCCGTGAGTACAGCTCCAGGGAGGCAAAGTCAATG | 4 |
| wt-156960-157080 | 1304 | TTCTCGATGAGGGAGGTGTTTAGTGAGATGAAGGTCTGCAGGGTGGCAATGCCGTCCAGCTCGATGGTTTTAAAGTGGTGGTAGTCGTTGTAGACGTGGATCTCGTTGCCGGACTGGAAG | 4 |
| wt-157080-157200 | 1305 | TAGTACTGGCTGGTCGCCTGGCACACCTCCGTCATCTTTTTTGTGAGGAAGATCTCGTTGTCGGTGCCCAGCTGTCCCTCGTAGGTCTTGGTGTCGTTGATAAAGCTGAAGGACACCAGG | 4 |
| wt-157200-157320 | 1306 | GGGCGCGAGTAGCACATGGTCTCGGAGCCAGGGACCCTCATGCTCTTGCGCAGGGTGACGGTGGCCTGGTTAACGGGCACGCACTGGGAGACTGAGATGACATCCCCCAGGCGCTTGGCC | 8 |
| wt-157320-157440 | 1307 | GCCACCGCCTTACCGTAGATGCTGGACATGACGGTGGTTGGATTAATCTTGGTTAGTTCTCTCAGCACCATGTTCTGCCTCTTCTGCTCCAGGCACCAGGCCCGCGCAAGGTCTCCCAGC | 4 |

| | | | |
|---|---|---|---|
| wt-157440-157560 | 1308 | ATGCGGTTGATCTGGCGGCGCAGGGAGTCGTAGGCAAATTGGATCTGGACGGTGGCGGGATTGTTGAGGGTGCCCAGGGACTTCCCGGGGGCCGTGGGGGGCACCGGTGTGGTGGCGTTC | 8 |
| wt-157560-157680 | 1309 | CCCGCATCCCGCCTCCGACGCCTCAGAACGGCGGCGGGGGTGCTCCCGCGGGCCGCGGATGGGGCTGGGGGCGATGGACTGCTGGGGGGTGAGGAAGTCGGAGTGGTAAGCTCCGTCAGG | 8 |
| wt-157680-157800 | 1310 | TTCTTGACGGTGGCCAACGAGCGCGGGGTCAGAGGTAGCCAAGCTAATAACAATCCTCCGCTCGTTATAAAATATGTAATGGCTTCCTGGCCCTTCGTGTAACGATCCTGGACGGCCTCG | 4 |
| wt-157800-157920 | 1311 | TACTTCTCATGCATGGTCTTGTTCACCTGCTCTTCGATGCACTTGAAGGCGTCCGGGAGCTCTATGCCCACGGTTGTGTTGGTCACGAAGCTAGAGGTGCCCTCGTCAGTCACAAAATGT | 4 |
| wt-157920-158040 | 1312 | ATTGACTTCCCTGTTTCTGTGGCGATGGTCGAGTCAAAGGTTTGCCAGTGTTGAAGCGGGCAGTAGGCTGTCCTGTTCTCGAGCTTCCAAGATAGCGTGTAAGTGCCCTTGTCCAGGAAG | 4 |
| wt-158040-158160 | 1313 | GCTCGGCGTTCGCCTTGCGGGTTCGTCCCTCGGTTGTCGTAGTCCACTATCTTGTAGTTAGTTCTCACGTGGAAGGAGTCTGCCCGCTCATGGAAGGTTTCCTTATTTTTCCCGTCATAG | 4 |
| wt-158160-158280 | 1314 | AAAGGGGACATTTCCACAGTCTGCCCGGTGGTGGTCACAAAGAAGTCGAAGGGGCTGTTGGACTTGGCCATCATGTCAGTTATCAGGCAGTTGACGGTAGTTCTTGTTCTGTAAGTCCAT | 4 |
| wt-158280-158400 | 1315 | ATCAACCACCCGGGGGCGTCATAGAGCTCCGTCTGGCTGGCGTAGCGGCGCACCCCGTTGGCCAGGCCCCCGGTGGGCTTTAGGTTGACGGTGATGTTAACTCCGTCGCGGTCTACATAC | 4 |
| wt-158400-158520 | 1316 | ACGCGCGTCAGCCCATCTTTTGTCATCTTGACCGCGTTGTAGCACTGGTAGATGGTATCCATCTGGTCAGTTTCGTAGCTGTCAACGGAGAACTTCTCCTCGTGCCGGTTGGTCACGGAG | 4 |
| wt-158520-158640 | 1317 | TCCGCGTACCAGCCATTGTAGATGAGAATGTTGGTCACTATCTTGGTGTAGGAGCGGACCTTAAACGAGTAGGGAATAATGTTGTCTTTAAACACCATCAACAGGCCCTCCGTGTGATTC | 4 |
| wt-158640-158760 | 1318 | TCCCGCGTGCCAAACGAGGGACACTGGATGTCCGAGGAGAAGCGGAACAGGTCGCCGTGGCTGGAGAGCTCGCAGACTCGGAAAGGAAAGCTGGTTTGCTGACGCGTGGCGGTAGGCTGC | 4 |
| wt-158760-158880 | 1319 | ACCGTGGTGGCGGGGGGTGCGGGCTGCTCTGGGGTCTGCGCACCGAGACGGCACGCCAGGGCGGCTAGCAGCACGACCACGCTTAGCACCCTACGCCGAGTCATCTCTCATTTGGAGGTG | 8 |
| wt-158880-159000 | 1320 | CAGGTAGAGAAGGGCATATAGATCCTTAAATACCCACCCCCTGCCCTTATACAGAAGAATTAGGGGGCGGTCAGAGTCGTACGTGAGGTAAAGCCCATCCGGGGGCAGGGCCTGGCCGGG | 4 |
| wt-159000-159120 | 1321 | GCTGACCGCGTCCGCCCGGCGCAGGATCAAGGACCGCCCCCAGGTCTTGTTGTAGAGGGACACGGTTAGGACGGCCTCGCGCAGCGCCCGGCACAGAATTTGCTGGCTAGATGCCAGTGA | 8 |
| wt-159120-159240 | 1322 | GCCCCCGGGTACGCTGTAGAAGCTGTTGAAGGAGGTCTCTATCCAGTCGCTCGGCTCGATGCCTGGCCATATCAGGGAAGTCAGGAATGCCTTCTGGTGGGGCAGCGTACCTGCGGCGTC | 4 |
| wt-159240-159360 | 1323 | ACAGCAGCGAGCCAGGGCCACGTTGCTGGGTGGGGAAAGAGCCCGCTCTCCTCCGCCAGGGGCCCCGTGATGAAGGTGTACAGGCTGTGCGTCAGCGCGTGCAGGTGCTCCGAGCTCAG | 8 |
| wt-159360-159480 | 1324 | GGTCTGGGTAAACAGGTGTGTTTTGATGTACTTGGAATTCTCAAAGGCGGCACCCTCGCCGGCGCGCCTGTCCTCCCAGGGACCCGAGACGAAGGCCCGTCTGTAGAGGAAGTGGTTGCG | 4 |
| wt-159480-159600 | 1325 | CATGCGGGCCAGCTCCCAGTAGACCACGTCCCCCCAGACGCGCAGGCACAGGGTCTCGGTCAGGGTCTCGCTCTGTTGCGCCAGGCAGGACTGCAGCTTGGCCAGACCCTCGGTGGCCAC | 8 |
| wt-159600-159720 | 1326 | CTGGCGCAGGTACTGCTCCTTGCGCTTGAGCGCGTCCGAGAGGGCGCCGGACGGGCCGGGCTCTCGTGCCCCAGCCGGCCGGGGCACCTCCGGGCTCTCCCGGGACGCCTCCTCCTCGCC | 12 |
| wt-159720-159840 | 1327 | TCGGCCCAACCGCTGCATGGCTCGGTTGAGCCGCGTGTACAGCTCGTTCCTCTTTTGCAGGATGGCCCGGTACTGGGGGTGCGCCGTGAAGGCGGCGGCGCAGTCCGCCTTCAGCGCCTC | 8 |
| wt-159840-159960 | 1328 | CACCGCGTCGCCCGAGGAGCTGTAGACCCCGCCGCAGAAGAGCCGCTCCGTGGCCCCGGGAGCCACGGCGTCAAACAGGTGAGTCAGCCTTGCCCCCGCCAGCGCCTCCTCGCAGGCCCC | 8 |
| wt-159960-160080 | 1329 | CCGCACCAGGGCCAGGCGACGCTCCCGGGCAAACAGGGCAGAGAGGCGGGAATGGCCGCCACCCTCCCCCTGCCCCGTTGCACCGATAGCATGGCCGCCAGAGTTCCAATAGAGGAGCTC | 8 |
| wt-160080-160200 | 1330 | CGAGAGCTCCGCCACCTCCGGGGGCACTGTCGAGAAGACGTTGTAGGTGTCCAGCGCTCTGGTCGCCCCCTCTGCCTCCGGCCGCCCCGGGCCCGGGACCGCGCCCTCCTCTGGGCCGCC | 12 |
| wt-160200-160320 | 1331 | CGGCCTCGCCTTCTCCTCAGCCTCCAACAGGTGCCCGAGCCCCGCCTGGCGGACTTCATTCTCAAACAGTCCCGAGACCGGCTCCGGATTCACCGGCACCGCCAGGTGGTTACAGGAGAC | 8 |
| wt-160320-160440 | 1332 | GTGGGTCCCCTCTGCCGTGGAAGGGTTGCCGTGGTTGGGCAGAACCATCAGCTCGCCCACACAGCGCCAGCAGGGCACAGAGGTGATGTAGAGGCGCGGGTCTGGGATGGGACTTACGCC | 8 |
| wt-160440-160560 | 1333 | CCGAAAGCGGCCCAGCAGATCCAGGGCCCGTTCCAGGCTCTCCAGCCCCATGGTGTGAGACATGCAATAAAACACGCTATTGATTCTCTTCATTAAAATCTCTATGTCATTTATTAGGCA | 4 |
| wt-160560-160680 | 1334 | CAAACTTACATCGACTTTATGCCCCCCGTAAAACTCCACAGAGTACGCGACTGAGGGGGTACGGAGAGGCGGGACCCGGGTACCCTTTCTACCAGGGGCGAGCAGCGCGGCAGAGGCCTC | 4 |
| wt-160800-160920 | 1336 | CACCACCGCAGCCACGTCCAGACTCCGGGGGTCCAGCCGGGTGCACACGCTCAGCTCAACCGCCAGGGCGTACACCTGGCTGTACGCCGCCGCCAGCAGCCCCGACATCGCCGCCCCAGG | 8 |
| wt-160920-161040 | 1337 | GGTCTCTAGACCTCGAGTCCGGGGAGAACGGTGGCCAGACGGCGCTTGCGTCTGCCCCCGGAGCCCTGCCCTCCTCCACCCAGCAGCAGCCCGGCCGAGGCCTGCGACGCGGTGCTGACC | 8 |
| wt-161040-161160 | 1338 | GGCTCGGCCACGCTGATAAAGTTGTCCTGGGCTGCCCCGGGCCCACCCCACACTCCCTCCAGAAAGTCCCGAGCGGCCTCCGCCGTCCACTCTATCCCGCTGGAGGCAATGGTCGCCAGG | 8 |
| wt-161160-161280 | 1339 | GTTTCTAGGACGCTGTCCGCCAGGACGGAGAAGCGGCCCAATAAGTACTCCGCGTCGTCCCTAGTCAGCGAGGCGCATGCCTCGCCCATGGCATCCACAAGGTTGCACACCACATCAAAC | 4 |
| wt-161280-161400 | 1340 | ACACAGTCTTCCTCCTGTTTTTGTGATATAATGGCCTCCAGGCCAGCCCTGATGTTCTCAATCTCATATGTGGTCGCGGCTTGGGTCCGGCGCTTCACGGTCAACCCTAGGGTGGGGGTG | 4 |
| wt-161400-161520 | 1341 | GCAAAGACAAACTTCTTCCGCATGGAAGAGCCCCCGGCCTGCTTGCGCAGCCCAGCCCCGGGGGCCTGCAGCAGGTTCCTGTCCACGCCCCGGCCCATAAAGTATCCCAGGTTCCCGGCC | 8 |
| wt-161520-161640 | 1342 | TGGAATATCTGGTTGTTGCCGTTGACCCCCGTGTACTTGTTGATGGTCACTGGCAGCGTGACAACCGGACGGGCCTTGCAGACCTGGCTAAGACAGTCTGTGGCCGCGCAGACCACCGTG | 4 |
| wt-161640-161760 | 1343 | GTCGCAGTAAGGGAGGAGGTGGCCTCCGCGTAGGCCGCTGCCGACTCCACCGCCCGCGTGCCCAGTACGTGGGGGTAGTCACGGGCGGGCACCGACTGCGTCCTCGGCACCAGTCCCTGA | 8 |
| wt-161760-161880 | 1344 | ATCAGGCTGATGTAGAACTGGGTCTGGCCGCACGCCTTCAGGATGGCGTTGTTGAGCCTCTGCTTGGCGTAAGTGACCAGGTTGCCAGGCACCACATCTATGACGTTGCTCTCTTCGTGG | 4 |
| wt-161880-162000 | 1345 | GCCCGGGAGCCCCCGTCCACAAAGAGGGCCAGGTCAGAGTACTCCTCCGCGCTGGCCCCGCTGGGGACAGGGACCGAGCGCCGCCTGGAAAAGTTGTGCCACAGGTACAGGCTTGAGAGC | 8 |
| wt-162000-162120 | 1346 | TTAGTGTCCGGGAATAGGGTCTTGTGGTAGGTGTTGAGGAATTTCATGTAGGGCCCGTTGATGATGTAGTTCTCCCTCCTGGTAGTGGACTTGATGAAGCTGTTCTGGAGGGCGGCATTC | 4 |
| wt-162120-162240 | 1347 | TCCCCCGTGAAGACCACCCTGTTCTTGATCTTGATGTTCCTGGGGCACAGCATCAGCACCTTGGACATGCGCACAGGCAGCCGCCGGCCGTACACCCGGCCCTGCAGGGCCGCGTCCAGG | 8 |
| wt-162240-162360 | 1348 | TCTGGCAGGTCGCAGGTGGGCTCCCCATGCACCACCTTGGCCTCCTTGGCCGTGAGGACCCCCTTGTCGATGGCCAGGCTCCTAAAGTTGGTGCACAGCGTCTGGTAGTGACCCTTTAGC | 4 |
| wt-162360-162480 | 1349 | CACTCTGGGGGGCTCTGGCCAAGCCCGGGGTTGTCATTCTCATAGCACATACAGATGGGCAGGGAGATGTCCTGCAGGATGGTCAGCAGTGAGCGGTAAAACAGCTGGGTGAAGATGGGG | 4 |
| wt-162480-162600 | 1350 | CAGGCGGGCTGCGCAAAGGGGTTGCACGAGTACTGCATCACGTGGTAGCAGCTCTTGACCAGGTCCTTGTAGGTGATGTTGTTCTTGGCCATGCTGTTCATAAACTGGACCACTTCGGCG | 4 |
| wt-162600-162720 | 1351 | TCCACCGCCGCATCCACGTCCTTGAACATCTTGACAAAGTCACGCGGGCCATGGGGCTCCTTCTCTAGCTTTCCTTCAGCGTCTATGCCCAGCCGAGACAGCCGCTCCAGCAGGTTCTGG | 4 |
| wt-162720-162840 | 1352 | TTCAGCTGCCAGTAGGTGTAGCGGGGCTCGTCGTCCGGCCGCTGCCCGTCGTCCTCCTTATCGATGAAGTTGAGAAAGTTGCCCAAAAAGTCCGTCTCGTTGTAGGAGCCCGAGGCCCCC | 4 |
| wt-162840-162960 | 1353 | GAGATCACATAGGGGTCCCTCCGCTGCGTGGACATGACGGGGGGGAAGCGGTCCCTCAGCCTAAAGAAGAGCGTGTTCAGGCACACGGCCGGGGCCCGGCCCTCGCAGAGCGAGCACATG | 8 |
| wt-162960-163080 | 1354 | GGACTGGCGGCCGCCCCCGCCACGTAGCTGCCCGTCTCCGGCACCGGGGTCAGAGAGCTCTTCTGTCCCTGGCAAAACTGCAGGTAGTAGGCATAGCGGGCAAGAAGGTTGGGCGAGAAG | 8 |
| wt-163080-163200 | 1355 | GAGGCCGCATAGACCAGGTGCTCCACAGCGTAGTTTCCCGGACCGTTGGTTCCGGTCACGTCTGGCCCACCCCAGCCCGAGAAGCAGGGTCGGCGGCAGGGGTCCCAGGTCCCCTCCTGC | 8 |
| wt-163200-163320 | 1356 | AGGGTCCCCAGGCCGTGGGTCATGTAGAAACTGTTAAAGAGACTCTCCTTGCCCTGACCGGTTGACTTCGAGACCCCCGAGACGTAGAGGACGGAATTGGTGGCAAAGATCTGCGTGGAC | 4 |
| wt-163560-163680 | 1359 | ACCTTGGCCACCTTGGCCTGGTCCTGCAGGCTCTGCTTCTCCAGCAGCTCCACCAGCTTGCCCACCCGTCGGACGCGCAGCGCCTGCGCCAGCCCGGTGTACAGCGCCTCGTGCATGCAG | 8 |
| wt-163680-163800 | 1360 | CGGCTGAGGTCCGAGTTGTAAAACTGGCGGAGCTGGGGCACGCCCTCTGGGAACACCTCCTTGTCGTAGAGCGGGACCCTAACGCTCGCAGACTGCCCCACCGCTACCTCCTGTTTTAAC | 4 |
| wt-163800-163920 | 1361 | GATGGAATGGCCACCAGGTTTCCGCTGTAGAGTCGCTCCTTGAAGGCCTCGGTTATTGCCACCGCCCCCAGGTAGGCAGAGGGATCTAGCCCTTCGGGGAAGAAGTCCCCCGGCTTGGAG | 4 |

| | | | |
|---|---|---|---|
| wt-163920-164040 | 1362 | CTTTCCCTCGGTAGGGCGCTGTAGGCGTCGTACCCAAACACCTCCCTGGTCTCGCCACAGAGGGCCTCGAGACCCGGCCCCTCAAAGATGGGGGGAACCATATGGGCATTGTGGAACACG | 4 |
| wt-164040-164160 | 1363 | TAGATGTCCCTGTGATAGGAGGTAGCGCGTAGGAGCCCGCAGTTGGGGTCGGGCCTCCTGTGCAGAGCCTTGACATTGATGCTGAAGCCCGGCTCCACGGTGATGCCGCAAAGGAGCGGC | 4 |
| wt-164160-164280 | 1364 | ACCGTCAGGCACCTGTGGCCCGCGTAGCCGGTCCCCAGTGTGGCCACCTCCCTAAGAGGGTAGGTGGCCAGGGGGTAAAAGTAGATGTAGCCGCACGGACCCGGCTGGCTCTGGCTGCCC | 8 |
| wt-164280-164400 | 1365 | AGATTATCCTCGCTAGTCTGTGCACCCTGCATGATGCCCAAGGTATCGCCCCGGCCTCCCAGTCCCACATTAAATGTTACACTTTACTCATCACGCAACACCCACTGTTTATTCATTTAC | 4 |
| wt-164400-164520 | 1366 | AAAGATTTCAGGAAGTCAGTCAGGCTGGCCAGGGCCCACGTCACGGGGAACTGACGTCTCAGCGATCTTGGCATGCCGCCCAGCCTCGCAAACCAGAGTCTGCGATAGAGGGCCAGGTAG | 4 |
| wt-164520-164640 | 1367 | TGGGCGATTGCCCCCAGCACGAAGGCGGCGCTCTTGTGGTCATCCAGGTAGTTTCGCACCGCAAACACCACTGTGTAGCACAGCACCACCCTGAGCCGCGACCAGTAGTCGTAGTGGTCG | 4 |
| wt-164640-164760 | 1368 | TTGTACACTGCGCGCAGGACGCTGATGATGAGCCGTACGTGCGTGTCTTTGCCCCCGATGTCGGCTGTCCTGCAGGCCAGCTCCGCGTACAGCTTCCTATCCTTCCTCAGGGAGGCCTTG | 4 |
| wt-164760-164880 | 1369 | ATGAGCCGGCAGAGGACCAGGGCTGGCAAAGGCAGGTCTTTCTCATCCCGGGTGAACACCGCGTACATGGCCCTGAACATGAGGTAGCTGGACTCAGCCACCTTGTCGTCCGGCGGCGAG | 4 |
| wt-164880-165000 | 1370 | GGCGCGACCCACGCCTCGACCGGGGTCCTCACAAACACAGAATCTGTAGACTTGGCTGGCCTCATGGTCTCGTCAGGCCAGCTCACGGGCTTCAGGCTTATATGATAAAATGGGCGTGGC | 4 |
| wt-165000-165120 | 1371 | AGAATAGTATAAGACGCGAGGCCTGGGTGAGGAGAGTCCAGAGCAATGGCCAGGTTCATCGCTCAGCTCCTCCTGTTGGCCTCCTGTGTGGCCGCCGGCCAGGCTGTCACCGCTTTCTTG | 4 |
| wt-165120-165240 | 1372 | GGTGAGCGAGTCACCCTGACCTCCTACTGGAGGAGGGTGAGCCTCGGTCCAGAGATTGAGGTCAGCTGGTTTAAACTGGGCCCAGGAGAGGAGCAGGTGCTTATTGGGCGCATGCACCAC | 4 |
| wt-165240-165360 | 1373 | GATGTCATCTTTATAGAGTGGCCTTTCAGGGGCTTCTTTGATATCCACAGAAGTGCCAACACCTTCTTTTTAGTAGTCACCGCTGCCAACATCTCCCATGACGGCAACTACCTGTGCCGC | 4 |
| wt-165360-165480 | 1374 | ATGAAACTGGGCGAGACCGAGGTCACCAAGCAGGAACACCTGAGCGTGGTGAAGCCTCTAACGCTGTCTGTCCACTCCGAAAGGTCTCAGTTCCCAGACTTCTCTGTCCTTACTGTGACA | 4 |
| wt-165480-165600 | 1375 | TGCACCGTGAATGCATTTCCCCATCCCCACGTCCAGTGGCTCATGCCCGAGGGCGTGGAGCCCGCACCAACTGCGGCAAATGGCGGTGTTATGAAGGAAAAGGATGGGAGCCTCTCTGTT | 4 |
| wt-165600-165720 | 1376 | GCTGTTGACCTGTCACTTCCCAAGCCCTGGCACCTGCCAGTGACCTGCGTTGGGAAAAATGACAAGGAGGAAGCCCACGGGGTTTATGTTTCTGGATACTTGTCGCAATAAACGCACTTG | 4 |
| wt-165720-165840 | 1377 | CCTATTTCACCTTGTTTTAGTGTGGCATTGGGGGGGTGGCATTGCGGGTGGATAGCCTCGCGACTCGTGGGAAAATGGGCGGAAGGGCACCGTGGGAAAATAGTTCCAGGTGACAGCAGC | 4 |
| wt-165840-165960 | 1378 | AGTGTGTGAAGATTGTCACAGCTGCTGGTTTGGAGAAAACGGGGGTGGGCGGTGATCAGGGAGAACAATTCCCCGGGGACACCTGCACGAGACCCCTGGGCTCTCAGGAACTCCGCCCAG | 4 |
| wt-165960-166080 | 1379 | GTCTTGCCAATTGGGGTGATCCTGTAGCGCCGCGGTTTCAGCATCACAGGTTATTTTGCCTGAAGCTTGCTGGGGCGTAAATCCCTCTCGCCTTGTTTCTCAGAGAGCATTTCAGGCCGG | 4 |
| wt-166080-166200 | 1380 | TTTTGCAGTCGCTGCTGCAGCTATGGGGTCCCTAGAAATGGTGCCAATGGGCGCGGGTCCCCCTAGCCCCGGCGGGGATCCGGATGGGTACGATGGCGGAAACAACTCCCAATATCCATC | 4 |
| wt-166200-166320 | 1381 | TGCTTCTGGCTCTTCTGGGAACACCCCCACCCCACCGAACGATGAGGAACGTGAATCTAATGAAGAGCCCCCACCGCCTTATGAGGACCCATATTGGGGCAATGGCGACCGTCACTCGGA | 4 |
| wt-166320-166440 | 1382 | CTATCAACCACTAGGAACCCAAGATCAAAGTCTGTACTTGGGATTGCAACACGACGGGAATGACGGGCTCCCTCCCCCTCCCTACTCTCCACGGGATGACTCATCTCAACACATATACGA | 4 |
| wt-166440-166560 | 1383 | AGAAGCGGGCAGAGGAAGGTAAGAGTGCCATCTATCTGTACTTTTATTTATTGCATCACAAGTCACATCAATAATAAGGGCGCCATCTAGCGGGAGATGTTATCCACACCATCCCAATTC | 4 |
| wt-166560-166680 | 1384 | ACATCTCAGGGACAACAGGTCAAAGTTCTTTGTTGACACCCCCAGCGCTGGCTCCAGGGGGTGGAAGCGTTGGATGCAGTCCTCCGCATCGGGGCGGACGCCTCCTCCCAACGCGTTTCT | 4 |
| wt-166680-166800 | 1385 | GCGGATCAGTCGCTGGCTGGTGGGCATCGGAGTCGGTGGGCGGTCCTCCACGGGGACACGCTCCTTCTTGGCCTTGTTCTTTGACCTTTTGGACATTCTTCTGAAGGAACGGCGGAGAGT | 4 |
| wt-166800-166920 | 1386 | AGCGTAGAATCCAGCCAGTGGTCTACCCGGTCGCATGGTGGCTTCTTAGATGAGGAGCAGGCATAAAAGTCCAAACAGGACACAGAGTACCACCAGGAGTAGTCTTAGTCTGCTGACGTC | 4 |
| wt-167040-167160 | 1388 | GCACAGCTGTTTTCGTGGACTTTTATACAGTAAGGACAAGGAAAGAAGGCCAGAGGAATGTGGAAAGATGAGCGAGGACAGGTGTGGAGGTTTTGGGCTAGCTCTTAGTTTCTGGGTGTG | 4 |
| wt-167160-167280 | 1389 | AGAGAGGGATTAAAGTGCTTATGCGCAAAGAATGTGTCAACAACAGGTGTTCCTGCCTCTGCTGGCATGAGTTAGGTGTGGCTTGGGCTGAATCCAAATGTGTATTGGCACAAGATGGAA | 4 |
| wt-167280-167400 | 1390 | AGCAAAGTTGCTGGAGTTACTGGGTGGGAGACAGGGATGTATGTGGTCCCCCGCTGGTATGCCAGTACCCTGTGGAAGTAAGGGGCCTCATCTGCCTGGTAGTTGTGTTGTGCAGAGGTC | 4 |
| wt-167400-167520 | 1391 | TGATGTGTGTAGGAGGGGTGGGTTCAACGCAGGGGCGTTGGTGGCGGAGTCTGGCAACGCCCGGGTCCTTGCTACCTGTGTGGTGTGTTAAGGGCTGGGTAAAGGTGTCTGCCAATTCTC | 4 |
| wt-167520-167640 | 1392 | GCATGTCCTCCTTTCCCCTTGTTTTGAAATAGAATATGAATGTGGCTTTTCAGCCTAGACAGACAGTGTGGCTAAGGGAGTGTGTGCCAGTTAAGGTGATTAGCTAAGGCATTCCCAGTA | 4 |
| wt-167640-167760 | 1393 | AATGGAGGGAGAGTCAGTCAGGCAAGCCTATGACATGGTAATGCCTAGAAGTAAAGAAAGGTTAGTCATAGTAGCTTAGCTGAACTGGGCCGTGGGGGTCGTCATCATCTCCACCGGAAC | 4 |
| wt-167760-167880 | 1394 | CAGAAGAACCCAAAAGCAGCGTAGGAAGGTGTGGATCACCGCCGCCATGGCCGGAATCATGACTATGACCGCCGCCTCCGTCTGTCATCAAAGGCGGGCCCTGGTCACCTCCTTTGTTTT | 4 |
| wt-167880-168000 | 1395 | CAACCTCTTCCGTCAATTGTGGAGGGCCTCCATCATTTCCAGCAGAGTCGCTAGGGCTATGAGGCAGCGGGTCATGTGGGCCATTGTCATCAGTGTTGTCAGGGTCCTGTGGGCCATTGT | 4 |
| wt-168000-168120 | 1396 | CATCAGTGTTGTCAGGGTCCTGAGGCAGCGGGTCATGTGGGCCATTGTCATCAGTGTTGTCAGGGTCCTGTGGGCCATTGTCATCAGTGTTGTCAGGGTCCTGTGGGCCATTGTCAGGAC | 4 |
| wt-168120-168240 | 1397 | CACCTCCAGGTGCGCCTAGGTTTTGAGAGCAGAGTGGGGGTCCGTCGCCGGCTCCACTCACGAGCAGGTGGTGTCTGCCCTCGTTGGAGTTAGAGTCAGATTCATGGCCAGAATCATCGG | 4 |
| wt-168240-168360 | 1398 | TAGCTTGTTGAGGGTGCGGGAGGGAGTCATCGTGGTGGTGTTCATCACTGTGTCGTTGTCCATGGTAATACATCCAGATTAAAATCGCCAGAAACAGGAGGAGCCAAAGGAGATCAACCA | 4 |
| wt-168360-168480 | 1399 | ATAGAGTCCACCAGTTTTGTTGTAGATAGAGAGCAATAATGAGCAGGATGAGGTCTAGGAAGAAGGCTAGGAAGAAGGCCAAAAGCTGCCAGATGGTGGCACCAAGTCGCCAGAGCATCT | 4 |
| wt-168480-168600 | 1400 | CCAATAAGTAGATCCAGATACCTAAGACTGCGTTGAAAAAAGAGTGTTAGGGTTGGAAAAGTGGGGGTGTGGTAAATAATTCCTAGGGAATGTTAGATCTTACCAAGTAAGCACCCGAAG | 4 |
| wt-168600-168720 | 1401 | ATGAACAGCACAATTCCAAGGAACAATGCCTGTCCGTGCAAATTCCAGAGAGCGATGAGCAGGAGGGTGACTGGGGAAAGAGGAGAAAGTGCGTTAGAGAAGGAAGAGTAAGGGAAAGGG | 4 |
| wt-168720-168840 | 1402 | GGTGTGGGGCAAAGGGTGTAATACTTACTCATCAGTAGGAGTATACAAAGGGCTCCAAGTGGACAGAGAAGGTCTCTTCTGAAGATAAAGATGATCAAAATTATAATTATAAGCATGAGA | 4 |
| wt-168840-168960 | 1403 | GCAAAGGAATAGAGGACAAGGAGGGCTCCTCCAGTCCAGTCACTCATAACGATGTACAGCCAAAACAGTAGCGCCAAGAGGAGGAGAAGGAGAGCAAGGCCTAGGGAAGAGGAGAGGGGG | 4 |
| wt-168960-169080 | 1404 | GGTCCTCGAGGGGGCCGTCGCGGGCCCGGTGGGCCCCTCTCAAGGTCGTGTTCCATCCTCAGGGCAGTGTGTCAGGAGCAAGGCAGTTGAGGAAAGAAGGGGGCAGAGCAGTGTGAGAGG | 8 |
| wt-169080-169200 | 1405 | CTTATGTAGGGCGGCTACGTCAGAGTAACGCGTGTTTCTTGGGATGTAGGCCCGGGGGGATTTGCGGGGTCTGCCGGAGGCAGTACGGGTACAGATTTCCCGAAAGCGGCGGTGTGTGTG | 4 |
| wt-169200-169320 | 1406 | TGCATGTAAGCGTAGAAAGGGGAAGTAGAAAGCGTGTGTTTGTGTTAGAAAAGCGGGTCCCCGGGGGGCAAGCTGTGGGAATGCGGTGGCCAAGTGCAACAGGAAATGGAAAGGCAGTGC | 4 |
| wt-169320-169440 | 1407 | GGCAATCAGAAGGGGGAGTGCGTAGTGTTGTGGGAAGCGGCAGTGTAATCTGCACAAAGAGGCGCGGGGCGCGCAACGTTGGGAGGTCGTTGGCGGCAGGCGGGAGGCCGTGCTTTAGGG | 4 |
| wt-169440-169560 | 1408 | GGGTTCAGGTGAGGCAAGGCTGTGGGGTAACCGTAGGGGAGGCGGGTGAGGCGGCTAAGAGGGCTAAGGGTCGGCGGGTGACGAAGCAGCAGACGGCGGATATGGGAATTTCAGAATGAG | 4 |
| wt-169560-169680 | 1409 | GTGGCGGATTCAGGCGAAAAGGGTGTGGGCTGTGCGAGTGTCATGAGGCAGGCGCGGAAAGTCGCTGCGGCTTGCTGGGGCATGGGGGGCCGCGCATTCCTGGAAAAAGTGGAGGGGGCG | 8 |
| wt-169680-169800 | 1410 | TGGCCTTCCCCCGCGGCCCCCCAGCCCCCCCGCACAGAGCGGCGCTACGGCGGGCGGGCGGCGGGGGGTCGGGGTCCGCGGGCTCCGGGGGCTGCGGGCGGTGGATGGCGGCGGACGTTC | 12 |
| wt-169800-169920 | 1411 | CGGGGATCGGGGGGGTCGGGGGGCGCCGCGCGGGCGCAGCCATGCGTGACCGTGATGAGGGGGCAGGGTCGCAGGGGGTGTGTCTGGTGGGGGCGGGAGCGGGGGGCGGCGCGGGAGCCT | 12 |
| wt-169920-170040 | 1412 | GCACGCCGTTGGAGGGTAGAATGACAGGGGGCGGGGACAGAGAGGCGGTCGCGCCCCCGGCCGCGCCAGCCAAGCCCCCAAGGGGGGCGGGGAGCGGGCAATGGAGCGTGACGAAGGGCC | 8 |
| wt-170040-170160 | 1413 | CCAGGGCTGACCCCGGCAAACGTGACCCGGGGCTCCGGGGTGACCCAGCCAAGCGTGACCAAGGGGCCCGTGGGTGACACAGGCAACCCTGACAAAGGCCCCCCAGGAAAGACCCCCGGG | 8 |

| | | | |
|---|---|---|---|
| wt-170160-170280 | 1414 | GGGCATCGGGGGGTGGGGCATGGGGGGCCGCGCATTCCTGGAAAAAGTGGAGGGGGCGTGGCCTTCCCCCGCGGCCCCCCAGCCCCCCCGCACAGAGCGGCGCTACGGCGGGCGGGCGGC | 12 |
| wt-170280-170400 | 1415 | GGGGGGTCGGGGTCCGCGGGCTCCGGGGGCTGCGGGCGGTGGATGGCGGCGGACGTTCCGGGGATCGGGGGGGTCGGGGGGCGCCGCGCGGGCGCAGCCATGCGTGACCGTGATGAGGGG | 12 |
| wt-170400-170520 | 1416 | GCAGGGTCGCAGGGGGTGTGTCTGGTGGGGGCGGGAGCGGGGGGCGGCGCGGGAGCCTGCACGCCGTTGGAGGGTAGAATGACAGGGGGCGGGGACAGAGAGGCGGTCGCGCCCCCGGCC | 12 |
| wt-170520-170640 | 1417 | GCGCCAGCCAAGCCCCCAAGGGGGGGCGGGGAGCGGGCAATGGAGCGTGACGAAGGGCCCCAGGGCTGACCCCGGCAAACGTGACCCGGGGCTCCGGGGTGACCCAGCCAAGCGTGACCAA | 8 |
| wt-170640-170760 | 1418 | GGGGCCCGTGGGTGACACAGGCAACCCTGACAAAGGCCCCCCAGGAAAGACCCCCGTGGGGCATGGGGGGCCGCGCATTCCTGGAAAAAGTGGAGGGGGCGTGGCCTTCCCCCGCGGCCC | 8 |
| wt-170760-170880 | 1419 | CCCAGCCCCCCCGCACAGAGCGGCGCTACGGCGGGCGGGCGGCGGGGGGTCGGGGTCCGCGGGCTCCGGGGGCTGCGGGCGGTGGATGGCGGCGGACGTTCCGGGGATCGGGGGGGTCGG | 12 |
| wt-170880-171000 | 1420 | GGGGCGCCGCGCGGGCGCAGCCATGCGTGACCGTGATGAGGGGGCAGGGTCGCAGGGGGTGTGTCTGGTGGGGGCGGGAGCGGGGGGCGGCGCGGGAGCCTGCACGCCGTTGGAGGGTAG | 12 |
| wt-171000-171120 | 1421 | AATGACAGGGGGCGGGGACAGAGAGGCGGTCGCGCCCCCGGCCGCGCCAGCCAAGCCCCCAAGGGGGGGCGGGGAGCGGGCAATGGAGCGTGACGAAGGGCCCCAGGGCTGACCCCGGCAA | 12 |
| wt-171120-171240 | 1422 | ACGTGACCCGGGGCTCCGGGGTGACCCAGCCAAGCGTGACCAAGGGGCCCGTGGGTGACACAGGCAACCCTGACAAAGGCCCCCCAGGAAAGACCCCCGGGGGGCATCGGGGGGTGGGGC | 8 |
| wt-171240-171360 | 1423 | ATGGGGGGCCGCGCATTCCTGGAAAAAGTGGAGGGGGCGTGGCCTTCCCCCGCGGCCCCCCAGCCCCCCCGCACAGAGCGGCGCTACGGCGGGCGGGCGGCGGGGGGTCGGGGTCCGCGG | 12 |
| wt-171360-171480 | 1424 | GCTCCGGGGGCTGCGGGCGGTGGATGGCGGCGGACGTTCCGGGGATCGGGGGGGTCGGGGGGCGCCGCGCGGGCGCAGCCATGCGTGACCGTGATGAGGGGGCAGGGTCGCAGGGGGTGT | 12 |
| wt-171480-171600 | 1425 | GTCTGGTGGGGGCGGGAGCGGGGGGCGGCGCGGGAGCCTGCACGCCGTTGGAGGGTAGAATGACAGGGGGCGGGGACAGAGAGGCGGTCGCGCCCCCGGCCGCGCCAGCCAAGCCCCCAA | 12 |
| wt-171600-171720 | 1426 | GGGGGGCGGGGAGCGGGCAATGGAGCGTGACGAAGGGCCCCAGGGCTGACCCCGGCAAACGTGACCCGGGGCTCCGGGGTGACCCAGCCAAGCGTGACCAAGGGGCCCGTGGGTGACACA | 8 |
| wt-171703-171823 | 1427 | GGCCCGTGGGTGACACAGGCAACCCTGACAAAGGCCCCCCAGGAAAGACCCCCGGGGGGCATCGGGGGGGGTGTTGGCGGGGGCATGGGGGGGTCGGATTTCGCCCTTATTGCCCTGTTT | 8 |
| wt-30-150 | 1428 | TTTCTTCTTGCCCGTTCTCTTTCTTAGTATGAATCCAGTATGCCTGCCTGTAATTGTTGCGCCCTACCTCTTTTGGCTGGCGGCTATTGCCGCCTCGTGTTTCACGGCCTCAGTTAGTAC | 4 |
| wt-150-270 | 1429 | CGTTGTGACCGCCACCGGCTTGGCCCTCTCACTTCTACTCTTGGCAGCAGTGGCCAGCTCATATGCCGCTGCACAAAGGAAACTGCTGACACCGGTGACAGTGCTTACTGCGGTTGTCAC | 4 |
| wt-270-390 | 1430 | TTGTGAGTACACACGCACCATTTACAATGCATGATGTTCGTGAGATTGATCTGTCTCTAACAGTTCACTTCCTCTGCTTTTCTCCTCAGTCTTTGCAATTTGCCTAACATGGAGGATTGA | 4 |
| wt-390-510 | 1431 | GGACCCACCTTTTAATTCTCTTCTGTTTGCATTGCTGGCCGCAGCTGGCGGACTACAAGGCATTTACGGTTAGTGTGCCTCTGTTATGAAATGCAGGTTTGACTTCATATGTATGCCTTG | 4 |
| wt-510-630 | 1432 | GCATGACGTCAACTTTACTTTTATTTCAGTTCTGGTGATGCTTGTGCTCCTGATACTAGCGTACAGAAGGAGATGGCGCCGTTTGACTGTTTGTGGCGGCATCATGTTTTTGGCATGTGT | 4 |
| wt-630-750 | 1433 | ACTTGTCCTCATCGTCGACGCTGTTTTGCAGCTGAGTCCCCTCCTTGGAGCTGTAACTGTGGTTTCCATGACGCTGCTGCTACTGGCTTTCGTCCTCTGGCTCTCTTCGCCAGGGGGCCT | 4 |
| wt-750-870 | 1434 | AGGTACTCTTGGTGCAGCCCTTTTAACATTGGCAGCAGGTAAGCCACACGTGTGACATTGCTTGCCTTTTTGCCACATGTTTTCTGGACACAGGACTAACCATGCCATCTCTGATTATAG | 4 |
| wt-870-990 | 1435 | CTCTGGCACTGCTAGCGTCACTGATTTTGGGCACACTTAACTTGACTACAATGTTCCTTCTCATGCTCCTATGGACACTTGGTAAGTTTTCCCTTCCTTTAACTCATTACTTGTTCTTTT | 4 |
| wt-990-1110 | 1436 | GTAATCGCAGCTCTAACTTGGCATCTCTTTTACAGTGGTTCTCCTGATTTGCTCTTCGTGCTCTTCATGTCCACTGAGCAAGATCCTTCTGGCACGACTGTTCCTATATGCTCTCGCACT | 4 |
| wt-1110-1230 | 1437 | CTTGTTGCTAGCCTCCGCGCTAATCGCTGGTGGCAGTATTTTGCAAACAAACTTCAAGAGTTTAAGCAGCACTGAATTTATACCCAGTGAGTATCTATTTGTTACTCCTGTTTAGTTGAA | 4 |
| wt-1230-1350 | 1438 | GAAAACAAGCTATTGGATTGTAACACACATTTTACGCTTTGTTCCTTAGATTTGTTCTGCATGTTATTACTGATTGTCGCTGGCATACTCTTCATTCTTGCTATCCTGACCGAATGGGGC | 4 |
| wt-1350-1470 | 1439 | AGTGGAAATAGAACATACGGTCCAGTTTTTATGTGCCTCGGTGGCCTGCTCACCATGGTAGCCGGCGCTGTGTGGCTGACGGTGATGTCTAACACGCTTTTGTCTGCCTGGATTCTTACA | 4 |
| wt-1470-1590 | 1440 | GCAGGATTCCTGATTTTCCTCATTGGTAAGTGTGACACCAACAGGTGTTGCCTTGTTATGTCACCGTTCTGACACATGACTTACATGGGTTTGGCTTTTGTAGGCTTTGCCCTCTTTGGG | 4 |
| wt-1590-1710 | 1441 | GTCATTAGATGCTGCCGCTACTGCTGCTACTACTGCCTTACACTGGAAAGTGAGGAGCGCCCACCGACCCCATATCGCAACACTGTATAAAGGTAAGTATTATTAAATTTTAGAGACACT | 4 |
| wt-1710-1830 | 1442 | ATCACGTGTAACTTGACGTGCAAGGATGGAAGAGAGGGGCAGGGAAACGCAAATGCCGGTTGCCCGGTATGGGGGCCCGTTTATTATGGTAAGGCTCTTCGGGCAAGATGGAGAGGCAAA | 4 |
| wt-1830-1950 | 1443 | CATACAGGAGGAAAGGCTATATGAGCTACTCTCTGACCCACGCTCCGCGCTCGGCCTAGACCCGGGGCCCCTGATTGCTGAGAACCTGCTGCTAGTGGCGCTGCGTGGCACCAACAACGA | 4 |
| wt-1950-2070 | 1444 | TCCCAGGCCTCAGCGTCAGGAGAGGGCCAGAGAACTGGCCCTCGTTGGCATTCTACTAGGAAACGGCGAGCAGGGTGAACACTTGGGCACGGAGAGTGCCCTGGAGGCCTCAGGCAACAA | 4 |
| wt-2070-2190 | 1445 | CTATGTGTATGCCTACGGACCAGACTGGATGGCAAGGCCTTCCACATGGTCCGCGGAAATCCAGCAATTCCTGCGACTCCTGGGCGCCACGTACGTGCTTCGCGTGGAGATGGGCAGGCA | 4 |
| wt-2190-2310 | 1446 | GTTTGGCTTCGAGGTGCATAGAAGCCGGCCCTCCTTCCGTCAGTTCCAGGCCATCAATCACCTTGTCCTGTTTGACAACGCCCTTCGCAAGTACGATTCCGGCCAGGTGGCGGCGGGCTT | 4 |
| wt-2310-2430 | 1447 | CCAGAGGGCCCTTCTGGTGGCCGGGCCAGAGACCGCTGACACGAGGCCGGACCTCCGCAAGCTGAATGAGTGGGTGTTTGGTGGCAGGGCTGCTGGTGGCAGACAGCTGGCCGACGAGCT | 8 |
| wt-2430-2550 | 1448 | AAAGATCGTGTCCGCGCTGCGAGACACTTACTCGGGCCACTTGGTCCTTCAGCCCACGGAGACCCTTGACACATGGAAGGTGTTGAGCAGGGACACACGAACCGCTCATAGTTTGGAGCA | 4 |
| wt-2550-2670 | 1449 | CGGATTCATTCATGCCGCGGGGACCATCCAGGCCAACTGCCCACAGCTGTTTATGAGACGCCAGCACCCCGGCCTCTTTCCCTTCGTTAATGCAATAGCATCATCGCTGGGCTGGTACTA | 4 |
| wt-2670-2790 | 1450 | CCAGACCGCCACCGGCCCCGGAGCAGATGCCAGGGCGGCGGCCCGGCGCCAACAGGCCTTTCAGACCAGGGCGGCGGCTGAATGCCATGCCAAAAGCGGGGTGCCGGTCGTGGCCGGCTT | 8 |
| wt-2790-2910 | 1451 | CTACAGGACCATCAACGCCACGCTCAAGGGAGGAGAGGGCCTACAGCCCACTATGTTTAACGGGGAGCTGGGGGCCATCAAGCACCAGGCACTTGACACTGTGAGGTATGACTACGGCCA | 4 |
| wt-2910-3030 | 1452 | CTATCTCATAATGTTGGGGCCATTCCAGCCATGGAGCGGACTGACGGCCCCTCCGTGCCCCTACGCCGAAAGTTCATGGGCACAGGCGGCCGTGCAGACGGCCCTCGAGCTGTTCTCGGC | 4 |
| wt-3030-3150 | 1453 | CCTGTACCCGGCCCCGTGCATCTCGGGCTACGCGCGCCCCCCGGGCCCCAGTGCTGTGATCGAGCATCTGGGGTCCCTAGTTCCAAAGGGGGGTCTGCTGTTGTTTCTGTCTCACCTACC | 8 |
| wt-3150-3270 | 1454 | GGATGATGTTAAGGACGGGCTCGGAGAAATGGGGCCGGCCAGGGCCACGGGACCTGGAATGCAGCAGTTTGTCAGCAGCTACTTCCTCAACCCCGCCTGTTCCAACGTCTTCATTACAGT | 4 |
| wt-3270-3390 | 1455 | GAGGCAGCGAGGGGAGAAGATCAACGGCCGTACCGTCCTCCAAGCGCTCGGACGCGCATGCGATATGGCAGGCTGCCAGCACTATGTGCTGGGCTCCACGGTTCCCCTCGGTGGACTCAA | 4 |
| wt-3390-3510 | 1456 | CTTTGTCAACGACCTGGCGTCCCCGGTTTCCACCGCCGAGATGATGGATGATTTCTCTCCCTTCTTCACCGTGGAGTTTCCCCCGATTCAAGAGGAGGGCGCAAGTTCTCCGGTACCCTT | 4 |
| wt-3510-3630 | 1457 | AGATGTGGACGAGAGCATGGACATCTCTCCGTCTTACGAGTTGCCCTGGCTCTCGCTGGAGTCATGCCTCACAAGCATCCTGTCACACCCCACCGTGGGAAGCAAGGAGCACTTGGTCAG | 4 |
| wt-3630-3750 | 1458 | GCACACGGACAGGGTCAGCGGAGGACGCGTGGCACAGCAGCCCGGGGTAGGTCCCCTGGACCTGCCGCTGGCGGACTACGCCTTCGTTGCCCACAGTCAGGTCTGGACCAGGCCCGGTGG | 8 |
| wt-3750-3870 | 1459 | GGCTCCTCCCTTGCCCTATCGTACCTGGGATCGAATGACAGAGAAGCTGCTTGTCTCCGCAAAACCCGGCGGAGAGAACGTTAAGGTTTCAGGTACCGTGATTACATTGGGAGAACAGGG | 4 |
| wt-3870-3990 | 1460 | GTACAAAGTGTCGTTGGATCTGAGGGAGGGAACCAGGCTGGCAATGGCTGAGGCGCTGCTGAACGCAGCATGTGCCCCAATCTTGGATCCGGAAGACGTCTTGCTCACCCTGCATCTACA | 4 |
| wt-3990-4110 | 1461 | CCTGGATCCGCGCCGGGCAGACAACTCGGCCGTGATGGAGGCTATGACGGCGGCGAGTGACTACGCGCGTGGCCTGGGCGTGAAGCTGACCTTTGGCTCGGCCTCCTGCCCCGAGACCGG | 8 |
| wt-4110-4230 | 1462 | CTCGTCCGCCTCCAACTTCATGACCGTGGTGGCCTCTGTCTCCGCCCCAGGGGAATTCTCGGGTCCTCTGATCACGCCAGTGCTTCAGAAGACGGGCAGTCTCCTGATTGCGGTGCGTTG | 4 |
| wt-4230-4350 | 1463 | CGGGGATGGCAAGATCCAGGGAGGGTCGCTGTTTGAGCAGCTCTTTAGCGACGTGGCCACGACCCCACGGGCACCCGAGGCGTTGTCTCTGAAGAATCTCTTCCGGGCAGTCCAGCAGCT | 4 |
| wt-4350-4470 | 1464 | GGTCAAGAGCGGCATCGTGCTGTCAGGGCATGACATCAGCGACGGGGGCCTGGTGACCTGCCTGGTGGAGATGGCCCTGGCCGGGCAGCGGGGAGTGACCATCACTATGCCGGTGGCCTC | 8 |

| | | | |
|---|---|---|---|
| wt-4470-4590 | 1465 | CGACTACCTCCCGGAGATGTTTGCAGAGCACCCCGGCCTGGTGTTTGAGGTGGAGGAGCGCAGCGTGGGTGAGGTGCTGCAGACCCTGCGCTCCATGAACATGTACCCGGCAGTCCTCGG | 4 |
| wt-4590-4710 | 1466 | TCGAGTGGGCGAGCAAGGTCCAGATCAAATGTTTGAGGTGCAGCACGGCCCAGAGACGGTGTTGCGCCAGTCGCTGCGCCTGCTGCTGGGAACCTGGTCATCCTTTGCCAGCGAGCAGTA | 4 |
| wt-4710-4830 | 1467 | CGAGTGCCTGCGACCAGATCGGATTAACCGGTCCATGCACGTGTCCGACTACGGCTATAACGAAGCACTGGCAGTCTCCCCGTTGACAGGAAAGAATCTCAGCCCACGCCGGTTGGTGAC | 4 |
| wt-4830-4950 | 1468 | AGAGCCTGACCCACGATGTCAGGTGGCCGTGCTATGCGCCCCGGGCACCAGGGGCCATGAAAGCCTCCTGGCGGCCTTCACGAATGCCGGATGCCTGTGCCGACGGGTGTTCTTTCGCGA | 8 |
| wt-4950-5070 | 1469 | GGTTAGGGACAACACGTTCCTCGACAAGTACGTGGGTCTGGCCATCGGAGGAGTTCATGGGGCCAGGGACTCTGCCCTGGCAGGCCGTGCCACCGTGGCGCTGATTAATCGTTTCCCCGC | 4 |
| wt-5070-5190 | 1470 | CCTGCGTGACGCTATTCTAAAGTTCCTCAACAGGCCAGATACGTTCTCGGTGGCCTTGGGGGAGCTGGGGGTGCAAGTTTTGGCTGGCCTGGGGGCCGTGGGGTCAACAGATAATCCACC | 4 |
| wt-5190-5310 | 1471 | CGCCCCTGGCGTGGAAGTTAATGTCCAGAGATCACCTCTGATTCTGGCCCCCAACGCCTCTGGCATGTTTGAGTCCCGCTGGCTGAACATTAGCATCCCGGCGACCACCAGCTCTGTCAT | 4 |
| wt-5310-5430 | 1472 | GCTGCGTGGCCTCCGGGGCTGCGTCCTGCCTTGTTGGGTGCAAGGCTCGTGCCTGGGCCTGCAATTTACTAACCTCGGGATGCCATATGTTTTGCAGAATGCCCACCAGATCGCCTGCCA | 4 |
| wt-5430-5550 | 1473 | CTTCCACAGCAATGGCACGGATGCCTGGCGCTTTGCTATGAATTATCCAAGAAACCCCACGGAGCAGGGCAACATTGCAGGGCTCTGTTCACGCGATGGTCGTCATCTGGCTCTCCTGTG | 4 |
| wt-5550-5670 | 1474 | TGACCCCTCACTTTGTACAGACTTTTGGCAATGGGAGCACATTCCCCCCGCCTTTGGGCACCCCACGGGGTGCTCCCCCTGGACACTTATGTTTCAAGCAGCTCACCTATGGTCACTCAG | 4 |
| wt-5670-5790 | 1475 | GCACGGTCGCCCCTCCGAGTGACCAGTCACCTTCCAGACTATGCATACACTGAATTTAGCCTGATATTGTCCCCCTAGCCCCGGGCCCAGCCCTCCTCAGAAAACTCTGCATGGAGAAGC | 4 |
| wt-5790-5910 | 1476 | TGGACGTGAACCTCCCCCCCAGACCTGTGTGCTGTATTTACAAACACTACAATAAACCCAATGTGCAAATGTGGTTTGTATGGCTACTTTGTGTTCCTAAAAAATGCAACAATAGAAGTG | 4 |
| wt-5910-6030 | 1477 | GAAACCCTCAGTCACGGGACATTAACCTCAACCACAAAATGGGGGTTGGAGAAAGTAACCACATATACTGGAGATGATTCATGGGCTGGGGGTTCCCGGACAATACACCCATCTGGAGTT | 4 |
| wt-6030-6150 | 1478 | CAACCTAATTACATGGTAGATAAATTAAGAGTCCCTCCTCACCACTCGAAACTATGGCAGACATTCTATAAGATAACGAGGAGAGATGAGGTGAGGGCAGAGGACATTGGGCAGGTGTGG | 4 |
| wt-6150-6270 | 1479 | GCCACGGGGCAGCTGGCCATATCCCCCGCACTACAGAAGTGTAAGCAAAGTGAAGGGCTCGGAAGGCAGGCGGGGCCTAGCAATGTCACAGCTAAATGCCCACCAGGGCACACACTCAAG | 4 |
| wt-6270-6390 | 1480 | CGGGGTCTCGGAGCTCCTAGGTCAGACCACGAAAGGTCAGCCTGCAAGGTGGATGGCGTGTTTTCTGAGGTTATCCCCGCTACGTGCAGTGCTGGGTGATAGAGACCCTAGAATGTGTCG | 4 |
| wt-6390-6510 | 1481 | AAATGACCAAGCGTCCCCGCAGCGGGGCTCCCAACACGGGTTCCCAGAGAGGGTAAAAGAGGGGGCCATAAAGCCCAGGGTGTAAAACACCGACCGCGCCACCAGATGGCACACGTGGGG | 4 |
| wt-6510-6630 | 1482 | GAAATGAGGGTTAGCATAGGCAACCCCCGCCTACACACCAACTATAGCAAACCCCGCCCCGTCACGGTGACGTAGTCTGTCTTGAGGAGATGTAGACTTGTAGACACTGCAAAACCTCAG | 4 |
| wt-6630-6750 | 1483 | GACCTACGCTGCCCTAGAGGTTTTGCTAGGGAGGAGACGTGTGTGGCTGTAGCCACCCGTCCCGGGTACAAGTCCCGGGTGGTGAGGACGGTGTCTGTGGTTGTCTTCCCAGACTCTGCT | 4 |
| wt-6750-6870 | 1484 | TTCTGCCGTCTTCGGTCAAGTACCAGCTGGTGGTCCGCATGTTTTGATCCAAACTTTTGTTTTAGGATTTATGCATCCATTATCCCGCAGTTCCACCTAAACGGGGCTTAACGTTGCATC | 4 |
| wt-6870-6990 | 1485 | CCAGAAGATGCACGCTTAACCCCGCCTACAACCGTGACGTAGCTGTTTACCAGCATGTATAGAGTTACGGTTCGCTACATCAAACAGGACAGCCGTTGCCCTAGTGGTTTCGGACACACC | 4 |
| wt-6990-7110 | 1486 | GCCAACGCTCAGTGCGGTGCTACCGACCCGAGGTCAAGTCCCGGGGGAGGAGAAGAGAGGCTTCCCGCCTAGAGCATTTGCAAGTCAGGATTCTCTAATCCCTCTGGGAGAAGGGTATTC | 4 |
| wt-7110-7230 | 1487 | GGCTTGTCCGCTATTTTTTTGTGGCTAGTTTTGCACCCACAACATGTAAGGGCCCGCTACCCCTACAACACAAAACAAACTATCTCCCCTAACCATCCTTTTGCCAATCAATTCTGTGAC | 4 |
| wt-7230-7350 | 1488 | AGGGTTTCCTGGACACCCAGTCTTAGTTCAGGTAGACACCCAGTTATGCAGTGCCACCAATTCCAACCATTTTTAAACCTCCTGGAATTCTATCATTAAACGGCATGCAGGAAAAGGACA | 4 |
| wt-7350-7470 | 1489 | AGCAGCGAAAATTCACGCCCCCTTGGGAGGTGGCGGCATATGCAAAGGATAGCACTCCCACTCTACTACTGGGTATCATATGCTGACTGTATATGCATGAGGATAGCATATGCTACCCGG | 4 |
| wt-7470-7590 | 1490 | ATACAGATTAGGATAGCATATACTACCCAGATATAGATTAGGATAGCATATGCTACCCAGATATAGATTAGGATAGCCTATGCTACCCAGATATAAATTAGGATAGCATATACTACCCAG | 4 |
| wt-7590-7710 | 1491 | ATATAGATTAGGATAGCATATGCTACCCAGATATAGATTAGGATAGCCTATGCTACCCAGATATAGATTAGGATAGCATATGCTACCCAGATATAGATTAGGATAGCATATGCTATCCAG | 4 |
| wt-7710-7830 | 1492 | ATATTTGGGTAGTATATGCTACCCAGATATAAATTAGGATAGCATATACTACCCTAATCTCTATTAGGATAGCATATGCTACCCGGATACAGATTAGGATAGCATATACTACCCAGATAT | 4 |
| wt-7830-7950 | 1493 | AGATTAGGATAGCATATGCTACCCAGATATAGATTAGGATAGCCTATGCTACCCAGATATAAATTAGGATAGCATATACTACCCAGATATAGATTAGGATAGCATATGCTACCCAGATAT | 4 |
| wt-7950-8070 | 1494 | AGATTAGGATAGCCTATGCTACCCAGATATAGATTAGGATAGCATATGCTATCCAGATATTTGGGTAGTATATGCTACCCATGGCAACATTAGCCCACCGTGCTCTCAGCGACCTCGTGA | 4 |
| wt-8070-8190 | 1495 | ATATGAGGACCAACAACCCTGTGCTTGGCGCTCAGGCGCAAGTGTGTGTAATTTGTCCTCCAGATCGCAGCAATCGCGCCCCTATCTTGGCCCGCCCACCTACTTATGCAGGTATTCCCC | 4 |
| wt-8190-8310 | 1496 | GGGGTGCCATTAGTGGTTTTGTGGGCAAGTGGTTTGACCGCAGTGGTTAGCGGGGTTACAATCAGCCAAGTTATTACACCCTTATTTTACAGTCCAAAACCGCAGGGCGGCGTGTGGGGG | 4 |
| wt-8310-8430 | 1497 | CTGACGCGTGCCCCCACTCCACAATTTCAAAAAAAAGAGTGGCCACTTGTCTTTGTTTATGGGCCCCATTGGCGTGGAGCCCCGTTTAATTTTCGGGGGTGTTAGAGACAACCAGTGGAG | 4 |
| wt-8430-8550 | 1498 | TCCGCTGCTGTCGGCGTCCACTCTCTTTCCCCTTGTTACAAATAGAGTGTAACAACATGGTTCACCTGTCTTGGTCCCTGCCTGGGACACATCTTAATAACCCCAGTATCATATTGCACT | 4 |
| wt-8550-8670 | 1499 | AGGATTATGTGTTGCCCATAGCCATAAATTCGTGTGAGATGGACATCCAGTCTTTACGGCTTGTCCCCACCCCATGGATTTCTATTGTTAAAGATATTCAGAATGTTTCATTCCTACACT | 4 |
| wt-8670-8790 | 1500 | AGTATTTATTGCCCAAGGGGTTTGTGAGGGTTATATTGGTGTCATAGCACAATGCCACCACTGAACCCCCCGTCCAAATTTTATTCTGGGGGCGTCACCTGAAACCTTGTTTTCGAGCAC | 4 |
| wt-8790-8910 | 1501 | CTCACATACACCTTACTGTTCACAACTCAGCAGTTATTCTATTAGCTAAACGAAGGAGAATGAAGAAGCAGGCGAAGATTCAGGAGAGTTCACTGCCCGCTCCTTGATCTTCAGCCACTG | 4 |
| wt-8910-9030 | 1502 | CCCTTGTGACTAAAATGGTTCACTACCCTCGTGGAATCCTGACCCCATGTAAATAAAACCGTGACAGCTCATGGGGTGGGAGATATCGCTGTTCCTTAGGACCCTTTTACTAACCCTAAT | 4 |
| wt-9030-9150 | 1503 | TCGATAGCATATGCTTCCCGTTGGGTAACATATGCTATTGAATTAGGGTTAGTCTGGATAGTATATACTACTACCCGGGAAGCATATGCTACCCGTTTAGGGTTAACAAGGGGGCCTTAT | 4 |
| wt-9150-9270 | 1504 | AAACACTATTGCTAATGCCCTCTTGAGGGTCCGCTTATCGGTAGCTACACAGGCCCCTCTGATTGACGTTGGTGTAGCCTCCCGTAGTCTTCCTGGGCCCCTGGGAGGTACATGTCCCCC | 4 |
| wt-9270-9390 | 1505 | AGCATTGGTGTAAGAGCTTCAGCCAAGAGTTACACATAAAGGCAATGTTGTGTTGCAGTCCACAGACTGCAAAGTCTGCTCCAGGATGAAAGCCACTCAGTGTTGGCAAATGTGCACATC | 4 |
| wt-9390-9510 | 1506 | CATTTATAAGGATGTCAACTACAGTCAGAGAACCCCTTTGTGTTTGGTCCCCCCCCGTGTCACATGTGGAACAGGGCCCAGTTGGCAAGTTGTACCAACCAACTGAAGGGATTACATGCA | 4 |
| wt-9510-9630 | 1507 | CTGCCCCGCGGGAAATACGTCCTACCCAGGAACCCGAAACAGTGTTTCCCAGAAGCTGTAAAAATAGAACGCCCTGGAACTGCCCCACTGTGCAATGCAGCTTTTAGCCATGCCATGCTC | 4 |
| wt-9630-9750 | 1508 | TATAAATCACTTCCCTATCTCAGGTAGGCCTGCACACCTTAGGTATGGAGCGAAGGTTAGTGGTCACTCTGCAGTGCCTGGTGCTGCTTTACCTGGCACCTGAGTGTGGAGGTACAGACC | 4 |
| wt-9750-9870 | 1509 | AATGTGACAATTTTCCCCAAATGTTGAGGGACCTAAGAGATGCCTTCAGTCGTGTTAAAACCTTTTTCCAGACAAAGGACGAGGTAGATAACCTTTTGCTCAAGGAGTCTCTGCTAGAGG | 4 |
| wt-9870-9990 | 1510 | ACTTTAAGGGCTACCTTGGATGCCAGGCCCTGTCAGAAATGATCCAATTCTACCTGGAGGAAGTCATGCCACAGGCTGAAAACCAGGACCCTGAAGCCAAAGACCATGTCAATTCTTTGG | 4 |
| wt-9990-10110 | 1511 | GTGAAAATCTAAAGACCCTACGGCTCCGCCTGCGCAGGTGCCACAGGTTCCTGCCGTGTGAGAACAAGAGTAAAGCTGTGGAACAGATAAAAAATGCCTTTAACAAGCTGCAGGAAAAAG | 4 |
| wt-10230-10350 | 1512 | AACCCCCCCTTTCGACTGTCATTTACAATAAAATGAAACCTTTTATTCTTGATTGCCTCTTGTGTTCTTGCCGCCCAGGTACCTTCCTGTGTTCTCCCCACGGGAAAAAGAATAGCTTCT | 4 |
| wt-10350-10470 | 1513 | GCAGAAGGCCATTGACGCAAGTTTTGCCCGTGGGGATTACCCGACCCAGCCACTTACAGCACATTTTGTTCTAGGTCCATCTTAGGAGCCCGGGCCAGCATTCTATCAGCTTAACGGGAA | 4 |
| wt-10470-10590 | 1514 | GAGAAGTGGGGAGGGCACTCGCCCACTAACCTTAACACCTGCAGCCTACAAAAGTACACTAGCTGTTTGCTCTATTCGCCACTAGAGACCGCCAAGATGCGAAACTACAGGCCCGGGCCC | 4 |
| wt-10590-10710 | 1515 | AGGCCTTGCAGGGCAGACGGTTAGGCTGACAAGGGGACAAGTGTGGCAGGTGGGCGGGAAGGGGCACAAGAATGCCGGCGAAACTGGACCACGGTCCACCCCGCCCTCAAGCGTCCGGGA | 8 |

| | | | |
|---|---|---|---|
| wt-10710-10830 | 1516 | GCCGGGCGGCTCGGCTAAGGAGGGCGGCCTTGCGAACAATTATTAGTAGCTACCAACAAGGGCCCCCAGATGCCCCCCACCAGTCACCCGGCCGTGTCCACTCACATATTCCACTCTTAT | 4 |
| wt-10830-10950 | 1517 | TTTTAAATTAATGTGTCCCAATTAGAAACCCAAGCGCAGAAATTAGTTGAGAGGCTAGTGTTTTAAACATGCACCCTAGGCCAGCCAGAGATAATGTCACAAGATTATCAAGTTGGTGTA | 4 |
| wt-10950-11070 | 1518 | AACACGCCGTGGGAAAAAATTTATGGTTCAGTGCGTCGAGTGCTATCTTTGGAACAGTAGAAAATTGAACCTTGTTGGCGGGAGAAGGAATAACGCCTTATCTGGGAGGAGCGACGGATT | 4 |
| wt-11070-11190 | 1519 | ATAGCCAATAAGAGAGCTCAAGACGCAGGGCTCGCAAAGTATAGTGGCCCCGTGGGACCTTAGAGGTGGAGCAACGTCTAAAGTGGTAATAACACCAGGCGGGGCTGGGCAAAGGGGTCC | 4 |
| wt-11190-11310 | 1520 | TACGGGCGGGATTAATTACGCCTTGCTTACGCAAGCTCAGTTAATTCGCCCACGACTTGAAAAATGTAGCCCTTAACCAATTGGCGGCCCCTAAGGGGGGGACTAAGGTCCCACTACAAA | 4 |
| wt-11310-11430 | 1521 | AACTCTGTGTTCTGCTGCAAATTTTAGATCAGATGGCATAGAGACAAGGACACCGAAGACCCCCAGAGCCCTCATCGCAGGGTTCTTACCATGCGGCCATGTAGGCCCACTTAACACTAC | 4 |
| wt-11430-11550 | 1522 | AAGACCTACGCCTCTCCATTCATCATGTAACCCACAAATCATCTAAACCGTAAGTCTAAGGGCCTCCTGAGGTTTTCTCAGGAGGCCCTAATGTATAATTAATCATGCATTTGATTTTAA | 4 |
| wt-11550-11670 | 1523 | AAAAGTAGGTTACACTCATTTTAGGCCAGACTTTATTTGCAGATTAATAATTTATGTGATTCTCCTTCCCTCTAGGACTGAAGAAACAGCCTCCTGCACGTGAGCATGTATCTGAAATAA | 4 |
| wt-11670-11790 | 1524 | TTATTATGTCATAAGTGTAATGATTAGAAAGTCATAAACCCACTTCCCTTTACATGAATCTGGGCACTGAATTTTGGGGTACTTCTAAAGACTAACGTGTTCGATTTCGGGGTCACTTCC | 4 |
| wt-11790-11910 | 1525 | CCTTTTATAAGTGTGTGAACAGTGATTTCAGTAAAACCTAAGAGATATTTGGTGTCACTTCCGCATTTTAAGTTTCAGAAAATTTTAAAATTAAAATTGAAATTTCTCTCAAAATAATTC | 4 |
| wt-11910-12030 | 1526 | CAATGAAAACTTCAAAGAATCTTATGTATGTAATTCTTTTGCCCCAAACTGGGCTTCAGATGCCTTCTATTGCACTCTCACAAAAACATTCTGGACACATGTGCCAGACGCCTGGGCCTC | 4 |
| wt-12030-12150 | 1527 | TAAGGCCCTCGGGTCCCCCTGGACCCCGGCCTCAGCAACCCTGCTGCTCCCCTCCTGCCACCCCAGCCTCCCCCCCTCCCCGTCCCCCTTCGCTCCTGTTCCTCCCCCGGTCCCCAGTAG | 8 |
| wt-12150-12270 | 1528 | GGCCGCCTGCCCCCCTGCACCCAGTACCTGCCCCTCTTGGCCACGCACCCCGGGCCAGGCCACCTTAGACCCGGCCAAGCCCCATCCCTGAAGACCCAGCGGCCATTCTCTCTGGTAACG | 8 |
| wt-12270-12390 | 1529 | AGCAGAGAAGAAGTAGAGGCCCGCGGCCATTGGGCCCAGATTGAGAGACCAGTCCAGGGGCCCGAGGTTGGAGCCAGCGGGCACCCGAGGTCCCAGCACCCGGTCCCTCCGGGGGGCAGA | 8 |
| wt-12390-12510 | 1530 | GACAGGCAGGGCCCCCCGGCAGCTGGCCCCGAGGAGGCGCCCGGAGTGGGGCCGGTCGGCTGGGCTGGCCGAGCCCGGGTCTGGGAGGTCTGGGGTGGCGAGCCTGCTGTCTCAGGAGGG | 12 |
| wt-12510-12630 | 1531 | GCCTGGCTCCGCCGGGTGGCCCTGGGGTAAGTCTGGGAGGCAGAGGGTCGGCCTAGGCCCGGGGAAGTGGAGGGGGATCGCCCGGGTCTCTGTTGGCAGAGTCCGGGCGATCCTCTGAGA | 8 |
| wt-12630-12750 | 1532 | CCCTCCGGGCCCGGACGGTCGCCCTCAGCCCCCCAGACAGACCCCAGGGTCTCCAGGCAGGGTCCGGCATCTTCAGGGGCAGCAGGCTCACCACCACAGGCCCCCCAGACCCGGGTCTCG | 8 |
| wt-12750-12870 | 1533 | GCCAGCCGAGCCGACCGGCCCCGCGCCTGGCGCCTCCTCGGGGCCAGCCGCCGGGGTTGGTTCTGCCCCTCTCTCTGTCCTTCAGAGGAACCAGGGACCTCGGGCACCCCAGAGCCCCTC | 8 |
| wt-12870-12990 | 1534 | GGGCCCGCCTCCAGGCGCCCTCCTGGTCTCCGCTCCCCTCTGAGCCCCGTTAAACCCAAAGAATGTCTGAGGGGAGCCACCCTCGGGGCCCAGGCCCCAGAGTCCAGAGGTCAGGGGCAC | 8 |
| wt-12990-13110 | 1535 | CTCAGGGTGCCTCCCCGGGTCCCAGGCCAGCCGGAGGGACCCCGGCAGCCCGGGCGGCCCCAGAGGCCGGTTCCTCGCCCCTTCCCCGGGCTTCAGAGCCCAGGATGTCCCCCAGAAGGG | 12 |
| wt-13110-13230 | 1536 | ACCCTAGGCGTCCCCTCTCCTCCCCTCCAGGCCCGAGCCTCTCCCTCGCGGAGAGGGGCCTCTTTGGGCCCTCAAGTCCAGCCCCACCGAGACCCGAGTGGCCCGGATCCCCCCACCGGC | 8 |
| wt-13230-13350 | 1537 | CCTTCTCTCTGTCCCCCTGCTCCTCTCCAACCTTCGCTCCACCCTAGACCCCAGCTTCTGGCCTCCCCGGGTCCACCAGGCCAGCCGGAGGGACCCCGGCAGCCCGGGCGAGTCGCCTTC | 8 |
| wt-13350-13470 | 1538 | CCTCTCCCCTGGCCTCTCCTTCCCGCCTCCCACCCGAGCCCCCTCAGCTTGCCTCCCCACCGGGTCCATCAGGCCGGCCGGAGGGACCCCGGCGGCCCGGTGTCAGTCCCCCCTGCAGCC | 12 |
| wt-13470-13590 | 1539 | GCCCAGTCTCTGCCTCCAGGCAAGGGCGCCAGCTTTTCTCCCCCCAGCCTGAGGCCCAGTCTCCTGTGCACTGTCTGTAAAGTCCAGCCTCCCACGCCCGTCCACGGCTCCCGGGCCCAG | 8 |
| wt-13590-13710 | 1540 | CCTCGTCCACCCCTCCCCACGGTGGACAGGCCCTCTGTCCACCCGGGCCATCCCCGCCCCCCTGTGTCCACCCCAGTCCCGTCCAGGGGGGACTTTATGTGACCCTTGGGCCTGGCTCCC | 8 |
| wt-13710-13830 | 1541 | CATAGACTCCCATGTAAGCCTGCCTCGAGTAGGTGCCTCCAGAGCCCCTTTTGCCCCCCTGGCGGCCCAGCCCGACCCCCGGGCGCCCCCAAACTTTGTCCAGATGTCCAGGGGTCCCCG | 8 |
| wt-13830-13950 | 1542 | AGGGTGAGGCCCAGCCCCCTCCCGCCCCTGTCCACTGCCCCGGTCCCCCCAGAAGCCCCCAAAAGTAGAGGCTCAGGCCATGCGCGCCCTGTCACCAGGCCTGCCAAAGAGCCAGATCTA | 8 |
| wt-13950-14070 | 1543 | AGGCCGGGAGAGGCAGCCCCAAAGCGGGTGCAGTAACAGGTAATCTCTGGTAGTGATTTGGACCCGAAATCTGACACTTTAGAGCTCTGGAGGACTTTAAAACTCTAAAAATCAAAACTT | 4 |
| wt-14070-14190 | 1544 | TAGAGGCGAATGGGCGCCATTTTGTCCCCACGCGCGCATAATGGCGGACCTAGGCCTAAAACCCCCAGGAAGCGGGTCTATGGTTGGCTGCGCTGCTGCTATCTTTAGAGGGGAAAAGAG | 4 |
| wt-14190-14310 | 1545 | GAATAAGCCCCCAGACAGGGGAGTGGGCTTGTTTGTGACTTCACCAAAGGTCAGGGCCCAAGGGGGTTCGCGTTGCTAGGCCACCTTCTCAGTCCAGCGCGTTTACGTAAGCCAGACAGC | 4 |
| wt-14310-14430 | 1546 | AGCCAATTGTCAGTTCTAGGGAGGGGGACCACTGCCCCTGGTATAAAGTGGTCCTGCAGCTATTTCTGGTCGCATCAGAGCGCCAGGAGTCCACACAAATGTAAGAGGGGGTCTTCTACC | 4 |
| wt-14430-14550 | 1547 | TCTCCCTAGCCCTCCGCCCCCTCCAAGGACTCGGGCCCAGTTTCTAACTTTTCCCCTTCCCTCCCTCGTCTTGCCCTGCGCCCGGGGCCACCTTCATCACCGTCGCTGACTCCGCCATCC | 8 |
| wt-14550-14670 | 1548 | AAGCCTAGGGGAGACCGAAGTGAAGGCCCTGGACCAACCCGGCCCGGGCCCCCCGGTATCGGGCCAGAGGTAAGTGGACTTTAATTTTTTCTGCTAAGCCCAACACTCCACCACACCCAG | 4 |
| wt-14670-14790 | 1549 | GCACACACTACACACACCCACCCGTCTCAGGGTCCCCTCGGACAGCTCCTAAGAAGGCACCGGTCGCCCAGTCCTACCAGAGGGGGCCAAGAACCCAGACGAGTCCGTAGAAGGGTCCTC | 4 |
| wt-14790-14910 | 1550 | GTCCAGCAAGAAGAGGAGGTGGTAAGCGGTTCACCTTCAGGGGTAAGTAACCTGACCTCTCCAGGGCTCACATAAAGGGAGGCTTAGTATACATGCTTCTTGCTTTTCACAGGAACCTGG | 4 |
| wt-14910-15030 | 1551 | GGGCTAGTCTGGGTGGGATTAGGCTGCCTCAAGTTGCATCAGCCAGGGCTTCATGCCCTCCTCAGTTCCCTAGTCCCCGGGCTTCAGGCCCCCTCCGTCCCCGTCCTCCAGAGACCCGGG | 8 |
| wt-15030-15150 | 1552 | CTTCAGGCCCTGCCTCTCCTGTTACCCTTTTAGAACCACAGCCTGGACACATGTGCCAGACGCCTTGGCCTCTAAGGCCCTCGGGTCCCCCTGGACCCCGGCCTCAGCAACCCTGCTGCT | 4 |
| wt-15150-15270 | 1553 | CCCCTCCTGCCACCCCAGCCTCCCCCCCTCCCCGTCCCCCTTCGCTCCTGATCCTCCCCCGGTCCCCAGTAGGGCCGCCTGCCCCCCTGCACCCAGTACCTGCCCCTCTTGGCCACGCAC | 8 |
| wt-15270-15390 | 1554 | CCCGGGCCAGGCCACCTTAGACCCGGCCAAGCCCCATCCCTGAAGACCCAGCGGCCATTCTCTCTGGTAACGAGCAGAGAAGAAGTAGAGGCCCGCGGCCATTGGGCCCAGATTGAGAGA | 4 |
| wt-15390-15510 | 1555 | CCAGTCCAGGGGCCCGAGGTTGGAGCCAGCGGGCACCCGAGGTCCCAGCACCCGGTCCCTCCGGGGGGCAGAGACAGGCAGGGCCCCCCGGCAGCTGGCCCCGAGGAGGCGCCCGGAGTG | 12 |
| wt-15510-15630 | 1556 | GGGCCGGTCGGCTGGGCTGGCCGAGCCCGGGTCTGGGAGGTCTGGGGTGGCGAGCCTGCTGTCTCAGGAGGGGCCTGGCTCCGCCGGGTGGCCCTGGGGTAAGTCTGGGAGGCAGAGGGT | 8 |
| wt-15630-15750 | 1557 | CGGCCTAGGCCCGGGGAAGTGGAGGGGGATCGCCCGGGTCTCTGTTGGCAGAGTCCGGGCGATCCTCTGAGACCCTCCGGGCCCGGACGGTCGCCCTCAGCCCCCCAGACAGACCCCAGG | 8 |
| wt-15750-15870 | 1558 | GTCTCCAGGCAGGGTCCGGCATCTTCAGGGGCAGCAGGCTCACCACCACAGGCCCCCCAGACCCGGGTCTCGGCCAGCCGAGCCGACCGGCCCCGCGCCTGGCGCCTCCTCGGGGCCAGC | 12 |
| wt-15870-15990 | 1559 | CGCCGGGGTTGGTTCTGCCCCTCTCTCTGTCCTTCAGAGGAACCAGGGACCTCGGGCACCCCAGAGCCCCTCGGGCCCGCCTCCAGGCGCCCTCCTGGTCTCCGCTCCCCTCTGAGCCCC | 8 |
| wt-15990-16110 | 1560 | GTTAAACCCAAAGAATGTCTGAGGGGAGCCACCCTCGGGGCCCAGGCCCCAGAGTCCAGAGGTCAGGGGCACCTCAGGGTGCCTCCCCGGGTCCCAGGCCAGCCGGAGGGACCCCGGCAG | 8 |
| wt-16110-16230 | 1561 | CCCGGGCGGCCCCAGAGGCCGGTTCCTCGCCCCTTCCCCGGGCTTCAGAGCCCAGGATGTCCCCCAGAAGGGACCCTAGGCGTCCCCTCTCCTCCCCTCCAGGCCCGAGCCTCTCCCTCG | 8 |
| wt-16230-16350 | 1562 | CGGAGAGGGGCCTCTTTGGGCCCTCAAGTCCAGCCCCACCGAGACCCGAGTGGCCCGGATCCCCCCACCGGCCCTTCTCTCTGTCCCCCTGCTCCTCTCCAACCTTCGCTCCACCCTAGA | 8 |
| wt-16350-16470 | 1563 | CCCCAGCTTCTGGCCTCCCCGGGTCCACCAGGCCAGCCGGAGGGACCCCGGCAGCCCGGGCGAGTCGCCTTCCCTCTCCCCTGGCCTCTCCTTCCCGCCTCCCACCCGAGCCCCCTCAGC | 12 |
| wt-16470-16590 | 1564 | TTGCCTCCCCACCGGGTCCATCAGGCCGGCCGGAGGGACCCCGGCGGCCCGGTGTCAGTCCCCCCTGCAGCCGCCCAGTCTCTGCCTCCAGGCAAGGGCGCCAGCTTTTCTCCCCCCAGC | 8 |
| wt-16590-16710 | 1565 | CTGAGGCCCAGTCTCCTGTGCACTGTCTGTAAAGTCCAGCCTCCCACGCCCGTCCACGGCTCCCGGGCCCAGCCTCGTCCACCCCTCCCCACGGTGGACAGGCCCTCTGTCCACCCGGGC | 8 |
| wt-16710-16830 | 1566 | CATCCCCGCCCCCCTGTGTCCACCCCAGTCCCGTCCAGGGGGGACTTTATGTGACCCTTGGGCCTGGCTCCCCATAGACTCCCATGTAAGCCTGCCTCGAGTAGGTGCCTCCAGAGCCCC | 8 |

| | | | |
|---|---|---|---|
| wt-16830-16950 | 1567 | TTTTGCCCCCCTGGCGGCCCAGCCCGACCCCCGGGCGCCCCCAAACTTTGTCCAGATGTCCAGGGGTCCCCGAGGGTGAGGCCCAGCCCCCTCCCGCCCCTGTCCACTGCCCCGGTCCCC | 8 |
| wt-16950-17070 | 1568 | CCAGAAGCCCCCAAAAGTAGAGGCTCAGGCCATGCGCGCCCTGTCACCAGGCCTGCCAAAGAGCCAGATCTAAGGCCGGGAGAGGCAGCCCCAAAGCGGGTGCAGTAACAGGTAATCTCT | 4 |
| wt-17070-17190 | 1569 | GGTAGTGATTTGGACCCGAAATCTGACACTTTAGAGCTCTGGAGGACTTTAAAACTCTAAAAATCAAAACTTTAGAGGCGAATGGGCGCCATTTTGTCCCCACGCGCGCATAATGGCGGA | 4 |
| wt-17190-17310 | 1570 | CCTAGGCCTAAAACCCCCAGGAAGCGGGTCTATGGTTGGCTGCGCTGCTGCTATCTTTAGAGGGGAAAAGAGGAATAAGCCCCCAGACAGGGGAGTGGGCTTGTTTGTGACTTCACCAAA | 4 |
| wt-17310-17430 | 1571 | GGTCAGGGCCCAAGGGGGTTCGCGTTGCTAGGCCACCTTCTCAGTCCAGCGCGTTTACGTAAGCCAGACAGCAGCCAATTGTCAGTTCTAGGGAGGGGGACCACTGCCCCTGGTATAAAG | 4 |
| wt-17430-17550 | 1572 | TGGTCCTGCAGCTATTTCTGGTCGCATCAGAGCGCCAGGAGTCCACACAAATGTAAGAGGGGGTCTTCTACCTCTCCCTAGCCCTCCGCCCCCTCCAAGGACTCGGGCCCAGTTTCTAAC | 4 |
| wt-17550-17670 | 1573 | TTTTCCCCTTCCCTCCCTCGTCTTGCCCTGCGCCCGGGGCCACCTTCATCACCGTCGCTGACTCCGCCATCCAAGCCTAGGGGAGACCGAAGTGAAGGCCCTGGACCAACCCGGCCCGGG | 8 |
| wt-17670-17790 | 1574 | CCCCCCGGTATCGGGCCAGAGGTAAGTGGACTTTAATTTTTTCTGCTAAGCCCAACACTCCACCACACCCAGGCACACACTACACACACCCACCCGTCTCAGGGTCCCCTCGGACAGCTC | 4 |
| wt-17790-17910 | 1575 | CTAAGAAGGCACCGGTCGCCCAGTCCTACCAGAGGGGGCCAAGAACCCAGACGAGTCCGTAGAAGGGTCCTCGTCCAGCAAGAAGAGGAGGTGGTAAGCGGTTCACCTTCAGGGGTAAGT | 4 |
| wt-17910-18030 | 1576 | AACCTGACCTCTCCAGGGCTCACATAAAGGGAGGCTTAGTATACATGCTTCTTGCTTTTCACAGGAACCTGGGGGCTAGTCTGGGTGGGATTAGGCTGCCTCAAGTTGCATCAGCCAGGG | 4 |
| wt-18030-18150 | 1577 | CTTCATGCCCTCCTCAGTTCCCTAGTCCCCGGGCTTCAGGCCCCCTCCGTCCCCGTCCTCCAGAGACCCGGGCTTCAGGCCCTGCCTCTCCTGTTACCCTTTTAGAACCACAGCCTGGAC | 4 |
| wt-18150-18270 | 1578 | ACATGTGCCAGACGCCTTGGCCTCTAAGGCCCTCGGGTCCCCCTGGACCCCGGCCTCAGCAACCCTGCTGCTCCCCTCCTGCCACCCCAGCCTCCCCCCCTCCCCGTCCCCCTTCGCTCC | 8 |
| wt-18270-18390 | 1579 | TGATCCTCCCCCGGTCCCCAGTAGGGCCGCCTGCCCCCCTGCACCCAGTACCTGCCCCTCTTGGCCACGCACCCCGGGCCAGGCCACCTTAGACCCGGCCAAGCCCCATCCCTGAAGACC | 8 |
| wt-18510-18630 | 1581 | CACCCGGTCCCTCCGGGGGGCAGAGACAGGCAGGGCCCCCCGGCAGCTGGCCCCGAGGAGGCGCCCGGAGTGGGGCCGGTCGGCTGGGCTGGCCGAGCCCGGGTCTGGGAGGTCTGGGGT | 12 |
| wt-18630-18750 | 1582 | GGCGAGCCTGCTGTCTCAGGAGGGGCCTGGCTCCGCCGGGTGGCCCTGGGGTAAGTCTGGGAGGCAGAGGGTCGGCCTAGGCCCGGGGAAGTGGAGGGGGATCGCCCGGGTCTCTGTTGG | 8 |
| wt-18750-18870 | 1583 | CAGAGTCCGGGCGATCCTCTGAGACCCTCCGGGCCCGGACGGTCGCCCTCAGCCCCCCAGACAGACCCCAGGGTCTCCAGGCAGGGTCCGGCATCTTCAGGGGCAGCAGGCTCACCACCA | 8 |
| wt-18870-18990 | 1584 | CAGGCCCCCCAGACCCGGGTCTCGGCCAGCCGAGCCGACCGGCCCCGCGCCTGGCGCCTCCTCGGGGCCAGCCGCCGGGGTTGGTTCTGCCCCTCTCTCTGTCCTTCAGAGGAACCAGGG | 12 |
| wt-18990-19110 | 1585 | ACCTCGGGCACCCCAGAGCCCCTCGGGCCCGCCTCCAGGCGCCCTCCTGGTCTCCGCTCCCCTCTGAGCCCCGTTAAACCCAAAGAATGTCTGAGGGGAGCCACCCTCGGGGCCCAGGCC | 8 |
| wt-19110-19230 | 1586 | CCAGAGTCCAGAGGTCAGGGGCACCTCAGGGTGCCTCCCCGGGTCCCAGGCCAGCCGGAGGGACCCCGGCAGCCCGGGCGGCCCCAGAGGCCGGTTCCTCGCCCCTTCCCCGGGCTTCAG | 12 |
| wt-19230-19350 | 1587 | AGCCCAGGATGTCCCCCAGAAGGGACCCTAGGCGTCCCCTCTCCTCCCCTCCAGGCCCGAGCCTCTCCCTCGCGGAGAGGGGCCTCTTTGGGCCCTCAAGTCCAGCCCCACCGAGACCCG | 8 |
| wt-19350-19470 | 1588 | AGTGGCCCGGATCCCCCCACCGGCCCTTCTCTCTGTCCCCCTGCTCCTCTCCAACCTTCGCTCCACCCTAGACCCCAGCTTCTGGCCTCCCCGGGTCCACCAGGCCAGCCGGAGGGACCC | 8 |
| wt-19470-19590 | 1589 | CGGCAGCCCGGGCGAGTCGCCTTCCCTCTCCCCTGGCCTCTCCTTCCCGCCTCCCACCCGAGCCCCCTCAGCTTGCCTCCCCACCGGGTCCATCAGGCCGGCCGGAGGGACCCCGGCGGC | 12 |
| wt-19590-19710 | 1590 | CCGGTGTCAGTCCCCCCTGCAGCCGCCCAGTCTCTGCCTCCAGGCAAGGGCGCCAGCTTTTCTCCCCCCAGCCTGAGGCCCAGTCTCCTGTGCACTGTCTGTAAAGTCCAGCCTCCCACG | 8 |
| wt-19710-19830 | 1591 | CCCGTCCACGGCTCCCGGGCCCAGCCTCGTCCACCCCTCCCCACGGTGGACAGGCCCTCTGTCCACCCGGGCCATCCCCGCCCCCCTGTGTCCACCCCAGTCCCGTCCAGGGGGGACTTT | 8 |
| wt-19830-19950 | 1592 | ATGTGACCCTTGGGCCTGGCTCCCCATAGACTCCCATGTAAGCCTGCCTCGAGTAGGTGCCTCCAGAGCCCCTTTTGCCCCCCTGGCGGCCCAGCCCGACCCCCGGGCGCCCCCAAACTT | 8 |
| wt-19950-20070 | 1593 | TGTCCAGATGTCCAGGGGTCCCCGAGGGTGAGGCCCAGCCCCCTCCCGCCCCTGTCCACTGCCCCGGTCCCCCCAGAAGCCCCCAAAAGTAGAGGCTCAGGCCATGCGCGCCCTGTCACC | 8 |
| wt-20070-20190 | 1594 | AGGCCTGCCAAAGAGCCAGATCTAAGGCCGGGAGAGGCAGCCCCAAAGCGGGTGCAGTAACAGGTAATCTCTGGTAGTGATTTGGACCCGAAATCTGACACTTTAGAGCTCTGGAGGACT | 4 |
| wt-20190-20310 | 1595 | TTAAAACTCTAAAAATCAAAACTTTAGAGGCGAATGGGCGCCATTTTGTCCCCACGCGCGCATAATGGCGGACCTAGGCCTAAAACCCCCAGGAAGCGGGTCTATGGTTGGCTGCGCTGC | 4 |
| wt-20310-20430 | 1596 | TGCTATCTTTAGAGGGGAAAAGAGGAATAAGCCCCCAGACAGGGGAGTGGGCTTGTTTGTGACTTCACCAAAGGTCAGGGCCCAAGGGGGTTCGCGTTGCTAGGCCACCTTCTCAGTCCA | 4 |
| wt-20430-20550 | 1597 | GCGCGTTTACGTAAGCCAGACAGCAGCCAATTGTCAGTTCTAGGGAGGGGGACCACTGCCCCTGGTATAAAGTGGTCCTGCAGCTATTTCTGGTCGCATCAGAGCGCCAGGAGTCCACAC | 4 |
| wt-20550-20670 | 1598 | AAATGTAAGAGGGGGTCTTCTACCTCTCCCTAGCCCTCCGCCCCCTCCAAGGACTCGGGCCCAGTTTCTAACTTTTCCCCTTCCCTCCCTCGTCTTGCCCTGCGCCCGGGGCCACCTTCA | 4 |
| wt-20670-20790 | 1599 | TCACCGTCGCTGACTCCGCCATCCAAGCCTAGGGGAGACCGAAGTGAAGGCCCTGGACCAACCCGGCCCGGGCCCCCCGGTATCGGGCCAGAGGTAAGTGGACTTTAATTTTTTCTGCTA | 4 |
| wt-20790-20910 | 1600 | AGCCCAACACTCCACCACACCCAGGCACACACTACACACACCCACCCGTCTCAGGGTCCCCTCGGACAGCTCCTAAGAAGGCACCGGTCGCCCAGTCCTACCAGAGGGGGCCAAGAACCC | 4 |
| wt-20910-21030 | 1601 | AGACGAGTCCGTAGAAGGGTCCTCGTCCAGCAAGAAGAGGAGGTGGTAAGCGGTTCACCTTCAGGGGTAAGTAACCTGACCTCTCCAGGGCTCACATAAAGGGAGGCTTAGTATACATGC | 4 |
| wt-21030-21150 | 1602 | TTCTTGCTTTTCACAGGAACCTGGGGGCTAGTCTGGGTGGGATTAGGCTGCCTCAAGTTGCATCAGCCAGGGCTTCATGCCCTCCTCAGTTCCCTAGTCCCCGGGCTTCAGGCCCCCTCC | 4 |
| wt-21150-21270 | 1603 | GTCCCCGTCCTCCAGAGACCCGGGCTTCAGGCCCTGCCTCTCCTGTTACCCTTTTAGAACCACAGCCTGGACACATGTGCCAGACGCCTTGGCCTCTAAGGCCCTCGGGTCCCCCTGGAC | 4 |
| wt-21270-21390 | 1604 | CCCGGCCTCAGCAACCCTGCTGCTCCCCTCCTGCCACCCCAGCCTCCCCCCCTCCCCGTCCCCCTTCGCTCCTGATCCTCCCCCGGTCCCCAGTAGGGCCGCCTGCCCCCCTGCACCCAG | 12 |
| wt-21510-21630 | 1606 | GGCCATTGGGCCCAGATTGAGAGACCAGTCCAGGGGCCCGAGGTTGGAGCCAGCGGGCACCCGAGGTCCCAGCACCCGGTCCCTCCGGGGGGCAGAGACAGGCAGGGCCCCCCGGCAGCT | 8 |
| wt-21630-21750 | 1607 | GGCCCCGAGGAGGCGCCCGGAGTGGGGCCGGTCGGCTGGGCTGGCCGAGCCCGGGTCTGGGAGGTCTGGGGTGGCGAGCCTGCTGTCTCAGGAGGGGCCTGGCTCCGCCGGGTGGCCCTG | 12 |
| wt-21750-21870 | 1608 | GGGTAAGTCTGGGAGGCAGAGGGTCGGCCTAGGCCCGGGGAAGTGGAGGGGGATCGCCCGGGTCTCTGTTGGCAGAGTCCGGGCGATCCTCTGAGACCCTCCGGGCCCGGACGGTCGCCC | 8 |
| wt-21870-21990 | 1609 | TCAGCCCCCCAGACAGACCCCAGGGTCTCCAGGCAGGGTCCGGCATCTTCAGGGGCAGCAGGCTCACCACCACAGGCCCCCCAGACCCGGGTCTCGGCCAGCCGAGCCGACCGGCCCCGC | 8 |
| wt-21990-22110 | 1610 | GCCTGGCGCCTCCTCGGGGCCAGCCGCCGGGGTTGGTTCTGCCCCTCTCTCTGTCCTTCAGAGGAACCAGGGACCTCGGGCACCCCAGAGCCCCTCGGGCCCGCCTCCAGGCGCCCTCCT | 8 |
| wt-22110-22230 | 1611 | GGTCTCCGCTCCCCTCTGAGCCCCGTTAAACCCAAAGAATGTCTGAGGGGAGCCACCCTCGGGGCCCAGGCCCCAGAGTCCAGAGGTCAGGGGCACCTCAGGGTGCCTCCCCGGGTCCCA | 8 |
| wt-22230-22350 | 1612 | GGCCAGCCGGAGGGACCCCGGCAGCCCGGGCGGCCCCAGAGGCCGGTTCCTCGCCCCTTCCCCGGGCTTCAGAGCCCAGGATGTCCCCCAGAAGGGACCCTAGGCGTCCCCTCTCCTCCC | 12 |
| wt-22350-22470 | 1613 | CTCCAGGCCCGAGCCTCTCCCTCGCGGAGAGGGGCCTCTTTGGGCCCTCAAGTCCAGCCCCACCGAGACCCGAGTGGCCCGGATCCCCCCACCGGCCCTTCTCTCTGTCCCCCTGCTCCT | 8 |
| wt-22470-22590 | 1614 | CTCCAACCTTCGCTCCACCCTAGACCCCAGCTTCTGGCCTCCCCGGGTCCACCAGGCCAGCCGGAGGGACCCCGGCAGCCCGGGCGAGTCGCCTTCCCTCTCCCCTGGCCTCTCCTTCCC | 8 |
| wt-22590-22710 | 1615 | GCCTCCCACCCGAGCCCCCTCAGCTTGCCTCCCCACCGGGTCCATCAGGCCGGCCGGAGGGACCCCGGCGGCCCGGTGTCAGTCCCCCCTGCAGCCGCCCAGTCTCTGCCTCCAGGCAAG | 12 |
| wt-22710-22830 | 1616 | GGCGCCAGCTTTTCTCCCCCCAGCCTGAGGCCCAGTCTCCTGTGCACTGTCTGTAAAGTCCAGCCTCCCACGCCCGTCCACGGCTCCCGGGCCCAGCCTCGTCCACCCCTCCCCACGGTG | 8 |
| wt-22830-22950 | 1617 | GACAGGCCCTCTGTCCACCCGGGCCATCCCCGCCCCCCTGTGTCCACCCCAGTCCCGTCCAGGGGGGACTTTATGTGACCCTTGGGCCTGGCTCCCCATAGACTCCCATGTAAGCCTGCC | 8 |
| wt-22950-23070 | 1618 | TCGAGTAGGTGCCTCCAGAGCCCCTTTTGCCCCCCTGGCGGCCCAGCCCGACCCCCGGGCGCCCCCAAACTTTGTCCAGATGTCCAGGGGTCCCCGAGGGTGAGGCCCAGCCCCCTCCCG | 8 |
| wt-23070-23190 | 1619 | CCCCTGTCCACTGCCCCGGTCCCCCCAGAAGCCCCCAAAAGTAGAGGCTCAGGCCATGCGCGCCCTGTCACCAGGCCTGCCAAAGAGCCAGATCTAAGGCCGGGAGAGGCAGCCCCAAAG | 8 |

| | | | |
|---|---|---|---|
| wt-23190-23310 | 1620 | CGGGTGCAGTAACAGGTAATCTCTGGTAGTGATTTGGACCCGAAATCTGACACTTTAGAGCTCTGGAGGACTTTAAAACTCTAAAAATCAAAACTTTAGAGGCGAATGGGCGCCATTTTG | 4 |
| wt-23310-23430 | 1621 | TCCCCACGCGCGCATAATGGCGGACCTAGGCCTAAAACCCCCAGGAAGCGGGTCTATGGTTGGCTGCGCTGCTGCTATCTTTAGAGGGGAAAAGAGGAATAAGCCCCCAGACAGGGGAGT | 4 |
| wt-23430-23550 | 1622 | GGGCTTGTTTGTGACTTCACCAAAGGTCAGGGCCCAAGGGGGTTCGCGTTGCTAGGCCACCTTCTCAGTCCAGCGCGTTTACGTAAGCCAGACAGCAGCCAATTGTCAGTTCTAGGGAGG | 4 |
| wt-23550-23670 | 1623 | GGGACCACTGCCCCTGGTATAAAGTGGTCCTGCAGCTATTTCTGGTCGCATCAGAGCGCCAGGAGTCCACACAAATGTAAGAGGGGGTCTTCTACCTCTCCCTAGCCCTCCGCCCCCTCC | 4 |
| wt-23670-23790 | 1624 | AAGGACTCGGGCCCAGTTTCTAACTTTTCCCCTTCCCTCCCTCGTCTTGCCCTGCGCCCGGGGCCACCTTCATCACCGTCGCTGACTCCGCCATCCAAGCCTAGGGGAGACCGAAGTGAA | 4 |
| wt-23790-23910 | 1625 | GGCCCTGGACCAACCCGGCCCGGGCCCCCCGGTATCGGGCCAGAGGTAAGTGGACTTTAATTTTTTCTGCTAAGCCCAACACTCCACCACACCCAGGCACACACTACACACACCCACCCG | 4 |
| wt-23910-24030 | 1626 | TCTCAGGGTCCCCTCGGACAGCTCCTAAGAAGGCACCGGTCGCCCAGTCCTACCAGAGGGGGCCAAGAACCCAGACGAGTCCGTAGAAGGGTCCTCGTCCAGCAAGAAGAGGAGGTGGTA | 4 |
| wt-24030-24150 | 1627 | AGCGGTTCACCTTCAGGGGTAAGTAACCTGACCTCTCCAGGGCTCACATAAAGGGAGGCTTAGTATACATGCTTCTTGCTTTTCACAGGAACCTGGGGGCTAGTCTGGGTGGGATTAGGC | 4 |
| wt-24270-24390 | 1629 | CCCTTTTAGAACCACAGCCTGGACACATGTGCCAGACGCCTTGGCCTCTAAGGCCCTCGGGTCCCCCTGGACCCCGGCCTCAGCAACCCTGCTGCTCCCCTCCTGCCACCCCAGCCTCCC | 8 |
| wt-24390-24510 | 1630 | CCCCTCCCCGTCCCCCTTCGCTCCTGATCCTCCCCCGGTCCCCAGTAGGGCCGCCTGCCCCCCTGCACCCAGTACCTGCCCCTCTTGGCCACGCACCCCGGGCCAGGCCACCTTAGACCC | 8 |
| wt-24510-24630 | 1631 | GGCCAAGCCCCATCCCTGAAGACCCAGCGGCCATTCTCTCTGGTAACGAGCAGAGAAGAAGTAGAGGCCCGCGGCCATTGGGCCCAGATTGAGAGACCAGTCCAGGGGCCCGAGGTTGGA | 4 |
| wt-24630-24750 | 1632 | GCCAGCGGGCACCCGAGGTCCCAGCACCCGGTCCCTCCGGGGGGCAGAGACAGGCAGGGCCCCCCGGCAGCTGGCCCCGAGGAGGCGCCCGGAGTGGGGCCGGTCGGCTGGGCTGGCCGA | 12 |
| wt-24750-24870 | 1633 | GCCCGGGTCTGGGAGGTCTGGGGTGGCGAGCCTGCTGTCTCAGGAGGGGCCTGGCTCCGCCGGGTGGCCCTGGGGTAAGTCTGGGAGGCAGAGGGTCGGCCTAGGCCCGGGGAAGTGGAG | 8 |
| wt-24870-24990 | 1634 | GGGGATCGCCCGGGTCTCTGTTGGCAGAGTCCGGGCGATCCTCTGAGACCCTCCGGGCCCGGACGGTCGCCCTCAGCCCCCCAGACAGACCCCAGGGTCTCCAGGCAGGGTCCGGCATCT | 8 |
| wt-24990-25110 | 1635 | TCAGGGGCAGCAGGCTCACCACCACAGGCCCCCCAGACCCGGGTCTCGGCCAGCCGAGCCGACCGGCCCCGCGCCTGGCGCCTCCTCGGGGCCAGCCGCCGGGGTTGGTTCTGCCCCTCT | 12 |
| wt-25110-25230 | 1636 | CTCTGTCCTTCAGAGGAACCAGGGACCTCGGGCACCCCAGAGCCCCTCGGGCCCGCCTCCAGGCGCCCTCCTGGTCTCCGCTCCCCTCTGAGCCCCGTTAAACCCAAAGAATGTCTGAGG | 8 |
| wt-25230-25350 | 1637 | GGAGCCACCCTCGGGGCCCAGGCCCCAGAGTCCAGAGGTCAGGGGCACCTCAGGGTGCCTCCCCGGGTCCCAGGCCAGCCGGAGGGACCCCGGCAGCCCGGGCGGCCCCAGAGGCCGGTT | 12 |
| wt-25350-25470 | 1638 | CCTCGCCCCTTCCCCGGGCTTCAGAGCCCAGGATGTCCCCCAGAAGGGACCCTAGGCGTCCCCTCTCCTCCCCTCCAGGCCCGAGCCTCTCCCTCGCGGAGAGGGGCCTCTTTGGGCCCT | 8 |
| wt-25470-25590 | 1639 | CAAGTCCAGCCCCACCGAGACCCGAGTGGCCCGGATCCCCCCACCGGCCCTTCTCTCTGTCCCCCTGCTCCTCTCCAACCTTCGCTCCACCCTAGACCCCAGCTTCTGGCCTCCCCGGGT | 8 |
| wt-25590-25710 | 1640 | CCACCAGGCCAGCCGGAGGGACCCCGGCAGCCCGGGCGAGTCGCCTTCCCTCTCCCCTGGCCTCTCCTTCCCGCCTCCCACCCGAGCCCCCTCAGCTTGCCTCCCCACCGGGTCCATCAG | 8 |
| wt-25710-25830 | 1641 | GCCGGCCGGAGGGACCCCGGCGGCCCGGTGTCAGTCCCCCCTGCAGCCGCCCAGTCTCTGCCTCCAGGCAAGGGCGCCAGCTTTTCTCCCCCCAGCCTGAGGCCCAGTCTCCTGTGCACT | 8 |
| wt-25830-25950 | 1642 | GTCTGTAAAGTCCAGCCTCCCACGCCCGTCCACGGCTCCCGGGCCCAGCCTCGTCCACCCCTCCCCACGGTGGACAGGCCCTCTGTCCACCCGGGCCATCCCCGCCCCCCTGTGTCCACC | 8 |
| wt-25950-26070 | 1643 | CCAGTCCCGTCCAGGGGGGACTTTATGTGACCCTTGGGCCTGGCTCCCCATAGACTCCCATGTAAGCCTGCCTCGAGTAGGTGCCTCCAGAGCCCCTTTTGCCCCCCTGGCGGCCCAGCC | 8 |
| wt-26070-26190 | 1644 | CGACCCCCGGGCGCCCCCAAACTTTGTCCAGATGTCCAGGGGTCCCCGAGGGTGAGGCCCAGCCCCTCCCGCCCCTGTCCACTGCCCCGGTCCCCCCAGAAGCCCCCAAAAGTAGAGGC | 8 |
| wt-26190-26310 | 1645 | TCAGGCCATGCGCGCCCTGTCACCAGGCCTGCCAAAGAGCCAGATCTAAGGCCGGGAGAGGCAGCCCCAAAGCGGGTGCAGTAACAGGTAATCTCTGGTAGTGATTTGGACCCGAAATCT | 4 |
| wt-26310-26430 | 1646 | GACACTTTAGAGCTCTGGAGGACTTTAAAACTCTAAAAATCAAAACTTTAGAGGCGAATGGGCGCCATTTTGTCCCCACGCGCGCATAATGGCGGACCTAGGCCTAAAACCCCCAGGAAG | 4 |
| wt-26430-26550 | 1647 | CGGGTCTATGGTTGGCTGCGCTGCTGCTATCTTTAGAGGGGAAAAGAGGAATAAGCCCCCAGACAGGGGAGTGGGCTTGTTTGTGACTTCACCAAAGGTCAGGGCCCAAGGGGGTTCGCG | 4 |
| wt-26550-26670 | 1648 | TTGCTAGGCCACCTTCTCAGTCCAGCGCGTTTACGTAAGCCAGACAGCAGCCAATTGTCAGTTCTAGGGAGGGGGACCACTGCCCCTGGTATAAAGTGGTCCTGCAGCTATTTCTGGTCG | 4 |
| wt-26670-26790 | 1649 | CATCAGAGCGCCAGGAGTCCACACAAATGTAAGAGGGGGTCTTCTACCTCTCCCTAGCCCTCCGCCCCCTCCAAGGACTCGGGCCCAGTTTCTAACTTTTCCCCTTCCCTCCCTCGTCTT | 4 |
| wt-26790-26910 | 1650 | GCCCTGCGCCCGGGGCCACCTTCATCACCGTCGCTGACTCCGCCATCCAAGCCTAGGGGAGACCGAAGTGAAGGCCCTGGACCAACCCGGCCCGGGCCCCCCGGTATCGGGCCAGAGGTA | 8 |
| wt-26910-27030 | 1651 | AGTGGACTTTAATTTTTTCTGCTAAGCCCAACACTCCACCACACCCAGGCACACACTACACACACCCACCCGTCTCAGGGTCCCCTCGGACAGCTCCTAAGAAGGCACCGGTCGCCCAGT | 4 |
| wt-27030-27150 | 1652 | CCTACCAGAGGGGGCCAAGAACCCAGACGAGTCCGTAGAAGGGTCCTCGTCCAGCAAGAAGAGGAGGTGGTAAGCGGTTCACCTTCAGGGGTAAGTAACCTGACCTCTCCAGGGCTCACA | 4 |
| wt-27150-27270 | 1653 | TAAAGGGAGGCTTAGTATACATGCTTCTTGCTTTTCACAGGAACCTGGGGGCTAGTCTGGGTGGGATTAGGCTGCCTCAAGTTGCATCAGCCAGGGCTTCATGCCCTCCTCAGTTCCCTA | 4 |
| wt-27270-27390 | 1654 | GTCCCCGGGCTTCAGGCCCCCTCCGTCCCCGTCCTCCAGAGACCCGGGCTTCAGGCCCTGCCTCTCCTGTTACCCTTTTAGAACCACAGCCTGGACACATGTGCCAGACGCCTTGGCCTC | 8 |
| wt-27390-27510 | 1655 | TAAGGCCCTCGGGTCCCCCTGGACCCCGGCCTCAGCAACCCTGCTGCTCCCCTCCTGCCACCCCAGCCTCCCCCCCTCCCCGTCCCCCTTCGCTCCTGATCCTCCCCCGGTCCCCAGTAG | 8 |
| wt-27510-27630 | 1656 | GGCCGCCTGCCCCCCTGCACCCAGTACCTGCCCCTCTTGGCCACGCACCCCGGGCCAGGCCACCTTAGACCCGGCCAAGCCCCATCCCTGAAGACCCAGCGGCCATTCTCTCTGGTAACG | 8 |
| wt-27630-27750 | 1657 | AGCAGAGAAGAAGTAGAGGCCCGCGGCCATTGGGCCCAGATTGAGAGACCAGTCCAGGGGCCCGAGGTTGGAGCCAGCGGGCACCCGAGGTCCCAGCACCCGGTCCCTCCGGGGGGCAGA | 8 |
| wt-27750-27870 | 1658 | GACAGGCAGGGCCCCCCGGCAGCTGGCCCCGAGGAGGCGCCCGGAGTGGGGCCGGTCGGCTGGGCTGGCCGAGCCCGGGTCTGGGAGGTCTGGGGTGGCGAGCCTGCTGTCTCAGGAGGG | 12 |
| wt-27870-27990 | 1659 | GCCTGGCTCCGCCGGGTGGCCCTGGGGTAAGTCTGGGAGGCAGAGGGTCGGCCTAGGCCCGGGGAAGTGGAGGGGGATCGCCCGGGTCTCTGTTGGCAGAGTCCGGGCGATCCTCTGAGA | 8 |
| wt-27990-28110 | 1660 | CCCTCCGGGCCCGGACGGTCGCCCTCAGCCCCCCAGACAGACCCCAGGGTCTCCAGGCAGGGTCCGGCATCTTCAGGGGCAGCAGGCTCACCACCACAGGCCCCCCAGACCCGGGTCTCG | 8 |
| wt-28110-28230 | 1661 | GCCAGCCGAGCCGACCGGCCCCGCGCCTGGCGCCTCCTCGGGGCCAGCCGCCGGGGTTGGTTCTGCCCCTCTCTCTGTCCTTCAGAGGAACCAGGGACCTCGGGCACCCCAGAGCCCCTC | 8 |
| wt-28230-28350 | 1662 | GGGCCCGCCTCCAGGCGCCCTCCTGGTCTCCGCTCCCCTCTGAGCCCCGTTAAACCCAAAGAATGTCTGAGGGGAGCCACCCTCGGGGCCCAGGCCCCAGAGTCCAGAGGTCAGGGGCAC | 8 |
| wt-28350-28470 | 1663 | CTCAGGGTGCCTCCCCGGGTCCCAGGCCAGCCGGAGGGACCCCGGCAGCCCGGGCGGCCCCAGAGGCCGGTTCCTCGCCCCTTCCCCGGGCTTCAGAGCCCAGGATGTCCCCCAGAAGGG | 12 |
| wt-28470-28590 | 1664 | ACCCTAGGCGTCCCCTCTCCTCCCCTCCAGGCCCGAGCCTCTCCCTCGCGGAGAGGGGCCTCTTTGGGCCCTCAAGTCCAGCCCCACCGAGACCCGAGTGGCCCGGATCCCCCCACCGGC | 8 |
| wt-28590-28710 | 1665 | CCTTCTCTCTGTCCCCCTGCTCCTCTCCAACCTTCGCTCCACCCTAGACCCCAGCTTCTGGCCTCCCCGGGTCCACCAGGCCAGCCGGAGGGACCCCGGCAGCCCGGGCGAGTCGCCTTC | 8 |
| wt-28710-28830 | 1666 | CCTCTCCCCTGGCCTCTCCTTCCCGCCTCCCACCCGAGCCCCCTCAGCTTGCCTCCCCACCGGGTCCATCAGGCCGGCCGGAGGGACCCCGGCGGCCCGGTGTCAGTCCCCCCTGCAGCC | 12 |
| wt-28830-28950 | 1667 | GCCCAGTCTCTGCCTCCAGGCAAGGGCGCCAGCTTTTCTCCCCCCAGCCTGAGGCCCAGTCTCCTGTGCACTGTCTGTAAAGTCCAGCCTCCCACGCCCGTCCACGGCTCCCGGGCCCAG | 8 |
| wt-28950-29070 | 1668 | CCTCGTCCACCCCTCCCCACGGTGGACAGGCCCTCTGTCCACCCGGGCCATCCCCGCCCCCCTGTGTCCACCCCAGTCCCGTCCAGGGGGGACTTTATGTGACCCTTGGGCCTGGCTCCC | 8 |
| wt-29070-29190 | 1669 | CATAGACTCCCATGTAAGCCTGCCTCGAGTAGGTGCCTCCAGAGCCCCTTTTGCCCCCCTGGCGGCCCAGCCCGACCCCCGGGCGCCCCCAAACTTTGTCCAGATGTCCAGGGGTCCCCG | 8 |
| wt-29190-29310 | 1670 | AGGGTGAGGCCCAGCCCCTCCCGCCCCTGTCCACTGCCCCGGTCCCCCCAGAAGCCCCCAAAAGTAGAGGCTCAGGCCATGCGCGCCCTGTCACCAGGCCTGCCAAAGAGCCAGATCTA | 8 |
| wt-29310-29430 | 1671 | AGGCCGGGAGAGGCAGCCCCAAAGCGGGTGCAGTAACAGGTAATCTCTGGTAGTGATTTGGACCCGAAATCTGACACTTTAGAGCTCTGGAGGACTTTAAAACTCTAAAAATCAAAACTT | 4 |

| | | | |
|---|---|---|---|
| wt-29430-29550 | 1672 | TAGAGGCGAATGGGCGCCATTTTGTCCCCACGCGCGCATAATGGCGGACCTAGGCCTAAAACCCCCAGGAAGCGGGTCTATGGTTGGCTGCGCTGCTGCTATCTTTAGAGGGGAAAAGAG | 4 |
| wt-29550-29670 | 1673 | GAATAAGCCCCCAGACAGGGGAGTGGGCTTGTTTGTGACTTCACCAAAGGTCAGGGCCCAAGGGGGTTCGCGTTGCTAGGCCACCTTCTCAGTCCAGCGCGTTTACGTAAGCCAGACAGC | 4 |
| wt-29670-29790 | 1674 | AGCCAATTGTCAGTTCTAGGGAGGGGGACCACTGCCCCTGGTATAAAGTGGTCCTGCAGCTATTTCTGGTCGCATCAGAGCGCCAGGAGTCCACACAAATGTAAGAGGGGGTCTTCTACC | 4 |
| wt-29790-29910 | 1675 | TCTCCCTAGCCCTCCGCCCCCTCCAAGGACTCGGGCCCAGTTTCTAACTTTTCCCCTTCCCTCCCTCGTCTTGCCCTGCGCCCGGGGCCACCTTCATCACCGTCGCTGACTCCGCCATCC | 8 |
| wt-29910-30030 | 1676 | AAGCCTAGGGGAGACCGAAGTGAAGGCCCTGGACCAACCCGGCCCGGGCCCCCCGGTATCGGGCCAGAGGTAAGTGGACTTTAATTTTTTCTGCTAAGCCCAACACTCCACCACACCCAG | 4 |
| wt-30030-30150 | 1677 | GCACACACTACACACACCCACCCGTCTCAGGGTCCCCTCGGACAGCTCCTAAGAAGGCACCGGTCGCCCAGTCCTACCAGAGGGGGCCAAGAACCCAGACGAGTCCGTAGAAGGGTCCTC | 4 |
| wt-30150-30270 | 1678 | GTCCAGCAAGAAGAGGAGGTGGTAAGCGGTTCACCTTCAGGGGTAAGTAACCTGACCTCTCCAGGGCTCACATAAAGGGAGGCTTAGTATACATGCTTCTTGCTTTTCACAGGAACCTGG | 4 |
| wt-30270-30390 | 1679 | GGGCTAGTCTGGGTGGGATTAGGCTGCCTCAAGTTGCATCAGCCAGGGCTTCATGCCCTCCTCAGTTCCCTAGTCCCCGGGCTTCAGGCCCCCTCCGTCCCCGTCCTCCAGAGACCCGGG | 8 |
| wt-30390-30510 | 1680 | CTTCAGGCCCTGCCTCTCCTGTTACCCTTTTAGAACCACAGCCTGGACACATGTGCCAGACGCCTTGGCCTCTAAGGCCCTCGGGTCCCCCTGGACCCCGGCCTCAGCAACCCTGCTGCT | 4 |
| wt-30510-30630 | 1681 | CCCCTCCTGCCACCCCAGCCTCCCCCCCTCCCCGTCCCCCTTCGCTCCTGATCCTCCCCCGGTCCCCAGTAGGGCCGCCTGCCCCCCTGCACCCAGTACCTGCCCCTCTTGGCCACGCAC | 8 |
| wt-30630-30750 | 1682 | CCCGGGCCAGGCCACCTTAGACCCGGCCAAGCCCCATCCCTGAAGACCCAGCGGCCATTCTCTCTGGTAACGAGCAGAGAAGAAGTAGAGGCCCGCGGCCATTGGGCCCAGATTGAGAGA | 4 |
| wt-30750-30870 | 1683 | CCAGTCCAGGGGCCCGAGGTTGGAGCCAGCGGGCACCCGAGGTCCCAGCACCCGGTCCCTCCGGGGGGCAGAGACAGGCAGGGCCCCCCGGCAGCTGGCCCCGAGGAGGCGCCCGGAGTG | 12 |
| wt-30870-30990 | 1684 | GGGCCGGTCGGCTGGGCTGGCCGAGCCCGGGTCTGGGAGGTCTGGGGTGGCGAGCCTGCTGTCTCAGGAGGGGCCTGGCTCCGCCGGGTGGCCCTGGGGTAAGTCTGGGAGGCAGAGGGT | 8 |
| wt-30990-31110 | 1685 | CGGCCTAGGCCCGGGGAAGTGGAGGGGGATCGCCCGGGTCTCTGTTGGCAGAGTCCGGGCGATCCTCTGAGACCCTCCGGGCCCGGACGGTCGCCCTCAGCCCCCCAGACAGACCCCAGG | 8 |
| wt-31110-31230 | 1686 | GTCTCCAGGCAGGGTCCGGCATCTTCAGGGGCAGCAGGCTCACCACCACAGGCCCCCCAGACCCGGGTCTCGGCCAGCCGAGCCGACCGGCCCCGCGCCTGGCGCCTCCTCGGGGCCAGC | 12 |
| wt-31230-31350 | 1687 | CGCCGGGGTTGGTTCTGCCCCTCTCTCTGTCCTTCAGAGGAACCAGGGACCTCGGGCACCCCAGAGCCCCTCGGGCCCGCCTCCAGGCGCCCTCCTGGTCTCCGCTCCCCTCTGAGCCCC | 8 |
| wt-31350-31470 | 1688 | GTTAAACCCAAAGAATGTCTGAGGGGAGCCACCCTCGGGGCCCAGGCCCCAGAGTCCAGAGGTCAGGGGCACCTCAGGGTGCCTCCCCGGGTCCCAGGCCAGCCGGAGGGACCCCGGCAG | 8 |
| wt-31470-31590 | 1689 | CCCGGGCGGCCCCAGAGGCCGGTTCCTCGCCCCTTCCCCGGGCTTCAGAGCCCAGGATGTCCCCCAGAAGGGACCCTAGGCGTCCCCTCTCCTCCCCTCCAGGCCCGAGCCTCTCCCTCG | 8 |
| wt-31590-31710 | 1690 | CGGAGAGGGGCCTCTTTGGGCCCTCAAGTCCAGCCCCACCGAGACCCGAGTGGCCCGGATCCCCCCACCGGCCCTTCTCTCTGTCCCCCTGCTCCTCTCCAACCTTCGCTCCACCCTAGA | 8 |
| wt-31710-31830 | 1691 | CCCCAGCTTCTGGCCTCCCCGGGTCCACCAGGCCAGCCGGAGGGACCCCGGCAGCCCGGGCGAGTCGCCTTCCCTCTCCCCTGGCCTCTCCTTCCCGCCTCCCACCCGAGCCCCCTCAGC | 12 |
| wt-31830-31950 | 1692 | TTGCCTCCCCACCGGGTCCATCAGGCCGGCCGGAGGGACCCCGGCGGCCCGGTGTCAGTCCCCCCTGCAGCCGCCCAGTCTCTGCCTCCAGGCAAGGGCGCCAGCTTTTCTCCCCCCAGC | 8 |
| wt-31950-32070 | 1693 | CTGAGGCCCAGTCTCCTGTGCACTGTCTGTAAAGTCCAGCCTCCCACGCCCGTCCACGGCTCCCGGGCCCAGCCTCGTCCACCCCTCCCCACGGTGGACAGGCCCTCTGTCCACCCGGGC | 8 |
| wt-32070-32190 | 1694 | CATCCCCGCCCCCCTGTGTCCACCCCAGTCCCGTCCAGGGGGGACTTTATGTGACCCTTGGGCCTGGCTCCCCATAGACTCCCATGTAAGCCTGCCTCGAGTAGGTGCCTCCAGAGCCCC | 8 |
| wt-32190-32310 | 1695 | TTTTGCCCCCCTGGCGGCCCAGCCCGACCCCCGGGCGCCCCCAAACTTTGTCCAGATGTCCAGGGGTCCCCGAGGGTGAGGCCCAGCCCCTCCCGCCCCTGTCCACTGCCCCGGTCCCC | 8 |
| wt-32310-32430 | 1696 | CCAGAAGCCCCCAAAAGTAGAGGCTCAGGCCATGCGCGCCCTGTCACCAGGCCTGCCAAAGAGCCAGATCTAAGGCCGGGAGAGGCAGCCCCAAAGCGGGTGCAGTAACAGGTAATCTCT | 4 |
| wt-32430-32550 | 1697 | GGTAGTGATTTGGACCCGAAATCTGACACTTTAGAGCTCTGGAGGACTTTAAAACTCTAAAAATCAAAACTTTAGAGGCGAATGGGCGCCATTTTGTCCCCACGCGCGCATAATGGCGGA | 4 |
| wt-32550-32670 | 1698 | CCTAGGCCTAAAACCCCCAGGAAGCGGGTCTATGGTTGGCTGCGCTGCTGCTATCTTTAGAGGGGAAAAGAGGAATAAGCCCCCAGACAGGGGAGTGGGCTTGTTTGTGACTTCACCAAA | 4 |
| wt-32670-32790 | 1699 | GGTCAGGGCCCAAGGGGGTTCGCGTTGCTAGGCCACCTTCTCAGTCCAGCGCGTTTACGTAAGCCAGACAGCAGCCAATTGTCAGTTCTAGGGAGGGGGACCACTGCCCCTGGTATAAAG | 4 |
| wt-32790-32910 | 1700 | TGGTCCTGCAGCTATTTCTGGTCGCATCAGAGCGCCAGGAGTCCACACAAATGTAAGAGGGGGTCTTCTACCTCTCCCTAGCCCTCCGCCCCCTCCAAGGACTCGGGCCCAGTTTCTAAC | 4 |
| wt-32910-33030 | 1701 | TTTTCCCCTTCCCTCCCTCGTCTTGCCCTGCGCCCGGGGCCACCTTCATCACCGTCGCTGACTCCGCCATCCAAGCCTAGGGGAGACCGAAGTGAAGGCCCTGGACCAACCCGGCCCGGG | 8 |
| wt-33030-33150 | 1702 | CCCCCCGGTATCGGGCCAGAGGTAAGTGGACTTTAATTTTTTCTGCTAAGCCCAACACTCCACCACACCCAGGCACACACTACACACACCCACCCGTCTCAGGGTCCCCTCGGACAGCTC | 4 |
| wt-33150-33270 | 1703 | CTAAGAAGGCACCGGTCGCCCAGTCCTACCAGAGGGGGCCAAGAACCCAGACGAGTCCGTAGAAGGGTCCTCGTCCAGCAAGAAGAGGAGGTGGTAAGCGGTTCACCTTCAGGGGTAAGT | 4 |
| wt-33270-33390 | 1704 | AACCTGACCTCTCCAGGGCTCACATAAAGGGAGGCTTAGTATACATGCTTCTTGCTTTTCACAGGAACCTGGGGGCTAGTCTGGGTGGGATTAGGCTGCCTCAAGTTGCATCAGCCAGGG | 4 |
| wt-33390-33510 | 1705 | CTTCATGCCCTCCTCAGTTCCCTAGTCCCCGGGCTTCAGGCCCCCTCCGTCCCCGTCCTCCAGAGACCCGGGCTTCAGGCCCTGCCTCTCCTGTTACCCTTTTAGAACCACAGCCTGGAC | 4 |
| wt-33510-33630 | 1706 | ACATGTGCCAGACGCCTTGGCCTCTAAGGCCCTCGGGTCCCCCTGGACCCCGGCCTCAGCAACCCTGCTGCTCCCCTCCTGCCACCCCAGCCTCCCCCCCTCCCCGTCCCCCTTCGCTCC | 8 |
| wt-33630-33750 | 1707 | TGATCCTCCCCCGGTCCCCAGTAGGGCCGCCTGCCCCCCTGCACCCAGTACCTGCCCCTCTTGGCCACGCACCCCGGGCCAGGCCACCTTAGACCCGGCCAAGCCCCATCCCTGAAGACC | 8 |
| wt-33870-33990 | 1709 | CACCCGGTCCCTCCGGGGGGCAGAGACAGGCAGGGCCCCCCGGCAGCTGGCCCCGAGGAGGCGCCCGGAGTGGGGCCGGTCGGCTGGGCTGGCCGAGCCCGGGTCTGGGAGGTCTGGGGT | 12 |
| wt-33990-34110 | 1710 | GGCGAGCCTGCTGTCTCAGGAGGGGCCTGGCTCCGCCGGGTGGCCCTGGGGTAAGTCTGGGAGGCAGAGGGTCGGCCTAGGCCCGGGGAAGTGGAGGGGGATCGCCCGGGTCTCTGTTGG | 8 |
| wt-34110-34230 | 1711 | CAGAGTCCGGGCGATCCTCTGAGACCCTCCGGGCCCGGACGGTCGCCCTCAGCCCCCCAGACAGACCCCAGGGTCTCCAGGCAGGGTCCGGCATCTTCAGGGGCAGCAGGCTCACCACCA | 8 |
| wt-34230-34350 | 1712 | CAGGCCCCCCAGACCCGGGTCTCGGCCAGCCGAGCCGACCGGCCCCGCGCCTGGCGCCTCCTCGGGGCCAGCCGCCGGGGTTGGTTCTGCCCCTCTCTCTGTCCTTCAGAGGAACCAGGG | 12 |
| wt-34350-34470 | 1713 | ACCTCGGGCACCCCAGAGCCCCTCGGGCCCGCCTCCAGGCGCCCTCCTGGTCTCCGCTCCCCTCTGAGCCCCGTTAAACCCAAAGAATGTCTGAGGGGAGCCACCCTCGGGGCCCAGGCC | 8 |
| wt-34470-34590 | 1714 | CCAGAGTCCAGAGGTCAGGGGCACCTCAGGGTGCCTCCCCGGGTCCCAGGCCAGCCGGAGGGACCCCGGCAGCCCGGGCGGCCCCAGAGGCCGGTTCCTCGCCCCTTCCCCGGGCTTCAG | 12 |
| wt-34590-34710 | 1715 | AGCCCAGGATGTCCCCCAGAAGGGACCCTAGGCGTCCCCTCTCCTCCCCTCCAGGCCCGAGCCTCTCCCTCGCGGAGAGGGGCCTCTTTGGGCCCTCAAGTCCAGCCCCACCGAGACCCG | 8 |
| wt-34710-34830 | 1716 | AGTGGCCCGGATCCCCCCACCGGCCCTTCTCTCTGTCCCCCTGCTCCTCTCCAACCTTCGCTCCACCCTAGACCCCAGCTTCTGGCCTCCCCGGGTCCACCAGGCCAGCCGGAGGGACCC | 8 |
| wt-34830-34950 | 1717 | CGGCAGCCCGGGCGAGTCGCCTTCCCTCTCCCCTGGCCTCTCCTTCCCGCCTCCCACCCGAGCCCCCTCAGCTTGCCTCCCCACCGGGTCCATCAGGCCGGCCGGAGGGACCCCGGCGGC | 12 |
| wt-34950-35070 | 1718 | CCGGTGTCAGTCCCCCCTGCAGCCGCCCAGTCTCTGCCTCCAGGCAAGGGCGCCAGCTTTTCTCCCCCCAGCCTGAGGCCCAGTCTCCTGTGCACTGTCTGTAAAGTCCAGCCTCCCACG | 8 |
| wt-35070-35190 | 1719 | CCCGTCCACGGCTCCCGGGCCCAGCCTCGTCCACCCCTCCCCACGGTGGACAGGCCCTCTGTCCACCCGGGCCATCCCCGCCCCCCTGTGTCCACCCCAGTCCCGTCCAGGGGGGACTTT | 8 |
| wt-35190-35310 | 1720 | ATGTGACCCTTGGGCCTGGCTCCCCATAGACTCCCATGTAAGCCTGCCTCGAGTAGGTGCCTCCAGAGCCCCTTTTGCCCCCCTGGCGGCCCAGCCCGACCCCCGGGCGCCCCCAAACTT | 8 |
| wt-35310-35430 | 1721 | TGTCCAGATGTCCAGGGGTCCCCGAGGGTGAGGCCCAGCCCCTCTCGCCCAAGCTGCTTTGATTCCTGGGATATTTTTGGGAATGGTGTTAACTTTCTCCCCTTGTATTTGCTATTCAA | 4 |
| wt-35430-35550 | 1722 | TCAACCTGATTCCCCCTGCTCATACCTCCACTTACAACCAAGCCACTACGGCCACGTCCCCGGCCTCCCGCTCGGGTAAGTGCTTTTTCATTTTTAGCCCCAGCCCCTCCTCTATAAGTT | 4 |
| wt-35550-35670 | 1723 | CTAGGCAAACCTCCAATCACCAGCCACCTTCCAATGTAGTCTCTTAGAGAGTGGCTGCTACGCATTAGAGACCACTTTGAGCCACCCACAGTAACCACCCAGCGCCAATCTGTCTACATA | 4 |

| | | | |
|---|---|---|---|
| wt-35670-35790 | 1724 | GAAGAAGAAGAGGATGAAGACTAAGTCACAGGCTTAGCCAGGTGATTTGTGAATTTCAGTTTATTTACTTTCTTCCAATCAAGCTTTCCCAGCCTCCGCTTGTTAGGTCCTAGTTATGGG | 4 |
| wt-35790-35910 | 1725 | TTTTCCATGGGGGACTTAGTATCCGTTCTATTAGATTAACGTGCAAGACGCTAAACTTAACCAAGGTCAGCCAAGGGACGCGTGTTATCCCAGGCTGCCCACCCTGAGGATTTCCCCCCA | 4 |
| wt-35910-36030 | 1726 | AAATCCTCCTACCCTCTCTTTATGCCATGTGTGTTGTTGGCTTGTGTTAGTGCTATGTAATGCGTTGCCGCCAGGTGGCAGCCTGTTTATAGATGTGCAGTACCCCTTAATGTTAGGTCT | 4 |
| wt-36030-36150 | 1727 | GCTTTAGGGCTGCCAGGTGGCGCAATCTAGGATTAATTCACCTGTATCCCTTTCCCTCCACCCGCAGTAACCCAGCACTGGCGTGTGACGTGGTGTAAAGTTTTGCCTGAACCTGTGGTT | 4 |
| wt-36150-36270 | 1728 | GGGCAGGTACATGCCAACAACCTTCTAAGCACCCGCGCTTGTGTTTTGCTTTATCTGCCGCCATCATGCCTACATTCTATCTTGCGTTACATGGGGGACAAACATATCATCTAATTGTTG | 4 |
| wt-36270-36390 | 1729 | ACACGGATAGTCTTGGAAACCCGTCACTCTCAGTAATTCCCTCGAATCCCTACCAGGAACAACTGTCAGACACTCCATTAATTCCACTAACAATCTTTGTTGGGGAAAACACGGGGGTGC | 4 |
| wt-36510-36630 | 1730 | CACCTCAGCGCAGGGATGCCTGGACACAAGAGCCATCACCTCTTGATAGGGATCCGCTAGGATATGACGTCGGGCATGGACCTCTAGCATCTGCTATGCGAATGCTTTGGATGGCTAATT | 4 |
| wt-36630-36750 | 1731 | ATATTGTAAGACAATCACGGGGTGACCGGGGCCTTATTTTGCCACAAGGCCCACAAACAGCCCCTCAGGCCAGGTTGGTCCAGCCACATGTCCCCCCTCTACGCCCGACAGCACCCACCA | 4 |
| wt-36750-36870 | 1732 | TTTTGTCACCTCTGTCACAACCGAGGCTTACCCCTCCACAACCACTCATGATGCCACCAAGGCCTACCCCTCCTACCCCTCTGCCACCTGCAACACTAACGGTGCCACCAAGGCCTACCC | 4 |
| wt-36870-36990 | 1733 | GTCCTACCACTCTGCCACCCACACCACTACTCACGGTACTACAAAGGCCTACCGAACTTCAACCCACACCATCACCACCACGCATGCATCTCCCTGTCTTGCATGTGCCAGACCAATCAA | 4 |
| wt-36990-37110 | 1734 | TGCACCCTCTTACTCATCAAAGCACCCCAAATGATCCAGATAGTCCAGAACCACGGTCCCCGACTGTATTTTATAACATTCCACCTATGCCATTACCCCCCTCACAATTGCCACCACCAG | 4 |
| wt-37110-37230 | 1735 | CAGCACCAGCACAGCCACCTCCAGGGGTCATCAACGACCAACAATTACATCATCTACCCTCGGGGCCACCATGGTGGCCACCCATCTGCGACCCCCCGCAACCCTCTAAGACTCAAGGCC | 4 |
| wt-37350-37470 | 1736 | GTCCTAGCATGCCTGAACTAAGTCCAGTCCTCGGTCTTCATCAGGGACAAGGGGCTGGGGACTCACCAACTCCTGGCCCATCCAATGCCGCCCCCGTTTGTAGAAATTCACACACGGCAA | 4 |
| wt-37470-37590 | 1737 | CCCCTAACGTTTCACCAATACATGAACCGGAGTCCCATAATAGCCCAGAGGCTCCCATTCTCTTCCCCGATGATTGGTATCCTCCATCTATAGACCCCGCAGACTTAGACGAAAGTTGGG | 4 |
| wt-37590-37710 | 1738 | ATTACATTTTTGAGACAACAGAATCTCCTAGCTCAGATGAAGATTATGTGGAGGGACCCAGTAAAAGACCTCGCCCCTCCATCCAGTAAAAACCCTTGCCCTCTCCAGCAACCAATGTAT | 4 |
| wt-37710-37830 | 1739 | CCCAAATAAATGTTACTTCTTTTGCTCTTAACCATTGACACGCCTGTCATTCTATCAATTAAACAAGGGAAAAAGGTTTAGCTATTCCACCAACACGACCCCAAGGAAGGCTTGCCAAAA | 4 |
| wt-37830-37950 | 1740 | TTGGTGCCTTGCTCTCAGCACTTTGCCAGCAACTTATAGCATGGTAGGCAGCTCAACTCGGCCCGTCTTACTGCCCAGCCTACTCTCCACTCCCAGTCCATGTTCGCACTCCTATGCATT | 4 |
| wt-37950-38070 | 1741 | TCCTGCCCTCCCACTTTTACCCCAGTCCCAACCCAAAACCACACACAACACATAGAATTGTTAGTTTAAACAGTTTATTGATAGGTGGCTGCTTTTAGCCTAATTGTGTATTGCTCTCGT | 4 |
| wt-38070-38190 | 1742 | TGCCAAAACCTGTTGTAAGGGCCGGCACCCGCAACATGGGGAAAACATAACCGCCGCCATCCCATGGGGAGGGTAGAGGCGGTTGACATGTAGGTGAGTAGTGTAAGAAGCATGGCGAAG | 4 |
| wt-38190-38310 | 1743 | TAGACAGGTTACTTTTAGAGTGTAGTGTACAGGGCCGGGCGCAACAGTGCCACCAACCCGGGGTCTGAGCATTCCATGGGCAGCAGGGACACTGCACTACCGCCAGGTCCTGGGGCAGCC | 4 |
| wt-38310-38430 | 1744 | GGGGTTCCTGGCGCTCCGGGGGCAGCCGGGCGGCCGCCGGTGGGTCCGCTGGGCCGCTGCCCCGCTCCGGGTGGGGGGTGGCCCCGCTGGGCACCGCTGCGCCGCCGCCAGGTCCTGGGG | 12 |
| wt-38430-38550 | 1745 | CAGCCGGGGTTCCTGGCGCTCCGGGGGCAGCCGGGCGGCCGCCGGTGGGTCCGCTGGGCCGCTGCCCCGCTCCGGGTGGGGGGTGGCCCCGCTGGGCACCGCTGCGCCGCCGCCAGGTCC | 12 |
| wt-38550-38670 | 1746 | TGGGGCAGCCGGGGTTCCTGGCGCTCCGGGGGCAGCCGGGCGGCCGCCGGTGGGTCCGCTGGGCCGCTGCCCCGCTCCGGGTGGGGGGTGGCCCCGCTGGGCACCGCTGCGCCGCCGCCA | 12 |
| wt-38670-38790 | 1747 | GGTCCTGGGGCAGCCGGGGTTCCTGGCGCTCCGGGGGCAGCCGGGCGGCCGCCGGTGGGTCCGCTGGGCCGCTGCCCCGCTCCGGGTGGGGGGTGGCCCCGCTGGGCACCGCTGCGCCGC | 12 |
| wt-38790-38910 | 1748 | CGCCAGGTCCTGGGGCAGCCGGGGTTCCTGGCGCTCCGGGGGCAGCCGGGCGGCCGCCGGTGGGTCCGCTGGGCCGCTGCCCCGCTCCGGGTGGGGGGTGGCCCCGCTGGGCACCGCTGC | 12 |
| wt-38910-39030 | 1749 | GCCGCCGCCAGGTCCTGGGGCAGCCGGGGTTCCTGGCGCTCCGGGGGCAGCCGGGCGGCCGCCGGTGGGTCCGCTGGGCCGCTGCCCCGCTCCGGGTGGGGGGTGGCCCCGCTGGGCACC | 12 |
| wt-39030-39150 | 1750 | GCTGCGCCGCCGCCAGGTCCTGGGGCAGCCGGGGTTCCTGGCGCTCCGGGGGCAGCCGGGCGGCCGCCGGTGGGTCCGCTGGGCCGCTGCCCCGCTCCGGGTGGGGGGTGGCCCCGCTGG | 12 |
| wt-39150-39270 | 1751 | GCACCGCTGCGCCGCCGCCAGGTCCTGGGGCAGCCGGGGTTCCTGGCGCTCCGGGGGCAGCCGGGCGGCCGCCGGTGGGTCCGCTGGGCCGCTGCCCCGCTCCGGGTGGGGGGTGGCCCC | 12 |
| wt-39270-39390 | 1752 | GCTGGGCACCGCTGCGCCGCCGCCAGGTCCTGGGGCAGCCGGGGTTCCTGGCGCTCCGGGGGCAGCCGGGCGGCCGCCGGTGGGTCCGCTGGGCCGCTGCCCCGCTCCGGGTGGGGGGTG | 12 |
| wt-39390-39510 | 1753 | GCCCCGCTGGGCACCGCTGCGCCGCCGCCAGGTCCTGGGGCAGCCGGGGTTCCTGGCGCTCCGGGGGCAGCCGGGCGGCCGCCGGTGGGTCCGCTGGGCCGCTGCCCCGCTCCGGGTGGG | 12 |
| wt-39510-39630 | 1754 | GGGTGGCCCCGCTGGGCACCGCTGCGCCGCCGCCAGGTCCTGGGGCAGCCGGGGTTCCTGGCGCTCCGGGGGCAGCCGGGCGGCCGCCGGTGGGTCCGCTGGGCCGCTGCCCCGCTCCGG | 12 |
| wt-39630-39750 | 1755 | GTGGGGGGTGGCCCCGCTGGGCACCGCTGCGCCGCCGCCAGGTCCTGGGGCAGCCGGGGTTCCTGGCGCTCCGGGGGCAGCCGGGCGGCCGCCGGTGGGTCCGCTGGGCCGCTGCCCCGC | 12 |
| wt-39750-39870 | 1756 | TCCGGGTGGGGGGTGGCCCCGCTGGGCACCGCTGCGCCGCCGCCAGGTCCTGGGGCAGCCGGGGTTCCTGGCGCTCCACTGCACCTGGAATGCAGGGTGGGGGCGTGGTCCCCTGGACCC | 12 |
| wt-39870-39990 | 1757 | CAGCCCCGCCGATCCCTCCCCCAGGGCGTACCCGGCTTGCCTGGTTCTGGGGCTCCTCTGGGGGTCGCTGCATCCGCCGGTAGGGTTCGAATGGGCGTGGTCCGCTTGCTCTGCTGGCCC | 8 |
| wt-39990-40110 | 1758 | GGTACGCCTGGATTGCCGGCTGGGGGCTGGGGTCCCGGGACGCCCCCTCCCTGCTCCCACCCGGTTCCCTCCCCCAGGGCGTGCCCCGCTTGCCTGGTCCTGGAGCTCATCCGGGGATGC | 8 |
| wt-40110-40230 | 1759 | TGCATCCGCTAGTCCGACCTGGGTGGGTGCGGTCCGCTGGCCCCACCCTGGGGGTAGCCGCCGGGTCTGCTGGTCCGGTGCACCTGGAAGGCAGGGGGGGGGGCAGTGAGGGAGGGGCGT | 8 |
| wt-40230-40350 | 1760 | GGTCCTGGGACCCCGCGCCGACTGGCAGGGGGTCCCCATGGCACAGGCCTAGGGGTCCAGGGGGCAGCCGCGGCCCAGCGCGCCCCGTTCACGGGGGAGGACCGCGGCCGAGCCACCAGG | 12 |
| wt-40350-40470 | 1761 | GGCCCGGCGGGGGTGGGGGGTGCGCTCCCAGGCCGGACCCTGGTGCCAGGCAGGGACCCCGCGCCACCCGCTTCATGGGGGGGGAGGCCGCCGCAAGGACGCCGGGCCGGCTGGGAGGTG | 12 |
| wt-40470-40590 | 1762 | TGCACCCCCCGAGCGTCTGGACGACGCTGGCGAGCCGGGCCGGCTCGCCTTCTTTTATCCTCTTTTTGGGGTCTCTGTGTAATACTTTAAGGTTTGCTCAGGAGTGGGGGCTTCTTATTG | 4 |
| wt-40590-40710 | 1763 | GTTAATTCAGGTGTGTCATTTTAGCCCGTTGGGTTTCATTAAGGTGTGTCACCAGGTGGGTGGTACCTGGAGGTTATTCTATTGGGATAACGAGAGGAGGAGGGGCTAGAGGTCCGCGAG | 4 |
| wt-40710-40830 | 1764 | ATTTGGGGTAGGCGGAGCCTCAGGAGGGTCCCCTCCATAGGGTTGAACCAGGAGGGGGAGGATTGGGCTCCGCCCCGATATACCTAGTGGGTGGAGCCTAGAGGTAGGTATCCATAGGGT | 4 |
| wt-40830-40950 | 1765 | TCCATTATCCTGGAGGTATCCTAAGCTCCGCCCCTATATACCAGGTGGGTGGAGCTAGGTAGGATTCAGCTAGGTTCCTACTGGGGTACCCCCCTACCCTACCTTAAGGTGCGCCACCCT | 4 |
| wt-40950-41070 | 1766 | TCCTCCTTCCGTTTTAATGGTAGAATAACCTATAGGTTATTAACCTAGTGGTGGAATAGGGTATTGCAGCTGGGTATATACCTATAGGTATATAGAACCTAGAGGAAGGGAACCCTATAG | 4 |
| wt-41070-41190 | 1767 | TGTAATCCCTCCCCCCCCTACCCCCCCCTCCCTTACGGTTGCCTGAGCCCATCCCCCACCCCAGCACCCCGGGGTGACGTGGCACCCCGCGTGCCTTACTGACTTGTCACCTTTGCACAT | 8 |
| wt-41190-41310 | 1768 | TTGGTCAGCTGACCGATGCTCGCCACTTCCTGGGTCATGACCTGGCCTGTGCCTTGTCCCGTGGACAATGTCCCTCCAGCGTGGTGGCTGCCTTTGGGATGCATCACTTTGAGCCACTAA | 4 |
| wt-41310-41430 | 1769 | GCCCCCGTTGCTCGCCTTGCCTGCCTCACCATGACACACTAAGCCCCTGCTAATCCATGAGCCCCGCCTTTAGGAAGCACCACGTCCCGGGGACGGAAGGGGACTTGGGGTGATTTTCTA | 4 |
| wt-41430-41550 | 1770 | TGTGGGGGTGGAAATATGAGCAAGAATAAGGACGGCTCCTTATTAACCTGATCAGCCCCGGAGTTGCCTGTTTCATCACTAACCCCGGGCCTGAAGAGGTTGACAAGAAGGGTCAAGGTT | 4 |
| wt-41550-41670 | 1771 | TCGTCTGTGTGTTGAAGGGCAGGGGCTGTTGGGTGCATCTGGAACGGCTTACCTCGGGTAACTGTTTGCCATTAAAAGGTTGGGGATTAGGTTTAGCCCCTTTAGCTGCCATTTCGAACC | 4 |
| wt-41670-41790 | 1772 | GGGGTGTGCAGATGCAGGTCTCCGGGTGGGCAGGCAGTACGAGATGTCACGTTGTGTTGTCTTTCCTCCCACCCCTGTCCTGGCTGTGGCAAATGCGACCCTCATAGAGTTGTGTTTCAG | 4 |
| wt-41790-41910 | 1773 | GTCTGTGTCCTGTTTTGCGGTGGGTTATTTCTTCCCTCAGTGTTTGCCAGCTTATTTCCCCAGTTTTCACGTACTGGGGCCTGTGGACACCTGAGGGAGCGGCCGTTGGTGGGTATGTGT | 4 |
| wt-41910-42030 | 1774 | TGGAATTGCTCCCACCCTCAATTTTCGCTTGCCTTCTTCCCTTGTTAACCTGATAGCATAGCCTCTAGGTTTCCTTGTAGGTCTGTTTGGGTTTGTTGGTTCACGTGGTGCTAACTTGAA | 4 |

| | | | |
|---|---|---|---|
| wt-42030-42150 | 1775 | TTTTTTGGTTTTCTAGTTCCCTCTTAATTACATTTGTGCCAGATCTTGTAGAGCAAGATGGCCTATTCAACAAGGGAGATACTGTTAGCCCTGTGTATACGGGACAGTCGTGTGCATGGA | 4 |
| wt-42150-42270 | 1776 | AATGGTACCCTGCATCCTGTGTTGGAGCTAGCAGCAAGAGAAACACCTCTCCGCCTTTCGCCAGAGGACACTGTAGTTCTGCGTTATCATGTGTTGCTTGAGGAGATAATTGAACGAAAT | 4 |
| wt-42270-42390 | 1777 | TCAGAGACATTTACAGAAACTTGGAACAGATTTATAACACACACCGAACATGTGGATCTGGATTTTAACTCAGTATTTTTAGAGATATTTCACCGTGGAGACCCAAGCCTTGGGCGCGCG | 4 |
| wt-42390-42510 | 1778 | TTGGCCTGGATGGCCTGGTGCATGCATGCCTGCAGGACATTGTGTTGTAACCAGTCTACTCCTTACTATGTTGTGGACCTGTCAGTTCGTGGGATGTTAGAAGCCAGCGAAGGCCTGGAT | 4 |
| wt-42510-42630 | 1779 | GGTTGGATTCATCAACAGGGCGGCTGGTCTACATTAATTGAAGACAACATTCCTGGATCCAGAAGGTTTAGCTGGACTTTGTTTCTTGCTGGACTGACTTTGAGTCTGTTAGTTATATGT | 4 |
| wt-42630-42750 | 1780 | AGTTATTTATTTATCTCCAGAGGAAGACACTAATCTATACATTTTCTCAGCACTTTATATGAATCAGGGTCATTGGGCCTGCGGGGAACTGAGCCAGTAGGATATTAGGCAAGGGTGACA | 4 |
| wt-42750-42870 | 1781 | CAGTGCCCATGCATTATAATTTAACCAAACAGTGGTCGTGAGTTTTAGGCCGGCCATGGGGGCTTACAAGAATAACATGCCAATGACCCGGCCCCCACTTTTAAATTCTGTTGCAGCAGA | 4 |
| wt-42870-42990 | 1782 | TAGCTGATACCCAATGTTATCTTTTGCGGCAGAAATTGAAAGTGCTGGCCATATCTACAATTGGGTGTCCTAGGTGGGATATACGCCTGTGGTGTTCTAACGGGAAGTGTGTAAGCACAC | 4 |
| wt-42990-43110 | 1783 | ACGTAATTTGCAAGCGGTGCTTCACGCTCTTCGTTAAAATAACACAAGGACAAGATACTAAAGAAATAACTGAGGTGAGTGTGGGAAGATGGGAATACTATGTGTTATGTTAACGGGTGA | 4 |
| wt-43110-43230 | 1784 | GAGCCTATACTGCAGCCCAGACTCGGGGGGAGGAGGAAATGGTAAGAGTTATACTCTACTTATCTTTTTTGACACTACATTTAACTGTTATGTAACAATGTTTGCTTATTTTCATGTTCA | 4 |
| wt-43230-43350 | 1785 | ATAAACGCTATGTTAATGATGAAGAACCTGTGTTCTTTGGAAGTGGGCCCAATGGGGTAGTAGGTTTTGGGAGGGTGCCGTGCTAGATATTTCAACTGCCACAGACCCCATTTTGTCCCA | 4 |
| wt-43350-43470 | 1786 | CCTGTTACCACATTCTAGGTCCTGCATCCAGTGGGCCAGGTGTCTCACCATGGCTCTTTCTAGGTGGATACCACAGTCCAGGCCCCCAAGGCTACCGTGCTAATTACCTCCTCATGTCCA | 4 |
| wt-43470-43590 | 1787 | CCCCCACCCTGTGTTACTGTCGCCTGATTATCCTGGCTTAGCAGCCTCCAAGTTTTACAAGACGTCCCATTGCCCTGCCCTTGGTCCAAGTCTCGCCGGTTTTCAGCAGCCTGTTGTAGC | 4 |
| wt-43590-43710 | 1788 | CTGCCCCCAAGTTTCGCAGGTTTCCCCCATGCTTCCACCCGTTAACCCAATAGCATGACAGCCAATCCAACACGAGGCAAGTTTTAAGAGTTAAAAGCAACTACTGTTTATTTTCCAAAA | 4 |
| wt-43710-43830 | 1789 | TGAGCTGGGTATAGTTGATGATCTGTAGGCGCAGCTCATCCCCACATTCCAGGTCCTTGATGGCCTCGTAGATGGCATCTTCGTCGACATTGACAGCCTTCTCATATACCGTGTCTCTGG | 4 |
| wt-43830-43950 | 1790 | GGCTGACCTTTATACAGAAGGCGTCCCCTACTAGGTCCACGGCCAGCTCGTAGGTGGGGCCTATGTTTTCACATAACAGTTTCAAGCAGGTCTCTGGGATGTGAAGGGAGGTGCCCTGGA | 4 |
| wt-43950-44070 | 1791 | GCAGGAGATGCATGATTAGGCGCCCTTTTCCATTTGTGCTGAAGATGGGGCAGATGGTGCCACAAAAGTGTCCGGTGACCAGGTAAGCGTAGAGAAGGCTGGGTTGGGAAAGTCCAGCCT | 4 |
| wt-44070-44190 | 1792 | TTACTGCACTGGGAGAGCTGCTGAGCAGAGACACATAGAAGGTCTTGTTGGGTATTATCTTGTGGACATTGTTGAAGAAGGAGAGCTGGGTGGAGCTAAACTCCTGAGGCACATGAACCT | 4 |
| wt-44190-44310 | 1793 | GGGACCTATTGATGCAGATCTCGCAGTGAGACCCCAGAGTCAGGCTGTGGCCGAAGGGAGACAGGCGAAGGCAGCGCCCGGGGGAGAGAGTGCACAGTGACAGTGGGAGAAACACGGCCT | 4 |
| wt-44310-44430 | 1794 | CTGAGACATGTATGGGGGTGTTCATCTCACGCAGAAAATCTTTGCCCAGCTCAAAGTTGGCAGAGATTCCCCTGAAGAAGTCCCGTAGTGAAAAATGGGATCTGTCTACACCATGTCTGG | 4 |
| wt-44430-44550 | 1795 | TGTGCCGGGAACATATTGATCGGGCCACACTGCCAACCCTTTCCATTCTTCCCAGCTCTGAGCGAGATTTTCCACACCTGGACACCGACTTCACGCTATGCGCCGAGGCCTTTGAGGCCG | 4 |
| wt-44550-44670 | 1796 | TGTAGTTTCTGTGGTGCGGATGCATTAGGCGGCGCAATGCGGGATCTGCCGGTCGCTGTTGGCGTGCATTCACGGCATCTGGGGTGACCGGGGCCATCGGGTTTACTTTTCACACGTAGA | 4 |
| wt-44670-44790 | 1797 | CCTGGGAAGTTTGATAGGACTGTACCAGGTCAAGGCCGTGGATGCGCAGGACCACGTCCAGTTCCTTAGTGACATCCACGAGGATTGTTTTGCCCACTCTGGCCACTTGTGTGGATTTAA | 4 |
| wt-44790-44910 | 1798 | ATATGTACACAAGCGTAATTAACGAGTCACAGACCCCCTGTTCCAGATTCTGACCGGCTGCAAGCGCTGCCTTAAAGGCCTGGAAGCTGGGTGGGTAAATCTGACCAAACAGCACGCTCG | 4 |
| wt-44910-45030 | 1799 | GATTCGTGATGCTGTGGTTGATGGCACACAGGGGGTCGCAGAACAGGTGCTTGTGGAAGTCTTGCGGGGTGCACATCTGCAGCCAGGCCCTTAGCCTGGGGCATGGCACATCCAGCAGCG | 4 |
| wt-45030-45150 | 1800 | TGTTTTGGGTCTTGATGAGGAACACGATCCTGTCTAGGATTTTGATGTTGTTGCCGAACGAGTCAAGAATCAGGCTCTTGAAGCGGTCAAGGGTGTCCTTGGCGTCCGGGTGGGCCCCGA | 4 |
| wt-45270-45390 | 1802 | CCAACTCCGCCAGCTCATCGCCGCTGTATTCCGCGTTTAAACCGATAGCATGGTGGCCTGGCCCCCCGAGCAGGTCCGTCCCCTGCCACGTACCTAATAGTAGTCCACAGTAGTCGGCCT | 4 |
| wt-45390-45510 | 1803 | TGGTTGTAATTTCAGGAGAGAGTCCTCCCTTTTCGGCCCTGAGAAATGGATGCTGAACTCGGTTTCTGGTAGGCAGGTGGCAGCACAGGGCGGTGTACAGGCCCCTGCCGACGTCCCCTG | 4 |
| wt-45510-45630 | 1804 | GGACATCCTGGGAATCTTTGCAGGTTCTGGGTCCAGGGAGGGTAAGAAAAGTGGGGGTGGTTCTGGGCCACATGGACTTGAAGCAGAAGTTGGCCGGGGACTGGCCGGTGAGGATGGATT | 4 |
| wt-45630-45750 | 1805 | TCAGAAACTCCAATTTGTAGTAGCCGAGGTTGGCATTTCTAATCATGTCAGAAGAGGACACAGGGAGGAAGCACCGGCAAATGTAAAAGTGAAGCTGGATGTCAATGGCAAGAATCCTGG | 4 |
| wt-45750-45870 | 1806 | AGGGCATGAAGAGGGAATCCAACCCCCCGGCCATGGGGAAGTATTTTATCAGGATGTGTAAAAAGTCCATGCCTGTGATGAGGCTAGAGATCCAGGCTCGTGGGGCATTTAGACAGTAGT | 4 |
| wt-45870-45990 | 1807 | AGCAGAGCAGGGCATAGTCCTCAAAGAAGGCCACGGGGGCATCTGAGTGATTGACCAGGGTGTCGAGCAGATCACAAACTCGGCAGGTGCTGGCTGGAGAGAGGGACTCGTAGGTGTGGA | 4 |
| wt-45990-46110 | 1808 | CGAGTGGTGGGTAGGCTATGCCTTCTTCCGCGTTGGCTGGAAGATAGGAGTGGGCCATCAAAAGGCCGACTGCCTCGAACTGGCTTTTCAGATTGTCCACGGTCCAGGGCACAAAGTCCT | 4 |
| wt-46110-46230 | 1809 | CCATCTTTGGAGTTCTGCCCGCGATCTGTGCCACCTCTGTTACGCCACTCCTCGTGAGGGGGCAGCTGGACAGTCTTTTTCCGGTCAGGGGGTTTGGCTCGTTTGCGCTCGTGACTTTGT | 4 |
| wt-46230-46350 | 1810 | GAGCCATGACACATCTGGGTGGCAAGGTGAGGTCTTCTGGGTTTTTAATACCGGGGTCGGCACCAGTTTCTGGGACACCGCCACAAGGACAAGGTGGGCTAGCAAGTTCTCGAGTCTACG | 4 |
| wt-46350-46470 | 1811 | AAGACTCCGGGGGCAGTCTTTTGAGTTTCTCGCCTATGATCCACCCCAATCTCGCCCCCCTAATTGCGCCATCTGCCTACGCGAGGCTGAACCTCCTGAATCACTGCATCTTTCTTGAGG | 4 |
| wt-46470-46590 | 1812 | CGTTTAAAGAAGAGAATAGTGGCCAGGGCCTCGGTGGGGTCCAGCGTGAGGTCTTATTTTTGAAAAGGGATATTATAAAACAGGTCATTGCTCGGATTGTGGCAGCCGATAGCACCCTAG | 4 |
| wt-46590-46710 | 1813 | ATCTAGTGAATCATGGCGAGCCCGGAAGAGAGGCTCCTAGACGAGCTCAATAACGTAATTGTGTCATTTCTGTGTGACTCTGGGTCTCTGGAAGTGGAGAGATGCTCCGGGGCGCATGTG | 4 |
| wt-46710-46830 | 1814 | TTCTCCAGGGGGCAGCTCCCAACCCCTCTGCACCGTGAAGCTGCGCCACGGACAGATTTACCACCTGGAGTTTGTCTACAAGTTCCTGGCCTTTAAGCTGAAGAACTGCAACTACCCCTCC | 4 |
| wt-46830-46950 | 1815 | TCGCCCGTGTTTGTGATATCCAACAACGGCCTGGCCACCACCCTGAGGTGCTTTTTGCACGAGCCGTCGGGTCTCAGATCGGGCCAGAGCGGCCCCTTGCCTGGGTCTCTCAACGGATGTT | 4 |
| wt-46950-47070 | 1816 | GACCTACCAAAGAACTCCATCATTATGCTGGGCCAGGATGACTTCATTAAGTTCAAAAGCCCCCTGGTCTTCCCTGCTGAGCTTGATCTCCTGAAATCTATGGTGGTCTGCCGGGCCTAC | 4 |
| wt-47070-47190 | 1817 | ATCACGGAACACCGGACGACGATGCAGTTTCTGGTGTTTCAGGCCGCCAACGCCCAGAAGGCCTCGCGGGTCATGGATATGATTAGTGATATGTCTCAGCAACTGTCTCGGTCTGGTCAA | 4 |
| wt-47190-47310 | 1818 | GTCGAGGATACGGGCGCCAGAGTCACAGGTGGAGGAGGTCCCAGGCCTGGCGTCACGCACTCGGGGTGTCTTGGGGACTCACACGTTAGGGGGCGCGGTGGTTGGGACTTGGATAACTTT | 4 |
| wt-47310-47430 | 1819 | TCAGAAGCTGAGACCGAAGACGAGGCGAGTTACGCTCCTTGGAGGGACAAAGACTCGTGGTCGGAATCCGAGGCGGCGCCGTGGAAGAAGGAACTCGTGAGGCACCCCATCCGCAGGCAC | 4 |
| wt-47430-47550 | 1820 | CGGACACGCGAGACTCGCCGTATGCGCGGGAGCCATTCACGGGTGGAACACGTGCCCCCCGAGACCCGGGAGACGGTGGTGGGGGGAGCATGGCGTTATTCTTGGCGCGCCACACCTTAT | 8 |
| wt-47550-47670 | 1821 | CTGGCACGGGTGCTGGCTGTCACGGCCGTGGCCCTGCTCCTGATGTTTCTGAGGTGGACCTGACGTTGCAGGCCCTTGGGGAGCGGGGGTTCTCCAGGCTCCTGGATCTGGGGCTGGCCT | 8 |
| wt-47670-47790 | 1822 | GCCTGGATCTGAGCTATGTGGAAATGAGGGAATTTGTGGTTTGGGGCAGGCCCCCAGCTTCTGAGGCGGCTGTGGCCTCTACGCCAGGCTCGCTTTTCCGAAGCCACTCGTCCGCCTACT | 4 |
| wt-47790-47910 | 1823 | GGTTGTCGGAGGTGGAGAGGCCCGGGGGCCTTGTCCGCTGGGCCAGGTCACAGACCAGCCCCTCATCCCTGACCCTCGCGCCCCATCTTGGCCCGTCCCTCTTGTCCCTTTCAGTGGTCA | 8 |
| wt-47910-48030 | 1824 | CCGGTGGTGGGTGTGGAGCCGTGGCCTTTTGCAACGCCTTTTTCCTAGCTTATTTTTTGGTTGTGCGGTCTGTTTTCCCCGCGTTTTCCGATAGAATAGCTGCCTGGATCTGCGACCGGT | 4 |
| wt-48030-48150 | 1825 | CCCCTTTCTGCGAAAACACCCGGGCCGTGGCCAGGGGTTACCGAGGCCTCGTGAAGAGGTTCTTGGCATTCGTGTTTGAGCGTAGTAGCTATGACCCCCCCTTGTTGAGGCAAAACTCTA | 4 |
| wt-48150-48270 | 1826 | GGCCTGTGGAGCGCTGCTTTGCCATCAAGAATTATGTCCCGGGCCTGGACTCACAAAGCTGTGTGACGGTCCCGAGCTTCTCCCGCTGGGCCCAGTCTCACGCCAGCGAGCTCGATCCCC | 4 |

| | | | |
|---|---|---|---|
| wt-48270-48390 | 1827 | GGGAGATTCGCGACAGAGTTACACCAGCGACTGCACCTTCGTTCGTGGCTGATCATGCCTCGGCTCTATTGGCCTCCCTCCAGAAGAAGGCCTCCGACACCCCCTGTGGGAATCCCATTC | 4 |
| wt-48390-48510 | 1828 | AGTGGATGTGGTACCGCCTGTTGGTAAACTCGTGCCTGAGGAGTGCCCACTGTCTTCTGCCTATACCTGCCGTCTCTGAGGGGGGGAGAAAGACGGGCGGGGGCGTAGGGGAGGAGCTCG | 4 |
| wt-48510-48630 | 1829 | TGGGGGCCGGGGGGCCCTGCCTGAGCCGGGATGTTTTCGTGGCGATCGTAAGCCGCAATGTTCTCTCGTGTCTGCTGAACGTGCCTGCCGCGGGTCCCCGGGCCTACAAGTGTTTCAGAT | 4 |
| wt-48630-48750 | 1830 | CCCACGCCTCCAGACCGGTGTCTGGCCCGGATTACCCTCCCTTGGCCGTGTTTTGCATGGACTGCGGTTACTGCTTGAACTTTGGAAAGCAGACAGGTGTAGGAGGCAGGCTCAATTCCT | 4 |
| wt-48750-48870 | 1831 | TTAGACCCACTCTCCAGTTTTATCCCCGTGACCAGAAGGAGAAGCATGTGCTGACCTGCCATGCCAGCGGCCGTGTGTACTGCTCCAACTGCGGCTCTGCGGCGGTGGGCTGCCAGAGGC | 4 |
| wt-48870-48990 | 1832 | TGGCTGAGCCACCGAGCGCCCGCTCGGGCTGGCGGCCCCGAATCCGGGCAGTGCTGCCGCACAACGCGGCCTACGAGCTCGACCGTGGCTCCCGCCTCTTGGATGCCATCATCCCCTGCT | 8 |
| wt-48990-49110 | 1833 | TGGGACCCGACCGCACTTGCATGCGGCCGGTGGTCCTGCGGGGGGTGACGGTCAGGCAGCTCCTGTATTTAACTTTGCGGACAGAGGCCAGAGCCGTTTGCTCCATCTGTCAGCAACGCC | 4 |
| wt-49110-49230 | 1834 | AAGCTCCAGAGGACGCCCGCGACGAGCCTCACCTGTTCTCCTCCTGTTTAGAGGTAGAATTGCCACCTGGTGAGCGGTGTGCGGGCTGCCGTCTCTATCAGACGCGTTATGGCACGCCGG | 4 |
| wt-49230-49350 | 1835 | CTGCCCAAGCCCACCCTCCAGGGGAGGCTGGAGGCGGATTTTCCAGACAGTCCCCTGCTTCCTAAATTTCAAGAGCTGAACCAGAATAATCTCCCCAATGATGTTTTTCGGGAGGCTCAA | 4 |
| wt-49350-49470 | 1836 | AGAAGTTACCTGGTATTTCTGACATCCCAGTTCTGCTACGAAGAGTACGTGCAGAGGACTTTTGGGGTGCCTCGGCGCCAACGCGCCATAGACAAGAGGCAGAGAGCCAGTGTGGCTGGG | 4 |
| wt-49470-49590 | 1837 | GCTGGTGCTCATGCACACCTTGGCGGGTCATCCGCCACCCCCGTCCAGCAGGCTCAGGCCGCCGCATCCGCTGGGACCGGGGCCTTGGCATCATCAGCGCCGTCCACGGCCGTAGCCCAG | 8 |
| wt-49590-49710 | 1838 | TCCGCGACCCCCTCTGTTTCTTCATCTATTAGCAGCCTCCGGGCCGCGACTTCGGGGGCGACTGCCGCCGCCTCCGCCGCCGCAGCCGTCGATACCGGGTCAGGTGGCGGGGGACAACCC | 8 |
| wt-49710-49830 | 1839 | CACGACACCGCCCCACGCGGGGCACGTAAGAAACAGTAGAGGGCACGAAACATGGTGTATGCACTTTATTAATAAACAATTACAGATACAAAAACTTGAGTCTCTCGAGGTCTGCGATGA | 4 |
| wt-49830-49950 | 1840 | GGCGGTGGGTGGAACGCTCCAGCTTGCGGCGAAGCTGGCTCACGAAGCGAGACAGTACTCGGCTAGCCTGACTAAGGGTGAGGCTATAACGCAGGTCCTGTTCCGGGGCGGCGGTGGATA | 4 |
| wt-49950-50070 | 1841 | GAGAGGAGGGGGATCCGGAGGGGACCACTAGGTCGCCGGAGGTCGACCCTCCTGTCACCACCTCCCTGATAATGTCTTCAATAGACAGAAATTGGGTGACCACTGAGGGAGTGTTCCACA | 4 |
| wt-50070-50190 | 1842 | GTAATGTTGTCTGGTCGCTAGATGGCGCGGGTGAGGCCACGCTTTGCGAAAACGAAAGTGCTTGAAAAGGCGCGGGATAGCGTGCGCTACCGGATGGCGGGTAATACATGCTATCCTTAC | 4 |
| wt-50190-50310 | 1843 | ATTTTGGCATTTTGGGCAGCTGGGAGGCGGCGGATGGGGGTGCTTCTTTTCGCACGGTGTATGTTTGGGGACCCGCATGCCGGTACTGGGATAGGCGCACCTCGGGCCGCGCGCCAGGCT | 4 |
| wt-50310-50430 | 1844 | CCGAGCCGGAATGCATTGGGGGCAATGGGATTGCGGGGGATTGTTGCTGTCTGCTCCTGACAGGGAGAGACACGCGCGGCGGAGATGCAGCCGACGGCGGGGCCGCGGTGGGCTGCCCCC | 8 |
| wt-50430-50550 | 1845 | GAGGACGGGCGCCGGCCGCCAGCGCCCCCGTGGCCTTTGGCACGGGCCTGGCACCCACCGCTTTAATTGTGGGGGTGGGCAGGGCAGCTGCATCTTGGGGCCTTTGTGCTTGCGTTTTTT | 8 |
| wt-50550-50670 | 1846 | GGGGGCGCGGTGCCAATGCACCAACTGGGGTGTGCGCCGGGGCGGCCAAGCCGGACCCCAGGGCGGGTGCCTGGGGGATGGGAAAGCCGGACGGCGCTTCTCCCGGGTCGAACGCTGGAG | 8 |
| wt-50670-50790 | 1847 | TAGCGGAGGCTGCTGCGCCGGCGGCCACCACGGGCGCACGGGGTCGCAGCCCGACGGCCGTGGGGAGGCGGGTGGCGGAGGGCCGAATCTCCGCGGCTTCTTCCCGGCCCCCCTGCTGTT | 12 |
| wt-50790-50910 | 1848 | TCTTCTCCCGTTGCATGATAGAATGGCCATAGGGTGGGTCCTGAGAGGAGGCTTGTGTGTCCTGGGGCTGGAGCCCAAAAGTCGTTAAAGATGCCGCTGATGGTGTGGGAGCTATGCCTC | 4 |
| wt-50910-51030 | 1849 | CCGTCGACTGGCCGGGCTTGTAGGGGGCTGAGGGTGGATAACTGGGCTTCTGTGAAGGCACCAACCCTGGAATCTGGATGGTATGTTTCTTCTGTGACCCCGAGGCAGTCGATGGTGTAG | 4 |
| wt-51030-51150 | 1850 | AGTGTGGAGACAATGTGTAGACGATGGGCCCTTGTTCAGAAGCCCAGGGACTTGAGGGGGGCTGTTGTGGTGCTGGTTGGGGAAGGAGCTCCAGGGAATCTTTGGGCCATGGCCTTGGGG | 4 |
| wt-51270-51390 | 1852 | CGTCCCGGGGGGAGGAGGTAGGAGACAGCCAGTCCCTGGGCCTGCCAGGGGCCACCTTCTCTAAGAGGGGGCTCTGTGGGCTGGGAGGGCCAGAGGCCTCAGATTCAGCAGTAGTGCTCC | 8 |
| wt-51390-51510 | 1853 | CCTTTTCCCCCTGGTCCGTCTCCCCTCCTCCCAACTGCTGGAGCCGGTCGGAGGAGGCCGGGGTGTTATCTGCTGACTGAAACCCGTCCCCGCTGACCAGTCCGTGCCCCACCCTTGGGG | 8 |
| wt-51510-51630 | 1854 | GGAAACCGGAGAACAGCTCCTGGACGTTGCGTGGATTCGGGGGAAGCTGGTATCCAACCGGCAGTGGAGGATCTTCGTGCTCGTAGAAGGAGGGGTTGAGTACATCGGTCGGCCATCGTG | 4 |
| wt-51630-51750 | 1855 | AGGCCCCGGCCGCGTTAAAGTAGAACTGCACGTCCGGCAGATTGTGCCGATAGGTGAAACACTTCCAGATGATGTTTTTTCTGTTGGCCAGGATGGCCACGGTGGGGGGCCTGGCCTCCT | 4 |
| wt-51750-51870 | 1856 | TAGGTTTGGCGGCCCTGGCCTCGGTGAGAAGCTCGCGTAGCCACACGGCCTGGCGTGCAAAGATGGACATCTCTGGCTCGAAAGACTCGGAGTAGCCGTCCAGGTCCTGCAGAAAATTCA | 4 |
| wt-51870-51990 | 1857 | GCGAGATGGTCTCCACCAGGGACCGGAAGGGCTCAGAGTGCCCGTCGCAGTAGAGGAGGGGAGCAACGACCCTGACCTGTCCCAGGGTCTTCAGGTTAAACAGATATTGAGAGGAGACAA | 4 |
| wt-51990-52110 | 1858 | AGAGAGTTAGGGGCCGACCGAGGAAGGCCGCCGCCACGGCCGCCTCAAAAACGGAGACGGGGATGGTGTCACCGGAGCCCCTCTTAGGACCGGTAATGGGAGTGCCATAAGGCATAAGAT | 4 |
| wt-52110-52230 | 1859 | TTCTCAGGGCCCGGCCGGTAACGGTGCCGTAGGAAGACGGGGTTTCGCGGGGGACCTCGAGTCCCTCCGCCCTGGGGAGCTCTTCTCCGCGTGTATAGGCCTGCTTCACAAAGTCGCGCA | 8 |
| wt-52230-52350 | 1860 | GGTAGTCCTGAAATGCGACCGGGCCCTCCAGCGGGCGCAATGAGTGCCAGAGCTGCTGAAGGGCCTCGGGGGCGAAGCACCGGCGTGCGAGGAGCAGCATGCAGGCTCGGGCGCGGGCCG | 8 |
| wt-52350-52470 | 1861 | TACTTTGGTTGTGGACCAGGCCCAAGAACTCGGGGTGCGGCCAGAGGGCGGCTCGGGTATCCATCTCCTCCCAGGCGTCCTGGAAGAAGATGAAGCCGGTGGGTGGACCGGCGATGCGGT | 4 |
| wt-52470-52590 | 1862 | GGCGGGTGAGGCGGCGCGCGTCTTCCCCGTCGTTGCTGCCGCGGGTGGTTGAGGGCATGCCCCCCCTCCCGGAGGCTGGACTCCTGACCAGCCTGTAGGTGAGGACCGAGTCCGACAGGA | 8 |
| wt-52590-52710 | 1863 | GGTCTCCCAAACCCCCATCTCTCGCTAGAGCCGAGACCAGGCCGAGTCCTGCGTAGAACGATGGGGCGCCCAGGAAGGCGGCAGCGTAGGCCGGATGTGTGCCGACCAGCAGCGCCATCA | 8 |
| wt-52710-52830 | 1864 | TCTCCCGTTGTTCCAATAGAATAACTTCCCGGTCTGTGGCCGGGGCTGGATAAGGGGGGTGATTCCTAGAGGCGATGAGACTGGCGTGCGCTAAAAGTGTCATGGCCACAATGGGGTTGT | 4 |
| wt-52830-52950 | 1865 | CTGCCAGGTCTTCCATCAGGGCTTTGGGCGCAGAGACGTATTCCCGAAGCAGCTCCCCGGCGTTGGACTCCACGTCGGGCCAGGTGTCCCAGTAGGAGTCGGCGGCGGCGGCGCTGAGGC | 8 |
| wt-52950-53070 | 1866 | GGGCGGAAGCTACACTGGCCAGGGTTCTTCTCCTCCTCTCTTGGTCATCCTGCGGGGGACCAATAGCTTGGGGGCGTCCGGCTGGGGTCAGGGAAAAGGCCTCTGGGTTCTCCAGCACGG | 4 |
| wt-53070-53190 | 1867 | TGGGCATGACATATTCCAGAAAGTTGTGGTAGACGGGGATGTAGTTGAGCGGCTCCTGGGTGTCTGCGGAGACGTAGGCCGGGTTAAGGGGGTCGCAGGGAGACTCTGTTTCCAGCCAGA | 4 |
| wt-53190-53310 | 1868 | GGGTGCCGGCGTATTTCGCCGGCCCTGCCGCCGCCAGAAATTGTGCCCGCCGGGTCGGGGCTCCATTGCCCCATCCAGTTGGTGGTGCCGAAATCGTGATGAGGAGGGGCAGGTTGTTGG | 8 |
| wt-53310-53430 | 1869 | TCAAGGGATGCTTAACGAAAACGGTAGGCTGGGCGGTCTCGTAAAAAGCCAGGAAACTCTGCTTGGCCGAGGCATAGCGCAGCAGCTTGTCCTTGAGGAGGGCATACTGGGAGCCAGCCG | 4 |
| wt-53430-53550 | 1870 | AGGCCCCAAGCGCCAGGCCCCTGGCAGCCTCCACCACGATCTTGAGCTGGCGCGGGTCGGTGTGGCCCCTGGCCTGGGTGACCAGATCCTGCAGCGTTCCCTGCAGCTGGGACTCTTCCT | 8 |
| wt-53550-53670 | 1871 | GGGCCTCCTGGATGATGGCCTCCAGTCGGGAGAGGCGCCTTTTCCAGTCTGCGACGGTCTCCTTGCCCCCCGCGACCCGCTTGGGGTCCAACGTGGCCAGAGCCAACCTCAGCTCCTCCA | 8 |
| wt-53670-53790 | 1872 | TGCCATCCATGGAGTTCTGGGCCATGCCCTCGACTTCCAGGAGCCGTGTTAGCTCATGAATTTCACCGTCAGCCGCAGCGGCTAGGTTCAGCCAGGCACCCGCGCCCCCAGCTAAGGCCA | 4 |
| wt-53790-53910 | 1873 | GGGCTCCTTCGGAAAGACCCCGCACGGCCTCGCAGATGCCCCGGATCCACTTGGCGGCTGCCAGGGATTTCCGGTAGGGCCATGAGCCGTTCCCGGCCGCTGCCCGGGCCAGGGCGGCCT | 8 |
| wt-53910-54030 | 1874 | CGAGGGGAGCCTGGACAGGGGCTTTGGGCGGGGAGGGAAGCAGGCTCCGGAGTTCATCGTCGGGGGCTTCGTCGCGTGACCTGGAGAGGACGGCCTCCAGAGCCGTGTGAAAGCCCCGCC | 8 |
| wt-54030-54150 | 1875 | GAGTGCTTGCCGCCATCTCGTGGGCCTTCGCCATCAGGGTCTGGCTCTCCCGGACCTGCTCTTCCAGCGCCCGGACCTCGGCCGCCTCGGCCTCGGTCAGCAGCTCCGAGAAGAAGTCCC | 8 |
| wt-54150-54270 | 1876 | CCGTGGCCTGGAGGAGATCGTCCCGCTCTCGCCTTGTCAGCAGCTGGGGCTTCTTAGGCCAGAGCGCCGAGTCCGAGGCCAGCCTGGGCGGGGCGGTTGCCTGGGGGATAGTTGGAGGAG | 8 |
| wt-54270-54390 | 1877 | GAGGCAGGTTAGCCTGGCCTGGGTCATTAGTGGCTTCGGGTAGCGTCCGATCCACGTACTCGCTCACGATGGCCGTCAGGGCAGCCTCGGCTGCTCGTCTTTTTTCCAGAAGCCCGGCCA | 4 |
| wt-54390-54510 | 1878 | GCCCCCGCTCGTACTCCGCGTAGGGGGCCTCCAGATCCGTGTTGACCACCGCTGATTTCATGTCCGGGGACTGCAGGGCCTGGCGCGTCTGCGCGAGGGCCGAACGGATGGCATCGGCCG | 8 |

| | | | |
|---|---|---|---|
| wt-54510-54630 | 1879 | CCGTCCTGGCGCGAAAGAGGGCCCCGGCCGCTTCCTCCGCTCCTCGCCCTCCTCCTCCTTCTTTGGCGGTAGCCGCGGGGGTGGCGGGCCAAGCGTCCAGTCTGGCCAGAGGGCCGGTCT | 8 |
| wt-54630-54750 | 1880 | CGATATCCGTGAACCAGCCGGGTTCCGCGGCCTCCATTCTCTCCGCCGCACCACCATCGTCCACGAGCAGGGATCGCAGTCTCTCCCTCCTCACCCTCGTTATTCCCAATAGCATAGCGG | 4 |
| wt-54750-54870 | 1881 | CAAGGATCTGTGTGAGGGAGTCCAAGATGTCCGTGTTTCTGGCTACTGCCGCCGCTGCTGCCGCGGCTGAGTCCGTATTGTCTGGCAGCAGGGAGGCCAGCAGGGTGTTCCAGTCATCGG | 4 |
| wt-54870-54990 | 1882 | GCGAAGTGGGAGCGGGCTCTGGGCGTGCCCCCAGCGCCTTCCTAATTCTGGCCCAGGCCTCATTCGCCTCTCGCGCTCGCCGCTCCTGCCTCTCCTTGTCTTCCTGTTCTCGGAGCTTCT | 8 |
| wt-54990-55110 | 1883 | CCTTTTCCTTGCGCCCGGTCTCCATAAGCTGCCGCAGCTTCTTCTCATACTGTCGCTTGAGCTCTTTGTTGGGGGCAGTGTCCAGAAAGGCCTCGAGCTGTTCCTCGGTGGCGGGCTTAA | 4 |
| wt-55110-55230 | 1884 | AGCCTTCGGCCTCCAGGCGCCAGGCCTGCACCTCCTTCTGTCTGAGCTGATCGTTGTTGTTATTCTTCAATTTCTGCAGGTAACTTAGGAAGCGTTTCTTGAGCTTCCCTGGGATGAGCG | 4 |
| wt-55230-55350 | 1885 | TTTGGGAGAGCTGATTCTGCAGCCCAGAGAGTAGTCTCAGGGCATCCTCTGGAGCCTGACCTGTGATCGTCGCATCATAGACCGCCAGTAGACCTGGGAGCAGATTCACCGCCGCGGCCG | 4 |
| wt-55350-55470 | 1886 | TCTCCTTTAAGGTGCTGTGAGTAGCAAAATTCTGCAAGGCCACTAGGCGCGCTGGCTCCAGCGTCAGCCGGTTGCCCATCTCGAATGTGTGCAGGGCCTCTGAGACCATGGGGTCCAGGA | 4 |
| wt-55470-55590 | 1887 | TGCGGTCAATGCCATCCTGCACCTCAGGGTCAAGGACCGGCAAGTCACGATAGAGGTGGTCTATGCTCTCCTCGAAGGAGGCAATGTAGTTATCGATGGTGTAGAAGGTGATGGATTTCA | 4 |
| wt-55590-55710 | 1888 | GGATGTTCATCAGGTACTTTTTGGAGCGAACAATCTGCTGTATAGTGTCACGTAGGCGGATGTACGTGGGGTTCTTTGCGGCCCCGACTATCGACCCTGCATTTGCGATGTACTTTTCTA | 4 |
| wt-55710-55830 | 1889 | TGACGGGGATGGTGAGGGCCGCGGTGTCGGCCAGCGGTGGCGTGGCTTCGGGGTTGTCGTGGTTGGCGGGTGTCGCAGAGGGAGAGGCGGGAGAGATGGGGGCGCCTGGGGCCGAGGCCA | 8 |
| wt-55830-55950 | 1890 | CACCGGCCAGGCCCAACATTGCCTCGATGTCGTCCAGGATGGTGCGGAGGCGCTTTTCGTTTTCTCTGGTGGTCTCGAGCTCCTTCTGTTTTTTCGCGACTGTCTCAAACTCTGGAAGGG | 4 |
| wt-55950-56070 | 1891 | GGGCAATGCTGGGGTCGTCCTCCTCAACTCGCTCCAGGGGCCAGGGGATACCGCTCATATCACTAAGGGCGGTGCCCAGGTAGAGGAGCTCGCGATAGTCCCATTCAATGGACGTGTACC | 4 |
| wt-56070-56190 | 1892 | GGATGTTTAGGAGAGGCAGGGAGGCGATGATCTGGCATGTGTGCCGCAGGTGTGTCAGGAGGTCGTCAAAATCCATCACTGTTGGGAGGCTTGGGTCCTCAAGGTAGGAGAGATAATCGG | 4 |
| wt-56190-56310 | 1893 | AGGCCGCCGAGGCCACCTTGTCCCTGATGTCCGCCGTACACCTGCGCACGTGCAGGGCCGCATTCTTGGACCGGACGGCCACGTTGTGGACAAAGGGGGGCACTGAGGCGGCGGGAGGGG | 8 |
| wt-56310-56430 | 1894 | CCCCATACTCTATCGCTGTCAACAGCGCCAAAAAGCGGACGTCCTCCTCATCTACCCCAGCCTGTTGTCTGGCCACGGCCGTTCGGGCGGCCTCCGCCAGGGATAGGAGGCGCTTCCAGC | 4 |
| wt-56430-56550 | 1895 | TTTCGTCGTCCAGGACCAAGGGGACGTCCACGTGCGGGCCCCTGTAGATGGAATTATCCTCGGGTTCTCCTCCTCCTTCCCCCGCCTCCTGATCTCCGCCCGAGAGCAGGTCGGTCAGGC | 4 |
| wt-56550-56670 | 1896 | GTCTGCGGGCCGCCTCCAGGTCAAATTTTCCATCGTCGCTCTCGGCCAGCTGGGGAATTTCAGCCAGCATCTTAGCACCGGCATCTACACGGACCGCGTCCTTCGTGGCCAGGGACGGCA | 4 |
| wt-56670-56790 | 1897 | GGCAGGCCTCCAGCTTTGCGGCCAGGTGCTTATGGAACTCTCCCGCTCTTCCCTTGTTTTCTGATAGCATGTTTGCGAGGTTTTGGATGTTAAGTTCGGAAGTGAGCAGTTGCTCCAGGT | 4 |
| wt-56790-56910 | 1898 | CCAGCGTGGGGACCTGCAGATGTCCCGACCAGTCCTTTAAGAATTCCAGCAGATTTAGCACAGACGATCGGTCCCTACTCCTTATTAGCCCCTGCTCGAGGACCACTGTCACAAGAAGAT | 4 |
| wt-56910-57030 | 1899 | AGTCTATCATGCTCAAGGCATCTGCCTCTGGCACTTCCCGGTTAGAGGCCGGGTCGTAGACGATGGCCTGTTCCTGGTAGGTATGTCCGGCTATTCTCGCAATGTTGCTCTCGAGGGGCA | 4 |
| wt-57030-57150 | 1900 | CAAAGTCCATCTCAGGAGTCTCTATGTCAAAGGTGGTCTGATAGTATTGGCTCCTGGCGGTGTCCAGTGTGATGGGGGACGTGGGGGCACTGGATCCCGATTCCAGGCTGTTGGAGAACA | 4 |
| wt-57150-57270 | 1901 | CTTCATCTTCGAACATGTCTTCATCCTCTGTGGTGGGGATATCGGAGGCTAAGTCGCTCTCCGCTTCTTCAGAGTCGGACATGGATAGGAAAGGCTCCTCTAGGTCAGACAGGTAGCGGA | 4 |
| wt-57270-57390 | 1902 | CGAGGCCAGAACCCCCAGATGCATCATCCCCAAAGGAGGGCTGCTGCCCGAAGGGAGGTGATGGGGATATCTCCGTTCCAGCCCTGTCAGCGGCCGGGGGATGGTTTTTTTCTGGTTCGA | 4 |
| wt-57390-57510 | 1903 | GTGTCGTGGCTGATGGTGGGAGCTGCTGAGCAGGAGGAGGAGCCGGGGTAGCTGATGGCAGGGGCTGCTGCTGAGGAGGAAGAGGAGAAGGAGCCCGGGCGGCTGATGGCGGGGGCTGCT | 8 |
| wt-57510-57630 | 1904 | GCTGAGGAAGAAGTGGAGAAGGAGCCGGGGCGGCTGATGGCGTGGGCTGCTGCTGAGGAAGAAGTGGAGAAGGAGCCGGGGCGGCTGATTGCGGGGGCTGCTGCTGAGTTGGAGGAGGAG | 8 |
| wt-57630-57750 | 1905 | AAAGAGTCGTGGTGGTGGGGGCTGCTGCTGCAGTCGGGGAAGGGGATGGGGTGGTCAGAGGGATTTTTGGGTTCGAGGGAGCTGCCTGTGGCAGAGGGATGGGTATTTGCAAAGGGAGGC | 4 |
| wt-57750-57870 | 1906 | GAGGAGATGGAGTGACTGAAGGAGCGATAGTTGAGACTGGCGCGGGGTGGGGTGTCGGGGAGGCGGGTGGTGATTGGTGAGGGATGGGGATTACTGGAGGGGGAAGGCGAGCTGCTGAAG | 4 |
| wt-57870-57990 | 1907 | GGGGGCGATGGGGCGGAACGTGGGTGCGTGGCAGCTGATCATCCTCTGTGTCAGTGGTGGAGGACAGAGGGAGGCGGCGGCCGGAGGTGGGCTTCTTGTGGGGGCTATCTTTGCCCAATC | 8 |
| wt-57990-58110 | 1908 | CCTTTTTCCTCTTGGGAGTCTGAGGCGCTGCGCCGCTCGACGCCCTTGGTGGCGTGGAGGGAGCGGGGACCCCGGGGGTGTGACCTAGGCCGGGGATGGGGATGAAGAGGGGAGGGCTGG | 8 |
| wt-58110-58230 | 1909 | AGGCCGGGGCCGCGGAGGCCGGGGCCGCGGAGGCCGGGGCCGCGGAGGCCGGGGCCGCGGAGGCCGGGGCCGCGGAGGCCGGGGCCGCAGAGGCCGGGGCCGCAGAGGCCGGGGCCGCAG | 16 |
| wt-58230-58350 | 1910 | AGGCCGGAGACGACGGCGGGGAGTTGGTCTTTGCAGGACTATACCTGGCGGCAGGGAATGAGTCGGATGTGAAAGATCGAGAGGGCAGTGGCCTGAGGTTATACGGTATTATTCGCCGTT | 4 |
| wt-58350-58470 | 1911 | CAAACGGTAGCATGACGGGAGGGCTGCTATCAGCACCGGGCGTCCCCGCCGCCTCCCCATCACTGGACACAAGCTCGGGCCCCACCAGGTCAAAGCCGCTGCCGTTGGCCTCATAAAAGT | 4 |
| wt-58470-58590 | 1912 | CATACACGCCATAGTGTTCCAGCATAAAGATGCGGGGGTCCTCTGTCTCAAAGGCCTCGGGTAGAAAATAGAGATGCACGCAAGTGTACTGGGCCCCTGGTGCCCCCACGTACTGCAGGA | 4 |
| wt-58590-58710 | 1913 | TGTCGTGCGCATAGGTGCTGACTCTGACATGGGCGGGGGTGCCCGGGGCCGCATCCTTCTGGCAGTGGGGGTCAAACAAGTAGAAGGAGCCATCTGTCTCGATGATGATGGCCCCCGCGT | 4 |
| wt-58710-58830 | 1914 | AGATGTCGCAGATGTAGAGGATGAACTGGGCCACCCCGTTGTAACTGCCGTGCAGGACCTCGGCCAGGGACTGAACAACTGCCGAGTTTGCGATCTGGGCAGGGAATAGGACGAGGCCAA | 4 |
| wt-58830-58950 | 1915 | AGATCTCCGCCGAGCGGTATATGTGCACGCGCCCACCGCCCCTCAGGACCACGGAGCTGGGCACGTCCGTCAACTGGGCCATCTCGTGCCCCTTGAGGATGCCGCTCTGGCGCATGAGGG | 8 |
| wt-58950-59070 | 1916 | CATCCAGCCGCGCCCCCTCGTCCAGGACCTCGTCCAGCTCAGGGCGGGAGGTCAGGGGGCGGCCGGCCAGGAAGCTCTTGACCAGGTAGAGGACGCAGTTGCTGACGCACTGGATGCCGG | 8 |
| wt-59070-59190 | 1917 | CAAAGCGGCCAAACTTGCAGTGGGCCTGGTTGCACGAGGCCGTGCCTAGGATGCGGAGGGCCGAGCCTCCACTCCCGCCCCCGGGGGCATTCACATCCATGGTCCTGATTCCGCGCACGG | 8 |
| wt-59190-59310 | 1918 | GGCCGGTTCCCCGGGTGCGCTGGCTTTGCCCCCAGTCGCCGTTACTCATCTTCGGCGGTGGGGCGGGGAGGACGCCTTGTCGCCCCCCTTCTGGTCCGGGGTCTTACGCGGCTGGCGGCG | 8 |
| wt-59310-59430 | 1919 | GCAGCCGCCGAGAGATAAGGGGGGTACGTGTGTGCCTCCGCCTCTCCTCTGTCTGGGCCGCCGCCGCCGCTTGCCCGCCTTGAAGGAGAGGGGGTAGTCCGCGGACTGCGTCTGCGGGGG | 8 |
| wt-59430-59550 | 1920 | CAGGAGGTCTCAACCTTCTGGGCTCGGGCCGCGGTGTCGATATCCGATGGCCTTTCCCTGTCTTCCTCGTATGCTCCTTCTCCTCCTCCTCCCGGCACGCCCCTGAGATCTGCCTCCCCT | 4 |
| wt-59550-59670 | 1921 | CCCTCTCCCTCGTCCTGGTCGGAAAAGTCTGAGGAGGAGAAGGAGAATGGGGAGGAGTCCAAAACGGCACGCCACCTGCCGTGGGGCGGTGGTGACAGGTCCCGGCTGGCCCGGCGCTTG | 8 |
| wt-59670-59790 | 1922 | CTCGCGTTCCTGCCGTTACCCAGGAGAATGGCCGCGAGTTTTTTGGCGGGGAGGATGCGGAATGGCGGGGGCGTTTGTCCCACGGGTGAGGGGGAATCGTCGGTTAGGGCCGGCACGAGG | 8 |
| wt-59790-59910 | 1923 | TGGTGGGTCTGGACCCGGGCCGTGCGAGCAAAGGCGGCGAGAACCGAGGGGCTTCTGGGGGTGACTGTGATCTGTTCCGGATTTAGGTCCATGGCGGGTGTGTATGTTTTAATAGGGGTG | 4 |
| wt-59910-60030 | 1924 | GTCTCTGGCGCGGCAGGATGATGGTCGAGGACGTCCACCAGGGCCTTGCAGATGCTCTTGCCTAGATACAGGATGTCGTCCATGCTGAGGGGAGGTGGGGTGTCTGCTCCCCCCTGCGGA | 4 |
| wt-60030-60150 | 1925 | AGCCGCCTGGGTGCGGGGATGAAGACAGGTGGTGGGCGGGCGTCTCGCCGGACTATGGCCTCGGCACGCTCGGCGTCGATGGCGGGTGGCTGGAACAGGCGGGCGAATGTGTAATCCCGG | 8 |
| wt-60150-60270 | 1926 | AACCGGTAGGCGACGCTGCGCCTGAGGGCGCCCGTCAGGCTGTATCCCAGCTCCAGGGCGTGCTCCACCCGCTCGTTGAGCTCCTCGAGATCCGGACGCAGGGGCTCGCTGGTGTGGGCC | 8 |
| wt-60270-60390 | 1927 | CAGAGGGGGTGATCCGCGATGCCCCGGCTCTCCCTGAGGGCCGGCACCAGGAGGCGCCTTCTGAGGGTGGCCGTGTCGGCCGTGGCCAGGGCCCACCTGGCGGCGGCGTCCCGGCACACA | 12 |
| wt-60390-60510 | 1928 | TCCTGGATGCCCTCCACGACGCTCTTTAGCGTCTGGAGGTCCGTGGAGTAGTGGCGGGGGGAGGATGAAACGCTCTTTTCCTTCACCGCTACCACCGCCTCCTCCTCCTCTTCCGTCGCC | 4 |
| wt-60510-60630 | 1929 | AGAGGGATCTGCACCCTCCCGGTCTCTGCGTCGTACAGGAGCGGGCGGGAGCACAGCCTCCAAGCTGCCCCCGTCAAGCGCGAGATGTCCTCCGAGAGGGTCTCACCCGAGACCAGAAAG | 8 |

| | | | |
|---|---|---|---|
| wt-60630-60750 | 1930 | CGGCGGGTGGCCAGGCCCAACGACTCCGCCGTCGTGCTGTATCTCAGGGTGAAGAGGAGTGAAAAGAGGGAGGTGGGCCAGGCAAGCGGTGGTGCTTCCGCCGCCCGCTCTGAAGCTGAG | 8 |
| wt-60750-60870 | 1931 | ATAGTCTCGGAGATGATGCCTGAGACCTCTCGGACGGCGTCCATGATCCTAAGGACTGCGTCGTGGGACGACAGCCCCCAGGGGCCCCCGCCCTCTTCGTCTTCTGCACCCTCGGCTCCT | 4 |
| wt-60870-60990 | 1932 | GCGTCCCCGGCCTTGCCTTCCCCCTCTAAGTTGAGGGGGCGCAGTCCGACCGCCTGGGGGGACTCCCCAGGCATCGGAGGGGCCCCGTCATAGATCTCCCAGACGGTGGCGTATATGAGC | 8 |
| wt-60990-61110 | 1933 | TCGAGAGGACGGCGGGCCCGGGTCAGCTCGGGGGAAGGGAGGGCCAGGTCGCTGCCGAAGGAGACCAGCCAGCGCAGGGCGGCCAGAGAGCGGGTTTTGGGCAGCTCGTTGGAGAGGACC | 8 |
| wt-61110-61230 | 1934 | CGGCGAAGGGCGGGCCAGATTTGGAACTCGATGAAGGCGGCCGGGAAGAAGGGGCTGTGGACATAGGCCGGATCCGCGCGCGCCGTTTGGCCGGCCCTCAGGGACCGGCAGTATGCCTCG | 8 |
| wt-61230-61350 | 1935 | ACGTCTGTCCGCGGGGCCGCCGCCACCGCTGCCGTCCACTGCCTTCTTCCCTGCTCGCCGGGGAGTAGGGGGGGCTTACAGGGGAGGGCCGGAGCCGGGGCCGGGGCCTGCCACAGGCGG | 12 |
| wt-61350-61470 | 1936 | CTGTAGCGGACCCATAGCAGAGACCTGAGGAGTTCGGATGAAAGGTCCCCCGCCACCTGCTCATACTCGGCCGCGGGAGGGGGGACGATGAAGATGCGCAGAGGGGTTACGGCGTCCCAA | 4 |
| wt-61470-61590 | 1937 | GGGTCCGCCGCCGCCCCCACACCCACAGCCGTCGCGGCGGGGGCGGCGGCGGGCGTAGAGGGGCCGCTGGTGCGCCGGGCTCGTCTGTCCACGGCCTCGGCCTCCGCCCGCAGGTAGGCC | 12 |
| wt-61590-61710 | 1938 | GCCCGGGCCACACGGGCGAAGCGGCTCGTGGGGCTCGCGGTGGGCAGCAGTCGGAAAAAGTGCAGGGCAAAGCCCGATAGACTCTCTAGGAGGGCGGCGGTGGCCTCGAGCCACCTCCAC | 8 |
| wt-61710-61830 | 1939 | CGCGAGCGGGACACCCGGGGCACAGAGGCCAGCATCATGGCGTAGTCCCCCGCCACGGTCTCGTTGAGCCCGGCCGAGAGCAGAACCGTGGCCACCTGCTCGATGGCGGCTGGAGAGAAG | 8 |
| wt-61830-61950 | 1940 | GATGCCCGGCTCCCCGCCGCCTCCTGCACACGAGCGGCCAGGGCCTCCATCTCTGCCGCCATCCCGGCCAGGAAGGCCTCGATGACCGAGTCTGGGACGCCGTAAGTCTGGTCCCAGAGC | 8 |
| wt-61950-62070 | 1941 | AGGGCCTCGTACACATAGTCGTAAAAGAGGGCCCCTGAGGGCTCCAAAAGCCGGAGCCGGGCGGCGTCAAAGGCCAGGACGGGCACAGCCGCGACGGGGGGCGTTTGTCCCCCGCTGGCC | 8 |
| wt-62070-62190 | 1942 | TCCGCGTACACGCCCAGGATCTCTACCGCCCGCCGCCGGGCCAGGGGCAGCGAGGCCACCACGCTGGAAAGTGACTCGGGGCGGTGAAAGAGACCACCACCGCTTTCTTCACCCTCTCCC | 8 |
| wt-62190-62310 | 1943 | CCGCCGGCCCCGCCCCCACTGTGCTCCACCAGCTCCACGGCCATGGCCTTGATGTCCGCGGCCGTGGGCTGACCCTGCCCTGCAGCCGCCCAGGGGTAGCGGTTGGTCTCCGCGTATACG | 8 |
| wt-62310-62430 | 1944 | GTGACCAGCCATCTCCCCAGCGTCGTTTTCGCCGCGTTAAAAGCGTAGAATGACAGCCCCTCCCGCGGGAAGGCGTCCCACCGGGCCAGATAAGTGTCGGCCACCAGCTCTTCCACGAAG | 4 |
| wt-62430-62550 | 1945 | GCAAAGGTGGCCGTTGGGCCAGAGACCGCGAGCACCTCCCCGCTGCCCTCTTCGATGATGCGCCGGTACGCGGCCGCCAGGGCCCGGGTCTCTGCGATGAGCCGAGAGCCGTCCAGCGGA | 8 |
| wt-62550-62670 | 1946 | TCGTCGGTGGCCGGAGAGGCTGTCGTGGGGGGCAGTGAGGATGCCAGCACGTCCAGGGCCGCCTCCAGATGGCCGAGGCCGAAGCTGCGCCTGGAAAAGGAGGCCGCCCGGAGTAGGTAG | 8 |
| wt-62670-62790 | 1947 | TAGGCGTGGTGGCGGAGGACCGCCGCCGGGTAAGCGTGGCCGCTCATGAGGGTGAGAGTATTTAAAAAATCGCGCACCAGCACCGGCTGGGCCAAATCCCCCAGTCCAAAGATCCCCAGC | 4 |
| wt-62790-62910 | 1948 | TCCAGAGGCATCAGCGCGCGCAGGCGGGCAGCGGGGTCGTCCCCAGACAGCAGCAACTGACGCGTCACGCGGGCGAGCCCCCCGTCCACCTCTGCCAGGGGTGGCTGGGCGTCTGCCCCT | 8 |
| wt-62910-63030 | 1949 | CCGCTACCGCCGCTGCTGTCACTCTCCATAGCGGACGCCATGAAGGTCCAGGGGTCCGTCGATCGCCGCCGTCTGCAACGCCGAATCGCGGGGCTGCTGCCCCCTCCGGCCCGGCGTCTA | 8 |
| wt-63030-63150 | 1950 | AATATTTCCCGGGGGTCCGAATTCACGCGGGACGTTCGTGGGCTGGTTGAGGAACACGCGCAGGCCTCCTCGCTGAGTGCGGCGGCCGTCTGGCGCGCAGGGCTGCTGGCCCCGGGGGAG | 8 |
| wt-63150-63270 | 1951 | GTGGCGGTCGCCGGGGGTGGCAGTGGAGGGGGGAGCTTCAGCTGGTCTGGGTGGCGGCCGCCAGTCTTTGGGGACTTTCTGATACACGCCAGCTCCTTCAACAACGCCGAGGCCACTGGA | 8 |
| wt-63270-63390 | 1952 | ACGCCCCTTTTCCAATTCAAGCAGAGTGACCCGTTCTCGGGCGTCGACGCGGTATTCACTCCTCTCTCCCTGTTTATCCTAATGAATCACGGCCGGGGTGTAGCCGCCCGGGTCGAGGCA | 4 |
| wt-63390-63510 | 1953 | GGTGGGGGCCTGACGCGGATGGCCAACCTGCTGTACGACAGCCCCGCAACCCTGGCTGACCTGGTCCCGGACTTTGGGCGGCTGGTGGCCGACCGCCGCTTCCACAACTTCATCACCCCT | 8 |
| wt-63510-63630 | 1954 | GTGGGCCCCCTGGTGGAGAATATAAAGAGCACCTATCTGAATAAAATCACCACGGTGGTCCACGGGCCTGTGGTCAGCAAGGCCATCCCTCGCAGCACCGTCAAGGTGACGGTGCCCCAG | 4 |
| wt-63630-63750 | 1955 | GAGGCCTTTGTGGATCTGGACGCGTGGCTCTCCGGCGGCGCCGGGGGTGGCGGTGGAGTATGCTTCGTCGGGGGGCTGGGCCTGCAGCCGTGCCCCGCCGATGCGCGCCTCTATGTCGCT | 8 |
| wt-63750-63870 | 1956 | CTGACCTATGAGGAAGCCGGGCCGCGGTTTACGTTTTTCCAGTCGTCCCGCGGCCACTGTCAGATCATGAATATCTTAAGAATTTATTACTCACCATCCATCATGCACCGCTATGCTGTG | 4 |
| wt-63870-63990 | 1957 | GTCCAGCCCCTACATATAGAGGAGCTAACCTTCGGGGCGGTTGCCTGTCTGGGGACATTTAGTGCTACTGACGGTTGGAGGAGGTCTGCCTTCAATTACCGTGGCTCTAGCCTCCCCGTG | 4 |
| wt-63990-64110 | 1958 | GTGGAGATTGACAGCTTTTATTCCAACGTCTCTGACTGGGAGGTGATTCTCTAGACTTAACGGGAGGAAACAGGAGGAGGAGGGGGACAAGAGCACAAAAGTGGTTCAGTGGACACCCAC | 4 |
| wt-64110-64230 | 1959 | CACACAGCATGGCAACGACCAGTCATGTCGAGCATGAGCTCCTCTCCAAATTGATTGATGAGTTAAAGGTCAAGGCCAACTCAGACCCCGAGGCTGATGTCCTGGCCGGGCGCCTGCTCC | 4 |
| wt-64230-64350 | 1960 | ACCGCCTTAAGGCCGAGTCAGTTACACACACAGTAGCCGAATATCTGGAGGTCTTCTCTGACAAATTCTACGATGAGGAATTCTTCCAGATGCACCGGGATGAGCTGGAGACCCGAGTCT | 4 |
| wt-64350-64470 | 1961 | CTGCTTTCGCGCAGAGCCCGGCCTACGAGCGCATCGTCTCCAGCGGCTACCTGTCGGCCCTGCGCTACTATGACACCTATCTGTATGTGGGGCGCAGCGGGAAGCAGGAGAGTGTGCAGC | 4 |
| wt-64470-64590 | 1962 | ACTTTTACATGCGGTTAGCCGGCTTCTGTGCCTCAACCACCTGCCTCTACGCGGGTCTCAGGGCAGCCCTGCAGCGGGCCAGGCCGGAGATTGAGAGTGACATGGAGGTGTTTGATTACT | 4 |
| wt-64590-64710 | 1963 | ACTTTGAGCACCTAACCTCCCAGACGGTGTGCTGCTCCACGCCCTTTATGCGCTTTGCCGGGGTGGAAAACTCCACTCTGGCCAGCTGCATCCTCACCACCCCCGACCTCAGCTCCGAGT | 4 |
| wt-64830-64950 | 1965 | TGATCAACAGCCACGTGGAGTACCACAACTATGGCTGCAAGAGGCCGGTCAGTGTGGCGGCCTACATGGAGCCCTGGCACAGCCAGATTTTCAAGTTTTTGGAAACGAAGCTGCCGGAGA | 4 |
| wt-64950-65070 | 1966 | ACCACGAGAGGTGCCCGGGCATCTTTACGGGGCTCTTTGTCCCCGAGCTCTTCTTCAAGCTTTTTAGGGACACGCCCTGGTCGGACTGGTACCTGTTTGACCCCAAGGACGCCGGGGACC | 4 |
| wt-65070-65190 | 1967 | TGGAGAGGCTCTACGGGGAGGAGTTTGAGCGCGAGTACTATCGGCTGGTGACAGCGGGCAAGTTTTGTGGGCGGGTCTCCATCAAGTCCCTGATGTTCTCTATCGTCAACTGCGCCGTCA | 4 |
| wt-65190-65310 | 1968 | AGGCCGGCAGCCCCTTCATCCTTTTGAAGGAGGCCTGCAACGCCCACTTTTGGCGCGACCTGCAGGGCGAGGCCATGAACGCCGCCAACCTGTGCGCCGAGGTGCTGCAGCCCTCGAGGA | 8 |
| wt-65310-65430 | 1969 | AGTCTGTGGCCACCTGCAATCTGGCCAACATCTGCCTCCCGCGCTGCCTGGTGAATGCGCCTCTGGCGGTGCGGGCACAGCGGGCCGACACGCAGGGGGATGAACTCCTGCTGGCCCTCC | 8 |
| wt-65430-65550 | 1970 | CTCGACTCTCAGTCACCCTACCTGGAGAGGGGGCAGTCGGTGATGGATTCTCGCTAGCCCGCCTCAGAGATGCCACCCAGTGTGCCACCTTTGTGGTGGCCTGCTCCATTCTTCAGGGAT | 4 |
| wt-65550-65670 | 1971 | CCCCCACTTATGATTCCAGGGATATGGCCTCCATGGGCCTCGGGGTGCAGGGCCTGGCCGATGTCTTTGCGGACCTGGGCTGGCAGTACACTGACCCTCCCTCTCGCTCGTTAAACAAGG | 4 |
| wt-65670-65790 | 1972 | AAATATTCGAACATATGTACTTTACGGCCCTCTGCACCAGTAGTCTGATTGGACTTCACACCAGGAAGATTTTTCCGGGTTTCAAACAGAGCAAGTATGCCGGGGGGTGGTTTCACTGGC | 4 |
| wt-65790-65910 | 1973 | ACGATTGGGCAGGAACAGACCTTTCTATTCCCAGGGAAATTTGGTCTCGCCTCTCTGAACGCATTGTGAGGGATGGGCTTTTCAATTCACAGTTTATCGCCCTGATGCCCACCTCAGGCT | 4 |
| wt-65910-66030 | 1974 | GTGCCCAGGTGACGGGCTGTTCGGACGCCTTCTACCCCTTCTATGCCAATGCGTCCACCAAGGTCACCAACAAGGAGGAGGCCCTTAGGCCAAACCGGTCTTTTTGGCGTCATGTGCGTC | 4 |
| wt-66030-66150 | 1975 | TGGATGACAGGGAAGCTTTGAATCTTGTCGGGGGCCGTGTCTCCTGCCTCCCGGAGGCTCTGCGGCAGCGCTACCTGCGTTTCCAAACGGCCTTTGATTACAACCAGGAGGACCTGATTC | 4 |
| wt-66150-66270 | 1976 | AGATGTCCCGGGACAGGGCCCCCTTTGTGGACCAGAGCCAATCTCACAGCCTGTTTTTGCGTGAGGAAGATGCCGCGCGGGCCAGCACGCTAGCCAACCTACTGGTGCGCAGCTACGAGC | 4 |
| wt-66270-66390 | 1977 | TGGGCCTGAAGACTATCATGTACTATTGTCGCATTGAGAAGGCCGCCGATCTGGGGGTGATGGAGTGTAAGGCCAGCGCGGCTCTGTCGGTGCCGCGGGAGGAACAGAATGAGCGGAGTC | 4 |
| wt-66390-66510 | 1978 | CCGCTGAGCAGATGCCGCCTCGTCCCATGGAACCGGCGCAGGTTGCGGGGCCGGTTGACATCATGAGCAAGGGCCCAGGGGAGGGACCAGGTGGGTGGTGTGTGCCCGGGGGATTGGAAG | 8 |
| wt-66510-66630 | 1979 | TGTGCTATAAGTACCGTCAGCTCTTCTCAGAGGATGATCTGTTGGAGACTGACGGTTTTACTGAACGAGCCTGTGAATCTTGCCAATAAACGTTTATTGCCATGTCCAAGTTGTTGTACG | 4 |
| wt-66630-66750 | 1980 | TGCGTGATCATGAGGGCTTTGCCTGCCTAACGGTCGAAACCCACCGCAACCGCTGGTTCGCGGCTCACATTGTCCTCACCAAGGACTGCGGGTGTCTCAAGCTACTCAATGAGAGGGACT | 4 |
| wt-66750-66870 | 1981 | TGGAGTTTTACAAGTTCCTCTTTACGTTCCTGGCCATGGCCGAGAAGCTTGTGAACTTTAACATTGATGAACTGGTCACCAGCTTCGAGAGCCACGACATTGATCACTACTACACCGAGC | 4 |

| | | | |
|---|---|---|---|
| wt-66870-66990 | 1982 | AGAAGGCCATGGAGAACGTCCACGGGGAGACTTATGCTAACATTTTAAACATGCTCTTTGATGGGGACAGGGCGGCGATGAACGCCTACGCAGAGGCCATCATGGCCGACGAGGCCCTGC | 4 |
| wt-66990-67110 | 1983 | AAGCCAAGATTTCCTGGCTCCGTGACAAGGTGGCGGCCGCCGTCACCCTGCCGGAGAAGATTCTTGTGTTCCTGCTGATTGAAGGCATCTTCTTCATTAGCTCCTTCTACAGCATAGCCC | 4 |
| wt-67110-67230 | 1984 | TGCTGCGGGTCCGGGGCCTAATGCCTGGCATCTGCCTGGCCAATAACTACATAAGTAGGGATGAGCTGCTCCACACCCGCGCTGCCTCCCTGTTATACAATAGCATGACAGCCAAGGCTG | 4 |
| wt-67230-67350 | 1985 | ACCGACCAAGGGCCACCTGGATCCAGGAGCTGTTTCGCACTGCGGTGGAGGTAGAGACTGCCTTCATCGAGGCTCGTGGAGAGGGGGTTACCTTGGTGGATGTGCGAGCCATAAAGCAGT | 4 |
| wt-67350-67470 | 1986 | TTCTGGAGGCCACGGCCGATCGCATCCTGGGTGACATTGGTCAGGCTCCCTTGTATGGCACACCACCCCCCAAGGACTGCCCGCTCACCTACATGACTAGCATCAAGCAAACTAATTTCT | 4 |
| wt-67470-67590 | 1987 | TTGAGCAAGAGAGTTCCGATTACACCATGCTGGTGGTAGATGACCTTTGAGTCAGGGTGGCTACTTGCTCAGGTTTCTGGGCATAAATTCTCCTGCCTGCCTCTGCTCTGGTACGTTGGC | 4 |
| wt-67590-67710 | 1988 | TTCTGCTGCTGCTTGTGATCATGGAAACCACTCAGACTCTCCGCTTTAAGACCAAGGCCCTAGCCGTCCTGTCCAAGTGCTATGACCATGCCCAGACTCATCTCAAGGGAGGAGTGCTGC | 4 |
| wt-67710-67830 | 1989 | AGGTAAACCTTCTGTCTGTAAACTATGGAGGCCCCCGGCTGGCCGCCGTGGCCAACGCAGGCACGGCCGGGCTAATCAGCTTCGAGGTCTCCCCTGACGCTGTGGCCGAGTGGCAGAATC | 4 |
| wt-67830-67950 | 1990 | ACCAGAGCCCAGAGGAGGCCCCGGCCGCCGTGTCATTTAGAAACCTTGCCTACGGGCGCACCTGTGTCCTGGGCAAGGAGCTGTTTGGCTCGGCTGTGGAGCAGGCTTCCCTGCAATTTT | 4 |
| wt-67950-68070 | 1991 | ACAAGCGGCCACAAGGGGGTTCCCGGCCTGAATTTGTTAAGCTCACTATGGAATATGATGATAAGGTGTCCAAGAGCCACCACACCTGCGCCCTGATGCCCTATATGCCCCCGGCCAGCG | 4 |
| wt-68070-68190 | 1992 | ACAGGCTGAGGAACGAGCAGATGATTGGGCAGGTGCTGTTGATGCCCAAGACGGCTTCCTCGTTGCAGAAGTGGGCACGCCAGCAAGGCTCAGGCGGCGTTAAGGTGACACTCAATCCGG | 4 |
| wt-68190-68310 | 1993 | ATCTCTACGTCACCACGTATACTTCTGGGGAGGCCTGCCTCACCCTAGACTACAAGCCTCTGAGTGTGGGGCCATACGAGGCCTTCACTGGCCCTGTGGCCAAGGCTCAGGACGTGGGGG | 4 |
| wt-68310-68430 | 1994 | CCGTTGAGGCCCACGTTGTCTGCTCGGTAGCAGCGGACTCGCTGGCGGCGGCGCTTAGCCTCTGCCGCATTCCGGCCGTTAGCGTGCCAATCTTGAGGTTTTACAGGTCTGGCATCATAG | 4 |
| wt-68430-68550 | 1995 | CTGTGGTGGCCGGCCTGCTGACGTCAGCGGGGGACCTGCCGTTGGATCTTAGTGTTATTTTATTTAACCACGCCTCCGAAGAGGCGGCCGCCAGTACGGCCTCTGAGCCAGAAGATAAAA | 4 |
| wt-68550-68670 | 1996 | GTCCCCGGGTGCAACCACTGGGCACAGGACTCCAACAACGCCCCAGACATACGGTCAGTCCATCTCCTTCACCTCCGCCACCTCCTAGGACCCCTACTTGGGAGAGTCCGGCAAGGCCAG | 4 |
| wt-68670-68790 | 1997 | AGACACCCTCGCCTGCCATTCCCAGCCACTCCAGCAACACCGCACTGGAGAGGCCTCTGGCTGTTCAGCTCGCGAGGAAAAGGACATCGTCGGAGGCCAGGCAGAAGCAGAAGCACCCCA | 4 |
| wt-68790-68910 | 1998 | AGAAAGTGAAGCAGGCCTTTAACCCCCTCATTTAACACCATGTTCTCGTGCAAGCAGCACCTGTCCCTGGGGGCCTGTGTCTTCTGTCTCGGCCTCCTGGCCAGCACCCCCTTCATTTGG | 4 |
| wt-68910-69030 | 1999 | TGCTTTGTCTTTGCCAACCTGCTCTCTCTGGAGATCTTCTCACCGTGGCAGACACACGTGTACAGGCTTGGATTCCCGACGGCATGCCTAATGGCCGTCCTCTGGACGCTGGTACCCGCC | 4 |
| wt-69030-69150 | 2000 | AAGCACGCGGTGAGGGCCGTCACTCCAGCCATCATGCTGAATATTGCCAGCGCCTTGATCTTCTTCTCCCTCAGAGTCTACTCGACCAGCACGTGGGTTTCTGCCCCCTGTCTCTTTCTG | 4 |
| wt-69150-69270 | 2001 | GCCAACCTGCCTCTCTTATGCCTGTGGCCCCGGCTGGCCATCGAGATTGTTTACATCTGCCCGGCTATACACCAAAGGTTCTTTGAACTTGGGTTGCTCTTGGCCTGCACCATCTTTGCC | 4 |
| wt-69270-69390 | 2002 | CTGTCCGTGGTCTCCAGGGCCCTGGAGGTGTCGGCTGTCTTCATGTCTCCATTTTTCATCTTTCTGGCTTTGGGCTCTGGAAGCCTGGCCGGTGCTCGGCGTAACCAGATTTACACCTCG | 4 |
| wt-69390-69510 | 2003 | GGTCTCGAGCGGAGACGCAGCATTTTCTGCGCCCGGGGAGATCATTCGGTGGCATCCCTGAAGGAGACCCTCCATAAATGCCCGTGGGATCTGCTGGCCATCTCTGCCTTGACCGTTCTT | 4 |
| wt-69510-69630 | 2004 | GTCGTCTGTGTGATGATTGTGTTGCATGTGCACGCAGAGGTGTTCTTTGGACTCTCTAGATACCTGCCCCTCTTTCTCTGTGGGGCGATGGCCTCCGGGGGGCTGTACCTGGCCATTCC | 4 |
| wt-69630-69750 | 2005 | AGCATCATTGCATGTGTCATGGCCACCCTCTGCACCCTGACATCTGTTGTGGTATATTTCCTCCATGAAACCCTTGGACCCCTGGGCAAGACCGTGCTGTTTATCTCAATCTTTGTCTAT | 4 |
| wt-69750-69870 | 2006 | TACTTTAGCGGGGTAGCGGCCCTGAGCGCAGCTATGCGCTACAAGCTTAAGAAGTTTGTGAACGGACCCCTGGTCCATCTCCGTGTGGTATACATGTGCTGTTTTGTCTTTACTTTTTGT | 4 |
| wt-69870-69990 | 2007 | GAATATCTGTTGGTGACATTCATTAAATCCTAACGACCGGAGTCCTGTCTCTTTGTGTTCTTGGGGGACTTGAGTTAGCTGTCTTTCCTCTTATTACATTGGGCTAACGGGAGGAAATGA | 4 |
| wt-69990-70110 | 2008 | ACCCAGGGGTGGCAGTGGATGGGGTCATTTATGGGCAAAACTCACAGGACATGTTTGGGGAGTTAGCATTGGCGTCGGGAAACACAGCTCTGGCAGTTATAACCGCACCAGCTAACAGGA | 4 |
| wt-70110-70230 | 2009 | CATGTTTGGGGGAGTTGGCATTGGCGTCAGGAGACACGGCTCTGTCAGTTATCACCGTACCATGAGTGCCATGTGTGTCCAGTGCCTAATCACCGTTCCTCATTTTGTGTGCCTCCTCAA | 4 |
| wt-70230-70350 | 2010 | ATGTTCCAGAAGTCGGCCACAGGGGAGGTGGCTGAATTAGGGCCTTTTCCCTCATTCCCCCATGAGACCCACGTGGCAGGCCTAGGGGCTACATTCGCCTCCCACGTTTCCCTTCGCGTG | 4 |
| wt-70350-70470 | 2011 | AGGCATCCGATATGACTGAATTTTCGCAGTCTCTTTTCCCTCTTCCCTTGTTATTCCCATAGAATTACAGTGAGGTTACACAGGTGGAGATTCAGTTTAACCATTTATTGATTTAATCCA | 4 |
| wt-70470-70590 | 2012 | GGAACAAAAAACAGTCCTAGTGACCCAGTGCCCGGAGAGAGAATGGCCCTGACAAGTCGGCTGCATGATGCACTTCGGCAGTCACGTGTGTGAGTCTCCACGGCCTCTGTCAAAAGGGAG | 4 |
| wt-70590-70710 | 2013 | CTTAGCGTGCCAGGGTTGTAATTCTTGATGTAGTGGCCCAGGAATTCAACTTCATCGTGTCTCCGTCTGCAGTTGGCGTTAATGTAGGCTGGGGCTACTGCCGCATATGCTGCCAAGAGA | 4 |
| wt-70710-70830 | 2014 | CAGAGGGGCTGCTTCACATATGAGCTGCTCAGGGTCTCCACCACCTTGTTTTGACGGGCCGTGGCACAGGTGATGTAGAAGAGTTGCTTCACAAAGTTGTAGTCTCGCGTGTTAGGAAGG | 4 |
| wt-70830-70950 | 2015 | AAGCAGGGTGCCAGCTCTTTGAGCTTGGTCAGGATCACCTTGCTAAGACTCATGGCGCAGGCCAGGAGGATGTCTTCCGCGGGAGCTAGGGGCAGGTCGCCGTGGTAGGTGATCTCCTGG | 4 |
| wt-70950-71070 | 2016 | AGCCAAAAGATGGTCTCTTCTAGCATGGCCACCAGGGTGCAGAGCCCCGCGTTCTGGATCGCCTGCATGCGTGCATCCAGCCATGTGTCCTTGTTGGTTGACTTGGTGAAAAACTCACGT | 4 |
| wt-71070-71190 | 2017 | AGTGTCTTGTAGCTCCTGCGCAGCTCGTGTCTGGGTTGCACTTTCTGCCAGGCTCCAATCTCTGGATGGGCGGCCACCGCCAGCATCGACTGTAGGAACGGGTCTTGGATGGGCTCTAGG | 4 |
| wt-71190-71310 | 2018 | GTCAGAGAGGCCAGGGGGCTGGGCAAGGTGACAAATGTAATCTTGGAGACAGGCTTAACCAGACTCATGTCAAACCACGGTTTGTTCGGCAGGGGCCTCTGGCTGCGTTCTTGCCTCGCC | 4 |
| wt-71430-71550 | 2020 | CTGGGCGCTCTTGCTCCCGGGGGCAGGTGTCGGCGCTTGCCATAACTTTCATCGGTGCAGCCATGGACCTCTCCGCGTCGCCTTTTGTGGCCTCTGGTGTAAGAGGAGTTGCCAGTCTCC | 4 |
| wt-71550-71670 | 2021 | TCCTTCTCGTCCTCGTCCCTGCACAGGGGTGAGCGATGCAATGTGACTGTCTTGTCCTGTAGGTCCCACTTCTTTCTGGGAATCACAAACGATGCCGAGGTAGGGGTTATGACCACGCTG | 4 |
| wt-71670-71790 | 2022 | GAGGGCCGTGCAGGTATGGCGTGGGCCGGAGTTGGATCTTCATCCTCCTCCTCTGAGGATGAAATCTCTCCATCTGTGGAGTGTTCTTCGCTGCCCTCCATAGGGTCCAGATCGCAGTCT | 4 |
| wt-71790-71910 | 2023 | GTGTTGGTGTCTGAGACCGCTTCGAGTTCCAGAATGTGGCTCTCTGCAGAGGGGAGACAAAAGGTGGAGACTGCCTTGAGCACCTCTGTCTCAGGCACCGGATGCCCCCGGCTCCACGGC | 4 |
| wt-71910-72030 | 2024 | CCCGGCCACTGGCCGGTGTAGCTTCTTACCTGCGGGATCCTCGTTGGAGGAAATGCTGCTAGTTCGGGAGAGTCTCTGAGAAGGAACCATCTTGTCTGTCTCTACGACGGGCTAGCTGGG | 4 |
| wt-72030-72150 | 2025 | ATGTAGTGCTGTCTTGACTGGCCTCAGCCCTATTTATGATTCTGGAGGCGGGCACGCTGATGGAGAAATGGGCGGTCGGTTGATTGGCCCCACAGCGACCGGCGAAGCACTGACTCATGA | 4 |
| wt-72150-72270 | 2026 | AGGTGACCGTGATGGCCTGTGATGTGTAGTAGAGTACCAGAAACACCCTCACATTCTTGGAGCTGGCCCTGTGGGTATGCCTCAGGCACGCAAAGTTCCTGCCCCGGGCATGGCACACCT | 4 |
| wt-72270-72390 | 2027 | GAACTAAGTTTGGCCCGGTTTGCTCAAACGTGACATGGAGAAACTGGGGGAATTTGTCTTCTGGCACAGCTGTTGCCAGGGTGCTCATGAGCGAGGGCCAGATGCAGGAGCTGACCCAGG | 4 |
| wt-72390-72510 | 2028 | CGACGAGATCCAGGCCCAGATGTCCCTCTATCATGGCGCAGACATTCTCCACGGTGGGGGGCAGGGTCTCGCGGGTCCTCTGGATTAGATAGTCACGCCCATCATCCGCGATGTGGTAGC | 4 |
| wt-72510-72630 | 2029 | AGAAGGTTTTGGGGGCCGGCCAGCCCACGTGCAGTGAGTGATGTAAGAGGTTTTGAATGTTGAGGGCATTCTTAACATAGCTGTGCTTGTCTTCCTCTTCCGGATGACAGACAAAGAGGC | 4 |
| wt-72630-72750 | 2030 | GCAGCTGCCGGCTAAGACCACCGCCCCTGTCCACCTTGTAGGTATGCGGCAGCCGGATGCACCGCCCGGCGTGATACACGCCGCTGTCAAAAAGCGGGGCCCCAATCTCTTTGATCTTGT | 4 |
| wt-72750-72870 | 2031 | GACGCATGCGGCGCAGGCAGGCCGTTAGGCCCATGAGCTTCTGCAGCACAGACACAAACCCTTGTACTGCGCTTGTTCCCACAATAGCATGGCCTCTAGGTAGGGGGGTGATGACGCGAA | 4 |
| wt-72870-72990 | 2032 | AGCCCAGTTTTCCCGTGCATATGCAAAAGGGGAGCACATCTTCCATATTATCCGGGTCGGCGGGTGGACAAGCTGATTTGAAAAAATAGACTGGGTGGGCCCTGGACACTGGACCCAGGC | 4 |
| wt-72990-73110 | 2033 | GGCGCATGAGGCGCAGTACCTCACGCCGCACGGTCCGGCACAGGTCATAGATTTCCTCCAGCGACCAGGGGGCCCCCTTGATCTTTAGATCCAGGTCCAAGACCAGGTTGCAGACCGGAA | 4 |

| | | | |
|---|---|---|---|
| wt-73110-73230 | 2034 | GCCGGGGATTAAAGTATTCATGCCGGGAGACAAAGAGCTGCTCGCTCAGGCTGTTCTGTGAATAGTACACTGGGGTGTAGGAGAGGGCCCTGGTGAGACACGTGTCTGGGAGGCGGCAGT | 4 |
| wt-73230-73350 | 2035 | TGGTCGGGGTGGAGACGACCTCCGCCAGGTGGGATGAGAAGGGGTCAGCGGCTGTCATTACAAAGTAGTGCCTGTCTGCAAAATGGCAGAGGAAGACCGGTAGCCGCTGCACCCTTCGAA | 4 |
| wt-73470-73590 | 2037 | AGCGCGCGAGCTCCACCAGCTCCCGGGCCACGTCCACACTGAGCCCGGCCCAGGCCCGCATGAGTCCGTCATCGGGGTCGGGGTCCCACGTGTATGGGGCCGGGGGCTCCATGCGGATTT | 8 |
| wt-73710-73830 | 2039 | ACCCAATTTCTCCCATGCTCGTTAACTGAGAGTATCGCTTAAAGGTTCCCTCGTTGAAGCACTGTGCGTGGGCCAAATAGACGTAGCGCACGAGATCGGCCGAGGCCAGGGGAAGGCGCC | 4 |
| wt-73830-73950 | 2040 | CCCTGTAGGCGTCTATCGTCCTTGCCACAGCGCGGATCTCTCGCGAGTCCCGCCGCAGTTTCTCGTGTGCAAAGTGGGCAAAAGCCTCGGTCTGCTCCGCCCATGCCGAGGAGCCAAAGA | 4 |
| wt-73950-74070 | 2041 | CCTCCCCCAGCTCGGCCAGGGACGTGACGGCGGCCAGGCTCTGACCAGACTCGGAAGTAAATAGCTCCGTGAGGTGCGCCAGGGTCTCAATCGTACAAGGAATGCCCCAAAAATAGTAAG | 4 |
| wt-74070-74190 | 2042 | CAGCCGTGACTAGCACGAACTGGGCCTCGTGGGAGCCAAAGGTGCTAATGAACCACCTGGCCGAGATGTTAACGCGGTAGATGCGGCGCAGACAGCCCACGATCTTGGGACGCAGCCACG | 4 |
| wt-74190-74310 | 2043 | CCACGCGGCCTCTGGCATCCCCCTGTGGCTGTTTCTTAGCGCTCAGTGTGAGCAGTTCCACGAGGGGCGTGAGCGAGCGCAGGGCCCCCGCGCGATCTAGGTAGGTGGATAGACGGTCCG | 8 |
| wt-74310-74430 | 2044 | CGGTGAGCGGCGTGAGGCCGCGCAGGAAGGGGAAGGCCTCCTCCGCCGGCAGGTGCAGCGTCAGAACCAGGCCGCAGCGGCTCTGTGAGGTCAGCCGCTTCTTGGGCAGGTGAAGCTGCA | 8 |
| wt-74430-74550 | 2045 | GTTCCACGAGAGAACCCGCCACGTGGTGGAGGGGCGAGGCGTTGTGGCACAAACAAAACAGGCGGAAGCCCTCGTCAGGCCGCGAGAGGATGGCATCGAGGATGGCCTCCGCAATGTCAG | 4 |
| wt-74550-74670 | 2046 | TGTTTGAGGCCACAAGGGCCTTGATGACGACGGGGGCGGACATTATTTAAGACCGGGAGGCCCCAACGGCGGGCTAAACAGAACGATGGCCTTCTATCTCCCAGACTGGTCGTGCTGCGG | 4 |
| wt-74790-74910 | 2048 | GGTGGGAGATTTGCTAAGAAATGAGCAGACGATGGGCTCACTGAGACAGGTTTATTTGCTCGCTGTTCAAGCCAATAGCATCACGGATCACCTGAAGCGCTTTGACGCCGTCCGCGTCCC | 4 |
| wt-74910-75030 | 2049 | TGAGAGCTGTCGTGGGGTGGTGGAGGCCCAGGTGGCCAAGCTTGAGGCCGTGCGCTCAGTCATCTGGAATACCATGATCTCTCTGGCTGTAAGCGGCATCGAGATGGACGAGAATGGGCT | 4 |
| wt-75030-75150 | 2050 | CAAGGCCCTGCTGGACAAACAGGCTGGCGACAGCCTGGCCCTGATGGAGATGGAGAAGGTGGCCACGGCGCTCAAGATGGACGAGACCGGTGCCTGGGCGCAAGAGATCTCGGCCGTTGT | 4 |
| wt-75150-75270 | 2051 | CTCATCGGTGACCGCCCCCTCAGCCTCGGCCCCTTTCATCAACTCCGCCTTTGAGCCCGAGGTGCCCACCCCCGTCCTTGCACCGCCTCCCGTGGTGCGGCAGCCGGAGCACTCTGGGCC | 8 |
| wt-75270-75390 | 2052 | CACGGAGCTCGCGTTAACGTAGCAACCAGACTCCACACCAAATAAACATTTTATTGGTAAAACAAGGGATATGAAGGTGTCATTGACCCGAGGATCCAAACCCCCTCCCCTGTCTCCCCT | 4 |
| wt-75390-75510 | 2053 | CGAGCGCCTCGCTCAGCCCACTATCACCCATGGCCAGGCTCGGCACCTCCTCGAAGGTGCAGCTGGCCCACCTAAAGAGAGATCTGGGGCCAAGGACCCCCGCGTCACTGTGGGGGCTGT | 8 |
| wt-75510-75630 | 2054 | AGAAGGAGGTGAGGTGGTGCTTGTGAAGGTAAACAAGCTGACAGAAGCGCCGGTACTTGTTAAGGAACACGGTCTGGTCACTAAAGTTGGTCAGGCTGACGTCCACCCCACCCCGGCGCC | 4 |
| wt-75630-75750 | 2055 | ACCTGCAGGGCTTCACTAGAATACCCTGCATGGCCAGGCCCGACCTGCCAAAGATTGTCGGCCTGTGGTGAGGGATAGAAGGGGGGGGCACGGTGAGTGTCACTGAGACGGTCTGATGGG | 4 |
| wt-75750-75870 | 2056 | GGAAGAGGGCCAGGTCCTTTGGCAAAGAGACGTCCAGGCCCACGTCCCCGGGGTACTGGGGGTGGTTGATGGGACCCTTGTCCTCCTCCATCTGGGGGGTGGCATATCTGAAGGCAGCCA | 4 |
| wt-75870-75990 | 2057 | GGTGGATTTTGAGCTCCGATGGACGCAGCGTGGAGTTGTAGCGCCGCTGATTCTGGAGGATTAGCCGGAGTTCCCCCGTGTAGCCGGGATCGATGATGCCAACATGAGACGTGACCGGAC | 4 |
| wt-75990-76110 | 2058 | GGGAGGTGCTGCCCCACAGCATGAGCCCATGACCCTCGGGTGGGCGGGCATAGAGGCCTAGGTCCACAGTTGTGGTCTTCATCGGGCGCAGCAGGATGGTGGTCTTGTTGACCAAGGTGA | 4 |
| wt-76110-76230 | 2059 | GCCGCCCTACACTAGCCTGCTGGAGCAACAGCTTGTCATTCTGGAAGGCGTAGCGTATGTGTGGACAGGCCTCCATGGTGATGATCTAACAGACAGGGACGGCGGCGCTATATATAAGAG | 4 |
| wt-76230-76350 | 2060 | CCCAAGACCCGGCTCTCTTTACTGCGAAATGGGGAAGGTCCTAAGAAAGCCGTTTGCAAAGGCTGTGCCACTGCTCTTCCTCGCCGCCACCTGGCTTCTGACCGGGGTGCTGCCGGCCGG | 4 |
| wt-76350-76470 | 2061 | CGCTTCCAGTCCCACAAACGCGGCGGCGGCTTCCCTGACTGAAGCCCAGGACCAGTTCTACTCCTACACATGTAATGCGGACACATTCTCGCCTTCTTTGACCAGCTTTGCCTCCATCTG | 4 |
| wt-76470-76590 | 2062 | GGCACTTCTGACGCTTGTCTTAGTCATTATAGCCTCAGCCATCTACCTGATGTACGTCTGCTTTAACAAGTTTGTGAACACGCTGCTGACGGATTAGATGGGGATATTTAAAAGGGGCAG | 4 |
| wt-76590-76710 | 2063 | CAATCTCGGCTGTTTGTACTTCTTCTCTGCTCGTTAAACCAATAGCATGTCAGCTCCACGCAAAGTCAGATTGCCTTCTGTTAAGGCTGTTGACATGAGCATGGAAGACATGGCCGCCCG | 4 |
| wt-76710-76830 | 2064 | CCTGGCTCGCCTGGAGTCTGAGAATAAGGCTCTGAAGCAACAGGTCCTCAGAGGGGGTGCCTGTGCCTCGTCTACCTCTGTTCCTTCTGCTCCAGTGCCTCCGCCTGAGCCGCTTACAGC | 4 |
| wt-76830-76950 | 2065 | TCGACAGCGAGAGGTAATGATTACGCAGGCCACGGGCCGTTTGGCGTCTCAGGCTATGAAGAAGATTGAAGACAAGGTTCGGAAATCTGTTGACGGTGTAACTACCCGCAATGAAATGGA | 4 |
| wt-76950-77070 | 2066 | AAATATATTGCAAAATCTGACCCTCCGCATTCAAGTATCTATGTTGGGTGCAAAAGGCCAACCCAGCCCTGGTGAGGGAACACGACCACGAGAATCAAACGACCCCAACGCCACCCGACG | 4 |
| wt-77070-77190 | 2067 | TGCCCGCTCCCGCTCCCGGGGACGTGAAGCAAAGAAAGTGCAAATTTCTGATTAATAAATTTTTATTGACTTTATACATAGGTCTCGGCGTCATCATATGGTGGGGTGGTGTAGGTATGG | 4 |
| wt-77190-77310 | 2068 | GATGTAGACAAGTTACGCCTAAAGGCGCAGTCCGCCATGACCAGCAGCAGCAGAAGGGTCAGCACAGCCAGAGAGGCCCACTGCAGTACTAGCATGGAGAGGTTTGAGAATCTGGGCTGG | 4 |
| wt-77310-77430 | 2069 | GACGTTGGCGGGACTGGCACGGTGGCTTGGGCTGTGGTAACCGGTGGGCTCGTAAAAGTCCAGCGGGGCCGCAGTTTGCTAGAAGTGCTGGGAGGTAGATAGGTGGTCGCATTGTATCTC | 4 |
| wt-77430-77550 | 2070 | GGTCTTGGCGTAGTTGAATCACCGCCGTAATCTGTGGTGGGCTCTGTACTTGTCCGGGCTCCATGTCCTGTGGTGTGCTTTCCACCGGTGGTAGAATTGGCCTTTCCACCTGTTGAGGTG | 4 |
| wt-77550-77670 | 2071 | ACCGTGGGAACCGCCGTCTTTTGGCCACTGGGGGCCTGGGGCGACGTTGCATTTTGGGGGGGCGTGCCTTTGGTGACATTAACCTCCCCCGGTTTTGTGGATGTGGAACTGTTTCCAGGG | 4 |
| wt-77670-77790 | 2072 | CCTGACGCTTGGCTGGTGGTGCCTGGGCGGGGTGCTGGCGAACTGGTGGACACATGATGTGTGCTGATAGAGGCTGGTGTCACCTGTGTTATATTTTCACCACCTGTTGGGTGAGCGGAG | 4 |
| wt-77790-77910 | 2073 | GTTAGTAAAGGCATATGTGACGTTGAATTGTCACTGGTGGAGGGGCTGAGTGTCTCTGGGTTTGAACTGGGTCTCAGTGACATGGAAGAGGTTGAACTTGAAGTTATGTTATGTTGGCCT | 4 |
| wt-77910-78030 | 2074 | GTGGTAACAGCACTGGTTGCATTTTTTGGTTGGCTGGTAACTACTGGGGTGGGACTTGTTCCTCCTAAGGTGTGGTTGGTGGCATTTGCCTGTGGACTTGTTTCTCCCACAGTAGGGCCG | 4 |
| wt-78030-78150 | 2075 | GTGGCATTTGGGGTTGGGGTAGTCACTGCTGAGGTGGGGCTTGTTTTTCCCAAGGTGGGGCTGGTGGCATTTGGGGTTGGGGTAGTCACTGCTGAGGTGGGGCTTGTTTTTCCCAAGGTG | 4 |
| wt-78150-78270 | 2076 | GGGCTGGTGGCATTTGGGGTTGGGGTAGTCACTGCTGAGGTAGGACTTGTTTTTCCCAAGGTGGGGCTGGTGGCATTTGGGGTTGGGGTAGTCACTGCTGGGGTGGGGCTGGTGGCATTT | 4 |
| wt-78270-78390 | 2077 | GGGGTTGGGGTAGTCACTGCTGGGGTGGGGCTGGTGGCATTTGGGGTTGGGGTAGTCACTGGTGAGGTGGAGCTGGTCATGTCGGGGGCCTTACTTTCTGTGCCGTTGTCCCATGGAGAT | 4 |
| wt-78390-78510 | 2078 | GGACTTGGTGTCACCGGTGATGCGCCTGACGTTGTGCCGGCTGGTGTTGGGCTGGTGACATCCGCGGTGGATACAGTGGGGCCTGTGCTTGCAGGTGCGGTGAGGTTGGTAGGCACGTGA | 4 |
| wt-78510-78630 | 2079 | GTAGAGCTGGGTAGACCTGTCGTTGTATTGGGATCAGCAAATCCAGTTGTATTCAAGGTAGGGGAGGTGGTGGTGCTCTCGGGTGCCTTGGAGAATATAACCTTGTGGGTTGTTGTGGTG | 4 |
| wt-78630-78750 | 2080 | GCATTGGTAGCCGTTCGTGTGATAATGAGTGTCTTGGGGGCCGTGCCAAGACCCGAGACAGTAATGTCAAATGTCCGATTGCTCGCAAATGCACCAGAAATATTTTCACAACCCGAAGGT | 4 |
| wt-78750-78870 | 2081 | GTCCCCGAGGTGAGAGTCCATTTGCACTTAAAGTCAGTTTCAGTGTTGTTTGGCCAGGCCCAAAAGGCAGTCACTGTAACATTTGGCGAGTTTGCGTCCTCAGAAGTGACCATTGGCACT | 4 |
| wt-78870-78990 | 2082 | GAATAGGTAGCATTGTCACCCACATATGTGATGTCTGTGGTGTTTGTCGGCATGTCCTGTGAAGCTGGAATCTCATCAGAGAACACAATGTTGGACTGAATGCAGTAATCTCCCCCGCTC | 4 |
| wt-78990-79110 | 2083 | GCCTTCGGTCCATTCCCAGAGTAAAACACGTACAGGATACTGTTATTGCCAAGAAATCGTGACACTGGACGTGGTGTCAGACGCAGGCTGTATGCATACCCTGTACCAGGTATTGGGGTG | 4 |
| wt-79110-79230 | 2084 | GCCACGGGACTCGTTGATGTGAGAATTCCGCCGCTGGGAACATGGCTCTCGTATCCACTGCAGGTGATGTTAAATTTGTTGTCTCCGGGCAGAACTTGTGAAATTTCGCCATCCTCCATA | 4 |
| wt-79230-79350 | 2085 | ATACACTCAATATCTATCTCATTACCGAGCATTTCTGTTTTTACGCTGAAATTCGAGTCTTGAGCTGACGTTGGCAAACTTAAGGGTAGCGTGACATCCAGCCCCTGTGCCCTCACTACT | 4 |
| wt-79350-79470 | 2086 | GCCGTTATATTGGTAGAATTACAGTTATCCCACTTTATGTATGGCACTGTTTCTGGTATCAGGTACACGGGGTTTTGCATTTCTGCATGGTGGCACCACATGGTTCCAAACACATCTTGA | 4 |
| wt-79470-79590 | 2087 | AAGTAGACATCTACAGATTCCAGGCTTACTTGTTGCTCCTCTCCGGTGGTGACGTTAATTGGAAGCTTCTTAGACCGCATAGTTAGAGCCAATTCTCCTGCACCAAGGAGCTCCAGTAGA | 4 |

| | | | |
|---|---|---|---|
| wt-79590-79710 | 2088 | AAGAGATTGGTGGCATTTTCTGAGCCACCAAATGCACCTCGAGGTTGGTAGACAGCCTTCGTATGGGGTGTCAGCTGGCCAAAGTCAAGATCAAGTTGATGCTTTTTGCCCCCGACATCG | 4 |
| wt-79710-79830 | 2089 | AAATTGATAGTTACATTGACATCTGCCGTGCAAACATTGCATGTGGGGTAAAATGGGAATTCCGGAATCTCAACATTGAAAAAACCAGGATCTTCACCCGTGAGATGGATCAGGCTCTGG | 4 |
| wt-79830-79950 | 2090 | ATGGTGTACTGACACACAAGCAAGGCTGCCTCCATTGTCTCGGCACCGATTTCTAGGCAGCATCCTCTTTAATAGGTACAAGGGGGGTGCGGTGTTGGTGAGTCACACTTTTGTTGCAGA | 4 |
| wt-79950-80070 | 2091 | CAAAATGGACAAGGACAGGCCGGGTCCCCCGGCCCTGGATGACAACATGGAAGAAGAAGTCCCATCTACCTCGGTTGTGCAGGAACAGGTATCGGCGGGAGATTGGGAAAATGTCCTCAT | 4 |
| wt-80070-80190 | 2092 | AGAGTTATCAGATAGCAGCTCAGAAAAGGAAGCAGAAGATGCCCACCTGGAGCCGGCCCAGAAGGGTACGAAGAGAAAGCGGGTCGATCATGATGCCGGTGGGTCAGCTCCAGCACGACC | 4 |
| wt-80190-80310 | 2093 | CATGCTCCCACCCCAGCCGGATCTCCCTGGGCGAGAAGCCATTCTCCGCAGGTTTCCACTAGATCTAAGAACACTTCTTCAAGCGATTGGAGCCGCGGCTACGGTGAGCATCCCTATGGC | 4 |
| wt-80310-80430 | 2094 | CTAAGTGTGTGATGTGTGTTTTTACCCATCACACAACAACAAGGTAAGTAATTTGTTGCCGTTGGTTTCAGCGCATCGACACACGAGCCATAGACCAGTTTTTCGGATCCCAGATTTCAA | 4 |
| wt-80430-80550 | 2095 | ATACCGAGATGTACATAATGTATGCCATGGCCATTCGACAGGCCATTAGAGATCGTCGGAGAAATCCAGCTTCTCGTAGAGATCAGGCCAAATGGAGACTGCAAACCCTGGCCGCCGGAT | 4 |
| wt-80550-80670 | 2096 | GGCCTATGGGTTACCAGGCATACAGCAGCTGGATGTACAGCTACACCGATCACCAGACGACTCCCACATTCGTACATCTCCAGGCGACACTTGGGTGCACAGGTGGCCGTAGGTGTCACG | 4 |
| wt-80670-80790 | 2097 | TGACCTTTTCCGCCGGCACCTTTAAGCTGCCGCGATGTACCCCCGGGGATCGCCAGTGGTTGTATGTTCAGAGCTCCGTGGGTAACATTGTACAGAGCTGTAATCCCCGCTACAGTATTT | 4 |
| wt-80790-80910 | 2098 | TCTTTGACTATATGGCTATACACAGGAGCCTCACGAAAATCTGGGAGGAAGTTTTAACACCTGACCAGCGTGTTTCATTTATGGAATTCCTAGGATTTTTGCAGAGAACGGATTTGTCCT | 4 |
| wt-80910-81030 | 2099 | ATATCAAGAGCTTTGTCAGCGATGCCCTGGGCACCACTAGTATCCAAACACCGTGGATCGATGACAATCCTAGCACGGAGACGGCACAGGCTTGGAATGCCGGCTTTCTCCGGGGTCGTG | 4 |
| wt-81030-81150 | 2100 | CGTATGGGATAGACTTGCTTAGAACTGAAGGGGAACATGTCGAAGGTGCTACCGGTGAAACGCGAGAAGAAAGTGAGGACACGGAGAGCGATGGAGATGATGAAGATCTTCCTTGTATAG | 4 |
| wt-81150-81270 | 2101 | TGTCCAGAGGTGGACCTAAGGTCAAACGACCCCCTATATTTATAAGACGTCTGCACAGGTTGCTGCTGATGAGAGCGGGCAAACGAACAGAACAGGGCAAGGAGGTACTGGAAAAGGCCC | 4 |
| wt-81270-81390 | 2102 | GTGGGAGCACTTATGGCACACCTAGGCCGCCTGTTCCGAAACCAAGACCAGAGGTCCCACAAAGCGACGAGACAGCTACCAGTCACGGGTCGGCGCAAGTCCCAGAACCCCCAACCATTC | 4 |
| wt-81390-81510 | 2103 | ACCTAGCAGCTCAGGGAATGGCATACCCATTACATGAACAACACGGCATGGCCCCGTGTCCGGTAGCACAGGCCCCACCTACGCCCTTGCCCCCTGTATCTCCAGGGGATCAACTCCCAG | 4 |
| wt-81510-81630 | 2104 | GTGTTTTTAGCGACGGGCGAGTGGCGTGTGCACCAGTACCCGCCCCGGCTGGGCCTATTGTCCGGCCCTGGGAGCCATCCCTGACACAGGCTGCGGGGCAGGCCTTTGCACCCGTTAGAC | 8 |
| wt-81630-81750 | 2105 | CACAACACATGCCAGTAGAACCCGTCCCTGTCCCGACAGTGGCACTTGAGCGACCAGTTTACCCCAAGCCAGTTCGTCCGGCACCTCCTAAGATTGCTATGCAGGGCCCCGGGGAAACTT | 4 |
| wt-81870-81990 | 2107 | TCAAACAGGCTAGTGTTGAGGTGCAGCCCCCCCAGTTGACCCAAGTATCCCCTCAGCAACCAATGGAGGGGCCGTTGGTACCAGAGCAGCAGATGTTCCCTGGTGCCCCCTTTAGCCAGG | 4 |
| wt-81990-82110 | 2108 | TTGCTGATGTGGTCCGGGCACCTGGGGTACCGGCGATGCAGCCACAGTACTTTGACCTCCCCTTAATTCAACCCATTAGCCAGGGGGCACCCGTGGCCCCGTTGAGGGCTAGTATGGGCC | 4 |
| wt-82110-82230 | 2109 | CGGTACCTCCGGTACCGGCAACACAGCCACAGTATTTTGACATCCCCTTAACTGAACCCATTAACCAGGGGGCATCCGCGGCCCATTTTCTCCCTCAGCAACCGATGGAGGGGCCGTTGG | 4 |
| wt-82230-82350 | 2110 | TACCTGAGCAGTGGATGTTCCCAGGTGCCGCCCTGAGCCAGAGTGTTAGGCCAGGGGTAGCGCAGTCACAATATTTTGACCTCCCCTTAACTCAACCCATTAACCATGGGGCACCCGCAG | 4 |
| wt-82350-82470 | 2111 | CCCATTTCCTCCATCAGCCACCAATGGAGGGGCCGTGGGTACCCGAGCAGTGGATGTTCCAAGGTGCCCCCCCTAGCCAAGGCACTGACGTGGTCCAACATCAGCTGGATGCTTTGGGGT | 4 |
| wt-82470-82590 | 2112 | ATACACTCCATGGTCTTAACCATCCCGGGGTTCCCGTGTCTCCTGCCGTTAACCAATATCATCTCAGCCAGGCTGCCTTTGGGTTACCTATTGATGAGGATGAGAGTGGCGAGGGGTCCG | 4 |
| wt-82590-82710 | 2113 | ATACCTCCGAGCCGTGTGAAGCTCTTGATTTGTCAATCCATGGCAGGCCCTGCCCTCAGGCCCCCGAGTGGCCTGTTCAAGAGGAGGGTGGCCAGGATGCCACCGAGGTTCTTGATTTGT | 4 |
| wt-82710-82830 | 2114 | CAATCCATGGCAGGCCCCGCCCTCGGACCCCCGAGTGGCCTGTTCAAGGGGAAGGTGGCCAAAATGTCACAGGCCCTGAAACTAGAAGGGTGGTGGTGTCAGCTGTTGTTCACATGTGTC | 4 |
| wt-82830-82950 | 2115 | AGGATGACGAGTTTCCGGATCTACAAGATCCTCCAGATGAGGCCTAAGCAAAGGTGTAGAAGTGTGTCCCCCTCCATTCCACCCACTGATAATACGCCCGACAATAAAGTTGATGATATT | 4 |
| wt-82950-83070 | 2116 | GAATTCCACACCTGCTTGTGTTTGTGATTTTATTTCATATTCCATGAGAGAGACCTCGCATATTTGCAGAAGGGTCACTGAAACATCTTATCTTAAAACAGTTACACCTGAATAATGAAG | 4 |
| wt-83070-83190 | 2117 | AAAGCGTGGCTCAGCAGAGCACAGCAAGCCGATGCCGGGGGGGCATCTGGCTCCGAGGACCCACCAGATTATGGAGATCAAGGTAATGTGACACAGGTGGGATCTGAGCCTATTTCACCT | 4 |
| wt-83190-83310 | 2118 | GAGATTGGCCCCTTTGAACTCTCTGCGGCCAGTGAGGATGATCCTCAATCTGGGCCAGTGGAAGAGAATTTAGATGCCGCTGCAAGAGAGGAAGAGGAACCTCATGAGCAGGAGCACAAT | 4 |
| wt-83310-83430 | 2119 | GGTGGTGACGATCCCTTGGATGTCCATACTCGCCAGCCTAGATTTGTGGATGTGAACCCAACGCAGGCTCCAGTGATCCAACTAGTCCACGCTGTCTATGATTCCATGTTGGTAAGAGGC | 4 |
| wt-83430-83550 | 2120 | ACCTAGAACATTTCCAGATGTTTCGCTTGGATTTTTTGGCCAGTCTTAATTGATTGTCATTGGTTTCAGCAATCGGACCTCCGGCCCCTAGGCAGTTTATTCCTTGAGCAAAACCTGAAC | 4 |
| wt-83550-83670 | 2121 | ATCGAAGAATTTATATGGATGTGCATGACCGTGCGTCACAGATGTCAGGCCATCAGAAAAAAACCATTACCAATTGTTAAGCAGAGGCGTTGGAAGCTCCTGTCATCTTGCAGATCCTGG | 4 |
| wt-83670-83790 | 2122 | CGTATGGGTTACCGCACGCATAACCTCAAAGTAAACAGTTTTGAGTCAGGGGGGGACAATGTCCACCCGGTCCTTGTGACTGCTACGCTAGGATGTGATGAGGGCACGCGGCATGCAACA | 4 |
| wt-83790-83910 | 2123 | ACGTACAGTGCTGGCATTGTACAGATACCACGAATATCAGACCAAAACCAAAAGATCGAAACAGCCTTCCTGATGGCACGTCGTGCTAGGTCACTTTCGGCAGAAAGATATACTTTGTTC | 4 |
| wt-83910-84030 | 2124 | TTTGATTTAGTATCCTCCGGAAACACCCTGTATGCTATATGGATTGGGCTGGGCACGAAAAACCGAGTTTCATTTATTGAGTTTGTAGGATGGTTATGCAAGAAGGACCACACTCATATA | 4 |
| wt-84030-84150 | 2125 | CGCGAATGGTTCCGCCAGTGCACCGGGAGACCCAAAGCAGCCAAGCCATGGTTAAGAGCGCATCCTGTCGCCATTCCTTATGATGATCCGTTAACAAACGAGGAGATTGATCTGGCCTAT | 4 |
| wt-84150-84270 | 2126 | GCCCGCGGGCAGGCCATGAATATTGAGGCTCCTAGACTGCCAGATGATCCTATAATTGTTGAGGATGACGACGAAAGTGAGGAAATTGAAGCTGAAAGCGACGAGGAGGAAGACAAGAGT | 4 |
| wt-84270-84390 | 2127 | GGAATGGAATCTCTTAAAAATATACCGCAAACACTGCCGTACAATCCAACAGTATACGGCAGGCCCGCGGTGTTTGACCGAAAGTCAGATGCAAAATCAACCAAAAAATGCAGGGCCATA | 4 |
| wt-84390-84510 | 2128 | GTAACTGACTTTAGTGTAATCAAGGCCATTGAAGAGGAACACAGAAAGAAGAAGGCAGCCAGAACAGAGCAGCCAAGAGCCACGCCTGAATCCCAGGCCCCCACAGTGGTCCTCCAGCGA | 4 |
| wt-84510-84630 | 2129 | CCACCCACGCAGCAAGAGCCTGGCCCCGTCGGCCCACTGAGTGTCCAGGCTCGGCTGGAGCCATGGCAACCTTTGCCTGGGCCCCAAGTGACAGCAGTTCTACTTCACGAAGAATCCATG | 4 |
| wt-84630-84750 | 2130 | CAGGGTGTCCAAGTACATGGTTCGATGCTAGACCTTCTTGAAAAAGACGATGAAGTCATGGAGCAGAGGGTTATGGCAACCCTACTGCCACCAGTACCACAACAGCCCCGGGCTGGCAGA | 4 |
| wt-84750-84870 | 2131 | AGAGGCCCTTGTGTCTTCACCGGTGACCTAGGCATAGAGAGTGATGAGCCCGCTTCGACAGAGCCGGTTCATGATCAGCTACTGCCTGCCCCAGGACCTGACCCTCTTGAAATCCAACCA | 4 |
| wt-84870-84990 | 2132 | CTAACGTCCCCCACCACGTCTCAACTTAGCAGTTCGGCACCAAGCTGCGCACAAACTCCATGGCCGGTGGTTCAGCCAAGTCAGACTCCAGATGACCCAACGAAACAGTCCCGGCCACCG | 4 |
| wt-84990-85110 | 2133 | GAAACAGCTGCCCCACGCCAGTGGCCAATGCCCCTGCGACCTATCCCTATGCGCCCCTTGCGGATGCAGCCAATCCCATTTAATCATCCAGTGGGACCCACTCCCCATCAGACACCTCAA | 4 |
| wt-85110-85230 | 2134 | GTGGAAATAACACCATATAAGCCCACTTGGGCTCAGATAGGGCACATTCCATATCAGCCTACACCAACGGGTCCTGCTACCATGCTGTTGCGCCAGTGGGCACCCGCCACCATGCAGACA | 4 |
| wt-85230-85350 | 2135 | CCACCGAGAGCGCCCACTCCCATGTCACCACCTGAGGTGCCACCCGTTCCCCGGCAGAGGCCTCGGGGGGCGCCCACTCCCACGCCACCTCCTCAGGTGCCGCCCGTTCCCCGGCAGAGG | 8 |
| wt-85350-85470 | 2136 | CCTCGGGGGGCGCCCACTCCCACGCCACCTCCTCAGGTGCTGCCCACTCCCATGCAGCTGGCACTAAGGGCTCCTGCTGGTCAGCAGGGGCCGACAAAGCAAATTTTGCGCCAATTGTTA | 4 |
| wt-85470-85590 | 2137 | ACGGGGGGCGTCAAGAAAGGGAGACCATCACTTAAGTTACAGGCCGCCCTTGAGCGTCAAGCCGCTGCGGGCTGGCAGCCTTCACCAGGGTCCGGCACCAGTGACAAGATTGTGCAGGCG | 4 |
| wt-85590-85710 | 2138 | CCTATTTTCTACCCACCCGTTTTGCAGCCCATACAGGTTATGGGGCAAGGGGGTTCCCCAACGGCCATGGCCGCCTCAGCGGTGACACAGGCACCCACGGAATATACCAGGGAAAGGAGG | 4 |
| wt-85710-85830 | 2139 | GGAGTGGGGCCTATGCCTCCCACCGATATTCCGCCGTCTAAACGAGCGAAGATCGAGGCCTATACAGAGCCCGAGATGCCGCACGGGGGGGCCTCGCACTCTCCCGTCGTTATCTTGGAG | 4 |

| | | | | |
|---|---|---|---|---|
| wt-85830-85950 | 2140 | AATGTCGGCCAGGGGCAACAGCAGACTCTGGAGTGCGGAGGAACTGCTAAACAGGAAAGGGACATGTTGGGGCTGGGGGACATTGCAGTTTCTTCCCCTTCCTCTTCTGAAACATCGAAC | | 4 |
| wt-85950-86070 | 2141 | GATGAGTGATTTCGCCCATGTAACAAGAACTGGGATGAACCCTGGGGCAACAGACTGCGGGGAGGAGGGGGGCAGTGATAAGTCATGACAATTTTAGATGAGGTAGAAATTTTGCATATT | | 4 |
| wt-86070-86190 | 2142 | TTCAGACCCACCATGGAATCATTTGAAGGACAGGGGGACTCTAGACAGTCACCCGACAATGAGCGGGGAGATAATGTACAGACTACCGGCGAGCATGATCAGGACCCTGGGCCGGGGCCT | | 4 |
| wt-86190-86310 | 2143 | CCATCCAGTGGGGCTTCTGAGAGATTGGTACCAGAAGAGTCATACTCAAGAGATCAGCAACCTTGGGGGCAAAGCAGGGGTGATGAAAACAGAGGCTGGATGCAGCGCATCAGGCGAAGG | | 4 |
| wt-86310-86430 | 2144 | CGGAGAAGACGGGCTGCCTTGTCCGGCCATCTTTTAGACACGGAAGACAATGTGCCGCCATGGTTGCCTCCACACGACATCACACCATATACCGCAAGGAATATCAGGGATGCTGCCTGC | | 4 |
| wt-86430-86550 | 2145 | CGGGCTGTCAAGGTGAGTATGCCTCTAACTGGGTTCATGGGGGCCATCTAAGGCCCACGTGTGACCCATGTTTCCATTAATTTTAGCAATCGCACCTGCAAGCGCTATCAAACCTGATAC | | 4 |
| wt-86550-86670 | 2146 | TCGATAGTGGGTTAGACACACAACACATCTTGTGCTTCGTGATGGCAGCCAGGCAGCGTCTTCAGGACATTCGACGTGGACCCTTGGTTGCGGAGGGCGGTGTGGGTTGGCGACATTGGC | | 4 |
| wt-86670-86790 | 2147 | TTCTAACATCTCCCAGCCAATCCTGGCCCATGGGATATCGTACAGCAACACTACGCACATTAACTCCCGTGCCTAACAGGGTTGGGGCTGACAGCATCATGTTAACTGCCACATTTGGAT | | 4 |
| wt-86790-86910 | 2148 | GCCAAAATGCGGCACGAACTCTAAACACCTTCTCTGCCACCGTGTGGACACCACCCCATGCTGGACCAAGAGAGCAAGAAAGATACGCTCGGGAAGCCGAGGTACGCTTCCTTCGTGGTA | | 4 |
| wt-86910-87030 | 2149 | AATGGCAGAGGCGGTACCGAAGAATCTATGATTTGATAGAACTGTGTGGCTCTCTGCACCACATCTGGCAAAACTTGCTCCAGACCGAGGAGAACCTTTTAGATTTCGTGCGTTTCATGG | | 4 |
| wt-87030-87150 | 2150 | GTGTCATGTCCAGCTGCAATAATCCAGCTGTGAATTACTGGTTTCACAAGACAATCGGAAACTTTAAGCCATATTACCCGTGGAATGCACCACCTAATGAAAATCCATATCACGCGCGGA | | 4 |
| wt-87150-87270 | 2151 | GAGGCATAAAAGAACACGTAATCCAGAACGCATTTCGAAAGGCACAAATACAGGGTTTATCAATGTTAGCAACGGGAGGTGAACCCAGAGGTGATGCTACTAGTGAAACGAGCAGTGATG | | 4 |
| wt-87270-87390 | 2152 | AGGACACCGGTAGACAGGGTTCGGACGTGGAGCTAGAGTCCTCGGACGATGAGCTGCCATATATCGATCCCAATATGGAGCCGGTTCAGCAGAGGCCCGTCATGTTTGTGAGCCGTGTGC | | 4 |
| wt-87390-87510 | 2153 | CTGCAAAGAAACCGAGGAAACTGCCTTGGCCCACGCCCAAGACGCACCCAGTGAAGCGCACAAATGTTAAGACCTCTGATAGATCTGATAAGGCAGAAGCACAAAGCACCCCTGAAAGGC | | 4 |
| wt-87510-87630 | 2154 | CGGGCCCTTCCGAACAATCATCAGTGACCGTGGAGCCCGCCCACCCGACCCCGGTGGAGATGCCAATGGTGATTCTCCATCAACCACCTCCAGTGCCCAAACCGGTTCCAGTCAAGCCTA | | 4 |
| wt-87630-87750 | 2155 | CGCCACCGCCTTCCCGTAGGAGAAGGGGAGCGTGTGTTGTGTACGACGATGATGTCATAGAGGTGATTGATGTTGAAACCACCGAAGATTCATCGTCAGTGTCACAGCCAAATAAGCCAC | | 4 |
| wt-87750-87870 | 2156 | ATCGGAAACATCAAGACGGCTTTCAACGTTCAGGCCGACGTCAAAAACGAGCCGCGCCTCCCACCGTGAGTCCTTCGGATACTGGGCCTCCTGCCGTGGGGCCTCCTGCCGCGGGGCCTC | | 4 |
| wt-87870-87990 | 2157 | CTGCCGCGGGGCCTCCTGCCGCGGGGCCTCCTGCCGCGGGGCCTCCTGCCGCGGGGCCTCCTGCCGCGGGGCCTCGCATACTGGCGCCTCTTTCCGCTGGGCCTCCTGCCGCGGGGCCTC | | 12 |
| wt-87990-88110 | 2158 | ACATAGTGACGCCTCCTTCCGCCCGGCCTCGTATAATGGCGCCTCCCGTCGTACGTATGTTTATGAGGGAGCGACAGCTCCCCCAGTCCACCGGCCGTAAACCTCAGTGCTTCTGGGAAA | | 4 |
| wt-88110-88230 | 2159 | TGCGGGCTGGTCGTGAAATTACACAAATGCAACAAGAACCAAGTTCACACCTGCAGTCCGCCACTCAGCCTACAACGCCTCGCCCATCATGGGCCCCATCAGTCTGCGCCCTCTCGGTGA | | 4 |
| wt-88230-88350 | 2160 | TGGATGCTGGTAAGGCCCAGCCCATAGAAAGTTCACACTTGAGTTCCATGTCGCCCACACAGCCGATATCGCACGAAGAACAACCCCGGTATGAGGATCCTGACGCTCCTCTGGATTTAA | | 4 |
| wt-88350-88470 | 2161 | GTTTACATCCAGACGTTGCTGCTCAACCAGCTCCCCAGGCTCCATACCAGGGATACCAGGAGCCGCCGGCCCCCCAGGCTCCATACCAGGGATACCAGGAGCCGCCGCCCCCCCAGGCTC | | 8 |
| wt-88470-88590 | 2162 | CATACCAGGGATACCAGGAGCCGCCGGCCCACGGGCTCCAATCATCTTCATATCCAGGATATGCGGGTCCCTGGACCCCAAGGTCTCAACATCCATGTTATAGGCACCCCTGGGCACCAT | | 4 |
| wt-88590-88710 | 2163 | GGTCTCAAGATCCTGTGCATGGGCACACCCAGGGTCCATGGGATCCCAGGGCACCACATCTCCCACCTCAGTGGGATGGATCTGCAGGACATGGCCAGGATCAGGTCTCCCAGTTCCCAC | | 4 |
| wt-88710-88830 | 2164 | ATCTGCAATCGGAGACAGGCCCACCACGTCTTCAACTTTCATTGGTGCCACTGGTCTCATCCTCTGCACCATCATGGTCATCTCCCCAGCCCCGAGCCCCCATACGCCCCATTCCAACAA | | 4 |
| wt-88830-88950 | 2165 | GATTCCCCCCTCCCCCTATGCCGTTACAAGATAGCATGGCCGTGGGGTGTGACTCATCAGGTACAGCATGCCCAAGCATGCCCTTTGCCAGTGATTACAGTCAAGGTGCATTTACCCCAC | | 4 |
| wt-88950-89070 | 2166 | TGGACATTAATGCCACCACGCCAAAAAGGCCTCGAGTAGAAGAAAGTTCTCACGGACCTGCCCGGTGTTCCCAAGCTACTGCTGAAGCACAGGAGATTCTCAGTGACAATTCTGAGATCT | | 4 |
| wt-89070-89190 | 2167 | CCGTGTTCCCAAAGGACGCGAAGCAGACTGACTACGATGCATCCACTGAAAGTGAGCTAGATTAAGGGGATCCAAGGTGACCCCTGTTAGCTATTTGATCTTTGACTGACACATAAACAT | | 4 |
| wt-89190-89310 | 2168 | GGTTTAAGGAATGAACACTCATGGTGTGAGACTGGAACTGTACTAAATTTGCTGACATATGTACAATGAGAGCCAAAAATTTGATAAACCTTAAAAGTCCCCCCATCTAATGATGTCCAG | | 4 |
| wt-89310-89430 | 2169 | TTCCCTTCTCCCACCCTGTACACCCCGACCCAAAGGGACTCAATGGCATTCAGATTTCTAGTTACCACAGGTAGAATATCGGGCGTTGGCCCATAAAAATAAGTGCATGGATATAGCTCT | | 4 |
| wt-89430-89550 | 2170 | GCACAGGCTTGGAAACACCCATTCCAGGTGTGCTTCTTTTTGGTGAAATAAAAACAGCATCCTTTATATGAAAATGTGTATTCTCTGGTGTTGCAGTATGTACAGTTAGCTTTGGTATAG | | 4 |
| wt-89550-89670 | 2171 | TTTTGGGGTACCTGAAATGTGTGCAGGGTGGGTGTCCAATGTGGCAGTTTTACCTCTTTGTCCCCATACTCCTGCTCGGCCGTCTTGTTAAAGTTAACCGGCGGTGGAGGATCCACCGGC | | 4 |
| wt-89670-89790 | 2172 | CAGACCTCTACATTTGGTTTGGGTACCCAGGTGATGGCCGCGGCTGCCACCCGCCCTCCTCCTCTTACCCTGGGTGGCAAAAAGTATGCCAGGAGTAGAACAATAACAAGTGCGATGGCG | | 4 |
| wt-89790-89910 | 2173 | GTAAACAATGGCACCCTCACCTGCTTAAATGAAACCATGGCAACCACTTCAAAGAGAGCCGACAGGAAGATATTTATTAATATTCCATTAGTAAACGAGGCGTGAAGCAGGCGTGGTTTC | | 4 |
| wt-89910-90030 | 2174 | AATAACGGGAGTTAGAAATTTAAGAGATCCTCGTGTAAAACATCTGGTGTCCGGGGGATAATGGAGTCAACATCCAGGCTTGGGCACATCTGCTTCAACAGGAGGCGCAGCCTGTCATTT | | 4 |
| wt-90030-90150 | 2175 | TCAGATGATTTGGCAGCAGCCACCTGCGGACAAAAATCAGGCGTTTAGATGGGGCATTTATGTTTGGGACGCTAGCCGCCTGGGCATTCGTGTTAGTATATACTGACCTCACGGTAGTGC | | 4 |
| wt-90150-90270 | 2176 | TGCAGCAGTTGCTTAAACTTGGCCCGGCATTTTCTGGAAGCCACCCGATTCTTGTATCGCTTTATTTCTAGTTCAGAATCGCATTCCTCCAGCTGCGAGCAAGGGAATGCGTTACTACAA | | 4 |
| wt-90270-90390 | 2177 | GTGGTGCCTAGTCAGTTGAAACAAGCCCCACCATCCGCTGCCGCCCCTCCATGAGCCCCACCGTCCGCTGCCGCCCCTCCTTGAGCCCCTCCTTACCGATTCTGGCTGTTGTGGTTTCCG | | 4 |
| wt-90390-90510 | 2178 | TGTGCGTCGTGCCGGGGCAGCCACTGGTGCAGGCTGTGGAACACCAATGTCTGCTAGCTGTTGTCCTTGGTTAGCCCCGGGGCAAGCAAACACCACTGCTGCTGCTGTTTGAACAGTAGA | | 4 |
| wt-90510-90630 | 2179 | ATTGTCTCCAGGTTGAGGTGCTTCTCCCCCGGCTTGGTTAGTCTGTTGATTCTGGGTTATGTCGGAGACTGGGAACAGCTGAGGTGCTGCATAAGCTTGATAAGCATTCTCAGGAGCAGG | | 4 |
| wt-90630-90750 | 2180 | CTGAGGGGCAGAAAACCACGACCCAGTCGGAGCGGTTGAAACATGATAGGCAGTTAGCTGGCCTTGTGGCAGAGGCTCTGGCAGCACCGGCCACAGCACACAAGGCAAAGGAGCTTGCGA | | 4 |
| wt-90750-90870 | 2181 | TGGCCCTCCCAGGTCCTGATAGACTCTGGTAGCTTGGTCAAAAGCTTGTACAAAAGGCACCTGGTATGGGTCAGGTGTAAATTTTACATCTTCAGAAGTCGAGTTTGGGTCCATCATCTT | | 4 |
| wt-90870-90990 | 2182 | CAGCAAAGATAGCAAAGGTGGCCGGCAAGGTGCAATGTTTAGTGAGTTACCTGTCTAACATCTCCCCTTTAAAGCCAAGGCACCAGCCTCCTCTGTGATGTCATGGTTTGGGACGTGCTA | | 4 |
| wt-90990-91110 | 2183 | AATTTAGGTGTGTCTATGAGGTACATTAGCAATGCCTGTGGCTCATGCATAGTTTCTAAAAGAGGAGGAGGCAGTTTTCAGAAGTGTCTAAAATAAGCTGGTGTCAAAAATAGACAGCCC | | 4 |
| wt-91110-91230 | 2184 | AGTTGAAATATGCATGGCATGCAGCAGACATTCATCATTTAGAAATGTATCCAAGATTTCATTAAGTTCGGGGGTCAGGGGGGAGTCCAGATTCAAATCCTCTGTCATGGACTCTAGTGT | | 4 |
| wt-91230-91350 | 2185 | TGTGGTCAGTTCGTCCAAATGGCCACGAGGGGGCGGGTGGCTCAGGTCCATCTGTCCACATATGGCTGCTTCCTCCTTCTGGGGAATAACAGTGTCAGCCATCTCCCTTAGGGCCTTCAC | | 4 |
| wt-91350-91470 | 2186 | GGCCTGACTGGTTTCTTCATCAGGGTCCTCCAACAGATGACTTGCCTCGGGGGTTACTGCGGGGGCCGGGTCAAGTGGCTGGGGCACCGGGGCTGGCGTTAGGGATCCGACCGGTTCATG | | 4 |
| wt-91470-91590 | 2187 | GACAGGTCCTGTGGGGGTGGGAGCCAAAGAGGCAGGCAGGGGCCGGTTGGCCCACGGGGATCCGGGTGGATGGAAGGGCCTGATCCTCTTTGGCTGACACACCTCTCGCCCCTCGAACAC | | 8 |
| wt-91590-91710 | 2188 | GTCAGATATGGCACTGCCCGCTTCCGGCTTTGGCAGGAACATACCTTCCCGGCTATCCCTGAGGCCCTTCTTCCTTTTAACGGGAGGAAGAAAGGTGGGCTTTGAGGGGTGGGGGAATAT | | 4 |
| wt-91710-91830 | 2189 | GGGTCTCTCATCGCTCTCTTGGTGGACCGCTGCTATCCAAGGCTGTTCAGGTTCCGCCGCGTTGGAAGGACATGGAGTTTGACCACGGTTGGGCCTGGATGTCCGGCGCGACTTTGGGGC | | 4 |
| wt-91830-91950 | 2190 | CCGCAGGCGCGGGGCCTCGGCCCTGGCCTCTTCCCGCTCGCTCTGCTCGGTGTCACTGTTGCCCGAGTCACTGCTGCTGGAACTGCTGTCACCGCAGTCGGCGCTTTGGGCACCGGGCTT | | 8 |

| | | | |
|---|---|---|---|
| wt-91950-92070 | 2191 | CAGGGGCATGGTCGGGCTCGGGAGACTTTCGAGTTCATCTGTAAAAGCATGAAACTGTCCGGACTCCGAGTAGCGGGCCTCGGTGTGAGAGGCACCCCCATCATTCCCCATGAGCTCCTC | 4 |
| wt-92070-92190 | 2192 | GTCCATCCTGTCGGCTCCGGACACGAGGATAGGAGTTTCCACTGCCTTGGACTTGGTTGACAGCAGGCACGCGGGAAGCACGCCGCTCACGTAGCTCCTCTGTCCGGCGTGGCTGGAGTA | 4 |
| wt-92190-92310 | 2193 | GGAGGCCCGGGGCAGTGTCTTAATCAGAGCCCTGACATCCTTAACATCGTCCGTCAGATGGCCTGTCTTGGACGAGACCATAGTCTGGAACATCTCCTCGAGGACGGGATAGGTGAACAC | 4 |
| wt-92310-92430 | 2194 | CCACTTGCAAAAGGCCTTGAACTTGGAGCTTAGGAGGCCTTCCTTCTCCATCCTGTTCAGGTGTTCCACTACCTGCTTGCCGGAGGCCATGATGGCCGCGCGGTCCACGCCCAGCACCTT | 4 |
| wt-92430-92550 | 2195 | GCTGTAGGTGTAGGCCCGCACCCGACTGTGTTTTAGGAGCTTGTACATAGCGGTGCCTATGGTGGCAGGAATCATCACCCGGTTGCTGGGGGCCTGGATGAAGAATCTGTCAGTGACCAC | 4 |
| wt-92550-92670 | 2196 | TATCAGGTGGTCTAACACGTAGCGCATCACTATAGGGCACGCGATGGAACATGCGTCGTTGCCGGCATTCTCAGCCCGTCTTCTTACCCTGTTGTTTCGGAGAATGGCCCAAAAATTGCA | 4 |
| wt-92670-92790 | 2197 | GATGTTGAGCGTGGCCATTAGCCCGCCCCATTCTCGCCCGTGGGCCTTGGCCTCATTTATAAATGCCTTGCATATTTTGTAGGATCTCAGAGTAATCTCCACACTCCCGGCTGTAAATTC | 4 |
| wt-92790-92910 | 2198 | CTTGTTGAGGACGTTGCAGTAGTCAGAGACCAGAGAGCCCAGCTGCTTTTTGATTTCAGGAGTTAGCCTCAGAAAGTCTTCCAAGCCATCCTTTTTAGGCCTCATGGCTAGTAGTAACAG | 4 |
| wt-92910-93030 | 2199 | AGGAAATGCCCGACCATTAAAATCTTTCCTCCATGAGCTTTACCTGAAACACTATCCCGAAGTGGGGGATGTGGTGCATCTACTGAACACCATCGGGGTCGACTGCGACCTCCCACCTAG | 4 |
| wt-93030-93150 | 2200 | CCACCCACTCCTGACAGCCCAGAGGGGGCTGTTCCTGGCAAGAGTCTTGCAGGCTGTACAGCAGCACAAGCTGCTGGAAGACACCATCGTCCCCAAGATCTTAAAGAAGCTGGCTTATTT | 4 |
| wt-93150-93270 | 2201 | CTTAGAGCTGCTAAGCTACTACTCCCCCAAGGATGAACAGCGTGACATCGCCGAGGTTCTTGACCACCTCAAGACGAATCGGGACCTGGGGCTGGACGACAGACTCTGGGCCCTGATTAG | 4 |
| wt-93270-93390 | 2202 | GAAACTGCGCCAAGACAGACACCATGCCTCTGTAAATGTCCTCATGCCAGGAAGCGACTACACAGCCGTGTCGCTGCAGTACTACGACGGCATCTCCATAGGTATGAGGAAGGTAATCGC | 4 |
| wt-93390-93510 | 2203 | GGATGTCTGCCGCAGTGGCTATGCCTCCATGCCCTCCATGACGGCCACGCACAACCTCTCCCACCAGCTCTTGATGGCGTCCGGGCCCAGTGAGGAACCGTGCGCCTGGCGCGGGTTCTT | 8 |
| wt-93510-93630 | 2204 | TAACCAGGTCCTCCTCTGGACTGTGGCCCTCTGCAAGTTTCGCAGATGCATTTACTATAACTACATTCAGGGATCTATAGCCACCATCTCCCAGCTTCTGCACCTCGAGATCAAGGCCCT | 4 |
| wt-93630-93750 | 2205 | CTGCAGCTGGATAATATCCCAGGATGGCATGCGCCTCTTTCAACACAGCAGGCCTCTCCTCACCCTCTGGGAGAGCGTGGCCGCAAATCAGGAGGTCACGGATGCCATTACCCTGCCTGA | 4 |
| wt-93750-93870 | 2206 | CTGCGCTGAATACATAGACCTACTAAAGCACACAAAACATGTCTTAGAAAACTGTTCTGCCATGCAATACAAATAAATTTCTCTTACCTGCGTCTGTTTGTGTAGTGAGGTGTTGTGTCC | 4 |
| wt-93870-93990 | 2207 | TGTATGGTATTCTACTTTAAAAAGGCCGGCTGACATGGATTACTGGTCTTTTATGAGCCATTGGCATGGGCGGGACAATCGCAATATAAAACCCTGACCATCACATGGGGCATTAGGCGA | 4 |
| wt-93990-94110 | 2208 | CTCTGCATCAGCATCGCTTAAGTATGAGTGGGCAGCAGAGAGGCTCGGTTATTTTGGTTCCTGAACATCTGGCTGGGGCATTAACTAAGCTTATGAGCGATTTTATCACAGGACAAGATG | 4 |
| wt-94110-94230 | 2209 | TCACTCTTTCTGGAGGAAATATTGCAGTCAAAATTCGCGATGCTATAAACCAGACCCCCGGGGGTGGTGATGTAGCTATACTTTCTTCCCTGTTTGCTTTATGGAATGCCCTCCCAACAT | 4 |
| wt-94230-94350 | 2210 | CTGGTAGACAATCCTCCAGGGACGATTTAATCCCAGCCGCCGTGCAGGCCTTAACCACGGCCCACAACTTATGTCTGGGTGTTATTCCAGGTGAGACCTCACACAAGGACACACCCGAGT | 4 |
| wt-94350-94470 | 2211 | CATTGCTCCGGGCTATCGTGACGGGTCTCCAAAAATTGTGGGTGGATTCGTGCGGATGTCCAGAGTGCCTACAATGTCTTAAGGGATTGAAGGCAATTAAGCCCGGCCTTTATGAAATCC | 4 |
| wt-94470-94590 | 2212 | CTAGGATAATACCACACACTAAGCAGTGTAGTCCTGTCAATCTCCTGAACATGTTGGTCCACAAGCTTGTGGCTTTACGTGGTCATGTGCAGCTTGCATACGACGCCCGTGTCCTGACGC | 4 |
| wt-94590-94710 | 2213 | CTGACTTTCACGAAATCCCTGACCTCGATGACTCCGATGCTGTTTTCGCACGCACCTTATTGGCAGCCTTATTTCACCTCAATATGTTCTTTATTCTCAAAGATTACATAACACAAGACT | 4 |
| wt-94710-94830 | 2214 | CCATGAGCTTGAAGCAGGCCCTCAGTGGTCATTGGATGTCTGCCACGGGCAACCCCCTGCCTGCAGCACCGGAAACCCTGCGAGACTACTTGGAAGCTTTCCGAAATTCGGATAATCACT | 4 |
| wt-94830-94950 | 2215 | TTTATCTCCCGACGACAGGGCCTTTAAACACCTTCCAATTTCCCGAAGAGCTTCTGGGGCGCGTTGTTGTTATTGATTCCTCTTTGTGTGCCGCCAGTCACGTTCAGGACGTTATCACCC | 4 |
| wt-94950-95070 | 2216 | ATGGTGTTGGGGCGGGTGTTCCTCGTCCTCGGTTTTCGGCCCTGCCTCCGGCCCCATCCCGCGAGCCCCAGCAGACATGCTCTCAGTTAACGAGCAGAGGGAATGAAAGCTCACGGCGAA | 4 |
| wt-95070-95190 | 2217 | ACTTGGGCCAGCCCGGGGGGACCTCCCCTGCTGTTCCCCCAGTTTGCCCCATCGTTTCCCTGACGGCCTCAGGGGCCAAGCAAAACCGCGGGGGCATGGGATCCTTGCACTTAGCCAAGC | 8 |
| wt-95190-95310 | 2218 | CTGAGGAAACCTCCCCCGCCGTCTCCCCAGTATGCCCCATCGCTTCCCCAGCGGCCTCCAGGTCCAAGCAGCACTGCGGGGTCACTGGATCCTCACAGGCCGCACCCAGCTTTTCTTCCG | 8 |
| wt-95310-95430 | 2219 | TTGCCCCAGTAGCATCTCTGTCTGGTGACCTTGAAGAGGAAGAGGAGGGGTCCCGAGAATCCCCATCCCTACCGTCCAGCAAAAAGGGGGACGAGGAATTTGAGGCCTGGCTTGAGGCTC | 4 |
| wt-95430-95550 | 2220 | AGGACGCAAATCTTGAGGATGTTCAGCGGGAGTTTTCCGGGCTGCGAGTAATTGGTGATGAGGACGAGGATGGTTCGGAGGATGGGGAATTTTCAGACCTGGATCTGTCTGACAGCGACC | 4 |
| wt-95550-95670 | 2221 | ATGAAGGGGATGAGGGTGGGGGGGCTGTTGGAGGGGGCAGGAGTCTGCACTCCCTGTATTCACTGAGCGTCGTCTAATAAAGATGTCTATTGATCTCTTTTAGTGTGAATCATGTCTGAC | 4 |
| wt-95670-95790 | 2222 | GAGGGGCCAGGTACAGGACCTGGAAATGGCCTAGGAGAGAAGGGAGACACATCTGGACCAGAAGGCTCCGGCGGCAGTGGACCTCAAAGAAGAGGGGGTGATAACCATGGACGAGGACGG | 4 |
| wt-95790-95910 | 2223 | GGAAGAGGACGAGGACGAGGAGGCGGAAGACCAGGAGCCCCGGGCGGCTCAGGATCAGGGCCAAGACATAGAGATGGTGTCCGGAGACCCCAAAAACGTCCAAGTTGCATTGGCTGCAAA | 4 |
| wt-95910-96030 | 2224 | GGGACCCACGGTGGAACAGGAGCAGGAGCAGGAGCGGGAGGGGCAGGAGCAGGAGGGGCAGGAGCAGGAGGAGGGGCAGGAGCAGGAGGAGGGGCAGGAGGGGCAGGAGGGGCAGGAGGG | 8 |
| wt-96030-96150 | 2225 | GCAGGAGCAGGAGGAGGGGCAGGAGCAGGAGGAGGGGCAGGAGGGGCAGGAGGGGCAGGAGCAGGAGGAGGGGCAGGAGCAGGAGGAGGGGCAGGAGGGGCAGGAGCAGGAGGAGGGGCA | 8 |
| wt-96150-96270 | 2226 | GGAGGGGCAGGAGGGGCAGGAGCAGGAGGAGGGGCAGGAGCAGGAGGAGGGGCAGGAGGGGCAGGAGCAGGAGGAGGGGCAGGAGGGGCAGGAGGGGCAGGAGCAGGAGGAGGGGCAGGA | 8 |
| wt-96270-96390 | 2227 | GCAGGAGGGGCAGGAGGGGCAGGAGGGGCAGGAGCAGGAGGGGCAGGAGCAGGAGGAGGGGCAGGAGGGGCAGGAGGGGCAGGAGCAGGAGGGGCAGGAGCAGGAGGGGCAGGAGCAGGA | 8 |
| wt-96510-96630 | 2228 | GCAGGAGGGGCAGGAGCAGGAGGGGCAGGAGGGGCAGGAGCAGGAGGGGCAGGAGGGGCAGGAGCAGGAGGGGCAGGAGGGGCAGGAGCAGGAGGAGGGGCAGGAGCAGGAGGGGCAGGA | 8 |
| wt-96630-96750 | 2229 | GCAGGAGGTGGAGGCCGGGGTCGAGGAGGCAGTGGAGGCCGGGGTCGAGGAGGTAGTGGAGGCCGGGGTCGAGGAGGTAGTGGAGGCCGCCGGGGTAGAGGACGTGAAAGAGCCAGGGGG | 8 |
| wt-96750-96870 | 2230 | GGAAGTCGTGAAAGAGCCAGGGGGAGAGGTCGTGGACGTGGAGAAAAGAGGCCCAGGAGTCCCAGTAGTCAGTCATCATCATCCGGGTCTCCACCGCGCAGGCCCCCTCCAGGTAGAAGG | 4 |
| wt-96870-96990 | 2231 | CCATTTTTCCACCCTGTAGGGGAAGCCGATTATTTTGAATACCACCAAGAAGGTGGCCCAGATGGTGAGCCTGACGTGCCCCCGGGAGCGATAGAGCAGGGCCCCGCAGATGACCCAGGA | 4 |
| wt-96990-97110 | 2232 | GAAGGCCCAAGCACTGGACCCCGGGGTCAGGGTGATGGAGGCAGGCGCAAAAAAGGAGGGTGGTTTGGAAAGCATCGTGGTCAAGGAGGTTCCAACCCGAAATTTGAGAACATTGCAGAA | 4 |
| wt-97110-97230 | 2233 | GGTTTAAGAGCTCTCCTGGCTAGGAGTCACGTAGAAAGGACTACCGACGAAGGAACTTGGGTCGCCGGTGTGTTCGTATATGGAGGTAGTAAGACCTCCCTTTACAACCTAAGGCGAGGA | 4 |
| wt-97230-97350 | 2234 | ACTGCCCTTGCTATTCCACAATGTCGTCTTACACCATTGAGTCGTCTCCCCTTTGGAATGGCCCCTGGACCCGGCCCACAACCTGGCCCGCTAAGGGAGTCCATTGTCTGTTATTTCATG | 4 |
| wt-97350-97470 | 2235 | GTCTTTTTACAAACTCATATATTTGCTGAGGTTTTGAAGGATGCGATTAAGGACCTTGTTATGACAAAGCCCGCTCCTACCTGCAATATCAGGGTGACTGTGTGCAGCTTTGACGATGGA | 4 |
| wt-97470-97590 | 2236 | GTAGATTTGCCTCCCTGGTTTCCACCTATGGTGGAAGGGGCTGCCGCGGAGGGTGATGACGGAGATGACGGAGATGAAGGAGGTGATGGAGATGAGGGTGAGGAAGGGCAGGAGTGATGT | 4 |
| wt-97590-97710 | 2237 | AACTTGTTAGGAGACGCCCTCAATCGTATTAAAAGCCGTGTATTCCCCCGCACTAAAGAATAAATCCCCAGTAGACATCATGCGTGCTGTTGGTGTATTTCTGGCCATCTGTCTTGTCAC | 4 |
| wt-97710-97830 | 2238 | CATTTTCGTCCTCCCAACATGGGGCAATTGGGCATACCCATGTTGTCACGTCACTCAGCTCCGCGCTCAACACCTTCTCGCGTTGGAAAACATTAGCGACATTTACCTGGTGAGCAATCA | 4 |
| wt-97830-97950 | 2239 | GACATGCGACGGCTTTAGCCTGGCCTCCTTAAATTCACCTAAGAATGGGAGCAACCAGCTGGTCATCAGCCGCTGCGCAAACGGACTCAACGTGGTCTCCTTCTTTATCTCCATCCTGAA | 4 |
| wt-97950-98070 | 2240 | GCGAAGCAGCTCCGCCCTCACGGGCCATCTCCGTGAGTTGTTAACCACCCTGGAGACTCTTTACGGTTCATTCTCAGTGGAAGACCTGTTTGGTGCCAACTTAAACAGATACGCATGGCA | 4 |
| wt-98070-98190 | 2241 | TCGCGGGGGCTAGACCTCTGGCTGGATGAGCACGTGTGGAAGAGGAAACAGGAGATTGGTGTGAAAGGAGAAAATCTGCTTCTCCCCGACTTATGGCTAGATTTCCTACAACTCAGCCCC | 4 |

| | | | |
|---|---|---|---|
| wt-98190-98310 | 2242 | ATCTTCCAGCGCAAGCTTGCTGCCGTTATTGCCTGTGTCCGACGCCTGCGGACTCAGGCCACCGTCTACCCAGAGGAGGACATGTGCATGGCCTGGGCCCGCTTTTGCGACCCCTCTGAT | 4 |
| wt-98310-98430 | 2243 | ATTAAGGTGGTTATTTTGGGCCAGGACCCCTATCACGGGGGTCAAGCAAACGGCCTGGCATTCAGCGTCGCATACGGCTTTCCAGTTCCCCCCAGCCTGAGGAACATCTACGCGGAGCTG | 4 |
| wt-98430-98550 | 2244 | CACCGGAGCCTGCCGGAGTTTTCTCCCCCAGATCACGGCTGTCTAGACGCGTGGGCCTCCCAGGGGGTGTTGCTACTCAACACCATCCTGACCGTGCAAAAGGGCAAGCCCGGCTCGCAC | 4 |
| wt-98550-98670 | 2245 | GCAGACATTGGCTGGGCGTGGTTTACTGACCACGTAATTTCATTGCTCTCTGAGCGGTTAAAAGCGTGCGTGTTTATGCTGTGGGGTGCGAAGGCGGGAGACAAAGCTTCACTAATCAAC | 4 |
| wt-98670-98790 | 2246 | TCCAAGAAGCATCTGGTTCTGACCTCTCAGCATCCCTCTCCCCTGGCCCAGAACAGCACCCGAAAGAGTGCCCAGCAGAAGTTCCTGGGCAACAACCACTTTGTCCTCGCTAACAACTTT | 4 |
| wt-98790-98910 | 2247 | TTGCGTGAGAAGGGGCTCGGTGAGATAGATTGGAGGCTGTAGAGGGGTCATCACTATGGCCATGTTTCTGAAGTCGCGTGGGGTCCGGTCTTGCAGGGACCGGCGCCTCTTGTCGGACGA | 4 |
| wt-98910-99030 | 2248 | GGAGGAAGAGACTTCACAGAGCAGCAGCTACACTCTGGGGTCTCAGGCCTCCCAGTCTATCCAGGAGGAGGACGTGAGTGACACTGATGAGTCTGACTACTCAGATGAAGACGAGGAGAT | 4 |
| wt-99030-99150 | 2249 | TGATTTGGAGGAAGAGTACCCCAGTGACGAAGACCCATCTGAGGGCAGTGATAGCGACCCCTCGTGGCATCCTTCAGATTCAGACGAGTCTGACTACAGCGAGAGCGACGAGGATGAAGC | 4 |
| wt-99150-99270 | 2250 | AACCCCCGGCTCTCAGGCCTCACGATCTTCAAGAGTCTCGCCATCTACCCAACAGTCTTCAGGTCTGACACCCACGCCTTCGTTCTCCCGACCACGCACCCGGGCACCTCCGAGGCCGCC | 4 |
| wt-99270-99390 | 2251 | GGCTCCCGCGCCGGTCAGGGGACGGGCCTCAGCACCTCCCAGGCCACCAGCCCCAGTTCAGCAATCCACCAAAGACAAGGGTCCCCATAGACCTACGCGACCTGTACTTAGAGGCCCAGC | 8 |
| wt-99390-99510 | 2252 | TCCACGCCGCCCCCCTCCACCTTCAAGTCCCAATACATACAATAAACACATGATGGAAACCACCCCCCCCATTAAGGGCAATAACAACTACAATTGGCCATGGCTGTAAATAAAATGTCA | 4 |
| wt-99510-99630 | 2253 | TAACCTGGAGTCTGCATGTCTGTTGTTTTATTCAGTAAACCAGTAGTGCGCGTGAGTTCTTTAGGGCATCCACGATGTAGCCGCTCGCGGGGTTCCCCTCCCCAGTGATCATCTCGGATA | 4 |
| wt-99630-99750 | 2254 | GGGGATTCCTGTCCATGACCACGCAATTAGAGTGCCGGGCCCGGGACAGCGCCACATACACATGGCCGGGTTTGATGTTTCTGTGGCTGCCGAAGCAGATGGCGACTTTGTTTAGGGACA | 4 |
| wt-99750-99870 | 2255 | GACCCTGGGCCTTGGCTATGGTCATGGCCAGCTTTGAGCTAATGCCATAGTCACGGATGCTGCAGAGGTTCAGGGACTTGTCCTCTATCGTCTCATACAGTTTGTTAGTATTGTGTTCCA | 4 |
| wt-99870-99990 | 2256 | GGCAGCACACGAAGCCTGCCTCATCCTTGACCATTAGCCTGGGCATGCGTGAACTGCCAGCGTCCTGAGGCTGCTGCTTTCCTCGGATGCCAAAGAAGACGTTGAGATGCGTGTAGCCCA | 4 |
| wt-99990-100110 | 2257 | GAAGCGTGTAGTTCTCGGTGGTGGAGGCGTAGTCCAGGAGGCCGTGAAGGAGAGGCTCGTCTGAGGTGAACTCTATGTTGTCGCGAATCAGCATGTTGTTGGTAAATGTGCAGAAGGGGA | 4 |
| wt-100110-100230 | 2258 | GGTCCCTGAACTCCCTTCCGCCATAGCGGACGGCCACATCCAGGCATTGCCTGAAATAGGCCCTGAGGTCATTATATATGTTTAACAGGGAGCAGAGGGGGGCAGAATTTGCGGCCGGGG | 4 |
| wt-100230-100350 | 2259 | GAGCCAGTACTCGGGCATAGAAGACAGCGGCGGGGCTCCGCTCCCCATCCCAGGCAACCTCCAGCGGCAGTTCGCCCAGCTCCATCCCAGCAGTCACCTCCGGATCCCACGTACGCCCGG | 8 |
| wt-100350-100470 | 2260 | GCAGGGGCACAGCACCAAGCTCCGCCACGTATTCCCCGTTTTCACAGAGAGAATGTCCTCCGTGGCTAAAAGCGTAGATGCCTCCGTAGATGAGTCGGGCCAGGAAGCTGTAGACATACT | 4 |
| wt-100470-100590 | 2261 | CGGGCTGCTCATGCCCGTGGGCCTCCACGAAGCTGTCCGCCTCGAGCGTGTCCATAAAGTCCCCGAAGGTGCCGGTATAGCCACAGATGGACTTTTTGGTCTTGCAGTTGACCGACACCG | 4 |
| wt-100590-100710 | 2262 | AGCTGTGCTTGACGTAGGTGACATTGTAGGTGACCTTGACCCGTTCTTCGTCCTGCTCGGTGCCCACCGGGACCATGTCTTGGTCGGCGAACTGCGAGTAGTTACCGAGGCGTGCATAAT | 4 |
| wt-100710-100830 | 2263 | TCTTTTGGAGCCAGGTGTGGGCCGTGAGGCCCGGAAGCCCGACCAGGGTCTTGTACTGGGCCAGGGGATCGAGGAAGACCTCGCACTCCACCGGGCAGGTAAACATGGTCACCCCGCCCC | 8 |
| wt-100830-100950 | 2264 | CATCTCCCCCGGTTCCCCGCGCGGCACGCCCCTGCCCGGCAGTCTTGAGCGTGGCATGGAGGGTGGTGAGGAAGGTCTTGACCTCGGCGTGGGAGAGGAAGAGCCGGGTCCAGCCCACGT | 8 |
| wt-100950-101070 | 2265 | ACTGCGCGGGGTCCATTATGGCCGCCCTGGGGACGACGAAGCGGTCGACGTAGGCCAGGATGTCCGGCGAGAGCTCGAGGCCGTACTCGAGGGTCTTCATGAGGTGTCCAAACTGGACGT | 4 |
| wt-101070-101190 | 2266 | CGGTGCAGCGCTTGTTGTTGATGAAGAGGGCCCAGTTGCGGGCCACGTCCACGTAGGTCGCGGCCCTGGGGTTGCCCACCAGGAAGGTGAGGATGTTGTCGCACTCGCGAATCTTGTTTA | 4 |
| wt-101190-101310 | 2267 | CCTGGGTCTCGTGGCTAAAGGAGGACTGAAAGGCGTCTGTCTGGGTGGGAGAGCCCACGCAGACGATGCAGGGAATGCGGCCCCGGCGGTAGAGTGGGGTACGCAGCCAGGCGTTGAAGA | 4 |
| wt-101310-101430 | 2268 | ACCAGTAGCAAAAGACCACGGCTGTTAGAATGTGCACGGAAAGCGTTCCAGCTTCGTCCACCACGATCACATTGGTGGTCCATAGCTGCCCCTGGTGCATGTCTCTCAGGACCTCAAAGG | 4 |
| wt-101430-101550 | 2269 | CGGGGCCAGAGACTCCCGAGTAGAGCCCCCTGGGCTTGGTTCGCCTGAACTCGGCGGCAATGTCGGAGAGTACCGGCCAGTATTTGGCCAGGTCCCGCCGCTGGAGTTCCTCTAGGGCGG | 8 |
| wt-101550-101670 | 2270 | CGTCCGTAGAGCGACCATGACTGCTGACCCGCTGCGTCATATTTATGTGGCGGCTCTTGAACCCAAAGGCGCTATAGACGGTTGGGCAGTAGGCTCGGAGTGTCTGGGAGAGGTTCTGTG | 4 |
| wt-101670-101790 | 2271 | CGGCCACGGTTGTGGCTCCCGTGACCAGGCAGTCCATCGTGTGGTGGAGGCAGCTAACGCTGGTGCTCTTGCCAGCCCCCGCCGTTCCCGTAATTACATAGGCTGAAAAGGGCAGGAAGG | 4 |
| wt-101790-101910 | 2272 | GGGGCTCCGAGAGCTCCGGGTCAAACTCGGGGGAGAACGTCTCCATATCCGGGAGTTGTTGGACGCGGCGCCTAGCCAGGGTCCCTATCCTCCTGACTATACGCCTCACGGAGGCGTCTG | 4 |
| wt-101910-102030 | 2273 | AGGTCATGTTCAACATGAACGTGGACGAGAGCGCCTCTGGCGCCCTCGGCTCCTCGGCCATTCCTGTTCACCCCACGCCGGCCTCGGTCCGACTTTTTGAGATCCTGCAGGGAAAGTACG | 4 |
| wt-102030-102150 | 2274 | CCTACGTCCAGGGACAGACCATCTACGCCAACCTCCGCAACCCCGGAGTCTTCTCGAGGCAGGTGTTTACCCATTTGTTTAAACGAGCCATCTCTCATTGCACGTACGATGACGTGCTAC | 4 |
| wt-102150-102270 | 2275 | ATGACTGGAACAAGTTCGAGGCCTGCATCCAGAAGCGATGGCCGAGCGATGACTCGTGTGCGAGCCGGTTTCGTGAGTCCACCTTCGAGTCGTGGTCCACGACCATGAAGCTGACCGTGC | 4 |
| wt-102270-102390 | 2276 | GTGACCTGCTGACCACCAACATCTACCGAGTGCTACACAGCCGCTCCGTGCTCTCCTATGAGCGTTATGTGGACTGGATCTGCGCCACCGGCATGGTGCCCGCCGTTAAGAAGCCCATAA | 4 |
| wt-102390-102510 | 2277 | CCCAAGAGCTCCACTCCAAGATAAAGAGCCTGAGGGACAGGTGCGTCTGTCGGGAATTGGGGCACGAGAGGACCATCAGGAGTATCGGGACGGAATTATATGAGGCAACGAAGGAAATAA | 4 |
| wt-102510-102630 | 2278 | TAGAGTCGCTCAACTCCACGTTCATCCCCCAGTTTACGGAGGTGACCATCGAGTACCTTCCGAGGAGCGACGAGTATGTGGCCTACTACTGTGGCCGCCGCATCAGGCTGCATGTGCTCT | 4 |
| wt-102630-102750 | 2279 | TCCCCCCGGCCATCTTTGCCGGAACGGTGACCTTCGACAGCCCGGTGCAGCGCCTCTACCAGAACATTTTCATGTGCTACCGCACGCTGGAGCATGCCAAGATCTGCCAGCTCCTGAACA | 4 |
| wt-102750-102870 | 2280 | CGGCCCCTCTCAAGGCCATCGTGGGCCACGGGGGGCGAGACATGTACAAGGACATCCTGGCCCATCTGGAGCAGAACTCACAGCGCAAGGACCCCAAGAAGGAGCTGCTGAACCTGCTGG | 4 |
| wt-102870-102990 | 2281 | TCAAGCTCTCGGAGAACAAGACCATCAGCGGGGTCACGGACGTGGTGGAGGAGTTCATAACGGATGCCTCCAACAACCTGGTGGACCGCAACCGTCTATTTGGCCAGCCCGGGGAGACAG | 4 |
| wt-102990-103110 | 2282 | CTGCACAGGGCCTAAAGAAAAAGGTCTCCAACACGGTGGTCAAGTGTCTGACTGATCAGATAAACGAGCAATTTGACCAGATTAATGGCCTAGAGAAGGAGAGGGAGCTCTATCTAAAGA | 4 |
| wt-103110-103230 | 2283 | AGATCCGCTCCATGGAGTCTCAGCTGCAGGCCTCCCTGGGTCCCGGCGGCAACAACCCAGCGGCGTCAGCCCCCGCCGCAGTTGCGGCAGAAGCCGCGTCTGTAGATATACTGACGGGCA | 8 |
| wt-103230-103350 | 2284 | GCACCGCCTCCGCAATCGAAAAGCTGTTCAACTCCCCGTCCGCCAGCCTGGGTGCCAGGGTGTCTGGTCACAATGAAAGCATCCTAAACAGTTTCGTTTCTCAATACATCCCCCCTTCGC | 4 |
| wt-103350-103470 | 2285 | GGGAAATGACTAAGGATCTGACTGAACTTTGGGAAAGCGAGCTGTTTAACACCTTCAAGTTAACACCCGTGGTTGATAATCAGGGGCAGCGTCTCTACGTCAGATACTCGTCAGACACGA | 4 |
| wt-103470-103590 | 2286 | TCTCTATATTATTGGGCCCCTTCACCTATCTGGTGGCAGAGCTTTCACCGGTGGAACTCGTGACAGATGTCTACGCCACCCTAGGCATCGTGGAGATCATCGACGAGCTCTACCGGAGCA | 4 |
| wt-103590-103710 | 2287 | GTCGCCTGGCCATCTACATCGAGGACCTCGGTCGAAAATACTGCCCCGCGAGCGCGACCGGGGGAGATCATGGCATCCGGCAAGCACCATCAGCCCGGGGGGACACGGAGCCTGACCATG | 8 |
| wt-103710-103830 | 2288 | CAAAAAGTAAGCCTGCGCGTGACCCCCCGCCTGGTGCTGGAAGTTAACCGCCATAACGCCATCTGCGTGGCCACCAACGTCCCTGAGTTCTACAATGCCAGGGGGGACCTTAACATCCGA | 4 |
| wt-103830-103950 | 2289 | GACCTCCGGGCCCACGTCAAGGCCCGGATGATCTCGTCCCAGTTTTGCGGCTACGTCCTCGTGAGTCTGCTGGACTCCGAGGACCAGGTCGACCACCTCAACATATTCCCCCACGTGTTC | 4 |
| wt-103950-104070 | 2290 | TCCGAGAGGATGATCCTGTACAAACCCAACAATGTGAACCTTATGGAGATGTGCGCCCTGCTCTCGATGATTGAGAATGCCAAGAGCCCCTCCATAGGCCTCTGCCGGGAGGTGCTGGGT | 4 |
| wt-104070-104190 | 2291 | CGCCTGACCCTCTTGCACTCCAAGTGCAACAATCTGGACTCTCTGTTTCTGTACAATGGGGCCAGGACGCTGCTGTCCACCCTGGTCAAGTACCACGACCTGGAGGAGGGGGCTGCCACC | 4 |
| wt-104190-104310 | 2292 | CCCGGGCCGTGGAATGAGGGCCTGAGTCTCTTTAAGCTGCACAAGGAGCTGAAGCGCGCCCCATCCGAAGCCCGGGACCTCATGCAGAGCCTCTTTCTGACCTCGGGGAAGATGGGGTGC | 4 |

| | | | |
|---|---|---|---|
| wt-104310-104430 | 2293 | CTGGCCAGGTCACCCAAGGATTACTGCGCGGATCTAAACAAGGAGGAAGATGCCAACTCGGGCTTCACATTTAACCTGTTTTATCAAGATTCTTTATTGACCAAGCATTTCCAGTGCCAG | 4 |
| wt-104430-104550 | 2294 | ACCGTCCTCCAGACCTTGAGACGCAAGTGCCTCGGGAGTGACACGGTCTCAAAAATAATTCCCTAGAATAAACTGAGAACAGTCATCAGTAAATCTGTCTCTCGCGTGATTTCCATAGGA | 4 |
| wt-104550-104670 | 2295 | ATGGTGTAGCCGGGGTGGAGGGCCGATATCACATCAAGCAGAAAGGCCATAATCTCTCGAAAGTAGGCGGTGGGGCTGAGACCATGCTCAGTGGCCGTCTGGCAGGGGGCCGGGCGCGCT | 4 |
| wt-104670-104790 | 2296 | CCGTCCTTGTCCAGGAGACACACGTGGCTTCCAGAGAGGCGCAGCCCAGCCCTCCGCAGCCGCTGAAGCCAGGCTCGCGGAAGAGCCCAAAACCTGTTTCGGCGCCGCCCGGGGGCCAGT | 8 |
| wt-105030-105150 | 2299 | TCTCCACCCATCACATCCCGGTCGAGCCGGCTGGCATCCTCCGAGTCTTTCACGCCTTGCAGGAAAGCCTAGGAGATACAGCAACAGAAAGCTATTAGCCGGTGGTTCCCCCACCATCAT | 4 |
| wt-105150-105270 | 2300 | TCTTCCTGTTAACGGGAAGAATAAGAGTTGGGCAAACCCCGGGGGCCGCGCTCTCCCACCCAGCCCCGCTTCTCACCTGTGCTAGTGGCTCCTCTGAAGGATGGGCGGAGGTTGGTGCCA | 4 |
| wt-105390-105510 | 2302 | TGCCATAGCGGTGAAGGGCCCGCAGCCAGGTTTGCGCGTCCCGCGCCTGCCCTCCGCCATCAGGCGTTCCCACGGGGGCGCCCCTGGCAGAGAGGTGGCAGCGGGCCAATTCGTAGAGCC | 8 |
| wt-105510-105630 | 2303 | ACCAAGTGGCATCAGCCTCAAGGATGGCTGTGGCCTCCGCGCGCCCGACCACCGTCGTCTCGTCCTCCCCCCCTCCCTCGCCGCCTTCCCGCGTGCAAACGTGGCGAGGGTTAATCTCCT | 8 |
| wt-105630-105750 | 2304 | TTCGGGTCGGGGGCCAGATTTGTTGTAGGAGCAGCGAGCCGCGTCGTTGCCCTGACCGCGCGTCGAGGCCCAGGAGGGCGTCTGCCAGGGGCGTCCCAGAGACTCCCAGGTTCAGGTCCA | 8 |
| wt-105750-105870 | 2305 | GTAGCAGGAGACCCTCGCTGTGTGGCGCCCGGTGCCAGAAGGCCGGCCTCGCCCGTCCCACATAATGGATGGGCAGGAAGGGAAAGCCCGGGACATAGGGCTGGAAATCTGAGCCCCCTG | 8 |
| wt-105870-105990 | 2306 | GGCAGAGTTCGGGGTCCAGGAGGTAGAAGATGGGCTTGGTGCCTCTGTGGTTGGCGTAGCAGGAGGCATAGATACTGCGGAGGAAGGCGTAGAGCCCGCCCCCGGCCATACTCCAAGAGT | 4 |
| wt-105990-106110 | 2307 | TGACAAGCCAGGACTCGAATCCCCCAGCCGGCTCAAGAATTTTCAGGCTGACGCGGTGCCGTCGGGCGTCCCCACCACGGCCGGTGGCCCCGTCGGACGACACCAGATCTACTTCATAAG | 4 |
| wt-106110-106230 | 2308 | TGACCGGTCGCAGGATGTCCCTAAAGGGGACGGGAGAGGGGTCGTCGGGAGTCTCGGTGGAATAGGTGAAAACATCCCCACGCGGTGTCCTGATGTATACGTCCAACTGTCCGGGAGACT | 4 |
| wt-106230-106350 | 2309 | CAGAGTGCCTCTGAGCATGGGGGCATGTCTGTTCCCCCTCCATCTCGGACCCGAAGCCATCAACAGGTGGGGGTTGTTGGTCCCGCCCATCATCCCCCGAGCAGCTTTGGCAGACCACCT | 4 |
| wt-106350-106470 | 2310 | GTGCTGGAAAGAGAGGCTGGAAGATGAGGCCCTGCTCATCCTCCACCCTGGCGGCGGACAAGAGTCTGCGGTCTCGGGTTCTAAATGAAAGGTCAAATAGGTCCTTCTCGGCGGCATCGG | 4 |
| wt-106470-106590 | 2311 | CGAGCATAGCAATGAGCCCCCCGCTGCGCCTGAGCTCCCGCTCCCATCGCAAAAAGTTGAGTTCGGTAGTCGAGGGCGCGTTGACCACGGGGGGCTCCAGGGAGCCTCCAAGCGGCGGCT | 8 |
| wt-106590-106710 | 2312 | GGCAGGCCTGCACCACGATCAGAGTCTCAACGTCCTCCCTTTTGATGGGCACGATGCCCACGACCCAAATCGCCCACCACCGCCCTGCGGTCTGGGTAACATTATAAAAGGTAACCGAGC | 4 |
| wt-106710-106830 | 2313 | TGACGCGGGCCCTGACGCTCTCCGCGGGTGTTTCCATCATTGTTTGAGATCTGAGGAGGACTGGACCCTTTAAAACATCCGGTCACGCCCTTTGCAAATTATTTAAAAGGTGAATGCTCA | 4 |
| wt-106830-106950 | 2314 | ACTGAGACCATCGCAATCATGAAGTCCTCCAAGAATGACACGTTCGTCTATAGAACGTGGGTCAAAACGCTTGTTGTGTACTTTGTGATGTTTGTCATGTCGGCGGTGGTCCCCATCACC | 4 |
| wt-106950-107070 | 2315 | GCCATGTTCCCCAACCTGGGGTACCCCTGCTACTTTAACGCACTGGTTGATTACGGGGCACTTAACCTGACCAATTACAACCTGGCCCACCACCTGACCCCCACGCTCTATCTGGAGCCG | 4 |
| wt-107070-107190 | 2316 | CCGGAGATGTTTGTCTACATCACACTGGTCTTTATCGCGGACTGCGTGGCTTTCATCTACTACGCCTGCGGCGAGGTGGCGCTAATCAAGGCCCGAAAAAAGGTCTCGGGTCTTACAGAC | 4 |
| wt-107190-107310 | 2317 | CTCTCGGCCTGGGTCTCGGCAGTGGGCTCCCCAACCGTGCTGTTTTTGGCCATCCTCAAGCTCTGGTCCATACAGGTCTTCATCCAGGTCCTTTCCTACAAGCACGTCTTTCTCTCGGCC | 4 |
| wt-107310-107430 | 2318 | TTTGTGTACTTTTTGCACTTTCTGGCCTCAGTTCTACACGCCTGCGCATGTGTGACCCGCTTCTCCCCGGTCTGGGTGGTCAAGGCCCAGGACAACTCTATTCCCCAGGACACCTTCTTG | 4 |
| wt-107430-107550 | 2319 | TGGTGGGTGGTCTTCTACCTGAAGCCCGTAGTTACAAACCTGTACCTGGGGTGCCTTGCCCTGGAGACGCTGGTCTTCTCGCTCAGCGTGTTCCTGGCCCTGGGCAACAGCTTTTACTTT | 4 |
| wt-107550-107670 | 2320 | ATGGTGGGGGACATGGTGCTGGGAGCCGTGAACCTCTTCCTCATCCTGCCCATCTTCTGGTACATTCTGACGGAGGTGTGGCTGGCCTCCTTCCTGCGGCACAACTTTGGCTTCTACTGC | 4 |
| wt-107670-107790 | 2321 | GGCATGTTCATCGCCTCCATCATCCTGATCCTGCCCTTGGTCAGGTACGAGGCCGTCTTTGTCTCCGCCAAGCTGCACACCACTGTGGCCATCAATGTGGCCATCATACCTATCCTGTGC | 4 |
| wt-107790-107910 | 2322 | TCGGTGGCCATGCTCATCAGGATATGCCGGATTTTCAAAAGCATGCGCCAGGGCACTGACTATGTCCCTGTCTCGGAGACGGTGGAACTGGAGCTAGAGTCAGAGCCGAGGCCTAGGCCC | 4 |
| wt-107910-108030 | 2323 | TCGCGCACGCCATCACCCGGGCGCAACCGCCGCCGCTCTTCTACGTCCTCATCTTCCTCCAGGTCAACCAGGAGACAGAGGCCCGTCTCTACCCAAGCCCTCGTCTCCTCCGTTTTACCG | 4 |
| wt-108030-108150 | 2324 | ATGACGACGGACAGCGAGGAGGAGATCTTCCCCTAATGCAATAAAAACTTAAAACACTGAGGTTACTTTCCCGTCATTCTTTCGGGGGAACGAGGGGAGGCGGGAATTGGGTTAAGATAG | 4 |
| wt-108150-108270 | 2325 | GGGCGAAGGGTGGGGGTGGGTGCAAGAATTGGGGCTGGGAATGGAGAGGGGAGTGGGCTAGGTGCCGACACCGGGGTGCCAAGATAATGGATTGAGTAAGCATGGGGCTCTGATCGGGTC | 4 |
| wt-108270-108390 | 2326 | CGCCGGGTTCTCAGGGGTGTAGTGGGTGGGCATTGCATATTTTTGCCGCGGTGCTGTTGGGCCTTGGACTCGGGGTGATCATCCGTACCATCACCCGCACCCGCACCCCAGTCCACAGCC | 4 |
| wt-108390-108510 | 2327 | ACCGGCCAAGGTCCTGGGCCTCCCACCACCGTTATGCCTCCCCCTTTACCCATTAATTACAAGAGATGTTAGTTTGGTTTTTTATTTGGCAAAAACAGCAATTCATCATTTTCAGAGTCC | 4 |
| wt-108510-108630 | 2328 | TCATCATATTCGAGCCCCTCGTTGGTTTCCCCGCAGGCCCTCCCTTCTTCGGCCGCTATTAGCTTAGTAGTCTCCAGGTTAAACTCCTCATAGTCATTATACAGGTTGATTATTCCCCCG | 4 |
| wt-108630-108750 | 2329 | TCCACGTCGCCTATGGAGTTGACTCGTCGTCGGCAAAGAGACCAGAGGGCACCCATGGCGCGGTGTCAAAAGTATTGTCTGCGTACGCTTTCCAGGAGCCAGCCGCGGTGCTCAAGGTCT | 4 |
| wt-108750-108870 | 2330 | TACGGATGACAGAGTCCGGCAGGACCACGGGTGTCACCAGCACCGCCACGGGAATCTCCACCGAGGCGTCCAGAAGCAGGTCTGAGCCGAGCGTGCAGGTCGCCGGGTCTAGAGGCGACC | 8 |
| wt-108870-108990 | 2331 | GTTTTCGAAAGAAGGCCGTCACAATGTTCACCCGGGGTGAGCAGTCTCTCCCGGGCTTGCCACCCCCACTGTGGCGGACGTAGTCTCCAACAATTTTGTATTGGAGGAGCACCTGGTAGA | 4 |
| wt-108990-109110 | 2332 | AGTAGTTGTGCCGTGGATTGATGAAGATGTTGACTGGGACCCGGTCTTTAATACCAATGCGCCCCGCATTTTCGCTTGGGTCCGTCATTACGTAGAGCATAGACTCCACCCCCCTGTTGG | 4 |
| wt-109110-109230 | 2333 | CAGCTAGGCTGTCTGCCACCAGGTCATGACCGGGGCCCAGTTTGCGCTTACGGACATCTTTAAGATTCCAGGCCTCATCCTGCGTCAACAGATAGTCACCCTCCGAGGGCAACCGCCCAT | 4 |
| wt-109230-109350 | 2334 | CCGGGACGTACTCCACGGTAGGACGAGCTATAGAATTGATAAATCTGATAAATGACCTCTTGCATGGCCTCTTGTAAAGCGCAGTGTAGGATGGGTAGATGGGGTCAAATTCTGACTTGG | 4 |
| wt-109350-109470 | 2335 | AAAAGAGGTACTTGAAGCGGCACTTAATCTCATAAATGCAGCTCCGGTCGGTGAACAGTATAAAGTCTCCCTGTGACTCCACATTGACGCAAAGATCCAGAGACACCCCAAAAATGCCAT | 4 |
| wt-109470-109590 | 2336 | CCGTGGGACTAATCATAAAGCCAAATTGACGGTTGGCGGATGCGTCCCCGCAGATGAGCTTACAGACAATGTCCTTGACCGTGTCCTCACACCGCAGGCCAAAGGCCACAGGTCCCCCAA | 4 |
| wt-109590-109710 | 2337 | AGTAGTGATTTGTGGAGATGGGAGCTGGCTCAAACACCTTGGTGGGTCCATTCTTAATGGTGGAGAGCAGCTTGGAAGAGGAAATTATGCCATTTCGCAATATGTCCCACATCAGGTTCT | 4 |
| wt-109710-109830 | 2338 | CAGACTGCCCCCTGGTCATGGACTCCACGTACGAGCAGAGAACAGTCCTCTGCTCGTCGGTGGCCTCCTGTAGCCCCCAGTAAATGGATTTCAGGGAGGGACCGTCCTTGCTGTCATTCT | 4 |
| wt-109830-109950 | 2339 | CTTGGACTAACGAGGAGACAAAGTCACAGAAGCCAGTTTCACCAGAGAACTCTTGTATTTGTTTACAGAGGCAATAGAGATAGACAAAGCGCATGGCCGGCATCTGAGGTGGACGGTCAA | 4 |
| wt-109950-110070 | 2340 | GGTTACGGACAAAGGCCTCAGTCTCCGGACTGCGGAGGAAGCGGGCAAACGTGTAGGAGGTCATCTCCTCCATGGGATCCTCGAGCTCATCCACGTCGGCCATCTGGACCAAAGAAGTCG | 4 |
| wt-110070-110190 | 2341 | TCTGCCAAGAGTTCAGCTACCAGACCTGGAAGATGAGGGTGCTCAAACCGTGGGCGACAGTTGAAGAAGTAGCTCTCCTTGAACCTCTTTTTAAGGCTCCGGCACCACTGCAAGAATTGA | 4 |
| wt-110190-110310 | 2342 | CTCATATGCTCCGCCGTGACATCCACGCACGGACTCTCGCCACACGAGGTCAGGCCCATGTCTAAGTTCAGGTTCCACATCTGCGACAGCACCTCCAACAGCACCACCTTTGGGGCTGCA | 4 |
| wt-110310-110430 | 2343 | AATTGCAAAAAGTAGAGCGGGTCGGATCGGTCAAATCCCATGTCAGGGTTGGGGTAGGGGATTTTGTGGGTGGAGTCAGCGAGGTGCATGATACCATAGAGCAGCGAGTAGCCGAGCGAC | 4 |
| wt-110550-110670 | 2345 | TCCTTGGCTATAGAAAATGGTTCCCTCTGGCAATAGAGGCGATAGAGCTGCCGGCCCTTAGAAGACTTTAGCCGCACATCCAGCATCTTGTTGCGGTCGTGGAGGGAAGCAGTCCCATAA | 4 |
| wt-110670-110790 | 2346 | TCAGTCAGGACCAGCCTACCCATGCCCCACATGGTGTCTGTGAAATCCACCAGGATGTTGCTGGGGCTAATGTCCGAATGGAAGAGGCCGCAGTGCCGATTCAGAAAGTAAACGGCATCT | 4 |
| wt-110790-110910 | 2347 | TTGAGGCCCTGAAAGCCCCGCACCAGGGGCTCAATACTACCATCATGCCAGTGGCCATAATCCTGGAGACTGCATCTGAACTGGGGCATAAACAGGGCGTGGCAGGACGTGCAGGCCGAC | 4 |

| | | | |
|---|---|---|---|
| wt-110910-111030 | 2348 | AGGTAGTCCACCAGGGCCTTGTCCTGCCCATCCTCGGCCGTGGCCTTCCCAATCTGAATCATGTCACACACCATGAGCTCGTGATACAGCTCCGTCACAGAGTCATAGAGTTTGACCGTG | 4 |
| wt-111030-111150 | 2349 | GCATTATCTGCATGTGCATACACGGCCCCGTAGCTCCCCCGCCCCAGCAGATACTCGCAGGTAATGGGGAGGTGATCACAGCGCGTCATGTTCTCCGGCAGCTTTACATAGAGGGTCTCC | 4 |
| wt-111150-111270 | 2350 | GTCATGTCATCAATGTTGGTCACCTTCAGGTGTTTGTGCTGAAAGGTGAAGTAATCAATGACAGTCACCTTCCCCAAAAAGGCCTGGGTCTCTCGAGGGGGTTCTGGGGAGACACTCAAC | 4 |
| wt-111270-111390 | 2351 | TCGCCACTGCTGGAGGAGTTCGTCGGGCTCAACTCCGCAGCCATATTCACATCCATGTTCCTCAAATGGCTCGAGGGCCTGTCGCAGCTCGTCTCTGGCCTCAAGCTCCTGCTCACGGAG | 4 |
| wt-111390-111510 | 2352 | CTCCTCCACCCGCTCTAGCTGCTTGTAGTTGATTTTTGGAAATTGAGTCTTGGTCGCGGTGACCACCCTCTGATAGGTAGAAATTAGCTGTTTGGACTCAAACGTCTCCCTTGCGTGGCG | 4 |
| wt-111510-111630 | 2353 | CAGGGACTCTAAGGCACCCCGAGCAGATGTAAACTGTGTTTCAAACAGAGCGTGGTCCCTCCCAAATCTGTCACGTGCGCTCACAGCCGCTCTCTTTTCTACCGAGGCTCTTAGTTGCTG | 4 |
| wt-111630-111750 | 2354 | GGCCACCAGATCTCGCTTAGAACTACTCATCTTCATAAGTCACCATGTCCGCAACTATGGAGCCCAGATCATACGTGGGGTAGAGTACGGTAGTTCCAGTGGAGGCTTCCCGGTAATTTC | 4 |
| wt-111750-111870 | 2355 | CCACAGCGTCCACCATATATCTTTCTGCCTCTCCCGTTAGAATTAGGCAAGGATCATACGTGTCCACCGGCCTTTTATACTGAGCGTTTAGGTTTTGTTTATGTAGCAAGCACAAAAGGC | 4 |
| wt-111870-111990 | 2356 | ACACACGAGTGATGCAAAAGGGTTCCTGAGGCAGCAGGCAGAGCTGTTTTGCCATTTTATTCAGGCGGCTAACGTCAAAGGGAGGAGCTATATCCTCACCCTTCCAGTCACGCACGTCCA | 4 |
| wt-111990-112110 | 2357 | AGTACAGGGCATACACACACCTGGTGAGGTGTGCCAGGAATGCCTCTATGTTGGCACATGGTGTATAAACCGCAGTGGGTAGCAGAATAGGGCCCCTTTTGCCCCGTGCCGCAGCGTAAA | 4 |
| wt-112110-112230 | 2358 | CGCAGTGACGCTCTTCGCAGTGGGACCTGGGGCCGTAGAAGAGGGCCCACATCCAAGGGAGTGGGTCTTCAGGCACCAGGGAGGTCCAGGTTTGGGAGTGGGCCAATATTTGCAAGGCCT | 4 |
| wt-112230-112350 | 2359 | GACCTATAACCTCATCTTTGTTCCAGGCCAGCGCAATTCGCATAAGGTCCCCATCAAACACCTCAAAACACAGACCCATGCCCATTTCAGGCTGAGAGGGCTCCATCCGGCTCGACCAAC | 4 |
| wt-112350-112470 | 2360 | CTTGTCCACCAAACTGCCATTCTTCTGGTAAACGGGGGTTGAGGGGCAAGAGCTCCAAAGCCAGGCTCGAGAAGTCATAGTCATCCTCGGCCACACGGCCGGAGCTCCGGGCCTCGTGCC | 4 |
| wt-112470-112590 | 2361 | AGGGCCTGTTGTCCTGGGGGAGGATATTGGACACGAGCAGGAAGCTCTTGAGTGGCGTCTCCACCAGCTTAAATTGCTCGGGCGTGTCCTGGCAGGCCTCCAGTGCCAGTTCCAGACACT | 4 |
| wt-112590-112710 | 2362 | GCCCATACCTGCGGGCGAGCATCGGGTCATCGGGCATATCGGCCTTGACCGCGTTGAACATGCTGTATGCCTCGCAGCGCGGCCGTCTGACCGAGAACCTAAGAAACGCCCTTCAGCAGG | 4 |
| wt-112710-112830 | 2363 | ACAGCACCACGCAAGGCTGCCTGGGTGCCGAGACCCCGAGTATTATGTACACAGGGGCCAAGTCAGACAGGTGGGCTCACCCTCTGGTGGGCACAATTCACGCCAGTAATTTATATTGCC | 4 |
| wt-112830-112950 | 2364 | CAATGCTTCGAGCATACTGCCGCCACTATGGCCCCAGGCCCGTGTTTGTAGCTTCTGATGAATCATTACCCATGTTCGGTGCGAGCCCTGCCCTTCACACCCCAGTCCAGGTCCAGATGT | 4 |
| wt-112950-113070 | 2365 | GCCTACTACCAGAGCTACGCGACACGTTACAGCGCCTGCTGCCACCACCCAATCTTGAAGACTCCGAGGCCTTGACGGAATTCAAGACCAGCGTGTCCTCTGCCCGTGCCATCCTTGAGG | 4 |
| wt-113070-113190 | 2366 | ACCCCAACTTTTTGGAGATGAGAGAGTTTGTCACCAGCCTGGCCAGCTTCCTGAGTGGTCAGTACAAGCACAAGCCCGCCCGCCTAGAAGCATTCCAGAAACAAGTAGTGTTACATTCTT | 4 |
| wt-113190-113310 | 2367 | TTTATTTTCTGATCTCAATCAAATCTTTAGAGATTACAGACACCATGTTTGACATCTTTCAAAGTGCTTTCGGATTGGAAGAAATGACGCTGGAGAAGCTGCACATTTTTAAGCAAAAAG | 4 |
| wt-113310-113430 | 2368 | CCAGCGTGTTTCTTATCCCCAGGCGCCACGGCAAGACCTGGATAGTCGTGGCCATCATCAGCCTCATCCTCTCGAATCTCTCCAACGTGCAAATAGGCTACGTGGCTCACCAGAAACATG | 4 |
| wt-113430-113550 | 2369 | TCGCGTCCGCCGTTTTCACTGAAATTATTGACACCTTGACCAAGAGCTTCGACTCCAAGCGTGTAGAGGTCAACAAGGAGACCAGCACCATCACGTTTAGGCACAGTGGGAAAATCTCCA | 4 |
| wt-113550-113670 | 2370 | GCACCGTAATGTGTGCCACCTGCTTCAATAAGAATGTAAGACCTGACGTTTCAGTACTTGGCAATTGTAGAGCATAGCCCGGCTGTAAAGGTCAGAAAATCGCAGCAGGGTCCAAGGTTG | 4 |
| wt-113670-113790 | 2371 | TGCTGTACATGGGACCTCTTTCCCATTAGCAAGAACCCCCTGCAGGACACGTGACATGTCCGGGTGCATTTTGGGTGGGTTAAATCTCAGTCCCACCACAAAGGGGGCATCCTCCGGTTT | 4 |
| wt-113790-113910 | 2372 | GAACATCAGACCCAACAAAGCCCGATGCCCAGTTATGGGTACGTAGTCGTTGTTCAGGGCCGTGCATGGCAGCAGACAAGGACAGGTGCCAGATGTGCCTGGGCTATCGTCCTCCGTCCA | 4 |
| wt-113910-114030 | 2373 | GCCACGCAGGATGTTCACGTGGGCCCCGGCACCATAGCATGTCACACATTCCCCGTTATCACATCTGGTTAGCAGGTTGATAAAATGGGTCAGTGATGGAAAGGTTGGCATATTGGGGCA | 4 |
| wt-114030-114150 | 2374 | GCACATCAGCATGTCCATGTTAACGAAAAACATGTACAGGGCCCCTTCTGCATACCAGGCACCACCCCGTCCCAGTGGGATGATCTCCGAGGGTGTGATATCTTGCAGTTCTTCTACTGT | 4 |
| wt-114150-114270 | 2375 | TTTAACGGCGGTTGAGGTGGTAAAGACGTGGGCCGTGGTCAGATCTGTGCAGGTGACTACAGGGTTACCCCTAATCTCCACAGGCACCGCCTCACCCACTGCATCTGAGAATACCCCAAA | 4 |
| wt-114270-114390 | 2376 | GTACATGAGAGTCAGGCTGTGTGGCCCCTGGACTGCCTTAGTGAAGAGAACCTCGGGCCTGGCCACGGTGGCTAGGGTTCCATTGATGTAGACGGTCACATAGGTGGGCTTCTTCTTGGG | 4 |
| wt-114390-114510 | 2377 | CTTCAGCACAATGAGGGTAACATTCATGTAGGTTTTAGGAGGTCCGGCTATCTGAGGCACGTACACAGCTGACACGGCGGTTGTGGCCGTATAGACTTTCATCTGGGGCGTAGAGGCATC | 4 |
| wt-114510-114630 | 2378 | GCTCAGCACCCAGAGGCACTCCTTGTTGAGGAACTTGCGAAGCTGTTCCCGGCTACTGTTCGCGGCGGATGCCATGACGTGCCAGAATATATCCCCTCTCCTCGGGGGTGAGTGCCAATT | 4 |
| wt-114630-114750 | 2379 | GGCCTTTAATAACAAAGCCCCCAGGCAGCACCAAAAATGCCTGCCCGTCCGATGTGGTGGCCAGGTGGACGCAGTGCCCGTCAGTTCCAAGGGCTACTAGCTGGGAAGCAGCCCCAACCA | 4 |
| wt-114750-114870 | 2380 | GCCCACCCGGGGGCCTGGAGTCGATCACCTTACCCCAGGCCGAGGCCCCTTCCTCATACAGCGGGTGGCTATCTATCCATAGGCAGGCATCCGGCGTCTTTGGTGCATTGGAGATAGTAG | 4 |
| wt-114870-114990 | 2381 | CTTTCACCCAACAACTTTCCCAACTAACCCGTGTCTGGACAGTGAAGAACGCTTCCCTGATCAGGTCTGAATTTTTATAGATACGGGAGTAGGAGGTGGGAATAACAACTGGGATTTCTT | 4 |
| wt-114990-115110 | 2382 | GTTGTGCTGTCCAGGCCTGCATGGCCAGTTTTTCCCTGAAGCTAGCAGAAATTCTGAGGGCCACTGAAATGAGGAAGCGAAACTCCCTCTCTGGAGCTCCCAAAATTGAAACCTCAGCAA | 4 |
| wt-115110-115230 | 2383 | GATCTGTTGCTGGGGAGGCATGGGTGACAGCTGTCATCCTGTGCAGTCTGCCCTGGGCACTCAGCTCTGGATATGTGACAACATAGAGAGCGTGGGGGCTAAAAATATGAGCAATTCCCC | 4 |
| wt-115230-115350 | 2384 | TGACCAGGGCCCTGGACTCACGAATGGCCCGACGGGTCTTAGAGAAAGAAACAGGCACCCTCGAGAGTGCCCCCGACCCGACCCCCACAGTGCCGCCAGTCCCTGCTCGGCCTCCGCCGC | 8 |
| wt-115350-115470 | 2385 | CTTCCCCACCGGCGCTGCCCCGGATGTTGCTGGGGTTCTCGAGGGCTGGGTGGTGCTTGGACACAGAGGTCTCAGCAGCCGCCTTGGTCTCGGCCCCGGCCCTAAGTCTGAGCCCCAGGC | 8 |
| wt-115470-115590 | 2386 | AAAGGGCCGGACTCCCAGCGTGGCCCAACCTCTGCTCCCCTCTATTCTCCTCTTGCGTTATCTCCAATAGAATTTGCTTGAGGTCATACGTTTTAGGGTGCTCGACCTGGGCCGCGGCCA | 4 |
| wt-115590-115710 | 2387 | CCGGCATATGCTCTATACCCGCCCCTCCGGGGGGCCCAGGATCTATAGGTATGGGCTGCATAGCCGCAGCAGACTCCTGGACCCCAGAGGCCTCTCTGATAAGATGCCCGTCGGTCAGAG | 4 |
| wt-115710-115830 | 2388 | CCCTTTTGGCCCCCTCAAAGAGAGACAGGTAATAAATCTGTAGCTCCCCAACCAGCCCTCCTTCATCGTAAAATCGAAGGGCGGCCACGTGGAAGGGGTTGTAGAGCTCTGGAAGGCCCT | 4 |
| wt-115830-115950 | 2389 | CATCGCAGTACACTGGCACACTGGTAAACGTGCCCCGATGGCTAGGCCGTCCGGGCAGCATGCCCCGAGCAGCAAACACGCGGCAGACCCTCGTGAGACCCGTCCGGTCACTGAAGAGAG | 4 |
| wt-115950-116070 | 2390 | TCTGGCACCAGGCCCCCTCGCAGTTTGGCACGCGATTGGGGCAAAGCTCTGCCATAACCGTGTCGGGAACAAATAGGTGCACGAGGAGGGGGGTCCCGAGGCCACTCAACACTTGGTTGT | 4 |
| wt-116070-116190 | 2391 | CAATGTGGACATCCATAGCTCTCTCATGCGTTTGGCTACAGCATCATAGCGCTTGTTTCTGGTGGATTTAAATAACAGGGCCCCGTAGACAGTCTTTTGTGAGTAAATAGAGATGATGAC | 4 |
| wt-116190-116310 | 2392 | ATGGATGTAGAGACTGAGGACCACATCCACCACCTTCTCGGAGGAGGCCCCCCTAAACAGCATCAGGCAGCAAGGGAACACAAAGGAAACCAGGGCCGGGATGTGAGGCCTCAGCGCCCC | 4 |
| wt-116310-116430 | 2393 | CTCCTGATCAAAGAGGGCCTCGCTGACCCCGGAGATGACATTCTCATTCAGAAAGTAGTGATAGAGGTGATTGACCACAGTCTTAACCAGGCCCTGGACTTGTTCAGGCTCCCACTTGTC | 4 |
| wt-116430-116550 | 2394 | CCGCTGGTCCTGTGTGTCTTGTCGGATCTCGGTCCAGGGCCTCAGCGCCGGCTGGAAATGCGGCCCCATGTAGTTGCCTGTAAGGGCGCACACCACTCCCTCATGGGTCTCAATCAGGGT | 4 |
| wt-116550-116670 | 2395 | GCACTCGCTGGATCCATCACATACGTGGTACTCGCCACAGCCCCAGCAGGCAAACACGGAGGCCATGCTCTCAGGTAACGGGAGATGGAACTCCAGCTTACTATACGAGCACAGGTGGCG | 4 |
| wt-116790-116910 | 2397 | ACCTCCTGCTCAAAGAGACCTCCCACCGCCACGTAGACGCGGGCCACCGTCCGGGGAAGGTCAGTGGGGTCCCAGCTCAGCAATTCTCCAAATTCTCTCTCCCCAATAGTGCCTCGCTTC | 4 |
| wt-116910-117030 | 2398 | TTATCCTGTCTTTCAGAGCATCCGGGGGCAGACATTTCACCTCTTGTTTGTGGACGAGGCTAACTTTATCAAGAAGGAGGCCCTGCCGGCGATCCTGGGCTTTATGCTTCAGAAGGATGC | 4 |
| wt-117030-117150 | 2399 | CAAGATTATCTTCATCTCGTCTGTGAACTCGGCTGACCAGGCCACCAGCTTTCTTTATAAGCTGAAGGATGCTCAGGAGCGGCTGCTGAACGTGGTAAGTTATGTGTGTCAGGAGCATCG | 4 |

| | | | |
|---|---|---|---|
| wt-117150-117270 | 2400 | GCAAGATTTTGACATGCAGGACAGCATGGTCTCATGCCCCTGCTTTCGCCTGCACATCCCGTCCTACATCACCATGGACAGCAACATCCGAGCAACCACCAACCTCTTTCTGGACGGGGC | 4 |
| wt-117270-117390 | 2401 | CTTTAGCACCGAGCTGATGGGTGACACCTCCTCGCTGAGCCAGGGTAGCCTGAGCCGCACTGTGCGTGACGATGCCATCAACCAGCTGGAGCTCTGCCGGGTTGACACCCTCAACCCCCG | 4 |
| wt-117390-117510 | 2402 | AGTAGCCGGACGCCTAGCCTCCTCCCTCTACGTGTACGTTGATCCGGCCTATACCAACAACACATCCGCATCAGGCACCGGAATCGCCGCCGTGACTCACGACAGGGCGGACCCTAACAG | 4 |
| wt-117510-117630 | 2403 | GGTCATCGTCCTGGGCCTGGAACACTTCTTCCTCAAGGACCTAACAGGGGACGCTGCCCTCCAGATCGCCACCTGCGTCGTGGCCCTCGTCTCCTCGATCGTCACCCTGCACCCCCACTT | 4 |
| wt-117630-117750 | 2404 | GGAGGAGGTGAAGGTAGCCGTGGAGGGCAACAGCAGTCAGGACTCTGCGGTGGCCATTGCCTCAATCATTGGGGAATCCTGCCCCCTCCCCTGCGCCTTCGTGCACACCAAGGACAAGAC | 4 |
| wt-117750-117870 | 2405 | GTCCAGCCTGCAGTGGCCCATGTACCTCCTGACTAATGAGAAGTCCAAGGCCTTTGAGAGGCTCATCTACGCAGTGAACACGGCCAGCCTTTCTGCCAGTCAGGTCACCGTCTCCAACAC | 4 |
| wt-117870-117990 | 2406 | CATCCAGCTCTCCTTCGATCCGGTCCTCTATCTCATCTCCCAGATCAGGGCCATCAAGCCCATCCCTCTCCGCGACGGTACCTACACCTACACCGGCAAGCAGCGCAACCTCTCTGACGA | 4 |
| wt-117990-118110 | 2407 | CGTGCTGGTTGCGCTAGTCATGGCTCATTTTCTCGCAACAACACAGAAGCACACGTTCAAGAAAGTTCATTAAACTTTATTGACTACACCAGTCCCTTGTAAAGCGACGGGTCTCGCGTG | 4 |
| wt-118110-118230 | 2408 | ACGGCATTCGTGAGCAGGGCTTCGTCCAGGGGCTTGTTCTTGGCGGACATCATTAGCCCAGCCGCAAATATCAGAATTAGCATCAGAAAAGTGAGCCCCACAAACACCAGTGTCCAGAGA | 4 |
| wt-118230-118350 | 2409 | GGAAGACCGTAAGATAAAGATGGCTGCCTCTCATCTGGAACGGTGGGAAGCTCAGCAGTTGTTTTTGTGGCATTGGACGTCCCTTTGGAGGACAGCGTGGGGGCCAAGGTGGTAGCGTTG | 4 |
| wt-118350-118470 | 2410 | GTAATACGGGTAGTAGCACTGGTGGTGGAGGAGGACCTGGTGGTGACATTGCTAGTCACACCCGTGGAGGTTCCTGTTCCGGCCTCGGTGGCAGTGATGTTCTGTGCAGTAACCTTAGTG | 4 |
| wt-118470-118590 | 2411 | GTGACATTGATGGTGGATGCGTTGGAAGTTGTTGGGACTGGTGTGACAGTTGTCCCAGTGAATGTCACCGTGGTTGTGTTGGTGCTCAGAATAGCAGTTGTGGTTATAGGGGCGCTAGTC | 4 |
| wt-118590-118710 | 2412 | GTGGTCAAGGTCGTAGACTGGTTTGTGCTAGGACCCGATGCCGACGGTGATGGTGTAGTCACAGCCGTTGTGCCTGTCACGTTCCCCGCCGAGGCCGTCGAACTGCCACTAGATGTCCAA | 4 |
| wt-118710-118830 | 2413 | ATAAGGCTTGTCTCACAGATGAGTATCATGGCCATAACAGCGCCTGCCTTGTCTCTGGCGTGTGCCATCGCGTCTGGACGCAGAAGGCCTCCCGGCCTCTTTTATAGCTAGTCTCCACAC | 4 |
| wt-118830-118950 | 2414 | CCAATACTCTACTGAACCATCACATACATGACCTCCTCGAGGTATGCAGGGAATGAGCGGTCCGTGAGCCGGTCAACACGACATTGCTTCCGTTTCATGCCTCCAGCTGCCCCTGACCAG | 4 |
| wt-118950-119070 | 2415 | TTAGGACCCTTGACGGATGTCTTTAACGGCGCGGTGCAGTTGGTCACCAATGACGGCCTAAAGGCCAACACATCCTTGAAGCAGGGCGTAGGAATGGTACCAAACTCGGGGCCCACCCCA | 4 |
| wt-119070-119190 | 2416 | TCAAAGACATAATATGTCTCATAGTGGCAGTGATGATGCATCACCACCACAGCACTCGCCAGGACCCTCTGCATATCTTGTACAAGGCGCCTTTCAACTCGGCCACTGGCTCTGGTGACG | 4 |
| wt-119190-119310 | 2417 | TTAAATGTCCTGTTCCTATTAGTCACAGCCTGTAGATTTGGGCACCCAGACTCAAAAAGTGCAGCTACATGAAGGGCAGCCGCCTCAAATCCACCATGACCCCCATGGCTGTCCGTGTTG | 4 |
| wt-119310-119430 | 2418 | TTGGGGTAATAAGTCACATTGTTAATGACCACGGCCGGGATAAGGGTGTAAACCTTGCAGAATGGATTGGTCGGACACCCATAAGACAGGGGCACCCCAAAATCACGCCCCTTACCCCGA | 4 |
| wt-119430-119550 | 2419 | AGCACCTTGGCCCCCACCGGCATAAAGCTGGGCAAAAAGAGTGGGTTAAAACCAAAGGCGAGTAGGGCCAGGAACGCCAAATAGCAGCAGTAATAGATGAAAACAAAGCTCAGCATGAAA | 4 |
| wt-119550-119670 | 2420 | CAGCGTGGAGGCTCAGCTAGGGTCTCTGCCTCTCCATCATAGACATCTTCCTTGAATCTCATTCTCTCACCGCATACCTCGCTCTTCATCCAGGAGGGGGCCATGGCTGCCATTCTACCA | 4 |
| wt-119670-119790 | 2421 | GTTAACGAGGAGAGAGAGAGAGTAGGTCCGCGGAAATTGGTGCCCCTCTCTGCCCTCCTGACGAGGCCATGGTGTCATCCATCTCCGCAGTCCGTTCTTCAGCTTTGGCATTGGTCCGGGTC | 4 |
| wt-119790-119910 | 2422 | CGGGTGGTCTGATTTTGATTCTGATCCTGGGTATTGGTCTTGGTCTCTCCTCCCCCATTGGCATGGATTGGCATAGGTGGGTGTGGCTCAGGCTCAGGTTCCGGCCCTGGGACGGCAGCA | 4 |
| wt-119910-120030 | 2423 | GCCGCCGGGACGGTGAAGTCGTGGAAGGTAGAGGCCCGTCCCTCCCGAGGTCGTGGGGCCGGAGCCTTATAAAAGACTTCCACCCTCTCCCCGCTGGCCAAGACACGCCGCTCGTGGACC | 8 |
| wt-120030-120150 | 2424 | ACGCCATCTTCCTCCCGGCTGATTGTGTGGCTGACGGTGCCGTGTTCCACCGCCACTTGTTCATCGACCATGGTACCCCCTTTATCTTAACCAGCAAGTGGCCGTCAGGGTCTCTTGAGA | 4 |
| wt-120150-120270 | 2425 | GTATGCCGCTGTGGCCAAGCGAGGCCCCAAATTAAATAGTGATGCCAAAGACTGTAGGTAGGTCATCATCACACGCATGCGTGATAAATCATCCGCCACTGACAGGTCATCCAGGTCTAT | 4 |
| wt-120270-120390 | 2426 | CCGGGCTATCTCATCCGGCACCATTTCCTGGAAGAGATTCAAGAGGTCGTGATGCTCATGCCGGATAAGGCCTCGGACCAGGCGCATACTGGCCCTGGGCAGCAGGGTCACCATGATGCA | 4 |
| wt-120390-120510 | 2427 | AAAGTAGAGACTCAGATTGTCCAGCAGGGCCAAGCCAAGGGGCCCTGGCACCTCCGGGAGGGCCAACTCGTAGTGGTGCCCCAGGTATGAAACAGAGCCAAGATGCATGTGTACATCGAG | 4 |
| wt-120510-120630 | 2428 | CATGTCTGCGTTCCCGGGAGCCTGCATGACAACCCGGGAGTACACGTTAAACAGGAGAATCTTCTGCAGCACCTCCTCTGCTATGGGCGTAGGCAGCACCATGGGGAAAACAATGTCCAC | 4 |
| wt-120630-120750 | 2429 | ATCATTGGACTCTAACTTCACGGTGGCATGCTCTCGTCCAAATACCGGGGGCATAACACTGAGGCTCCCGGTCCCATGCCACTGGAAAAAGGGCTGGTACTTGTTCTTAATGGCGTAGGT | 4 |
| wt-120750-120870 | 2430 | CTGACCTGGAACAATCTTGGTGAGTATCAAACTGTCCACGCTAACCTCATCCAGCACGGCCAGGGTGCAATCAGACAGGTAGTTGTACATGGACACGTAGTCCGGGACCGTCTCTAGAGA | 4 |
| wt-120870-120990 | 2431 | GTACACCTGACCCAAGCCCAATCCCTGCACATTCTGCGTCCCGTGAGTGGAAGCCAGGGGTAAGATGCAGCCAATCCTCTGTTGCATCTTGGCAATCTCATCGGTATACAGACGAGAGGA | 4 |
| wt-120990-121110 | 2432 | GAGAGACACTACCACTTTCAAATCCATCTTTATTGACAATTATCAAAAAACCACCTTATTTCCAAACTTTAATATTCTTCGTACCGGCGCCACCTCTTCAATTATATAGTGTCCGTAATG | 4 |
| wt-121110-121230 | 2433 | GATGGGGGCGTGGGTCTGTTTGACAGACATAAACTCATCGATGAGTGCCCGGGAGGAGGCTGAGAGTGCGGGGAATGCCTCCTGCAGAAAGCTGCAGGGCTGCTCCAGAAACACGTCAGT | 4 |
| wt-121230-121350 | 2434 | GCCAGCAATCACTACAAACTGCACCTCTGTGTTGCTGGTGGCTGGGTGCCCTCCAAGTCGCTGGCTGTACTCGTTGACCATGTTGTAGAGTCCCCTGTTGTTGCGCAGAAGCTCCTCCTT | 4 |
| wt-121350-121470 | 2435 | GTTGAAAAATGCCCGGCAGGGGCTGTAGAGGCCCGGGACGGCCGTCTGGCGATAGGAGGAGTTGTACATGATGTCACCCAGAGAACCCAGCTGAGATGCCCAGGGATTCACAGTGCTCCG | 4 |
| wt-121470-121590 | 2436 | GTATTCATAGGCGGCATCCGGGCGAGAATGGTCATAGATGAGCCCCTCGGCAACCTCCTGATTGTAGTTTTCACAGGAGACCACACAGGCGGCCCGTCCCCTTGGAGAGTTGGACTTTTG | 4 |
| wt-121590-121710 | 2437 | AAAATAAGCCACGTCTGCCGTGACCGGTGTTACGATAATCTCACAGGTGGCCTGCTGGCCGTGGCAGAGTCCTGGAGCTCCATTAACATTAGTCATACCTGCCAGGTATGTCCTGGGGTC | 4 |
| wt-121710-121830 | 2438 | CCGAAGCAGCGTCCCATTGCGCTGAGCGCCCACCTTGGCCTTGATGTAGTCATTGACTTGCTGGTTGCCAAAGGCCTCGGCCGGAAAGACGCTAAAGAAGTCTTGGGTGTGGATACCCAT | 4 |
| wt-121830-121950 | 2439 | GTCAGTAGTGATGGCCGCCACCCTGGCCGGAGTCATGGTCGAGCTATAACTAAGCCCGGTGTCGATGGAGGCCATCTCGTGATGCACCTCAAAGGTTACCGCGTCCACCCTGGCCTCCCG | 4 |
| wt-121950-122070 | 2440 | GCGGCTAACATTTGGGGTCCCAATGAACATGGATGTTGAGGCCCTGGAGCTAAACAATATGTTTTCAGAGAGGATCTCATCGGTCCTGACCACGGTCATGGCCACCCCTGGGTGGATCTT | 4 |
| wt-122070-122190 | 2441 | GAGCTTGGCCTGGGCAATATAGGCCATGGGGGACATCTTGATGTGCATGGCGGTCATTCCACTGATTGAAACGAGGGAAGGAAGACATTCGGCCGCGTATTTGCCCATGGGCGAGCGGTG | 4 |
| wt-122190-122310 | 2442 | CCACTCCCGGTACTCTGCAAAGAGCTGCTCTGGCCGGTTGAAGGCTTCCACGGCCCGCTGCTGAGGATTGCGCATAACAAAGGTGGCAACATCCTGGTGCATGGTGGCAGCCACTCGCGG | 4 |
| wt-122310-122430 | 2443 | GTCCCCGTAAAACATATGGAAAGGAATGGCGTGAAAGAGACACTGGGTGACGGCCCGGGTCCTCTCGGAGAAGGCAAAGGCCACCAGCCCGTTCACCAAAACAGTCTGCTCTGTCCGCTT | 4 |
| wt-122430-122550 | 2444 | GTCGGCGGGATTCGGGGCCAGCTGCTGCGTAACGTCATTGTCCACCGACACACGCACGGCACGGGTGAAAGTGGGGCAGGTCATGAATGAGGCGCTGAGGTCCCTGATCATGCCCACGGT | 4 |
| wt-122550-122670 | 2445 | GGGGCGGAGGTCGGAGATCTCCAGCAGATCCCTGAGCGTCCCATTCTCCAAATTGTCGAGGATGTCCTCGTCCCTGGTAAAATGGTGGCTGAAGGCTGGCCCGTTGTAGGCCAGGGTCTG | 4 |
| wt-122670-122790 | 2446 | GGCCACGTGCTGAAAGTCCACCCCGAGGCCGCACATGTGGGCATTGGTGCAGGTTGGGAGGAAAACGTAGTAAAAGATCTTTTCCAGCACATCCGCATGCCCCTCATCTACATAAGGGCC | 4 |
| wt-122790-122910 | 2447 | TAGGTGCAGACGGAAATCGTGGTCGTGGTCTCCGTTAACCCGGTAGCCGTACAAGGCCACAAATTGGGCAGCCATCTCATCCATGTTTCCAACCCTCTCAATAAACTGGGGCGCGGCCAG | 4 |
| wt-122910-123030 | 2448 | GGTGTCAGCGTAAACCTCATTTCCGATAATAATCTGGGGGGCCCGGTCACTAACGGTGAGAAGATGGGTGAAAATGTCTGTGTAGGCCACCGGGGGGAGCAGGTTAGGGTCCAGGAGAGC | 4 |
| wt-123030-123150 | 2449 | GCAGACATACTGACCCACGCTCTCATCCCCCACAACATCTGACCCGGCCAGGCGCATCAGGGCCTGCTCTAGGGCTATAAGTTCCCCATAGATTTTTCTATACATGGAATAGGCCTCCTT | 4 |
| wt-123150-123270 | 2450 | GGAGATGGCGTTATTTCCCAGGTGGCGGCAGATGAACTTGATCATGGAAAAGCTGTTCACAAAGGCAAGCCTCCCTGACCGTTCCCAGTAGGTGTTGATGCACAGGGACACCAAAGGCAC | 4 |

| | | | |
|---|---|---|---|
| wt-123270-123390 | 2451 | GTTCATGACAAACTTTTCCTCAAACCCGTGGATCATAGCCTCGACTACGTAGAAGAAGGCTGGATAGGCAGTGTCATAGGCAGTATCCTGCACAGTCTCAATAACGGCCTGATCCACCAC | 4 |
| wt-123390-123510 | 2452 | GTGGGCCAGAGATGTGGCGGTCTCAAACTGCTGCCCCCGGGCCTCTTGGAATGCAGCTGGGGCCAGGGGAGTCGGCAGGTTACCCACCATTAGCCGGTGCACAGCCCTGTGCCTGGCCCT | 8 |
| wt-123510-123630 | 2453 | CTCCCCGGCATCCCTGCCAATGTAAATATCATAAAGGGGGTGCAGCTCCAGCCGCAGCAGGTCATAATTGGACGGGTGGAGGAAGTCTTCGGTGGGCAGCCCGCACTTGAGAGCTATATC | 4 |
| wt-123630-123750 | 2454 | TGTCACGGGGGCTGCATACTTGTTATCATAGAACTCGTCCACAATAACAAGCACATTCATGTGATTGGGCCTCCTGTGTTGCAGGGAGTAGGTCTCGCGCCTGTCTCGCGGGGCCGGGGC | 4 |
| wt-123750-123870 | 2455 | CGCGTTGAGGCTGTTTAGGGTATGGGCGGGTGTGTGGAGTCGGGGGTGACAGAGAACCTTGAGAGCATTCTGTAGGTTAAACGCGAGGAGAAGGTTATTCTTGTTTACGATCCATGCCTC | 4 |
| wt-123870-123990 | 2456 | CACCGGTAGCTGCTGTGTGGGGTTGTCCAGCATTTTGATGGCGGCGGAGGTCGTGTACTTGGGATTGGGCATAAACAGGCCCACTGGGAAATAGTAGCTGTACTGCATTCTTCTGTTGAG | 4 |
| wt-123990-124110 | 2457 | GGGGTATGGGGACTGAGTGTCATTGTACATCTTTTGCAGGCTTTCCACGGCCACCGCGTGGTTGCCCAGCTTGATGACGGCGGCTGAGATCGGCACCCGGGGCTGATCCTCGACCCCTGC | 4 |
| wt-124110-124230 | 2458 | GGCCACAGCCGGCAGGTCAGACTTGGTGCTTCCGGCTTTTTCCGGTGAGTCCACGATCCTAGCCATGAAATGCTCAAACGTACGCATCACGCGCCCGTAGCTCACGGCAGTGACCAGGTT | 4 |
| wt-124230-124350 | 2459 | CTCCCCCCGTACCACAAAAGAAGCATAGCTCGAGGGCCCCATAATCTGGTTGTCGGCCTCCTCACCCAGGAAGGTCAAGAGCTGGCGCAGAACGTTGTCGGTGACAATAAACACCCCCCC | 4 |
| wt-124350-124470 | 2460 | CACTGGCTCTCCCCCCTTGGCGGTCGTGTAGGTACTGACCCCCTTGAGCACGCTCTCCCCGGACACGGCCGCTACCATCTCAGAGAGACGGCTTCGCACGTACTGAGAAAACCCGGAGCC | 4 |
| wt-124470-124590 | 2461 | CATGTTCTCGGCCCGGTCCAGGAAGAAGGAGTGCTCCAGCAGATGCCTCTTGAACATGGCAATGAGGTCAGACTTGACAGTCTTGGAGAACCCCCTCTCAGTGAAGGTGGGATCCGCCAG | 4 |
| wt-124590-124710 | 2462 | GGTCTGCAGGATAAACATGGGAGGGGCATGGCGAAGCTTCACACTCAGGACGGTGTTAATGAGGCCCCTCTCCAGGGCATCGACCCCAAACTGTAGGGCCGAGGCCACGGTCTTGACAGC | 4 |
| wt-124710-124830 | 2463 | CCCCACGTACTCTGCGTACTCGACCGGGGTCTCGGGGATACTATGCAGGATCTCCAGATCCAGCATGGACAGTTCCATTTCCGTACTAATGTGGTGTTTGTGGCAATTTTTGACCACAAT | 4 |
| wt-124830-124950 | 2464 | GAATGTCCGCTGCTTGCTGGGTCTCCTTCCGTCCCCGTGAGCAATGGTGGGGACGGAGATTCGAAATTGAATCTTGCCATCCGTCATACGACTCAGGTCTTTGAATTCCGTGTTCACACA | 4 |
| wt-124950-125070 | 2465 | GGACACGGCCAGTGCCGTCTCCAGGAAGCGAACATATTGGATGGCGTTCGTGTAGACCCCGAGTAGCACCTCAAACTTGATGCCCGCCTCTCTGGCATCCTTGCCCACCAGCAGGTCAAA | 4 |
| wt-125070-125190 | 2466 | GCTATGAAACAACCCCTCAGCCGCTGACTGCCGCAGGTTCGAGAGCAGGTCGGCATCCACCGTCAGATAGGGGAAGGGTCTGTTTTCCACACCCTCATTTGAGGCCATGACACAAGGTAA | 4 |
| wt-125190-125310 | 2467 | GAGGGAGATGGGGGGAGGTCTCGAGGGCTTCTCTTCACAGCTGGGTCTCTTTTACGCCCTGGCCTGCAACCGCAGCCCACCCACACTTCCCGAGGATGCTACCCTTCTAATCAAATGGTT | 4 |
| wt-125310-125430 | 2468 | GGACACGGCCCTGGGCAGGGAGGCCACCTTTTACGCGTGTCGGGCTATGCGTCGGCTTCTACTCGGCGTTATCCGAATGAATGACTGCCAGGAGCTGCCACCCGGTTTAATAATTCTGAG | 4 |
| wt-125430-125550 | 2469 | TCCGGGCACCGTCCCTGGCCCCCTTGGAGTCCAGAGTCTGGAGCATACAGACTGCGAAATATGGTCCTCTGCCCACCCTGACCACGCTGCCCACCTCCCGGTGCCCAGGGTCATCACATA | 4 |
| wt-125550-125670 | 2470 | CACCGACTGCCCGGGTTCCATAAACACGAGCTCAATGTTTCGCCTTATCATCCGCTACTTGTCTCATCACCAATTTGAGCGCTGCTTCGAGCAGTTCTGCCGCGTGGTCCCGCGTCGCTT | 4 |
| wt-125670-125790 | 2471 | CCTAGGGACCTGTAAGCGAAACTCTGCAAAGATGCTGGCTCATCTGAATCAGGTTACCAGGATCCCCCCCTGTCCGCCCTTCAGCGGGCGGGAGGCCAGACTCAAGTTCCACTTCTTCTC | 4 |
| wt-125790-125910 | 2472 | CTGGAGCACATTCATGCTGTCATGGCCAAACAATGCCACACTCCGGGAGATCAGGACGAGGGCCGCCACCAACCTCACCCACCACCCACATCTAGTGGATACTCTGTACCACGCCTCTCC | 4 |
| wt-125910-126030 | 2473 | GCAGACCCCATTTCTGACACGCAGCGGTGCTCTATACCGCTTCGTCACCTGTTGCAACTGCACCCTGCCCAATATCTCCATCCAGCAGTGCAAGGCCGGGGACAGACCGGGGGACCTGGA | 4 |
| wt-126030-126150 | 2474 | GATCATTCTACAGAGTAACGGCGGAGGGAGGCCCGCGAGCTTCCAGTTCCCCTCCTCCCCAACTGGCTCCCTATTGCGATGCATAGTTGCTGCGTCCCTGCTGCCGGAGGTGTCCGTGGG | 4 |
| wt-126150-126270 | 2475 | GCACCAGGAGCTGTCTCCGCTGCGGTCCAGAAGCCAGGGAGGGCAGACGGATGTCAGGTCGGGCCCGGACCCGGCCCGGAGACTGGTGGCCCTCCTGCGAAGGGAAGATGGGGCACCTAA | 8 |
| wt-126270-126390 | 2476 | AGACCCCCCTCTGGGACCGTTTGGACACCCCCGGGGGCCCGGCCCGGCCAAGAGCGAAGACGAGGAGTCTGAGCGTCGAGACGCCCCTCCACCCCCGCTCGATTCCAGCTTCCAAGCTTC | 8 |
| wt-126390-126510 | 2477 | CCGGTTGGTGCCCGTGGGGCCTGGGTTTCGCCTGCTCGTGTTCAACACCAATCGGGTGATCAACACTAAATTGGTGTGCTCAGAGCCCCTGGTGAAGATGCGAGTTTGCAATGTCCCCCG | 4 |
| wt-126510-126630 | 2478 | CCTCATCAACAACTTTGTAGCCCGCAAGTACGTGGTGAAAGAGACGGCGTTCACCGTCAGTCTATTCTTTACGGACGGGGTGGGGGCCAACCTAGCCATCAATGTCAATATCAGTGGCAC | 4 |
| wt-126630-126750 | 2479 | CTATCTGAGCTTCCTATTGGCCATGACGTCACTGCGGTGCTTCCTGCCTGTGGAGGCTATTTATCCCGCGGCCGTGTCAAACTGGAACTCGACTCTAGATCTCCATGGGCTGGAAAATCA | 4 |
| wt-126750-126870 | 2480 | GAGCCTAGTCAGAGAGAACCGAAGCGGGGTCTTTTGGACTACCAACTTTCCCTCGGTGGTGTCCTGCCGGGACGGTCTCAACGTGTCCTGGTTTAAGGCCGCAACTGCCACCATATCTCG | 4 |
| wt-126870-126990 | 2481 | AGTGCACGGGCAGACATTGGAGCAGCACCTGATCCGTGAAATCACCCCCATCGTGACGCATCGAGAGGCAAAAATCTCCCGGATTAAAAACCGGCTCTTTACCCTGCTAGAGCTACGCAA | 4 |
| wt-126990-127110 | 2482 | TCGGAGTCAGATTCAAGTGCTGCACAAGCGTTTCCTGGAAGGCCTGCTAGACTGCGCCTCCCTCCTGCGCCTGGATCCCAGCTGTATCAACCGAATCGCCTCCGAGGGCCTGTTTGATTT | 4 |
| wt-127110-127230 | 2483 | CTCCAAGAGAAGCATCGCCCACTCCAAAAACCGACACGAGTGCGCGCTTCTGGGTCACAGACATTCGGCGAACGTGACAAAGCTGGTGGTAAACGAGCGCAAGACCCGCCTGGACATACT | 4 |
| wt-127230-127350 | 2484 | GGGCCGTAACGCTAACTTTTTAACGAGGTGTAAGCATCAGGTTAATCTAAGACAGTCACCTATTTTCCTGACCCTCCTGAGGCACATCCGCCGACGTCTGGGCCTGGGCCGTGCTTCCGT | 4 |
| wt-127350-127470 | 2485 | AAAACGAGAGATTACCCTTCTCCTGGCCCACCTGCGCAAAAAGACAGCCCCCATCCACTGCCGTGATGCTCAAGTGTAAGCAGCCCGGGGCCCGCTTCATTCACGGGGCCGTGCACCTGC | 4 |
| wt-127470-127590 | 2486 | CATCGGGACAGATTGTCTTCCACACCATCCACAGCCCCACTCTTGCCTCGGCGCTGGGACTGCCTGGGGAAAATGTACCCATCCCGGCCCTCTTCCGTGCCTCGGGCCTCAACGTCCGTG | 4 |
| wt-127590-127710 | 2487 | AGAGCCTACCCATGACCAACATGAGGGCACCGATCATCTCGCTGGCTCGCCTCATCCTGGCCCCCAACCCCTATATCCTAGAGGGACAGCTGACGGTGGGCATGACACAGGACAACGGCA | 4 |
| wt-127710-127830 | 2488 | TTCCCGTGCTTTTTGCCAGGCCTGTCATTGAGGTAAAAAGCGGGCCTGAGTCCAACATTAAAGCCTCCTCGCAACTTATGATAGCAGAAGACTCCTGCCTGAATCAGATCGCCCCCTTTT | 4 |
| wt-127830-127950 | 2489 | CCGCATCAGAGCACCCCGCCTTCTCCATGGTTGAGTCCGTAAAACGAGTCCGGGTCGATGAGGGAGCAAACACCCGGCGCACCATCCGGGATATTCTGGAGATCCCCGTGACTGTGCTCT | 4 |
| wt-127950-128070 | 2490 | CATCCCTGCAACTGTCTCCCACCAAGTCCATCCTGAAAAAGGCACCGGAGCCCCCACCTCCGGAGCCCCAAGCCACCTTCGATGCCACCCCCTATGCCCGCATCTTTTACGACATCGGGC | 4 |
| wt-128070-128190 | 2491 | GACAGGTGCCCAAGCTGGGCAATGCCCCCGCCGCGCAGGTCAGCAACGTGCTCATCGCCAACCGCTCCCACAACTCTCTAAGGCTGGTGCCCAATCCGGACTTGCTGCCTCTCCAGCATT | 4 |
| wt-128190-128310 | 2492 | TGTACCTCAAGCACGTAGTGCTAAAGAGTCTGAATCTGGAGAATATAGTGCAGGACTTTGAGGCCATCTTCACCTCCCCGTCTGATACCATCAGTGAGGCTGAAACCAAGGCCTTTGAGA | 4 |
| wt-128310-128430 | 2493 | AGCTGGTGGAGCAAGCCAAAAACACCGTAGAGAACATAGTCTTTTGCCTCAACAGCATCTGTTCCACCTCTACACTCCCAGATGTCGTCCCCGATGTCAATAACCCAAACATTAGCCTGG | 4 |
| wt-128430-128550 | 2494 | CTCTAGAGAAGTATTTTCTCATGTTCCCTCCCTCAGGCACCATTATGAGAAATGTCAGATTCGCCACCCCCATCGTCCGGCTCTTGTGCCAAGGGGCTGAGCTTGGCACCATGGCACAGT | 4 |
| wt-128550-128670 | 2495 | TTCTAGGAAAGTACATCAAGGTCAAGAAGGAAACTGGAATGTACACACTGGTCAAGCTTTATTACCTGCTGCGCATCTAAAGGAAAAACATAACAATCTTGTGAACCAGAAAGATACCCA | 4 |
| wt-128670-128790 | 2496 | GAGCAAAAGCAATAAAGTACAGGATTATTGCCAAAACAACGTGTGCTCTTTCTTCATACAGGCCCGCAATTTCCATGACAGTCCCGTTGGTGGTCAGCAGCAGATAGTGAACGTGGAGGT | 4 |
| wt-128790-128910 | 2497 | TGTCAAAATCAAAGTAGTTGGAGCTCAAGATGGAGTTTTGGACTTCCTGGGAGGTGATGTAGGTTGTAGTTTCCAGGCCTTCCTTTTCATCATAACTGAGCAGGGCAAAGCCACAAAAAA | 4 |
| wt-128910-129030 | 2498 | TGCAGGATTTCTGCGTCCTGGTAAAATTCTGGATCTTTGGAATCTGGCGGGGCTCCCCAGCCACAGCACCCTGCGAACATTTATTCATTATAACGGGGGAGAGAAAGAGAGAGCTGCTGA | 4 |
| wt-129030-129150 | 2499 | GATAGGTGGTGCTGGCCTCGTATAGCGCCGAGCCTCGGACCTCACGGTCACTAGAGATTATGAATGTCACATTGATGAGCGGGATAATCATCAGAACTTTGTCGAGCCTGTCCACGCATT | 4 |
| wt-129150-129270 | 2500 | TGTAGGCGGGGAGATGCCACGCATCCCTGTCTTCTCGCTCCAAAGAGAGCCGCCCAAGAAACCCATCCACAGCATTTGAAACGGCCGCCTGGTCCAGCGTTGCCTCCTGGGGGGCCATGC | 4 |
| wt-129270-129390 | 2501 | TCAGCAGCTTGTCTCGTGTGAGGTCAAATCGTAGGCTGAGGTAGCACGGTGAGAAGAGCCCGCTCTCTGTCCCCAGGGCTAGCCCCCGCAAAACCTCCCCAATCTCTAGGGCCGAGCACA | 4 |

| | | | |
|---|---|---|---|
| wt-129390-129510 | 2502 | GGGCGGTGGACAGCAGTTGGTATAGGGCAAGGTTGGGCCCCTGGGTAGTCACGTTCAGCCGCAACTCGCGTAGCACCACGTGGCTGCCGATAAACAGGGTCTCTCTCATCACGGTATGCA | 4 |
| wt-129510-129630 | 2503 | GGGGCTGGAAAAGGGGGTGGCGGTTGTAGGCCGAGAGAAGCACAGATGTGGCGCCTCCAATGAGGCCACTGTAAACCCCGGCCTTGGGGTAGCCGACGGTGGCTAACCTCAGCGCGTACT | 4 |
| wt-129630-129750 | 2504 | CCTGTTTCTCAGTAGTCAGGTGACCCAGCTCCTCCATCTTGACCGTGGCCATCAGCATGGCGGCCAAGCGCTCCAGCCCGTAGGATTGCATGCCCTTGACAGTGGCCCCATAACATATGC | 4 |
| wt-129750-129870 | 2505 | CGATGATGTCTTTCAGGACAGTCAGCTCAAAGAAGCTCTTGGCCAACCAGCGGAGGTCCACGCAGCCATTGCCAGTCTCACCCACAGCATGACCCACCTTAAAGAAGGCCACAGAAACCT | 4 |
| wt-129870-129990 | 2506 | CAAACATGGTAGTCAGCGTTTCCGTGTCCAGTTCCGGCTCCCGGCAGCCTCCCTTCATCTCCAGCAGGACCAGTTTCTGGAGAACGTAGCGAGCGTAGCTGGCGGCTGTCATGGTGACGG | 4 |
| wt-129990-130110 | 2507 | CTCGGGAAAACATATCCTTCAGGTTGGGTACAAAGTAGTTGTGAAAGTTGGCATAATGCACAAAGGTTGTAACAATCACCAGGGAATAGTCCCCGCTTTGGGCACTGGTTAAGGATGGGT | 4 |
| wt-130110-130230 | 2508 | AACTAAAAGGCCCCCTCAGATCCGGCAGGTCCTTCGTCTTGCCAAAGACCAGGCTCAACACATGCTCATCTCCCTTCTCGGTCACTCGCTTGTAGGTGCCCATCAGAAATTTAGAAGTCA | 4 |
| wt-130230-130350 | 2509 | TGGCCCCCGTGTACTGAAACTTGTCCCCGTTGATGGACAGGGCCACATAAGACAAGTGACAGCGCAGCTGATAAAAGACATAGCTGTGTGGCCGCGTGTTGGGCAGCATGGTGCCAATAT | 4 |
| wt-130350-130470 | 2510 | AGTAGAAGAGCTGCTTCTCAAGGGGGGCACTAAGCATGCAGGCAGGGGAATTCAGGCCGCTAATGACTCCGGGATGGACCTTAGATGCATCCACTTGCATGGATCCTTCAGAGACAGCAG | 4 |
| wt-130470-130590 | 2511 | GGATATCGACAGGCTCGGCCAGCGCAATACCAAGGGTACCAGACGTCTTGTAAATTAACTTGTAGCGGTTAAGCATAGACGCCAAATCTTCGGTGACATTTGCCTCTCTCCACAGCGCCT | 4 |
| wt-130590-130710 | 2512 | CTGGGCTAAGGCCTGGGACCTTTGCCATCAGTTCGGTCCATGGGATGGTGTAATGCGAAGCATGCCCCTCTATGTCCAGGTGCAGCTTAACCTCGCTGAGGCTGGCAGCCCCCACCTCCC | 4 |
| wt-130710-130830 | 2513 | ATAGCAACACCAGGCAAAAAACACAGAGCAACTGCATCCTAGTCCCGATTTCCCCTCTCAAAATCAGAGATCACCTTGCTCAGACCAGCCCAATCGAAAAACTGAGATCGTATTGCCGGA | 4 |
| wt-130830-130950 | 2514 | TTCTTCAATGCCTGCATGTAAATCTCCGTCCAGCATCCAGGTAAATCGTCCTGAAACTCTGAGAGGTCCACAAGCACAAACTGAAGGTAGGCTAGCGTTCGGGTGAACGCAAGACAAACT | 4 |
| wt-130950-131070 | 2515 | TCCAACAACACCGCGTCGGCTCGGAAAGGCTGTATGACTTCCTTAAGTACACTAAAGATGCTGTTCTTATACAGCTTCTCGGCCACACCACTTCGAATTATGGGGGTGTGGCTTTGATGA | 4 |
| wt-131070-131190 | 2516 | CATACTGTCGTGATTGTTGTTAGACCGGCACATACCTTCACAATGTCCTCGGGGGCAAAATACTGTGTTAGGAGCCAGGCACAGTAAACGGCGTGATATGCATCGTTGACACTCTTCAGG | 4 |
| wt-131190-131310 | 2517 | TAGCCAGCATCCAGTCCTGACTCATGTTTCCTCCCTCGCTTCTTCAGGCGGCGCATGTTCTCCTCCACGTTTAACTTCATCCAGACTATGGTGTCCCCCGGGTCTGCGGTAAACGTGGCC | 4 |
| wt-131310-131430 | 2518 | AAAACTTGAATAAAGTCACTATAGGAGAGAAGCTGGCTCCGGAGCAGCATTAGAGGGAAAACCACGGAGGCCGACAGCAAATGGCGATCATGCAAAATCCAACAATCCAGGGGCGCGACT | 4 |
| wt-131430-131550 | 2519 | GACCTGGCACCAGACTCGGTAACCAGCAAGCTCCGCTTCCTAGAGGCCAAGACTCTGAAAGGGGTGGTAAATTTCATCTGGCATGCTAAAACCTCAGCCGACGTGTCTTCCCTTCCATGC | 4 |
| wt-131550-131670 | 2520 | CTCGCCCGAGTCACATTCTTGTGCATGGCCTTAATGGCATTTTCATACACATGAGTCCAGTACCGCATCGGTTCAGGGACTACAATGGTCAGGTCCCCAAAGACAGCCTTCAAATGATTC | 4 |
| wt-131670-131790 | 2521 | AGCATAGTAGTCTTTCCCACACCAGGGGCACCTTCCAAAAATAGGGAACAGGCAGGTTTGATTACTGGTACATGATTTGTTAGGTGGGTCACAATTGGAACCCGCATGCTCTCCTTCCTC | 4 |
| wt-131790-131910 | 2522 | TGAGCCTTGGCCTGGCGGGTGTCTTGGGCATCATCCAGATTCAGAACATTCATCACACTCCCACTTAGCCGCTTCAGCTGGGCAGCATGCTTGGATAACTTACTAAACTCGCGCCCATGG | 4 |
| wt-131910-132030 | 2523 | GCGGCCAGGTGTTCGAAGAGACCAGAAGGCTTACCCTTGCCACCATTCTTTTGTTTTAACGCGGAATGAGAAGAGGGCCTGCGGAAATTAGACTCATCCTCAGACTCACAGTCAGATTTG | 4 |
| wt-132030-132150 | 2524 | TCATCGAGCCCAAGGCCGGCCAGGCCCTCCTCAAAGCCTTTCTGGTACATGAAGCTCCGGCTCGTGGAGTCCGCACCTCCTTCTGTGCACGAAGTTTTGCGGAACCAGGAGAAGGGGTCT | 4 |
| wt-132150-132270 | 2525 | GGCGTCTTGCTGGGGCCACACTCCCGGCTACGGGGCTTCGGGGTAGGGGCAGTAGGCTTTTGGTGTGCGGGTGCTGGTGGCTGGGCTCCCCTGGGCAGGGTAAAGGGGCACGATGTGTGC | 8 |
| wt-132270-132390 | 2526 | CGGCTACCCGGAGAGTTTCCAGTATTAGATGTCACGGCAGCCTGGGTCCGGCACGGCACCCTCTCCCCAGACAGTCCGGTCGGAGCCATCAAGGGGGGCCAGTGGGTGGGCACCTGGTAG | 8 |
| wt-132390-132510 | 2527 | AGGCCGTCGTCATCTTCCTCACCTGCCCCTGAGTCACTACCGGTTGGGGTAAGAACTGAGGGGGCAAAGTCATCAATCTCAGCGTAAAAGTTTTCGTGTCTTTCGTTTTCAGGGGACTCA | 4 |
| wt-132510-132630 | 2528 | TCCTCCTGACATTTTCGCCAGCCGCCGGGCGGGCCGGCCTCCTTTCCTGGAAATCCAGCCATGGATCCCACCCGGGGTCTGTGTGCCCTCTCCACACACGACCTGGCAAAATTTCACAGT | 4 |
| wt-132630-132750 | 2529 | CTTCCCCCGGCTAGAAAGGCGGCAGGTAAGCGAGCGCACCTTCGGTGTTACTCCAAGCTGCTCTCTCTTAAGAGCTGGGAGCAACTGGCCTCTTTTTTGTCTCTGCCCCCGGGACCCACG | 4 |
| wt-132750-132870 | 2530 | TTTACAGACTTTAGACTATTTTTCGAAGTCACCCTGGGTCGGAGAATCGCAGATTGCGTTGTGGTAGCTCTGCAGCCTTATCCCCGGTGTTATATTGTAGAATTTAAGACGGCCATGAGC | 4 |
| wt-132870-132990 | 2531 | AACACGGCCAACCCGCAAAGCGTTACTCGCAAGGCACAGAGGCTAGAGGGCACCGCCCAGTTGTGTGACTGTGCCAATTTTCTTCGCACGTCCTGCCCCCCCGTGCTGGGCAGTCAGGGC | 4 |
| wt-132990-133110 | 2532 | CTGGAAGTCTTGGCGGCGTTGGTATTTAAAAACCAGCGATCCCTGAGAACGCTCCAGGTAGAGTTTCCAGCCCTGGGCCAAAAGACCCTCCCCACCTCCACCACCGGCCTGCTAAACCTC | 4 |
| wt-133110-133230 | 2533 | CTCTCCCGCTGGCAGGATGGCGCTCTCCGGGCACGTCTTGATAGACCCCGCCCGACTGCCCAGGGACACAGGCCCCGAACTCATGTGGGCCCCAAGCCTTCGCAACTCACTGCGCGTGTC | 8 |
| wt-133230-133350 | 2534 | CCCCGAAGCGCTCGAGCTGGCAGAGCGGGAGGCCGAAAGGGCCAGGTCGGAGCGGTGGGACAGGTGTGCCCAGGTGCTCAAAAATAGGCTGCTCCGCGTGGAGCTGGACGGCATCATGCG | 8 |
| wt-133350-133470 | 2535 | TGACCACCTGGCCAGGGCGGAGGAGATCCGCCAGGACCTGGATGCTGTAGTGGCCTTCTCTGATGGCCTGGAGAGCATGCAGGTCAGGTCCCCCTCCACGGGAGGGCGCTCTGCGCCAGC | 8 |
| wt-133470-133590 | 2536 | CCCGCCCTCCCCATCCCCAGCCCAGCCGTTCACTCGGCTCACCGGGAACGCCCAGTATGCAGTCTCAATCTCTCCCACGGACCCCCCTCTGATGGTGGCCGGCAGCCTGGCTCAAACGCT | 8 |
| wt-133590-133710 | 2537 | GCTTGGTAATCTGTACGGGAACATCAACCAGTGGGTACCGTCCTTCGGACCCTGGTACAGGACCATGTCGGCTAATGCCATGCAGCGGCGCGTGTTCCCTAAGCAGCTGAGGGGCAACCT | 4 |
| wt-133710-133830 | 2538 | GAACTTTACCAACTCCGTCTCCCTAAAGCTGATGACAGAAGTGGTGGCGGTGCTTGAGGGCACCACCCAGGACTTTTTCTCAGACGTCAGGCACCTGCCAGACCTCCAGGCTGCCCTGAT | 4 |
| wt-133830-133950 | 2539 | CCTCTCGGTGGCCTACCTGCTACTCCAGGGGGGCTCCTCACACCAGCAGCGCCCCCTCCCTGCCTCACGGGAAGAGCTGCTGGAGCTGGGCCCGGAGAGCCTAGAGAAAATCATCGCCGA | 8 |
| wt-133950-134070 | 2540 | CCTCAAGGCCAAGTCACCCGGCGGAAATTTTATGATTTTAACAAGCGGAAACAAGGAAGCGCGCCAGTCAATAGCCCCTCTCAACCGACAGGCGGCATATCCACCCGGCACATTCGCGGA | 4 |
| wt-134070-134190 | 2541 | CAATAAGATTTACAACCTGTTTGTGGGAGCGGGACTACTGCCCACGACGGCCGCGCTGAACGTGCCCGGGGCGGCGGGTCGGGACCGGGACCTGGTGTACCGGATCGCCAACCAGATCTT | 4 |
| wt-134310-134430 | 2543 | CACCCGGCGTCTACACCTATCGTCCCTGCTCCCCCAGGCAATGCAGCGGCGCAAGCCTGCCATGGCGTCAGCTGGTATGCCGGGCGCCTATCCAGTCCAGACGCTTTTCCGCCACGGGGA | 8 |
| wt-134430-134550 | 2544 | GCTCTTCCGCTTCATCTGGGCCCACTACGTGAGGCCCACGGTGGCGGCAGACCCCCAGGCCTCCATCAGCTCTCTTTTCCCCGGGCTGGTTTTGCTGGCCCTGGAGCTGAAGTTGATGGA | 4 |
| wt-134550-134670 | 2545 | TGGGCAGGCTCCCTCCCATTATGCCATAAACCTGACCGGACAAAAGTTTGACACCCTCTTTGAGATTATCAACCAGAAGCTTTTATTTCACGACCCGGCTGCCATGCTGGCGGCGCGCAC | 4 |
| wt-134670-134790 | 2546 | ACAGCTGCGTCTAGCCTTCGAGGACGGCGTCGGTGTTGCCCTGGGGCGCCCCTCGCCCATGCTTGCGGCGCGGGAGATCCTGGAGCGTCAGTTCTCAGCCTCGGATGACTACGACCGGCT | 8 |
| wt-134790-134910 | 2547 | GTACTTCCTGACGCTGGGCTACCTGGCCTCCCCGGTGGCCCCAAGCTGAGCCAGTTCCTCGCACTGGAGTGGGTCATTGGCAAAAAGGTAAATAAACTCATCGCACGGGGGTTTTGCCTC | 4 |
| wt-135150-135270 | 2550 | GTTAGGGAGGACACTGCCAAGGTTGAAGAGCAGATTTCCGTCAGCCAGGGTGACCTGGCTCAGGTCCGGCGCCCTGCGCAGTCCAAGCTGCGCCCACACACATGCACAGACGGCCCCTGT | 4 |
| wt-135270-135390 | 2551 | GACATCAGGCCGGTCATGCAAAAACAGACAAAGAGACCGTGAGCGGTTACCGGGGCGCAGGGCCTCTGCCGGGAAGCCCACCCGGGCCAGGGCCCGGTAAAGCAGGTACCAGTATTCATC | 4 |
| wt-135390-135510 | 2552 | CGGCACCTTGCGTGCCAACACACGATTCGTGCGGTTTCCAGTATTTATCACGGCTTCCCGCCACAGGTAAAAGTTAACACTTAGGGTCAGCAGCTTGGTCAGGGATAGGTGCAAAAACCT | 4 |
| wt-135510-135630 | 2553 | GAGCTCGTCCTCGCGCAGAGCGCAAAGCGGCCAGTTCTTTAGCATCTTCAGGAGGAGCCCGTGAATCCCAGGTGTCATTCGCGCGTCATCCCCGCGCACCCCCAGTCCCATTAACATAGC | 4 |
| wt-135630-135750 | 2554 | GGGCACAATGGTGCAGGCACCGTCTGTATACGTCTGCGGCTTCGTGGAGCGCCCGGACGCCCCACCCAAGGACGCCTGCCTTCACCTGGATCCCCTCACCGTCAAGAGCCAGCTCCCTCT | 8 |
| wt-135750-135870 | 2555 | GAAGAAGCCCTTGCCACTCACGGTGGAACACCTGCCGGATGCTCCGGTCGGCTCAGTCTTTGGCCTTTACCAGAGCCGAGCGGGTCTCTTTAGCGCAGCCTCGATTACCTCTGGGGACTT | 4 |

| | | | |
|---|---|---|---|
| wt-135870-135990 | 2556 | CCTGTCCCTGCTGGACTCAATTTACCACGATTGCGATATTGCACAGAGTCAGCGCCTGCCCCTCCCTCGAGAACCCAAGGTGGAGGCTCTGCACGCCTGGCTCCCCTCACTGTCACTGGC | 4 |
| wt-135990-136110 | 2557 | CTCCCTCCACCCAGACATACCCCAAACCACCGCAGATGGAGGCAAGCTGTCCTTCTTTGACCACGTGTCTATCTGTGCCCTGGGTCGTCGGCGCGGCACCACGGCAGTCTACGGTACAGA | 4 |
| wt-136110-136230 | 2558 | CCTTGCGTGGGTCCTGAAGCACTTTAGTGACCTGGAACCGTCTATCGCCGCCCAGATTGAGAATGACGCCAATGCCGCAAAGCGTGAATCCGGATGCCCGGAAGACCACCCTCTGCCCCT | 4 |
| wt-136230-136350 | 2559 | CACGAAGCTCATAGCTAAGGCAATCGATGCTGGATTTCTGAGAAACCGCGTGGAGACTCTGAGGCAGGACAGGGGTGTGGCCAATATCCCAGCCGAGTCGTATTTAAAGGCCAGCGACGC | 4 |
| wt-136350-136470 | 2560 | CCCGGACCTACAAAAGCCGGACAAGGCACTTCAGAGCCCACCACCGGCCTCCACAGACCCAGCCACCATGCTATCAGGTAACGCAGGAGAAGGAGCAACAGCCTGCGGAGGTTCGGCCGC | 4 |
| wt-136470-136590 | 2561 | CGCGGGCCAGGACCTCATCAGCGTCCCCCGCAACACCTTTATGACACTGCTTCAGACCAACCTGGACAACAAACCGCCGAGGCAGACCCCGCTACCCTACGCGGCCCCGCTGCCCCCCTT | 8 |
| wt-136590-136710 | 2562 | TTCCCACCAGGCAATAGCCACCGCGCCTTCCTACGGTCCTGGGGCCGGAGCGGTCGCCCCGGCCGGCGGCTACTTTACCTCCCCAGGAGGTTACTACGCCGGGCCCGCGGGCGGGGACCC | 8 |
| wt-136710-136830 | 2563 | GGGTGCCTTCTTGGCGATGGACGCTCACACCTACCACCCCCACCCACACCCCCCTCCGGCCTACTTTGGCTTGCCGGGCCTCTTTGGCCCCCCTCCACCCGTGCCTCCTTACTACGGATC | 8 |
| wt-137070-137190 | 2566 | CTACACCCACTCGGGGCCTTACGGATTTCAGCCTCATCAAAGCTACGAAGTGCCCAGATACGTCCCTCATCCGCCCCCACCACCAACTTCTCACCAGGCAGCTCAGGCGCAGCCTCCACC | 4 |
| wt-137190-137310 | 2567 | CCCGGGCACACAGGCCCCCGAAGCCCACTGTGTGGCCGAGTCCACGATCCCTGAGGCGGGAGCAGCCGGGAACTCTGGACCCCGGGAGGACACCAACCCTCAGCAGCCCACCACCGAGGG | 8 |
| wt-137310-137430 | 2568 | CCACCACCGCGGAAAGAAACTGGTGCAGGCCTCTGCGTCCGGAGTGGCTCAGTCTAAGGAGCCCACCACCCCCAAGGCCAAGTCTGTGTCAGCCCACCTCAAGTCCATCTTTTGCGAGGA | 4 |
| wt-137430-137550 | 2569 | ATTGCTGAATAAACGCGTGGCTTGAAAGTAAACTTTATTGCGTGTTAGTACCTGTCCATTCACAGGGGTATCCAGCCCTTGCGCCGCCTCCCCCAGCCCGCCAGCCACCCCAGACAGGAG | 4 |
| wt-137550-137670 | 2570 | ATGATAATGATGAGGAGCACCGGAGCCACCACAGCACAAGTGATTAGGAGCAGGGCCCAGTGCACCCAGGTGGTCTTAGGGCGCCAGGGATCGATTGGAAAAGGGCCCAGGGTCACTGGC | 4 |
| wt-137670-137790 | 2571 | TTATGCGTGGGACGTTTAGAAACAGGCCGCCTATGGGGCCTGTGACTGGTGCTTGTGGTGTGGGAGACTAATGTGGTGGGGGCTATGGTAGTGGCTGGGATAACAGTAAGATGCATACGC | 4 |
| wt-137790-137910 | 2572 | TGAGTGAGGGTCCGGTTGGCATGGTATTGGTCGTCTTCTTCCCCTGCAGAGTAATTGCAGTGGACCCCGGAGGCCACACTGCAATTTCTCAGTGTCACATTGCACGTGTAGTAACCTGCA | 4 |
| wt-137910-138030 | 2573 | TGCGCAAGGGTCACATTGGGGATTATCAGAGAGACGGAGGTGTTGGAGTCATTTACCCATTCTAGGGTAAGGCTATAATTGTAACCCCCGTTAGTTATATGAGTTCCGTTGTTGGAAGTA | 4 |
| wt-138030-138150 | 2574 | GCTACGGCCAAGGGCAGTTGTCCATCCCCGGGAGTGTATCCCCGGCCCAACTCGATCCGAGAGACCGACTCATTGCTAGGAACGCTGCAGGTGAGATTCACTCTAGCACCTGCATGGGCG | 4 |
| wt-138150-138270 | 2575 | GTGACATTTTCAAATTTAACCAGATCTGAGAAAAATGCACAAACAGACCCCACACAGCAGCACAATAGAAGCACTAAATGAGTCATTCCTAAACTGTCAGTTTTAAAACTCCCTGCTTCT | 4 |
| wt-138270-138390 | 2576 | CAGGCCTAAATATGTGGTGGGGTGTGCTTAGGATCACTTTCATATTCTGCAACAACAGCCATACCCGGAAGAGGAGCTGCCGGTTGCCATTTTTCAAGCTGCTAAACCACGAGTGGCAGC | 4 |
| wt-138390-138510 | 2577 | AGGCCTAAGAAGCTCCTCAGCAACATGGAGACCTCGAAGGGAAACTGGCAGGAGCAGGGAGTCACGTAGGCACTAGCCTCTTCATGTGAGGTAAGAGATCGCTAAAAATGGGATCAGGGT | 4 |
| wt-138510-138630 | 2578 | ATGTAAACCGAGTTTTGCGGGGGATGGTGAGCCAGACACGGCGGGTGGGGGAAGGAGCTGACACGATTGAGTAGAAAGGGCCAAAAATACACCAGCTATAAGGAATTGCTCAGGCCAAAG | 4 |
| wt-138630-138750 | 2579 | TTGTTCCTCAGGTGGCTTTAGGCCTAATGTAGGCAATTGCGTGCCTAGAACATTGCTAATGTGCCCTGGGTTTCCTGCCTTCATGCAAATATTCTACCTCCCCCGGCCTGGTGCAAAATG | 4 |
| wt-138750-138870 | 2580 | TCTGCCTCAGAATACTAACAGCTAATCCAAGCTAACATTCTATCAGTAAACGGGCAGAAAACTGATAAGGACCGCGGAGTTTGGCCCTCCGCGGTGTCCGGTGGTCCTCACACGTGCCCT | 4 |
| wt-138870-138990 | 2581 | CCCCCCGGGCCGATGGCTGAGGCCCGGAATATGCAAGTGCATCTTTCTAACCAGTAGGGGCCTCCACCTAGGTGCTTTGTTAATCTTTAGTGGGAACTAGTGGGAGTGCTGTGCCTCGGG | 4 |
| wt-138990-139110 | 2582 | TACCCCTATCCTATAGGTCCTACCGGAGCTCCTTGTCTTGATAATCCCTGTAAACACACACCACCTAAGAACAAGGCATTGTTAACCTTTGGTGGAACCTAGTGTTAGTGTTGTGCTGTA | 4 |
| wt-139110-139230 | 2583 | AATAAGTGTCCAGCGCACCACTAGTCACCAGGTGTCACCGGAGGCTACTTGCCTCAGTGCCACTTTTACCTTCTCAAATCTATACGGGGGGGGGGGGGGCTCTGTAACATTTGGTGGGAC | 4 |
| wt-139230-139350 | 2584 | CTGATGCTGCTGGTGTGCTGTAAATAAGTGCCTAGCACATCACGTAGGCACCAGGTGTCACCAGGGCTACTTGCCTCGGCATCTCCTCACCGGAGAAGGGGTTAACAAACCCGTGGGGGG | 4 |
| wt-139350-139470 | 2585 | TCTTAGTGGAAGTGACGTGCTGTGAATACAGGTCCATAGCACCGCTATCCACTATGTCTCGCCCGGGCTATATGTCGCCTTACCTCCCCTATATAGTCACGACCCCACCGAACCAGGCAT | 4 |
| wt-139470-139590 | 2586 | GATGTAGAATAAAATTTTATGCATCATCTTCTAATCTGTGCCGCTTGGAGGGAAACATGACCACCTGAAGTCTGTTAACCAGGTCAGTGGTTTTGTTTCCTTGATAGAGACACAAGGACT | 4 |
| wt-139590-139710 | 2587 | GCCAGCCCCATTGGGGAGGGGGGGTGGGTACGGGAGAGTTTGGGCTCGTTTAAACAAAGTCTCATCTGATGCTCTGTGGCACCTCAAGGTGAATATAGCTGCCCATCGACGTATCGCTGG | 4 |
| wt-139710-139830 | 2588 | AAACCGGTGGGCCGCTGTTCACCTAAAGTGACGCAAGGTCTGTCAGCCGCCAGGGTCCGTTTACCAGGCTTTCAGGTGTGGAATTTAGATAGAGTGGGTGTGTGCTCTTGTTTAATTACA | 4 |
| wt-139830-139950 | 2589 | CCAAGATCACCACCCTCTATCCATATCCCACAATTGATAAACCTCCGCATGTCCAACCACCACGTTGAACAGGATGTGGCACCCTAAGAGGACGCAGGCATACAAGGTTATTACCCAGTC | 4 |
| wt-139950-140070 | 2590 | CTTGTATGCCTGGTGTCCCCTTAGTGGGACGCAGGCCTAGGTAGCATCATTTACACTAAAAGCAGTGACCTTGTTGGTACTTTAAGGTTGGTCCAATCCATAGGCTTTTTTTGTGAAAAC | 4 |
| wt-140070-140190 | 2591 | CCGGGGATCGGACTAGCCTTAGAGTAACTCAAGGCCAAGCATTTCACACCTGCAAATGCACCATGTAACCACAGATCTAAACTGAAAGTTGCAGCTTTAGATGGCAAGGAAACTTGGGTT | 4 |
| wt-140190-140310 | 2592 | TCAGGCATAGAAAGCCTGGCTCACTATAGCAGCCCATGTTTGTTCCAGGGTGGGGGAAAGGCACGTGCCCTTAGAAAACTTAGCTGCAAAAATTCTATTGTGTTGGGAGAGCCTCTATAT | 4 |
| wt-140310-140430 | 2593 | CTAAAGGCCTTTCCTCACAATACAAATGTTACTAACGTCTGCCCTCTGGAGACCTGCTATGTGGCTAGACGTATGGCCTACCCAAGACGTTGGGGGTCTCGGGTAGGCCATGATTCTTCC | 4 |
| wt-140430-140550 | 2594 | AGGCATAGGTTACAACCAGTCACTGCTATCAAGCCTACTCAGTTCCCAACGCAGCACATACCCCCCGCCTCTCCTGCCATGAGGACTTATGGCAGTGTTTACTGTTCTGCTTTTACTCTT | 4 |
| wt-140550-140670 | 2595 | GGACCAGGCTGTCATTCTATCAGAATAACAGGGGAAGCAAGGCCCCCTGCTTCAGCGGGACACGTGTTTCTAGAATCTCGGAGCCAATAACTACCTGCCCCTCTAATCTGTATGCTGCAT | 4 |
| wt-140670-140790 | 2596 | GAAAAACCACATACACGTGATGTAAGTTTAGCCAGTTTATTGTTACACCAATGCCCCGAAAGTCTCCCCCTGTCCCTTTGGGTCTCAGGACCCAGCCCTGGAGCTCGGGGGCGGCCGGGT | 4 |
| wt-140790-140910 | 2597 | GGCCCACCGGGTCCGCTGGGTCCGCTGCCCCGCTCCGGCGGGGGGTGGCCGGCTGCAGCCGGGTCCGGGGTTCCGGCCCTGGAGCTCGGGGGGCGGCCGGGTGGCCCACCGGGTCCGCTG | 12 |
| wt-140910-141030 | 2598 | GGTCCGCTGCCCCGCTCCGGCGGGGGGTGGCCGGCTGCAGCCGGGTCCGGGGTTCCGGCCCTGGAGCTCGGGGGGCGGCCGGGTGGCCCACCGGGTCCGCTGGGTCCGCTGCCCCGCTCC | 12 |
| wt-141030-141150 | 2599 | GGCGGGGGGTGGCCGGCTGCAGCCGGGTCCGGGGTTCCGGCCCTGGAGCTCGGGGGGCGGCCGGGTGGCCCACCGGGTCCGCTGGGTCCGCTGCCCCGCTCCGGCGGGGGGTGGCCGGCT | 12 |
| wt-141150-141270 | 2600 | GCAGCCGGGTCCGGGGTTCCGGCCCTGGAGCTCGGGGGGCGGCCGGGTGGCCCACCGGGTCCGCTGGGTCCGCTGCCCCGCTCCGGCGGGGGGTGGCCGGCTGCAGCCGGGTCCGGGGTT | 12 |
| wt-141270-141390 | 2601 | CCGGCCCTGGAGCTCGGGGGGCGGCCGGGTGGCCCACCGGGTCCGCTGGGTCCGCTGCCCCGCTCCGGCGGGGGGTGGCCGGCTGCAGCCGGGTCCGGGGTTCCGGCCCTGGAGCTCGGG | 12 |
| wt-141390-141510 | 2602 | GGGCGGCCGGGTGGCCCACCGGGTCCGCTGGGTCCGCTGCCCCGCTCCGGCGGGGGGTGGCCGGCTGCAGCCGGGTCCGGGGTTCCGGCCCTGGAGCTCGGGGGGCGGCCGGGTGGCCCA | 12 |
| wt-141510-141630 | 2603 | CCGGGTCCGCTGGGTCCGCTGCCCCGCTCCGGCGGGGGGTGGCCGGCTGCAGCCGGGTCCGGGGTTCCGGCCCTGGAGCTCGGGGGGCGGCCGGGTGGCCCACCGGGTCCGCTGGGTCCG | 12 |
| wt-141630-141750 | 2604 | CTGCCCCGCTCCGGCGGGGGGTGGCCGGCTGCAGCCGGGTCCGGGGTTCCGGCCCTGGAGCTCGGGGGGCGGCCGGGTGGCCCACCGGGTCCGCTGGGTCCGCTGCCCCGCTCCGGCGGG | 12 |
| wt-141750-141870 | 2605 | GGGTGGCCGGCTGCAGCCGGGTCCGGGGTTCCGGCCCTGGAGCTCGGGGGGCGGCCGGGTGGCCCACCGGGTCCGCTGGGTCCGCTGCCCCGCTCCGGCGGGGGGTGGCCGGCTGCAGCC | 12 |
| wt-141870-141990 | 2606 | GGGTCCGGGGTTCCGGCCCTGGAGCTCGGGGGGCGGCCGGGTGGCCCACCGGGTCCGCTGGGTCCGCTGCCCCGCTCCGGCGGGGGGTGGCCGGCTGCAGCCGGGTCCGGGGTTCCGGCC | 12 |
| wt-141990-142110 | 2607 | CTGGAGCTCGGGGGGCGGCCGGGTGGCCCACCGGGTCCGCTGGGTCCGCTGCCCCGCTCCGGCGGGGGGTGGCCGGCTGCAGCCGGGTCCGGGGTTCCGGCCCTGGAGCTCGGGGGGCGG | 12 |
| wt-142110-142230 | 2608 | CCGGGTGGCCCACCGGGTCCGCTGGGTCCGCTGCCCCGCTCCGGCGGGGGGTGGCCGGCTGCAGCCGGGTCCGGGGTTCCGGCCCTGGAGCTCGGGGGGCGGCCGGGTGGCCCACCGGGT | 12 |

| | | | | |
|---|---|---|---|---|
| wt-142230-142350 | 2609 | CCGCTGGGTCCGCTGCCCCGCTCCGGCGGGGGGTGGCCGGCTGCAGCCGGGTCCGGGGTTCCGGCCCTGGAGCTCGGGGGGCGGCCGGGTGGCCCACCGGGTCCGCTGGGTCCGCTGCCC | | 12 |
| wt-142350-142470 | 2610 | CGCTCCGGCGGGGGGTGGCCGGCTGCAGCCGGGTCCGGGGTTCCGGCCCTGGAGCTCGGGGGGCGGCCGGGTGGCCCACCGGGTCCGCTGGGTCCGCTGCCCCGCTCCGGCGGGGGGTGG | | 12 |
| wt-142470-142590 | 2611 | CCGGCTGCAGCCGGGTCCGGGGTTCCGGCCCTGGAGCTCGGGGGGCGGCCGGGTGGCCCACCGGGTCCGCTGGGTCCGCTGCCCCGCTCCGGCGGGGGGTGGCCGGCTGCAGCCGGGTCC | | 12 |
| wt-142590-142710 | 2612 | GGGGTTCCGGCCCTGGAGCTCGGGGGGCGGCCGGGTGGCCCACCGGGTCCGCTGGGTCCGCTGCCCCGCTCCGGCGGGGGGTGGCCGGCTGCAGCCGGGTCCGGGGTTCCGGCCCTGGAG | | 12 |
| wt-142710-142830 | 2613 | CTCGGGGGGCGGCCGGGTGGCCCACCGGGTCCGCTGGGTCCGCTGCCCCGCTCCGGCGGGGGGTGGCCGGCTGCAGCCGGGTCCGGGGTTCCGGCCCTGGAGCTCGGGGGGCGGCCGGGT | | 12 |
| wt-142830-142950 | 2614 | GGCCCACCGGGTCCGCTGGGTCCGCTGCCCCGCTCCGGCGGGGGGTGGCCGGCTGCAGCCGGGTCCGGGGTTCCGGCCCTGGAGCTCGGGGGGCGGCCGGGTGGCCCACCGGGTCCGCTG | | 12 |
| wt-142950-143070 | 2615 | GGTCCGCTGCCCCGCTCCGGCGGGGGGTGGCCGGCTGCAGCCGGGTCCGGGGTTCCGGCCCTGGAGCTCGGGGGGCGGCCGGGTGGCCCACCGGGTCCGCTGGGTCCGCTGCCCCGCTCC | | 12 |
| wt-143070-143190 | 2616 | GGCGGGGGGTGGCCGGCTGCAGCCGGGTCCGGGGTTCCGGCCCTGGAGCTCGGGGGGCGGCCGGGTGGCCCACCGGGTCCGCTGGGTCCGCTGCCCCGCTCCGGCGGGGGGTGGCCGGCT | | 12 |
| wt-143190-143310 | 2617 | GCAGCCGGGTCCGGGGTTCCGGCCCTGGAGCTCGGGGGGCGGCCGGGTGGCCCACCGGGTCCGCTGGGTCCGCTGCCCCGCTCCGGCGGGGATGGGGGTGCGCTCCCAGGCCGGACCCTG | | 12 |
| wt-143310-143430 | 2618 | GTGCCAGGCAGGGACCCCGCGCCACCCGCTTCATGGGGGGGGAGGCCGCCGCAAGGACGCCGGGCCGGCTGGGAGGTGTGCACCCCCCGAGCGTCTGGACGACGCTGGCGAGCCGGGCCA | | 12 |
| wt-143430-143550 | 2619 | GCTCGCCTTCTTTTATCCTCTTTTTGGGGTCTCTGTGCAATACCTTAAGGTTTGCTCAGGAGTGGGGGGCTTCTCATTGGTTAATTCAGGTGTGTGATTTTAGCCCGTTGGGTTACATTA | | 4 |
| wt-143550-143670 | 2620 | AGGTGTGTAACCAGGTGGGTGGTACCTGGAGGTCATTCTATTGGGATAACGAGAGGAGGAGGGGCTAGAGGCCCGCGAGATTTGGGGTAGGCGGAGCCTCAGGAGGGTCCCCTCCATAGG | | 4 |
| wt-143670-143790 | 2621 | GTTGAACCAGGAGGGGGAGGATCGGGCTCCGCCCCGATATACCTAGTGGGTGGAGCCTAGAGGTAGGTATCCATAGGGTTCCATTATCCTGGAGGTATCCTAAGCTCCGCCCCTATATAC | | 4 |
| wt-143790-143910 | 2622 | CAGGTGGGTGGAGCTAGGTAGGATTCAGCTAGGTTCCTACTGGGGTACCCCCCTACCCTACCTTAAGGTGCGCCACCCTTCCTCCTTCCGTTTTAATGGTAGAATAACCTATAGGTTATT | | 4 |
| wt-143910-144030 | 2623 | AACCTAGTGGTGGAATAGGGTATTGCAGCTGGGTATATACCTATAGGTATATAGAACCTAGAGGAAGGGAACCCTATAGTGTAATCCCTCCCCCCCCTACCCCCCCCTCCCTTACGGTTG | | 4 |
| wt-144030-144150 | 2624 | CCTGAGCCCATCCCCCACCCCAGCACCCCGGGGTGACGTGGCACCCCGCGTGCCTTACTGACTTGTCACCTTTGCACATTTGGTCAGCTGACCGATGCTCGCCACTTCCTGGGTCATGAC | | 4 |
| wt-144150-144270 | 2625 | CTGGCCTGTGCCTTGTCCCATGGACAATGTCCCTCCAGCGTGGTGGCTGCCTTTGGGATGCATCACTTTGAGCCACTAAGCCCCCGTTGCTCGCCTTGCCTGCCTCACCATGACACACTA | | 4 |
| wt-144270-144390 | 2626 | AGCCCCTGCTAATCCATGAGCCCCGCCTTTAGGAAGCACCACGTCCCGGGGACGGAAGCTGGATTTTGGCCAGTCTTCAATTTTGGGGAGTGGTTTTGTGTGAGCCGGAAGTTGGCAATG | | 4 |
| wt-144390-144510 | 2627 | GGGTGAGGGTGGCGCTGGTTAAGCTGACGACCTCCCAAGGTCTCTCACCCTGGGTACACAGGTGGGGCGGCAGCCTCTAACTTTGGCTGTGGCCTCTATTTCCTCCCTTTCCTAGCCAGG | | 4 |
| wt-144510-144630 | 2628 | GCCATGTGTTCCTGCATGTCTACTTGCCTCCTGTGGTGGCAGAGCTTGGCCCTGGGCCCAACCCCCGCCTTGGGAGCCTGTAGGGGCCAACACCCTTGGTTTGTTTGTGTTCCTGTTTGC | | 4 |
| wt-144630-144750 | 2629 | TGGCAACTTACTGGCAGCCGAGCAGATTCTAATGGGCGCCCGCCTTCTTTCTCTCTTGTTTTATTAATAGAATCTCAGCCAGGACCTATACCTGAGACTTCAAAGTCTGGTCCTGGGTTC | | 4 |
| wt-144750-144870 | 2630 | TGAGACCCCCAAGATTTGTCATGCACACCTGCACACCTGTTGGTATTGGGTTTCTATTCTTGAGTGTGAAAGTTTGTAAAAAAATTCATAAAATGTCACTAATTCCTCTTACCTGTTTAG | | 4 |
| wt-144870-144990 | 2631 | GGTATTGTGCAATTCTTCAGCCTGCCTATTTTCAATTTGCCTAAGGTGGCAATTTAAGATGTGGTTAATTAACCATTTTCCTGTCTGACACCACTGCATGGGCAACCGGGTTCCATGGCA | | 4 |
| wt-144990-145110 | 2632 | CATTTAGAGATAAACATAGATGTCTTGTCTTGCTCATGTGCAGAGGAGGGGGTGTTGGTGTGCAATATAGTTTCTGGATTCCAAATTGAGTTGGGGGTGCTATTTTCACTATGGAATTAA | | 4 |
| wt-145110-145230 | 2633 | ATTACTGACATTAGACAGTGGACACCGGGCTATATGTGGGGATGTCTGTGGCTTGTCATTTCCTCTTAGAAGGTAATCCCCCATCTTAACTTCCCTTTAAATTGTGATGCAAGCCCTGGG | | 4 |
| wt-145230-145350 | 2634 | TTATTTATAGAATGATTATCTAGGTTTGATAGTCTGAAGGCTGGGCAGAGAATGTTTGTAATTTTTATTCACCTTCTTTACCCCCCACGAGTATCCAGTTCTAGAAGATCTCCTGATATC | | 4 |
| wt-145350-145470 | 2635 | CCGGGCTGCCATTATTCCCTTGAGTGTTATAGCTTCCTCTTAACTTAAGCAAGAGCTCCAGGATGTTAGCTTTTTTGGTGGGGCTGGTTGTCAGGAAGAGGTTCCAGTGTTGTCCTTTAT | | 4 |
| wt-145470-145590 | 2636 | TTTTAGATGTTAGCTTTGTGTTAGGTTAGTATGGGCTGGGTATTCACTAGTGAAGGCAACTAACACAGTTAGACGTGCTAGTTGTGCCCACTGGTGTTTATCCGGTCCCAAATGTCACCA | | 4 |
| wt-145590-145710 | 2637 | CAGAACACAGGGGGCTGGATTTGGCAGCAGCACTTGTGCTTTTGTTGATTTTTACCCGTGTATCAGAGTGGGGGATGCTAGCCAATTTAGCTTCCCCTCCCCTTAACAGGGGGTCTCGCG | | 4 |
| wt-145710-145830 | 2638 | GGGTGCCAATTGTCGCCTGCCTTCCCCCGCTTCCCCTTGTTAACTTATAGCATGATAGGTAGGTCACCTAACGTGGAAGCCTGGTGGGTGATCCTTCCTCGGTAGGGAGCGCTTAGGGCT | | 4 |
| wt-145830-145950 | 2639 | GTTGAGCTCAACAGCCCCACCTGGGTAAAATGTATGTTCTAAAGAGTTACCCAATTATAACAAAACTGTTGTAGGGTAACGAAGACCTGATGGAAGTGGTATTGTTGCCGTTGAAAGACG | | 4 |
| wt-145950-146070 | 2640 | GGTGTCCTGGCTCAAGTTCGCACTTCCTATACAGTGTTAAAGCCTTGTATCGGAAGTTTGGGCTTCGTCCCAGTGTACTCGATAATGTCGACTGCTGCGAAAGGTTTGGACCGTCTTCCA | | 4 |
| wt-146070-146190 | 2641 | GTAGGTGTTGGGGGTCCCAAATCACGAGGTTAGGCAGGTGCACTTGGCTCTTTAGGAGGGACCCTTAAGCCAGACAATGTAGTGCCCCTTTTTTTTGCAAATTGGCCTTATTATTAATTT | | 4 |
| wt-146190-146310 | 2642 | CTTGTTAACACTAATTCTGTTCTATGACCCTGTGTTTTTCAGATGCCGTTGAACGTGTCACTGAGCTGAATTTGGACGCAGCTACTTGACCTTTGCCCCCGTGCCTCCAGCGCTGATAAG | | 4 |
| wt-146310-146430 | 2643 | TGCTGCGTCCACTTTGTGTTACAGGTGGGCCAAACCTCCAGAATATCAATTGGTGGGGCCTTGGTGGGCTGCATAAGGCAGTAGGTTTGAGGTGACCTACTTGGACCATGTGGATCCAGT | | 4 |
| wt-146430-146550 | 2644 | GTCCTGATCCTGGACCTTGACTATGAAACAATTCTAAAAAAATGCATCATAGTCCAGTGTCCAGGGACAGTGCACTCGGAAGTCTCATCATCTCCGTTTGTGTGTTTAGTGTGGCCAGTA | | 4 |
| wt-146550-146670 | 2645 | CGGCCACCCCTGTGCCACGCCCTGGCATGCTGCTGACATCTGGCCGCCAATTTCAGCGGGCCCTTTTCCCCCTTGTTCACCCCATAGCAAGAAGGGTAGGTTACATGGGTATTTTCCCAT | | 4 |
| wt-146670-146790 | 2646 | CAGCACCTGACTGGCCGGTGCAATTAGAGGAGAGGGCAACAACGCAAGGCTGTTGTTTTATTTGGGTTACAAGAGCTGCGGCGGTCGATGGGTTCACTGATTACGGTTTCCTAGATTGTA | | 4 |
| wt-146790-146910 | 2647 | CAGATGAACTAGAACTGTCACAATCTATGGGGTCGTAGACAGTGTGCTTACCAGACTTCCATGGAAGATGTGAATTTGCTGCTAGCTATATGGGTGGTGCTATGGGCTCCCTAGGGACTC | | 4 |
| wt-146910-147030 | 2648 | ATGTAGTGGGGCTTTGTGATAGCTAATGAATGTGGCAGCTGTTGTTTGTACTGGACCCTGAATTGGAAACAGTAACTTGGATTCTGTAACACTTCATGGGTCCCGTAGTGACAACTATGC | | 4 |
| wt-147030-147150 | 2649 | TGAATATCTTGAATATGGGAGGAGGGGGGCTTTGGGTTCCATTGTGTGCCCTTTCCTGGCCAACGTGAGGGTCCTAGTGTTATAGGGCGTGGCAGTTTTCTTGAGGGCTAATAACCCGGG | | 4 |
| wt-147150-147270 | 2650 | TGAGGCGGTTGTCACAGGTGCTAGACCCTGGAGTTGAACCAGTACCACTCGGTTACAAAGTCATGGTCTAGTAGTTGTGACCCTGCAAAGCTACGTGGGGATGAGCAGCCAGGGACTTTG | | 4 |
| wt-147270-147390 | 2651 | GTTGGCAAGCAGACAGGCGGCGCATTGGAACCCCAGAGGAGTGTCCCGGGGCCACCTCTTTGGTTCTGTACATATTTTGTTATTGTACATAACCATGGAGTTGGCTGTGGTGCACTCCAT | | 4 |
| wt-147390-147510 | 2652 | CTGGTAAGGGGGGCTGGTGCGGACGCCTGTGTTTAGTCTATGCCAATGTTTACCTGCCTTGGGTTACTATTCCAAACGACCACACCTTTGAGGACACCTGGAGCCCTGATCATTCTCGGCT | | 4 |
| wt-147510-147630 | 2653 | TTTACTGCCACCTGGCTTCTGTTGGGTCAGACAGTTTGGTGCGCTAGTTGTGTGCTTAGCAGCAACGCACACCAGGCTGACTGCCTTAGCAGTGTGGCCCTTTATTGTGGCATCCTAAGG | | 4 |
| wt-147630-147750 | 2654 | AGGGATTCTGGAGTGCCTTTCGCGTGAAGCATGCCCTGAGACGTACTCGAGTTAGGACTTAATCGCTCCTGTGCCGCTGGATGAGGGAGCGCCAATTTGTACATCCTAGCTCTGGCCATA | | 4 |
| wt-147750-147870 | 2655 | GAGTTAGCCCACCCTTGTGTCTCCCTTTGGCCTTTGCGGTGCCAATTTCCGGTGGTTTCCCTTTTCCGCCCGTTTATCCAATAGCATGTAAGAGAGGTTGCCTAGATTTGGCAACTTTGA | | 4 |
| wt-147870-147990 | 2656 | GGGAACGTTCCGTGTAGCTGGTGACCTAACACCCGCCCATCACCACCGGACAGATTCTGAACTTGTCCTGTGGTGTTTGGTGTGGTTTTGGGGTACGCAGGAGTACGTTGGAATGCTTTG | | 4 |
| wt-147990-148110 | 2657 | GAGCCGAGAGGGATGGGCCCGCTTGTGCGCTTATGTGTTACACGGTGCCAATAACCGGCCCGGTGCGGCTGCCCCGTGACCCGTGGGCCTTACCTTCCTGGCCATCGGGGGACCCTGGTG | | 8 |
| wt-148110-148230 | 2658 | CTAGGGTCCCTTGTGTTGCTTTCTGCCATAGGGGGGAAAGCATCGCCTTCAGAATTGGCTGCTCCGTTGGAACATTTGAGGCCTACTGTATCCGTGTCCTGACAACATTCCCCGCAAACA | | 4 |
| wt-148230-148350 | 2659 | TGACATGGGTTAATTTAAACATGTTTTGTTTGCTTGGGAATGCTCTTAGGGCCTGGAAGCTTGTCATTGGATTCATCGTTTCCTGAACTACAGGCGTAGGGCCTATTGTAGCAGGCATGT | | 4 |

| | | | |
|---|---|---|---|
| wt-148350-148470 | 2660 | CTTCATTCCTGCGTACCGAATGGCATGAAGGCACAGCCTGTTACCATTGGCACCTTTTTTCCATGTAAACCTCCGTGATCCTGGGTCCTTTGGAGACTCAAGTGTGAATTTGTTTTGGTG | 4 |
| wt-148470-148590 | 2661 | TTCGGCGCCAGGGTATCTCGACGTTGGAATGTCAACTCAACTTGGGCACCTCGATAACCGGCTCGTGGCTCGTACAGACGATTGTTTGGCTCTGTAACTTGCCAGGGACGGCTGACGATG | 4 |
| wt-148590-148710 | 2662 | TGTTTAGTCTGCCACTTGCATCCGGCGCTTTGGTTACTCGGGAGACTAATGGGGGGTGTGGTATGGCACAGGCTGGGGGTGAGTCTGGGGATGTCCCTGGGCGTTGCTGCAGCCCATTCG | 4 |
| wt-148710-148830 | 2663 | CCCTCTGGGGATGAGATGTTCAGGGGTGGCCGGTACCCTACGCTGCCGATTTACATAATATAAATTGTAAATGCTGCAGTAGTAGGGATCTGGACGCGCGACCTGCTACTCTTCGGAAAC | 4 |
| wt-148830-148950 | 2664 | GCCAACCCAGGAGCGTCGCCTCTGGCCCCATACTCCCGCCATGCGACTGCTCGCCCCCTCCCAGGCCTCCCTGGTGAGCCCTTGCCGCTCCCCGCATTCCTGCTTTCGGCGCCCCTGCGG | 8 |
| wt-148950-149070 | 2665 | ATCCCGATGACAGCAGGCCTTTCCTTCCCCCGTTAATGAAAAGAATGACAGTGAGGTTGTGACAGAAGGACAGCTTTATTCAGTTTACAGAGTGCCCTCGGAGGCTACGATATTCCCGTT | 4 |
| wt-149070-149190 | 2666 | AAATGTCTTGTTGATTCTCTCAAAGGTGGGGAGGGAGGAGCTCTCCACAACAATGTTCCCTGGCAGCGTGAGCGCGCAGCCCTGCCGTTGGATGTATCTTCTCATGATGGTGCTGATAGA | 4 |
| wt-149190-149310 | 2667 | GGGGTCTCCGGCGTAGATGAAAAAGGCCTGGGCCATGCTCTGGCCGGTCACGATCGTTATGGGGTTGTTGGAAATGTTCCGGACCGTCAGCTTGAGGGTCTGGCCCGGCTTCCACTCCTG | 4 |
| wt-149310-149430 | 2668 | TGGGTAGACGTAGAAGACCGGGTTGGAGGAGTGGGACACGACAACGGCCGTAATCTTGGAGCTCAGGGGGGCCTCGTAGGTGTTGTTGTATTCCAGCTCCGTGATGAAATTAGGAGGAAT | 4 |
| wt-149430-149550 | 2669 | AATCACAGGGGAGCCAAAGTAGCGGATGTCTGTGGATTCCCCGTCCCAGCGCCAGTGGCTCTTAGGGTAGGGGTTGTAACGGAAGGCAATAATCACATCATCCAATAGGGTCATGCCCAC | 4 |
| wt-149550-149670 | 2670 | CTTGACGTTCAGCGGGCCCTCTCGTTTCAGGTCCGGCGTGTCCACGGAGACTCGGACGTAGCCCTTACCGCGGCGTATGGCGTTTACCGGACACACCTTCCCCGGGAATGTGTGAATACG | 4 |
| wt-149670-149790 | 2671 | GGCGTATGACTTTAGAAATGGGGGCGTGTGCTGCGCCAGCAGGTAAGGCAGGCACTCGTCCTGGCTGGTGACGGGAGAGCCACTGAGGAAGATCTGGGGCTCGCTGGTGTTTAGCTTGTC | 4 |
| wt-149790-149910 | 2672 | CCCGCTCTGGGTGCAGGAGCGTGTCAGCTGAATGTCGCTCTGCCCGGGCAGAATCTGCAGGTAGAGGTAGGGGTTCTTGACCAATCTGATGGGCACAATGTACCAGGTAAACTTCCCTTT | 4 |
| wt-149910-150030 | 2673 | CTCTATGAACAGGCTGCGCGGATTCAGGACGCTTAGCACGATGTCCTGGTCAGAGTGCATAACGAAGAAGGGCTTGAGGAATACCTCGTTGTCTTCCGCTCCAAAGAACAAAAACGCGAC | 4 |
| wt-150030-150150 | 2674 | CGTAAAGTAGCGGCTGCCGTAGGTGGTCGTGTTGAAGGAGAAAGAAGGTAACTTGAAGCTGAGTATCTGGCCCACCGAGGGGCAGGGAGGCAGCTCTTGGCACTGCGCGTCCAGCTGCAA | 4 |
| wt-150150-150270 | 2675 | TACCTGCTTGTTGGTGACGCGGACGTATGAGGGGAAGATCTCGTACTTCCACACGCCTCTCATGAACGACGTGTCTGGTTTTTCAGTGGGCCGCAGGCGGCGGAGGCTGTTCCTGAACGA | 4 |
| wt-150270-150390 | 2676 | CGAGCGCCGGGACGCTAGTGCTGCATGGGCTCCTCCGGGGTAAGCTTCGGCCATGGCCGGAGCTCGTCGACGGGCAAGGTGAGAGTCGGGGGGCGGGCGACGGTGCGGCCCCAATACAAC | 8 |
| wt-150390-150510 | 2677 | TCTCCGCTCGTTAGCTGGTAGAATATCCGCCCGGCGTCTAGGTTGTCACTTCGCTCGGCCGGCCAGAAGAGCGCAAGTCCAAGTCTGGTGCTGGGGCCGATGTGCAGCGGTTTGTGCCCG | 4 |
| wt-150510-150630 | 2678 | CAGTTGTAGACTGTCATTTTTATGGGCGAGTGGGCGGTCCACACGCGCGGGCGCAGCACCCATTGGTCGCACGCCGCCTCCTGGAATGTAAACCCCCAGAGAGAGGGCGTGCCGCCCTGG | 4 |
| wt-150630-150750 | 2679 | AGATGGCCCTGTGCCATCACATGTATTTCCTCCTTGGGTGGAACAACGGCGTCGTGCTCCGGGTGGAGGGGGAATAGCGTCCAGGCATCTTTCAGGGTCACGAGACCGGGGTCCATGCTC | 4 |
| wt-150750-150870 | 2680 | AGAGAACAGCCCTCCCGGGCGGTGGGCGGCCCGGGCTCCAGCAGAACGTCGCAGACCCAGCCCTCCTCGGCCCTGTCCACCTGTATGTCCAGGTGCACGGACCCGGAGGCTGCGTCTCGT | 8 |
| wt-150870-150990 | 2681 | GACATGGCCAGGCCTGGTGCCAGCCGACCACGTCCCGTGTCCCAGCCGAGGCCGCGCCAGAGCAGAGCCCGGGACTGACTCAGGGCCACATCCCCTCGGCCCGCGGACGCCGCCTCGCCA | 8 |
| wt-150990-151110 | 2682 | GCCCCCGGGCCTTCATGGGCCCGCTTTCTACCTCTCTCCGGCACCCCAGCCTGGTCAGCCGCAGAGGAAGCATGACCTTGGGGTGGGACGGGGCAGGCGTGATCCTGGGCGCAATCTTTG | 8 |
| wt-151110-151230 | 2683 | CCGATCCCCACACCTTCACTCCTTGTTAGGTTGATAGAATGTCGGTACCACGCCACGGGGGGCGGGCCCGCATAGGGAAAAGCCAGGGAGAGCGATGTGGGCGAGGATGGGCTCAGGCGG | 4 |
| wt-151230-151350 | 2684 | CCCCAGACACGCAATTTGCCCCCCTGGGCGGCCGCAGCCTGCCCCTCGGCGGCCCGTGCCCCAGCTCCGTCACGGGGGGCGCATAGGAGGGGTATATCTAGGATAGCCGCACCTACACAA | 8 |
| wt-151350-151470 | 2685 | ATGAGACACAGACACAGGTCGTGAGGATTTAGGCAACGCAGGCTTGTCTTTATAGTTACAAACATGGGAGCGTGCACCTGGAAGATGCAGCTGGGGTAGATCTTTACATCTTTACAGGGC | 4 |
| wt-151470-151590 | 2686 | GCAGCGGCCGCCAGACACTGAAGGGCAGAGTTCACGGCGGGCACCTCCCAGAGGGAGCCCACCAGCCCGTACCTGGCCACGGCCAGGGCCTCGTAGGCCGAGACGGGCAGCCGGAGCTTG | 8 |
| wt-151590-151710 | 2687 | TGGTACTGTCCCTCCGGCAGGTGGAGTGGGACACAGTTAGAGAACATTAGTCCTCTGGTCCCTATCTCCACCCGCCAGGCCTGTGTGTCAGTTTGCAGGGCCATCCTCGCACTCAGGTGG | 4 |
| wt-151710-151830 | 2688 | ACTGGCTAGGCACCCTTCTGAAGTATCTGGCGGTGACTGTCACCTGGTTCTTGAGAGAGTCCATAAAATGGCTGAAGCTCCAGGCGTATAGTATAATGAGCAACAGGGCCAAACAGGCGG | 4 |
| wt-151830-151950 | 2689 | CGGGGCCTGGGTAGTAGCGGGCAACGAGAGACTCTGTGCAATCAAACCCCAGGCTCCCGGCCTCACCCAGGAAGAGCAGCGGCAGGGACAGCATAAACCAGGAGAAGGCGCAAATGAGTC | 4 |
| wt-151950-152070 | 2690 | CGGTGAAGGTGACGTTGCATATCAGGCGCGGCTTCCTTCCGAATTTTGTGCGCAAAAGTTTCCAGATGATGATAACTGTGAGGAGGACGATCAGGACTGCCGCCAGTAGGTAGCAGCCGG | 4 |
| wt-152070-152190 | 2691 | CTTTCAGTCCTTGGACGGCCGTGTGCATGCCTTTGGTGGGGCCTTCCCTGCACATGTTGGGGCCTCTGTTGAGATTGGCGTCGGGGCCCATGGTAATGAGGAGGATGATAATCAGCAGGA | 4 |
| wt-152190-152310 | 2692 | GTACCAGACAAAACACGCCCATCAGGTACAGGCACACATTTCTGTGGGAGGTTCTCTTGGGCGTTCGGCTGAACAATGCTAGGGTCTTCTCCAACGCCATACCCAAGTGAGTCCATACGG | 4 |
| wt-152310-152430 | 2693 | AGCACATCAGGCCCAAGAACATCATGTTCTGGGTCAAAAGGCAGAGACCGGTAGACGAGAACTCCTGAATCATTTTTCCCAGCACCCAGAGCAGCAGTTCTATGAGAAGAGCTATCAGCC | 4 |
| wt-152430-152550 | 2694 | AGACATCCATTCGGTGAACCAATTTTCTTACAAAGATGATAAACAAGATGCCAGCCAGTGTTAGCAGAATCAGCAGGACGAGCAGCAGGCTTGTCATGCCGCTGAGGAAGGCGCTGTAGG | 4 |
| wt-152550-152670 | 2695 | ATTTAGTGCACGCATCTTCCGTTGCATTGACGGAAGTCATGTTGGCCACCAGGGTCCCCACGGTGGACCCGGGGGCCATGGTGGAGAGCATCTTGCTGGTCAGAGCCAGACTGGGTGGTG | 4 |
| wt-152670-152790 | 2696 | TCTGCAGCAAAAGAGGAACTTGCCCAGGCAGTCAGTTATTTTGCATGCCACCTCCCTGCCTGGTGGACTTCCAGACTATTTTCTGCATTCGCCCTTGCGTGTCCATTGTTGCAAGGAGCG | 4 |
| wt-152790-152910 | 2697 | ATTTGGAGAAAATAAACTGTGAGTTTCACAGATCCACGGGCCACGCTCCCCTGGGGGCTTCATGATCCCACCGCCTTTCCCGATGATGATGACAACCGCGGCTGTCTGAAGCGGCTGACG | 4 |
| wt-152910-153030 | 2698 | AAATCGGTTGAGATTCTGATGAGAGGCTTGGGGGGGTCTTTGCCCTCAAGGCGAGGCTCCTTCTCCTAGGAATGCCGAGCCCCCTGCACTAGCTTCGCTCCACTGGGGATCTTTGCCAGC | 4 |
| wt-153030-153150 | 2699 | CTTCATACTAGATTCAGCGATCCCCCGGTTGGGAATCTTCGCCAGCCCCCCGTCCTGCTATCCCGCTCGTCGCCGCGCCTCCCATGCTAAGGGCCCCCTTCCTTTCCCTTGACTTTGGGG | 4 |
| wt-153150-153270 | 2700 | ATATTCGGAGTCTGCTCTCGCCGCTCTCTTCTCTCGTTTAAACGAGAGAATAGTAGTAGGGTCCAGTCTCAGGCCCCCTCACTTTGGGTCTTAGAATGGTGGCCGGGCTGTAAAATTCTG | 4 |
| wt-153270-153390 | 2701 | GAGGACGGAGAGGGCGGCCCCGGAGTTGTTATCAAAGAGGCACTGGAGGATGTTGGCCGCTCCTTGGAGCAGCTTGTCGAAATAATGATCCACGGCCACGGGAACGCCGTGCCGCTCGGC | 4 |
| wt-153390-153510 | 2702 | GTAGGCCGGGTCCTCGGCCATCTCCGTCTTTCTCGCCCCCTTCACTCCCCCCTTGGGCTCCACAAAGACGTACTGGATGCGGTCGTGGATCTGGGGCAGTTCCTCGTTGCGCTCGACGAA | 4 |
| wt-153510-153630 | 2703 | CTTCTGGTAGACGGCCAGGTGAGGCATCTGGGTGCTCTTGTAGGCTGAGAGCTTGCGGCTGAGCTCCGTTGAAAAGCAGAGCTCCCCCATGGGGACCCTGCCTTCACGGAGGTCTGTGTA | 4 |
| wt-153630-153750 | 2704 | GGCCTGGTTTAGGATGTCAATGACGGGCAAAAAGCCCACAGGTAGCCCTTGTGTAAATGACTCTTGGAAGGGCCGGTGGGAGAGGAGGCTGGCCGCCTCCTTTACCCGGGCATCCGCCAG | 4 |
| wt-153750-153870 | 2705 | CACCAGGTCGAGCACGCGCCGGCAGCGTGTCTGCACAAACTTGCAGGCCGTCTTCCGGACGAGCTCCACCCCCTTCATCAGGGTCTTGCCGTCCGTCAGCACCCCCACATATCTCTTCTT | 4 |
| wt-153870-153990 | 2706 | TGTAATCAGCATCAGGCAGGAGAAGGTCTTCTCGGCCTCCAGGGAGATGGGGGCCACAAACAGGCTCCGGGTGGTGTGGGCGGCCAGGGCATCGGCAAAGCGCAGGGTCTCGCTCTCTGA | 4 |
| wt-153990-154110 | 2707 | AAACCCCCGGCACTCGATAAACAGCGAGTCCGTGTCCCCGTAGATGACTCGAAGCTGGCCCTCGGGGTTGAGGGGCGCCCAGGCGTCCGGGGAGGGGGCCAGGGCCTGCAGGTTGGCGGG | 8 |
| wt-154110-154230 | 2708 | GCTCAGGGCCTCCACGAAGGCCTTGGCCCGCTCCAACATCGTGCGGCCCTGCAGCGTCACCGTCTCGGCGATGGAGAGGCAGGGAAAGAGGCCGTTGGCCACCCCGGTGAAGCCGTAGAC | 8 |
| wt-154230-154350 | 2709 | GGCGTTGCACGTGCACTTGATGGCCAGCTGCTGCTTGTCGAGGATGGTCCTTTGGCGCGGATCCTCGCAGGCCGCCAGCAGCTTCTTGATGGCCTTGCGCTTGGCCAGCCAGGAGGTCAA | 4 |
| wt-154350-154470 | 2710 | CAGACTAGCCAAGAAGGACTCGTGCACGTGCTTCTTTACAAAGTGGTAGACGCCCCCCGTGAGCCTGAAGGACTCATAGTCTTCTCCCGGGCGCAGGCCGGCTAGCCTGTGCTCTTCTCC | 4 |

| | | | |
|---|---|---|---|
| wt-154470-154590 | 2711 | CGGCGTTATCATGGTAGAATAACAGAGATTATGAGCCTGAATGATGCTCGGGTAGAGGCTGGCAAAGTCCACCACCAGAACCGGGGAGTTGTAGAATCCGGACAGGGGCTGGATGACGGT | 4 |
| wt-154590-154710 | 2712 | GGCCCCCTGGTAGCCGTCCCGGTCAGAGGCCGAGGGCATGGGCAGGATAAAGTTTTCCTTTTGGGCGGCCGCCAGGAGGCAGGAGAACACGCGGATCTGCTGCCCATCGTCCAGCACCCG | 8 |
| wt-154710-154830 | 2713 | CCTGCAGGGGATGTGAGCGATCTTGGCAATCTCTGCCACCTCCACGTGGATCACGAAATGGTTTAGCAGATCCATGACCAGGGCCGAGTCCTGCACGCAGTACATGCCGAGCCGCCTGCG | 4 |
| wt-154830-154950 | 2714 | CCCCTCGGGGCCCGCTGCAAAGAGGCGAGGAATCTCCTTGTAATGCACATCCTCCTTCTTGGCCCCCAGTAGGTGCCTGGCTACTGTGTCCAGCTTGTAGTCTGAGAGGCTGAGCTTGTC | 4 |
| wt-154950-155070 | 2715 | CCGGCACACGGCGTACATGTCGATGGGGATGAGGCCGGTGATGCGGACCTTGGTGTTGGCCCGCAAGAAGCCCTTGCCCGCATCATGGGGTCGCCTGACCTCGCAGACGCCCCCAGCCCT | 8 |
| wt-155070-155190 | 2716 | AATTTTGCCCAGAGAGGCTGGGTTGATGCTGTAGATGTGCCTGGCTCTGTCCAGAATGTAGGGCCAGTCAAAGTTGGCCACGTTGTAGCCGGTCACAATCTCCACGCTGAGGTCTCTGAT | 4 |
| wt-155190-155310 | 2717 | GAGCTGGAAGAAGGCGTAGAGCATGTCCAGCTCCGATGGGAACTCGTAGACCTCAACCCCCTCTATGTCTTCGCAGGTGCCCAGCGTCAGCAGGATGCGCCTATAGCGCCCGGCCTCCTC | 4 |
| wt-155310-155430 | 2718 | CCCTGTCGACCAGAGGACGCAGGATATCTGCAGGATCAGGTCAGCCTCGTTGGTGGCCGTGGGGAAGCCCTCCTCCCCCAGACACTCGATATCGAAGGCCAGGGCCTGGTAGGAGGGCCA | 4 |
| wt-155430-155550 | 2719 | GGAGCTGTCTTCACGCCGGACCGAGAGGTCGCCCACCTCACAGTCGTACTCGAGCTCGGCGTACGAGTCCCGGTGCTGGAGGCGGGGGATGGCGCGGCGGCAGCTGTACCAGCCAAAGGT | 8 |
| wt-155550-155670 | 2720 | GACAAAGTCATTGTCCAGGACAAAGCGGCGCGTGGCATCCACGTTGGCCTCAAAGATCCGACACCCGTGCTTGTCTTGCAGCCACGTGGCCACGTGACACACACTGTTGGGATGGGAGAG | 4 |
| wt-155790-155910 | 2722 | GAGGACTGCAAACTCCACGTCCAGACCCTGAGGCGCGCTGGCGTAGAAGTAGGCCTGCTGCCCAAACACGTTCACACACACGCTGGCCCCATCGGCCTTGCGCCGGCCCAGTAGCTTGAT | 4 |
| wt-155910-156030 | 2723 | GACGATGCCACATGGCACCACATACCCCTGTTTATCCGATGGAATGACGGCGCATTTCTCGTGCGTGTACACCGTCTCGAGTATGTCGTAGACATGGAAGTCCAGAGGGCTTCCGTGGGT | 4 |
| wt-156030-156150 | 2724 | GTCTGCCTCCGGCCTTGCCGTGCCCTCTTGGGCACGCTGGCGCCACCACATGCCCTTTCCATCCTCGTCACCCCCCACCACCGTCAGGGAGTCTTGGTAGAAGCACAGGGGGGGCTGAGG | 8 |
| wt-156150-156270 | 2725 | CCCCCGCACATCCACCACCCCTGCGGCGCCTGGTGTCTGGAAACACTTGGGAATGAGACGCAGGTACTCCTTGTCAGGCTTTTTCAGAAGGCCTTTATTAGGTCTTAGGAAAGGGTTATA | 4 |
| wt-156270-156390 | 2726 | GAAGAGTCCCCCAGACATGGTTAAAACTCAGTCTCTGCCTCCCCAAGCAGTGCGGCGGCGGTCTCTGGATCGTGATAGCGTCTTCTGCGTAGGCCTGGAAAACGGTCCCTGGCTGCCTGC | 4 |
| wt-156390-156510 | 2727 | AATGCTCTGCTGGCCACTGAGGGTCCGGCCGCCCTCTGAGCTGCTCTCTTTTGCTCCTGGTTTTGCTCATGCAGCGCTAACATGATGGCTTGTAATTCTGTCTTACTAATGGGATTAATG | 4 |
| wt-156510-156630 | 2728 | CCTGGACCCTCACCAGAGGCATGTTGCTGAGCGAGCTCGTCGATCCCGGGGTAGAGCATCTGCACCGGCTGCTGCGACATCTGGCGCGTGCGCCTCGTGAGGGAAATAACCAGGATCACC | 4 |
| wt-156630-156750 | 2729 | ACCCCCGCCACCAGGACCAGAATGAGCATGCCGCCGAAGGGGTTTTTGAAGAAGGAGATGAAACCAGAGACCAGGCTGCTAAACAAACCCCCCACCGTGCTGACTAGGTTGGTGATGGAC | 4 |
| wt-156750-156870 | 2730 | TGACCCACGCTACCCAGACTGTCCATAAGTTCCCCCAGGCCGTCCACGAATTGATTTCTTCCGTTTGACACTGCATTGTCCAAATCCTTCCGCAGGCCGGCGATGTTTTGCGCCTGGAAG | 4 |
| wt-156870-156990 | 2731 | TTGTACTCCCGGAAGATGCCCTCCAGGTCAAAGACGTTGGAGGCACGCTGTTCGTCCCGTGAGTACAGCTCCAGGGAGGCAAAGTCAATGTTCTCGATGAGGGAGGTGTTTAGTGAGATG | 4 |
| wt-156990-157110 | 2732 | AAGGTCTGCAGGGTGGCAATGCCGTCCAGCTCGATGGTTTTAAAGTGGTGGTAGTCGTTGTAGACGTGGATCTCGTTGCCGGACTGGAAGTAGTACTGGCTGGTCGCCTGGCACACCTCC | 4 |
| wt-157110-157230 | 2733 | GTCATCTTTTTTGTGAGGAAGATCTCGTTGTCGGTGCCCAGCTGTCCCTCGTAGGTCTTGGTGTCGTTGATAAAGCTGAAGGACACCAGGGGGGCGCGAGTAGCACATGGTCTCGGAGCCA | 4 |
| wt-157230-157350 | 2734 | GGGACCCTCATGCTCTTGCGCAGGGTGACGGTGGCCTGGTTAACGGGCACGCACTGGGAGACTGAGATGACATCCCCCAGGCGCTTGGCCGCCACCGCCTTACCGTAGATGCTGGACATG | 4 |
| wt-157350-157470 | 2735 | ACGGTGGTTGGATTAATCTTGGTTAGTTCTCTCAGCACCATGTTCTGCCTCTTCTGCTCCAGGCACCAGGCCCGCGCAAGGTCTCCCAGCATGCGGTTGATCTGGCGGCGCAGGGAGTCG | 4 |
| wt-157470-157590 | 2736 | TAGGCAAATTGGATCTGGACGGTGGCGGGATTGTTGAGGGTGCCCAGGGACTTCCCGGGGGCCGTGGGGGGCACCGGTGTGGTGGCGTTCCCCGCATCCCGCCTCCGACGCCTCAGAACG | 8 |
| wt-157590-157710 | 2737 | GCGGCGGGGGTGCTCCCGCGGGCCGCGGATGGGGCTGGGGGCGATGGACTGCTGGGGGGTGAGGAAGTCGGAGTGGTAAGCTCCGTCAGGTTCTTGACGGTGGCCAACGAGCGCGGGGTC | 8 |
| wt-157710-157830 | 2738 | AGAGGTAGCCAAGCTAATAACAATCCTCCGCTCGTTATAAAATATGTAATGGCTTCCTGGCCCTTCGTGTAACGATCCTGGACGGCCTCGTACTTCTCATGCATGGTCTTGTTCACCTGC | 4 |
| wt-157830-157950 | 2739 | TCTTCGATGCACTTGAAGGCGTCCGGGAGCTCTATGCCCACGGTTGTGTTGGTCACGAAGCTAGAGGTGCCCTCGTCAGTCACAAAATGTATTGACTTCCCTGTTTCTGTGGCGATGGTC | 4 |
| wt-157950-158070 | 2740 | GAGTCAAAGGTTTGCCAGTGTTGAAGCGGGCAGTAGGCTGTCCTGTTCTCGAGCTTCCAAGATAGCGTGTAAGTGCCCTTGTCCAGGAAGGCTCGGCGTTCGCCTTGCGGGTTCGTCCCT | 4 |
| wt-158070-158190 | 2741 | CGGTTGTCGTAGTCCACTATCTTGTAGTTAGTTCTCACGTGGAAGGAGTCTGCCCGCTCATGGAAGGTTTCCTTATTTTTCCCGTCATAGAAAGGGGACATTTCCACAGTCTGCCCGGTG | 4 |
| wt-158190-158310 | 2742 | GTGGTCACAAAGAAGTCGAAGGGGCTGTTGGACTTGGCCATCATGTCAGTTATCAGGCAGTTGACGGTAGTTCTTGTTCTGTAAGTCCATATCAACCACCCGGGGGCGTCATAGAGCTCC | 4 |
| wt-158310-158430 | 2743 | GTCTGGCTGGCGTAGCGGCGCACCCCGTTGGCCAGGCCCCCGGTGGGCTTTAGGTTGACGGTGATGTTAACTCCGTCGCGGTCTACATACACGCGCGTCAGCCCATCTTTTGTCATCTTG | 4 |
| wt-158430-158550 | 2744 | ACCGCGTTGTAGCACTGGTAGATGGTATCCATCTGGTCAGTTTCGTAGCTGTCAACGGAGAACTTCTCCTCGTGCCGGTTGGTCACGGAGTCCGCGTACCAGCCATTGTAGATGAGAATG | 4 |
| wt-158550-158670 | 2745 | TTGGTCACTATCTTGGTGTAGGAGCGGACCTTAAACGAGTAGGGAATAATGTTGTCTTTAAACACCATCAACAGGCCCTCCGTGTGATTCTCCCGCGTGCCAAACGAGGGACACTGGATG | 4 |
| wt-158670-158790 | 2746 | TCCGAGGAGAAGCGGAACAGGTCGCCGTGGCTGGAGAGCTCGCAGACTCGGAAAGGAAAGCTGGTTTGCTGACGCGTGGCGGTAGGCTGCACCGTGGTGGCGGGGGGTGCGGGCTGCTCT | 8 |
| wt-158790-158910 | 2747 | GGGGTCTGCGCACCGAGACGGCACGCCAGGGCGGCTAGCAGCACGACCACGCTTAGCACCCTACGCCGAGTCATCTCTCATTTGGAGGTGCAGGTAGAGAAGGGCATATAGATCCTTAAA | 4 |
| wt-158910-159030 | 2748 | TACCCACCCCCTGCCCTTATACAGAAGAATTAGGGGGCGGTCAGAGTCGTACGTGAGGTAAAGCCCATCCGGGGGCAGGGCCTGGCCGGGGCTGACCGCGTCCGCCCGGCGCAGGATCAA | 8 |
| wt-159030-159150 | 2749 | GGACCGCCCCCAGGTCTTGTTGTAGAGGGACACGGTTAGGACGGCCTCGCGCAGCGCCCGGCACAGAATTTGCTGGCTAGATGCCAGTGAGCCCCCGGGTACGCTGTAGAAGCTGTTGAA | 4 |
| wt-159150-159270 | 2750 | GGAGGTCTCTATCCAGTCGCTCGGCTCGATGCCTGGCCATATCAGGGAAGTCAGGAATGCCTTCTGGTGGGGCAGCGTACCTGCGGCGTCACAGCAGCGAGCCAGGGCCACGTTGCTGGG | 4 |
| wt-159270-159390 | 2751 | TGGGGGAAAGAGCCCGCTCTCCTCCGCCAGGGGCCCCGTGATGAAGGTGTACAGGCTGTGCGTCAGCGCGTGCAGGTGCTCCGAGCTCAGGGTCTGGGTAAACAGGTGTGTTTTGATGTA | 4 |
| wt-159390-159510 | 2752 | CTTGGAATTCTCAAAGGCGGCACCCTCGCCGGCGCGCCTGTCCTCCCAGGGACCCGAGACGAAGGCCCGTCTGTAGAGGAAGTGGTTGCGCATGCGGGCCAGCTCCCAGTAGACCACGTC | 8 |
| wt-159510-159630 | 2753 | CCCCCAGACGCGCAGGCACAGGGTCTCGGTCAGGGTCTCGCTCTGTTGCGCCAGGCAGGACTGCAGCTTGGCCAGACCCTCGGTGGCCACCTGGCGCAGGTACTGCTCCTTGCGCTTGAG | 8 |
| wt-159630-159750 | 2754 | CGCGTCCGAGAGGGCGCCGGACGGGCCGGGCTCTCGTGCCCCAGCCGGCCGGGGCACCTCCGGGCTCTCCCGGGACGCCTCCTCCTCGCCTCGGCCCAACCGCTGCATGGCTCGGTTGAG | 12 |
| wt-159750-159870 | 2755 | CCGCGTGTACAGCTCGTTCCTCTTTTGCAGGATGGCCCGGTACTGGGGGTGCGCCGTGAAGGCGGCGGCGCAGTCCGCCTTCAGCGCCTCCACCGCGTCGCCCGAGGAGCTGTAGACCCC | 8 |
| wt-159870-159990 | 2756 | GCCGCAGAAGAGCCGCTCCGTGGCCCCGGGAGCCACGGCGTCAAACAGGTGAGTCAGCCTTGCCCCCGCCAGCGCCTCCTCGCAGGCCCCCCGCACCAGGGCCAGGCGACGCTCCCGGGC | 12 |
| wt-159990-160110 | 2757 | AAACAGGGCAGAGAGGCGGGAATGGCCGCCACCCTCCCCCTGCCCCGTTGCACCGATAGCATGGCCGCCAGAGTTCCAATAGAGGAGCTCCGAGAGCTCCGCCACCTCCGGGGGCACTGT | 8 |
| wt-160110-160230 | 2758 | CGAGAAGACGTTGTAGGTGTCCAGCGCTCTGGTCGCCCCCTCTGCCTCCGGCCGCCCCGGGCCCGGGACCGCGCCCTCCTCTGGGCCGCCCGGCCTCGCCTTCTCCTCAGCCTCCAACAG | 8 |
| wt-160230-160350 | 2759 | GTGCCCGAGCCCCGCCTGGCGGACTTCATTCTCAAACAGTCCCGAGACCGGCTCCGGATTCACCGGCACCGCCAGGTGGTTACAGGAGACGTGGGTCCCCTCTGCCGTGGAAGGGTTGCC | 8 |
| wt-160350-160470 | 2760 | GTGGTTGGGCAGAACCATCAGCTCGCCCACACAGCGCCAGCAGGGCACAGAGGTGATGTAGAGGCGCGGGTCTGGGATGGGACTTACGCCCCGAAAGCGGCCCAGCAGATCCAGGGCCCG | 8 |
| wt-160470-160590 | 2761 | TTCCAGGCTCTCCAGCCCCATGGTGTGAGACATGCAATAAAACACGCTATTGATTCTCTTCATTAAAATCTCTATGTCATTTATTAGGCACAAACTTACATCGACTTTATGCCCCCCGTA | 4 |
| wt-160590-160710 | 2762 | AAACTCCACAGAGTACGCGACTGAGGGGGTACGGAGAGGCGGGACCCGGGTACCCTTTCTACCAGGGGCGAGCAGCGCGGCAGAGGCCTCTCTCGAGTTCTCTAGCAGGTGCACCAGCTC | 4 |

| | | | |
|---|---|---|---|
| wt-160710-160830 | 2763 | CAGGGACAGGGCGCTGCATGCACGGTCATTCTGCCGTCTCAAACGGGGAAGGAGGATGGCCTCCAGCTCGGCCAGCAGGCCGGCGTTGCGCACCACCGCAGCCACGTCCAGACTCCGGGG | 8 |
| wt-160830-160950 | 2764 | GTCCAGCCGGGTGCACACGCTCAGCTCAACCGCCAGGGCGTACACCTGGCTGTACGCCGCCGCCAGCAGCCCCGACATCGCCGCCCCAGGGGTCTCTAGACCTCGAGTCCGGGGAGAACG | 8 |
| wt-160950-161070 | 2765 | GTGGCCAGACGGCGCTTGCGTCTGCCCCCGGAGCCCTGCCCTCCTCCACCCAGCAGCAGCCCGGCCGAGGCCTGCGACGCGGTGCTGACCGGCTCGGCCACGCTGATAAAGTTGTCCTGG | 8 |
| wt-161070-161190 | 2766 | GCTGCCCCGGGCCCACCCCACACTCCCTCCAGAAAGTCCCGAGCGGCCTCCGCCGTCCACTCTATCCCGCTGGAGGCAATGGTCGCCAGGGTTTCTAGGACGCTGTCCGCCAGGACGGAG | 8 |
| wt-161190-161310 | 2767 | AAGCGGCCCAATAAGTACTCCGCGTCGTCCCTAGTCAGCGAGGCGCATGCCTCGCCCATGGCATCCACAAGGTTGCACACCACATCAAACACACAGTCTTCCTCCTGTTTTTGTGATATA | 4 |
| wt-161310-161430 | 2768 | ATGGCCTCCAGGCCAGCCCTGATGTTCTCAATCTCATATGTGGTCGCGGCTTGGGTCCGGCGCTTCACGGTCAACCCTAGGGTGGGGGTGGCAAAGACAAACTTCTTCCGCATGGAAGAG | 4 |
| wt-161430-161550 | 2769 | CCCCCGGCCTGCTTGCGCAGCCCAGCCCCGGGGGCCTGCAGCAGGTTCCTGTCCACGCCCCGGCCCATAAAGTATCCCAGGTTCCCGGCCTGGAATATCTGGTTGTTGCCGTTGACCCCC | 8 |
| wt-161550-161670 | 2770 | GTGTACTTGTTGATGGTCACTGGCAGCGTGACAACCGGACGGGCCTTGCAGACCTGGCTAAGACAGTCTGTGGCCGCGCAGACCACCGTGGTCGCAGTAAGGGAGGAGGTGGCCTCCGCG | 4 |
| wt-161670-161790 | 2771 | TAGGCCGCTGCCGACTCCACCGCCCGCGTGCCCAGTACGTGGGGGTAGTCACGGGCGGGCACCGACTGCGTCCTCGGCACCAGTCCCTGAATCAGGCTGATGTAGAACTGGGTCTGGCCG | 8 |
| wt-161790-161910 | 2772 | CACGCCTTCAGGATGGCGTTGTTGAGCCTCTGCTTGGCGTAAGTGACCAGGTTGCCAGGCACCACATCTATGACGTTGCTCTCTTCGTGGGCCCGGGAGCCCCCGTCCACAAAGAGGGCC | 4 |
| wt-161910-162030 | 2773 | AGGTCAGAGTACTCCTCCGCGCTGGCCCCGCTGGGGACAGGGACCGAGCGCCGCCTGGAAAAGTTGTGCCACAGGTACAGGCTTGAGAGCTTAGTGTCCGGGAATAGGGTCTTGTGGTAG | 4 |
| wt-162030-162150 | 2774 | GTGTTGAGGAATTTCATGTAGGGCCCGTTGATGATGTAGTTCTCCCTCCTGGTAGTGGACTTGATGAAGCTGTTCTGGAGGGCGGCATTCTCCCCCGTGAAGACCACCCTGTTCTTGATC | 4 |
| wt-162150-162270 | 2775 | TTGATGTTCCTGGGGCACAGCATCAGCACCTTGGACATGCGCACAGGCAGCCGCCGGCCGTACACCCGGCCCTGCAGGGCCGCGTCCAGGTCTGGCAGGTCGCAGGTGGGCTCCCCATGC | 8 |
| wt-162270-162390 | 2776 | ACCACCTTGGCCTCCTTGGCCGTGAGGACCCCCTTGTCGATGGCCAGGCTCCTAAAGTTGGTGCACAGCGTCTGGTAGTGACCCTTTAGCCACTCTGGGGGGCTCTGGCCAAGCCCGGGG | 4 |
| wt-162390-162510 | 2777 | TTGTCATTCTCATAGCACATACAGATGGGCAGGGAGATGTCCTGCAGGATGGTCAGCAGTGAGCGGTAAAACAGCTGGGTGAAGATGGGGCAGGCGGGCTGCGCAAAGGGGTTGCACGAG | 4 |
| wt-162510-162630 | 2778 | TACTGCATCACGTGGTAGCAGCTCTTGACCAGGTCCTTGTAGGTGATGTTGTTCTTGGCCATGCTGTTCATAAACTGGACCACTTCGGCGTCCACCGCCGCATCCACGTCCTTGAACATC | 4 |
| wt-162630-162750 | 2779 | TTGACAAAGTCACGCGGGCCATGGGGCTCCTTCTCTAGCTTTCCTTCAGCGTCTATGCCCAGCCGAGACAGCCGCTCCAGCAGGTTCTGGTTCAGCTGCCAGTAGGTGTAGCGGGGCTCG | 4 |
| wt-162750-162870 | 2780 | TCGTCCGGCCGCTGCCCGTCGTCCTCCTTATCGATGAAGTTGAGAAAGTTGCCCAAAAAGTCCGTCTCGTTGTAGGAGCCCGAGGCCCCCGAGATCACATAGGGGTCCCTCCGCTGCGTG | 4 |
| wt-162870-162990 | 2781 | GACATGACGGGGGGGAAGCGGTCCCTCAGCCTAAAGAAGAGCGTGTTCAGGCACACGGCCGGGGCCCGGCCCTCGCAGAGCGAGCACATGGGACTGGCGGCCGCCCCCGCCACGTAGCTG | 8 |
| wt-162990-163110 | 2782 | CCCGTCTCCGGCACCGGGGTCAGAGAGCTCTTCTGTCCCTGGCAAAACTGCAGGTAGTAGGCATAGCGGGCAAGAAGGTTGGGCGAGAAGGAGGCCGCATAGACCAGGTGCTCCACAGCG | 4 |
| wt-163110-163230 | 2783 | TAGTTTCCCGGACCGTTGGTTCCGGTCACGTCTGGCCCACCCCAGCCCGAGAAGCAGGGTCGGCGGCAGGGGTCCCAGGTCCCCTCCTGCAGGGTCCCCAGGCCGTGGGTCATGTAGAAA | 8 |
| wt-163230-163350 | 2784 | CTGTTAAAGAGACTCTCCTTGCCCTGACCGGTTGACTTCGAGACCCCCGAGACGTAGAGGACGGAATTGGTGGCAAAGATCTGCGTGGACACGTGGGGGGCCAGGCTGGCATTATATCGG | 4 |
| wt-163350-163470 | 2785 | TGTAACGCAGCCACACGGGCCTCTGGACCCTCACAGTCGGCAAACAGGGGCCACGAGTCGTAGTTGAGGCTGGCCGGGGTCTCGTGCGAGGCCTCCAGCATGGCGGGTGCGTAGCTCACC | 8 |
| wt-163470-163590 | 2786 | GCCAGCTCGCATGCCGCGCTGTCCACAATCATTAAGGCTCCCGAGTCCGGGTGACTGATGGTTGAGGCTGGGAACTCCTTGAGGGGGGCCACCTTGGCCACCTTGGCCTGGTCCTGCAGG | 4 |
| wt-163590-163710 | 2787 | CTCTGCTTCTCCAGCAGCTCCACCAGCTTGCCCACCCGTCGGACGCGCAGCGCCTGCGCCAGCCCGGTGTACAGCGCCTCGTGCATGCAGCGGCTGAGGTCCGAGTTGTAAAACTGGCGG | 8 |
| wt-163710-163830 | 2788 | AGCTGGGGCACGCCCTCTGGGAACACCTCCTTGTCGTAGAGCGGGACCCTAACGCTCGCAGACTGCCCCACCGCTACCTCCTGTTTTAACGATGGAATGGCCACCAGGTTTCCGCTGTAG | 4 |
| wt-163830-163950 | 2789 | AGTCGCTCCTTGAAGGCCTCGGTTATTGCCACCGCCCCCAGGTAGGCAGAGGGATCTAGCCCTTCGGGGAAGAAGTCCCCCGGCTTGGAGCTTTCCCTCGGTAGGGCGCTGTAGGCGTCG | 4 |
| wt-163950-164070 | 2790 | TACCCAAACACCTCCCTGGTCTCGCCACAGAGGGCCTCGAGACCCGGCCCCTCAAAGATGGGGGGAACCATATGGGCATTGTGGAACACGTAGATGTCCCTGTGATAGGAGGTAGCGCGT | 4 |
| wt-164070-164190 | 2791 | AGGAGCCCGCAGTTGGGGTCGGGCCTCCTGTGCAGAGCCTTGACATTGATGCTGAAGCCCGGCTCCACGGTGATGCCGCAAAGGAGCGGCACCGTCAGGCACCTGTGGCCCGCGTAGCCG | 8 |
| wt-164190-164310 | 2792 | GTCCCCAGTGTGGCCACCTCCCTAAGAGGGTAGGTGGCCAGGGGGTAAAAGTAGATGTAGCCGCACGGACCCGGCTGGCTCTGGCTGCCCAGATTATCCTCGCTAGTCTGTGCACCCTGC | 4 |
| wt-164310-164430 | 2793 | ATGATGCCCAAGGTATCGCCCCGGCCTCCCAGTCCCACATTAAATGTTACACTTTACTCATCACGCAACACCCACTGTTTATTCATTTACAAAGATTTCAGGAAGTCAGTCAGGCTGGCC | 4 |
| wt-164430-164550 | 2794 | AGGGCCCACGTCACGGGGAACTGACGTCTCAGCGATCTTGGCATGCCGCCCAGCCTCGCAAACCAGAGTCTGCGATAGAGGGCCAGGTAGTGGGCGATTGCCCCCAGCACGAAGGCGGCG | 8 |
| wt-164550-164670 | 2795 | CTCTTGTGGTCATCCAGGTAGTTTCGCACCGCAAACACCACTGTGTAGCACAGCACCACCCTGAGCCGCGACCAGTAGTCGTAGTGGTCGTTGTACACTGCGCGCAGGACGCTGATGATG | 4 |
| wt-164670-164790 | 2796 | AGCCGTACGTGCGTGTCTTTGCCCCCGATGTCGGCTGTCCTGCAGGCCAGCTCCGCGTACAGCTTCCTATCCTTCCTCAGGGAGGCCTTGATGAGCCGGCAGAGGACCAGGGCTGGCAAA | 4 |
| wt-164790-164910 | 2797 | GGCAGGTCTTTCTCATCCCGGGTGAACACCGCGTACATGGCCCTGAACATGAGGTAGCTGGACTCAGCCACCTTGTCGTCCGGCGGCGAGGGCGCGACCCACGCCTCGACCGGGGTCCTC | 8 |
| wt-164910-165030 | 2798 | ACAAACACAGAATCTGTAGACTTGGCTGGCCTCATGGTCTCGTCAGGCCAGCTCACGGGCTTCAGGCTTATATGATAAAATGGGCGTGGCAGAATAGTATAAGACGCGAGGCCTGGGTGA | 4 |
| wt-165030-165150 | 2799 | GGAGAGTCCAGAGCAATGGCCAGGTTCATCGCTCAGCTCCTCCTGTTGGCCTCCTGTGTGGCCGCCGGCCAGGCTGTCACCGCTTTCTTGGGTGAGCGAGTCACCCTGACCTCCTACTGG | 4 |
| wt-165150-165270 | 2800 | AGGAGGGTGAGCCTCGGTCCAGAGATTGAGGTCAGCTGGTTTAAACTGGGCCCAGGAGAGGAGCAGGTGCTTATTGGGCGCATGCACCACGATGTCATCTTTATAGAGTGGCCTTTCAGG | 4 |
| wt-165270-165390 | 2801 | GGCTTCTTTGATATCCACAGAAGTGCCAACACCTTCTTTTTAGTAGTCACCGCTGCCAACATCTCCCATGACGGCAACTACCTGTGCCGCATGAAACTGGGCGAGACCGAGGTCACCAAG | 4 |
| wt-165390-165510 | 2802 | CAGGAACACCTGAGCGTGGTGAAGCCTCTAACGCTGTCTGTCCACTCCGAAAGGTCTCAGTTCCCAGACTTCTCTGTCCTTACTGTGACATGCACCGTGAATGCATTTCCCCATCCCCAC | 4 |
| wt-165510-165630 | 2803 | GTCCAGTGGCTCATGCCCGAGGGCGTGGAGCCCGCACCAACTGCGGCAAATGGCGGTGTTATGAAGGAAAAGGATGGGAGCCTCTCTGTTGCTGTTGACCTGTCACTTCCCAAGCCCTGG | 4 |
| wt-165630-165750 | 2804 | CACCTGCCAGTGACCTGCGTTGGGAAAAATGACAAGGAGGAAGCCCACGGGGTTTATGTTTCTGGATACTTGTCGCAATAAACGCACTTGCCTATTTCACCTTGTTTTAGTGTGGCATTG | 4 |
| wt-165750-165870 | 2805 | GGGGGGTGGCATTGCGGGTGGATAGCCTCGCGACTCGTGGGAAAATGGGCGGAAGGGCACCGTGGGAAAATAGTTCCAGGTGACAGCAGCAGTGTGTGAAGATTGTCACAGCTGCTGGTT | 4 |
| wt-165870-165990 | 2806 | TGGAGAAAACGGGGGTGGGCGGTGATCAGGGAGAACAATTCCCCGGGGACACCTGCACGAGACCCCTGGGCTCTCAGGAACTCCGCCCAGGTCTTGCCAATTGGGGTGATCCTGTAGCGC | 4 |
| wt-165990-166110 | 2807 | CGCGGTTTCAGCATCACAGGTTATTTTGCCTGAAGCTTGCTGGGGCGTAAATCCCTCTCGCCTTGTTTCTCAGAGAGCATTTCAGGCCGGTTTTGCAGTCGCTGCTGCAGCTATGGGGTC | 4 |
| wt-166110-166230 | 2808 | CCTAGAAATGGTGCCAATGGGCGCGGGTCCCCCTAGCCCCGGCGGGGATCCGGATGGGTACGATGGCGGAAACAACTCCCAATATCCATCTGCTTCTGGCTCTTCTGGGAACACCCCCAC | 4 |
| wt-166230-166350 | 2809 | CCCACCGAACGATGAGGAACGTGAATCTAATGAAGAGCCCCCACCGCCTTATGAGGACCCATATTGGGGCAATGGCGACCGTCACTCGGACTATCAACCACTAGGAACCCAAGATCAAAG | 4 |
| wt-166350-166470 | 2810 | TCTGTACTTGGGATTGCAACACGACGGGAATGACGGGCTCCCTCCCCCTCCCTACTCTCCACGGGATGACTCATCTCAACACATATACGAAGAAGCGGGCAGAGGAAGGTAAGAGTGCCA | 4 |
| wt-166470-166590 | 2811 | TCTATCTGTACTTTTATTTATTGCATCACAAGTCACATCAATAATAAGGGCGCCATCTAGCGGGAGATGTTATCCACACCATCCCAATTCACATCTCAGGGACAACAGGTCAAAGTTCTT | 4 |
| wt-166590-166710 | 2812 | TGTTGACACCCCCAGCGCTGGCTCCAGGGGGTGGAAGCGTTGGATGCAGTCCTCCGCATCGGGGCGGACGCCTCCTCCCAACGCGTTTCTGCGGATCAGTCGCTGGCTGGTGGGCATCGG | 8 |
| wt-166710-166830 | 2813 | AGTCGGTGGGCGGTCCTCCACGGGGACACGCTCCTTCTTGGCCTTGTTCTTTGACCTTTTGGACATTCTTCTGAAGGAACGGCGGAGAGTAGCGTAGAATCCAGCCAGTGGTCTACCCGG | 4 |

| | | | |
|---|---|---|---|
| wt-166830-166950 | 2814 | TCGCATGGTGGCTTCTTAGATGAGGAGCAGGCATAAAAGTCCAAACAGGACACAGAGTACCACCAGGAGTAGTCTTAGTCTGCTGACGTCTGGGTCCTCGGGGCAGGGGTGGCTAGGCCT | 4 |
| wt-166950-167070 | 2815 | GGTCTCCGTAGAAGAGCCGGGCAGGCCGCAGGCAGAGGACTGCTGCTCTAGCAAAGCACGCTCCAGGACGTGTACCATCTCGAGAGTGAGGCACAGCTGTTTTCGTGGACTTTTATACAG | 4 |
| wt-167070-167190 | 2816 | TAAGGACAAGGAAAGAAGGCCAGAGGAATGTGGAAAGATGAGCGAGGACAGGTGTGGAGGTTTTGGGCTAGCTCTTAGTTTCTGGGTGTGAGAGAGGGATTAAAGTGCTTATGCGCAAAG | 4 |
| wt-167190-167310 | 2817 | AATGTGTCAACAACAGGTGTTCCTGCCTCTGCTGGCATGAGTTAGGTGTGGCTTGGGCTGAATCCAAATGTGTATTGGCACAAGATGGAAAGCAAAGTTGCTGGAGTTACTGGGTGGGAG | 4 |
| wt-167310-167430 | 2818 | ACAGGGATGTATGTGGTCCCCCGCTGGTATGCCAGTACCCTGTGGAAGTAAGGGGCCTCATCTGCCTGGTAGTTGTGTTGTGCAGAGGTCTGATGTGTGTAGGAGGGGTGGGTTCAACGC | 4 |
| wt-167430-167550 | 2819 | AGGGGCGTTGGTGGCGGAGTCTGGCAACGCCCGGGTCCTTGCTACCTGTGTGGTGTGTTAAGGGCTGGGTAAAGGTGTCTGCCAATTCTCGCATGTCCTCCTTTCCCCTTGTTTTGAAAT | 4 |
| wt-167550-167670 | 2820 | AGAATATGAATGTGGCTTTTCAGCCTAGACAGACAGTGTGGCTAAGGGAGTGTGTGCCAGTTAAGGTGATTAGCTAAGGCATTCCCAGTAAATGGAGGGAGAGTCAGTCAGGCAAGCCTA | 4 |
| wt-167670-167790 | 2821 | TGACATGGTAATGCCTAGAAGTAAAGAAAGGTTAGTCATAGTAGCTTAGCTGAACTGGGCCGTGGGGGTCGTCATCATCTCCACCGGAACCAGAAGAACCCAAAAGCAGCGTAGGAAGGT | 4 |
| wt-167790-167910 | 2822 | GTGGATCACCGCCGCCATGGCCGGAATCATGACTATGACCGCCGCCTCCGTCTGTCATCAAAGGCGGGCCCTGGTCACCTCCTTTGTTTTCAACCTCTTCCGTCAATTGTGGAGGGCCTC | 4 |
| wt-167910-168030 | 2823 | CATCATTTCCAGCAGAGTCGCTAGGGCTATGAGGCAGCGGGTCATGTGGGCCATTGTCATCAGTGTTGTCAGGGTCCTGTGGGCCATTGTCATCAGTGTTGTCAGGGTCCTGAGGCAGCG | 4 |
| wt-168030-168150 | 2824 | GGTCATGTGGGCCATTGTCATCAGTGTTGTCAGGGTCCTGTGGGCCATTGTCATCAGTGTTGTCAGGGTCCTGTGGGCCATTGTCAGGACCACCTCCAGGTGCGCCTAGGTTTTGAGAGC | 4 |
| wt-168150-168270 | 2825 | AGAGTGGGGGTCCGTCGCCGGCTCCACTCACGAGCAGGTGGTGTCTGCCCTCGTTGGAGTTAGAGTCAGATTCATGGCCAGAATCATCGGTAGCTTGTTGAGGGTGCGGGAGGGAGTCAT | 4 |
| wt-168270-168390 | 2826 | CGTGGTGGTGTTCATCACTGTGTCGTTGTCCATGGTAATACATCCAGATTAAAATCGCCAGAAACAGGAGGAGCCAAAGGAGATCAACCAATAGAGTCCACCAGTTTTGTTGTAGATAGA | 4 |
| wt-168390-168510 | 2827 | GAGCAATAATGAGCAGGATGAGGTCTAGGAAGAAGGCTAGGAAGAAGGCCAAAAGCTGCCAGATGGTGGCACCAAGTCGCCAGAGCATCTCCAATAAGTAGATCCAGATACCTAAGACTG | 4 |
| wt-168510-168630 | 2828 | CGTTGAAAAAAGAGTGTTAGGGTTGGAAAAGTGGGGGTGTGGTAAATAATTCCTAGGGAATGTTAGATCTTACCAAGTAAGCACCCGAAGATGAACAGCACAATTCCAAGGAACAATGCC | 4 |
| wt-168630-168750 | 2829 | TGTCCGTGCAAATTCCAGAGAGCGATGAGCAGGAGGGTGACTGGGGAAAGAGGAGAAAGTGCGTTAGAGAAGGAAGAGTAAGGGAAAGGGGGTGTGGGGCAAAGGGTGTAATACTTACTC | 4 |
| wt-168750-168870 | 2830 | ATCAGTAGGAGTATACAAAGGGCTCCAAGTGGACAGAGAAGGTCTCTTCTGAAGATAAAGATGATCAAAATTATAATTATAAGCATGAGAGCAAAGGAATAGAGGACAAGGAGGGCTCCT | 4 |
| wt-168870-168990 | 2831 | CCAGTCCAGTCACTCATAACGATGTACAGCCAAAACAGTAGCGCCAAGAGGAGGAGAAGGAGAGCAAGGCCTAGGGAAGAGGAGAGGGGGGGTCCTCGAGGGGGCCGTCGCGGGCCCGGT | 4 |
| wt-168990-169110 | 2832 | GGGCCCCTCTCAAGGTCGTGTTCCATCCTCAGGGCAGTGTGTCAGGAGCAAGGCAGTTGAGGAAAGAAGGGGGCAGAGCAGTGTGAGAGGCTTATGTAGGGCGGCTACGTCAGAGTAACG | 4 |
| wt-169110-169230 | 2833 | CGTGTTTCTTGGGATGTAGGCCCGGGGGGATTTGCGGGGTCTGCCGGAGGCAGTACGGGTACAGATTTCCCGAAAGCGGCGGTGTGTGTGTGCATGTAAGCGTAGAAAGGGGAAGTAGAA | 4 |
| wt-169230-169350 | 2834 | AGCGTGTGTTTGTGTTAGAAAAGCGGGTCCCCGGGGGGCAAGCTGTGGGAATGCGGTGGCCAAGTGCAACAGGAAATGGAAAGGCAGTGCGGCAATCAGAAGGGGGAGTGCGTAGTGTTG | 4 |
| wt-169470-169590 | 2836 | CCGTAGGGGAGGCGGGTGAGGCGGCTAAGAGGGCTAAGGGTCGGCGGGTGACGAAGCAGCAGACGGCGGATATGGGAATTTCAGAATGAGGTGGCGGATTCAGGCGAAAAGGGTGTGGGC | 4 |
| wt-169590-169710 | 2837 | TGTGCGAGTGTCATGAGGCAGGCGCGGAAAGTCGCTGCGGCTTGCTGGGGCATGGGGGGCCGCGCATTCCTGGAAAAAGTGGAGGGGGCGTGGCCTTCCCCCGCGGCCCCCCAGCCCCCC | 8 |
| wt-169710-169830 | 2838 | CGCACAGAGCGGCGCTACGGCGGGCGGGCGGCGGGGGGTCGGGGTCCGCGGGCTCCGGGGGCTGCGGGCGGTGGATGGCGGCGGACGTTCCGGGGATCGGGGGGGTCGGGGGGCGCCGCG | 12 |
| wt-169830-169950 | 2839 | CGGGCGCAGCCATGCGTGACCGTGATGAGGGGGCAGGGTCGCAGGGGGTGTGTCTGGTGGGGGCGGGAGCGGGGGGCGGCGCGGGAGCCTGCACGCCGTTGGAGGGTAGAATGACAGGGG | 8 |
| wt-169950-170070 | 2840 | GCGGGGACAGAGAGGCGGTCGCGCCCCCGGCCGCGCCAGCCAAGCCCCCAAGGGGGGCGGGGAGCGGGCAATGGAGCGTGACGAAGGGCCCCAGGGCTGACCCCGGCAAACGTGACCCGG | 12 |
| wt-170070-170190 | 2841 | GGCTCCGGGGTGACCCAGCCAAGCGTGACCAAGGGGCCCGTGGGTGACACAGGCAACCCTGACAAAGGCCCCCCAGGAAAGACCCCCGGGGGGCATCGGGGGGTGGGGCATGGGGGGCCG | 8 |
| wt-170190-170310 | 2842 | CGCATTCCTGGAAAAAGTGGAGGGGGCGTGGCCTTCCCCCGCGGCCCCCCAGCCCCCCCGCACAGAGCGGCGCTACGGCGGGCGGGCGGCGGGGGGTCGGGGTCCGCGGGCTCCGGGGGC | 12 |
| wt-170310-170430 | 2843 | TGCGGGCGGTGGATGGCGGCGGACGTTCCGGGGATCGGGGGGGTCGGGGGGCGCCGCGCGGGCGCAGCCATGCGTGACCGTGATGAGGGGGCAGGGTCGCAGGGGGTGTGTCTGGTGGGG | 12 |
| wt-170430-170550 | 2844 | GCGGGAGCGGGGGGCGGCGCGGGAGCCTGCACGCCGTTGGAGGGTAGAATGACAGGGGGCGGGGACAGAGAGGCGGTCGCGCCCCCGGCCGCGCCAGCCAAGCCCCCAAGGGGGGCGGGG | 12 |
| wt-170550-170670 | 2845 | AGCGGGCAATGGAGCGTGACGAAGGGCCCCAGGGCTGACCCCGGCAAACGTGACCCGGGGCTCCGGGGTGACCCAGCCAAGCGTGACCAAGGGGCCCGTGGGTGACACAGGCAACCCTGA | 8 |
| wt-170670-170790 | 2846 | CAAAGGCCCCCCAGGAAAGACCCCCGTGGGGCATGGGGGGCCGCGCATTCCTGGAAAAAGTGGAGGGGGCGTGGCCTTCCCCCGCGGCCCCCCAGCCCCCCCGCACAGAGCGGCGCTACG | 8 |
| wt-170790-170910 | 2847 | GCGGGCGGGCGGCGGGGGGTCGGGGTCCGCGGGCTCCGGGGGCTGCGGGCGGTGGATGGCGGCGGACGTTCCGGGGATCGGGGGGGTCGGGGGGCGCCGCGCGGGCGCAGCCATGCGTGA | 12 |
| wt-170910-171030 | 2848 | CCGTGATGAGGGGGCAGGGTCGCAGGGGGTGTGTCTGGTGGGGGCGGGAGCGGGGGGCGGCGCGGGAGCCTGCACGCCGTTGGAGGGTAGAATGACAGGGGGCGGGGACAGAGAGGCGGT | 8 |
| wt-171030-171150 | 2849 | CGCGCCCCCGGCCGCGCCAGCCAAGCCCCCAAGGGGGGCGGGGAGCGGGCAATGGAGCGTGACGAAGGGCCCCAGGGCTGACCCCGGCAAACGTGACCCGGGGCTCCGGGGTGACCCAGC | 12 |
| wt-171150-171270 | 2850 | CAAGCGTGACCAAGGGGCCCGTGGGTGACACAGGCAACCCTGACAAAGGCCCCCCAGGAAAGACCCCCGGGGGGCATCGGGGGGTGGGGCATGGGGGGCCGCGCATTCCTGGAAAAAGTG | 8 |
| wt-171270-171390 | 2851 | GAGGGGGCGTGGCCTTCCCCCGCGGCCCCCCAGCCCCCCCGCACAGAGCGGCGCTACGGCGGGCGGGCGGCGGGGGGTCGGGGTCCGCGGGCTCCGGGGGCTGCGGGCGGTGGATGGCGG | 16 |
| wt-171390-171510 | 2852 | CGGACGTTCCGGGGATCGGGGGGGTCGGGGGGCGCCGCGCGGGCGCAGCCATGCGTGACCGTGATGAGGGGGCAGGGTCGCAGGGGGTGTGTCTGGTGGGGGCGGGAGCGGGGGGCGGCG | 12 |
| wt-171510-171630 | 2853 | CGGGAGCCTGCACGCCGTTGGAGGGTAGAATGACAGGGGGCGGGGACAGAGAGGCGGTCGCGCCCCCGGCCGCGCCAGCCAAGCCCCCAAGGGGGGCGGGGAGCGGGCAATGGAGCGTGA | 8 |
| wt-171630-171750 | 2854 | CGAAGGGCCCCAGGGCTGACCCCGGCAAACGTGACCCGGGGCTCCGGGGTGACCCAGCCAAGCGTGACCAAGGGGCCCGTGGGTGACACAGGCAACCCTGACAAAGGCCCCCCAGGAAAG | 8 |
| wt-171703-171823 | 2855 | GGCCCGTGGGTGACACAGGCAACCCTGACAAAGGCCCCCCAGGAAAGACCCCCGGGGGGCATCGGGGGGGGTGTTGGCGGGGGCATGGGGGGGTCGGATTTCGCCCTTATTGCCCTGTTT | 8 |
| wt-60-180 | 2856 | GAATCCAGTATGCCTGCCTGTAATTGTTGCGCCCTACCTCTTTTGGCTGGCGGCTATTGCCGCCTCGTGTTTCACGGCCTCAGTTAGTACCGTTGTGACCGCCACCGGCTTGGCCCTCTC | 4 |
| wt-180-300 | 2857 | ACTTCTACTCTTGGCAGCAGTGGCCAGCTCATATGCCGCTGCACAAAGGAAACTGCTGACACCGGTGACAGTGCTTACTGCGGTTGTCACTTGTGAGTACACACGCACCATTTACAATGC | 4 |
| wt-300-420 | 2858 | ATGATGTTCGTGAGATTGATCTGTCTCTAACAGTTCACTTCCTCTGCTTTTCTCCTCAGTCTTTGCAATTTGCCTAACATGGAGGATTGAGGACCCACCTTTTAATTCTCTTCTGTTTGC | 4 |
| wt-420-540 | 2859 | ATTGCTGGCCGCAGCTGGCGGACTACAAGGCATTTACGGTTAGTGTGCCTCTGTTATGAAATGCAGGTTTGACTTCATATGTATGCCTTGGCATGACGTCAACTTTACTTTTATTTCAGT | 4 |
| wt-540-660 | 2860 | TCTGGTGATGCTTGTGCTCCTGATACTAGCGTACAGAAGGAGATGGCGCCGTTTGACTGTTTGTGGCGGCATCATGTTTTTGGCATGTGTACTTGTCCTCATCGTCGACGCTGTTTTGCA | 4 |
| wt-660-780 | 2861 | GCTGAGTCCCCTCCTTGGAGCTGTAACTGTGGTTTCCATGACGCTGCTGCTACTGGCTTTCGTCCTCTGGCTCTCTTCGCCAGGGGGCCTAGGTACTCTTGGTGCAGCCCTTTTAACATT | 4 |
| wt-780-900 | 2862 | GGCAGCAGGTAAGCCACACGTGTGACATTGCTTGCCTTTTTGCCACATGTTTTCTGGACACAGGACTAACCATGCCATCTCTGATTATAGCTCTGGCACTGCTAGCGTCACTGATTTTGG | 4 |
| wt-900-1020 | 2863 | GCACACTTAACTTGACTACAATGTTCCTTCTCATGCTCCTATGGACACTTGGTAAGTTTTCCCTTCCTTTAACTCATTACTTGTTCTTTTGTAATCGCAGCTCTAACTTGGCATCTCTTT | 4 |
| wt-1020-1140 | 2864 | TACAGTGGTTCTCCTGATTTGCTCTTCGTGCTCTTCATGTCCACTGAGCAAGATCCTTCTGGCACGACTGTTCCTATATGCTCTCGCACTCTTGTTGCTAGCCTCCGCGCTAATCGCTGG | 4 |
| wt-1140-1260 | 2865 | TGGCAGTATTTTGCAAACAAACTTCAAGAGTTTAAGCAGCACTGAATTTATACCCAGTGAGTATCTATTTGTTACTCCTGTTTAGTTGAAGAAAACAAGCTATTGGATTGTAACACACAT | 4 |

| | | | | |
|---|---|---|---|---|
| wt-1260-1380 | 2866 | TTTACGCTTTGTTCCTTAGATTTGTTCTGCATGTTATTACTGATTGTCGCTGGCATACTCTTCATTCTTGCTATCCTGACCGAATGGGGCAGTGGAAATAGAACATACGGTCCAGTTTTT | | 4 |
| wt-1380-1500 | 2867 | ATGTGCCTCGGTGGCCTGCTCACCATGGTAGCCGGCGCTGTGTGGCTGACGGTGATGTCTAACACGCTTTTGTCTGCCTGGATTCTTACAGCAGGATTCCTGATTTTCCTCATTGGTAAG | | 4 |
| wt-1500-1620 | 2868 | TGTGACACCAACAGGTGTTGCCTTGTTATGTCACCGTTCTGACACATGACTTACATGGGTTTGGCTTTTGTAGGCTTTGCCCTCTTTGGGGTCATTAGATGCTGCCGCTACTGCTGCTAC | | 4 |
| wt-1620-1740 | 2869 | TACTGCCTTACACTGGAAAGTGAGGAGCGCCCACCGACCCCATATCGCAACACTGTATAAAGGTAAGTATTATTAAATTTTAGAGACACTATCACGTGTAACTTGACGTGCAAGGATGGA | | 4 |
| wt-1740-1860 | 2870 | AGAGAGGGGCAGGGAAACGCAAATGCCGGTTGCCCGGTATGGGGGCCCGTTTATTATGGTAAGGCTCTTCGGGCAAGATGGAGAGGCAAACATACAGGAGGAAAGGCTATATGAGCTACT | | 4 |
| wt-1860-1980 | 2871 | CTCTGACCCACGCTCCGCGCTCGGCCTAGACCCGGGGCCCCTGATTGCTGAGAACCTGCTGCTAGTGGCGCTGCGTGGCACCAACAACGATCCCAGGCCTCAGCGTCAGGAGAGGGCCAG | | 8 |
| wt-1980-2100 | 2872 | AGAACTGGCCCTCGTTGGCATTCTACTAGGAAACGGCGAGCAGGGTGAACACTTGGGCACGGAGAGTGCCCTGGAGGCCTCAGGCAACAACTATGTGTATGCCTACGGACCAGACTGGAT | | 4 |
| wt-2100-2220 | 2873 | GGCAAGGCCTTCCACATGGTCCGCGGAAATCCAGCAATTCCTGCGACTCCTGGGCGCCACGTACGTGCTTCGCGTGGAGATGGGCAGGCAGTTTGGCTTCGAGGTGCATAGAAGCCGGCC | | 4 |
| wt-2220-2340 | 2874 | CTCCTTCCGTCAGTTCCAGGCCATCAATCACCTTGTCCTGTTTGACAACGCCCTTCGCAAGTACGATTCCGGCCAGGTGGCGGCGGGCTTCCAGAGGGCCCTTCTGGTGGCCGGGCCAGA | | 4 |
| wt-2340-2460 | 2875 | GACCGCTGACACGAGGCCGGACCTCCGCAAGCTGAATGAGTGGGTGTTTGGTGGCAGGGCTGCTGGTGGCAGACAGCTGGCCGACGAGCTAAAGATCGTGTCCGCGCTGCGAGACACTTA | | 4 |
| wt-2460-2580 | 2876 | CTCGGGCCACTTGGTCCTTCAGCCCACGGAGACCCTTGACACATGGAAGGTGTTGAGCAGGGACACACGAACCGCTCATAGTTTGGAGCACGGATTCATTCATGCCGCGGGGACCATCCA | | 4 |
| wt-2580-2700 | 2877 | GGCCAACTGCCCACAGCTGTTTATGAGACGCCAGCACCCCGGCCTCTTTCCCTTCGTTAATGCAATAGCATCATCGCTGGGCTGGTACTACCAGACCGCCACCGGCCCCGGAGCAGATGC | | 4 |
| wt-2700-2820 | 2878 | CAGGGCGGCGGCCCGGCGCCAACAGGCCTTTCAGACCAGGGCGGCGGCTGAATGCCATGCCAAAAGCGGGGTGCCGGTCGTGGCCGGCTTCTACAGGACCATCAACGCCACGCTCAAGGG | | 8 |
| wt-2820-2940 | 2879 | AGGAGAGGGCCTACAGCCCACTATGTTTAACGGGGAGCTGGGGGCCATCAAGCACCAGGCACTTGACACTGTGAGGTATGACTACGGCCACTATCTCATAATGTTGGGGCCATTCCAGCC | | 4 |
| wt-2940-3060 | 2880 | ATGGAGCGGACTGACGGCCCCTCCGTGCCCCTACGCCGAAAGTTCATGGGCACAGGCGGCCGTGCAGACGGCCCTCGAGCTGTTCTCGGCCCTGTACCCGGCCCCGTGCATCTCGGGCTA | | 8 |
| wt-3060-3180 | 2881 | CGCGCGCCCCCCGGGCCCCAGTGCTGTGATCGAGCATCTGGGGTCCCTAGTTCCAAAGGGGGGTCTGCTGTTGTTTCTGTCTCACCTACCGGATGATGTTAAGGACGGGCTCGGAGAAAT | | 4 |
| wt-3180-3300 | 2882 | GGGGCCGGCCAGGGCCACGGGACCTGGAATGCAGCAGTTTGTCAGCAGCTACTTCCTCAACCCCGCCTGTTCCAACGTCTTCATTACAGTGAGGCAGCGAGGGGAGAAGATCAACGGCCG | | 4 |
| wt-3300-3420 | 2883 | TACCGTCCTCCAAGCGCTCGGACGCGCATGCGATATGGCAGGCTGCCAGCACTATGTGCTGGGCTCCACGGTTCCCCTCGGTGGACTCAACTTTGTCAACGACCTGGCGTCCCCGGTTTC | | 4 |
| wt-3420-3540 | 2884 | CACCGCCGAGATGATGGATGATTTCTCTCCCTTCTTCACCGTGGAGTTTCCCCCGATTCAAGAGGAGGGCGCAAGTTCTCCGGTACCCTTAGATGTGGACGAGAGCATGGACATCTCTCC | | 4 |
| wt-3540-3660 | 2885 | GTCTTACGAGTTGCCCTGGCTCTCGCTGGAGTCATGCCTCACAAGCATCCTGTCACACCCCACCGTGGGAAGCAAGGAGCACTTGGTCAGGCACACGGACAGGGTCAGCGGAGGACGCGT | | 4 |
| wt-3660-3780 | 2886 | GGCACAGCAGCCCGGGGTAGGTCCCCTGGACCTGCCGCTGGCGGACTACGCCTTCGTTGCCCACAGTCAGGTCTGGACCAGGCCCGGTGGGGCTCCTCCCTTGCCCTATCGTACCTGGGA | | 8 |
| wt-3780-3900 | 2887 | TCGAATGACAGAGAAGCTGCTTGTCTCCGCAAAACCCGGCGGAGAGAACGTTAAGGTTTCAGGTACCGTGATTACATTGGGAGAACAGGGGTACAAAGTGTCGTTGGATCTGAGGGAGGG | | 4 |
| wt-3900-4020 | 2888 | AACCAGGCTGGCAATGGCTGAGGCGCTGCTGAACGCAGCATGTGCCCCAATCTTGGATCCGGAAGACGTCTTGCTCACCCTGCATCTACACCTGGATCCGCGCCGGGCAGACAACTCGGC | | 4 |
| wt-4020-4140 | 2889 | CGTGATGGAGGCTATGACGGCGGCGAGTGACTACGCGCGTGGCCTGGGCGTGAAGCTGACCTTTGGCTCGGCCTCCTGCCCCGAGACCGGCTCGTCCGCCTCCAACTTCATGACCGTGGT | | 8 |
| wt-4140-4260 | 2890 | GGCCTCTGTCTCCGCCCCAGGGGAATTCTCGGGTCCTCTGATCACGCCAGTGCTTCAGAAGACGGGCAGTCTCCTGATTGCGGTGCGTTGCGGGGATGGCAAGATCCAGGGAGGGTCGCT | | 4 |
| wt-4260-4380 | 2891 | GTTTGAGCAGCTCTTTAGCGACGTGGCCACGACCCCACGGGCACCCGAGGCGTTGTCTCTGAAGAATCTCTTCCGGGCAGTCCAGCAGCTGGTCAAGAGCGGCATCGTGCTGTCAGGGCA | | 4 |
| wt-4380-4500 | 2892 | TGACATCAGCGACGGGGGCCTGGTGACCTGCCTGGTGGAGATGGCCCTGGCCGGGCAGCGGGGAGTGACCATCACTATGCCGGTGGCCTCCGACTACCTCCCGGAGATGTTTGCAGAGCA | | 8 |
| wt-4500-4620 | 2893 | CCCCGGCCTGGTGTTTGAGGTGGAGGAGCGCAGCGTGGGTGAGGTGCTGCAGACCCTGCGCTCCATGAACATGTACCCGGCAGTCCTCGGTCGAGTGGGCGAGCAAGGTCCAGATCAAAT | | 4 |
| wt-4620-4740 | 2894 | GTTTGAGGTGCAGCACGGCCCAGAGACGGTGTTGCGCCAGTCGCTGCGCCTGCTGCTGGGAACCTGGTCATCCTTTGCCAGCGAGCAGTACGAGTGCCTGCGACCAGATCGGATTAACCG | | 4 |
| wt-4740-4860 | 2895 | GTCCATGCACGTGTCCGACTACGGCTATAACGAAGCACTGGCAGTCTCCCCGTTGACAGGAAAGAATCTCAGCCCACGCCGGTTGGTGACAGAGCCTGACCCACGATGTCAGGTGGCCGT | | 4 |
| wt-4860-4980 | 2896 | GCTATGCGCCCCGGGCACCAGGGGCCATGAAAGCCTCCTGGCGGCCTTCACGAATGCCGGATGCCTGTGCCGACGGGTGTTCTTTCGCGAGGTTAGGGACAACACGTTCCTCGACAAGTA | | 4 |
| wt-4980-5100 | 2897 | CGTGGGTCTGGCCATCGGAGGAGTTCATGGGGCCAGGGACTCTGCCCTGGCAGGCCGTGCCACCGTGGCGCTGATTAATCGTTTCCCCGCCCTGCGTGACGCTATTCTAAAGTTCCTCAA | | 4 |
| wt-5100-5220 | 2898 | CAGGCCAGATACGTTCTCGGTGGCCTTGGGGGAGCTGGGGGTGCAAGTTTTGGCTGGCCTGGGGGCCGTGGGGTCAACAGATAATCCACCCGCCCCTGGCGTGGAAGTTAATGTCCAGAG | | 4 |
| wt-5220-5340 | 2899 | ATCACCTCTGATTCTGGCCCCCAACGCCTCTGGCATGTTTGAGTCCCGCTGGCTGAACATTAGCATCCCGGCGACCACCAGCTCTGTCATGCTGCGTGGCCTCCGGGGCTGCGTCCTGCC | | 4 |
| wt-5340-5460 | 2900 | TTGTTGGGTGCAAGGCTCGTGCCTGGGCCTGCAATTTACTAACCTCGGGATGCCATATGTTTTGCAGAATGCCCACCAGATCGCCTGCCACTTCCACAGCAATGGCACGGATGCCTGGCG | | 4 |
| wt-5460-5580 | 2901 | CTTTGCTATGAATTATCCAAGAAACCCCACGGAGCAGGGCAACATTGCAGGGCTCTGTTCACGCGATGGTCGTCATCTGGCTCTCCTGTGTGACCCCTCACTTTGTACAGACTTTTGGCA | | 4 |
| wt-5580-5700 | 2902 | ATGGGAGCACATTCCCCCCGCCTTTGGGCACCCCACGGGGTGCTCCCCCTGGACACTTATGTTTCAAGCAGCTCACCTATGGTCACTCAGGCACGGTCGCCCCTCCGAGTGACCAGTCAC | | 4 |
| wt-5700-5820 | 2903 | CTTCCAGACTATGCATACACTGAATTTAGCCTGATATTGTCCCCCTAGCCCCGGGCCCAGCCCTCCTCAGAAAACTCTGCATGGAGAAGCTGGACGTGAACCTCCCCCCCAGACCTGTGT | | 4 |
| wt-5820-5940 | 2904 | GCTGTATTTACAAACACTACAATAAACCCAATGTGCAAATGTGGTTTGTATGGCTACTTTGTGTTCCTAAAAAATGCAACAATAGAAGTGGAAACCCTCAGTCACGGGACATTAACCTCA | | 4 |
| wt-5940-6060 | 2905 | ACCACAAAATGGGGGTTGGAGAAAGTAACCACATATACTGGAGATGATTCATGGGCTGGGGGTTCCCGGACAATACACCCATCTGGAGTTCAACCTAATTACATGGTAGATAAATTAAGA | | 4 |
| wt-6060-6180 | 2906 | GTCCCTCCTCACCACTCGAAACTATGGCAGACATTCTATAAGATAACGAGGAGAGATGAGGTGAGGGCAGAGGACATTGGGCAGGTGTGGGCCACGGGGCAGCTGGCCATATCCCCCGCA | | 4 |
| wt-6180-6300 | 2907 | CTACAGAAGTGTAAGCAAAGTGAAGGGCTCGGAAGGCAGGCGGGGCCTAGCAATGTCACAGCTAAATGCCCACCAGGGCACACACTCAAGCGGGGTCTCGGAGCTCCTAGGTCAGACCAC | | 4 |
| wt-6300-6420 | 2908 | GAAAGGTCAGCCTGCAAGGTGGATGGCGTGTTTTCTGAGGTTATCCCCGCTACGTGCAGTGCTGGGTGATAGAGACCCTAGAATGTGTCGAAATGACCAAGCGTCCCCGCAGCGGGGCTC | | 4 |
| wt-6420-6540 | 2909 | CCAACACGGGTTCCCAGAGAGGGTAAAAGAGGGGGCCATAAAGCCCAGGGTGTAAAACACCGACCGCGCCACCAGATGGCACACGTGGGGGAAATGAGGGTTAGCATAGGCAACCCCCGC | | 4 |
| wt-6540-6660 | 2910 | CTACACACCAACTATAGCAAACCCCGCCCCGTCACGGTGACGTAGTCTGTCTTGAGGAGATGTAGACTTGTAGACACTGCAAAACCTCAGGACCTACGCTGCCCTAGAGGTTTTGCTAGG | | 4 |
| wt-6660-6780 | 2911 | GAGGAGACGTGTGTGGCTGTAGCCACCCGTCCCGGGTACAAGTCCCGGGTGGTGAGGACGGTGTCTGTGGTTGTCTTCCCAGACTCTGCTTTCTGCCGTCTTCGGTCAAGTACCAGCTGG | | 4 |
| wt-6780-6900 | 2912 | TGGTCCGCATGTTTTGATCCAAACTTTTGTTTTAGGATTTATGCATCCATTATCCCGCAGTTCCACCTAAACGGGGCTTAACGTTGCATCCCAGAAGATGCACGCTTAACCCCGCCTACA | | 4 |
| wt-6900-7020 | 2913 | ACCGTGACGTAGCTGTTTACCAGCATGTATAGAGTTACGGTTCGCTACATCAAACAGGACAGCCGTTGCCCTAGTGGTTTCGGACACACCGCCAACGCTCAGTGCGGTGCTACCGACCCG | | 4 |
| wt-7020-7140 | 2914 | AGGTCAAGTCCCGGGGGAGGAGAAGAGAGGCTTCCCGCCTAGAGCATTTGCAAGTCAGGATTCTCTAATCCCTCTGGGAGAAGGGTATTCGGCTTGTCCGCTATTTTTTTGTGGCTAGTT | | 4 |
| wt-7140-7260 | 2915 | TTGCACCCACAACATGTAAGGGCCCGCTACCCCTACAACACAAAACAAACTATCTCCCCTAACCATCCTTTTGCCAATCAATTCTGTGACAGGGTTTCCTGGACACCCAGTCTTAGTTCA | | 4 |
| wt-7260-7380 | 2916 | GGTAGACACCCAGTTATGCAGTGCCACCAATTCCAACCATTTTTAAACCTCCTGGAATTCTATCATTAAACGGCATGCAGGAAAAGGACAAGCAGCGAAAATTCACGCCCCCTTGGGAGG | | 4 |

| | | | |
|---|---|---|---|
| wt-7380-7500 | 2917 | TGGCGGCATATGCAAAGGATAGCACTCCCACTCTACTACTGGGTATCATATGCTGACTGTATATGCATGAGGATAGCATATGCTACCCGGATACAGATTAGGATAGCATATACTACCCAG | 4 |
| wt-7500-7620 | 2918 | ATATAGATTAGGATAGCATATGCTACCCAGATATAGATTAGGATAGCCTATGCTACCCAGATATAAATTAGGATAGCATATACTACCCAGATATAGATTAGGATAGCATATGCTACCCAG | 4 |
| wt-7620-7740 | 2919 | ATATAGATTAGGATAGCCTATGCTACCCAGATATAGATTAGGATAGCATATGCTACCCAGATATAGATTAGGATAGCATATGCTATCCAGATATTTGGGTAGTATATGCTACCCAGATAT | 4 |
| wt-7740-7860 | 2920 | AAATTAGGATAGCATATACTACCCTAATCTCTATTAGGATAGCATATGCTACCCGGATACAGATTAGGATAGCATATACTACCCAGATATAGATTAGGATAGCATATGCTACCCAGATAT | 4 |
| wt-7860-7980 | 2921 | AGATTAGGATAGCCTATGCTACCCAGATATAAATTAGGATAGCATATACTACCCAGATATAGATTAGGATAGCATATGCTACCCAGATATAGATTAGGATAGCCTATGCTACCCAGATAT | 4 |
| wt-7980-8100 | 2922 | AGATTAGGATAGCATATGCTATCCAGATATTTGGGTAGTATATGCTACCCATGGCAACATTAGCCCACCGTGCTCTCAGCGACCTCGTGAATATGAGGACCAACAACCCTGTGCTTGGCG | 4 |
| wt-8100-8220 | 2923 | CTCAGGCGCAAGTGTGTGTAATTTGTCCTCCAGATCGCAGCAATCGCGCCCCTATCTTGGCCCGCCCACCTACTTATGCAGGTATTCCCCGGGGTGCCATTAGTGGTTTTGTGGGCAAGT | 4 |
| wt-8220-8340 | 2924 | GGTTTGACCGCAGTGGTTAGCGGGGTTACAATCAGCCAAGTTATTACACCCTTATTTTACAGTCCAAAACCGCAGGGCGGCGTGTGGGGGCTGACGCGTGCCCCCACTCCACAATTTCAA | 4 |
| wt-8340-8460 | 2925 | AAAAAAGAGTGGCCACTTGTCTTTGTTTATGGGCCCCATTGGCGTGGAGCCCCGTTTAATTTTCGGGGGTGTTAGAGACAACCAGTGGAGTCCGCTGCTGTCGGCGTCCACTCTCTTTCC | 4 |
| wt-8460-8580 | 2926 | CCTTGTTACAAATAGAGTGTAACAACATGGTTCACCTGTCTTGGTCCCTGCCTGGGACACATCTTAATAACCCCAGTATCATATTGCACTAGGATTATGTGTTGCCCATAGCCATAAATT | 4 |
| wt-8580-8700 | 2927 | CGTGTGAGATGGACATCCAGTCTTTACGGCTTGTCCCCACCCCATGGATTTCTATTGTTAAAGATATTCAGAATGTTTCATTCCTACACTAGTATTTATTGCCCAAGGGGTTTGTGAGGG | 4 |
| wt-8700-8820 | 2928 | TTATATTGGTGTCATAGCACAATGCCACCACTGAACCCCCCGTCCAAATTTTATTCTGGGGGCGTCACCTGAAACCTTGTTTTCGAGCACCTCACATACACCTTACTGTTCACAACTCAG | 4 |
| wt-8820-8940 | 2929 | CAGTTATTCTATTAGCTAAACGAAGGAGAATGAAGAAGCAGGCGAAGATTCAGGAGAGTTCACTGCCCGCTCCTTGATCTTCAGCCACTGCCCTTGTGACTAAAATGGTTCACTACCCTC | 4 |
| wt-8940-9060 | 2930 | GTGGAATCCTGACCCCATGTAAATAAAACCGTGACAGCTCATGGGGTGGGAGATATCGCTGTTCCTTAGGACCCTTTTACTAACCCTAATTCGATAGCATATGCTTCCCGTTGGGTAACA | 4 |
| wt-9060-9180 | 2931 | TATGCTATTGAATTAGGGTTAGTCTGGATAGTATATACTACTACCCGGGAAGCATATGCTACCCGTTTAGGGTTAACAAGGGGGCCTTATAAACACTATTGCTAATGCCCTCTTGAGGGT | 4 |
| wt-9180-9300 | 2932 | CCGCTTATCGGTAGCTACACAGGCCCCTCTGATTGACGTTGGTGTAGCCTCCCGTAGTCTTCCTGGGCCCCTGGGAGGTACATGTCCCCCAGCATTGGTGTAAGAGCTTCAGCCAAGAGT | 4 |
| wt-9300-9420 | 2933 | TACACATAAAGGCAATGTTGTGTTGCAGTCCACAGACTGCAAAGTCTGCTCCAGGATGAAAGCCACTCAGTGTTGGCAAATGTGCACATCCATTTATAAGGATGTCAACTACAGTCAGAG | 4 |
| wt-9420-9540 | 2934 | AACCCCTTTGTGTTTGGTCCCCCCCCGTGTCACATGTGGAACAGGGCCCAGTTGGCAAGTTGTACCAACCAACTGAAGGGATTACATGCACTGCCCCGCGGGAAATACGTCCTACCCAGG | 4 |
| wt-9540-9660 | 2935 | AACCCGAAACAGTGTTTCCCAGAAGCTGTAAAAATAGAACGCCCTGGAACTGCCCCACTGTGCAATGCAGCTTTTAGCCATGCCATGCTCTATAAATCACTTCCCTATCTCAGGTAGGCC | 4 |
| wt-9660-9780 | 2936 | TGCACACCTTAGGTATGGAGCGAAGGTTAGTGGTCACTCTGCAGTGCCTGGTGCTGCTTTACCTGGCACCTGAGTGTGGAGGTACAGACCAATGTGACAATTTTCCCCAAATGTTGAGGG | 4 |
| wt-9780-9900 | 2937 | ACCTAAGAGATGCCTTCAGTCGTGTTAAAACCTTTTTCCAGACAAAGGACGAGGTAGATAACCTTTTGCTCAAGGAGTCTCTGCTAGAGGACTTTAAGGGCTACCTTGGATGCCAGGCCC | 4 |
| wt-9900-10020 | 2938 | TGTCAGAAATGATCCAATTCTACCTGGAGGAAGTCATGCCACAGGCTGAAAACCAGGACCCTGAAGCCAAAGACCATGTCAATTCTTTGGGTGAAAATCTAAAGACCCTACGGCTCCGCC | 4 |
| wt-10020-10140 | 2939 | TGCGCAGGTGCCACAGGTTCCTGCCGTGTGAGAACAAGAGTAAAGCTGTGGAACAGATAAAAAATGCCTTTAACAAGCTGCAGGAAAAAGGAATTTACAAAGCCATGAGTGAATTTGACA | 4 |
| wt-10260-10380 | 2940 | AAATGAAACCTTTTATTCTTGATTGCCTCTTGTGTTCTTGCCGCCCAGGTACCTTCCTGTGTTCTCCCCACGGGAAAAAGAATAGCTTCTGCAGAAGGCCATTGACGCAAGTTTTGCCCG | 4 |
| wt-10380-10500 | 2941 | TGGGGATTACCCGACCCAGCCACTTACAGCACATTTTGTTCTAGGTCCATCTTAGGAGCCCGGGCCAGCATTCTATCAGCTTAACGGGAAGAGAAGTGGGGAGGGCACTCGCCCACTAAC | 4 |
| wt-10500-10620 | 2942 | CTTAACACCTGCAGCCTACAAAAGTACACTAGCTGTTTGCTCTATTCGCCACTAGAGACCGCCAAGATGCGAAACTACAGGCCCGGGCCCAGGCCTTGCAGGGCAGACGGTTAGGCTGAC | 4 |
| wt-10620-10740 | 2943 | AAGGGGACAAGTGTGGCAGGTGGGCGGGAAGGGGCACAAGAATGCCGGCGAAACTGGACCACGGTCCACCCCGCCCTCAAGCGTCCGGGAGCCGGGCGGCTCGGCTAAGGAGGGCGGCCT | 8 |
| wt-10740-10860 | 2944 | TGCGAACAATTATTAGTAGCTACCAACAAGGGCCCCCAGATGCCCCCCACCAGTCACCCGGCCGTGTCCACTCACATATTCCACTCTTATTTTTAAATTAATGTGTCCCAATTAGAAACC | 4 |
| wt-10860-10980 | 2945 | CAAGCGCAGAAATTAGTTGAGAGGCTAGTGTTTTAAACATGCACCCTAGGCCAGCCAGAGATAATGTCACAAGATTATCAAGTTGGTGTAAACACGCCGTGGGAAAAAATTTATGGTTCA | 4 |
| wt-10980-11100 | 2946 | GTGCGTCGAGTGCTATCTTTGGAACAGTAGAAAATTGAACCTTGTTGGCGGGAGAAGGAATAACGCCTTATCTGGGAGGAGCGACGGATTATAGCCAATAAGAGAGCTCAAGACGCAGGG | 4 |
| wt-11100-11220 | 2947 | CTCGCAAAGTATAGTGGCCCCGTGGGACCTTAGAGGTGGAGCAACGTCTAAAGTGGTAATAACACCAGGCGGGGCTGGGCAAAGGGGTCCTACGGGCGGGATTAATTACGCCTTGCTTAC | 4 |
| wt-11220-11340 | 2948 | GCAAGCTCAGTTAATTCGCCCACGACTTGAAAAATGTAGCCCTTAACCAATTGGCGGCCCCTAAGGGGGGGACTAAGGTCCCACTACAAAAACTCTGTGTTCTGCTGCAAATTTTAGATC | 4 |
| wt-11340-11460 | 2949 | AGATGGCATAGAGACAAGGACACCGAAGACCCCCAGAGCCCTCATCGCAGGGTTCTTACCATGCGGCCATGTAGGCCCACTTAACACTACAAGACCTACGCCTCTCCATTCATCATGTAA | 4 |
| wt-11460-11580 | 2950 | CCCACAAATCATCTAAACCGTAAGTCTAAGGGCCTCCTGAGGTTTTCTCAGGAGGCCCTAATGTATAATTAATCATGCATTTGATTTTAAAAAAGTAGGTTACACTCATTTTAGGCCAGA | 4 |
| wt-11580-11700 | 2951 | CTTTATTTGCAGATTAATAATTTATGTGATTCTCCTTCCCTCTAGGACTGAAGAAACAGCCTCCTGCACGTGAGCATGTATCTGAAATAATTATTATGTCATAAGTGTAATGATTAGAAA | 4 |
| wt-11700-11820 | 2952 | GTCATAAACCCACTTCCCTTTACATGAATCTGGGCACTGAATTTTGGGGTACTTCTAAAGACTAACGTGTTCGATTTCGGGGTCACTTCCCCTTTTATAAGTGTGTGAACAGTGATTTCA | 4 |
| wt-11820-11940 | 2953 | GTAAAACCTAAGAGATATTTGGTGTCACTTCCGCATTTTAAGTTTCAGAAAATTTTAAAATTAAAATTGAAATTTCTCTCAAAATAATTCCAATGAAAACTTCAAAGAATCTTATGTATG | 4 |
| wt-11940-12060 | 2954 | TAATTCTTTTGCCCCAAACTGGGCTTCAGATGCCTTCTATTGCACTCTCACAAAAACATTCTGGACACATGTGCCAGACGCCTGGGCCTCTAAGGCCCTCGGGTCCCCCTGGACCCCGGC | 4 |
| wt-12060-12180 | 2955 | CTCAGCAACCCTGCTGCTCCCCTCCTGCCACCCCAGCCTCCCCCCCTCCCCGTCCCCCTTCGCTCCTGTTCCTCCCCCGGTCCCCAGTAGGGCCGCCTGCCCCCCTGCACCCAGTACCTG | 8 |
| wt-12180-12300 | 2956 | CCCCTCTTGGCCACGCACCCCGGGCCAGGCCACCTTAGACCCGGCCAAGCCCCATCCCTGAAGACCCAGCGGCCATTCTCTCTGGTAACGAGCAGAGAAGAAGTAGAGGCCCGCGGCCAT | 8 |
| wt-12300-12420 | 2957 | TGGGCCCAGATTGAGAGACCAGTCCAGGGGCCCGAGGTTGGAGCCAGCGGGCACCCGAGGTCCCAGCACCCGGTCCCTCCGGGGGGCAGAGACAGGCAGGGCCCCCCGGCAGCTGGCCCC | 8 |
| wt-12420-12540 | 2958 | GAGGAGGCGCCCGGAGTGGGGCCGGTCGGCTGGGCTGGCCGAGCCCGGGTCTGGGAGGTCTGGGGTGGCGAGCCTGCTGTCTCAGGAGGGGCCTGGCTCCGCCGGGTGGCCCTGGGGTAA | 12 |
| wt-12540-12660 | 2959 | GTCTGGGAGGCAGAGGGTCGGCCTAGGCCCGGGGAAGTGGAGGGGGATCGCCCGGGTCTCTGTTGGCAGAGTCCGGGCGATCCTCTGAGACCCTCCGGGCCCGGACGGTCGCCCTCAGCC | 8 |
| wt-12660-12780 | 2960 | CCCCAGACAGACCCCAGGGTCTCCAGGCAGGGTCCGGCATCTTCAGGGGCAGCAGGCTCACCACCACAGGCCCCCCAGACCCGGGTCTCGGCCAGCCGAGCCGACCGGCCCCGCGCCTGG | 8 |
| wt-12780-12900 | 2961 | CGCCTCCTCGGGGCCAGCCGCCGGGGTTGGTTCTGCCCCTCTCTCTGTCCTTCAGAGGAACCAGGGACCTCGGGCACCCCAGAGCCCCTCGGGCCCGCCTCCAGGCGCCCTCCTGGTCTC | 8 |
| wt-12900-13020 | 2962 | CGCTCCCCTCTGAGCCCCGTTAAACCCAAAGAATGTCTGAGGGGAGCCACCCTCGGGGCCCAGGCCCCAGAGTCCAGAGGTCAGGGGCACCTCAGGGTGCCTCCCCGGGTCCCAGGCCAG | 8 |
| wt-13020-13140 | 2963 | CCGGAGGGACCCCGGCAGCCCGGGCGGCCCCAGAGGCCGGTTCCTCGCCCCTTCCCCGGGCTTCAGAGCCCAGGATGTCCCCCAGAAGGGACCCTAGGCGTCCCCTCTCCTCCCCTCCAG | 8 |
| wt-13140-13260 | 2964 | GCCCGAGCCTCTCCCTCGCGGAGAGGGGCCTCTTTGGGCCCTCAAGTCCAGCCCCACCGAGACCCGAGTGGCCCGGATCCCCCCACCGGCCCTTCTCTCTGTCCCCCTGCTCCTCTCCAA | 8 |
| wt-13260-13380 | 2965 | CCTTCGCTCCACCCTAGACCCCAGCTTCTGGCCTCCCCGGGTCCACCAGGCCAGCCGGAGGGACCCCGGCAGCCCGGGCGAGTCGCCTTCCCTCTCCCCTGGCCTCTCCTTCCCGCCTCC | 8 |
| wt-13380-13500 | 2966 | CACCCGAGCCCCCTCAGCTTGCCTCCCCACCGGGTCCATCAGGCCGGCCGGAGGGACCCCGGCGGCCCGGTGTCAGTCCCCCCTGCAGCCGCCCAGTCTCTGCCTCCAGGCAAGGGCGCC | 12 |
| wt-13500-13620 | 2967 | AGCTTTTCTCCCCCCAGCCTGAGGCCCAGTCTCCTGTGCACTGTCTGTAAAGTCCAGCCTCCCACGCCCGTCCACGGCTCCCGGGCCCAGCCTCGTCCACCCCTCCCCACGGTGGACAGG | 8 |

| | | | |
|---|---|---|---|
| wt-13620-13740 | 2968 | CCCTCTGTCCACCCGGGCCATCCCCGCCCCCCTGTGTCCACCCCAGTCCCGTCCAGGGGGGACTTTATGTGACCCTTGGGCCTGGCTCCCCATAGACTCCCATGTAAGCCTGCCTCGAGT | 8 |
| wt-13740-13860 | 2969 | AGGTGCCTCCAGAGCCCCTTTTGCCCCCCTGGCGGCCCAGCCCGACCCCCGGGCGCCCCCAAACTTTGTCCAGATGTCCAGGGGTCCCCGAGGGTGAGGCCCAGCCCCCTCCCGCCCCTG | 8 |
| wt-13860-13980 | 2970 | TCCACTGCCCCGGTCCCCCCAGAAGCCCCCAAAAGTAGAGGCTCAGGCCATGCGCGCCCTGTCACCAGGCCTGCCAAAGAGCCAGATCTAAGGCCGGGAGAGGCAGCCCCAAAGCGGGTG | 8 |
| wt-13980-14100 | 2971 | CAGTAACAGGTAATCTCTGGTAGTGATTTGGACCCGAAATCTGACACTTTAGAGCTCTGGAGGACTTTAAAACTCTAAAAATCAAAACTTTAGAGGCGAATGGGCGCCATTTTGTCCCCA | 4 |
| wt-14100-14220 | 2972 | CGCGCGCATAATGGCGGACCTAGGCCTAAAACCCCCAGGAAGCGGGTCTATGGTTGGCTGCGCTGCTGCTATCTTTAGAGGGGAAAAGAGGAATAAGCCCCCAGACAGGGGAGTGGGCTT | 4 |
| wt-14220-14340 | 2973 | GTTTGTGACTTCACCAAAGGTCAGGGCCCAAGGGGGGTTCGCGTTGCTAGGCCACCTTCTCAGTCCAGCGCGTTTACGTAAGCCAGACAGCAGCCAATTGTCAGTTCTAGGGAGGGGGACC | 4 |
| wt-14340-14460 | 2974 | ACTGCCCCTGGTATAAAGTGGTCCTGCAGCTATTTCTGGTCGCATCAGAGCGCCAGGAGTCCACACAAATGTAAGAGGGGGTCTTCTACCTCTCCCTAGCCCTCCGCCCCCTCCAAGGAC | 4 |
| wt-14460-14580 | 2975 | TCGGGCCCAGTTTCTAACTTTTCCCCTTCCCTCCCTCGTCTTGCCCTGCGCCCGGGGCCACCTTCATCACCGTCGCTGACTCCGCCATCCAAGCCTAGGGGAGACCGAAGTGAAGGCCCT | 4 |
| wt-14580-14700 | 2976 | GGACCAACCCGGCCCGGGCCCCCCGGTATCGGGCCAGAGGTAAGTGGACTTTAATTTTTTCTGCTAAGCCCAACACTCCACCACACCCAGGCACACACTACACACACCCACCCGTCTCAG | 4 |
| wt-14700-14820 | 2977 | GGTCCCCTCGGACAGCTCCTAAGAAGGCACCGGTCGCCCAGTCCTACCAGAGGGGGCCAAGAACCCAGACGAGTCCGTAGAAGGGTCCTCGTCCAGCAAGAAGAGGAGGTGGTAAGCGGT | 4 |
| wt-14820-14940 | 2978 | TCACCTTCAGGGGTAAGTAACCTGACCTCTCCAGGGCTCACATAAAGGGAGGCTTAGTATACATGCTTCTTGCTTTTCACAGGAACCTGGGGGCTAGTCTGGGTGGGATTAGGCTGCCTC | 4 |
| wt-14940-15060 | 2979 | AAGTTGCATCAGCCAGGGCTTCATGCCCTCCTCAGTTCCCTAGTCCCCGGGCTTCAGGCCCCCTCCGTCCCCGTCCTCCAGAGACCCGGGCTTCAGGCCCTGCCTCTCCTGTTACCCTTT | 4 |
| wt-15060-15180 | 2980 | TAGAACCACAGCCTGGACACATGTGCCAGACGCCTTGGCCTCTAAGGCCCTCGGGTCCCCCTGGACCCCGGCCTCAGCAACCCTGCTGCTCCCCTCCTGCCACCCCAGCCTCCCCCCCTC | 8 |
| wt-15180-15300 | 2981 | CCCGTCCCCCTTCGCTCCTGATCCTCCCCCGGTCCCCAGTAGGGCCGCCTGCCCCCCTGCACCCAGTACCTGCCCCTCTTGGCCACGCACCCCGGGCCAGGCCACCTTAGACCCGGCCAA | 8 |
| wt-15300-15420 | 2982 | GCCCCATCCCTGAAGACCCAGCGGCCATTCTCTCTGGTAACGAGCAGAGAAGAAGTAGAGGCCCGCGGCCATTGGGCCCAGATTGAGAGACCAGTCCAGGGGCCCGAGGTTGGAGCCAGC | 4 |
| wt-15420-15540 | 2983 | GGGCACCCGAGGTCCCAGCACCCGGTCCCTCCGGGGGGCAGAGACAGGCAGGGCCCCCCGGCAGCTGGCCCCGAGGAGGCGCCCGGAGTGGGGCCGGTCGGCTGGGCTGGCCGAGCCCGG | 12 |
| wt-15540-15660 | 2984 | GTCTGGGAGGTCTGGGGTGGCGAGCCTGCTGTCTCAGGAGGGGCCTGGCTCCGCCGGGTGGCCCTGGGGTAAGTCTGGGAGGCAGAGGGTCGGCCTAGGCCCGGGGAAGTGGAGGGGGAT | 8 |
| wt-15660-15780 | 2985 | CGCCCGGGTCTCTGTTGGCAGAGTCCGGGCGATCCTCTGAGACCCTCCGGGCCCGGACGGTCGCCCTCAGCCCCCCAGACAGACCCCAGGGTCTCCAGGCAGGGTCCGGCATCTTCAGGG | 8 |
| wt-15780-15900 | 2986 | GCAGCAGGCTCACCACCACAGGCCCCCCAGACCCGGGTCTCGGCCAGCCGAGCCGACCGGCCCCGCGCCTGGCGCCTCCTCGGGGCCAGCCGCCGGGGTTGGTTCTGCCCCTCTCTCTGT | 12 |
| wt-15900-16020 | 2987 | CCTTCAGAGGAACCAGGGACCTCGGGCACCCCAGAGCCCCTCGGGCCCGCCTCCAGGCGCCCTCCTGGTCTCCGCTCCCCTCTGAGCCCCGTTAAACCCAAAGAATGTCTGAGGGGAGCC | 8 |
| wt-16020-16140 | 2988 | ACCCTCGGGGCCCAGGCCCCAGAGTCCAGAGGTCAGGGGCACCTCAGGGTGCCTCCCCGGGTCCCAGGCCAGCCGGAGGGACCCCGGCAGCCCGGGCGGCCCCAGAGGCCGGTTCCTCGC | 12 |
| wt-16140-16260 | 2989 | CCCTTCCCCGGGCTTCAGAGCCCAGGATGTCCCCCAGAAGGGACCCTAGGCGTCCCCTCTCCTCCCCTCCAGGCCCGAGCCTCTCCCTCGCGGAGAGGGGCCTCTTTGGGCCCTCAAGTC | 8 |
| wt-16260-16380 | 2990 | CAGCCCCACCGAGACCCGAGTGGCCCGGATCCCCCCACCGGCCCTTCTCTCTGTCCCCCTGCTCCTCTCCAACCTTCGCTCCACCCTAGACCCCAGCTTCTGGCCTCCCCGGGTCCACCA | 8 |
| wt-16380-16500 | 2991 | GGCCAGCCGGAGGGACCCCGGCAGCCCGGGCGAGTCGCCTTCCCTCTCCCCTGGCCTCTCCTTCCCGCCTCCCACCCGAGCCCCCTCAGCTTGCCTCCCCACCGGGTCCATCAGGCCGGC | 12 |
| wt-16500-16620 | 2992 | CGGAGGGACCCCGGCGGCCCGGTGTCAGTCCCCCCTGCAGCCGCCCAGTCTCTGCCTCCAGGCAAGGGCGCCAGCTTTTCTCCCCCCAGCCTGAGGCCCAGTCTCCTGTGCACTGTCTGT | 8 |
| wt-16620-16740 | 2993 | AAAGTCCAGCCTCCCACGCCCGTCCACGGCTCCCGGGCCCAGCCTCGTCCACCCCTCCCCACGGTGGACAGGCCCTCTGTCCACCCGGGCCATCCCCGCCCCCCTGTGTCCACCCCAGTC | 8 |
| wt-16740-16860 | 2994 | CCGTCCAGGGGGGACTTTATGTGACCCTTGGGCCTGGCTCCCCATAGACTCCCATGTAAGCCTGCCTCGAGTAGGTGCCTCCAGAGCCCCTTTTGCCCCCCTGGCGGCCCAGCCCGACCC | 8 |
| wt-16860-16980 | 2995 | CCGGGCGCCCCCAAACTTTGTCCAGATGTCCAGGGGTCCCCGAGGGTGAGGCCCAGCCCCCTCCCGCCCCTGTCCACTGCCCCGGTCCCCCCAGAAGCCCCCAAAAGTAGAGGCTCAGGC | 8 |
| wt-16980-17100 | 2996 | CATGCGCGCCCTGTCACCAGGCCTGCCAAAGAGCCAGATCTAAGGCCGGGAGAGGCAGCCCCAAAGCGGGTGCAGTAACAGGTAATCTCTGGTAGTGATTTGGACCCGAAATCTGACACT | 4 |
| wt-17100-17220 | 2997 | TTAGAGCTCTGGAGGACTTTAAAACTCTAAAAATCAAAACTTTAGAGGCGAATGGGCGCCATTTTGTCCCCACGCGCGCATAATGGCGGACCTAGGCCTAAAACCCCCAGGAAGCGGGTC | 4 |
| wt-17220-17340 | 2998 | TATGGTTGGCTGCGCTGCTGCTATCTTTAGAGGGGAAAAGAGGAATAAGCCCCCAGACAGGGGAGTGGGCTTGTTTGTGACTTCACCAAAGGTCAGGGCCCAAGGGGGGTTCGCGTTGCTA | 4 |
| wt-17340-17460 | 2999 | GGCCACCTTCTCAGTCCAGCGCGTTTACGTAAGCCAGACAGCAGCCAATTGTCAGTTCTAGGGAGGGGGACCACTGCCCCTGGTATAAAGTGGTCCTGCAGCTATTTCTGGTCGCATCAG | 4 |
| wt-17460-17580 | 3000 | AGCGCCAGGAGTCCACACAAATGTAAGAGGGGGTCTTCTACCTCTCCCTAGCCCTCCGCCCCCTCCAAGGACTCGGGCCCAGTTTCTAACTTTTCCCCTTCCCTCCCTCGTCTTGCCCTG | 4 |
| wt-17580-17700 | 3001 | CGCCCGGGGCCACCTTCATCACCGTCGCTGACTCCGCCATCCAAGCCTAGGGGAGACCGAAGTGAAGGCCCTGGACCAACCCGGCCCGGGCCCCCCGGTATCGGGCCAGAGGTAAGTGGA | 8 |
| wt-17700-17820 | 3002 | CTTTAATTTTTTCTGCTAAGCCCAACACTCCACCACACCCAGGCACACACTACACACACCCACCCGTCTCAGGGTCCCCTCGGACAGCTCCTAAGAAGGCACCGGTCGCCCAGTCCTACC | 4 |
| wt-17820-17940 | 3003 | AGAGGGGGCCAAGAACCCAGACGAGTCCGTAGAAGGGTCCTCGTCCAGCAAGAAGAGGAGGTGGTAAGCGGTTCACCTTCAGGGGTAAGTAACCTGACCTCTCCAGGGCTCACATAAAGG | 4 |
| wt-17940-18060 | 3004 | GAGGCTTAGTATACATGCTTCTTGCTTTTCACAGGAACCTGGGGGCTAGTCTGGGTGGGATTAGGCTGCCTCAAGTTGCATCAGCCAGGGCTTCATGCCCTCCTCAGTTCCCTAGTCCCC | 4 |
| wt-18060-18180 | 3005 | GGGCTTCAGGCCCCCTCCGTCCCCGTCCTCCAGAGACCCGGGCTTCAGGCCCTGCCTCTCCTGTTACCCTTTTAGAACCACAGCCTGGACACATGTGCCAGACGCCTTGGCCTCTAAGGC | 4 |
| wt-18180-18300 | 3006 | CCTCGGGTCCCCCTGGACCCCGGCCTCAGCAACCCTGCTGCTCCCCTCCTGCCACCCCAGCCTCCCCCCCTCCCCGTCCCCCTTCGCTCCTGATCCTCCCCCGGTCCCCAGTAGGGCCGC | 12 |
| wt-18300-18420 | 3007 | CTGCCCCCCTGCACCCAGTACCTGCCCCTCTTGGCCACGCACCCCGGGCCAGGCCACCTTAGACCCGGCCAAGCCCCATCCCTGAAGACCCAGCGGCCATTCTCTCTGGTAACGAGCAGA | 8 |
| wt-18420-18540 | 3008 | GAAGAAGTAGAGGCCCGCGGCCATTGGGCCCAGATTGAGAGACCAGTCCAGGGGCCCGAGGTTGGAGCCAGCGGGCACCCGAGGTCCCAGCACCCGGTCCCTCCGGGGGGCAGAGACAGG | 8 |
| wt-18540-18660 | 3009 | CAGGGCCCCCCGGCAGCTGGCCCCGAGGAGGCGCCCGGAGTGGGGCCGGTCGGCTGGGCTGGCCGAGCCCGGGTCTGGGAGGTCTGGGGTGGCGAGCCTGCTGTCTCAGGAGGGGCCTGG | 12 |
| wt-18660-18780 | 3010 | CTCCGCCGGGTGGCCCTGGGGTAAGTCTGGGAGGCAGAGGGTCGGCCTAGGCCCGGGGAAGTGGAGGGGGATCGCCCGGGTCTCTGTTGGCAGAGTCCGGGCGATCCTCTGAGACCCTCC | 8 |
| wt-18780-18900 | 3011 | GGGCCCGGACGGTCGCCCTCAGCCCCCCAGACAGACCCCAGGGTCTCCAGGCAGGGTCCGGCATCTTCAGGGGCAGCAGGCTCACCACCACAGGCCCCCCAGACCCGGGTCTCGGCCAGC | 8 |
| wt-18900-19020 | 3012 | CGAGCCGACCGGCCCCGCGCCTGGCGCCTCCTCGGGGCCAGCCGCCGGGGTTGGTTCTGCCCCTCTCTCTGTCCTTCAGAGGAACCAGGGACCTCGGGCACCCCAGAGCCCCTCGGGCCC | 12 |
| wt-19020-19140 | 3013 | GCCTCCAGGCGCCCTCCTGGTCTCCGCTCCCCTCTGAGCCCCGTTAAACCCAAAGAATGTCTGAGGGGAGCCACCCTCGGGGCCCAGGCCCCAGAGTCCAGAGGTCAGGGGCACCTCAGG | 8 |
| wt-19140-19260 | 3014 | GTGCCTCCCCGGGTCCCAGGCCAGCCGGAGGGACCCCGGCAGCCCGGGCGGCCCCAGAGGCCGGTTCCTCGCCCCTTCCCCGGGCTTCAGAGCCCAGGATGTCCCCCAGAAGGGACCCTA | 12 |
| wt-19260-19380 | 3015 | GGCGTCCCCTCTCCTCCCCTCCAGGCCCGAGCCTCTCCCTCGCGGAGAGGGGCCTCTTTGGGCCCTCAAGTCCAGCCCCACCGAGACCCGAGTGGCCCGGATCCCCCCACCGGCCCTTCT | 8 |
| wt-19380-19500 | 3016 | CTCTGTCCCCCTGCTCCTCTCCAACCTTCGCTCCACCCTAGACCCCAGCTTCTGGCCTCCCCGGGTCCACCAGGCCAGCCGGAGGGACCCCGGCAGCCCGGGCGAGTCGCCTTCCCTCTC | 8 |
| wt-19500-19620 | 3017 | CCCTGGCCTCTCCTTCCCGCCTCCCACCCGAGCCCCCTCAGCTTGCCTCCCCACCGGGTCCATCAGGCCGGCCGGAGGGACCCCGGCGGCCCGGTGTCAGTCCCCCCTGCAGCCGCCCAG | 12 |
| wt-19620-19740 | 3018 | TCTCTGCCTCCAGGCAAGGGCGCCAGCTTTTCTCCCCCCAGCCTGAGGCCCAGTCTCCTGTGCACTGTCTGTAAAGTCCAGCCTCCCACGCCCGTCCACGGCTCCCGGGCCCAGCCTCGT | 8 |

| | | | |
|---|---|---|---|
| wt-19740-19860 | 3019 | CCACCCCTCCCCACGGTGGACAGGCCCTCTGTCCACCCGGGCCATCCCCGCCCCCCTGTGTCCACCCCAGTCCCGTCCAGGGGGGACTTTATGTGACCCTTGGGCCTGGCTCCCCATAGA | 8 |
| wt-19860-19980 | 3020 | CTCCCATGTAAGCCTGCCTCGAGTAGGTGCCTCCAGAGCCCCTTTTGCCCCCCTGGCGGCCCAGCCCGACCCCCGGGCGCCCCCAAACTTTGTCCAGATGTCCAGGGGTCCCCGAGGGTG | 8 |
| wt-19980-20100 | 3021 | AGGCCCAGCCCCCTCCCGCCCCTGTCCACTGCCCCGGTCCCCCAGAAGCCCCCAAAAGTAGAGGCTCAGGCCATGCGCGCCCTGTCACCAGGCCTGCCAAAGAGCCAGATCTAAGGCCG | 8 |
| wt-20100-20220 | 3022 | GGAGAGGCAGCCCCAAAGCGGGTGCAGTAACAGGTAATCTCTGGTAGTGATTTGGACCCGAAATCTGACACTTTAGAGCTCTGGAGGACTTTAAAACTCTAAAAATCAAAACTTTAGAGG | 4 |
| wt-20220-20340 | 3023 | CGAATGGGCGCCATTTTGTCCCCACGCGCGCATAATGGCGGACCTAGGCCTAAAACCCCCAGGAAGCGGGTCTATGGTTGGCTGCGCTGCTGCTATCTTTAGAGGGGAAAAGAGGAATAA | 4 |
| wt-20340-20460 | 3024 | GCCCCCAGACAGGGGAGTGGGCTTGTTTGTGACTTCACCAAAGGTCAGGGCCCAAGGGGGTTCGCGTTGCTAGGCCACCTTCTCAGTCCAGCGCGTTTACGTAAGCCAGACAGCAGCCAA | 4 |
| wt-20460-20580 | 3025 | TTGTCAGTTCTAGGGAGGGGGACCACTGCCCCTGGTATAAAGTGGTCCTGCAGCTATTTCTGGTCGCATCAGAGCGCCAGGAGTCCACACAAATGTAAGAGGGGGTCTTCTACCTCTCCC | 4 |
| wt-20700-20820 | 3027 | AGGGGAGACCGAAGTGAAGGCCCTGGACCAACCCGGCCCGGGCCCCCCGGTATCGGGCCAGAGGTAAGTGGACTTTAATTTTTTCTGCTAAGCCCAACACTCCACCACACCCAGGCACAC | 4 |
| wt-20820-20940 | 3028 | ACTACACACACCCACCCGTCTCAGGGTCCCCTCGGACAGCTCCTAAGAAGGCACCGGTCGCCCAGTCCTACCAGAGGGGGCCAAGAACCCAGACGAGTCCGTAGAAGGGTCCTCGTCCAG | 4 |
| wt-20940-21060 | 3029 | CAAGAAGAGGAGGTGGTAAGCGGTTCACCTTCAGGGGTAAGTAACCTGACCTCTCCAGGGCTCACATAAAGGGAGGCTTAGTATACATGCTTCTTGCTTTTCACAGGAACCTGGGGGCTA | 4 |
| wt-21300-21420 | 3032 | CTGCCACCCCAGCCTCCCCCCCTCCCCGTCCCCCTTCGCTCCTGATCCTCCCCCGGTCCCCAGTAGGGCCGCCTGCCCCCCTGCACCCAGTACCTGCCCCTCTTGGCCACGCACCCCGGG | 12 |
| wt-21420-21540 | 3033 | CCAGGCCACCTTAGACCCGGCCAAGCCCCATCCCTGAAGACCCAGCGGCCATTCTCTCTGGTAACGAGCAGAGAAGAAGTAGAGGCCCGCGGCCATTGGGCCCAGATTGAGAGACCAGTC | 4 |
| wt-21540-21660 | 3034 | CAGGGGCCCGAGGTTGGAGCCAGCGGGCACCCGAGGTCCCAGCACCCGGTCCCTCCGGGGGGCAGAGACAGGCAGGGCCCCCCGGCAGCTGGCCCCGAGGAGGCGCCCGGAGTGGGGCCG | 12 |
| wt-21660-21780 | 3035 | GTCGGCTGGGCTGGCCGAGCCCGGGTCTGGGAGGTCTGGGGTGGCGAGCCTGCTGTCTCAGGAGGGGCCTGGCTCCGCCGGGTGGCCCTGGGGTAAGTCTGGGAGGCAGAGGGTCGGCCT | 8 |
| wt-21780-21900 | 3036 | AGGCCCGGGGAAGTGGAGGGGGATCGCCCGGGTCTCTGTTGGCAGAGTCCGGGCGATCCTCTGAGACCCTCCGGGCCCGGACGGTCGCCCTCAGCCCCCCAGACAGACCCCAGGGTCTCC | 8 |
| wt-21900-22020 | 3037 | AGGCAGGGTCCGGCATCTTCAGGGGCAGCAGGCTCACCACCACAGGCCCCCCAGACCCGGGTCTCGGCCAGCCGAGCCGACCGGCCCCGCGCCTGGCGCCTCCTCGGGGCCAGCCGCCGG | 12 |
| wt-22020-22140 | 3038 | GGTTGGTTCTGCCCCTCTCTCTGTCCTTCAGAGGAACCAGGGACCTCGGGCACCCCAGAGCCCCTCGGGCCCGCCTCCAGGCGCCCTCCTGGTCTCCGCTCCCCTCTGAGCCCCGTTAAA | 8 |
| wt-22140-22260 | 3039 | CCCAAAGAATGTCTGAGGGGAGCCACCCTCGGGGCCCAGGCCCCAGAGTCCAGAGGTCAGGGGCACCTCAGGGTGCCTCCCCGGGTCCCAGGCCAGCCGGAGGGACCCCGGCAGCCCGGG | 8 |
| wt-22260-22380 | 3040 | CGGCCCCAGAGGCCGGTTCCTCGCCCCTTCCCCGGGCTTCAGAGCCCAGGATGTCCCCCAGAAGGGACCCTAGGCGTCCCCTCTCCTCCCCTCCAGGCCCGAGCCTCTCCCTCGCGGAGA | 8 |
| wt-22380-22500 | 3041 | GGGGCCTCTTTGGGCCCTCAAGTCCAGCCCCACCGAGACCCGAGTGGCCCGGATCCCCCCACCGGCCCTTCTCTCTGTCCCCCTGCTCCTCTCCAACCTTCGCTCCACCCTAGACCCCAG | 8 |
| wt-22500-22620 | 3042 | CTTCTGGCCTCCCCGGGTCCACCAGGCCAGCCGGAGGGACCCCGGCAGCCCGGGCGAGTCGCCTTCCCTCTCCCCTGGCCTCTCCTTCCCGCCTCCCACCCGAGCCCCCTCAGCTTGCCT | 8 |
| wt-22620-22740 | 3043 | CCCCACCGGGTCCATCAGGCCGGCCGGAGGGACCCCGGCGGCCCGGTGTCAGTCCCCCCTGCAGCCGCCCAGTCTCTGCCTCCAGGCAAGGGCGCCAGCTTTTCTCCCCCCAGCCTGAGG | 8 |
| wt-22740-22860 | 3044 | CCCAGTCTCCTGTGCACTGTCTGTAAAGTCCAGCCTCCCACGCCCGTCCACGGCTCCCGGGCCCAGCCTCGTCCACCCCTCCCCACGGTGGACAGGCCCTCTGTCCACCCGGGCCATCCC | 8 |
| wt-22860-22980 | 3045 | CGCCCCCCTGTGTCCACCCCAGTCCCGTCCAGGGGGGACTTTATGTGACCCTTGGGCCTGGCTCCCCATAGACTCCCATGTAAGCCTGCCTCGAGTAGGTGCCTCCAGAGCCCCTTTTGC | 4 |
| wt-22980-23100 | 3046 | CCCCCTGGCGGCCCAGCCCGACCCCCGGGCGCCCCCAAACTTTGTCCAGATGTCCAGGGGTCCCCGAGGGTGAGGCCCAGCCCCCTCCCGCCCCTGTCCACTGCCCCGGTCCCCCAGAA | 12 |
| wt-23100-23220 | 3047 | GCCCCCAAAAGTAGAGGCTCAGGCCATGCGCGCCCTGTCACCAGGCCTGCCAAAGAGCCAGATCTAAGGCCGGGAGAGGCAGCCCCAAAGCGGGTGCAGTAACAGGTAATCTCTGGTAGT | 4 |
| wt-23220-23340 | 3048 | GATTTGGACCCGAAATCTGACACTTTAGAGCTCTGGAGGACTTTAAAACTCTAAAAATCAAAACTTTAGAGGCGAATGGGCGCCATTTTGTCCCCACGCGCGCATAATGGCGGACCTAGG | 4 |
| wt-23340-23460 | 3049 | CCTAAAACCCCCAGGAAGCGGGTCTATGGTTGGCTGCGCTGCTGCTATCTTTAGAGGGGAAAAGAGGAATAAGCCCCCAGACAGGGGAGTGGGCTTGTTTGTGACTTCACCAAAGGTCAG | 4 |
| wt-23460-23580 | 3050 | GGCCCAAGGGGGTTCGCGTTGCTAGGCCACCTTCTCAGTCCAGCGCGTTTACGTAAGCCAGACAGCAGCCAATTGTCAGTTCTAGGGAGGGGGACCACTGCCCCTGGTATAAAGTGGTCC | 4 |
| wt-23580-23700 | 3051 | TGCAGCTATTTCTGGTCGCATCAGAGCGCCAGGAGTCCACACAAATGTAAGAGGGGGTCTTCTACCTCTCCCTAGCCCTCCGCCCCCTCCAAGGACTCGGGCCCAGTTTCTAACTTTTCC | 4 |
| wt-23700-23820 | 3052 | CCTTCCCTCCCTCGTCTTGCCCTGCGCCCGGGGCCACCTTCATCACCGTCGCTGACTCCGCCATCCAAGCCTAGGGGAGACCGAAGTGAAGGCCCTGGACCAACCCGGCCCGGGCCCCCC | 8 |
| wt-23820-23940 | 3053 | GGTATCGGGCCAGAGGTAAGTGGACTTTAATTTTTTCTGCTAAGCCCAACACTCCACCACACCCAGGCACACACTACACACACCCACCCGTCTCAGGGTCCCCTCGGACAGCTCCTAAGA | 4 |
| wt-23940-24060 | 3054 | AGGCACCGGTCGCCCAGTCCTACCAGAGGGGGCCAAGAACCCAGACGAGTCCGTAGAAGGGTCCTCGTCCAGCAAGAAGAGGAGGTGGTAAGCGGTTCACCTTCAGGGGTAAGTAACCTG | 4 |
| wt-24060-24180 | 3055 | ACCTCTCCAGGGCTCACATAAAGGGAGGCTTAGTATACATGCTTCTTGCTTTTCACAGGAACCTGGGGGCTAGTCTGGGTGGGATTAGGCTGCCTCAAGTTGCATCAGCCAGGGCTTCAT | 4 |
| wt-24180-24300 | 3056 | GCCCTCCTCAGTTCCCTAGTCCCCGGGCTTCAGGCCCCCTCCGTCCCCGTCCTCCAGAGACCCGGGCTTCAGGCCCTGCCTCTCCTGTTACCCTTTTAGAACCACAGCCTGGACACATGT | 4 |
| wt-24300-24420 | 3057 | GCCAGACGCCTTGGCCTCTAAGGCCCTCGGGTCCCCCTGGACCCCGGCCTCAGCAACCCTGCTGCTCCCCTCCTGCCACCCCAGCCTCCCCCCCTCCCCGTCCCCCTTCGCTCCTGATCC | 8 |
| wt-24420-24540 | 3058 | TCCCCCGGTCCCCAGTAGGGCCGCCTGCCCCCCTGCACCCAGTACCTGCCCCTCTTGGCCACGCACCCCGGGCCAGGCCACCTTAGACCCGGCCAAGCCCCATCCCTGAAGACCCAGCGG | 8 |
| wt-24660-24780 | 3060 | GTCCCTCCGGGGGGCAGAGACAGGCAGGGCCCCCCGGCAGCTGGCCCCGAGGAGGCGCCCGGAGTGGGGCCGGTCGGCTGGGCTGGCCGAGCCCGGGTCTGGGAGGTCTGGGGTGGCGAG | 12 |
| wt-24780-24900 | 3061 | CCTGCTGTCTCAGGAGGGGCCTGGCTCCGCCGGGTGGCCCTGGGGTAAGTCTGGGAGGCAGAGGGTCGGCCTAGGCCCGGGGAAGTGGAGGGGGATCGCCCGGGTCTCTGTTGGCAGAGT | 8 |
| wt-24900-25020 | 3062 | CCGGGCGATCCTCTGAGACCCTCCGGGCCCGGACGGTCGCCCTCAGCCCCCCAGACAGACCCCAGGGTCTCCAGGCAGGGTCCGGCATCTTCAGGGGCAGCAGGCTCACCACCACAGGCC | 8 |
| wt-25020-25140 | 3063 | CCCCAGACCCGGGTCTCGGCCAGCCGAGCCGACCGGCCCCGCGCCTGGCGCCTCCTCGGGGCCAGCCGCCGGGGTTGGTTCTGCCCCTCTCTCTGTCCTTCAGAGGAACCAGGGACCTCG | 8 |
| wt-25140-25260 | 3064 | GGCACCCCAGAGCCCCTCGGGCCCGCCTCCAGGCGCCCTCCTGGTCTCCGCTCCCCTCTGAGCCCCGTTAAACCCAAAGAATGTCTGAGGGGAGCCACCCTCGGGGCCCAGGCCCCAGAG | 8 |
| wt-25260-25380 | 3065 | TCCAGAGGTCAGGGGCACCTCAGGGTGCCTCCCCGGGTCCCAGGCCAGCCGGAGGGACCCCGGCAGCCCGGGCGGCCCCAGAGGCCGGTTCCTCGCCCCTTCCCCGGGCTTCAGAGCCCA | 12 |
| wt-25380-25500 | 3066 | GGATGTCCCCCAGAAGGGACCCTAGGCGTCCCCTCTCCTCCCCTCCAGGCCCGAGCCTCTCCCTCGCGGAGAGGGGCCTCTTTGGGCCCTCAAGTCCAGCCCCACCGAGACCCGAGTGGC | 8 |
| wt-25500-25620 | 3067 | CCGGATCCCCCCACCGGCCCTTCTCTCTGTCCCCCTGCTCCTCTCCAACCTTCGCTCCACCCTAGACCCCAGCTTCTGGCCTCCCCGGGTCCACCAGGCCAGCCGGAGGGACCCCGGCAG | 8 |
| wt-25620-25740 | 3068 | CCCGGGCGAGTCGCCTTCCCTCTCCCCTGGCCTCTCCTTCCCGCCTCCCACCCGAGCCCCCTCAGCTTGCCTCCCCACCGGGTCCATCAGGCCGGCCGGAGGGACCCCGGCGGCCCGGTG | 12 |
| wt-25740-25860 | 3069 | TCAGTCCCCCCTGCAGCCGCCCAGTCTCTGCCTCCAGGCAAGGGCGCCAGCTTTTCTCCCCCCAGCCTGAGGCCCAGTCTCCTGTGCACTGTCTGTAAAGTCCAGCCTCCCACGCCCGTC | 8 |
| wt-25860-25980 | 3070 | CACGGCTCCCGGGCCCAGCCTCGTCCACCCCTCCCCACGGTGGACAGGCCCTCTGTCCACCCGGGCCATCCCCGCCCCCCTGTGTCCACCCCAGTCCCGTCCAGGGGGGACTTTATGTGA | 8 |
| wt-25980-26100 | 3071 | CCCTTGGGCCTGGCTCCCCATAGACTCCCATGTAAGCCTGCCTCGAGTAGGTGCCTCCAGAGCCCCTTTTGCCCCCCTGGCGGCCCAGCCCGACCCCCGGGCGCCCCCAAACTTTGTCCA | 8 |
| wt-26100-26220 | 3072 | GATGTCCAGGGGTCCCCGAGGGTGAGGCCCAGCCCCCTCCCGCCCCTGTCCACTGCCCCGGTCCCCCAGAAGCCCCCAAAAGTAGAGGCTCAGGCCATGCGCGCCCTGTCACCAGGCCT | 8 |
| wt-26220-26340 | 3073 | GCCAAAGAGCCAGATCTAAGGCCGGGAGAGGCAGCCCCAAAGCGGGTGCAGTAACAGGTAATCTCTGGTAGTGATTTGGACCCGAAATCTGACACTTTAGAGCTCTGGAGGACTTTAAAA | 4 |

| | | | |
|---|---|---|---|
| wt-26340-26460 | 3074 | CTCTAAAAATCAAAACTTTAGAGGCGAATGGGCGCCATTTTGTCCCCACGCGCGCATAATGGCGGACCTAGGCCTAAAACCCCCAGGAAGCGGGTCTATGGTTGGCTGCGCTGCTGCTAT | 4 |
| wt-26460-26580 | 3075 | CTTTAGAGGGGAAAAGAGGAATAAGCCCCCAGACAGGGGAGTGGGCTTGTTTGTGACTTCACCAAAGGTCAGGGCCCAAGGGGGTTCGCGTTGCTAGGCCACCTTCTCAGTCCAGCGCGT | 4 |
| wt-26580-26700 | 3076 | TTACGTAAGCCAGACAGCAGCCAATTGTCAGTTCTAGGGAGGGGGACCACTGCCCCTGGTATAAAGTGGTCCTGCAGCTATTTCTGGTCGCATCAGAGCGCCAGGAGTCCACACAAATGT | 4 |
| wt-26700-26820 | 3077 | AAGAGGGGGTCTTCTACCTCTCCCTAGCCCTCCGCCCCCTCCAAGGACTCGGGCCCAGTTTCTAACTTTTCCCCTTCCCTCCCTCGTCTTGCCCTGCGCCCGGGGCCACCTTCATCACCG | 4 |
| wt-26820-26940 | 3078 | TCGCTGACTCCGCCATCCAAGCCTAGGGGAGACCGAAGTGAAGGCCCTGGACCAACCCGGCCCGGGCCCCCCGGTATCGGGCCAGAGGTAAGTGGACTTTAATTTTTTCTGCTAAGCCCA | 4 |
| wt-26940-27060 | 3079 | ACACTCCACCACACCCAGGCACACACTACACACACCCACCCGTCTCAGGGTCCCCTCGGACAGCTCCTAAGAAGGCACCGGTCGCCCAGTCCTACCAGAGGGGGCCAAGAACCCAGACGA | 4 |
| wt-27060-27180 | 3080 | GTCCGTAGAAGGGTCCTCGTCCAGCAAGAAGAGGAGGTGGTAAGCGGTTCACCTTCAGGGGTAAGTAACCTGACCTCTCCAGGGCTCACATAAAGGGAGGCTTAGTATACATGCTTCTTG | 4 |
| wt-27180-27300 | 3081 | CTTTTCACAGGAACCTGGGGGCTAGTCTGGGTGGGATTAGGCTGCCTCAAGTTGCATCAGCCAGGGCTTCATGCCCTCCTCAGTTCCCTAGTCCCCGGGCTTCAGGCCCCCTCCGTCCCC | 4 |
| wt-27420-27540 | 3083 | CTCAGCAACCCTGCTGCTCCCCTCCTGCCACCCCAGCCTCCCCCCCTCCCCGTCCCCCTTCGCTCCTGATCCTCCCCCGGTCCCCAGTAGGGCCGCCTGCCCCCCTGCACCCAGTACCTG | 8 |
| wt-27540-27660 | 3084 | CCCCTCTTGGCCACGCACCCCGGGCCAGGCCACCTTAGACCCGGCCAAGCCCCATCCCTGAAGACCCAGCGGCCATTCTCTCTGGTAACGAGCAGAGAAGAAGTAGAGGCCCGCGGCCAT | 8 |
| wt-27660-27780 | 3085 | TGGGCCCAGATTGAGAGACCAGTCCAGGGGCCCGAGGTTGGAGCCAGCGGGCACCCGAGGTCCCAGCACCCGGTCCCTCCGGGGGGCAGAGACAGGCAGGGCCCCCCGGCAGCTGGCCCC | 8 |
| wt-27780-27900 | 3086 | GAGGAGGCGCCCGGAGTGGGGCCGGTCGGCTGGGCTGGCCGAGCCCGGGTCTGGGAGGTCTGGGGTGGCGAGCCTGCTGTCTCAGGAGGGGCCTGGCTCCGCCGGGTGGCCCTGGGGTAA | 12 |
| wt-27900-28020 | 3087 | GTCTGGGAGGCAGAGGGTCGGCCTAGGCCCGGGGAAGTGGAGGGGGATCGCCCGGGTCTCTGTTGGCAGAGTCCGGGCGATCCTCTGAGACCCTCCGGGCCCGGACGGTCGCCCTCAGCC | 8 |
| wt-28020-28140 | 3088 | CCCCAGACAGACCCCAGGGTCTCCAGGCAGGGTCCGGCATCTTCAGGGGCAGCAGGCTCACCACCACAGGCCCCCCAGACCCGGGTCTCGGCCAGCCGAGCCGACCGGCCCCGCGCCTGG | 8 |
| wt-28140-28260 | 3089 | CGCCTCCTCGGGGCCAGCCGCCGGGGTTGGTTCTGCCCCTCTCTCTGTCCTTCAGAGGAACCAGGGACCTCGGGCACCCCAGAGCCCCTCGGGCCCGCCTCCAGGCGCCCTCCTGGTCTC | 8 |
| wt-28260-28380 | 3090 | CGCTCCCCTCTGAGCCCCGTTAAACCCAAAGAATGTCTGAGGGGAGCCACCCTCGGGGCCCAGGCCCCAGAGTCAGAGGTCAGGGGCACCTCAGGGTGCCTCCCCGGGTCCCAGGCCAG | 8 |
| wt-28380-28500 | 3091 | CCGGAGGGACCCCGGCAGCCCGGGCGGCCCCAGAGGCCGGTTCCTCGCCCCTTCCCCGGGCTTCAGAGCCCAGGATGTCCCCAGAAGGGACCCTAGGCGTCCCCTCTCCTCCCCTCCAG | 8 |
| wt-28500-28620 | 3092 | GCCCGAGCCTCTCCCTCGCGGAGAGGGGCCTCTTTGGGCCCTCAAGTCCAGCCCCACCGAGACCCGAGTGGCCCGGATCCCCCCACCGGCCCTTCTCTCTGTCCCCCTGCTCCTCTCCAA | 8 |
| wt-28620-28740 | 3093 | CCTTCGCTCCACCCTAGACCCCAGCTTCTGGCCTCCCCGGGTCCACCAGGCCAGCCGGAGGGACCCCGGCAGCCCGGGCGAGTCGCCTTCCCTCTCCCCTGGCCTCTCCTTCCCGCCTCC | 8 |
| wt-28740-28860 | 3094 | CACCCGAGCCCCCTCAGCTTGCCTCCCCACCGGGTCCATCAGGCCGGCCGGAGGGACCCCGGCGGCCCGGTGTCAGTCCCCCCTGCAGCCGCCCAGTCTCTGCCTCCAGGCAAGGGCGCC | 12 |
| wt-28860-28980 | 3095 | AGCTTTTCTCCCCCCAGCCTGAGGCCCAGTCTCCTGTGCACTGTCTGTAAAGTCCAGCCTCCCACGCCCGTCCACGGCTCCCGGGCCCAGCCTCGTCCACCCCTCCCCACGGTGGACAGG | 8 |
| wt-28980-29100 | 3096 | CCCTCTGTCCACCCGGGCCATCCCCGCCCCCCTGTGTCCACCCCAGTCCCGTCCAGGGGGGACTTTATGTGACCCTTGGGCCTGGCTCCCCATAGACTCCCATGTAAGCCTGCCTCGAGT | 8 |
| wt-29100-29220 | 3097 | AGGTGCCTCCAGAGCCCCTTTTGCCCCCCTGGCGGCCCAGCCCGACCCCCGGGCGCCCCCAAACTTTGTCCAGATGTCCAGGGGTCCCCGAGGGTGAGGCCCAGCCCCCTCCCGCCCCTG | 8 |
| wt-29220-29340 | 3098 | TCCACTGCCCCGGTCCCCCCAGAAGCCCCCAAAAGTAGAGGCTCAGGCCATGCGCGCCCTGTCACCAGGCCTGCCAAAGAGCCAGATCTAAGGCCGGGAGAGGCAGCCCCAAAGCGGGTG | 8 |
| wt-29340-29460 | 3099 | CAGTAACAGGTAATCTCTGGTAGTGATTTGGACCCGAAATCTGACACTTTAGAGCTCTGGAGGACTTTAAAACTCTAAAAATCAAAACTTTAGAGGCGAATGGGCGCCATTTTGTCCCCA | 4 |
| wt-29460-29580 | 3100 | CGCGCGCATAATGGCGGACCTAGGCCTAAAACCCCCAGGAAGCGGGTCTATGGTTGGCTGCGCTGCTGCTATCTTTAGAGGGGAAAAGAGGAATAAGCCCCCAGACAGGGGAGTGGGCTT | 4 |
| wt-29580-29700 | 3101 | GTTTGTGACTTCACCAAAGGTCAGGGCCCAAGGGGGTTCGCGTTGCTAGGCCACCTTCTCAGTCCAGCGCGTTTACGTAAGCCAGACAGCAGCCAATTGTCAGTTCTAGGGAGGGGGACC | 4 |
| wt-29700-29820 | 3102 | ACTGCCCCTGGTATAAAGTGGTCCTGCAGCTATTTCTGGTCGCATCAGAGCGCCAGGAGTCCACACAAATGTAAGAGGGGGTCTTCTACCTCTCCCTAGCCCTCCGCCCCCTCCAAGGAC | 4 |
| wt-29820-29940 | 3103 | TCGGGCCCAGTTTCTAACTTTTCCCCTTCCCTCCCTCGTCTTGCCCTGCGCCCGGGGCCACCTTCATCACCGTCGCTGACTCCGCCATCCAAGCCTAGGGGAGACCGAAGTGAAGGCCCT | 4 |
| wt-29940-30060 | 3104 | GGACCAACCCGGCCCGGGCCCCCCGGTATCGGGCCAGAGGTAAGTGGACTTTAATTTTTTCTGCTAAGCCCAACACTCCACCACACCCAGGCACACACTACACACACCCACCCGTCTCAG | 4 |
| wt-30060-30180 | 3105 | GGTCCCCTCGGACAGCTCCTAAGAAGGCACCGGTCGCCCAGTCCTACCAGAGGGGGCCAAGAACCCAGACGAGTCCGTAGAAGGGTCCTCGTCCAGCAAGAAGAGGAGGTGGTAAGCGGT | 4 |
| wt-30180-30300 | 3106 | TCACCTTCAGGGGTAAGTAACCTGACCTCTCCAGGGCTCACATAAAGGGAGGCTTAGTATACATGCTTCTTGCTTTTCACAGGAACCTGGGGGCTAGTCTGGGTGGGATTAGGCTGCCTC | 4 |
| wt-30300-30420 | 3107 | AAGTTGCATCAGCCAGGGCTTCATGCCCTCCTCAGTTCCCTAGTCCCCGGGCTTCAGGCCCCCTCCGTCCCCGTCCTCCAGAGACCCGGGCTTCAGGCCCTGCCTCTCCTGTTACCCTTT | 4 |
| wt-30420-30540 | 3108 | TAGAACCACAGCCTGGACACATGTGCCAGACGCCTTGGCCTCTAAGGCCCTCGGGTCCCCTGGACCCCGGCCTCAGCAACCCTGCTGCTCCCCTCCTGCCACCCCAGCCTCCCCCCCTC | 8 |
| wt-30540-30660 | 3109 | CCCGTCCCCCTTCGCTCCTGATCCTCCCCCGGTCCCCAGTAGGGCCGCCTGCCCCCCTGCACCCAGTACCTGCCCCTCTTGGCCACGCACCCCGGGCCAGGCCACCTTAGACCCGGCCAA | 8 |
| wt-30660-30780 | 3110 | GCCCCATCCCTGAAGACCCAGCGGCCATTCTCTCTGGTAACGAGCAGAGAAGAAGTAGAGGCCCGCGGCCATTGGGCCCAGATTGAGAGACCAGTCCAGGGGCCCGAGGTTGGAGCCAGC | 4 |
| wt-30780-30900 | 3111 | GGGCACCCGAGGTCCCAGCACCCGGTCCCTCCGGGGGGCAGAGACAGGCAGGGCCCCCCGGCAGCTGGCCCCGAGGAGGCGCCCGGAGTGGGGCCGGTCGGCTGGGCTGGCCGAGCCCGG | 12 |
| wt-30900-31020 | 3112 | GTCTGGGAGGTCTGGGGTGGCGAGCCTGCTGTCTCAGGAGGGGCCTGGCTCCGCCGGGTGGCCCTGGGGTAAGTCTGGGAGGCAGAGGGTCGGCCTAGGCCCGGGGAAGTGGAGGGGGAT | 8 |
| wt-31020-31140 | 3113 | CGCCCGGGTCTCTGTTGGCAGAGTCCGGGCGATCCTCTGAGACCCTCCGGGCCCGGACGGTCGCCCTCAGCCCCCCAGACAGACCCCAGGGTCTCCAGGCAGGGTCCGGCATCTTCAGGG | 8 |
| wt-31140-31260 | 3114 | GCAGCAGGCTCACCACCACAGGCCCCCCAGACCCGGGTCTCGGCCAGCCGAGCCGACCGGCCCCGCGCCTGGCGCCTCCTCGGGGCCAGCCGCCGGGGTTGGTTCTGCCCCTCTCTCTGT | 12 |
| wt-31260-31380 | 3115 | CCTTCAGAGGAACCAGGGACCTCGGGCACCCCAGAGCCCCTCGGGCCCGCCTCCAGGCGCCCTCCTGGTCTCCGCTCCCCTCTGAGCCCCGTTAAACCCAAAGAATGTCTGAGGGGAGCC | 8 |
| wt-31380-31500 | 3116 | ACCCTCGGGGCCCAGGCCCCAGAGTCAGAGGTCAGGGGCACCTCAGGGTGCCTCCCCGGGTCCCAGGCCAGCCGGAGGGACCCCGGCAGCCCGGGCGGCCCCAGAGGCCGGTTCCTCGC | 12 |
| wt-31500-31620 | 3117 | CCCTTCCCCGGGCTTCAGAGCCCAGGATGTCCCCAGAAGGGACCCTAGGCGTCCCCTCTCCTCCCCTCCAGGCCCGAGCCTCTCCCTCGCGGAGAGGGGCCTCTTTGGGCCCTCAAGTC | 8 |
| wt-31620-31740 | 3118 | CAGCCCCACCGAGACCCGAGTGGCCCGGATCCCCCCACCGGCCCTTCTCTCTGTCCCCCTGCTCCTCTCCAACCTTCGCTCCACCCTAGACCCCAGCTTCTGGCCTCCCCGGGTCCACCA | 8 |
| wt-31740-31860 | 3119 | GGCCAGCCGGAGGGACCCCGGCAGCCCGGGCGAGTCGCCTTCCCTCTCCCCTGGCCTCTCCTTCCCGCCTCCCACCCGAGCCCCCTCAGCTTGCCTCCCCACCGGGTCCATCAGGCCGGC | 12 |
| wt-31860-31980 | 3120 | CGGAGGGACCCCGGCGGCCCGGTGTCAGTCCCCCCTGCAGCCGCCCAGTCTCTGCCTCCAGGCAAGGGCGCCAGCTTTTCTCCCCCCAGCCTGAGGCCCAGTCTCCTGTGCACTGTCTGT | 8 |
| wt-31980-32100 | 3121 | AAAGTCCAGCCTCCCACGCCCGTCCACGGCTCCCGGGCCCAGCCTCGTCCACCCCTCCCCACGGTGGACAGGCCCTCTGTCCACCCGGGCCATCCCCGCCCCCCTGTGTCCACCCCAGTC | 8 |
| wt-32100-32220 | 3122 | CCGTCCAGGGGGGACTTTATGTGACCCTTGGGCCTGGCTCCCCATAGACTCCCATGTAAGCCTGCCTCGAGTAGGTGCCTCCAGAGCCCCTTTTGCCCCCCTGGCGGCCCAGCCCGACCC | 8 |
| wt-32220-32340 | 3123 | CCGGGCGCCCCCAAACTTTGTCCAGATGTCCAGGGGTCCCCGAGGGTGAGGCCCAGCCCCCTCCCGCCCCTGTCCACTGCCCCGGTCCCCCCAGAAGCCCCCAAAAGTAGAGGCTCAGGC | 8 |
| wt-32340-32460 | 3124 | CATGCGCGCCCTGTCACCAGGCCTGCCAAAGAGCCAGATCTAAGGCCGGGAGAGGCAGCCCCAAAGCGGGTGCAGTAACAGGTAATCTCTGGTAGTGATTTGGACCCGAAATCTGACACT | 4 |
| wt-32460-32580 | 3125 | TTAGAGCTCTGGAGGACTTTAAAACTCTAAAAATCAAAACTTTAGAGGCGAATGGGCGCCATTTTGTCCCCACGCGCGCATAATGGCGGACCTAGGCCTAAAACCCCCAGGAAGCGGGTC | 4 |

| | | | |
|---|---|---|---|
| wt-32580-32700 | 3126 | TATGGTTGGCTGCGCTGCTGCTATCTTTAGAGGGGAAAAGAGGAATAAGCCCCCAGACAGGGGAGTGGGCTTGTTTGTGACTTCACCAAAGGTCAGGGCCCAAGGGGGTTCGCGTTGCTA | 4 |
| wt-32700-32820 | 3127 | GGCCACCTTCTCAGTCCAGCGCGTTTACGTAAGCCAGACAGCAGCCAATTGTCAGTTCTAGGGAGGGGGACCACTGCCCCTGGTATAAAGTGGTCCTGCAGCTATTTCTGGTCGCATCAG | 4 |
| wt-32820-32940 | 3128 | AGCGCCAGGAGTCCACACAAATGTAAGAGGGGGTCTTCTACCTCTCCCTAGCCCTCCGCCCCCTCCAAGGACTCGGGCCCAGTTTCTAACTTTTCCCCTTCCCTCCCTCGTCTTGCCCTG | 4 |
| wt-32940-33060 | 3129 | CGCCCGGGGCCACCTTCATCACCGTCGCTGACTCCGCCATCCAAGCCTAGGGGAGACCGAAGTGAAGGCCCTGGACCAACCCGGCCCGGGCCCCCGGTATCGGGCCAGAGGTAAGTGGA | 8 |
| wt-33060-33180 | 3130 | CTTTAATTTTTTCTGCTAAGCCCAACACTCCACCACACCCAGGCACACACTACACACACCCACCCGTCTCAGGGTCCCCTCGGACAGCTCCTAAGAAGGCACCGGTCGCCCAGTCCTACC | 4 |
| wt-33180-33300 | 3131 | AGAGGGGGCCAAGAACCCAGACGAGTCCGTAGAAGGGTCCTCGTCCAGCAAGAAGAGGAGGTGGTAAGCGGTTCACCTTCAGGGGTAAGTAACCTGACCTCTCCAGGGCTCACATAAAGG | 4 |
| wt-33300-33420 | 3132 | GAGGCTTAGTATACATGCTTCTTGCTTTTCACAGGAACCTGGGGGCTAGTCTGGGTGGGATTAGGCTGCCTCAAGTTGCATCAGCCAGGGCTTCATGCCCTCCTCAGTTCCCTAGTCCCC | 4 |
| wt-33420-33540 | 3133 | GGGCTTCAGGCCCCCTCCGTCCCCGTCCTCCAGAGACCCGGGCTTCAGGCCCTGCCTCTCCTGTTACCCTTTTAGAACCACAGCCTGGACACATGTGCCAGACGCCTTGGCCTCTAAGGC | 4 |
| wt-33540-33660 | 3134 | CCTCGGGTCCCCCTGGACCCCGGCCTCAGCAACCCTGCTGCTCCCCTCCTGCCACCCCAGCCTCCCCCCCTCCCCGTCCCCCTTCGCTCCTGATCCTCCCCCGGTCCCCAGTAGGGCCGC | 12 |
| wt-33660-33780 | 3135 | CTGCCCCCCTGCACCCAGTACCTGCCCCTCTTGGCCACGCACCCCGGGCCAGGCCACCTTAGACCCGGCCAAGCCCCATCCCTGAAGACCCAGCGGCCATTCTCTCTGGTAACGAGCAGA | 8 |
| wt-33780-33900 | 3136 | GAAGAAGTAGAGGCCCGCGGCCATTGGGCCCAGATTGAGAGACCAGTCCAGGGGCCCGAGGTTGGAGCCAGCGGGCACCCGAGGTCCCAGCACCCGGTCCCTCCGGGGGGCAGAGACAGG | 8 |
| wt-33900-34020 | 3137 | CAGGGCCCCCCGGCAGCTGGCCCCGAGGAGGCGCCCGGAGTGGGGCCGGTCGGCTGGGCTGGCCGAGCCCGGGTCTGGGAGGTCTGGGGTGGCGAGCCTGCTGTCTCAGGAGGGGCCTGG | 12 |
| wt-34020-34140 | 3138 | CTCCGCCGGGTGGCCCTGGGGTAAGTCTGGGAGGCAGAGGGTCGGCCTAGGCCCGGGGAAGTGGAGGGGGATCGCCCGGGTCTCTGTTGGCAGAGTCCGGGCGATCCTCTGAGACCCTCC | 8 |
| wt-34140-34260 | 3139 | GGGCCCGGACGGTCGCCCTCAGCCCCCCAGACAGACCCCAGGGTCTCCAGGCAGGGTCCGGCATCTTCAGGGGCAGCAGGCTCACCACCACAGGCCCCCCAGACCCGGGTCTCGGCCAGC | 8 |
| wt-34260-34380 | 3140 | CGAGCCGACCGGCCCCGCGCCTGGCGCCTCCTCGGGGCCAGCCGCCGGGGTTGGTTCTGCCCCTCTCTCTGTCCTTCAGAGGAACCAGGGACCTCGGGCACCCCAGAGCCCCTCGGGCCC | 12 |
| wt-34380-34500 | 3141 | GCCTCCAGGCGCCCTCCTGGTCTCCGCTCCCCTCTGAGCCCCGTTAAACCCAAAGAATGTCTGAGGGGAGCCACCCTCGGGGCCCAGGCCCCAGAGTCCAGAGGTCAGGGGCACCTCAGG | 8 |
| wt-34500-34620 | 3142 | GTGCCTCCCCGGGTCCCAGGCCAGCCGGAGGGACCCCGGCAGCCCGGGCGGCCCCAGAGGCCGGTTCCTCGCCCCTTCCCCGGGCTTCAGAGCCCAGGATGTCCCCCAGAAGGGACCCTA | 12 |
| wt-34620-34740 | 3143 | GGCGTCCCCTCTCCTCCCCTCCAGGCCCGAGCCTCTCCCTCGCGGAGAGGGGCCTCTTTGGGCCCTCAAGTCCAGCCCCACCGAGACCCGAGTGGCCCGGATCCCCCCACCGGCCCTTCT | 8 |
| wt-34740-34860 | 3144 | CTCTGTCCCCCTGCTCCTCTCCAACCTTCGCTCCACCCTAGACCCCAGCTTCTGGCCTCCCCGGGTCCACCAGGCCAGCCGGAGGGACCCCGGCAGCCCGGGCGAGTCGCCTTCCCTCTC | 8 |
| wt-34860-34980 | 3145 | CCCTGGCCTCTCCTTCCCGCCTCCCACCCGAGCCCCCTCAGCTTGCCTCCCCACCGGGTCCATCAGGCCGGCCGGAGGGACCCCGGCGGCCCGGTGTCAGTCCCCCCTGCAGCCGCCCAG | 12 |
| wt-34980-35100 | 3146 | TCTCTGCCTCCAGGCAAGGGCGCCAGCTTTTCTCCCCCCAGCCTGAGGCCCAGTCTCCTGTGCACTGTCTGTAAAGTCCAGCCTCCCACGCCCGTCCACGGCTCCCGGGCCCAGCCTCGT | 8 |
| wt-35100-35220 | 3147 | CCACCCCTCCCCACGGTGGACAGGCCCTCTGTCCACCCGGGCCATCCCCGCCCCCCTGTGTCCACCCCAGTCCCGTCCAGGGGGGACTTTATGTGACCCTTGGGCCTGGCTCCCCATAGA | 8 |
| wt-35220-35340 | 3148 | CTCCCATGTAAGCCTGCCTCGAGTAGGTGCCTCCAGAGCCCCTTTTGCCCCCCTGGCGGCCCAGCCCGACCCCCGGGCGCCCCCAAACTTTGTCCAGATGTCCAGGGGTCCCCGAGGGTG | 8 |
| wt-35340-35460 | 3149 | AGGCCCAGCCCCCTCTCGCCCAAGCTGCTTTGATTCCTGGGATATTTTTGGGAATGGTGTTAACTTTCTCCCCTTGTATTTGCTATTCAATCAACCTGATTCCCCCTGCTCATACCTCCA | 4 |
| wt-35460-35580 | 3150 | CTTACAACCAAGCCACTACGGCCACGTCCCCGGCCTCCCGCTCGGGTAAGTGCTTTTTCATTTTTAGCCCCAGCCCCTCCTCTATAAGTTCTAGGCAAACCTCCAATCACCAGCCACCTT | 4 |
| wt-35580-35700 | 3151 | CCAATGTAGTCTCTTAGAGAGTGGCTGCTACGCATTAGAGACCACTTTGAGCCACCCACAGTAACCACCCAGCGCCAATCTGTCTACATAGAAGAAGAAGAGGATGAAGACTAAGTCACA | 4 |
| wt-35700-35820 | 3152 | GGCTTAGCCAGGTGATTTGTGAATTTCAGTTTATTTACTTTCTTCCAATCAAGCTTTCCCAGCCTCCGCTTGTTAGGTCCTAGTTATGGGTTTTCCATGGGGGACTTAGTATCCGTTCTA | 4 |
| wt-35820-35940 | 3153 | TTAGATTAACGTGCAAGACGCTAAACTTAACCAAGGTCAGCCAAGGGACGCGTGTTATCCCAGGCTGCCCACCCTGAGGATTTCCCCCCAAAATCCTCCTACCCTCTCTTTATGCCATGT | 4 |
| wt-35940-36060 | 3154 | GTGTTGTTGGCTTGTGTTAGTGCTATGTAATGCGTTGCCGCCAGGTGGCAGCCTGTTTATAGATGTGCAGTACCCCTTAATGTTAGGTCTGCTTTAGGGCTGCCAGGTGGCGCAATCTAG | 4 |
| wt-36060-36180 | 3155 | GATTAATTCACCTGTATCCCTTTCCCTCCACCCGCAGTAACCCAGCACTGGCGTGTGACGTGGTGTAAAGTTTTGCCTGAACCTGTGGTTGGGCAGGTACATGCCAACAACCTTCTAAGC | 4 |
| wt-36180-36300 | 3156 | ACCCGCGCTTGTGTTTTGCTTTATCTGCCGCCATCATGCCTACATTCTATCTTGCGTTACATGGGGGACAAACATATCATCTAATTGTTGACACGGATAGTCTTGGAAACCCGTCACTCT | 4 |
| wt-36300-36420 | 3157 | CAGTAATTCCCTCGAATCCCTACCAGGAACAACTGTCAGACACTCCATTAATTCCACTAACAATCTTTGTTGGGGAAAACACGGGGGTGCCCCCACCACTCCCACCACCCCCCCCACCAC | 4 |
| wt-36540-36660 | 3158 | AGCCATCACCTCTTGATAGGGATCCGCTAGGATATGACGTCGGGCATGGACCTCTAGCATCTGCTATGCGAATGCTTTGGATGGCTAATTATATTGTAAGACAATCACGGGGTGACCGGG | 4 |
| wt-36660-36780 | 3159 | GCCTTATTTTGCCACAAGGCCCACAAACAGCCCCTCAGGCCAGGTTGGTCCAGCCACATGTCCCCCCTCTACGCCCGACAGCACCCACCATTTTGTCACCTCTGTCACAACCGAGGCTTA | 4 |
| wt-36780-36900 | 3160 | CCCCTCCACAACCACTCATGATGCCACCAAGGCCTACCCCTCCTACCCCTCTGCCACCTGCAACACTAACGGTGCCACCAAGGCCTACCCGTCCTACCACTCTGCCACCCACACCACTAC | 4 |
| wt-36900-37020 | 3161 | TCACGGTACTACAAAGGCCTACCGAACTTCAACCCACACCATCACCACCACGCATGCATCTCCCTGTCTTGCATGTGCCAGACCAATCAATGCACCCTCTTACTCATCAAAGCACCCCAA | 4 |
| wt-37020-37140 | 3162 | ATGATCCAGATAGTCCAGAACCACGGTCCCCGACTGTATTTTATAACATTCCACCTATGCCATTACCCCCCTCACAATTGCCACCACCAGCAGCACCAGCACAGCCACCTCCAGGGGTCA | 4 |
| wt-37140-37260 | 3163 | TCAACGACCAACAATTACATCATCTACCCTCGGGGCCACCATGGTGGCCACCCATCTGCGACCCCCCGCAACCCTCTAAGACTCAAGGCCAGAGCCGGGGACAGAGCAGGGGGAGGGGCA | 4 |
| wt-37260-37380 | 3164 | GGGGCAGGGGCAGGGGCAGGGGCAAGGGCAAGTCCAGGGACAAGCAACGCAAGCCCGGTGGACCTTGGAGACCAGAGCCAAACACCTCCAGTCCTAGCATGCCTGAACTAAGTCCAGTCC | 4 |
| wt-37380-37500 | 3165 | TCGGTCTTCATCAGGGACAAGGGGCTGGGGACTCACCAACTCCTGGCCCATCCAATGCCGCCCCCGTTTGTAGAAATTCACACACGGCAACCCCTAACGTTTCACCAATACATGAACCGG | 4 |
| wt-37500-37620 | 3166 | AGTCCCATAATAGCCCAGAGGCTCCCATTCTCTTCCCCGATGATTGGTATCCTCCATCTATAGACCCCGCAGACTTAGACGAAAGTTGGGATTACATTTTTGAGACAACAGAATCTCCTA | 4 |
| wt-37620-37740 | 3167 | GCTCAGATGAAGATTATGTGGAGGGACCCAGTAAAAGACCTCGCCCCTCCATCCAGTAAAAACCCTTGCCCTCTCCAGCAACCAATGTATCCCAAATAAATGTTACTTCTTTTGCTCTTA | 4 |
| wt-37740-37860 | 3168 | ACCATTGACACGCCTGTCATTCTATCAATTAAACAAGGGAAAAAGGTTTAGCTATTCCACCAACACGACCCCAAGGAAGGCTTGCCAAAATTGGTGCCTTGCTCTCAGCACTTTGCCAGC | 4 |
| wt-37860-37980 | 3169 | AACTTATAGCATGGTAGGCAGCTCAACTCGGCCCGTCTTACTGCCCAGCCTACTCTCCACTCCCAGTCCATGTTCGCACTCCTATGCATTTCCTGCCCTCCCACTTTTACCCCAGTCCCA | 4 |
| wt-37980-38100 | 3170 | ACCCAAAACCACACACAACACATAGAATTGTTAGTTTAAACAGTTTATTGATAGGTGGCTGCTTTTAGCCTAATTGTGTATTGCTCTCGTTGCCAAAACCTGTTGTAAGGGCCGGCACCC | 4 |
| wt-38100-38220 | 3171 | GCAACATGGGGAAAACATAACCGCCGCCATCCCATGGGGAGGGTAGAGGCGGTTGACATGTAGGTGAGTAGTGTAAGAAGCATGGCGAAGTAGACAGGTTACTTTTAGAGTGTAGTGTAC | 4 |
| wt-38220-38340 | 3172 | AGGGCCGGGCGCAACAGTGCCACCAACCCGGGGTCTGAGCATTCCATGGGCAGCAGGGACACTGCACTACCGCCAGGTCCTGGGGCAGCCGGGGTTCCTGGCGCTCCGGGGGCAGCCGGG | 8 |
| wt-38340-38460 | 3173 | CGGCCGCCGGTGGGTCCGCTGGGCCGCTGCCCCGCTCCGGGTGGGGGGTGGCCCCGCTGGGCACCGCTGCGCCGCCGCCAGGTCCTGGGGCAGCCGGGGTTCCTGGCGCTCCGGGGGCAG | 12 |
| wt-38460-38580 | 3174 | CCGGGCGGCCGCCGGTGGGTCCGCTGGGCCGCTGCCCCGCTCCGGGTGGGGGGTGGCCCCGCTGGGCACCGCTGCGCCGCCGCCAGGTCCTGGGGCAGCCGGGGTTCCTGGCGCTCCGGG | 12 |
| wt-38580-38700 | 3175 | GGCAGCCGGGCGGCCGCCGGTGGGTCCGCTGGGCCGCTGCCCCGCTCCGGGTGGGGGGTGGCCCCGCTGGGCACCGCTGCGCCGCCGCCAGGTCCTGGGGCAGCCGGGGTTCCTGGCGCT | 12 |
| wt-38700-38820 | 3176 | CCGGGGGCAGCCGGGCGGCCGCCGGTGGGTCCGCTGGGCCGCTGCCCCGCTCCGGGTGGGGGGTGGCCCCGCTGGGCACCGCTGCGCCGCCGCCAGGTCCTGGGGCAGCCGGGGTTCCTG | 12 |

| | | | |
|---|---|---|---|
| wt-38820-38940 | 3177 | GCGCTCCGGGGGCAGCCGGGCGGCCGCCGGTGGGTCCGCTGGGCCGCTGCCCCGCTCCGGGTGGGGGTGGCCCCGCTGGGCACCGCTGCGCCGCCGCCAGGTCCTGGGGCAGCCGGGGT | 16 |
| wt-38940-39060 | 3178 | TCCTGGCGCTCCGGGGGCAGCCGGGCGGCCGCCGGTGGGTCCGCTGGGCCGCTGCCCCGCTCCGGGTGGGGGTGGCCCCGCTGGGCACCGCTGCGCCGCCGCCAGGTCCTGGGGCAGCC | 12 |
| wt-39060-39180 | 3179 | GGGGTTCCTGGCGCTCCGGGGGCAGCCGGGCGGCCGCCGGTGGGTCCGCTGGGCCGCTGCCCCGCTCCGGGTGGGGGTGGCCCCGCTGGGCACCGCTGCGCCGCCGCCAGGTCCTGGGG | 12 |
| wt-39180-39300 | 3180 | CAGCCGGGGTTCCTGGCGCTCCGGGGGCAGCCGGGCGGCCGCCGGTGGGTCCGCTGGGCCGCTGCCCCGCTCCGGGTGGGGGTGGCCCCGCTGGGCACCGCTGCGCCGCCGCCAGGTCC | 12 |
| wt-39300-39420 | 3181 | TGGGGCAGCCGGGGTTCCTGGCGCTCCGGGGGCAGCCGGGCGGCCGCCGGTGGGTCCGCTGGGCCGCTGCCCCGCTCCGGGTGGGGGTGGCCCCGCTGGGCACCGCTGCGCCGCCGCCA | 12 |
| wt-39420-39540 | 3182 | GGTCCTGGGGCAGCCGGGGTTCCTGGCGCTCCGGGGGCAGCCGGGCGGCCGCCGGTGGGTCCGCTGGGCCGCTGCCCCGCTCCGGGTGGGGGTGGCCCCGCTGGGCACCGCTGCGCCGC | 12 |
| wt-39540-39660 | 3183 | CGCCAGGTCCTGGGGCAGCCGGGGTTCCTGGCGCTCCGGGGGCAGCCGGGCGGCCGCCGGTGGGTCCGCTGGGCCGCTGCCCCGCTCCGGGTGGGGGTGGCCCCGCTGGGCACCGCTGC | 12 |
| wt-39660-39780 | 3184 | GCCGCCGCCAGGTCCTGGGGCAGCCGGGGTTCCTGGCGCTCCGGGGGCAGCCGGGCGGCCGCCGGTGGGTCCGCTGGGCCGCTGCCCCGCTCCGGGTGGGGGTGGCCCCGCTGGGCACC | 12 |
| wt-39780-39900 | 3185 | GCTGCGCCGCCGCCAGGTCCTGGGGCAGCCGGGGTTCCTGGCGCTCCACTGCACCTGGAATGCAGGGTGGGGGCGTGGTCCCCTGGACCCCAGCCCCGCCGATCCCTCCCCCAGGGCGTA | 12 |
| wt-39900-40020 | 3186 | CCCGGCTTGCCTGGTTCTGGGGCTCCTCTGGGGGTCGCTGCATCCGCCGGTAGGGTTCGAATGGGCGTGGTCCGCTTGCTCTGCTGGCCCGGTACGCCTGGATTGCCGGCTGGGGGCTGG | 8 |
| wt-40020-40140 | 3187 | GGTCCCGGGACGCCCCCTCCCTGCTCCCACCCGGTTCCCTCCCCCAGGGCGTGCCCCGCTTGCCTGGTCCTGGAGCTCATCCGGGGATGCTGCATCCGCTAGTCCGACCTGGGTGGGTGC | 8 |
| wt-40140-40260 | 3188 | GGTCCGCTGGCCCCACCCTGGGGGTAGCCGCCGGGTCTGCTGGTCCGGTGCACCTGGAAGGCAGGGGGGGGGGCAGTGAGGGAGGGGCGTGGTCCTGGGACCCCGCGCCGACTGGCAGGG | 12 |
| wt-40260-40380 | 3189 | GGTCCCCATGGCACAGGCCTAGGGGTCCAGGGGGCAGCCGCGGCCCAGCGCGCCCCGTTCACGGGGGAGGACCGCGGCCGAGCCACCAGGGGCCCGGCGGGGGTGGGGGGTGCGCTCCCA | 12 |
| wt-40380-40500 | 3190 | GGCCGGACCCTGGTGCCAGGCAGGGACCCCGCGCCACCCGCTTCATGGGGGGGGAGGCCGCCGCAAGGACGCCGGGCCGGCTGGGAGGTGTGCACCCCCCGAGCGTCTGGACGACGCTGG | 12 |
| wt-40500-40620 | 3191 | CGAGCCGGGCCGGCTCGCCTTCTTTTATCCTCTTTTTGGGGTCTCTGTGTAATACTTTAAGGTTTGCTCAGGAGTGGGGGCTTCTTATTGGTTAATTCAGGTGTGTCATTTTAGCCCGTT | 4 |
| wt-40620-40740 | 3192 | GGGTTTCATTAAGGTGTGTCACCAGGTGGGTGGTACCTGGAGGTTATTCTATTGGGATAACGAGAGGAGGAGGGGCTAGAGGTCCGCGAGATTTGGGGTAGGCGGAGCCTCAGGAGGGTC | 4 |
| wt-40740-40860 | 3193 | CCCTCCATAGGGTTGAACCAGGAGGGGGAGGATTGGGCTCCGCCCCGATATACCTAGTGGGTGGAGCCTAGAGGTAGGTATCCATAGGGTTCCATTATCCTGGAGGTATCCTAAGCTCCG | 4 |
| wt-40860-40980 | 3194 | CCCCTATATACCAGGTGGGTGGAGCTAGGTAGGATTCAGCTAGGTTCCTACTGGGGTACCCCCCTACCCTACCTTAAGGTGCGCCACCCTTCCTCCTTCCGTTTTAATGGTAGAATAACC | 4 |
| wt-40980-41100 | 3195 | TATAGGTTATTAACCTAGTGGTGGAATAGGGTATTGCAGCTGGGTATATACCTATAGGTATATAGAACCTAGAGGAAGGGAACCCTATAGTGTAATCCCTCCCCCCCCTACCCCCCCCTC | 4 |
| wt-41100-41220 | 3196 | CCTTACGGTTGCCTGAGCCCATCCCCCACCCCAGCACCCCGGGGTGACGTGGCACCCCGCGTGCCTTACTGACTTGTCACCTTTGCACATTTGGTCAGCTGACCGATGCTCGCCACTTCC | 4 |
| wt-41220-41340 | 3197 | TGGGTCATGACCTGGCCTGTGCCTTGTCCCGTGGACAATGTCCCTCCAGCGTGGTGGCTGCCTTTGGGATGCATCACTTTGAGCCACTAAGCCCCCGTTGCTCGCCTTGCCTGCCTCACC | 4 |
| wt-41340-41460 | 3198 | ATGACACACTAAGCCCCTGCTAATCCATGAGCCCCGCCTTTAGGAAGCACCACGTCCCGGGGACGGAAGGGGACTTGGGGTGATTTTCTATGTGGGGGTGGAAATATGAGCAAGAATAAG | 4 |
| wt-41460-41580 | 3199 | GACGGCTCCTTATTAACCTGATCAGCCCCGGAGTTGCCTGTTTCATCACTAACCCCGGGCCTGAAGAGGTTGACAAGAAGGGTCAAGGTTTCGTCTGTGTGTTGAAGGGCAGGGGCTGTT | 4 |
| wt-41580-41700 | 3200 | GGGTGCATCTGGAACGGCTTACCTCGGGTAACTGTTTGCCATTAAAAGGTTGGGGATTAGGTTTAGCCCCTTTAGCTGCCATTTCGAACCGGGGTGTGCAGATGCAGGTCTCCGGGTGGG | 4 |
| wt-41700-41820 | 3201 | CAGGCAGTACGAGATGTCACGTTGTGTTGTCTTTCCTCCCACCCCTGTCCTGGCTGTGGCAAATGCGACCCTCATAGAGTTGTGTTTCAGGTCTGTGTCCTGTTTTGCGGTGGGTTATTT | 4 |
| wt-41820-41940 | 3202 | CTTCCCTCAGTGTTTGCCAGCTTATTTCCCCAGTTTTCACGTACTGGGGCCTGTGGACACCTGAGGGAGCGGCCGTTGGTGGGTATGTGTTGGAATTGCTCCCACCCTCAATTTTCGCTT | 4 |
| wt-41940-42060 | 3203 | GCCTTCTTCCCTTGTTAACCTGATAGCATAGCCTCTAGGTTTCCTTGTAGGTCTGTTTGGGTTTGTTGGTTCACGTGGTGCTAACTTGAATTTTTTGGTTTTCTAGTTCCCTCTTAATTA | 4 |
| wt-42060-42180 | 3204 | CATTTGTGCCAGATCTTGTAGAGCAAGATGGCCTATTCAACAAGGGAGATACTGTTAGCCCTGTGTATACGGGACAGTCGTGTGCATGGAAATGGTACCCTGCATCCTGTGTTGGAGCTA | 4 |
| wt-42180-42300 | 3205 | GCAGCAAGAGAAACACCTCTCCGCCTTTCGCCAGAGGACACTGTAGTTCTGCGTTATCATGTGTTGCTTGAGGAGATAATTGAACGAAATTCAGAGACATTTACAGAAACTTGGAACAGA | 4 |
| wt-42300-42420 | 3206 | TTTATAACACACACCGAACATGTGGATCTGGATTTTAACTCAGTATTTTTAGAGATATTTCACCGTGGAGACCCAAGCCTTGGGCGCGCGTTGGCCTGGATGGCCTGGTGCATGCATGCC | 4 |
| wt-42420-42540 | 3207 | TGCAGGACATTGTGTTGTAACCAGTCTACTCCTTACTATGTTGTGGACCTGTCAGTTCGTGGGATGTTAGAAGCCAGCGAAGGCCTGGATGGTTGGATTCATCAACAGGGCGGCTGGTCT | 4 |
| wt-42540-42660 | 3208 | ACATTAATTGAAGACAACATTCCTGGATCCAGAAGGTTTAGCTGGACTTTGTTTCTTGCTGGACTGACTTTGAGTCTGTTAGTTATATGTAGTTATTTATTTATCTCCAGAGGAAGACAC | 4 |
| wt-42660-42780 | 3209 | TAATCTATACATTTTCTCAGCACTTTATATGAATCAGGGTCATTGGGCCTGCGGGGAACTGAGCCAGTAGGATATTAGGCAAGGGTGACACAGTGCCCATGCATTATAATTTAACCAAAC | 4 |
| wt-42780-42900 | 3210 | AGTGGTCGTGAGTTTTAGGCCGGCCATGGGGGCTTACAAGAATAACATGCCAATGACCCGGCCCCCACTTTTAAATTCTGTTGCAGCAGATAGCTGATACCCAATGTTATCTTTTGCGGC | 4 |
| wt-42900-43020 | 3211 | AGAAATTGAAAGTGCTGGCCATATCTACAATTGGGTGTCCTAGGTGGGATATACGCCTGTGGTGTTCTAACGGGAAGTGTGTAAGCACACACGTAATTTGCAAGCGGTGCTTCACGCTCT | 4 |
| wt-43020-43140 | 3212 | TCGTTAAAATAACACAAGGACAAGATACTAAAGAAATAACTGAGGTGAGTGTGGGAAGATGGGAATACTATGTGTTATGTTAACGGGTGAGAGCCTATACTGCAGCCCAGACTCGGGGGG | 4 |
| wt-43140-43260 | 3213 | AGGAGGAAATGGTAAGAGTTATACTCTACTTATCTTTTTTGACACTACATTTAACTGTTATGTAACAATGTTTGCTTATTTTCATGTTCAATAAACGCTATGTTAATGATGAAGAACCTG | 4 |
| wt-43260-43380 | 3214 | TGTTCTTTGGAAGTGGGCCCAATGGGGTAGTAGGTTTTGGGAGGGTGCCGTGCTAGATATTTCAACTGCCACAGACCCCATTTTGTCCCACCTGTTACCACATTCTAGGTCCTGCATCCA | 4 |
| wt-43380-43500 | 3215 | GTGGGCCAGGTGTCTCACCATGGCTCTTTCTAGGTGGATACCACAGTCCAGGCCCCCAAGGCTACCGTGCTAATTACCTCCTCATGTCCACCCCCACCCTGTGTTACTGTCGCCTGATTA | 4 |
| wt-43500-43620 | 3216 | TCCTGGCTTAGCAGCCTCCAAGTTTTACAAGACGTCCCATTGCCCTGCCCTTGGTCCAAGTCTCGCCGGTTTTCAGCAGCCTGTTGTAGCCTGCCCCCAAGTTTCGCAGGTTTCCCCCAT | 4 |
| wt-43620-43740 | 3217 | GCTTCCACCCGTTAACCCAATAGCATGACAGCCAATCCAACACGAGGCAAGTTTTAAGAGTTAAAAGCAACTACTGTTTATTTTCCAAAATGAGCTGGGTATAGTTGATGATCTGTAGGC | 4 |
| wt-43740-43860 | 3218 | GCAGCTCATCCCCACATTCCAGGTCCTTGATGGCCTCGTAGATGGCATCTTCGTCGACATTGACAGCCTTCTCATATACCGTGTCTCTGGGGCTGACCTTTATACAGAAGGCGTCCCCTA | 4 |
| wt-43860-43980 | 3219 | CTAGGTCCACGGCCAGCTCGTAGGTGGGGCCTATGTTTTCACATAACAGTTTCAAGCAGGTCTCTGGGATGTGAAGGGAGGTGCCCTGGAGCAGGAGATGCATGATTAGGCGCCCTTTTC | 4 |
| wt-43980-44100 | 3220 | CATTTGTGCTGAAGATGGGGCAGATGGTGCCACAAAAGTGTCCGGTGACCAGGTAAGCGTAGAGAAGGCTGGGTTGGGAAAGTCCAGCCTTTACTGCACTGGGAGAGCTGCTGAGCAGAG | 4 |
| wt-44100-44220 | 3221 | ACACATAGAAGGTCTTGTTGGGTATTATCTTGTGGACATTGTTGAAGAAGGAGAGCTGGGTGGAGCTAAACTCCTGAGGCACATGAACCTGGGACCTATTGATGCAGATCTCGCAGTGAG | 4 |
| wt-44220-44340 | 3222 | ACCCCAGAGTCAGGCTGTGGCCGAAGGGAGACAGGCGAAGGCAGCGCCCGGGGGAGAGAGTGCACAGTGACAGTGGGAGAAACACGGCCTCTGAGACATGTATGGGGGTGTTCATCTCAC | 4 |
| wt-44340-44460 | 3223 | GCAGAAAATCTTTGCCCAGCTCAAAGTTGGCAGAGATTCCCCTGAAGAAGTCCCGTAGTGAAAAATGGGATCTGTCTACACCATGTCTGGTGTGCCGGGAACATATTGATCGGCCACAC | 4 |
| wt-44460-44580 | 3224 | TGCCAACCCTTTCCATTCTTCCCAGCTCTGAGCGAGATTTTCCACACCTGGACACCGACTTCACGCTATGCGCCGAGGCCTTTGAGGCCGTGTAGTTTCTGTGGTGCGGATGCATTAGGC | 4 |
| wt-44580-44700 | 3225 | GGCGCAATGCGGGATCTGCCGGTCGCTGTTGGCGTGCATTCACGGCATCTGGGGTGACCGGGGCCATCGGGTTTACTTTTCACACGTAGACCTGGGAAGTTTGATAGGACTGTACCAGGT | 4 |
| wt-44700-44820 | 3226 | CAAGGCCGTGGATGCGCAGGACCACGTCCAGTTCCTTAGTGACATCCACGAGGATTGTTTTGCCCACTCTGGCCACTTGTGTGGATTTAAATATGTACACAAGCGTAATTAACGAGTCAC | 4 |
| wt-44820-44940 | 3227 | AGACCCCCTGTTCCAGATTCTGACCGGCTGCAAGCGCTGCCTTAAAGGCCTGGAAGCTGGGTGGGTAAATCTGACCAAACAGCACGCTCGGATTCGTGATGCTGTGGTTGATGGCACACA | 4 |

| | | | |
|---|---|---|---|
| wt-44940-45060 | 3228 | GGGGGTCGCAGAACAGGTGCTTGTGGAAGTCTTGCGGGGTGCACATCTGCAGCCAGGCCCTTAGCCTGGGGCATGGCACATCCAGCAGCGTGTTTTGGGTCTTGATGAGGAACACGATCC | 4 |
| wt-45060-45180 | 3229 | TGTCTAGGATTTTGATGTTGTTGCCGAACGAGTCAAGAATCAGGCTCTTGAAGCGGTCAAGGGTGTCCTTGGCGTCCGGGTGGGCCCCGAGGCTCTCGCAGAGTGGGCAGATGGTCCGTG | 4 |
| wt-45180-45300 | 3230 | AGGCATTCTTGTGCCTTAGTCCAAACATGGGGGCCAGGAGGCAGGGGCCCTGCGAATGGTCGCCAGCCTCCGGTCTGGTGATGGCCAGGGCCAACTCCGCCAGCTCATCGCCGCTGTATT | 4 |
| wt-45300-45420 | 3231 | CCGCGTTTAAACCGATAGCATGGTGGCCTGGCCCCCCGAGCAGGTCCGTCCCCTGCCACGTACCTAATAGTAGTCCACAGTAGTCGGCCTTGGTTGTAATTTCAGGAGAGAGTCCTCCCT | 4 |
| wt-45420-45540 | 3232 | TTTCGGCCCTGAGAAATGGATGCTGAACTCGGTTTCTGGTAGGCAGGTGGCAGCACAGGGCGGTGTACAGGCCCCTGCCGACGTCCCCTGGGACATCCTGGGAATCTTTGCAGGTTCTGG | 4 |
| wt-45540-45660 | 3233 | GTCCAGGGAGGGTAAGAAAAGTGGGGGTGGTTCTGGGCCACATGGACTTGAAGCAGAAGTTGGCCGGGGACTGGCCGGTGAGGATGGATTTCAGAAACTCCAATTTGTAGTAGCCGAGGT | 4 |
| wt-45660-45780 | 3234 | TGGCATTTCTAATCATGTCAGAAGAGGACACAGGGAGGAAGCACCGGCAAATGTAAAAGTGAAGCTGGATGTCAATGGCAAGAATCCTGGAGGGCATGAAGAGGGAATCCAACCCCCCGG | 4 |
| wt-45780-45900 | 3235 | CCATGGGGAAGTATTTTATCAGGATGTGTAAAAAGTCCATGCCTGTGATGAGGCTAGAGATCCAGGCTCGTGGGGCATTTAGACAGTAGTAGCAGAGCAGGGCATAGTCCTCAAAGAAGG | 4 |
| wt-45900-46020 | 3236 | CCACGGGGGCATCTGAGTGATTGACCAGGGTGTCGAGCAGATCACAAACTCGGCAGGTGCTGGCTGGAGAGAGGGACTCGTAGGTGTGGACGAGTGGTGGGTAGGCTATGCCTTCTTCCG | 4 |
| wt-46020-46140 | 3237 | CGTTGGCTGGAAGATAGGAGTGGGCCATCAAAAGGCCGACTGCCTCGAACTGGCTTTTCAGATTGTCCACGGTCCAGGGCACAAAGTCCTCCATCTTTGGAGTTCTGCCCGCGATCTGTG | 4 |
| wt-46140-46260 | 3238 | CCACCTCTGTTACGCCACTCCTCGTGAGGGGGCAGCTGGACAGTCTTTTTCCGGTCAGGGGGTTTGGCTCGTTTGCGCTCGTGACTTTGTGAGCCATGACACATCTGGGTGGCAAGGTGA | 4 |
| wt-46260-46380 | 3239 | GGTCTTCTGGGTTTTTAATACCGGGGTCGGCACCAGTTTCTGGGACACCGCCACAAGGACAAGGTGGGCTAGCAAGTTCTCGAGTCTACGAAGACTCCGGGGGCAGTCTTTTGAGTTTCT | 4 |
| wt-46380-46500 | 3240 | CGCCTATGATCCACCCCAATCTCGCCCCCCTAATTGCGCCATCTGCCTACGCGAGGCTGAACCTCCTGAATCACTGCATCTTTCTTGAGGCGTTTAAAGAAGAGAATAGTGGCCAGGGCC | 4 |
| wt-46500-46620 | 3241 | TCGGTGGGGTCCAGCGTGAGGTCTTATTTTTGAAAAGGGATATTATAAAACAGGTCATTGCTCGGATTGTGGCAGCCGATAGCACCCTAGATCTAGTGAATCATGGCGAGCCCGGAAGAG | 4 |
| wt-46620-46740 | 3242 | AGGCTCCTAGACGAGCTCAATAACGTAATTGTGTCATTTCTGTGTGACTCTGGGTCTCTGGAAGTGGAGAGATGCTCCGGGGCGCATGTGTTCTCCAGGGGCAGCTCCCAACCCCTCTGC | 4 |
| wt-46740-46860 | 3243 | ACCGTGAAGCTGCGCCACGGACAGATTTACCACCTGGAGTTTGTCTACAAGTTCCTGGCCTTTAAGCTGAAGAACTGCAACTACCCCTCCTCGCCCGTGTTTGTGATATCCAACAACGGC | 4 |
| wt-46860-46980 | 3244 | CTGGCCACCACCCTGAGGTGCTTTTTGCACGAGCCGTCGGGTCTCAGATCGGGCCAGAGCGGCCCTTGCCTGGGTCTCTCAACGGATGTTGACCTACCAAAGAACTCCATCATTATGCTG | 4 |
| wt-46980-47100 | 3245 | GGCCAGGATGACTTCATTAAGTTCAAAAGCCCCCTGGTCTTCCCTGCTGAGCTTGATCTCCTGAAATCTATGGTGGTCTGCCGGGCCTACATCACGGAACACCGGACGACGATGCAGTTT | 4 |
| wt-47100-47220 | 3246 | CTGGTGTTTCAGGCCGCCAACGCCCAGAAGGCCTCGCGGGTCATGGATATGATTAGTGATATGTCTCAGCAACTGTCTCGGTCTGGTCAAGTCGAGGATACGGGCGCCAGAGTCACAGGT | 4 |
| wt-47220-47340 | 3247 | GGAGGAGGTCCCAGGCCTGGCGTCACGCACTCGGGGTGTCTTGGGGACTCACACGTTAGGGGGCGCGGTGGTTGGGACTTGGATAACTTTTCAGAAGCTGAGACCGAAGACGAGGCGAGT | 4 |
| wt-47340-47460 | 3248 | TACGCTCCTTGGAGGGACAAAGACTCGTGGTCGGAATCCGAGGCGGCGCCGTGGAAGAAGGAACTCGTGAGGCACCCCATCCGCAGGCACCGGACACGCGAGACTCGCCGTATGCGCGGG | 8 |
| wt-47460-47580 | 3249 | AGCCATTCACGGGTGGAACACGTGCCCCCCGAGACCCGGGAGACGGTGGTGGGGGGAGCATGGCGTTATTCTTGGCGCGCCACACCTTATCTGGCACGGGTGCTGGCTGTCACGGCCGTG | 8 |
| wt-47580-47700 | 3250 | GCCCTGCTCCTGATGTTTCTGAGGTGGACCTGACGTTGCAGGCCCTTGGGGAGCGGGGGTTCTCCAGGCTCCTGGATCTGGGGCTGGCCTGCCTGGATCTGAGCTATGTGGAAATGAGGG | 4 |
| wt-47700-47820 | 3251 | AATTTGTGGTTTGGGGCAGGCCCCCAGCTTCTGAGGCGGCTGTGGCCTCTACGCCAGGCTCGCTTTTCCGAAGCCACTCGTCCGCCTACTGGTTGTCGGAGGTGGAGAGGCCCGGGGGCC | 8 |
| wt-47940-48060 | 3253 | GCAACGCCTTTTTCCTAGCTTATTTTTTGGTTGTGCGGTCTGTTTTCCCCGCGTTTTCCGATAGAATAGCTGCCTGGATCTGCGACCGGTCCCCTTTCTGCGAAAACACCCGGGCCGTGG | 4 |
| wt-48060-48180 | 3254 | CCAGGGGTTACCGAGGCCTCGTGAAGAGGTTCTTGGCATTCGTGTTTGAGCGTAGTAGCTATGACCCCCCCTTGTTGAGGCAAAACTCTAGGCCTGTGGAGCGCTGCTTTGCCATCAAGA | 4 |
| wt-48180-48300 | 3255 | ATTATGTCCCGGGCCTGGACTCACAAAGCTGTGTGACGGTCCCGAGCTTCTCCCGCTGGGCCCAGTCTCACGCCAGCGAGCTCGATCCCCGGGAGATTCGCGACAGAGTTACACCAGCGA | 4 |
| wt-48300-48420 | 3256 | CTGCACCTTCGTTCGTGGCTGATCATGCCTCGGCTCTATTGGCCTCCCTCCAGAAGAAGGCCTCCGACACCCCCTGTGGGAATCCCATTCAGTGGATGTGGTACCGCCTGTTGGTAAACT | 4 |
| wt-48420-48540 | 3257 | CGTGCCTGAGGAGTGCCCACTGTCTTCTGCCTATACCTGCCGTCTCTGAGGGGGGGAGAAAGACGGGCGGGGGCGTAGGGGAGGAGCTCGTGGGGGCCGGGGGGCCCTGCCTGAGCCGGG | 8 |
| wt-48540-48660 | 3258 | ATGTTTTCGTGGCGATCGTAAGCCGCAATGTTCTCTCGTGTCTGCTGAACGTGCCTGCCGCGGGTCCCCGGGCCTACAAGTGTTTCAGATCCCACGCCTCCAGACCGGTGTCTGGCCCGG | 4 |
| wt-48660-48780 | 3259 | ATTACCCTCCCTTGGCCGTGTTTTGCATGGACTGCGGTTACTGCTTGAACTTTGGAAAGCAGACAGGTGTAGGAGGCAGGCTCAATTCCTTTAGACCCACTCTCCAGTTTTATCCCCGTG | 4 |
| wt-48780-48900 | 3260 | ACCAGAAGGAGAAGCATGTGCTGACCTGCCATGCCAGCGGCCGTGTGTACTGCTCCAACTGCGGCTCTGCGGCGGTGGGCTGCCAGAGGCTGGCTGAGCCACCGAGCGCCCGCTCGGGCT | 8 |
| wt-48900-49020 | 3261 | GGCGGCCCCGAATCCGGGCAGTGCTGCCGCACAACGCGGCCTACGAGCTCGACCGTGGCTCCCGCCTCTTGGATGCCATCATCCCCTGCTTGGGACCCGACCGCACTTGCATGCGGCCGG | 8 |
| wt-49020-49140 | 3262 | TGGTCCTGCGGGGGGTGACGGTCAGGCAGCTCCTGTATTTAACTTTGCGGACAGAGGCCAGAGCCGTTTGCTCCATCTGTCAGCAACGCCAAGCTCCAGAGGACGCCCGCGACGAGCCTC | 4 |
| wt-49140-49260 | 3263 | ACCTGTTCTCCTCCTGTTTAGAGGTAGAATTGCCACCTGGTGAGCGGTGTGCGGGCTGCCGTCTCTATCAGACGCGTTATGGCACGCCGGCTGCCCAAGCCCACCCTCCAGGGGAGGCTG | 4 |
| wt-49260-49380 | 3264 | GAGGCGGATTTTCCAGACAGTCCCCTGCTTCCTAAATTTCAAGAGCTGAACCAGAATAATCTCCCCAATGATGTTTTTCGGGAGGCTCAAAGAAGTTACCTGGTATTTCTGACATCCCAG | 4 |
| wt-49380-49500 | 3265 | TTCTGCTACGAAGAGTACGTGCAGAGGACTTTTGGGGTGCCTCGGCGCCAACGCGCCATAGACAAGAGGCAGAGAGCCAGTGTGGCTGGGGCTGGTGCTCATGCACACCTTGGCGGGTCA | 4 |
| wt-49500-49620 | 3266 | TCCGCCACCCCCGTCCAGCAGGCTCAGGCCGCCGCATCCGCTGGGACCGGGGCCTTGGCATCATCAGCGCCGTCCACGGCCGTAGCCCAGTCCGCGACCCCCTCTGTTTCTTCATCTATT | 8 |
| wt-49620-49740 | 3267 | AGCAGCCTCCGGGCCGCGACTTCGGGGGCGACTGCCGCCGCCTCCGCCGCCGCAGCCGTCGATACCGGGTCAGGTGGCGGGGGACAACCCCACGACACCGCCCCACGCGGGGCACGTAAG | 12 |
| wt-49740-49860 | 3268 | AAACAGTAGAGGGCACGAAACATGGTGTATGCACTTTATTAATAAACAATTACAGATACAAAAACTTGAGTCTCTCGAGGTCTGCGATGAGGCGGTGGGTGGAACGCTCCAGCTTGCGGC | 4 |
| wt-49860-49980 | 3269 | GAAGCTGGCTCACGAAGCGAGACAGTACTCGGCTAGCCTGACTAAGGGTGAGGCTATAACGCAGGTCCTGTTCCGGGGCGGCGGTGGATAGAGAGGAGGGGGATCCGGAGGGGACCACTA | 4 |
| wt-49980-50100 | 3270 | GGTCGCCGGAGGTCGACCCTCCTGTCACCACCTCCCTGATAATGTCTTCAATAGACAGAAATTGGGTGACCACTGAGGGAGTGTTCCACAGTAATGTTGTCTGGTCGCTAGATGGCGCGG | 4 |
| wt-50100-50220 | 3271 | GTGAGGCCACGCTTTGCGAAAACGAAAGTGCTTGAAAAGGCGCGGGATAGCGTGCGCTACCGGATGGCGGGTAATACATGCTATCCTTACATTTTGGCATTTTGGGCAGCTGGGAGGCGG | 4 |
| wt-50220-50340 | 3272 | CGGATGGGGGTGCTTCTTTTCGCACGGTGTATGTTTGGGGACCCGCATGCCGGTACTGGGATAGGCGCACCTCGGGCCGCGCGCCAGGCTCCGAGCCGGAATGCATTGGGGGCAATGGGA | 8 |
| wt-50340-50460 | 3273 | TTGCGGGGGATTGTTGCTGTCTGCTCCTGACAGGGAGAGACACGCGCGGCGGAGATGCAGCCGACGGCGGGGCCGCGGTGGGCTGCCCCCGAGGACGGGCGCCGGCCGCCAGCGCCCCCG | 12 |
| wt-50460-50580 | 3274 | TGGCCTTTGGCACGGGCCTGGCACCCACCGCTTTAATTGTGGGGGTGGGCAGGGCAGCTGCATCTTGGGGCCTTTGTGCTTGCGTTTTTTGGGGGCGCGGTGCCAATGCACCAACTGGGG | 4 |
| wt-50580-50700 | 3275 | TGTGCGCCGGGGCGGCCAAGCCGGACCCCAGGGCGGGTGCCTGGGGGATGGGAAAGCCGGACGGCGCTTCTCCCGGGTCGAACGCTGGAGTAGCGGAGGCTGCTGCGCCGGCGGCCACCA | 12 |
| wt-50700-50820 | 3276 | CGGGCGCACGGGGTCGCAGCCCGACGGCCGTGGGGAGGCGGGTGGCGGAGGGCCGAATCTCCGCGGCTTCTTCCCGGCCCCCCTGCTGTTTCTTCTCCCGTTGCATGATAGAATGGCCAT | 8 |
| wt-50820-50940 | 3277 | AGGGTGGGTCCTGAGAGGAGGCTTGTGTGTCCTGGGGCTGGAGCCCAAAAGTCGTTAAAGATGCCGCTGATGGTGTGGGAGCTATGCCTCCCGTCGACTGGCCGGGCTTGTAGGGGGCTG | 4 |
| wt-50940-51060 | 3278 | AGGGTGGATAACTGGGCTTCTGTGAAGGCACCAACCCTGGAATCTGGATGGTATGTTTCTTCTGTGACCCCGAGGCAGTCGATGGTGTAGAGTGTGGAGACAATGTGTAGACGATGGGCC | 4 |
| wt-51060-51180 | 3279 | CTTGTTCAGAAGCCCAGGGACTTGAGGGGGGCTGTTGTGGTGCTGGTTGGGGAAGGAGCTCCAGGGAATCTTTGGCCATGGCCTTGGGGAGCTTCCCGGAGACCGGTCTGGGCTCTCGG | 4 |

| | | | |
|---|---|---|---|
| wt-51180-51300 | 3280 | AAGCCCTCGTTTCGGCCCCGAAATAGGGCCTTGCCATCAATCGGGGGCCTGGGAGAGTGATGGGGGCGGCCAATCCCGGGGTAACTGTCACGTCCCGGGGGGAGGAGGTAGGAGACAGCC | 8 |
| wt-51300-51420 | 3281 | AGTCCCTGGGCCTGCCAGGGGCCACCTTCTCTAAGAGGGGGCTCTGTGGGCTGGGAGGGCCAGAGGCCTCAGATTCAGCAGTAGTGCTCCCCTTTTCCCCCTGGTCCGTCTCCCCTCCTC | 8 |
| wt-51420-51540 | 3282 | CCAACTGCTGGAGCCGGTCGGAGGAGGCCGGGGTGTTATCTGCTGACTGAAACCCGTCCCCGCTGACCAGTCCGTGCCCCACCCTTGGGGGGAAACCGGAGAACAGCTCCTGGACGTTGC | 8 |
| wt-51540-51660 | 3283 | GTGGATTCGGGGGAAGCTGGTATCCAACCGGCAGTGGAGGATCTTCGTGCTCGTAGAAGGAGGGGTTGAGTACATCGGTCGGCCATCGTGAGGCCCCGGCCGCGTTAAAGTAGAACTGCA | 4 |
| wt-51660-51780 | 3284 | CGTCCGGCAGATTGTGCCGATAGGTGAAACACTTCCAGATGATGTTTTTTCTGTTGGCCAGGATGGCCACGGTGGGGGGCCTGGCCTCCTTAGGTTTGGCGGCCCTGGCCTCGGTGAGAA | 4 |
| wt-51780-51900 | 3285 | GCTCGCGTAGCCACACGGCCTGGCGTGCAAAGATGGACATCTCTGGCTCGAAAGACTCGGAGTAGCCGTCCAGGTCCTGCAGAAAATTCAGCGAGATGGTCTCCACCAGGGACCGGAAGG | 4 |
| wt-51900-52020 | 3286 | GCTCAGAGTGCCCGTCGCAGTAGAGGAGGGGAGCAACGACCCTGACCTGTCCCAGGGTCTTCAGGTTAAACAGATATTGAGAGGAGACAAAGAGAGTTAGGGGCCGACCGAGGAAGGCCG | 4 |
| wt-52020-52140 | 3287 | CCGCCACGGCCGCCTCAAAAACGGAGACGGGGATGGTGTCACCGGAGCCCCTCTTAGGACCGGTAATGGGAGTGCCATAAGGCATAAGATTTCTCAGGGCCCGGCCGGTAACGGTGCCGT | 4 |
| wt-52140-52260 | 3288 | AGGAAGACGGGGTTTCGCGGGGGACCTCGAGTCCCTCCGCCCTGGGGAGCTCTTCTCCGCGTGTATAGGCCTGCTTCACAAAGTCGCGCAGGTAGTCCTGAAATGCGACCGGGCCCTCCA | 4 |
| wt-52260-52380 | 3289 | GCGGGCGCAATGAGTGCCAGAGCTGCTGAAGGGCCTCGGGGGCGAAGCACCGGCGTGCGAGGAGCAGCATGCAGGCTCGGGCGCGGGCCGTACTTTGGTTGTGGACCAGGCCCAAGAACT | 8 |
| wt-52380-52500 | 3290 | CGGGGTGCGGCCAGAGGGCGGCTCGGGTATCCATCTCCTCCCAGGCGTCCTGGAAGAAGATGAAGCCGGTGGGTGGACCGGCGATGCGGTGGCGGGTGAGGCGGCGCGCGTCTTCCCCGT | 8 |
| wt-52500-52620 | 3291 | CGTTGCTGCCGCGGGTGGTTGAGGGCATGCCCCCCCTCCCGGAGGCTGGACTCCTGACCAGCCTGTAGGTGAGGACCGAGTCCGACAGGAGGTCTCCCAAACCCCCATCTCTCGCTAGAG | 8 |
| wt-52620-52740 | 3292 | CCGAGACCAGGCCGAGTCCTGCGTAGAACGATGGGGCGCCCAGGAAGGCGGCAGCGTAGGCCGGATGTGTGCCGACCAGCAGCGCCATCATCTCCCGTTGTTCCAATAGAATAACTTCCC | 4 |
| wt-52740-52860 | 3293 | GGTCTGTGGCCGGGGCTGGATAAGGGGGGTGATTCCTAGAGGCGATGAGACTGGCGTGCGCTAAAAGTGTCATGGCCACAATGGGGTTGTCTGCCAGGTCTTCCATCAGGGCTTTGGGCG | 4 |
| wt-52860-52980 | 3294 | CAGAGACGTATTCCCGAAGCAGCTCCCCGGCGTTGGACTCCACGTCGGGCCAGGTGTCCCAGTAGGAGTCGGCGGCGGCGGCGCTGAGGCGGGCGGAAGCTACACTGGCCAGGGTTCTTC | 8 |
| wt-52980-53100 | 3295 | TCCTCCTCTCTTGGTCATCCTGCGGGGGACCAATAGCTTGGGGGCGTCCGGCTGGGGTCAGGGAAAAGGCCTCTGGGTTCTCCAGCACGGTGGGCATGACATATTCCAGAAAGTTGTGGT | 4 |
| wt-53100-53220 | 3296 | AGACGGGGATGTAGTTGAGCGGCTCCTGGGTGTCTGCGGAGACGTAGGCCGGGTTAAGGGGGTCGCAGGGAGACTCTGTTTCCAGCCAGAGGGTGCCGGCGTATTTCGCCGGCCCTGCCG | 8 |
| wt-53220-53340 | 3297 | CCGCCAGAAATTGTGCCCGCCGGGTCGGGGCTCCATTGCCCCATCCAGTTGGTGGTGCCGAAATCGTGATGAGGAGGGGCAGGTTGTTGGTCAAGGGATGCTTAACGAAAACGGTAGGCT | 4 |
| wt-53460-53580 | 3299 | CCACCACGATCTTGAGCTGGCGCGGGTCGGTGTGGCCCCTGGCCTGGGTGACCAGATCCTGCAGCGTTCCCTGCAGCTGGGACTCTTCCTGGGCCTCCTGGATGATGGCCTCCAGTCGGG | 8 |
| wt-53820-53940 | 3302 | CGCAGATGCCCCGGATCCACTTGGCGGCTGCCAGGGATTTCCGGTAGGGCCATGAGCCGTTCCCGGCCGCTGCCCGGGCCAGGGCGGCCTCGAGGGGAGCCTGGACAGGGGCTTTGGGCG | 8 |
| wt-53940-54060 | 3303 | GGGAGGGAAGCAGGCTCCGGAGTTCATCGTCGGGGGCTTCGTCGCGTGACCTGGAGAGGACGGCCTCCAGAGCCGTGTGAAAGCCCCGCCGAGTGCTTGCCGCCATCTCGTGGGCCTTCG | 8 |
| wt-54060-54180 | 3304 | CCATCAGGGTCTGGCTCTCCCGGACCTGCTCTTCCAGCGCCCGGACCTCGGCCGCCTCGGCCTCGGTCAGCAGCTCCGAGAAGAAGTCCCCCGTGGCCTGGAGGAGATCGTCCCGCTCTC | 8 |
| wt-54180-54300 | 3305 | GCCTTGTCAGCAGCTGGGGCTTCTTAGGCCAGAGCGCCGAGTCCGAGGCCAGCCTGGGCGGGGCGGTTGCCTGGGGGATAGTTGGAGGAGGAGGCAGGTTAGCCTGGCCTGGGTCATTAG | 8 |
| wt-54300-54420 | 3306 | TGGCTTCGGGTAGCGTCCGATCCACGTACTCGCTCACGATGGCCGTCAGGGCAGCCTCGGCTGCTCGTCTTTTTTCCAGAAGCCCGGCCAGCCCCCGCTCGTACTCCGCGTAGGGGGCCT | 8 |
| wt-54420-54540 | 3307 | CCAGATCCGTGTTGACCACCGCTGATTTCATGTCCGGGGACTGCAGGGCCTGGCGCGTCTGCGCGAGGGCCGAACGGATGGCATCGGCCGCCGTCCTGGCGCGAAAGAGGGCCCCGGCCG | 8 |
| wt-54540-54660 | 3308 | CTTCCTCCGCTCCTCGCCCTCCTCCTCCTTCTTTGGCGGTAGCCGCGGGGGTGGCGGGCCAAGCGTCCAGTCTGGCCAGAGGGCCGGTCTCGATATCCGTGAACCAGCCGGGTTCCGCGG | 8 |
| wt-54660-54780 | 3309 | CCTCCATTCTCTCCGCCGCACCACCATCGTCCACGAGCAGGGATCGCAGTCTCTCCCTCCTCACCCTCGTTATTCCCAATAGCATAGCGGCAAGGATCTGTGTGAGGGAGTCCAAGATGT | 4 |
| wt-54780-54900 | 3310 | CCGTGTTTCTGGCTACTGCCGCCGCTGCTGCCGCGGCTGAGTCCGTATTGTCTGGCAGCAGGGAGGCCAGCAGGGTGTTCCAGTCATCGGGCGAAGTGGGAGCGGGCTCTGGGCGTGCCC | 8 |
| wt-54900-55020 | 3311 | CCAGCGCCTTCCTAATTCTGGCCCAGGCCTCATTCGCCTCTCGCGCTCGCCGCTCCTGCCTCTCCTTGTCTTCCTGTTCTCGGAGCTTCTCCTTTTCCTTGCGCCCGGTCTCCATAAGCT | 4 |
| wt-55020-55140 | 3312 | GCCGCAGCTTCTTCTCATACTGTCGCTTGAGCTCTTTGTTGGGGGCAGTGTCCAGAAAGGCCTCGAGCTGTTCCTCGGTGGCGGGCTTAAAGCCTTCGGCCTCCAGGCGCCAGGCCTGCA | 4 |
| wt-55140-55260 | 3313 | CCTCCTTCTGTCTGAGCTGATCGTTGTTGTTATTCTTCAATTTCTGCAGGTAACTTAGGAAGCGTTTCTTGAGCTTCCCTGGGATGAGCGTTTGGGAGAGCTGATTCTGCAGCCCAGAGA | 4 |
| wt-55260-55380 | 3314 | GTAGTCTCAGGGCATCCTCTGGAGCCTGACCTGTGATCGTCGCATCATAGACCGCCAGTAGACCTGGGAGCAGATTCACCGCCGCGGCCGTCTCCTTTAAGGTGCTGTGAGTAGCAAAAT | 4 |
| wt-55380-55500 | 3315 | TCTGCAAGGCCACTAGGCGCGCTGGCTCCAGCGTCAGCCGGTTGCCCATCTCGAATGTGTGCAGGGCCTCTGAGACCATGGGGTCCAGGATGCGGTCAATGCCATCCTGCACCTCAGGGT | 4 |
| wt-55500-55620 | 3316 | CAAGGACCGGCAAGTCACGATAGAGGTGGTCTATGCTCTCCTCGAAGGAGGCAATGTAGTTATCGATGGTGTAGAAGGTGATGGATTTCAGGATGTTCATCAGGTACTTTTTGGAGCGAA | 4 |
| wt-55620-55740 | 3317 | CAATCTGCTGTATAGTGTCACGTAGGCGGATGTACGTGGGGTTCTTTGCGGCCCCGACTATCGACCCTGCATTTGCGATGTACTTTTCTATGACGGGGATGGTGAGGGCCGCGGTGTCGG | 4 |
| wt-55740-55860 | 3318 | CCAGCGGTGGCGTGGCTTCGGGGTTGTCGTGGTTGGCGGGTGTCGCAGAGGGAGAGGCGGGAGAGATGGGGGCGCCTGGGGCCGAGGCCACACCGGCCAGGCCCAACATTGCCTCGATGT | 8 |
| wt-55860-55980 | 3319 | CGTCCAGGATGGTGCGGAGGCGCTTTTCGTTTTCTCTGGTGGTCTCGAGCTCCTTCTGTTTTTTCGCGACTGTCTCAAACTCTGGAAGGGGGGCAATGCTGGGGTCGTCCTCCTCAACTC | 4 |
| wt-55980-56100 | 3320 | GCTCCAGGGGCCAGGGGATACCGCTCATATCACTAAGGGCGGTGCCCAGGTAGAGGAGCTCGCGATAGTCCCATTCAATGGACGTGTACCGGATGTTTAGGAGAGGCAGGGAGGCGATGA | 4 |
| wt-56100-56220 | 3321 | TCTGGCATGTGTGCCGCAGGTGTGTCAGGAGGTCGTCAAAATCCATCACTGTTGGGAGGCTTGGGTCCTCAAGGTAGGAGAGATAATCGGAGGCCGCCGAGGCCACCTTGTCCCTGATGT | 4 |
| wt-56220-56340 | 3322 | CCGCCGTACACCTGCGCACGTGCAGGGCCGCATTCTTGGACCGGACGGCCACGTTGTGGACAAAGGGGGGCACTGAGGCGGCGGGAGGGGCCCCATACTCTATCGCTGTCAACAGCGCCA | 8 |
| wt-56460-56580 | 3324 | CGTGCGGGCCCCTGTAGATGGAATTATCCTCGGGTTCTCCTCCTCCTTCCCCCGCCTCCTGATCTCCGCCCGAGAGCAGGTCGGTCAGGCGTCTGCGGGCCGCCTCCAGGTCAAATTTTC | 4 |
| wt-56700-56820 | 3326 | TATGGAACTCTCCCGCTCTTCCCTTGTTTTCTGATAGCATGTTTGCGAGGTTTTGGATGTTAAGTTCGGAAGTGAGCAGTTGCTCCAGGTCCAGCGTGGGGACCTGCAGATGTCCCGACC | 4 |
| wt-56820-56940 | 3327 | AGTCCTTTAAGAATTCCAGCAGATTTAGCACAGACGATCGGTCCCTACTCCTTATTAGCCCCTGCTCGAGGACCACTGTCACAAGAAGATAGTCTATCATGCTCAAGGCATCTGCCTCTG | 4 |
| wt-56940-57060 | 3328 | GCACTTCCCGGTTAGAGGCCGGGTCGTAGACGATGGCCTGTTCCTGGTAGGTATGTCCGGCTATTCTCGCAATGTTGCTCTCGAGGGGCACAAAGTCCATCTCAGGAGTCTCTATGTCAA | 4 |
| wt-57060-57180 | 3329 | AGGTGGTCTGATAGTATTGGCTCCTGGCGGTGTCCAGTGTGATGGGGGACGTGGGGGCACTGGATCCCGATTCCAGGCTGTTGGAGAACACTTCATCTTCGAACATGTCTTCATCCTCTG | 4 |
| wt-57180-57300 | 3330 | TGGTGGGGATATCGGAGGCTAAGTCGCTCTCCGCTTCTTCAGAGTCGGACATGGATAGGAAAGGCTCCTCTAGGTCAGACAGGTAGCGGACGAGGCCAGAACCCCCAGATGCATCATCCC | 4 |
| wt-57300-57420 | 3331 | CAAAGGAGGGCTGCTGCCCGAAGGGAGGTGATGGGGATATCTCCGTTCCAGCCCTGTCAGCGGCCGGGGGATGGTTTTTTTCTGGTTCGAGTGTCGTGGCTGATGGTGGGAGCTGCTGAG | 4 |
| wt-57420-57540 | 3332 | CAGGAGGAGGAGCCGGGGTAGCTGATGGCAGGGGCTGCTGCTGAGGAGGAAGAGGAGAAGGAGCCCGGGCGGCTGATGGCGGGGGCTGCTGCTGAGGAAGAAGTGGAGAAGGAGCCGGGG | 8 |
| wt-57540-57660 | 3333 | CGGCTGATGGCGTGGGCTGCTGCTGAGGAAGAAGTGGAGAAGGAGCCGGGGCGGCTGATTGCGGGGGCTGCTGCTGAGTTGGAGGAGGAGAAAGAGTCGTGGTGGTGGGGGCTGCTGCTG | 8 |
| wt-57660-57780 | 3334 | CAGTCGGGGAAGGGGATGGGGTGGTCAGAGGGATTTTTGGGTTCGAGGGAGCTGCCTGTGGCAGAGGGATGGGTATTTGCAAAGGGAGGCGAGGAGATGGAGTGACTGAAGGAGCGATAG | 4 |
| wt-57780-57900 | 3335 | TTGAGACTGGCGCGGGGTGGGGTGTCGGGGAGGCGGGTGGTGATTGGTGAGGGATGGGGATTACTGGAGGGGGAAGGCGAGCTGCTGAAGGGGGGCGATGGGGCGGAACGTGGGTGCGTG | 8 |

| | | | |
|---|---|---|---|
| wt-57900-58020 | 3336 | GCAGCTGATCATCCTCTGTGTCAGTGGTGGAGGACAGAGGGAGGCGGCGGCCGGAGGTGGGCTTCTTGTGGGGGCTATCTTTGCCCAATCCCTTTTTCCTCTTGGGAGTCTGAGGCGCTG | 4 |
| wt-58020-58140 | 3337 | CGCCGCTCGACGCCCTTGGTGGCGTGGAGGGAGCGGGGACCCCGGGGGTGTGACCTAGGCCGGGGATGGGGATGAAGAGGGGAGGGCTGGAGGCCGGGGCCGCGGAGGCCGGGGCCGCGG | 12 |
| wt-58140-58260 | 3338 | AGGCCGGGGCCGCGGAGGCCGGGGCCGCGGAGGCCGGGGCCGCGGAGGCCGGGGCCGCAGAGGCCGGGGCCGCAGAGGCCGGGGCCGCAGAGGCCGGAGACGACGGCGGGGAGTTGGTCT | 12 |
| wt-58260-58380 | 3339 | TTGCAGGACTATACCTGGCGGCAGGGAATGAGTCGGATGTGAAAGATCGAGAGGGCAGTGGCCTGAGGTTATACGGTATTATTCGCCGTTCAAACGGTAGCATGACGGGAGGGCTGCTAT | 4 |
| wt-58380-58500 | 3340 | CAGCACCGGGCGTCCCCGCCGCCTCCCCATCACTGGACACAAGCTCGGGCCCCACCAGGTCAAAGCCGCTGCCGTTGGCCTCATAAAAGTCATACACGCCATAGTGTTCCAGCATAAAGA | 4 |
| wt-58500-58620 | 3341 | TGCGGGGGTCCTCTGTCTCAAAGGCCTCGGGTAGAAAATAGAGATGCACGCAAGTGTACTGGGCCCCTGGTGCCCCCACGTACTGCAGGATGTCGTGCGCATAGGTGCTGACTCTGACAT | 4 |
| wt-58620-58740 | 3342 | GGGCGGGGGTGCCCGGGGCCGCATCCTTCTGGCAGTGGGGGTCAAACAAGTAGAAGGAGCCATCTGTCTCGATGATGATGGCCCCCGCGTAGATGTCGCAGATGTAGAGGATGAACTGGG | 4 |
| wt-58740-58860 | 3343 | CCACCCCGTTGTAACTGCCGTGCAGGACCTCGGCCAGGGACTGAACAACTGCCGAGTTTGCGATCTGGGCAGGGAATAGGACGAGGCCAAAGATCTCCGCCGAGCGGTATATGTGCACGC | 4 |
| wt-58860-58980 | 3344 | GCCCACCGCCCCTCAGGACCACGGAGCTGGGCACGTCCGTCAACTGGGCCATCTCGTGCCCCTTGAGGATGCCGCTCTGGCGCATGAGGGCATCCAGCCGCGCCCCCTCGTCCAGGACCT | 8 |
| wt-58980-59100 | 3345 | CGTCCAGCTCAGGGCGGGAGGTCAGGGGGCGGCCGGCCAGGAAGCTCTTGACCAGGTAGAGGACGCAGTTGCTGACGCACTGGATGCCGGCAAAGCGGCCAAACTTGCAGTGGGCCTGGT | 8 |
| wt-59100-59220 | 3346 | TGCACGAGGCCGTGCCTAGGATGCGGAGGGCCGAGCCTCCACTCCCGCCCCCGGGGGCATTCACATCCATGGTCCTGATTCCGCGCACGGGGCCGGTTCCCCGGGTGCGCTGGCTTTGCC | 8 |
| wt-59220-59340 | 3347 | CCCAGTCGCCGTTACTCATCTTCGGCGGTGGGGCGGGGAGGACGCCTTGTCGCCCCCCTTCTGGTCCGGGGTCTTACGCGGCTGGCGGCGGCAGCCGCCGAGAGATAAGGGGGGTACGTG | 8 |
| wt-59340-59460 | 3348 | TGTGCCTCCGCCTCTCCTCTGTCTGGGCCGCCGCCGCCGCTTGCCCGCCTTGAAGGAGAGGGGGTAGTCCGCGGACTGCGTCTGCGGGGGCAGGAGGTCTCAACCTTCTGGGCTCGGGCC | 8 |
| wt-59460-59580 | 3349 | GCGGTGTCGATATCCGATGGCCTTTCCCTGTCTTCCTCGTATGCTCCTTCTCCTCCTCCTCCCGGCACGCCCCTGAGATCTGCCTCCCCTCCCTCTCCCTCGTCCTGGTCGGAAAAGTCT | 4 |
| wt-59580-59700 | 3350 | GAGGAGGAGAAGGAGAATGGGGAGGAGTCCAAAACGGCACGCCACCTGCCGTGGGGCGGTGGTGACAGGTCCCGGCTGGCCCGGCGCTTGCTCGCGTTCCTGCCGTTACCCAGGAGAATG | 8 |
| wt-59700-59820 | 3351 | GCCGCGAGTTTTTTGGCGGGGAGGATGCGGAATGGCGGGGGCGTTTGTCCCACGGGTGAGGGGGAATCGTCGGTTAGGGCCGGCACGAGGTGGTGGGTCTGGACCCGGGCCGTGCGAGCA | 8 |
| wt-59820-59940 | 3352 | AAGGCGGCGAGAACCGAGGGGCTTCTGGGGGTGACTGTGATCTGTTCCGGATTTAGGTCCATGGCGGGTGTGTATGTTTTAATAGGGGTGGTCTCTGGCGCGGCAGGATGATGGTCGAGG | 4 |
| wt-59940-60060 | 3353 | ACGTCCACCAGGGCCTTGCAGATGCTCTTGCCTAGATACAGGATGTCGTCCATGCTGAGGGGAGGTGGGGTGTCTGCTCCCCCCTGCGGAAGCCGCCTGGGTGCGGGGATGAAGACAGGT | 4 |
| wt-60060-60180 | 3354 | GGTGGGCGGGCGTCTCGCCGGACTATGGCCTCGGCACGCTCGGCGTCGATGGCGGGTGGCTGGAACAGGCGGGCGAATGTGTAATCCCGGAACCGGTAGGCGACGCTGCGCCTGAGGGCG | 8 |
| wt-60180-60300 | 3355 | CCCGTCAGGCTGTATCCCAGCTCCAGGGCGTGCTCCACCCGCTCGTTGAGCTCCTCGAGATCCGGACGCAGGGGCTCGCTGGTGTGGGCCCAGAGGGGGTGATCCGCGATGCCCCGGCTC | 8 |
| wt-60300-60420 | 3356 | TCCCTGAGGGCCGGCACCAGGAGGCGCCTTCTGAGGGTGGCCGTGTCGGCCGTGGCCAGGGCCCACCTGGCGGCGGCGTCCCGGCACACATCCTGGATGCCCTCCACGACGCTCTTTAGC | 8 |
| wt-60420-60540 | 3357 | GTCTGGAGGTCCGTGGAGTAGTGGCGGGGGGAGGATGAAACGCTCTTTTCCTTCACCGCTACCACCGCCTCCTCCTCCTCTTCCGTCGCCAGAGGGATCTGCACCCTCCCGGTCTCTGCG | 4 |
| wt-60540-60660 | 3358 | TCGTACAGGAGCGGGCGGGAGCACAGCCTCCAAGCTGCCCCCGTCAAGCGCGAGATGTCCTCCGAGAGGGTCTCACCCGAGACCAGAAAGCGGCGGGTGGCCAGGCCCAACGACTCCGCC | 8 |
| wt-60660-60780 | 3359 | GTCGTGCTGTATCTCAGGGTGAAGAGGAGTGAAAAGAGGGAGGTGGGCCAGGCAAGCGGTGGTGCTTCCGCCGCCCGCTCTGAAGCTGAGATAGTCTCGGAGATGATGCCTGAGACCTCT | 4 |
| wt-60780-60900 | 3360 | CGGACGGCGTCCATGATCCTAAGGACTGCGTCGTGGGACGACAGCCCCCAGGGGCCCCCGCCCTCTTCGTCTTCTGCACCCTCGGCTCCTGCGTCCCCGGCCTTGCCTTCCCCCTCTAAG | 8 |
| wt-60900-61020 | 3361 | TTGAGGGGGCGCAGTCCGACCGCCTGGGGGGACTCCCCAGGCATCGGAGGGGCCCCGTCATAGATCTCCCAGACGGTGGCGTATATGAGCTCGAGAGGACGGCGGGCCCGGGTCAGCTCG | 8 |
| wt-61020-61140 | 3362 | GGGGAAGGGAGGGCCAGGTCGCTGCCGAAGGAGACCAGCCAGCGCAGGGCGGCCAGAGAGCGGGTTTTGGGCAGCTCGTTGGAGAGGACCCGGCGAAGGGCGGGCCAGATTTGGAACTCG | 8 |
| wt-61140-61260 | 3363 | ATGAAGGCGGCCGGGAAGAAGGGGCTGTGGACATAGGCCGGATCCGCGCGCGCCGTTTGGCCGGCCCTCAGGGACCGGCAGTATGCCTCGACGTCTGTCCGCGGGGCCGCCGCCACCGCT | 8 |
| wt-61260-61380 | 3364 | GCCGTCCACTGCCTTCTTCCCTGCTCGCCGGGGAGTAGGGGGGGCTTACAGGGGAGGGCCGGAGCCGGGGCCGGGGCCTGCCACAGGCGGCTGTAGCGGACCCATAGCAGAGACCTGAGG | 8 |
| wt-61380-61500 | 3365 | AGTTCGGATGAAAGGTCCCCCGCCACCTGCTCATACTCGGCCGCGGGAGGGGGGACGATGAAGATGCGCAGAGGGGTTACGGCGTCCCAAGGGTCCGCCGCCGCCCCCACACCCACAGCC | 8 |
| wt-61500-61620 | 3366 | GTCGCGGCGGGGGCGGCGGCGGGCGTAGAGGGGCCGCTGGTGCGCCGGGCTCGTCTGTCCACGGCCTCGGCCTCCGCCCGCAGGTAGGCCGCCCGGGCCACACGGGCGAAGCGGCTCGTG | 12 |
| wt-61620-61740 | 3367 | GGGCTCGCGGTGGGCAGCAGTCGGAAAAAGTGCAGGGCAAAGCCCGATAGACTCTCTAGGAGGGCGGCGGTGGCCTCGAGCCACCTCCACCGCGAGCGGGACACCCGGGGCACAGAGGCC | 8 |
| wt-61740-61860 | 3368 | AGCATCATGGCGTAGTCCCCCGCCACGGTCTCGTTGAGCCCGGCCGAGAGCAGAACCGTGGCCACCTGCTCGATGGCGGCTGGAGAGAAGGATGCCCGGCTCCCCGCCGCCTCCTGCACA | 8 |
| wt-61860-61980 | 3369 | CGAGCGGCCAGGGCCTCCATCTCTGCCGCCATCCCGGCCAGGAAGGCCTCGATGACCGAGTCTGGGACGCCGTAAGTCTGGTCCCAGAGCAGGGCCTCGTACACATAGTCGTAAAAGAGG | 4 |
| wt-61980-62100 | 3370 | GCCCCTGAGGGCTCCAAAAGCCGGAGCCGGGCGGCGTCAAAGGCCAGGACGGGCACAGCCGCGACGGGGGGCGTTTGTCCCCCGCTGGCCTCCGCGTACACGCCCAGGATCTCTACCGCC | 8 |
| wt-62100-62220 | 3371 | CGCCGCCGGGCCAGGGGCAGCGAGGCCACCACGCTGGAAAGTGACTCGGGGCGGTGAAAGAGACCACCACCGCTTTCTTCACCCTCTCCCCCGCCGGCCCCGCCCCCACTGTGCTCCACC | 8 |
| wt-62220-62340 | 3372 | AGCTCCACGGCCATGGCCTTGATGTCCGCGGCCGTGGGCTGACCCTGCCCTGCAGCCGCCCAGGGGTAGCGGTTGGTCTCCGCGTATACGGTGACCAGCCATCTCCCCAGCGTCGTTTTC | 8 |
| wt-62460-62580 | 3374 | AGCACCTCCCCGCTGCCCTCTTCGATGATGCGCCGGTACGCGGCCGCCAGGGCCCGGGTCTCTGCGATGAGCCGAGAGCCGTCCAGCGGATCGTCGGTGGCCGGAGAGGCTGTCGTGGGG | 8 |
| wt-62580-62700 | 3375 | GGCAGTGAGGATGCCAGCACGTCCAGGGCCGCCTCCAGATGGCCGAGGCCGAAGCTGCGCCTGGAAAAGGAGGCCGCCCGGAGTAGGTAGTAGGCGTGGTGGCGGAGGACCGCCGCCGGG | 8 |
| wt-62700-62820 | 3376 | TAAGCGTGGCCGCTCATGAGGGTGAGAGTATTTAAAAAATCGCGCACCAGCACCGGCTGGGCCAAATCCCCCAGTCCAAAGATCCCCAGCTCCAGAGGCATCAGCGCGCGCAGGCGGGCA | 4 |
| wt-62820-62940 | 3377 | GCGGGGTCGTCCCCAGACAGCAGCAACTGACGCGTCACGCGGGCGAGCCCCCCGTCCACCTCTGCCAGGGGTGGCTGGGCGTCTGCCCCTCCGCTACCGCCGCTGCTGTCACTCTCCATA | 8 |
| wt-62940-63060 | 3378 | GCGGACGCCATGAAGGTCCAGGGGTCCGTCGATCGCCGCCGTCTGCAACGCCGAATCGCGGGGCTGCTGCCCCCTCCGGCCCGGCGTCTAAATATTTCCCGGGGGTCCGAATTCACGCGG | 8 |
| wt-63060-63180 | 3379 | GACGTTCGTGGGCTGGTTGAGGAACACGCGCAGGCCTCCTCGCTGAGTGCGGCGGCCGTCTGGCGCGCAGGGCTGCTGGCCCCGGGGGAGGTGGCGGTCGCCGGGGGTGGCAGTGGAGGG | 12 |
| wt-63180-63300 | 3380 | GGGAGCTTCAGCTGGTCTGGGTGGCGGCCGCCAGTCTTTGGGGACTTTCTGATACACGCCAGCTCCTTCAACAACGCCGAGGCCACTGGAACGCCCCTTTTCCAATTCAAGCAGAGTGAC | 4 |
| wt-63420-63540 | 3382 | CTGTACGACAGCCCCGCAACCCTGGCTGACCTGGTCCCGGACTTTGGGCGGCTGGTGGCCGACCGCCGCTTCCACAACTTCATCACCCCTGTGGGCCCCCTGGTGGAGAATATAAAGAGC | 4 |
| wt-63540-63660 | 3383 | ACCTATCTGAATAAAATCACCACGGTGGTCCACGGGCCTGTGGTCAGCAAGGCCATCCCTCGCAGCACCGTCAAGGTGACGGTGCCCCAGGAGGCCTTTGTGGATCTGGACGCGTGGCTC | 4 |
| wt-63660-63780 | 3384 | TCCGGCGGCGCCGGGGGTGGCGGTGGAGTATGCTTCGTCGGGGGGCTGGGCCTGCAGCCGTGCCCCGCCGATGCGCGCCTCTATGTCGCTCTGACCTATGAGGAAGCCGGGCCGCGGTTT | 8 |
| wt-63780-63900 | 3385 | ACGTTTTTCCAGTCGTCCCGCGGCCACTGTCAGATCATGAATATCTTAAGAATTTATTACTCACCATCCATCATGCACCGCTATGCTGTGGTCCAGCCCCTACATATAGAGGAGCTAACC | 4 |
| wt-63900-64020 | 3386 | TTCGGGGCGGTTGCCTGTCTGGGGACATTTAGTGCTACTGACGGTTGGAGGAGGTCTGCCTTCAATTACCGTGGCTCTAGCCTCCCCGTGGTGGAGATTGACAGCTTTTATTCCAACGTC | 4 |
| wt-64020-64140 | 3387 | TCTGACTGGGAGGTGATTCTCTAGACTTAACGGGAGGAAACAGGAGGAGGAGGGGGACAAGAGCACAAAAGTGGTTCAGTGGACACCCACCACACAGCATGGCAACGACCAGTCATGTCG | 4 |
| wt-64140-64260 | 3388 | AGCATGAGCTCCTCTCCAAATTGATTGATGAGTTAAAGGTCAAGGCCAACTCAGACCCCGAGGCTGATGTCCTGGCCGGGCGCCTGCTCCACCGCCTTAAGGCCGAGTCAGTTACACACA | 4 |

| | | | | |
|---|---|---|---|---|
| wt-64260-64380 | 3389 | CAGTAGCCGAATATCTGGAGGTCTTCTCTGACAAATTCTACGATGAGGAATTCTTCCAGATGCACCGGGATGAGCTGGAGACCCGAGTCTCTGCTTTCGCGCAGAGCCCGGCCTACGAGC | | 4 |
| wt-64380-64500 | 3390 | GCATCGTCTCCAGCGGCTACCTGTCGGCCCTGCGCTACTATGACACCTATCTGTATGTGGGGCGCAGCGGGAAGCAGGAGAGTGTGCAGCACTTTTACATGCGGTTAGCCGGCTTCTGTG | | 4 |
| wt-64500-64620 | 3391 | CCTCAACCACCTGCCTCTACGCGGGTCTCAGGGCAGCCCTGCAGCGGGCCAGGCCGGAGATTGAGAGTGACATGGAGGTGTTTGATTACTACTTTGAGCACCTAACCTCCCAGACGGTGT | | 4 |
| wt-64620-64740 | 3392 | GCTGCTCCACGCCCTTTATGCGCTTTGCCGGGGTGGAAAACTCCACTCTGGCCAGCTGCATCCTCACCACCCCCGACCTCAGCTCCGAGTGGGACGTGACCCAGGCCCTCTATAGGCACC | | 4 |
| wt-64740-64860 | 3393 | TGGGGCGCTACCTCTTTCAGCGAGCCGGGGTGGGTGTAGGGGTGACGGGGGCTGGCCAGGATGGGAAACACATCAGCCTCCTGATGAGGATGATCAACAGCCACGTGGAGTACCACAACT | | 4 |
| wt-64860-64980 | 3394 | ATGGCTGCAAGAGGCCGGTCAGTGTGGCGGCCTACATGGAGCCCTGGCACAGCCAGATTTTCAAGTTTTTGGAAACGAAGCTGCCGGAGAACCACGAGAGGTGCCCGGGCATCTTTACGG | | 4 |
| wt-64980-65100 | 3395 | GGCTCTTTGTCCCCGAGCTCTTCTTCAAGCTTTTTAGGGACACGCCCTGGTCGGACTGGTACCTGTTTGACCCCAAGGACGCCGGGGACCTGGAGAGGCTCTACGGGGAGGAGTTTGAGC | | 4 |
| wt-65100-65220 | 3396 | GCGAGTACTATCGGCTGGTGACAGCGGGCAAGTTTTGTGGGCGGGTCTCCATCAAGTCCCTGATGTTCTCTATCGTCAACTGCGCCGTCAAGGCCGGCAGCCCCTTCATCCTTTTGAAGG | | 4 |
| wt-65220-65340 | 3397 | AGGCCTGCAACGCCCACTTTTGGCGCGACCTGCAGGGCGAGGCCATGAACGCCGCCAACCTGTGCGCCGAGGTGCTGCAGCCCTCGAGGAAGTCTGTGGCCACCTGCAATCTGGCCAACA | | 8 |
| wt-65340-65460 | 3398 | TCTGCCTCCCGCGCTGCCTGGTGAATGCGCCTCTGGCGGTGCGGGCACAGCGGGCCGACACGCAGGGGGATGAACTCCTGCTGGCCCTCCCTCGACTCTCAGTCACCCTACCTGGAGAGG | | 8 |
| wt-65460-65580 | 3399 | GGGCAGTCGGTGATGGATTCTCGCTAGCCCGCCTCAGAGATGCCACCCAGTGTGCCACCTTTGTGGTGGCCTGCTCCATTCTTCAGGGATCCCCCACTTATGATTCCAGGGATATGGCCT | | 4 |
| wt-65580-65700 | 3400 | CCATGGGCCTCGGGGTGCAGGGCCTGGCCGATGTCTTTGCGGACCTGGGCTGGCAGTACACTGACCCTCCCTCTCGCTCGTTAAACAAGGAAATATTCGAACATATGTACTTTACGGCCC | | 4 |
| wt-65700-65820 | 3401 | TCTGCACCAGTAGTCTGATTGGACTTCACACCAGGAAGATTTTTCCGGGTTTCAAACAGAGCAAGTATGCCGGGGGGTGGTTTCACTGGCACGATTGGGCAGGAACAGACCTTTCTATTC | | 4 |
| wt-65820-65940 | 3402 | CCAGGGAAATTTGGTCTCGCCTCTCTGAACGCATTGTGAGGGATGGGCTTTTCAATTCACAGTTTATCGCCCTGATGCCCACCTCAGGCTGTGCCCAGGTGACGGGCTGTTCGGACGCCT | | 4 |
| wt-65940-66060 | 3403 | TCTACCCCTTCTATGCCAATGCGTCCACCAAGGTCACCAACAAGGAGGAGGCCCTTAGGCCAAACCGGTCTTTTTGGCGTCATGTGCGTCTGGATGACAGGGAAGCTTTGAATCTTGTCG | | 4 |
| wt-66060-66180 | 3404 | GGGGCCGTGTCTCCTGCCTCCCGGAGGCTCTGCGGCAGCGCTACCTGCGTTTCCAAACGGCCTTTGATTACAACCAGGAGGACCTGATTCAGATGTCCCGGGACAGGGCCCCCTTTGTGG | | 4 |
| wt-66180-66300 | 3405 | ACCAGAGCCAATCTCACAGCCTGTTTTTGCGTGAGGAAGATGCCGCGCGGGCCAGCACGCTAGCCAACCTACTGGTGCGCAGCTACGAGCTGGGCCTGAAGACTATCATGTACTATTGTC | | 4 |
| wt-66300-66420 | 3406 | GCATTGAGAAGGCCGCCGATCTGGGGGTGATGGAGTGTAAGGCCAGCGCGGCTCTGTCGGTGCCGCGGGAGGAACAGAATGAGCGGAGTCCCGCTGAGCAGATGCCGCCTCGTCCCATGG | | 8 |
| wt-66420-66540 | 3407 | AACCGGCGCAGGTTGCGGGGCCGGTTGACATCATGAGCAAGGGCCCAGGGGAGGGACCAGGTGGGTGGTGTGTGCCCGGGGGATTGGAAGTGTGCTATAAGTACCGTCAGCTCTTCTCAG | | 4 |
| wt-66540-66660 | 3408 | AGGATGATCTGTTGGAGACTGACGGTTTTACTGAACGAGCCTGTGAATCTTGCCAATAAACGTTTATTGCCATGTCCAAGTTGTTGTACGTGCGTGATCATGAGGGCTTTGCCTGCCTAA | | 4 |
| wt-66660-66780 | 3409 | CGGTCGAAACCCACCGCAACCGCTGGTTCGCGGCTCACATTGTCCTCACCAAGGACTGCGGGTGTCTCAAGCTACTCAATGAGAGGGACTTGGAGTTTTACAAGTTCCTCTTTACGTTCC | | 4 |
| wt-66780-66900 | 3410 | TGGCCATGGCCGAGAAGCTTGTGAACTTTAACATTGATGAACTGGTCACCAGCTTCGAGAGCCACGACATTGATCACTACTACACCGAGCAGAAGGCCATGGAGAACGTCCACGGGGAGA | | 4 |
| wt-66900-67020 | 3411 | CTTATGCTAACATTTTAAACATGCTCTTTGATGGGGACAGGGCGGCGATGAACGCCTACGCAGAGGCCATCATGGCCGACGAGGCCCTGCAAGCCAAGATTTCCTGGCTCCGTGACAAGG | | 4 |
| wt-67020-67140 | 3412 | TGGCGGCCGCCGTCACCCTGCCGGAGAAGATTCTTGTGTTCCTGCTGATTGAAGGCATCTTCTTCATTAGCTCCTTCTACAGCATAGCCCTGCTGCGGGTCCGGGGCCTAATGCCTGGCA | | 4 |
| wt-67140-67260 | 3413 | TCTGCCTGGCCAATAACTACATAAGTAGGGATGAGCTGCTCCACACCCGCGCTGCCTCCCTGTTATACAATAGCATGACAGCCAAGGCTGACCGACCAAGGGCCACCTGGATCCAGGAGC | | 4 |
| wt-67260-67380 | 3414 | TGTTTCGCACTGCGGTGGAGGTAGAGACTGCCTTCATCGAGGCTCGTGGAGAGGGGGTTACCTTGGTGGATGTGCGAGCCATAAAGCAGTTTCTGGAGGCCACGGCCGATCGCATCCTGG | | 4 |
| wt-67380-67500 | 3415 | GTGACATTGGTCAGGCTCCCTTGTATGGCACACCACCCCCCAAGGACTGCCCGCTCACCTACATGACTAGCATCAAGCAAACTAATTTCTTTGAGCAAGAGAGTTCCGATTACACCATGC | | 4 |
| wt-67500-67620 | 3416 | TGGTGGTAGATGACCTTTGAGTCAGGGTGGCTACTTGCTCAGGTTTCTGGGCATAAATTCTCCTGCCTGCCTCTGCTCTGGTACGTTGGCTTCTGCTGCTGCTTGTGATCATGGAAACCA | | 4 |
| wt-67620-67740 | 3417 | CTCAGACTCTCCGCTTTAAGACCAAGGCCCTAGCCGTCCTGTCCAAGTGCTATGACCATGCCCAGACTCATCTCAAGGGAGGAGTGCTGCAGGTAAACCTTCTGTCTGTAAACTATGGAG | | 4 |
| wt-67740-67860 | 3418 | GCCCCCGGCTGGCCGCCGTGGCCAACGCAGGCACGGCCGGGCTAATCAGCTTCGAGGTCTCCCCTGACGCTGTGGCCGAGTGGCAGAATCACCAGAGCCCAGAGGAGGCCCCGGCCGCCG | | 8 |
| wt-67860-67980 | 3419 | TGTCATTTAGAAACCTTGCCTACGGGCGCACCTGTGTCCTGGGCAAGGAGCTGTTTGGCTCGGCTGTGGAGCAGGCTTCCCTGCAATTTTACAAGCGGCCACAAGGGGGTTCCCGGCCTG | | 4 |
| wt-67980-68100 | 3420 | AATTTGTTAAGCTCACTATGGAATATGATGATAAGGTGTCCAAGAGCCACCACACCTGCGCCCTGATGCCCTATATGCCCCCGGCCAGCGACAGGCTGAGGAACGAGCAGATGATTGGGC | | 4 |
| wt-68100-68220 | 3421 | AGGTGCTGTTGATGCCCAAGACGGCTTCCTCGTTGCAGAAGTGGGCACGCCAGCAAGGCTCAGGCGGCGTTAAGGTGACACTCAATCCGGATCTCTACGTCACCACGTATACTTCTGGGG | | 4 |
| wt-68220-68340 | 3422 | AGGCCTGCCTCACCCTAGACTACAAGCCTCTGAGTGTGGGGCCATACGAGGCCTTCACTGGCCCTGTGGCCAAGGCTCAGGACGTGGGGGCCGTTGAGGCCCACGTTGTCTGCTCGGTAG | | 4 |
| wt-68340-68460 | 3423 | CAGCGGACTCGCTGGCGGCGGCGCTTAGCCTCTGCCGCATTCCGGCCGTTAGCGTGCCAATCTTGAGGTTTTACAGGTCTGGCATCATAGCTGTGGTGGCCGGCCTGCTGACGTCAGCGG | | 4 |
| wt-68460-68580 | 3424 | GGGACCTGCCGTTGGATCTTAGTGTTATTTTATTTAACCACGCCTCCGAAGAGGCGGCCGCCAGTACGGCCTCTGAGCCAGAAGATAAAAGTCCCCGGGTGCAACCACTGGGCACAGGAC | | 4 |
| wt-68580-68700 | 3425 | TCCAACAACGCCCCAGACATACGGTCAGTCCATCTCCTTCACCTCCGCCACCTCCTAGGACCCCTACTTGGGAGAGTCCGGCAAGGCCAGAGACACCCTCGCCTGCCATTCCCAGCCACT | | 4 |
| wt-68700-68820 | 3426 | CCAGCAACACCGCACTGGAGAGGCCTCTGGCTGTTCAGCTCGCGAGGAAAAGGACATCGTCGGAGGCCAGGCAGAAGCAGAAGCACCCCAAGAAAGTGAAGCAGGCCTTTAACCCCCTCA | | 4 |
| wt-68820-68940 | 3427 | TTTAACACCATGTTCTCGTGCAAGCAGCACCTGTCCCTGGGGGCCTGTGTCTTCTGTCTCGGCCTCCTGGCCAGCACCCCCTTCATTTGGTGCTTTGTCTTTGCCAACCTGCTCTCTCTG | | 4 |
| wt-68940-69060 | 3428 | GAGATCTTCTCACCGTGGCAGACACACGTGTACAGGCTTGGATTCCCGACGGCATGCCTAATGGCCGTCCTCTGGACGCTGGTACCCGCCAAGCACGCGGTGAGGGCCGTCACTCCAGCC | | 4 |
| wt-69060-69180 | 3429 | ATCATGCTGAATATTGCCAGCGCCTTGATCTTCTTCTCCCTCAGAGTCTACTCGACCAGCACGTGGGTTTCTGCCCCCTGTCTCTTTCTGGCCAACCTGCCTCTCTTATGCCTGTGGCCC | | 4 |
| wt-69180-69300 | 3430 | CGGCTGGCCATCGAGATTGTTTACATCTGCCCGGCTATACACCAAAGGTTCTTTGAACTTGGGTTGCTCTTGGCCTGCACCATCTTTGCCCTGTCCGTGGTCTCCAGGGCCCTGGAGGTG | | 4 |
| wt-69300-69420 | 3431 | TCGGCTGTCTTCATGTCTCCATTTTTCATCTTTCTGGCTTTGGGCTCTGGAAGCCTGGCCGGTGCTCGGCGTAACCAGATTTACACCTCGGGTCTCGAGCGGAGACGCAGCATTTTCTGC | | 4 |
| wt-69420-69540 | 3432 | GCCCGGGGAGATCATTCGGTGGCATCCCTGAAGGAGACCCTCCATAAATGCCCGTGGGATCTGCTGGCCATCTCTGCCTTGACCGTTCTTGTCGTCTGTGTGATGATTGTGTTGCATGTG | | 4 |
| wt-69540-69660 | 3433 | CACGCAGAGGTGTTCTTTGGACTCTCTAGATACCTGCCCCTCTTTCTCTGTGGGGCGATGGCCTCCGGGGGGCTGTACCTGGGCCATTCCAGCATCATTGCATGTGTCATGGCCACCCTC | | 4 |
| wt-69660-69780 | 3434 | TGCACCCTGACATCTGTTGTGGTATATTTCCTCCATGAAACCCTTGGACCCCTGGGCAAGACCGTGCTGTTTATCTCAATCTTTGTCTATTACTTTAGCGGGGTAGCGGCCCTGAGCGCA | | 4 |
| wt-69780-69900 | 3435 | GCTATGCGCTACAAGCTTAAGAAGTTTGTGAACGGACCCCTGGTCCATCTCCGTGTGGTATACATGTGCTGTTTTGTCTTTACTTTTTGTGAATATCTGTTGGTGACATTCATTAAATCC | | 4 |
| wt-69900-70020 | 3436 | TAACGACCGGAGTCCTGTCTCTTTGTGTTCTTGGGGGACTTGAGTTAGCTGTCTTTCCTCTTATTACATTGGGCTAACGGGAGGAAATGAACCCAGGGGTGGCAGTGGATGGGGTCATTT | | 4 |
| wt-70020-70140 | 3437 | ATGGGCAAAACTCACAGGACATGTTTGGGGAGTTAGCATTGGCGTCGGGAAACACAGCTCTGGCAGTTATAACCGCACCAGCTAACAGGACATGTTTGGGGGAGTTGGCATTGGCGTCAG | | 4 |
| wt-70140-70260 | 3438 | GAGACACGGCTCTGTCAGTTATCACCGTACCATGAGTGCCATGTGTGTCCAGTGCCTAATCACCGTTCCTCATTTTGTGTGCCTCCTCAAATGTTCCAGAAGTCGGCCACAGGGGAGGTG | | 4 |
| wt-70260-70380 | 3439 | GCTGAATTAGGGCCTTTTCCCTCATTCCCCCATGAGACCCACGTGGCAGGCCTAGGGGCTACATTCGCCTCCCACGTTTCCCTTCGCGTGAGGCATCCGATATGACTGAATTTTCGCAGT | | 4 |

| | | | |
|---|---|---|---|
| wt-70380-70500 | 3440 | CTCTTTTCCCTCTTCCCTTGTTATTCCCATAGAATTACAGTGAGGTTACACAGGTGGAGATTCAGTTTAACCATTTATTGATTTAATCCAGGAACAAAAAACAGTCCTAGTGACCCAGTG | 4 |
| wt-70500-70620 | 3441 | CCCGGAGAGAGAATGGCCCTGACAAGTCGGCTGCATGATGCACTTCGGCAGTCACGTGTGTGAGTCTCCACGGCCTCTGTCAAAAGGGAGCTTAGCGTGCCAGGGTTGTAATTCTTGATG | 4 |
| wt-70620-70740 | 3442 | TAGTGGCCCAGGAATTCAACTTCATCGTGTCTCCGTCTGCAGTTGGCGTTAATGTAGGCTGGGGCTACTGCCGCATATGCTGCCAAGAGACAGAGGGGCTGCTTCACATATGAGCTGCTC | 4 |
| wt-70740-70860 | 3443 | AGGGTCTCCACCACCTTGTTTTGACGGGCCGTGGCACAGGTGATGTAGAAGAGTTGCTTCACAAAGTTGTAGTCTCGCGTGTTAGGAAGGAAGCAGGGTGCCAGCTCTTTGAGCTTGGTC | 4 |
| wt-70860-70980 | 3444 | AGGATCACCTTGCTAAGACTCATGGCGCAGGCCAGGAGGATGTCTTCCGCGGGAGCTAGGGGCAGGTCGCCGTGGTAGGTGATCTCCTGGAGCCAAAAGATGGTCTCTTCTAGCATGGCC | 4 |
| wt-70980-71100 | 3445 | ACCAGGGTGCAGAGCCCCGCGTTCTGGATCGCCTGCATGCGTGCATCCAGCCATGTGTCCTTGTTGGTTGACTTGGTGAAAAACTCACGTAGTGTCTTGTAGCTCCTGCGCAGCTCGTGT | 4 |
| wt-71100-71220 | 3446 | CTGGGTTGCACTTTCTGCCAGGCTCCAATCTCTGGATGGGCGGCCACCGCCAGCATCGACTGTAGGAACGGGTCTTGGATGGGCTCTAGGGTCAGAGAGGCCAGGGGGCTGGGCAAGGTG | 4 |
| wt-71220-71340 | 3447 | ACAAATGTAATCTTGGAGACAGGCTTAACCAGACTCATGTCAAACCACGGTTTGTTCGGCAGGGGCCTCTGGCTGCGTTCTTGCCTCGCCTGCTTCCTTGTGCTCCTGCCGGCCCCTCGA | 4 |
| wt-71340-71460 | 3448 | GATTCTGACCGGGGACCTCTGGTTGCTCTGTTGCTTCGGGGAGCTCTTGGAGACCTCGGTGCTCTAGGCACCCTGGGGGCCCTTGGGGCTCTGGGCGCTCTTGCTCCCGGGGGCAGGTGT | 8 |
| wt-71460-71580 | 3449 | CGGCGCTTGCCATAACTTTCATCGGTGCAGCCATGGACCTCTCCGCGTCGCCTTTTGTGGCCTCTGGTGTAAGAGGAGTTGCCAGTCTCCTCCTTCTCGTCCTCGTCCCTGCACAGGGGT | 4 |
| wt-71580-71700 | 3450 | GAGCGATGCAATGTGACTGTCTTGTCCTGTAGGTCCCACTTCTTTCTGGGAATCACAAACGATGCCGAGGTAGGGGTTATGACCACGCTGGAGGGCCGTGCAGGTATGGCGTGGGCCGGA | 4 |
| wt-71700-71820 | 3451 | GTTGGATCTTCATCCTCCTCCTCTGAGGATGAAATCTCTCCATCTGTGGAGTGTTCTTCGCTGCCCTCCATAGGGTCCAGATCGCAGTCTGTGTTGGTGTCTGAGACCGCTTCGAGTTCC | 4 |
| wt-71820-71940 | 3452 | AGAATGTGGCTCTCTGCAGAGGGGAGACAAAAGGTGGAGACTGCCTTGAGCACCTCTGTCTCAGGCACCGGATGCCCCCGGCTCCACGGCCCCGGCCACTGGCCGGTGTAGCTTCTTACC | 4 |
| wt-71940-72060 | 3453 | TGCGGGATCCTCGTTGGAGGAAATGCTGCTAGTTCGGGAGAGTCTCTGAGAAGGAACCATCTTGTCTGTCTCTACGACGGGCTAGCTGGGATGTAGTGCTGTCTTGACTGGCCTCAGCCC | 4 |
| wt-72060-72180 | 3454 | TATTTATGATTCTGGAGGCGGGCACGCTGATGGAGAAATGGGCGGTCGGTTGATTGGCCCCACAGCGACCGGCGAAGCACTGACTCATGAAGGTGACCGTGATGGCCTGTGATGTGTAGT | 4 |
| wt-72180-72300 | 3455 | AGAGTACCAGAAACACCCTCACATTCTTGGAGCTGGCCCTGTGGGTATGCCTCAGGCACGCAAAGTTCCTGCCCCGGGCATGGCACACCTGAACTAAGTTTGGCCCGGTTTGCTCAAACG | 4 |
| wt-72300-72420 | 3456 | TGACATGGAGAAACTGGGGGAATTTGTCTTCTGGCACAGCTGTTGCCAGGGTGCTCATGAGCGAGGGCCAGATGCAGGAGCTGACCCAGGCGACGAGATCCAGGCCCAGATGTCCCTCTA | 4 |
| wt-72420-72540 | 3457 | TCATGGCGCAGACATTCTCCACGGTGGGGGGCAGGGTCTCGCGGGTCCTCTGGATTAGATAGTCACGCCCATCATCCGCGATGTGGTAGCAGAAGGTTTTGGGGGCCGGCCAGCCCACGT | 4 |
| wt-72540-72660 | 3458 | GCAGTGAGTGATGTAAGAGGTTTTGAATGTTGAGGGCATTCTTAACATAGCTGTGCTTGTCTTCCTCTTCCGGATGACAGACAAAGAGGCGCAGCTGCCGGCTAAGACCACCGCCCCTGT | 4 |
| wt-72660-72780 | 3459 | CCACCTTGTAGGTATGCGGCAGCCGGATGCACCGCCCGGCGTGATACACGCCGCTGTCAAAAAGCGGGGCCCCAATCTCTTTGATCTTGTGACGCATGCGGCGCAGGCAGGCCGTTAGGC | 4 |
| wt-72780-72900 | 3460 | CCATGAGCTTCTGCAGCACAGACACAAACCCTTGTACTGCGCTTGTTCCCACAATAGCATGGCCTCTAGGTAGGGGGGTGATGACGCGAAAGCCCAGTTTTCCCGTGCATATGCAAAAGG | 4 |
| wt-72900-73020 | 3461 | GGAGCACATCTTCCATATTATCCGGGTCGGCGGGTGGACAAGCTGATTTGAAAAAATAGACTGGGTGGGCCCTGGACACTGGACCCAGGCGGCGCATGAGGCGCAGTACCTCACGCCGCA | 4 |
| wt-73020-73140 | 3462 | CGGTCCGGCACAGGTCATAGATTTCCTCCAGCGACCAGGGGGCCCCCTTGATCTTTAGATCCAGGTCCAAGACCAGGTTGCAGACCGGAAGCCGGGGATTAAAGTATTCATGCCGGGAGA | 4 |
| wt-73140-73260 | 3463 | CAAAGAGCTGCTCGCTCAGGCTGTTCTGTGAATAGTACACTGGGGTGTAGGAGAGGGCCCTGGTGAGACACGTGTCTGGGAGGCGGCAGTTGGTCGGGGTGGAGACGACCTCCGCCAGGT | 4 |
| wt-73260-73380 | 3464 | GGGATGAGAAGGGGTCAGCGGCTGTCATTACAAAGTAGTGCCTGTCTGCAAAATGGCAGAGGAAGACCGGTAGCCGCTGCACCCTTCGAAGGACGGTGGGTGGGAGGAATTGTTCCTTGG | 4 |
| wt-73380-73500 | 3465 | GATTCCACTGGCCCCGGCAGGTGGCCTGGCCGGCCAAGCATAGAAACCCTTGAAGCGTGGGGGGGTATGTGGGACCCTCATCCGCGTGCCAGCGCGCGAGCTCCACCAGCTCCCGGGCCA | 8 |
| wt-73500-73620 | 3466 | CGTCCACACTGAGCCCGGCCCAGGCCCGCATGAGTCCGTCATCGGGGTCGGGGTCCCACGTGTATGGGGCCGGGGGCTCCATGCGGATTTTCAGCTGCTGGACACGCACATGCTCAGCCA | 8 |
| wt-73620-73740 | 3467 | GGTAAGTCTCCCGGGTGAAGTAGGTGCGCATGTGCTCCGCAAAGCCCCTGTCCAGGAGCGAGGGGAGCACGACGCCCCCCGAAGGCAGACACCCAATTTCTCCCATGCTCGTTAACTGAG | 4 |
| wt-73740-73860 | 3468 | AGTATCGCTTAAAGGTTCCCTCGTTGAAGCACTGTGCGTGGGCCAAATAGACGTAGCGCACGAGATCGGCCGAGGCCAGGGGAAGGCGCCCCCTGTAGGCGTCTATCGTCCTTGCCACAG | 4 |
| wt-73860-73980 | 3469 | CGCGGATCTCTCGCGAGTCCCGCCGCAGTTTCTCGTGTGCAAAGTGGGCAAAAGCCTCGGTCTGCTCCGCCCATGCCGAGGAGCCAAAGACCTCCCCCAGCTCGGCCAGGGACGTGACGG | 8 |
| wt-73980-74100 | 3470 | CGGCCAGGCTCTGACCAGACTCGGAAGTAAATAGCTCCGTGAGGTGCGCCAGGGTCTCAATCGTACAAGGAATGCCCCAAAAATAGTAAGCAGCCGTGACTAGCACGAACTGGGCCTCGT | 4 |
| wt-74100-74220 | 3471 | GGGAGCCAAAGGTGCTAATGAACCACCTGGCCGAGATGTTAACGCGGTAGATGCGGCGCAGACAGCCCACGATCTTGGGACGCAGCCACGCCACGCGGCCTCTGGCATCCCCCTGTGGCT | 4 |
| wt-74220-74340 | 3472 | GTTTCTTAGCGCTCAGTGTGAGCAGTTCCACGAGGGGCGTGAGCGAGCGCAGGGCCCCCGCGCGATCTAGGTAGGTGGATAGACGGTCCGCGGTGAGCGGCGTGAGGCCGCGCAGGAAGG | 8 |
| wt-74340-74460 | 3473 | GGAAGGCCTCCTCCGCCGGCAGGTGCAGCGTCAGAACCAGGCCGCAGCGGCTCTGTGAGGTCAGCCGCTTCTTGGGCAGGTGAAGCTGCAGTTCCACGAGAGAACCCGCCACGTGGTGGA | 8 |
| wt-74460-74580 | 3474 | GGGGCGAGGCGTTGTGGCACAAACAAAACAGGCGGAAGCCCTCGTCAGGCCGCGAGAGGATGGCATCGAGGATGGCCTCCGCAATGTCAGTGTTTGAGGCCACAAGGGCCTTGATGACGA | 4 |
| wt-74580-74700 | 3475 | CGGGGGCGGACATTATTTAAGACCGGGAGGCCCCAACGGCGGGCTAAACAGAACGATGGCCTTCTATCTCCCAGACTGGTCGTGCTGCGGGCTCTGGCTCTTTGGCCGGCCCAGGAATAG | 4 |
| wt-74700-74820 | 3476 | ATACAGCCAGCTCCCTGAGGAGCCGGAGACCTTTGAGTGCCCGGACCGCTGGCGAGCCGAGATAGATCTGGGCCTGCCCCCTGGTGTGCAGGTGGGAGATTTGCTAAGAAATGAGCAGAC | 4 |
| wt-74820-74940 | 3477 | GATGGGCTCACTGAGACAGGTTTATTTGCTCGCTGTTCAAGCCAATAGCATCACGGATCACCTGAAGCGCTTTGACGCCGTCCGCGTCCCTGAGAGCTGTCGTGGGGTGGTGGAGGCCCA | 4 |
| wt-74940-75060 | 3478 | GGTGGCCAAGCTTGAGGCCGTGCGCTCAGTCATCTGGAATACCATGATCTCTCTGGCTGTAAGCGGCATCGAGATGGACGAGAATGGGCTCAAGGCCCTGCTGGACAAACAGGCTGGCGA | 4 |
| wt-75060-75180 | 3479 | CAGCCTGGCCCTGATGGAGATGGAGAAGGTGGCCACGGCGCTCAAGATGGACGAGACCGGTGCCTGGGCGCAAGAGATCTCGGCCGTTGTCTCATCGGTGACCGCCCCCTCAGCCTCGGC | 8 |
| wt-75180-75300 | 3480 | CCCTTTCATCAACTCCGCCTTTGAGCCCGAGGTGCCCACCCCCGTCCTTGCACCGCCTCCCGTGGTGCGGCAGCCGGAGCACTCTGGGCCCACGGAGCTCGCGTTAACGTAGCAACCAGA | 8 |
| wt-75300-75420 | 3481 | CTCCACACCAAATAAACATTTTATTGGTAAAACAAGGGATATGAAGGTGTCATTGACCCGAGGATCCAAACCCCCTCCCCTGTCTCCCCTCGAGCGCCTCGCTCAGCCCACTATCACCCA | 4 |
| wt-75420-75540 | 3482 | TGGCCAGGCTCGGCACCTCCTCGAAGGTGCAGCTGGCCCACCTAAAGAGAGATCTGGGGCCAAGGACCCCCGCGTCACTGTGGGGGCTGTAGAAGGAGGTGAGGTGGTGCTTGTGAAGGT | 4 |
| wt-75540-75660 | 3483 | AAACAAGCTGACAGAAGCGCCGGTACTTGTTAAGGAACACGGTCTGGTCACTAAAGTTGGTCAGGCTGACGTCCACCCCACCCCGGCGCCACCTGCAGGGCTTCACTAGAATACCCTGCA | 4 |
| wt-75660-75780 | 3484 | TGGCCAGGCCCGACCTGCCAAAGATTGTCGGCCTGTGGTGAGGGATAGAAGGGGGGGGCACGGTGAGTGTCACTGAGACGGTCTGATGGGGGAAGAGGGCCAGGTCCTTTGGCAAAGAGA | 4 |
| wt-75780-75900 | 3485 | CGTCCAGGCCCACGTCCCCGGGGTACTGGGGGTGGTTGATGGGACCCTTGTCCTCCTCCATCTGGGGGGTGGCATATCTGAAGGCAGCCAGGTGGATTTTGAGCTCCGATGGACGCAGCG | 4 |
| wt-75900-76020 | 3486 | TGGAGTTGTAGCGCCGCTGATTCTGGAGGATTAGCCGGAGTTCCCCCGTGTAGCCGGGATCGATGATGCCAACATGAGACGTGACCGGACGGGAGGTGCTGCCCCACAGCATGAGCCCAT | 4 |
| wt-76020-76140 | 3487 | GACCCTCGGGTGGGCGGGCATAGAGGCCTAGGTCCACAGTTGTGGTCTTCATCGGGCGCAGCAGGATGGTGGTCTTGTTGACCAAGGTGAGCCGCCCTACACTAGCCTGCTGGAGCAACA | 4 |
| wt-76140-76260 | 3488 | GCTTGTCATTCTGGAAGGCGTAGCGTATGTGTGGACAGGCCTCCATGGTGATGATCTAACAGACAGGGACGGCGGCGCTATATATAAGAGCCCAAGACCCGGCTCTCTTTACTGCGAAAT | 4 |
| wt-76260-76380 | 3489 | GGGGAAGGTCCTAAGAAAGCCGTTTGCAAAGGCTGTGCCACTGCTCTTCCTCGCCGCCACCTGGCTTCTGACCGGGGTGCTGCCGGCCGGCGCTTCCAGTCCCACAAACGCGGCGGCGGC | 8 |
| wt-76380-76500 | 3490 | TTCCCTGACTGAAGCCCAGGACCAGTTCTACTCCTACACATGTAATGCGGACACATTCTCGCCTTCTTTGACCAGCTTTGCCTCCATCTGGGCACTTCTGACGCTTGTCTTAGTCATTAT | 4 |

| | | | |
|---|---|---|---|
| wt-76500-76620 | 3491 | AGCCTCAGCCATCTACCTGATGTACGTCTGCTTTAACAAGTTTGTGAACACGCTGCTGACGGATTAGATGGGGATATTTAAAAGGGGCAGCAATCTCGGCTGTTTGTACTTCTTCTCTGC | 4 |
| wt-76620-76740 | 3492 | TCGTTAAACCAATAGCATGTCAGCTCCACGCAAAGTCAGATTGCCTTCTGTTAAGGCTGTTGACATGAGCATGGAAGACATGGCCGCCCGCCTGGCTCGCCTGGAGTCTGAGAATAAGGC | 4 |
| wt-76740-76860 | 3493 | TCTGAAGCAACAGGTCCTCAGAGGGGGTGCCTGTGCCTCGTCTACCTCTGTTCCTTCTGCTCCAGTGCCTCCGCCTGAGCCGCTTACAGCTCGACAGCGAGAGGTAATGATTACGCAGGC | 4 |
| wt-76860-76980 | 3494 | CACGGGCCGTTTGGCGTCTCAGGCTATGAAGAAGATTGAAGACAAGGTTCGGAAATCTGTTGACGGTGTAACTACCCGCAATGAAATGGAAAATATATTGCAAAATCTGACCCTCCGCAT | 4 |
| wt-76980-77100 | 3495 | TCAAGTATCTATGTTGGGTGCAAAAGGCCAACCCAGCCCTGGTGAGGGAACACGACCACGAGAATCAAACGACCCCAACGCCACCCGACGTGCCCGCTCCCGCTCCCGGGGACGTGAAGC | 4 |
| wt-77100-77220 | 3496 | AAAGAAAGTGCAAATTTCTGATTAATAAATTTTTATTGACTTTATACATAGGTCTCGGCGTCATCATATGGTGGGGTGGTGTAGGTATGGGATGTAGACAAGTTACGCCTAAAGGCGCAG | 4 |
| wt-77220-77340 | 3497 | TCCGCCATGACCAGCAGCAGCAGAAGGGTCAGCACAGCCAGAGAGGCCCACTGCAGTACTAGCATGGAGAGGTTTGAGAATCTGGGCTGGGACGTTGGCGGGACTGGCACGGTGGCTTGG | 4 |
| wt-77340-77460 | 3498 | GCTGTGGTAACCGGTGGGCTCGTAAAAGTCCAGCGGGGCCGCAGTTTGCTAGAAGTGCTGGGAGGTAGATAGGTGGTCGCATTGTATCTCGGTCTTGGCGTAGTTGAATCACCGCCGTAA | 4 |
| wt-77460-77580 | 3499 | TCTGTGGTGGGCTCTGTACTTGTCCGGGCTCCATGTCCTGTGGTGTGCTTTCCACCGGTGGTAGAATTGGCCTTTCCACCTGTTGAGGTGACCGTGGGAACCGCCGTCTTTTGGCCACTG | 4 |
| wt-77580-77700 | 3500 | GGGGCCTGGGGCGACGTTGCATTTTGGGGGGGCGTGCCTTTGGTGACATTAACCTCCCCCGGTTTTGTGGATGTGGAACTGTTTCCAGGGCCTGACGCTTGGCTGGTGGTGCCTGGGCGG | 4 |
| wt-77700-77820 | 3501 | GGTGCTGGCGAACTGGTGGACACATGATGTGTGCTGATAGAGGCTGGTGTCACCTGTGTTATATTTTCACCACCTGTTGGGTGAGCGGAGGTTAGTAAAGGCATATGTGACGTTGAATTG | 4 |
| wt-77820-77940 | 3502 | TCACTGGTGGAGGGGCTGAGTGTCTCTGGGTTTGAACTGGGTCTCAGTGACATGGAAGAGGTTGAACTTGAAGTTATGTTATGTTGGCCTGTGGTAACAGCACTGGTTGCATTTTTTGGT | 4 |
| wt-77940-78060 | 3503 | TGGCTGGTAACTACTGGGGTGGGACTTGTTCCTCCTAAGGTGTGGTTGGTGGCATTTGCCTGTGGACTTGTTTCTCCCACAGTAGGGCCGGTGGCATTTGGGGTTGGGGTAGTCACTGCT | 4 |
| wt-78060-78180 | 3504 | GAGGTGGGGCTTGTTTTTCCCAAGGTGGGGCTGGTGGCATTTGGGGTTGGGGTAGTCACTGCTGAGGTGGGGCTTGTTTTTCCCAAGGTGGGGCTGGTGGCATTTGGGGTTGGGGTAGTC | 4 |
| wt-78180-78300 | 3505 | ACTGCTGAGGTAGGACTTGTTTTTCCCAAGGTGGGGCTGGTGGCATTTGGGGTTGGGGTAGTCACTGCTGGGGTGGGGCTGGTGGCATTTGGGGTTGGGGTAGTCACTGCTGGGGTGGGG | 4 |
| wt-78300-78420 | 3506 | CTGGTGGCATTTGGGGTTGGGGTAGTCACTGGTGAGGTGGAGCTGGTCATGTCGGGGGCCTTACTTTCTGTGCCGTTGTCCCATGGAGATGGACTTGGTGTCACCGGTGATGCGCCTGAC | 4 |
| wt-78420-78540 | 3507 | GTTGTGCCGGCTGGTGTTGGGCTGGTGACATCCGCGGTGGATACAGTGGGGCCTGTGCTTGCAGGTGCGGTGAGGTTGGTAGGCACGTGAGTAGAGCTGGGTAGACCTGTCGTTGTATTG | 4 |
| wt-78540-78660 | 3508 | GGATCAGCAAATCCAGTTGTATTCAAGGTAGGGGAGGTGGTGGTGCTCTCGGGTGCCTTGGAGAATATAACCTTGTGGGTTGTTGTGGTGGCATTGGTAGCCGTTCGTGTGATAATGAGT | 4 |
| wt-78660-78780 | 3509 | GTCTTGGGGGCCGTGCCAAGACCCGAGACAGTAATGTCAAATGTCCGATTGCTCGCAAATGCACCAGAAATATTTTCACAACCCGAAGGTGTCCCCGAGGTGAGAGTCCATTTGCACTTA | 4 |
| wt-78780-78900 | 3510 | AAGTCAGTTTCAGTGTTGTTTGGCCAGGCCCAAAAGGCAGTCACTGTAACATTTGGCGAGTTTGCGTCCTCAGAAGTGACCATTGGCACTGAATAGGTAGCATTGTCACCCACATATGTG | 4 |
| wt-78900-79020 | 3511 | ATGTCTGTGGTGTTTGTCGGCATGTCCTGTGAAGCTGGAATCTCATCAGAGAACACAATGTTGGACTGAATGCAGTAATCTCCCCCGCTCGCCTTCGGTCCATTCCCAGAGTAAAACACG | 4 |
| wt-79020-79140 | 3512 | TACAGGATACTGTTATTGCCAAGAAATCGTGACACTGGACGTGGTGTCAGACGCAGGCTGTATGCATACCCTGTACCAGGTATTGGGGTGGCCACGGGACTCGTTGATGTGAGAATTCCG | 4 |
| wt-79140-79260 | 3513 | CCGCTGGGAACATGGCTCTCGTATCCACTGCAGGTGATGTTAAATTTGTTGTCTCCGGGCAGAACTTGTGAAATTTCGCCATCCTCCATAATACACTCAATATCTATCTCATTACCGAGC | 4 |
| wt-79260-79380 | 3514 | ATTTCTGTTTTTACGCTGAAATTCGAGTCTTGAGCTGACGTTGGCAAACTTAAGGGTAGCGTGACATCCAGCCCCTGTGCCCTCACTACTGCCGTTATATTGGTAGAATTACAGTTATCC | 4 |
| wt-79380-79500 | 3515 | CACTTTATGTATGGCACTGTTTCTGGTATCAGGTACACGGGGTTTTGCATTTCTGCATGGTGGCACCACATGGTTCCAAACACATCTTGAAAGTAGACATCTACAGATTCCAGGCTTACT | 4 |
| wt-79500-79620 | 3516 | TGTTGCTCCTCTCCGGTGGTGACGTTAATTGGAAGCTTCTTAGACCGCATAGTTAGAGCCAATTCTCCTGCACCAAGGAGCTCCAGTAGAAAGAGATTGGTGGCATTTTCTGAGCCACCA | 4 |
| wt-79620-79740 | 3517 | AATGCACCTCGAGGTTGGTAGACAGCCTTCGTATGGGGTGTCAGCTGGCCAAAGTCAAGATCAAGTTGATGCTTTTTGCCCCCGACATCGAAATTGATAGTTACATTGACATCTGCCGTG | 4 |
| wt-79740-79860 | 3518 | CAAACATTGCATGTGGGGTAAAATGGGAATTCCGGAATCTCAACATTGAAAAAACCAGGATCTTCACCCGTGAGATGGATCAGGCTCTGGATGGTGTACTGACACACAAGCAAGGCTGCC | 4 |
| wt-79860-79980 | 3519 | TCCATTGTCTCGGCACCGATTTCTAGGCAGCATCCTCTTTAATAGGTACAAGGGGGGTGCGGTGTTGGTGAGTCACACTTTTGTTGCAGACAAAATGGACAAGGACAGGCCGGGTCCCCC | 4 |
| wt-79980-80100 | 3520 | GGCCCTGGATGACAACATGGAAGAAGAAGTCCCATCTACCTCGGTTGTGCAGGAACAGGTATCGGCGGGAGATTGGGAAAATGTCCTCATAGAGTTATCAGATAGCAGCTCAGAAAAGGA | 4 |
| wt-80100-80220 | 3521 | AGCAGAAGATGCCCACCTGGAGCCGGCCCAGAAGGGTACGAAGAGAAAGCGGGTCGATCATGATGCCGGTGGGTCAGCTCCAGCACGACCCATGCTCCCACCCCAGCCGGATCTCCCTGG | 4 |
| wt-80220-80340 | 3522 | GCGAGAAGCCATTCTCCGCAGGTTTCCACTAGATCTAAGAACACTTCTTCAAGCGATTGGAGCCGCGGCTACGGTGAGCATCCCTATGGCCTAAGTGTGTGATGTGTGTTTTTACCCATC | 4 |
| wt-80340-80460 | 3523 | ACACAACAACAAGGTAAGTAATTTGTTGCCGTTGGTTTCAGCGCATCGACACACGAGCCATAGACCAGTTTTTCGGATCCCAGATTTCAAATACCGAGATGTACATAATGTATGCCATGG | 4 |
| wt-80460-80580 | 3524 | CCATTCGACAGGCCATTAGAGATCGTCGGAGAAATCCAGCTTCTCGTAGAGATCAGGCCAAATGGAGACTGCAAACCCTGGCCGCCGGATGGCCTATGGGTTACCAGGCATACAGCAGCT | 4 |
| wt-80580-80700 | 3525 | GGATGTACAGCTACACCGATCACCAGACGACTCCCACATTCGTACATCTCCAGGCGACACTTGGGTGCACAGGTGGCCGTAGGTGTCACGTGACCTTTTCCGCCGGCACCTTTAAGCTGC | 4 |
| wt-80700-80820 | 3526 | CGCGATGTACCCCCGGGGATCGCCAGTGGTTGTATGTTCAGAGCTCCGTGGGTAACATTGTACAGAGCTGTAATCCCCGCTACAGTATTTTCTTTGACTATATGGCTATACACAGGAGCC | 4 |
| wt-80820-80940 | 3527 | TCACGAAAATCTGGGAGGAAGTTTTAACACCTGACCAGCGTGTTTCATTTATGGAATTCCTAGGATTTTTGCAGAGAACGGATTTGTCCTATATCAAGAGCTTTGTCAGCGATGCCCTGG | 4 |
| wt-80940-81060 | 3528 | GCACCACTAGTATCCAAACACCGTGGATCGATGACAATCCTAGCACGGAGACGGCACAGGCTTGGAATGCCGGCTTTCTCCGGGGTCGTGCGTATGGGATAGACTTGCTTAGAACTGAAG | 4 |
| wt-81060-81180 | 3529 | GGGAACATGTCGAAGGTGCTACCGGTGAAACGCGAGAAGAAAGTGAGGACACGGAGAGCGATGGAGATGATGAAGATCTTCCTTGTATAGTGTCCAGAGGTGGACCTAAGGTCAAACGAC | 4 |
| wt-81180-81300 | 3530 | CCCCTATATTTATAAGACGTCTGCACAGGTTGCTGCTGATGAGAGCGGGCAAACGAACAGAACAGGGCAAGGAGGTACTGGAAAAGGCCCGTGGGAGCACTTATGGCACACCTAGGCCGC | 4 |
| wt-81300-81420 | 3531 | CTGTTCCGAAACCAAGACCAGAGGTCCCACAAAGCGACGAGACAGCTACCAGTCACGGGTCGGCGCAAGTCCCAGAACCCCCAACCATTCACCTAGCAGCTCAGGGAATGGCATACCCAT | 4 |
| wt-81420-81540 | 3532 | TACATGAACAACACGGCATGGCCCCGTGTCCGGTAGCACAGGCCCCACCTACGCCCTTGCCCCCTGTATCTCCAGGGGATCAACTCCCAGGTGTTTTTAGCGACGGGCGAGTGGCGTGTG | 4 |
| wt-81540-81660 | 3533 | CACCAGTACCCGCCCCGGCTGGGCCTATTGTCCGGCCCTGGGAGCCATCCCTGACACAGGCTGCGGGGCAGGCCTTTGCACCCGTTAGACCACAACACATGCCAGTAGAACCCGTCCCTG | 8 |
| wt-81660-81780 | 3534 | TCCCGACAGTGGCACTTGAGCGACCAGTTTACCCCAAGCCAGTTCGTCCGGCACCTCCTAAGATTGCTATGCAGGGCCCCGGGGAAACTTCTGGCATTAGACGCGCGCGGGAGCGTTGGA | 4 |
| wt-81780-81900 | 3535 | GGCCCGCACCTTGGACGCCAAATCCACCCCGTTCTCCCAGTCAGATGTCCGTGCGTGACCGTCTGGCTCGTTTGCGTGCTGAGGCACAGGTCAAACAGGCTAGTGTTGAGGTGCAGCCCC | 4 |
| wt-81900-82020 | 3536 | CCCAGTTGACCCAAGTATCCCCTCAGCAACCAATGGAGGGGCCGTTGGTACCAGAGCAGCAGATGTTCCCTGGTGCCCCCTTTAGCCAGGTTGCTGATGTGGTCCGGGCACCTGGGGTAC | 4 |
| wt-82020-82140 | 3537 | CGGCGATGCAGCCACAGTACTTTGACCTCCCCTTAATTCAACCCATTAGCCAGGGGGCACCCGTGGCCCCGTTGAGGGCTAGTATGGGCCCGGTACCTCCGGTACCGGCAACACAGCCAC | 4 |
| wt-82140-82260 | 3538 | AGTATTTTGACATCCCCTTAACTGAACCCATTAACCAGGGGGCATCCGCGGCCCATTTTCTCCCTCAGCAACCGATGGAGGGGCCGTTGGTACCTGAGCAGTGGATGTTCCCAGGTGCCG | 4 |
| wt-82260-82380 | 3539 | CCCTGAGCCAGAGTGTTAGGCCAGGGGTAGCGCAGTCACAATATTTTGACCTCCCCTTAACTCAACCCATTAACCATGGGGCACCCGCAGCCCATTTCCTCCATCAGCCACCAATGGAGG | 4 |
| wt-82380-82500 | 3540 | GGCCGTGGGTACCCGAGCAGTGGATGTTCCAAGGTGCCCCCCCTAGCCAAGGCACTGACGTGGTCCAACATCAGCTGGATGCTTTGGGGTATACACTCCATGGTCTTAACCATCCCGGGG | 4 |
| wt-82500-82620 | 3541 | TTCCCGTGTCTCCTGCCGTTAACCAATATCATCTCAGCCAGGCTGCCTTTGGGTTACCTATTGATGAGGATGAGAGTGGCGAGGGGTCCGATACCTCCGAGCCGTGTGAAGCTCTTGATT | 4 |

| | | | |
|---|---|---|---|
| wt-82620-82740 | 3542 | TGTCAATCCATGGCAGGCCCTGCCCTCAGGCCCCCGAGTGGCCTGTTCAAGAGGAGGGTGGCCAGGATGCCACCGAGGTTCTTGATTTGTCAATCCATGGCAGGCCCCGCCCTCGGACCC | 4 |
| wt-82740-82860 | 3543 | CCGAGTGGCCTGTTCAAGGGGAAGGTGGCCAAAATGTCACAGGCCCTGAAACTAGAAGGGTGGTGGTGTCAGCTGTTGTTCACATGTGTCAGGATGACGAGTTTCCGGATCTACAAGATC | 4 |
| wt-82860-82980 | 3544 | CTCCAGATGAGGCCTAAGCAAAGGTGTAGAAGTGTGTCCCCCTCCATTCCACCCACTGATAATACGCCCGACAATAAAGTTGATGATATTGAATTCCACACCTGCTTGTGTTTGTGATTT | 4 |
| wt-82980-83100 | 3545 | TATTTCATATTCCATGAGAGAGACCTCGCATATTTGCAGAAGGGTCACTGAAACATCTTATCTTAAAACAGTTACACCTGAATAATGAAGAAAGCGTGGCTCAGCAGAGCACAGCAAGCC | 4 |
| wt-83100-83220 | 3546 | GATGCCGGGGGGGCATCTGGCTCCGAGGACCCACCAGATTATGGAGATCAAGGTAATGTGACACAGGTGGGATCTGAGCCTATTTCACCTGAGATTGGCCCCTTTGAACTCTCTGCGGCC | 4 |
| wt-83220-83340 | 3547 | AGTGAGGATGATCCTCAATCTGGGCCAGTGGAAGAGAATTTAGATGCCGCTGCAAGAGAGGAAGAGGAACCTCATGAGCAGGAGCACAATGGTGGTGACGATCCCTTGGATGTCCATACT | 4 |
| wt-83340-83460 | 3548 | CGCCAGCCTAGATTTGTGGATGTGAACCCAACGCAGGCTCCAGTGATCCAACTAGTCCACGCTGTCTATGATTCCATGTTGGTAAGAGGCACCTAGAACATTTCCAGATGTTTCGCTTGG | 4 |
| wt-83460-83580 | 3549 | ATTTTTTGGCCAGTCTTAATTGATTGTCATTGGTTTCAGCAATCGGACCTCCGGCCCCTAGGCAGTTTATTCCTTGAGCAAAACCTGAACATCGAAGAATTTATATGGATGTGCATGACC | 4 |
| wt-83580-83700 | 3550 | GTGCGTCACAGATGTCAGGCCATCAGAAAAAAACCATTACCAATTGTTAAGCAGAGGCGTTGGAAGCTCCTGTCATCTTGCAGATCCTGGCGTATGGGTTACCGCACGCATAACCTCAAA | 4 |
| wt-83700-83820 | 3551 | GTAAACAGTTTTGAGTCAGGGGGGGACAATGTCCACCCGGTCCTTGTGACTGCTACGCTAGGATGTGATGAGGGCACGCGGCATGCAACAACGTACAGTGCTGGCATTGTACAGATACCA | 4 |
| wt-83820-83940 | 3552 | CGAATATCAGACCAAAACCAAAAGATCGAAACAGCCTTCCTGATGGCACGTCGTGCTAGGTCACTTTCGGCAGAAAGATATACTTTGTTCTTTGATTTAGTATCCTCCGGAAACACCCTG | 4 |
| wt-83940-84060 | 3553 | TATGCTATATGGATTGGGCTGGGCACGAAAAACCGAGTTTCATTTATTGAGTTTGTAGGATGGTTATGCAAGAAGGACCACACTCATATACGCGAATGGTTCCGCCAGTGCACCGGGAGA | 4 |
| wt-84060-84180 | 3554 | CCCAAAGCAGCCAAGCCATGGTTAAGAGCGCATCCTGTCGCCATTCCTTATGATGATCCGTTAACAAACGAGGAGATTGATCTGGCCTATGCCCGCGGGCAGGCCATGAATATTGAGGCT | 4 |
| wt-84180-84300 | 3555 | CCTAGACTGCCAGATGATCCTATAATTGTTGAGGATGACGACGAAAGTGAGGAAATTGAAGCTGAAAGCGACGAGGAGGAAGACAAGAGTGGAATGGAATCTCTTAAAAATATACCGCAA | 4 |
| wt-84300-84420 | 3556 | ACACTGCCGTACAATCCAACAGTATACGGCAGGCCCGCGGTGTTTGACCGAAAGTCAGATGCAAAATCAACCAAAAAATGCAGGGCCATAGTAACTGACTTTAGTGTAATCAAGGCCATT | 4 |
| wt-84420-84540 | 3557 | GAAGAGGAACACAGAAAGAAGAAGGCAGCCAGAACAGAGCAGCCAAGAGCCACGCCTGAATCCCAGGCCCCCACAGTGGTCCTCCAGCGACCACCCACGCAGCAAGAGCCTGGCCCCGTC | 4 |
| wt-84540-84660 | 3558 | GGCCCACTGAGTGTCCAGGCTCGGCTGGAGCCATGGCAACCTTTGCCTGGGCCCCAAGTGACAGCAGTTCTACTTCACGAAGAATCCATGCAGGGTGTCCAAGTACATGGTTCGATGCTA | 4 |
| wt-84660-84780 | 3559 | GACCTTCTTGAAAAAGACGATGAAGTCATGGAGCAGAGGGTTATGGCAACCCTACTGCCACCAGTACCACAACAGCCCCGGGCTGGCAGAAGAGGCCCTTGTGTCTTCACCGGTGACCTA | 4 |
| wt-84780-84900 | 3560 | GGCATAGAGAGTGATGAGCCCGCTTCGACAGAGCCGGTTCATGATCAGCTACTGCCTGCCCCAGGACCTGACCCTCTTGAAATCCAACCACTAACGTCCCCCACCACGTCTCAACTTAGC | 4 |
| wt-84900-85020 | 3561 | AGTTCGGCACCAAGCTGCGCACAAACTCCATGGCCGGTGGTTCAGCCAAGTCAGACTCCAGATGACCCAACGAAACAGTCCCGGCCACCGGAAACAGCTGCCCCACGCCAGTGGCCAATG | 4 |
| wt-85020-85140 | 3562 | CCCCTGCGACCTATCCCTATGCGCCCCTTGCGGATGCAGCCAATCCCATTTAATCATCCAGTGGGACCCACTCCCCATCAGACACCTCAAGTGGAAATAACACCATATAAGCCCACTTGG | 4 |
| wt-85140-85260 | 3563 | GCTCAGATAGGGCACATTCCATATCAGCCTACACCAACGGGTCCTGCTACCATGCTGTTGCGCCAGTGGGCACCCGCCACCATGCAGACACCACCGAGAGCGCCCACTCCCATGTCACCA | 4 |
| wt-85260-85380 | 3564 | CCTGAGGTGCCACCCGTTCCCCGGCAGAGGCCTCGGGGGGCGCCCACTCCCACGCCACCTCCTCAGGTGCCGCCCGTTCCCCGGCAGAGGCCTCGGGGGGCGCCCACTCCCACGCCACCT | 12 |
| wt-85380-85500 | 3565 | CCTCAGGTGCTGCCCACTCCCATGCAGCTGGCACTAAGGGCTCCTGCTGGTCAGCAGGGGCCGACAAAGCAAATTTTGCGCCAATTGTTAACGGGGGGCGTCAAGAAAGGGAGACCATCA | 4 |
| wt-85500-85620 | 3566 | CTTAAGTTACAGGCCGCCCTTGAGCGTCAAGCCGCTGCGGGCTGGCAGCCTTCACCAGGGTCCGGCACCAGTGACAAGATTGTGCAGGCGCCTATTTTCTACCCACCCGTTTTGCAGCCC | 4 |
| wt-85620-85740 | 3567 | ATACAGGTTATGGGGCAAGGGGGTTCCCCAACGGCCATGGCCGCCTCAGCGGTGACACAGGCACCCACGGAATATACCAGGGAAAGGAGGGGAGTGGGGCCTATGCCTCCCACCGATATT | 4 |
| wt-85740-85860 | 3568 | CCGCCGTCTAAACGAGCGAAGATCGAGGCCTATACAGAGCCCGAGATGCCGCACGGGGGGGCCTCGCACTCTCCCGTCGTTATCTTGGAGAATGTCGGCCAGGGGCAACAGCAGACTCTG | 4 |
| wt-85860-85980 | 3569 | GAGTGCGGAGGAACTGCTAAACAGGAAAGGGACATGTTGGGGCTGGGGGACATTGCAGTTTCTTCCCCTTCCTCTTCTGAAACATCGAACGATGAGTGATTTCGCCCATGTAACAAGAAC | 4 |
| wt-85980-86100 | 3570 | TGGGATGAACCCTGGGGCAACAGACTGCGGGGAGGAGGGGGGCAGTGATAAGTCATGACAATTTTAGATGAGGTAGAAATTTTGCATATTTTCAGACCCACCATGGAATCATTTGAAGGA | 4 |
| wt-86100-86220 | 3571 | CAGGGGGACTCTAGACAGTCACCCGACAATGAGCGGGGAGATAATGTACAGACTACCGGCGAGCATGATCAGGACCCTGGGCCGGGGCCTCCATCCAGTGGGGCTTCTGAGAGATTGGTA | 4 |
| wt-86220-86340 | 3572 | CCAGAAGAGTCATACTCAAGAGATCAGCAACCTTGGGGGCAAAGCAGGGGTGATGAAAACAGAGGCTGGATGCAGCGCATCAGGCGAAGGCGGAGAAGACGGGCTGCCTTGTCCGGCCAT | 4 |
| wt-86340-86460 | 3573 | CTTTTAGACACGGAAGACAATGTGCCGCCATGGTTGCCTCCACACGACATCACACCATATACCGCAAGGAATATCAGGGATGCTGCCTGCCGGGCTGTCAAGGTGAGTATGCCTCTAACT | 4 |
| wt-86460-86580 | 3574 | GGGTTCATGGGGGCCATCTAAGGCCCACGTGTGACCCATGTTTCCATTAATTTTAGCAATCGCACCTGCAAGCGCTATCAAACCTGATACTCGATAGTGGGTTAGACACACAACACATCT | 4 |
| wt-86580-86700 | 3575 | TGTGCTTCGTGATGGCAGCCAGGCAGCGTCTTCAGGACATTCGACGTGGACCCTTGGTTGCGGAGGGCGGTGTGGGTTGGCGACATTGGCTTCTAACATCTCCCAGCCAATCCTGGCCCA | 4 |
| wt-86700-86820 | 3576 | TGGGATATCGTACAGCAACACTACGCACATTAACTCCCGTGCCTAACAGGGTTGGGGCTGACAGCATCATGTTAACTGCCACATTTGGATGCCAAAATGCGGCACGAACTCTAAACACCT | 4 |
| wt-86820-86940 | 3577 | TCTCTGCCACCGTGTGGACACCACCCCATGCTGGACCAAGAGAGCAAGAAAGATACGCTCGGGAAGCCGAGGTACGCTTCCTTCGTGGTAAATGGCAGAGGCGGTACCGAAGAATCTATG | 4 |
| wt-86940-87060 | 3578 | ATTTGATAGAACTGTGTGGCTCTCTGCACCACATCTGGCAAAACTTGCTCCAGACCGAGGAGAACCTTTTAGATTTCGTGCGTTTCATGGGTGTCATGTCCAGCTGCAATAATCCAGCTG | 4 |
| wt-87060-87180 | 3579 | TGAATTACTGGTTTCACAAGACAATCGGAAACTTTAAGCCATATTACCCGTGGAATGCACCACCTAATGAAAATCCATATCACGCGCGGAGAGGCATAAAAGAACACGTAATCCAGAACG | 4 |
| wt-87180-87300 | 3580 | CATTTCGAAAGGCACAAATACAGGGTTTATCAATGTTAGCAACGGGAGGTGAACCCAGAGGTGATGCTACTAGTGAAACGAGCAGTGATGAGGACACCGGTAGACAGGGTTCGGACGTGG | 4 |
| wt-87300-87420 | 3581 | AGCTAGAGTCCTCGGACGATGAGCTGCCATATATCGATCCCAATATGGAGCCGGTTCAGCAGAGGCCCGTCATGTTTGTGAGCCGTGTGCCTGCAAAGAAACCGAGGAAACTGCCTTGGC | 4 |
| wt-87420-87540 | 3582 | CCACGCCCAAGACGCACCCAGTGAAGCGCACAAATGTTAAGACCTCTGATAGATCTGATAAGGCAGAAGCACAAAGCACCCCTGAAAGGCCGGGCCCTTCCGAACAATCATCAGTGACCG | 4 |
| wt-87540-87660 | 3583 | TGGAGCCCGCCCACCCGACCCCGGTGGAGATGCCAATGGTGATTCTCCATCAACCACCTCCAGTGCCCAAACCGGTTCCAGTCAAGCCTACGCCACCGCCTTCCCGTAGGAGAAGGGGAG | 4 |
| wt-87660-87780 | 3584 | CGTGTGTTGTGTACGACGATGATGTCATAGAGGTGATTGATGTTGAAACCACCGAAGATTCATCGTCAGTGTCACAGCCAAATAAGCCACATCGGAAACATCAAGACGGCTTTCAACGTT | 4 |
| wt-87780-87900 | 3585 | CAGGCCGACGTCAAAAACGAGCCGCGCCTCCCACCGTGAGTCCTTCGGATACTGGGCCTCCTGCCGTGGGGCCTCCTGCCGCGGGGCCTCCTGCCGCGGGGCCTCCTGCCGCGGGGCCTC | 8 |
| wt-87900-88020 | 3586 | CTGCCGCGGGGCCTCCTGCCGCGGGGCCTCCTGCCGCGGGGCCTCGCATACTGGCGCCTCTTTCCGCTGGGCCTCCTGCCGCGGGGCCTCACATAGTGACGCCTCCTTCCGCCCGGCCTC | 12 |
| wt-88020-88140 | 3587 | GTATAATGGCGCCTCCCGTCGTACGTATGTTTATGAGGGAGCGACAGCTCCCCCAGTCCACCGGCCGTAAACCTCAGTGCTTCTGGGAAATGCGGGCTGGTCGTGAAATTACACAAATGC | 4 |
| wt-88140-88260 | 3588 | AACAAGAACCAAGTTCACACCTGCAGTCCGCCACTCAGCCTACAACGCCTCGCCCATCATGGGCCCCATCAGTCTGCGCCCTCTCGGTGATGGATGCTGGTAAGGCCCAGCCCATAGAAA | 4 |
| wt-88260-88380 | 3589 | GTTCACACTTGAGTTCCATGTCGCCCACACAGCCGATATCGCACGAAGAACAACCCCGGTATGAGGATCCTGACGCTCCTCTGGATTTAAGTTTACATCCAGACGTTGCTGCTCAACCAG | 4 |
| wt-88380-88500 | 3590 | CTCCCCAGGCTCCATACCAGGGATACCAGGAGCCGCCGGCCCCCCAGGCTCCATACCAGGGATACCAGGAGCCGCCGCCCCCCCAGGCTCCATACCAGGGATACCAGGAGCCGCCGGCCC | 8 |
| wt-88500-88620 | 3591 | ACGGGCTCCAATCATCTTCATATCCAGGATATGCGGGTCCCTGGACCCCAAGGTCTCAACATCCATGTTATAGGCACCCCTGGGCACCATGGTCTCAAGATCCTGTGCATGGGCACACCC | 4 |
| wt-88620-88740 | 3592 | AGGGTCCATGGGATCCCAGGGCACCACATCTCCCACCTCAGTGGGATGGATCTGCAGGACATGGCCAGGATCAGGTCTCCCAGTTCCCACATCTGCAATCGGAGACAGGCCCACCACGTC | 4 |

| | | | |
|---|---|---|---|
| wt-88740-88860 | 3593 | TTCAACTTTCATTGGTGCCACTGGTCTCATCCTCTGCACCATCATGGTCATCTCCCCAGCCCCGAGCCCCCATACGCCCCATTCCAACAAGATTCCCCCCTCCCCCTATGCCGTTACAAG | 4 |
| wt-88860-88980 | 3594 | ATAGCATGGCCGTGGGGTGTGACTCATCAGGTACAGCATGCCCAAGCATGCCCTTTGCCAGTGATTACAGTCAAGGTGCATTTACCCCACTGGACATTAATGCCACCACGCCAAAAAGGC | 4 |
| wt-88980-89100 | 3595 | CTCGAGTAGAAGAAAGTTCTCACGGACCTGCCCGGTGTTCCCAAGCTACTGCTGAAGCACAGGAGATTCTCAGTGACAATTCTGAGATCTCCGTGTTCCCAAAGGACGCGAAGCAGACTG | 4 |
| wt-89100-89220 | 3596 | ACTACGATGCATCCACTGAAAGTGAGCTAGATTAAGGGGATCCAAGGTGACCCCTGTTAGCTATTTGATCTTTGACTGACACATAAACATGGTTTAAGGAATGAACACTCATGGTGTGAG | 4 |
| wt-89220-89340 | 3597 | ACTGGAACTGTACTAAATTTGCTGACATATGTACAATGAGAGCCAAAAATTTGATAAACCTTAAAAGTCCCCCCATCTAATGATGTCCAGTTCCCTTCTCCCACCCTGTACACCCCGACC | 4 |
| wt-89340-89460 | 3598 | CAAAGGGACTCAATGGCATTCAGATTTCTAGTTACCACAGGTAGAATATCGGGCGTTGGCCCATAAAAATAAGTGCATGGATATAGCTCTGCACAGGCTTGGAAACACCCATTCCAGGTG | 4 |
| wt-89460-89580 | 3599 | TGCTTCTTTTTGGTGAAATAAAAACAGCATCCTTTATATGAAAATGTGTATTCTCTGGTGTTGCAGTATGTACAGTTAGCTTTGGTATAGTTTTGGGGTACCTGAAATGTGTGCAGGGTG | 4 |
| wt-89580-89700 | 3600 | GGTGTCCAATGTGGCAGTTTTACCTCTTTGTCCCCATACTCCTGCTCGGCCGTCTTGTTAAAGTTAACCGGCGGTGGAGGATCCACCGGCCAGACCTCTACATTTGGTTTGGGTACCCAG | 4 |
| wt-89700-89820 | 3601 | GTGATGGCCGCGGCTGCCACCCGCCCTCCTCCTCTTACCCTGGGTGGCAAAAAGTATGCCAGGAGTAGAACAATAACAAGTGCGATGGCGGTAAACAATGGCACCCTCACCTGCTTAAAT | 4 |
| wt-89820-89940 | 3602 | GAAACCATGGCAACCACTTCAAAGAGAGCCGACAGGAAGATATTTATTAATATTCCATTAGTAAACGAGGCGTGAAGCAGGCGTGGTTTCAATAACGGGAGTTAGAAATTTAAGAGATCC | 4 |
| wt-89940-90060 | 3603 | TCGTGTAAAACATCTGGTGTCCGGGGGATAATGGAGTCAACATCCAGGCTTGGGCACATCTGCTTCAACAGGAGGCGCAGCCTGTCATTTTCAGATGATTTGGCAGCAGCCACCTGCGGA | 4 |
| wt-90060-90180 | 3604 | CAAAAATCAGGCGTTTAGATGGGGCATTTATGTTTGGGACGCTAGCCGCCTGGGCATTCGTGTTAGTATATACTGACCTCACGGTAGTGCTGCAGCAGTTGCTTAAACTTGGCCCGGCAT | 4 |
| wt-90180-90300 | 3605 | TTTCTGGAAGCCACCCGATTCTTGTATCGCTTTATTTCTAGTTCAGAATCGCATTCCTCCAGCTGCGAGCAAGGGAATGCGTTACTACAAGTGGTGCCTAGTCAGTTGAAACAAGCCCCA | 4 |
| wt-90300-90420 | 3606 | CCATCCGCTGCCGCCCCTCCATGAGCCCCACCGTCCGCTGCCGCCCCTCCTTGAGCCCCTCCTTACCGATTCTGGCTGTTGTGGTTTCCGTGTGCGTCGTGCCGGGGCAGCCACTGGTGC | 8 |
| wt-90420-90540 | 3607 | AGGCTGTGGAACACCAATGTCTGCTAGCTGTTGTCCTTGGTTAGCCCCGGGGCAAGCAAACACCACTGCTGCTGCTGTTTGAACAGTAGAATTGTCTCCAGGTTGAGGTGCTTCTCCCCC | 4 |
| wt-90540-90660 | 3608 | GGCTTGGTTAGTCTGTTGATTCTGGGTTATGTCGGAGACTGGGAACAGCTGAGGTGCTGCATAAGCTTGATAAGCATTCTCAGGAGCAGGCTGAGGGGCAGAAAACCACGACCCAGTCGG | 4 |
| wt-90660-90780 | 3609 | AGCGGTTGAAACATGATAGGCAGTTAGCTGGCCTTGTGGCAGAGGCTCTGGCAGCACCGGCCACAGCACACAAGGCAAAGGAGCTTGCGATGGCCCTCCCAGGTCCTGATAGACTCTGGT | 4 |
| wt-90780-90900 | 3610 | AGCTTGGTCAAAAGCTTGTACAAAAGGCACCTGGTATGGGTCAGGTGTAAATTTTACATCTTCAGAAGTCGAGTTTGGGTCCATCATCTTCAGCAAAGATAGCAAAGGTGGCCGGCAAGG | 4 |
| wt-90900-91020 | 3611 | TGCAATGTTTAGTGAGTTACCTGTCTAACATCTCCCCTTTAAAGCCAAGGCACCAGCCTCCTCTGTGATGTCATGGTTTGGGACGTGCTAAATTTAGGTGTGTCTATGAGGTACATTAGC | 4 |
| wt-91020-91140 | 3612 | AATGCCTGTGGCTCATGCATAGTTTCTAAAAGAGGAGGAGGCAGTTTTCAGAAGTGTCTAAAATAAGCTGGTGTCAAAAATAGACAGCCCAGTTGAAATATGCATGGCATGCAGCAGACA | 4 |
| wt-91140-91260 | 3613 | TTCATCATTTAGAAATGTATCCAAGATTTCATTAAGTTCGGGGGTCAGGGGGGAGTCCAGATTCAAATCCTCTGTCATGGACTCTAGTGTTGTGGTCAGTTCGTCCAAATGGCCACGAGG | 4 |
| wt-91260-91380 | 3614 | GGGCGGGTGGCTCAGGTCCATCTGTCCACATATGGCTGCTTCCTCCTTCTGGGGAATAACAGTGTCAGCCATCTCCCTTAGGGCCTTCACGGCCTGACTGGTTTCTTCATCAGGGTCCTC | 4 |
| wt-91380-91500 | 3615 | CAACAGATGACTTGCCTCGGGGGTTACTGCGGGGGCCGGGTCAAGTGGCTGGGGCACCGGGGCTGGCGTTAGGGATCCGACCGGTTCATGGACAGGTCCTGTGGGGGTGGGAGCCAAAGA | 8 |
| wt-91500-91620 | 3616 | GGCAGGCAGGGGCCGGTTGGCCCACGGGGATCCGGGTGGATGGAAGGGCCTGATCCTCTTTGGCTGACACACCTCTCGCCCCTCGAACACGTCAGATATGGCACTGCCCGCTTCCGGCTT | 8 |
| wt-91620-91740 | 3617 | TGGCAGGAACATACCTTCCCGGCTATCCCTGAGGCCCTTCTTCCTTTTAACGGGAGGAAGAAAGGTGGGCTTTGAGGGGTGGGGGAATATGGGTCTCTCATCGCTCTCTTGGTGGACCGC | 4 |
| wt-91740-91860 | 3618 | TGCTATCCAAGGCTGTTCAGGTTCCGCCGCGTTGGAAGGACATGGAGTTTGACCACGGTTGGGCCTGGATGTCCGGCGCGACTTTGGGGCCCGCAGGCGCGGGGCCTCGGCCCTGGCCTC | 8 |
| wt-91980-92100 | 3620 | GAGTTCATCTGTAAAAGCATGAAACTGTCCGGACTCCGAGTAGCGGGCCTCGGTGTGAGAGGCACCCCCATCATTCCCCATGAGCTCCTCGTCCATCCTGTCGGCTCCGGACACGAGGAT | 4 |
| wt-92100-92220 | 3621 | AGGAGTTTCCACTGCCTTGGACTTGGTTGACAGCAGGCACGCGGGAAGCACGCCGCTCACGTAGCTCCTCTGTCCGGCGTGGCTGGAGTAGGAGGCCCGGGGCAGTGTCTTAATCAGAGC | 4 |
| wt-92220-92340 | 3622 | CCTGACATCCTTAACATCGTCCGTCAGATGGCCTGTCTTGGACGAGACCATAGTCTGGAACATCTCCTCGAGGACGGGATAGGTGAACACCCACTTGCAAAAGGCCTTGAACTTGGAGCT | 4 |
| wt-92340-92460 | 3623 | TAGGAGGCCTTCCTTCTCCATCCTGTTCAGGTGTTCCACTACCTGCTTGCCGGAGGCCATGATGGCCGCGCGGTCCACGCCCAGCACCTTGCTGTAGGTGTAGGCCCGCACCCGACTGTG | 4 |
| wt-92460-92580 | 3624 | TTTTAGGAGCTTGTACATAGCGGTGCCTATGGTGGCAGGAATCATCACCCGGTTGCTGGGGGCCTGGATGAAGAATCTGTCAGTGACCACTATCAGGTGGTCTAACACGTAGCGCATCAC | 4 |
| wt-92580-92700 | 3625 | TATAGGGCACGCGATGGAACATGCGTCGTTGCCGGCATTCTCAGCCCGTCTTCTTACCCTGTTGTTTCGGAGAATGGCCCAAAAATTGCAGATGTTGAGCGTGGCCATTAGCCCGCCCCA | 4 |
| wt-92700-92820 | 3626 | TTCTCGCCCGTGGGCCTTGGCCTCATTTATAAATGCCTTGCATATTTTGTAGGATCTCAGAGTAATCTCCACACTCCCGGCTGTAAATTCCTTGTTGAGGACGTTGCAGTAGTCAGAGAC | 4 |
| wt-92820-92940 | 3627 | CAGAGAGCCCAGCTGCTTTTTGATTTCAGGAGTTAGCCTCAGAAAGTCTTCCAAGCCATCCTTTTTAGGCCTCATGGCTAGTAGTAACAGAGGAAATGCCCGACCATTAAAATCTTTCCT | 4 |
| wt-92940-93060 | 3628 | CCATGAGCTTTACCTGAAACACTATCCCGAAGTGGGGGATGTGGTGCATCTACTGAACACCATCGGGGTCGACTGCGACCTCCCACCTAGCCACCCACTCCTGACAGCCCAGAGGGGGCT | 4 |
| wt-93060-93180 | 3629 | GTTCCTGGCAAGAGTCTTGCAGGCTGTACAGCAGCACAAGCTGCTGGAAGACACCATCGTCCCCAAGATCTTAAAGAAGCTGGCTTATTTCTTAGAGCTGCTAAGCTACTACTCCCCCAA | 4 |
| wt-93180-93300 | 3630 | GGATGAACAGCGTGACATCGCCGAGGTTCTTGACCACCTCAAGACGAATCGGGACCTGGGGCTGGACGACAGACTCTGGGCCCTGATTAGGAAACTGCGCCAAGACAGACACCATGCCTC | 4 |
| wt-93300-93420 | 3631 | TGTAAATGTCCTCATGCCAGGAAGCGACTACACAGCCGTGTCGCTGCAGTACTACGACGGCATCTCCATAGGTATGAGGAAGGTAATCGCGGATGTCTGCCGCAGTGGCTATGCCTCCAT | 4 |
| wt-93420-93540 | 3632 | GCCCTCCATGACGGCCACGCACAACCTCTCCCACCAGCTCTTGATGGCGTCCGGGCCCAGTGAGGAACCGTGCGCCTGGCGCGGGTTCTTTAACCAGGTCCTCCTCTGGACTGTGGCCCT | 8 |
| wt-93540-93660 | 3633 | CTGCAAGTTTCGCAGATGCATTTACTATAACTACATTCAGGGATCTATAGCCACCATCTCCCAGCTTCTGCACCTCGAGATCAAGGCCCTCTGCAGCTGGATAATATCCCAGGATGGCAT | 4 |
| wt-93660-93780 | 3634 | GCGCCTCTTTCAACACAGCAGGCCTCTCCTCACCCTCTGGGAGAGCGTGGCCGCAAATCAGGAGGTCACGGATGCCATTACCCTGCCTGACTGCGCTGAATACATAGACCTACTAAAGCA | 4 |
| wt-93780-93900 | 3635 | CACAAAACATGTCTTAGAAAACTGTTCTGCCATGCAATACAAATAAATTTCTCTTACCTGCGTCTGTTTGTGTAGTGAGGTGTTGTGTCCTGTATGGTATTCTACTTTAAAAAGGCCGGC | 4 |
| wt-93900-94020 | 3636 | TGACATGGATTACTGGTCTTTTATGAGCCATTGGCATGGGCGGGACAATCGCAATATAAAACCCTGACCATCACATGGGGCATTAGGCGACTCTGCATCAGCATCGCTTAAGTATGAGTG | 4 |
| wt-94020-94140 | 3637 | GGCAGCAGAGAGGCTCGGTTATTTTGGTTCCTGAACATCTGGCTGGGGCATTAACTAAGCTTATGAGCGATTTTATCACAGGACAAGATGTCACTCTTTCTGGAGGAAATATTGCAGTCA | 4 |
| wt-94140-94260 | 3638 | AAATTCGCGATGCTATAAACCAGACCCCCGGGGGTGGTGATGTAGCTATACTTTCTTCCCTGTTTGCTTTATGGAATGCCCTCCCAACATCTGGTAGACAATCCTCCAGGGACGATTTAA | 4 |
| wt-94260-94380 | 3639 | TCCCAGCCGCCGTGCAGGCCTTAACCACGGCCCACAACTTATGTCTGGGTGTTATTCCAGGTGAGACCTCACACAAGGACACACCCGAGTCATTGCTCCGGGCTATCGTGACGGGTCTCC | 4 |
| wt-94380-94500 | 3640 | AAAAATTGTGGGTGGATTCGTGCGGATGTCCAGAGTGCCTACAATGTCTTAAGGGATTGAAGGCAATTAAGCCCGGCCTTTATGAAATCCCTAGGATAATACCACACACTAAGCAGTGTA | 4 |
| wt-94500-94620 | 3641 | GTCCTGTCAATCTCCTGAACATGTTGGTCCACAAGCTTGTGGCTTTACGTGGTCATGTGCAGCTTGCATACGACGCCCGTGTCCTGACGCCTGACTTTCACGAAATCCCTGACCTCGATG | 4 |
| wt-94620-94740 | 3642 | ACTCCGATGCTGTTTTCGCACGCACCTTATTGGCAGCCTTATTTCACCTCAATATGTTCTTTATTCTCAAAGATTACATAACACAAGACTCCATGAGCTTGAAGCAGGCCCTCAGTGGTC | 4 |
| wt-94740-94860 | 3643 | ATTGGATGTCTGCCACGGGCAACCCCCTGCCTGCAGCACCGGAAACCCTGCGAGACTACTTGGAAGCTTTCCGAAATTCGGATAATCACTTTTATCTCCCGACGACAGGGCCTTTAAACA | 4 |
| wt-94860-94980 | 3644 | CCTTCCAATTTCCCGAAGAGCTTCTGGGGCGCGTTGTTGTTATTGATTCCTCTTTGTGTGCCGCCAGTCACGTTCAGGACGTTATCACCCATGGTGTTGGGGCGGGTGTTCCTCGTCCTC | 4 |

| | | | |
|---|---|---|---|
| wt-94980-95100 | 3645 | GGTTTTCGGCCCTGCCTCCGGCCCCATCCCGCGAGCCCCAGCAGACATGCTCTCAGTTAACGAGCAGAGGGAATGAAAGCTCACGGCGAAACTTGGGCCAGCCCGGGGGGACCTCCCCTG | 8 |
| wt-95100-95220 | 3646 | CTGTTCCCCCAGTTTGCCCCATCGTTTCCCTGACGGCCTCAGGGGCCAAGCAAAACCGCGGGGGCATGGGATCCTTGCACTTAGCCAAGCCTGAGGAAACCTCCCCCGCCGTCTCCCCAG | 4 |
| wt-95220-95340 | 3647 | TATGCCCCATCGCTTCCCCAGCGGCCTCCAGGTCCAAGCAGCACTGCGGGGTCACTGGATCCTCACAGGCCGCACCCAGCTTTTCTTCCGTTGCCCCAGTAGCATCTCTGTCTGGTGACC | 4 |
| wt-95340-95460 | 3648 | TTGAAGAGGAAGAGGAGGGGTCCCGAGAATCCCCATCCCTACCGTCCAGCAAAAAGGGGGACGAGGAATTTGAGGCCTGGCTTGAGGCTCAGGACGCAAATCTTGAGGATGTTCAGCGGG | 4 |
| wt-95460-95580 | 3649 | AGTTTTCCGGGCTGCGAGTAATTGGTGATGAGGACGAGGATGGTTCGGAGGATGGGGAATTTTCAGACCTGGATCTGTCTGACAGCGACCATGAAGGGGATGAGGGTGGGGGGGCTGTTG | 4 |
| wt-95580-95700 | 3650 | GAGGGGGCAGGAGTCTGCACTCCCTGTATTCACTGAGCGTCGTCTAATAAAGATGTCTATTGATCTCTTTTAGTGTGAATCATGTCTGACGAGGGGCCAGGTACAGGACCTGGAAATGGC | 4 |
| wt-95700-95820 | 3651 | CTAGGAGAGAAGGGAGACACATCTGGACCAGAAGGCTCCGGCGGCAGTGGACCTCAAAGAAGAGGGGGTGATAACCATGGACGAGGACGGGGAAGAGGACGAGGACGAGGAGGCGGAAGA | 4 |
| wt-95820-95940 | 3652 | CCAGGAGCCCCGGGCGGCTCAGGATCAGGGCCAAGACATAGAGATGGTGTCCGGAGACCCCAAAAACGTCCAAGTTGCATTGGCTGCAAAGGGACCCACGGTGGAACAGGAGCAGGAGCA | 4 |
| wt-95940-96060 | 3653 | GGAGCGGGAGGGGCAGGAGCAGGAGGGGCAGGAGCAGGAGGAGGGGCAGGAGCAGGAGGAGGGGCAGGAGGGGCAGGAGGGGCAGGAGGGGCAGGAGCAGGAGGAGGGGCAGGAGCAGGA | 8 |
| wt-96060-96180 | 3654 | GGAGGGGCAGGAGGGGCAGGAGGGGCAGGAGCAGGAGGAGGGGCAGGAGCAGGAGGAGGGGCAGGAGGGGCAGGAGCAGGAGGAGGGGCAGGAGGGGCAGGAGGGGCAGGAGCAGGAGGA | 8 |
| wt-96180-96300 | 3655 | GGGGCAGGAGCAGGAGGAGGGGCAGGAGGGGCAGGAGCAGGAGGAGGGGCAGGAGGGGCAGGAGGGGCAGGAGCAGGAGGAGGGGCAGGAGCAGGAGGGGCAGGAGGGGCAGGAGGGGCA | 8 |
| wt-96540-96660 | 3656 | GGGGCAGGAGCAGGAGGGGCAGGAGGGGCAGGAGCAGGAGGGGCAGGAGGGGCAGGAGCAGGAGGAGGGGCAGGAGCAGGAGGGGCAGGAGCAGGAGGTGGAGGCCGGGGTCGAGGAGGC | 8 |
| wt-96660-96780 | 3657 | AGTGGAGGCCGGGGTCGAGGAGGTAGTGGAGGCCGGGGTCGAGGAGGTAGTGGAGGCCGCCGGGGTAGAGGACGTGAAAGAGCCAGGGGGGGAAGTCGTGAAAGAGCCAGGGGGAGAGGT | 8 |
| wt-96780-96900 | 3658 | CGTGGACGTGGAGAAAAGAGGCCCAGGAGTCCCAGTAGTCAGTCATCATCATCCGGGTCTCCACCGCGCAGGCCCCCTCCAGGTAGAAGGCCATTTTTCCACCCTGTAGGGGAAGCCGAT | 4 |
| wt-96900-97020 | 3659 | TATTTTGAATACCACCAAGAAGGTGGCCCAGATGGTGAGCCTGACGTGCCCCCGGGAGCGATAGAGCAGGGCCCCGCAGATGACCCAGGAGAAGGCCCAAGCACTGGACCCCGGGGTCAG | 4 |
| wt-97020-97140 | 3660 | GGTGATGGAGGCAGGCGCAAAAAAGGAGGGTGGTTTGGAAAGCATCGTGGTCAAGGAGGTTCCAACCCGAAATTTGAGAACATTGCAGAAGGTTTAAGAGCTCTCCTGGCTAGGAGTCAC | 4 |
| wt-97140-97260 | 3661 | GTAGAAAGGACTACCGACGAAGGAACTTGGGTCGCCGGTGTGTTCGTATATGGAGGTAGTAAGACCTCCCTTTACAACCTAAGGCGAGGAACTGCCCTTGCTATTCCACAATGTCGTCTT | 4 |
| wt-97260-97380 | 3662 | ACACCATTGAGTCGTCTCCCCTTTGGAATGGCCCCTGGACCCGGCCCACAACCTGGCCCGCTAAGGGAGTCCATTGTCTGTTATTTCATGGTCTTTTTACAAACTCATATATTTGCTGAG | 4 |
| wt-97380-97500 | 3663 | GTTTTGAAGGATGCGATTAAGGACCTTGTTATGACAAAGCCCGCTCCTACCTGCAATATCAGGGTGACTGTGTGCAGCTTTGACGATGGAGTAGATTTGCCTCCCTGGTTTCCACCTATG | 4 |
| wt-97500-97620 | 3664 | GTGGAAGGGGCTGCCGCGGAGGGTGATGACGGAGATGACGGAGATGAAGGAGGTGATGGAGATGAGGGTGAGGAAGGGCAGGAGTGATGTAACTTGTTAGGAGACGCCCTCAATCGTATT | 4 |
| wt-97620-97740 | 3665 | AAAAGCCGTGTATTCCCCCGCACTAAAGAATAAATCCCCAGTAGACATCATGCGTGCTGTTGGTGTATTTCTGGCCATCTGTCTTGTCACCATTTTCGTCCTCCCAACATGGGGCAATTG | 4 |
| wt-97740-97860 | 3666 | GGCATACCCATGTTGTCACGTCACTCAGCTCCGCGCTCAACACCTTCTCGCGTTGGAAAACATTAGCGACATTTACCTGGTGAGCAATCAGACATGCGACGGCTTTAGCCTGGCCTCCTT | 4 |
| wt-97860-97980 | 3667 | AAATTCACCTAAGAATGGGAGCAACCAGCTGGTCATCAGCCGCTGCGCAAACGGACTCAACGTGGTCTCCTTCTTTATCTCCATCCTGAAGCGAAGCAGCTCCGCCCTCACGGGCCATCT | 4 |
| wt-97980-98100 | 3668 | CCGTGAGTTGTTAACCACCCTGGAGACTCTTTACGGTTCATTCTCAGTGGAAGACCTGTTTGGTGCCAACTTAAACAGATACGCATGGCATCGCGGGGGCTAGACCTCTGGCTGGATGAG | 4 |
| wt-98100-98220 | 3669 | CACGTGTGGAAGAGGAAACAGGAGATTGGTGTGAAAGGAGAAAATCTGCTTCTCCCCGACTTATGGCTAGATTTCCTACAACTCAGCCCCATCTTCCAGCGCAAGCTTGCTGCCGTTATT | 4 |
| wt-98220-98340 | 3670 | GCCTGTGTCCGACGCCTGCGGACTCAGGCCACCGTCTACCCAGAGGAGGACATGTGCATGGCCTGGGCCCGCTTTTGCGACCCCTCTGATATTAAGGTGGTTATTTTGGGCCAGGACCCC | 4 |
| wt-98340-98460 | 3671 | TATCACGGGGGTCAAGCAAACGGCCTGGCATTCAGCGTCGCATACGGCTTTCCAGTTCCCCCAGCCTGAGGAACATCTACGCGGAGCTGCACCGGAGCCTGCCGGAGTTTTCTCCCCCA | 4 |
| wt-98460-98580 | 3672 | GATCACGGCTGTCTAGACGCGTGGGCCTCCCAGGGGGTGTTGCTACTCAACACCATCCTGACCGTGCAAAAGGGCAAGCCCGGCTCGCACGCAGACATTGGCTGGGCGTGGTTTACTGAC | 4 |
| wt-98580-98700 | 3673 | CACGTAATTTCATTGCTCTCTGAGCGGTTAAAAGCGTGCGTGTTTATGCTGTGGGGTGCGAAGGCGGGAGACAAAGCTTCACTAATCAACTCCAAGAAGCATCTGGTTCTGACCTCTCAG | 4 |
| wt-98700-98820 | 3674 | CATCCCTCTCCCCTGGCCCAGAACAGCACCCGAAAGAGTGCCCAGCAGAAGTTCCTGGGCAACAACCACTTTGTCCTCGCTAACAACTTTTTGCGTGAGAAGGGGCTCGGTGAGATAGAT | 4 |
| wt-98820-98940 | 3675 | TGGAGGCTGTAGAGGGGTCATCACTATGGCCATGTTTCTGAAGTCGCGTGGGGTCCGGTCTTGCAGGGACCGGCGCCTCTTGTCGGACGAGGAGGAAGAGACTTCACAGAGCAGCAGCTA | 4 |
| wt-98940-99060 | 3676 | CACTCTGGGGTCTCAGGCCTCCCAGTCTATCCAGGAGGAGGACGTGAGTGACACTGATGAGTCTGACTACTCAGATGAAGACGAGGAGATTGATTTGGAGGAAGAGTACCCCAGTGACGA | 4 |
| wt-99060-99180 | 3677 | AGACCCATCTGAGGGCAGTGATAGCGACCCCTCGTGGCATCCTTCAGATTCAGACGAGTCTGACTACAGCGAGAGCGACGAGGATGAAGCAACCCCCGGCTCTCAGGCCTCACGATCTTC | 4 |
| wt-99180-99300 | 3678 | AAGAGTCTCGCCATCTACCCAACAGTCTTCAGGTCTGACACCCACGCCTTCGTTCTCCCGACCACGCACCCGGGCACCTCCGAGGCCGCCGGCTCCCGCGCCGGTCAGGGGACGGGCCTC | 8 |
| wt-99300-99420 | 3679 | AGCACCTCCCAGGCCACCAGCCCCAGTTCAGCAATCCACCAAAGACAAGGGTCCCCATAGACCTACGCGACCTGTACTTAGAGGCCCAGCTCCACGCCGCCCCCCTCCACCTTCAAGTCC | 4 |
| wt-99420-99540 | 3680 | CAATACATACAATAAACACATGATGGAAACCACCCCCCCATTAAGGGCAATAACAACTACAATTGGCCATGGCTGTAAATAAAATGTCATAACCTGGAGTCTGCATGTCTGTTGTTTTA | 4 |
| wt-99540-99660 | 3681 | TTCAGTAAACCAGTAGTGCGCGTGAGTTCTTTAGGGCATCCACGATGTAGCCGCTCGCGGGGTTCCCCTCCCCAGTGATCATCTCGGATAGGGGATTCCTGTCCATGACCACGCAATTAG | 4 |
| wt-99660-99780 | 3682 | AGTGCCGGGCCCGGGACAGCGCCACATACACATGGCCGGGTTTGATGTTTCTGTGGCTGCCGAAGCAGATGGCGACTTTGTTTAGGGACAGACCCTGGGCCTTGGCTATGGTCATGGCCA | 4 |
| wt-99780-99900 | 3683 | GCTTTGAGCTAATGCCATAGTCACGGATGCTGCAGAGGTTCAGGGACTTGTCCTCTATCGTCTCATACAGTTTGTTAGTATTGTGTTCCAGGCAGCACACGAAGCCTGCCTCATCCTTGA | 4 |
| wt-99900-100020 | 3684 | CCATTAGCCTGGGCATGCGTGAACTGCCAGCGTCCTGAGGCTGCTGCTTTCCTCGGATGCCAAAGAAGACGTTGAGATGCGTGTAGCCCAGAAGCGTGTAGTTCTCGGTGGTGGAGGCGT | 4 |
| wt-100020-100140 | 3685 | AGTCCAGGAGGCCGTGAAGGAGAGGCTCGTCTGAGGTGAACTCTATGTTGTCGCGAATCAGCATGTTGTTGGTAAATGTGCAGAAGGGGAGGTCCCTGAACTCCCTTCCGCCATAGCGGA | 4 |
| wt-100140-100260 | 3686 | CGGCCACATCCAGGCATTGCCTGAAATAGGCCCTGAGGTCATTATATATGTTTAACAGGGAGCAGAGGGGGGCAGAATTTGCGGCCGGGGGAGCCAGTACTCGGGCATAGAAGACAGCGG | 4 |
| wt-100260-100380 | 3687 | CGGGGCTCCGCTCCCCATCCCAGGCAACCTCCAGCGGCAGTTCGCCCAGCTCCATCCCAGCAGTCACCTCCGGATCCCACGTACGCCCGGGCAGGGGCACAGCACCAAGCTCCGCCACGT | 8 |
| wt-100380-100500 | 3688 | ATTCCCCGTTTTCACAGAGAGAATGTCCTCCGTGGCTAAAAGCGTAGATGCCTCCGTAGATGAGTCGGGCCAGGAAGCTGTAGACATACTCGGGCTGCTCATGCCCGTGGGCCTCCACGA | 4 |
| wt-100500-100620 | 3689 | AGCTGTCCGCCTCGAGCGTGTCCATAAAGTCCCCGAAGGTGCCGGTATAGCCACAGATGGACTTTTTGGTCTTGCAGTTGACCGACACCGAGCTGTGCTTGACGTAGGTGACATTGTAGG | 4 |
| wt-100620-100740 | 3690 | TGACCTTGACCCGTTCTTCGTCCTGCTCGGTGCCCACCGGGACCATGTCTTGGTCGGCGAACTGCGAGTAGTTACCGAGGCGTGCATAATTCTTTTGGAGCCAGGTGTGGGCCGTGAGGC | 4 |
| wt-100740-100860 | 3691 | CCGGAAGCCCGACCAGGGTCTTGTACTGGGCCAGGGGATCGAGGAAGACCTCGCACTCCACCGGGCAGGTAAACATGGTCACCCCGCCCCCATCTCCCCCGGTTCCCCGCGCGGCACGCC | 8 |
| wt-100860-100980 | 3692 | CCTGCCCGGCAGTCTTGAGCGTGGCATGGAGGGTGGTGAGGAAGGTCTTGACCTCGGCGTGGGAGAGGAAGAGCCGGGTCCAGCCCACGTACTGCGCGGGGTCCATTATGGCCGCCCTGG | 8 |
| wt-100980-101100 | 3693 | GGACGACGAAGCGGTCGACGTAGGCCAGGATGTCCGGCGAGAGCTCGAGGCCGTACTCGAGGGTCTTCATGAGGTGTCCAAACTGGACGTCGGTGCAGCGCTTGTTGTTGATGAAGAGGG | 4 |
| wt-101100-101220 | 3694 | CCCAGTTGCGGGCCACGTCCACGTAGGTCGCGGCCCTGGGGTTGCCCACCAGGAAGGTGAGGATGTTGTCGCACTCGCGAATCTTGTTTACCTGGGTCTCGTGGCTAAAGGAGGACTGAA | 4 |
| wt-101220-101340 | 3695 | AGGCGTCTGTCTGGGTGGGAGAGCCCACGCAGACGATGCAGGGAATGCGGCCCCGGCGGTAGAGTGGGGTACGCAGCCAGGCGTTGAAGAACCAGTAGCAAAAGACCACGGCTGTTAGAA | 4 |

| | | | |
|---|---|---|---|
| wt-101340-101460 | 3696 | TGTGCACGGAAAGCGTTCCAGCTTCGTCCACCACGATCACATTGGTGGTCCATAGCTGCCCCTGGTGCATGTCTCTCAGGACCTCAAAGGCGGGGCCAGAGACTCCCGAGTAGAGCCCCC | 4 |
| wt-101460-101580 | 3697 | TGGGCTTGGTTCGCCTGAACTCGGCGGCAATGTCGGAGAGTACCGGCCAGTATTTGGCCAGGTCCCGCCGCTGGAGTTCCTCTAGGGCGGCGTCCGTAGAGCGACCATGACTGCTGACCC | 4 |
| wt-101580-101700 | 3698 | GCTGCGTCATATTTATGTGGCGGCTCTTGAACCCAAAGGCGCTATAGACGGTTGGGCAGTAGGCTCGGAGTGTCTGGGAGAGGTTCTGTGCGGCCACGGTTGTGGCTCCCGTGACCAGGC | 4 |
| wt-101700-101820 | 3699 | AGTCCATCGTGTGGTGGAGGCAGCTAACGCTGGTGCTCTTGCCAGCCCCCGCCGTTCCCGTAATTACATAGGCTGAAAAGGGCAGGAAGGGGGGCTCCGAGAGCTCCGGGTCAAACTCGG | 4 |
| wt-101820-101940 | 3700 | GGGAGAACGTCTCCATATCCGGGAGTTGTTGGACGCGGCGCCTAGCCAGGGTCCCTATCCTCCTGACTATACGCCTCACGGAGGCGTCTGAGGTCATGTTCAACATGAACGTGGACGAGA | 4 |
| wt-101940-102060 | 3701 | GCGCCTCTGGCGCCCTCGGCTCCTCGGCCATTCCTGTTCACCCCACGCCGGCCTCGGTCCGACTTTTTGAGATCCTGCAGGGAAAGTACGCCTACGTCCAGGGACAGACCATCTACGCCA | 4 |
| wt-102060-102180 | 3702 | ACCTCCGCAACCCCGGAGTCTTCTCGAGGCAGGTGTTTACCCATTTGTTTAAACGAGCCATCTCTCATTGCACGTACGATGACGTGCTACATGACTGGAACAAGTTCGAGGCCTGCATCC | 4 |
| wt-102180-102300 | 3703 | AGAAGCGATGGCCGAGCGATGACTCGTGTGCGAGCCGGTTTCGTGAGTCCACCTTCGAGTCGTGGTCCACGACCATGAAGCTGACCGTGCGTGACCTGCTGACCACCAACATCTACCGAG | 4 |
| wt-102300-102420 | 3704 | TGCTACACAGCCGCTCCGTGCTCTCCTATGAGCGTTATGTGGACTGGATCTGCGCCACCGGCATGGTGCCCGCCGTTAAGAAGCCCATAACCCAAGAGCTCCACTCCAAGATAAAGAGCC | 4 |
| wt-102420-102540 | 3705 | TGAGGGACAGGTGCGTCTGTCGGGAATTGGGGCACGAGAGGACCATCAGGAGTATCGGGACGGAATTATATGAGGCAACGAAGGAAATAATAGAGTCGCTCAACTCCACGTTCATCCCCC | 4 |
| wt-102540-102660 | 3706 | AGTTTACGGAGGTGACCATCGAGTACCTTCCGAGGAGCGACGAGTATGTGGCCTACTACTGTGGCCGCCGCATCAGGCTGCATGTGCTCTTCCCCCCGGCCATCTTTGCCGGAACGGTGA | 4 |
| wt-102660-102780 | 3707 | CCTTCGACAGCCCGGTGCAGCGCCTCTACCAGAACATTTTCATGTGCTACCGCACGCTGGAGCATGCCAAGATCTGCCAGCTCCTGAACACGGCCCCTCTCAAGGCCATCGTGGGCCACG | 4 |
| wt-102780-102900 | 3708 | GGGGGCGAGACATGTACAAGGACATCCTGGCCCATCTGGAGCAGAACTCACAGCGCAAGGACCCCAAGAAGGAGCTGCTGAACCTGCTGGTCAAGCTCTCGGAGAACAAGACCATCAGCG | 4 |
| wt-102900-103020 | 3709 | GGGTCACGGACGTGGTGGAGGAGTTCATAACGGATGCCTCCAACAACCTGGTGGACCGCAACCGTCTATTTGGCCAGCCCGGGGAGACAGCTGCACAGGGCCTAAAGAAAAAGGTCTCCA | 4 |
| wt-103020-103140 | 3710 | ACACGGTGGTCAAGTGTCTGACTGATCAGATAAACGAGCAATTTGACCAGATTAATGGCCTAGAGAAGGAGAGGGAGCTCTATCTAAAGAAGATCCGCTCCATGGAGTCTCAGCTGCAGG | 4 |
| wt-103140-103260 | 3711 | CCTCCCTGGGTCCCGGCGGCAACAACCCAGCGGCGTCAGCCCCCGCCGCAGTTGCGGCAGAAGCCGCGTCTGTAGATATACTGACGGGCAGCACCGCCTCCGCAATCGAAAAGCTGTTCA | 8 |
| wt-103260-103380 | 3712 | ACTCCCCGTCCGCCAGCCTGGGTGCCAGGGTGTCTGGTCACAATGAAAGCATCCTAAACAGTTTCGTTTCTCAATACATCCCCCCTTCGCGGGAAATGACTAAGGATCTGACTGAACTTT | 4 |
| wt-103380-103500 | 3713 | GGGAAAGCGAGCTGTTTAACACCTTCAAGTTAACACCCGTGGTTGATAATCAGGGGCAGCGTCTCTACGTCAGATACTCGTCAGACACGATCTCTATATTATTGGGCCCCTTCACCTATC | 4 |
| wt-103500-103620 | 3714 | TGGTGGCAGAGCTTTCACCGGTGGAACTCGTGACAGATGTCTACGCCACCCTAGGCATCGTGGAGATCATCGACGAGCTCTACCGGAGCAGTCGCCTGGCCATCTACATCGAGGACCTCG | 4 |
| wt-103620-103740 | 3715 | GTCGAAAATACTGCCCCGCGAGCGCGACCGGGGGAGATCATGGCATCCGGCAAGCACCATCAGCCCGGGGGGACACGGAGCCTGACCATGCAAAAAGTAAGCCTGCGCGTGACCCCCCGC | 8 |
| wt-103740-103860 | 3716 | CTGGTGCTGGAAGTTAACCGCCATAACGCCATCTGCGTGGCCACCAACGTCCCTGAGTTCTACAATGCCAGGGGGGACCTTAACATCCGAGACCTCCGGGCCCACGTCAAGGCCCGGATG | 4 |
| wt-103860-103980 | 3717 | ATCTCGTCCCAGTTTTGCGGCTACGTCCTCGTGAGTCTGCTGGACTCCGAGGACCAGGTCGACCACCTCAACATATTCCCCCACGTGTTCTCCGAGAGGATGATCCTGTACAAACCCAAC | 4 |
| wt-103980-104100 | 3718 | AATGTGAACCTTATGGAGATGTGCGCCCTGCTCTCGATGATTGAGAATGCCAAGAGCCCCTCCATAGGCCTCTGCCGGGAGGTGCTGGGTCGCCTGACCCTCTTGCACTCCAAGTGCAAC | 4 |
| wt-104100-104220 | 3719 | AATCTGGACTCTCTGTTTCTGTACAATGGGGCCAGGACGCTGCTGTCCACCCTGGTCAAGTACCACGACCTGGAGGAGGGGGCTGCCACCCCCGGGCCGTGGAATGAGGGCCTGAGTCTC | 4 |
| wt-104220-104340 | 3720 | TTTAAGCTGCACAAGGAGCTGAAGCGCGCCCCATCCGAAGCCCGGGACCTCATGCAGAGCCTCTTTCTGACCTCGGGGAAGATGGGGTGCCTGGCCAGGTCACCCAAGGATTACTGCGCG | 4 |
| wt-104340-104460 | 3721 | GATCTAAACAAGGAGGAAGATGCCAACTCGGGCTTCACATTTAACCTGTTTTATCAAGATTCTTTATTGACCAAGCATTTCCAGTGCCAGACCGTCCTCCAGACCTTGAGACGCAAGTGC | 4 |
| wt-104460-104580 | 3722 | CTCGGGAGTGACACGGTCTCAAAAATAATTCCCTAGAATAAACTGAGAACAGTCATCAGTAAATCTGTCTCTCGCGTGATTTCCATAGGAATGGTGTAGCCGGGGTGGAGGGCCGATATC | 4 |
| wt-104580-104700 | 3723 | ACATCAAGCAGAAAGGCCATAATCTCTCGAAAGTAGGCGGTGGGGCTGAGACCATGCTCAGTGGCCGTCTGGCAGGGGGCCGGGCGCGCTCCGTCCTTGTCCAGGAGACACACGTGGCTT | 4 |
| wt-104700-104820 | 3724 | CCAGAGAGGCGCAGCCCAGCCCTCCGCAGCCGCTGAAGCCAGGCTCGCGGAAGAGCCCAAAACCTGTTTCGGCGCCGCCCGGGGGCCAGTCTCCGGGTCAGGTCGCGGACCAGGGTCAAC | 8 |
| wt-104820-104940 | 3725 | AGGTGGTCGTGGGATGGCGGGGCCTTGTCTGCCTCGGGTCTCGCCGCTAGTTGGTCCAGGGTCCAGGAGAAGGCTTCGTGCCAAACCAAAAAGGGCCCCGAGTGCTCCCTACATCCACCC | 4 |
| wt-105060-105180 | 3727 | CTGGCATCCTCCGAGTCTTTCACGCCTTGCAGGAAAGCCTAGGAGATACAGCAACAGAAAGCTATTAGCCGGTGGTTCCCCCACCATCATTCTTCCTGTTAACGGGAAGAATAAGAGTTG | 4 |
| wt-105180-105300 | 3728 | GGCAAACCCCGGGGGCCGCGCTCTCCCACCCAGCCCCGCTTCTCACCTGTGCTAGTGGCTCCTCTGAAGGATGGGCGGAGGTTGGTGCCACAAAGCCCAGGATGAACTCGTCTGCATAAG | 4 |
| wt-105300-105420 | 3729 | CCCAGGTCAGTCCTAGGTCAGCGGCCGCGTGTAGGAGAACCCGGGTGACGGCGGTGTAGAGGCCCCCGAGTGCCCGTCGCGTGTCTGAGGTGCCATAGCGGTGAAGGGCCCGCAGCCAGG | 8 |
| wt-105420-105540 | 3730 | TTTGCGCGTCCCGCGCCTGCCCTCCGCCATCAGGCGTTCCCACGGGGGCGCCCCTGGCAGAGAGGTGGCAGCGGGCCAATTCGTAGAGCCACCAAGTGGCATCAGCCTCAAGGATGGCTG | 8 |
| wt-105540-105660 | 3731 | TGGCCTCCGCGCGCCCGACCACCGTCGTCTCGTCCTCCCCCCCTCCCTCGCCGCCTTCCCGCGTGCAAACGTGGCGAGGGTTAATCTCCTTTCGGGTCGGGGGCCAGATTTGTTGTAGGA | 8 |
| wt-105660-105780 | 3732 | GCAGCGAGCCGCGTCGTTGCCCTGACCGCGCGTCGAGGCCCAGGAGGGCGTCTGCCAGGGGCGTCCCAGAGACTCCCAGGTTCAGGTCCAGTAGCAGGAGACCCTCGCTGTGTGGCGCCC | 8 |
| wt-105780-105900 | 3733 | GGTGCCAGAAGGCCGGCCTCGCCCGTCCCACATAATGGATGGGCAGGAAGGGAAAGCCCGGGACATAGGGCTGGAAATCTGAGCCCCCTGGGCAGAGTTCGGGGTCCAGGAGGTAGAAGA | 4 |
| wt-105900-106020 | 3734 | TGGGCTTGGTGCCTCTGTGGTTGGCGTAGCAGGAGGCATAGATACTGCGGAGGAAGGCGTAGAGCCCGCCCCCGGCCATACTCCAAGAGTTGACAAGCCAGGACTCGAATCCCCCAGCCG | 4 |
| wt-106020-106140 | 3735 | GCTCAAGAATTTTCAGGCTGACGCGGTGCCGTCGGGCGTCCCCACCACGGCCGGTGGCCCCGTCGGACGACACCAGATCTACTTCATAAGTGACCGGTCGCAGGATGTCCCTAAAGGGGA | 4 |
| wt-106140-106260 | 3736 | CGGGAGAGGGGTCGTCGGGAGTCTCGGTGGAATAGGTGAAAACATCCCCACGCGGTGTCCTGATGTATACGTCCAACTGTCCGGGAGACTCAGAGTGCCTCTGAGCATGGGGGCATGTCT | 4 |
| wt-106260-106380 | 3737 | GTTCCCCCTCCATCTCGGACCCGAAGCCATCAACAGGTGGGGGTTGTTGGTCCCGCCCATCATCCCCCGAGCAGCTTTGGCAGACCACCTGTGCTGGAAAGAGAGGCTGGAAGATGAGGC | 4 |
| wt-106380-106500 | 3738 | CCTGCTCATCCTCCACCCTGGCGGCGGACAAGAGTCTGCGGTCTCGGGTTCTAAATGAAAGGTCAAATAGGTCCTTCTCGGCGGCATCGGCGAGCATAGCAATGAGCCCCCCGCTGCGCC | 4 |
| wt-106500-106620 | 3739 | TGAGCTCCCGCTCCCATCGCAAAAAGTTGAGTTCGGTAGTCGAGGGCGCGTTGACCACGGGGGGCTCCAGGGAGCCTCCAAGCGGCGGCTGGCAGGCCTGCACCACGATCAGAGTCTCAA | 4 |
| wt-106620-106740 | 3740 | CGTCCTCCCTTTTGATGGGCACGATGCCCACGACCCAAATCGCCCACCACCGCCCTGCGGTCTGGGTAACATTATAAAAGGTAACCGAGCTGACGCGGGCCCTGACGCTCTCCGCGGGTG | 4 |
| wt-106740-106860 | 3741 | TTTCCATCATTGTTTGAGATCTGAGGAGGACTGGACCCTTTAAAACATCCGGTCACGCCCTTTGCAAATTATTTAAAAGGTGAATGCTCAACTGAGACCATCGCAATCATGAAGTCCTCC | 4 |
| wt-106860-106980 | 3742 | AAGAATGACACGTTCGTCTATAGAACGTGGGTCAAAACGCTTGTTGTGTACTTTGTGATGTTTGTCATGTCGGCGGTGGTCCCCATCACCGCCATGTTCCCCAACCTGGGGTACCCCTGC | 4 |
| wt-106980-107100 | 3743 | TACTTTAACGCACTGGTTGATTACGGGGCACTTAACCTGACCAATTACAACCTGGCCCACCACCTGACCCCCACGCTCTATCTGGAGCCGCCGGAGATGTTTGTCTACATCACACTGGTC | 4 |
| wt-107100-107220 | 3744 | TTTATCGCGGACTGCGTGGCTTTCATCTACTACGCCTGCGGCGAGGTGGCGCTAATCAAGGCCCGAAAAAAGGTCTCGGGTCTTACAGACCTCTCGGCCTGGGTCTCGGCAGTGGGCTCC | 4 |
| wt-107220-107340 | 3745 | CCAACCGTGCTGTTTTTGGCCATCCTCAAGCTCTGGTCCATACAGGTCTTCATCCAGGTCCTTTCCTACAAGCACGTCTTTCTCTCGGCCTTTGTGTACTTTTTGCACTTTCTGGCCTCA | 4 |
| wt-107340-107460 | 3746 | GTTCTACACGCCTGCGCATGTGTGACCCGCTTCTCCCCGGTCTGGGTGGTCAAGGCCCAGGACAACTCTATTCCCCAGGACACCTTCTTGTGGTGGGTGGTCTTCTACCTGAAGCCCGTA | 4 |
| wt-107460-107580 | 3747 | GTTACAAACCTGTACCTGGGGTGCCTTGCCCTGGAGACGCTGGTCTTCTCGCTCAGCGTGTTCCTGGCCCTGGGCAACAGCTTTTACTTTATGGTGGGGGACATGGTGCTGGGAGCCGTG | 4 |

| | | | | |
|---|---|---|---|---|
| wt-107580-107700 | 3748 | AACCTCTTCCTCATCCTGCCCATCTTCTGGTACATTCTGACGGAGGTGTGGCTGGCCTCCTTCCTGCGGCACAACTTTGGCTTCTACTGCGGCATGTTCATCGCCTCCATCATCCTGATC | | 4 |
| wt-107700-107820 | 3749 | CTGCCCTTGGTCAGGTACGAGGCCGTCTTTGTCTCCGCCAAGCTGCACACCACTGTGGCCATCAATGTGGCCATCATACCTATCCTGTGCTCGGTGGCCATGCTCATCAGGATATGCCGG | | 4 |
| wt-107820-107940 | 3750 | ATTTTCAAAAGCATGCGCCAGGGCACTGACTATGTCCCTGTCTCGGAGACGGTGGAACTGGAGCTAGAGTCAGAGCCGAGGCCTAGGCCCTCGCGCACGCCATCACCCGGGCGCAACCGC | | 4 |
| wt-107940-108060 | 3751 | CGCCGCTCTTCTACGTCCTCATCTTCCTCCAGGTCAACCAGGAGACAGAGGCCCGTCTCTACCCAAGCCCTCGTCTCCTCCGTTTTACCGATGACGACGGACAGCGAGGAGGAGATCTTC | | 4 |
| wt-108060-108180 | 3752 | CCCTAATGCAATAAAAACTTAAAACACTGAGGTTACTTTCCCGTCATTCTTTCGGGGGAACGAGGGGAGGCGGGAATTGGGTTAAGATAGGGGCGAAGGGTGGGGGTGGGTGCAAGAATT | | 4 |
| wt-108180-108300 | 3753 | GGGGCTGGGAATGGAGAGGGGAGTGGGCTAGGTGCCGACACCGGGGTGCCAAGATAATGGATTGAGTAAGCATGGGGCTCTGATCGGGTCCGCCGGGTTCTCAGGGGTGTAGTGGGTGGG | | 4 |
| wt-108300-108420 | 3754 | CATTGCATATTTTTGCCGCGGTGCTGTTGGGCCTTGGACTCGGGGTGATCATCCGTACCATCACCCGCACCCGCACCCCAGTCCACAGCCACCGGCCAAGGTCCTGGGCCTCCCACCACC | | 4 |
| wt-108420-108540 | 3755 | GTTATGCCTCCCCCTTTACCCATTAATTACAAGAGATGTTAGTTTGGTTTTTTATTTGGCAAAAACAGCAATTCATCATTTTCAGAGTCCTCATCATATTCGAGCCCCTCGTTGGTTTCC | | 4 |
| wt-108540-108660 | 3756 | CCGCAGGCCCTCCCTTCTTCGGCCGCTATTAGCTTAGTAGTCTCCAGGTTAAACTCCTCATAGTCATTATACAGGTTGATTATTCCCCCGTCCACGTCGCCTATGGAGTTGACTCGTCGT | | 4 |
| wt-108660-108780 | 3757 | CGGCAAAGAGACCAGAGGGCACCCATGGCGCGGTGTCAAAAGTATTGTCTGCGTACGCTTTCCAGGAGCCAGCCGCGGTGCTCAAGGTCTTACGGATGACAGAGTCCGGCAGGACCACGG | | 4 |
| wt-108780-108900 | 3758 | GTGTCACCAGCACCGCCACGGGAATCTCCACCGAGGCGTCCAGAAGCAGGTCTGAGCCGAGCGTGCAGGTCGCCGGGTCTAGAGGCGACCGTTTTCGAAAGAAGGCCGTCACAATGTTCA | | 4 |
| wt-108900-109020 | 3759 | CCCGGGGTGAGCAGTCTCTCCCGGGCTTGCCACCCCCACTGTGGCGGACGTAGTCTCCAACAATTTTGTATTGGAGGAGCACCTGGTAGAAGTAGTTGTGCCGTGGATTGATGAAGATGT | | 4 |
| wt-109020-109140 | 3760 | TGACTGGGACCCGGTCTTTAATACCAATGCGCCCCGCATTTTCGCTTGGGTCCGTCATTACGTAGAGCATAGACTCCACCCCCCTGTTGGCAGCTAGGCTGTCTGCCACCAGGTCATGAC | | 4 |
| wt-109140-109260 | 3761 | CGGGGCCCAGTTTGCGCTTACGGACATCTTTAAGATTCCAGGCCTCATCCTGCGTCAACAGATAGTCACCCTCCGAGGGCAACCGCCCATCCGGGACGTACTCCACGGTAGGACGAGCTA | | 4 |
| wt-109260-109380 | 3762 | TAGAATTGATAAATCTGATAAATGACCTCTTGCATGGCCTCTTGTAAAGCGCAGTGTAGGATGGGTAGATGGGGTCAAATTCTGACTTGGAAAAGAGGTACTTGAAGCGGCACTTAATCT | | 4 |
| wt-109380-109500 | 3763 | CATAAATGCAGCTCCGGTCGGTGAACAGTATAAAGTCTCCCTGTGACTCCACATTGACGCAAAGATCCAGAGACACCCCAAAAATGCCATCCGTGGGACTAATCATAAAGCCAAATTGAC | | 4 |
| wt-109500-109620 | 3764 | GGTTGGCGGATGCGTCCCCGCAGATGAGCTTACAGACAATGTCCTTGACCGTGTCCTCACACCGCAGGCCAAAGGCCACAGGTCCCCCAAAGTAGTGATTTGTGGAGATGGGAGCTGGCT | | 4 |
| wt-109620-109740 | 3765 | CAAACACCTTGGTGGGTCCATTCTTAATGGTGGAGAGCAGCTTGGAAGAGGAAATTATGCCATTTCGCAATATGTCCCACATCAGGTTCTCAGACTGCCCCCTGGTCATGGACTCCACGT | | 4 |
| wt-109740-109860 | 3766 | ACGAGCAGAGAACAGTCCTCTGCTCGTCGGTGGCCTCCTGTAGCCCCCAGTAAATGGATTTCAGGGAGGGACCGTCCTTGCTGTCATTCTCTTGGACTAACGAGGAGACAAAGTCACAGA | | 4 |
| wt-109860-109980 | 3767 | AGCCAGTTTCACCAGAGAACTCTTGTATTTGTTTACAGAGGCAATAGAGATAGACAAAGCGCATGGCCGGCATCTGAGGTGGACGGTCAAGGTTACGGACAAAGGCCTCAGTCTCCGGAC | | 4 |
| wt-109980-110100 | 3768 | TGCGGAGGAAGCGGGCAAACGTGTAGGAGGTCATCTCCTCCATGGGATCCTCGAGCTCATCCACGTCGGCCATCTGGACCAAAGAAGTCGTCTGCCAAGAGTTCAGCTACCAGACCTGGA | | 4 |
| wt-110100-110220 | 3769 | AGATGAGGGTGCTCAAACCGTGGGCGACAGTTGAAGAAGTAGCTCTCCTTGAACCTCTTTTTAAGGCTCCGGCACCACTGCAAGAATTGACTCATATGCTCCGCCGTGACATCCACGCAC | | 4 |
| wt-110220-110340 | 3770 | GGACTCTCGCCACACGAGGTCAGGCCCATGTCTAAGTTCAGGTTCCACATCTGCGACAGCACCTCCAACAGCACCACCTTTGGGGCTGCAAATTGCAAAAAGTAGAGCGGGTCGGATCGG | | 4 |
| wt-110340-110460 | 3771 | TCAAATCCCATGTCAGGGTTGGGGTAGGGGATTTTGTGGGTGGAGTCAGCGAGGTGCATGATACCATAGAGCAGCGAGTAGCCGAGCGACTGCAGATCCAGGCGAAGGGCCGTCTGCGCC | | 4 |
| wt-110460-110580 | 3772 | CCCACGGGGCCACACGCCGAGGGGTCAGGGATGTGCCCAGCCCCCCTCAAGATGTAGCACTTGCTCAAAAGGCAGAGGGGCTTATAGGTGTCCTTGGCTATAGAAAATGGTTCCCTCTGG | | 4 |
| wt-110580-110700 | 3773 | CAATAGAGGCGATAGAGCTGCCGGCCCTTAGAAGACTTTAGCCGCACATCCAGCATCTTGTTGCGGTCGTGGAGGGAAGCAGTCCCATAATCAGTCAGGACCAGCCTACCCATGCCCCAC | | 4 |
| wt-110700-110820 | 3774 | ATGGTGTCTGTGAAATCCACCAGGATGTTGCTGGGGCTAATGTCCGAATGGAAGAGGCCGCAGTGCCGATTCAGAAAGTAAACGGCATCTTTGAGGCCCTGAAAGCCCCGCACCAGGGGC | | 4 |
| wt-110820-110940 | 3775 | TCAATACTACCATCATGCCAGTGGCCATAATCCTGGAGACTGCATCTGAACTGGGGCATAAACAGGGCGTGGCAGGACGTGCAGGCCGACAGGTAGTCCACCAGGGCCTTGTCCTGCCCA | | 4 |
| wt-110940-111060 | 3776 | TCCTCGGCCGTGGCCTTCCCAATCTGAATCATGTCACACACCATGAGCTCGTGATACAGCTCCGTCACAGAGTCATAGAGTTTGACCGTGGCATTATCTGCATGTGCATACACGGCCCCG | | 4 |
| wt-111060-111180 | 3777 | TAGCTCCCCCGCCCCAGCAGATACTCGCAGGTAATGGGGAGGTGATCACAGCGCGTCATGTTCTCCGGCAGCTTTACATAGAGGGTCTCCGTCATGTCATCAATGTTGGTCACCTTCAGG | | 4 |
| wt-111180-111300 | 3778 | TGTTTGTGCTGAAAGGTGAAGTAATCAATGACAGTCACCTTCCCCAAAAAGGCCTGGGTCTCTCGAGGGGGTTCTGGGGAGACACTCAACTCGCCACTGCTGGAGGAGTTCGTCGGGCTC | | 4 |
| wt-111300-111420 | 3779 | AACTCCGCAGCCATATTCACATCCATGTTCCTCAAATGGCTCGAGGGCCTGTCGCAGCTCGTCTCTGGCCTCAAGCTCCTGCTCACGGAGCTCCTCCACCCGCTCTAGCTGCTTGTAGTT | | 4 |
| wt-111420-111540 | 3780 | GATTTTTGGAAATTGAGTCTTGGTCGCGGTGACCACCCTCTGATAGGTAGAAATTAGCTGTTTGGACTCAAACGTCTCCCTTGCGTGGCGCAGGGACTCTAAGGCACCCCGAGCAGATGT | | 4 |
| wt-111540-111660 | 3781 | AAACTGTGTTTCAAACAGAGCGTGGTCCCTCCCAAATCTGTCACGTGCGCTCACAGCCGCTCTCTTTTCTACCGAGGCTCTTAGTTGCTGGGCCACCAGATCTCGCTTAGAACTACTCAT | | 4 |
| wt-111660-111780 | 3782 | CTTCATAAGTCACCATGTCCGCAACTATGGAGCCCAGATCATACGTGGGGTAGAGTACGGTAGTTCCAGTGGAGGCTTCCCGGTAATTTCCCACAGCGTCCACCATATATCTTTCTGCCT | | 4 |
| wt-111780-111900 | 3783 | CTCCCGTTAGAATTAGGCAAGGATCATACGTGTCCACCGGCCTTTTATACTGAGCGTTTAGGTTTTGTTTATGTAGCAAGCACAAAAGGCACACACGAGTGATGCAAAAGGGTTCCTGAG | | 4 |
| wt-111900-112020 | 3784 | GCAGCAGGCAGAGCTGTTTTGCCATTTTATTCAGGCGGCTAACGTCAAAGGGAGGAGCTATATCCTCACCCTTCCAGTCACGCACGTCCAAGTACAGGGCATACACACACCTGGTGAGGT | | 4 |
| wt-112020-112140 | 3785 | GTGCCAGGAATGCCTCTATGTTGGCACATGGTGTATAAACCGCAGTGGGTAGCAGAATAGGGCCCCTTTTGCCCCGTGCCGCAGCGTAAACGCAGTGACGCTCTTCGCAGTGGGACCTGG | | 4 |
| wt-112140-112260 | 3786 | GGCCGTAGAAGAGGGCCCACATCCAAGGGAGTGGGTCTTCAGGCACCAGGGAGGTCCAGGTTTGGGAGTGGGCCAATATTTGCAAGGCCTGACCTATAACCTCATCTTTGTTCCAGGCCA | | 4 |
| wt-112260-112380 | 3787 | GCGCAATTCGCATAAGGTCCCCATCAAACACCTCAAAACACAGACCCATGCCCATTTCAGGCTGAGAGGGCTCCATCCGGCTCGACCAACCTTGTCCACCAAACTGCCATTCTTCTGGTA | | 4 |
| wt-112380-112500 | 3788 | AACGGGGGTTGAGGGGCAAGAGCTCCAAAGCCAGGCTCGAGAAGTCATAGTCATCCTCGGCCACACGGCCGGAGCTCCGGGCCTCGTGCCAGGGCCTGTTGTCCTGGGGGAGGATATTGG | | 4 |
| wt-112500-112620 | 3789 | ACACGAGCAGGAAGCTCTTGAGTGGCGTCTCCACCAGCTTAAATTGCTCGGGCGTGTCCTGGCAGGCCTCCAGTGCCAGTTCCAGACACTGCCCATACCTGCGGGCGAGCATCGGGTCAT | | 4 |
| wt-112620-112740 | 3790 | CGGGCATATCGGCCTTGACCGCGTTGAACATGCTGTATGCCTCGCAGCGCGGCCGTCTGACCGAGAACCTAAGAAACGCCCTTCAGCAGGACAGCACCACGCAAGGCTGCCTGGGTGCCG | | 4 |
| wt-112740-112860 | 3791 | AGACCCCGAGTATTATGTACACAGGGGCCAAGTCAGACAGGTGGGCTCACCCTCTGGTGGGCACAATTCACGCCAGTAATTTATATTGCCCAATGCTTCGAGCATACTGCCGCCACTATG | | 4 |
| wt-112860-112980 | 3792 | GCCCCAGGCCCGTGTTTGTAGCTTCTGATGAATCATTACCCATGTTCGGTGCGAGCCCTGCCCTTCACACCCCAGTCCAGGTCCAGATGTGCCTACTACCAGAGCTACGCGACACGTTAC | | 4 |
| wt-112980-113100 | 3793 | AGCGCCTGCTGCCACCACCCAATCTTGAAGACTCCGAGGCCTTGACGGAATTCAAGACCAGCGTGTCCTCTGCCCGTGCCATCCTTGAGGACCCCAACTTTTTGGAGATGAGAGAGTTTG | | 4 |
| wt-113100-113220 | 3794 | TCACCAGCCTGGCCAGCTTCCTGAGTGGTCAGTACAAGCACAAGCCCGCCCGCCTAGAAGCATTCCAGAAACAAGTAGTGTTACATTCTTTTTATTTTCTGATCTCAATCAAATCTTTAG | | 4 |
| wt-113220-113340 | 3795 | AGATTACAGACACCATGTTTGACATCTTTCAAAGTGCTTTCGGATTGGAAGAAATGACGCTGGAGAAGCTGCACATTTTTAAGCAAAAAGCCAGCGTGTTTCTTATCCCCAGGCGCCACG | | 4 |
| wt-113340-113460 | 3796 | GCAAGACCTGGATAGTCGTGGCCATCATCAGCCTCATCCTCTCGAATCTCTCCAACGTGCAAATAGGCTACGTGGCTCACCAGAAACATGTCGCGTCCGCCGTTTTCACTGAAATTATTG | | 4 |
| wt-113460-113580 | 3797 | ACACCTTGACCAAGAGCTTCGACTCCAAGCGTGTAGAGGTCAACAAGGAGACCAGCACCATCACGTTTAGGCACAGTGGGAAAATCTCCAGCACCGTAATGTGTGCCACCTGCTTCAATA | | 4 |
| wt-113580-113700 | 3798 | AGAATGTAAGACCTGACGTTTCAGTACTTGGCAATTGTAGAGCATAGCCCGGCTGTAAAGGTCAGAAAATCGCAGCAGGGTCCAAGGTTGTGCTGTACATGGGACCTCTTTCCCATTAGC | | 4 |

| | | | |
|---|---|---|---|
| wt-113700-113820 | 3799 | AAGAACCCCCTGCAGGACACGTGACATGTCCGGGTGCATTTTGGGTGGGTTAAATCTCAGTCCCACCACAAAGGGGGCATCCTCCGGTTTGAACATCAGACCCAACAAAGCCCGATGCCC | 4 |
| wt-113820-113940 | 3800 | AGTTATGGGTACGTAGTCGTTGTTCAGGGCCGTGCATGGCAGCAGACAAGGACAGGTGCCAGATGTGCCTGGGCTATCGTCCTCCGTCCAGCCACGCAGGATGTTCACGTGGGCCCCGGC | 4 |
| wt-113940-114060 | 3801 | ACCATAGCATGTCACACATTCCCCGTTATCACATCTGGTTAGCAGGTTGATAAAATGGGTCAGTGATGGAAAGGTTGGCATATTGGGGCAGCACATCAGCATGTCCATGTTAACGAAAAA | 4 |
| wt-114060-114180 | 3802 | CATGTACAGGGCCCCTTCTGCATACCAGGCACCACCCCGTCCCAGTGGGATGATCTCCGAGGGTGTGATATCTTGCAGTTCTTCTACTGTTTTAACGGCGGTTGAGGTGGTAAAGACGTG | 4 |
| wt-114180-114300 | 3803 | GGCCGTGGTCAGATCTGTGCAGGTGACTACAGGGTTACCCCTAATCTCCACAGGCACCGCCTCACCCACTGCATCTGAGAATACCCCAAAGTACATGAGAGTCAGGCTGTGTGGCCCCTG | 4 |
| wt-114300-114420 | 3804 | GACTGCCTTAGTGAAGAGAACCTCGGGCCTGGCCACGGTGGCTAGGGTTCCATTGATGTAGACGGTCACATAGGTGGGCTTCTTCTTGGGCTTCAGCACAATGAGGGTAACATTCATGTA | 4 |
| wt-114420-114540 | 3805 | GGTTTTAGGAGGTCCGGCTATCTGAGGCACGTACACAGCTGACACGGCGGTTGTGGCCGTATAGACTTTCATCTGGGGCGTAGAGGCATCGCTCAGCACCCAGAGGCACTCCTTGTTGAG | 4 |
| wt-114540-114660 | 3806 | GAACTTGCGAAGCTGTTCCCGGCTACTGTTCGCGGCGGATGCCATGACGTGCCAGAATATATCCCCTCTCCTCGGGGGTGAGTGCCAATTGGCCTTTAATAACAAAGCCCCCAGGCAGCA | 4 |
| wt-114780-114900 | 3808 | TACCCCAGGCCGAGGCCCCTTCCTCATACAGCGGGTGGCTATCTATCCATAGGCAGGCATCCGGCGTCTTTGGTGCATTGGAGATAGTAGCTTTCACCCAACAACTTTCCCAACTAACCC | 4 |
| wt-114900-115020 | 3809 | GTGTCTGGACAGTGAAGAACGCTTCCCTGATCAGGTCTGAATTTTTATAGATACGGGAGTAGGAGGTGGGAATAACAACTGGGATTTCTTGTTGTGCTGTCCAGGCCTGCATGGCCAGTT | 4 |
| wt-115020-115140 | 3810 | TTTCCCTGAAGCTAGCAGAAATTCTGAGGGCCACTGAAATGAGGAAGCGAAACTCCCTCTCTGGAGCTCCCAAAATTGAAACCTCAGCAAGATCTGTTGCTGGGGAGGCATGGGTGACAG | 4 |
| wt-115140-115260 | 3811 | CTGTCATCCTGTGCAGTCTGCCCTGGGCACTCAGCTCTGGATATGTGACAACATAGAGAGCGTGGGGGCTAAAAATATGAGCAATTCCCCTGACCAGGGCCCTGGACTCACGAATGGCCC | 4 |
| wt-115260-115380 | 3812 | GACGGGTCTTAGAGAAAGAAACAGGCACCCTCGAGAGTGCCCCCGACCCGACCCCCACAGTGCCGCCAGTCCCTGCTCGGCCTCCGCCGCCTTCCCCACCGGCGCTGCCCCGGATGTTGC | 8 |
| wt-115380-115500 | 3813 | TGGGGTTCTCGAGGGCTGGGTGGTGCTTGGACACAGAGGTCTCAGCAGCCGCCTTGGTCTCGGCCCCGGCCCTAAGTCTGAGCCCCAGGCAAAGGGCCGGACTCCCAGCGTGGCCCAACC | 8 |
| wt-115500-115620 | 3814 | TCTGCTCCCCTCTATTCTCCTCTTGCGTTATCTCCAATAGAATTTGCTTGAGGTCATACGTTTTAGGGTGCTCGACCTGGGCCGCGGCCACCGGCATATGCTCTATACCCGCCCCTCCGG | 4 |
| wt-115620-115740 | 3815 | GGGGCCCAGGATCTATAGGTATGGGCTGCATAGCCGCAGCAGACTCCTGGACCCCAGAGGCCTCTCTGATAAGATGCCCGTCGGTCAGAGCCCTTTTGGCCCCCTCAAAGAGAGACAGGT | 4 |
| wt-115740-115860 | 3816 | AATAAATCTGTAGCTCCCCAACCAGCCCTCCTTCATCGTAAAATCGAAGGGCGGCCACGTGGAAGGGGTTGTAGAGCTCTGGAAGGCCCTCATCGCAGTACACTGGCACACTGGTAAACG | 4 |
| wt-115860-115980 | 3817 | TGCCCCGATGGCTAGGCCGTCCGGGCAGCATGCCCCGAGCAGCAAACACGCGGCAGACCCTCGTGAGACCCGTCCGGTCACTGAAGAGAGTCTGGCACCAGGCCCCCTCGCAGTTTGGCA | 8 |
| wt-115980-116100 | 3818 | CGCGATTGGGGCAAAGCTCTGCCATAACCGTGTCGGGAACAAATAGGTGCACGAGGAGGGGGGTCCCGAGGCCACTCAACACTTGGTTGTCAATGTGGACATCCATAGCTCTCTCATGCG | 4 |
| wt-116100-116220 | 3819 | TTTGGCTACAGCATCATAGCGCTTGTTTCTGGTGGATTTAAATAACAGGGCCCCGTAGACAGTCTTTTGTGAGTAAATAGAGATGATGACATGGATGTAGAGACTGAGGACCACATCCAC | 4 |
| wt-116220-116340 | 3820 | CACCTTCTCGGAGGAGGCCCCCCTAAACAGCATCAGGCAGCAAGGGAACACAAAGGAAACCAGGGCCGGGATGTGAGGCCTCAGCGCCCCCTCCTGATCAAAGAGGGCCTCGCTGACCCC | 4 |
| wt-116340-116460 | 3821 | GGAGATGACATTCTCATTCAGAAAGTAGTGATAGAGGTGATTGACCACAGTCTTAACCAGGCCCTGGACTTGTTCAGGCTCCCACTTGTCCCGCTGGTCCTGTGTGTCTTGTCGGATCTC | 4 |
| wt-116460-116580 | 3822 | GGTCCAGGGCCTCAGCGCCGGCTGGAAATGCGGCCCCATGTAGTTGCCTGTAAGGGCGCACACCACTCCCTCATGGGTCTCAATCAGGGTGCACTCGCTGGATCCATCACATACGTGGTA | 4 |
| wt-116580-116700 | 3823 | CTCGCCACAGCCCCAGCAGGCAAACACGGAGGCCATGCTCTCAGGTAACGGGAGATGGAACTCCAGCTTACTATACGAGCACAGGTGGCGAGGATTGGGCTCATCCGTGCCCCCCTCCCC | 4 |
| wt-116700-116820 | 3824 | CCGCGGGAGGCTCAATCGGCCTTGGTCTGACATTCCACCCCGGCCAGGTCCAGGAGGGTGCAAATATTCTCCAGGCGCTGCACCTCAGAGACCTCCTGCTCAAAGAGACCTCCCACCGCC | 4 |
| wt-116820-116940 | 3825 | ACGTAGACGCGGGCCACCGTCCGGGGAAGGTCAGTGGGGTCCCAGCTCAGCAATTCTCCAAATTCTCTCTCCCCAATAGTGCCTCGCTTCTTATCCTGTCTTTCAGAGCATCCGGGGGCA | 4 |
| wt-116940-117060 | 3826 | GACATTTCACCTCTTGTTTGTGGACGAGGCTAACTTTATCAAGAAGGAGGCCCTGCCGGCGATCCTGGGCTTTATGCTTCAGAAGGATGCCAAGATTATCTTCATCTCGTCTGTGAACTC | 4 |
| wt-117060-117180 | 3827 | GGCTGACCAGGCCACCAGCTTTCTTTATAAGCTGAAGGATGCTCAGGAGCGGCTGCTGAACGTGGTAAGTTATGTGTGTCAGGAGCATCGGCAAGATTTTGACATGCAGGACAGCATGGT | 4 |
| wt-117180-117300 | 3828 | CTCATGCCCCTGCTTTCGCCTGCACATCCCGTCCTACATCACCATGGACAGCAACATCCGAGCAACCACCAACCTCTTTCTGGACGGGGCCTTTAGCACCGAGCTGATGGGTGACACCTC | 4 |
| wt-117420-117540 | 3830 | CGTGTACGTTGATCCGGCCTATACCAACAACACATCCGCATCAGGCACCGGAATCGCCGCCGTGACTCACGACAGGGCGGACCCTAACAGGGTCATCGTCCTGGGCCTGGAACACTTCTT | 4 |
| wt-117660-117780 | 3832 | CAGCAGTCAGGACTCTGCGGTGGCCATTGCCTCAATCATTGGGGAATCCTGCCCCCTCCCCTGCGCCTTCGTGCACACCAAGGACAAGACGTCCAGCCTGCAGTGGCCCATGTACCTCCT | 4 |
| wt-117780-117900 | 3833 | GACTAATGAGAAGTCCAAGGCCTTTGAGAGGCTCATCTACGCAGTGAACACGGCCAGCCTTTCTGCCAGTCAGGTCACCGTCTCCAACACCATCCAGCTCTCCTTCGATCCGGTCCTCTA | 4 |
| wt-117900-118020 | 3834 | TCTCATCTCCCAGATCAGGGCCATCAAGCCCATCCCTCTCCGCGACGGTACCTACACCTACACCGGCAAGCAGCGCAACCTCTCTGACGACGTGCTGGTTGCGCTAGTCATGGCTCATTT | 4 |
| wt-118020-118140 | 3835 | TCTCGCAACAACACAGAAGCACACGTTCAAGAAAGTTCATTAAACTTTATTGACTACACCAGTCCCTTGTAAAGCGACGGGTCTCGCGTGACGGCATTCGTGAGCAGGGCTTCGTCCAGG | 4 |
| wt-118140-118260 | 3836 | GGCTTGTTCTTGGCGGACATCATTAGCCCAGCCGCAAATATCAGAATTAGCATCAGAAAAGTGAGCCCCACAAACACCAGTGTCCAGAGAGGAAGACCGTAAGATAAAGATGGCTGCCTC | 4 |
| wt-118260-118380 | 3837 | TCATCTGGAACGGTGGGAAGCTCAGCAGTTGTTTTTGTGGCATTGGACGTCCCTTTGGAGGACAGCGTGGGGGCCAAGGTGGTAGCGTTGGTAATACGGGTAGTAGCACTGGTGGTGGAG | 4 |
| wt-118380-118500 | 3838 | GAGGACCTGGTGGTGACATTGCTAGTCACACCCGTGGAGGTTCCTGTTCCGGCCTCGGTGGCAGTGATGTTCTGTGCAGTAACCTTAGTGGTGACATTGATGGTGGATGCGTTGGAAGTT | 4 |
| wt-118500-118620 | 3839 | GTTGGGACTGGTGTGACAGTTGTCCCAGTGAATGTCACCGTGGTTGTGTTGGTGCTCAGAATAGCAGTTGTGGTTATAGGGGCGCTAGTCGTGGTCAAGGTCGTAGACTGGTTTGTGCTA | 4 |
| wt-118620-118740 | 3840 | GGACCCGATGCCGACGGTGATGGTGTAGTCACAGCCGTTGTGCCTGTCACGTTCCCCGCCGAGGCCGTCGAACTGCCACTAGATGTCCAAATAAGGCTTGTCTCACAGATGAGTATCATG | 4 |
| wt-118740-118860 | 3841 | GCCATAACAGCGCCTGCCTTGTCTCTGGCGTGTGCCATCGCGTCTGGACGCAGAAGGCCTCCCGGCCTCTTTTATAGCTAGTCTCCACACCCAATACTCTACTGAACCATCACATACATG | 4 |
| wt-118860-118980 | 3842 | ACCTCCTCGAGGTATGCAGGGAATGAGCGGTCCGTGAGCCGGTCAACACGACATTGCTTCCGTTTCATGCCTCCAGCTGCCCCTGACCAGTTAGGACCCTTGACGGATGTCTTTAACGGC | 4 |
| wt-118980-119100 | 3843 | GCGGTGCAGTTGGTCACCAATGACGGCCTAAAGGCCAACACATCCTTGAAGCAGGGCGTAGGAATGGTACCAAACTCGGGGCCCACCCCATCAAAGACATAATATGTCTCATAGTGGCAG | 4 |
| wt-119100-119220 | 3844 | TGATGATGCATCACCACCACAGCACTCGCCAGGACCCTCTGCATATCTTGTACAAGGCGCCTTTCAACTCGGCCACTGGCTCTGGTGACGTTAAATGTCCTGTTCCTATTAGTCACAGCC | 4 |
| wt-119220-119340 | 3845 | TGTAGATTTGGGCACCCAGACTCAAAAAGTGCAGCTACATGAAGGGCAGCCGCCTCAAATCCACCATGACCCCCATGGCTGTCCGTGTTGTTGGGGTAATAAGTCACATTGTTAATGACC | 4 |
| wt-119340-119460 | 3846 | ACGGCCGGGATAAGGGTGTAAACCTTGCAGAATGGATTGGTCGGACACCCATAAGACAGGGGCACCCCAAAATCACGCCCCTTACCCCGAAGCACCTTGGCCCCCACCGGCATAAAGCTG | 4 |
| wt-119460-119580 | 3847 | GGCAAAAAGAGTGGGTTAAAACCAAAGGCGAGTAGGGCCAGGAACGCCAAATAGCAGCAGTAATAGATGAAAACAAAGCTCAGCATGAAACAGCGTGGAGGCTCAGCTAGGGTCTCTGCC | 4 |
| wt-119580-119700 | 3848 | TCTCCATCATAGACATCTTCCTTGAATCTCATTCTCTCACCGCATACCTCGCTCTTCATCCAGGAGGGGGCCATGGCTGCCATTCTACCAGTTAACGAGGAGAGAGAGAGAGTAGGTCCGCG | 4 |
| wt-119700-119820 | 3849 | GAAATTGGTGCCCCTCTCTGCCCTCCTGACGAGGCCATGGTGTCATCCATCTCCGCAGTCCGTTCTTCAGCTTTGGCATTGGTCCGGGTCCGGGTGGTCTGATTTTGATTCTGATCCTGG | 4 |
| wt-119820-119940 | 3850 | GTATTGGTCTTGGTCTCTCCTCCCCCATTGGCATGGATTGGCATAGGTGGGTGTGGCTCAGGCTCAGGTTCCGGCCCTGGGACGGCAGCAGCCGCCGGGACGGTGAAGTCGTGGAAGGTA | 4 |
| wt-119940-120060 | 3851 | GAGGCCCGTCCCTCCCGAGGTCGTGGGGCCGGAGCCTTATAAAAGACTTCCACCCTCTCCCCGCTGGCCAAGACACGCCGCTCGTGGACCACGCCATCTTCCTCCCGGCTGATTGTGTGG | 8 |
| wt-120060-120180 | 3852 | CTGACGGTGCCGTGTTCCACCGCCACTTGTTCATCGACCATGGTACCCCCTTTATCTTAACCAGCAAGTGGCCGTCAGGGTCTCTTGAGAGTATGCCGCTGTGGCCAAGCGAGGCCCCAA | 4 |

| | | | |
|---|---|---|---|
| wt-120180-120300 | 3853 | ATTAAATAGTGATGCCAAAGACTGTAGGTAGGTCATCATCACACGCATGCGTGATAAATCATCCGCCACTGACAGGTCATCCAGGTCTATCCGGGCTATCTCATCCGGCACCATTTCCTG | 4 |
| wt-120300-120420 | 3854 | GAAGAGATTCAAGAGGTCGTGATGCTCATGCCGGATAAGGCCTCGGACCAGGCGCATACTGGCCCTGGGCAGCAGGGTCACCATGATGCAAAAGTAGAGACTCAGATTGTCCAGCAGGGC | 4 |
| wt-120420-120540 | 3855 | CAAGCCAAGGGGCCCTGGCACCTCCGGGAGGGCCAACTCGTAGTGGTGCCCCAGGTATGAAACAGAGCCAAGATGCATGTGTACATCGAGCATGTCTGCGTTCCCGGGAGCCTGCATGAC | 4 |
| wt-120540-120660 | 3856 | AACCCGGGAGTACACGTTAAACAGGAGAATCTTCTGCAGCACCTCCTCTGCTATGGGCGTAGGCAGCACCATGGGGAAAACAATGTCCACATCATTGGACTCTAACTTCACGGTGGCATG | 4 |
| wt-120660-120780 | 3857 | CTCTCGTCCAAATACCGGGGGCATAACACTGAGGCTCCCGGTCCCATGCCACTGGAAAAAGGGCTGGTACTTGTTCTTAATGGCGTAGGTCTGACCTGGAACAATCTTGGTGAGTATCAA | 4 |
| wt-120780-120900 | 3858 | ACTGTCCACGCTAACCTCATCCAGCACGGCCAGGGTGCAATCAGACAGGTAGTTGTACATGGACACGTAGTCCGGGACCGTCTCTAGAGAGTACACCTGACCCAAGCCCAATCCCTGCAC | 4 |
| wt-120900-121020 | 3859 | ATTCTGCGTCCCGTGAGTGGAAGCCAGGGGTAAGATGCAGCCAATCCTCTGTTGCATCTTGGCAATCTCATCGGTATACAGACGAGAGGAGAGAGACACTACCACTTTCAAATCCATCTT | 4 |
| wt-121020-121140 | 3860 | TATTGACAATTATCAAAAAACCACCTTATTTCCAAACTTTAATATTCTTCGTACCGGCGCCACCTCTTCAATTATATAGTGTCCGTAATGGATGGGGGCGTGGGTCTGTTTGACAGACAT | 4 |
| wt-121140-121260 | 3861 | AAACTCATCGATGAGTGCCCGGGAGGAGGCTGAGAGTGCGGGGAATGCCTCCTGCAGAAAGCTGCAGGGCTGCTCCAGAAACACGTCAGTGCCAGCAATCACTACAAACTGCACCTCTGT | 4 |
| wt-121260-121380 | 3862 | GTTGCTGGTGGCTGGGTGCCCTCCAAGTCGCTGGCTGTACTCGTTGACCATGTTGTAGAGTCCCCTGTTGTTGCGCAGAAGCTCCTCCTTGTTGAAAAATGCCCGGCAGGGGCTGTAGAG | 4 |
| wt-121380-121500 | 3863 | GCCCGGGACGGCCGTCTGGCGATAGGAGGAGTTGTACATGATGTCACCCAGAGAACCCAGCTGAGATGCCCAGGGATTCACAGTGCTCCGGTATTCATAGGCGGCATCCGGGCGAGAATG | 4 |
| wt-121500-121620 | 3864 | GTCATAGATGAGCCCCTCGGCAACCTCCTGATTGTAGTTTTCACAGGAGACCACACAGGCGGCCCGTCCCCTTGGAGAGTTGGACTTTTGAAAATAAGCCACGTCTGCCGTGACCGGTGT | 4 |
| wt-121620-121740 | 3865 | TACGATAATCTCACAGGTGGCCTGCTGGCCGTGGCAGAGTCCTGGAGCTCCATTAACATTAGTCATACCTGCCAGGTATGTCCTGGGGTCCCGAAGCAGCGTCCCATTGCGCTGAGCGCC | 4 |
| wt-121740-121860 | 3866 | CACCTTGGCCTTGATGTAGTCATTGACTTGCTGGTTGCCAAAGGCCTCGGCCGGAAAGACGCTAAAGAAGTCTTGGGTGTGGATACCCATGTCAGTAGTGATGGCCGCCACCCTGGCCGG | 4 |
| wt-121860-121980 | 3867 | AGTCATGGTCGAGCTATAACTAAGCCCGGTGTCGATGGAGGCCATCTCGTGATGCACCTCAAAGGTTACCGCGTCCACCCTGGCCTCCCGGCGGCTAACATTTGGGGTCCCAATGAACAT | 4 |
| wt-121980-122100 | 3868 | GGATGTTGAGGCCCTGGAGCTAAACAATATGTTTTCAGAGAGGATCTCATCGGTCCTGACCACGGTCATGGCCACCCCTGGGTGGATCTTGAGCTTGGCCTGGGCAATATAGGCCATGGG | 4 |
| wt-122100-122220 | 3869 | GGACATCTTGATGTGCATGGCGGTCATTCCACTGATTGAAACGAGGGAAGGAAGACATTCGGCCGCGTATTTGCCCATGGGCGAGCGGTGCCACTCCCGGTACTCTGCAAAGAGCTGCTC | 4 |
| wt-122220-122340 | 3870 | TGGCCGGTTGAAGGCTTCCACGGCCCGCTGCTGAGGATTGCGCATAACAAAGGTGGCAACATCCTGGTGCATGGTGGCAGCCACTCGCGGGTCCCCGTAAAACATATGGAAAGGAATGGC | 4 |
| wt-122340-122460 | 3871 | GTGAAAGAGACACTGGGTGACGGCCCGGGTCCTCTCGGAGAAGGCAAAGGCCACCAGCCCGTTCACCAAAACAGTCTGCTCTGTCCGCTTGTCGGCGGGATTCGGGGCCAGCTGCTGCGT | 4 |
| wt-122460-122580 | 3872 | AACGTCATTGTCCACCGACACACGCACGGCACGGGTGAAAGTGGGGCAGGTCATGAATGAGGCGCTGAGGTCCCTGATCATGCCCACGGTGGGGCGGAGGTCGGAGATCTCCAGCAGATC | 4 |
| wt-122580-122700 | 3873 | CCTGAGCGTCCCATTCTCCAAATTGTCGAGGATGTCCTCGTCCCTGGTAAAATGGTGGCTGAAGGCTGGCCCGTTGTAGGCCAGGGTCTGGGCCACGTGCTGAAAGTCCACCCCGAGGCC | 4 |
| wt-122700-122820 | 3874 | GCACATGTGGGCATTGGTGCAGGTTGGGAGGAAAACGTAGTAAAAGATCTTTTCCAGCACATCCGCATGCCCCTCATCTACATAAGGGCCTAGGTGCAGACGGAAATCGTGGTCGTGGTC | 4 |
| wt-122820-122940 | 3875 | TCCGTTAACCCGGTAGCCGTACAAGGCCACAAATTGGGCAGCCATCTCATCCATGTTTCCAACCCTCTCAATAAACTGGGGCGCGGCCAGGGTGTCAGCGTAAACCTCATTTCCGATAAT | 4 |
| wt-122940-123060 | 3876 | AATCTGGGGGGCCCGGTCACTAACGGTGAGAAGATGGGTGAAAATGTCTGTGTAGGCCACCGGGGGGAGCAGGTTAGGGTCCAGGAGAGCGCAGACATACTGACCCACGCTCTCATCCCC | 4 |
| wt-123060-123180 | 3877 | CACAACATCTGACCCGGCCAGGCGCATCAGGGCCTGCTCTAGGGCTATAAGTTCCCCATAGATTTTTCTATACATGGAATAGGCCTCCTTGGAGATGGCGTTATTTCCCAGGTGGCGGCA | 4 |
| wt-123180-123300 | 3878 | GATGAACTTGATCATGGAAAAGCTGTTCACAAAGGCAAGCCTCCCTGACCGTTCCCAGTAGGTGTTGATGCACAGGGACACCAAAGGCACGTTCATGACAAACTTTTCCTCAAACCCGTG | 4 |
| wt-123300-123420 | 3879 | GATCATAGCCTCGACTACGTAGAAGAAGGCTGGATAGGCAGTGTCATAGGCAGTATCCTGCACAGTCTCAATAACGGCCTGATCCACCACGTGGGCCAGAGATGTGGCGGTCTCAAACTG | 4 |
| wt-123540-123660 | 3881 | ATAAAGGGGGTGCAGCTCCAGCCGCAGCAGGTCATAATTGGACGGGTGGAGGAAGTCTTCGGTGGGCAGCCCGCACTTGAGAGCTATATCTGTCACGGGGGCTGCATACTTGTTATCATA | 4 |
| wt-123660-123780 | 3882 | GAACTCGTCCACAATAACAAGCACATTCATGTGATTGGGCCTCCTGTGTTGCAGGGAGTAGGTCTCGCGCCTGTCTCGCGGGGCCGGGGCCGCGTTGAGGCTGTTTAGGGTATGGGCGGG | 4 |
| wt-123780-123900 | 3883 | TGTGTGGAGTCGGGGGTGACAGAGAACCTTGAGAGCATTCTGTAGGTTAAACGCGAGGAGAAGGTTATTCTTGTTTACGATCCATGCCTCCACCGGTAGCTGCTGTGTGGGGTTGTCCAG | 4 |
| wt-123900-124020 | 3884 | CATTTTGATGGCGGCGGAGGTCGTGTACTTGGGATTGGGCATAAACAGGCCCACTGGGAAATAGTAGCTGTACTGCATTCTTCTGTTGAGGGGGTATGGGGACTGAGTGTCATTGTACAT | 4 |
| wt-124020-124140 | 3885 | CTTTTGCAGGCTTTCCACGGCCACCGCGTGGTTGCCCAGCTTGATGACGGCGGCTGAGATCGGCACCCGGGGCTGATCCTCGACCCCTGCGGCCACAGCCGGCAGGTCAGACTTGGTGCT | 8 |
| wt-124140-124260 | 3886 | TCCGGCTTTTTCCGGTGAGTCCACGATCCTAGCCATGAAATGCTCAAACGTACGCATCACGCGCCCGTAGCTCACGGCAGTGACCAGGTTCTCCCCCCGTACCACAAAAGAAGCATAGCT | 4 |
| wt-124260-124380 | 3887 | CGAGGGCCCCATAATCTGGTTGTCGGCCTCCTCACCCAGGAAGGTCAAGAGCTGGCGCAGAACGTTGTCGGTGACAATAAACACCCCCCCCACTGGCTCTCCCCCCTTGGCGGTCGTGTA | 4 |
| wt-124380-124500 | 3888 | GGTACTGACCCCCTTGAGCACGCTCTCCCCGGACACGGCCGCTACCATCTCAGAGAGACGGCTTCGCACGTACTGAGAAAACCCGGAGCCCATGTTCTCGGCCCGGTCCAGGAAGAAGGA | 4 |
| wt-124500-124620 | 3889 | GTGCTCCAGCAGATGCCTCTTGAACATGGCAATGAGGTCAGACTTGACAGTCTTGGAGAACCCCCTCTCAGTGAAGGTGGGATCCGCCAGGGTCTGCAGGATAAACATGGGAGGGGCATG | 4 |
| wt-124620-124740 | 3890 | GCGAAGCTTCACACTCAGGACGGTGTTAATGAGGCCCCTCTCCAGGGCATCGACCCCAAACTGTAGGGCCGAGGCCACGGTCTTGACAGCCCCCACGTACTCTGCGTACTCGACCGGGGT | 4 |
| wt-124740-124860 | 3891 | CTCGGGGATACTATGCAGGATCTCCAGATCCAGCATGGACAGTTCCATTTCCGTACTAATGTGGTGTTTGTGGCAATTTTTGACCACAATGAATGTCCGCTGCTTGCTGGGTCTCCTTCC | 4 |
| wt-124860-124980 | 3892 | GTCCCCGTGAGCAATGGTGGGGACGGAGATTCGAAATTGAATCTTGCCATCCGTCATACGACTCAGGTCTTTGAATTCCGTGTTCACACAGGACACGGCCAGTGCCGTCTCCAGGAAGCG | 4 |
| wt-124980-125100 | 3893 | AACATATTGGATGGCGTTCGTGTAGACCCCGAGTAGCACCTCAAACTTGATGCCCGCCTCTCTGGCATCCTTGCCCACCAGCAGGTCAAAGCTATGAAACAACCCCTCAGCCGCTGACTG | 4 |
| wt-125100-125220 | 3894 | CCGCAGGTTCGAGAGCAGGTCGGCATCCACCGTCAGATAGGGGAAGGGTCTGTTTTCCACACCCTCATTTGAGGCCATGACACAAGGTAAGAGGGAGATGGGGGGAGGTCTCGAGGGCTT | 4 |
| wt-125220-125340 | 3895 | CTCTTCACAGCTGGGTCTCTTTTACGCCCTGGCCTGCAACCGCAGCCCACCCACACTTCCCGAGGATGCTACCCTTCTAATCAAATGGTTGGACACGGCCCTGGGCAGGGAGGCCACCTT | 4 |
| wt-125340-125460 | 3896 | TTACGCGTGTCGGGCTATGCGTCGGCTTCTACTCGGCGTTATCCGAATGAATGACTGCCAGGAGCTGCCACCCGGTTTAATAATTCTGAGTCCGGGCACCGTCCCTGGCCCCCTTGGAGT | 4 |
| wt-125460-125580 | 3897 | CCAGAGTCTGGAGCATACAGACTGCGAAATATGGTCCTCTGCCCACCCTGACCACGCTGCCCACCTCCCGGTGCCCAGGGTCATCACATACACCGACTGCCCGGGTTCCATAAACACGAG | 4 |
| wt-125580-125700 | 3898 | CTCAATGTTTCGCCTTATCATCCGCTACTTGTCTCATCACCAATTTGAGCGCTGCTTCGAGCAGTTCTGCCGCGTGGTCCCGCGTCGCTTCCTAGGGACCTGTAAGCGAAACTCTGCAAA | 4 |
| wt-125700-125820 | 3899 | GATGCTGGCTCATCTGAATCAGGTTACCAGGATCCCCCCCTGTCCGCCCTTCAGCGGGCGGGAGGCCAGACTCAAGTTCCACTTCTTCTCCTGGAGCACATTCATGCTGTCATGGCCAAA | 4 |
| wt-125820-125940 | 3900 | CAATGCCACACTCCGGGAGATCAGGACGAGGGCCGCCACCAACCTCACCCACCACCCACATCTAGTGGATACTCTGTACCACGCCTCTCCGCAGACCCCATTTCTGACACGCAGCGGTGC | 4 |
| wt-125940-126060 | 3901 | TCTATACCGCTTCGTCACCTGTTGCAACTGCACCCTGCCCAATATCTCCATCCAGCAGTGCAAGGCCGGGGACAGACCGGGGGACCTGGAGATCATTCTACAGAGTAACGGCGGAGGGAG | 4 |
| wt-126060-126180 | 3902 | GCCCGCGAGCTTCCAGTTCCCCTCCTCCCCAACTGGCTCCCTATTGCGATGCATAGTTGCTGCGTCCCTGCTGCCGGAGGTGTCCGTGGGGCACCAGGAGCTGTCTCCGCTGCGGTCCAG | 8 |
| wt-126180-126300 | 3903 | AAGCCAGGGAGGGCAGACGGATGTCAGGTCGGGCCCGGACCCGGCCCGGAGACTGGTGGCCCTCCTGCGAAGGGAAGATGGGGCACCTAAAGACCCCCCTCTGGGACCGTTTGGACACCC | 8 |
| wt-126300-126420 | 3904 | CCGGGGGCCCGGCCCGGCCAAGAGCGAAGACGAGGAGTCTGAGCGTCGAGACGCCCCTCCACCCCCGCTCGATTCCAGCTTCCAAGCTTCCCGGTTGGTGCCCGTGGGGCCTGGGTTTCG | 8 |

| | | | |
|---|---|---|---|
| wt-126420-126540 | 3905 | CCTGCTCGTGTTCAACACCAATCGGGTGATCAACACTAAATTGGTGTGCTCAGAGCCCCTGGTGAAGATGCGAGTTTGCAATGTCCCCCGCCTCATCAACAACTTTGTAGCCCGCAAGTA | 4 |
| wt-126540-126660 | 3906 | CGTGGTGAAAGAGACGGCGTTCACCGTCAGTCTATTCTTTACGGACGGGGTGGGGGCCAACCTAGCCATCAATGTCAATATCAGTGGCACCTATCTGAGCTTCCTATTGGCCATGACGTC | 4 |
| wt-126660-126780 | 3907 | ACTGCGGTGCTTCCTGCCTGTGGAGGCTATTTATCCCGCGGCCGTGTCAAACTGGAACTCGACTCTAGATCTCCATGGGCTGGAAAATCAGAGCCTAGTCAGAGAGAACCGAAGCGGGGT | 4 |
| wt-126780-126900 | 3908 | CTTTTGGACTACCAACTTTCCCTCGGTGGTGTCCTGCCGGGACGGTCTCAACGTGTCCTGGTTTAAGGCCGCAACTGCCACCATATCTCGAGTGCACGGGCAGACATTGGAGCAGCACCT | 4 |
| wt-126900-127020 | 3909 | GATCCGTGAAATCACCCCCATCGTGACGCATCGAGAGGCAAAAATCTCCCGGATTAAAAACCGGCTCTTTACCCTGCTAGAGCTACGCAATCGGAGTCAGATTCAAGTGCTGCACAAGCG | 4 |
| wt-127020-127140 | 3910 | TTTCCTGGAAGGCCTGCTAGACTGCGCCTCCCTCCTGCGCCTGGATCCCAGCTGTATCAACCGAATCGCCTCCGAGGGCCTGTTTGATTTCTCCAAGAGAAGCATCGCCCACTCCAAAAA | 4 |
| wt-127140-127260 | 3911 | CCGACACGAGTGCGCGCTTCTGGGTCACAGACATTCGGCGAACGTGACAAAGCTGGTGGTAAACGAGCGCAAGACCCGCCTGGACATACTGGGCCGTAACGCTAACTTTTTAACGAGGTG | 4 |
| wt-127260-127380 | 3912 | TAAGCATCAGGTTAATCTAAGACAGTCACCTATTTTCCTGACCCTCCTGAGGCACATCCGCCGACGTCTGGGCCTGGGCCGTGCTTCCGTAAAACGAGAGATTACCCTTCTCCTGGCCCA | 4 |
| wt-127380-127500 | 3913 | CCTGCGCAAAAAGACAGCCCCCATCCACTGCCGTGATGCTCAAGTGTAAGCAGCCCGGGGCCCGCTTCATTCACGGGGCCGTGCACCTGCCATCGGGACAGATTGTCTTCCACACCATCC | 4 |
| wt-127500-127620 | 3914 | ACAGCCCCACTCTTGCCTCGGCGCTGGGACTGCCTGGGGAAAATGTACCCATCCCGGCCCTCTTCCGTGCCTCGGGCCTCAACGTCCGTGAGAGCCTACCCATGACCAACATGAGGGCAC | 4 |
| wt-127620-127740 | 3915 | CGATCATCTCGCTGGCTCGCCTCATCCTGGCCCCCAACCCCTATATCCTAGAGGGACAGCTGACGGTGGGCATGACACAGGACAACGGCATTCCCGTGCTTTTTGCCAGGCCTGTCATTG | 4 |
| wt-127740-127860 | 3916 | AGGTAAAAAGCGGGCCTGAGTCCAACATTAAAGCCTCCTCGCAACTTATGATAGCAGAAGACTCCTGCCTGAATCAGATCGCCCCCTTTTCCGCATCAGAGCACCCCGCCTTCTCCATGG | 4 |
| wt-127860-127980 | 3917 | TTGAGTCCGTAAAACGAGTCCGGGTCGATGAGGGAGCAAACACCCGGCGCACCATCCGGGATATTCTGGAGATCCCCGTGACTGTGCTCTCATCCCTGCAACTGTCTCCCACCAAGTCCA | 4 |
| wt-128100-128220 | 3919 | CCGCGCAGGTCAGCAACGTGCTCATCGCCAACCGCTCCCACAACTCTCTAAGGCTGGTGCCCAATCCGGACTTGCTGCCTCTCCAGCATTTGTACCTCAAGCACGTAGTGCTAAAGAGTC | 4 |
| wt-128220-128340 | 3920 | TGAATCTGGAGAATATAGTGCAGGACTTTGAGGCCATCTTCACCTCCCCGTCTGATACCATCAGTGAGGCTGAAACCAAGGCCTTTGAGAAGCTGGTGGAGCAAGCCAAAAACACCGTAG | 4 |
| wt-128340-128460 | 3921 | AGAACATAGTCTTTTGCCTCAACAGCATCTGTTCCACCTCTACACTCCCAGATGTCGTCCCCGATGTCAATAACCCAAACATTAGCCTGGCTCTAGAGAAGTATTTTCTCATGTTCCCTC | 4 |
| wt-128460-128580 | 3922 | CCTCAGGCACCATTATGAGAAATGTCAGATTCGCCACCCCCATCGTCCGGCTCTTGTGCCAAGGGGCTGAGCTTGGCACCATGGCACAGTTTCTAGGAAAGTACATCAAGGTCAAGAAGG | 4 |
| wt-128580-128700 | 3923 | AAACTGGAATGTACACACTGGTCAAGCTTTATTACCTGCTGCGCATCTAAAGGAAAAACATAACAATCTTGTGAACCAGAAAGATACCCAGAGCAAAAGCAATAAAGTACAGGATTATTG | 4 |
| wt-128700-128820 | 3924 | CCAAAACAACGTGTGCTCTTTCTTCATACAGGCCCGCAATTTCCATGACAGTCCCGTTGGTGGTCAGCAGCAGATAGTGAACGTGGAGGTTGTCAAAATCAAAGTAGTTGGAGCTCAAGA | 4 |
| wt-128820-128940 | 3925 | TGGAGTTTTGGACTTCCTGGGAGGTGATGTAGGTTGTAGTTTCCAGGCCTTCCTTTTCATCATAACTGAGCAGGGCAAAGCCACAAAAAATGCAGGATTTCTGCGTCCTGGTAAAATTCT | 4 |
| wt-128940-129060 | 3926 | GGATCTTTGGAATCTGGCGGGGCTCCCCAGCCACAGCACCCTGCGAACATTTATTCATTATAACGGGGGAGAGAAAGAGAGAGCTGCTGAGATAGGTGGTGCTGGCCTCGTATAGCGCCG | 4 |
| wt-129060-129180 | 3927 | AGCCTCGGACCTCACGGTCACTAGAGATTATGAATGTCACATTGATGAGCGGGATAATCATCAGAACTTTGTCGAGCCTGTCCACGCATTTGTAGGCGGGGAGATGCCACGCATCCCTGT | 4 |
| wt-129180-129300 | 3928 | CTTCTCGCTCCAAAGAGAGCCGCCCAAGAAACCCATCCACAGCATTTGAAACGGCCGCCTGGTCCAGCGTTGCCTCCTGGGGGGCCATGCTCAGCAGCTTGTCTCGTGTGAGGTCAAATC | 4 |
| wt-129300-129420 | 3929 | GTAGGCTGAGGTAGCACGGTGAGAAGAGCCCGCTCTCTGTCCCCAGGGCTAGCCCCCGCAAAACCTCCCCAATCTCTAGGGCCGAGCACAGGGCGGTGGACAGCAGTTGGTATAGGGCAA | 4 |
| wt-129420-129540 | 3930 | GGTTGGGCCCCTGGGTAGTCACGTTCAGCCGCAACTCGCGTAGCACCACGTGGCTGCCGATAAACAGGGTCTCTCTCATCACGGTATGCAGGGGCTGGAAAAGGGGGTGGCGGTTGTAGG | 4 |
| wt-129540-129660 | 3931 | CCGAGAGAAGCACAGATGTGGCGCCTCCAATGAGGCCACTGTAAACCCCGGCCTTGGGGTAGCCGACGGTGGCTAACCTCAGCGCGTACTCCTGTTTCTCAGTAGTCAGGTGACCCAGCT | 4 |
| wt-129660-129780 | 3932 | CCTCCATCTTGACCGTGGCCATCAGCATGGCGGCCAAGCGCTCCAGCCCGTAGGATTGCATGCCCTTGACAGTGGCCCCATAACATATGCCGATGATGTCTTTCAGGACAGTCAGCTCAA | 4 |
| wt-129780-129900 | 3933 | AGAAGCTCTTGGCCAACCAGCGGAGGTCCACGCAGCCATTGCCAGTCTCACCCACAGCATGACCCACCTTAAAGAAGGCCACAGAAACCTCAAACATGGTAGTCAGCGTTTCCGTGTCCA | 4 |
| wt-129900-130020 | 3934 | GTTCCGGCTCCCGGCAGCCTCCCTTCATCTCCAGCAGGACCAGTTTCTGGAGAACGTAGCGAGCGTAGCTGGCGGCTGTCATGGTGACGGCTCGGGAAAACATATCCTTCAGGTTGGGTA | 4 |
| wt-130020-130140 | 3935 | CAAAGTAGTTGTGAAAGTTGGCATAATGCACAAAGGTTGTAACAATCACCAGGGAATAGTCCCCGCTTTGGGCACTGGTTAAGGATGGGTAACTAAAAGGCCCCCTCAGATCCGGCAGGT | 4 |
| wt-130140-130260 | 3936 | CCTTCGTCTTGCCAAAGACCAGGCTCAACACATGCTCATCTCCCTTCTCGGTCACTCGCTTGTAGGTGCCCATCAGAAATTTAGAAGTCATGGCCCCCGTGTACTGAAACTTGTCCCCGT | 4 |
| wt-130260-130380 | 3937 | TGATGGACAGGGCCACATAAGACAAGTGACAGCGCAGCTGATAAAAGACATAGCTGTGTGGCCGCGTGTTGGGCAGCATGGTGCCAATATAGTAGAAGAGCTGCTTCTCAAGGGGGGCAC | 4 |
| wt-130380-130500 | 3938 | TAAGCATGCAGGCAGGGGAATTCAGGCCGCTAATGACTCCGGGATGGACCTTAGATGCATCCACTTGCATGGATCCTTCAGAGACAGCAGGGATATCGACAGGCTCGGCCAGCGCAATAC | 4 |
| wt-130500-130620 | 3939 | CAAGGGTACCAGACGTCTTGTAAATTAACTTGTAGCGGTTAAGCATAGACGCCAAATCTTCGGTGACATTTGCCTCTCTCCACAGCGCCTCTGGGCTAAGGCCTGGGACCTTTGCCATCA | 4 |
| wt-130620-130740 | 3940 | GTTCGGTCCATGGGATGGTGTAATGCGAAGCATGCCCCTCTATGTCCAGGTGCAGCTTAACCTCGCTGAGGCTGGCAGCCCCCACCTCCCATAGCAACACCAGGCAAAAAACACAGAGCA | 4 |
| wt-130740-130860 | 3941 | ACTGCATCCTAGTCCCGATTTCCCCTCTCAAAATCAGAGATCACCTTGCTCAGACCAGCCCAATCGAAAAACTGAGATCGTATTGCCGGATTCTTCAATGCCTGCATGTAAATCTCCGTC | 4 |
| wt-130860-130980 | 3942 | CAGCATCCAGGTAAATCGTCCTGAAACTCTGAGAGGTCCACAAGCACAAACTGAAGGTAGGCTAGCGTTCGGGTGAACGCAAGACAAACTTCCAACAACACCGCGTCGGCTCGGAAAGGC | 4 |
| wt-130980-131100 | 3943 | TGTATGACTTCCTTAAGTACACTAAAGATGCTGTTCTTATACAGCTTCTCGGCCACACCACTTCGAATTATGGGGGTGTGGCTTTGATGACATACTGTCGTGATTGTTGTTAGACCGGCA | 4 |
| wt-131100-131220 | 3944 | CATACCTTCACAATGTCCTCGGGGGCAAAATACTGTGTTAGGAGCCAGGCACAGTAAACGGCGTGATATGCATCGTTGACACTCTTCAGGTAGCCAGCATCCAGTCCTGACTCATGTTTC | 4 |
| wt-131220-131340 | 3945 | CTCCCTCGCTTCTTCAGGCGGCGCATGTTCTCCTCCACGTTTAACTTCATCCAGACTATGGTGTCCCCCGGGTCTGCGGTAAACGTGGCCAAAACTTGAATAAAGTCACTATAGGAGAGA | 4 |
| wt-131340-131460 | 3946 | AGCTGGCTCCGGAGCAGCATTAGAGGGAAAACCACGGAGGCCGACAGCAAATGGCGATCATGCAAAATCCAACAATCCAGGGGCGCGACTGACCTGGCACCAGACTCGGTAACCAGCAAG | 4 |
| wt-131460-131580 | 3947 | CTCCGCTTCCTAGAGGCCAAGACTCTGAAAGGGGTGGTAAATTTCATCTGGCATGCTAAAACCTCAGCCGACGTGTCTTCCCTTCCATGCCTCGCCCGAGTCACATTCTTGTGCATGGCC | 4 |
| wt-131580-131700 | 3948 | TTAATGGCATTTTCATACACATGAGTCCAGTACCGCATCGGTTCAGGGACTACAATGGTCAGGTCCCCAAAGACAGCCTTCAAATGATTCAGCATAGTAGTCTTTCCCACACCAGGGGCA | 4 |
| wt-131700-131820 | 3949 | CCTTCCAAAAATAGGGAACAGGCAGGTTTGATTACTGGTACATGATTTGTTAGGTGGGTCACAATTGGAACCCGCATGCTCTCCTTCCTCTGAGCCTTGGCCTGGCGGGTGTCTTGGGCA | 4 |
| wt-131820-131940 | 3950 | TCATCCAGATTCAGAACATTCATCACACTCCCACTTAGCCGCTTCAGCTGGGCAGCATGCTTGGATAACTTACTAAACTCGCGCCCATGGGCGGCCAGGTGTTCGAAGAGACCAGAAGGC | 4 |
| wt-131940-132060 | 3951 | TTACCCTTGCCACCATTCTTTTGTTTTAACGCGGAATGAGAAGAGGGCCTGCGGAAATTAGACTCATCCTCAGACTCACAGTCAGATTTGTCATCGAGCCCAAGGCCGGCCAGGCCCTCC | 4 |
| wt-132060-132180 | 3952 | TCAAAGCCTTTCTGGTACATGAAGCTCCGGCTCGTGGAGTCCGCACCTCCTTCTGTGCACGAAGTTTTGCGGAACCAGGAGAAGGGGTCTGGCGTCTTGCTGGGGCCACACTCCCGGCTA | 4 |
| wt-132180-132300 | 3953 | CGGGGCTTCGGGGTAGGGGCAGTAGGCTTTTGGTGTGCGGGTGCTGGTGGCTGGGCTCCCCTGGGCAGGGTAAAGGGGCACGATGTGTGCCGGCTACCCGGAGAGTTTCCAGTATTAGAT | 4 |
| wt-132300-132420 | 3954 | GTCACGGCAGCCTGGGTCCGGCACGGCACCCTCTCCCCAGACAGTCCGGTCGGAGCCATCAAGGGGGGCCAGTGGGTGGGCACCTGGTAGAGGCCGTCGTCATCTTCCTCACCTGCCCCT | 8 |
| wt-132420-132540 | 3955 | GAGTCACTACCGGTTGGGGTAAGAACTGAGGGGGCAAAGTCATCAATCTCAGCGTAAAAGTTTTCGTGTCTTTCGTTTTCAGGGGACTCATCCTCCTGACATTTTCGCCAGCCGCCGGGC | 4 |
| wt-132540-132660 | 3956 | GGGCCGGCCTCCTTTCCTGGAAATCCAGCCATGGATCCCACCCGGGGTCTGTGTGCCCTCTCCACACACGACCTGGCAAAATTTCACAGTCTTCCCCCGGCTAGAAAGGCGGCAGGTAAG | 4 |

| | | | |
|---|---|---|---|
| wt-132660-132780 | 3957 | CGAGCGCACCTTCGGTGTTACTCCAAGCTGCTCTCTCTTAAGAGCTGGGAGCAACTGGCCTCTTTTTTGTCTCTGCCCCCGGGACCCACGTTTACAGACTTTAGACTATTTTTCGAAGTC | 4 |
| wt-132780-132900 | 3958 | ACCCTGGGTCGGAGAATCGCAGATTGCGTTGTGGTAGCTCTGCAGCCTTATCCCCGGTGTTATATTGTAGAATTTAAGACGGCCATGAGCAACACGGCCAACCCGCAAAGCGTTACTCGC | 4 |
| wt-132900-133020 | 3959 | AAGGCACAGAGGCTAGAGGGCACCGCCCAGTTGTGTGACTGTGCCAATTTTCTTCGCACGTCCTGCCCCCCCGTGCTGGGCAGTCAGGGCCTGGAAGTCTTGGCGGCGTTGGTATTTAAA | 4 |
| wt-133020-133140 | 3960 | AACCAGCGATCCCTGAGAACGCTCCAGGTAGAGTTTCCAGCCCTGGGCCAAAAGACCCTCCCCACCTCCACCACCGGCCTGCTAAACCTCCTCTCCCGCTGGCAGGATGGCGCTCTCCGG | 4 |
| wt-133140-133260 | 3961 | GCACGTCTTGATAGACCCCGCCCGACTGCCCAGGGACACAGGCCCCGAACTCATGTGGGCCCCAAGCCTTCGCAACTCACTGCGCGTGTCCCCCGAAGCGCTCGAGCTGGCAGAGCGGGA | 8 |
| wt-133260-133380 | 3962 | GGCCGAAAGGGCCAGGTCGGAGCGGTGGGACAGGTGTGCCCAGGTGCTCAAAAATAGGCTGCTCCGCGTGGAGCTGGACGGCATCATGCGTGACCACCTGGCCAGGGCGGAGGAGATCCG | 8 |
| wt-133380-133500 | 3963 | CCAGGACCTGGATGCTGTAGTGGCCTTCTCTGATGGCCTGGAGAGCATGCAGGTCAGGTCCCCCTCCACGGGAGGGCGCTCTGCGCCAGCCCCGCCCTCCCCATCCCCAGCCCAGCCGTT | 8 |
| wt-133500-133620 | 3964 | CACTCGGCTCACCGGGAACGCCCAGTATGCAGTCTCAATCTCTCCCACGGACCCCCCTCTGATGGTGGCCGGCAGCCTGGCTCAAACGCTGCTTGGTAATCTGTACGGGAACATCAACCA | 4 |
| wt-133620-133740 | 3965 | GTGGGTACCGTCCTTCGGACCCTGGTACAGGACCATGTCGGCTAATGCCATGCAGCGGCGCGTGTTCCCTAAGCAGCTGAGGGGCAACCTGAACTTTACCAACTCCGTCTCCCTAAAGCT | 4 |
| wt-133740-133860 | 3966 | GATGACAGAAGTGGTGGCGGTGCTTGAGGGCACCACCCAGGACTTTTTCTCAGACGTCAGGCACCTGCCAGACCTCCAGGCTGCCCTGATCCTCTCGGTGGCCTACCTGCTACTCCAGGG | 4 |
| wt-133860-133980 | 3967 | GGGCTCCTCACACCAGCAGCGCCCCCTCCCTGCCTCACGGGAAGAGCTGCTGGAGCTGGGCCCGGAGAGCCTAGAGAAAATCATCGCCGACCTCAAGGCCAAGTCACCCGGCGGAAATTT | 4 |
| wt-133980-134100 | 3968 | TATGATTTTAACAAGCGGAAACAAGGAAGCGCGCCAGTCAATAGCCCCTCTCAACCGACAGGCGGCATATCCACCCGGCACATTCGCGGACAATAAGATTTACAACCTGTTTGTGGGAGC | 4 |
| wt-134100-134220 | 3969 | GGGACTACTGCCCACGACGGCCGCGCTGAACGTGCCCGGGGCGGCGGGTCGGGACCGGGACCTGGTGTACCGGATCGCCAACCAGATCTTTGGGGAGGATGTGCCCCCCTTCTCATCTCA | 8 |
| wt-134220-134340 | 3970 | CCAGTGGAACCTGCGCGTAGGTTTAGCCGCACTCGAGGCCCTGATGCTCGTCTACACGCTCTGCGAGACCGCCAACCTGGCCGAGGCGGCCACCCGGCGTCTACACCTATCGTCCCTGCT | 8 |
| wt-134340-134460 | 3971 | CCCCCAGGCAATGCAGCGGCGCAAGCCTGCCATGGCGTCAGCTGGTATGCCGGGCGCCTATCCAGTCCAGACGCTTTTCCGCCACGGGGAGCTCTTCCGCTTCATCTGGGCCCACTACGT | 8 |
| wt-134460-134580 | 3972 | GAGGCCCACGGTGGCGGCAGACCCCCAGGCCTCCATCAGCTCTCTTTTCCCCGGGCTGGTTTTGCTGGCCCTGGAGCTGAAGTTGATGGATGGGCAGGCTCCCTCCCATTATGCCATAAA | 4 |
| wt-134580-134700 | 3973 | CCTGACCGGACAAAAGTTTGACACCCTCTTTGAGATTATCAACCAGAAGCTTTTATTTCACGACCCGGCTGCCATGCTGGCGGCGCGCACACAGCTGCGTCTAGCCTTCGAGGACGGCGT | 4 |
| wt-134700-134820 | 3974 | CGGTGTTGCCCTGGGGCGCCCCTCGCCCATGCTTGCGGCGCGGGAGATCCTGGAGCGTCAGTTCTCAGCCTCGGATGACTACGACCGGCTGTACTTCCTGACGCTGGGCTACCTGGCCTC | 8 |
| wt-134820-134940 | 3975 | CCCGGTGGCCCCAAGCTGAGCCAGTTCCTCGCACTGGAGTGGGTCATTGGCAAAAAGGTAAATAAACTCATCGCACGGGGGTTTTGCCTCCTTCTCGTCTCTTGTTTCGGGTAGGGGAGT | 4 |
| wt-134940-135060 | 3976 | AAGGCCGTGCCAGGCCGCCATGCTCAGGGCCACGGCGTGCCAGAGGCCCTCGTAGTCGTGCGCATCCGAGAGGATGGCACGGTCCAGAAGCAGATAGCCGGCCAGGCAGAGGAAGGCCAC | 8 |
| wt-135060-135180 | 3977 | AAAGAGGGGGCGAAGGCGTGCCCGAACCCGGGTTTCATGCTCGTCTGCACCCCAGTGGACAAGGCAGTAGAGGACACCCACCACCAGGCGGTTAGGGAGGACACTGCCAAGGTTGAAGAG | 4 |
| wt-135180-135300 | 3978 | CAGATTTCCGTCAGCCAGGGTGACCTGGCTCAGGTCCGGCGCCCTGCGCAGTCCAAGCTGCGCCCACACACATGCACAGACGGCCCCTGTGACATCAGGCCGGTCATGCAAAAACAGACA | 4 |
| wt-135420-135540 | 3980 | GCGGTTTCCAGTATTTATCACGGCTTCCCGCCACAGGTAAAAGTTAACACTTAGGGTCAGCAGCTTGGTCAGGGATAGGTGCAAAAACCTGAGCTCGTCCTCGCGCAGAGCGCAAAGCGG | 4 |
| wt-135540-135660 | 3981 | CCAGTTCTTTAGCATCTTCAGGAGGAGCCCGTGAATCCCAGGTGTCATTCGCGCGTCATCCCCGCGCACCCCCAGTCCCATTAACATAGCGGGCACAATGGTGCAGGCACCGTCTGTATA | 4 |
| wt-135660-135780 | 3982 | CGTCTGCGGCTTCGTGGAGCGCCCGGACGCCCCACCCAAGGACGCCTGCCTTCACCTGGATCCCCTCACCGTCAAGAGCCAGCTCCCTCTGAAGAAGCCCTTGCCACTCACGGTGGAACA | 8 |
| wt-135780-135900 | 3983 | CCTGCCGGATGCTCCGGTCGGCTCAGTCTTTGGCCTTTACCAGAGCCGAGCGGGTCTCTTTAGCGCAGCCTCGATTACCTCTGGGGACTTCCTGTCCCTGCTGGACTCAATTTACCACGA | 4 |
| wt-135900-136020 | 3984 | TTGCGATATTGCACAGAGTCAGCGCCTGCCCCTCCCTCGAGAACCCAAGGTGGAGGCTCTGCACGCCTGGCTCCCCTCACTGTCACTGGCCTCCCTCCACCCAGACATACCCCAAACCAC | 4 |
| wt-136020-136140 | 3985 | CGCAGATGGAGGCAAGCTGTCCTTCTTTGACCACGTGTCTATCTGTGCCCTGGGTCGTCGGCGCGGCACCACGGCAGTCTACGGTACAGACCTTGCGTGGGTCCTGAAGCACTTTAGTGA | 4 |
| wt-136140-136260 | 3986 | CCTGGAACCGTCTATCGCCGCCCAGATTGAGAATGACGCCAATGCCGCAAAGCGTGAATCCGGATGCCCGGAAGACCACCCTCTGCCCCTCACGAAGCTCATAGCTAAGGCAATCGATGC | 4 |
| wt-136260-136380 | 3987 | TGGATTTCTGAGAAACCGCGTGGAGACTCTGAGGCAGGACAGGGGTGTGGCCAATATCCCAGCCGAGTCGTATTTAAAGGCCAGCGACGCCCCGGACCTACAAAAGCCGGACAAGGCACT | 4 |
| wt-136380-136500 | 3988 | TCAGAGCCCACCACCGGCCTCCACAGACCCAGCCACCATGCTATCAGGTAACGCAGGAGAAGGAGCAACAGCCTGCGGAGGTTCGGCCGCCGCGGGCCAGGACCTCATCAGCGTCCCCCG | 8 |
| wt-136500-136620 | 3989 | CAACACCTTTATGACACTGCTTCAGACCAACCTGGACAACAAACCGCCGAGGCAGACCCCGCTACCCTACGCGGCCCCGCTGCCCCCCTTTTCCCACCAGGCAATAGCCACCGCGCCTTC | 4 |
| wt-136620-136740 | 3990 | CTACGGTCCTGGGGCCGGAGCGGTCGCCCCGGCCGGCGGCTACTTTACCTCCCCAGGAGGTTACTACGCCGGGCCCGCGGGCGGGGACCCGGGTGCCTTCTTGGCGATGGACGCTCACAC | 8 |
| wt-136740-136860 | 3991 | CTACCACCCCCACCCACACCCCCCTCCGGCCTACTTTGGCTTGCCGGGCCTCTTTGGCCCCCCTCCACCCGTGCCTCCTTACTACGGATCCCACTTGCGGGCAGACTACGTCCCCGCTCC | 8 |
| wt-136860-136980 | 3992 | CTCGCGATCCAACAAGCGGAAAAGAGACCCCGAGGAGGATGAAGAAGGCGGGGGGCTATTCCCGGGGGAGGACGCCACCCTCTACCGCAAGGACATAGCGGGCCTCTCCAAGAGTGTGAA | 4 |
| wt-137100-137220 | 3994 | GCCTCATCAAAGCTACGAAGTGCCCAGATACGTCCCTCATCCGCCCCCACCACCAACTTCTCACCAGGCAGCTCAGGCGCAGCCTCCACCCCCGGGCACACAGGCCCCCGAAGCCCACTG | 8 |
| wt-137220-137340 | 3995 | TGTGGCCGAGTCCACGATCCCTGAGGCGGGAGCAGCCGGGAACTCTGGACCCCGGGAGGACACCAACCCTCAGCAGCCCACCACCGAGGGCCACCACCGCGGAAAGAAACTGGTGCAGGC | 8 |
| wt-137340-137460 | 3996 | CTCTGCGTCCGGAGTGGCTCAGTCTAAGGAGCCCACCACCCCCAAGGCCAAGTCTGTGTCAGCCCACCTCAAGTCCATCTTTTGCGAGGAATTGCTGAATAAACGCGTGGCTTGAAAGTA | 4 |
| wt-137460-137580 | 3997 | AACTTTATTGCGTGTTAGTACCTGTCCATTCACAGGGGTATCCAGCCCTTGCGCCGCCTCCCCCAGCCCGCCAGCCACCCCAGACAGGAGATGATAATGATGAGGAGCACCGGAGCCACC | 4 |
| wt-137580-137700 | 3998 | ACAGCACAAGTGATTAGGAGCAGGGCCCAGTGCACCCAGGTGGTCTTAGGGCGCCAGGGATCGATTGGAAAAGGGCCCAGGGTCACTGGCTTATGCGTGGGACGTTTAGAAACAGGCCGC | 4 |
| wt-137700-137820 | 3999 | CTATGGGGCCTGTGACTGGTGCTTGTGGTGTGGGAGACTAATGTGGTGGGGGCTATGGTAGTGGCTGGGATAACAGTAAGATGCATACGCTGAGTGAGGGTCCGGTTGGCATGGTATTGG | 4 |
| wt-137820-137940 | 4000 | TCGTCTTCTTCCCCTGCAGAGTAATTGCAGTGGACCCCGGAGGCCACACTGCAATTTCTCAGTGTCACATTGCACGTGTAGTAACCTGCATGCGCAAGGGTCACATTGGGGATTATCAGA | 4 |
| wt-137940-138060 | 4001 | GAGACGGAGGTGTTGGAGTCATTTACCCATTCTAGGGTAAGGCTATAATTGTAACCCCCGTTAGTTATATGAGTTCCGTTGTTGGAAGTAGCTACGGCCAAGGGCAGTTGTCCATCCCCG | 4 |
| wt-138060-138180 | 4002 | GGAGTGTATCCCCGGCCCAACTCGATCCGAGAGACCGACTCATTGCTAGGAACGCTGCAGGTGAGATTCACTCTAGCACCTGCATGGGCGGTGACATTTTCAAATTTAACCAGATCTGAG | 4 |
| wt-138180-138300 | 4003 | AAAAATGCACAAACAGACCCCACACAGCAGCACAATAGAAGCACTAAATGAGTCATTCCTAAACTGTCAGTTTTAAAACTCCCTGCTTCTCAGGCCTAAATATGTGGTGGGGTGTGCTTA | 4 |
| wt-138300-138420 | 4004 | GGATCACTTTCATATTCTGCAACAACAGCCATACCCGGAAGAGGAGCTGCCGGTTGCCATTTTTCAAGCTGCTAAACCACGAGTGGCAGCAGGCCTAAGAAGCTCCTCAGCAACATGGAG | 4 |
| wt-138420-138540 | 4005 | ACCTCGAAGGGAAACTGGCAGGAGCAGGGAGTCACGTAGGCACTAGCCTCTTCATGTGAGGTAAGAGATCGCTAAAAATGGGATCAGGGTATGTAAACCGAGTTTTGCGGGGGATGGTGA | 4 |
| wt-138540-138660 | 4006 | GCCAGACACGGCGGGTGGGGGAAGGAGCTGACACGATTGAGTAGAAAGGGCCAAAAATACACCAGCTATAAGGAATTGCTCAGGCCAAAGTTGTTCCTCAGGTGGCTTTAGGCCTAATGT | 4 |
| wt-138660-138780 | 4007 | AGGCAATTGCGTGCCTAGAACATTGCTAATGTGCCCTGGGTTTCCTGCCTTCATGCAAATATTCTACCTCCCCCGGCCTGGTGCAAAATGTCTGCCTCAGAATACTAACAGCTAATCCAA | 4 |
| wt-138780-138900 | 4008 | GCTAACATTCTATCAGTAAACGGGCAGAAAACTGATAAGGACCGCGGAGTTTGGCCCTCCGCGGTGTCCGGTGGTCCTCACACGTGCCCTCCCCCCGGGCCGATGGCTGAGGCCCGGAAT | 4 |
| wt-138900-139020 | 4009 | ATGCAAGTGCATCTTTCTAACCAGTAGGGGCCTCCACCTAGGTGCTTTGTTAATCTTTAGTGGGAACTAGTGGGAGTGCTGTGCCTCGGGTACCCCTATCCTATAGGTCCTACCGGAGCT | 4 |

| | | | |
|---|---|---|---|
| wt-139020-139140 | 4010 | CCTTGTCTTGATAATCCCTGTAAACACACACCACCTAAGAACAAGGCATTGTTAACCTTTGGTGGAACCTAGTGTTAGTGTTGTGCTGTAAATAAGTGTCCAGCGCACCACTAGTCACCA | 4 |
| wt-139140-139260 | 4011 | GGTGTCACCGGAGGCTACTTGCCTCAGTGCCACTTTTACCTTCTCAAATCTATACGGGGGGGGGGGGGGCTCTGTAACATTTGGTGGGACCTGATGCTGCTGGTGTGCTGTAAATAAGTG | 4 |
| wt-139260-139380 | 4012 | CCTAGCACATCACGTAGGCACCAGGTGTCACCAGGGCTACTTGCCTCGGCATCTCCTCACCGGAGAAGGGGTTAACAAACCCGTGGGGGGTCTTAGTGGAAGTGACGTGCTGTGAATACA | 4 |
| wt-139380-139500 | 4013 | GGTCCATAGCACCGCTATCCACTATGTCTCGCCCGGGCTATATGTCGCCTTACCTCCCCTATATAGTCACGACCCCACCGAACCAGGCATGATGTAGAATAAAATTTTATGCATCATCTT | 4 |
| wt-139500-139620 | 4014 | CTAATCTGTGCCGCTTGGAGGGAAACATGACCACCTGAAGTCTGTTAACCAGGTCAGTGGTTTTGTTTCCTTGATAGAGACACAAGGACTGCCAGCCCCATTGGGGAGGGGGGGTGGGTA | 4 |
| wt-139620-139740 | 4015 | CGGGAGAGTTTGGGCTCGTTTAAACAAAGTCTCATCTGATGCTCTGTGGCACCTCAAGGTGAATATAGCTGCCCATCGACGTATCGCTGGAAACCGGTGGGCCGCTGTTCACCTAAAGTG | 4 |
| wt-139740-139860 | 4016 | ACGCAAGGTCTGTCAGCCGCCAGGGTCCGTTTACCAGGCTTTCAGGTGTGGAATTTAGATAGAGTGGGTGTGTGCTCTTGTTTAATTACACCAAGATCACCACCCTCTATCCATATCCCA | 4 |
| wt-139860-139980 | 4017 | CAATTGATAAACCTCCGCATGTCCAACCACCACGTTGAACAGGATGTGGCACCCTAAGAGGACGCAGGCATACAAGGTTATTACCCAGTCCTTGTATGCCTGGTGTCCCCTTAGTGGGAC | 4 |
| wt-139980-140100 | 4018 | GCAGGCCTAGGTAGCATCATTTACACTAAAAGCAGTGACCTTGTTGGTACTTTAAGGTTGGTCCAATCCATAGGCTTTTTTTGTGAAAACCCGGGGATCGGACTAGCCTTAGAGTAACTC | 4 |
| wt-140100-140220 | 4019 | AAGGCCAAGCATTTCACACCTGCAAATGCACCATGTAACCACAGATCTAAACTGAAAGTTGCAGCTTTAGATGGCAAGGAAACTTGGGTTTCAGGCATAGAAAGCCTGGCTCACTATAGC | 4 |
| wt-140220-140340 | 4020 | AGCCCATGTTTGTTCCAGGGTGGGGGAAAGGCACGTGCCCTTAGAAAACTTAGCTGCAAAAATTCTATTGTGTTGGGAGAGCCTCTATATCTAAAGGCCTTTCCTCACAATACAAATGTT | 4 |
| wt-140340-140460 | 4021 | ACTAACGTCTGCCCTCTGGAGACCTGCTATGTGGCTAGACGTATGGCCTACCCAAGACGTTGGGGGTCTCGGGTAGGCCATGATTCTTCCAGGCATAGGTTACAACCAGTCACTGCTATC | 4 |
| wt-140460-140580 | 4022 | AAGCCTACTCAGTTCCCAACGCAGCACATACCCCCCGCCTCTCCTGCCATGAGGACTTATGGCAGTGTTTACTGTTCTGCTTTTACTCTTGGACCAGGCTGTCATTCTATCAGAATAACA | 4 |
| wt-140580-140700 | 4023 | GGGGAAGCAAGGCCCCCTGCTTCAGCGGGACACGTGTTTCTAGAATCTCGGAGCCAATAACTACCTGCCCCTCTAATCTGTATGCTGCATGAAAAACCACATACACGTGATGTAAGTTTA | 4 |
| wt-140700-140820 | 4024 | GCCAGTTTATTGTTACACCAATGCCCCGAAAGTCTCCCCCTGTCCCTTTGGGTCTCAGGACCCAGCCCTGGAGCTCGGGGGCGGCCGGGTGGCCCACCGGGTCCGCTGGGTCCGCTGCCC | 8 |
| wt-140820-140940 | 4025 | CGCTCCGGCGGGGGGTGGCCGGCTGCAGCCGGGTCCGGGGTTCCGGCCCTGGAGCTCGGGGGGCGGCCGGGTGGCCCACCGGGTCCGCTGGGTCCGCTGCCCCGCTCCGGCGGGGGGTGG | 12 |
| wt-140940-141060 | 4026 | CCGGCTGCAGCCGGGTCCGGGGTTCCGGCCCTGGAGCTCGGGGGGCGGCCGGGTGGCCCACCGGGTCCGCTGGGTCCGCTGCCCCGCTCCGGCGGGGGGTGGCCGGCTGCAGCCGGGTCC | 12 |
| wt-141060-141180 | 4027 | GGGGTTCCGGCCCTGGAGCTCGGGGGGCGGCCGGGTGGCCCACCGGGTCCGCTGGGTCCGCTGCCCCGCTCCGGCGGGGGGTGGCCGGCTGCAGCCGGGTCCGGGGTTCCGGCCCTGGAG | 12 |
| wt-141180-141300 | 4028 | CTCGGGGGGCGGCCGGGTGGCCCACCGGGTCCGCTGGGTCCGCTGCCCCGCTCCGGCGGGGGGTGGCCGGCTGCAGCCGGGTCCGGGGTTCCGGCCCTGGAGCTCGGGGGGCGGCCGGGT | 12 |
| wt-141300-141420 | 4029 | GGCCCACCGGGTCCGCTGGGTCCGCTGCCCCGCTCCGGCGGGGGGTGGCCGGCTGCAGCCGGGTCCGGGGTTCCGGCCCTGGAGCTCGGGGGGCGGCCGGGTGGCCCACCGGGTCCGCTG | 12 |
| wt-141420-141540 | 4030 | GGTCCGCTGCCCCGCTCCGGCGGGGGGTGGCCGGCTGCAGCCGGGTCCGGGGTTCCGGCCCTGGAGCTCGGGGGGCGGCCGGGTGGCCCACCGGGTCCGCTGGGTCCGCTGCCCCGCTCC | 12 |
| wt-141540-141660 | 4031 | GGCGGGGGGTGGCCGGCTGCAGCCGGGTCCGGGGTTCCGGCCCTGGAGCTCGGGGGGCGGCCGGGTGGCCCACCGGGTCCGCTGGGTCCGCTGCCCCGCTCCGGCGGGGGGTGGCCGGCT | 12 |
| wt-141660-141780 | 4032 | GCAGCCGGGTCCGGGGTTCCGGCCCTGGAGCTCGGGGGGCGGCCGGGTGGCCCACCGGGTCCGCTGGGTCCGCTGCCCCGCTCCGGCGGGGGGTGGCCGGCTGCAGCCGGGTCCGGGGTT | 12 |
| wt-141780-141900 | 4033 | CCGGCCCTGGAGCTCGGGGGGCGGCCGGGTGGCCCACCGGGTCCGCTGGGTCCGCTGCCCCGCTCCGGCGGGGGGTGGCCGGCTGCAGCCGGGTCCGGGGTTCCGGCCCTGGAGCTCGGG | 12 |
| wt-141900-142020 | 4034 | GGGCGGCCGGGTGGCCCACCGGGTCCGCTGGGTCCGCTGCCCCGCTCCGGCGGGGGGTGGCCGGCTGCAGCCGGGTCCGGGGTTCCGGCCCTGGAGCTCGGGGGGCGGCCGGGTGGCCCA | 12 |
| wt-142020-142140 | 4035 | CCGGGTCCGCTGGGTCCGCTGCCCCGCTCCGGCGGGGGGTGGCCGGCTGCAGCCGGGTCCGGGGTTCCGGCCCTGGAGCTCGGGGGGCGGCCGGGTGGCCCACCGGGTCCGCTGGGTCCG | 12 |
| wt-142140-142260 | 4036 | CTGCCCCGCTCCGGCGGGGGGTGGCCGGCTGCAGCCGGGTCCGGGGTTCCGGCCCTGGAGCTCGGGGGGCGGCCGGGTGGCCCACCGGGTCCGCTGGGTCCGCTGCCCCGCTCCGGCGGG | 12 |
| wt-142260-142380 | 4037 | GGGTGGCCGGCTGCAGCCGGGTCCGGGGTTCCGGCCCTGGAGCTCGGGGGGCGGCCGGGTGGCCCACCGGGTCCGCTGGGTCCGCTGCCCCGCTCCGGCGGGGGGTGGCCGGCTGCAGCC | 12 |
| wt-142380-142500 | 4038 | GGGTCCGGGGTTCCGGCCCTGGAGCTCGGGGGGCGGCCGGGTGGCCCACCGGGTCCGCTGGGTCCGCTGCCCCGCTCCGGCGGGGGGTGGCCGGCTGCAGCCGGGTCCGGGGTTCCGGCC | 12 |
| wt-142500-142620 | 4039 | CTGGAGCTCGGGGGGCGGCCGGGTGGCCCACCGGGTCCGCTGGGTCCGCTGCCCCGCTCCGGCGGGGGGTGGCCGGCTGCAGCCGGGTCCGGGGTTCCGGCCCTGGAGCTCGGGGGGCGG | 12 |
| wt-142620-142740 | 4040 | CCGGGTGGCCCACCGGGTCCGCTGGGTCCGCTGCCCCGCTCCGGCGGGGGGTGGCCGGCTGCAGCCGGGTCCGGGGTTCCGGCCCTGGAGCTCGGGGGGCGGCCGGGTGGCCCACCGGGT | 12 |
| wt-142740-142860 | 4041 | CCGCTGGGTCCGCTGCCCCGCTCCGGCGGGGGGTGGCCGGCTGCAGCCGGGTCCGGGGTTCCGGCCCTGGAGCTCGGGGGGCGGCCGGGTGGCCCACCGGGTCCGCTGGGTCCGCTGCCC | 12 |
| wt-142860-142980 | 4042 | CGCTCCGGCGGGGGGTGGCCGGCTGCAGCCGGGTCCGGGGTTCCGGCCCTGGAGCTCGGGGGGCGGCCGGGTGGCCCACCGGGTCCGCTGGGTCCGCTGCCCCGCTCCGGCGGGGGGTGG | 12 |
| wt-142980-143100 | 4043 | CCGGCTGCAGCCGGGTCCGGGGTTCCGGCCCTGGAGCTCGGGGGGCGGCCGGGTGGCCCACCGGGTCCGCTGGGTCCGCTGCCCCGCTCCGGCGGGGGGTGGCCGGCTGCAGCCGGGTCC | 12 |
| wt-143100-143220 | 4044 | GGGGTTCCGGCCCTGGAGCTCGGGGGGCGGCCGGGTGGCCCACCGGGTCCGCTGGGTCCGCTGCCCCGCTCCGGCGGGGGGTGGCCGGCTGCAGCCGGGTCCGGGGTTCCGGCCCTGGAG | 12 |
| wt-143220-143340 | 4045 | CTCGGGGGGCGGCCGGGTGGCCCACCGGGTCCGCTGGGTCCGCTGCCCCGCTCCGGCGGGGATGGGGGTGCGCTCCCAGGCCGGACCCTGGTGCCAGGCAGGGACCCCGCGCCACCCGCT | 12 |
| wt-143340-143460 | 4046 | TCATGGGGGGGGAGGCCGCCGCAAGGACGCCGGGCCGGCTGGGAGGTGTGCACCCCCCGAGCGTCTGGACGACGCTGGCGAGCCGGGCCAGCTCGCCTTCTTTTATCCTCTTTTTGGGGT | 8 |
| wt-143460-143580 | 4047 | CTCTGTGCAATACCTTAAGGTTTGCTCAGGAGTGGGGGGCTTCTCATTGGTTAATTCAGGTGTGTGATTTTAGCCCGTTGGGTTACATTAAGGTGTGTAACCAGGTGGGTGGTACCTGGA | 4 |
| wt-143580-143700 | 4048 | GGTCATTCTATTGGGATAACGAGAGGAGGAGGGGCTAGAGGCCCGCGAGATTTGGGGTAGGCGGAGCCTCAGGAGGGTCCCCTCCATAGGGTTGAACCAGGAGGGGGAGGATCGGGCTCC | 4 |
| wt-143700-143820 | 4049 | GCCCCGATATACCTAGTGGGTGGAGCCTAGAGGTAGGTATCCATAGGGTTCCATTATCCTGGAGGTATCCTAAGCTCCGCCCCTATATACCAGGTGGGTGGAGCTAGGTAGGATTCAGCT | 4 |
| wt-143820-143940 | 4050 | AGGTTCCTACTGGGGTACCCCCCTACCCTACCTTAAGGTGCGCCACCCTTCCTCCTTCCGTTTTAATGGTAGAATAACCTATAGGTTATTAACCTAGTGGTGGAATAGGGTATTGCAGCT | 4 |
| wt-143940-144060 | 4051 | GGGTATATACCTATAGGTATATAGAACCTAGAGGAAGGGAACCCTATAGTGTAATCCCTCCCCCCCCTACCCCCCCCTCCCTTACGGTTGCCTGAGCCCATCCCCCACCCCAGCACCCCG | 4 |
| wt-144060-144180 | 4052 | GGGTGACGTGGCACCCCGCGTGCCTTACTGACTTGTCACCTTTGCACATTTGGTCAGCTGACCGATGCTCGCCACTTCCTGGGTCATGACCTGGCCTGTGCCTTGTCCCATGGACAATGT | 4 |
| wt-144180-144300 | 4053 | CCCTCCAGCGTGGTGGCTGCCTTTGGGATGCATCACTTTGAGCCACTAAGCCCCCGTTGCTCGCCTTGCCTGCCTCACCATGACACACTAAGCCCCTGCTAATCCATGAGCCCCGCCTTT | 4 |
| wt-144300-144420 | 4054 | AGGAAGCACCACGTCCCGGGGACGGAAGCTGGATTTTGGCCAGTCTTCAATTTTGGGGAGTGGTTTTGTGTGAGCCGGAAGTTGGCAATGGGGTGAGGGTGGCGCTGGTTAAGCTGACGA | 4 |
| wt-144420-144540 | 4055 | CCTCCCAAGGTCTCTCACCCTGGGTACACAGGTGGGGCGGCAGCCTCTAACTTTGGCTGTGGCCTCTATTTCCTCCCTTTCCTAGCCAGGGCCATGTGTTCCTGCATGTCTACTTGCCTC | 4 |
| wt-144540-144660 | 4056 | CTGTGGTGGCAGAGCTTGGCCCTGGGCCCAACCCCCGCCTTGGGAGCCTGTAGGGGCCAACACCCTTGGTTTGTTTGTGTTCCTGTTTGCTGGCAACTTACTGGCAGCCGAGCAGATTCT | 4 |
| wt-144660-144780 | 4057 | AATGGGCGCCCGCCTTCTTTCTCTCTTGTTTTATTAATAGAATCTCAGCCAGGACCTATACCTGAGACTTCAAAGTCTGGTCCTGGGTTCTGAGACCCCCAAGATTTGTCATGCACACCT | 4 |
| wt-144780-144900 | 4058 | GCACACCTGTTGGTATTGGGTTTCTATTCTTGAGTGTGAAAGTTTGTAAAAAAATTCATAAAATGTCACTAATTCCTCTTACCTGTTTAGGGTATTGTGCAATTCTTCAGCCTGCCTATT | 4 |
| wt-144900-145020 | 4059 | TTCAATTTGCCTAAGGTGGCAATTTAAGATGTGGTTAATTAACCATTTTCCTGTCTGACACCACTGCATGGGCAACCGGGTTCCATGGCACATTTAGAGATAAACATAGATGTCTTGTCT | 4 |
| wt-145020-145140 | 4060 | TGCTCATGTGCAGAGGAGGGGGTGTTGGTGTGCAATATAGTTTCTGGATTCCAAATTGAGTTGGGGGTGCTATTTTCACTATGGAATTAAATTACTGACATTAGACAGTGGACACCGGGC | 4 |

| | | | |
|---|---|---|---|
| wt-145140-145260 | 4061 | TATATGTGGGGATGTCTGTGGCTTGTCATTTCCTCTTAGAAGGTAATCCCCCATCTTAACTTCCCTTTAAATTGTGATGCAAGCCCTGGGTTATTTATAGAATGATTATCTAGGTTTGAT | 4 |
| wt-145260-145380 | 4062 | AGTCTGAAGGCTGGGCAGAGAATGTTTGTAATTTTTATTCACCTTCTTTACCCCCCACGAGTATCCAGTTCTAGAAGATCTCCTGATATCCCGGGCTGCCATTATTCCCTTGAGTGTTAT | 4 |
| wt-145380-145500 | 4063 | AGCTTCCTCTTAACTTAAGCAAGAGCTCCAGGATGTTAGCTTTTTTGGTGGGGCTGGTTGTCAGGAAGAGGTTCCAGTGTTGTCCTTTATTTTTAGATGTTAGCTTTGTGTTAGGTTAGT | 4 |
| wt-145500-145620 | 4064 | ATGGGCTGGGTATTCACTAGTGAAGGCAACTAACACAGTTAGACGTGCTAGTTGTGCCCACTGGTGTTTATCCGGTCCCAAATGTCACCACAGAACACAGGGGGCTGGATTTGGCAGCAG | 4 |
| wt-145620-145740 | 4065 | CACTTGTGCTTTTGTTGATTTTTACCCGTGTATCAGAGTGGGGGATGCTAGCCAATTTAGCTTCCCCTCCCCTTAACAGGGGGTCTCGCGGGGTGCCAATTGTCGCCTGCCTTCCCCCGC | 4 |
| wt-145740-145860 | 4066 | TTCCCCTTGTTAACTTATAGCATGATAGGTAGGTCACCTAACGTGGAAGCCTGGTGGGTGATCCTTCCTCGGTAGGGAGCGCTTAGGGCTGTTGAGCTCAACAGCCCCACCTGGGTAAAA | 4 |
| wt-145860-145980 | 4067 | TGTATGTTCTAAAGAGTTACCCAATTATAACAAAACTGTTGTAGGGTAACGAAGACCTGATGGAAGTGGTATTGTTGCCGTTGAAAGACGGGTGTCCTGGCTCAAGTTCGCACTTCCTAT | 4 |
| wt-145980-146100 | 4068 | ACAGTGTTAAAGCCTTGTATCGGAAGTTTGGGCTTCGTCCCAGTGTACTCGATAATGTCGACTGCTGCGAAAGGTTTGGACCGTCTTCCAGTAGGTGTTGGGGGTCCCAAATCACGAGGT | 4 |
| wt-146100-146220 | 4069 | TAGGCAGGTGCACTTGGCTCTTTAGGAGGGACCCTTAAGCCAGACAATGTAGTGCCCCTTTTTTTTGCAAATTGGCCTTATTATTAATTTCTTGTTAACACTAATTCTGTTCTATGACCC | 4 |
| wt-146220-146340 | 4070 | TGTGTTTTTCAGATGCCGTTGAACGTGTCACTGAGCTGAATTTGGACGCAGCTACTTGACCTTTGCCCCCGTGCCTCCAGCGCTGATAAGTGCTGCGTCCACTTTGTGTTACAGGTGGGC | 4 |
| wt-146340-146460 | 4071 | CAAACCTCCAGAATATCAATTGGTGGGGCCTTGGTGGGCTGCATAAGGCAGTAGGTTTGAGGTGACCTACTTGGACCATGTGGATCCAGTGTCCTGATCCTGGACCTTGACTATGAAACA | 4 |
| wt-146460-146580 | 4072 | ATTCTAAAAAAATGCATCATAGTCCAGTGTCCAGGGACAGTGCACTCGGAAGTCTCATCATCTCCGTTTGTGTGTTTAGTGTGGCCAGTACGGCCACCCCTGTGCCACGCCCTGGCATGC | 4 |
| wt-146580-146700 | 4073 | TGCTGACATCTGGCCGCCAATTTCAGCGGGCCCTTTTCCCCCTTGTTCACCCCATAGCAAGAAGGGTAGGTTACATGGGTATTTTCCCATCAGCACCTGACTGGCCGGTGCAATTAGAGG | 4 |
| wt-146700-146820 | 4074 | AGAGGGCAACAACGCAAGGCTGTTGTTTTATTTGGGTTACAAGAGCTGCGGCGGTCGATGGGTTCACTGATTACGGTTTCCTAGATTGTACAGATGAACTAGAACTGTCACAATCTATGG | 4 |
| wt-146820-146940 | 4075 | GGTCGTAGACAGTGTGCTTACCAGACTTCCATGGAAGATGTGAATTTGCTGCTAGCTATATGGGTGGTGCTATGGGCTCCCTAGGGACTCATGTAGTGGGGCTTTGTGATAGCTAATGAA | 4 |
| wt-146940-147060 | 4076 | TGTGGCAGCTGTTGTTTGTACTGGACCCTGAATTGGAAACAGTAACTTGGATTCTGTAACACTTCATGGGTCCCGTAGTGACAACTATGCTGAATATCTTGAATATGGGAGGAGGGGGGC | 4 |
| wt-147060-147180 | 4077 | TTTGGGTTCCATTGTGTGCCCTTTCCTGGCCAACGTGAGGGTCCTAGTGTTATAGGGCGTGGCAGTTTTCTTGAGGGCTAATAACCCGGGTGAGGCGGTTGTCACAGGTGCTAGACCCTG | 4 |
| wt-147180-147300 | 4078 | GAGTTGAACCAGTACCACTCGGTTACAAAGTCATGGTCTAGTAGTTGTGACCCTGCAAAGCTACGTGGGGATGAGCAGCCAGGGACTTTGGTTGGCAAGCAGACAGGCGGCGCATTGGAA | 4 |
| wt-147300-147420 | 4079 | CCCCAGAGGAGTGTCCCGGGGCCACCTCTTTGGTTCTGTACATATTTTGTTATTGTACATAACCATGGAGTTGGCTGTGGTGCACTCCATCTGGTAAGGGGGCTGGTGCGGACGCCTGTG | 4 |
| wt-147420-147540 | 4080 | TTTAGTCTATGCCAATGTTTACCTGCCTTGGGTTACTATTCCAAACGACCACACCTTTGAGGACACCTGGAGCCCTGATCATTCTCGGCTTTTACTGCCACCTGGCTTCTGTTGGGTCAG | 4 |
| wt-147540-147660 | 4081 | ACAGTTTGGTGCGCTAGTTGTGTGCTTAGCAGCAACGCACACCAGGCTGACTGCCTTAGCAGTGTGGCCCTTTATTGTGGCATCCTAAGGAGGGATTCTGGAGTGCCTTTCGCGTGAAGC | 4 |
| wt-147660-147780 | 4082 | ATGCCCTGAGACGTACTCGAGTTAGGACTTAATCGCTCCTGTGCCGCTGGATGAGGGAGCGCCAATTTGTACATCCTAGCTCTGGCCATAGAGTTAGCCCACCCTTGTGTCTCCCTTTGG | 4 |
| wt-147780-147900 | 4083 | CCTTTGCGGTGCCAATTTCCGGTGGTTTCCCTTTTCCGCCCGTTTATCCAATAGCATGTAAGAGAGGTTGCCTAGATTTGGCAACTTTGAGGGAACGTTCCGTGTAGCTGGTGACCTAAC | 4 |
| wt-147900-148020 | 4084 | ACCCGCCCATCACCACCGGACAGATTCTGAACTTGTCCTGTGGTGTTTGGTGTGGTTTTGGGGTACGCAGGAGTACGTTGGAATGCTTTGGAGCCGAGAGGGATGGGCCCGCTTGTGCGC | 4 |
| wt-148020-148140 | 4085 | TTATGTGTTACACGGTGCCAATAACCGGCCCGGTGCGGCTGCCCCGTGACCCGTGGGCCTTACCTTCCTGGCCATCGGGGGACCCTGGTGCTAGGGTCCCTTGTGTTGCTTTCTGCCATA | 4 |
| wt-148140-148260 | 4086 | GGGGGGAAAGCATCGCCTTCAGAATTGGCTGCTCCGTTGGAACATTTGAGGCCTACTGTATCCGTGTCCTGACAACATTCCCCGCAAACATGACATGGGTTAATTTAAACATGTTTTGTT | 4 |
| wt-148260-148380 | 4087 | TGCTTGGGAATGCTCTTAGGGCCTGGAAGCTTGTCATTGGATTCATCGTTTCCTGAACTACAGGCGTAGGGCCTATTGTAGCAGGCATGTCTTCATTCCTGCGTACCGAATGGCATGAAG | 4 |
| wt-148380-148500 | 4088 | GCACAGCCTGTTACCATTGGCACCTTTTTTCCATGTAAACCTCCGTGATCCTGGGTCCTTTGGAGACTCAAGTGTGAATTTGTTTTGGTGTTCGGCGCCAGGGTATCTCGACGTTGGAAT | 4 |
| wt-148500-148620 | 4089 | GTCAACTCAACTTGGGCACCTCGATAACCGGCTCGTGGCTCGTACAGACGATTGTTTGGCTCTGTAACTTGCCAGGGACGGCTGACGATGTGTTTAGTCTGCCACTTGCATCCGGCGCTT | 4 |
| wt-148620-148740 | 4090 | TGGTTACTCGGGAGACTAATGGGGGGTGTGGTATGGCACAGGCTGGGGGTGAGTCTGGGGATGTCCCTGGGCGTTGCTGCAGCCCATTCGCCCTCTGGGGATGAGATGTTCAGGGGTGGC | 4 |
| wt-148740-148860 | 4091 | CGGTACCCTACGCTGCCGATTTACATAATATAAATTGTAAATGCTGCAGTAGTAGGGATCTGGACGCGCGACCTGCTACTCTTCGGAAACGCCAACCCAGGAGCGTCGCCTCTGGCCCCA | 4 |
| wt-148860-148980 | 4092 | TACTCCCGCCATGCGACTGCTCGCCCCCTCCCAGGCCTCCCTGGTGAGCCCTTGCCGCTCCCCGCATTCCTGCTTTCGGCGCCCCTGCGGATCCCGATGACAGCAGGCCTTTCCTTCCCC | 8 |
| wt-148980-149100 | 4093 | CGTTAATGAAAAGAATGACAGTGAGGTTGTGACAGAAGGACAGCTTTATTCAGTTTACAGAGTGCCCTCGGAGGCTACGATATTCCCGTTAAATGTCTTGTTGATTCTCTCAAAGGTGGG | 4 |
| wt-149100-149220 | 4094 | GAGGGAGGAGCTCTCCACAACAATGTTCCCTGGCAGCGTGAGCGCGCAGCCCTGCCGTTGGATGTATCTTCTCATGATGGTGCTGATAGAGGGGTCTCCGGCGTAGATGAAAAAGGCCTG | 4 |
| wt-149220-149340 | 4095 | GGCCATGCTCTGGCCGGTCACGATCGTTATGGGGTTGTTGGAAATGTTCCGGACCGTCAGCTTGAGGGTCTGGCCCGGCTTCCACTCCTGTGGGTAGACGTAGAAGACCGGGTTGGAGGA | 4 |
| wt-149340-149460 | 4096 | GTGGGACACGACAACGGCCGTAATCTTGGAGCTCAGGGGGGCCTCGTAGGTGTTGTTGTATTCCAGCTCCGTGATGAAATTAGGAGGAATAATCACAGGGGAGCCAAAGTAGCGGATGTC | 4 |
| wt-149460-149580 | 4097 | TGTGGATTCCCCGTCCCAGCGCCAGTGGCTCTTAGGGTAGGGGTTGTAACGGAAGGCAATAATCACATCATCCAATAGGGTCATGCCCACCTTGACGTTCAGCGGGCCCTCTCGTTTCAG | 4 |
| wt-149580-149700 | 4098 | GTCCGGCGTGTCCACGGAGACTCGGACGTAGCCCTTACCGCGGCGTATGGCGTTTACCGGACACACCTTCCCCGGGAATGTGTGAATACGGGCGTATGACTTTAGAAATGGGGGCGTGTG | 4 |
| wt-149700-149820 | 4099 | CTGCGCCAGCAGGTAAGGCAGGCACTCGTCCTGGCTGGTGACGGGAGAGCCACTGAGGAAGATCTGGGGCTCGCTGGTGTTTAGCTTGTCCCCGCTCTGGGTGCAGGAGCGTGTCAGCTG | 4 |
| wt-149820-149940 | 4100 | AATGTCGCTCTGCCCGGGCAGAATCTGCAGGTAGAGGTAGGGGTTCTTGACCAATCTGATGGGCACAATGTACCAGGTAAACTTCCCTTTCTCTATGAACAGGCTGCGCGGATTCAGGAC | 4 |
| wt-149940-150060 | 4101 | GCTTAGCACGATGTCCTGGTCAGAGTGCATAACGAAGAAGGGCTTGAGGAATACCTCGTTGTCTTCCGCTCCAAAGAACAAAAACGCGACCGTAAAGTAGCGGCTGCCGTAGGTGGTCGT | 4 |
| wt-150060-150180 | 4102 | GTTGAAGGAGAAAGAAGGTAACTTGAAGCTGAGTATCTGGCCCACCGAGGGGCAGGGAGGCAGCTCTTGGCACTGCGCGTCCAGCTGCAATACCTGCTTGTTGGTGACGCGGACGTATGA | 4 |
| wt-150180-150300 | 4103 | GGGGAAGATCTCGTACTTCCACACGCCTCTCATGAACGACGTGTCTGGTTTTTCAGTGGGCCGCAGGCGGCGGAGGCTGTTCCTGAACGACGAGCGCCGGGACGCTAGTGCTGCATGGGC | 4 |
| wt-150300-150420 | 4104 | TCCTCCGGGGTAAGCTTCGGCCATGGCCGGAGCTCGTCGACGGGCAAGGTGAGAGTCGGGGGGCGGGCGACGGTGCGGCCCCAATACAACTCTCCGCTCGTTAGCTGGTAGAATATCCGC | 8 |
| wt-150420-150540 | 4105 | CCGGCGTCTAGGTTGTCACTTCGCTCGGCCGGCCAGAAGAGCGCAAGTCCAAGTCTGGTGCTGGGGCCGATGTGCAGCGGTTTGTGCCCGCAGTTGTAGACTGTCATTTTTATGGGCGAG | 4 |
| wt-150540-150660 | 4106 | TGGGCGGTCCACACGCGCGGGCGCAGCACCCATTGGTCGCACGCCGCCTCCTGGAATGTAAACCCCCAGAGAGAGGGCGTGCCGCCCTGGAGATGGCCCTGTGCCATCACATGTATTTCC | 8 |
| wt-150660-150780 | 4107 | TCCTTGGGTGGAACAACGGCGTCGTGCTCCGGGTGGAGGGGGAATAGCGTCCAGGCATCTTTCAGGGTCACGAGACCGGGGTCCATGCTCAGAGAACAGCCCTCCCGGGCGGTGGGCGGC | 8 |
| wt-150780-150900 | 4108 | CCGGGCTCCAGCAGAACGTCGCAGACCCAGCCCTCCTCGGCCCTGTCCACCTGTATGTCCAGGTGCACGGACCCGGAGGCTGCGTCTCGTGACATGGCCAGGCCTGGTGCCAGCCGACCA | 8 |
| wt-150900-151020 | 4109 | CGTCCCGTGTCCCAGCCGAGGCCGCGCCAGAGCAGAGCCCGGGACTGACTCAGGGCCACATCCCCTCGGCCCGCGGACGCCGCCTCGCCAGCCCCCGGGCCTTCATGGGCCCGCTTTCTA | 8 |
| wt-151020-151140 | 4110 | CCTCTCTCCGGCACCCCAGCCTGGTCAGCCGCAGAGGAAGCATGACCTTGGGGTGGGACGGGGCAGGCGTGATCCTGGGCGCAATCTTTGCCGATCCCCACACCTTCACTCCTTGTTAGG | 4 |
| wt-151140-151260 | 4111 | TTGATAGAATGTCGGTACCACGCCACGGGGGGCGGGCCCGCATAGGGAAAAGCCAGGGAGAGCGATGTGGGCGAGGATGGGCTCAGGCGGCCCCAGACACGCAATTTGCCCCCCTGGGCG | 8 |

| | | | |
|---|---|---|---|
| wt-151260-151380 | 4112 | GCCGCAGCCTGCCCCTCGGCGGCCCGTGCCCCAGCTCCGTCACGGGGGGCGCATAGGAGGGGTATATCTAGGATAGCCGCACCTACACAAATGAGACACAGACACAGGTCGTGAGGATTT | 4 |
| wt-151380-151500 | 4113 | AGGCAACGCAGGCTTGTCTTTATAGTTACAAACATGGGAGCGTGCACCTGGAAGATGCAGCTGGGGTAGATCTTTACATCTTTACAGGGCGCAGCGGCCGCCAGACACTGAAGGGCAGAG | 4 |
| wt-151500-151620 | 4114 | TTCACGGCGGGCACCTCCCAGAGGGAGCCCACCAGCCCGTACCTGGCCACGGCCAGGGCCTCGTAGGCCGAGACGGGCAGCCGGAGCTTGTGGTACTGTCCCTCCGGCAGGTGGAGTGGG | 8 |
| wt-151620-151740 | 4115 | ACACAGTTAGAGAACATTAGTCCTCTGGTCCCTATCTCCACCCGCCAGGCCTGTGTGTCAGTTTGCAGGGCCATCCTCGCACTCAGGTGGACTGGCTAGGCACCCTTCTGAAGTATCTGG | 4 |
| wt-151740-151860 | 4116 | CGGTGACTGTCACCTGGTTCTTGAGAGAGTCCATAAAATGGCTGAAGCTCCAGGCGTATAGTATAATGAGCAACAGGGCCAAACAGGCGGCGGGGCCTGGGTAGTAGCGGGCAACGAGAG | 4 |
| wt-151860-151980 | 4117 | ACTCTGTGCAATCAAACCCCAGGCTCCCGGCCTCACCCAGGAAGAGCAGCGGCAGGGACAGCATAAACCAGGAGAAGGCGCAAATGAGTCCGGTGAAGGTGACGTTGCATATCAGGCGCG | 4 |
| wt-151980-152100 | 4118 | GCTTCCTTCCGAATTTTGTGCGCAAAAGTTTCCAGATGATGATAACTGTGAGGAGGACGATCAGGACTGCCGCCAGTAGGTAGCAGCCGGCTTTCAGTCCTTGGACGGCCGTGTGCATGC | 4 |
| wt-152100-152220 | 4119 | CTTTGGTGGGGCCTTCCCTGCACATGTTGGGGCCTCTGTTGAGATTGGCGTCGGGGCCCATGGTAATGAGGAGGATGATAATCAGCAGGAGTACCAGACAAAACACGCCCATCAGGTACA | 4 |
| wt-152220-152340 | 4120 | GGCACACATTTCTGTGGGAGGTTCTCTTGGGCGTTCGGCTGAACAATGCTAGGGTCTTCTCCAACGCCATACCCAAGTGAGTCCATACGGAGCACATCAGGCCCAAGAACATCATGTTCT | 4 |
| wt-152340-152460 | 4121 | GGGTCAAAAGGCAGAGACCGGTAGACGAGAACTCCTGAATCATTTTTCCCAGCACCCAGAGCAGCAGTTCTATGAGAAGAGCTATCAGCCAGACATCCATTCGGTGAACCAATTTTCTTA | 4 |
| wt-152460-152580 | 4122 | CAAAGATGATAAACAAGATGCCAGCCAGTGTTAGCAGAATCAGCAGGACGAGCAGCAGGCTTGTCATGCCGCTGAGGAAGGCGCTGTAGGATTTAGTGCACGCATCTTCCGTTGCATTGA | 4 |
| wt-152580-152700 | 4123 | CGGAAGTCATGTTGGCCACCAGGGTCCCCACGGTGGACCCGGGGGCCATGGTGGAGAGCATCTTGCTGGTCAGAGCCAGACTGGGTGGTGTCTGCAGCAAAAGAGGAACTTGCCCAGGCA | 4 |
| wt-152700-152820 | 4124 | GTCAGTTATTTTGCATGCCACCTCCCTGCCTGGTGGACTTCCAGACTATTTTCTGCATTCGCCCTTGCGTGTCCATTGTTGCAAGGAGCGATTTGGAGAAAATAAACTGTGAGTTTCACA | 4 |
| wt-152820-152940 | 4125 | GATCCACGGGCCACGCTCCCCTGGGGGCTTCATGATCCCACCGCCTTTCCCGATGATGATGACAACCGCGGCTGTCTGAAGCGGCTGACGAAATCGGTTGAGATTCTGATGAGAGGCTTG | 4 |
| wt-152940-153060 | 4126 | GGGGGGTCTTTGCCCTCAAGGCGAGGCTCCTTCTCCTAGGAATGCCGAGCCCCCTGCACTAGCTTCGCTCCACTGGGGATCTTTGCCAGCCTTCATACTAGATTCAGCGATCCCCCGGTT | 4 |
| wt-153060-153180 | 4127 | GGGAATCTTCGCCAGCCCCCCGTCCTGCTATCCCGCTCGTCGCCGCGCCTCCCATGCTAAGGGCCCCCTTCCTTTCCCTTGACTTTGGGGATATTCGGAGTCTGCTCTCGCCGCTCTCTT | 4 |
| wt-153180-153300 | 4128 | CTCTCGTTTAAACGAGAGAATAGTAGTAGGGTCCAGTCTCAGGCCCCCTCACTTTGGGTCTTAGAATGGTGGCCGGGCTGTAAAATTCTGGAGGACGGAGAGGGCGGCCCCGGAGTTGTT | 4 |
| wt-153300-153420 | 4129 | ATCAAAGAGGCACTGGAGGATGTTGGCCGCTCCTTGGAGCAGCTTGTCGAAATAATGATCCACGGCCACGGGAACGCCGTGCCGCTCGGCGTAGGCCGGGTCCTCGGCCATCTCCGTCTT | 4 |
| wt-153420-153540 | 4130 | TCTCGCCCCCTTCACTCCCCCCTTGGGCTCCACAAAGACGTACTGGATGCGGTCGTGGATCTGGGGCAGTTCCTCGTTGCGCTCGACGAACTTCTGGTAGACGGCCAGGTGAGGCATCTG | 4 |
| wt-153540-153660 | 4131 | GGTGCTCTTGTAGGCTGAGAGCTTGCGGCTGAGCTCCGTTGAAAAGCAGAGCTCCCCCATGGGGACCCTGCCTTCACGGAGGTCTGTGTAGGCCTGGTTTAGGATGTCAATGACGGGCAA | 4 |
| wt-153780-153900 | 4133 | CTGCACAAACTTGCAGGCCGTCTTCCGGACGAGCTCCACCCCCTTCATCAGGGTCTTGCCGTCCGTCAGCACCCCCACATATCTCTTCTTTGTAATCAGCATCAGGCAGGAGAAGGTCTT | 4 |
| wt-153900-154020 | 4134 | CTCGGCCTCCAGGGAGATGGGGGCCACAAACAGGCTCCGGGTGGTGTGGGCGGCCAGGGCATCGGCAAAGCGCAGGGTCTCGCTCTCTGAAAACCCCCGGCACTCGATAAACAGCGAGTC | 8 |
| wt-154020-154140 | 4135 | CGTGTCCCCGTAGATGACTCGAAGCTGGCCCTCGGGGTTGAGGGGCGCCCAGGCGTCCGGGGAGGGGGCCAGGGCCTGCAGGTTGGCGGGGCTCAGGGCCTCCACGAAGGCCTTGGCCCG | 8 |
| wt-154140-154260 | 4136 | CTCCAACATCGTGCGGCCCTGCAGCGTCACCGTCTCGGCGATGGAGAGGCAGGGAAAGAGGCCGTTGGCCACCCCGGTGAAGCCGTAGACGGCGTTGCACGTGCACTTGATGGCCAGCTG | 8 |
| wt-154260-154380 | 4137 | CTGCTTGTCGAGGATGGTCCTTTGGCGCGGATCCTCGCAGGCCGCCAGCAGCTTCTTGATGGCCTTGCGCTTGGCCAGCCAGGAGGTCAACAGACTAGCCAAGAAGGACTCGTGCACGTG | 4 |
| wt-154380-154500 | 4138 | CTTCTTTACAAAGTGGTAGACGCCCCCCGTGAGCCTGAAGGACTCATAGTCTTCTCCCGGGCGCAGGCCGGCTAGCCTGTGCTCTTCTCCCGGCGTTATCATGGTAGAATAACAGAGATT | 4 |
| wt-154500-154620 | 4139 | ATGAGCCTGAATGATGCTCGGGTAGAGGCTGGCAAAGTCCACCACCAGAACCGGGGAGTTGTAGAATCCGGACAGGGGCTGGATGACGGTGGCCCCCTGGTAGCCGTCCCGGTCAGAGGC | 4 |
| wt-154620-154740 | 4140 | CGAGGGCATGGGCAGGATAAAGTTTTCCTTTTGGGCGGCCGCCAGGAGGCAGGAGAACACGCGGATCTGCTGCCCATCGTCCAGCACCCGCCTGCAGGGGATGTGAGCGATCTTGGCAAT | 4 |
| wt-154860-154980 | 4142 | AATCTCCTTGTAATGCACATCCTCCTTCTTGGCCCCCAGTAGGTGCCTGGCTACTGTGTCCAGCTTGTAGTCTGAGAGGCTGAGCTTGTCCCGGCACACGGCGTACATGTCGATGGGGAT | 4 |
| wt-154980-155100 | 4143 | GAGGCCGGTGATGCGGACCTTGGTGTTGGCCCGCAAGAAGCCCTTGCCCGCATCATGGGGTCGCCTGACCTCGCAGACGCCCCCAGCCCTAATTTTGCCCAGAGAGGCTGGGTTGATGCT | 4 |
| wt-155100-155220 | 4144 | GTAGATGTGCCTGGCTCTGTCCAGAATGTAGGGCCAGTCAAAGTTGGCCACGTTGTAGCCGGTCACAATCTCCACGCTGAGGTCTCTGATGAGCTGGAAGAAGGCGTAGAGCATGTCCAG | 4 |
| wt-155220-155340 | 4145 | CTCCGATGGGAACTCGTAGACCTCAACCCCCTCTATGTCTTCGCAGGTGCCCAGCGTCAGCAGGATGCGCCTATAGCGCCCGGCCTCCTCCCCTGTCGACCAGAGGACGCAGGATATCTG | 4 |
| wt-155340-155460 | 4146 | CAGGATCAGGTCAGCCTCGTTGGTGGCCGTGGGGAAGCCCTCCTCCCCCAGACACTCGATATCGAAGGCCAGGGCCTGGTAGGAGGGCCAGGAGCTGTCTTCACGCCGGACCGAGAGGTC | 8 |
| wt-155580-155700 | 4148 | CGTGGCATCCACGTTGGCCTCAAAGATCCGACACCCGTGCTTGTCTTGCAGCCACGTGGCCACGTGACACACACTGTTGGGATGGGAGAGGGTGATCTTGTGGTAGTCGCCGGCATGGTT | 4 |
| wt-155700-155820 | 4149 | GCCGTAGCCCATAATGGAACGGCGCGTGACCTTCTCCACCGAGACCCGGCAGGGGGTCCTGCGGTCGAAGGTGCTGGCCTTGAGGGCGCTGAGGACTGCAAACTCCACGTCCAGACCCTG | 8 |
| wt-155820-155940 | 4150 | AGGCGCGCTGGCGTAGAAGTAGGCCTGCTGCCCAAACACGTTCACACACACGCTGGCCCCATCGGCCTTGCGCCGGCCCAGTAGCTTGATGACGATGCCACATGGCACCACATACCCCTG | 4 |
| wt-155940-156060 | 4151 | TTTATCCGATGGAATGACGGCGCATTTCTCGTGCGTGTACACCGTCTCGAGTATGTCGTAGACATGGAAGTCCAGAGGGCTTCCGTGGGTGTCTGCCTCCGGCCTTGCCGTGCCCTCTTG | 4 |
| wt-156060-156180 | 4152 | GGCACGCTGGCGCCACCACATGCCCTTTCCATCCTCGTCACCCCCCACCACCGTCAGGGAGTCTTGGTAGAAGCACAGGGGGGGCTGAGGCCCCCGCACATCCACCACCCCTGCGGCGCC | 8 |
| wt-156180-156300 | 4153 | TGGTGTCTGGAAACACTTGGGAATGAGACGCAGGTACTCCTTGTCAGGCTTTTTCAGAAGGCCTTTATTAGGTCTTAGGAAAGGGTTATAGAAGAGTCCCCCAGACATGGTTAAAACTCA | 4 |
| wt-156420-156540 | 4155 | GCCCTCTGAGCTGCTCTCTTTTGCTCCTGGTTTTGCTCATGCAGCGCTAACATGATGGCTTGTAATTCTGTCTTACTAATGGGATTAATGCCTGGACCCTCACCAGAGGCATGTTGCTGA | 4 |
| wt-156540-156660 | 4156 | GCGAGCTCGTCGATCCCGGGGTAGAGCATCTGCACCGGCTGCTGCGACATCTGGCGCGTGCGCCTCGTGAGGGAAATAACCAGGATCACCACCCCCGCCACCAGGACCAGAATGAGCATG | 4 |
| wt-156660-156780 | 4157 | CCGCCGAAGGGGTTTTTGAAGAAGGAGATGAAACCAGAGACCAGGCTGCTAAACAAACCCCCCACCGTGCTGACTAGGTTGGTGATGGACTGACCCACGCTACCCAGACTGTCCATAAGT | 4 |
| wt-156780-156900 | 4158 | TCCCCCAGGCCGTCCACGAATTGATTTCTTCCGTTTGACACTGCATTGTCCAAATCCTTCCGCAGGCCGGCGATGTTTTGCGCCTGGAAGTTGTACTCCCGGAAGATGCCCTCCAGGTCA | 4 |
| wt-156900-157020 | 4159 | AAGACGTTGGAGGCACGCTGTTCGTCCCGTGAGTACAGCTCCAGGGAGGCAAAGTCAATGTTCTCGATGAGGGAGGTGTTTAGTGAGATGAAGGTCTGCAGGGTGGCAATGCCGTCCAGC | 4 |
| wt-157020-157140 | 4160 | TCGATGGTTTTAAAGTGGTGGTAGTCGTTGTAGACGTGGATCTCGTTGCCGGACTGGAAGTAGTACTGGCTGGTCGCCTGGCACACCTCCGTCATCTTTTTTGTGAGGAAGATCTCGTTG | 4 |
| wt-157140-157260 | 4161 | TCGGTGCCCAGCTGTCCCTCGTAGGTCTTGGTGTCGTTGATAAAGCTGAAGGACACCAGGGGGCGCGAGTAGCACATGGTCTCGGAGCCAGGGACCCTCATGCTCTTGCGCAGGGTGACG | 4 |
| wt-157260-157380 | 4162 | GTGGCCTGGTTAACGGGCACGCACTGGGAGACTGAGATGACATCCCCCAGGCGCTTGGCCGCCACCGCCTTACCGTAGATGCTGGACATGACGGTGGTTGGATTAATCTTGGTTAGTTCT | 4 |
| wt-157380-157500 | 4163 | CTCAGCACCATGTTCTGCCTCTTCTGCTCCAGGCACCAGGCCCGCGCAAGGTCTCCCAGCATGCGGTTGATCTGGCGGCGCAGGGAGTCGTAGGCAAATTGGATCTGGACGGTGGCGGGA | 4 |
| wt-157500-157620 | 4164 | TTGTTGAGGGTGCCCAGGGACTTCCCGGGGGCCGTGGGGGGCACCGGTGTGGTGGCGTTCCCCGCATCCCGCCTCCGACGCCTCAGAACGGCGGCGGGGGTGCTCCCGCGGGCCGCGGAT | 12 |
| wt-157620-157740 | 4165 | GGGGCTGGGGGCGATGGACTGCTGGGGGGTGAGGAAGTCGGAGTGGTAAGCTCCGTCAGGTTCTTGACGGTGGCCAACGAGCGCGGGGTCAGAGGTAGCCAAGCTAATAACAATCCTCCG | 4 |
| wt-157740-157860 | 4166 | CTCGTTATAAAATATGTAATGGCTTCCTGGCCCTTCGTGTAACGATCCTGGACGGCCTCGTACTTCTCATGCATGGTCTTGTTCACCTGCTCTTCGATGCACTTGAAGGCGTCCGGGAGC | 4 |

| | | | |
|---|---|---|---|
| wt-157860-157980 | 4167 | TCTATGCCCACGGTTGTGTTGGTCACGAAGCTAGAGGTGCCCTCGTCAGTCACAAAATGTATTGACTTCCCTGTTTCTGTGGCGATGGTCGAGTCAAAGGTTTGCCAGTGTTGAAGCGGG | 4 |
| wt-157980-158100 | 4168 | CAGTAGGCTGTCCTGTTCTCGAGCTTCCAAGATAGCGTGTAAGTGCCCTTGTCCAGGAAGGCTCGGCGTTCGCCTTGCGGGTTCGTCCCTCGGTTGTCGTAGTCCACTATCTTGTAGTTA | 4 |
| wt-158100-158220 | 4169 | GTTCTCACGTGGAAGGAGTCTGCCCGCTCATGGAAGGTTTCCTTATTTTTCCCGTCATAGAAAGGGGACATTTCCACAGTCTGCCCGGTGGTGGTCACAAAGAAGTCGAAGGGGCTGTTG | 4 |
| wt-158220-158340 | 4170 | GACTTGGCCATCATGTCAGTTATCAGGCAGTTGACGGTAGTTCTTGTTCTGTAAGTCCATATCAACCACCCGGGGGCGTCATAGAGCTCCGTCTGGCTGGCGTAGCGGCGCACCCCGTTG | 4 |
| wt-158340-158460 | 4171 | GCCAGGCCCCCGGTGGGCTTTAGGTTGACGGTGATGTTAACTCCGTCGCGGTCTACATACACGCGCGTCAGCCCATCTTTTGTCATCTTGACCGCGTTGTAGCACTGGTAGATGGTATCC | 4 |
| wt-158460-158580 | 4172 | ATCTGGTCAGTTTCGTAGCTGTCAACGGAGAACTTCTCCTCGTGCCGGTTGGTCACGGAGTCCGCGTACCAGCCATTGTAGATGAGAATGTTGGTCACTATCTTGGTGTAGGAGCGGACC | 4 |
| wt-158580-158700 | 4173 | TTAAACGAGTAGGGAATAATGTTGTCTTTAAACACCATCAACAGGCCCTCCGTGTGATTCTCCCGCGTGCCAAACGAGGGACACTGGATGTCCGAGGAGAAGCGGAACAGGTCGCCGTGG | 4 |
| wt-158700-158820 | 4174 | CTGGAGAGCTCGCAGACTCGGAAAGGAAAGCTGGTTTGCTGACGCGTGGCGGTAGGCTGCACCGTGGTGGCGGGGGGTGCGGGCTGCTCTGGGGTCTGCGCACCGAGACGGCACGCCAGG | 8 |
| wt-158820-158940 | 4175 | GCGGCTAGCAGCACGACCACGCTTAGCACCCTACGCCGAGTCATCTCTCATTTGGAGGTGCAGGTAGAGAAGGGCATATAGATCCTTAAATACCCACCCCCTGCCCTTATACAGAAGAAT | 4 |
| wt-158940-159060 | 4176 | TAGGGGGCGGTCAGAGTCGTACGTGAGGTAAAGCCCATCCGGGGGCAGGGCCTGGCCGGGGCTGACCGCGTCCGCCCGGCGCAGGATCAAGGACCGCCCCCAGGTCTTGTTGTAGAGGGA | 8 |
| wt-159060-159180 | 4177 | CACGGTTAGGACGGCCTCGCGCAGCGCCCGGCACAGAATTTGCTGGCTAGATGCCAGTGAGCCCCCGGGTACGCTGTAGAAGCTGTTGAAGGAGGTCTCTATCCAGTCGCTCGGCTCGAT | 4 |
| wt-159180-159300 | 4178 | GCCTGGCCATATCAGGGAAGTCAGGAATGCCTTCTGGTGGGGCAGCGTACCTGCGGCGTCACAGCAGCGAGCCAGGGCCACGTTGCTGGGTGGGGGAAAGAGCCCGCTCTCCTCCGCCAG | 8 |
| wt-159300-159420 | 4179 | GGGCCCCGTGATGAAGGTGTACAGGCTGTGCGTCAGCGCGTGCAGGTGCTCCGAGCTCAGGGTCTGGGTAAACAGGTGTGTTTTGATGTACTTGGAATTCTCAAAGGCGGCACCCTCGCC | 4 |
| wt-159420-159540 | 4180 | GGCGCGCCTGTCCTCCCAGGGACCCGAGACGAAGGCCCGTCTGTAGAGGAAGTGGTTGCGCATGCGGGCCAGCTCCCAGTAGACCACGTCCCCCCAGACGCGCAGGCACAGGGTCTCGGT | 8 |
| wt-159540-159660 | 4181 | CAGGGTCTCGCTCTGTTGCGCCAGGCAGGACTGCAGCTTGGCCAGACCCTCGGTGGCCACCTGGCGCAGGTACTGCTCCTTGCGCTTGAGCGCGTCCGAGAGGGCGCCGGACGGGCCGGG | 8 |
| wt-159660-159780 | 4182 | CTCTCGTGCCCCAGCCGGCCGGGGCACCTCCGGGCTCTCCCGGGACGCCTCCTCCTCGCCTCGGCCCAACCGCTGCATGGCTCGGTTGAGCCGCGTGTACAGCTCGTTCCTCTTTTGCAG | 8 |
| wt-159780-159900 | 4183 | GATGGCCCGGTACTGGGGGTGCGCCGTGAAGGCGGCGGCGCAGTCCGCCTTCAGCGCCTCCACCGCGTCGCCCGAGGAGCTGTAGACCCCGCCGCAGAAGAGCCGCTCCGTGGCCCCGGG | 12 |
| wt-159900-160020 | 4184 | AGCCACGGCGTCAAACAGGTGAGTCAGCCTTGCCCCCGCCAGCGCCTCCTCGCAGGCCCCCCGCACCAGGGCCAGGCGACGCTCCCGGGCAAACAGGGCAGAGAGGCGGGAATGGCCGCC | 8 |
| wt-160020-160140 | 4185 | ACCCTCCCCCTGCCCCGTTGCACCGATAGCATGGCCGCCAGAGTTCCAATAGAGGAGCTCCGAGAGCTCCGCCACCTCCGGGGGCACTGTCGAGAAGACGTTGTAGGTGTCCAGCGCTCT | 4 |
| wt-160140-160260 | 4186 | GGTCGCCCCCTCTGCCTCCGGCCGCCCCGGGCCCGGGACCGCGCCCTCCTCTGGGCCGCCCGGCCTCGCCTTCTCCTCAGCCTCCAACAGGTGCCCGAGCCCCGCCTGGCGGACTTCATT | 12 |
| wt-160380-160500 | 4188 | ACAGCGCCAGCAGGGCACAGAGGTGATGTAGAGGCGCGGGTCTGGGATGGGACTTACGCCCCGAAAGCGGCCCAGCAGATCCAGGGCCCGTTCCAGGCTCTCCAGCCCCATGGTGTGAGA | 8 |
| wt-160500-160620 | 4189 | CATGCAATAAAACACGCTATTGATTCTCTTCATTAAAATCTCTATGTCATTTATTAGGCACAAACTTACATCGACTTTATGCCCCCCGTAAAACTCCACAGAGTACGCGACTGAGGGGGT | 4 |
| wt-160620-160740 | 4190 | ACGGAGAGGCGGGACCCGGGTACCCTTTCTACCAGGGGCGAGCAGCGCGGCAGAGGCCTCTCTCGAGTTCTCTAGCAGGTGCACCAGCTCCAGGGACAGGGCGCTGCATGCACGGTCATT | 8 |
| wt-160740-160860 | 4191 | CTGCCGTCTCAAACGGGGAAGGAGGATGGCCTCCAGCTCGGCCAGCAGGCCGGCGTTGCGCACCACCGCAGCCACGTCCAGACTCCGGGGGTCCAGCCGGGTGCACACGCTCAGCTCAAC | 8 |
| wt-160860-160980 | 4192 | CGCCAGGGCGTACACCTGGCTGTACGCCGCCGCCAGCAGCCCCGACATCGCCGCCCCAGGGGTCTCTAGACCTCGAGTCCGGGGAGAACGGTGGCCAGACGGCGCTTGCGTCTGCCCCCG | 8 |
| wt-160980-161100 | 4193 | GAGCCCTGCCCTCCTCCACCCAGCAGCAGCCCGGCCGAGGCCTGCGACGCGGTGCTGACCGGCTCGGCCACGCTGATAAAGTTGTCCTGGGCTGCCCCGGGCCCACCCCACACTCCCTCC | 8 |
| wt-161100-161220 | 4194 | AGAAAGTCCCGAGCGGCCTCCGCCGTCCACTCTATCCCGCTGGAGGCAATGGTCGCCAGGGTTTCTAGGACGCTGTCCGCCAGGACGGAGAAGCGGCCCAATAAGTACTCCGCGTCGTCC | 4 |
| wt-161220-161340 | 4195 | CTAGTCAGCGAGGCGCATGCCTCGCCCATGGCATCCACAAGGTTGCACACCACATCAAACACACAGTCTTCCTCCTGTTTTTGTGATATAATGGCCTCCAGGCCAGCCCTGATGTTCTCA | 4 |
| wt-161340-161460 | 4196 | ATCTCATATGTGGTCGCGGCTTGGGTCCGGCGCTTCACGGTCAACCCTAGGGTGGGGGTGGCAAAGACAAACTTCTTCCGCATGGAAGAGCCCCCGGCCTGCTTGCGCAGCCCAGCCCCG | 4 |
| wt-161460-161580 | 4197 | GGGGCCTGCAGCAGGTTCCTGTCCACGCCCCGGCCCATAAAGTATCCCAGGTTCCCGGCCTGGAATATCTGGTTGTTGCCGTTGACCCCCGTGTACTTGTTGATGGTCACTGGCAGCGTG | 4 |
| wt-161580-161700 | 4198 | ACAACCGGACGGGCCTTGCAGACCTGGCTAAGACAGTCTGTGGCCGCGCAGACCACCGTGGTCGCAGTAAGGGAGGAGGTGGCCTCCGCGTAGGCCGCTGCCGACTCCACCGCCCGCGTG | 8 |
| wt-161700-161820 | 4199 | CCCAGTACGTGGGGGTAGTCACGGGCGGGCACCGACTGCGTCCTCGGCACCAGTCCCTGAATCAGGCTGATGTAGAACTGGGTCTGGCCGCACGCCTTCAGGATGGCGTTGTTGAGCCTC | 4 |
| wt-161820-161940 | 4200 | TGCTTGGCGTAAGTGACCAGGTTGCCAGGCACCACATCTATGACGTTGCTCTCTTCGTGGGCCCGGGAGCCCCCGTCCACAAAGAGGGCCAGGTCAGAGTACTCCTCCGCGCTGGCCCCG | 4 |
| wt-161940-162060 | 4201 | CTGGGGACAGGGACCGAGCGCCGCCTGGAAAAGTTGTGCCACAGGTACAGGCTTGAGAGCTTAGTGTCCGGGAATAGGGTCTTGTGGTAGGTGTTGAGGAATTTCATGTAGGGCCCGTTG | 4 |
| wt-162060-162180 | 4202 | ATGATGTAGTTCTCCCTCCTGGTAGTGGACTTGATGAAGCTGTTCTGGAGGGCGGCATTCTCCCCCGTGAAGACCACCCTGTTCTTGATCTTGATGTTCCTGGGGCACAGCATCAGCACC | 4 |
| wt-162180-162300 | 4203 | TTGGACATGCGCACAGGCAGCCGCCGGCCGTACACCCGGCCCTGCAGGGCCGCGTCCAGGTCTGGCAGGTCGCAGGTGGGCTCCCCATGCACCACCTTGGCCTCCTTGGCCGTGAGGACC | 8 |
| wt-162300-162420 | 4204 | CCCTTGTCGATGGCCAGGCTCCTAAAGTTGGTGCACAGCGTCTGGTAGTGACCCTTTAGCCACTCTGGGGGGCTCTGGCCAAGCCCGGGGTTGTCATTCTCATAGCACATACAGATGGGC | 4 |
| wt-162420-162540 | 4205 | AGGGAGATGTCCTGCAGGATGGTCAGCAGTGAGCGGTAAAACAGCTGGGTGAAGATGGGGCAGGCGGGCTGCGCAAAGGGGTTGCACGAGTACTGCATCACGTGGTAGCAGCTCTTGACC | 4 |
| wt-162540-162660 | 4206 | AGGTCCTTGTAGGTGATGTTGTTCTTGGCCATGCTGTTCATAAACTGGACCACTTCGGCGTCCACCGCCGCATCCACGTCCTTGAACATCTTGACAAAGTCACGCGGGCCATGGGGCTCC | 4 |
| wt-162660-162780 | 4207 | TTCTCTAGCTTTCCTTCAGCGTCTATGCCCAGCCGAGACAGCCGCTCCAGCAGGTTCTGGTTCAGCTGCCAGTAGGTGTAGCGGGGCTCGTCGTCCGGCCGCTGCCCGTCGTCCTCCTTA | 4 |
| wt-162780-162900 | 4208 | TCGATGAAGTTGAGAAAGTTGCCCAAAAAGTCCGTCTCGTTGTAGGAGCCCGAGGCCCCCGAGATCACATAGGGGTCCCTCCGCTGCGTGGACATGACGGGGGGGAAGCGGTCCCTCAGC | 4 |
| wt-162900-163020 | 4209 | CTAAAGAAGAGCGTGTTCAGGCACACGGCCGGGGCCCGGCCCTCGCAGAGCGAGCACATGGGACTGGCGGCCGCCCCCGCCACGTAGCTGCCCGTCTCCGGCACCGGGGTCAGAGAGCTC | 8 |
| wt-163020-163140 | 4210 | TTCTGTCCCTGGCAAAACTGCAGGTAGTAGGCATAGCGGGCAAGAAGGTTGGGCGAGAAGGAGGCCGCATAGACCAGGTGCTCCACAGCGTAGTTTCCCGGACCGTTGGTTCCGGTCACG | 4 |
| wt-163140-163260 | 4211 | TCTGGCCCACCCCAGCCCGAGAAGCAGGGTCGGCGGCAGGGGTCCCAGGTCCCCTCCTGCAGGGTCCCCAGGCCGTGGGTCATGTAGAAACTGTTAAAGAGACTCTCCTTGCCCTGACCG | 8 |
| wt-163260-163380 | 4212 | GTTGACTTCGAGACCCCCGAGACGTAGAGGACGGAATTGGTGGCAAAGATCTGCGTGGACACGTGGGGGGCCAGGCTGGCATTATATCGGTGTAACGCAGCCACACGGGCCTCTGGACCC | 4 |
| wt-163380-163500 | 4213 | TCACAGTCGGCAAACAGGGGCCACGAGTCGTAGTTGAGGCTGGCCGGGGTCTCGTGCGAGGCCTCCAGCATGGCGGGTGCGTAGCTCACCGCCAGCTCGCATGCCGCGCTGTCCACAATC | 8 |
| wt-163500-163620 | 4214 | ATTAAGGCTCCCGAGTCCGGGTGACTGATGGTTGAGGCTGGGAACTCCTTGAGGGGGGCCACCTTGGCCACCTTGGCCTGGTCCTGCAGGCTCTGCTTCTCCAGCAGCTCCACCAGCTTG | 4 |
| wt-163620-163740 | 4215 | CCCACCCGTCGGACGCGCAGCGCCTGCGCCAGCCCGGTGTACAGCGCCTCGTGCATGCAGCGGCTGAGGTCCGAGTTGTAAAACTGGCGGAGCTGGGGCACGCCCTCTGGGAACACCTCC | 8 |
| wt-163740-163860 | 4216 | TTGTCGTAGAGCGGGACCCTAACGCTCGCAGACTGCCCCACCGCTACCTCCTGTTTTAACGATGGAATGGCCACCAGGTTTCCGCTGTAGAGTCGCTCCTTGAAGGCCTCGGTTATTGCC | 4 |
| wt-163980-164100 | 4218 | AGGGCCTCGAGACCCGGCCCCTCAAAGATGGGGGGAACCATATGGGCATTGTGGAACACGTAGATGTCCCTGTGATAGGAGGTAGCGCGTAGGAGCCCGCAGTTGGGGTCGGGCCTCCTG | 4 |
| wt-164100-164220 | 4219 | TGCAGAGCCTTGACATTGATGCTGAAGCCCGGCTCCACGGTGATGCCGCAAAGGAGCGGCACCGTCAGGCACCTGTGGCCCGCGTAGCCGGTCCCCAGTGTGGCCACCTCCCTAAGAGGG | 8 |

| | | | |
|---|---|---|---|
| wt-164220-164340 | 4220 | TAGGTGGCCAGGGGGTAAAAGTAGATGTAGCCGCACGGACCCGGCTGGCTCTGGCTGCCCAGATTATCCTCGCTAGTCTGTGCACCCTGCATGATGCCCAAGGTATCGCCCCGGCCTCCC | 4 |
| wt-164340-164460 | 4221 | AGTCCCACATTAAATGTTACACTTTACTCATCACGCAACACCCACTGTTTATTCATTTACAAAGATTTCAGGAAGTCAGTCAGGCTGGCCAGGGCCCACGTCACGGGGAACTGACGTCTC | 4 |
| wt-164460-164580 | 4222 | AGCGATCTTGGCATGCCGCCCAGCCTCGCAAACCAGAGTCTGCGATAGAGGGCCAGGTAGTGGGCGATTGCCCCCAGCACGAAGGCGGCGCTCTTGTGGTCATCCAGGTAGTTTCGCACC | 4 |
| wt-164580-164700 | 4223 | GCAAACACCACTGTGTAGCACAGCACCACCCTGAGCCGCGACCAGTAGTCGTAGTGGTCGTTGTACACTGCGCGCAGGACGCTGATGATGAGCCGTACGTGCGTGTCTTTGCCCCCGATG | 4 |
| wt-164700-164820 | 4224 | TCGGCTGTCCTGCAGGCCAGCTCCGCGTACAGCTTCCTATCCTTCCTCAGGGAGGCCTTGATGAGCCGGCAGAGGACCAGGGCTGGCAAAGGCAGGTCTTTCTCATCCCGGGTGAACACC | 4 |
| wt-164820-164940 | 4225 | GCGTACATGGCCCTGAACATGAGGTAGCTGGACTCAGCCACCTTGTCGTCCGGCGGCGAGGGCGCGACCCACGCCTCGACCGGGGTCCTCACAAACACAGAATCTGTAGACTTGGCTGGC | 4 |
| wt-164940-165060 | 4226 | CTCATGGTCTCGTCAGGCCAGCTCACGGGCTTCAGGCTTATATGATAAAATGGGCGTGGCAGAATAGTATAAGACGCGAGGCCTGGGTGAGGAGAGTCCAGAGCAATGGCCAGGTTCATC | 4 |
| wt-165060-165180 | 4227 | GCTCAGCTCCTCCTGTTGGCCTCCTGTGTGGCCGCCGGCCAGGCTGTCACCGCTTTCTTGGGTGAGCGAGTCACCCTGACCTCCTACTGGAGGAGGGTGAGCCTCGGTCCAGAGATTGAG | 4 |
| wt-165180-165300 | 4228 | GTCAGCTGGTTTAAACTGGGCCCAGGAGAGGAGCAGGTGCTTATTGGGCGCATGCACCACGATGTCATCTTTATAGAGTGGCCTTTCAGGGGCTTCTTTGATATCCACAGAAGTGCCAAC | 4 |
| wt-165300-165420 | 4229 | ACCTTCTTTTTAGTAGTCACCGCTGCCAACATCTCCCATGACGGCAACTACCTGTGCCGCATGAAACTGGGCGAGACCGAGGTCACCAAGCAGGAACACCTGAGCGTGGTGAAGCCTCTA | 4 |
| wt-165420-165540 | 4230 | ACGCTGTCTGTCCACTCCGAAAGGTCTCAGTTCCCAGACTTCTCTGTCCTTACTGTGACATGCACCGTGAATGCATTTCCCCATCCCCACGTCCAGTGGCTCATGCCCGAGGGCGTGGAG | 4 |
| wt-165540-165660 | 4231 | CCCGCACCAACTGCGGCAAATGGCGGTGTTATGAAGGAAAAGGATGGGAGCCTCTCTGTTGCTGTTGACCTGTCACTTCCCAAGCCCTGGCACCTGCCAGTGACCTGCGTTGGGAAAAAT | 4 |
| wt-165660-165780 | 4232 | GACAAGGAGGAAGCCCACGGGGTTTATGTTTCTGGATACTTGTCGCAATAAACGCACTTGCCTATTTCACCTTGTTTTAGTGTGGCATTGGGGGGGTGGCATTGCGGGTGGATAGCCTCG | 4 |
| wt-165780-165900 | 4233 | CGACTCGTGGGAAAATGGGCGGAAGGGCACCGTGGGAAAATAGTTCCAGGTGACAGCAGCAGTGTGTGAAGATTGTCACAGCTGCTGGTTTGGAGAAAACGGGGGTGGGCGGTGATCAGG | 4 |
| wt-165900-166020 | 4234 | GAGAACAATTCCCCGGGGACACCTGCACGAGACCCCTGGGCTCTCAGGAACTCCGCCCAGGTCTTGCCAATTGGGGTGATCCTGTAGCGCCGCGGTTTCAGCATCACAGGTTATTTTGCC | 4 |
| wt-166020-166140 | 4235 | TGAAGCTTGCTGGGGCGTAAATCCCTCTCGCCTTGTTTCTCAGAGAGCATTTCAGGCCGGTTTTGCAGTCGCTGCTGCAGCTATGGGGTCCCTAGAAATGGTGCCAATGGGCGCGGGTCC | 4 |
| wt-166140-166260 | 4236 | CCCTAGCCCCGGCGGGGATCCGGATGGGTACGATGGCGGAAACAACTCCCAATATCCATCTGCTTCTGGCTCTTCTGGGAACACCCCCACCCCACCGAACGATGAGGAACGTGAATCTAA | 4 |
| wt-166260-166380 | 4237 | TGAAGAGCCCCCACCGCCTTATGAGGACCCATATTGGGGCAATGGCGACCGTCACTCGGACTATCAACCACTAGGAACCCAAGATCAAAGTCTGTACTTGGGATTGCAACACGACGGGAA | 4 |
| wt-166380-166500 | 4238 | TGACGGGCTCCCTCCCCCTCCCTACTCTCCACGGGATGACTCATCTCAACACATATACGAAGAAGCGGGCAGAGGAAGGTAAGAGTGCCATCTATCTGTACTTTTATTTATTGCATCACA | 4 |
| wt-166500-166620 | 4239 | AGTCACATCAATAATAAGGGCGCCATCTAGCGGGAGATGTTATCCACACCATCCCAATTCACATCTCAGGGACAACAGGTCAAAGTTCTTTGTTGACACCCCCAGCGCTGGCTCCAGGGG | 4 |
| wt-166620-166740 | 4240 | GTGGAAGCGTTGGATGCAGTCCTCCGCATCGGGGCGGACGCCTCCTCCCAACGCGTTTCTGCGGATCAGTCGCTGGCTGGTGGGCATCGGAGTCGGTGGGCGGTCCTCCACGGGGACACG | 8 |
| wt-166740-166860 | 4241 | CTCCTTCTTGGCCTTGTTCTTTGACCTTTTGGACATTCTTCTGAAGGAACGGCGGAGAGTAGCGTAGAATCCAGCCAGTGGTCTACCCGGTCGCATGGTGGCTTCTTAGATGAGGAGCAG | 4 |
| wt-166860-166980 | 4242 | GCATAAAAGTCCAAACAGGACACAGAGTACCACCAGGAGTAGTCTTAGTCTGCTGACGTCTGGGTCCTCGGGGCAGGGGTGGCTAGGCCTGGTCTCCGTAGAAGAGCCGGGCAGGCCGCA | 4 |
| wt-166980-167100 | 4243 | GGCAGAGGACTGCTGCTCTAGCAAAGCACGCTCCAGGACGTGTACCATCTCGAGAGTGAGGCACAGCTGTTTTCGTGGACTTTTATACAGTAAGGACAAGGAAAGAAGGCCAGAGGAATG | 4 |
| wt-167100-167220 | 4244 | TGGAAAGATGAGCGAGGACAGGTGTGGAGGTTTTGGGCTAGCTCTTAGTTTCTGGGTGTGAGAGAGGGATTAAAGTGCTTATGCGCAAAGAATGTGTCAACAACAGGTGTTCCTGCCTCT | 4 |
| wt-167220-167340 | 4245 | GCTGGCATGAGTTAGGTGTGGCTTGGGCTGAATCCAAATGTGTATTGGCACAAGATGGAAAGCAAAGTTGCTGGAGTTACTGGGTGGGAGACAGGGATGTATGTGGTCCCCCGCTGGTAT | 4 |
| wt-167340-167460 | 4246 | GCCAGTACCCTGTGGAAGTAAGGGGCCTCATCTGCCTGGTAGTTGTGTTGTGCAGAGGTCTGATGTGTGTAGGAGGGGTGGGTTCAACGCAGGGGCGTTGGTGGCGGAGTCTGGCAACGC | 4 |
| wt-167460-167580 | 4247 | CCGGGTCCTTGCTACCTGTGTGGTGTGTTAAGGGCTGGGTAAAGGTGTCTGCCAATTCTCGCATGTCCTCCTTTCCCCTTGTTTTGAAATAGAATATGAATGTGGCTTTTCAGCCTAGAC | 4 |
| wt-167580-167700 | 4248 | AGACAGTGTGGCTAAGGGAGTGTGTGCCAGTTAAGGTGATTAGCTAAGGCATTCCCAGTAAATGGAGGGAGAGTCAGTCAGGCAAGCCTATGACATGGTAATGCCTAGAAGTAAAGAAAG | 4 |
| wt-167700-167820 | 4249 | GTTAGTCATAGTAGCTTAGCTGAACTGGGCCGTGGGGGTCGTCATCATCTCCACCGGAACCAGAAGAACCCAAAAGCAGCGTAGGAAGGTGTGGATCACCGCCGCCATGGCCGGAATCAT | 4 |
| wt-167820-167940 | 4250 | GACTATGACCGCCGCCTCCGTCTGTCATCAAAGGCGGGCCCTGGTCACCTCCTTTGTTTTCAACCTCTTCCGTCAATTGTGGAGGGCCTCCATCATTTCCAGCAGAGTCGCTAGGGCTAT | 4 |
| wt-167940-168060 | 4251 | GAGGCAGCGGGTCATGTGGGCCATTGTCATCAGTGTTGTCAGGGTCCTGTGGGCCATTGTCATCAGTGTTGTCAGGGTCCTGAGGCAGCGGGTCATGTGGGCCATTGTCATCAGTGTTGT | 4 |
| wt-168060-168180 | 4252 | CAGGGTCCTGTGGGCCATTGTCATCAGTGTTGTCAGGGTCCTGTGGGCCATTGTCAGGACCACCTCCAGGTGCGCCTAGGTTTTGAGAGCAGAGTGGGGGTCCGTCGCCGGCTCCACTCA | 4 |
| wt-168180-168300 | 4253 | CGAGCAGGTGGTGTCTGCCCTCGTTGGAGTTAGAGTCAGATTCATGGCCAGAATCATCGGTAGCTTGTTGAGGGTGCGGGAGGGAGTCATCGTGGTGGTGTTCATCACTGTGTCGTTGTC | 4 |
| wt-168300-168420 | 4254 | CATGGTAATACATCCAGATTAAAATCGCCAGAAACAGGAGGAGCCAAAGGAGATCAACCAATAGAGTCCACCAGTTTTGTTGTAGATAGAGAGCAATAATGAGCAGGATGAGGTCTAGGA | 4 |
| wt-168420-168540 | 4255 | AGAAGGCTAGGAAGAAGGCCAAAAGCTGCCAGATGGTGGCACCAAGTCGCCAGAGCATCTCCAATAAGTAGATCCAGATACCTAAGACTGCGTTGAAAAAAGAGTGTTAGGGTTGGAAAA | 4 |
| wt-168540-168660 | 4256 | GTGGGGGTGTGGTAAATAATTCCTAGGGAATGTTAGATCTTACCAAGTAAGCACCCGAAGATGAACAGCACAATTCCAAGGAACAATGCCTGTCCGTGCAAATTCCAGAGAGCGATGAGC | 4 |
| wt-168660-168780 | 4257 | AGGAGGGTGACTGGGGAAAGAGGAGAAAGTGCGTTAGAGAAGGAAGAGTAAGGGAAAGGGGGTGTGGGGCAAAGGGTGTAATACTTACTCATCAGTAGGAGTATACAAAGGGCTCCAAGT | 4 |
| wt-168780-168900 | 4258 | GGACAGAGAAGGTCTCTTCTGAAGATAAAGATGATCAAAATTATAATTATAAGCATGAGAGCAAAGGAATAGAGGACAAGGAGGGCTCCTCCAGTCCAGTCACTCATAACGATGTACAGC | 4 |
| wt-168900-169020 | 4259 | CAAAACAGTAGCGCCAAGAGGAGGAGAAGGAGAGCAAGGCCTAGGGAAGAGGAGAGGGGGGGTCCTCGAGGGGGCCGTCGCGGGCCCGGTGGGCCCCTCTCAAGGTCGTGTTCCATCCTC | 8 |
| wt-169020-169140 | 4260 | AGGGCAGTGTGTCAGGAGCAAGGCAGTTGAGGAAAGAAGGGGGCAGAGCAGTGTGAGAGGCTTATGTAGGGCGGCTACGTCAGAGTAACGCGTGTTTCTTGGGATGTAGGCCCGGGGGGA | 4 |
| wt-169140-169260 | 4261 | TTTGCGGGGTCTGCCGGAGGCAGTACGGGTACAGATTTCCCGAAAGCGGCGGTGTGTGTGTGCATGTAAGCGTAGAAAGGGGAAGTAGAAAGCGTGTGTTTGTGTTAGAAAAGCGGGTCC | 4 |
| wt-169260-169380 | 4262 | CCGGGGGGCAAGCTGTGGGAATGCGGTGGCCAAGTGCAACAGGAAATGGAAAGGCAGTGCGGCAATCAGAAGGGGGAGTGCGTAGTGTTGTGGGAAGCGGCAGTGTAATCTGCACAAAGA | 4 |
| wt-169380-169500 | 4263 | GGCGCGGGGCGCGCAACGTTGGGAGGTCGTTGGCGGCAGGCGGGAGGCCGTGCTTTAGGGGGGGTTCAGGTGAGGCAAGGCTGTGGGGTAACCGTAGGGGAGGCGGGTGAGGCGGCTAAGA | 8 |
| wt-169500-169620 | 4264 | GGGCTAAGGGTCGGCGGGTGACGAAGCAGCAGACGGCGGATATGGGAATTTCAGAATGAGGTGGCGGATTCAGGCGAAAAGGGTGTGGGCTGTGCGAGTGTCATGAGGCAGGCGCGGAAA | 4 |
| wt-169620-169740 | 4265 | GTCGCTGCGGCTTGCTGGGGCATGGGGGGCCGCGCATTCCTGGAAAAAGTGGAGGGGGCGTGGCCTTCCCCCGCGGCCCCCCAGCCCCCCCGCACAGAGCGGCGCTACGGCGGGCGGGCG | 12 |
| wt-169740-169860 | 4266 | GCGGGGGGTCGGGGTCCGCGGGCTCCGGGGGCTGCGGGCGGTGGATGGCGGCGGACGTTCCGGGGATCGGGGGGGTCGGGGGGCGCCGCGCGGGCGCAGCCATGCGTGACCGTGATGAGG | 12 |
| wt-169860-169980 | 4267 | GGGCAGGGTCGCAGGGGGTGTGTCTGGTGGGGGCGGGAGCGGGGGGCGGCGCGGGAGCCTGCACGCCGTTGGAGGGTAGAATGACAGGGGGCGGGGACAGAGAGGCGGTCGCGCCCCCGG | 12 |
| wt-169980-170100 | 4268 | CCGCGCCAGCCAAGCCCCCAAGGGGGGCGGGGAGCGGGCAATGGAGCGTGACGAAGGGCCCCAGGGCTGACCCCGGCAAACGTGACCCGGGGCTCCGGGGTGACCCAGCCAAGCGTGACC | 8 |
| wt-170100-170220 | 4269 | AAGGGGCCCGTGGGTGACACAGGCAACCCTGACAAAGGCCCCCCAGGAAAGACCCCCGGGGGGCATCGGGGGGTGGGGCATGGGGGGCCGCGCATTCCTGGAAAAAGTGGAGGGGGCGTG | 8 |
| wt-170220-170340 | 4270 | GCCTTCCCCCGCGGCCCCCCAGCCCCCCCGCACAGAGCGGCGCTACGGCGGGCGGGCGGCGGGGGGTCGGGGTCCGCGGGCTCCGGGGGCTGCGGGCGGTGGATGGCGGCGGACGTTCCG | 12 |

| | | | |
|---|---|---|---|
| wt-170340-170460 | 4271 | GGGATCGGGGGGGTCGGGGGGCGCCGCGCGGGCGCAGCCATGCGTGACCGTGATGAGGGGGCAGGGTCGCAGGGGGTGTGTCTGGTGGGGGCGGGAGCGGGGGGCGGCGCGGGAGCCTGC | 12 |
| wt-170460-170580 | 4272 | ACGCCGTTGGAGGGTAGAATGACAGGGGGCGGGGACAGAGAGGCGGTCGCGCCCCCGGCCGCGCCAGCCAAGCCCCCAAGGGGGGCGGGGAGCGGGCAATGGAGCGTGACGAAGGGCCCC | 8 |
| wt-170580-170700 | 4273 | AGGGCTGACCCCGGCAAACGTGACCCGGGGCTCCGGGGTGACCCAGCCAAGCGTGACCAAGGGGCCCGTGGGTGACACAGGCAACCCTGACAAAGGCCCCCCAGGAAAGACCCCCGTGGG | 8 |
| wt-170700-170820 | 4274 | GCATGGGGGGCCGCGCATTCCTGGAAAAAGTGGAGGGGGCGTGGCCTTCCCCCGCGGCCCCCCAGCCCCCCCGCACAGAGCGGCGCTACGGCGGGCGGGCGGCGGGGGGTCGGGGTCCGC | 12 |
| wt-170820-170940 | 4275 | GGGCTCCGGGGGCTGCGGGCGGTGGATGGCGGCGGACGTTCCGGGGATCGGGGGGGTCGGGGGGCGCCGCGCGGGCGCAGCCATGCGTGACCGTGATGAGGGGGCAGGGTCGCAGGGGGT | 12 |
| wt-170940-171060 | 4276 | GTGTCTGGTGGGGGCGGGAGCGGGGGGCGGCGCGGGAGCCTGCACGCCGTTGGAGGGTAGAATGACAGGGGGCGGGGACAGAGAGGCGGTCGCGCCCCCGGCCGCGCCAGCCAAGCCCCC | 12 |
| wt-171060-171180 | 4277 | AAGGGGGGCGGGGAGCGGGCAATGGAGCGTGACGAAGGGCCCCAGGGCTGACCCCGGCAAACGTGACCCGGGGCTCCGGGGTGACCCAGCCAAGCGTGACCAAGGGGCCCGTGGGTGACA | 8 |
| wt-171180-171300 | 4278 | CAGGCAACCCTGACAAAGGCCCCCCAGGAAAGACCCCCGGGGGGCATCGGGGGGTGGGGCATGGGGGGCCGCGCATTCCTGGAAAAAGTGGAGGGGGCGTGGCCTTCCCCCGCGGCCCCC | 8 |
| wt-171300-171420 | 4279 | CAGCCCCCCCGCACAGAGCGGCGCTACGGCGGGCGGGCGGCGGGGGGTCGGGGTCCGCGGGCTCCGGGGGCTGCGGGCGGTGGATGGCGGCGGACGTTCCGGGGATCGGGGGGGTCGGGG | 12 |
| wt-171420-171540 | 4280 | GGCGCCGCGCGGGCGCAGCCATGCGTGACCGTGATGAGGGGGCAGGGTCGCAGGGGGTGTGTCTGGTGGGGGCGGGAGCGGGGGGCGGCGCGGGAGCCTGCACGCCGTTGGAGGGTAGAA | 12 |
| wt-171540-171660 | 4281 | TGACAGGGGGCGGGGACAGAGAGGCGGTCGCGCCCCCGGCCGCGCCAGCCAAGCCCCCAAGGGGGGCGGGGAGCGGGCAATGGAGCGTGACGAAGGGCCCCAGGGCTGACCCCGGCAAAC | 12 |
| wt-171660-171780 | 4282 | GTGACCCGGGGCTCCGGGGTGACCCAGCCAAGCGTGACCAAGGGGCCCGTGGGTGACACAGGCAACCCTGACAAAGGCCCCCCAGGAAAGACCCCCGGGGGGCATCGGGGGGGGTGTTGG | 8 |
| wt-171703-171823 | 4283 | GGCCCGTGGGTGACACAGGCAACCCTGACAAAGGCCCCCCAGGAAAGACCCCCGGGGGGCATCGGGGGGGGTGTTGGCGGGGGCATGGGGGGGTCGGATTTCGCCCTTATTGCCCTGTTT | 8 |
| wt-90-210 | 4284 | GCCCTACCTCTTTTGGCTGGCGGCTATTGCCGCCTCGTGTTTCACGGCCTCAGTTAGTACCGTTGTGACCGCCACCGGCTTGGCCCTCTCACTTCTACTCTTGGCAGCAGTGGCCAGCTC | 4 |
| wt-210-330 | 4285 | ATATGCCGCTGCACAAAGGAAACTGCTGACACCGGTGACAGTGCTTACTGCGGTTGTCACTTGTGAGTACACACGCACCATTTACAATGCATGATGTTCGTGAGATTGATCTGTCTCTAA | 4 |
| wt-330-450 | 4286 | CAGTTCACTTCCTCTGCTTTTCTCCTCAGTCTTTGCAATTTGCCTAACATGGAGGATTGAGGACCCACCTTTTAATTCTCTTCTGTTTGCATTGCTGGCCGCAGCTGGCGGACTACAAGG | 4 |
| wt-450-570 | 4287 | CATTTACGGTTAGTGTGCCTCTGTTATGAAATGCAGGTTTGACTTCATATGTATGCCTTGGCATGACGTCAACTTTACTTTTATTTCAGTTCTGGTGATGCTTGTGCTCCTGATACTAGC | 4 |
| wt-570-690 | 4288 | GTACAGAAGGAGATGGCGCCGTTTGACTGTTTGTGGCGGCATCATGTTTTTGGCATGTGTACTTGTCCTCATCGTCGACGCTGTTTTGCAGCTGAGTCCCCTCCTTGGAGCTGTAACTGT | 4 |
| wt-690-810 | 4289 | GGTTTCCATGACGCTGCTGCTACTGGCTTTCGTCCTCTGGCTCTCTTCGCCAGGGGGCCTAGGTACTCTTGGTGCAGCCCTTTTAACATTGGCAGCAGGTAAGCCACACGTGTGACATTG | 4 |
| wt-810-930 | 4290 | CTTGCCTTTTTGCCACATGTTTTCTGGACACAGGACTAACCATGCCATCTCTGATTATAGCTCTGGCACTGCTAGCGTCACTGATTTTGGGCACACTTAACTTGACTACAATGTTCCTTC | 4 |
| wt-930-1050 | 4291 | TCATGCTCCTATGGACACTTGGTAAGTTTTCCCTTCCTTTAACTCATTACTTGTTCTTTTGTAATCGCAGCTCTAACTTGGCATCTCTTTTACAGTGGTTCTCCTGATTTGCTCTTCGTG | 4 |
| wt-1050-1170 | 4292 | CTCTTCATGTCCACTGAGCAAGATCCTTCTGGCACGACTGTTCCTATATGCTCTCGCACTCTTGTTGCTAGCCTCCGCGCTAATCGCTGGTGGCAGTATTTTGCAAACAAACTTCAAGAG | 4 |
| wt-1170-1290 | 4293 | TTTAAGCAGCACTGAATTTATACCCAGTGAGTATCTATTTGTTACTCCTGTTTAGTTGAAGAAAACAAGCTATTGGATTGTAACACACATTTTACGCTTTGTTCCTTAGATTTGTTCTGC | 4 |
| wt-1290-1410 | 4294 | ATGTTATTACTGATTGTCGCTGGCATACTCTTCATTCTTGCTATCCTGACCGAATGGGGCAGTGGAAATAGAACATACGGTCCAGTTTTTATGTGCCTCGGTGGCCTGCTCACCATGGTA | 4 |
| wt-1410-1530 | 4295 | GCCGGCGCTGTGTGGCTGACGGTGATGTCTAACACGCTTTTGTCTGCCTGGATTCTTACAGCAGGATTCCTGATTTTCCTCATTGGTAAGTGTGACACCAACAGGTGTTGCCTTGTTATG | 4 |
| wt-1530-1650 | 4296 | TCACCGTTCTGACACATGACTTACATGGGTTTGGCTTTTGTAGGCTTTGCCCTCTTTGGGGTCATTAGATGCTGCCGCTACTGCTGCTACTACTGCCTTACACTGGAAAGTGAGGAGCGC | 4 |
| wt-1650-1770 | 4297 | CCACCGACCCCATATCGCAACACTGTATAAAGGTAAGTATTATTAAATTTTAGAGACACTATCACGTGTAACTTGACGTGCAAGGATGGAAGAGAGGGGCAGGGAAACGCAAATGCCGGT | 4 |
| wt-1770-1890 | 4298 | TGCCCGGTATGGGGGCCCGTTTATTATGGTAAGGCTCTTCGGGCAAGATGGAGAGGCAAACATACAGGAGGAAAGGCTATATGAGCTACTCTCTGACCCACGCTCCGCGCTCGGCCTAGA | 4 |
| wt-1890-2010 | 4299 | CCCGGGGCCCCTGATTGCTGAGAACCTGCTGCTAGTGGCGCTGCGTGGCACCAACAACGATCCCAGGCCTCAGCGTCAGGAGAGGGCCAGAGAACTGGCCCTCGTTGGCATTCTACTAGG | 4 |
| wt-2010-2130 | 4300 | AAACGGCGAGCAGGGTGAACACTTGGGCACGGAGAGTGCCCTGGAGGCCTCAGGCAACAACTATGTGTATGCCTACGGACCAGACTGGATGGCAAGGCCTTCCACATGGTCCGCGGAAAT | 4 |
| wt-2130-2250 | 4301 | CCAGCAATTCCTGCGACTCCTGGGCGCCACGTACGTGCTTCGCGTGGAGATGGGCAGGCAGTTTGGCTTCGAGGTGCATAGAAGCCGGCCCTCCTTCCGTCAGTTCCAGGCCATCAATCA | 4 |
| wt-2250-2370 | 4302 | CCTTGTCCTGTTTGACAACGCCCTTCGCAAGTACGATTCCGGCCAGGTGGCGGCGGGCTTCCAGAGGGCCCTTCTGGTGGCCGGGCCAGAGACCGCTGACACGAGGCCGGACCTCCGCAA | 8 |
| wt-2370-2490 | 4303 | GCTGAATGAGTGGGTGTTTGGTGGCAGGGCTGCTGGTGGCAGACAGCTGGCCGACGAGCTAAAGATCGTGTCCGCGCTGCGAGACACTTACTCGGGCCACTTGGTCCTTCAGCCCACGGA | 4 |
| wt-2490-2610 | 4304 | GACCCTTGACACATGGAAGGTGTTGAGCAGGGACACACGAACCGCTCATAGTTTGGAGCACGGATTCATTCATGCCGCGGGGACCATCCAGGCCAACTGCCCACAGCTGTTTATGAGACG | 4 |
| wt-2610-2730 | 4305 | CCAGCACCCCGGCCTCTTTCCCTTCGTTAATGCAATAGCATCATCGCTGGGCTGGTACTACCAGACCGCCACCGGCCCCGGAGCAGATGCCAGGGCGGCGGCCCGGCGCCAACAGGCCTT | 8 |
| wt-2730-2850 | 4306 | TCAGACCAGGGCGGCGGCTGAATGCCATGCCAAAAGCGGGGTGCCGGTCGTGGCCGGCTTCTACAGGACCATCAACGCCACGCTCAAGGGAGGAGAGGGCCTACAGCCCACTATGTTTAA | 4 |
| wt-2850-2970 | 4307 | CGGGGAGCTGGGGGCCATCAAGCACCAGGCACTTGACACTGTGAGGTATGACTACGGCCACTATCTCATAATGTTGGGGCCATTCCAGCCATGGAGCGGACTGACGGCCCCTCCGTGCCC | 4 |
| wt-2970-3090 | 4308 | CTACGCCGAAAGTTCATGGGCACAGGCGGCCGTGCAGACGGCCCTCGAGCTGTTCTCGGCCCTGTACCCGGCCCCGTGCATCTCGGGCTACGCGCGCCCCCCGGGCCCCAGTGCTGTGAT | 8 |
| wt-3090-3210 | 4309 | CGAGCATCTGGGGTCCCTAGTTCCAAAGGGGGGTCTGCTGTTGTTTCTGTCTCACCTACCGGATGATGTTAAGGACGGGCTCGGAGAAATGGGGCCGGCCAGGGCCACGGGACCTGGAAT | 4 |
| wt-3210-3330 | 4310 | GCAGCAGTTTGTCAGCAGCTACTTCCTCAACCCCGCCTGTTCCAACGTCTTCATTACAGTGAGGCAGCGAGGGGAGAAGATCAACGGCCGTACCGTCCTCCAAGCGCTCGGACGCGCATG | 4 |
| wt-3330-3450 | 4311 | CGATATGGCAGGCTGCCAGCACTATGTGCTGGGCTCCACGGTTCCCCTCGGTGGACTCAACTTTGTCAACGACCTGGCGTCCCCGGTTTCCACCGCCGAGATGATGGATGATTTCTCTCC | 4 |
| wt-3450-3570 | 4312 | CTTCTTCACCGTGGAGTTTCCCCCGATTCAAGAGGAGGGCGCAAGTTCTCCGGTACCCTTAGATGTGGACGAGAGCATGGACATCTCTCCGTCTTACGAGTTGCCCTGGCTCTCGCTGGA | 4 |
| wt-3690-3810 | 4314 | CCTGCCGCTGGCGGACTACGCCTTCGTTGCCCACAGTCAGGTCTGGACCAGGCCCGGTGGGGCTCCTCCCTTGCCCTATCGTACCTGGGATCGAATGACAGAGAAGCTGCTTGTCTCCGC | 4 |
| wt-3810-3930 | 4315 | AAAACCCGGCGGAGAGAACGTTAAGGTTTCAGGTACCGTGATTACATTGGGAGAACAGGGGTACAAAGTGTCGTTGGATCTGAGGGAGGGAACCAGGCTGGCAATGGCTGAGGCGCTGCT | 4 |
| wt-3930-4050 | 4316 | GAACGCAGCATGTGCCCCAATCTTGGATCCGGAAGACGTCTTGCTCACCCTGCATCTACACCTGGATCCGCGCCGGGCAGACAACTCGGCCGTGATGGAGGCTATGACGGCGGCGAGTGA | 4 |
| wt-4050-4170 | 4317 | CTACGCGCGTGGCCTGGGCGTGAAGCTGACCTTTGGCTCGGCCTCCTGCCCCGAGACCGGCTCGTCCGCCTCCAACTTCATGACCGTGGTGGCCTCTGTCTCCGCCCCAGGGGAATTCTC | 8 |
| wt-4170-4290 | 4318 | GGGTCCTCTGATCACGCCAGTGCTTCAGAAGACGGGCAGTCTCCTGATTGCGGTGCGTTGCGGGGATGGCAAGATCCAGGGAGGGTCGCTGTTTGAGCAGCTCTTTAGCGACGTGGCCAC | 4 |
| wt-4290-4410 | 4319 | GACCCCACGGGCACCCGAGGCGTTGTCTCTGAAGAATCTCTTCCGGGCAGTCCAGCAGCTGGTCAAGAGCGGCATCGTGCTGTCAGGGCATGACATCAGCGACGGGGGCCTGGTGACCTG | 4 |
| wt-4410-4530 | 4320 | CCTGGTGGAGATGGCCCTGGCCGGGCAGCGGGGAGTGACCATCACTATGCCGGTGGCCTCCGACTACCTCCCGGAGATGTTTGCAGAGCACCCCGGCCTGGTGTTTGAGGTGGAGGAGCG | 8 |
| wt-4530-4650 | 4321 | CAGCGTGGGTGAGGTGCTGCAGACCCTGCGCTCCATGAACATGTACCCGGCAGTCCTCGGTCGAGTGGGCGAGCAAGGTCCAGATCAAATGTTTGAGGTGCAGCACGGCCCAGAGACGGT | 4 |
| wt-4650-4770 | 4322 | GTTGCGCCAGTCGCTGCGCCTGCTGCTGGGAACCTGGTCATCCTTTGCCAGCGAGCAGTACGAGTGCCTGCGACCAGATCGGATTAACCGGTCCATGCACGTGTCCGACTACGGCTATAA | 4 |

| | | | |
|---|---|---|---|
| wt-4770-4890 | 4323 | CGAAGCACTGGCAGTCTCCCCGTTGACAGGAAAGAATCTCAGCCCACGCCGGTTGGTGACAGAGCCTGACCCACGATGTCAGGTGGCCGTGCTATGCGCCCCGGGCACCAGGGGCCATGA | 4 |
| wt-4890-5010 | 4324 | AAGCCTCCTGGCGGCCTTCACGAATGCCGGATGCCTGTGCCGACGGGTGTTCTTTCGCGAGGTTAGGGACAACACGTTCCTCGACAAGTACGTGGGTCTGGCCATCGGAGGAGTTCATGG | 4 |
| wt-5010-5130 | 4325 | GGCCAGGGACTCTGCCCTGGCAGGCCGTGCCACCGTGGCGCTGATTAATCGTTTCCCCGCCCTGCGTGACGCTATTCTAAAGTTCCTCAACAGGCCAGATACGTTCTCGGTGGCCTTGGG | 4 |
| wt-5130-5250 | 4326 | GGAGCTGGGGGTGCAAGTTTTGGCTGGCCTGGGGGCCGTGGGGTCAACAGATAATCCACCCGCCCCTGGCGTGGAAGTTAATGTCCAGAGATCACCTCTGATTCTGGCCCCCAACGCCTC | 4 |
| wt-5250-5370 | 4327 | TGGCATGTTTGAGTCCCGCTGGCTGAACATTAGCATCCCGGCGACCACCAGCTCTGTCATGCTGCGTGGCCTCCGGGGCTGCGTCCTGCCTTGTTGGGTGCAAGGCTCGTGCCTGGGCCT | 4 |
| wt-5370-5490 | 4328 | GCAATTTACTAACCTCGGGATGCCATATGTTTTGCAGAATGCCCACCAGATCGCCTGCCACTTCCACAGCAATGGCACGGATGCCTGGCGCTTTGCTATGAATTATCCAAGAAACCCCAC | 4 |
| wt-5490-5610 | 4329 | GGAGCAGGGCAACATTGCAGGGCTCTGTTCACGCGATGGTCGTCATCTGGCTCTCCTGTGTGACCCCTCACTTTGTACAGACTTTTGGCAATGGGAGCACATTCCCCCCGCCTTTGGGCA | 4 |
| wt-5610-5730 | 4330 | CCCCACGGGGTGCTCCCCCTGGACACTTATGTTTCAAGCAGCTCACCTATGGTCACTCAGGCACGGTCGCCCCTCCGAGTGACCAGTCACCTTCCAGACTATGCATACACTGAATTTAGC | 4 |
| wt-5730-5850 | 4331 | CTGATATTGTCCCCCTAGCCCCGGGCCCAGCCCTCCTCAGAAAACTCTGCATGGAGAAGCTGGACGTGAACCTCCCCCCCAGACCTGTGTGCTGTATTTACAAACACTACAATAAACCCA | 4 |
| wt-5850-5970 | 4332 | ATGTGCAAATGTGGTTTGTATGGCTACTTTGTGTTCCTAAAAAATGCAACAATAGAAGTGGAAACCCTCAGTCACGGGACATTAACCTCAACCACAAAATGGGGGTTGGAGAAAGTAACC | 4 |
| wt-5970-6090 | 4333 | ACATATACTGGAGATGATTCATGGGCTGGGGGTTCCCGGACAATACACCCATCTGGAGTTCAACCTAATTACATGGTAGATAAATTAAGAGTCCCTCCTCACCACTCGAAACTATGGCAG | 4 |
| wt-6090-6210 | 4334 | ACATTCTATAAGATAACGAGGAGAGATGAGGTGAGGGCAGAGGACATTGGGCAGGTGTGGGCCACGGGGCAGCTGGCCATATCCCCCGCACTACAGAAGTGTAAGCAAAGTGAAGGGCTC | 4 |
| wt-6210-6330 | 4335 | GGAAGGCAGGCGGGGCCTAGCAATGTCACAGCTAAATGCCCACCAGGGCACACACTCAAGCGGGGTCTCGGAGCTCCTAGGTCAGACCACGAAAGGTCAGCCTGCAAGGTGGATGGCGTG | 4 |
| wt-6330-6450 | 4336 | TTTTCTGAGGTTATCCCCGCTACGTGCAGTGCTGGGTGATAGAGACCCTAGAATGTGTCGAAATGACCAAGCGTCCCCGCAGCGGGGCTCCCAACACGGGTTCCCAGAGAGGGTAAAAGA | 4 |
| wt-6450-6570 | 4337 | GGGGGCCATAAAGCCCAGGGTGTAAAACACCGACCGCGCCACCAGATGGCACACGTGGGGGAAATGAGGGTTAGCATAGGCAACCCCCGCCTACACACCAACTATAGCAAACCCCGCCCC | 4 |
| wt-6570-6690 | 4338 | GTCACGGTGACGTAGTCTGTCTTGAGGAGATGTAGACTTGTAGACACTGCAAAACCTCAGGACCTACGCTGCCCTAGAGGTTTTGCTAGGGAGGAGACGTGTGTGGCTGTAGCCACCCGT | 4 |
| wt-6690-6810 | 4339 | CCCGGGTACAAGTCCCGGGTGGTGAGGACGGTGTCTGTGGTTGTCTTCCCAGACTCTGCTTTCTGCCGTCTTCGGTCAAGTACCAGCTGGTGGTCCGCATGTTTTGATCCAAACTTTTGT | 4 |
| wt-6810-6930 | 4340 | TTTAGGATTTATGCATCCATTATCCCGCAGTTCCACCTAAACGGGGCTTAACGTTGCATCCCAGAAGATGCACGCTTAACCCCGCCTACAACCGTGACGTAGCTGTTTACCAGCATGTAT | 4 |
| wt-6930-7050 | 4341 | AGAGTTACGGTTCGCTACATCAAACAGGACAGCCGTTGCCCTAGTGGTTTCGGACACACCGCCAACGCTCAGTGCGGTGCTACCGACCCGAGGTCAAGTCCCGGGGGAGGAGAAGAGAGG | 4 |
| wt-7050-7170 | 4342 | CTTCCCGCCTAGAGCATTTGCAAGTCAGGATTCTCTAATCCCTCTGGGAGAAGGGTATTCGGCTTGTCCGCTATTTTTTTGTGGCTAGTTTTGCACCCACAACATGTAAGGGCCCGCTAC | 4 |
| wt-7170-7290 | 4343 | CCCTACAACACAAAACAAACTATCTCCCCTAACCATCCTTTTGCCAATCAATTCTGTGACAGGGTTTCCTGGACACCCAGTCTTAGTTCAGGTAGACACCCAGTTATGCAGTGCCACCAA | 4 |
| wt-7290-7410 | 4344 | TTCCAACCATTTTTAAACCTCCTGGAATTCTATCATTAAACGGCATGCAGGAAAAGGACAAGCAGCGAAAATTCACGCCCCCTTGGGAGGTGGCGGCATATGCAAAGGATAGCACTCCCA | 4 |
| wt-7410-7530 | 4345 | CTCTACTACTGGGTATCATATGCTGACTGTATATGCATGAGGATAGCATATGCTACCCGGATACAGATTAGGATAGCATATACTACCCAGATATAGATTAGGATAGCATATGCTACCCAG | 4 |
| wt-7530-7650 | 4346 | ATATAGATTAGGATAGCCTATGCTACCCAGATATAAATTAGGATAGCATATACTACCCAGATATAGATTAGGATAGCATATGCTACCCAGATATAGATTAGGATAGCCTATGCTACCCAG | 4 |
| wt-7650-7770 | 4347 | ATATAGATTAGGATAGCATATGCTACCCAGATATAGATTAGGATAGCATATGCTATCCAGATATTTGGGTAGTATATGCTACCCAGATATAAATTAGGATAGCATATACTACCCTAATCT | 4 |
| wt-7770-7890 | 4348 | CTATTAGGATAGCATATGCTACCCGGATACAGATTAGGATAGCATATACTACCCAGATATAGATTAGGATAGCATATGCTACCCAGATATAGATTAGGATAGCCTATGCTACCCAGATAT | 4 |
| wt-7890-8010 | 4349 | AAATTAGGATAGCATATACTACCCAGATATAGATTAGGATAGCATATGCTACCCAGATATAGATTAGGATAGCCTATGCTACCCAGATATAGATTAGGATAGCATATGCTATCCAGATAT | 4 |
| wt-8010-8130 | 4350 | TTGGGTAGTATATGCTACCCATGGCAACATTAGCCCACCGTGCTCTCAGCGACCTCGTGAATATGAGGACCAACAACCCTGTGCTTGGCGCTCAGGCGCAAGTGTGTGTAATTTGTCCTC | 4 |
| wt-8130-8250 | 4351 | CAGATCGCAGCAATCGCGCCCCTATCTTGGCCCGCCCACCTACTTATGCAGGTATTCCCCGGGGTGCCATTAGTGGTTTTGTGGGCAAGTGGTTTGACCGCAGTGGTTAGCGGGGTTACA | 4 |
| wt-8250-8370 | 4352 | ATCAGCCAAGTTATTACACCCTTATTTTACAGTCCAAAACCGCAGGGCGGCGTGTGGGGGCTGACGCGTGCCCCCACTCCACAATTTCAAAAAAAAGAGTGGCCACTTGTCTTTGTTTAT | 4 |
| wt-8370-8490 | 4353 | GGGCCCCATTGGCGTGGAGCCCCGTTTAATTTTCGGGGGTGTTAGAGACAACCAGTGGAGTCCGCTGCTGTCGGCGTCCACTCTCTTTCCCCTTGTTACAAATAGAGTGTAACAACATGG | 4 |
| wt-8490-8610 | 4354 | TTCACCTGTCTTGGTCCCTGCCTGGGACACATCTTAATAACCCCAGTATCATATTGCACTAGGATTATGTGTTGCCCATAGCCATAAATTCGTGTGAGATGGACATCCAGTCTTTACGGC | 4 |
| wt-8610-8730 | 4355 | TTGTCCCCACCCCATGGATTTCTATTGTTAAAGATATTCAGAATGTTTCATTCCTACACTAGTATTTATTGCCCAAGGGGTTTGTGAGGGTTATATTGGTGTCATAGCACAATGCCACCA | 4 |
| wt-8730-8850 | 4356 | CTGAACCCCCCGTCCAAATTTTATTCTGGGGGCGTCACCTGAAACCTTGTTTTCGAGCACCTCACATACACCTTACTGTTCACAACTCAGCAGTTATTCTATTAGCTAAACGAAGGAGAA | 4 |
| wt-8850-8970 | 4357 | TGAAGAAGCAGGCGAAGATTCAGGAGAGTTCACTGCCCGCTCCTTGATCTTCAGCCACTGCCCTTGTGACTAAAATGGTTCACTACCCTCGTGGAATCCTGACCCCATGTAAATAAAACC | 4 |
| wt-8970-9090 | 4358 | GTGACAGCTCATGGGGTGGGAGATATCGCTGTTCCTTAGGACCCTTTTACTAACCCTAATTCGATAGCATATGCTTCCCGTTGGGTAACATATGCTATTGAATTAGGGTTAGTCTGGATA | 4 |
| wt-9090-9210 | 4359 | GTATATACTACTACCCGGGAAGCATATGCTACCCGTTTAGGGTTAACAAGGGGGCCTTATAAACACTATTGCTAATGCCCTCTTGAGGGTCCGCTTATCGGTAGCTACACAGGCCCCTCT | 4 |
| wt-9210-9330 | 4360 | GATTGACGTTGGTGTAGCCTCCCGTAGTCTTCCTGGGCCCCTGGGAGGTACATGTCCCCCAGCATTGGTGTAAGAGCTTCAGCCAAGAGTTACACATAAAGGCAATGTTGTGTTGCAGTC | 4 |
| wt-9330-9450 | 4361 | CACAGACTGCAAAGTCTGCTCCAGGATGAAAGCCACTCAGTGTTGGCAAATGTGCACATCCATTTATAAGGATGTCAACTACAGTCAGAGAACCCCTTTGTGTTTGGTCCCCCCCCGTGT | 4 |
| wt-9450-9570 | 4362 | CACATGTGGAACAGGGCCCAGTTGGCAAGTTGTACCAACCAACTGAAGGGATTACATGCACTGCCCCGCGGGAAATACGTCCTACCCAGGAACCCGAAACAGTGTTTCCCAGAAGCTGTA | 4 |
| wt-9570-9690 | 4363 | AAAATAGAACGCCCTGGAACTGCCCCACTGTGCAATGCAGCTTTTAGCCATGCCATGCTCTATAAATCACTTCCCTATCTCAGGTAGGCCTGCACACCTTAGGTATGGAGCGAAGGTTAG | 4 |
| wt-9690-9810 | 4364 | TGGTCACTCTGCAGTGCCTGGTGCTGCTTTACCTGGCACCTGAGTGTGGAGGTACAGACCAATGTGACAATTTTCCCCAAATGTTGAGGGACCTAAGAGATGCCTTCAGTCGTGTTAAAA | 4 |
| wt-9810-9930 | 4365 | CCTTTTTCCAGACAAAGGACGAGGTAGATAACCTTTTGCTCAAGGAGTCTCTGCTAGAGGACTTTAAGGGCTACCTTGGATGCCAGGCCCTGTCAGAAATGATCCAATTCTACCTGGAGG | 4 |
| wt-9930-10050 | 4366 | AAGTCATGCCACAGGCTGAAAACCAGGACCCTGAAGCCAAAGACCATGTCAATTCTTTGGGTGAAAATCTAAAGACCCTACGGCTCCGCCTGCGCAGGTGCCACAGGTTCCTGCCGTGTG | 4 |
| wt-10170-10290 | 4367 | CAATTAAAGCCAGGTGATAATTCCATACCCTGGAAGCAGGAGATGGGTGCATTTCACCCCAACCCCCCCTTTCGACTGTCATTTACAATAAAATGAAACCTTTTATTCTTGATTGCCTCT | 4 |
| wt-10290-10410 | 4368 | TGTGTTCTTGCCGCCCAGGTACCTTCCTGTGTTCTCCCCACGGGAAAAAGAATAGCTTCTGCAGAAGGCCATTGACGCAAGTTTTGCCCGTGGGGATTACCCGACCCAGCCACTTACAGC | 4 |
| wt-10410-10530 | 4369 | ACATTTTGTTCTAGGTCCATCTTAGGAGCCCGGGCCAGCATTCTATCAGCTTAACGGGAAGAGAAGTGGGGAGGGCACTCGCCCACTAACCTTAACACCTGCAGCCTACAAAAGTACACT | 4 |
| wt-10530-10650 | 4370 | AGCTGTTTGCTCTATTCGCCACTAGAGACCGCCAAGATGCGAAACTACAGGCCCGGGCCCAGGCCTTGCAGGGCAGACGGTTAGGCTGACAAGGGGACAAGTGTGGCAGGTGGGCGGGAA | 4 |
| wt-10650-10770 | 4371 | GGGGCACAAGAATGCCGGCGAAACTGGACCACGGTCCACCCCGCCCTCAAGCGTCCGGGAGCCGGGCGGCTCGGCTAAGGAGGGCGGCCTTGCGAACAATTATTAGTAGCTACCAACAAG | 4 |
| wt-10770-10890 | 4372 | GGCCCCCAGATGCCCCCCACCAGTCACCCGGCCGTGTCCACTCACATATTCCACTCTTATTTTTAAATTAATGTGTCCCAATTAGAAACCCAAGCGCAGAAATTAGTTGAGAGGCTAGTG | 4 |
| wt-10890-11010 | 4373 | TTTTAAACATGCACCCTAGGCCAGCCAGAGATAATGTCACAAGATTATCAAGTTGGTGTAAACACGCCGTGGGAAAAAATTTATGGTTCAGTGCGTCGAGTGCTATCTTTGGAACAGTAG | 4 |

| | | | |
|---|---|---|---|
| wt-11010-11130 | 4374 | AAAATTGAACCTTGTTGGCGGGAGAAGGAATAACGCCTTATCTGGGAGGAGCGACGGATTATAGCCAATAAGAGAGCTCAAGACGCAGGGCTCGCAAAGTATAGTGGCCCCGTGGACCT | 4 |
| wt-11130-11250 | 4375 | TAGAGGTGGAGCAACGTCTAAAGTGGTAATAACACCAGGCGGGGCTGGGCAAAGGGGTCCTACGGGCGGGATTAATTACGCCTTGCTTACGCAAGCTCAGTTAATTCGCCCACGACTTGA | 4 |
| wt-11250-11370 | 4376 | AAAATGTAGCCCTTAACCAATTGGCGGCCCCTAAGGGGGGGACTAAGGTCCCACTACAAAAACTCTGTGTTCTGCTGCAAATTTTAGATCAGATGGCATAGAGACAAGGACACCGAAGAC | 4 |
| wt-11370-11490 | 4377 | CCCCAGAGCCCTCATCGCAGGGTTCTTACCATGCGGCCATGTAGGCCCACTTAACACTACAAGACCTACGCCTCTCCATTCATCATGTAACCCACAAATCATCTAAACCGTAAGTCTAAG | 4 |
| wt-11490-11610 | 4378 | GGCCTCCTGAGGTTTTCTCAGGAGGCCCTAATGTATAATTAATCATGCATTTGATTTTAAAAAAGTAGGTTACACTCATTTTAGGCCAGACTTTATTTGCAGATTAATAATTTATGTGAT | 4 |
| wt-11610-11730 | 4379 | TCTCCTTCCCTCTAGGACTGAAGAAACAGCCTCCTGCACGTGAGCATGTATCTGAAATAATTATTATGTCATAAGTGTAATGATTAGAAAGTCATAAACCCACTTCCCTTTACATGAATC | 4 |
| wt-11730-11850 | 4380 | TGGGCACTGAATTTTGGGGTACTTCTAAAGACTAACGTGTTCGATTTCGGGGTCACTTCCCCTTTTATAAGTGTGTGAACAGTGATTTCAGTAAAACCTAAGAGATATTTGGTGTCACTT | 4 |
| wt-11850-11970 | 4381 | CCGCATTTTAAGTTTCAGAAAATTTTAAAATTAAAATTGAAATTTCTCTCAAAATAATTCCAATGAAAACTTCAAAGAATCTTATGTATGTAATTCTTTTGCCCCAAACTGGGCTTCAGA | 4 |
| wt-11970-12090 | 4382 | TGCCTTCTATTGCACTCTCACAAAAACATTCTGGACACATGTGCCAGACGCCTGGGCCTCTAAGGCCCTCGGGTCCCCCTGGACCCCGGCCTCAGCAACCCTGCTGCTCCCCTCCTGCCA | 4 |
| wt-12090-12210 | 4383 | CCCCAGCCTCCCCCCCTCCCCGTCCCCCTTCGCTCCTGTTCCTCCCCCGGTCCCCAGTAGGGCCGCCTGCCCCCCTGCACCCAGTACCTGCCCCTCTTGGCCACGCACCCCGGGCCAGGC | 12 |
| wt-12210-12330 | 4384 | CACCTTAGACCCGGCCAAGCCCCATCCCTGAAGACCCAGCGGCCATTCTCTCTGGTAACGAGCAGAGAAGAAGTAGAGGCCCGCGGCCATTGGGCCCAGATTGAGAGACCAGTCCAGGGG | 4 |
| wt-12330-12450 | 4385 | CCCGAGGTTGGAGCCAGCGGGCACCCGAGGTCCCAGCACCCGGTCCCTCCGGGGGGCAGAGACAGGCAGGGCCCCCCGGCAGCTGGCCCCGAGGAGGCGCCCGGAGTGGGGCCGGTCGGC | 12 |
| wt-12450-12570 | 4386 | TGGGCTGGCCGAGCCCGGGTCTGGGAGGTCTGGGGTGGCGAGCCTGCTGTCTCAGGAGGGGCCTGGCTCCGCCGGGTGGCCCTGGGGTAAGTCTGGGAGGCAGAGGGTCGGCCTAGGCCC | 8 |
| wt-12570-12690 | 4387 | GGGGAAGTGGAGGGGGATCGCCCGGGTCTCTGTTGGCAGAGTCCGGGCGATCCTCTGAGACCCTCCGGGCCCGGACGGTCGCCCTCAGCCCCCCAGACAGACCCCAGGGTCTCCAGGCAG | 8 |
| wt-12690-12810 | 4388 | GGTCCGGCATCTTCAGGGGCAGCAGGCTCACCACCACAGGCCCCCCAGACCCGGGTCTCGGCCAGCCGAGCCGACCGGCCCCGCGCCTGGCGCCTCCTCGGGGCCAGCCGCCGGGGTTGG | 12 |
| wt-12810-12930 | 4389 | TTCTGCCCCTCTCTCTGTCCTTCAGAGGAACCAGGGACCTCGGGCACCCCAGAGCCCCTCGGGCCCGCCTCCAGGCGCCCTCCTGGTCTCCGCTCCCCTCTGAGCCCCGTTAAACCCAAA | 8 |
| wt-12930-13050 | 4390 | GAATGTCTGAGGGGAGCCACCCTCGGGGCCCAGGCCCCAGAGTCCAGAGGTCAGGGGCACCTCAGGGTGCCTCCCCGGGTCCCAGGCCAGCCGGAGGGACCCCGGCAGCCCGGGCGGCCC | 12 |
| wt-13050-13170 | 4391 | CAGAGGCCGGTTCCTCGCCCCTTCCCCGGGCTTCAGAGCCCAGGATGTCCCCCAGAAGGGACCCTAGGCGTCCCCTCTCCTCCCCTCCAGGCCCGAGCCTCTCCCTCGCGGAGAGGGGCC | 8 |
| wt-13170-13290 | 4392 | TCTTTGGGCCCTCAAGTCCAGCCCCACCGAGACCCGAGTGGCCCGGATCCCCCCACCGGCCCTTCTCTCTGTCCCCCTGCTCCTCTCCAACCTTCGCTCCACCCTAGACCCCAGCTTCTG | 8 |
| wt-13290-13410 | 4393 | GCCTCCCCGGGTCCACCAGGCCAGCCGGAGGGACCCCGGCAGCCCGGGCGAGTCGCCTTCCCTCTCCCCTGGCCTCTCCTTCCCGCCTCCCACCCGAGCCCCCTCAGCTTGCCTCCCCAC | 12 |
| wt-13410-13530 | 4394 | CGGGTCCATCAGGCCGGCCGGAGGGACCCCGGCGGCCCGGTGTCAGTCCCCCCTGCAGCCGCCCAGTCTCTGCCTCCAGGCAAGGGCGCCAGCTTTTCTCCCCCCAGCCTGAGGCCCAGT | 8 |
| wt-13530-13650 | 4395 | CTCCTGTGCACTGTCTGTAAAGTCCAGCCTCCCACGCCCGTCCACGGCTCCCGGGCCCAGCCTCGTCCACCCCTCCCCACGGTGGACAGGCCCTCTGTCCACCCGGGCCATCCCCGCCCC | 8 |
| wt-13650-13770 | 4396 | CCTGTGTCCACCCCAGTCCCGTCCAGGGGGGACTTTATGTGACCCTTGGGCCTGGCTCCCCATAGACTCCCATGTAAGCCTGCCTCGAGTAGGTGCCTCCAGAGCCCCTTTTGCCCCCCT | 4 |
| wt-13770-13890 | 4397 | GGCGGCCCAGCCCGACCCCCGGGCGCCCCCAAACTTTGTCCAGATGTCCAGGGGTCCCCGAGGGTGAGGCCCAGCCCCCTCCCGCCCCTGTCCACTGCCCCGGTCCCCCCAGAAGCCCCC | 12 |
| wt-13890-14010 | 4398 | AAAAGTAGAGGCTCAGGCCATGCGCGCCCTGTCACCAGGCCTGCCAAAGAGCCAGATCTAAGGCCGGGAGAGGCAGCCCCAAAGCGGGTGCAGTAACAGGTAATCTCTGGTAGTGATTTG | 4 |
| wt-14010-14130 | 4399 | GACCCGAAATCTGACACTTTAGAGCTCTGGAGGACTTTAAAACTCTAAAAATCAAAACTTTAGAGGCGAATGGGCGCCATTTTGTCCCCACGCGCGCATAATGGCGGACCTAGGCCTAAA | 4 |
| wt-14130-14250 | 4400 | ACCCCCAGGAAGCGGGTCTATGGTTGGCTGCGCTGCTGCTATCTTTAGAGGGGAAAAGAGGAATAAGCCCCCAGACAGGGGAGTGGGCTTGTTTGTGACTTCACCAAAGGTCAGGGCCCA | 4 |
| wt-14250-14370 | 4401 | AGGGGGTTCGCGTTGCTAGGCCACCTTCTCAGTCCAGCGCGTTTACGTAAGCCAGACAGCAGCCAATTGTCAGTTCTAGGGAGGGGGACCACTGCCCCTGGTATAAAGTGGTCCTGCAGC | 4 |
| wt-14370-14490 | 4402 | TATTTCTGGTCGCATCAGAGCGCCAGGAGTCCACACAAATGTAAGAGGGGGTCTTCTACCTCTCCCTAGCCCTCCGCCCCCTCCAAGGACTCGGGCCCAGTTTCTAACTTTTCCCCTTCC | 4 |
| wt-14490-14610 | 4403 | CTCCCTCGTCTTGCCCTGCGCCCGGGGCCACCTTCATCACCGTCGCTGACTCCGCCATCCAAGCCTAGGGGAGACCGAAGTGAAGGCCCTGGACCAACCCGGCCCGGGCCCCCCGGTATC | 8 |
| wt-14610-14730 | 4404 | GGGCCAGAGGTAAGTGGACTTTAATTTTTTCTGCTAAGCCCAACACTCCACCACACCCAGGCACACACTACACACACCCACCCGTCTCAGGGTCCCCTCGGACAGCTCCTAAGAAGGCAC | 4 |
| wt-14730-14850 | 4405 | CGGTCGCCCAGTCCTACCAGAGGGGGCCAAGAACCCAGACGAGTCCGTAGAAGGGTCCTCGTCCAGCAAGAAGAGGAGGTGGTAAGCGGTTCACCTTCAGGGGTAAGTAACCTGACCTCT | 4 |
| wt-14850-14970 | 4406 | CCAGGGCTCACATAAAGGGAGGCTTAGTATACATGCTTCTTGCTTTTCACAGGAACCTGGGGGCTAGTCTGGGTGGGATTAGGCTGCCTCAAGTTGCATCAGCCAGGGCTTCATGCCCTC | 4 |
| wt-14970-15090 | 4407 | CTCAGTTCCCTAGTCCCCGGGCTTCAGGCCCCCTCCGTCCCCGTCCTCCAGAGACCCGGGCTTCAGGCCCTGCCTCTCCTGTTACCCTTTTAGAACCACAGCCTGGACACATGTGCCAGA | 4 |
| wt-15090-15210 | 4408 | CGCCTTGGCCTCTAAGGCCCTCGGGTCCCCCTGGACCCCGGCCTCAGCAACCCTGCTGCTCCCCTCCTGCCACCCCAGCCTCCCCCCCTCCCCGTCCCCCTTCGCTCCTGATCCTCCCCC | 8 |
| wt-15210-15330 | 4409 | GGTCCCCAGTAGGGCCGCCTGCCCCCCTGCACCCAGTACCTGCCCCTCTTGGCCACGCACCCCGGGCCAGGCCACCTTAGACCCGGCCAAGCCCCATCCCTGAAGACCCAGCGGCCATTC | 8 |
| wt-15330-15450 | 4410 | TCTCTGGTAACGAGCAGAGAAGAAGTAGAGGCCCGCGGCCATTGGGCCCAGATTGAGAGACCAGTCCAGGGGCCCGAGGTTGGAGCCAGCGGGCACCCGAGGTCCCAGCACCCGGTCCCT | 8 |
| wt-15450-15570 | 4411 | CCGGGGGGCAGAGACAGGCAGGGCCCCCCGGCAGCTGGCCCCGAGGAGGCGCCCGGAGTGGGGCCGGTCGGCTGGGCTGGCCGAGCCCGGGTCTGGGAGGTCTGGGGTGGCGAGCCTGCT | 12 |
| wt-15570-15690 | 4412 | GTCTCAGGAGGGGCCTGGCTCCGCCGGGTGGCCCTGGGGTAAGTCTGGGAGGCAGAGGGTCGGCCTAGGCCCGGGGAAGTGGAGGGGGATCGCCCGGGTCTCTGTTGGCAGAGTCCGGGC | 8 |
| wt-15690-15810 | 4413 | GATCCTCTGAGACCCTCCGGGCCCGGACGGTCGCCCTCAGCCCCCCAGACAGACCCCAGGGTCTCCAGGCAGGGTCCGGCATCTTCAGGGGCAGCAGGCTCACCACCACAGGCCCCCCAG | 8 |
| wt-15810-15930 | 4414 | ACCCGGGTCTCGGCCAGCCGAGCCGACCGGCCCCGCGCCTGGCGCCTCCTCGGGGCCAGCCGCCGGGGTTGGTTCTGCCCCTCTCTCTGTCCTTCAGAGGAACCAGGGACCTCGGGCACC | 8 |
| wt-15930-16050 | 4415 | CCAGAGCCCCTCGGGCCCGCCTCCAGGCGCCCTCCTGGTCTCCGCTCCCCTCTGAGCCCCGTTAAACCCAAAGAATGTCTGAGGGGAGCCACCCTCGGGGCCCAGGCCCCAGAGTCCAGA | 8 |
| wt-16050-16170 | 4416 | GGTCAGGGGCACCTCAGGGTGCCTCCCCGGGTCCCAGGCCAGCCGGAGGGACCCCGGCAGCCCGGGCGGCCCCAGAGGCCGGTTCCTCGCCCCTTCCCCGGGCTTCAGAGCCCAGGATGT | 12 |
| wt-16170-16290 | 4417 | CCCCCAGAAGGGACCCTAGGCGTCCCCTCTCCTCCCCTCCAGGCCCGAGCCTCTCCCTCGCGGAGAGGGGCCTCTTTGGGCCCTCAAGTCCAGCCCCACCGAGACCCGAGTGGCCCGGAT | 8 |
| wt-16290-16410 | 4418 | CCCCCCACCGGCCCTTCTCTCTGTCCCCCTGCTCCTCTCCAACCTTCGCTCCACCCTAGACCCCAGCTTCTGGCCTCCCCGGGTCCACCAGGCCAGCCGGAGGGACCCCGGCAGCCCGGG | 8 |
| wt-16410-16530 | 4419 | CGAGTCGCCTTCCCTCTCCCCTGGCCTCTCCTTCCCGCCTCCCACCCGAGCCCCCTCAGCTTGCCTCCCCACCGGGTCCATCAGGCCGGCCGGAGGGACCCCGGCGGCCCGGTGTCAGTC | 8 |
| wt-16530-16650 | 4420 | CCCCCTGCAGCCGCCCAGTCTCTGCCTCCAGGCAAGGGCGCCAGCTTTTCTCCCCCCAGCCTGAGGCCCAGTCTCCTGTGCACTGTCTGTAAAGTCCAGCCTCCCACGCCCGTCCACGGC | 8 |
| wt-16650-16770 | 4421 | TCCCGGGCCCAGCCTCGTCCACCCCTCCCCACGGTGGACAGGCCCTCTGTCCACCCGGGCCATCCCCGCCCCCCTGTGTCCACCCCAGTCCCGTCCAGGGGGGACTTTATGTGACCCTTG | 8 |
| wt-16770-16890 | 4422 | GGCCTGGCTCCCCATAGACTCCCATGTAAGCCTGCCTCGAGTAGGTGCCTCCAGAGCCCCTTTTGCCCCCCTGGCGGCCCAGCCCGACCCCCGGGCGCCCCCAAACTTTGTCCAGATGTC | 8 |
| wt-16890-17010 | 4423 | CAGGGGTCCCCGAGGGTGAGGCCCAGCCCCCTCCCGCCCCTGTCCACTGCCCCGGTCCCCCCAGAAGCCCCCAAAAGTAGAGGCTCAGGCCATGCGCGCCCTGTCACCAGGCCTGCCAAA | 8 |
| wt-17010-17130 | 4424 | GAGCCAGATCTAAGGCCGGGAGAGGCAGCCCCAAAGCGGGTGCAGTAACAGGTAATCTCTGGTAGTGATTTGGACCCGAAATCTGACACTTTAGAGCTCTGGAGGACTTTAAAACTCTAA | 4 |

| | | | |
|---|---|---|---|
| wt-17130-17250 | 4425 | AAATCAAAACTTTAGAGGCGAATGGGCGCCATTTTGTCCCCACGCGCGCATAATGGCGGACCTAGGCCTAAAACCCCCAGGAAGCGGGTCTATGGTTGGCTGCGCTGCTGCTATCTTTAG | 4 |
| wt-17250-17370 | 4426 | AGGGGAAAAGAGGAATAAGCCCCCAGACAGGGGAGTGGGCTTGTTTGTGACTTCACCAAAGGTCAGGGCCCAAGGGGGTTCGCGTTGCTAGGCCACCTTCTCAGTCCAGCGCGTTTACGT | 4 |
| wt-17370-17490 | 4427 | AAGCCAGACAGCAGCCAATTGTCAGTTCTAGGGAGGGGGACCACTGCCCCTGGTATAAAGTGGTCCTGCAGCTATTTCTGGTCGCATCAGAGCGCCAGGAGTCCACACAAATGTAAGAGG | 4 |
| wt-17610-17730 | 4429 | ACTCCGCCATCCAAGCCTAGGGGAGACCGAAGTGAAGGCCCTGGACCAACCCGGCCCGGGCCCCCCGGTATCGGGCCAGAGGTAAGTGGACTTTAATTTTTTCTGCTAAGCCCAACACTC | 4 |
| wt-17730-17850 | 4430 | CACCACACCCAGGCACACACTACACACACCCACCCGTCTCAGGGTCCCCTCGGACAGCTCCTAAGAAGGCACCGGTCGCCCAGTCCTACCAGAGGGGGCCAAGAACCCAGACGAGTCCGT | 4 |
| wt-17850-17970 | 4431 | AGAAGGGTCCTCGTCCAGCAAGAAGAGGAGGTGGTAAGCGGTTCACCTTCAGGGGTAAGTAACCTGACCTCTCCAGGGCTCACATAAAGGGAGGCTTAGTATACATGCTTCTTGCTTTTC | 4 |
| wt-17970-18090 | 4432 | ACAGGAACCTGGGGGCTAGTCTGGGTGGGATTAGGCTGCCTCAAGTTGCATCAGCCAGGGCTTCATGCCCTCCTCAGTTCCCTAGTCCCCGGGCTTCAGGCCCCCTCCGTCCCCGTCCTC | 4 |
| wt-18210-18330 | 4434 | AACCCTGCTGCTCCCCTCCTGCCACCCCAGCCTCCCCCCCTCCCCGTCCCCCTTCGCTCCTGATCCTCCCCCGGTCCCCAGTAGGGCCGCCTGCCCCCCTGCACCCAGTACCTGCCCCTC | 8 |
| wt-18330-18450 | 4435 | TTGGCCACGCACCCCGGGCCAGGCCACCTTAGACCCGGCCAAGCCCCATCCCTGAAGACCCAGCGGCCATTCTCTCTGGTAACGAGCAGAGAAGAAGTAGAGGCCCGCGGCCATTGGGCC | 8 |
| wt-18450-18570 | 4436 | CAGATTGAGAGACCAGTCCAGGGGCCCGAGGTTGGAGCCAGCGGGCACCCGAGGTCCCAGCACCCGGTCCCTCCGGGGGGCAGAGACAGGCAGGGCCCCCCGGCAGCTGGCCCCGAGGAG | 8 |
| wt-18570-18690 | 4437 | GCGCCCGGAGTGGGGCCGGTCGGCTGGGCTGGCCGAGCCCGGGTCTGGGAGGTCTGGGGTGGCGAGCCTGCTGTCTCAGGAGGGGCCTGGCTCCGCCGGGTGGCCCTGGGGTAAGTCTGG | 12 |
| wt-18690-18810 | 4438 | GAGGCAGAGGGTCGGCCTAGGCCCGGGGAAGTGGAGGGGGATCGCCCGGGTCTCTGTTGGCAGAGTCCGGGCGATCCTCTGAGACCCTCCGGGCCCGGACGGTCGCCCTCAGCCCCCCAG | 8 |
| wt-18810-18930 | 4439 | ACAGACCCCAGGGTCTCCAGGCAGGGTCCGGCATCTTCAGGGGCAGCAGGCTCACCACCACAGGCCCCCCAGACCCGGGTCTCGGCCAGCCGAGCCGACCGGCCCCGCGCCTGGCGCCTC | 8 |
| wt-18930-19050 | 4440 | CTCGGGGCCAGCCGCCGGGGTTGGTTCTGCCCCTCTCTCTGTCCTTCAGAGGAACCAGGGACCTCGGGCACCCCAGAGCCCCTCGGGCCCGCCTCCAGGCGCCCTCCTGGTCTCCGCTCC | 8 |
| wt-19050-19170 | 4441 | CCTCTGAGCCCCGTTAAACCCAAAGAATGTCTGAGGGGAGCCACCCTCGGGGCCCAGGCCCCAGAGTCCAGAGGTCAGGGGCACCTCAGGGTGCCTCCCCGGGTCCCAGGCCAGCCGGAG | 8 |
| wt-19170-19290 | 4442 | GGACCCCGGCAGCCCGGGCGGCCCCAGAGGCCGGTTCCTCGCCCCTTCCCCGGGCTTCAGAGCCCAGGATGTCCCCCAGAAGGGACCCTAGGCGTCCCCTCTCCTCCCCTCCAGGCCCGA | 8 |
| wt-19290-19410 | 4443 | GCCTCTCCCTCGCGGAGAGGGGCCTCTTTGGGCCCTCAAGTCCAGCCCCACCGAGACCCGAGTGGCCCGGATCCCCCCACCGGCCCTTCTCTCTGTCCCCCTGCTCCTCTCCAACCTTCG | 8 |
| wt-19410-19530 | 4444 | CTCCACCCTAGACCCCAGCTTCTGGCCTCCCCGGGTCCACCAGGCCAGCCGGAGGGACCCCGGCAGCCCGGGCGAGTCGCCTTCCCTCTCCCCTGGCCTCTCCTTCCCGCCTCCCACCCG | 8 |
| wt-19530-19650 | 4445 | AGCCCCCTCAGCTTGCCTCCCCACCGGGTCCATCAGGCCGGCCGGAGGGACCCCGGCGGCCCGGTGTCAGTCCCCCCTGCAGCCGCCCAGTCTCTGCCTCCAGGCAAGGGCGCCAGCTTT | 8 |
| wt-19650-19770 | 4446 | TCTCCCCCCAGCCTGAGGCCCAGTCTCCTGTGCACTGTCTGTAAAGTCCAGCCTCCCACGCCCGTCCACGGCTCCCGGGCCCAGCCTCGTCCACCCCTCCCCACGGTGGACAGGCCCTCT | 8 |
| wt-19770-19890 | 4447 | GTCCACCCGGGCCATCCCCGCCCCCCTGTGTCCACCCCAGTCCCGTCCAGGGGGGACTTTATGTGACCCTTGGGCCTGGCTCCCCATAGACTCCCATGTAAGCCTGCCTCGAGTAGGTGC | 8 |
| wt-19890-20010 | 4448 | CTCCAGAGCCCCTTTTGCCCCCCTGGCGGCCCAGCCCGACCCCCGGGCGCCCCCAAACTTTGTCCAGATGTCCAGGGGTCCCCGAGGGTGAGGCCCAGCCCCCTCCCGCCCCTGTCCACT | 8 |
| wt-20010-20130 | 4449 | GCCCCGGTCCCCCCAGAAGCCCCCAAAAGTAGAGGCTCAGGCCATGCGCGCCCTGTCACCAGGCCTGCCAAAGAGCCAGATCTAAGGCCGGGAGAGGCAGCCCCAAAGCGGGTGCAGTAA | 8 |
| wt-20130-20250 | 4450 | CAGGTAATCTCTGGTAGTGATTTGGACCCGAAATCTGACACTTTAGAGCTCTGGAGGACTTTAAAACTCTAAAAATCAAAACTTTAGAGGCGAATGGGCGCCATTTTGTCCCCACGCGCG | 4 |
| wt-20250-20370 | 4451 | CATAATGGCGGACCTAGGCCTAAAACCCCCAGGAAGCGGGTCTATGGTTGGCTGCGCTGCTGCTATCTTTAGAGGGGAAAAGAGGAATAAGCCCCCAGACAGGGGAGTGGGCTTGTTTGT | 4 |
| wt-20370-20490 | 4452 | GACTTCACCAAAGGTCAGGGCCCAAGGGGGTTCGCGTTGCTAGGCCACCTTCTCAGTCCAGCGCGTTTACGTAAGCCAGACAGCAGCCAATTGTCAGTTCTAGGGAGGGGGACCACTGCC | 4 |
| wt-20490-20610 | 4453 | CCTGGTATAAAGTGGTCCTGCAGCTATTTCTGGTCGCATCAGAGCGCCAGGAGTCCACACAAATGTAAGAGGGGGGTCTTCTACCTCTCCCTAGCCCTCCGCCCCCTCCAAGGACTCGGGC | 4 |
| wt-20610-20730 | 4454 | CCAGTTTCTAACTTTTCCCCTTCCCTCCCTCGTCTTGCCCTGCGCCCGGGGCCACCTTCATCACCGTCGCTGACTCCGCCATCCAAGCCTAGGGGAGACCGAAGTGAAGGCCCTGGACCA | 4 |
| wt-20730-20850 | 4455 | ACCCGGCCCGGGCCCCCCGGTATCGGGCCAGAGGTAAGTGGACTTTAATTTTTTCTGCTAAGCCCAACACTCCACCACACCCAGGCACACACTACACACACCCACCCGTCTCAGGGTCCC | 4 |
| wt-20850-20970 | 4456 | CTCGGACAGCTCCTAAGAAGGCACCGGTCGCCCAGTCCTACCAGAGGGGGCCAAGAACCCAGACGAGTCCGTAGAAGGGTCCTCGTCCAGCAAGAAGAGGAGGTGGTAAGCGGTTCACCT | 4 |
| wt-20970-21090 | 4457 | TCAGGGGTAAGTAACCTGACCTCTCCAGGGCTCACATAAAGGGAGGCTTAGTATACATGCTTCTTGCTTTTCACAGGAACCTGGGGGCTAGTCTGGGTGGGATTAGGCTGCCTCAAGTTG | 4 |
| wt-21090-21210 | 4458 | CATCAGCCAGGGCTTCATGCCCTCCTCAGTTCCCTAGTCCCCGGGCTTCAGGCCCCCTCCGTCCCCGTCCTCCAGAGACCCGGGCTTCAGGCCCTGCCTCTCCTGTTACCCTTTTAGAAC | 4 |
| wt-21210-21330 | 4459 | CACAGCCTGGACACATGTGCCAGACGCCTTGGCCTCTAAGGCCCTCGGGTCCCCCTGGACCCCGGCCTCAGCAACCCTGCTGCTCCCCTCCTGCCACCCCAGCCTCCCCCCCTCCCCGTC | 8 |
| wt-21330-21450 | 4460 | CCCCTTCGCTCCTGATCCTCCCCCGGTCCCCAGTAGGGCCGCCTGCCCCCCTGCACCCAGTACCTGCCCCTCTTGGCCACGCACCCCGGGCCAGGCCACCTTAGACCCGGCCAAGCCCCA | 8 |
| wt-21450-21570 | 4461 | TCCCTGAAGACCCAGCGGCCATTCTCTCTGGTAACGAGCAGAGAAGAAGTAGAGGCCCGCGGCCATTGGGCCCAGATTGAGAGACCAGTCCAGGGGCCCGAGGTTGGAGCCAGCGGGCAC | 4 |
| wt-21570-21690 | 4462 | CCGAGGTCCCAGCACCCGGTCCCTCCGGGGGGCAGAGACAGGCAGGGCCCCCCGGCAGCTGGCCCCGAGGAGGCGCCCGGAGTGGGGCCGGTCGGCTGGGCTGGCCGAGCCCGGGTCTGG | 12 |
| wt-21690-21810 | 4463 | GAGGTCTGGGGTGGCGAGCCTGCTGTCTCAGGAGGGGCCTGGCTCCGCCGGGTGGCCCTGGGGTAAGTCTGGGAGGCAGAGGGTCGGCCTAGGCCCGGGGAAGTGGAGGGGGATCGCCCG | 8 |
| wt-21810-21930 | 4464 | GGTCTCTGTTGGCAGAGTCCGGGCGATCCTCTGAGACCCTCCGGGCCCGGACGGTCGCCCTCAGCCCCCCAGACAGACCCCAGGGTCTCCAGGCAGGGTCCGGCATCTTCAGGGGCAGCA | 8 |
| wt-21930-22050 | 4465 | GGCTCACCACCACAGGCCCCCCAGACCCGGGTCTCGGCCAGCCGAGCCGACCGGCCCCGCGCCTGGCGCCTCCTCGGGGCCAGCCGCCGGGGTTGGTTCTGCCCCTCTCTCTGTCCTTCA | 12 |
| wt-22050-22170 | 4466 | GAGGAACCAGGGACCTCGGGCACCCCAGAGCCCCTCGGGCCCGCCTCCAGGCGCCCTCCTGGTCTCCGCTCCCCTCTGAGCCCCGTTAAACCCAAAGAATGTCTGAGGGGAGCCACCCTC | 8 |
| wt-22170-22290 | 4467 | GGGGCCCAGGCCCCAGAGTCCAGAGGTCAGGGGCACCTCAGGGTGCCTCCCCGGGTCCCAGGCCAGCCGGAGGGACCCCGGCAGCCCGGGCGGCCCCAGAGGCCGGTTCCTCGCCCCTTC | 12 |
| wt-22290-22410 | 4468 | CCCGGGCTTCAGAGCCCAGGATGTCCCCCAGAAGGGACCCTAGGCGTCCCCTCTCCTCCCCTCCAGGCCCGAGCCTCTCCCTCGCGGAGAGGGGCCTCTTTGGGCCCTCAAGTCCAGCCC | 8 |
| wt-22410-22530 | 4469 | CACCGAGACCCGAGTGGCCCGGATCCCCCCACCGGCCCTTCTCTCTGTCCCCCTGCTCCTCTCCAACCTTCGCTCCACCCTAGACCCCAGCTTCTGGCCTCCCCGGGTCCACCAGGCCAG | 8 |
| wt-22530-22650 | 4470 | CCGGAGGGACCCCGGCAGCCCGGGCGAGTCGCCTTCCCTCTCCCCTGGCCTCTCCTTCCCGCCTCCCACCCGAGCCCCCTCAGCTTGCCTCCCCACCGGGTCCATCAGGCCGGCCGGAGG | 12 |
| wt-22650-22770 | 4471 | GACCCCGGCGGCCCGGTGTCAGTCCCCCCTGCAGCCGCCCAGTCTCTGCCTCCAGGCAAGGGCGCCAGCTTTTCTCCCCCCAGCCTGAGGCCCAGTCTCCTGTGCACTGTCTGTAAAGTC | 8 |
| wt-22770-22890 | 4472 | CAGCCTCCCACGCCCGTCCACGGCTCCCGGGCCCAGCCTCGTCCACCCCTCCCCACGGTGGACAGGCCCTCTGTCCACCCGGGCCATCCCCGCCCCCCTGTGTCCACCCCAGTCCCGTCC | 12 |
| wt-22890-23010 | 4473 | AGGGGGGACTTTATGTGACCCTTGGGCCTGGCTCCCCATAGACTCCCATGTAAGCCTGCCTCGAGTAGGTGCCTCCAGAGCCCCTTTTGCCCCCCTGGCGGCCCAGCCCGACCCCCGGGC | 8 |
| wt-23010-23130 | 4474 | GCCCCCAAACTTTGTCCAGATGTCCAGGGGTCCCCGAGGGTGAGGCCCAGCCCCCTCCCGCCCCTGTCCACTGCCCCGGTCCCCCCAGAAGCCCCCAAAAGTAGAGGCTCAGGCCATGCG | 8 |
| wt-23130-23250 | 4475 | CGCCCTGTCACCAGGCCTGCCAAAGAGCCAGATCTAAGGCCGGGAGAGGCAGCCCCAAAGCGGGTGCAGTAACAGGTAATCTCTGGTAGTGATTTGGACCCGAAATCTGACACTTTAGAG | 4 |
| wt-23250-23370 | 4476 | CTCTGGAGGACTTTAAAACTCTAAAAATCAAAACTTTAGAGGCGAATGGGCGCCATTTTGTCCCCACGCGCGCATAATGGCGGACCTAGGCCTAAAACCCCCAGGAAGCGGGTCTATGGT | 4 |
| wt-23370-23490 | 4477 | TGGCTGCGCTGCTGCTATCTTTAGAGGGGAAAAGAGGAATAAGCCCCCAGACAGGGGAGTGGGCTTGTTTGTGACTTCACCAAAGGTCAGGGCCCAAGGGGGTTCGCGTTGCTAGGCCAC | 4 |

| | | | |
|---|---|---|---|
| wt-23490-23610 | 4478 | CTTCTCAGTCCAGCGCGTTTACGTAAGCCAGACAGCAGCCAATTGTCAGTTCTAGGGAGGGGGACCACTGCCCCTGGTATAAAGTGGTCCTGCAGCTATTTCTGGTCGCATCAGAGCGCC | 4 |
| wt-23610-23730 | 4479 | AGGAGTCCACACAAATGTAAGAGGGGGTCTTCTACCTCTCCCTAGCCCTCCGCCCCCTCCAAGGACTCGGGCCCAGTTTCTAACTTTTCCCCTTCCCTCCCTCGTCTTGCCCTGCGCCCG | 4 |
| wt-23850-23970 | 4481 | TTTTTTCTGCTAAGCCCAACACTCCACCACACCCAGGCACACACTACACACACCCACCCGTCTCAGGGTCCCCTCGGACAGCTCCTAAGAAGGCACCGGTCGCCCAGTCCTACCAGAGGG | 4 |
| wt-23970-24090 | 4482 | GGCCAAGAACCCAGACGAGTCCGTAGAAGGGTCCTCGTCCAGCAAGAAGAGGAGGTGGTAAGCGGTTCACCTTCAGGGGTAAGTAACCTGACCTCTCCAGGGCTCACATAAAGGGAGGCT | 4 |
| wt-24090-24210 | 4483 | TAGTATACATGCTTCTTGCTTTTCACAGGAACCTGGGGGCTAGTCTGGGTGGGATTAGGCTGCCTCAAGTTGCATCAGCCAGGGCTTCATGCCCTCCTCAGTTCCCTAGTCCCCGGGCTT | 4 |
| wt-24330-24450 | 4485 | GTCCCCCTGGACCCCGGCCTCAGCAACCCTGCTGCTCCCCTCCTGCCACCCCAGCCTCCCCCCCTCCCCGTCCCCCTTCGCTCCTGATCCTCCCCCGGTCCCCAGTAGGGCCGCCTGCCC | 12 |
| wt-24570-24690 | 4487 | GTAGAGGCCCGCGGCCATTGGGCCCAGATTGAGAGACCAGTCCAGGGGCCCGAGGTTGGAGCCAGCGGGCACCCGAGGTCCCAGCACCCGGTCCCTCCGGGGGGCAGAGACAGGCAGGGC | 8 |
| wt-24690-24810 | 4488 | CCCCCGGCAGCTGGCCCCGAGGAGGCGCCCGGAGTGGGGCCGGTCGGCTGGGCTGGCCGAGCCCGGGTCTGGGAGGTCTGGGGTGGCGAGCCTGCTGTCTCAGGAGGGGCCTGGCTCCGC | 12 |
| wt-24810-24930 | 4489 | CGGGTGGCCCTGGGGTAAGTCTGGGAGGCAGAGGGTCGGCCTAGGCCCGGGGAAGTGGAGGGGGATCGCCCGGGTCTCTGTTGGCAGAGTCCGGGCGATCCTCTGAGACCCTCCGGGCCC | 8 |
| wt-24930-25050 | 4490 | GGACGGTCGCCCTCAGCCCCCCAGACAGACCCCAGGGTCTCCAGGCAGGGTCCGGCATCTTCAGGGGCAGCAGGCTCACCACCACAGGCCCCCCAGACCCGGGTCTCGGCCAGCCGAGCC | 8 |
| wt-25050-25170 | 4491 | GACCGGCCCCGCGCCTGGCGCCTCCTCGGGGCCAGCCGCCGGGGTTGGTTCTGCCCCTCTCTCTGTCCTTCAGAGGAACCAGGGACCTCGGGCACCCCAGAGCCCCTCGGGCCCGCCTCC | 12 |
| wt-25170-25290 | 4492 | AGGCGCCCTCCTGGTCTCCGCTCCCCTCTGAGCCCCGTTAAACCCAAAGAATGTCTGAGGGGAGCCACCCTCGGGGCCCAGGCCCCAGAGTCCAGAGGTCAGGGGCACCTCAGGGTGCCT | 8 |
| wt-25290-25410 | 4493 | CCCCGGGTCCCAGGCCAGCCGGAGGGACCCCGGCAGCCCGGGCGGCCCCAGAGGCCGGTTCCTCGCCCCTTCCCCGGGCTTCAGAGCCCAGGATGTCCCCCAGAAGGGACCCTAGGCGTC | 12 |
| wt-25410-25530 | 4494 | CCCTCTCCTCCCCTCCAGGCCCGAGCCTCTCCCTCGCGGAGAGGGGCCTCTTTGGGCCCTCAAGTCCAGCCCCACCGAGACCCGAGTGGCCCGGATCCCCCCACCGGCCCTTCTCTCTGT | 8 |
| wt-25530-25650 | 4495 | CCCCCTGCTCCTCTCCAACCTTCGCTCCACCCTAGACCCCAGCTTCTGGCCTCCCCGGGTCCACCAGGCCAGCCGGAGGGACCCCGGCAGCCCGGGCGAGTCGCCTTCCCTCTCCCCTGG | 8 |
| wt-25650-25770 | 4496 | CCTCTCCTTCCCGCCTCCCACCCGAGCCCCCTCAGCTTGCCTCCCCACCGGGTCCATCAGGCCGGCCGGAGGGACCCCGGCGGCCCGGTGTCAGTCCCCCCTGCAGCCGCCCAGTCTCTG | 12 |
| wt-25770-25890 | 4497 | CCTCCAGGCAAGGGCGCCAGCTTTTCTCCCCCCAGCCTGAGGCCCAGTCTCCTGTGCACTGTCTGTAAAGTCCAGCCTCCCACGCCCGTCCACGGCTCCCGGGCCCAGCCTCGTCCACCC | 8 |
| wt-25890-26010 | 4498 | CTCCCCACGGTGGACAGGCCCTCTGTCCACCCGGGCCATCCCCGCCCCCCTGTGTCCACCCCAGTCCCGTCCAGGGGGGACTTTATGTGACCCTTGGGCCTGGCTCCCCATAGACTCCCA | 8 |
| wt-26010-26130 | 4499 | TGTAAGCCTGCCTCGAGTAGGTGCCTCCAGAGCCCCTTTTGCCCCCCTGGCGGCCCAGCCCGACCCCCGGGCGCCCCCAAACTTTGTCCAGATGTCCAGGGGTCCCCGAGGGTGAGGCCC | 8 |
| wt-26130-26250 | 4500 | AGCCCCCTCCCGCCCCTGTCCACTGCCCCGGTCCCCCCAGAAGCCCCCAAAAGTAGAGGCTCAGGCCATGCGCGCCCTGTCACCAGGCCTGCCAAAGAGCCAGATCTAAGGCCGGGAGAG | 8 |
| wt-26250-26370 | 4501 | GCAGCCCCAAAGCGGGTGCAGTAACAGGTAATCTCTGGTAGTGATTTGGACCCGAAATCTGACACTTTAGAGCTCTGGAGGACTTTAAAACTCTAAAAATCAAAACTTTAGAGGCGAATG | 4 |
| wt-26370-26490 | 4502 | GGCGCCATTTTGTCCCCACGCGCGCATAATGGCGGACCTAGGCCTAAAACCCCCAGGAAGCGGGTCTATGGTTGGCTGCGCTGCTGCTATCTTTAGAGGGGAAAAGAGGAATAAGCCCCC | 4 |
| wt-26490-26610 | 4503 | AGACAGGGGAGTGGGCTTGTTTGTGACTTCACCAAAGGTCAGGGCCCAAGGGGGTTCGCGTTGCTAGGCCACCTTCTCAGTCCAGCGCGTTTACGTAAGCCAGACAGCAGCCAATTGTCA | 4 |
| wt-26610-26730 | 4504 | GTTCTAGGGAGGGGGACCACTGCCCCTGGTATAAAGTGGTCCTGCAGCTATTTCTGGTCGCATCAGAGCGCCAGGAGTCCACACAAATGTAAGAGGGGGTCTTCTACCTCTCCCTAGCCC | 4 |
| wt-26850-26970 | 4506 | GACCGAAGTGAAGGCCCTGGACCAACCCGGCCCGGGCCCCCCGGTATCGGGCCAGAGGTAAGTGGACTTTAATTTTTTCTGCTAAGCCCAACACTCCACCACACCCAGGCACACACTACA | 4 |
| wt-26970-27090 | 4507 | CACACCCACCCGTCTCAGGGTCCCCTCGGACAGCTCCTAAGAAGGCACCGGTCGCCCAGTCCTACCAGAGGGGGCCAAGAACCCAGACGAGTCCGTAGAAGGGTCCTCGTCCAGCAAGAA | 4 |
| wt-27090-27210 | 4508 | GAGGAGGTGGTAAGCGGTTCACCTTCAGGGGTAAGTAACCTGACCTCTCCAGGGCTCACATAAAGGGAGGCTTAGTATACATGCTTCTTGCTTTTCACAGGAACCTGGGGGCTAGTCTGG | 4 |
| wt-27210-27330 | 4509 | GTGGGATTAGGCTGCCTCAAGTTGCATCAGCCAGGGCTTCATGCCCTCCTCAGTTCCCTAGTCCCCGGGCTTCAGGCCCCCTCCGTCCCCGTCCTCCAGAGACCCGGGCTTCAGGCCCTG | 8 |
| wt-27330-27450 | 4510 | CCTCTCCTGTTACCCTTTTAGAACCACAGCCTGGACACATGTGCCAGACGCCTTGGCCTCTAAGGCCCTCGGGTCCCCCTGGACCCCGGCCTCAGCAACCCTGCTGCTCCCCTCCTGCCA | 4 |
| wt-27450-27570 | 4511 | CCCCAGCCTCCCCCCCTCCCCGTCCCCCTTCGCTCCTGATCCTCCCCCGGTCCCCAGTAGGGCCGCCTGCCCCCCTGCACCCAGTACCTGCCCCTCTTGGCCACGCACCCCGGGCCAGGC | 12 |
| wt-27570-27690 | 4512 | CACCTTAGACCCGGCCAAGCCCCATCCCTGAAGACCCAGCGGCCATTCTCTCTGGTAACGAGCAGAGAAGAAGTAGAGGCCCGCGGCCATTGGGCCCAGATTGAGAGACCAGTCCAGGGG | 4 |
| wt-27690-27810 | 4513 | CCCGAGGTTGGAGCCAGCGGGCACCCGAGGTCCCAGCACCCGGTCCCTCCGGGGGGCAGAGACAGGCAGGGCCCCCCGGCAGCTGGCCCCGAGGAGGCGCCCGGAGTGGGGCCGGTCGGC | 12 |
| wt-27810-27930 | 4514 | TGGGCTGGCCGAGCCCGGGTCTGGGAGGTCTGGGGTGGCGAGCCTGCTGTCTCAGGAGGGGCCTGGCTCCGCCGGGTGGCCCTGGGGTAAGTCTGGGAGGCAGAGGGTCGGCCTAGGCCC | 8 |
| wt-27930-28050 | 4515 | GGGGAAGTGGAGGGGGATCGCCCGGGTCTCTGTTGGCAGAGTCCGGGCGATCCTCTGAGACCCTCCGGGCCCGGACGGTCGCCCTCAGCCCCCCAGACAGACCCCAGGGTCTCCAGGCAG | 8 |
| wt-28050-28170 | 4516 | GGTCCGGCATCTTCAGGGGCAGCAGGCTCACCACCACAGGCCCCCCAGACCCGGGTCTCGGCCAGCCGAGCCGACCGGCCCCGCGCCTGGCGCCTCCTCGGGGCCAGCCGCCGGGGTTGG | 12 |
| wt-28170-28290 | 4517 | TTCTGCCCCTCTCTCTGTCCTTCAGAGGAACCAGGGACCTCGGGCACCCCAGAGCCCCTCGGGCCCGCCTCCAGGCGCCCTCCTGGTCTCCGCTCCCCTCTGAGCCCCGTTAAACCCAAA | 8 |
| wt-28290-28410 | 4518 | GAATGTCTGAGGGGAGCCACCCTCGGGGCCCAGGCCCCAGAGTCCAGAGGTCAGGGGCACCTCAGGGTGCCTCCCCGGGTCCCAGGCCAGCCGGAGGGACCCCGGCAGCCCGGGCGGCCC | 12 |
| wt-28410-28530 | 4519 | CAGAGGCCGGTTCCTCGCCCCTTCCCCGGGCTTCAGAGCCCAGGATGTCCCCCAGAAGGGACCCTAGGCGTCCCCTCTCCTCCCCTCCAGGCCCGAGCCTCTCCCTCGCGGAGAGGGGCC | 8 |
| wt-28530-28650 | 4520 | TCTTTGGGCCCTCAAGTCCAGCCCCACCGAGACCCGAGTGGCCCGGATCCCCCCACCGGCCCTTCTCTCTGTCCCCCTGCTCCTCTCCAACCTTCGCTCCACCCTAGACCCCAGCTTCTG | 8 |
| wt-28650-28770 | 4521 | GCCTCCCCGGGTCCACCAGGCCAGCCGGAGGGACCCCGGCAGCCCGGGCGAGTCGCCTTCCCTCTCCCCTGGCCTCTCCTTCCCGCCTCCCACCCGAGCCCCCTCAGCTTGCCTCCCCAC | 12 |
| wt-28770-28890 | 4522 | CGGGTCCATCAGGCCGGCCGGAGGGACCCCGGCGGCCCGGTGTCAGTCCCCCCTGCAGCCGCCCAGTCTCTGCCTCCAGGCAAGGGCGCCAGCTTTTCTCCCCCCAGCCTGAGGCCCAGT | 8 |
| wt-28890-29010 | 4523 | CTCCTGTGCACTGTCTGTAAAGTCCAGCCTCCCACGCCCGTCCACGGCTCCCGGGCCCAGCCTCGTCCACCCCTCCCCACGGTGGACAGGCCCTCTGTCCACCCGGGCCATCCCCGCCCC | 8 |
| wt-29010-29130 | 4524 | CCTGTGTCCACCCCAGTCCCGTCCAGGGGGGACTTTATGTGACCCTTGGGCCTGGCTCCCCATAGACTCCCATGTAAGCCTGCCTCGAGTAGGTGCCTCCAGAGCCCCTTTTGCCCCCCT | 4 |
| wt-29130-29250 | 4525 | GGCGGCCCAGCCCGACCCCCGGGCGCCCCCAAACTTTGTCCAGATGTCCAGGGGTCCCCGAGGGTGAGGCCCAGCCCCCTCCCGCCCCTGTCCACTGCCCCGGTCCCCCCAGAAGCCCCC | 12 |
| wt-29250-29370 | 4526 | AAAAGTAGAGGCTCAGGCCATGCGCGCCCTGTCACCAGGCCTGCCAAAGAGCCAGATCTAAGGCCGGGAGAGGCAGCCCCAAAGCGGGTGCAGTAACAGGTAATCTCTGGTAGTGATTTG | 4 |
| wt-29370-29490 | 4527 | GACCCGAAATCTGACACTTTAGAGCTCTGGAGGACTTTAAAACTCTAAAAATCAAAACTTTAGAGGCGAATGGGCGCCATTTTGTCCCCACGCGCGCATAATGGCGGACCTAGGCCTAAA | 4 |
| wt-29490-29610 | 4528 | ACCCCCAGGAAGCGGGTCTATGGTTGGCTGCGCTGCTGCTATCTTTAGAGGGGAAAAGAGGAATAAGCCCCCAGACAGGGGAGTGGGCTTGTTTGTGACTTCACCAAAGGTCAGGGCCCA | 4 |
| wt-29610-29730 | 4529 | AGGGGGTTCGCGTTGCTAGGCCACCTTCTCAGTCCAGCGCGTTTACGTAAGCCAGACAGCAGCCAATTGTCAGTTCTAGGGAGGGGGACCACTGCCCCTGGTATAAAGTGGTCCTGCAGC | 4 |
| wt-29730-29850 | 4530 | TATTTCTGGTCGCATCAGAGCGCCAGGAGTCCACACAAATGTAAGAGGGGGTCTTCTACCTCTCCCTAGCCCTCCGCCCCCTCCAAGGACTCGGGCCCAGTTTCTAACTTTTCCCCTTCC | 4 |
| wt-29850-29970 | 4531 | CTCCCTCGTCTTGCCCTGCGCCCGGGGCCACCTTCATCACCGTCGCTGACTCCGCCATCCAAGCCTAGGGGAGACCGAAGTGAAGGCCCTGGACCAACCCGGCCCGGGCCCCCCGGTATC | 8 |
| wt-29970-30090 | 4532 | GGGCCAGAGGTAAGTGGACTTTAATTTTTTCTGCTAAGCCCAACACTCCACCACACCCAGGCACACACTACACACACCCACCCGTCTCAGGGTCCCCTCGGACAGCTCCTAAGAAGGCAC | 4 |

| | | | |
|---|---|---|---|
| wt-30090-30210 | 4533 | CGGTCGCCCAGTCCTACCAGAGGGGGCCAAGAACCCAGACGAGTCCGTAGAAGGGTCCTCGTCCAGCAAGAAGAGGAGGTGGTAAGCGGTTCACCTTCAGGGGTAAGTAACCTGACCTCT | 4 |
| wt-30210-30330 | 4534 | CCAGGGCTCACATAAAGGGAGGCTTAGTATACATGCTTCTTGCTTTTCACAGGAACCTGGGGGCTAGTCTGGGTGGGATTAGGCTGCCTCAAGTTGCATCAGCCAGGGCTTCATGCCCTC | 4 |
| wt-30330-30450 | 4535 | CTCAGTTCCCTAGTCCCCGGGCTTCAGGCCCCCTCCGTCCCCGTCCTCCAGAGACCCGGGCTTCAGGCCCTGCCTCTCCTGTTACCCTTTTAGAACCACAGCCTGGACACATGTGCCAGA | 4 |
| wt-30450-30570 | 4536 | CGCCTTGGCCTCTAAGGCCCTCGGGTCCCCCTGGACCCCGGCCTCAGCAACCCTGCTGCTCCCCTCCTGCCACCCCAGCCTCCCCCCCTCCCCGTCCCCCTTCGCTCCTGATCCTCCCCC | 8 |
| wt-30570-30690 | 4537 | GGTCCCCAGTAGGGCCGCCTGCCCCCCTGCACCCAGTACCTGCCCCTCTTGGCCACGCACCCCGGGCCAGGCCACCTTAGACCCGGCCAAGCCCCATCCCTGAAGACCCAGCGGCCATTC | 8 |
| wt-30690-30810 | 4538 | TCTCTGGTAACGAGCAGAGAAGAAGTAGAGGCCCGCGGCCATTGGGCCCAGATTGAGAGACCAGTCCAGGGGCCCGAGGTTGGAGCCAGCGGGCACCCGAGGTCCCAGCACCCGGTCCCT | 8 |
| wt-30810-30930 | 4539 | CCGGGGGGCAGAGACAGGCAGGGCCCCCCGGCAGCTGGCCCCGAGGAGGCGCCCGGAGTGGGGCCGGTCGGCTGGGCTGGCCGAGCCCGGGTCTGGGAGGTCTGGGGTGGCGAGCCTGCT | 12 |
| wt-30930-31050 | 4540 | GTCTCAGGAGGGGCCTGGCTCCGCCGGGTGGCCCTGGGGTAAGTCTGGGAGGCAGAGGGTCGGCCTAGGCCCGGGGAAGTGGAGGGGGATCGCCCGGGTCTCTGTTGGCAGAGTCCGGGC | 8 |
| wt-31050-31170 | 4541 | GATCCTCTGAGACCCTCCGGGCCCGGACGGTCGCCCTCAGCCCCCAGACAGACCCCAGGGTCTCCAGGCAGGGTCCGGCATCTTCAGGGGCAGCAGGCTCACCACCACAGGCCCCCCAG | 8 |
| wt-31170-31290 | 4542 | ACCCGGGTCTCGGCCAGCCGAGCCGACCGGCCCCGCGCCTGGCGCCTCCTCGGGGCCAGCCGCCGGGGTTGGTTCTGCCCCTCTCTCTGTCCTTCAGAGGAACCAGGGACCTCGGGCACC | 8 |
| wt-31290-31410 | 4543 | CCAGAGCCCCTCGGGCCCGCCTCCAGGCGCCCTCCTGGTCTCCGCTCCCCTCTGAGCCCCGTTAAACCCAAAGAATGTCTGAGGGGAGCCACCCTCGGGGCCCAGGCCCCAGAGTCCAGA | 8 |
| wt-31410-31530 | 4544 | GGTCAGGGGCACCTCAGGGTGCCTCCCCGGGTCCCAGGCCAGCCGGAGGGACCCCGGCAGCCCGGGCGGCCCCAGAGGCCGGTTCCTCGCCCCTTCCCCGGGCTTCAGAGCCCAGGATGT | 12 |
| wt-31530-31650 | 4545 | CCCCCAGAAGGGACCCTAGGCGTCCCCTCTCCTCCCCTCCAGGCCCGAGCCTCTCCCTCGCGGAGAGGGGCCTCTTTGGGCCCTCAAGTCCAGCCCCACCGAGACCCGAGTGGCCCGGAT | 8 |
| wt-31650-31770 | 4546 | CCCCCCACCGGCCCTTCTCTCTGTCCCCCTGCTCCTCTCCAACCTTCGCTCCACCCTAGACCCCAGCTTCTGGCCTCCCCGGGTCCACCAGGCCAGCCGGAGGGACCCCGGCAGCCCGGG | 8 |
| wt-31770-31890 | 4547 | CGAGTCGCCTTCCCTCTCCCCTGGCCTCTCCTTCCCGCCTCCCACCCGAGCCCCCTCAGCTTGCCTCCCCACCGGGTCCATCAGGCCGGCCGGAGGGACCCCGGCGGCCCGGTGTCAGTC | 8 |
| wt-31890-32010 | 4548 | CCCCCTGCAGCCGCCCAGTCTCTGCCTCCAGGCAAGGGCGCCAGCTTTTCTCCCCCCAGCCTGAGGCCCAGTCTCCTGTGCACTGTCTGTAAAGTCCAGCCTCCCACGCCCGTCCACGGC | 8 |
| wt-32010-32130 | 4549 | TCCCGGGCCCAGCCTCGTCCACCCCTCCCCACGGTGGACAGGCCCTCTGTCCACCCGGGCCATCCCCGCCCCCCTGTGTCCACCCCAGTCCCGTCCAGGGGGGACTTTATGTGACCCTTG | 8 |
| wt-32130-32250 | 4550 | GGCCTGGCTCCCCATAGACTCCCATGTAAGCCTGCCTCGAGTAGGTGCCTCCAGAGCCCCTTTTGCCCCCCTGGCGGCCCAGCCCGACCCCCGGGCGCCCCCAAACTTTGTCCAGATGTC | 8 |
| wt-32250-32370 | 4551 | CAGGGGTCCCCGAGGGTGAGGCCCAGCCCCCTCCCGCCCCTGTCCACTGCCCCGGTCCCCCCAGAAGCCCCCAAAAGTAGAGGCTCAGGCCATGCGCGCCCTGTCACCAGGCCTGCCAAA | 8 |
| wt-32370-32490 | 4552 | GAGCCAGATCTAAGGCCGGGAGAGGCAGCCCCAAAGCGGGTGCAGTAACAGGTAATCTCTGGTAGTGATTTGGACCCGAAATCTGACACTTTAGAGCTCTGGAGGACTTTAAAACTCTAA | 4 |
| wt-32490-32610 | 4553 | AAATCAAAACTTTAGAGGCGAATGGGCGCCATTTTGTCCCCACGCGCGCATAATGGCGGACCTAGGCCTAAAACCCCCAGGAAGCGGGTCTATGGTTGGCTGCGCTGCTGCTATCTTTAG | 4 |
| wt-32610-32730 | 4554 | AGGGGAAAAGAGGAATAAGCCCCCAGACAGGGGAGTGGGCTTGTTTGTGACTTCACCAAAGGTCAGGGCCCAAGGGGGTTCGCGTTGCTAGGCCACCTTCTCAGTCCAGCGCGTTTACGT | 4 |
| wt-32730-32850 | 4555 | AAGCCAGACAGCAGCCAATTGTCAGTTCTAGGGAGGGGGACCACTGCCCCTGGTATAAAGTGGTCCTGCAGCTATTTCTGGTCGCATCAGAGCGCCAGGAGTCCACACAAATGTAAGAGG | 4 |
| wt-32970-33090 | 4557 | ACTCCGCCATCCAAGCCTAGGGGAGACCGAAGTGAAGGCCCTGGACCAACCCGGCCCGGGCCCCCCGGTATCGGGCCAGAGGTAAGTGGACTTTAATTTTTTCTGCTAAGCCCAACACTC | 4 |
| wt-33090-33210 | 4558 | CACCACACCCAGGCACACACTACACACACCCACCCGTCTCAGGGTCCCCTCGGACAGCTCCTAAGAAGGCACCGGTCGCCCAGTCCTACCAGAGGGGGCCAAGAACCCAGACGAGTCCGT | 4 |
| wt-33210-33330 | 4559 | AGAAGGGTCCTCGTCCAGCAAGAAGAGGAGGTGGTAAGCGGTTCACCTTCAGGGGTAAGTAACCTGACCTCTCCAGGGCTCACATAAAGGGAGGCTTAGTATACATGCTTCTTGCTTTTC | 4 |
| wt-33330-33450 | 4560 | ACAGGAACCTGGGGGCTAGTCTGGGTGGGATTAGGCTGCCTCAAGTTGCATCAGCCAGGGCTTCATGCCCTCCTCAGTTCCCTAGTCCCCGGGCTTCAGGCCCCCTCCGTCCCCGTCCTC | 4 |
| wt-33570-33690 | 4562 | AACCCTGCTGCTCCCCTCCTGCCACCCCAGCCTCCCCCCCTCCCCGTCCCCCTTCGCTCCTGATCCTCCCCCGGTCCCCAGTAGGGCCGCCTGCCCCCCTGCACCCAGTACCTGCCCCTC | 8 |
| wt-33690-33810 | 4563 | TTGGCCACGCACCCCGGGCCAGGCCACCTTAGACCCGGCCAAGCCCCATCCCTGAAGACCCAGCGGCCATTCTCTCTGGTAACGAGCAGAGAAGAAGTAGAGGCCCGCGGCCATTGGGCC | 8 |
| wt-33810-33930 | 4564 | CAGATTGAGAGACCAGTCCAGGGGCCCGAGGTTGGAGCCAGCGGGCACCCGAGGTCCCAGCACCCGGTCCCTCCGGGGGGCAGAGACAGGCAGGGCCCCCCGGCAGCTGGCCCCGAGGAG | 8 |
| wt-33930-34050 | 4565 | GCGCCCGGAGTGGGGCCGGTCGGCTGGGCTGGCCGAGCCCGGGTCTGGGAGGTCTGGGGTGGCGAGCCTGCTGTCTCAGGAGGGGCCTGGCTCCGCCGGGTGGCCCTGGGGTAAGTCTGG | 12 |
| wt-34050-34170 | 4566 | GAGGCAGAGGGTCGGCCTAGGCCCGGGGAAGTGGAGGGGGATCGCCCGGGTCTCTGTTGGCAGAGTCCGGGCGATCCTCTGAGACCCTCCGGGCCCGGACGGTCGCCCTCAGCCCCCCAG | 8 |
| wt-34170-34290 | 4567 | ACAGACCCCAGGGTCTCCAGGCAGGGTCCGGCATCTTCAGGGGCAGCAGGCTCACCACCACAGGCCCCCCAGACCCGGGTCTCGGCCAGCCGAGCCGACCGGCCCCGCGCCTGGCGCCTC | 8 |
| wt-34290-34410 | 4568 | CTCGGGGCCAGCCGCCGGGGTTGGTTCTGCCCCTCTCTCTGTCCTTCAGAGGAACCAGGGACCTCGGGCACCCCAGAGCCCCTCGGGCCCGCCTCCAGGCGCCCTCCTGGTCTCCGCTCC | 8 |
| wt-34410-34530 | 4569 | CCTCTGAGCCCCGTTAAACCCAAAGAATGTCTGAGGGGAGCCACCCTCGGGGCCCAGGCCCCAGAGTCCAGAGGTCAGGGGCACCTCAGGGTGCCTCCCCGGGTCCCAGGCCAGCCGGAG | 8 |
| wt-34530-34650 | 4570 | GGACCCCGGCAGCCCGGGCGGCCCCAGAGGCCGGTTCCTCGCCCCTTCCCCGGGCTTCAGAGCCCAGGATGTCCCCCAGAAGGGACCCTAGGCGTCCCCTCTCCTCCCCTCCAGGCCCGA | 8 |
| wt-34650-34770 | 4571 | GCCTCTCCCTCGCGGAGAGGGGCCTCTTTGGGCCCTCAAGTCCAGCCCCACCGAGACCCGAGTGGCCCGGATCCCCCCACCGGCCCTTCTCTCTGTCCCCCTGCTCCTCTCCAACCTTCG | 8 |
| wt-34770-34890 | 4572 | CTCCACCCTAGACCCCAGCTTCTGGCCTCCCCGGGTCCACCAGGCCAGCCGGAGGGACCCCGGCAGCCCGGGCGAGTCGCCTTCCCTCTCCCCTGGCCTCTCCTTCCCGCCTCCCACCCG | 8 |
| wt-34890-35010 | 4573 | AGCCCCCTCAGCTTGCCTCCCCACCGGGTCCATCAGGCCGGCCGGAGGGACCCCGGCGGCCCGGTGTCAGTCCCCCCTGCAGCCGCCCAGTCTCTGCCTCCAGGCAAGGGCGCCAGCTTT | 8 |
| wt-35010-35130 | 4574 | TCTCCCCCCAGCCTGAGGCCCAGTCTCCTGTGCACTGTCTGTAAAGTCCAGCCTCCCACGCCCGTCCACGGCTCCCGGGCCCAGCCTCGTCCACCCCTCCCCACGGTGGACAGGCCCTCT | 8 |
| wt-35130-35250 | 4575 | GTCCACCCGGGCCATCCCCGCCCCCCTGTGTCCACCCCAGTCCCGTCCAGGGGGGACTTTATGTGACCCTTGGGCCTGGCTCCCCATAGACTCCCATGTAAGCCTGCCTCGAGTAGGTGC | 8 |
| wt-35250-35370 | 4576 | CTCCAGAGCCCCTTTTGCCCCCCTGGCGGCCCAGCCCGACCCCCGGGCGCCCCCAAACTTTGTCCAGATGTCCAGGGGTCCCCGAGGGTGAGGCCCAGCCCCCTCTCGCCCAAGCTGCTT | 8 |
| wt-35370-35490 | 4577 | TGATTCCTGGGATATTTTTGGGAATGGTGTTAACTTTCTCCCCTTGTATTTGCTATTCAATCAACCTGATTCCCCCTGCTCATACCTCCACTTACAACCAAGCCACTACGGCCACGTCCC | 4 |
| wt-35490-35610 | 4578 | CGGCCTCCCGCTCGGGTAAGTGCTTTTTCATTTTTAGCCCCAGCCCCTCCTCTATAAGTTCTAGGCAAACCTCCAATCACCAGCCACCTTCCAATGTAGTCTCTTAGAGAGTGGCTGCTA | 4 |
| wt-35610-35730 | 4579 | CGCATTAGAGACCACTTTGAGCCACCCACAGTAACCACCCAGCGCCAATCTGTCTACATAGAAGAAGAAGAGGATGAAGACTAAGTCACAGGCTTAGCCAGGTGATTTGTGAATTTCAGT | 4 |
| wt-35730-35850 | 4580 | TTATTTACTTTCTTCCAATCAAGCTTTCCCAGCCTCCGCTTGTTAGGTCCTAGTTATGGGTTTTCCATGGGGGACTTAGTATCCGTTCTATTAGATTAACGTGCAAGACGCTAAACTTAA | 4 |
| wt-35850-35970 | 4581 | CCAAGGTCAGCCAAGGGACGCGTGTTATCCCAGGCTGCCCACCCTGAGGATTTCCCCCCAAAATCCTCCTACCCTCTCTTTATGCCATGTGTGTTGTTGGCTTGTGTTAGTGCTATGTAA | 4 |
| wt-35970-36090 | 4582 | TGCGTTGCCGCCAGGTGGCAGCCTGTTTATAGATGTGCAGTACCCCTTAATGTTAGGTCTGCTTTAGGGCTGCCAGGTGGCGCAATCTAGGATTAATTCACCTGTATCCCTTTCCCTCCA | 4 |
| wt-36090-36210 | 4583 | CCCGCAGTAACCCAGCACTGGCGTGTGACGTGGTGTAAAGTTTTGCCTGAACCTGTGGTTGGGCAGGTACATGCCAACAACCTTCTAAGCACCCGCGCTTGTGTTTTGCTTTATCTGCCG | 4 |
| wt-36210-36330 | 4584 | CCATCATGCCTACATTCTATCTTGCGTTACATGGGGGACAAACATATCATCTAATTGTTGACACGGATAGTCTTGGAAACCCGTCACTCTCAGTAATTCCCTCGAATCCCTACCAGGAAC | 4 |
| wt-36570-36690 | 4585 | GATATGACGTCGGGCATGGACCTCTAGCATCTGCTATGCGAATGCTTTGGATGGCTAATTATATTGTAAGACAATCACGGGGTGACCGGGGCCTTATTTTGCCACAAGGCCCACAAACAG | 4 |

| | | | |
|---|---|---|---|
| wt-36690-36810 | 4586 | CCCCTCAGGCCAGGTTGGTCCAGCCACATGTCCCCCCTCTACGCCCGACAGCACCCACCATTTTGTCACCTCTGTCACAACCGAGGCTTACCCCTCCACAACCACTCATGATGCCACCAA | 4 |
| wt-36810-36930 | 4587 | GGCCTACCCCTCCTACCCCTCTGCCACCTGCAACACTAACGGTGCCACCAAGGCCTACCCGTCCTACCACTCTGCCACCCACACCACTACTCACGGTACTACAAAGGCCTACCGAACTTC | 4 |
| wt-36930-37050 | 4588 | AACCCACACCATCACCACCACGCATGCATCTCCCTGTCTTGCATGTGCCAGACCAATCAATGCACCCTCTTACTCATCAAAGCACCCCAAATGATCCAGATAGTCCAGAACCACGGTCCC | 4 |
| wt-37050-37170 | 4589 | CGACTGTATTTTATAACATTCCACCTATGCCATTACCCCCCTCACAATTGCCACCACCAGCAGCACCAGCACAGCCACCTCCAGGGGTCATCAACGACCAACAATTACATCATCTACCCT | 4 |
| wt-37290-37410 | 4590 | AGTCCAGGGACAAGCAACGCAAGCCCGGTGGACCTTGGAGACCAGAGCCAAACACCTCCAGTCCTAGCATGCCTGAACTAAGTCCAGTCCTCGGTCTTCATCAGGGACAAGGGGCTGGGG | 4 |
| wt-37410-37530 | 4591 | ACTCACCAACTCCTGGCCCATCCAATGCCGCCCCCGTTTGTAGAAATTCACACACGGCAACCCCTAACGTTTCACCAATACATGAACCGGAGTCCCATAATAGCCCAGAGGCTCCCATTC | 4 |
| wt-37530-37650 | 4592 | TCTTCCCCGATGATTGGTATCCTCCATCTATAGACCCCGCAGACTTAGACGAAAGTTGGGATTACATTTTTGAGACAACAGAATCTCCTAGCTCAGATGAAGATTATGTGGAGGGACCCA | 4 |
| wt-37650-37770 | 4593 | GTAAAAGACCTCGCCCCTCCATCCAGTAAAAACCCTTGCCCTCTCCAGCAACCAATGTATCCCAAATAAATGTTACTTCTTTTGCTCTTAACCATTGACACGCCTGTCATTCTATCAATT | 4 |
| wt-37770-37890 | 4594 | AAACAAGGGAAAAAGGTTTAGCTATTCCACCAACACGACCCCAAGGAAGGCTTGCCAAAATTGGTGCCTTGCTCTCAGCACTTTGCCAGCAACTTATAGCATGGTAGGCAGCTCAACTCG | 4 |
| wt-37890-38010 | 4595 | GCCCGTCTTACTGCCCAGCCTACTCTCCACTCCCAGTCCATGTTCGCACTCCTATGCATTTCCTGCCCTCCCACTTTTACCCCAGTCCCAACCCAAAACCACACACAACACATAGAATTG | 4 |
| wt-38010-38130 | 4596 | TTAGTTTAAACAGTTTATTGATAGGTGGCTGCTTTTAGCCTAATTGTGTATTGCTCTCGTTGCCAAAACCTGTTGTAAGGGCCGGCACCCGCAACATGGGGAAAACATAACCGCCGCCAT | 4 |
| wt-38130-38250 | 4597 | CCCATGGGGAGGGTAGAGGCGGTTGACATGTAGGTGAGTAGTGTAAGAAGCATGGCGAAGTAGACAGGTTACTTTTAGAGTGTAGTGTACAGGGCCGGGCGCAACAGTGCCACCAACCCG | 4 |
| wt-38250-38370 | 4598 | GGGTCTGAGCATTCCATGGGCAGCAGGGACACTGCACTACCGCCAGGTCCTGGGGCAGCCGGGGTTCCTGGCGCTCCGGGGGCAGCCGGGCGGCCGCCGGTGGGTCCGCTGGGCCGCTGC | 12 |
| wt-38370-38490 | 4599 | CCCGCTCCGGGTGGGGGGTGGCCCCGCTGGGCACCGCTGCGCCGCCGCCAGGTCCTGGGGCAGCCGGGGTTCCTGGCGCTCCGGGGGCAGCCGGGCGGCCGCCGGTGGGTCCGCTGGGCC | 12 |
| wt-38490-38610 | 4600 | GCTGCCCCGCTCCGGGTGGGGGGTGGCCCCGCTGGGCACCGCTGCGCCGCCGCCAGGTCCTGGGGCAGCCGGGGTTCCTGGCGCTCCGGGGGCAGCCGGGCGGCCGCCGGTGGGTCCGCT | 12 |
| wt-38610-38730 | 4601 | GGGCCGCTGCCCCGCTCCGGGTGGGGGGTGGCCCCGCTGGGCACCGCTGCGCCGCCGCCAGGTCCTGGGGCAGCCGGGGTTCCTGGCGCTCCGGGGGCAGCCGGGCGGCCGCCGGTGGGT | 12 |
| wt-38730-38850 | 4602 | CCGCTGGGCCGCTGCCCCGCTCCGGGTGGGGGGTGGCCCCGCTGGGCACCGCTGCGCCGCCGCCAGGTCCTGGGGCAGCCGGGGTTCCTGGCGCTCCGGGGGCAGCCGGGCGGCCGCCGG | 16 |
| wt-38850-38970 | 4603 | TGGGTCCGCTGGGCCGCTGCCCCGCTCCGGGTGGGGGGTGGCCCCGCTGGGCACCGCTGCGCCGCCGCCAGGTCCTGGGGCAGCCGGGGTTCCTGGCGCTCCGGGGGCAGCCGGGCGGCC | 12 |
| wt-38970-39090 | 4604 | GCCGGTGGGTCCGCTGGGCCGCTGCCCCGCTCCGGGTGGGGGGTGGCCCCGCTGGGCACCGCTGCGCCGCCGCCAGGTCCTGGGGCAGCCGGGGTTCCTGGCGCTCCGGGGGCAGCCGGG | 12 |
| wt-39090-39210 | 4605 | CGGCCGCCGGTGGGTCCGCTGGGCCGCTGCCCCGCTCCGGGTGGGGGGTGGCCCCGCTGGGCACCGCTGCGCCGCCGCCAGGTCCTGGGGCAGCCGGGGTTCCTGGCGCTCCGGGGGCAG | 12 |
| wt-39210-39330 | 4606 | CCGGGCGGCCGCCGGTGGGTCCGCTGGGCCGCTGCCCCGCTCCGGGTGGGGGGTGGCCCCGCTGGGCACCGCTGCGCCGCCGCCAGGTCCTGGGGCAGCCGGGGTTCCTGGCGCTCCGGG | 12 |
| wt-39330-39450 | 4607 | GGCAGCCGGGCGGCCGCCGGTGGGTCCGCTGGGCCGCTGCCCCGCTCCGGGTGGGGGGTGGCCCCGCTGGGCACCGCTGCGCCGCCGCCAGGTCCTGGGGCAGCCGGGGTTCCTGGCGCT | 12 |
| wt-39450-39570 | 4608 | CCGGGGGCAGCCGGGCGGCCGCCGGTGGGTCCGCTGGGCCGCTGCCCCGCTCCGGGTGGGGGGTGGCCCCGCTGGGCACCGCTGCGCCGCCGCCAGGTCCTGGGGCAGCCGGGGTTCCTG | 12 |
| wt-39570-39690 | 4609 | GCGCTCCGGGGGCAGCCGGGCGGCCGCCGGTGGGTCCGCTGGGCCGCTGCCCCGCTCCGGGTGGGGGGTGGCCCCGCTGGGCACCGCTGCGCCGCCGCCAGGTCCTGGGGCAGCCGGGGT | 16 |
| wt-39690-39810 | 4610 | TCCTGGCGCTCCGGGGGCAGCCGGGCGGCCGCCGGTGGGTCCGCTGGGCCGCTGCCCCGCTCCGGGTGGGGGGTGGCCCCGCTGGGCACCGCTGCGCCGCCGCCAGGTCCTGGGGCAGCC | 12 |
| wt-39810-39930 | 4611 | GGGGTTCCTGGCGCTCCACTGCACCTGGAATGCAGGGTGGGGGCGTGGTCCCCTGGACCCCAGCCCCGCCGATCCCTCCCCCAGGGCGTACCCGGCTTGCCTGGTTCTGGGGCTCCTCTG | 8 |
| wt-39930-40050 | 4612 | GGGGTCGCTGCATCCGCCGGTAGGGTTCGAATGGGCGTGGTCCGCTTGCTCTGCTGGCCCGGTACGCCTGGATTGCCGGCTGGGGGCTGGGGTCCCGGGACGCCCCCTCCCTGCTCCCAC | 8 |
| wt-40050-40170 | 4613 | CCGGTTCCCTCCCCCAGGGCGTGCCCCGCTTGCCTGGTCCTGGAGCTCATCCGGGGATGCTGCATCCGCTAGTCCGACCTGGGTGGGTGCGGTCCGCTGGCCCCACCCTGGGGGTAGCCG | 8 |
| wt-40170-40290 | 4614 | CCGGGTCTGCTGGTCCGGTGCACCTGGAAGGCAGGGGGGGGGGCAGTGAGGGAGGGGCGTGGTCCTGGGACCCCGCGCCGACTGGCAGGGGGTCCCCATGGCACAGGCCTAGGGGTCCAG | 8 |
| wt-40290-40410 | 4615 | GGGGCAGCCGCGGCCCAGCGCGCCCCGTTCACGGGGGAGGACCGCGGCCGAGCCACCAGGGGCCCGGCGGGGGTGGGGGGTGCGCTCCCAGGCCGGACCCTGGTGCCAGGCAGGGACCCC | 12 |
| wt-40410-40530 | 4616 | GCGCCACCCGCTTCATGGGGGGGGAGGCCGCCGCAAGGACGCCGGGCCGGCTGGGAGGTGTGCACCCCCCGAGCGTCTGGACGACGCTGGCGAGCCGGGCCGGCTCGCCTTCTTTTATCC | 8 |
| wt-40530-40650 | 4617 | TCTTTTTGGGGTCTCTGTGTAATACTTTAAGGTTTGCTCAGGAGTGGGGGCTTCTTATTGGTTAATTCAGGTGTGTCATTTTAGCCCGTTGGGTTTCATTAAGGTGTGTCACCAGGTGGG | 4 |
| wt-40650-40770 | 4618 | TGGTACCTGGAGGTTATTCTATTGGGATAACGAGAGGAGGAGGGGCTAGAGGTCCGCGAGATTTGGGGTAGGCGGAGCCTCAGGAGGGTCCCCTCCATAGGGTTGAACCAGGAGGGGGAG | 4 |
| wt-40770-40890 | 4619 | GATTGGGCTCCGCCCCGATATACCTAGTGGGTGGAGCCTAGAGGTAGGTATCCATAGGGTTCCATTATCCTGGAGGTATCCTAAGCTCCGCCCCTATATACCAGGTGGGTGGAGCTAGGT | 4 |
| wt-40890-41010 | 4620 | AGGATTCAGCTAGGTTCCTACTGGGGTACCCCCCTACCCTACCTTAAGGTGCGCCACCCTTCCTCCTTCCGTTTTAATGGTAGAATAACCTATAGGTTATTAACCTAGTGGTGGAATAGG | 4 |
| wt-41010-41130 | 4621 | GTATTGCAGCTGGGTATATACCTATAGGTATATAGAACCTAGAGGAAGGGAACCCTATAGTGTAATCCCTCCCCCCCCTACCCCCCCCTCCCTTACGGTTGCCTGAGCCCATCCCCCACC | 4 |
| wt-41130-41250 | 4622 | CCAGCACCCCGGGGTGACGTGGCACCCCGCGTGCCTTACTGACTTGTCACCTTTGCACATTTGGTCAGCTGACCGATGCTCGCCACTTCCTGGGTCATGACCTGGCCTGTGCCTTGTCCC | 4 |
| wt-41250-41370 | 4623 | GTGGACAATGTCCCTCCAGCGTGGTGGCTGCCTTTGGGATGCATCACTTTGAGCCACTAAGCCCCCGTTGCTCGCCTTGCCTGCCTCACCATGACACACTAAGCCCCTGCTAATCCATGA | 4 |
| wt-41370-41490 | 4624 | GCCCCGCCTTTAGGAAGCACCACGTCCCGGGGACGGAAGGGGACTTGGGGTGATTTTCTATGTGGGGGTGGAAATATGAGCAAGAATAAGGACGGCTCCTTATTAACCTGATCAGCCCCG | 4 |
| wt-41490-41610 | 4625 | GAGTTGCCTGTTTCATCACTAACCCCGGGCCTGAAGAGGTTGACAAGAAGGGTCAAGGTTTCGTCTGTGTGTTGAAGGGCAGGGGCTGTTGGGTGCATCTGGAACGGCTTACCTCGGGTA | 4 |
| wt-41610-41730 | 4626 | ACTGTTTGCCATTAAAAGGTTGGGGATTAGGTTTAGCCCCTTTAGCTGCCATTTCGAACCGGGGTGTGCAGATGCAGGTCTCCGGGTGGGCAGGCAGTACGAGATGTCACGTTGTGTTGT | 4 |
| wt-41730-41850 | 4627 | CTTTCCTCCCACCCCTGTCCTGGCTGTGGCAAATGCGACCCTCATAGAGTTGTGTTTCAGGTCTGTGTCCTGTTTTGCGGTGGGTTATTTCTTCCCTCAGTGTTTGCCAGCTTATTTCCC | 4 |
| wt-41850-41970 | 4628 | CAGTTTTCACGTACTGGGGCCTGTGGACACCTGAGGGAGCGGCCGTTGGTGGGTATGTGTTGGAATTGCTCCCACCCTCAATTTTCGCTTGCCTTCTTCCCTTGTTAACCTGATAGCATA | 4 |
| wt-41970-42090 | 4629 | GCCTCTAGGTTTCCTTGTAGGTCTGTTTGGGTTTGTTGGTTCACGTGGTGCTAACTTGAATTTTTTGGTTTTCTAGTTCCCTCTTAATTACATTTGTGCCAGATCTTGTAGAGCAAGATG | 4 |
| wt-42090-42210 | 4630 | GCCTATTCAACAAGGGAGATACTGTTAGCCCTGTGTATACGGGACAGTCGTGTGCATGGAAATGGTACCCTGCATCCTGTGTTGGAGCTAGCAGCAAGAGAAACACCTCTCCGCCTTTCG | 4 |
| wt-42210-42330 | 4631 | CCAGAGGACACTGTAGTTCTGCGTTATCATGTGTTGCTTGAGGAGATAATTGAACGAAATTCAGAGACATTTACAGAAACTTGGAACAGATTTATAACACACACCGAACATGTGGATCTG | 4 |
| wt-42330-42450 | 4632 | GATTTTAACTCAGTATTTTTAGAGATATTTCACCGTGGAGACCCAAGCCTTGGGCGCGCGTTGGCCTGGATGGCCTGGTGCATGCATGCCTGCAGGACATTGTGTTGTAACCAGTCTACT | 4 |
| wt-42450-42570 | 4633 | CCTTACTATGTTGTGGACCTGTCAGTTCGTGGGATGTTAGAAGCCAGCGAAGGCCTGGATGGTTGGATTCATCAACAGGGCGGCTGGTCTACATTAATTGAAGACAACATTCCTGGATCC | 4 |
| wt-42570-42690 | 4634 | AGAAGGTTTAGCTGGACTTTGTTTCTTGCTGGACTGACTTTGAGTCTGTTAGTTATATGTAGTTATTTATTTATCTCCAGAGGAAGACACTAATCTATACATTTTCTCAGCACTTTATAT | 4 |
| wt-42690-42810 | 4635 | GAATCAGGGTCATTGGGCCTGCGGGGAACTGAGCCAGTAGGATATTAGGCAAGGGTGACACAGTGCCCATGCATTATAATTTAACCAAACAGTGGTCGTGAGTTTTAGGCCGGCCATGGG | 4 |
| wt-42810-42930 | 4636 | GGCTTACAAGAATAACATGCCAATGACCCGGCCCCCACTTTTAAATTCTGTTGCAGCAGATAGCTGATACCCAATGTTATCTTTTGCGGCAGAAATTGAAAGTGCTGGCCATATCTACAA | 4 |

| | | | |
|---|---|---|---|
| wt-42930-43050 | 4637 | TTGGGTGTCCTAGGTGGGATATACGCCTGTGGTGTTCTAACGGGAAGTGTGTAAGCACACACGTAATTTGCAAGCGGTGCTTCACGCTCTTCGTTAAAATAACACAAGGACAAGATACTA | 4 |
| wt-43050-43170 | 4638 | AAGAAATAACTGAGGTGAGTGTGGGAAGATGGGAATACTATGTGTTATGTTAACGGGTGAGAGCCTATACTGCAGCCCAGACTCGGGGGGAGGAGGAAATGGTAAGAGTTATACTCTACT | 4 |
| wt-43170-43290 | 4639 | TATCTTTTTTGACACTACATTTAACTGTTATGTAACAATGTTTGCTTATTTTCATGTTCAATAAACGCTATGTTAATGATGAAGAACCTGTGTTCTTTGGAAGTGGGCCCAATGGGGTAG | 4 |
| wt-43290-43410 | 4640 | TAGGTTTTGGGAGGGTGCCGTGCTAGATATTTCAACTGCCACAGACCCCATTTTGTCCCACCTGTTACCACATTCTAGGTCCTGCATCCAGTGGGCCAGGTGTCTCACCATGGCTCTTTC | 4 |
| wt-43410-43530 | 4641 | TAGGTGGATACCACAGTCCAGGCCCCCAAGGCTACCGTGCTAATTACCTCCTCATGTCCACCCCCACCCTGTGTTACTGTCGCCTGATTATCCTGGCTTAGCAGCCTCCAAGTTTTACAA | 4 |
| wt-43530-43650 | 4642 | GACGTCCCATTGCCCTGCCCTTGGTCCAAGTCTCGCCGGTTTTCAGCAGCCTGTTGTAGCCTGCCCCCAAGTTTCGCAGGTTTCCCCCATGCTTCCACCCGTTAACCCAATAGCATGACA | 4 |
| wt-43650-43770 | 4643 | GCCAATCCAACACGAGGCAAGTTTTAAGAGTTAAAAGCAACTACTGTTTATTTTCCAAAATGAGCTGGGTATAGTTGATGATCTGTAGGCGCAGCTCATCCCCACATTCCAGGTCCTTGA | 4 |
| wt-43770-43890 | 4644 | TGGCCTCGTAGATGGCATCTTCGTCGACATTGACAGCCTTCTCATATACCGTGTCTCTGGGGCTGACCTTTATACAGAAGGCGTCCCCTACTAGGTCCACGGCCAGCTCGTAGGTGGGGC | 4 |
| wt-43890-44010 | 4645 | CTATGTTTTCACATAACAGTTTCAAGCAGGTCTCTGGGATGTGAAGGGAGGTGCCCTGGAGCAGGAGATGCATGATTAGGCGCCCTTTTCCATTTGTGCTGAAGATGGGGCAGATGGTGC | 4 |
| wt-44010-44130 | 4646 | CACAAAAGTGTCCGGTGACCAGGTAAGCGTAGAGAAGGCTGGGTTGGGAAAGTCCAGCCTTTACTGCACTGGGAGAGCTGCTGAGCAGAGACACATAGAAGGTCTTGTTGGGTATTATCT | 4 |
| wt-44130-44250 | 4647 | TGTGGACATTGTTGAAGAAGGAGAGCTGGGTGGAGCTAAACTCCTGAGGCACATGAACCTGGGACCTATTGATGCAGATCTCGCAGTGAGACCCCAGAGTCAGGCTGTGGCCGAAGGGAG | 4 |
| wt-44250-44370 | 4648 | ACAGGCGAAGGCAGCGCCCGGGGGAGAGAGTGCACAGTGACAGTGGGAGAAACACGGCCTCTGAGACATGTATGGGGGTGTTCATCTCACGCAGAAAATCTTTGCCCAGCTCAAAGTTGG | 4 |
| wt-44370-44490 | 4649 | CAGAGATTCCCCTGAAGAAGTCCCGTAGTGAAAAATGGGATCTGTCTACACCATGTCTGGTGTGCCGGGAACATATTGATCGGGCCACACTGCCAACCCTTTCCATTCTTCCCAGCTCTG | 4 |
| wt-44490-44610 | 4650 | AGCGAGATTTTCCACACCTGGACACCGACTTCACGCTATGCGCCGAGGCCTTTGAGGCCGTGTAGTTTCTGTGGTGCGGATGCATTAGGCGGCGCAATGCGGGATCTGCCGGTCGCTGTT | 4 |
| wt-44610-44730 | 4651 | GGCGTGCATTCACGGCATCTGGGGTGACCGGGGCCATCGGGTTTACTTTTCACACGTAGACCTGGGAAGTTTGATAGGACTGTACCAGGTCAAGGCCGTGGATGCGCAGGACCACGTCCA | 4 |
| wt-44730-44850 | 4652 | GTTCCTTAGTGACATCCACGAGGATTGTTTTGCCCACTCTGGCCACTTGTGTGGATTTAAATATGTACACAAGCGTAATTAACGAGTCACAGACCCCCTGTTCCAGATTCTGACCGGCTG | 4 |
| wt-44850-44970 | 4653 | CAAGCGCTGCCTTAAAGGCCTGGAAGCTGGGTGGGTAAATCTGACCAAACAGCACGCTCGGATTCGTGATGCTGTGGTTGATGGCACACAGGGGGTCGCAGAACAGGTGCTTGTGGAAGT | 4 |
| wt-44970-45090 | 4654 | CTTGCGGGGTGCACATCTGCAGCCAGGCCCTTAGCCTGGGGCATGGCACATCCAGCAGCGTGTTTTGGGTCTTGATGAGGAACACGATCCTGTCTAGGATTTTGATGTTGTTGCCGAACG | 4 |
| wt-45090-45210 | 4655 | AGTCAAGAATCAGGCTCTTGAAGCGGTCAAGGGTGTCCTTGGCGTCCGGGTGGGCCCCGAGGCTCTCGCAGAGTGGGCAGATGGTCCGTGAGGCATTCTTGTGCCTTAGTCCAAACATGG | 4 |
| wt-45210-45330 | 4656 | GGGCCAGGAGGCAGGGGCCCTGCGAATGGTCGCCAGCCTCCGGTCTGGTGATGGCCAGGGCCAACTCCGCCAGCTCATCGCCGCTGTATTCCGCGTTTAAACCGATAGCATGGTGGCCTG | 8 |
| wt-45330-45450 | 4657 | GCCCCCCGAGCAGGTCCGTCCCCTGCCACGTACCTAATAGTAGTCCACAGTAGTCGGCCTTGGTTGTAATTTCAGGAGAGAGTCCTCCCTTTTCGGCCCTGAGAAATGGATGCTGAACTC | 4 |
| wt-45450-45570 | 4658 | GGTTTCTGGTAGGCAGGTGGCAGCACAGGGCGGTGTACAGGCCCCTGCCGACGTCCCCTGGGACATCCTGGGAATCTTTGCAGGTTCTGGGTCCAGGGAGGGTAAGAAAAGTGGGGGTGG | 4 |
| wt-45570-45690 | 4659 | TTCTGGGCCACATGGACTTGAAGCAGAAGTTGGCCGGGGACTGGCCGGTGAGGATGGATTTCAGAAACTCCAATTTGTAGTAGCCGAGGTTGGCATTTCTAATCATGTCAGAAGAGGACA | 4 |
| wt-45690-45810 | 4660 | CAGGGAGGAAGCACCGGCAAATGTAAAAGTGAAGCTGGATGTCAATGGCAAGAATCCTGGAGGGCATGAAGAGGGAATCCAACCCCCCGGCCATGGGGAAGTATTTTATCAGGATGTGTA | 4 |
| wt-45810-45930 | 4661 | AAAAGTCCATGCCTGTGATGAGGCTAGAGATCCAGGCTCGTGGGGCATTTAGACAGTAGTAGCAGAGCAGGGCATAGTCCTCAAAGAAGGCCACGGGGGCATCTGAGTGATTGACCAGGG | 4 |
| wt-45930-46050 | 4662 | TGTCGAGCAGATCACAAACTCGGCAGGTGCTGGCTGGAGAGAGGGACTCGTAGGTGTGGACGAGTGGTGGGTAGGCTATGCCTTCTTCCGCGTTGGCTGGAAGATAGGAGTGGGCCATCA | 4 |
| wt-46050-46170 | 4663 | AAAGGCCGACTGCCTCGAACTGGCTTTTCAGATTGTCCACGGTCCAGGGCACAAAGTCCTCCATCTTTGGAGTTCTGCCCGCGATCTGTGCCACCTCTGTTACGCCACTCCTCGTGAGGG | 4 |
| wt-46170-46290 | 4664 | GGCAGCTGGACAGTCTTTTTCCGGTCAGGGGGTTTGGCTCGTTTGCGCTCGTGACTTTGTGAGCCATGACACATCTGGGTGGCAAGGTGAGGTCTTCTGGGTTTTTAATACCGGGGTCGG | 4 |
| wt-46290-46410 | 4665 | CACCAGTTTCTGGGACACCGCCACAAGGACAAGGTGGGCTAGCAAGTTCTCGAGTCTACGAAGACTCCGGGGGCAGTCTTTTGAGTTTCTCGCCTATGATCCACCCCAATCTCGCCCCCC | 4 |
| wt-46410-46530 | 4666 | TAATTGCGCCATCTGCCTACGCGAGGCTGAACCTCCTGAATCACTGCATCTTTCTTGAGGCGTTTAAAGAAGAGAATAGTGGCCAGGGCCTCGGTGGGGTCCAGCGTGAGGTCTTATTTT | 4 |
| wt-46530-46650 | 4667 | TGAAAAGGGATATTATAAAACAGGTCATTGCTCGGATTGTGGCAGCCGATAGCACCCTAGATCTAGTGAATCATGGCGAGCCCGGAAGAGAGGCTCCTAGACGAGCTCAATAACGTAATT | 4 |
| wt-46650-46770 | 4668 | GTGTCATTTCTGTGTGACTCTGGGTCTCTGGAAGTGGAGAGATGCTCCGGGGCGCATGTGTTCTCCAGGGGCAGCTCCCAACCCCTCTGCACCGTGAAGCTGCGCCACGGACAGATTTAC | 4 |
| wt-46770-46890 | 4669 | CACCTGGAGTTTGTCTACAAGTTCCTGGCCTTTAAGCTGAAGAACTGCAACTACCCCTCCTCGCCCGTGTTTGTGATATCCAACAACGGCCTGGCCACCACCCTGAGGTGCTTTTTGCAC | 4 |
| wt-46890-47010 | 4670 | GAGCCGTCGGGTCTCAGATCGGGCCAGAGCGGCCCTTGCCTGGGTCTCTCAACGGATGTTGACCTACCAAAGAACTCCATCATTATGCTGGGCCAGGATGACTTCATTAAGTTCAAAAGC | 4 |
| wt-47010-47130 | 4671 | CCCCTGGTCTTCCCTGCTGAGCTTGATCTCCTGAAATCTATGGTGGTCTGCCGGGCCTACATCACGGAACACCGGACGACGATGCAGTTTCTGGTGTTTCAGGCCGCCAACGCCCAGAAG | 4 |
| wt-47130-47250 | 4672 | GCCTCGCGGGTCATGGATATGATTAGTGATATGTCTCAGCAACTGTCTCGGTCTGGTCAAGTCGAGGATACGGGCGCCAGAGTCACAGGTGGAGGAGGTCCCAGGCCTGGCGTCACGCAC | 4 |
| wt-47250-47370 | 4673 | TCGGGGTGTCTTGGGGACTCACACGTTAGGGGGCGCGGTGGTTGGGACTTGGATAACTTTTCAGAAGCTGAGACCGAAGACGAGGCGAGTTACGCTCCTTGGAGGGACAAAGACTCGTGG | 4 |
| wt-47370-47490 | 4674 | TCGGAATCCGAGGCGGCGCCGTGGAAGAAGGAACTCGTGAGGCACCCCATCCGCAGGCACCGGACACGCGAGACTCGCCGTATGCGCGGGAGCCATTCACGGGTGGAACACGTGCCCCCC | 8 |
| wt-47610-47730 | 4676 | TGACGTTGCAGGCCCTTGGGGAGCGGGGGTTCTCCAGGCTCCTGGATCTGGGGCTGGCCTGCCTGGATCTGAGCTATGTGGAAATGAGGGAATTTGTGGTTTGGGGCAGGCCCCCAGCTT | 4 |
| wt-47730-47850 | 4677 | CTGAGGCGGCTGTGGCCTCTACGCCAGGCTCGCTTTTCCGAAGCCACTCGTCCGCCTACTGGTTGTCGGAGGTGGAGAGGCCCGGGGGCCTTGTCCGCTGGGCCAGGTCACAGACCAGCC | 8 |
| wt-47850-47970 | 4678 | CCTCATCCCTGACCCTCGCGCCCCATCTTGGCCCGTCCCTCTTGTCCCTTTCAGTGGTCACCGGTGGTGGGTGTGGAGCCGTGGCCTTTTGCAACGCCTTTTTCCTAGCTTATTTTTTGG | 4 |
| wt-47970-48090 | 4679 | TTGTGCGGTCTGTTTTCCCCGCGTTTTCCGATAGAATAGCTGCCTGGATCTGCGACCGGTCCCCTTTCTGCGAAAACACCCGGGCCGTGGCCAGGGGTTACCGAGGCCTCGTGAAGAGGT | 4 |
| wt-48090-48210 | 4680 | TCTTGGCATTCGTGTTTGAGCGTAGTAGCTATGACCCCCCCTTGTTGAGGCAAAACTCTAGGCCTGTGGAGCGCTGCTTTGCCATCAAGAATTATGTCCCGGGCCTGGACTCACAAAGCT | 4 |
| wt-48210-48330 | 4681 | GTGTGACGGTCCCGAGCTTCTCCCGCTGGGCCCAGTCTCACGCCAGCGAGCTCGATCCCCGGGAGATTCGCGACAGAGTTACACCAGCGACTGCACCTTCGTTCGTGGCTGATCATGCCT | 4 |
| wt-48330-48450 | 4682 | CGGCTCTATTGGCCTCCCTCCAGAAGAAGGCCTCCGACACCCCCTGTGGGAATCCCATTCAGTGGATGTGGTACCGCCTGTTGGTAAACTCGTGCCTGAGGAGTGCCCACTGTCTTCTGC | 4 |
| wt-48450-48570 | 4683 | CTATACCTGCCGTCTCTGAGGGGGGGAGAAAGACGGGCGGGGGCGTAGGGGAGGAGCTCGTGGGGGCCGGGGGGCCCTGCCTGAGCCGGGATGTTTTCGTGGCGATCGTAAGCCGCAATG | 8 |
| wt-48570-48690 | 4684 | TTCTCTCGTGTCTGCTGAACGTGCCTGCCGCGGGTCCCCGGGCCTACAAGTGTTTCAGATCCCACGCCTCCAGACCGGTGTCTGGCCCGGATTACCCTCCCTTGGCCGTGTTTTGCATGG | 4 |
| wt-48690-48810 | 4685 | ACTGCGGTTACTGCTTGAACTTTGGAAAGCAGACAGGTGTAGGAGGCAGGCTCAATTCCTTTAGACCCACTCTCCAGTTTTATCCCCGTGACCAGAAGGAGAAGCATGTGCTGACCTGCC | 4 |
| wt-48810-48930 | 4686 | ATGCCAGCGGCCGTGTGTACTGCTCCAACTGCGGCTCTGCGGCGGTGGGCTGCCAGAGGCTGGCTGAGCCACCGAGCGCCCGCTCGGGCTGGCGGCCCCGAATCCGGGCAGTGCTGCCGC | 8 |
| wt-48930-49050 | 4687 | ACAACGCGGCCTACGAGCTCGACCGTGGCTCCCGCCTCTTGGATGCCATCATCCCCTGCTTGGGACCCGACCGCACTTGCATGCGGCCGGTGGTCCTGCGGGGGGTGACGGTCAGGCAGC | 8 |
| wt-49050-49170 | 4688 | TCCTGTATTTAACTTTGCGGACAGAGGCCAGAGCCGTTTGCTCCATCTGTCAGCAACGCCAAGCTCCAGAGGACGCCCGCGACGAGCCTCACCTGTTCTCCTCCTGTTTAGAGGTAGAAT | 4 |

| | | | |
|---|---|---|---|
| wt-49170-49290 | 4689 | TGCCACCTGGTGAGCGGTGTGCGGGCTGCCGTCTCTATCAGACGCGTTATGGCACGCCGGCTGCCCAAGCCCACCCTCCAGGGGAGGCTGGAGGCGGATTTTCCAGACAGTCCCCTGCTT | 8 |
| wt-49290-49410 | 4690 | CCTAAATTTCAAGAGCTGAACCAGAATAATCTCCCCAATGATGTTTTTCGGGAGGCTCAAAGAAGTTACCTGGTATTTCTGACATCCCAGTTCTGCTACGAAGAGTACGTGCAGAGGACT | 4 |
| wt-49410-49530 | 4691 | TTTGGGGTGCCTCGGCGCCAACGCGCCATAGACAAGAGGCAGAGAGCCAGTGTGGCTGGGGCTGGTGCTCATGCACACCTTGGCGGGTCATCCGCCACCCCCGTCCAGCAGGCTCAGGCC | 8 |
| wt-49530-49650 | 4692 | GCCGCATCCGCTGGGACCGGGGCCTTGGCATCATCAGCGCCGTCCACGGCCGTAGCCCAGTCCGCGACCCCCTCTGTTTCTTCATCTATTAGCAGCCTCCGGGCCGCGACTTCGGGGGCG | 8 |
| wt-49650-49770 | 4693 | ACTGCCGCCGCCTCCGCCGCCGCAGCCGTCGATACCGGGTCAGGTGGCGGGGGACAACCCCACGACACCGCCCCACGCGGGGCACGTAAGAAACAGTAGAGGGCACGAAACATGGTGTAT | 8 |
| wt-49770-49890 | 4694 | GCACTTTATTAATAAACAATTACAGATACAAAAACTTGAGTCTCTCGAGGTCTGCGATGAGGCGGTGGGTGGAACGCTCCAGCTTGCGGCGAAGCTGGCTCACGAAGCGAGACAGTACTC | 4 |
| wt-50010-50130 | 4696 | CCTCCCTGATAATGTCTTCAATAGACAGAAATTGGGTGACCACTGAGGGAGTGTTCCACAGTAATGTTGTCTGGTCGCTAGATGGCGCGGGTGAGGCCACGCTTTGCGAAAACGAAAGTG | 4 |
| wt-50130-50250 | 4697 | CTTGAAAAGGCGCGGGATAGCGTGCGCTACCGGATGGCGGGTAATACATGCTATCCTTACATTTTGGCATTTTGGGCAGCTGGGAGGCGGCGGATGGGGGTGCTTCTTTTCGCACGGTGT | 4 |
| wt-50370-50490 | 4699 | CAGGGAGAGACACGCGCGGCGGAGATGCAGCCGACGGCGGGGCCGCGGTGGGCTGCCCCCGAGGACGGGCGCCGGCCGCCAGCGCCCCCGTGGCCTTTGGCACGGGCCTGGCACCCACCG | 12 |
| wt-50610-50730 | 4701 | GGGCGGGTGCCTGGGGGATGGGAAAGCCGGACGGCGCTTCTCCCGGGTCGAACGCTGGAGTAGCGGAGGCTGCTGCGCCGGCGGCCACCACGGGCGCACGGGGTCGCAGCCCGACGGCCG | 12 |
| wt-50730-50850 | 4702 | TGGGGAGGCGGGTGGCGGAGGGCCGAATCTCCGCGGCTTCTTCCCGGCCCCCCTGCTGTTTCTTCTCCCGTTGCATGATAGAATGGCCATAGGGTGGGTCCTGAGAGGAGGCTTGTGTGT | 4 |
| wt-50850-50970 | 4703 | CCTGGGGCTGGAGCCCAAAAGTCGTTAAAGATGCCGCTGATGGTGTGGGAGCTATGCCTCCCGTCGACTGGCCGGGCTTGTAGGGGGCTGAGGGTGGATAACTGGGCTTCTGTGAAGGCA | 4 |
| wt-50970-51090 | 4704 | CCAACCCTGGAATCTGGATGGTATGTTTCTTCTGTGACCCCGAGGCAGTCGATGGTGTAGAGTGTGGAGACAATGTGTAGACGATGGGCCCTTGTTCAGAAGCCCAGGGACTTGAGGGGG | 4 |
| wt-51090-51210 | 4705 | GCTGTTGTGGTGCTGGTTGGGGAAGGAGCTCCAGGGAATCTTTGGGCCATGGCCTTGGGGAGCTTCCCGGAGACCGGTCTGGGCTCTCGGAAGCCCTCGTTTCGGCCCCGAAATAGGGCC | 4 |
| wt-51210-51330 | 4706 | TTGCCATCAATCGGGGGCCTGGGAGAGTGATGGGGGCGGCCAATCCCGGGGTAACTGTCACGTCCCGGGGGGAGGAGGTAGGAGACAGCCAGTCCCTGGGCCTGCCAGGGGCCACCTTCT | 8 |
| wt-51330-51450 | 4707 | CTAAGAGGGGGCTCTGTGGGCTGGGAGGGCCAGAGGCCTCAGATTCAGCAGTAGTGCTCCCCTTTTCCCCCTGGTCCGTCTCCCCTCCTCCCAACTGCTGGAGCCGGTCGGAGGAGGCCG | 8 |
| wt-51450-51570 | 4708 | GGGTGTTATCTGCTGACTGAAACCCGTCCCCGCTGACCAGTCCGTGCCCCACCCTTGGGGGGAAACCGGAGAACAGCTCCTGGACGTTGCGTGGATTCGGGGGAAGCTGGTATCCAACCG | 4 |
| wt-51570-51690 | 4709 | GCAGTGGAGGATCTTCGTGCTCGTAGAAGGAGGGGTTGAGTACATCGGTCGGCCATCGTGAGGCCCCGGCCGCGTTAAAGTAGAACTGCACGTCCGGCAGATTGTGCCGATAGGTGAAAC | 4 |
| wt-51690-51810 | 4710 | ACTTCCAGATGATGTTTTTTCTGTTGGCCAGGATGGCCACGGTGGGGGGCCTGGCCTCCTTAGGTTTGGCGGCCCTGGCCTCGGTGAGAAGCTCGCGTAGCCACACGGCCTGGCGTGCAA | 4 |
| wt-51810-51930 | 4711 | AGATGGACATCTCTGGCTCGAAAGACTCGGAGTAGCCGTCCAGGTCCTGCAGAAAATTCAGCGAGATGGTCTCCACCAGGGACCGGAAGGGCTCAGAGTGCCCGTCGCAGTAGAGGAGGG | 4 |
| wt-51930-52050 | 4712 | GAGCAACGACCCTGACCTGTCCCAGGGTCTTCAGGTTAAACAGATATTGAGAGGAGACAAAGAGAGTTAGGGGCCGACCGAGGAAGGCCGCCGCCACGGCCGCCTCAAAAACGGAGACGG | 4 |
| wt-52050-52170 | 4713 | GGATGGTGTCACCGGAGCCCCTCTTAGGACCGGTAATGGGAGTGCCATAAGGCATAAGATTTCTCAGGGCCCGGCCGGTAACGGTGCCGTAGGAAGACGGGGTTTCGCGGGGGACCTCGA | 4 |
| wt-52170-52290 | 4714 | GTCCCTCCGCCCTGGGGAGCTCTTCTCCGCGTGTATAGGCCTGCTTCACAAAGTCGCGCAGGTAGTCCTGAAATGCGACCGGGCCCTCCAGCGGGCGCAATGAGTGCCAGAGCTGCTGAA | 4 |
| wt-52290-52410 | 4715 | GGGCCTCGGGGGCGAAGCACCGGCGTGCGAGGAGCAGCATGCAGGCTCGGGCGCGGGCCGTACTTTGGTTGTGGACCAGGCCCAAGAACTCGGGGTGCGGCCAGAGGGCGGCTCGGGTAT | 8 |
| wt-52410-52530 | 4716 | CCATCTCCTCCCAGGCGTCCTGGAAGAAGATGAAGCCGGTGGGTGGACCGGCGATGCGGTGGCGGGTGAGGCGGCGCGCGTCTTCCCCGTCGTTGCTGCCGCGGGTGGTTGAGGGCATGC | 8 |
| wt-52530-52650 | 4717 | CCCCCCTCCCGGAGGCTGGACTCCTGACCAGCCTGTAGGTGAGGACCGAGTCCGACAGGAGGTCTCCCAAACCCCCATCTCTCGCTAGAGCCGAGACCAGGCCGAGTCCTGCGTAGAACG | 8 |
| wt-52650-52770 | 4718 | ATGGGGCGCCCAGGAAGGCGGCAGCGTAGGCCGGATGTGTGCCGACCAGCAGCGCCATCATCTCCCGTTGTTCCAATAGAATAACTTCCCGGTCTGTGGCCGGGGCTGGATAAGGGGGGT | 4 |
| wt-52770-52890 | 4719 | GATTCCTAGAGGCGATGAGACTGGCGTGCGCTAAAAGTGTCATGGCCACAATGGGGTTGTCTGCCAGGTCTTCCATCAGGGCTTTGGGCGCAGAGACGTATTCCCGAAGCAGCTCCCCGG | 4 |
| wt-52890-53010 | 4720 | CGTTGGACTCCACGTCGGGCCAGGTGTCCCAGTAGGAGTCGGCGGCGGCGGCGCTGAGGCGGGCGGAAGCTACACTGGCCAGGGTTCTTCTCCTCCTCTCTTGGTCATCCTGCGGGGGAC | 8 |
| wt-53010-53130 | 4721 | CAATAGCTTGGGGGCGTCCGGCTGGGGTCAGGGAAAAGGCCTCTGGGTTCTCCAGCACGGTGGGCATGACATATTCCAGAAAGTTGTGGTAGACGGGGATGTAGTTGAGCGGCTCCTGGG | 4 |
| wt-53130-53250 | 4722 | TGTCTGCGGAGACGTAGGCCGGGTTAAGGGGGTCGCAGGGAGACTCTGTTTCCAGCCAGAGGGTGCCGGCGTATTTCGCCGGCCCTGCCGCCGCCAGAAATTGTGCCCGCCGGGTCGGGG | 8 |
| wt-53250-53370 | 4723 | CTCCATTGCCCCATCCAGTTGGTGGTGCCGAAATCGTGATGAGGAGGGGCAGGTTGTTGGTCAAGGGATGCTTAACGAAAACGGTAGGCTGGGCGGTCTCGTAAAAAGCCAGGAAACTCT | 4 |
| wt-53370-53490 | 4724 | GCTTGGCCGAGGCATAGCGCAGCAGCTTGTCCTTGAGGAGGGCATACTGGGAGCCAGCCGAGGCCCCAAGCGCCAGGCCCCTGGCAGCCTCCACCACGATCTTGAGCTGGCGCGGGTCGG | 8 |
| wt-53610-53730 | 4726 | CCTTGCCCCCCGCGACCCGCTTGGGGTCCAACGTGGCCAGAGCCAACCTCAGCTCCTCCATGCCATCCATGGAGTTCTGGGCCATGCCCTCGACTTCCAGGAGCCGTGTTAGCTCATGAA | 4 |
| wt-53730-53850 | 4727 | TTTCACCGTCAGCCGCAGCGGCTAGGTTCAGCCAGGCACCCGCGCCCCCAGCTAAGGCCAGGGCTCCTTCGGAAAGACCCCGCACGGCCTCGCAGATGCCCCGGATCCACTTGGCGGCTG | 8 |
| wt-53850-53970 | 4728 | CCAGGGATTTCCGGTAGGGCATGAGCCGTTCCCGGCCGCTGCCCGGGCCAGGGCGGCCTCGAGGGGAGCCTGGACAGGGGCTTTGGGCGGGGAGGGAAGCAGGCTCCGGAGTTCATCGT | 8 |
| wt-53970-54090 | 4729 | CGGGGGCTTCGTCGCGTGACCTGGAGAGGACGGCCTCCAGAGCCGTGTGAAAGCCCCGCCGAGTGCTTGCCGCCATCTCGTGGGCCTTCGCCATCAGGGTCTGGCTCTCCCGGACCTGCT | 8 |
| wt-54090-54210 | 4730 | CTTCCAGCGCCCGGACCTCGGCCGCCTCGGCCTCGGTCAGCAGCTCCGAGAAGAAGTCCCCCGTGGCCTGGAGGAGATCGTCCCGCTCTCGCCTTGTCAGCAGCTGGGGCTTCTTAGGCC | 8 |
| wt-54210-54330 | 4731 | AGAGCGCCGAGTCCGAGGCCAGCCTGGGCGGGGCGGTTGCCTGGGGGATAGTTGGAGGAGGAGGCAGGTTAGCCTGGCCTGGGTCATTAGTGGCTTCGGGTAGCGTCCGATCCACGTACT | 8 |
| wt-54450-54570 | 4733 | TGTCCGGGGACTGCAGGGCCTGGCGCGTCTGCGCGAGGGCCGAACGGATGGCATCGGCCGCCGTCCTGGCGCGAAAGAGGGCCCCGGCCGCTTCCTCCGCTCCTCGCCCTCCTCCTCCTT | 8 |
| wt-54570-54690 | 4734 | CTTTGGCGGTAGCCGCGGGGGTGGCGGGCCAAGCGTCCAGTCTGGCCAGAGGGCCGGTCTCGATATCCGTGAACCAGCCGGGTTCCGCGGCCTCCATTCTCTCCGCCGCACCACCATCGT | 8 |
| wt-54690-54810 | 4735 | CCACGAGCAGGGATCGCAGTCTCTCCCTCCTCACCCTCGTTATTCCCAATAGCATAGCGGCAAGGATCTGTGTGAGGGAGTCCAAGATGTCCGTGTTTCTGGCTACTGCCGCCGCTGCTG | 4 |
| wt-54810-54930 | 4736 | CCGCGGCTGAGTCCGTATTGTCTGGCAGCAGGGAGGCCAGCAGGGTGTTCCAGTCATCGGGCGAAGTGGGAGCGGGCTCTGGGCGTGCCCCCAGCGCCTTCCTAATTCTGGCCCAGGCCT | 8 |
| wt-54930-55050 | 4737 | CATTCGCCTCTCGCGCTCGCCGCTCCTGCCTCTCCTTGTCTTCCTGTTCTCGGAGCTTCTCCTTTTCCTTGCGCCCGGTCTCCATAAGCTGCCGCAGCTTCTTCTCATACTGTCGCTTGA | 4 |
| wt-55050-55170 | 4738 | GCTCTTTGTTGGGGGCAGTGTCCAGAAAGGCCTCGAGCTGTTCCTCGGTGGCGGGCTTAAAGCCTTCGGCCTCCAGGCGCCAGGCCTGCACCTCCTTCTGTCTGAGCTGATCGTTGTTGT | 4 |
| wt-55170-55290 | 4739 | TATTCTTCAATTTCTGCAGGTAACTTAGGAAGCGTTTCTTGAGCTTCCCTGGGATGAGCGTTTGGGAGAGCTGATTCTGCAGCCCAGAGAGTAGTCTCAGGGCATCCTCTGGAGCCTGAC | 4 |
| wt-55290-55410 | 4740 | CTGTGATCGTCGCATCATAGACCGCCAGTAGACCTGGGAGCAGATTCACCGCCGCGGCCGTCTCCTTTAAGGTGCTGTGAGTAGCAAAATTCTGCAAGGCCACTAGGCGCGCTGGCTCCA | 4 |
| wt-55410-55530 | 4741 | GCGTCAGCCGGTTGCCCATCTCGAATGTGTGCAGGGCCTCTGAGACCATGGGGTCCAGGATGCGGTCAATGCCATCCTGCACCTCAGGGTCAAGGACCGGCAAGTCACGATAGAGGTGGT | 4 |
| wt-55530-55650 | 4742 | CTATGCTCTCCTCGAAGGAGGCAATGTAGTTATCGATGGTGTAGAAGGTGATGGATTTCAGGATGTTCATCAGGTACTTTTTGGAGCGAACAATCTGCTGTATAGTGTCACGTAGGCGGA | 4 |
| wt-55650-55770 | 4743 | TGTACGTGGGGTTCTTTGCGGCCCCGACTATCGACCCTGCATTTGCGATGTACTTTTCTATGACGGGGATGGTGAGGGCCGCGGTGTCGGCCAGCGGTGGCGTGGCTTCGGGGTTGTCGT | 4 |
| wt-55770-55890 | 4744 | GGTTGGCGGGTGTCGCAGAGGGAGAGGCGGGAGAGATGGGGGCGCCTGGGGCCGAGGCCACACCGGCCAGGCCCAACATTGCCTCGATGTCGTCCAGGATGGTGCGGAGGCGCTTTTCGT | 8 |

| | | | |
|---|---|---|---|
| wt-55890-56010 | 4745 | TTTCTCTGGTGGTCTCGAGCTCCTTCTGTTTTTTCGCGACTGTCTCAAACTCTGGAAGGGGGGCAATGCTGGGGTCGTCCTCCTCAACTCGCTCCAGGGGCCAGGGGATACCGCTCATAT | 4 |
| wt-56010-56130 | 4746 | CACTAAGGGCGGTGCCCAGGTAGAGGAGCTCGCGATAGTCCCATTCAATGGACGTGTACCGGATGTTTAGGAGAGGCAGGGAGGCGATGATCTGGCATGTGTGCCGCAGGTGTGTCAGGA | 4 |
| wt-56130-56250 | 4747 | GGTCGTCAAAATCCATCACTGTTGGGAGGCTTGGGTCCTCAAGGTAGGAGAGATAATCGGAGGCCGCCGAGGCCACCTTGTCCCTGATGTCCGCCGTACACCTGCGCACGTGCAGGGCCG | 4 |
| wt-56250-56370 | 4748 | CATTCTTGGACCGGACGGCCACGTTGTGGACAAAGGGGGGCACTGAGGCGGCGGGAGGGGCCCCATACTCTATCGCTGTCAACAGCGCCAAAAAGCGGACGTCCTCCTCATCTACCCCAG | 4 |
| wt-56610-56730 | 4751 | CAGCCAGCATCTTAGCACCGGCATCTACACGGACCGCGTCCTTCGTGGCCAGGGACGGCAGGCAGGCCTCCAGCTTTGCGGCCAGGTGCTTATGGAACTCTCCCGCTCTTCCCTTGTTTT | 4 |
| wt-56730-56850 | 4752 | CTGATAGCATGTTTGCGAGGTTTTGGATGTTAAGTTCGGAAGTGAGCAGTTGCTCCAGGTCCAGCGTGGGGACCTGCAGATGTCCCGACCAGTCCTTTAAGAATTCCAGCAGATTTAGCA | 4 |
| wt-56850-56970 | 4753 | CAGACGATCGGTCCCTACTCCTTATTAGCCCCTGCTCGAGGACCACTGTCACAAGAAGATAGTCTATCATGCTCAAGGCATCTGCCTCTGGCACTTCCCGGTTAGAGGCCGGGTCGTAGA | 4 |
| wt-56970-57090 | 4754 | CGATGGCCTGTTCCTGGTAGGTATGTCCGGCTATTCTCGCAATGTTGCTCTCGAGGGGCACAAAGTCCATCTCAGGAGTCTCTATGTCAAAGGTGGTCTGATAGTATTGGCTCCTGGCGG | 4 |
| wt-57090-57210 | 4755 | TGTCCAGTGTGATGGGGGACGTGGGGGCACTGGATCCCGATTCCAGGCTGTTGGAGAACACTTCATCTTCGAACATGTCTTCATCCTCTGTGGTGGGGATATCGGAGGCTAAGTCGCTCT | 4 |
| wt-57210-57330 | 4756 | CCGCTTCTTCAGAGTCGGACATGGATAGGAAAGGCTCCTCTAGGTCAGACAGGTAGCGGACGAGGCCAGAACCCCCAGATGCATCATCCCCAAAGGAGGGCTGCTGCCCGAAGGGAGGTG | 4 |
| wt-57330-57450 | 4757 | ATGGGGATATCTCCGTTCCAGCCCTGTCAGCGGCCGGGGGATGGTTTTTTTCTGGTTCGAGTGTCGTGGCTGATGGTGGGAGCTGCTGAGCAGGAGGAGGAGCCGGGGTAGCTGATGGCA | 4 |
| wt-57450-57570 | 4758 | GGGGCTGCTGCTGAGGAGGAAGAGGAGAAGGAGCCCGGGCGGCTGATGGCGGGGGCTGCTGCTGAGGAAGAAGTGGAGAAGGAGCCGGGGCGGCTGATGGCGTGGGCTGCTGCTGAGGAA | 8 |
| wt-57570-57690 | 4759 | GAAGTGGAGAAGGAGCCGGGGCGGCTGATTGCGGGGGCTGCTGCTGAGTTGGAGGAGGAGAAAGAGTCGTGGTGGTGGGGGCTGCTGCTGCAGTCGGGGAAGGGGATGGGGTGGTCAGAG | 8 |
| wt-57690-57810 | 4760 | GGATTTTTGGGTTCGAGGGAGCTGCCTGTGGCAGAGGGATGGGTATTTGCAAAGGGAGGCGAGGAGATGGAGTGACTGAAGGAGCGATAGTTGAGACTGGCGCGGGGTGGGGTGTCGGGG | 4 |
| wt-57810-57930 | 4761 | AGGCGGGTGGTGATTGGTGAGGGATGGGGATTACTGGAGGGGGAAGGCGAGCTGCTGAAGGGGGGCGATGGGGCGGAACGTGGGTGCGTGGCAGCTGATCATCCTCTGTGTCAGTGGTGG | 4 |
| wt-57930-58050 | 4762 | AGGACAGAGGGAGGCGGCGGCCGGAGGTGGGCTTCTTGTGGGGGCTATCTTTGCCCAATCCCTTTTTCCTCTTGGGAGTCTGAGGCGCTGCGCCGCTCGACGCCCTTGGTGGCGTGGAGG | 8 |
| wt-58050-58170 | 4763 | GAGCGGGGACCCCGGGGGTGTGACCTAGGCCGGGGATGGGGATGAAGAGGGGAGGGCTGGAGGCCGGGGCCGCGGAGGCCGGGGCCGCGGAGGCCGGGGCCGCGGAGGCCGGGGCCGCGG | 12 |
| wt-58170-58290 | 4764 | AGGCCGGGGCCGCGGAGGCCGGGGCCGCAGAGGCCGGGGCCGCAGAGGCCGGGGCCGCAGAGGCCGGAGACGACGGCGGGGAGTTGGTCTTTGCAGGACTATACCTGGCGGCAGGGAATG | 12 |
| wt-58290-58410 | 4765 | AGTCGGATGTGAAAGATCGAGAGGGCAGTGGCCTGAGGTTATACGGTATTATTCGCCGTTCAAACGGTAGCATGACGGGAGGGCTGCTATCAGCACCGGGCGTCCCCGCCGCCTCCCCAT | 4 |
| wt-58410-58530 | 4766 | CACTGGACACAAGCTCGGGCCCCACCAGGTCAAAGCCGCTGCCGTTGGCCTCATAAAAGTCATACACGCCATAGTGTTCCAGCATAAAGATGCGGGGGTCCTCTGTCTCAAAGGCCTCGG | 4 |
| wt-58530-58650 | 4767 | GTAGAAAATAGAGATGCACGCAAGTGTACTGGGCCCCTGGTGCCCCCACGTACTGCAGGATGTCGTGCGCATAGGTGCTGACTCTGACATGGGCGGGGGTGCCCGGGGCCGCATCCTTCT | 4 |
| wt-58650-58770 | 4768 | GGCAGTGGGGGTCAAACAAGTAGAAGGAGCCATCTGTCTCGATGATGATGGCCCCCGCGTAGATGTCGCAGATGTAGAGGATGAACTGGGCCACCCCGTTGTAACTGCCGTGCAGGACCT | 4 |
| wt-58770-58890 | 4769 | CGGCCAGGGACTGAACAACTGCCGAGTTTGCGATCTGGGCAGGGAATAGGACGAGGCCAAAGATCTCCGCCGAGCGGTATATGTGCACGCGCCCACCGCCCCTCAGGACCACGGAGCTGG | 4 |
| wt-58890-59010 | 4770 | GCACGTCCGTCAACTGGGCCATCTCGTGCCCCTTGAGGATGCCGCTCTGGCGCATGAGGGCATCCAGCCGCGCCCCCTCGTCCAGGACCTCGTCCAGCTCAGGGCGGGAGGTCAGGGGGC | 8 |
| wt-59130-59250 | 4772 | CCGAGCCTCCACTCCCGCCCCCGGGGGCATTCACATCCATGGTCCTGATTCCGCGCACGGGGCCGGTTCCCCGGGTGCGCTGGCTTTGCCCCCAGTCGCCGTTACTCATCTTCGGCGGTG | 8 |
| wt-59250-59370 | 4773 | GGGCGGGGAGGACGCCTTGTCGCCCCCCTTCTGGTCCGGGGTCTTACGCGGCTGGCGGCGGCAGCCGCCGAGAGATAAGGGGGGTACGTGTGTGCCTCCGCCTCTCCTCTGTCTGGGCCG | 8 |
| wt-59370-59490 | 4774 | CCGCCGCCGCTTGCCCGCCTTGAAGGAGAGGGGGTAGTCCGCGGACTGCGTCTGCGGGGGCAGGAGGTCTCAACCTTCTGGGCTCGGGCCGCGGTGTCGATATCCGATGGCCTTTCCCTG | 8 |
| wt-59490-59610 | 4775 | TCTTCCTCGTATGCTCCTTCTCCTCCTCCTCCCGGCACGCCCCTGAGATCTGCCTCCCCTCCCTCTCCCTCGTCCTGGTCGGAAAAGTCTGAGGAGGAGAAGGAGAATGGGGAGGAGTCC | 4 |
| wt-59610-59730 | 4776 | AAAACGGCACGCCACCTGCCGTGGGGCGGTGGTGACAGGTCCCGGCTGGCCCGGCGCTTGCTCGCGTTCCTGCCGTTACCCAGGAGAATGGCCGCGAGTTTTTTGGCGGGGAGGATGCGG | 8 |
| wt-59730-59850 | 4777 | AATGGCGGGGGCGTTTGTCCCACGGGTGAGGGGGAATCGTCGGTTAGGGCCGGCACGAGGTGGTGGGTCTGGACCCGGGCCGTGCGAGCAAAGGCGGCGAGAACCGAGGGGCTTCTGGGG | 8 |
| wt-59850-59970 | 4778 | GTGACTGTGATCTGTTCCGGATTTAGGTCCATGGCGGGTGTGTATGTTTTAATAGGGGTGGTCTCTGGCGCGGCAGGATGATGGTCGAGGACGTCCACCAGGGCCTTGCAGATGCTCTTG | 4 |
| wt-59970-60090 | 4779 | CCTAGATACAGGATGTCGTCCATGCTGAGGGGAGGTGGGGTGTCTGCTCCCCCCTGCGGAAGCCGCCTGGGTGCGGGGATGAAGACAGGTGGTGGGCGGGCGTCTCGCCGGACTATGGCC | 8 |
| wt-60090-60210 | 4780 | TCGGCACGCTCGGCGTCGATGGCGGGTGGCTGGAACAGGCGGGCGAATGTGTAATCCCGGAACCGGTAGGCGACGCTGCGCCTGAGGGCGCCCGTCAGGCTGTATCCCAGCTCCAGGGCG | 8 |
| wt-60210-60330 | 4781 | TGCTCCACCCGCTCGTTGAGCTCCTCGAGATCCGGACGCAGGGGCTCGCTGGTGTGGGCCCAGAGGGGGTGATCCGCGATGCCCCGGCTCTCCCTGAGGGCCGGCACCAGGAGGCGCCTT | 8 |
| wt-60330-60450 | 4782 | CTGAGGGTGGCCGTGTCGGCCGTGGCCAGGGCCCACCTGGCGGCGGCGTCCCGGCACACATCCTGGATGCCCTCCACGACGCTCTTTAGCGTCTGGAGGTCCGTGGAGTAGTGGCGGGGG | 8 |
| wt-60450-60570 | 4783 | GAGGATGAAACGCTCTTTTCCTTCACCGCTACCACCGCCTCCTCCTCCTCTTCCGTCGCCAGAGGGATCTGCACCCTCCCGGTCTCTGCGTCGTACAGGAGCGGGCGGGAGCACAGCCTC | 4 |
| wt-60690-60810 | 4785 | GAAAAGAGGGAGGTGGGCCAGGCAAGCGGTGGTGCTTCCGCCGCCCGCTCTGAAGCTGAGATAGTCTCGGAGATGATGCCTGAGACCTCTCGGACGGCGTCCATGATCCTAAGGACTGCG | 4 |
| wt-60810-60930 | 4786 | TCGTGGGACGACAGCCCCCAGGGGCCCCCGCCCTCTTCGTCTTCTGCACCCTCGGCTCCTGCGTCCCCGGCCTTGCCTTCCCCCTCTAAGTTGAGGGGGCGCAGTCCGACCGCCTGGGGG | 8 |
| wt-60930-61050 | 4787 | GACTCCCCAGGCATCGGAGGGGCCCCGTCATAGATCTCCCAGACGGTGGCGTATATGAGCTCGAGAGGACGGCGGGCCCGGGTCAGCTCGGGGGAAGGGAGGGCCAGGTCGCTGCCGAAG | 8 |
| wt-61050-61170 | 4788 | GAGACCAGCCAGCGCAGGGCGGCCAGAGAGCGGGTTTTGGGCAGCTCGTTGGAGAGGACCCGGCGAAGGGCGGGCCAGATTTGGAACTCGATGAAGGCGGCCGGGAAGAAGGGGCTGTGG | 8 |
| wt-61170-61290 | 4789 | ACATAGGCCGGATCCGCGCGCGCCGTTTGGCCGGCCCTCAGGGACCGGCAGTATGCCTCGACGTCTGTCCGCGGGGCCGCCGCCACCGCTGCCGTCCACTGCCTTCTTCCCTGCTCGCCG | 8 |
| wt-61290-61410 | 4790 | GGGAGTAGGGGGGGCTTACAGGGGAGGGCCGGAGCCGGGGCCGGGGCCTGCCACAGGCGGCTGTAGCGGACCCATAGCAGAGACCTGAGGAGTTCGGATGAAAGGTCCCCCGCCACCTGC | 8 |
| wt-61410-61530 | 4791 | TCATACTCGGCCGCGGGAGGGGGGACGATGAAGATGCGCAGAGGGGTTACGGCGTCCCAAGGGTCCGCCGCCGCCCCCACACCCACAGCCGTCGCGGCGGGGGCGGCGGCGGGCGTAGAG | 8 |
| wt-61530-61650 | 4792 | GGGCCGCTGGTGCGCCGGGCTCGTCTGTCCACGGCCTCGGCCTCCGCCCGCAGGTAGGCCGCCCGGGCCACACGGGCGAAGCGGCTCGTGGGGCTCGCGGTGGGCAGCAGTCGGAAAAAG | 12 |
| wt-61650-61770 | 4793 | TGCAGGGCAAAGCCCGATAGACTCTCTAGGAGGGCGGCGGTGGCCTCGAGCCACCTCCACCGCGAGCGGGACACCCGGGGCACAGAGGCCAGCATCATGGCGTAGTCCCCCGCCACGGTC | 8 |
| wt-61770-61890 | 4794 | TCGTTGAGCCCGGCCGAGAGCAGAACCGTGGCCACCTGCTCGATGGCGGCTGGAGAGAAGGATGCCCGGCTCCCCGCCGCCTCCTGCACACGAGCGGCCAGGGCCTCCATCTCTGCCGCC | 8 |
| wt-61890-62010 | 4795 | ATCCCGGCCAGGAAGGCCTCGATGACCGAGTCTGGGACGCCGTAAGTCTGGTCCCAGAGCAGGGCCTCGTACACATAGTCGTAAAAGAGGGCCCCTGAGGGCTCCAAAAGCCGGAGCCGG | 4 |
| wt-62010-62130 | 4796 | GCGGCGTCAAAGGCCAGGACGGGCACAGCCGCGACGGGGGGCGTTTGTCCCCCGCTGGCCTCCGCGTACACGCCCAGGATCTCTACCGCCCGCCGCCGGGCCAGGGGCAGCGAGGCCACC | 12 |
| wt-62130-62250 | 4797 | ACGCTGGAAAGTGACTCGGGGCGGTGAAAGAGACCACCACCGCTTTCTTCACCCTCTCCCCCGCCGGCCCCGCCCCCACTGTGCTCCACCAGCTCCACGGCCATGGCCTTGATGTCCGCG | 8 |
| wt-62370-62490 | 4799 | TCCCGCGGGAAGGCGTCCCACCGGGCCAGATAAGTGTCGGCCACCAGCTCTTCCACGAAGGCAAAGGTGGCCGTTGGGCCAGAGACCGCGAGCACCTCCCCGCTGCCCTCTTCGATGATG | 8 |
| wt-62490-62610 | 4800 | CGCCGGTACGCGGCCGCCAGGGCCCGGGTCTCTGCGATGAGCCGAGAGCCGTCCAGCGGATCGTCGGTGGCCGGAGAGGCTGTCGTGGGGGGCAGTGAGGATGCCAGCACGTCCAGGGCC | 8 |

| | | | |
|---|---|---|---|
| wt-62610-62730 | 4801 | GCCTCCAGATGGCCGAGGCCGAAGCTGCGCCTGGAAAAGGAGGCCGCCCGGAGTAGGTAGTAGGCGTGGTGGCGGAGGACCGCCGCCGGGTAAGCGTGGCCGCTCATGAGGGTGAGAGTA | 8 |
| wt-62730-62850 | 4802 | TTTAAAAAATCGCGCACCAGCACCGGCTGGGCCAAATCCCCCAGTCCAAAGATCCCCAGCTCCAGAGGCATCAGCGCGCGCAGGCGGGCAGCGGGGTCGTCCCCAGACAGCAGCAACTGA | 4 |
| wt-62850-62970 | 4803 | CGCGTCACGCGGGCGAGCCCCCCGTCCACCTCTGCCAGGGGTGGCTGGGCGTCTGCCCCTCCGCTACCGCCGCTGCTGTCACTCTCCATAGCGGACGCCATGAAGGTCCAGGGGTCCGTC | 8 |
| wt-62970-63090 | 4804 | GATCGCCGCCGTCTGCAACGCCGAATCGCGGGGCTGCTGCCCCCTCCGGCCCGGCGTCTAAATATTTCCCGGGGGTCCGAATTCACGCGGGACGTTCGTGGGCTGGTTGAGGAACACGCG | 8 |
| wt-63090-63210 | 4805 | CAGGCCTCCTCGCTGAGTGCGGCGGCCGTCTGGCGCGCAGGGCTGCTGGCCCCGGGGGAGGTGGCGGTCGCCGGGGGTGGCAGTGGAGGGGGGAGCTTCAGCTGGTCTGGGTGGCGGCCG | 12 |
| wt-63210-63330 | 4806 | CCAGTCTTTGGGGACTTTCTGATACACGCCAGCTCCTTCAACAACGCCGAGGCCACTGGAACGCCCCTTTTCCAATTCAAGCAGAGTGACCCGTTCTCGGGCGTCGACGCGGTATTCACT | 4 |
| wt-63330-63450 | 4807 | CCTCTCTCCCTGTTTATCCTAATGAATCACGGCCGGGGTGTAGCCGCCCGGGTCGAGGCAGGTGGGGGCCTGACGCGGATGGCCAACCTGCTGTACGACAGCCCCGCAACCCTGGCTGAC | 8 |
| wt-63450-63570 | 4808 | CTGGTCCCGGACTTTGGGCGGCTGGTGGCCGACCGCCGCTTCCACAACTTCATCACCCCTGTGGGCCCCCTGGTGGAGAATATAAAGAGCACCTATCTGAATAAAATCACCACGGTGGTC | 4 |
| wt-63570-63690 | 4809 | CACGGGCCTGTGGTCAGCAAGGCCATCCCTCGCAGCACCGTCAAGGTGACGGTGCCCCAGGAGGCCTTTGTGGATCTGGACGCGTGGCTCTCCGGCGGCGCCGGGGGTGGCGGTGGAGTA | 8 |
| wt-63690-63810 | 4810 | TGCTTCGTCGGGGGGCTGGGCCTGCAGCCGTGCCCCGCCGATGCGCGCCTCTATGTCGCTCTGACCTATGAGGAAGCCGGGCCGCGGTTTACGTTTTTCCAGTCGTCCCGCGGCCACTGT | 8 |
| wt-63810-63930 | 4811 | CAGATCATGAATATCTTAAGAATTTATTACTCACCATCCATCATGCACCGCTATGCTGTGGTCCAGCCCCTACATATAGAGGAGCTAACCTTCGGGGCGGTTGCCTGTCTGGGGACATTT | 4 |
| wt-63930-64050 | 4812 | AGTGCTACTGACGGTTGGAGGAGGTCTGCCTTCAATTACCGTGGCTCTAGCCTCCCCGTGGTGGAGATTGACAGCTTTTATTCCAACGTCTCTGACTGGGAGGTGATTCTCTAGACTTAA | 4 |
| wt-64050-64170 | 4813 | CGGGAGGAAACAGGAGGAGGAGGGGGACAAGAGCACAAAAGTGGTTCAGTGGACACCCACCACACAGCATGGCAACGACCAGTCATGTCGAGCATGAGCTCCTCTCCAAATTGATTGATG | 4 |
| wt-64170-64290 | 4814 | AGTTAAAGGTCAAGGCCAACTCAGACCCCGAGGCTGATGTCCTGGCCGGGCGCCTGCTCCACCGCCTTAAGGCCGAGTCAGTTACACACACAGTAGCCGAATATCTGGAGGTCTTCTCTG | 4 |
| wt-64290-64410 | 4815 | ACAAATTCTACGATGAGGAATTCTTCCAGATGCACCGGGATGAGCTGGAGACCCGAGTCTCTGCTTTCGCGCAGAGCCCGGCCTACGAGCGCATCGTCTCCAGCGGCTACCTGTCGGCCC | 4 |
| wt-64410-64530 | 4816 | TGCGCTACTATGACACCTATCTGTATGTGGGGCGCAGCGGGAAGCAGGAGAGTGTGCAGCACTTTTACATGCGGTTAGCCGGCTTCTGTGCCTCAACCACCTGCCTCTACGCGGGTCTCA | 4 |
| wt-64530-64650 | 4817 | GGGCAGCCCTGCAGCGGGCCAGGCCGGAGATTGAGAGTGACATGGAGGTGTTTGATTACTACTTTGAGCACCTAACCTCCCAGACGGTGTGCTGCTCCACGCCCTTTATGCGCTTTGCCG | 4 |
| wt-64650-64770 | 4818 | GGGTGGAAAACTCCACTCTGGCCAGCTGCATCCTCACCACCCCCGACCTCAGCTCCGAGTGGGACGTGACCCAGGCCCTCTATAGGCACCTGGGGCGCTACCTCTTTCAGCGAGCCGGGG | 8 |
| wt-64770-64890 | 4819 | TGGGTGTAGGGGTGACGGGGGCTGGCCAGGATGGGAAACACATCAGCCTCCTGATGAGGATGATCAACAGCCACGTGGAGTACCACAACTATGGCTGCAAGAGGCCGGTCAGTGTGGCGG | 4 |
| wt-64890-65010 | 4820 | CCTACATGGAGCCCTGGCACAGCCAGATTTTCAAGTTTTTGGAAACGAAGCTGCCGGAGAACCACGAGAGGTGCCCGGGCATCTTTACGGGGCTCTTTGTCCCCGAGCTCTTCTTCAAGC | 4 |
| wt-65010-65130 | 4821 | TTTTTAGGGACACGCCCTGGTCGGACTGGTACCTGTTTGACCCCAAGGACGCCGGGGACCTGGAGAGGCTCTACGGGGAGGAGTTTGAGCGCGAGTACTATCGGCTGGTGACAGCGGGCA | 4 |
| wt-65130-65250 | 4822 | AGTTTTGTGGGCGGGTCTCCATCAAGTCCCTGATGTTCTCTATCGTCAACTGCGCCGTCAAGGCCGGCAGCCCCTTCATCCTTTTGAAGGAGGCCTGCAACGCCCACTTTTGGCGCGACC | 4 |
| wt-65250-65370 | 4823 | TGCAGGGCGAGGCCATGAACGCCGCCAACCTGTGCGCCGAGGTGCTGCAGCCCTCGAGGAAGTCTGTGGCCACCTGCAATCTGGCCAACATCTGCCTCCCGCGCTGCCTGGTGAATGCGC | 8 |
| wt-65370-65490 | 4824 | CTCTGGCGGTGCGGGCACAGCGGGCCGACACGCAGGGGGATGAACTCCTGCTGGCCCTCCCTCGACTCTCAGTCACCCTACCTGGAGAGGGGGCAGTCGGTGATGGATTCTCGCTAGCCC | 8 |
| wt-65490-65610 | 4825 | GCCTCAGAGATGCCACCCAGTGTGCCACCTTTGTGGTGGCCTGCTCCATTCTTCAGGGATCCCCCACTTATGATTCCAGGGATATGGCCTCCATGGGCCTCGGGGTGCAGGGCCTGGCCG | 4 |
| wt-65610-65730 | 4826 | ATGTCTTTGCGGACCTGGGCTGGCAGTACACTGACCCTCCCTCTCGCTCGTTAAACAAGGAAATATTCGAACATATGTACTTTACGGCCCTCTGCACCAGTAGTCTGATTGGACTTCACA | 4 |
| wt-65730-65850 | 4827 | CCAGGAAGATTTTTCCGGGTTTCAAACAGAGCAAGTATGCCGGGGGGTGGTTTCACTGGCACGATTGGGCAGGAACAGACCTTTCTATTCCCAGGGAAATTTGGTCTCGCCTCTCTGAAC | 4 |
| wt-65850-65970 | 4828 | GCATTGTGAGGGATGGGCTTTTCAATTCACAGTTTATCGCCCTGATGCCCACCTCAGGCTGTGCCCAGGTGACGGGCTGTTCGGACGCCTTCTACCCCTTCTATGCCAATGCGTCCACCA | 4 |
| wt-65970-66090 | 4829 | AGGTCACCAACAAGGAGGAGGCCCTTAGGCCAAACCGGTCTTTTTGGCGTCATGTGCGTCTGGATGACAGGGAAGCTTTGAATCTTGTCGGGGGCCGTGTCTCCTGCCTCCCGGAGGCTC | 4 |
| wt-66090-66210 | 4830 | TGCGGCAGCGCTACCTGCGTTTCCAAACGGCCTTTGATTACAACCAGGAGGACCTGATTCAGATGTCCCGGGACAGGGCCCCCTTTGTGGACCAGAGCCAATCTCACAGCCTGTTTTTGC | 4 |
| wt-66210-66330 | 4831 | GTGAGGAAGATGCCGCGCGGGCCAGCACGCTAGCCAACCTACTGGTGCGCAGCTACGAGCTGGGCCTGAAGACTATCATGTACTATTGTCGCATTGAGAAGGCCGCCGATCTGGGGGTGA | 4 |
| wt-66330-66450 | 4832 | TGGAGTGTAAGGCCAGCGCGGCTCTGTCGGTGCCGCGGGAGGAACAGAATGAGCGGAGTCCCGCTGAGCAGATGCCGCCTCGTCCCATGGAACCGGCGCAGGTTGCGGGGCCGGTTGACA | 8 |
| wt-66450-66570 | 4833 | TCATGAGCAAGGGCCCAGGGGAGGGACCAGGTGGGTGGTGTGTGCCCGGGGGATTGGAAGTGTGCTATAAGTACCGTCAGCTCTTCTCAGAGGATGATCTGTTGGAGACTGACGGTTTTA | 4 |
| wt-66570-66690 | 4834 | CTGAACGAGCCTGTGAATCTTGCCAATAAACGTTTATTGCCATGTCCAAGTTGTTGTACGTGCGTGATCATGAGGGCTTTGCCTGCCTAACGGTCGAAACCCACCGCAACCGCTGGTTCG | 4 |
| wt-66690-66810 | 4835 | CGGCTCACATTGTCCTCACCAAGGACTGCGGGTGTCTCAAGCTACTCAATGAGAGGGACTTGGAGTTTTACAAGTTCCTCTTTACGTTCCTGGCCATGGCCGAGAAGCTTGTGAACTTTA | 4 |
| wt-66810-66930 | 4836 | ACATTGATGAACTGGTCACCAGCTTCGAGAGCCACGACATTGATCACTACTACACCGAGCAGAAGGCCATGGAGAACGTCCACGGGGAGACTTATGCTAACATTTTAAACATGCTCTTTG | 4 |
| wt-66930-67050 | 4837 | ATGGGGACAGGGCGGCGATGAACGCCTACGCAGAGGCCATCATGGCCGACGAGGCCCTGCAAGCCAAGATTTCCTGGCTCCGTGACAAGGTGGCGGCCGCCGTCACCCTGCCGGAGAAGA | 8 |
| wt-67050-67170 | 4838 | TTCTTGTGTTCCTGCTGATTGAAGGCATCTTCTTCATTAGCTCCTTCTACAGCATAGCCCTGCTGCGGGTCCGGGGCCTAATGCCTGGCATCTGCCTGGCCAATAACTACATAAGTAGGG | 4 |
| wt-67170-67290 | 4839 | ATGAGCTGCTCCACACCCGCGCTGCCTCCCTGTTATACAATAGCATGACAGCCAAGGCTGACCGACCAAGGGCCACCTGGATCCAGGAGCTGTTTCGCACTGCGGTGGAGGTAGAGACTG | 4 |
| wt-67290-67410 | 4840 | CCTTCATCGAGGCTCGTGGAGAGGGGGTTACCTTGGTGGATGTGCGAGCCATAAAGCAGTTTCTGGAGGCCACGGCCGATCGCATCCTGGGTGACATTGGTCAGGCTCCCTTGTATGGCA | 4 |
| wt-67410-67530 | 4841 | CACCACCCCCCAAGGACTGCCCGCTCACCTACATGACTAGCATCAAGCAAACTAATTTCTTTGAGCAAGAGAGTTCCGATTACACCATGCTGGTGGTAGATGACCTTTGAGTCAGGGTGG | 4 |
| wt-67530-67650 | 4842 | CTACTTGCTCAGGTTTCTGGGCATAAATTCTCCTGCCTGCCTCTGCTCTGGTACGTTGGCTTCTGCTGCTGCTTGTGATCATGGAAACCACTCAGACTCTCCGCTTTAAGACCAAGGCCC | 4 |
| wt-67650-67770 | 4843 | TAGCCGTCCTGTCCAAGTGCTATGACCATGCCCAGACTCATCTCAAGGGAGGAGTGCTGCAGGTAAACCTTCTGTCTGTAAACTATGGAGGCCCCCGGCTGGCCGCCGTGGCCAACGCAG | 4 |
| wt-67890-68010 | 4845 | CCTGTGTCCTGGGCAAGGAGCTGTTTGGCTCGGCTGTGGAGCAGGCTTCCCTGCAATTTTACAAGCGGCCACAAGGGGGTTCCCGGCCTGAATTTGTTAAGCTCACTATGGAATATGATG | 4 |
| wt-68010-68130 | 4846 | ATAAGGTGTCCAAGAGCCACCACACCTGCGCCCTGATGCCCTATATGCCCCCGGCCAGCGACAGGCTGAGGAACGAGCAGATGATTGGGCAGGTGCTGTTGATGCCCAAGACGGCTTCCT | 4 |
| wt-68130-68250 | 4847 | CGTTGCAGAAGTGGGCACGCCAGCAAGGCTCAGGCGGCGTTAAGGTGACACTCAATCCGGATCTCTACGTCACCACGTATACTTCTGGGGAGGCCTGCCTCACCCTAGACTACAAGCCTC | 4 |
| wt-68250-68370 | 4848 | TGAGTGTGGGGCCATACGAGGCCTTCACTGGCCCTGTGGCCAAGGCTCAGGACGTGGGGGCCGTTGAGGCCCACGTTGTCTGCTCGGTAGCAGCGGACTCGCTGGCGGCGGCGCTTAGCC | 8 |
| wt-68370-68490 | 4849 | TCTGCCGCATTCCGGCCGTTAGCGTGCCAATCTTGAGGTTTTACAGGTCTGGCATCATAGCTGTGGTGGCCGGCCTGCTGACGTCAGCGGGGGACCTGCCGTTGGATCTTAGTGTTATTT | 4 |
| wt-68490-68610 | 4850 | TATTTAACCACGCCTCCGAAGAGGCGGCCGCCAGTACGGCCTCTGAGCCAGAAGATAAAAGTCCCCGGGTGCAACCACTGGGCACAGGACTCCAACAACGCCCCAGACATACGGTCAGTC | 4 |
| wt-68610-68730 | 4851 | CATCTCCTTCACCTCCGCCACCTCCTAGGACCCCTACTTGGGAGAGTCCGGCAAGGCCAGAGACACCCTCGCCTGCCATTCCCAGCCACTCCAGCAACACCGCACTGGAGAGGCCTCTGG | 4 |
| wt-68730-68850 | 4852 | CTGTTCAGCTCGCGAGGAAAAGGACATCGTCGGAGGCCAGGCAGAAGCAGAAGCACCCCAAGAAAGTGAAGCAGGCCTTTAACCCCCTCATTTAACACCATGTTCTCGTGCAAGCAGCAC | 4 |

| | | | | |
|---|---|---|---|---|
| wt-68850-68970 | 4853 | CTGTCCCTGGGGGCCTGTGTCTTCTGTCTCGGCCTCCTGGCCAGCACCCCCTTCATTTGGTGCTTTGTCTTTGCCAACCTGCTCTCTCTGGAGATCTTCTCACCGTGGCAGACACACGTG | | 4 |
| wt-68970-69090 | 4854 | TACAGGCTTGGATTCCCGACGGCATGCCTAATGGCCGTCCTCTGGACGCTGGTACCCGCCAAGCACGCGGTGAGGGCCGTCACTCCAGCCATCATGCTGAATATTGCCAGCGCCTTGATC | | 4 |
| wt-69090-69210 | 4855 | TTCTTCTCCCTCAGAGTCTACTCGACCAGCACGTGGGTTTCTGCCCCCTGTCTCTTTCTGGCCAACCTGCCTCTCTTATGCCTGTGGCCCCGGCTGGCCATCGAGATTGTTTACATCTGC | | 4 |
| wt-69210-69330 | 4856 | CCGGCTATACACCAAAGGTTCTTTGAACTTGGGTTGCTCTTGGCCTGCACCATCTTTGCCCTGTCCGTGGTCTCCAGGGCCCTGGAGGTGTCGGCTGTCTTCATGTCTCCATTTTTCATC | | 4 |
| wt-69330-69450 | 4857 | TTTCTGGCTTTGGGCTCTGGAAGCCTGGCCGGTGCTCGGCGTAACCAGATTTACACCTCGGGTCTCGAGCGGAGACGCAGCATTTTCTGCGCCCGGGGAGATCATTCGGTGGCATCCCTG | | 4 |
| wt-69450-69570 | 4858 | AAGGAGACCCTCCATAAATGCCCGTGGGATCTGCTGGCCATCTCTGCCTTGACCGTTCTTGTCGTCTGTGTGATGATTGTGTTGCATGTGCACGCAGAGGTGTTCTTTGGACTCTCTAGA | | 4 |
| wt-69570-69690 | 4859 | TACCTGCCCCTCTTTCTCTGTGGGGCGATGGCCTCCGGGGGGCTGTACCTGGGCCATTCCAGCATCATTGCATGTGTCATGGCCACCCTCTGCACCCTGACATCTGTTGTGGTATATTTC | | 4 |
| wt-69690-69810 | 4860 | CTCCATGAAACCCTTGGACCCCTGGGCAAGACCGTGCTGTTTATCTCAATCTTTGTCTATTACTTTAGCGGGGTAGCGGCCCTGAGCGCAGCTATGCGCTACAAGCTTAAGAAGTTTGTG | | 4 |
| wt-69810-69930 | 4861 | AACGGACCCCTGGTCCATCTCCGTGTGGTATACATGTGCTGTTTTGTCTTTACTTTTTGTGAATATCTGTTGGTGACATTCATTAAATCCTAACGACCGGAGTCCTGTCTCTTTGTGTTC | | 4 |
| wt-69930-70050 | 4862 | TTGGGGGACTTGAGTTAGCTGTCTTTCCTCTTATTACATTGGGCTAACGGGAGGAAATGAACCCAGGGGTGGCAGTGGATGGGGTCATTTATGGGCAAAACTCACAGGACATGTTTGGGG | | 4 |
| wt-70050-70170 | 4863 | AGTTAGCATTGGCGTCGGGAAACACAGCTCTGGCAGTTATAACCGCACCAGCTAACAGGACATGTTTGGGGGAGTTGGCATTGGCGTCAGGAGACACGGCTCTGTCAGTTATCACCGTAC | | 4 |
| wt-70170-70290 | 4864 | CATGAGTGCCATGTGTGTCCAGTGCCTAATCACCGTTCCTCATTTTGTGTGCCTCCTCAAATGTTCCAGAAGTCGGCCACAGGGGAGGTGGCTGAATTAGGGCCTTTTCCCTCATTCCCC | | 4 |
| wt-70290-70410 | 4865 | CATGAGACCCACGTGGCAGGCCTAGGGGCTACATTCGCCTCCCACGTTTCCCTTCGCGTGAGGCATCCGATATGACTGAATTTTCGCAGTCTCTTTTCCCTCTTCCCTTGTTATTCCCAT | | 4 |
| wt-70410-70530 | 4866 | AGAATTACAGTGAGGTTACACAGGTGGAGATTCAGTTTAACCATTTATTGATTTAATCCAGGAACAAAAAACAGTCCTAGTGACCCAGTGCCCGGAGAGAGAATGGCCCTGACAAGTCGG | | 4 |
| wt-70530-70650 | 4867 | CTGCATGATGCACTTCGGCAGTCACGTGTGTGAGTCTCCACGGCCTCTGTCAAAAGGGAGCTTAGCGTGCCAGGGTTGTAATTCTTGATGTAGTGGCCCAGGAATTCAACTTCATCGTGT | | 4 |
| wt-70650-70770 | 4868 | CTCCGTCTGCAGTTGGCGTTAATGTAGGCTGGGGCTACTGCCGCATATGCTGCCAAGAGACAGAGGGGCTGCTTCACATATGAGCTGCTCAGGGTCTCCACCACCTTGTTTTGACGGGCC | | 4 |
| wt-70770-70890 | 4869 | GTGGCACAGGTGATGTAGAAGAGTTGCTTCACAAAGTTGTAGTCTCGCGTGTTAGGAAGGAAGCAGGGTGCCAGCTCTTTGAGCTTGGTCAGGATCACCTTGCTAAGACTCATGGCGCAG | | 4 |
| wt-70890-71010 | 4870 | GCCAGGAGGATGTCTTCCGCGGGAGCTAGGGGCAGGTCGCCGTGGTAGGTGATCTCCTGGAGCCAAAAGATGGTCTCTTCTAGCATGGCCACCAGGGTGCAGAGCCCCGCGTTCTGGATC | | 4 |
| wt-71010-71130 | 4871 | GCCTGCATGCGTGCATCCAGCCATGTGTCCTTGTTGGTTGACTTGGTGAAAAACTCACGTAGTGTCTTGTAGCTCCTGCGCAGCTCGTGTCTGGGTTGCACTTTCTGCCAGGCTCCAATC | | 4 |
| wt-71130-71250 | 4872 | TCTGGATGGGCGGCCACCGCCAGCATCGACTGTAGGAACGGGTCTTGGATGGGCTCTAGGGTCAGAGAGGCCAGGGGGCTGGGCAAGGTGACAAATGTAATCTTGGAGACAGGCTTAACC | | 4 |
| wt-71250-71370 | 4873 | AGACTCATGTCAAACCACGGTTTGTTCGGCAGGGGCCTCTGGCTGCGTTCTTGCCTCGCCTGCTTCCTTGTGCTCCTGCCGGCCCCTCGAGATTCTGACCGGGGACCTCTGGTTGCTCTG | | 4 |
| wt-71370-71490 | 4874 | TTGCTTCGGGGAGCTCTTGGAGACCTCGGTGCTCTAGGCACCCTGGGGGCCCTTGGGGCTCTGGGCGCTCTTGCTCCCGGGGGCAGGTGTCGGCGCTTGCCATAACTTTCATCGGTGCAG | | 8 |
| wt-71490-71610 | 4875 | CCATGGACCTCTCCGCGTCGCCTTTTGTGGCCTCTGGTGTAAGAGGAGTTGCCAGTCTCCTCCTTCTCGTCCTCGTCCCTGCACAGGGGTGAGCGATGCAATGTGACTGTCTTGTCCTGT | | 4 |
| wt-71610-71730 | 4876 | AGGTCCCACTTCTTTCTGGGAATCACAAACGATGCCGAGGTAGGGGTTATGACCACGCTGGAGGGCCGTGCAGGTATGGCGTGGGCCGGAGTTGGATCTTCATCCTCCTCCTCTGAGGAT | | 4 |
| wt-71730-71850 | 4877 | GAAATCTCTCCATCTGTGGAGTGTTCTTCGCTGCCCTCCATAGGGTCCAGATCGCAGTCTGTGTTGGTGTCTGAGACCGCTTCGAGTTCCAGAATGTGGCTCTCTGCAGAGGGGAGACAA | | 4 |
| wt-71850-71970 | 4878 | AAGGTGGAGACTGCCTTGAGCACCTCTGTCTCAGGCACCGGATGCCCCCGGCTCCACGGCCCCGGCCACTGGCCGGTGTAGCTTCTTACCTGCGGGATCCTCGTTGGAGGAAATGCTGCT | | 4 |
| wt-71970-72090 | 4879 | AGTTCGGGAGAGTCTCTGAGAAGGAACCATCTTGTCTGTCTCTACGACGGGCTAGCTGGGATGTAGTGCTGTCTTGACTGGCCTCAGCCCTATTTATGATTCTGGAGGCGGGCACGCTGA | | 4 |
| wt-72090-72210 | 4880 | TGGAGAAATGGGCGGTCGGTTGATTGGCCCCACAGCGACCGGCGAAGCACTGACTCATGAAGGTGACCGTGATGGCCTGTGATGTGTAGTAGAGTACCAGAAACACCCTCACATTCTTGG | | 4 |
| wt-72210-72330 | 4881 | AGCTGGCCCTGTGGGTATGCCTCAGGCACGCAAAGTTCCTGCCCCGGGCATGGCACACCTGAACTAAGTTTGGCCCGGTTTGCTCAAACGTGACATGGAGAAACTGGGGGAATTTGTCTT | | 4 |
| wt-72330-72450 | 4882 | CTGGCACAGCTGTTGCCAGGGTGCTCATGAGCGAGGGCCAGATGCAGGAGCTGACCCAGGCGACGAGATCCAGGCCCAGATGTCCCTCTATCATGGCGCAGACATTCTCCACGGTGGGGG | | 4 |
| wt-72450-72570 | 4883 | GCAGGGTCTCGCGGGTCCTCTGGATTAGATAGTCACGCCCATCATCCGCGATGTGGTAGCAGAAGGTTTTGGGGGCCGGCCAGCCCACGTGCAGTGAGTGATGTAAGAGGTTTTGAATGT | | 4 |
| wt-72570-72690 | 4884 | TGAGGGCATTCTTAACATAGCTGTGCTTGTCTTCCTCTTCCGGATGACAGACAAAGAGGCGCAGCTGCCGGCTAAGACCACCGCCCCTGTCCACCTTGTAGGTATGCGGCAGCCGGATGC | | 4 |
| wt-72690-72810 | 4885 | ACCGCCCGGCGTGATACACGCCGCTGTCAAAAAGCGGGGCCCCAATCTCTTTGATCTTGTGACGCATGCGGCGCAGGCAGGCCGTTAGGCCCATGAGCTTCTGCAGCACAGACACAAACC | | 4 |
| wt-72810-72930 | 4886 | CTTGTACTGCGCTTGTTCCCACAATAGCATGGCCTCTAGGTAGGGGGGTGATGACGCGAAAGCCCAGTTTTCCCGTGCATATGCAAAAGGGGAGCACATCTTCCATATTATCCGGGTCGG | | 4 |
| wt-72930-73050 | 4887 | CGGGTGGACAAGCTGATTTGAAAAAATAGACTGGGTGGGCCCTGGACACTGGACCCAGGCGGCGCATGAGGCGCAGTACCTCACGCCGCACGGTCCGGCACAGGTCATAGATTTCCTCCA | | 4 |
| wt-73050-73170 | 4888 | GCGACCAGGGGGCCCCCTTGATCTTTAGATCCAGGTCCAAGACCAGGTTGCAGACCGGAAGCCGGGGATTAAAGTATTCATGCCGGGAGACAAAGAGCTGCTCGCTCAGGCTGTTCTGTG | | 4 |
| wt-73170-73290 | 4889 | AATAGTACACTGGGGTGTAGGAGAGGGCCCTGGTGAGACACGTGTCTGGGAGGCGGCAGTTGGTCGGGGTGGAGACGACCTCCGCCAGGTGGGATGAGAAGGGGTCAGCGGCTGTCATTA | | 4 |
| wt-73290-73410 | 4890 | CAAAGTAGTGCCTGTCTGCAAAATGGCAGAGGAAGACCGGTAGCCGCTGCACCCTTCGAAGGACGGTGGGTGGGAGGAATTGTTCCTTGGGATTCCACTGGCCCCGGCAGGTGGCCTGGC | | 4 |
| wt-73410-73530 | 4891 | CGGCCAAGCATAGAAACCCTTGAAGCGTGGGGGGGTATGTGGGACCCTCATCCGCGTGCCAGCGCGCGAGCTCCACCAGCTCCCGGGCCACGTCCACACTGAGCCCGGCCCAGGCCCGCA | | 8 |
| wt-73530-73650 | 4892 | TGAGTCCGTCATCGGGGTCGGGGTCCCACGTGTATGGGGCCGGGGGCTCCATGCGGATTTTCAGCTGCTGGACACGCACATGCTCAGCCAGGTAAGTCTCCCGGGTGAAGTAGGTGCGCA | | 4 |
| wt-73650-73770 | 4893 | TGTGCTCCGCAAAGCCCCTGTCCAGGAGCGAGGGGAGCACGACGCCCCCCGAAGGCAGACACCCAATTTCTCCCATGCTCGTTAACTGAGAGTATCGCTTAAAGGTTCCCTCGTTGAAGC | | 4 |
| wt-73770-73890 | 4894 | ACTGTGCGTGGGCCAAATAGACGTAGCGCACGAGATCGGCCGAGGCCAGGGGAAGGCGCCCCCTGTAGGCGTCTATCGTCCTTGCCACAGCGCGGATCTCTCGCGAGTCCCGCCGCAGTT | | 8 |
| wt-73890-74010 | 4895 | TCTCGTGTGCAAAGTGGGCAAAAGCCTCGGTCTGCTCCGCCCATGCCGAGGAGCCAAAGACCTCCCCCAGCTCGGCCAGGGACGTGACGGCGGCCAGGCTCTGACCAGACTCGGAAGTAA | | 4 |
| wt-74010-74130 | 4896 | ATAGCTCCGTGAGGTGCGCCAGGGTCTCAATCGTACAAGGAATGCCCCAAAAATAGTAAGCAGCCGTGACTAGCACGAACTGGGCCTCGTGGGAGCCAAAGGTGCTAATGAACCACCTGG | | 4 |
| wt-74130-74250 | 4897 | CCGAGATGTTAACGCGGTAGATGCGGCGCAGACAGCCCACGATCTTGGGACGCAGCCACGCCACGCGGCCTCTGGCATCCCCCTGTGGCTGTTTCTTAGCGCTCAGTGTGAGCAGTTCCA | | 4 |
| wt-74250-74370 | 4898 | CGAGGGGCGTGAGCGAGCGCAGGGCCCCCGCGCGATCTAGGTAGGTGGATAGACGGTCCGCGGTGAGCGGCGTGAGGCCGCGCAGGAAGGGGAAGGCCTCCTCCGCCGGCAGGTGCAGCG | | 8 |
| wt-74370-74490 | 4899 | TCAGAACCAGGCCGCAGCGGCTCTGTGAGGTCAGCCGCTTCTTGGGCAGGTGAAGCTGCAGTTCCACGAGAGAACCCGCCACGTGGTGGAGGGGCGAGGCGTTGTGGCACAAACAAAACA | | 4 |
| wt-74490-74610 | 4900 | GGCGGAAGCCCTCGTCAGGCCGCGAGAGGATGGCATCGAGGATGGCCTCCGCAATGTCAGTGTTTGAGGCCACAAGGGCCTTGATGACGACGGGGGCGGACATTATTTAAGACCGGGAGG | | 4 |
| wt-74610-74730 | 4901 | CCCCAACGGCGGGCTAAACAGAACGATGGCCTTCTATCTCCCAGACTGGTCGTGCTGCGGGCTCTGGCTCTTTGGCCGGCCCAGGAATAGATACAGCCAGCTCCCTGAGGAGCCGGAGAC | | 4 |
| wt-74730-74850 | 4902 | CTTTGAGTGCCCGGACCGCTGGCGAGCCGAGATAGATCTGGGCCTGCCCCCTGGTGTGCAGGTGGGAGATTTGCTAAGAAATGAGCAGACGATGGGCTCACTGAGACAGGTTTATTTGCT | | 4 |
| wt-74850-74970 | 4903 | CGCTGTTCAAGCCAATAGCATCACGGATCACCTGAAGCGCTTTGACGCCGTCCGCGTCCCTGAGAGCTGTCGTGGGGTGGTGGAGGCCCAGGTGGCCAAGCTTGAGGCCGTGCGCTCAGT | | 4 |

| | | | |
|---|---|---|---|
| wt-74970-75090 | 4904 | CATCTGGAATACCATGATCTCTCTGGCTGTAAGCGGCATCGAGATGGACGAGAATGGGCTCAAGGCCCTGCTGGACAAACAGGCTGGCGACAGCCTGGCCCTGATGGAGATGGAGAAGGT | 4 |
| wt-75090-75210 | 4905 | GGCCACGGCGCTCAAGATGGACGAGACCGGTGCCTGGGCGCAAGAGATCTCGGCCGTTGTCTCATCGGTGACCGCCCCCTCAGCCTCGGCCCCTTTCATCAACTCCGCCTTTGAGCCCGA | 8 |
| wt-75210-75330 | 4906 | GGTGCCCACCCCCGTCCTTGCACCGCCTCCCGTGGTGCGGCAGCCGGAGCACTCTGGGCCCACGGAGCTCGCGTTAACGTAGCAACCAGACTCCACACCAAATAAACATTTTATTGGTAA | 4 |
| wt-75330-75450 | 4907 | AACAAGGGATATGAAGGTGTCATTGACCCGAGGATCCAAACCCCCTCCCCTGTCTCCCCTCGAGCGCCTCGCTCAGCCCACTATCACCCATGGCCAGGCTCGGCACCTCCTCGAAGGTGC | 4 |
| wt-75450-75570 | 4908 | AGCTGGCCCACCTAAAGAGAGATCTGGGGCCAAGGACCCCCGCGTCACTGTGGGGGCTGTAGAAGGAGGTGAGGTGGTGCTTGTGAAGGTAAACAAGCTGACAGAAGCGCCGGTACTTGT | 4 |
| wt-75570-75690 | 4909 | TAAGGAACACGGTCTGGTCACTAAAGTTGGTCAGGCTGACGTCCACCCCACCCCGGCGCCACCTGCAGGGCTTCACTAGAATACCCTGCATGGCCAGGCCCGACCTGCCAAAGATTGTCG | 4 |
| wt-75810-75930 | 4911 | GGTGGTTGATGGGACCCTTGTCCTCCTCCATCTGGGGGGTGGCATATCTGAAGGCAGCCAGGTGGATTTTGAGCTCCGATGGACGCAGCGTGGAGTTGTAGCGCCGCTGATTCTGGAGGA | 4 |
| wt-75930-76050 | 4912 | TTAGCCGGAGTTCCCCCGTGTAGCCGGGATCGATGATGCCAACATGAGACGTGACCGGACGGGAGGTGCTGCCCCACAGCATGAGCCCATGACCCTCGGGTGGGCGGGCATAGAGGCCTA | 4 |
| wt-76050-76170 | 4913 | GGTCCACAGTTGTGGTCTTCATCGGGCGCAGCAGGATGGTGGTCTTGTTGACCAAGGTGAGCCGCCCTACACTAGCCTGCTGGAGCAACAGCTTGTCATTCTGGAAGGCGTAGCGTATGT | 4 |
| wt-76170-76290 | 4914 | GTGGACAGGCCTCCATGGTGATGATCTAACAGACAGGGACGGCGGCGCTATATATAAGAGCCCAAGACCCGGCTCTCTTTACTGCGAAATGGGGAAGGTCCTAAGAAAGCCGTTTGCAAA | 4 |
| wt-76290-76410 | 4915 | GGCTGTGCCACTGCTCTTCCTCGCCGCCACCTGGCTTCTGACCGGGGTGCTGCCGGCCGGCGCTTCCAGTCCCACAAACGCGGCGGCGGCTTCCCTGACTGAAGCCCAGGACCAGTTCTA | 8 |
| wt-76410-76530 | 4916 | CTCCTACACATGTAATGCGGACACATTCTCGCCTTCTTTGACCAGCTTTGCCTCCATCTGGGCACTTCTGACGCTTGTCTTAGTCATTATAGCCTCAGCCATCTACCTGATGTACGTCTG | 4 |
| wt-76530-76650 | 4917 | CTTTAACAAGTTTGTGAACACGCTGCTGACGGATTAGATGGGGATATTTAAAAGGGGCAGCAATCTCGGCTGTTTGTACTTCTTCTCTGCTCGTTAAACCAATAGCATGTCAGCTCCACG | 4 |
| wt-76650-76770 | 4918 | CAAAGTCAGATTGCCTTCTGTTAAGGCTGTTGACATGAGCATGGAAGACATGGCCGCCCGCCTGGCTCGCCTGGAGTCTGAGAATAAGGCTCTGAAGCAACAGGTCCTCAGAGGGGGTGC | 4 |
| wt-76770-76890 | 4919 | CTGTGCCTCGTCTACCTCTGTTCCTTCTGCTCCAGTGCCTCCGCCTGAGCCGCTTACAGCTCGACAGCGAGAGGTAATGATTACGCAGGCCACGGGCCGTTTGGCGTCTCAGGCTATGAA | 4 |
| wt-76890-77010 | 4920 | GAAGATTGAAGACAAGGTTCGGAAATCTGTTGACGGTGTAACTACCCGCAATGAAATGGAAAATATATTGCAAAATCTGACCCTCCGCATTCAAGTATCTATGTTGGGTGCAAAAGGCCA | 4 |
| wt-77010-77130 | 4921 | ACCCAGCCCTGGTGAGGGAACACGACCACGAGAATCAAACGACCCCAACGCCACCCGACGTGCCCGCTCCCGCTCCCGGGGACGTGAAGCAAAGAAAGTGCAAATTTCTGATTAATAAAT | 4 |
| wt-77130-77250 | 4922 | TTTTATTGACTTTATACATAGGTCTCGGCGTCATCATATGGTGGGGTGGTGTAGGTATGGGATGTAGACAAGTTACGCCTAAAGGCGCAGTCCGCCATGACCAGCAGCAGCAGAAGGGTC | 4 |
| wt-77250-77370 | 4923 | AGCACAGCCAGAGAGGCCCACTGCAGTACTAGCATGGAGAGGTTTGAGAATCTGGGCTGGGACGTTGGCGGGACTGGCACGGTGGCTTGGGCTGTGGTAACCGGTGGGCTCGTAAAAGTC | 4 |
| wt-77370-77490 | 4924 | CAGCGGGGCCGCAGTTTGCTAGAAGTGCTGGGAGGTAGATAGGTGGTCGCATTGTATCTCGGTCTTGGCGTAGTTGAATCACCGCCGTAATCTGTGGTGGGCTCTGTACTTGTCCGGGCT | 4 |
| wt-77490-77610 | 4925 | CCATGTCCTGTGGTGTGCTTTCCACCGGTGGTAGAATTGGCCTTTCCACCTGTTGAGGTGACCGTGGGAACCGCCGTCTTTTGGCCACTGGGGGCCTGGGGCGACGTTGCATTTTGGGGG | 4 |
| wt-77610-77730 | 4926 | GGCGTGCCTTTGGTGACATTAACCTCCCCCGGTTTTGTGGATGTGGAACTGTTTCCAGGGCCTGACGCTTGGCTGGTGGTGCCTGGGCGGGGTGCTGGCGAACTGGTGGACACATGATGT | 4 |
| wt-77730-77850 | 4927 | GTGCTGATAGAGGCTGGTGTCACCTGTGTTATATTTTCACCACCTGTTGGGTGAGCGGAGGTTAGTAAAGGCATATGTGACGTTGAATTGTCACTGGTGGAGGGGCTGAGTGTCTCTGGG | 4 |
| wt-77850-77970 | 4928 | TTTGAACTGGGTCTCAGTGACATGGAAGAGGTTGAACTTGAAGTTATGTTATGTTGGCCTGTGGTAACAGCACTGGTTGCATTTTTTGGTTGGCTGGTAACTACTGGGGTGGGACTTGTT | 4 |
| wt-77970-78090 | 4929 | CCTCCTAAGGTGTGGTTGGTGGCATTTGCCTGTGGACTTGTTTCTCCCACAGTAGGGCCGGTGGCATTTGGGGTTGGGGTAGTCACTGCTGAGGTGGGGCTTGTTTTTCCCAAGGTGGGG | 4 |
| wt-78090-78210 | 4930 | CTGGTGGCATTTGGGGTTGGGGTAGTCACTGCTGAGGTGGGGCTTGTTTTTCCCAAGGTGGGGCTGGTGGCATTTGGGGTTGGGGTAGTCACTGCTGAGGTAGGACTTGTTTTTCCCAAG | 4 |
| wt-78210-78330 | 4931 | GTGGGGCTGGTGGCATTTGGGGTTGGGGTAGTCACTGCTGGGGTGGGGCTGGTGGCATTTGGGGTTGGGGTAGTCACTGCTGGGGTGGGGCTGGTGGCATTTGGGGTTGGGGTAGTCACT | 4 |
| wt-78330-78450 | 4932 | GGTGAGGTGGAGCTGGTCATGTCGGGGGCCTTACTTTCTGTGCCGTTGTCCCATGGAGATGGACTTGGTGTCACCGGTGATGCGCCTGACGTTGTGCCGGCTGGTGTTGGGCTGGTGACA | 4 |
| wt-78450-78570 | 4933 | TCCGCGGTGGATACAGTGGGGCCTGTGCTTGCAGGTGCGGTGAGGTTGGTAGGCACGTGAGTAGAGCTGGGTAGACCTGTCGTTGTATTGGGATCAGCAAATCCAGTTGTATTCAAGGTA | 4 |
| wt-78570-78690 | 4934 | GGGGAGGTGGTGGTGCTCTCGGGTGCCTTGGAGAATATAACCTTGTGGGTTGTTGTGGTGGCATTGGTAGCCGTTCGTGTGATAATGAGTGTCTTGGGGGCCGTGCCAAGACCCGAGACA | 4 |
| wt-78690-78810 | 4935 | GTAATGTCAAATGTCCGATTGCTCGCAAATGCACCAGAAATATTTTCACAACCCGAAGGTGTCCCCGAGGTGAGAGTCCATTTGCACTTAAAGTCAGTTTCAGTGTTGTTTGGCCAGGCC | 4 |
| wt-78810-78930 | 4936 | CAAAAGGCAGTCACTGTAACATTTGGCGAGTTTGCGTCCTCAGAAGTGACCATTGGCACTGAATAGGTAGCATTGTCACCCACATATGTGATGTCTGTGGTGTTTGTCGGCATGTCCTGT | 4 |
| wt-78930-79050 | 4937 | GAAGCTGGAATCTCATCAGAGAACACAATGTTGGACTGAATGCAGTAATCTCCCCCGCTCGCCTTCGGTCCATTCCCAGAGTAAAACACGTACAGGATACTGTTATTGCCAAGAAATCGT | 4 |
| wt-79050-79170 | 4938 | GACACTGGACGTGGTGTCAGACGCAGGCTGTATGCATACCCTGTACCAGGTATTGGGGTGGCCACGGGACTCGTTGATGTGAGAATTCCGCCGCTGGGAACATGGCTCTCGTATCCACTG | 4 |
| wt-79170-79290 | 4939 | CAGGTGATGTTAAATTTGTTGTCTCCGGGCAGAACTTGTGAAATTTCGCCATCCTCCATAATACACTCAATATCTATCTCATTACCGAGCATTTCTGTTTTTACGCTGAAATTCGAGTCT | 4 |
| wt-79290-79410 | 4940 | TGAGCTGACGTTGGCAAACTTAAGGGTAGCGTGACATCCAGCCCCTGTGCCCTCACTACTGCCGTTATATTGGTAGAATTACAGTTATCCCACTTTATGTATGGCACTGTTTCTGGTATC | 4 |
| wt-79410-79530 | 4941 | AGGTACACGGGGTTTTGCATTTCTGCATGGTGGCACCACATGGTTCCAAACACATCTTGAAAGTAGACATCTACAGATTCCAGGCTTACTTGTTGCTCCTCTCCGGTGGTGACGTTAATT | 4 |
| wt-79530-79650 | 4942 | GGAAGCTTCTTAGACCGCATAGTTAGAGCCAATTCTCCTGCACCAAGGAGCTCCAGTAGAAAGAGATTGGTGGCATTTTCTGAGCCACCAAATGCACCTCGAGGTTGGTAGACAGCCTTC | 4 |
| wt-79650-79770 | 4943 | GTATGGGGTGTCAGCTGGCCAAAGTCAAGATCAAGTTGATGCTTTTTGCCCCCGACATCGAAATTGATAGTTACATTGACATCTGCCGTGCAAACATTGCATGTGGGGTAAAATGGGAAT | 4 |
| wt-79770-79890 | 4944 | TCCGGAATCTCAACATTGAAAAAACCAGGATCTTCACCCGTGAGATGGATCAGGCTCTGGATGGTGTACTGACACACAAGCAAGGCTGCCTCCATTGTCTCGGCACCGATTTCTAGGCAG | 4 |
| wt-79890-80010 | 4945 | CATCCTCTTTAATAGGTACAAGGGGGGTGCGGTGTTGGTGAGTCACACTTTTGTTGCAGACAAAATGGACAAGGACAGGCCGGGTCCCCCGGCCCTGGATGACAACATGGAAGAAGAAGT | 4 |
| wt-80010-80130 | 4946 | CCCATCTACCTCGGTTGTGCAGGAACAGGTATCGGCGGGAGATTGGGAAAATGTCCTCATAGAGTTATCAGATAGCAGCTCAGAAAAGGAAGCAGAAGATGCCCACCTGGAGCCGGCCCA | 4 |
| wt-80130-80250 | 4947 | GAAGGGTACGAAGAGAAAGCGGGTCGATCATGATGCCGGTGGGTCAGCTCCAGCACGACCCATGCTCCCACCCCAGCCGGATCTCCCTGGGCGAGAAGCCATTCTCCGCAGGTTTCCACT | 4 |
| wt-80250-80370 | 4948 | AGATCTAAGAACACTTCTTCAAGCGATTGGAGCCGCGGCTACGGTGAGCATCCCTATGGCCTAAGTGTGTGATGTGTGTTTTTACCCATCACACAACAACAAGGTAAGTAATTTGTTGCC | 4 |
| wt-80370-80490 | 4949 | GTTGGTTTCAGCGCATCGACACACGAGCCATAGACCAGTTTTTCGGATCCCAGATTTCAAATACCGAGATGTACATAATGTATGCCATGGCCATTCGACAGGCCATTAGAGATCGTCGGA | 4 |
| wt-80490-80610 | 4950 | GAAATCCAGCTTCTCGTAGAGATCAGGCCAAATGGAGACTGCAAACCCTGGCCGCCGGATGGCCTATGGGTTACCAGGCATACAGCAGCTGGATGTACAGCTACACCGATCACCAGACGA | 4 |
| wt-80610-80730 | 4951 | CTCCCACATTCGTACATCTCCAGGCGACACTTGGGTGCACAGGTGGCCGTAGGTGTCACGTGACCTTTTCCGCCGGCACCTTTAAGCTGCCGCGATGTACCCCCGGGGATCGCCAGTGGT | 4 |
| wt-80730-80850 | 4952 | TGTATGTTCAGAGCTCCGTGGGTAACATTGTACAGAGCTGTAATCCCCGCTACAGTATTTTCTTTGACTATATGGCTATACACAGGAGCCTCACGAAAATCTGGGAGGAAGTTTTAACAC | 4 |
| wt-80850-80970 | 4953 | CTGACCAGCGTGTTTCATTTATGGAATTCCTAGGATTTTTGCAGAGAACGGATTTGTCCTATATCAAGAGCTTTGTCAGCGATGCCCTGGGCACCACTAGTATCCAAACACCGTGGATCG | 4 |
| wt-80970-81090 | 4954 | ATGACAATCCTAGCACGGAGACGGCACAGGCTTGGAATGCCGGCTTTCTCCGGGGTCGTGCGTATGGGATAGACTTGCTTAGAACTGAAGGGGAACATGTCGAAGGTGCTACCGGTGAAA | 4 |
| wt-81090-81210 | 4955 | CGCGAGAAGAAAGTGAGGACACGGAGAGCGATGGAGATGATGAAGATCTTCCTTGTATAGTGTCCAGAGGTGGACCTAAGGTCAAACGACCCCCTATATTTATAAGACGTCTGCACAGGT | 4 |

| | | | |
|---|---|---|---|
| wt-81210-81330 | 4956 | TGCTGCTGATGAGAGCGGGCAAACGAACAGAACAGGGCAAGGAGGTACTGGAAAAGGCCCGTGGGAGCACTTATGGCACACCTAGGCCGCCTGTTCCGAAACCAAGACCAGAGGTCCCAC | 4 |
| wt-81330-81450 | 4957 | AAAGCGACGAGACAGCTACCAGTCACGGGTCGGCGCAAGTCCCAGAACCCCCAACCATTCACCTAGCAGCTCAGGGAATGGCATACCCATTACATGAACAACACGGCATGGCCCCGTGTC | 4 |
| wt-81570-81690 | 4959 | TCCGGCCCTGGGAGCCATCCCTGACACAGGCTGCGGGGCAGGCCTTTGCACCCGTTAGACCACAACACATGCCAGTAGAACCCGTCCCTGTCCCGACAGTGGCACTTGAGCGACCAGTTT | 4 |
| wt-81690-81810 | 4960 | ACCCCAAGCCAGTTCGTCCGGCACCTCCTAAGATTGCTATGCAGGGCCCCGGGGAAACTTCTGGCATTAGACGCGCGCGGGAGCGTTGGAGGCCCGCACCTTGGACGCCAAATCCACCCC | 4 |
| wt-81810-81930 | 4961 | GTTCTCCCAGTCAGATGTCCGTGCGTGACCGTCTGGCTCGTTTGCGTGCTGAGGCACAGGTCAAACAGGCTAGTGTTGAGGTGCAGCCCCCCCAGTTGACCCAAGTATCCCCTCAGCAAC | 4 |
| wt-81930-82050 | 4962 | CAATGGAGGGGCCGTTGGTACCAGAGCAGCAGATGTTCCCTGGTGCCCCCTTTAGCCAGGTTGCTGATGTGGTCCGGGCACCTGGGGTACCGGCGATGCAGCCACAGTACTTTGACCTCC | 4 |
| wt-82050-82170 | 4963 | CCTTAATTCAACCCATTAGCCAGGGGGCACCCGTGGCCCCGTTGAGGGCTAGTATGGGCCCGGTACCTCCGGTACCGGCAACACAGCCACAGTATTTTGACATCCCCTTAACTGAACCCA | 4 |
| wt-82170-82290 | 4964 | TTAACCAGGGGGCATCCGCGGCCCATTTTCTCCCTCAGCAACCGATGGAGGGGCCGTTGGTACCTGAGCAGTGGATGTTCCCAGGTGCCGCCCTGAGCCAGAGTGTTAGGCCAGGGGTAG | 4 |
| wt-82290-82410 | 4965 | CGCAGTCACAATATTTTGACCTCCCCTTAACTCAACCCATTAACCATGGGGCACCCGCAGCCCATTTCCTCCATCAGCCACCAATGGAGGGGCCGTGGGTACCCGAGCAGTGGATGTTCC | 4 |
| wt-82410-82530 | 4966 | AAGGTGCCCCCCCTAGCCAAGGCACTGACGTGGTCCAACATCAGCTGGATGCTTTGGGGTATACACTCCATGGTCTTAACCATCCCGGGGTTCCCGTGTCTCCTGCCGTTAACCAATATC | 4 |
| wt-82530-82650 | 4967 | ATCTCAGCCAGGCTGCCTTTGGGTTACCTATTGATGAGGATGAGAGTGGCGAGGGGTCCGATACCTCCGAGCCGTGTGAAGCTCTTGATTTGTCAATCCATGGCAGGCCCTGCCCTCAGG | 4 |
| wt-82770-82890 | 4969 | AAAATGTCACAGGCCCTGAAACTAGAAGGGTGGTGGTGTCAGCTGTTGTTCACATGTGTCAGGATGACGAGTTTCCGGATCTACAAGATCCTCCAGATGAGGCCTAAGCAAAGGTGTAGA | 4 |
| wt-82890-83010 | 4970 | AGTGTGTCCCCCTCCATTCCACCCACTGATAATACGCCCGACAATAAAGTTGATGATATTGAATTCCACACCTGCTTGTGTTTGTGATTTTATTTCATATTCCATGAGAGAGACCTCGCA | 4 |
| wt-83010-83130 | 4971 | TATTTGCAGAAGGGTCACTGAAACATCTTATCTTAAAACAGTTACACCTGAATAATGAAGAAAGCGTGGCTCAGCAGAGCACAGCAAGCCGATGCCGGGGGGGCATCTGGCTCCGAGGAC | 4 |
| wt-83130-83250 | 4972 | CCACCAGATTATGGAGATCAAGGTAATGTGACACAGGTGGGATCTGAGCCTATTTCACCTGAGATTGGCCCCTTTGAACTCTCTGCGGCCAGTGAGGATGATCCTCAATCTGGGCCAGTG | 4 |
| wt-83250-83370 | 4973 | GAAGAGAATTTAGATGCCGCTGCAAGAGAGGAAGAGGAACCTCATGAGCAGGAGCACAATGGTGGTGACGATCCCTTGGATGTCCATACTCGCCAGCCTAGATTTGTGGATGTGAACCCA | 4 |
| wt-83370-83490 | 4974 | ACGCAGGCTCCAGTGATCCAACTAGTCCACGCTGTCTATGATTCCATGTTGGTAAGAGGCACCTAGAACATTTCCAGATGTTTCGCTTGGATTTTTTGGCCAGTCTTAATTGATTGTCAT | 4 |
| wt-83490-83610 | 4975 | TGGTTTCAGCAATCGGACCTCCGGCCCCTAGGCAGTTTATTCCTTGAGCAAAACCTGAACATCGAAGAATTTATATGGATGTGCATGACCGTGCGTCACAGATGTCAGGCCATCAGAAAA | 4 |
| wt-83610-83730 | 4976 | AAACCATTACCAATTGTTAAGCAGAGGCGTTGGAAGCTCCTGTCATCTTGCAGATCCTGGCGTATGGGTTACCGCACGCATAACCTCAAAGTAAACAGTTTTGAGTCAGGGGGGGACAAT | 4 |
| wt-83730-83850 | 4977 | GTCCACCCGGTCCTTGTGACTGCTACGCTAGGATGTGATGAGGGCACGCGGCATGCAACAACGTACAGTGCTGGCATTGTACAGATACCACGAATATCAGACCAAAACCAAAAGATCGAA | 4 |
| wt-83850-83970 | 4978 | ACAGCCTTCCTGATGGCACGTCGTGCTAGGTCACTTTCGGCAGAAAGATATACTTTGTTCTTTGATTTAGTATCCTCCGGAAACACCCTGTATGCTATATGGATTGGGCTGGGCACGAAA | 4 |
| wt-83970-84090 | 4979 | AACCGAGTTTCATTTATTGAGTTTGTAGGATGGTTATGCAAGAAGGACCACACTCATATACGCGAATGGTTCCGCCAGTGCACCGGGAGACCCAAAGCAGCCAAGCCATGGTTAAGAGCG | 4 |
| wt-84090-84210 | 4980 | CATCCTGTCGCCATTCCTTATGATGATCCGTTAACAAACGAGGAGATTGATCTGGCCTATGCCCGCGGGCAGGCCATGAATATTGAGGCTCCTAGACTGCCAGATGATCCTATAATTGTT | 4 |
| wt-84210-84330 | 4981 | GAGGATGACGACGAAAGTGAGGAAATTGAAGCTGAAAGCGACGAGGAGGAAGACAAGAGTGGAATGGAATCTCTTAAAAATATACCGCAAACACTGCCGTACAATCCAACAGTATACGGC | 4 |
| wt-84330-84450 | 4982 | AGGCCCGCGGTGTTTGACCGAAAGTCAGATGCAAAATCAACCAAAAAATGCAGGGCCATAGTAACTGACTTTAGTGTAATCAAGGCCATTGAAGAGGAACACAGAAAGAAGAAGGCAGCC | 4 |
| wt-84450-84570 | 4983 | AGAACAGAGCAGCCAAGAGCCACGCCTGAATCCCAGGCCCCCACAGTGGTCCTCCAGCGACCACCCACGCAGCAAGAGCCTGGCCCCGTCGGCCCACTGAGTGTCCAGGCTCGGCTGGAG | 8 |
| wt-84570-84690 | 4984 | CCATGGCAACCTTTGCCTGGGCCCCAAGTGACAGCAGTTCTACTTCACGAAGAATCCATGCAGGGTGTCCAAGTACATGGTTCGATGCTAGACCTTCTTGAAAAAGACGATGAAGTCATG | 4 |
| wt-84690-84810 | 4985 | GAGCAGAGGGTTATGGCAACCCTACTGCCACCAGTACCACAACAGCCCCGGGCTGGCAGAAGAGGCCCTTGTGTCTTCACCGGTGACCTAGGCATAGAGAGTGATGAGCCCGCTTCGACA | 4 |
| wt-84810-84930 | 4986 | GAGCCGGTTCATGATCAGCTACTGCCTGCCCCAGGACCTGACCCTCTTGAAATCCAACCACTAACGTCCCCCACCACGTCTCAACTTAGCAGTTCGGCACCAAGCTGCGCACAAACTCCA | 4 |
| wt-84930-85050 | 4987 | TGGCCGGTGGTTCAGCCAAGTCAGACTCCAGATGACCCAACGAAACAGTCCCGGCCACCGGAAACAGCTGCCCCACGCCAGTGGCCAATGCCCCTGCGACCTATCCCTATGCGCCCCTTG | 4 |
| wt-85050-85170 | 4988 | CGGATGCAGCCAATCCCATTTAATCATCCAGTGGGACCCACTCCCCATCAGACACCTCAAGTGGAAATAACACCATATAAGCCCACTTGGGCTCAGATAGGGCACATTCCATATCAGCCT | 4 |
| wt-85170-85290 | 4989 | ACACCAACGGGTCCTGCTACCATGCTGTTGCGCCAGTGGGCACCCGCCACCATGCAGACACCACCGAGAGCGCCCACTCCCATGTCACCACCTGAGGTGCCACCCGTTCCCCGGCAGAGG | 8 |
| wt-85290-85410 | 4990 | CCTCGGGGGGCGCCCACTCCCACGCCACCTCCTCAGGTGCCGCCCGTTCCCCGGCAGAGGCCTCGGGGGGCGCCCACTCCCACGCCACCTCCTCAGGTGCTGCCCACTCCCATGCAGCTG | 12 |
| wt-85410-85530 | 4991 | GCACTAAGGGCTCCTGCTGGTCAGCAGGGGCCGACAAAGCAAATTTTGCGCCAATTGTTAACGGGGGGCGTCAAGAAAGGGAGACCATCACTTAAGTTACAGGCCGCCCTTGAGCGTCAA | 4 |
| wt-85530-85650 | 4992 | GCCGCTGCGGGCTGGCAGCCTTCACCAGGGTCCGGCACCAGTGACAAGATTGTGCAGGCGCCTATTTTCTACCCACCCGTTTTGCAGCCCATACAGGTTATGGGGCAAGGGGGTTCCCCA | 4 |
| wt-85650-85770 | 4993 | ACGGCCATGGCCGCCTCAGCGGTGACACAGGCACCCACGGAATATACCAGGGAAAGGAGGGGAGTGGGGCCTATGCCTCCCACCGATATTCCGCCGTCTAAACGAGCGAAGATCGAGGCC | 4 |
| wt-85770-85890 | 4994 | TATACAGAGCCCGAGATGCCGCACGGGGGGGCCTCGCACTCTCCCGTCGTTATCTTGGAGAATGTCGGCCAGGGGCAACAGCAGACTCTGGAGTGCGGAGGAACTGCTAAACAGGAAAGG | 4 |
| wt-85890-86010 | 4995 | GACATGTTGGGGCTGGGGGACATTGCAGTTTCTTCCCCTTCCTCTTCTGAAACATCGAACGATGAGTGATTTCGCCCATGTAACAAGAACTGGGATGAACCCTGGGGCAACAGACTGCGG | 4 |
| wt-86010-86130 | 4996 | GGAGGAGGGGGGCAGTGATAAGTCATGACAATTTTAGATGAGGTAGAAATTTTGCATATTTTCAGACCCACCATGGAATCATTTGAAGGACAGGGGGACTCTAGACAGTCACCCGACAAT | 4 |
| wt-86130-86250 | 4997 | GAGCGGGGAGATAATGTACAGACTACCGGCGAGCATGATCAGGACCCTGGGCCGGGGCCTCCATCCAGTGGGGCTTCTGAGAGATTGGTACCAGAAGAGTCATACTCAAGAGATCAGCAA | 4 |
| wt-86250-86370 | 4998 | CCTTGGGGGCAAAGCAGGGGTGATGAAAACAGAGGCTGGATGCAGCGCATCAGGCGAAGGCGGAGAAGACGGGCTGCCTTGTCCGGCCATCTTTTAGACACGGAAGACAATGTGCCGCCA | 4 |
| wt-86370-86490 | 4999 | TGGTTGCCTCCACACGACATCACACCATATACCGCAAGGAATATCAGGGATGCTGCCTGCCGGGCTGTCAAGGTGAGTATGCCTCTAACTGGGTTCATGGGGGCCATCTAAGGCCCACGT | 4 |
| wt-86490-86610 | 5000 | GTGACCCATGTTTCCATTAATTTTAGCAATCGCACCTGCAAGCGCTATCAAACCTGATACTCGATAGTGGGTTAGACACACAACACATCTTGTGCTTCGTGATGGCAGCCAGGCAGCGTC | 4 |
| wt-86610-86730 | 5001 | TTCAGGACATTCGACGTGGACCCTTGGTTGCGGAGGGCGGTGTGGGTTGGCGACATTGGCTTCTAACATCTCCCAGCCAATCCTGGCCCATGGGATATCGTACAGCAACACTACGCACAT | 4 |
| wt-86730-86850 | 5002 | TAACTCCCGTGCCTAACAGGGTTGGGGCTGACAGCATCATGTTAACTGCCACATTTGGATGCCAAAATGCGGCACGAACTCTAAACACCTTCTCTGCCACCGTGTGGACACCACCCCATG | 4 |
| wt-86850-86970 | 5003 | CTGGACCAAGAGAGCAAGAAAGATACGCTCGGGAAGCCGAGGTACGCTTCCTTCGTGGTAAATGGCAGAGGCGGTACCGAAGAATCTATGATTTGATAGAACTGTGTGGCTCTCTGCACC | 4 |
| wt-86970-87090 | 5004 | ACATCTGGCAAAACTTGCTCCAGACCGAGGAGAACCTTTTAGATTTCGTGCGTTTCATGGGTGTCATGTCCAGCTGCAATAATCCAGCTGTGAATTACTGGTTTCACAAGACAATCGGAA | 4 |
| wt-87090-87210 | 5005 | ACTTTAAGCCATATTACCCGTGGAATGCACCACCTAATGAAAATCCATATCACGCGCGGAGAGGCATAAAAGAACACGTAATCCAGAACGCATTTCGAAAGGCACAAATACAGGGTTTAT | 4 |
| wt-87210-87330 | 5006 | CAATGTTAGCAACGGGAGGTGAACCCAGAGGTGATGCTACTAGTGAAACGAGCAGTGATGAGGACACCGGTAGACAGGGTTCGGACGTGGAGCTAGAGTCCTCGGACGATGAGCTGCCAT | 4 |
| wt-87330-87450 | 5007 | ATATCGATCCCAATATGGAGCCGGTTCAGCAGAGGCCCGTCATGTTTGTGAGCCGTGTGCCTGCAAAGAAACCGAGGAAACTGCCTTGGCCCACGCCCAAGACGCACCCAGTGAAGCGCA | 4 |
| wt-87450-87570 | 5008 | CAAATGTTAAGACCTCTGATAGATCTGATAAGGCAGAAGCACAAAGCACCCCTGAAAGGCCGGGCCCTTCCGAACAATCATCAGTGACCGTGGAGCCCGCCCACCCGACCCCGGTGGAGA | 4 |

| | | | |
|---|---|---|---|
| wt-87570-87690 | 5009 | TGCCAATGGTGATTCTCCATCAACCACCTCCAGTGCCCAAACCGGTTCCAGTCAAGCCTACGCCACCGCCTTCCCGTAGGAGAAGGGGAGCGTGTGTTGTGTACGACGATGATGTCATAG | 4 |
| wt-87690-87810 | 5010 | AGGTGATTGATGTTGAAACCACCGAAGATTCATCGTCAGTGTCACAGCCAAATAAGCCACATCGGAAACATCAAGACGGCTTTCAACGTTCAGGCCGACGTCAAAAACGAGCCGCGCCTC | 4 |
| wt-87810-87930 | 5011 | CCACCGTGAGTCCTTCGGATACTGGGCCTCCTGCCGTGGGGCCTCCTGCCGCGGGGCCTCCTGCCGCGGGGCCTCCTGCCGCGGGGCCTCCTGCCGCGGGGCCTCCTGCCGCGGGGCCTC | 12 |
| wt-87930-88050 | 5012 | CTGCCGCGGGGCCTCGCATACTGGCGCCTCTTTCCGCTGGGCCTCCTGCCGCGGGGCCTCACATAGTGACGCCTCCTTCCGCCCGGCCTCGTATAATGGCGCCTCCCGTCGTACGTATGT | 8 |
| wt-88050-88170 | 5013 | TTATGAGGGAGCGACAGCTCCCCCAGTCCACCGGCCGTAAACCTCAGTGCTTCTGGGAAATGCGGGCTGGTCGTGAAATTACACAAATGCAACAAGAACCAAGTTCACACCTGCAGTCCG | 4 |
| wt-88170-88290 | 5014 | CCACTCAGCCTACAACGCCTCGCCCATCATGGGCCCCATCAGTCTGCGCCCTCTCGGTGATGGATGCTGGTAAGGCCCAGCCCATAGAAAGTTCACACTTGAGTTCCATGTCGCCCACAC | 4 |
| wt-88290-88410 | 5015 | AGCCGATATCGCACGAAGAACAACCCCGGTATGAGGATCCTGACGCTCCTCTGGATTTAAGTTTACATCCAGACGTTGCTGCTCAACCAGCTCCCCAGGCTCCATACCAGGGATACCAGG | 4 |
| wt-88410-88530 | 5016 | AGCCGCCGGCCCCCCAGGCTCCATACCAGGGATACCAGGAGCCGCCGCCCCCCCAGGCTCCATACCAGGGATACCAGGAGCCGCCGGCCCACGGGCTCCAATCATCTTCATATCCAGGAT | 8 |
| wt-88530-88650 | 5017 | ATGCGGGTCCCTGGACCCCAAGGTCTCAACATCCATGTTATAGGCACCCCTGGGCACCATGGTCTCAAGATCCTGTGCATGGGCACACCCAGGGTCCATGGGATCCCAGGGCACCACATC | 4 |
| wt-88650-88770 | 5018 | TCCCACCTCAGTGGGATGGATCTGCAGGACATGGCCAGGATCAGGTCTCCCAGTTCCCACATCTGCAATCGGAGACAGGCCCACCACGTCTTCAACTTTCATTGGTGCCACTGGTCTCAT | 4 |
| wt-88770-88890 | 5019 | CCTCTGCACCATCATGGTCATCTCCCCAGCCCCGAGCCCCCATACGCCCCATTCCAACAAGATTCCCCCCTCCCCCTATGCCGTTACAAGATAGCATGGCCGTGGGGTGTGACTCATCAG | 4 |
| wt-88890-89010 | 5020 | GTACAGCATGCCCAAGCATGCCCTTTGCCAGTGATTACAGTCAAGGTGCATTTACCCCACTGGACATTAATGCCACCACGCCAAAAAGGCCTCGAGTAGAAGAAAGTTCTCACGGACCTG | 4 |
| wt-89010-89130 | 5021 | CCCGGTGTTCCCAAGCTACTGCTGAAGCACAGGAGATTCTCAGTGACAATTCTGAGATCTCCGTGTTCCCAAAGGACGCGAAGCAGACTGACTACGATGCATCCACTGAAAGTGAGCTAG | 4 |
| wt-89130-89250 | 5022 | ATTAAGGGGATCCAAGGTGACCCCTGTTAGCTATTTGATCTTTGACTGACACATAAACATGGTTTAAGGAATGAACACTCATGGTGTGAGACTGGAACTGTACTAAATTTGCTGACATAT | 4 |
| wt-89250-89370 | 5023 | GTACAATGAGAGCCAAAAATTTGATAAACCTTAAAAGTCCCCCCATCTAATGATGTCCAGTTCCCTTCTCCCACCCTGTACACCCCGACCCAAAGGGACTCAATGGCATTCAGATTTCTA | 4 |
| wt-89370-89490 | 5024 | GTTACCACAGGTAGAATATCGGGCGTTGGCCCATAAAAATAAGTGCATGGATATAGCTCTGCACAGGCTTGGAAACACCCATTCCAGGTGTGCTTCTTTTTGGTGAAATAAAAACAGCAT | 4 |
| wt-89490-89610 | 5025 | CCTTTATATGAAAATGTGTATTCTCTGGTGTTGCAGTATGTACAGTTAGCTTTGGTATAGTTTTGGGGTACCTGAAATGTGTGCAGGGTGGGTGTCCAATGTGGCAGTTTTACCTCTTTG | 4 |
| wt-89610-89730 | 5026 | TCCCCATACTCCTGCTCGGCCGTCTTGTTAAAGTTAACCGGCGGTGGAGGATCCACCGGCCAGACCTCTACATTTGGTTTGGGTACCCAGGTGATGGCCGCGGCTGCCACCCGCCCTCCT | 4 |
| wt-89730-89850 | 5027 | CCTCTTACCCTGGGTGGCAAAAAGTATGCCAGGAGTAGAACAATAACAAGTGCGATGGCGGTAAACAATGGCACCCTCACCTGCTTAAATGAAACCATGGCAACCACTTCAAAGAGAGCC | 4 |
| wt-89850-89970 | 5028 | GACAGGAAGATATTTATTAATATTCCATTAGTAAACGAGGCGTGAAGCAGGCGTGGTTTCAATAACGGGAGTTAGAAATTTAAGAGATCCTCGTGTAAAACATCTGGTGTCCGGGGGATA | 4 |
| wt-89970-90090 | 5029 | ATGGAGTCAACATCCAGGCTTGGGCACATCTGCTTCAACAGGAGGCGCAGCCTGTCATTTTCAGATGATTTGGCAGCAGCCACCTGCGGACAAAAATCAGGCGTTTAGATGGGGCATTTA | 4 |
| wt-90090-90210 | 5030 | TGTTTGGGACGCTAGCCGCCTGGGCATTCGTGTTAGTATATACTGACCTCACGGTAGTGCTGCAGCAGTTGCTTAAACTTGGCCCGGCATTTTCTGGAAGCCACCCGATTCTTGTATCGC | 4 |
| wt-90210-90330 | 5031 | TTTATTTCTAGTTCAGAATCGCATTCCTCCAGCTGCGAGCAAGGGAATGCGTTACTACAAGTGGTGCCTAGTCAGTTGAAACAAGCCCCACCATCCGCTGCCGCCCCTCCATGAGCCCCA | 4 |
| wt-90330-90450 | 5032 | CCGTCCGCTGCCGCCCCTCCTTGAGCCCCTCCTTACCGATTCTGGCTGTTGTGGTTTCCGTGTGCGTCGTGCCGGGGCAGCCACTGGTGCAGGCTGTGGAACACCAATGTCTGCTAGCTG | 4 |
| wt-90450-90570 | 5033 | TTGTCCTTGGTTAGCCCCGGGGCAAGCAAACACCACTGCTGCTGCTGTTTGAACAGTAGAATTGTCTCCAGGTTGAGGTGCTTCTCCCCCGGCTTGGTTAGTCTGTTGATTCTGGGTTAT | 4 |
| wt-90570-90690 | 5034 | GTCGGAGACTGGGAACAGCTGAGGTGCTGCATAAGCTTGATAAGCATTCTCAGGAGCAGGCTGAGGGGCAGAAAACCACGACCCAGTCGGAGCGGTTGAAACATGATAGGCAGTTAGCTG | 4 |
| wt-90690-90810 | 5035 | GCCTTGTGGCAGAGGCTCTGGCAGCACCGGCCACAGCACACAAGGCAAAGGAGCTTGCGATGGCCCTCCCAGGTCCTGATAGACTCTGGTAGCTTGGTCAAAAGCTTGTACAAAAGGCAC | 4 |
| wt-90810-90930 | 5036 | CTGGTATGGGTCAGGTGTAAATTTTACATCTTCAGAAGTCGAGTTTGGGTCCATCATCTTCAGCAAAGATAGCAAAGGTGGCCGGCAAGGTGCAATGTTTAGTGAGTTACCTGTCTAACA | 4 |
| wt-90930-91050 | 5037 | TCTCCCCTTTAAAGCCAAGGCACCAGCCTCCTCTGTGATGTCATGGTTTGGGACGTGCTAAATTTAGGTGTGTCTATGAGGTACATTAGCAATGCCTGTGGCTCATGCATAGTTTCTAAA | 4 |
| wt-91050-91170 | 5038 | AGAGGAGGAGGCAGTTTTCAGAAGTGTCTAAAATAAGCTGGTGTCAAAAATAGACAGCCCAGTTGAAATATGCATGGCATGCAGCAGACATTCATCATTTAGAAATGTATCCAAGATTTC | 4 |
| wt-91170-91290 | 5039 | ATTAAGTTCGGGGGTCAGGGGGGAGTCCAGATTCAAATCCTCTGTCATGGACTCTAGTGTTGTGGTCAGTTCGTCCAAATGGCCACGAGGGGGCGGGTGGCTCAGGTCCATCTGTCCACA | 4 |
| wt-91290-91410 | 5040 | TATGGCTGCTTCCTCCTTCTGGGGAATAACAGTGTCAGCCATCTCCCTTAGGGCCTTCACGGCCTGACTGGTTTCTTCATCAGGGTCCTCCAACAGATGACTTGCCTCGGGGGTTACTGC | 4 |
| wt-91410-91530 | 5041 | GGGGGCCGGGTCAAGTGGCTGGGGCACCGGGGCTGGCGTTAGGGATCCGACCGGTTCATGGACAGGTCCTGTGGGGGTGGGAGCCAAAGAGGCAGGCAGGGGCCGGTTGGCCCACGGGGA | 8 |
| wt-91530-91650 | 5042 | TCCGGGTGGATGGAAGGGCCTGATCCTCTTTGGCTGACACACCTCTCGCCCCTCGAACACGTCAGATATGGCACTGCCCGCTTCCGGCTTTGGCAGGAACATACCTTCCCGGCTATCCCT | 4 |
| wt-91650-91770 | 5043 | GAGGCCCTTCTTCCTTTTAACGGGAGGAAGAAAGGTGGGCTTTGAGGGGTGGGGGAATATGGGTCTCTCATCGCTCTCTTGGTGGACCGCTGCTATCCAAGGCTGTTCAGGTTCCGCCGC | 4 |
| wt-91770-91890 | 5044 | GTTGGAAGGACATGGAGTTTGACCACGGTTGGGCCTGGATGTCCGGCGCGACTTTGGGGCCCGCAGGCGCGGGGCCTCGGCCCTGGCCTCTTCCCGCTCGCTCTGCTCGGTGTCACTGTT | 8 |
| wt-91890-92010 | 5045 | GCCCGAGTCACTGCTGCTGGAACTGCTGTCACCGCAGTCGGCGCTTTGGGCACCGGGCTTCAGGGGCATGGTCGGGCTCGGGAGACTTTCGAGTTCATCTGTAAAAGCATGAAACTGTCC | 4 |
| wt-92010-92130 | 5046 | GGACTCCGAGTAGCGGGCCTCGGTGTGAGAGGCACCCCCATCATTCCCCATGAGCTCCTCGTCCATCCTGTCGGCTCCGGACACGAGGATAGGAGTTTCCACTGCCTTGGACTTGGTTGA | 4 |
| wt-92130-92250 | 5047 | CAGCAGGCACGCGGGAAGCACGCCGCTCACGTAGCTCCTCTGTCCGGCGTGGCTGGAGTAGGAGGCCCGGGGCAGTGTCTTAATCAGAGCCCTGACATCCTTAACATCGTCCGTCAGATG | 4 |
| wt-92250-92370 | 5048 | GCCTGTCTTGGACGAGACCATAGTCTGGAACATCTCCTCGAGGACGGGATAGGTGAACACCCACTTGCAAAAGGCCTTGAACTTGGAGCTTAGGAGGCCTTCCTTCTCCATCCTGTTCAG | 4 |
| wt-92370-92490 | 5049 | GTGTTCCACTACCTGCTTGCCGGAGGCCATGATGGCCGCGCGGTCCACGCCCAGCACCTTGCTGTAGGTGTAGGCCCGCACCCGACTGTGTTTTAGGAGCTTGTACATAGCGGTGCCTAT | 4 |
| wt-92490-92610 | 5050 | GGTGGCAGGAATCATCACCCGGTTGCTGGGGGCCTGGATGAAGAATCTGTCAGTGACCACTATCAGGTGGTCTAACACGTAGCGCATCACTATAGGGCACGCGATGGAACATGCGTCGTT | 4 |
| wt-92610-92730 | 5051 | GCCGGCATTCTCAGCCCGTCTTCTTACCCTGTTGTTTCGGAGAATGGCCCAAAAATTGCAGATGTTGAGCGTGGCCATTAGCCCGCCCCATTCTCGCCCGTGGGCCTTGGCCTCATTTAT | 4 |
| wt-92730-92850 | 5052 | AAATGCCTTGCATATTTTGTAGGATCTCAGAGTAATCTCCACACTCCCGGCTGTAAATTCCTTGTTGAGGACGTTGCAGTAGTCAGAGACCAGAGAGCCCAGCTGCTTTTTGATTTCAGG | 4 |
| wt-92850-92970 | 5053 | AGTTAGCCTCAGAAAGTCTTCCAAGCCATCCTTTTTAGGCCTCATGGCTAGTAGTAACAGAGGAAATGCCCGACCATTAAAATCTTTCCTCCATGAGCTTTACCTGAAACACTATCCCGA | 4 |
| wt-92970-93090 | 5054 | AGTGGGGGATGTGGTGCATCTACTGAACACCATCGGGGTCGACTGCGACCTCCCACCTAGCCACCCACTCCTGACAGCCCAGAGGGGGCTGTTCCTGGCAAGAGTCTTGCAGGCTGTACA | 4 |
| wt-93090-93210 | 5055 | GCAGCACAAGCTGCTGGAAGACACCATCGTCCCCAAGATCTTAAAGAAGCTGGCTTATTTCTTAGAGCTGCTAAGCTACTACTCCCCCAAGGATGAACAGCGTGACATCGCCGAGGTTCT | 4 |
| wt-93210-93330 | 5056 | TGACCACCTCAAGACGAATCGGGACCTGGGGCTGGACGACAGACTCTGGGCCCTGATTAGGAAACTGCGCCAAGACAGACACCATGCCTCTGTAAATGTCCTCATGCCAGGAAGCGACTA | 4 |
| wt-93330-93450 | 5057 | CACAGCCGTGTCGCTGCAGTACTACGACGGCATCTCCATAGGTATGAGGAAGGTAATCGCGGATGTCTGCCGCAGTGGCTATGCCTCCATGCCCTCCATGACGGCCACGCACAACCTCTC | 4 |
| wt-93450-93570 | 5058 | CCACCAGCTCTTGATGGCGTCCGGGCCCAGTGAGGAACCGTGCGCCTGGCGCGGGTTCTTTAACCAGGTCCTCCTCTGGACTGTGGCCCTCTGCAAGTTTCGCAGATGCATTTACTATAA | 4 |
| wt-93570-93690 | 5059 | CTACATTCAGGGATCTATAGCCACCATCTCCCAGCTTCTGCACCTCGAGATCAAGGCCCTCTGCAGCTGGATAATATCCCAGGATGGCATGCGCCTCTTTCAACACAGCAGGCCTCTCCT | 4 |

| | | | |
|---|---|---|---|
| wt-93690-93810 | 5060 | CACCCTCTGGGAGAGCGTGGCCGCAAATCAGGAGGTCACGGATGCCATTACCCTGCCTGACTGCGCTGAATACATAGACCTACTAAAGCACACAAAACATGTCTTAGAAAACTGTTCTGC | 4 |
| wt-93810-93930 | 5061 | CATGCAATACAAATAAATTTCTCTTACCTGCGTCTGTTTGTGTAGTGAGGTGTTGTGTCCTGTATGGTATTCTACTTTAAAAAGGCCGGCTGACATGGATTACTGGTCTTTTATGAGCCA | 4 |
| wt-93930-94050 | 5062 | TTGGCATGGGCGGGACAATCGCAATATAAAACCCTGACCATCACATGGGGCATTAGGCGACTCTGCATCAGCATCGCTTAAGTATGAGTGGGCAGCAGAGAGGCTCGGTTATTTTGGTTC | 4 |
| wt-94050-94170 | 5063 | CTGAACATCTGGCTGGGGCATTAACTAAGCTTATGAGCGATTTTATCACAGGACAAGATGTCACTCTTTCTGGAGGAAATATTGCAGTCAAAATTCGCGATGCTATAAACCAGACCCCCG | 4 |
| wt-94170-94290 | 5064 | GGGGTGGTGATGTAGCTATACTTTCTTCCCTGTTTGCTTTATGGAATGCCCTCCCAACATCTGGTAGACAATCCTCCAGGGACGATTTAATCCCAGCCGCCGTGCAGGCCTTAACCACGG | 4 |
| wt-94290-94410 | 5065 | CCCACAACTTATGTCTGGGTGTTATTCCAGGTGAGACCTCACACAAGGACACACCCGAGTCATTGCTCCGGGCTATCGTGACGGGTCTCCAAAAATTGTGGGTGGATTCGTGCGGATGTC | 4 |
| wt-94410-94530 | 5066 | CAGAGTGCCTACAATGTCTTAAGGGATTGAAGGCAATTAAGCCCGGCCTTTATGAAATCCCTAGGATAATACCACACACTAAGCAGTGTAGTCCTGTCAATCTCCTGAACATGTTGGTCC | 4 |
| wt-94530-94650 | 5067 | ACAAGCTTGTGGCTTTACGTGGTCATGTGCAGCTTGCATACGACGCCCGTGTCCTGACGCCTGACTTTCACGAAATCCCTGACCTCGATGACTCCGATGCTGTTTTCGCACGCACCTTAT | 4 |
| wt-94650-94770 | 5068 | TGGCAGCCTTATTTCACCTCAATATGTTCTTTATTCTCAAAGATTACATAACACAAGACTCCATGAGCTTGAAGCAGGCCCTCAGTGGTCATTGGATGTCTGCCACGGGCAACCCCCTGC | 4 |
| wt-94770-94890 | 5069 | CTGCAGCACCGGAAACCCTGCGAGACTACTTGGAAGCTTTCCGAAATTCGGATAATCACTTTTATCTCCCGACGACAGGGCCTTTAAACACCTTCCAATTTCCCGAAGAGCTTCTGGGGC | 4 |
| wt-94890-95010 | 5070 | GCGTTGTTGTTATTGATTCCTCTTTGTGTGCCGCCAGTCACGTTCAGGACGTTATCACCCATGGTGTTGGGGCGGGTGTTCCTCGTCCTCGGTTTTCGGCCCTGCCTCCGGCCCCATCCC | 4 |
| wt-95010-95130 | 5071 | GCGAGCCCCAGCAGACATGCTCTCAGTTAACGAGCAGAGGGAATGAAAGCTCACGGCGAAACTTGGGCCAGCCCGGGGGGACCTCCCCTGCTGTTCCCCCAGTTTGCCCCATCGTTTCCC | 4 |
| wt-95130-95250 | 5072 | TGACGGCCTCAGGGGCCAAGCAAAACCGCGGGGGCATGGGATCCTTGCACTTAGCCAAGCCTGAGGAAACCTCCCCCGCCGTCTCCCCAGTATGCCCCATCGCTTCCCCAGCGGCCTCCA | 8 |
| wt-95250-95370 | 5073 | GGTCCAAGCAGCACTGCGGGGTCACTGGATCCTCACAGGCCGCACCCAGCTTTTCTTCCGTTGCCCCAGTAGCATCTCTGTCTGGTGACCTTGAAGAGGAAGAGGAGGGGTCCCGAGAAT | 4 |
| wt-95370-95490 | 5074 | CCCCATCCCTACCGTCCAGCAAAAAGGGGGACGAGGAATTTGAGGCCTGGCTTGAGGCTCAGGACGCAAATCTTGAGGATGTTCAGCGGGAGTTTTCCGGGCTGCGAGTAATTGGTGATG | 4 |
| wt-95490-95610 | 5075 | AGGACGAGGATGGTTCGGAGGATGGGGAATTTTCAGACCTGGATCTGTCTGACAGCGACCATGAAGGGGATGAGGGTGGGGGGGCTGTTGGAGGGGGCAGGAGTCTGCACTCCCTGTATT | 4 |
| wt-95610-95730 | 5076 | CACTGAGCGTCGTCTAATAAAGATGTCTATTGATCTCTTTTAGTGTGAATCATGTCTGACGAGGGGCCAGGTACAGGACCTGGAAATGGCCTAGGAGAGAAGGGAGACACATCTGGACCA | 4 |
| wt-95730-95850 | 5077 | GAAGGCTCCGGCGGCAGTGGACCTCAAAGAAGAGGGGGTGATAACCATGGACGAGGACGGGGAAGAGGACGAGGACGAGGAGGCGGAAGACCAGGAGCCCCGGGCGGCTCAGGATCAGGG | 8 |
| wt-95850-95970 | 5078 | CCAAGACATAGAGATGGTGTCCGGAGACCCCAAAAACGTCCAAGTTGCATTGGCTGCAAAGGGACCCACGGTGGAACAGGAGCAGGAGCAGGAGCGGGAGGGGCAGGAGCAGGAGGGGCA | 4 |
| wt-95970-96090 | 5079 | GGAGCAGGAGGAGGGGCAGGAGCAGGAGGAGGGGCAGGAGGGGCAGGAGGGGCAGGAGGGGCAGGAGCAGGAGGAGGGGCAGGAGCAGGAGGAGGGGCAGGAGGGGCAGGAGGGGCAGGA | 8 |
| wt-96090-96210 | 5080 | GCAGGAGGAGGGGCAGGAGCAGGAGGAGGGGCAGGAGGGGCAGGAGCAGGAGGAGGGGCAGGAGGGGCAGGAGGGGCAGGAGCAGGAGGAGGGGCAGGAGCAGGAGGAGGGGCAGGAGGG | 8 |
| wt-96210-96330 | 5081 | GCAGGAGCAGGAGGAGGGGCAGGAGGGGCAGGAGGGGCAGGAGCAGGAGGAGGGGCAGGAGCAGGAGGGGCAGGAGGGGCAGGAGGGGCAGGAGCAGGAGGGGCAGGAGCAGGAGGAGGG | 8 |
| wt-96450-96570 | 5082 | GGAGGGGCAGGAGGGGCAGGAGCAGGAGGAGGGGCAGGAGGGGCAGGAGCAGGAGGAGGGGCAGGAGGGGCAGGAGCAGGAGGGGCAGGAGGGGCAGGAGCAGGAGGGGCAGGAGGGGCA | 8 |
| wt-96570-96690 | 5083 | GGAGCAGGAGGGGCAGGAGGGGCAGGAGCAGGAGGAGGGGCAGGAGCAGGAGGGGCAGGAGCAGGAGGTGGAGGCCGGGGTCGAGGAGGCAGTGGAGGCCGGGGTCGAGGAGGTAGTGGA | 8 |
| wt-96690-96810 | 5084 | GGCCGGGGTCGAGGAGGTAGTGGAGGCCGCCGGGGTAGAGGACGTGAAAGAGCCAGGGGGGGAAGTCGTGAAAGAGCCAGGGGGAGAGGTCGTGGACGTGGAGAAAAGAGGCCCAGGAGT | 8 |
| wt-96810-96930 | 5085 | CCCAGTAGTCAGTCATCATCATCCGGGTCTCCACCGCGCAGGCCCCCTCCAGGTAGAAGGCCATTTTTCCACCCTGTAGGGGAAGCCGATTATTTTGAATACCACCAAGAAGGTGGCCCA | 4 |
| wt-96930-97050 | 5086 | GATGGTGAGCCTGACGTGCCCCCGGGAGCGATAGAGCAGGGCCCCGCAGATGACCCAGGAGAAGGCCCAAGCACTGGACCCCGGGGTCAGGGTGATGGAGGCAGGCGCAAAAAAGGAGGG | 8 |
| wt-97050-97170 | 5087 | TGGTTTGGAAAGCATCGTGGTCAAGGAGGTTCCAACCCGAAATTTGAGAACATTGCAGAAGGTTTAAGAGCTCTCCTGGCTAGGAGTCACGTAGAAAGGACTACCGACGAAGGAACTTGG | 4 |
| wt-97170-97290 | 5088 | GTCGCCGGTGTGTTCGTATATGGAGGTAGTAAGACCTCCCTTTACAACCTAAGGCGAGGAACTGCCCTTGCTATTCCACAATGTCGTCTTACACCATTGAGTCGTCTCCCCTTTGGAATG | 4 |
| wt-97290-97410 | 5089 | GCCCCTGGACCCGGCCCACAACCTGGCCCGCTAAGGGAGTCCATTGTCTGTTATTTCATGGTCTTTTTACAAACTCATATATTTGCTGAGGTTTTGAAGGATGCGATTAAGGACCTTGTT | 4 |
| wt-97410-97530 | 5090 | ATGACAAAGCCCGCTCCTACCTGCAATATCAGGGTGACTGTGTGCAGCTTTGACGATGGAGTAGATTTGCCTCCCTGGTTTCCACCTATGGTGGAAGGGGCTGCCGCGGAGGGTGATGAC | 4 |
| wt-97530-97650 | 5091 | GGAGATGACGGAGATGAAGGAGGTGATGGAGATGAGGGTGAGGAAGGGCAGGAGTGATGTAACTTGTTAGGAGACGCCCTCAATCGTATTAAAAGCCGTGTATTCCCCCGCACTAAAGAA | 4 |
| wt-97650-97770 | 5092 | TAAATCCCCAGTAGACATCATGCGTGCTGTTGGTGTATTTCTGGCCATCTGTCTTGTCACCATTTTCGTCCTCCCAACATGGGGCAATTGGGCATACCCATGTTGTCACGTCACTCAGCT | 4 |
| wt-97770-97890 | 5093 | CCGCGCTCAACACCTTCTCGCGTTGGAAAACATTAGCGACATTTACCTGGTGAGCAATCAGACATGCGACGGCTTTAGCCTGGCCTCCTTAAATTCACCTAAGAATGGGAGCAACCAGCT | 4 |
| wt-97890-98010 | 5094 | GGTCATCAGCCGCTGCGCAAACGGACTCAACGTGGTCTCCTTCTTTATCTCCATCCTGAAGCGAAGCAGCTCCGCCCTCACGGGCCATCTCCGTGAGTTGTTAACCACCCTGGAGACTCT | 4 |
| wt-98010-98130 | 5095 | TTACGGTTCATTCTCAGTGGAAGACCTGTTTGGTGCCAACTTAAACAGATACGCATGGCATCGCGGGGGCTAGACCTCTGGCTGGATGAGCACGTGTGGAAGAGGAAACAGGAGATTGGT | 4 |
| wt-98130-98250 | 5096 | GTGAAAGGAGAAAATCTGCTTCTCCCCGACTTATGGCTAGATTTCCTACAACTCAGCCCCATCTTCCAGCGCAAGCTTGCTGCCGTTATTGCCTGTGTCCGACGCCTGCGGACTCAGGCC | 4 |
| wt-98250-98370 | 5097 | ACCGTCTACCCAGAGGAGGACATGTGCATGGCCTGGGCCCGCTTTTGCGACCCCTCTGATATTAAGGTGGTTATTTTGGGCCAGGACCCCTATCACGGGGGTCAAGCAAACGGCCTGGCA | 4 |
| wt-98370-98490 | 5098 | TTCAGCGTCGCATACGGCTTTCCAGTTCCCCCCAGCCTGAGGAACATCTACGCGGAGCTGCACCGGAGCCTGCCGGAGTTTTCTCCCCCAGATCACGGCTGTCTAGACGCGTGGGCCTCC | 4 |
| wt-98490-98610 | 5099 | CAGGGGGTGTTGCTACTCAACACCATCCTGACCGTGCAAAAGGGCAAGCCCGGCTCGCACGCAGACATTGGCTGGGCGTGGTTTACTGACCACGTAATTTCATTGCTCTCTGAGCGGTTA | 4 |
| wt-98610-98730 | 5100 | AAAGCGTGCGTGTTTATGCTGTGGGGTGCGAAGGCGGGAGACAAAGCTTCACTAATCAACTCCAAGAAGCATCTGGTTCTGACCTCTCAGCATCCCTCTCCCCTGGCCCAGAACAGCACC | 4 |
| wt-98730-98850 | 5101 | CGAAAGAGTGCCCAGCAGAAGTTCCTGGGCAACAACCACTTTGTCCTCGCTAACAACTTTTTGCGTGAGAAGGGGCTCGGTGAGATAGATTGGAGGCTGTAGAGGGGTCATCACTATGGC | 4 |
| wt-98850-98970 | 5102 | CATGTTTCTGAAGTCGCGTGGGGTCCGGTCTTGCAGGGACCGGCGCCTCTTGTCGGACGAGGAGGAAGAGACTTCACAGAGCAGCAGCTACACTCTGGGGTCTCAGGCCTCCCAGTCTAT | 4 |
| wt-98970-99090 | 5103 | CCAGGAGGAGGACGTGAGTGACACTGATGAGTCTGACTACTCAGATGAAGACGAGGAGATTGATTTGGAGGAAGAGTACCCCAGTGACGAAGACCCATCTGAGGGCAGTGATAGCGACCC | 4 |
| wt-99090-99210 | 5104 | CTCGTGGCATCCTTCAGATTCAGACGAGTCTGACTACAGCGAGAGCGACGAGGATGAAGCAACCCCCGGCTCTCAGGCCTCACGATCTTCAAGAGTCTCGCCATCTACCCAACAGTCTTC | 4 |
| wt-99210-99330 | 5105 | AGGTCTGACACCCACGCCTTCGTTCTCCCGACCACGCACCCGGGCACCTCCGAGGCCGCCGGCTCCCGCGCCGGTCAGGGGACGGGCCTCAGCACCTCCCAGGCCACCAGCCCCAGTTCA | 8 |
| wt-99330-99450 | 5106 | GCAATCCACCAAAGACAAGGGTCCCCATAGACCTACGCGACCTGTACTTAGAGGCCCAGCTCCACGCCGCCCCCTCCACCTTCAAGTCCCAATACATACAATAAACACATGATGGAAAC | 4 |
| wt-99450-99570 | 5107 | CACCCCCCCCATTAAGGGCAATAACAACTACAATTGGCCATGGCTGTAAATAAAATGTCATAACCTGGAGTCTGCATGTCTGTTGTTTTATTCAGTAAACCAGTAGTGCGCGTGAGTTCT | 4 |
| wt-99570-99690 | 5108 | TTAGGGCATCCACGATGTAGCCGCTCGCGGGGTTCCCCTCCCCAGTGATCATCTCGGATAGGGGATTCCTGTCCATGACCACGCAATTAGAGTGCCGGGCCCGGGACAGCGCCACATACA | 4 |
| wt-99690-99810 | 5109 | CATGGCCCGGGTTTGATGTTTCTGTGGCTGCCGAAGCAGATGGCGACTTTGTTTAGGGACAGACCCTGGGCCTTGGCTATGGTCATGGCCAGCTTTGAGCTAATGCCATAGTCACGGATGC | 4 |
| wt-99810-99930 | 5110 | TGCAGAGGTTCAGGGACTTGTCCTCTATCGTCTCATACAGTTTGTTAGTATTGTGTTCCAGGCAGCACACGAAGCCTGCCTCATCCTTGACCATTAGCCTGGGCATGCGTGAACTGCCAG | 4 |

| | | | |
|---|---|---|---|
| wt-99930-100050 | 5111 | CGTCCTGAGGCTGCTGCTTTCCTCGGATGCCAAAGAAGACGTTGAGATGCGTGTAGCCCAGAAGCGTGTAGTTCTCGGTGGTGGAGGCGTAGTCCAGGAGGCCGTGAAGGAGAGGCTCGT | 4 |
| wt-100050-100170 | 5112 | CTGAGGTGAACTCTATGTTGTCGCGAATCAGCATGTTGTTGGTAAATGTGCAGAAGGGGAGGTCCCTGAACTCCCTTCCGCCATAGCGGACGGCCACATCCAGGCATTGCCTGAAATAGG | 4 |
| wt-100170-100290 | 5113 | CCCTGAGGTCATTATATATGTTTAACAGGGAGCAGAGGGGGGCAGAATTTGCGGCCGGGGGAGCCAGTACTCGGGCATAGAAGACAGCGGCGGGGCTCCGCTCCCCATCCCAGGCAACCT | 4 |
| wt-100290-100410 | 5114 | CCAGCGGCAGTTCGCCCAGCTCCATCCCAGCAGTCACCTCCGGATCCCACGTACGCCCGGGCAGGGGCACAGCACCAAGCTCCGCCACGTATTCCCCGTTTTCACAGAGAGAATGTCCTC | 4 |
| wt-100410-100530 | 5115 | CGTGGCTAAAAGCGTAGATGCCTCCGTAGATGAGTCGGGCCAGGAAGCTGTAGACATACTCGGGCTGCTCATGCCCGTGGGCCTCCACGAAGCTGTCCGCCTCGAGCGTGTCCATAAAGT | 4 |
| wt-100530-100650 | 5116 | CCCCGAAGGTGCCGGTATAGCCACAGATGGACTTTTTGGTCTTGCAGTTGACCGACACCGAGCTGTGCTTGACGTAGGTGACATTGTAGGTGACCTTGACCCGTTCTTCGTCCTGCTCGG | 4 |
| wt-100650-100770 | 5117 | TGCCCACCGGGACCATGTCTTGGTCGGCGAACTGCGAGTAGTTACCGAGGCGTGCATAATTCTTTTGGAGCCAGGTGTGGGCCGTGAGGCCCGGAAGCCCGACCAGGGTCTTGTACTGGG | 4 |
| wt-100770-100890 | 5118 | CCAGGGGATCGAGGAAGACCTCGCACTCCACCGGGCAGGTAAACATGGTCACCCCGCCCCCATCTCCCCCGGTTCCCCGCGCGGCACGCCCCTGCCCGGCAGTCTTGAGCGTGGCATGGA | 8 |
| wt-100890-101010 | 5119 | GGGTGGTGAGGAAGGTCTTGACCTCGGCGTGGGAGAGGAAGAGCCGGGTCCAGCCCACGTACTGCGCGGGGTCCATTATGGCCGCCCTGGGGACGACGAAGCGGTCGACGTAGGCCAGGA | 8 |
| wt-101010-101130 | 5120 | TGTCCGGCGAGAGCTCGAGGCCGTACTCGAGGGTCTTCATGAGGTGTCCAAACTGGACGTCGGTGCAGCGCTTGTTGTTGATGAAGAGGGCCCAGTTGCGGGCCACGTCCACGTAGGTCG | 4 |
| wt-101130-101250 | 5121 | CGGCCCTGGGGTTGCCCACCAGGAAGGTGAGGATGTTGTCGCACTCGCGAATCTTGTTTACCTGGGTCTCGTGGCTAAAGGAGGACTGAAAGGCGTCTGTCTGGGTGGGAGAGCCCACGC | 4 |
| wt-101250-101370 | 5122 | AGACGATGCAGGGAATGCGGCCCCGGCGGTAGAGTGGGGTACGCAGCCAGGCGTTGAAGAACCAGTAGCAAAAGACCACGGCTGTTAGAATGTGCACGGAAAGCGTTCCAGCTTCGTCCA | 4 |
| wt-101370-101490 | 5123 | CCACGATCACATTGGTGGTCCATAGCTGCCCCTGGTGCATGTCTCTCAGGACCTCAAAGGCGGGGCCAGAGACTCCCGAGTAGAGCCCCCTGGGCTTGGTTCGCCTGAACTCGGCGGCAA | 4 |
| wt-101490-101610 | 5124 | TGTCGGAGAGTACCGGCCAGTATTTGGCCAGGTCCCGCCGCTGGAGTTCCTCTAGGGCGGCGTCCGTAGAGCGACCATGACTGCTGACCCGCTGCGTCATATTTATGTGGCGGCTCTTGA | 4 |
| wt-101610-101730 | 5125 | ACCCAAAGGCGCTATAGACGGTTGGGCAGTAGGCTCGGAGTGTCTGGGAGAGGTTCTGTGCGGCCACGGTTGTGGCTCCCGTGACCAGGCAGTCCATCGTGTGGTGGAGGCAGCTAACGC | 4 |
| wt-101730-101850 | 5126 | TGGTGCTCTTGCCAGCCCCCGCCGTTCCCGTAATTACATAGGCTGAAAAGGGCAGGAAGGGGGGCTCCGAGAGCTCCGGGTCAAACTCGGGGGAGAACGTCTCCATATCCGGGAGTTGTT | 4 |
| wt-101970-102090 | 5128 | TTCCTGTTCACCCCACGCCGGCCTCGGTCCGACTTTTTGAGATCCTGCAGGGAAAGTACGCCTACGTCCAGGGACAGACCATCTACGCCAACCTCCGCAACCCCGGAGTCTTCTCGAGGC | 4 |
| wt-102090-102210 | 5129 | AGGTGTTTACCCATTTGTTTAAACGAGCCATCTCTCATTGCACGTACGATGACGTGCTACATGACTGGAACAAGTTCGAGGCCTGCATCCAGAAGCGATGGCCGAGCGATGACTCGTGTG | 4 |
| wt-102210-102330 | 5130 | CGAGCCGGTTTCGTGAGTCCACCTTCGAGTCGTGGTCCACGACCATGAAGCTGACCGTGCGTGACCTGCTGACCACCAACATCTACCGAGTGCTACACAGCCGCTCCGTGCTCTCCTATG | 4 |
| wt-102330-102450 | 5131 | AGCGTTATGTGGACTGGATCTGCGCCACCGGCATGGTGCCCGCCGTTAAGAAGCCCATAACCCAAGAGCTCCACTCCAAGATAAAGAGCCTGAGGGACAGGTGCGTCTGTCGGGAATTGG | 4 |
| wt-102450-102570 | 5132 | GGCACGAGAGGACCATCAGGAGTATCGGGACGGAATTATATGAGGCAACGAAGGAAATAATAGAGTCGCTCAACTCCACGTTCATCCCCCAGTTTACGGAGGTGACCATCGAGTACCTTC | 4 |
| wt-102690-102810 | 5134 | AGAACATTTTCATGTGCTACCGCACGCTGGAGCATGCCAAGATCTGCCAGCTCCTGAACACGGCCCCTCTCAAGGCCATCGTGGGCCACGGGGGGCGAGACATGTACAAGGACATCCTGG | 4 |
| wt-102810-102930 | 5135 | CCCATCTGGAGCAGAACTCACAGCGCAAGGACCCCAAGAAGGAGCTGCTGAACCTGCTGGTCAAGCTCTCGGAGAACAAGACCATCAGCGGGGTCACGGACGTGGTGGAGGAGTTCATAA | 4 |
| wt-102930-103050 | 5136 | CGGATGCCTCCAACAACCTGGTGGACCGCAACCGTCTATTTGGCCAGCCCGGGGAGACAGCTGCACAGGGCCTAAAGAAAAAGGTCTCCAACACGGTGGTCAAGTGTCTGACTGATCAGA | 4 |
| wt-103050-103170 | 5137 | TAAACGAGCAATTTGACCAGATTAATGGCCTAGAGAAGGAGAGGGAGCTCTATCTAAAGAAGATCCGCTCCATGGAGTCTCAGCTGCAGGCCTCCCTGGGTCCCGGCGGCAACAACCCAG | 4 |
| wt-103170-103290 | 5138 | CGGCGTCAGCCCCCGCCGCAGTTGCGGCAGAAGCCGCGTCTGTAGATATACTGACGGGCAGCACCGCCTCCGCAATCGAAAAGCTGTTCAACTCCCCGTCCGCCAGCCTGGGTGCCAGGG | 8 |
| wt-103290-103410 | 5139 | TGTCTGGTCACAATGAAAGCATCCTAAACAGTTTCGTTTCTCAATACATCCCCCCTTCGCGGGAAATGACTAAGGATCTGACTGAACTTTGGGAAAGCGAGCTGTTTAACACCTTCAAGT | 4 |
| wt-103410-103530 | 5140 | TAACACCCGTGGTTGATAATCAGGGGCAGCGTCTCTACGTCAGATACTCGTCAGACACGATCTCTATATTATTGGGCCCCTTCACCTATCTGGTGGCAGAGCTTTCACCGGTGGAACTCG | 4 |
| wt-103530-103650 | 5141 | TGACAGATGTCTACGCCACCCTAGGCATCGTGGAGATCATCGACGAGCTCTACCGGAGCAGTCGCCTGGCCATCTACATCGAGGACCTCGGTCGAAAATACTGCCCCGCGAGCGCGACCG | 4 |
| wt-103650-103770 | 5142 | GGGGAGATCATGGCATCCGGCAAGCACCATCAGCCCGGGGGGACACGGAGCCTGACCATGCAAAAAGTAAGCCTGCGCGTGACCCCCCGCCTGGTGCTGGAAGTTAACCGCCATAACGCC | 4 |
| wt-103770-103890 | 5143 | ATCTGCGTGGCCACCAACGTCCCTGAGTTCTACAATGCCAGGGGGGACCTTAACATCCGAGACCTCCGGGCCCACGTCAAGGCCCGGATGATCTCGTCCCAGTTTTGCGGCTACGTCCTC | 4 |
| wt-103890-104010 | 5144 | GTGAGTCTGCTGGACTCCGAGGACCAGGTCGACCACCTCAACATATTCCCCCACGTGTTCTCCGAGAGGATGATCCTGTACAAACCCAACAATGTGAACCTTATGGAGATGTGCGCCCTG | 4 |
| wt-104010-104130 | 5145 | CTCTCGATGATTGAGAATGCCAAGAGCCCCTCCATAGGCCTCTGCCGGGAGGTGCTGGGTCGCCTGACCCTCTTGCACTCCAAGTGCAACAATCTGGACTCTCTGTTTCTGTACAATGGG | 4 |
| wt-104130-104250 | 5146 | GCCAGGACGCTGCTGTCCACCCTGGTCAAGTACCACGACCTGGAGGAGGGGGCTGCCACCCCCGGGCCGTGGAATGAGGGCCTGAGTCTCTTTAAGCTGCACAAGGAGCTGAAGCGCGCC | 8 |
| wt-104250-104370 | 5147 | CCATCCGAAGCCCGGGACCTCATGCAGAGCCTCTTTCTGACCTCGGGGAAGATGGGGTGCCTGGCCAGGTCACCCAAGGATTACTGCGCGGATCTAAACAAGGAGGAAGATGCCAACTCG | 4 |
| wt-104370-104490 | 5148 | GGCTTCACATTTAACCTGTTTTATCAAGATTCTTTATTGACCAAGCATTTCCAGTGCCAGACCGTCCTCCAGACCTTGAGACGCAAGTGCCTCGGGAGTGACACGGTCTCAAAAATAATT | 4 |
| wt-104490-104610 | 5149 | CCCTAGAATAAACTGAGAACAGTCATCAGTAAATCTGTCTCTCGCGTGATTTCCATAGGAATGGTGTAGCCGGGGTGGAGGGCCGATATCACATCAAGCAGAAAGGCCATAATCTCTCGA | 4 |
| wt-104610-104730 | 5150 | AAGTAGGCGGTGGGGCTGAGACCATGCTCAGTGGCCGTCTGGCAGGGGGCCGGGCGCGCTCCGTCCTTGTCCAGGAGACACACGTGGCTTCCAGAGAGGCGCAGCCCAGCCCTCCGCAGC | 8 |
| wt-104730-104850 | 5151 | CGCTGAAGCCAGGCTCGCGGAAGAGCCCAAAACCTGTTTCGGCGCCGCCCGGGGGCCAGTCTCCGGGTCAGGTCGCGGACCAGGGTCAACAGGTGGTCGTGGGATGGCGGGGCCTTGTCT | 8 |
| wt-104850-104970 | 5152 | GCCTCGGGTCTCGCCGCTAGTTGGTCCAGGGTCCAGGAGAAGGCTTCGTGCCAAACCAAAAAGGGCCCCGAGTGCTCCCTACATCCACCCACGTAAAGATCCCCCTGAAAGATGGCCATC | 4 |
| wt-104970-105090 | 5153 | AGTAGGCACCCGGGCCCGCGTCGAGCCTTCACCCGAATGTGTCTGCGGGCCACGGTGGCCTCTCCACCCATCACATCCCGGTCGAGCCGGCTGGCATCCTCCGAGTCTTTCACGCCTTGC | 8 |
| wt-105090-105210 | 5154 | AGGAAAGCCTAGGAGATACAGCAACAGAAAGCTATTAGCCGGTGGTTCCCCCACCATCATTCTTCCTGTTAACGGGAAGAATAAGAGTTGGGCAAACCCCGGGGGCCGCGCTCTCCCACC | 4 |
| wt-105210-105330 | 5155 | CAGCCCCGCTTCTCACCTGTGCTAGTGGCTCCTCTGAAGGATGGGCGGAGGTTGGTGCCACAAAGCCCAGGATGAACTCGTCTGCATAAGCCCAGGTCAGTCCTAGGTCAGCGGCCGCGT | 4 |
| wt-105330-105450 | 5156 | GTAGGAGAACCCGGGTGACGGCGGTGTAGAGGCCCCCGAGTGCCCGTCGCGTGTCTGAGGTGCCATAGCGGTGAAGGGCCCGCAGCCAGGTTTGCGCGTCCCGCGCCTGCCCTCCGCCAT | 8 |
| wt-105450-105570 | 5157 | CAGGCGTTCCCACGGGGGCGCCCCTGGCAGAGAGGTGGCAGCGGGCCAATTCGTAGAGCCACCAAGTGGCATCAGCCTCAAGGATGGCTGTGGCCTCCGCGCGCCCGACCACCGTCGTCT | 8 |
| wt-105570-105690 | 5158 | CGTCCTCCCCCCCTCCCTCGCCGCCTTCCCGCGTGCAAACGTGGCGAGGGTTAATCTCCTTTCGGGTCGGGGGCCAGATTTGTTGTAGGAGCAGCGAGCCGCGTCGTTGCCCTGACCGCG | 8 |
| wt-105690-105810 | 5159 | CGTCGAGGCCCAGGAGGGCGTCTGCCAGGGGCGTCCCAGAGACTCCCAGGTTCAGGTCCAGTAGCAGGAGACCCTCGCTGTGTGGCGCCCGGTGCCAGAAGGCCGGCCTCGCCCGTCCCA | 8 |
| wt-105810-105930 | 5160 | CATAATGGATGGGCAGGAAGGGAAAGCCCGGGACATAGGGCTGGAAATCTGAGCCCCCTGGGCAGAGTTCGGGGTCCAGGAGGTAGAAGATGGGCTTGGTGCCTCTGTGGTTGGCGTAGC | 4 |
| wt-105930-106050 | 5161 | AGGAGGCATAGATACTGCGGAGGAAGGCGTAGAGCCCGCCCCCGGCCATACTCCAAGAGTTGACAAGCCAGGACTCGAATCCCCCAGCCGGCTCAAGAATTTTCAGGCTGACGCGGTGCC | 4 |
| wt-106050-106170 | 5162 | GTCGGGCGTCCCCACCACGGCCGGTGGCCCCGTCGGACGACACCAGATCTACTTCATAAGTGACCGGTCGCAGGATGTCCCTAAAGGGGACGGGAGAGGGGTCGTCGGGAGTCTCGGTGG | 8 |
| wt-106170-106290 | 5163 | AATAGGTGAAAACATCCCCACGCGGTGTCCTGATGTATACGTCCAACTGTCCGGGAGACTCAGAGTGCCTCTGAGCATGGGGGCATGTCTGTTCCCCCTCCATCTCGGACCCGAAGCCAT | 4 |

| | | | |
|---|---|---|---|
| wt-106290-106410 | 5164 | CAACAGGTGGGGGTTGTTGGTCCCGCCCATCATCCCCCGAGCAGCTTTGGCAGACCACCTGTGCTGGAAAGAGAGGCTGGAAGATGAGGCCCTGCTCATCCTCCACCCTGGCGGCGGACA | 4 |
| wt-106410-106530 | 5165 | AGAGTCTGCGGTCTCGGGTTCTAAATGAAAGGTCAAATAGGTCCTTCTCGGCGGCATCGGCGAGCATAGCAATGAGCCCCCCGCTGCGCCTGAGCTCCCGCTCCCATCGCAAAAAGTTGA | 4 |
| wt-106530-106650 | 5166 | GTTCGGTAGTCGAGGGCGCGTTGACCACGGGGGGCTCCAGGGAGCCTCCAAGCGGCGGCTGGCAGGCCTGCACCACGATCAGAGTCTCAACGTCCTCCCTTTTGATGGGCACGATGCCCA | 8 |
| wt-106650-106770 | 5167 | CGACCCAAATCGCCCACCACCGCCCTGCGGTCTGGGTAACATTATAAAAGGTAACCGAGCTGACGCGGGCCCTGACGCTCTCCGCGGGTGTTTCCATCATTGTTTGAGATCTGAGGAGGA | 4 |
| wt-106770-106890 | 5168 | CTGGACCCTTTAAAACATCCGGTCACGCCCTTTGCAAATTATTTAAAAGGTGAATGCTCAACTGAGACCATCGCAATCATGAAGTCCTCCAAGAATGACACGTTCGTCTATAGAACGTGG | 4 |
| wt-106890-107010 | 5169 | GTCAAAACGCTTGTTGTGTACTTTGTGATGTTTGTCATGTCGGCGGTGGTCCCCATCACCGCCATGTTCCCCAACCTGGGGTACCCCTGCTACTTTAACGCACTGGTTGATTACGGGGCA | 4 |
| wt-107010-107130 | 5170 | CTTAACCTGACCAATTACAACCTGGCCCACCACCTGACCCCCACGCTCTATCTGGAGCCGCCGGAGATGTTTGTCTACATCACACTGGTCTTTATCGCGGACTGCGTGGCTTTCATCTAC | 4 |
| wt-107130-107250 | 5171 | TACGCCTGCGGCGAGGTGGCGCTAATCAAGGCCCGAAAAAAGGTCTCGGGTCTTACAGACCTCTCGGCCTGGGTCTCGGCAGTGGGCTCCCCAACCGTGCTGTTTTTGGCCATCCTCAAG | 4 |
| wt-107250-107370 | 5172 | CTCTGGTCCATACAGGTCTTCATCCAGGTCCTTTCCTACAAGCACGTCTTTCTCTCGGCCTTTGTGTACTTTTTGCACTTTCTGGCCTCAGTTCTACACGCCTGCGCATGTGTGACCCGC | 4 |
| wt-107370-107490 | 5173 | TTCTCCCCGGTCTGGGTGGTCAAGGCCCAGGACAACTCTATTCCCCAGGACACCTTCTTGTGGTGGGTGGTCTTCTACCTGAAGCCCGTAGTTACAAACCTGTACCTGGGGTGCCTTGCC | 4 |
| wt-107490-107610 | 5174 | CTGGAGACGCTGGTCTTCTCGCTCAGCGTGTTCCTGGCCCTGGGCAACAGCTTTTACTTTATGGTGGGGGACATGGTGCTGGGAGCCGTGAACCTCTTCCTCATCCTGCCCATCTTCTGG | 4 |
| wt-107610-107730 | 5175 | TACATTCTGACGGAGGTGTGGCTGGCCTCCTTCCTGCGGCACAACTTTGGCTTCTACTGCGGCATGTTCATCGCCTCCATCATCCTGATCCTGCCCTTGGTCAGGTACGAGGCCGTCTTT | 4 |
| wt-107730-107850 | 5176 | GTCTCCGCCAAGCTGCACACCACTGTGGCCATCAATGTGGCCATCATACCTATCCTGTGCTCGGTGGCCATGCTCATCAGGATATGCCGGATTTTCAAAAGCATGCGCCAGGGCACTGAC | 4 |
| wt-107970-108090 | 5178 | AGGTCAACCAGGAGACAGAGGCCCGTCTCTACCCAAGCCCTCGTCTCCTCCGTTTTACCGATGACGACGGACAGCGAGGAGGAGATCTTCCCCTAATGCAATAAAAACTTAAAACACTGA | 4 |
| wt-108090-108210 | 5179 | GGTTACTTTCCCGTCATTCTTTCGGGGGAACGAGGGGAGGCGGGAATTGGGTTAAGATAGGGGCGAAGGGTGGGGGTGGGTGCAAGAATTGGGGCTGGGAATGGAGAGGGGAGTGGGCTA | 4 |
| wt-108210-108330 | 5180 | GGTGCCGACACCGGGGTGCCAAGATAATGGATTGAGTAAGCATGGGGCTCTGATCGGGTCCGCCGGGTTCTCAGGGGTGTAGTGGGTGGGCATTGCATATTTTTGCCGCGGTGCTGTTGG | 4 |
| wt-108330-108450 | 5181 | GCCTTGGACTCGGGGTGATCATCCGTACCATCACCCGCACCCGCACCCCAGTCCACAGCCACCGGCCAAGGTCCTGGGCCTCCCACCACCGTTATGCCTCCCCCTTTACCCATTAATTAC | 4 |
| wt-108450-108570 | 5182 | AAGAGATGTTAGTTTGGTTTTTTATTTGGCAAAAACAGCAATTCATCATTTTCAGAGTCCTCATCATATTCGAGCCCCTCGTTGGTTTCCCCGCAGGCCCTCCCTTCTTCGGCCGCTATT | 4 |
| wt-108570-108690 | 5183 | AGCTTAGTAGTCTCCAGGTTAAACTCCTCATAGTCATTATACAGGTTGATTATTCCCCCGTCCACGTCGCCTATGGAGTTGACTCGTCGTCGGCAAAGAGACCAGAGGGCACCCATGGCG | 4 |
| wt-108690-108810 | 5184 | CGGTGTCAAAAGTATTGTCTGCGTACGCTTTCCAGGAGCCAGCCGCGGTGCTCAAGGTCTTACGGATGACAGAGTCCGGCAGGACCACGGGTGTCACCAGCACCGCCACGGGAATCTCCA | 4 |
| wt-108810-108930 | 5185 | CCGAGGCGTCCAGAAGCAGGTCTGAGCCGAGCGTGCAGGTCGCCGGGTCTAGAGGCGACCGTTTTCGAAAGAAGGCCGTCACAATGTTCACCCGGGGTGAGCAGTCTCTCCCGGGCTTGC | 4 |
| wt-108930-109050 | 5186 | CACCCCCACTGTGGCGGACGTAGTCTCCAACAATTTTGTATTGGAGGAGCACCTGGTAGAAGTAGTTGTGCCGTGGATTGATGAAGATGTTGACTGGGACCCGGTCTTTAATACCAATGC | 4 |
| wt-109050-109170 | 5187 | GCCCCGCATTTTCGCTTGGGTCCGTCATTACGTAGAGCATAGACTCCACCCCCCTGTTGGCAGCTAGGCTGTCTGCCACCAGGTCATGACCGGGGCCCAGTTTGCGCTTACGGACATCTT | 4 |
| wt-109170-109290 | 5188 | TAAGATTCCAGGCCTCATCCTGCGTCAACAGATAGTCACCCTCCGAGGGCAACCGCCCATCCGGGACGTACTCCACGGTAGGACGAGCTATAGAATTGATAAATCTGATAAATGACCTCT | 4 |
| wt-109290-109410 | 5189 | TGCATGGCCTCTTGTAAAGCGCAGTGTAGGATGGGTAGATGGGGTCAAATTCTGACTTGGAAAAGAGGTACTTGAAGCGGCACTTAATCTCATAAATGCAGCTCCGGTCGGTGAACAGTA | 4 |
| wt-109410-109530 | 5190 | TAAAGTCTCCCTGTGACTCCACATTGACGCAAAGATCCAGAGACACCCCAAAAATGCCATCCGTGGGACTAATCATAAAGCCAAATTGACGGTTGGCGGATGCGTCCCCGCAGATGAGCT | 4 |
| wt-109530-109650 | 5191 | TACAGACAATGTCCTTGACCGTGTCCTCACACCGCAGGCCAAAGGCCACAGGTCCCCCAAAGTAGTGATTTGTGGAGATGGGAGCTGGCTCAAACACCTTGGTGGGTCCATTCTTAATGG | 4 |
| wt-109650-109770 | 5192 | TGGAGAGCAGCTTGGAAGAGGAAATTATGCCATTTCGCAATATGTCCCACATCAGGTTCTCAGACTGCCCCCTGGTCATGGACTCCACGTACGAGCAGAGAACAGTCCTCTGCTCGTCGG | 4 |
| wt-109770-109890 | 5193 | TGGCCTCCTGTAGCCCCCAGTAAATGGATTTCAGGGAGGGACCGTCCTTGCTGTCATTCTCTTGGACTAACGAGGAGACAAAGTCACAGAAGCCAGTTTCACCAGAGAACTCTTGTATTT | 4 |
| wt-109890-110010 | 5194 | GTTTACAGAGGCAATAGAGATAGACAAAGCGCATGGCCGGCATCTGAGGTGGACGGTCAAGGTTACGGACAAAGGCCTCAGTCTCCGGACTGCGGAGGAAGCGGGCAAACGTGTAGGAGG | 4 |
| wt-110010-110130 | 5195 | TCATCTCCTCCATGGGATCCTCGAGCTCATCCACGTCGGCCATCTGGACCAAAGAAGTCGTCTGCCAAGAGTTCAGCTACCAGACCTGGAAGATGAGGGTGCTCAAACCGTGGGCGACAG | 4 |
| wt-110130-110250 | 5196 | TTGAAGAAGTAGCTCTCCTTGAACCTCTTTTTAAGGCTCCGGCACCACTGCAAGAATTGACTCATATGCTCCGCCGTGACATCCACGCACGGACTCTCGCCACACGAGGTCAGGCCCATG | 4 |
| wt-110250-110370 | 5197 | TCTAAGTTCAGGTTCCACATCTGCGACAGCACCTCCAACAGCACCACCTTTGGGGCTGCAAATTGCAAAAAGTAGAGCGGGTCGGATCGGTCAAATCCCATGTCAGGGTTGGGGTAGGGG | 4 |
| wt-110490-110610 | 5199 | ATGTGCCCAGCCCCCCTCAAGATGTAGCACTTGCTCAAAAGGCAGAGGGGCTTATAGGTGTCCTTGGCTATAGAAAATGGTTCCCTCTGGCAATAGAGGCGATAGAGCTGCCGGCCCTTA | 4 |
| wt-110610-110730 | 5200 | GAAGACTTTAGCCGCACATCCAGCATCTTGTTGCGGTCGTGGAGGGAAGCAGTCCCATAATCAGTCAGGACCAGCCTACCCATGCCCCACATGGTGTCTGTGAAATCCACCAGGATGTTG | 4 |
| wt-110730-110850 | 5201 | CTGGGGCTAATGTCCGAATGGAAGAGGCCGCAGTGCCGATTCAGAAAGTAAACGGCATCTTTGAGGCCCTGAAAGCCCCGCACCAGGGGCTCAATACTACCATCATGCCAGTGGCCATAA | 4 |
| wt-110850-110970 | 5202 | TCCTGGAGACTGCATCTGAACTGGGGCATAAACAGGGCGTGGCAGGACGTGCAGGCCGACAGGTAGTCCACCAGGGCCTTGTCCTGCCCATCCTCGGCCGTGGCCTTCCCAATCTGAATC | 4 |
| wt-110970-111090 | 5203 | ATGTCACACACCATGAGCTCGTGATACAGCTCCGTCACAGAGTCATAGAGTTTGACCGTGGCATTATCTGCATGTGCATACACGGCCCCGTAGCTCCCCCGCCCCAGCAGATACTCGCAG | 4 |
| wt-111090-111210 | 5204 | GTAATGGGGAGGTGATCACAGCGCGTCATGTTCTCCGGCAGCTTTACATAGAGGGTCTCCGTCATGTCATCAATGTTGGTCACCTTCAGGTGTTTGTGCTGAAAGGTGAAGTAATCAATG | 4 |
| wt-111210-111330 | 5205 | ACAGTCACCTTCCCCAAAAAGGCCTGGGTCTCTCGAGGGGGTTCTGGGGAGACACTCAACTCGCCACTGCTGGAGGAGTTCGTCGGGCTCAACTCCGCAGCCATATTCACATCCATGTTC | 4 |
| wt-111330-111450 | 5206 | CTCAAATGGCTCGAGGGCCTGTCGCAGCTCGTCTCTGGCCTCAAGCTCCTGCTCACGGAGCTCCTCCACCCGCTCTAGCTGCTTGTAGTTGATTTTTGGAAATTGAGTCTTGGTCGCGGT | 4 |
| wt-111450-111570 | 5207 | GACCACCCTCTGATAGGTAGAAATTAGCTGTTTGGACTCAAACGTCTCCCTTGCGTGGCGCAGGGACTCTAAGGCACCCCGAGCAGATGTAAACTGTGTTTCAAACAGAGCGTGGTCCCT | 4 |
| wt-111570-111690 | 5208 | CCCAAATCTGTCACGTGCGCTCACAGCCGCTCTCTTTTCTACCGAGGCTCTTAGTTGCTGGGCCACCAGATCTCGCTTAGAACTACTCATCTTCATAAGTCACCATGTCCGCAACTATGG | 4 |
| wt-111690-111810 | 5209 | AGCCCAGATCATACGTGGGGTAGAGTACGGTAGTTCCAGTGGAGGCTTCCCGGTAATTTCCCACAGCGTCCACCATATATCTTTCTGCCTCTCCCGTTAGAATTAGGCAAGGATCATACG | 4 |
| wt-111810-111930 | 5210 | TGTCCACCGGCCTTTTATACTGAGCGTTTAGGTTTTGTTTATGTAGCAAGCACAAAAGGCACACACGAGTGATGCAAAAGGGTTCCTGAGGCAGCAGGCAGAGCTGTTTTGCCATTTTAT | 4 |
| wt-111930-112050 | 5211 | TCAGGCGGCTAACGTCAAAGGGAGGAGCTATATCCTCACCCTTCCAGTCACGCACGTCCAAGTACAGGGCATACACACACCTGGTGAGGTGTGCCAGGAATGCCTCTATGTTGGCACATG | 4 |
| wt-112050-112170 | 5212 | GTGTATAAACCGCAGTGGGTAGCAGAATAGGGCCCCTTTTGCCCCGTGCCGCAGCGTAAACGCAGTGACGCTCTTCGCAGTGGGACCTGGGGCCGTAGAAGAGGGCCCACATCCAAGGGA | 4 |
| wt-112170-112290 | 5213 | GTGGGTCTTCAGGCACCAGGGAGGTCCAGGTTTGGGAGTGGGCCAATATTTGCAAGGCCTGACCTATAACCTCATCTTTGTTCCAGGCCAGCGCAATTCGCATAAGGTCCCCATCAAACA | 4 |
| wt-112290-112410 | 5214 | CCTCAAAACACAGACCCATGCCCATTTCAGGCTGAGAGGGCTCCATCCGGCTCGACCAACCTTGTCCACCAAACTGCCATTCTTCTGGTAAACGGGGGTTGAGGGGCAAGAGCTCCAAAG | 4 |
| wt-112410-112530 | 5215 | CCAGGCTCGAGAAGTCATAGTCATCCTCGGCCACACGGCCGGAGCTCCGGGCCTCGTGCCAGGGCCTGTTGTCCTGGGGGAGGATATTGGACACGAGCAGGAAGCTCTTGAGTGGCGTCT | 4 |
| wt-112530-112650 | 5216 | CCACCAGCTTAAATTGCTCGGGCGTGTCCTGGCAGGCCTCCAGTGCCAGTTCCAGACACTGCCCATACCTGCGGGCGAGCATCGGGTCATCGGGCATATCGGCCTTGACCGCGTTGAACA | 4 |

| | | | | |
|---|---|---|---|---|
| wt-112650-112770 | 5217 | TGCTGTATGCCTCGCAGCGCGGCCGTCTGACCGAGAACCTAAGAAACGCCCTTCAGCAGGACAGCACCACGCAAGGCTGCCTGGGTGCCGAGACCCCGAGTATTATGTACACAGGGGCCA | | 4 |
| wt-112770-112890 | 5218 | AGTCAGACAGGTGGGCTCACCCTCTGGTGGGCACAATTCACGCCAGTAATTTATATTGCCCAATGCTTCGAGCATACTGCCGCCACTATGGCCCCAGGCCCGTGTTTGTAGCTTCTGATG | | 4 |
| wt-112890-113010 | 5219 | AATCATTACCCATGTTCGGTGCGAGCCCTGCCCTTCACACCCCAGTCCAGGTCCAGATGTGCCTACTACCAGAGCTACGCGACACGTTACAGCGCCTGCTGCCACCACCCAATCTTGAAG | | 4 |
| wt-113010-113130 | 5220 | ACTCCGAGGCCTTGACGGAATTCAAGACCAGCGTGTCCTCTGCCCGTGCCATCCTTGAGGACCCCAACTTTTTGGAGATGAGAGAGTTTGTCACCAGCCTGGCCAGCTTCCTGAGTGGTC | | 4 |
| wt-113130-113250 | 5221 | AGTACAAGCACAAGCCCGCCCGCCTAGAAGCATTCCAGAAACAAGTAGTGTTACATTCTTTTTATTTTCTGATCTCAATCAAATCTTTAGAGATTACAGACACCATGTTTGACATCTTTC | | 4 |
| wt-113250-113370 | 5222 | AAAGTGCTTTCGGATTGGAAGAAATGACGCTGGAGAAGCTGCACATTTTTAAGCAAAAAGCCAGCGTGTTTCTTATCCCCAGGCGCCACGGCAAGACCTGGATAGTCGTGGCCATCATCA | | 4 |
| wt-113370-113490 | 5223 | GCCTCATCCTCTCGAATCTCTCCAACGTGCAAATAGGCTACGTGGCTCACCAGAAACATGTCGCGTCCGCCGTTTTCACTGAAATTATTGACACCTTGACCAAGAGCTTCGACTCCAAGC | | 4 |
| wt-113490-113610 | 5224 | GTGTAGAGGTCAACAAGGAGACCAGCACCATCACGTTTAGGCACAGTGGGAAAATCTCCAGCACCGTAATGTGTGCCACCTGCTTCAATAAGAATGTAAGACCTGACGTTTCAGTACTTG | | 4 |
| wt-113610-113730 | 5225 | GCAATTGTAGAGCATAGCCCGGCTGTAAAGGTCAGAAAATCGCAGCAGGGTCCAAGGTTGTGCTGTACATGGGACCTCTTTCCCATTAGCAAGAACCCCCTGCAGGACACGTGACATGTC | | 4 |
| wt-113730-113850 | 5226 | CGGGTGCATTTTGGGTGGGTTAAATCTCAGTCCCACCACAAAGGGGGCATCCTCCGGTTTGAACATCAGACCCAACAAAGCCCGATGCCCAGTTATGGGTACGTAGTCGTTGTTCAGGGC | | 4 |
| wt-113850-113970 | 5227 | CGTGCATGGCAGCAGACAAGGACAGGTGCCAGATGTGCCTGGGCTATCGTCCTCCGTCCAGCCACGCAGGATGTTCACGTGGGCCCCGGCACCATAGCATGTCACACATTCCCCGTTATC | | 4 |
| wt-113970-114090 | 5228 | ACATCTGGTTAGCAGGTTGATAAAATGGGTCAGTGATGGAAAGGTTGGCATATTGGGGCAGCACATCAGCATGTCCATGTTAACGAAAAACATGTACAGGGCCCCTTCTGCATACCAGGC | | 4 |
| wt-114090-114210 | 5229 | ACCACCCCGTCCCAGTGGGATGATCTCCGAGGGTGTGATATCTTGCAGTTCTTCTACTGTTTTAACGGCGGTTGAGGTGGTAAAGACGTGGGCCGTGGTCAGATCTGTGCAGGTGACTAC | | 4 |
| wt-114210-114330 | 5230 | AGGGTTACCCCTAATCTCCACAGGCACCGCCTCACCCACTGCATCTGAGAATACCCCAAAGTACATGAGAGTCAGGCTGTGTGGCCCCTGGACTGCCTTAGTGAAGAGAACCTCGGGCCT | | 4 |
| wt-114330-114450 | 5231 | GGCCACGGTGGCTAGGGTTCCATTGATGTAGACGGTCACATAGGTGGGCTTCTTCTTGGGCTTCAGCACAATGAGGGTAACATTCATGTAGGTTTTAGGAGGTCCGGCTATCTGAGGCAC | | 4 |
| wt-114450-114570 | 5232 | GTACACAGCTGACACGGCGGTTGTGGCCGTATAGACTTTCATCTGGGGCGTAGAGGCATCGCTCAGCACCCAGAGGCACTCCTTGTTGAGGAACTTGCGAAGCTGTTCCCGGCTACTGTT | | 4 |
| wt-114570-114690 | 5233 | CGCGGCGGATGCCATGACGTGCCAGAATATATCCCCTCTCCTCGGGGGTGAGTGCCAATTGGCCTTTAATAACAAAGCCCCCAGGCAGCACCAAAAATGCCTGCCCGTCCGATGTGGTGG | | 4 |
| wt-114690-114810 | 5234 | CCAGGTGGACGCAGTGCCCGTCAGTTCCAAGGGCTACTAGCTGGGAAGCAGCCCCAACCAGCCCACCCGGGGGCCTGGAGTCGATCACCTTACCCCAGGCCGAGGCCCCTTCCTCATACA | | 8 |
| wt-114810-114930 | 5235 | GCGGGTGGCTATCTATCCATAGGCAGGCATCCGGCGTCTTTGGTGCATTGGAGATAGTAGCTTTCACCCAACAACTTTCCCAACTAACCCGTGTCTGGACAGTGAAGAACGCTTCCCTGA | | 4 |
| wt-114930-115050 | 5236 | TCAGGTCTGAATTTTTATAGATACGGGAGTAGGAGGTGGGAATAACAACTGGGATTTCTTGTTGTGCTGTCCAGGCCTGCATGGCCAGTTTTTCCCTGAAGCTAGCAGAAATTCTGAGGG | | 4 |
| wt-115050-115170 | 5237 | CCACTGAAATGAGGAAGCGAAACTCCCTCTCTGGAGCTCCCAAAATTGAAACCTCAGCAAGATCTGTTGCTGGGGAGGCATGGGTGACAGCTGTCATCCTGTGCAGTCTGCCCTGGGCAC | | 4 |
| wt-115170-115290 | 5238 | TCAGCTCTGGATATGTGACAACATAGAGAGCGTGGGGGCTAAAAATATGAGCAATTCCCCTGACCAGGGCCCTGGACTCACGAATGGCCCGACGGGTCTTAGAGAAAGAAACAGGCACCC | | 4 |
| wt-115290-115410 | 5239 | TCGAGAGTGCCCCCGACCCGACCCCCACAGTGCCGCCAGTCCCTGCTCGGCCTCCGCCGCCTTCCCCACCGGCGCTGCCCCGGATGTTGCTGGGGTTCTCGAGGGCTGGGTGGTGCTTGG | | 8 |
| wt-115410-115530 | 5240 | ACACAGAGGTCTCAGCAGCCGCCTTGGTCTCGGCCCCGGCCCTAAGTCTGAGCCCCAGGCAAAGGGCCGGACTCCCAGCGTGGCCCAACCTCTGCTCCCCTCTATTCTCCTCTTGCGTTA | | 4 |
| wt-115530-115650 | 5241 | TCTCCAATAGAATTTGCTTGAGGTCATACGTTTTAGGGTGCTCGACCTGGGCCGCGGCCACCGGCATATGCTCTATACCCGCCCCTCCGGGGGGCCCAGGATCTATAGGTATGGGCTGCA | | 4 |
| wt-115650-115770 | 5242 | TAGCCGCAGCAGACTCCTGGACCCCAGAGGCCTCTCTGATAAGATGCCCGTCGGTCAGAGCCCTTTTGGCCCCCTCAAAGAGAGACAGGTAATAAATCTGTAGCTCCCCAACCAGCCCTC | | 4 |
| wt-115770-115890 | 5243 | CTTCATCGTAAAATCGAAGGGCGGCCACGTGGAAGGGGTTGTAGAGCTCTGGAAGGCCCTCATCGCAGTACACTGGCACACTGGTAAACGTGCCCCGATGGCTAGGCCGTCCGGGCAGCA | | 4 |
| wt-115890-116010 | 5244 | TGCCCCGAGCAGCAAACACGCGGCAGACCCTCGTGAGACCCGTCCGGTCACTGAAGAGAGTCTGGCACCAGGCCCCCTCGCAGTTTGGCACGCGATTGGGGCAAAGCTCTGCCATAACCG | | 4 |
| wt-116010-116130 | 5245 | TGTCGGGAACAAATAGGTGCACGAGGAGGGGGGTCCCGAGGCCACTCAACACTTGGTTGTCAATGTGGACATCCATAGCTCTCTCATGCGTTTGGCTACAGCATCATAGCGCTTGTTTCT | | 4 |
| wt-116130-116250 | 5246 | GGTGGATTTAAATAACAGGGCCCCGTAGACAGTCTTTTGTGAGTAAATAGAGATGATGACATGGATGTAGAGACTGAGGACCACATCCACCACCTTCTCGGAGGAGGCCCCCCTAAACAG | | 4 |
| wt-116250-116370 | 5247 | CATCAGGCAGCAAGGGAACACAAAGGAAACCAGGGCCGGGATGTGAGGCCTCAGCGCCCCCTCCTGATCAAAGAGGGCCTCGCTGACCCCGGAGATGACATTCTCATTCAGAAAGTAGTG | | 4 |
| wt-116370-116490 | 5248 | ATAGAGGTGATTGACCACAGTCTTAACCAGGCCCTGGACTTGTTCAGGCTCCCACTTGTCCCGCTGGTCCTGTGTGTCTTGTCGGATCTCGGTCCAGGGCCTCAGCGCCGGCTGGAAATG | | 4 |
| wt-116490-116610 | 5249 | CGGCCCCATGTAGTTGCCTGTAAGGGCGCACACCACTCCCTCATGGGTCTCAATCAGGGTGCACTCGCTGGATCCATCACATACGTGGTACTCGCCACAGCCCCAGCAGGCAAACACGGA | | 4 |
| wt-116610-116730 | 5250 | GGCCATGCTCTCAGGTAACGGGAGATGGAACTCCAGCTTACTATACGAGCACAGGTGGCGAGGATTGGGCTCATCCGTGCCCCCCTCCCCCCGCGGGAGGCTCAATCGGCCTTGGTCTGA | | 4 |
| wt-116850-116970 | 5252 | TCAGTGGGGTCCCAGCTCAGCAATTCTCCAAATTCTCTCTCCCCAATAGTGCCTCGCTTCTTATCCTGTCTTTCAGAGCATCCGGGGGCAGACATTTCACCTCTTGTTTGTGGACGAGGC | | 4 |
| wt-116970-117090 | 5253 | TAACTTTATCAAGAAGGAGGCCCTGCCGGCGATCCTGGGCTTTATGCTTCAGAAGGATGCCAAGATTATCTTCATCTCGTCTGTGAACTCGGCTGACCAGGCCACCAGCTTTCTTTATAA | | 4 |
| wt-117090-117210 | 5254 | GCTGAAGGATGCTCAGGAGCGGCTGCTGAACGTGGTAAGTTATGTGTGTCAGGAGCATCGGCAAGATTTTGACATGCAGGACAGCATGGTCTCATGCCCCTGCTTTCGCCTGCACATCCC | | 4 |
| wt-117210-117330 | 5255 | GTCCTACATCACCATGGACAGCAACATCCGAGCAACCACCAACCTCTTTCTGGACGGGGCCTTTAGCACCGAGCTGATGGGTGACACCTCCTCGCTGAGCCAGGGTAGCCTGAGCCGCAC | | 4 |
| wt-117330-117450 | 5256 | TGTGCGTGACGATGCCATCAACCAGCTGGAGCTCTGCCGGGTTGACACCCTCAACCCCCGAGTAGCCGGACGCCTAGCCTCCTCCCTCTACGTGTACGTTGATCCGGCCTATACCAACAA | | 4 |
| wt-117450-117570 | 5257 | CACATCCGCATCAGGCACCGGAATCGCCGCCGTGACTCACGACAGGGCGGACCCTAACAGGGTCATCGTCCTGGGCCTGGAACACTTCTTCCTCAAGGACCTAACAGGGGACGCTGCCCT | | 4 |
| wt-117690-117810 | 5259 | CTCAATCATTGGGGAATCCTGCCCCCTCCCCTGCGCCTTCGTGCACACCAAGGACAAGACGTCCAGCCTGCAGTGGCCCATGTACCTCCTGACTAATGAGAAGTCCAAGGCCTTTGAGAG | | 4 |
| wt-117810-117930 | 5260 | GCTCATCTACGCAGTGAACACGGCCAGCCTTTCTGCCAGTCAGGTCACCGTCTCCAACACCATCCAGCTCTCCTTCGATCCGGTCCTCTATCTCATCTCCCAGATCAGGGCCATCAAGCC | | 4 |
| wt-117930-118050 | 5261 | CATCCCTCTCCGCGACGGTACCTACACCTACACCGGCAAGCAGCGCAACCTCTCTGACGACGTGCTGGTTGCGCTAGTCATGGCTCATTTTCTCGCAACAACACAGAAGCACACGTTCAA | | 4 |
| wt-118050-118170 | 5262 | GAAAGTTCATTAAACTTTATTGACTACACCAGTCCCTTGTAAAGCGACGGGTCTCGCGTGACGGCATTCGTGAGCAGGGCTTCGTCCAGGGGCTTGTTCTTGGCGGACATCATTAGCCCA | | 4 |
| wt-118170-118290 | 5263 | GCCGCAAATATCAGAATTAGCATCAGAAAAGTGAGCCCCACAAACACCAGTGTCCAGAGAGGAAGACCGTAAGATAAAGATGGCTGCCTCTCATCTGGAACGGTGGGAAGCTCAGCAGTT | | 4 |
| wt-118290-118410 | 5264 | GTTTTTGTGGCATTGGACGTCCCTTTGGAGGACAGCGTGGGGGCCAAGGTGGTAGCGTTGGTAATACGGGTAGTAGCACTGGTGGTGGAGGAGGACCTGGTGGTGACATTGCTAGTCACA | | 4 |
| wt-118410-118530 | 5265 | CCCGTGGAGGTTCCTGTTCCGGCCTCGGTGGCAGTGATGTTCTGTGCAGTAACCTTAGTGGTGACATTGATGGTGGATGCGTTGGAAGTTGTTGGGACTGGTGTGACAGTTGTCCCAGTG | | 4 |
| wt-118530-118650 | 5266 | AATGTCACCGTGGTTGTGTTGGTGCTCAGAATAGCAGTTGTGGTTATAGGGGCGCTAGTCGTGGTCAAGGTCGTAGACTGGTTTGTGCTAGGACCCGATGCCGACGGTGATGGTGTAGTC | | 4 |
| wt-118650-118770 | 5267 | ACAGCCGTTGTGCCTGTCACGTTCCCCGCCGAGGCCGTCGAACTGCCACTAGATGTCCAAATAAGGCTTGTCTCACAGATGAGTATCATGGCCATAACAGCGCCTGCCTTGTCTCTGGCG | | 4 |
| wt-118770-118890 | 5268 | TGTGCCATCGCGTCTGGACGCAGAAGGCCTCCCGGCCTCTTTTATAGCTAGTCTCCACACCCAATACTCTACTGAACCATCACATACATGACCTCCTCGAGGTATGCAGGGAATGAGCGG | | 4 |
| wt-118890-119010 | 5269 | TCCGTGAGCCGGTCAACACGACATTGCTTCCGTTTCATGCCTCCAGCTGCCCCTGACCAGTTAGGACCCTTGACGGATGTCTTTAACGGCGCGGTGCAGTTGGTCACCAATGACGGCCTA | | 4 |

| | | | |
|---|---|---|---|
| wt-119010-119130 | 5270 | AAGGCCAACACATCCTTGAAGCAGGGCGTAGGAATGGTACCAAACTCGGGGCCCACCCCATCAAAGACATAATATGTCTCATAGTGGCAGTGATGATGCATCACCACCACAGCACTCGCC | 4 |
| wt-119130-119250 | 5271 | AGGACCCTCTGCATATCTTGTACAAGGCGCCTTTCAACTCGGCCACTGGCTCTGGTGACGTTAAATGTCCTGTTCCTATTAGTCACAGCCTGTAGATTTGGGCACCCAGACTCAAAAAGT | 4 |
| wt-119250-119370 | 5272 | GCAGCTACATGAAGGGCAGCCGCCTCAAATCCACCATGACCCCCATGGCTGTCCGTGTTGTTGGGGTAATAAGTCACATTGTTAATGACCACGGCCGGGATAAGGGTGTAAACCTTGCAG | 4 |
| wt-119370-119490 | 5273 | AATGGATTGGTCGGACACCCATAAGACAGGGGCACCCCAAAATCACGCCCCTTACCCCGAAGCACCTTGGCCCCCACCGGCATAAAGCTGGGCAAAAAGAGTGGGTTAAAACCAAAGGCG | 4 |
| wt-119490-119610 | 5274 | AGTAGGGCCAGGAACGCCAAATAGCAGCAGTAATAGATGAAAACAAAGCTCAGCATGAAACAGCGTGGAGGCTCAGCTAGGGTCTCTGCCTCTCCATCATAGACATCTTCCTTGAATCTC | 4 |
| wt-119610-119730 | 5275 | ATTCTCTCACCGCATACCTCGCTCTTCATCCAGGAGGGGGCCATGGCTGCCATTCTACCAGTTAACGAGGAGAGAGAGAGAGTAGGTCCGCGGAAATTGGTGCCCCTCTCTGCCCTCCTGAC | 4 |
| wt-119730-119850 | 5276 | GAGGCCATGGTGTCATCCATCTCCGCAGTCCGTTCTTCAGCTTTGGCATTGGTCCGGGTCCGGGTGGTCTGATTTTGATTCTGATCCTGGGTATTGGTCTTGGTCTCTCCTCCCCCATTG | 4 |
| wt-119850-119970 | 5277 | GCATGGATTGGCATAGGTGGGTGTGGCTCAGGCTCAGGTTCCGGCCCTGGGACGGCAGCAGCCGCCGGGACGGTGAAGTCGTGGAAGGTAGAGGCCCGTCCCTCCCGAGGTCGTGGGGCC | 8 |
| wt-119970-120090 | 5278 | GGAGCCTTATAAAAGACTTCCACCCTCTCCCCGCTGGCCAAGACACGCCGCTCGTGGACCACGCCATCTTCCTCCCGGCTGATTGTGTGGCTGACGGTGCCGTGTTCCACCGCCACTTGT | 4 |
| wt-120090-120210 | 5279 | TCATCGACCATGGTACCCCCTTTATCTTAACCAGCAAGTGGCCGTCAGGGTCTCTTGAGAGTATGCCGCTGTGGCCAAGCGAGGCCCCAAATTAAATAGTGATGCCAAAGACTGTAGGTA | 4 |
| wt-120210-120330 | 5280 | GGTCATCATCACACGCATGCGTGATAAATCATCCGCCACTGACAGGTCATCCAGGTCTATCCGGGCTATCTCATCCGGCACCATTTCCTGGAAGAGATTCAAGAGGTCGTGATGCTCATG | 4 |
| wt-120330-120450 | 5281 | CCGGATAAGGCCTCGGACCAGGCGCATACTGGCCCTGGGCAGCAGGGTCACCATGATGCAAAAGTAGAGACTCAGATTGTCCAGCAGGGCCAAGCCAAGGGGCCCTGGCACCTCCGGGAG | 4 |
| wt-120450-120570 | 5282 | GGCCAACTCGTAGTGGTGCCCCAGGTATGAAACAGAGCCAAGATGCATGTGTACATCGAGCATGTCTGCGTTCCCGGGAGCCTGCATGACAACCCGGGAGTACACGTTAAACAGGAGAAT | 4 |
| wt-120570-120690 | 5283 | CTTCTGCAGCACCTCCTCTGCTATGGGCGTAGGCAGCACCATGGGGAAAACAATGTCCACATCATTGGACTCTAACTTCACGGTGGCATGCTCTCGTCCAAATACCGGGGGCATAACACT | 4 |
| wt-120690-120810 | 5284 | GAGGCTCCCGGTCCCATGCCACTGGAAAAAGGGCTGGTACTTGTTCTTAATGGCGTAGGTCTGACCTGGAACAATCTTGGTGAGTATCAAACTGTCCACGCTAACCTCATCCAGCACGGC | 4 |
| wt-120810-120930 | 5285 | CAGGGTGCAATCAGACAGGTAGTTGTACATGGACACGTAGTCCGGGACCGTCTCTAGAGAGTACACCTGACCCAAGCCCAATCCCTGCACATTCTGCGTCCCGTGAGTGGAAGCCAGGGG | 4 |
| wt-120930-121050 | 5286 | TAAGATGCAGCCAATCCTCTGTTGCATCTTGGCAATCTCATCGGTATACAGACGAGAGGAGAGAGACACTACCACTTTCAAATCCATCTTTATTGACAATTATCAAAAAACCACCTTATT | 4 |
| wt-121050-121170 | 5287 | TCCAAACTTTAATATTCTTCGTACCGGCGCCACCTCTTCAATTATATAGTGTCCGTAATGGATGGGGGCGTGGGTCTGTTTGACAGACATAAACTCATCGATGAGTGCCCGGGAGGAGGC | 4 |
| wt-121170-121290 | 5288 | TGAGAGTGCGGGGAATGCCTCCTGCAGAAAGCTGCAGGGCTGCTCCAGAAACACGTCAGTGCCAGCAATCACTACAAACTGCACCTCTGTGTTGCTGGTGGCTGGGTGCCCTCCAAGTCG | 4 |
| wt-121290-121410 | 5289 | CTGGCTGTACTCGTTGACCATGTTGTAGAGTCCCCTGTTGTTGCGCAGAAGCTCCTCCTTGTTGAAAAATGCCCGGCAGGGGCTGTAGAGGCCCGGGACGGCCGTCTGGCGATAGGAGGA | 4 |
| wt-121410-121530 | 5290 | GTTGTACATGATGTCACCCAGAGAACCCAGCTGAGATGCCCAGGGATTCACAGTGCTCCGGTATTCATAGGCGGCATCCGGGCGAGAATGGTCATAGATGAGCCCCTCGGCAACCTCCTG | 4 |
| wt-121530-121650 | 5291 | ATTGTAGTTTTCACAGGAGACCACACAGGCGGCCCGTCCCCTTGGAGAGTTGGACTTTTGAAAATAAGCCACGTCTGCCGTGACCGGTGTTACGATAATCTCACAGGTGGCCTGCTGGCC | 4 |
| wt-121650-121770 | 5292 | GTGGCAGAGTCCTGGAGCTCCATTAACATTAGTCATACCTGCCAGGTATGTCCTGGGGTCCCGAAGCAGCGTCCCATTGCGCTGAGCGCCCACCTTGGCCTTGATGTAGTCATTGACTTG | 4 |
| wt-121770-121890 | 5293 | CTGGTTGCCAAAGGCCTCGGCCGGAAAGACGCTAAAGAAGTCTTGGGTGTGGATACCCATGTCAGTAGTGATGGCCGCCACCCTGGCCGGAGTCATGGTCGAGCTATAACTAAGCCCGGT | 4 |
| wt-121890-122010 | 5294 | GTCGATGGAGGCCATCTCGTGATGCACCTCAAAGGTTACCGCGTCCACCCTGGCCTCCCGGCGGCTAACATTTGGGGTCCCAATGAACATGGATGTTGAGGCCCTGGAGCTAAACAATAT | 4 |
| wt-122010-122130 | 5295 | GTTTTCAGAGAGGATCTCATCGGTCCTGACCACGGTCATGGCCACCCCTGGGTGGATCTTGAGCTTGGCCTGGGCAATATAGGCCATGGGGGACATCTTGATGTGCATGGCGGTCATTCC | 4 |
| wt-122130-122250 | 5296 | ACTGATTGAAACGAGGGAAGGAAGACATTCGGCCGCGTATTTGCCCATGGGCGAGCGGTGCCACTCCCGGTACTCTGCAAAGAGCTGCTCTGGCCGGTTGAAGGCTTCCACGGCCCGCTG | 4 |
| wt-122250-122370 | 5297 | CTGAGGATTGCGCATAACAAAGGTGGCAACATCCTGGTGCATGGTGGCAGCCACTCGCGGGTCCCCGTAAAACATATGGAAAGGAATGGCGTGAAAGAGACACTGGGTGACGGCCCGGGT | 4 |
| wt-122370-122490 | 5298 | CCTCTCGGAGAAGGCAAAGGCCACCAGCCCGTTCACCAAAACAGTCTGCTCTGTCCGCTTGTCGGCGGGATTCGGGGCCAGCTGCTGCGTAACGTCATTGTCCACCGACACACGCACGGC | 4 |
| wt-122490-122610 | 5299 | ACGGGTGAAAGTGGGGCAGGTCATGAATGAGGCGCTGAGGTCCCTGATCATGCCCACGGTGGGGCGGAGGTCGGAGATCTCCAGCAGATCCCTGAGCGTCCCATTCTCCAAATTGTCGAG | 4 |
| wt-122610-122730 | 5300 | GATGTCCTCGTCCCTGGTAAAATGGTGGCTGAAGGCTGGCCCGTTGTAGGCCAGGGTCTGGGCCACGTGCTGAAAGTCCACCCCGAGGCCGCACATGTGGGCATTGGTGCAGGTTGGGAG | 4 |
| wt-122730-122850 | 5301 | GAAAACGTAGTAAAAGATCTTTTCCAGCACATCCGCATGCCCCTCATCTACATAAGGGCCTAGGTGCAGACGGAAATCGTGGTCGTGGTCTCCGTTAACCCGGTAGCCGTACAAGGCCAC | 4 |
| wt-122850-122970 | 5302 | AAATTGGGCAGCCATCTCATCCATGTTTCCAACCCTCTCAATAAACTGGGGCGCGGCCAGGGTGTCAGCGTAAACCTCATTTCCGATAATAATCTGGGGGGCCCGGTCACTAACGGTGAG | 4 |
| wt-122970-123090 | 5303 | AAGATGGGTGAAAATGTCTGTGTAGGCCACCGGGGGGAGCAGGTTAGGGTCCAGGAGAGCGCAGACATACTGACCCACGCTCTCATCCCCCACAACATCTGACCCGGCCAGGCGCATCAG | 4 |
| wt-123090-123210 | 5304 | GGCCTGCTCTAGGGCTATAAGTTCCCCATAGATTTTTCTATACATGGAATAGGCCTCCTTGGAGATGGCGTTATTTCCCAGGTGGCGGCAGATGAACTTGATCATGGAAAAGCTGTTCAC | 4 |
| wt-123210-123330 | 5305 | AAAGGCAAGCCTCCCTGACCGTTCCCAGTAGGTGTTGATGCACAGGGACACCAAAGGCACGTTCATGACAAACTTTTCCTCAAACCCGTGGATCATAGCCTCGACTACGTAGAAGAAGGC | 4 |
| wt-123330-123450 | 5306 | TGGATAGGCAGTGTCATAGGCAGTATCCTGCACAGTCTCAATAACGGCCTGATCCACCACGTGGGCCAGAGATGTGGCGGTCTCAAACTGCTGCCCCCGGGCCTCTTGGAATGCAGCTGG | 4 |
| wt-123450-123570 | 5307 | GGCCAGGGGAGTCGGCAGGTTACCCACCATTAGCCGGTGCACAGCCCTGTGCCTGGCCCTCTCCCCGGCATCCCTGCCAATGTAAATATCATAAAGGGGGTGCAGCTCCAGCCGCAGCAG | 4 |
| wt-123570-123690 | 5308 | GTCATAATTGGACGGGTGGAGGAAGTCTTCGGTGGGCAGCCCGCACTTGAGAGCTATATCTGTCACGGGGGCTGCATACTTGTTATCATAGAACTCGTCCACAATAACAAGCACATTCAT | 4 |
| wt-123810-123930 | 5310 | GAGAGCATTCTGTAGGTTAAACGCGAGGAGAAGGTTATTCTTGTTTACGATCCATGCCTCCACCGGTAGCTGCTGTGTGGGGTTGTCCAGCATTTTGATGGCGGCGGAGGTCGTGTACTT | 4 |
| wt-123930-124050 | 5311 | GGGATTGGGCATAAACAGGCCCACTGGGAAATAGTAGCTGTACTGCATTCTTCTGTTGAGGGGGTATGGGGACTGAGTGTCATTGTACATCTTTTGCAGGCTTTCCACGGCCACCGCGTG | 4 |
| wt-124170-124290 | 5313 | AGCCATGAAATGCTCAAACGTACGCATCACGCGCCCGTAGCTCACGGCAGTGACCAGGTTCTCCCCCCGTACCACAAAAGAAGCATAGCTCGAGGGCCCCATAATCTGGTTGTCGGCCTC | 4 |
| wt-124290-124410 | 5314 | CTCACCCAGGAAGGTCAAGAGCTGGCGCAGAACGTTGTCGGTGACAATAAACACCCCCCCCACTGGCTCTCCCCCCTTGGCGGTCGTGTAGGTACTGACCCCCTTGAGCACGCTCTCCCC | 4 |
| wt-124410-124530 | 5315 | GGACACGGCCGCTACCATCTCAGAGAGACGGCTTCGCACGTACTGAGAAAACCCGGAGCCCATGTTCTCGGCCCGGTCCAGGAAGAAGGAGTGCTCCAGCAGATGCCTCTTGAACATGGC | 4 |
| wt-124530-124650 | 5316 | AATGAGGTCAGACTTGACAGTCTTGGAGAACCCCCTCTCAGTGAAGGTGGGATCCGCCAGGGTCTGCAGGATAAACATGGGAGGGGCATGGCGAAGCTTCACACTCAGGACGGTGTTAAT | 4 |
| wt-124650-124770 | 5317 | GAGGCCCCTCTCCAGGGCATCGACCCCAAACTGTAGGGCCGAGGCCACGGTCTTGACAGCCCCCACGTACTCTGCGTACTCGACCGGGGTCTCGGGGATACTATGCAGGATCTCCAGATC | 4 |
| wt-124770-124890 | 5318 | CAGCATGGACAGTTCCATTTCCGTACTAATGTGGTGTTTGTGGCAATTTTTGACCACAATGAATGTCCGCTGCTTGCTGGGTCTCCTTCCGTCCCCGTGAGCAATGGTGGGGACGGAGAT | 4 |
| wt-124890-125010 | 5319 | TCGAAATTGAATCTTGCCATCCGTCATACGACTCAGGTCTTTGAATTCCGTGTTCACACAGGACACGGCCAGTGCCGTCTCCAGGAAGCGAACATATTGGATGGCGTTCGTGTAGACCCC | 4 |
| wt-125010-125130 | 5320 | GAGTAGCACCTCAAACTTGATGCCCGCCTCTCTGGCATCCTTGCCCACCAGCAGGTCAAAGCTATGAAACAACCCCTCAGCCGCTGACTGCCGCAGGTTCGAGAGCAGGTCGGCATCCAC | 4 |
| wt-125130-125250 | 5321 | CGTCAGATAGGGGAAGGGTCTGTTTTCCACACCCTCATTTGAGGCCATGACACAAGGTAAGAGGGAGATGGGGGGAGGTCTCGAGGGCTTCTCTTCACAGCTGGGTCTCTTTTACGCCCT | 4 |
| wt-125250-125370 | 5322 | GGCCTGCAACCGCAGCCCACCCACACTTCCCGAGGATGCTACCCTTCTAATCAAATGGTTGGACACGGCCCTGGGCAGGGAGGCCACCTTTTACGCGTGTCGGGCTATGCGTCGGCTTCT | 4 |

| | | | |
|---|---|---|---|
| wt-125370-125490 | 5323 | ACTCGGCGTTATCCGAATGAATGACTGCCAGGAGCTGCCACCCGGTTTAATAATTCTGAGTCCGGGCACCGTCCCTGGCCCCCTTGGAGTCCAGAGTCTGGAGCATACAGACTGCGAAAT | 4 |
| wt-125490-125610 | 5324 | ATGGTCCTCTGCCCACCCTGACCACGCTGCCCACCTCCCGGTGCCCAGGGTCATCACATACACCGACTGCCCGGGTTCCATAAACACGAGCTCAATGTTTCGCCTTATCATCCGCTACTT | 4 |
| wt-125610-125730 | 5325 | GTCTCATCACCAATTTGAGCGCTGCTTCGAGCAGTTCTGCCGCGTGGTCCCGCGTCGCTTCCTAGGGACCTGTAAGCGAAACTCTGCAAAGATGCTGGCTCATCTGAATCAGGTTACCAG | 4 |
| wt-125730-125850 | 5326 | GATCCCCCCCTGTCCGCCCTTCAGCGGGCGGGAGGCCAGACTCAAGTTCCACTTCTTCTCCTGGAGCACATTCATGCTGTCATGGCCAAACAATGCCACACTCCGGGAGATCAGGACGAG | 4 |
| wt-125850-125970 | 5327 | GGCCGCCACCAACCTCACCCACCACCCACATCTAGTGGATACTCTGTACCACGCCTCTCCGCAGACCCCATTTCTGACACGCAGCGGTGCTCTATACCGCTTCGTCACCTGTTGCAACTG | 4 |
| wt-125970-126090 | 5328 | CACCCTGCCCAATATCTCCATCCAGCAGTGCAAGGCCGGGGACAGACCGGGGGACCTGGAGATCATTCTACAGAGTAACGGCGGAGGGAGGCCCGCGAGCTTCCAGTTCCCCTCCTCCCC | 4 |
| wt-126090-126210 | 5329 | AACTGGCTCCCTATTGCGATGCATAGTTGCTGCGTCCCTGCTGCCGGAGGTGTCCGTGGGGCACCAGGAGCTGTCTCCGCTGCGGTCCAGAAGCCAGGGAGGGCAGACGGATGTCAGGTC | 4 |
| wt-126210-126330 | 5330 | GGGCCCGGACCCGGCCCGGAGACTGGTGGCCCTCCTGCGAAGGGAAGATGGGGCACCTAAAGACCCCCCTCTGGGACCGTTTGGACACCCCCGGGGGCCCGGCCCGGCCAAGAGCGAAGA | 8 |
| wt-126330-126450 | 5331 | CGAGGAGTCTGAGCGTCGAGACGCCCCTCCACCCCCGCTCGATTCCAGCTTCCAAGCTTCCCGGTTGGTGCCCGTGGGGCCTGGGTTTCGCCTGCTCGTGTTCAACACCAATCGGGTGAT | 4 |
| wt-126450-126570 | 5332 | CAACACTAAATTGGTGTGCTCAGAGCCCCTGGTGAAGATGCGAGTTTGCAATGTCCCCCGCCTCATCAACAACTTTGTAGCCCGCAAGTACGTGGTGAAAGAGACGGCGTTCACCGTCAG | 4 |
| wt-126570-126690 | 5333 | TCTATTCTTTACGGACGGGGTGGGGGCCAACCTAGCCATCAATGTCAATATCAGTGGCACCTATCTGAGCTTCCTATTGGCCATGACGTCACTGCGGTGCTTCCTGCCTGTGGAGGCTAT | 4 |
| wt-126690-126810 | 5334 | TTATCCCGCGGCCGTGTCAAACTGGAACTCGACTCTAGATCTCCATGGGCTGGAAAATCAGAGCCTAGTCAGAGAGAACCGAAGCGGGGTCTTTTGGACTACCAACTTTCCCTCGGTGGT | 4 |
| wt-126810-126930 | 5335 | GTCCTGCCGGGACGGTCTCAACGTGTCCTGGTTTAAGGCCGCAACTGCCACCATATCTCGAGTGCACGGGCAGACATTGGAGCAGCACCTGATCCGTGAAATCACCCCCATCGTGACGCA | 4 |
| wt-126930-127050 | 5336 | TCGAGAGGCAAAAATCTCCCGGATTAAAAACCGGCTCTTTACCCTGCTAGAGCTACGCAATCGGAGTCAGATTCAAGTGCTGCACAAGCGTTTCCTGGAAGGCCTGCTAGACTGCGCCTC | 4 |
| wt-127050-127170 | 5337 | CCTCCTGCGCCTGGATCCCAGCTGTATCAACCGAATCGCCTCCGAGGGCCTGTTTGATTTCTCCAAGAGAAGCATCGCCCACTCCAAAAACCGACACGAGTGCGCGCTTCTGGGTCACAG | 4 |
| wt-127170-127290 | 5338 | ACATTCGGCGAACGTGACAAAGCTGGTGGTAAACGAGCGCAAGACCCGCCTGGACATACTGGGCCGTAACGCTAACTTTTTAACGAGGTGTAAGCATCAGGTTAATCTAAGACAGTCACC | 4 |
| wt-127290-127410 | 5339 | TATTTTCCTGACCCTCCTGAGGCACATCCGCCGACGTCTGGGCCTGGGCCGTGCTTCCGTAAAACGAGAGATTACCCTTCTCCTGGCCCACCTGCGCAAAAAGACAGCCCCCATCCACTG | 4 |
| wt-127410-127530 | 5340 | CCGTGATGCTCAAGTGTAAGCAGCCCGGGGCCCGCTTCATTCACGGGGCCGTGCACCTGCCATCGGGACAGATTGTCTTCCACACCATCCACAGCCCCACTCTTGCCTCGGCGCTGGGAC | 4 |
| wt-127530-127650 | 5341 | TGCCTGGGGAAAATGTACCCATCCCGGCCCTCTTCCGTGCCTCGGGCCTCAACGTCCGTGAGAGCCTACCCATGACCAACATGAGGGCACCGATCATCTCGCTGGCTCGCCTCATCCTGG | 4 |
| wt-127650-127770 | 5342 | CCCCCAACCCCTATATCCTAGAGGGACAGCTGACGGTGGGCATGACACAGGACAACGGCATTCCCGTGCTTTTTGCCAGGCCTGTCATTGAGGTAAAAAGCGGGCCTGAGTCCAACATTA | 4 |
| wt-127770-127890 | 5343 | AAGCCTCCTCGCAACTTATGATAGCAGAAGACTCCTGCCTGAATCAGATCGCCCCCTTTTCCGCATCAGAGCACCCCGCCTTCTCCATGGTTGAGTCCGTAAAACGAGTCCGGGTCGATG | 4 |
| wt-127890-128010 | 5344 | AGGGAGCAAACACCCGGCGCACCATCCGGGATATTCTGGAGATCCCCGTGACTGTGCTCTCATCCCTGCAACTGTCTCCCACCAAGTCCATCCTGAAAAAGGCACCGGAGCCCCCACCTC | 4 |
| wt-128010-128130 | 5345 | CGGAGCCCCAAGCCACCTTCGATGCCACCCCCTATGCCCGCATCTTTTACGACATCGGGCGACAGGTGCCCAAGCTGGGCAATGCCCCCGCCGCGCAGGTCAGCAACGTGCTCATCGCCA | 8 |
| wt-128130-128250 | 5346 | ACCGCTCCCACAACTCTCTAAGGCTGGTGCCCAATCCGGACTTGCTGCCTCTCCAGCATTTGTACCTCAAGCACGTAGTGCTAAAGAGTCTGAATCTGGAGAATATAGTGCAGGACTTTG | 4 |
| wt-128250-128370 | 5347 | AGGCCATCTTCACCTCCCCGTCTGATACCATCAGTGAGGCTGAAACCAAGGCCTTTGAGAAGCTGGTGGAGCAAGCCAAAAACACCGTAGAGAACATAGTCTTTTGCCTCAACAGCATCT | 4 |
| wt-128370-128490 | 5348 | GTTCCACCTCTACACTCCCAGATGTCGTCCCCGATGTCAATAACCCAAACATTAGCCTGGCTCTAGAGAAGTATTTTCTCATGTTCCCTCCCTCAGGCACCATTATGAGAAATGTCAGAT | 4 |
| wt-128490-128610 | 5349 | TCGCCACCCCCATCGTCCGGCTCTTGTGCCAAGGGGCTGAGCTTGGCACCATGGCACAGTTTCTAGGAAAGTACATCAAGGTCAAGAAGGAAACTGGAATGTACACACTGGTCAAGCTTT | 4 |
| wt-128610-128730 | 5350 | ATTACCTGCTGCGCATCTAAAGGAAAAACATAACAATCTTGTGAACCAGAAAGATACCCAGAGCAAAAGCAATAAAGTACAGGATTATTGCCAAAACAACGTGTGCTCTTTCTTCATACA | 4 |
| wt-128730-128850 | 5351 | GGCCCGCAATTTCCATGACAGTCCCGTTGGTGGTCAGCAGCAGATAGTGAACGTGGAGGTTGTCAAAATCAAAGTAGTTGGAGCTCAAGATGGAGTTTTGGACTTCCTGGGAGGTGATGT | 4 |
| wt-128850-128970 | 5352 | AGGTTGTAGTTTCCAGGCCTTCCTTTTCATCATAACTGAGCAGGGCAAAGCCACAAAAAATGCAGGATTTCTGCGTCCTGGTAAAATTCTGGATCTTTGGAATCTGGCGGGGCTCCCCAG | 4 |
| wt-128970-129090 | 5353 | CCACAGCACCCTGCGAACATTTATTCATTATAACGGGGGAGAGAAAGAGAGAGCTGCTGAGATAGGTGGTGCTGGCCTCGTATAGCGCCGAGCCTCGGACCTCACGGTCACTAGAGATTA | 4 |
| wt-129090-129210 | 5354 | TGAATGTCACATTGATGAGCGGGATAATCATCAGAACTTTGTCGAGCCTGTCCACGCATTTGTAGGCGGGGAGATGCCACGCATCCCTGTCTTCTCGCTCCAAAGAGAGCCGCCCAAGAA | 4 |
| wt-129210-129330 | 5355 | ACCCATCCACAGCATTTGAAACGGCCGCCTGGTCCAGCGTTGCCTCCTGGGGGGCCATGCTCAGCAGCTTGTCTCGTGTGAGGTCAAATCGTAGGCTGAGGTAGCACGGTGAGAAGAGCC | 4 |
| wt-129330-129450 | 5356 | CGCTCTCTGTCCCCAGGGCTAGCCCCCGCAAAACCTCCCCAATCTCTAGGGCCGAGCACAGGGCGGTGGACAGCAGTTGGTATAGGGCAAGGTTGGGCCCCTGGGTAGTCACGTTCAGCC | 4 |
| wt-129450-129570 | 5357 | GCAACTCGCGTAGCACCACGTGGCTGCCGATAAACAGGGTCTCTCTCATCACGGTATGCAGGGGCTGGAAAAGGGGGTGGCGGTTGTAGGCCGAGAGAAGCACAGATGTGGCGCCTCCAA | 4 |
| wt-129570-129690 | 5358 | TGAGGCCACTGTAAACCCCGGCCTTGGGGTAGCCGACGGTGGCTAACCTCAGCGCGTACTCCTGTTTCTCAGTAGTCAGGTGACCCAGCTCCTCCATCTTGACCGTGGCCATCAGCATGG | 4 |
| wt-129690-129810 | 5359 | CGGCCAAGCGCTCCAGCCCGTAGGATTGCATGCCCTTGACAGTGGCCCCATAACATATGCCGATGATGTCTTTCAGGACAGTCAGCTCAAAGAAGCTCTTGGCCAACCAGCGGAGGTCCA | 4 |
| wt-129810-129930 | 5360 | CGCAGCCATTGCCAGTCTCACCCACAGCATGACCCACCTTAAAGAAGGCCACAGAAACCTCAAACATGGTAGTCAGCGTTTCCGTGTCCAGTTCCGGCTCCCGGCAGCCTCCCTTCATCT | 4 |
| wt-129930-130050 | 5361 | CCAGCAGGACCAGTTTCTGGAGAACGTAGCGAGCGTAGCTGGCGGCTGTCATGGTGACGGCTCGGGAAAACATATCCTTCAGGTTGGGTACAAAGTAGTTGTGAAAGTTGGCATAATGCA | 4 |
| wt-130050-130170 | 5362 | CAAAGGTTGTAACAATCACCAGGGAATAGTCCCCGCTTTGGGCACTGGTTAAGGATGGGTAACTAAAAGGCCCCCTCAGATCCGGCAGGTCCTTCGTCTTGCCAAAGACCAGGCTCAACA | 4 |
| wt-130170-130290 | 5363 | CATGCTCATCTCCCTTCTCGGTCACTCGCTTGTAGGTGCCCATCAGAAATTTAGAAGTCATGGCCCCCGTGTACTGAAACTTGTCCCCGTTGATGGACAGGGCCACATAAGACAAGTGAC | 4 |
| wt-130290-130410 | 5364 | AGCGCAGCTGATAAAAGACATAGCTGTGTGGCCGCGTGTTGGGCAGCATGGTGCCAATATAGTAGAAGAGCTGCTTCTCAAGGGGGGCACTAAGCATGCAGGCAGGGGAATTCAGGCCGC | 4 |
| wt-130410-130530 | 5365 | TAATGACTCCGGGATGGACCTTAGATGCATCCACTTGCATGGATCCTTCAGAGACAGCAGGGATATCGACAGGCTCGGCCAGCGCAATACCAAGGGTACCAGACGTCTTGTAAATTAACT | 4 |
| wt-130530-130650 | 5366 | TGTAGCGGTTAAGCATAGACGCCAAATCTTCGGTGACATTTGCCTCTCTCCACAGCGCCTCTGGGCTAAGGCCTGGGACCTTTGCCATCAGTTCGGTCCATGGGATGGTGTAATGCGAAG | 4 |
| wt-130650-130770 | 5367 | CATGCCCCTCTATGTCCAGGTGCAGCTTAACCTCGCTGAGGCTGGCAGCCCCCACCTCCCATAGCAACACCAGGCAAAAAACACAGAGCAACTGCATCCTAGTCCCGATTTCCCCTCTCA | 4 |
| wt-130770-130890 | 5368 | AAATCAGAGATCACCTTGCTCAGACCAGCCCAATCGAAAAACTGAGATCGTATTGCCGGATTCTTCAATGCCTGCATGTAAATCTCCGTCCAGCATCCAGGTAAATCGTCCTGAAACTCT | 4 |
| wt-130890-131010 | 5369 | GAGAGGTCCACAAGCACAAACTGAAGGTAGGCTAGCGTTCGGGTGAACGCAAGACAAACTTCCAACAACACCGCGTCGGCTCGGAAAGGCTGTATGACTTCCTTAAGTACACTAAAGATG | 4 |
| wt-131010-131130 | 5370 | CTGTTCTTATACAGCTTCTCGGCCACACCACTTCGAATTATGGGGGTGTGGCTTTGATGACATACTGTCGTGATTGTTGTTAGACCGGCACATACCTTCACAATGTCCTCGGGGGCAAAA | 4 |
| wt-131130-131250 | 5371 | TACTGTGTTAGGAGCCAGGCACAGTAAACGGCGTGATATGCATCGTTGACACTCTTCAGGTAGCCAGCATCCAGTCCTGACTCATGTTTCCTCCCTCGCTTCTTCAGGCGGCGCATGTTC | 4 |
| wt-131250-131370 | 5372 | TCCTCCACGTTTAACTTCATCCAGACTATGGTGTCCCCCGGGTCTGCGGTAAACGTGGCCAAAACTTGAATAAAGTCACTATAGGAGAGAAGCTGGCTCCGGAGCAGCATTAGAGGGAAA | 4 |
| wt-131370-131490 | 5373 | ACCACGGAGGCCGACAGCAAATGGCGATCATGCAAAATCCAACAATCCAGGGGCGCGACTGACCTGGCACCAGACTCGGTAACCAGCAAGCTCCGCTTCCTAGAGGCCAAGACTCTGAAA | 4 |

| | | | | |
|---|---|---|---|---|
| wt-131490-131610 | 5374 | GGGGTGGTAAATTTCATCTGGCATGCTAAAACCTCAGCCGACGTGTCTTCCCTTCCATGCCTCGCCCGAGTCACATTCTTGTGCATGGCCTTAATGGCATTTTCATACACATGAGTCCAG | | 4 |
| wt-131610-131730 | 5375 | TACCGCATCGGTTCAGGGACTACAATGGTCAGGTCCCCAAAGACAGCCTTCAAATGATTCAGCATAGTAGTCTTTCCCACACCAGGGGCACCTTCCAAAAATAGGGAACAGGCAGGTTTG | | 4 |
| wt-131730-131850 | 5376 | ATTACTGGTACATGATTTGTTAGGTGGGTCACAATTGGAACCCGCATGCTCTCCTTCCTCTGAGCCTTGGCCTGGCGGGTGTCTTGGGCATCATCCAGATTCAGAACATTCATCACACTC | | 4 |
| wt-131850-131970 | 5377 | CCACTTAGCCGCTTCAGCTGGGCAGCATGCTTGGATAACTTACTAAACTCGCGCCCATGGGCGGCCAGGTGTTCGAAGAGACCAGAAGGCTTACCCTTGCCACCATTCTTTTGTTTTAAC | | 4 |
| wt-131970-132090 | 5378 | GCGGAATGAGAAGAGGGCCTGCGGAAATTAGACTCATCCTCAGACTCACAGTCAGATTTGTCATCGAGCCCAAGGCCGGCCAGGCCCTCCTCAAAGCCTTTCTGGTACATGAAGCTCCGG | | 4 |
| wt-132090-132210 | 5379 | CTCGTGGAGTCCGCACCTCCTTCTGTGCACGAAGTTTTGCGGAACCAGGAGAAGGGGTCTGGCGTCTTGCTGGGGCCACACTCCCGGCTACGGGGCTTCGGGGTAGGGGCAGTAGGCTTT | | 4 |
| wt-132210-132330 | 5380 | TGGTGTGCGGGTGCTGGTGGCTGGGCTCCCCTGGGCAGGGTAAAGGGGCACGATGTGTGCCGGCTACCCGGAGAGTTTCCAGTATTAGATGTCACGGCAGCCTGGGTCCGGCACGGCACC | | 8 |
| wt-132330-132450 | 5381 | CTCTCCCCAGACAGTCCGGTCGGAGCCATCAAGGGGGGCCAGTGGGTGGGCACCTGGTAGAGGCCGTCGTCATCTTCCTCACCTGCCCCTGAGTCACTACCGGTTGGGGTAAGAACTGAG | | 4 |
| wt-132450-132570 | 5382 | GGGGCAAAGTCATCAATCTCAGCGTAAAAGTTTTCGTGTCTTTCGTTTTCAGGGGACTCATCCTCCTGACATTTTCGCCAGCCGCCGGGCGGGCCGGCCTCCTTTCCTGGAAATCCAGCC | | 4 |
| wt-132570-132690 | 5383 | ATGGATCCCACCCGGGGTCTGTGTGCCCTCTCCACACACGACCTGGCAAAATTTCACAGTCTTCCCCCGGCTAGAAAGGCGGCAGGTAAGCGAGCGCACCTTCGGTGTTACTCCAAGCTG | | 4 |
| wt-132690-132810 | 5384 | CTCTCTCTTAAGAGCTGGGAGCAACTGGCCTCTTTTTTGTCTCTGCCCCCGGGACCCACGTTTACAGACTTTAGACTATTTTTCGAAGTCACCCTGGGTCGGAGAATCGCAGATTGCGTT | | 4 |
| wt-132810-132930 | 5385 | GTGGTAGCTCTGCAGCCTTATCCCCGGTGTTATATTGTAGAATTTAAGACGGCCATGAGCAACACGGCCAACCCGCAAAGCGTTACTCGCAAGGCACAGAGGCTAGAGGGCACCGCCCAG | | 4 |
| wt-132930-133050 | 5386 | TTGTGTGACTGTGCCAATTTTCTTCGCACGTCCTGCCCCCCCGTGCTGGGCAGTCAGGGCCTGGAAGTCTTGGCGGCGTTGGTATTTAAAAACCAGCGATCCCTGAGAACGCTCCAGGTA | | 4 |
| wt-133050-133170 | 5387 | GAGTTTCCAGCCCTGGGCCAAAAGACCCTCCCCACCTCCACCACCGGCCTGCTAAACCTCCTCTCCCGCTGGCAGGATGGCGCTCTCCGGGCACGTCTTGATAGACCCCGCCCGACTGCC | | 8 |
| wt-133170-133290 | 5388 | CAGGGACACAGGCCCCGAACTCATGTGGGCCCCAAGCCTTCGCAACTCACTGCGCGTGTCCCCCGAAGCGCTCGAGCTGGCAGAGCGGGAGGCCGAAAGGGCCAGGTCGGAGCGGTGGGA | | 8 |
| wt-133290-133410 | 5389 | CAGGTGTGCCCAGGTGCTCAAAAATAGGCTGCTCCGCGTGGAGCTGGACGGCATCATGCGTGACCACCTGGCCAGGGCGGAGGAGATCCGCCAGGACCTGGATGCTGTAGTGGCCTTCTC | | 4 |
| wt-133410-133530 | 5390 | TGATGGCCTGGAGAGCATGCAGGTCAGGTCCCCCTCCACGGGAGGGCGCTCTGCGCCAGCCCCGCCCTCCCCATCCCCAGCCCAGCCGTTCACTCGGCTCACCGGGAACGCCCAGTATGC | | 8 |
| wt-133530-133650 | 5391 | AGTCTCAATCTCTCCCACGGACCCCCCTCTGATGGTGGCCGGCAGCCTGGCTCAAACGCTGCTTGGTAATCTGTACGGGAACATCAACCAGTGGGTACCGTCCTTCGGACCCTGGTACAG | | 4 |
| wt-133650-133770 | 5392 | GACCATGTCGGCTAATGCCATGCAGCGGCGCGTGTTCCCTAAGCAGCTGAGGGGCAACCTGAACTTTACCAACTCCGTCTCCCTAAAGCTGATGACAGAAGTGGTGGCGGTGCTTGAGGG | | 4 |
| wt-133770-133890 | 5393 | CACCACCCAGGACTTTTTCTCAGACGTCAGGCACCTGCCAGACCTCCAGGCTGCCCTGATCCTCTCGGTGGCCTACCTGCTACTCCAGGGGGGCTCCTCACACCAGCAGCGCCCCCTCCC | | 8 |
| wt-133890-134010 | 5394 | TGCCTCACGGGAAGAGCTGCTGGAGCTGGGCCCGGAGAGCCTAGAGAAAATCATCGCCGACCTCAAGGCCAAGTCACCCGGCGGAAATTTTATGATTTTAACAAGCGGAAACAAGGAAGC | | 4 |
| wt-134010-134130 | 5395 | GCGCCAGTCAATAGCCCCTCTCAACCGACAGGCGGCATATCCACCCGGCACATTCGCGGACAATAAGATTTACAACCTGTTTGTGGGAGCGGGACTACTGCCCACGACGGCCGCGCTGAA | | 4 |
| wt-134130-134250 | 5396 | CGTGCCCGGGGCGGCGGGTCGGGACCGGGACCTGGTGTACCGGATCGCCAACCAGATCTTTGGGGAGGATGTGCCCCCCTTCTCATCTCACCAGTGGAACCTGCGCGTAGGTTTAGCCGC | | 8 |
| wt-134250-134370 | 5397 | ACTCGAGGCCCTGATGCTCGTCTACACGCTCTGCGAGACCGCCAACCTGGCCGAGGCGGCCACCCGGCGTCTACACCTATCGTCCCTGCTCCCCCAGGCAATGCAGCGGCGCAAGCCTGC | | 8 |
| wt-134370-134490 | 5398 | CATGGCGTCAGCTGGTATGCCGGGCGCCTATCCAGTCCAGACGCTTTTCCGCCACGGGGAGCTCTTCCGCTTCATCTGGGCCCACTACGTGAGGCCCACGGTGGCGGCAGACCCCCAGGC | | 8 |
| wt-134490-134610 | 5399 | CTCCATCAGCTCTCTTTTCCCCGGGCTGGTTTTGCTGGCCCTGGAGCTGAAGTTGATGGATGGGCAGGCTCCCTCCCATTATGCCATAAACCTGACCGGACAAAAGTTTGACACCCTCTT | | 4 |
| wt-134610-134730 | 5400 | TGAGATTATCAACCAGAAGCTTTTATTTCACGACCCGGCTGCCATGCTGGCGGCGCGCACACAGCTGCGTCTAGCCTTCGAGGACGGCGTCGGTGTTGCCCTGGGGCGCCCCTCGCCCAT | | 4 |
| wt-134730-134850 | 5401 | GCTTGCGGCGCGGGAGATCCTGGAGCGTCAGTTCTCAGCCTCGGATGACTACGACCGGCTGTACTTCCTGACGCTGGGCTACCTGGCCTCCCCGGTGGCCCCAAGCTGAGCCAGTTCCTC | | 8 |
| wt-134850-134970 | 5402 | GCACTGGAGTGGGTCATTGGCAAAAAGGTAAATAAACTCATCGCACGGGGGTTTTGCCTCCTTCTCGTCTCTTGTTTCGGGTAGGGGAGTAAGGCCGTGCCAGGCCGCCATGCTCAGGGC | | 4 |
| wt-134970-135090 | 5403 | CACGGCGTGCCAGAGGCCCTCGTAGTCGTGCGCATCCGAGAGGATGGCACGGTCCAGAAGCAGATAGCCGGCCAGGCAGAGGAAGGCCACAAAGAGGGGGCGAAGGCGTGCCCGAACCCG | | 8 |
| wt-135090-135210 | 5404 | GGTTTCATGCTCGTCTGCACCCCAGTGGACAAGGCAGTAGAGGACACCCACCACCAGGCGGTTAGGGAGGACACTGCCAAGGTTGAAGAGCAGATTTCCGTCAGCCAGGGTGACCTGGCT | | 4 |
| wt-135210-135330 | 5405 | CAGGTCCGGCGCCCTGCGCAGTCCAAGCTGCGCCCACACACATGCACAGACGGCCCCTGTGACATCAGGCCGGTCATGCAAAAACAGACAAAGAGACCGTGAGCGGTTACCGGGGCGCAG | | 4 |
| wt-135330-135450 | 5406 | GGCCTCTGCCGGGAAGCCCACCCGGGCCAGGGCCCGGTAAAGCAGGTACCAGTATTCATCCGGCACCTTGCGTGCCAACACACGATTCGTGCGGTTTCCAGTATTTATCACGGCTTCCCG | | 4 |
| wt-135450-135570 | 5407 | CCACAGGTAAAAGTTAACACTTAGGGTCAGCAGCTTGGTCAGGGATAGGTGCAAAAACCTGAGCTCGTCCTCGCGCAGAGCGCAAAGCGGCCAGTTCTTTAGCATCTTCAGGAGGAGCCC | | 4 |
| wt-135570-135690 | 5408 | GTGAATCCCAGGTGTCATTCGCGCGTCATCCCCGCGCACCCCCAGTCCCATTAACATAGCGGGCACAATGGTGCAGGCACCGTCTGTATACGTCTGCGGCTTCGTGGAGCGCCCGGACGC | | 4 |
| wt-135690-135810 | 5409 | CCCACCCAAGGACGCCTGCCTTCACCTGGATCCCCTCACCGTCAAGAGCCAGCTCCCTCTGAAGAAGCCCTTGCCACTCACGGTGGAACACCTGCCGGATGCTCCGGTCGGCTCAGTCTT | | 4 |
| wt-135810-135930 | 5410 | TGGCCTTTACCAGAGCCGAGCGGGTCTCTTTAGCGCAGCCTCGATTACCTCTGGGGACTTCCTGTCCCTGCTGGACTCAATTTACCACGATTGCGATATTGCACAGAGTCAGCGCCTGCC | | 4 |
| wt-135930-136050 | 5411 | CCTCCCTCGAGAACCCAAGGTGGAGGCTCTGCACGCCTGGCTCCCCTCACTGTCACTGGCCTCCCTCCACCCAGACATACCCCAAACCACCGCAGATGGAGGCAAGCTGTCCTTCTTTGA | | 4 |
| wt-136050-136170 | 5412 | CCACGTGTCTATCTGTGCCCTGGGTCGTCGGCGCGGCACCACGGCAGTCTACGGTACAGACCTTGCGTGGGTCCTGAAGCACTTTAGTGACCTGGAACCGTCTATCGCCGCCCAGATTGA | | 4 |
| wt-136170-136290 | 5413 | GAATGACGCCAATGCCGCAAAGCGTGAATCCGGATGCCCGGAAGACCACCCTCTGCCCCTCACGAAGCTCATAGCTAAGGCAATCGATGCTGGATTTCTGAGAAACCGCGTGGAGACTCT | | 4 |
| wt-136290-136410 | 5414 | GAGGCAGGACAGGGGTGTGGCCAATATCCCAGCCGAGTCGTATTTAAAGGCCAGCGACGCCCCGGACCTACAAAAGCCGGACAAGGCACTTCAGAGCCCACCACCGGCCTCCACAGACCC | | 4 |
| wt-136410-136530 | 5415 | AGCCACCATGCTATCAGGTAACGCAGGAGAAGGAGCAACAGCCTGCGGAGGTTCGGCCGCCGCGGGCCAGGACCTCATCAGCGTCCCCCGCAACACCTTTATGACACTGCTTCAGACCAA | | 4 |
| wt-136530-136650 | 5416 | CCTGGACAACAAACCGCCGAGGCAGACCCCGCTACCCTACGCGGCCCCGCTGCCCCCCTTTTCCCACCAGGCAATAGCCACCGCGCCTTCCTACGGTCCTGGGGCCGGAGCGGTCGCCCC | | 8 |
| wt-136650-136770 | 5417 | GGCCGGCGGCTACTTTACCTCCCCAGGAGGTTACTACGCCGGGCCCGCGGGCGGGGACCCGGGTGCCTTCTTGGCGATGGACGCTCACACCTACCACCCCCACCCACACCCCCCTCCGGC | | 8 |
| wt-136770-136890 | 5418 | CTACTTTGGCTTGCCGGGCCTCTTTGGCCCCCCTCCACCCGTGCCTCCTTACTACGGATCCCACTTGCGGGCAGACTACGTCCCCGCTCCCTCGCGATCCAACAAGCGGAAAAGAGACCC | | 4 |
| wt-136890-137010 | 5419 | CGAGGAGGATGAAGAAGGCGGGGGGCTATTCCCGGGGGAGGACGCCACCCTCTACCGCAAGGACATAGCGGGCCTCTCCAAGAGTGTGAATGAGTTACAGCACACGCTACAGGCCCTGCG | | 4 |
| wt-137010-137130 | 5420 | CCGGGAGACGCTGTCCTACGGCCACACCGGAGTCGGATACTGCCCCCAGCAGGGCCCCTGCTACACCCACTCGGGGCCTTACGGATTTCAGCCTCATCAAAGCTACGAAGTGCCCAGATA | | 4 |
| wt-137130-137250 | 5421 | CGTCCCTCATCCGCCCCCACCACCAACTTCTCACCAGGCAGCTCAGGCGCAGCCTCCACCCCGGGCACACAGGCCCCCGAAGCCCACTGTGTGGCCGAGTCCACGATCCCTGAGGCGGG | | 8 |
| wt-137250-137370 | 5422 | AGCAGCCGGGAACTCTGGACCCCGGGAGGACACCAACCCTCAGCAGCCCACCACCGAGGGCCACCACCGCGGAAAGAAACTGGTGCAGGCCTCTGCGTCCGGAGTGGCTCAGTCTAAGGA | | 8 |
| wt-137370-137490 | 5423 | GCCCACCACCCCCAAGGCCAAGTCTGTGTCAGCCCACCTCAAGTCCATCTTTTGCGAGGAATTGCTGAATAAACGCGTGGCTTGAAAGTAAACTTTATTGCGTGTTAGTACCTGTCCATT | | 4 |
| wt-137490-137610 | 5424 | CACAGGGGTATCCAGCCCTTGCGCCGCCTCCCCCAGCCCGCCAGCCACCCCAGACAGGAGATGATAATGATGAGGAGCACCGGAGCCACCACAGCACAAGTGATTAGGAGCAGGGCCCAG | | 4 |

| | | | |
|---|---|---|---|
| wt-137610-137730 | 5425 | TGCACCCAGGTGGTCTTAGGGCGCCAGGGATCGATTGGAAAAGGGCCCAGGGTCACTGGCTTATGCGTGGGACGTTTAGAAACAGGCCGCCTATGGGGCCTGTGACTGGTGCTTGTGGTG | 4 |
| wt-137730-137850 | 5426 | TGGGAGACTAATGTGGTGGGGGCTATGGTAGTGGCTGGGATAACAGTAAGATGCATACGCTGAGTGAGGGTCCGGTTGGCATGGTATTGGTCGTCTTCTTCCCCTGCAGAGTAATTGCAG | 4 |
| wt-137850-137970 | 5427 | TGGACCCCGGAGGCCACACTGCAATTTCTCAGTGTCACATTGCACGTGTAGTAACCTGCATGCGCAAGGGTCACATTGGGGATTATCAGAGAGACGGAGGTGTTGGAGTCATTTACCCAT | 4 |
| wt-137970-138090 | 5428 | TCTAGGGTAAGGCTATAATTGTAACCCCCGTTAGTTATATGAGTTCCGTTGTTGGAAGTAGCTACGGCCAAGGGCAGTTGTCCATCCCCGGGAGTGTATCCCCGGCCCAACTCGATCCGA | 4 |
| wt-138090-138210 | 5429 | GAGACCGACTCATTGCTAGGAACGCTGCAGGTGAGATTCACTCTAGCACCTGCATGGGCGGTGACATTTTCAAATTTAACCAGATCTGAGAAAAATGCACAAACAGACCCCACACAGCAG | 4 |
| wt-138210-138330 | 5430 | CACAATAGAAGCACTAAATGAGTCATTCCTAAACTGTCAGTTTTAAAACTCCCTGCTTCTCAGGCCTAAATATGTGGTGGGGTGTGCTTAGGATCACTTTCATATTCTGCAACAACAGCC | 4 |
| wt-138330-138450 | 5431 | ATACCCGGAAGAGGAGCTGCCGGTTGCCATTTTTCAAGCTGCTAAACCACGAGTGGCAGCAGGCCTAAGAAGCTCCTCAGCAACATGGAGACCTCGAAGGGAAACTGGCAGGAGCAGGGA | 4 |
| wt-138450-138570 | 5432 | GTCACGTAGGCACTAGCCTCTTCATGTGAGGTAAGAGATCGCTAAAAATGGGATCAGGGTATGTAAACCGAGTTTTGCGGGGGATGGTGAGCCAGACACGGCGGGTGGGGAAGGAGCTG | 4 |
| wt-138570-138690 | 5433 | ACACGATTGAGTAGAAAGGGCCAAAAATACACCAGCTATAAGGAATTGCTCAGGCCAAAGTTGTTCCTCAGGTGGCTTTAGGCCTAATGTAGGCAATTGCGTGCCTAGAACATTGCTAAT | 4 |
| wt-138690-138810 | 5434 | GTGCCCTGGGTTTCCTGCCTTCATGCAAATATTCTACCTCCCCCGGCCTGGTGCAAAATGTCTGCCTCAGAATACTAACAGCTAATCCAAGCTAACATTCTATCAGTAAACGGGCAGAAA | 4 |
| wt-138810-138930 | 5435 | ACTGATAAGGACCGCGGAGTTTGGCCCTCCGCGGTGTCCGGTGGTCCTCACACGTGCCCTCCCCCCGGGCCGATGGCTGAGGCCCGGAATATGCAAGTGCATCTTTCTAACCAGTAGGGG | 4 |
| wt-138930-139050 | 5436 | CCTCCACCTAGGTGCTTTGTTAATCTTTAGTGGGAACTAGTGGGAGTGCTGTGCCTCGGGTACCCCTATCCTATAGGTCCTACCGGAGCTCCTTGTCTTGATAATCCCTGTAAACACACA | 4 |
| wt-139050-139170 | 5437 | CCACCTAAGAACAAGGCATTGTTAACCTTTGGTGGAACCTAGTGTTAGTGTTGTGCTGTAAATAAGTGTCCAGCGCACCACTAGTCACCAGGTGTCACCGGAGGCTACTTGCCTCAGTGC | 4 |
| wt-139170-139290 | 5438 | CACTTTTACCTTCTCAAATCTATACGGGGGGGGGGGGGGCTCTGTAACATTTGGTGGGACCTGATGCTGCTGGTGTGCTGTAAATAAGTGCCTAGCACATCACGTAGGCACCAGGTGTCA | 4 |
| wt-139290-139410 | 5439 | CCAGGGCTACTTGCCTCGGCATCTCCTCACCGGAGAAGGGGTTAACAAACCCGTGGGGGGTCTTAGTGGAAGTGACGTGCTGTGAATACAGGTCCATAGCACCGCTATCCACTATGTCTC | 4 |
| wt-139410-139530 | 5440 | GCCCGGGCTATATGTCGCCTTACCTCCCCTATATAGTCACGACCCCACCGAACCAGGCATGATGTAGAATAAAATTTTATGCATCATCTTCTAATCTGTGCCGCTTGGAGGGAAACATGA | 4 |
| wt-139530-139650 | 5441 | CCACCTGAAGTCTGTTAACCAGGTCAGTGGTTTTGTTTCCTTGATAGAGACACAAGGACTGCCAGCCCCATTGGGGAGGGGGGGTGGGTACGGGAGAGTTTGGGCTCGTTTAAACAAAGT | 4 |
| wt-139650-139770 | 5442 | CTCATCTGATGCTCTGTGGCACCTCAAGGTGAATATAGCTGCCCATCGACGTATCGCTGGAAACCGGTGGGCCGCTGTTCACCTAAAGTGACGCAAGGTCTGTCAGCCGCCAGGGTCCGT | 4 |
| wt-139770-139890 | 5443 | TTACCAGGCTTTCAGGTGTGGAATTTAGATAGAGTGGGTGTGTGCTCTTGTTTAATTACACCAAGATCACCACCCTCTATCCATATCCCACAATTGATAAACCTCCGCATGTCCAACCAC | 4 |
| wt-139890-140010 | 5444 | CACGTTGAACAGGATGTGGCACCCTAAGAGGACGCAGGCATACAAGGTTATTACCCAGTCCTTGTATGCCTGGTGTCCCCTTAGTGGGACGCAGGCCTAGGTAGCATCATTTACACTAAA | 4 |
| wt-140010-140130 | 5445 | AGCAGTGACCTTGTTGGTACTTTAAGGTTGGTCCAATCCATAGGCTTTTTTTGTGAAAACCCGGGGATCGGACTAGCCTTAGAGTAACTCAAGGCCAAGCATTTCACACCTGCAAATGCA | 4 |
| wt-140130-140250 | 5446 | CCATGTAACCACAGATCTAAACTGAAAGTTGCAGCTTTAGATGGCAAGGAAACTTGGGTTTCAGGCATAGAAAGCCTGGCTCACTATAGCAGCCCATGTTTGTTCCAGGGTGGGGGAAAG | 4 |
| wt-140250-140370 | 5447 | GCACGTGCCCTTAGAAAACTTAGCTGCAAAAATTCTATTGTGTTGGGAGAGCCTCTATATCTAAAGGCCTTTCCTCACAATACAAATGTTACTAACGTCTGCCCTCTGGAGACCTGCTAT | 4 |
| wt-140370-140490 | 5448 | GTGGCTAGACGTATGGCCTACCCAAGACGTTGGGGGTCTCGGGTAGGCCATGATTCTTCCAGGCATAGGTTACAACCAGTCACTGCTATCAAGCCTACTCAGTTCCCAACGCAGCACATA | 4 |
| wt-140490-140610 | 5449 | CCCCCCGCCTCTCCTGCCATGAGGACTTATGGCAGTGTTTACTGTTCTGCTTTTACTCTTGGACCAGGCTGTCATTCTATCAGAATAACAGGGGAAGCAAGGCCCCCTGCTTCAGCGGGA | 4 |
| wt-140610-140730 | 5450 | CACGTGTTTCTAGAATCTCGGAGCCAATAACTACCTGCCCCTCTAATCTGTATGCTGCATGAAAAACCACATACACGTGATGTAAGTTTAGCCAGTTTATTGTTACACCAATGCCCCGAA | 4 |
| wt-140730-140850 | 5451 | AGTCTCCCCCTGTCCCTTTGGGTCTCAGGACCCAGCCCTGGAGCTCGGGGGCGGCCGGGTGGCCCACCGGGTCCGCTGGGTCCGCTGCCCCGCTCCGGCGGGGGGTGGCCGGCTGCAGCC | 12 |
| wt-140850-140970 | 5452 | GGGTCCGGGGTTCCGGCCCTGGAGCTCGGGGGGCGGCCGGGTGGCCCACCGGGTCCGCTGGGTCCGCTGCCCCGCTCCGGCGGGGGGTGGCCGGCTGCAGCCGGGTCCGGGGTTCCGGCC | 12 |
| wt-140970-141090 | 5453 | CTGGAGCTCGGGGGGCGGCCGGGTGGCCCACCGGGTCCGCTGGGTCCGCTGCCCCGCTCCGGCGGGGGGTGGCCGGCTGCAGCCGGGTCCGGGGTTCCGGCCCTGGAGCTCGGGGGGCGG | 12 |
| wt-141090-141210 | 5454 | CCGGGTGGCCCACCGGGTCCGCTGGGTCCGCTGCCCCGCTCCGGCGGGGGGTGGCCGGCTGCAGCCGGGTCCGGGGTTCCGGCCCTGGAGCTCGGGGGGCGGCCGGGTGGCCCACCGGGT | 12 |
| wt-141210-141330 | 5455 | CCGCTGGGTCCGCTGCCCCGCTCCGGCGGGGGGTGGCCGGCTGCAGCCGGGTCCGGGGTTCCGGCCCTGGAGCTCGGGGGGCGGCCGGGTGGCCCACCGGGTCCGCTGGGTCCGCTGCCC | 12 |
| wt-141330-141450 | 5456 | CGCTCCGGCGGGGGGTGGCCGGCTGCAGCCGGGTCCGGGGTTCCGGCCCTGGAGCTCGGGGGGCGGCCGGGTGGCCCACCGGGTCCGCTGGGTCCGCTGCCCCGCTCCGGCGGGGGGTGG | 12 |
| wt-141450-141570 | 5457 | CCGGCTGCAGCCGGGTCCGGGGTTCCGGCCCTGGAGCTCGGGGGGCGGCCGGGTGGCCCACCGGGTCCGCTGGGTCCGCTGCCCCGCTCCGGCGGGGGGTGGCCGGCTGCAGCCGGGTCC | 12 |
| wt-141570-141690 | 5458 | GGGGTTCCGGCCCTGGAGCTCGGGGGGCGGCCGGGTGGCCCACCGGGTCCGCTGGGTCCGCTGCCCCGCTCCGGCGGGGGGTGGCCGGCTGCAGCCGGGTCCGGGGTTCCGGCCCTGGAG | 12 |
| wt-141690-141810 | 5459 | CTCGGGGGGCGGCCGGGTGGCCCACCGGGTCCGCTGGGTCCGCTGCCCCGCTCCGGCGGGGGGTGGCCGGCTGCAGCCGGGTCCGGGGTTCCGGCCCTGGAGCTCGGGGGGCGGCCGGGT | 12 |
| wt-141810-141930 | 5460 | GGCCCACCGGGTCCGCTGGGTCCGCTGCCCCGCTCCGGCGGGGGGTGGCCGGCTGCAGCCGGGTCCGGGGTTCCGGCCCTGGAGCTCGGGGGGCGGCCGGGTGGCCCACCGGGTCCGCTG | 12 |
| wt-141930-142050 | 5461 | GGTCCGCTGCCCCGCTCCGGCGGGGGGTGGCCGGCTGCAGCCGGGTCCGGGGTTCCGGCCCTGGAGCTCGGGGGGCGGCCGGGTGGCCCACCGGGTCCGCTGGGTCCGCTGCCCCGCTCC | 12 |
| wt-142050-142170 | 5462 | GGCGGGGGGTGGCCGGCTGCAGCCGGGTCCGGGGTTCCGGCCCTGGAGCTCGGGGGGCGGCCGGGTGGCCCACCGGGTCCGCTGGGTCCGCTGCCCCGCTCCGGCGGGGGGTGGCCGGCT | 12 |
| wt-142170-142290 | 5463 | GCAGCCGGGTCCGGGGTTCCGGCCCTGGAGCTCGGGGGGCGGCCGGGTGGCCCACCGGGTCCGCTGGGTCCGCTGCCCCGCTCCGGCGGGGGGTGGCCGGCTGCAGCCGGGTCCGGGGTT | 12 |
| wt-142290-142410 | 5464 | CCGGCCCTGGAGCTCGGGGGGCGGCCGGGTGGCCCACCGGGTCCGCTGGGTCCGCTGCCCCGCTCCGGCGGGGGGTGGCCGGCTGCAGCCGGGTCCGGGGTTCCGGCCCTGGAGCTCGGG | 12 |
| wt-142410-142530 | 5465 | GGGCGGCCGGGTGGCCCACCGGGTCCGCTGGGTCCGCTGCCCCGCTCCGGCGGGGGGTGGCCGGCTGCAGCCGGGTCCGGGGTTCCGGCCCTGGAGCTCGGGGGGCGGCCGGGTGGCCCA | 12 |
| wt-142530-142650 | 5466 | CCGGGTCCGCTGGGTCCGCTGCCCCGCTCCGGCGGGGGGTGGCCGGCTGCAGCCGGGTCCGGGGTTCCGGCCCTGGAGCTCGGGGGGCGGCCGGGTGGCCCACCGGGTCCGCTGGGTCCG | 12 |
| wt-142650-142770 | 5467 | CTGCCCCGCTCCGGCGGGGGGTGGCCGGCTGCAGCCGGGTCCGGGGTTCCGGCCCTGGAGCTCGGGGGGCGGCCGGGTGGCCCACCGGGTCCGCTGGGTCCGCTGCCCCGCTCCGGCGGG | 12 |
| wt-142770-142890 | 5468 | GGGTGGCCGGCTGCAGCCGGGTCCGGGGTTCCGGCCCTGGAGCTCGGGGGGCGGCCGGGTGGCCCACCGGGTCCGCTGGGTCCGCTGCCCCGCTCCGGCGGGGGGTGGCCGGCTGCAGCC | 12 |
| wt-142890-143010 | 5469 | GGGTCCGGGGTTCCGGCCCTGGAGCTCGGGGGGCGGCCGGGTGGCCCACCGGGTCCGCTGGGTCCGCTGCCCCGCTCCGGCGGGGGGTGGCCGGCTGCAGCCGGGTCCGGGGTTCCGGCC | 12 |
| wt-143010-143130 | 5470 | CTGGAGCTCGGGGGGCGGCCGGGTGGCCCACCGGGTCCGCTGGGTCCGCTGCCCCGCTCCGGCGGGGGGTGGCCGGCTGCAGCCGGGTCCGGGGTTCCGGCCCTGGAGCTCGGGGGGCGG | 12 |
| wt-143130-143250 | 5471 | CCGGGTGGCCCACCGGGTCCGCTGGGTCCGCTGCCCCGCTCCGGCGGGGGGTGGCCGGCTGCAGCCGGGTCCGGGGTTCCGGCCCTGGAGCTCGGGGGGCGGCCGGGTGGCCCACCGGGT | 12 |
| wt-143250-143370 | 5472 | CCGCTGGGTCCGCTGCCCCGCTCCGGCGGGGATGGGGGTGCGCTCCCAGGCCGGACCCTGGTGCCAGGCAGGGACCCCGCGCCACCCGCTTCATGGGGGGGGAGGCCGCCGCAAGGACGC | 12 |
| wt-143370-143490 | 5473 | CGGGCCGGCTGGGAGGTGTGCACCCCCCGAGCGTCTGGACGACGCTGGCGAGCCGGGCCAGCTCGCCTTCTTTTATCCTCTTTTTGGGGTCTCTGTGCAATACCTTAAGGTTTGCTCAGG | 4 |
| wt-143490-143610 | 5474 | AGTGGGGGGCTTCTCATTGGTTAATTCAGGTGTGTGATTTTAGCCCGTTGGGTTACATTAAGGTGTGTAACCAGGTGGGTGGTACCTGGAGGTCATTCTATTGGGATAACGAGAGGAGGA | 4 |
| wt-143610-143730 | 5475 | GGGGCTAGAGGCCCGCGAGATTTGGGGTAGGCGGAGCCTCAGGAGGGTCCCCTCCATAGGGTTGAACCAGGAGGGGGAGGATCGGGCTCCGCCCCGATATACCTAGTGGGTGGAGCCTAG | 8 |

| | | | |
|---|---|---|---|
| wt-143730-143850 | 5476 | AGGTAGGTATCCATAGGGTTCCATTATCCTGGAGGTATCCTAAGCTCCGCCCCTATATACCAGGTGGGTGGAGCTAGGTAGGATTCAGCTAGGTTCCTACTGGGGTACCCCCCTACCCTA | 4 |
| wt-143850-143970 | 5477 | CCTTAAGGTGCGCCACCCTTCCTCCTTCCGTTTTAATGGTAGAATAACCTATAGGTTATTAACCTAGTGGTGGAATAGGGTATTGCAGCTGGGTATATACCTATAGGTATATAGAACCTA | 4 |
| wt-143970-144090 | 5478 | GAGGAAGGGAACCCTATAGTGTAATCCCTCCCCCCCCTACCCCCCCCTCCCTTACGGTTGCCTGAGCCCATCCCCCACCCCAGCACCCCGGGGTGACGTGGCACCCCGCGTGCCTTACTG | 8 |
| wt-144090-144210 | 5479 | ACTTGTCACCTTTGCACATTTGGTCAGCTGACCGATGCTCGCCACTTCCTGGGTCATGACCTGGCCTGTGCCTTGTCCCATGGACAATGTCCCTCCAGCGTGGTGGCTGCCTTTGGGATG | 4 |
| wt-144210-144330 | 5480 | CATCACTTTGAGCCACTAAGCCCCCGTTGCTCGCCTTGCCTGCCTCACCATGACACACTAAGCCCCTGCTAATCCATGAGCCCCGCCTTTAGGAAGCACCACGTCCCGGGGACGGAAGCT | 4 |
| wt-144330-144450 | 5481 | GGATTTTGGCCAGTCTTCAATTTTGGGGAGTGGTTTTGTGTGAGCCGGAAGTTGGCAATGGGGTGAGGGTGGCGCTGGTTAAGCTGACGACCTCCCAAGGTCTCTCACCCTGGGTACACA | 4 |
| wt-144450-144570 | 5482 | GGTGGGGCGGCAGCCTCTAACTTTGGCTGTGGCCTCTATTTCCTCCCTTTCCTAGCCAGGGCCATGTGTTCCTGCATGTCTACTTGCCTCCTGTGGTGGCAGAGCTTGGCCCTGGGCCCA | 4 |
| wt-144570-144690 | 5483 | ACCCCCGCCTTGGGAGCCTGTAGGGGCCAACACCCTTGGTTTGTTTGTGTTCCTGTTTGCTGGCAACTTACTGGCAGCCGAGCAGATTCTAATGGGCGCCCGCCTTCTTTCTCTCTTGTT | 4 |
| wt-144690-144810 | 5484 | TTATTAATAGAATCTCAGCCAGGACCTATACCTGAGACTTCAAAGTCTGGTCCTGGGTTCTGAGACCCCCAAGATTTGTCATGCACACCTGCACACCTGTTGGTATTGGGTTTCTATTCT | 4 |
| wt-144810-144930 | 5485 | TGAGTGTGAAAGTTTGTAAAAAAATTCATAAAATGTCACTAATTCCTCTTACCTGTTTAGGGTATTGTGCAATTCTTCAGCCTGCCTATTTTCAATTTGCCTAAGGTGGCAATTTAAGAT | 4 |
| wt-144930-145050 | 5486 | GTGGTTAATTAACCATTTTCCTGTCTGACACCACTGCATGGGCAACCGGGTTCCATGGCACATTTAGAGATAAACATAGATGTCTTGTCTTGCTCATGTGCAGAGGAGGGGGTGTTGGTG | 4 |
| wt-145050-145170 | 5487 | TGCAATATAGTTTCTGGATTCCAAATTGAGTTGGGGGTGCTATTTTCACTATGGAATTAAATTACTGACATTAGACAGTGGACACCGGGCTATATGTGGGGATGTCTGTGGCTTGTCATT | 4 |
| wt-145170-145290 | 5488 | TCCTCTTAGAAGGTAATCCCCCATCTTAACTTCCCTTTAAATTGTGATGCAAGCCCTGGGTTATTTATAGAATGATTATCTAGGTTTGATAGTCTGAAGGCTGGGCAGAGAATGTTTGTA | 4 |
| wt-145290-145410 | 5489 | ATTTTTATTCACCTTCTTTACCCCCCACGAGTATCCAGTTCTAGAAGATCTCCTGATATCCCGGGCTGCCATTATTCCCTTGAGTGTTATAGCTTCCTCTTAACTTAAGCAAGAGCTCCA | 4 |
| wt-145410-145530 | 5490 | GGATGTTAGCTTTTTTGGTGGGGCTGGTTGTCAGGAAGAGGTTCCAGTGTTGTCCTTTATTTTTAGATGTTAGCTTTGTGTTAGGTTAGTATGGGCTGGGTATTCACTAGTGAAGGCAAC | 4 |
| wt-145530-145650 | 5491 | TAACACAGTTAGACGTGCTAGTTGTGCCCACTGGTGTTTATCCGGTCCCAAATGTCACCACAGAACACAGGGGGCTGGATTTGGCAGCAGCACTTGTGCTTTTGTTGATTTTTACCCGTG | 4 |
| wt-145650-145770 | 5492 | TATCAGAGTGGGGGATGCTAGCCAATTTAGCTTCCCCTCCCCTTAACAGGGGGTCTCGCGGGGTGCCAATTGTCGCCTGCCTTCCCCCGCTTCCCCTTGTTAACTTATAGCATGATAGGT | 4 |
| wt-145770-145890 | 5493 | AGGTCACCTAACGTGGAAGCCTGGTGGGTGATCCTTCCTCGGTAGGGAGCGCTTAGGGCTGTTGAGCTCAACAGCCCCACCTGGGTAAAATGTATGTTCTAAAGAGTTACCCAATTATAA | 4 |
| wt-145890-146010 | 5494 | CAAAACTGTTGTAGGGTAACGAAGACCTGATGGAAGTGGTATTGTTGCCGTTGAAAGACGGGTGTCCTGGCTCAAGTTCGCACTTCCTATACAGTGTTAAAGCCTTGTATCGGAAGTTTG | 4 |
| wt-146010-146130 | 5495 | GGCTTCGTCCCAGTGTACTCGATAATGTCGACTGCTGCGAAAGGTTTGGACCGTCTTCCAGTAGGTGTTGGGGGTCCCAAATCACGAGGTTAGGCAGGTGCACTTGGCTCTTTAGGAGGG | 4 |
| wt-146130-146250 | 5496 | ACCCTTAAGCCAGACAATGTAGTGCCCCTTTTTTTTGCAAATTGGCCTTATTATTAATTTCTTGTTAACACTAATTCTGTTCTATGACCCTGTGTTTTTCAGATGCCGTTGAACGTGTCA | 4 |
| wt-146250-146370 | 5497 | CTGAGCTGAATTTGGACGCAGCTACTTGACCTTTGCCCCCGTGCCTCCAGCGCTGATAAGTGCTGCGTCCACTTTGTGTTACAGGTGGGCCAAACCTCCAGAATATCAATTGGTGGGGCC | 4 |
| wt-146370-146490 | 5498 | TTGGTGGGCTGCATAAGGCAGTAGGTTTGAGGTGACCTACTTGGACCATGTGGATCCAGTGTCCTGATCCTGGACCTTGACTATGAAACAATTCTAAAAAAATGCATCATAGTCCAGTGT | 4 |
| wt-146490-146610 | 5499 | CCAGGGACAGTGCACTCGGAAGTCTCATCATCTCCGTTTGTGTGTTTAGTGTGGCCAGTACGGCCACCCCTGTGCCACGCCCTGGCATGCTGCTGACATCTGGCCGCCAATTTCAGCGGG | 4 |
| wt-146610-146730 | 5500 | CCCTTTTCCCCCTTGTTCACCCCATAGCAAGAAGGGTAGGTTACATGGGTATTTTCCCATCAGCACCTGACTGGCCGGTGCAATTAGAGGAGAGGGCAACAACGCAAGGCTGTTGTTTTA | 4 |
| wt-146730-146850 | 5501 | TTTGGGTTACAAGAGCTGCGGCGGTCGATGGGTTCACTGATTACGGTTTCCTAGATTGTACAGATGAACTAGAACTGTCACAATCTATGGGGTCGTAGACAGTGTGCTTACCAGACTTCC | 4 |
| wt-146850-146970 | 5502 | ATGGAAGATGTGAATTTGCTGCTAGCTATATGGGTGGTGCTATGGGCTCCCTAGGGACTCATGTAGTGGGGCTTTGTGATAGCTAATGAATGTGGCAGCTGTTGTTTGTACTGGACCCTG | 4 |
| wt-146970-147090 | 5503 | AATTGGAAACAGTAACTTGGATTCTGTAACACTTCATGGGTCCCGTAGTGACAACTATGCTGAATATCTTGAATATGGGAGGAGGGGGGCTTTGGGTTCCATTGTGTGCCCTTTCCTGGC | 4 |
| wt-147090-147210 | 5504 | CAACGTGAGGGTCCTAGTGTTATAGGGCGTGGCAGTTTTCTTGAGGGCTAATAACCCGGGTGAGGCGGTTGTCACAGGTGCTAGACCCTGGAGTTGAACCAGTACCACTCGGTTACAAAG | 4 |
| wt-147210-147330 | 5505 | TCATGGTCTAGTAGTTGTGACCCTGCAAAGCTACGTGGGGATGAGCAGCCAGGGACTTTGGTTGGCAAGCAGACAGGCGGCGCATTGGAACCCCAGAGGAGTGTCCCGGGGCCACCTCTT | 4 |
| wt-147330-147450 | 5506 | TGGTTCTGTACATATTTTGTTATTGTACATAACCATGGAGTTGGCTGTGGTGCACTCCATCTGGTAAGGGGGCTGGTGCGGACGCCTGTGTTTAGTCTATGCCAATGTTTACCTGCCTTG | 4 |
| wt-147450-147570 | 5507 | GGTTACTATTCCAAACGACCACACCTTTGAGGACACCTGGAGCCCTGATCATTCTCGGCTTTTACTGCCACCTGGCTTCTGTTGGGTCAGACAGTTTGGTGCGCTAGTTGTGTGCTTAGC | 4 |
| wt-147570-147690 | 5508 | AGCAACGCACACCAGGCTGACTGCCTTAGCAGTGTGGCCCTTTATTGTGGCATCCTAAGGAGGGATTCTGGAGTGCCTTTCGCGTGAAGCATGCCCTGAGACGTACTCGAGTTAGGACTT | 4 |
| wt-147690-147810 | 5509 | AATCGCTCCTGTGCCGCTGGATGAGGGAGCGCCAATTTGTACATCCTAGCTCTGGCCATAGAGTTAGCCCACCCTTGTGTCTCCCTTTGGCCTTTGCGGTGCCAATTTCCGGTGGTTTCC | 4 |
| wt-147810-147930 | 5510 | CTTTTCCGCCCGTTTATCCAATAGCATGTAAGAGAGGTTGCCTAGATTTGGCAACTTTGAGGGAACGTTCCGTGTAGCTGGTGACCTAACACCCGCCCATCACCACCGGACAGATTCTGA | 4 |
| wt-147930-148050 | 5511 | ACTTGTCCTGTGGTGTTTGGTGTGGTTTTGGGGTACGCAGGAGTACGTTGGAATGCTTTGGAGCCGAGAGGGATGGGCCCGCTTGTGCGCTTATGTGTTACACGGTGCCAATAACCGGCC | 4 |
| wt-148050-148170 | 5512 | CGGTGCGGCTGCCCCGTGACCCGTGGGCCTTACCTTCCTGGCCATCGGGGGACCCTGGTGCTAGGGTCCCTTGTGTTGCTTTCTGCCATAGGGGGGAAAGCATCGCCTTCAGAATTGGCT | 4 |
| wt-148170-148290 | 5513 | GCTCCGTTGGAACATTTGAGGCCTACTGTATCCGTGTCCTGACAACATTCCCCGCAAACATGACATGGGTTAATTTAAACATGTTTTGTTTGCTTGGGAATGCTCTTAGGGCCTGGAAGC | 4 |
| wt-148290-148410 | 5514 | TTGTCATTGGATTCATCGTTTCCTGAACTACAGGCGTAGGGCCTATTGTAGCAGGCATGTCTTCATTCCTGCGTACCGAATGGCATGAAGGCACAGCCTGTTACCATTGGCACCTTTTTT | 4 |
| wt-148410-148530 | 5515 | CCATGTAAACCTCCGTGATCCTGGGTCCTTTGGAGACTCAAGTGTGAATTTGTTTTGGTGTTCGGCGCCAGGGTATCTCGACGTTGGAATGTCAACTCAACTTGGGCACCTCGATAACCG | 4 |
| wt-148530-148650 | 5516 | GCTCGTGGCTCGTACAGACGATTGTTTGGCTCTGTAACTTGCCAGGGACGGCTGACGATGTGTTTAGTCTGCCACTTGCATCCGGCGCTTTGGTTACTCGGGAGACTAATGGGGGGTGTG | 4 |
| wt-148650-148770 | 5517 | GTATGGCACAGGCTGGGGGTGAGTCTGGGGATGTCCCTGGGCGTTGCTGCAGCCCATTCGCCCTCTGGGGATGAGATGTTCAGGGGTGGCCGGTACCCTACGCTGCCGATTTACATAATA | 4 |
| wt-148770-148890 | 5518 | TAAATTGTAAATGCTGCAGTAGTAGGGATCTGGACGCGCGACCTGCTACTCTTCGGAAACGCCAACCCAGGAGCGTCGCCTCTGGCCCCATACTCCCGCCATGCGACTGCTCGCCCCCTC | 4 |
| wt-148890-149010 | 5519 | CCAGGCCTCCCTGGTGAGCCCTTGCCGCTCCCCGCATTCCTGCTTTCGGCGCCCCTGCGGATCCCGATGACAGCAGGCCTTTCCTTCCCCCGTTAATGAAAAGAATGACAGTGAGGTTGT | 4 |
| wt-149010-149130 | 5520 | GACAGAAGGACAGCTTTATTCAGTTTACAGAGTGCCCTCGGAGGCTACGATATTCCCGTTAAATGTCTTGTTGATTCTCTCAAAGGTGGGGAGGGAGGAGCTCTCCACAACAATGTTCCC | 4 |
| wt-149130-149250 | 5521 | TGGCAGCGTGAGCGCGCAGCCCTGCCGTTGGATGTATCTTCTCATGATGGTGCTGATAGAGGGGTCTCCGGCGTAGATGAAAAAGGCCTGGGCCATGCTCTGGCCGGTCACGATCGTTAT | 4 |
| wt-149250-149370 | 5522 | GGGGTTGTTGGAAATGTTCCGGACCGTCAGCTTGAGGGTCTGGCCCGGCTTCCACTCCTGTGGGTAGACGTAGAAGACCGGGTTGGAGGAGTGGGACACGACAACGGCCGTAATCTTGGA | 4 |
| wt-149370-149490 | 5523 | GCTCAGGGGGGCCTCGTAGGTGTTGTTGTATTCCAGCTCCGTGATGAAATTAGGAGGAATAATCACAGGGGAGCCAAAGTAGCGGATGTCTGTGGATTCCCCGTCCCAGCGCCAGTGGCT | 4 |
| wt-149490-149610 | 5524 | CTTAGGGTAGGGGTTGTAACGGAAGGCAATAATCACATCATCCAATAGGGTCATGCCCACCTTGACGTTCAGCGGGCCCTCTCGTTTCAGGTCCGGCGTGTCCACGGAGACTCGGACGTA | 4 |
| wt-149610-149730 | 5525 | GCCCTTACCGCGGCGTATGGCGTTTACCGGACACACCTTCCCCGGGAATGTGTGAATACGGGCGTATGACTTTAGAAATGGGGGCGTGTGCTGCGCCAGCAGGTAAGGCAGGCACTCGTC | 4 |
| wt-149730-149850 | 5526 | CTGGCTGGTGACGGGAGAGCCACTGAGGAAGATCTGGGGCTCGCTGGTGTTTAGCTTGTCCCCGCTCTGGGTGCAGGAGCGTGTCAGCTGAATGTCGCTCTGCCCGGGCAGAATCTGCAG | 4 |

| | | | |
|---|---|---|---|
| wt-149850-149970 | 5527 | GTAGAGGTAGGGGTTCTTGACCAATCTGATGGGCACAATGTACCAGGTAAACTTCCCTTTCTCTATGAACAGGCTGCGCGGATTCAGGACGCTTAGCACGATGTCCTGGTCAGAGTGCAT | 4 |
| wt-149970-150090 | 5528 | AACGAAGAAGGGCTTGAGGAATACCTCGTTGTCTTCCGCTCCAAAGAACAAAAACGCGACCGTAAAGTAGCGGCTGCCGTAGGTGGTCGTGTTGAAGGAGAAAGAAGGTAACTTGAAGCT | 4 |
| wt-150090-150210 | 5529 | GAGTATCTGGCCCACCGAGGGGCAGGGAGGCAGCTCTTGGCACTGCGCGTCCAGCTGCAATACCTGCTTGTTGGTGACGCGGACGTATGAGGGGAAGATCTCGTACTTCCACACGCCTCT | 4 |
| wt-150210-150330 | 5530 | CATGAACGACGTGTCTGGTTTTTCAGTGGGCCGCAGGCGGCGGAGGCTGTTCCTGAACGACGAGCGCCGGGACGCTAGTGCTGCATGGGCTCCTCCGGGGTAAGCTTCGGCCATGGCCGG | 8 |
| wt-150450-150570 | 5532 | GGCCAGAAGAGCGCAAGTCCAAGTCTGGTGCTGGGGCCGATGTGCAGCGGTTTGTGCCCGCAGTTGTAGACTGTCATTTTTATGGGCGAGTGGGCGGTCCACACGCGCGGGCGCAGCACC | 4 |
| wt-150570-150690 | 5533 | CATTGGTCGCACGCCGCCTCCTGGAATGTAAACCCCCAGAGAGAGGGCGTGCCGCCCTGGAGATGGCCCTGTGCCATCACATGTATTTCCTCCTTGGGTGGAACAACGGCGTCGTGCTCC | 4 |
| wt-150690-150810 | 5534 | GGGTGGAGGGGGAATAGCGTCCAGGCATCTTTCAGGGTCACGAGACCGGGGTCCATGCTCAGAGAACAGCCCTCCCGGGCGGTGGGCGGCCCGGGCTCCAGCAGAACGTCGCAGACCCAG | 8 |
| wt-150810-150930 | 5535 | CCCTCCTCGGCCCTGTCCACCTGTATGTCCAGGTGCACGGACCCGGAGGCTGCGTCTCGTGACATGGCCAGGCCTGGTGCCAGCCGACCACGTCCCGTGTCCCAGCCGAGGCCGCGCCAG | 8 |
| wt-150930-151050 | 5536 | AGCAGAGCCCGGGACTGACTCAGGGCCACATCCCCTCGGCCCGCGGACGCCGCCTCGCCAGCCCCCGGGCCTTCATGGGCCCGCTTTCTACCTCTCTCCGGCACCCCAGCCTGGTCAGCC | 8 |
| wt-151050-151170 | 5537 | GCAGAGGAAGCATGACCTTGGGGTGGGACGGGGCAGGCGTGATCCTGGGCGCAATCTTTGCCGATCCCCACACCTTCACTCCTTGTTAGGTTGATAGAATGTCGGTACCACGCCACGGGG | 4 |
| wt-151170-151290 | 5538 | GGCGGGCCCGCATAGGGAAAAGCCAGGGAGAGCGATGTGGGCGAGGATGGGCTCAGGCGGCCCCAGACACGCAATTTGCCCCCCTGGGCGGCCGCAGCCTGCCCCTCGGCGGCCCGTGCC | 8 |
| wt-151290-151410 | 5539 | CCAGCTCCGTCACGGGGGGCGCATAGGAGGGGTATATCTAGGATAGCCGCACCTACACAAATGAGACACAGACACAGGTCGTGAGGATTTAGGCAACGCAGGCTTGTCTTTATAGTTACA | 4 |
| wt-151410-151530 | 5540 | AACATGGGAGCGTGCACCTGGAAGATGCAGCTGGGGTAGATCTTTACATCTTTACAGGGCGCAGCGGCCGCCAGACACTGAAGGGCAGAGTTCACGGCGGGCACCTCCCAGAGGGAGCCC | 4 |
| wt-151530-151650 | 5541 | ACCAGCCCGTACCTGGCCACGGCCAGGGCCTCGTAGGCCGAGACGGGCAGCCGGAGCTTGTGGTACTGTCCCTCCGGCAGGTGGAGTGGGACACAGTTAGAGAACATTAGTCCTCTGGTC | 4 |
| wt-151650-151770 | 5542 | CCTATCTCCACCCGCCAGGCCTGTGTGTCAGTTTGCAGGGCCATCCTCGCACTCAGGTGGACTGGCTAGGCACCCTTCTGAAGTATCTGGCGGTGACTGTCACCTGGTTCTTGAGAGAGT | 4 |
| wt-151770-151890 | 5543 | CCATAAAATGGCTGAAGCTCCAGGCGTATAGTATAATGAGCAACAGGGCCAAACAGGCGGCGGGGCCTGGGTAGTAGCGGGCAACGAGAGACTCTGTGCAATCAAACCCCAGGCTCCCGG | 4 |
| wt-151890-152010 | 5544 | CCTCACCCAGGAAGAGCAGCGGCAGGGACAGCATAAACCAGGAGAAGGCGCAAATGAGTCCGGTGAAGGTGACGTTGCATATCAGGCGCGGCTTCCTTCCGAATTTTGTGCGCAAAAGTT | 4 |
| wt-152010-152130 | 5545 | TCCAGATGATGATAACTGTGAGGAGGACGATCAGGACTGCCGCCAGTAGGTAGCAGCCGGCTTTCAGTCCTTGGACGGCCGTGTGCATGCCTTTGGTGGGGCCTTCCCTGCACATGTTGG | 4 |
| wt-152130-152250 | 5546 | GGCCTCTGTTGAGATTGGCGTCGGGGCCCATGGTAATGAGGAGGATGATAATCAGCAGGAGTACCAGACAAAACACGCCCATCAGGTACAGGCACACATTTCTGTGGGAGGTTCTCTTGG | 4 |
| wt-152250-152370 | 5547 | GCGTTCGGCTGAACAATGCTAGGGTCTTCTCCAACGCCATACCCAAGTGAGTCCATACGGAGCACATCAGGCCCAAGAACATCATGTTCTGGGTCAAAAGGCAGAGACCGGTAGACGAGA | 4 |
| wt-152370-152490 | 5548 | ACTCCTGAATCATTTTTCCCAGCACCCAGAGCAGCAGTTCTATGAGAAGAGCTATCAGCCAGACATCCATTCGGTGAACCAATTTTCTTACAAAGATGATAAACAAGATGCCAGCCAGTG | 4 |
| wt-152490-152610 | 5549 | TTAGCAGAATCAGCAGGACGAGCAGCAGGCTTGTCATGCCGCTGAGGAAGGCGCTGTAGGATTTAGTGCACGCATCTTCCGTTGCATTGACGGAAGTCATGTTGGCCACCAGGGTCCCCA | 4 |
| wt-152610-152730 | 5550 | CGGTGGACCCGGGGGCCATGGTGGAGAGCATCTTGCTGGTCAGAGCCAGACTGGGTGGTGTCTGCAGCAAAAGAGGAACTTGCCCAGGCAGTCAGTTATTTTGCATGCCACCTCCCTGCC | 4 |
| wt-152730-152850 | 5551 | TGGTGGACTTCCAGACTATTTTCTGCATTCGCCCTTGCGTGTCCATTGTTGCAAGGAGCGATTTGGAGAAAATAAACTGTGAGTTTCACAGATCCACGGGCCACGCTCCCCTGGGGGCTT | 4 |
| wt-152850-152970 | 5552 | CATGATCCCACCGCCTTTCCCGATGATGATGACAACCGCGGCTGTCTGAAGCGGCTGACGAAATCGGTTGAGATTCTGATGAGAGGCTTGGGGGGGTCTTTGCCCTCAAGGCGAGGCTCC | 4 |
| wt-152970-153090 | 5553 | TTCTCCTAGGAATGCCGAGCCCCCTGCACTAGCTTCGCTCCACTGGGGATCTTTGCCAGCCTTCATACTAGATTCAGCGATCCCCCGGTTGGGAATCTTCGCCAGCCCCCCGTCCTGCTA | 4 |
| wt-153090-153210 | 5554 | TCCCGCTCGTCGCCGCGCCTCCCATGCTAAGGGCCCCCTTCCTTTCCCTTGACTTTGGGGATATTCGGAGTCTGCTCTCGCCGCTCTCTTCTCTCGTTTAAACGAGAGAATAGTAGTAGG | 4 |
| wt-153210-153330 | 5555 | GTCCAGTCTCAGGCCCCCTCACTTTGGGTCTTAGAATGGTGGCCGGGCTGTAAAATTCTGGAGGACGGAGAGGGCGGCCCCGGAGTTGTTATCAAAGAGGCACTGGAGGATGTTGGCCGC | 4 |
| wt-153450-153570 | 5557 | CACAAAGACGTACTGGATGCGGTCGTGGATCTGGGGCAGTTCCTCGTTGCGCTCGACGAACTTCTGGTAGACGGCCAGGTGAGGCATCTGGGTGCTCTTGTAGGCTGAGAGCTTGCGGCT | 4 |
| wt-153570-153690 | 5558 | GAGCTCCGTTGAAAAGCAGAGCTCCCCCATGGGGACCCTGCCTTCACGGAGGTCTGTGTAGGCCTGGTTTAGGATGTCAATGACGGGCAAAAAGCCCACAGGTAGCCCTTGTGTAAATGA | 4 |
| wt-153690-153810 | 5559 | CTCTTGGAAGGGCCGGTGGGAGAGGAGGCTGGCCGCCTCCTTTACCCGGGCATCCGCCAGCACCAGGTCGAGCACGCGCCGGCAGCGTGTCTGCACAAACTTGCAGGCCGTCTTCCGGAC | 8 |
| wt-153810-153930 | 5560 | GAGCTCCACCCCCTTCATCAGGGTCTTGCCGTCCGTCAGCACCCCCACATATCTCTTCTTTGTAATCAGCATCAGGCAGGAGAAGGTCTTCTCGGCCTCCAGGGAGATGGGGGCCACAAA | 4 |
| wt-154050-154170 | 5562 | CTCGGGGTTGAGGGGCGCCCAGGCGTCCGGGGAGGGGGCCAGGGCCTGCAGGTTGGCGGGGCTCAGGGCCTCCACGAAGGCCTTGGCCCGCTCCAACATCGTGCGGCCCTGCAGCGTCAC | 8 |
| wt-154290-154410 | 5564 | ATCCTCGCAGGCCGCCAGCAGCTTCTTGATGGCCTTGCGCTTGGCCAGCCAGGAGGTCAACAGACTAGCCAAGAAGGACTCGTGCACGTGCTTCTTTACAAAGTGGTAGACGCCCCCCGT | 4 |
| wt-154410-154530 | 5565 | GAGCCTGAAGGACTCATAGTCTTCTCCCGGGCGCAGGCCGGCTAGCCTGTGCTCTTCTCCCGGCGTTATCATGGTAGAATAACAGAGATTATGAGCCTGAATGATGCTCGGGTAGAGGCT | 4 |
| wt-154530-154650 | 5566 | GGCAAAGTCCACCACCAGAACCGGGGAGTTGTAGAATCCGGACAGGGGCTGGATGACGGTGGCCCCCTGGTAGCCGTCCCGGTCAGAGGCCGAGGGCATGGGCAGGATAAAGTTTTCCTT | 4 |
| wt-154650-154770 | 5567 | TTGGGCGGCCGCCAGGAGGCAGGAGAACACGCGGATCTGCTGCCCATCGTCCAGCACCCGCCTGCAGGGGATGTGAGCGATCTTGGCAATCTCTGCCACCTCCACGTGGATCACGAAATG | 4 |
| wt-154770-154890 | 5568 | GTTTAGCAGATCCATGACCAGGGCCGAGTCCTGCACGCAGTACATGCCGAGCCGCCTGCGCCCCTCGGGGCCCGCTGCAAAGAGGCGAGGAATCTCCTTGTAATGCACATCCTCCTTCTT | 4 |
| wt-154890-155010 | 5569 | GGCCCCCAGTAGGTGCCTGGCTACTGTGTCCAGCTTGTAGTCTGAGAGGCTGAGCTTGTCCCGGCACACGGCGTACATGTCGATGGGGATGAGGCCGGTGATGCGGACCTTGGTGTTGGC | 4 |
| wt-155010-155130 | 5570 | CCGCAAGAAGCCCTTGCCCGCATCATGGGGTCGCCTGACCTCGCAGACGCCCCCAGCCCTAATTTTGCCCAGAGAGGCTGGGTTGATGCTGTAGATGTGCCTGGCTCTGTCCAGAATGTA | 4 |
| wt-155130-155250 | 5571 | GGGCCAGTCAAAGTTGGCCACGTTGTAGCCGGTCACAATCTCCACGCTGAGGTCTCTGATGAGCTGGAAGAAGGCGTAGAGCATGTCCAGCTCCGATGGGAACTCGTAGACCTCAACCCC | 4 |
| wt-155250-155370 | 5572 | CTCTATGTCTTCGCAGGTGCCCAGCGTCAGCAGGATGCGCCTATAGCGCCCGGCCTCCTCCCCTGTCGACCAGAGGACGCAGGATATCTGCAGGATCAGGTCAGCCTCGTTGGTGGCCGT | 4 |
| wt-155370-155490 | 5573 | GGGGAAGCCCTCCTCCCCCAGACACTCGATATCGAAGGCCAGGGCCTGGTAGGAGGGCCAGGAGCTGTCTTCACGCCGGACCGAGAGGTCGCCCACCTCACAGTCGTACTCGAGCTCGGC | 8 |
| wt-155490-155610 | 5574 | GTACGAGTCCCGGTGCTGGAGGCGGGGGATGGCGCGGCGGCAGCTGTACCAGCCAAAGGTGACAAAGTCATTGTCCAGGACAAAGCGGCGCGTGGCATCCACGTTGGCCTCAAAGATCCG | 4 |
| wt-155610-155730 | 5575 | ACACCCGTGCTTGTCTTGCAGCCACGTGGCCACGTGACACACACTGTTGGGATGGGAGAGGGTGATCTTGTGGTAGTCGCCGGCATGGTTGCCGTAGCCCATAATGGAACGGCGCGTGAC | 4 |
| wt-155730-155850 | 5576 | CTTCTCCACCGAGACCCGGCAGGGGGTCCTGCGGTCGAAGGTGCTGGCCTTGAGGGCGCTGAGGACTGCAAACTCCACGTCCAGACCCTGAGGCGCGCTGGCGTAGAAGTAGGCCTGCTG | 8 |
| wt-155850-155970 | 5577 | CCCAAACACGTTCACACACACGCTGGCCCCATCGGCCTTGCGCCGGCCCAGTAGCTTGATGACGATGCCACATGGCACCACATACCCCTGTTTATCCGATGGAATGACGGCGCATTTCTC | 4 |
| wt-155970-156090 | 5578 | GTGCGTGTACACCGTCTCGAGTATGTCGTAGACATGGAAGTCCAGAGGGCTTCCGTGGGTGTCTGCCTCCGGCCTTGCCGTGCCCTCTTGGGCACGCTGGCGCCACCACATGCCCTTTCC | 4 |
| wt-156210-156330 | 5580 | CAGGTACTCCTTGTCAGGCTTTTTCAGAAGGCCTTTATTAGGTCTTAGGAAAGGGTTATAGAAGAGTCCCCCAGACATGGTTAAAACTCAGTCTCTGCCTCCCCAAGCAGTGCGGCGGCG | 4 |
| wt-156330-156450 | 5581 | GTCTCTGGATCGTGATAGCGTCTTCTGCGTAGGCCTGGAAAACGGTCCCTGGCTGCCTGCAATGCTCTGCTGGCCACTGAGGGTCCGGCCGCCCTCTGAGCTGCTCTCTTTTGCTCCTGG | 4 |
| wt-156450-156570 | 5582 | TTTTGCTCATGCAGCGCTAACATGATGGCTTGTAATTCTGTCTTACTAATGGGATTAATGCCTGGACCCTCACCAGAGGCATGTTGCTGAGCGAGCTCGTCGATCCCGGGGTAGAGCATC | 4 |

| | | | |
|---|---|---|---|
| wt-156570-156690 | 5583 | TGCACCGGCTGCTGCGACATCTGGCGCGTGCGCCTCGTGAGGGAAATAACCAGGATCACCACCCCCGCCACCAGGACCAGAATGAGCATGCCGCCGAAGGGGTTTTTGAAGAAGGAGATG | 4 |
| wt-156690-156810 | 5584 | AAACCAGAGACCAGGCTGCTAAACAAACCCCCCACCGTGCTGACTAGGTTGGTGATGGACTGACCCACGCTACCCAGACTGTCCATAAGTTCCCCCAGGCCGTCCACGAATTGATTTCTT | 4 |
| wt-156810-156930 | 5585 | CCGTTTGACACTGCATTGTCCAAATCCTTCCGCAGGCCGGCGATGTTTTGCGCCTGGAAGTTGTACTCCCGGAAGATGCCCTCCAGGTCAAAGACGTTGGAGGCACGCTGTTCGTCCCGT | 4 |
| wt-156930-157050 | 5586 | GAGTACAGCTCCAGGGAGGCAAAGTCAATGTTCTCGATGAGGGAGGTGTTTAGTGAGATGAAGGTCTGCAGGGTGGCAATGCCGTCCAGCTCGATGGTTTTAAAGTGGTGGTAGTCGTTG | 4 |
| wt-157050-157170 | 5587 | TAGACGTGGATCTCGTTGCCGGACTGGAAGTAGTACTGGCTGGTCGCCTGGCACACCTCCGTCATCTTTTTTGTGAGGAAGATCTCGTTGTCGGTGCCCAGCTGTCCCTCGTAGGTCTTG | 4 |
| wt-157170-157290 | 5588 | GTGTCGTTGATAAAGCTGAAGGACACCAGGGGGCGCGAGTAGCACATGGTCTCGGAGCCAGGGACCCTCATGCTCTTGCGCAGGGTGACGGTGGCCTGGTTAACGGGCACGCACTGGGAG | 4 |
| wt-157290-157410 | 5589 | ACTGAGATGACATCCCCCAGGCGCTTGGCCGCCACCGCCTTACCGTAGATGCTGGACATGACGGTGGTTGGATTAATCTTGGTTAGTTCTCTCAGCACCATGTTCTGCCTCTTCTGCTCC | 4 |
| wt-157410-157530 | 5590 | AGGCACCAGGCCCGCGCAAGGTCTCCCAGCATGCGGTTGATCTGGCGGCGCAGGGAGTCGTAGGCAAATTGGATCTGGACGGTGGCGGGATTGTTGAGGGTGCCCAGGGACTTCCCGGGG | 8 |
| wt-157530-157650 | 5591 | GCCGTGGGGGGCACCGGTGTGGTGGCGTTCCCCGCATCCCGCCTCCGACGCCTCAGAACGGCGGCGGGGGTGCTCCCGCGGGCCGCGGATGGGGCTGGGGGCGATGGACTGCTGGGGGGT | 12 |
| wt-157650-157770 | 5592 | GAGGAAGTCGGAGTGGTAAGCTCCGTCAGGTTCTTGACGGTGGCCAACGAGCGCGGGGTCAGAGGTAGCCAAGCTAATAACAATCCTCCGCTCGTTATAAAATATGTAATGGCTTCCTGG | 4 |
| wt-157770-157890 | 5593 | CCCTTCGTGTAACGATCCTGGACGGCCTCGTACTTCTCATGCATGGTCTTGTTCACCTGCTCTTCGATGCACTTGAAGGCGTCCGGGAGCTCTATGCCCACGGTTGTGTTGGTCACGAAG | 4 |
| wt-157890-158010 | 5594 | CTAGAGGTGCCCTCGTCAGTCACAAAATGTATTGACTTCCCTGTTTCTGTGGCGATGGTCGAGTCAAAGGTTTGCCAGTGTTGAAGCGGGCAGTAGGCTGTCCTGTTCTCGAGCTTCCAA | 4 |
| wt-158010-158130 | 5595 | GATAGCGTGTAAGTGCCCTTGTCCAGGAAGGCTCGGCGTTCGCCTTGCGGGTTCGTCCCTCGGTTGTCGTAGTCCACTATCTTGTAGTTAGTTCTCACGTGGAAGGAGTCTGCCCGCTCA | 4 |
| wt-158130-158250 | 5596 | TGGAAGGTTTCCTTATTTTTCCCGTCATAGAAAGGGGACATTTCCACAGTCTGCCCGGTGGTGGTCACAAAGAAGTCGAAGGGGCTGTTGGACTTGGCCATCATGTCAGTTATCAGGCAG | 4 |
| wt-158250-158370 | 5597 | TTGACGGTAGTTCTTGTTCTGTAAGTCCATATCAACCACCCGGGGGCGTCATAGAGCTCCGTCTGGCTGGCGTAGCGGCGCACCCCGTTGGCCAGGCCCCCGGTGGGCTTTAGGTTGACG | 4 |
| wt-158370-158490 | 5598 | GTGATGTTAACTCCGTCGCGGTCTACATACACGCGCGTCAGCCCATCTTTTGTCATCTTGACCGCGTTGTAGCACTGGTAGATGGTATCCATCTGGTCAGTTTCGTAGCTGTCAACGGAG | 4 |
| wt-158490-158610 | 5599 | AACTTCTCCTCGTGCCGGTTGGTCACGGAGTCCGCGTACCAGCCATTGTAGATGAGAATGTTGGTCACTATCTTGGTGTAGGAGCGGACCTTAAACGAGTAGGGAATAATGTTGTCTTTA | 4 |
| wt-158610-158730 | 5600 | AACACCATCAACAGGCCCTCCGTGTGATTCTCCCGCGTGCCAAACGAGGGACACTGGATGTCCGAGGAGAAGCGGAACAGGTCGCCGTGGCTGGAGAGCTCGCAGACTCGGAAAGGAAAG | 4 |
| wt-158730-158850 | 5601 | CTGGTTTGCTGACGCGTGGCGGTAGGCTGCACCGTGGTGGCGGGGGGTGCGGGCTGCTCTGGGGTCTGCGCACCGAGACGGCACGCCAGGGCGGCTAGCAGCACGACCACGCTTAGCACC | 8 |
| wt-158850-158970 | 5602 | CTACGCCGAGTCATCTCTCATTTGGAGGTGCAGGTAGAGAAGGGCATATAGATCCTTAAATACCCACCCCCTGCCCTTATACAGAAGAATTAGGGGGCGGTCAGAGTCGTACGTGAGGTA | 4 |
| wt-158970-159090 | 5603 | AAGCCCATCCGGGGGCAGGGCCTGGCCGGGGCTGACCGCGTCCGCCCGGCGCAGGATCAAGGACCGCCCCCAGGTCTTGTTGTAGAGGGACACGGTTAGGACGGCCTCGCGCAGCGCCCG | 8 |
| wt-159090-159210 | 5604 | GCACAGAATTTGCTGGCTAGATGCCAGTGAGCCCCCGGGTACGCTGTAGAAGCTGTTGAAGGAGGTCTCTATCCAGTCGCTCGGCTCGATGCCTGGCCATATCAGGGAAGTCAGGAATGC | 4 |
| wt-159210-159330 | 5605 | CTTCTGGTGGGGCAGCGTACCTGCGGCGTCACAGCAGCGAGCCAGGGCCACGTTGCTGGGTGGGGGAAAGAGCCCGCTCTCCTCCGCCAGGGGCCCCGTGATGAAGGTGTACAGGCTGTG | 8 |
| wt-159330-159450 | 5606 | CGTCAGCGCGTGCAGGTGCTCCGAGCTCAGGGTCTGGGTAAACAGGTGTGTTTTGATGTACTTGGAATTCTCAAAGGCGGCACCCTCGCCGGCGCGCCTGTCCTCCCAGGGACCCGAGAC | 4 |
| wt-159450-159570 | 5607 | GAAGGCCCGTCTGTAGAGGAAGTGGTTGCGCATGCGGGCCAGCTCCCAGTAGACCACGTCCCCCAGACGCGCAGGCACAGGGTCTCGGTCAGGGTCTCGCTCTGTTGCGCCAGGCAGGA | 8 |
| wt-159570-159690 | 5608 | CTGCAGCTTGGCCAGACCCTCGGTGGCCACCTGGCGCAGGTACTGCTCCTTGCGCTTGAGCGCGTCCGAGAGGGCGCCGGACGGGCCGGGCTCTCGTGCCCCAGCCGGCCGGGGCACCTC | 8 |
| wt-159690-159810 | 5609 | CGGGCTCTCCCGGGACGCCTCCTCCTCGCCTCGGCCCAACCGCTGCATGGCTCGGTTGAGCCGCGTGTACAGCTCGTTCCTCTTTTGCAGGATGGCCCGGTACTGGGGGTGCGCCGTGAA | 8 |
| wt-159810-159930 | 5610 | GGCGGCGGCGCAGTCCGCCTTCAGCGCCTCCACCGCGTCGCCCGAGGAGCTGTAGACCCCGCCGCAGAAGAGCCGCTCCGTGGCCCCGGGAGCCACGGCGTCAAACAGGTGAGTCAGCCT | 8 |
| wt-159930-160050 | 5611 | TGCCCCCGCCAGCGCCTCCTCGCAGGCCCCCCGCACCAGGGCCAGGCGACGCTCCCGGGCAAACAGGGCAGAGAGGCGGGAATGGCCGCCACCCTCCCCCTGCCCCGTTGCACCGATAGC | 12 |
| wt-160050-160170 | 5612 | ATGGCCGCCAGAGTTCCAATAGAGGAGCTCCGAGAGCTCCGCCACCTCCGGGGGCACTGTCGAGAAGACGTTGTAGGTGTCCAGCGCTCTGGTCGCCCCCTCTGCCTCCGGCCGCCCCGG | 8 |
| wt-160170-160290 | 5613 | GCCCGGGACCGCGCCCTCCTCTGGGCCGCCCGGCCTCGCCTTCTCCTCAGCCTCCAACAGGTGCCCGAGCCCCGCCTGGCGGACTTCATTCTCAAACAGTCCCGAGACCGGCTCCGGATT | 8 |
| wt-160410-160530 | 5615 | GAGGCGCGGGTCTGGGATGGGACTTACGCCCCGAAAGCGGCCCAGCAGATCCAGGGCCCGTTCCAGGCTCTCCAGCCCCATGGTGTGAGACATGCAATAAAACACGCTATTGATTCTCTT | 4 |
| wt-160530-160650 | 5616 | CATTAAAATCTCTATGTCATTTATTAGGCACAAACTTACATCGACTTTATGCCCCCCGTAAAACTCCACAGAGTACGCGACTGAGGGGGTACGGAGAGGCGGGACCCGGGTACCCTTTCT | 4 |
| wt-160770-160890 | 5618 | CTCCAGCTCGGCCAGCAGGCCGGCGTTGCGCACCACCGCAGCCACGTCCAGACTCCGGGGGTCCAGCCGGGTGCACACGCTCAGCTCAACCGCCAGGGCGTACACCTGGCTGTACGCCGC | 8 |
| wt-160890-161010 | 5619 | CGCCAGCAGCCCCGACATCGCCGCCCCAGGGGTCTCTAGACCTCGAGTCCGGGGAGAACGGTGGCCAGACGGCGCTTGCGTCTGCCCCCGGAGCCCTGCCCTCTCCACCCAGCAGCAGC | 8 |
| wt-161010-161130 | 5620 | CCGGCCGAGGCCTGCGACGCGGTGCTGACCGGCTCGGCCACGCTGATAAAGTTGTCCTGGGCTGCCCCGGGCCCACCCCACACTCCCTCCAGAAAGTCCCGAGCGGCCTCCGCCGTCCAC | 8 |
| wt-161130-161250 | 5621 | TCTATCCCGCTGGAGGCAATGGTCGCCAGGGTTTCTAGGACGCTGTCCGCCAGGACGGAGAAGCGGCCCAATAAGTACTCCGCGTCGTCCCTAGTCAGCGAGGCGCATGCCTCGCCCATG | 4 |
| wt-161250-161370 | 5622 | GCATCCACAAGGTTGCACACCACATCAAACACACAGTCTTCCTCCTGTTTTTGTGATATAATGGCCTCCAGGCCAGCCCTGATGTTCTCAATCTCATATGTGGTCGCGGCTTGGGTCCGG | 4 |
| wt-161370-161490 | 5623 | CGCTTCACGGTCAACCCTAGGGTGGGGGTGGCAAAGACAAACTTCTTCCGCATGGAAGAGCCCCCGGCCTGCTTGCGCAGCCCAGCCCCGGGGGCCTGCAGCAGGTTCCTGTCCACGCCC | 8 |
| wt-161490-161610 | 5624 | CGGCCCATAAAGTATCCCAGGTTCCCGGCCTGGAATATCTGGTTGTTGCCGTTGACCCCCGTGTACTTGTTGATGGTCACTGGCAGCGTGACAACCGGACGGGCCTTGCAGACCTGGCTA | 4 |
| wt-161610-161730 | 5625 | AGACAGTCTGTGGCCGCGCAGACCACCGTGGTCGCAGTAAGGGAGGAGGTGGCCTCCGCGTAGGCCGCTGCCGACTCCACCGCCCGCGTGCCCAGTACGTGGGGGTAGTCACGGGCGGGC | 8 |
| wt-161730-161850 | 5626 | ACCGACTGCGTCCTCGGCACCAGTCCCTGAATCAGGCTGATGTAGAACTGGGTCTGGCCGCACGCCTTCAGGATGGCGTTGTTGAGCCTCTGCTTGGCGTAAGTGACCAGGTTGCCAGGC | 4 |
| wt-161850-161970 | 5627 | ACCACATCTATGACGTTGCTCTCTTCGTGGGCCCGGGAGCCCCCGTCCACAAAGAGGGCCAGGTCAGAGTACTCCTCCGCGCTGGCCCCGCTGGGGACAGGGACCGAGCGCCGCCTGGAA | 8 |
| wt-161970-162090 | 5628 | AAGTTGTGCCACAGGTACAGGCTTGAGAGCTTAGTGTCCGGGAATAGGGTCTTGTGGTAGGTGTTGAGGAATTTCATGTAGGGCCCGTTGATGATGTAGTTCTCCCTCCTGGTAGTGGAC | 4 |
| wt-162090-162210 | 5629 | TTGATGAAGCTGTTCTGGAGGGCGGCATTCTCCCCCGTGAAGACCACCCTGTTCTTGATCTTGATGTTCCTGGGGCACAGCATCAGCACCTTGGACATGCGCACAGGCAGCCCGCCGGCCG | 4 |
| wt-162210-162330 | 5630 | TACACCCGGCCCTGCAGGGCCGCGTCCAGGTCTGGCAGGTCGCAGGTGGGCTCCCCATGCACCACCTTGGCCTCCTTGGCCGTGAGGACCCCCTTGTCGATGGCCAGGCTCCTAAAGTTG | 8 |
| wt-162330-162450 | 5631 | GTGCACAGCGTCTGGTAGTGACCCTTTAGCCACTCTGGGGGGCTCTGGCCAAGCCCGGGGTTGTCATTCTCATAGCACATACAGATGGGCAGGGAGATGTCCTGCAGGATGGTCAGCAGT | 4 |
| wt-162450-162570 | 5632 | GAGCGGTAAAACAGCTGGGTGAAGATGGGGCAGGCGGGCTGCGCAAAGGGGTTGCACGAGTACTGCATCACGTGGTAGCAGCTCTTGACCAGGTCCTTGTAGGTGATGTTGTTCTTGGCC | 4 |
| wt-162570-162690 | 5633 | ATGCTGTTCATAAACTGGACCACTTCGGCGTCCACCGCCGCATCCACGTCCTTGAACATCTTGACAAAGTCACGCGGGCCATGGGGCTCCTTCTCTAGCTTTCCTTCAGCGTCTATGCCC | 4 |
| wt-162690-162810 | 5634 | AGCCGAGACAGCCGCTCCAGCAGGTTCTGGTTCAGCTGCCAGTAGGTGTAGCGGGGCTCGTCGTCCGGCCGCTGCCCGTCGTCCTCCTTATCGATGAAGTTGAGAAAGTTGCCCAAAAAG | 4 |
| wt-162930-163050 | 5636 | GGGGCCCGGCCCTCGCAGAGCGAGCACATGGGACTGGCGGCCGCCCCCGCCACGTAGCTGCCCGTCTCCGGCACCGGGGTCAGAGAGCTCTTCTGTCCCTGGCAAAACTGCAGGTAGTAG | 8 |

| | | | |
|---|---|---|---|
| wt-163050-163170 | 5637 | GCATAGCGGGCAAGAAGGTTGGGCGAGAAGGAGGCCGCATAGACCAGGTGCTCCACAGCGTAGTTTCCCGGACCGTTGGTTCCGGTCACGTCTGGCCCACCCCAGCCCGAGAAGCAGGGT | 4 |
| wt-163170-163290 | 5638 | CGGCGGCAGGGGTCCCAGGTCCCCTCCTGCAGGGTCCCCAGGCCGTGGGTCATGTAGAAACTGTTAAAGAGACTCTCCTTGCCCTGACCGGTTGACTTCGAGACCCCCGAGACGTAGAGG | 4 |
| wt-163290-163410 | 5639 | ACGGAATTGGTGGCAAAGATCTGCGTGGACACGTGGGGGGCCAGGCTGGCATTATATCGGTGTAACGCAGCCACACGGGCCTCTGGACCCTCACAGTCGGCAAACAGGGGCCACGAGTCG | 4 |
| wt-163530-163650 | 5641 | GTTGAGGCTGGGAACTCCTTGAGGGGGGCCACCTTGGCCACCTTGGCCTGGTCCTGCAGGCTCTGCTTCTCCAGCAGCTCCACCAGCTTGCCCACCCGTCGGACGCGCAGCGCCTGCGCC | 8 |
| wt-163650-163770 | 5642 | AGCCCGGTGTACAGCGCCTCGTGCATGCAGCGGCTGAGGTCCGAGTTGTAAAACTGGCGGAGCTGGGGCACGCCCTCTGGGAACACCTCCTTGTCGTAGAGCGGGACCCTAACGCTCGCA | 4 |
| wt-163770-163890 | 5643 | GACTGCCCCACCGCTACCTCCTGTTTTAACGATGGAATGGCCACCAGGTTTCCGCTGTAGAGTCGCTCCTTGAAGGCCTCGGTTATTGCCACCGCCCCCAGGTAGGCAGAGGGATCTAGC | 4 |
| wt-164010-164130 | 5645 | GGGGGAACCATATGGGCATTGTGGAACACGTAGATGTCCCTGTGATAGGAGGTAGCGCGTAGGAGCCCGCAGTTGGGGTCGGGCCTCCTGTGCAGAGCCTTGACATTGATGCTGAAGCCC | 4 |
| wt-164250-164370 | 5647 | CCGCACGGACCCGGCTGGCTCTGGCTGCCCAGATTATCCTCGCTAGTCTGTGCACCCTGCATGATGCCCAAGGTATCGCCCCGGCCTCCCAGTCCCACATTAAATGTTACACTTTACTCA | 4 |
| wt-164370-164490 | 5648 | TCACGCAACACCCACTGTTTATTCATTTACAAAGATTTCAGGAAGTCAGTCAGGCTGGCCAGGGCCCACGTCACGGGGAACTGACGTCTCAGCGATCTTGGCATGCCGCCCAGCCTCGCA | 4 |
| wt-164490-164610 | 5649 | AACCAGAGTCTGCGATAGAGGGCCAGGTAGTGGGCGATTGCCCCCAGCACGAAGGCGGCGCTCTTGTGGTCATCCAGGTAGTTTCGCACCGCAAACACCACTGTGTAGCACAGCACCACC | 4 |
| wt-164610-164730 | 5650 | CTGAGCCGCGACCAGTAGTCGTAGTGGTCGTTGTACACTGCGCGCAGGACGCTGATGATGAGCCGTACGTGCGTGTCTTTGCCCCCGATGTCGGCTGTCCTGCAGGCCAGCTCCGCGTAC | 4 |
| wt-164730-164850 | 5651 | AGCTTCCTATCCTTCCTCAGGGAGGCCTTGATGAGCCGGCAGAGGACCAGGGCTGGCAAAGGCAGGTCTTTCTCATCCCGGGTGAACACCGCGTACATGGCCCTGAACATGAGGTAGCTG | 4 |
| wt-164850-164970 | 5652 | GACTCAGCCACCTTGTCGTCCGGCGGCGAGGGCGCGACCCACGCCTCGACCGGGGTCCTCACAAACACAGAATCTGTAGACTTGGCTGGCCTCATGGTCTCGTCAGGCCAGCTCACGGGC | 8 |
| wt-164970-165090 | 5653 | TTCAGGCTTATATGATAAAATGGGCGTGGCAGAATAGTATAAGACGCGAGGCCTGGGTGAGGAGAGTCCAGAGCAATGGCCAGGTTCATCGCTCAGCTCCTCCTGTTGGCCTCCTGTGTG | 4 |
| wt-165090-165210 | 5654 | GCCGCCGGCCAGGCTGTCACCGCTTTCTTGGGTGAGCGAGTCACCCTGACCTCCTACTGGAGGAGGGTGAGCCTCGGTCCAGAGATTGAGGTCAGCTGGTTTAAACTGGGCCCAGGAGAG | 4 |
| wt-165210-165330 | 5655 | GAGCAGGTGCTTATTGGGCGCATGCACCACGATGTCATCTTTATAGAGTGGCCTTTCAGGGGCTTCTTTGATATCCACAGAAGTGCCAACACCTTCTTTTTAGTAGTCACCGCTGCCAAC | 4 |
| wt-165330-165450 | 5656 | ATCTCCCATGACGGCAACTACCTGTGCCGCATGAAACTGGGCGAGACCGAGGTCACCAAGCAGGAACACCTGAGCGTGGTGAAGCCTCTAACGCTGTCTGTCCACTCCGAAAGGTCTCAG | 4 |
| wt-165450-165570 | 5657 | TTCCCAGACTTCTCTGTCCTTACTGTGACATGCACCGTGAATGCATTTCCCCATCCCCACGTCCAGTGGCTCATGCCCGAGGGCGTGGAGCCCGCACCAACTGCGGCAAATGGCGGTGTT | 4 |
| wt-165570-165690 | 5658 | ATGAAGGAAAAGGATGGGAGCCTCTCTGTTGCTGTTGACCTGTCACTTCCCAAGCCCTGGCACCTGCCAGTGACCTGCGTTGGGAAAAATGACAAGGAGGAAGCCCACGGGGTTTATGTT | 4 |
| wt-165690-165810 | 5659 | TCTGGATACTTGTCGCAATAAACGCACTTGCCTATTTCACCTTGTTTTAGTGTGGCATTGGGGGGGTGGCATTGCGGGTGGATAGCCTCGCGACTCGTGGGAAAATGGGCGGAAGGGCAC | 4 |
| wt-165810-165930 | 5660 | CGTGGGAAAATAGTTCCAGGTGACAGCAGCAGTGTGTGAAGATTGTCACAGCTGCTGGTTTGGAGAAAACGGGGGTGGGCGGTGATCAGGGAGAACAATTCCCCGGGGACACCTGCACGA | 4 |
| wt-165930-166050 | 5661 | GACCCCTGGGCTCTCAGGAACTCCGCCCAGGTCTTGCCAATTGGGGTGATCCTGTAGCGCCGCGGTTTCAGCATCACAGGTTATTTTGCCTGAAGCTTGCTGGGGCGTAAATCCCTCTCG | 4 |
| wt-166050-166170 | 5662 | CCTTGTTTCTCAGAGAGCATTTCAGGCCGGTTTTGCAGTCGCTGCTGCAGCTATGGGGTCCCTAGAAATGGTGCCAATGGGCGCGGGTCCCCCTAGCCCCGGCGGGGATCCGGATGGGTA | 4 |
| wt-166170-166290 | 5663 | CGATGGCGGAAACAACTCCCAATATCCATCTGCTTCTGGCTCTTCTGGGAACACCCCCACCCCACCGAACGATGAGGAACGTGAATCTAATGAAGAGCCCCCACCGCCTTATGAGGACCC | 4 |
| wt-166290-166410 | 5664 | ATATTGGGGCAATGGCGACCGTCACTCGGACTATCAACCACTAGGAACCCAAGATCAAAGTCTGTACTTGGGATTGCAACACGACGGGAATGACGGGCTCCCTCCCCCTCCCTACTCTCC | 4 |
| wt-166410-166530 | 5665 | ACGGGATGACTCATCTCAACACATATACGAAGAAGCGGGCAGAGGAAGGTAAGAGTGCCATCTATCTGTACTTTTATTTATTGCATCACAAGTCACATCAATAATAAGGGCGCCATCTAG | 4 |
| wt-166530-166650 | 5666 | CGGGAGATGTTATCCACACCATCCCAATTCACATCTCAGGGACAACAGGTCAAAGTTCTTTGTTGACACCCCCAGCGCTGGCTCCAGGGGGTGGAAGCGTTGGATGCAGTCCTCCGCATC | 4 |
| wt-166650-166770 | 5667 | GGGGCGGACGCCTCCTCCCAACGCGTTTCTGCGGATCAGTCGCTGGCTGGTGGGCATCGGAGTCGGTGGGCGGTCCTCCACGGGGACACGCTCCTTCTTGGCCTTGTTCTTTGACCTTTT | 4 |
| wt-166770-166890 | 5668 | GGACATTCTTCTGAAGGAACGGCGGAGAGTAGCGTAGAATCCAGCCAGTGGTCTACCCGGTCGCATGGTGGCTTCTTAGATGAGGAGCAGGCATAAAAGTCCAAACAGGACACAGAGTAC | 4 |
| wt-166890-167010 | 5669 | CACCAGGAGTAGTCTTAGTCTGCTGACGTCTGGGTCCTCGGGGCAGGGGTGGCTAGGCCTGGTCTCCGTAGAAGAGCCGGGCAGGCCGCAGGCAGAGGACTGCTGCTCTAGCAAAGCACG | 4 |
| wt-167010-167130 | 5670 | CTCCAGGACGTGTACCATCTCGAGAGTGAGGCACAGCTGTTTTCGTGGACTTTTATACAGTAAGGACAAGGAAAGAAGGCCAGAGGAATGTGGAAAGATGAGCGAGGACAGGTGTGGAGG | 4 |
| wt-167130-167250 | 5671 | TTTTGGGCTAGCTCTTAGTTTCTGGGTGTGAGAGAGGGATTAAAGTGCTTATGCGCAAAGAATGTGTCAACAACAGGTGTTCCTGCCTCTGCTGGCATGAGTTAGGTGTGGCTTGGGCTG | 4 |
| wt-167250-167370 | 5672 | AATCCAAATGTGTATTGGCACAAGATGGAAAGCAAAGTTGCTGGAGTTACTGGGTGGGAGACAGGGATGTATGTGGTCCCCCGCTGGTATGCCAGTACCCTGTGGAAGTAAGGGGCCTCA | 4 |
| wt-167370-167490 | 5673 | TCTGCCTGGTAGTTGTGTTGTGCAGAGGTCTGATGTGTGTAGGAGGGGTGGGTTCAACGCAGGGGCGTTGGTGGCGGAGTCTGGCAACGCCCGGGTCCTTGCTACCTGTGTGGTGTGTTA | 4 |
| wt-167490-167610 | 5674 | AGGGCTGGGTAAAGGTGTCTGCCAATTCTCGCATGTCCTCCTTTCCCCTTGTTTTGAAATAGAATATGAATGTGGCTTTTCAGCCTAGACAGACAGTGTGGCTAAGGGAGTGTGTGCCAG | 4 |
| wt-167610-167730 | 5675 | TTAAGGTGATTAGCTAAGGCATTCCCAGTAAATGGAGGGAGAGTCAGTCAGGCAAGCCTATGACATGGTAATGCCTAGAAGTAAAGAAAGGTTAGTCATAGTAGCTTAGCTGAACTGGGC | 4 |
| wt-167730-167850 | 5676 | CGTGGGGGTCGTCATCATCTCCACCGGAACCAGAAGAACCCAAAAGCAGCGTAGGAAGGTGTGGATCACCGCCGCCATGGCCGGAATCATGACTATGACCGCCGCCTCCGTCTGTCATCA | 4 |
| wt-167850-167970 | 5677 | AAGGCGGGCCCTGGTCACCTCCTTTGTTTTCAACCTCTTCCGTCAATTGTGGAGGGCCTCCATCATTTCCAGCAGAGTCGCTAGGGCTATGAGGCAGCGGGTCATGTGGGCCATTGTCAT | 4 |
| wt-167970-168090 | 5678 | CAGTGTTGTCAGGGTCCTGTGGGCCATTGTCATCAGTGTTGTCAGGGTCCTGAGGCAGCGGGTCATGTGGGCCATTGTCATCAGTGTTGTCAGGGTCCTGTGGGCCATTGTCATCAGTGT | 4 |
| wt-168090-168210 | 5679 | TGTCAGGGTCCTGTGGGCCATTGTCAGGACCACCTCCAGGTGCGCCTAGGTTTTGAGAGCAGAGTGGGGGTCCGTCGCCGGCTCCACTCACGAGCAGGTGGTGTCTGCCCTCGTTGGAGT | 4 |
| wt-168210-168330 | 5680 | TAGAGTCAGATTCATGGCCAGAATCATCGGTAGCTTGTTGAGGGTGCGGGAGGGAGTCATCGTGGTGGTGTTCATCACTGTGTCGTTGTCCATGGTAATACATCCAGATTAAAATCGCCA | 4 |
| wt-168330-168450 | 5681 | GAAACAGGAGGAGCCAAAGGAGATCAACCAATAGAGTCCACCAGTTTTGTTGTAGATAGAGAGCAATAATGAGCAGGATGAGGTCTAGGAAGAAGGCTAGGAAGAAGGCCAAAAGCTGCC | 4 |
| wt-168450-168570 | 5682 | AGATGGTGGCACCAAGTCGCCAGAGCATCTCCAATAAGTAGATCCAGATACCTAAGACTGCGTTGAAAAAAGAGTGTTAGGGTTGGAAAAGTGGGGGTGTGGTAAATAATTCCTAGGGAA | 4 |
| wt-168570-168690 | 5683 | TGTTAGATCTTACCAAGTAAGCACCCGAAGATGAACAGCACAATTCCAAGGAACAATGCCTGTCCGTGCAAATTCCAGAGAGCGATGAGCAGGAGGGTGACTGGGGAAAGAGGAGAAAGT | 4 |
| wt-168690-168810 | 5684 | GCGTTAGAGAAGGAAGAGTAAGGGAAAGGGGGTGTGGGGCAAAGGGTGTAATACTTACTCATCAGTAGGAGTATACAAAGGGCTCCAAGTGGACAGAGAAGGTCTCTTCTGAAGATAAAG | 4 |
| wt-168810-168930 | 5685 | ATGATCAAAATTATAATTATAAGCATGAGAGCAAAGGAATAGAGGACAAGGAGGGCTCCTCCAGTCCAGTCACTCATAACGATGTACAGCCAAAACAGTAGCGCCAAGAGGAGGAGAAGG | 4 |
| wt-168930-169050 | 5686 | AGAGCAAGGCCTAGGGAAGAGGAGAGGGGGGGTCCTCGAGGGGGCCGTCGCGGGCCCGGTGGGCCCCTCTCAAGGTCGTGTTCCATCCTCAGGGCAGTGTGTCAGGAGCAAGGCAGTTGA | 8 |
| wt-169050-169170 | 5687 | GGAAAGAAGGGGGCAGAGCAGTGTGAGAGGCTTATGTAGGGCGGCTACGTCAGAGTAACGCGTGTTTCTTGGGATGTAGGCCCGGGGGGATTTGCGGGGTCTGCCGGAGGCAGTACGGGT | 4 |
| wt-169170-169290 | 5688 | ACAGATTTCCCGAAAGCGGCGGTGTGTGTGTGCATGTAAGCGTAGAAAGGGGAAGTAGAAAGCGTGTGTTTGTGTTAGAAAAGCGGGTCCCCGGGGGGCAAGCTGTGGGAATGCGGTGGC | 4 |
| wt-169290-169410 | 5689 | CAAGTGCAACAGGAAATGGAAAGGCAGTGCGGCAATCAGAAGGGGGAGTGCGTAGTGTTGTGGGAAGCGGCAGTGTAATCTGCACAAAGAGGCGCGGGGCGCGCAACGTTGGGAGGTCGT | 4 |
| wt-169410-169530 | 5690 | TGGCGGCAGGCGGGAGGCCGTGCTTTAGGGGGGTTCAGGTGAGGCAAGGCTGTGGGGTAACCGTAGGGGAGGCGGGTGAGGCGGCTAAGAGGGCTAAGGGTCGGCGGGTGACGAAGCAGC | 8 |

| | | | | |
|---|---|---|---|---|
| wt-169530-169650 | 5691 | AGACGGCGGATATGGGAATTTCAGAATGAGGTGGCGGATTCAGGCGAAAAGGGTGTGGGCTGTGCGAGTGTCATGAGGCAGGCGCGGAAAGTCGCTGCGGCTTGCTGGGGCATGGGGGGC | | 4 |
| wt-169650-169770 | 5692 | CGCGCATTCCTGGAAAAAGTGGAGGGGGCGTGGCCTTCCCCCGCGGCCCCCCAGCCCCCCCGCACAGAGCGGCGCTACGGCGGGCGGGCGGCGGGGGGTCGGGGTCCGCGGGCTCCGGGG | | 12 |
| wt-169770-169890 | 5693 | GCTGCGGGCGGTGGATGGCGGCGGACGTTCCGGGGATCGGGGGGGTCGGGGGGCGCCGCGCGGGCGCAGCCATGCGTGACCGTGATGAGGGGGCAGGGTCGCAGGGGGTGTGTCTGGTGG | | 12 |
| wt-169890-170010 | 5694 | GGGCGGGAGCGGGGGGCGGCGCGGGAGCCTGCACGCCGTTGGAGGGTAGAATGACAGGGGGCGGGGACAGAGAGGCGGTCGCGCCCCCGGCCGCGCCAGCCAAGCCCCCAAGGGGGGCGG | | 12 |
| wt-170010-170130 | 5695 | GGAGCGGGCAATGGAGCGTGACGAAGGGCCCCAGGGCTGACCCCGGCAAACGTGACCCGGGGCTCCGGGGTGACCCAGCCAAGCGTGACCAAGGGGCCCGTGGGTGACACAGGCAACCCT | | 8 |
| wt-170130-170250 | 5696 | GACAAAGGCCCCCCAGGAAAGACCCCCGGGGGGCATCGGGGGGTGGGGCATGGGGGGCCGCGCATTCCTGGAAAAAGTGGAGGGGGCGTGGCCTTCCCCCGCGGCCCCCCAGCCCCCCCG | | 12 |
| wt-170250-170370 | 5697 | CACAGAGCGGCGCTACGGCGGGCGGGCGGCGGGGGGTCGGGGTCCGCGGGCTCCGGGGGCTGCGGGCGGTGGATGGCGGCGGACGTTCCGGGGATCGGGGGGGTCGGGGGGCGCCGCGCG | | 12 |
| wt-170370-170490 | 5698 | GGCGCAGCCATGCGTGACCGTGATGAGGGGGCAGGGTCGCAGGGGGTGTGTCTGGTGGGGGCGGGAGCGGGGGGCGGCGCGGGAGCCTGCACGCCGTTGGAGGGTAGAATGACAGGGGGC | | 8 |
| wt-170490-170610 | 5699 | GGGGACAGAGAGGCGGTCGCGCCCCCGGCCGCGCCAGCCAAGCCCCCAAGGGGGGCGGGGAGCGGGCAATGGAGCGTGACGAAGGGCCCCAGGGCTGACCCCGGCAAACGTGACCCGGGG | | 12 |
| wt-170610-170730 | 5700 | CTCCGGGGTGACCCAGCCAAGCGTGACCAAGGGGCCCGTGGGTGACACAGGCAACCCTGACAAAGGCCCCCCAGGAAAGACCCCCGTGGGGCATGGGGGGCCGCGCATTCCTGGAAAAAG | | 8 |
| wt-170730-170850 | 5701 | TGGAGGGGGCGTGGCCTTCCCCCGCGGCCCCCCAGCCCCCCCGCACAGAGCGGCGCTACGGCGGGCGGGCGGCGGGGGGTCGGGGTCCGCGGGCTCCGGGGGCTGCGGGCGGTGGATGGC | | 12 |
| wt-170850-170970 | 5702 | GGCGGACGTTCCGGGGATCGGGGGGGTCGGGGGGCGCCGCGCGGGCGCAGCCATGCGTGACCGTGATGAGGGGGCAGGGTCGCAGGGGGTGTGTCTGGTGGGGGCGGGAGCGGGGGGCGG | | 12 |
| wt-170970-171090 | 5703 | CGCGGGAGCCTGCACGCCGTTGGAGGGTAGAATGACAGGGGGCGGGGACAGAGAGGCGGTCGCGCCCCCGGCCGCGCCAGCCAAGCCCCCAAGGGGGGCGGGGAGCGGGCAATGGAGCGT | | 12 |
| wt-171090-171210 | 5704 | GACGAAGGGCCCCAGGGCTGACCCCGGCAAACGTGACCCGGGGCTCCGGGGTGACCCAGCCAAGCGTGACCAAGGGGCCCGTGGGTGACACAGGCAACCCTGACAAAGGCCCCCCAGGAA | | 8 |
| wt-171210-171330 | 5705 | AGACCCCCGGGGGGCATCGGGGGGTGGGGCATGGGGGGCCGCGCATTCCTGGAAAAAGTGGAGGGGGCGTGGCCTTCCCCCGCGGCCCCCCAGCCCCCCCGCACAGAGCGGCGCTACGGC | | 12 |
| wt-171330-171450 | 5706 | GGGCGGGCGGCGGGGGGTCGGGGTCCGCGGGCTCCGGGGGCTGCGGGCGGTGGATGGCGGCGGACGTTCCGGGGATCGGGGGGGTCGGGGGGCGCCGCGCGGGCGCAGCCATGCGTGACC | | 12 |
| wt-171450-171570 | 5707 | GTGATGAGGGGGCAGGGTCGCAGGGGGTGTGTCTGGTGGGGGCGGGAGCGGGGGGCGGCGCGGGAGCCTGCACGCCGTTGGAGGGTAGAATGACAGGGGGCGGGGACAGAGAGGCGGTCG | | 8 |
| wt-171570-171690 | 5708 | CGCCCCCGGCCGCGCCAGCCAAGCCCCCAAGGGGGGCGGGGAGCGGGCAATGGAGCGTGACGAAGGGCCCCAGGGCTGACCCCGGCAAACGTGACCCGGGGCTCCGGGGTGACCCAGCCA | | 12 |
| wt-171690-171810 | 5709 | AGCGTGACCAAGGGGCCCGTGGGTGACACAGGCAACCCTGACAAAGGCCCCCCAGGAAAGACCCCCGGGGGGCATCGGGGGGGGTGTTGGCGGGGGCATGGGGGGGTCGGATTTCGCCCT | | 8 |
| wt-171703-171823 | 5710 | GGCCCGTGGGTGACACAGGCAACCCTGACAAAGGCCCCCCAGGAAAGACCCCCGGGGGGCATCGGGGGGGGTGTTGGCGGGGGCATGGGGGGGTCGGATTTCGCCCTTATTGCCCTGTTT | | 8 |
| AG876-0-120 | 172765 | AGAATTTGTCTTGCTCTATTCACCGTTACTTTTCTTCTTGCCCGTTCTCTTTCTTAGTATGAATCCAGTATGCCTGCCTGTAATTGTTGCGCCCTACCTGTTTTGGCTGGCGGCTATTGC | | 4 |
| AG876-5760-5880 | 345530 | CCCTCCTCAGAAAACTCTGCATGGAGAAGCTGGACGTGAACCTTCCCCCCCCCCCCGACCTGTGTGCTGTATTTACAAACACTACAATAAACCCAATGTGCAAATGTGGTTTGTATGGCT | | 4 |
| AG876-5790-5910 | 518295 | TGGACGTGAACCTTCCCCCCCCCCCCGACCTGTGTGCTGTATTTACAAACACTACAATAAACCCAATGTGCAAATGTGGTTTGTATGGCTACTTTGTGTTCCTAAAAAATGCAACAATAG | | 4 |
| AG876-5820-5940 | 691060 | TGTGTGCTGTATTTACAAACACTACAATAAACCCAATGTGCAAATGTGGTTTGTATGGCTACTTTGTGTTCCTAAAAAATGCAACAATAGAAGTGGAAACCCTCAGTCACGGGACATTAA | | 4 |
| AG876-7500-7620 | 863825 | CCCGGATATAGATTAGGATAGCATATACTACCCAAATAGAGATTAGGATAGCATATGCTACCCAAATAGAGATTAGGATAGCATATGCTACCCGGATATAGATTAGGATAGCATATACTA | | 4 |
| AG876-7530-7650 | 1036590 | CCCAAATAGAGATTAGGATAGCATATGCTACCCAAATAGAGATTAGGATAGCATATGCTACCCGGATATAGATTAGGATAGCATATACTACCCAAATAGAGATTAGGATAGCATATGCTA | | 4 |
| AG876-7560-7680 | 1209355 | CCCAAATAGAGATTAGGATAGCATATGCTACCCGGATATAGATTAGGATAGCATATACTACCCAAATAGAGATTAGGATAGCATATGCTACCCAAATAGAGATTAGGATAGCATATGCTA | | 4 |
| AG876-7590-7710 | 1382120 | CCCGGATATAGATTAGGATAGCATATACTACCCAAATAGAGATTAGGATAGCATATGCTACCCAAATAGAGATTAGGATAGCATATGCTACCCGGATATAGATTAGGATAGCATATGCTA | | 4 |
| AG876-7620-7740 | 1554885 | CCCAAATAGAGATTAGGATAGCATATGCTACCCAAATAGAGATTAGGATAGCATATGCTACCCGGATATAGATTAGGATAGCATATGCTACCCAAATAGAGATTGGGATAGCATATGCTA | | 4 |
| AG876-7650-7770 | 1727650 | CCCAAATAGAGATTAGGATAGCATATGCTACCCGGATATAGATTAGGATAGCATATGCTACCCAAATAGAGATTGGGATAGCATATGCTACCCGGATATAGATTAGGATAGCATATGCTA | | 4 |
| AG876-7680-7800 | 1900415 | CCCGGATATAGATTAGGATAGCATATGCTACCCAAATAGAGATTGGGATAGCATATGCTACCCGGATATAGATTAGGATAGCATATGCTACCCAAATAGAGATTAGGATAGCATATGCTA | | 4 |
| AG876-7710-7830 | 2073180 | CCCAAATAGAGATTGGGATAGCATATGCTACCCGGATATAGATTAGGATAGCATATGCTACCCAAATAGAGATTAGGATAGCATATGCTACCCGGATATAGATTAGGATAGCATATGCTA | | 4 |
| AG876-7740-7860 | 2245945 | CCCGGATATAGATTAGGATAGCATATGCTACCCAAATAGAGATTAGGATAGCATATGCTACCCGGATATAGATTAGGATAGCATATGCTACCCAAATAGAGATTGGGATAGCATATGCTA | | 4 |
| AG876-7770-7890 | 2418710 | CCCAAATAGAGATTAGGATAGCATATGCTACCCGGATATAGATTAGGATAGCATATGCTACCCAAATAGAGATTGGGATAGCATATGCTATCCTAATCTCTATCTGGGTAGCATATACTA | | 4 |
| AG876-7800-7920 | 2591475 | CCCGGATATAGATTAGGATAGCATATGCTACCCAAATAGAGATTGGGATAGCATATGCTATCCTAATCTCTATCTGGGTAGCATATACTATCCTAATCTCTATATGGGTAGCATATGCTA | | 4 |
| AG876-7830-7950 | 2764240 | CCCAAATAGAGATTGGGATAGCATATGCTATCCTAATCTCTATCTGGGTAGCATATACTATCCTAATCTCTATATGGGTAGCATATGCTATCCTAATCTATATCTGGGTAGCATATACTA | | 4 |
| AG876-7860-7980 | 2937005 | TCCTAATCTCTATCTGGGTAGCATATACTATCCTAATCTCTATATGGGTAGCATATGCTATCCTAATCTATATCTGGGTAGCATATACTATCCTAATCTCTATATGGGTAGCATATGCTA | | 4 |
| AG876-7890-8010 | 3109770 | TCCTAATCTCTATATGGGTAGCATATGCTATCCTAATCTATATCTGGGTAGCATATACTATCCTAATCTCTATATGGGTAGCATATGCTATCCTAATCTATATCTGGGTAGCATATGCTA | | 4 |
| AG876-7920-8040 | 3282535 | TCCTAATCTATATCTGGGTAGCATATACTATCCTAATCTCTATATGGGTAGCATATGCTATCCTAATCTATATCTGGGTAGCATATGCTATCCATCGCAACATTAGCCCACCGTGCTCTC | | 4 |
| AG876-7950-8070 | 3455300 | TCCTAATCTCTATATGGGTAGCATATGCTATCCTAATCTATATCTGGGTAGCATATGCTATCCATCGCAACATTAGCCCACCGTGCTCTCAACGACCTCGTGAATATGAGGACCAACAAC | | 4 |
| AG876-7980-8100 | 3628065 | TCCTAATCTATATCTGGGTAGCATATGCTATCCATCGCAACATTAGCCCACCGTGCTCTCAACGACCTCGTGAATATGAGGACCAACAACCCTGTGCTTGGCGCTCAGGCGCAAGTGTGT | | 4 |
| AG876-8010-8130 | 3800830 | TCCATCGCAACATTAGCCCACCGTGCTCTCAACGACCTCGTGAATATGAGGACCAACAACCCTGTGCTTGGCGCTCAGGCGCAAGTGTGTGTAACTTGTCTTCCAGATCGCAGCAATCGC | | 4 |
| AG876-10140-10260 | 3973595 | AGAAGCATACATGACAATTAAAGCCAGGTGATAATTCCATACCCTGGAAGCAGGAGATGGGTGCATTTCACCCCAACCCCCCCTTTCGACTGTCATTTACAATAAAATGAAACCTTTTAT | | 4 |
| AG876-10170-10290 | 4146360 | ATAATTCCATACCCTGGAAGCAGGAGATGGGTGCATTTCACCCCAACCCCCCCTTTCGACTGTCATTTACAATAAAATGAAACCTTTTATTCTTGATTGCCTCTTGTGTTCTTGCCGCCC | | 4 |
| AG876-14040-14160 | 4319125 | AAATCAAAACTTTAGAGGCGAATGGGCGCCATTTTGTCCCCACGCGCGCATAATGGCGGACCTAGGCCTAAAACCCCCAGGAAGCGGGTCTATGGTTGGCTGCGCTGCTGCTATCTTTAG | | 4 |
| AG876-14460-14580 | 4491890 | TTTTTCCCCTTCCCTCCCTCGTCTTGCCCTGCGCCTGGGGCCACCTTCATCACCGTCGCTGACTCCGCCATCCAAGCCTAGGGGAGACCGAAGTGAAGTCCCTGGACCAGCCCGGCCCGG | | 8 |
| AG876-15060-15180 | 4664655 | CACATGTGCCAGACGCCTGGGCCTCTAAGGCCCTCGGGTCCCCCTGGACCCCGGCCTCAGCAACCCTGCTGCTCCCCTCCTGCCACCCCAGCCTCCCCCCCTCCCCGTCCCCCTTCGCTC | | 8 |
| AG876-15870-15990 | 4837420 | CCCCTCTCTCTGTCCTTCAGAGGAACCAGGGACCTCGGGCACCCCAGAGCCCCTCGGGCCCGCCTCCAGGCGCCCTCCTGGTCTCCGCTCCCCTCTGAGCCCCGTTAAACCCAAAGAATG | | 8 |
| AG876-15900-16020 | 5010185 | GACCTCGGGCACCCCAGAGCCCCTCGGGCCCGCCTCCAGGCGCCCTCCTGGTCTCCGCTCCCCTCTGAGCCCCGTTAAACCCAAAGAATGTCTGAGGGGAGCCACCCTCGGGGCCCAGGC | | 8 |
| AG876-16200-16320 | 5182950 | AGCCTCTCCCTCGCGGAGAGGGGCCTCTTCGAGCCCTCAAGTCCAGTCCCACCGAGACCCGAGTGGCCCGGATCCCCCACCGGCCCTTCTCTTTCCCCCTACTCCTCTCCAACCTTCGCT | | 8 |
| AG876-16260-16380 | 5355715 | GAGTGGCCCGGATCCCCCACCGGCCCTTCTCTTTCCCCCTACTCCTCTCCAACCTTCGCTCCACCCTAGACCCCAGCTTCTGGCCTCCCCGGGTCCACCAGGCCAGCCGGAGGGACCCCG | | 8 |

| | | | |
|---|---|---|---|
| AG876-16410-16530 | 5528480 | CTGGCCTCTCCTTCCCGCCTCCCACCCGAGCCCCCTCAGCTTGCCTCCCCACCGGGTCCATCAGGCCGGCCGGAGGGACCCCGGCGGCCCGGTGTCAGTTCCCCCTGCAGCCGCCCAGTC | 12 |
| AG876-16560-16680 | 5701245 | TCCCCCCAGCCTGAGGCCCAGGCTCCTGTGCACTGTCTGTAAAGTCCAGCCTCCCACGCCCGTCCACGGCTCCCAGGCCCAGCCCCGTCCACCCCTCCCCACGTTGGACAGGCCCTCTGT | 8 |
| AG876-16590-16710 | 5874010 | CACTGTCTGTAAAGTCCAGCCTCCCACGCCCGTCCACGGCTCCCAGGCCCAGCCCCGTCCACCCCTCCCCACGTTGGACAGGCCCTCTGTCCACCCGGGCCATCCCCGCCCCCCTGTGTC | 8 |
| AG876-16770-16890 | 6046775 | CCCATGTAAGCCTGCCTCGAGTAGGTGCCTCCAGAGCCCCTTTTGCCCCCCTGGCGGCCCAGCCCGACCCCCGGGCGCCCCCAAACTTTGTCCAGATGTCCAGGGGTCCCCGAGGGCGAG | 8 |
| AG876-16800-16920 | 6219540 | CCAGAGCCCCTTTTGCCCCCCTGGCGGCCCAGCCCGACCCCCGGGCGCCCCCAAACTTTGTCCAGATGTCCAGGGGTCCCCGAGGGCGAGGCCCAGCCCCCTCCCGCCCCTGTCCACTGC | 8 |
| AG876-16830-16950 | 6392305 | AGCCCGACCCCCGGGCGCCCCCAAACTTTGTCCAGATGTCCAGGGGTCCCCGAGGGCGAGGCCCAGCCCCCTCCCGCCCCTGTCCACTGCCCCGGTCCCCCCAGAAGCCCCCAAAAGTAG | 8 |
| AG876-16860-16980 | 6565070 | TCCAGATGTCCAGGGGTCCCCGAGGGCGAGGCCCAGCCCCCTCCCGCCCCTGTCCACTGCCCCGGTCCCCCCAGAAGCCCCCAAAAGTAGAGGCTCAGGCCATGCGCGCCCTGTCACCAG | 8 |
| AG876-16890-17010 | 6737835 | GCCCAGCCCCCTCCCGCCCCTGTCCACTGCCCCGGTCCCCCCAGAAGCCCCCAAAAGTAGAGGCTCAGGCCATGCGCGCCCTGTCACCAGGCCTGCCAAAGAGCCAGATCTAAGGCCGGG | 8 |
| AG876-17790-17910 | 7083365 | CCAGTCCTACCAGAGGGGGCCAAGAACCCAGACGAGTCCGTAGAAGGGTCCTCGTCCAGCAAGAAGAGGAGGTGGTAAGCGGTTCACCTTCAGGGGTAAGTAACCTGACCTCTCCAGGGC | 4 |
| AG876-17820-17940 | 7256130 | GACGAGTCCGTAGAAGGGTCCTCGTCCAGCAAGAAGAGGAGGTGGTAAGCGGTTCACCTTCAGGGGTAAGTAACCTGACCTCTCCAGGGCTCACATAAAGGGAGGCTTAGTATACATGCT | 4 |
| AG876-19020-19140 | 7428895 | GTCTCCGCTCCCCTCTGAGCCCCGTTAAACCCAAAGAATGTCTGAGGGGAGCCACCCTCGGGGCCCAGGCCCCAGAGTCCAGAGGTCAGGGGCACCTCAGGGTGCCTCCCCGGGTCCCAG | 8 |
| AG876-19290-19410 | 7601660 | GGGCCTCTTCGAGCCCTCAAGTCCAGTCCCACCGAGACCCGAGTGGCCCGGATCCCCCACCGGCCCTTCTCTTTCCCCCTACTCCTCTCCAACCTTCGCTCCACCCTAGACCCCAGCTTC | 8 |
| AG876-19320-19440 | 7774425 | ACCGAGACCCGAGTGGCCCGGATCCCCCACCGGCCCTTCTCTTTCCCCCTACTCCTCTCCAACCTTCGCTCCACCCTAGACCCCAGCTTCTGGCCTCCCCGGGTCCACCAGGCCAGCCGG | 8 |
| AG876-19800-19920 | 7947190 | GGGACTTTATGTGACCCTTGGGCCTGGCTCCCCATAGACTCCCATGTAAGCCTGCCTCGAGTAGGTGCCTCCAGAGCCCCTTTTGCCCCCCTGGCGGCCCAGCCCGACCCCCGGGCGCCC | 8 |
| AG876-19830-19950 | 8119955 | CCCATAGACTCCCATGTAAGCCTGCCTCGAGTAGGTGCCTCCAGAGCCCCTTTTGCCCCCCTGGCGGCCCAGCCCGACCCCCGGGCGCCCCCAAACTTTGTCCAGATGTCCAGGGGTCCC | 8 |
| AG876-20700-20820 | 8292720 | CCTGGACCAGCCCGGCCCGGGCCCCCCGGTATCGGGCCAGAGGTAAGTGGACTTTAATTTTTTCTGCTAAGCCCAACACTCCACCACACCCAGGCACACACTACACACACCCACCCGTCT | 4 |
| AG876-21480-21600 | 8638250 | GAGGCCCGCGGCCATTGGGCCCAGATTGAGAGACCAGTCCAGGGGCCCGAGGTTGGAGCCAGCGGGCACCCGAGGTCCCAGCACCCGGTCCCTCCGGGGGGCAGAGACAGGCAGGGCCCC | 8 |
| AG876-21510-21630 | 8811015 | AGACCAGTCCAGGGGCCCGAGGTTGGAGCCAGCGGGCACCCGAGGTCCCAGCACCCGGTCCCTCCGGGGGGCAGAGACAGGCAGGGCCCCCCGGCAGCTGGCCCCGAGGAGGCGCCCGGA | 12 |
| AG876-22110-22230 | 8983780 | CCCGTTAAACCCAAAGAATGTCTGAGGGGAGCCACCCTCGGGGCCCAGGCCCCAGAGTCCAGAGGTCAGGGGCACCTCAGGGTGCCTCCCCGGGTCCCAGGCCAGCCGGAGGGACCCCGG | 8 |
| AG876-22140-22260 | 9156545 | GCCACCCTCGGGGCCCAGGCCCCAGAGTCCAGAGGTCAGGGGCACCTCAGGGTGCCTCCCCGGGTCCCAGGCCAGCCGGAGGGACCCCGGCAGCCCGGGCGGCCCCAGAGGCCGGTTCCT | 12 |
| AG876-22170-22290 | 9329310 | AGAGGTCAGGGGCACCTCAGGGTGCCTCCCCGGGTCCCAGGCCAGCCGGAGGGACCCCGGCAGCCCGGGCGGCCCCAGAGGCCGGTTCCTCGCCCCTTCCCCGGGCTTCAGAGCCCAGGG | 12 |
| AG876-22200-22320 | 9502075 | CGGGTCCCAGGCCAGCCGGAGGGACCCCGGCAGCCCGGGCGGCCCCAGAGGCCGGTTCCTCGCCCCTTCCCCGGGCTTCAGAGCCCAGGGTGTCCCCCAGAAGGGACCCTAGGCGTCCCC | 12 |
| AG876-22230-22350 | 9674840 | CAGCCCGGGCGGCCCCAGAGGCCGGTTCCTCGCCCCTTCCCCGGGCTTCAGAGCCCAGGGTGTCCCCCAGAAGGGACCCTAGGCGTCCCCTCTCCTCCCCTCCAGGCCCGAGCCTCTCCC | 8 |
| AG876-22380-22500 | 9847605 | GTCCAGTCCCACCGAGACCCGAGTGGCCCGGATCCCCCACCGGCCCTTCTCTTTCCCCCTACTCCTCTCCAACCTTCGCTCCACCCTAGACCCCAGCTTCTGGCCTCCCCGGGTCCACCA | 8 |
| AG876-22410-22530 | 10020370 | GATCCCCCACCGGCCCTTCTCTTTCCCCCTACTCCTCTCCAACCTTCGCTCCACCCTAGACCCCAGCTTCTGGCCTCCCCGGGTCCACCAGGCCAGCCGGAGGGACCCCGGCAGCCCGGG | 8 |
| AG876-22440-22560 | 10193135 | ACTCCTCTCCAACCTTCGCTCCACCCTAGACCCCAGCTTCTGGCCTCCCCGGGTCCACCAGGCCAGCCGGAGGGACCCCGGCAGCCCGGGCGAGTCGCCTTCCCTCTCCCCTGGCCTCTC | 8 |
| AG876-22890-23010 | 10365900 | GGCCTGGCTCCCCATAGACTCCCATGTAAGCCTGCCTCGAGTAGGTGCCTCCAGAGCCCCTTTTGCCCCCCTGGCGGCCCAGCCCGACCCCCGGGCGCCCCCAAACTTTGTCCAGATGTC | 8 |
| AG876-23310-23430 | 10538665 | CCTAGGCCTAAAACCCCCAGGAAGCGGGTCTATGGTTGGCTGCGCTGCTGCTATCTTTAGAGGGGAAAAGAGGAATAAGCCCCCAGACAGGGGAGTGGGCTTGTTTGTGACTTCACCAAA | 4 |
| AG876-23670-23790 | 10711430 | TTTTTCCCCTTCCCTCCCTCGTCTTGCCCTGCGCCTGGGGCCACCTTCATCACCGTCGCTGACTCCGCCATCCAAGCCTAGGGGAGACCGAAGTGAAGTCCCTGGACCAGCCCGGCCCGG | 8 |
| AG876-24330-24450 | 10884195 | CAACCCTGCTGCTCCCCTCCTGCCACCCCAGCCTCCCCCCCTCCCCGTCCCCCTTCGCTCCTTATCCTCCCCCGGTCCCCAGTAGGGCCGCCTGCCCCCCTGCACCCAGTACCTGCCCCT | 8 |
| AG876-24870-24990 | 11056960 | GCAGAGTCCGGGCGATCCTCTGAGACCCTCCGGGCCCGGACGGCCGCCCTCGGCCCCCCAGACAGACCCCAGGGTCTCCAGGCAGGGTCCGGCATCTCCAGGGGCAGCAGGCTCACCACC | 8 |
| AG876-25830-25950 | 11229725 | CGTCCACGGCTCCCAGGCCCAGCCCCGTCCACCCCTCCCCACGTTGGACAGGCCCTCTGTCCACCCGGGCCATCCCCGCCCCCCTGTGTCCACCCCAGTCCCGTCCAGGGGGGACTTTAT | 8 |
| AG876-25860-25980 | 11402490 | ACCCCTCCCCACGTTGGACAGGCCCTCTGTCCACCCGGGCCATCCCCGCCCCCCTGTGTCCACCCCAGTCCCGTCCAGGGGGGACTTTATGTGACCCTTGGGCCTGGCTCCCCATAGACT | 8 |
| AG876-26070-26190 | 11575255 | TCCAGATGTCCAGGGGTCCCCGAGGGCGAGGCCCAGCCCCCTCCCGCCCCTGTCCACTGCCCCGGTCCCCCCAGAAGCCCCCAAAAGTAGAGGCTCAGGCCATGCGCGCCCTGTCACCAG | 8 |
| AG876-27000-27120 | 11748020 | CCAGTCCTACCAGAGGGGGCCAAGAACCCAGACGAGTCCGTAGAAGGGTCCTCGTCCAGCAAGAAGAGGAGGTGGTAAGCGGTTCACCTTCAGGGGTAAGTAACCTGACCTCTCCAGGGC | 4 |
| AG876-28110-28230 | 12093550 | CCGGGCGCCTCCTCGGAGCCAGCCCCCGGGGTTGGTTCTGCCCCTCTCTCTGTCCTTCAGAGGAACCAGGGACCTCGGGCACCCCAGAGCCCCTCGGGCCCGCCTCCAGGCGCCCTCCTG | 8 |
| AG876-29160-29280 | 12266315 | CGAGGGCGAGGCCCAGCCCCCTCCCGCCCCTGTCCACTGCCCCGGTCCCCCCAGAAGCCCCCAAAAGTAGAGGCTCAGGCCATGCGCGCCCTGTCACCAGGCCTGCCAAAGAGCCAGATC | 8 |
| AG876-29490-29610 | 12439080 | TGCGCTGCTGCTATCTTTAGAGGGGAAAAGAGGAATAAGCCCCCAGACAGGGGAGTGGGCTTGTTTGTGACTTCACCAAAGGTCAGGGCCCAAGGGGGGTTCGCGTTGCTAGGCCACCTTC | 4 |
| AG876-30390-30510 | 12611845 | TTTTAGAACCACAGCCTGGACACATGTGCCAGACGCCTGGGCCTCTAAGGCCCTCGGGTCCCCCTGGACCCCGGCCTCAGCAACCCTGCTGCTCCCCTCCTGCCACCCCAGCCTCCCCCC | 8 |
| AG876-30420-30540 | 12784610 | AGACGCCTGGGCCTCTAAGGCCCTCGGGTCCCCCTGGACCCCGGCCTCAGCAACCCTGCTGCTCCCCTCCTGCCACCCCAGCCTCCCCCCCTCCCCGTCCCCCTTCGCTCCTTATCCTCC | 8 |
| AG876-30450-30570 | 12957375 | CCCCTGGACCCCGGCCTCAGCAACCCTGCTGCTCCCCTCCTGCCACCCCAGCCTCCCCCCCTCCCCGTCCCCCTTCGCTCCTTATCCTCCCCCGGTCCCCAGTAGGGCCGCCTGCCCCCC | 12 |
| AG876-30480-30600 | 13130140 | GCTCCCCTCCTGCCACCCCAGCCTCCCCCCCTCCCCGTCCCCCTTCGCTCCTTATCCTCCCCCGGTCCCCAGTAGGGCCGCCTGCCCCCCTGCACCCAGTACCTGCCCCTCTTGGCCACG | 8 |
| AG876-30900-31020 | 13302905 | GCTGTCTCAGGAGGGGCCTGGCTCCGCCGGGTGGCCCTGGGGTAAGTCTGGGAGGCAGAGGGACGGCCTAGGCCCGGGGAAGTGGAGGGGGATCGCCCGGGTCTCTGTTGGCAGAGTCCG | 8 |
| AG876-32610-32730 | 13475670 | GGGAGTGGGCTTGTTTGTGACTTCACCAAAGGTCAGGGCCCAAGGGGGGTTCGCGTTGCTAGGCCACCTTCTCAGTCCAGCGCGTTTACGTAAGCCAGACAGCAGCCAATTGTCAGTTCTA | 4 |
| AG876-32640-32760 | 13648435 | GGTCAGGGCCCAAGGGGGGTTCGCGTTGCTAGGCCACCTTCTCAGTCCAGCGCGTTTACGTAAGCCAGACAGCAGCCAATTGTCAGTTCTAGGGAGGGGGACCACTGCCCCTGGTATAAAG | 4 |
| AG876-32850-32970 | 13821200 | CCCTCCAAGGACTCGGGCCCAGTTTCTAACTTTTTCCCCTTCCCTCCCTCGTCTTGCCCTGCGCCTGGGGCCACCTTCATCACCGTCGCTGACTCCGCCATCCAAGCCTAGGGGAGACCG | 4 |
| AG876-32880-33000 | 13993965 | TTTTTCCCCTTCCCTCCCTCGTCTTGCCCTGCGCCTGGGGCCACCTTCATCACCGTCGCTGACTCCGCCATCCAAGCCTAGGGGAGACCGAAGTGAAGTCCCTGGACCAGCCCGGCCCGG | 8 |
| AG876-32940-33060 | 14166730 | GACTCCGCCATCCAAGCCTAGGGGAGACCGAAGTGAAGTCCCTGGACCAGCCCGGCCCGGGCCCCCCGGTATCGGGCCAGAGGTAAGTGGACTTTAATTTTTTCTGCTAAGCCCAACACT | 4 |
| AG876-33690-33810 | 14339495 | TAGACCCGGCCAAGCCCCATCCCTGAAGACCCAGCGGCCATTCTCTCTGGTAACGAGCAGAGAAGAAGTAGAGGCCCGCGGCCATTGGGCCCAGATTGAGAGACCAGTCCAGGGGCCCGA | 4 |
| AG876-34200-34320 | 14685025 | ACAGGCCCCCCAGACCCGGGTCTCGGCCAGCCGAGCCGACCGGCCCCGCGCCGGGCGCCTCCTCGGAGCCAGCCCCCGGGGTTGGTTCTGCCCCTCTCTCTGTCCTTCAGAGGAACCAGG | 8 |
| AG876-34230-34350 | 14857790 | CCGAGCCGACCGGCCCCGCGCCGGGCGCCTCCTCGGAGCCAGCCCCCGGGGTTGGTTCTGCCCCTCTCTCTGTCCTTCAGAGGAACCAGGGACCTCGGGCACCCCAGAGCCCCTCGGGCC | 12 |

| | | | | |
|---|---|---|---|---|
| AG876-34260-34380 | 15030555 | CCTCGGAGCCAGCCCCCGGGGTTGGTTCTGCCCCTCTCTCTGTCCTTCAGAGGAACCAGGGACCTCGGGCACCCCAGAGCCCCTCGGGCCCGCCTCCAGGCGCCCTCCTGGTCTCCGCTC | | 8 |
| AG876-35490-35610 | 15203320 | TTTAGCCCCAGCCCCGCCTCTATAAGTTCTAGGCAAACCTCCAATCACCAGCCACCTTCCAATGTAGTCTCTTAGAGAGTGGCTGCTACGCATTAGCGAGAGATTTGACCCACACCCAGT | | 4 |
| AG876-35520-35640 | 15376085 | AGGCAAACCTCCAATCACCAGCCACCTTCCAATGTAGTCTCTTAGAGAGTGGCTGCTACGCATTAGCGAGAGATTTGACCCACACCCAGTAGCCACCCGACGCCAATCTGTCTACATAGA | | 4 |
| AG876-35550-35670 | 15548850 | AATGTAGTCTCTTAGAGAGTGGCTGCTACGCATTAGCGAGAGATTTGACCCACACCCAGTAGCCACCCGACGCCAATCTGTCTACATAGAAGAAGAAGAGGATGAAGACTAAGTCACAGG | | 4 |
| AG876-35580-35700 | 15721615 | CATTAGCGAGAGATTTGACCCACACCCAGTAGCCACCCGACGCCAATCTGTCTACATAGAAGAAGAAGAGGATGAAGACTAAGTCACAGGCTTAGCCAGGTGATTTGTGAATTTCAGTTT | | 4 |
| AG876-35610-35730 | 15894380 | AGCCACCCGACGCCAATCTGTCTACATAGAAGAAGAAGAGGATGAAGACTAAGTCACAGGCTTAGCCAGGTGATTTGTGAATTTCAGTTTATTTACTTTCTTCCAATCAAGCTTTCCCAG | | 4 |
| AG876-35760-35880 | 16067145 | TTCCATGGGGGACCTAGTATCCGTTCTATTAGATTAACTCGCAAGATACTAAAATTAACCAAGGTCAGCCCAAGTGACGCGTGTTATGCCAGGCTGCCCACCCTGAGGATTTCCCCCAAA | | 4 |
| AG876-35790-35910 | 16239910 | AGATTAACTCGCAAGATACTAAAATTAACCAAGGTCAGCCCAAGTGACGCGTGTTATGCCAGGCTGCCCACCCTGAGGATTTCCCCCAAAAATGGGCTGTTAGTAGGGTCCTCCTACCCC | | 4 |
| AG876-35820-35940 | 16412675 | AAGGTCAGCCCAAGTGACGCGTGTTATGCCAGGCTGCCCACCCTGAGGATTTCCCCCAAAAATGGGCTGTTAGTAGGGTCCTCCTACCCCCTCTTTAGGCCAGGTGTGTTATTTTGTTGT | | 4 |
| AG876-35850-35970 | 16585440 | AGGCTGCCCACCCTGAGGATTTCCCCCAAAAATGGGCTGTTAGTAGGGTCCTCCTACCCCCTCTTTAGGCCAGGTGTGTTATTTTGTTGTGTTAGTGCTATGTAATGTGTTGCCGCCAGG | | 4 |
| AG876-35880-36000 | 16758205 | AATGGGCTGTTAGTAGGGTCCTCCTACCCCCTCTTTAGGCCAGGTGTGTTATTTTGTTGTGTTAGTGCTATGTAATGTGTTGCCGCCAGGTGGCAGCTTGTTTGTAGATGTGCAGCGCCC | | 4 |
| AG876-35910-36030 | 16930970 | CTCTTTAGGCCAGGTGTGTTATTTTGTTGTGTTAGTGCTATGTAATGTGTTGCCGCCAGGTGGCAGCTTGTTTGTAGATGTGCAGCGCCCCTTAATGGTAGGTCTGCTTTAGGGCTGCCA | | 4 |
| AG876-35940-36060 | 17103735 | GTTAGTGCTATGTAATGTGTTGCCGCCAGGTGGCAGCTTGTTTGTAGATGTGCAGCGCCCCTTAATGGTAGGTCTGCTTTAGGGCTGCCAGGTGGCGCAAACTAGGATTAATTCACCTGT | | 4 |
| AG876-35970-36090 | 17276500 | TGGCAGCTTGTTTGTAGATGTGCAGCGCCCCTTAATGGTAGGTCTGCTTTAGGGCTGCCAGGTGGCGCAAACTAGGATTAATTCACCTGTATCCCTTTCCCTCCACCCGCAGTAACTCAG | | 4 |
| AG876-36270-36390 | 17449265 | GAAACCCGTCACTCTCCGTAATTCCCACGAATCCTTACCAGGAACAACTATCAAATAATCCATTAATTCAACTACAAATCGTTGTCGGGGAAAACACTGGGGCACCCGCACCGCCCCAAC | | 4 |
| AG876-36300-36420 | 17622030 | ATCCTTACCAGGAACAACTATCAAATAATCCATTAATTCAACTACAAATCGTTGTCGGGGAAAACACTGGGGCACCCGCACCGCCCCAACCACCCCCCCCACCCCCTCCACCTCCTCCAC | | 4 |
| AG876-36330-36450 | 17794795 | CATTAATTCAACTACAAATCGTTGTCGGGGAAAACACTGGGGCACCCGCACCGCCCCAACCACCCCCCCCACCCCCTCCACCTCCTCCACCTGAGCGCAGGGATGCCTGGACACAAGAGC | | 4 |
| AG876-36390-36510 | 18140325 | CACCCCCCCCACCCCCTCCACCTCCTCCACCTGAGCGCAGGGATGCCTGGACACAAGAGCCCTTACCACTGGATATGAATCCCCTGGGCAGTGACGCCAGTCAAGGCCCTCTGGCATCTT | | 4 |
| AG876-36420-36540 | 18313090 | CTGAGCGCAGGGATGCCTGGACACAAGAGCCCTTACCACTGGATATGAATCCCCTGGGCAGTGACGCCAGTCAAGGCCCTCTGGCATCTTCCATAAGAATGCTTTGCATGGCTCAATACC | | 4 |
| AG876-36450-36570 | 18485855 | CCTTACCACTGGATATGAATCCCCTGGGCAGTGACGCCAGTCAAGGCCCTCTGGCATCTTCCATAAGAATGCTTTGCATGGCTCAATACCTTCTTAGAAACGCACGGGGCCAACAAGGCC | | 4 |
| AG876-36480-36600 | 18658620 | GTGACGCCAGTCAAGGCCCTCTGGCATCTTCCATAAGAATGCTTTGCATGGCTCAATACCTTCTTAGAAACGCACGGGGCCAACAAGGCCTTCTTAGGCCACTAGGACCACAAACACGCT | | 4 |
| AG876-36510-36630 | 18831385 | CCATAAGAATGCTTTGCATGGCTCAATACCTTCTTAGAAACGCACGGGGCCAACAAGGCCTTCTTAGGCCACTAGGACCACAAACACGCTCTCAGGTCACCTTGGAACGTCAACCTGTCC | | 4 |
| AG876-36540-36660 | 19004150 | TTCTTAGAAACGCACGGGGCCAACAAGGCCTTCTTAGGCCACTAGGACCACAAACACGCTCTCAGGTCACCTTGGAACGTCAACCTGTCCACAACCCTCGCCAGGAGGCACCCATCATTT | | 4 |
| AG876-36570-36690 | 19176915 | TTCTTAGGCCACTAGGACCACAAACACGCTCTCAGGTCACCTTGGAACGTCAACCTGTCCACAACCCTCGCCAGGAGGCACCCATCATTTTGTTACAGAGTCCCGCGCCCCCCCGATTCA | | 4 |
| AG876-36600-36720 | 19349680 | CTCAGGTCACCTTGGAACGTCAACCTGTCCACAACCCTCGCCAGGAGGCACCCATCATTTTGTTACAGAGTCCCGCGCCCCCCCGATTCACACCAGTGCCCATGGTAGCCTTAGGACATA | | 4 |
| AG876-36630-36750 | 19522445 | ACAACCCTCGCCAGGAGGCACCCATCATTTTGTTACAGAGTCCCGCGCCCCCCCGATTCACACCAGTGCCCATGGTAGCCTTAGGACATACTCTTCAACCCACACCACCACCAAGGCCTA | | 4 |
| AG876-36660-36780 | 19695210 | TGTTACAGAGTCCCGCGCCCCCCCGATTCACACCAGTGCCCATGGTAGCCTTAGGACATACTCTTCAACCCACACCACCACCAAGGCCTACTCTTCCTCAACCCAGAATACCACTGATAA | | 4 |
| AG876-36690-36810 | 19867975 | CACCAGTGCCCATGGTAGCCTTAGGACATACTCTTCAACCCACACCACCACCAAGGCCTACTCTTCCTCAACCCAGAATACCACTGATAATACCACCAAGGCATACTAATCAACCAGCCA | | 4 |
| AG876-36720-36840 | 20040740 | CTCTTCAACCCACACCACCACCAAGGCCTACTCTTCCTCAACCCAGAATACCACTGATAATACCACCAAGGCATACTAATCAACCAGCCACAACACCACCCACGGCGCCACAAAGGCTCA | | 4 |
| AG876-36750-36870 | 20213505 | CTCTTCCTCAACCCAGAATACCACTGATAATACCACCAAGGCATACTAATCAACCAGCCACAACACCACCCACGGCGCCACAAAGGCTCACACTAGGGCATCAACTAAGTCTACCACCGC | | 4 |
| AG876-36780-36900 | 20386270 | TACCACCAAGGCATACTAATCAACCAGCCACAACACCACCCACGGCGCCACAAAGGCTCACACTAGGGCATCAACTAAGTCTACCACCGCATCCTCCCCCGCATCAGAGCACCCCACATT | | 4 |
| AG876-36810-36930 | 20559035 | CAACACCACCCACGGCGCCACAAAGGCTCACACTAGGGCATCAACTAAGTCTACCACCGCATCCTCCCCCGCATCAGAGCACCCCACATTGTAGTTCTGATAGTACAGGACTCCCTCCAC | | 4 |
| AG876-36840-36960 | 20731800 | CACTAGGGCATCAACTAAGTCTACCACCGCATCCTCCCCCGCATCAGAGCACCCCACATTGTAGTTCTGATAGTACAGGACTCCCTCCACCTCCCACATCTTACAGCATTCCTTCTATGA | | 4 |
| AG876-36870-36990 | 20904565 | ATCCTCCCCCGCATCAGAGCACCCCACATTGTAGTTCTGATAGTACAGGACTCCCTCCACCTCCCACATCTTACAGCATTCCTTCTATGACATTATCCCCCGAACCATTGCCGCCACCAG | | 4 |
| AG876-36900-37020 | 21077330 | GTAGTTCTGATAGTACAGGACTCCCTCCACCTCCCACATCTTACAGCATTCCTTCTATGACATTATCCCCCGAACCATTGCCGCCACCAGCAGCACCAGCACACCCCCTTCCAGGTGTCA | | 4 |
| AG876-36930-37050 | 21250095 | CTCCCACATCTTACAGCATTCCTTCTATGACATTATCCCCCGAACCATTGCCGCCACCAGCAGCACCAGCACACCCCCTTCCAGGTGTCATTTATGACCAACAAGCACTTCCTCCAACTC | | 4 |
| AG876-36960-37080 | 21422860 | CATTATCCCCCGAACCATTGCCGCCACCAGCAGCACCAGCACACCCCCTTCCAGGTGTCATTTATGACCAACAAGCACTTCCTCCAACTCCAGGGCCACCATGGTGGCCACCTGTCCGCG | | 4 |
| AG876-36990-37110 | 21595625 | CAGCACCAGCACACCCCCTTCCAGGTGTCATTTATGACCAACAAGCACTTCCTCCAACTCCAGGGCCACCATGGTGGCCACCTGTCCGCGACCCCACGCCAACCACTCAGACTCCACCAA | | 4 |
| AG876-37020-37140 | 21768390 | TTTATGACCAACAAGCACTTCCTCCAACTCCAGGGCCACCATGGTGGCCACCTGTCCGCGACCCCACGCCAACCACTCAGACTCCACCAACAAATACGAAACAGGGCCCGGACCAGGGCC | | 4 |
| AG876-37050-37170 | 21941155 | CAGGGCCACCATGGTGGCCACCTGTCCGCGACCCCACGCCAACCACTCAGACTCCACCAACAAATACGAAACAGGGCCCGGACCAGGGCCAGGGAAGGGGCAGGTGGCGGGGCAGGGGCA | | 8 |
| AG876-37080-37200 | 22113920 | ACCCCACGCCAACCACTCAGACTCCACCAACAAATACGAAACAGGGCCCGGACCAGGGCCAGGGAAGGGGCAGGTGGCGGGGCAGGGGCAGAAGCAAGGGCAGGGGCAGAATGCACAAAC | | 4 |
| AG876-37110-37230 | 22286685 | CAAATACGAAACAGGGCCCGGACCAGGGCCAGGGAAGGGGCAGGTGGCGGGGCAGGGGCAGAAGCAAGGGCAGGGGCAGAATGCACAAACTTCCTGAACCACGGAGACCAGGGCCAGACA | | 4 |
| AG876-37140-37260 | 22459450 | AGGGAAGGGGCAGGTGGCGGGGCAGGGGCAGAAGCAAGGGCAGGGGCAGAATGCACAAACTTCCTGAACCACGGAGACCAGGGCCAGACACTTCCAGTCCTAGTATGCCTCAATTAAGTC | | 4 |
| AG876-37170-37290 | 22632215 | GAAGCAAGGGCAGGGGCAGAATGCACAAACTTCCTGAACCACGGAGACCAGGGCCAGACACTTCCAGTCCTAGTATGCCTCAATTAAGTCCGGTTGTCAGTCTTCATCAGGGACAGGGGC | | 4 |
| AG876-37200-37320 | 22804980 | TTCCTGAACCACGGAGACCAGGGCCAGACACTTCCAGTCCTAGTATGCCTCAATTAAGTCCGGTTGTCAGTCTTCATCAGGGACAGGGGCCTGAGAACTCACCAACCCCCGGCCCGTCTA | | 4 |
| AG876-37230-37350 | 22977745 | CTTCCAGTCCTAGTATGCCTCAATTAAGTCCGGTTGTCAGTCTTCATCAGGGACAGGGGCCTGAGAACTCACCAACCCCCGGCCCGTCTACGGCCGGCCCCGTCTGTAGAGTGACACCTT | | 4 |
| AG876-37260-37380 | 23150510 | CGGTTGTCAGTCTTCATCAGGGACAGGGGCCTGAGAACTCACCAACCCCCGGCCCGTCTACGGCCGGCCCCGTCTGTAGAGTGACACCTTCAGCAACCCCTGACATTTCACCAATACATG | | 4 |
| AG876-37290-37410 | 23323275 | CTGAGAACTCACCAACCCCCGGCCCGTCTACGGCCGGCCCCGTCTGTAGAGTGACACCTTCAGCAACCCCTGACATTTCACCAATACATGAACCAGAGTCCTCTGATAGTGAAGAACCCC | | 4 |
| AG876-37320-37440 | 23496040 | CGGCCGGCCCCGTCTGTAGAGTGACACCTTCAGCAACCCCTGACATTTCACCAATACATGAACCAGAGTCCTCTGATAGTGAAGAACCCCCCTTTCTCTTCCCCAGTGATTGGTATCCTC | | 4 |
| AG876-37350-37470 | 23668805 | CAGCAACCCCTGACATTTCACCAATACATGAACCAGAGTCCTCTGATAGTGAAGAACCCCCCTTTCTCTTCCCCAGTGATTGGTATCCTCCAACGTTAGAACCTGCAGAATTAGATGAAA | | 4 |
| AG876-37380-37500 | 23841570 | AACCAGAGTCCTCTGATAGTGAAGAACCCCCCTTTCTCTTCCCCAGTGATTGGTATCCTCCAACGTTAGAACCTGCAGAATTAGATGAAAGTTGGGAGGGCATTTTTGAAACAACAGAAT | | 4 |

| | | | | |
|---|---|---|---|---|
| AG876-37410-37530 | 24014335 | CCTTTCTCTTCCCCAGTGATTGGTATCCTCCAACGTTAGAACCTGCAGAATTAGATGAAAGTTGGGAGGGCATTTTTGAAACAACAGAATCTCATAGCTCTGATGAAGAGAATGTAGGGG | | 4 |
| AG876-37440-37560 | 24187100 | CAACGTTAGAACCTGCAGAATTAGATGAAAGTTGGGAGGGCATTTTTGAAACAACAGAATCTCATAGCTCTGATGAAGAGAATGTAGGGGGGCCTAGTAAAAGACCTCGCACCTCCACTC | | 4 |
| AG876-37470-37590 | 24359865 | GTTGGGAGGGCATTTTTGAAACAACAGAATCTCATAGCTCTGATGAAGAGAATGTAGGGGGGCCTAGTAAAAGACCTCGCACCTCCACTCAGTAAAAGACCTTACTCTCTCCAGTAATCA | | 4 |
| AG876-37500-37620 | 24532630 | CTCATAGCTCTGATGAAGAGAATGTAGGGGGGCCTAGTAAAAGACCTCGCACCTCCACTCAGTAAAAGACCTTACTCTCTCCAGTAATCAATGTATCCCAAATAAATGTTAATGATTTTG | | 4 |
| AG876-37530-37650 | 24705395 | GGCCTAGTAAAAGACCTCGCACCTCCACTCAGTAAAAGACCTTACTCTCTCCAGTAATCAATGTATCCCAAATAAATGTTAATGATTTTGTTCTTAACTATTGACCCGCCTGTCATTCTA | | 4 |
| AG876-37560-37680 | 24878160 | AGTAAAAGACCTTACTCTCTCCAGTAATCAATGTATCCCAAATAAATGTTAATGATTTTGTTCTTAACTATTGACCCGCCTGTCATTCTATTAATTAAACAAGGGAAGCTATGTTTAGCT | | 4 |
| AG876-37590-37710 | 25050925 | ATGTATCCCAAATAAATGTTAATGATTTTGTTCTTAACTATTGACCCGCCTGTCATTCTATTAATTAAACAAGGGAAGCTATGTTTAGCTATTCCACCAACACTGCACCAATGGACAGCC | | 4 |
| AG876-37620-37740 | 25223690 | TTCTTAACTATTGACCCGCCTGTCATTCTATTAATTAAACAAGGGAAGCTATGTTTAGCTATTCCACCAACACTGCACCAATGGACAGCCAAAATTGGTGCCTTGCTCACAGCTCTTTTT | | 4 |
| AG876-37650-37770 | 25396455 | TTAATTAAACAAGGGAAGCTATGTTTAGCTATTCCACCAACACTGCACCAATGGACAGCCAAAATTGGTGCCTTGCTCACAGCTCTTTTTGCCAGTGGCTTACAGCCCAGTAGGCAGCTC | | 4 |
| AG876-37680-37800 | 25569220 | ATTCCACCAACACTGCACCAATGGACAGCCAAAATTGGTGCCTTGCTCACAGCTCTTTTTGCCAGTGGCTTACAGCCCAGTAGGCAGCTCAGAAAAGTTTAGCTATTCCACCAACCCGGC | | 4 |
| AG876-37710-37830 | 25741985 | AAAATTGGTGCCTTGCTCACAGCTCTTTTTGCCAGTGGCTTACAGCCCAGTAGGCAGCTCAGAAAAGTTTAGCTATTCCACCAACCCGGCCCCAATGAATGCTTGCCAAAATTGGGGCCT | | 4 |
| AG876-37740-37860 | 25914750 | GCCAGTGGCTTACAGCCCAGTAGGCAGCTCAGAAAAGTTTAGCTATTCCACCAACCCGGCCCCAATGAATGCTTGCCAAAATTGGGGCCTTGCTCTCAGCACTTTGCCAGCGACTTATAG | | 4 |
| AG876-37770-37890 | 26087515 | AGAAAAGTTTAGCTATTCCACCAACCCGGCCCCAATGAATGCTTGCCAAAATTGGGGCCTTGCTCTCAGCACTTTGCCAGCGACTTATAGCATGGTAGGCCGCTCAACTCGGCCTGTCTT | | 4 |
| AG876-37800-37920 | 26260280 | CCCAATGAATGCTTGCCAAAATTGGGGCCTTGCTCTCAGCACTTTGCCAGCGACTTATAGCATGGTAGGCCGCTCAACTCGGCCTGTCTTACTGCCCAGCCTACTCTCCACTTCCAGTCC | | 4 |
| AG876-37830-37950 | 26433045 | TGCTCTCAGCACTTTGCCAGCGACTTATAGCATGGTAGGCCGCTCAACTCGGCCTGTCTTACTGCCCAGCCTACTCTCCACTTCCAGTCCATGTTCGCACTCCTATGCATTTCCTTCCCT | | 4 |
| AG876-38370-38490 | 26605810 | GGGTGGGGGGTGGCCCCGCTGGGCACCGCTGCGCCGCCGCCAGGTCCAGGGGCAGCCGGGGTTCCTGGCGCTCCGGGGGCAGCCGGGCGGCCGCCGGTGGGTCCGCTGGGCCGCTGCCCC | | 12 |
| AG876-38400-38520 | 26778575 | GCGCCGCCGCCAGGTCCAGGGGCAGCCGGGGTTCCTGGCGCTCCGGGGGCAGCCGGGCGGCCGCCGGTGGGTCCGCTGGGCCGCTGCCCCGCTCCGGGTGGGGGGTGGCCCCGCTGGGCA | | 12 |
| AG876-38430-38550 | 26951340 | GTTCCTGGCGCTCCGGGGGCAGCCGGGCGGCCGCCGGTGGGTCCGCTGGGCCGCTGCCCCGCTCCGGGTGGGGGGTGGCCCCGCTGGGCACCGCTGCGCCGCCGCCAGGTCCTGGGGCAG | | 12 |
| AG876-38760-38880 | 27124105 | CCGCTGGGCACCGCTGTGCCGCCGCCAGGTCCTGGGGCAGCCGGGGTTCCTGGCGCTCCGGGGGCAGCCGGGCGGCCGCCGGTGGGTCCGCTGGGCCGCTGCCCCGCTCCGGGTGGGGGG | | 12 |
| AG876-38790-38910 | 27296870 | CCTGGGGCAGCCGGGGTTCCTGGCGCTCCGGGGGCAGCCGGGCGGCCGCCGGTGGGTCCGCTGGGCCGCTGCCCCGCTCCGGGTGGGGGGTGGCCCCGCTGGGCACCGCTGCGCCGCCGC | | 16 |
| AG876-38910-39030 | 27469635 | CAGGTCCAGGGGCAGCCGGGGTTCCTGGCGCTCCGGGGGCAGCCGGGCGGCCGCCGGTGGGTCCGCTGGGCCGCTGCCCCGCTCCGGGTGGGGGGTGGCCCCGCTGGGCACCGCTGCGCC | | 12 |
| AG876-39270-39390 | 27642400 | CCGCTGCGCCGCCGCCAGGTCCAGGGGCAGCCGGGGTTCCTGGCGCTCCGGGGGCAGCCGGGCGGCCGCCGGTGGGTCCGCTGGGCCGCTGCCCCGCTCCGGGTGGGGGGTGGCCCCGCT | | 12 |
| AG876-39300-39420 | 27815165 | CCGGGGTTCCTGGCGCTCCGGGGGCAGCCGGGCGGCCGCCGGTGGGTCCGCTGGGCCGCTGCCCCGCTCCGGGTGGGGGGTGGCCCCGCTGGGCACCGCTGCGCCGCCGCCAGGTCCTGG | | 12 |
| AG876-39510-39630 | 27987930 | CCGCTGGGCACCGCTGTGCCGCCGCCAGGTCCTGGGGCAGCCGGGGTTCCTGGCGCTCCGGGGGCAGCCGGGCGGCCGCCGGTGGGTCCGCTGGGCCGCTGCCCCGCTCCGGGTGGGGGG | | 12 |
| AG876-39540-39660 | 28160695 | CCTGGGGCAGCCGGGGTTCCTGGCGCTCCGGGGGCAGCCGGGCGGCCGCCGGTGGGTCCGCTGGGCCGCTGCCCCGCTCCGGGTGGGGGGTGGCCCCGCTGGGCACCGCTGCGCCGCCGC | | 16 |
| AG876-39570-39690 | 28333460 | GGGGCAGCCGGGCGGCCGCCGGTGGGTCCGCTGGGCCGCTGCCCCGCTCCGGGTGGGGGGTGGCCCCGCTGGGCACCGCTGCGCCGCCGCCAGGTCCAGGGGCAGCCGGGGTTCCTGGCG | | 12 |
| AG876-39600-39720 | 28506225 | CTGGGCCGCTGCCCCGCTCCGGGTGGGGGGTGGCCCCGCTGGGCACCGCTGCGCCGCCGCCAGGTCCAGGGGCAGCCGGGGTTCCTGGCGCTCCGGGGGCAGCCGGGCGGCCGCCGGTGG | | 12 |
| AG876-39630-39750 | 28678990 | TGGCCCCGCTGGGCACCGCTGCGCCGCCGCCAGGTCCAGGGGCAGCCGGGGTTCCTGGCGCTCCGGGGGCAGCCGGGCGGCCGCCGGTGGGTCCGCTGGGCCGCTGCCCCGCTCCGGGTG | | 12 |
| AG876-39660-39780 | 28851755 | CAGGTCCAGGGGCAGCCGGGGTTCCTGGCGCTCCGGGGGCAGCCGGGCGGCCGCCGGTGGGTCCGCTGGGCCGCTGCCCCGCTCCGGGTGGGGGGTGGCCCCGCTGGGCACCGCTGCGCC | | 12 |
| AG876-39690-39810 | 29024520 | CTCCGGGGGCAGCCGGGCGGCCGCCGGTGGGTCCGCTGGGCCGCTGCCCCGCTCCGGGTGGGGGGTGGCCCCGCTGGGCACCGCTGCGCCGCCGCCAGGTCCTGGGGCAGCCGGGGTTCC | | 12 |
| AG876-39810-39930 | 29197285 | TGGCGCTCCGGGGGCAGCCGGGCGGCCGCCGGTGGGTCCGCTGGGCCGCTGCCCCGCTCCGGGTGGGGGGTGGCCCCGCTGGGCACCGCTGCGCCGCCGCCAGGTCCTGGGGCAGCCGGG | | 16 |
| AG876-39840-39960 | 29370050 | GGTGGGTCCGCTGGGCCGCTGCCCCGCTCCGGGTGGGGGGTGGCCCCGCTGGGCACCGCTGCGCCGCCGCCAGGTCCTGGGGCAGCCGGGGTTCCTGGCGCTCCACTGCACCTGGAATGC | | 12 |
| AG876-78240-78360 | 29542815 | TTGGGGTAGTCACTGCTGGGGTGGGGCTGGTGGCATTTGGGGTTGGGGTAGTCACTGCTGAGGTGGGACTTGTTTTTCCCAAGGTGGGGCTGGTGGCATTTGGGGTTGGGGTAGTCACTG | | 4 |
| AG876-78270-78390 | 29715580 | TGGCATTTGGGGTTGGGGTAGTCACTGCTGAGGTGGGACTTGTTTTTCCCAAGGTGGGGCTGGTGGCATTTGGGGTTGGGGTAGTCACTGCTGGGGTGGGGCTGGTGGCATTTGGGGTTG | | 4 |
| AG876-80040-80160 | 29888345 | CCGGGTCTCCCGGCCCCGGATGACAACATAGAAGAAGTACCATCTACCTCGGGTGTTCAGGAACGGGCGTCTGAGGGAGATTGGGAAAATGTCCTCATAGAGATATCAGATAGCAGCTCA | | 4 |
| AG876-80070-80190 | 30061110 | GAAGAAGTACCATCTACCTCGGGTGTTCAGGAACGGGCGTCTGAGGGAGATTGGGAAAATGTCCTCATAGAGATATCAGATAGCAGCTCAGAAGAGGAAGCAGAAGATGCCCACCTGGAG | | 4 |
| AG876-80100-80220 | 30233875 | GAACGGGCGTCTGAGGGAGATTGGGAAAATGTCCTCATAGAGATATCAGATAGCAGCTCAGAAGAGGAAGCAGAAGATGCCCACCTGGAGCCATCCCAGAGGGGTAAAAAGAGAAAACGG | | 4 |
| AG876-80130-80250 | 30406640 | GTCCTCATAGAGATATCAGATAGCAGCTCAGAAGAGGAAGCAGAAGATGCCCACCTGGAGCCATCCCAGAGGGGTAAAAAGAGAAAACGGGTCGATGATGATGCCGGTGGTTCAGCTCCA | | 4 |
| AG876-80160-80280 | 30579405 | GAAGAGGAAGCAGAAGATGCCCACCTGGAGCCATCCCAGAGGGGTAAAAAGAGAAAACGGGTCGATGATGATGCCGGTGGTTCAGCTCCAGCACAACACGTGCCCCCCCCTCAGCTGGAT | | 4 |
| AG876-80190-80310 | 30752170 | CCATCCCAGAGGGGTAAAAAGAGAAAACGGGTCGATGATGATGCCGGTGGTTCAGCTCCAGCACAACACGTGCCCCCCCCTCAGCTGGATCACCCTGGTCGAGAAGCCATTCTCTACAGG | | 4 |
| AG876-80220-80340 | 30924935 | GTCGATGATGATGCCGGTGGTTCAGCTCCAGCACAACACGTGCCCCCCCCTCAGCTGGATCACCCTGGTCGAGAAGCCATTCTCTACAGGTTTCCGCTAGATTTAAGAAGGTTTATTCAA | | 4 |
| AG876-80250-80370 | 31097700 | GCACAACACGTGCCCCCCCCTCAGCTGGATCACCCTGGTCGAGAAGCCATTCTCTACAGGTTTCCGCTAGATTTAAGAAGGTTTATTCAAGCAATTGGAGCCGCAGCTACGGTGAGCTTC | | 4 |
| AG876-80280-80400 | 31270465 | CACCCTGGTCGAGAAGCCATTCTCTACAGGTTTCCGCTAGATTTAAGAAGGTTTATTCAAGCAATTGGAGCCGCAGCTACGGTGAGCTTCCCTATGGCCCAGGTGTGTGATGTGTGTTTT | | 4 |
| AG876-80310-80430 | 31443230 | TTTCCGCTAGATTTAAGAAGGTTTATTCAAGCAATTGGAGCCGCAGCTACGGTGAGCTTCCCTATGGCCCAGGTGTGTGATGTGTGTTTTTGCCCATCGCACAACAAGGTAAGTGATTTG | | 4 |
| AG876-80340-80460 | 31615995 | GCAATTGGAGCCGCAGCTACGGTGAGCTTCCCTATGGCCCAGGTGTGTGATGTGTGTTTTTGCCCATCGCACAACAAGGTAAGTGATTTGTTGCCGTTGGTTTCAGCACCCCGACACGCG | | 4 |
| AG876-80370-80490 | 31788760 | CCTATGGCCCAGGTGTGTGATGTGTGTTTTTGCCCATCGCACAACAAGGTAAGTGATTTGTTGCCGTTGGTTTCAGCACCCCGACACGCGAGCCATAGACCAGTTTTTCGGATCCCAGAT | | 4 |
| AG876-80400-80520 | 31961525 | TGCCCATCGCACAACAAGGTAAGTGATTTGTTGCCGTTGGTTTCAGCACCCCGACACGCGAGCCATAGACCAGTTTTTCGGATCCCAGATTTCAAATACCGACCTGTACGTAATGTATGC | | 4 |
| AG876-80430-80550 | 32134290 | TTGCCGTTGGTTTCAGCACCCCGACACGCGAGCCATAGACCAGTTTTTCGGATCCCAGATTTCAAATACCGACCTGTACGTAATGTATGCCATGGCCATTCGACAGGCCATTAGAGATCG | | 4 |
| AG876-80460-80580 | 32307055 | AGCCATAGACCAGTTTTTCGGATCCCAGATTTCAAATACCGACCTGTACGTAATGTATGCCATGGCCATTCGACAGGCCATTAGAGATCGTCGGAGGAATCCAGCTTCTCGTAGAAGTCA | | 4 |
| AG876-80580-80700 | 32479820 | GGTCAAATGGAGAATGACCACCCTGGCCGCTGGTTGGCCTATGGGTTACCAGGCATACAGTAGCTGGATGTACAGCTACACCGATCCCCAGGTGACTGCCACGATCATACATCTGCAGGC | | 4 |
| AG876-80610-80730 | 32652585 | TGGTTGGCCTATGGGTTACCAGGCATACAGTAGCTGGATGTACAGCTACACCGATCCCCAGGTGACTGCCACGATCATACATCTGCAGGCGACTCTTGGGTGCGCAAGTGGCCGTAGGTG | | 4 |

| | | | |
|---|---|---|---|
| AG876-80640-80760 | 32825350 | TAGCTGGATGTACAGCTACACCGATCCCCAGGTGACTGCCACGATCATACATCTGCAGGCGACTCTTGGGTGCGCAAGTGGCCGTAGGTGTCATGTGACCTTTTCTGCCGGCACCTTTAG | 4 |
| AG876-80670-80790 | 32998115 | GGTGACTGCCACGATCATACATCTGCAGGCGACTCTTGGGTGCGCAAGTGGCCGTAGGTGTCATGTGACCTTTTCTGCCGGCACCTTTAGGCCGCCGCGATGTAGTCCCGGGGATCGCCA | 4 |
| AG876-80700-80820 | 33170880 | GACTCTTGGGTGCGCAAGTGGCCGTAGGTGTCATGTGACCTTTTCTGCCGGCACCTTTAGGCCGCCGCGATGTAGTCCCGGGGATCGCCAGTGGTTGTATGTTCAGAGCCGCGTGGGTGA | 4 |
| AG876-80730-80850 | 33343645 | TCATGTGACCTTTTCTGCCGGCACCTTTAGGCCGCCGCGATGTAGTCCCGGGGATCGCCAGTGGTTGTATGTTCAGAGCCGCGTGGGTGACCTTGTGCAGAGTTCTAATCCATGCTACAG | 4 |
| AG876-80760-80880 | 33516410 | GCCGCCGCGATGTAGTCCCGGGGATCGCCAGTGGTTGTATGTTCAGAGCCGCGTGGGTGACCTTGTGCAGAGTTCTAATCCATGCTACAGTATTTTCTTTGACTACATGGCCATACATAG | 4 |
| AG876-80790-80910 | 33689175 | GTGGTTGTATGTTCAGAGCCGCGTGGGTGACCTTGTGCAGAGTTCTAATCCATGCTACAGTATTTTCTTTGACTACATGGCCATACATAGGAGCCTCACGAAAATCTGGGATGAAGTGGT | 4 |
| AG876-80820-80940 | 33861940 | CCTTGTGCAGAGTTCTAATCCATGCTACAGTATTTTCTTTGACTACATGGCCATACATAGGAGCCTCACGAAAATCTGGGATGAAGTGGTAACACCTGATCAGCGTGTTACATTTATGGA | 4 |
| AG876-80850-80970 | 34034705 | TATTTTCTTTGACTACATGGCCATACATAGGAGCCTCACGAAAATCTGGGATGAAGTGGTAACACCTGATCAGCGTGTTACATTTATGGAATTCCTGGGCTTTTTGCAACGCACGGAGTT | 4 |
| AG876-80880-81000 | 34207470 | GAGCCTCACGAAAATCTGGGATGAAGTGGTAACACCTGATCAGCGTGTTACATTTATGGAATTCCTGGGCTTTTTGCAACGCACGGAGTTGGTCTACATCAAGAGCTTTGTCAGCTATGC | 4 |
| AG876-80910-81030 | 34380235 | AACACCTGATCAGCGTGTTACATTTATGGAATTCCTGGGCTTTTTGCAACGCACGGAGTTGGTCTACATCAAGAGCTTTGTCAGCTATGCCTTGGGCACCACTAGTATCGAAACACCGTG | 4 |
| AG876-80940-81060 | 34553000 | ATTCCTGGGCTTTTTGCAACGCACGGAGTTGGTCTACATCAAGAGCTTTGTCAGCTATGCCTTGGGCACCACTAGTATCGAAACACCGTGGATGGATGAGAATCCTAGCACAGAGACGGC | 4 |
| AG876-80970-81090 | 34725765 | GGTCTACATCAAGAGCTTTGTCAGCTATGCCTTGGGCACCACTAGTATCGAAACACCGTGGATGGATGAGAATCCTAGCACAGAGACGGCACAGGCTTGGAATGCCGGCTTGCTCCGGGG | 4 |
| AG876-81000-81120 | 34898530 | CTTGGGCACCACTAGTATCGAAACACCGTGGATGGATGAGAATCCTAGCACAGAGACGGCACAGGCTTGGAATGCCGGCTTGCTCCGGGGGCGTGCGTACGGGCAAGACTTGCTTAGAAC | 4 |
| AG876-81030-81150 | 35071295 | GATGGATGAGAATCCTAGCACAGAGACGGCACAGGCTTGGAATGCCGGCTTGCTCCGGGGGCGTGCGTACGGGCAAGACTTGCTTAGAACTGAAGGAGAACATGGCGAAGGTGCTACCTG | 4 |
| AG876-81060-81180 | 35244060 | ACAGGCTTGGAATGCCGGCTTGCTCCGGGGGCGTGCGTACGGGCAAGACTTGCTTAGAACTGAAGGAGAACATGGCGAAGGTGCTACCTGTGAAACACGGGAAGAAAGTGAGGACACAGA | 4 |
| AG876-81090-81210 | 35416825 | GCGTGCGTACGGGCAAGACTTGCTTAGAACTGAAGGAGAACATGGCGAAGGTGCTACCTGTGAAACACGGGAAGAAAGTGAGGACACAGAGAGCGATGGGGATGATGAAGAACTTCCTCG | 4 |
| AG876-81120-81240 | 35589590 | TGAAGGAGAACATGGCGAAGGTGCTACCTGTGAAACACGGGAAGAAAGTGAGGACACAGAGAGCGATGGGGATGATGAAGAACTTCCTCGTGTAGTGTCCAGGGATGGAACTAAGCACAG | 4 |
| AG876-81150-81270 | 35762355 | TGAAACACGGGAAGAAAGTGAGGACACAGAGAGCGATGGGGATGATGAAGAACTTCCTCGTGTAGTGTCCAGGGATGGAACTAAGCACAGACGACCCCCTATATTTTTAAGACGCCTGCA | 4 |
| AG876-81180-81300 | 35935120 | GAGCGATGGGGATGATGAAGAACTTCCTCGTGTAGTGTCCAGGGATGGAACTAAGCACAGACGACCCCCTATATTTTTAAGACGCCTGCACAGGTTGCTGTTGATGAGAGCGGGCAAAGG | 4 |
| AG876-81210-81330 | 36107885 | TGTAGTGTCCAGGGATGGAACTAAGCACAGACGACCCCCTATATTTTTAAGACGCCTGCACAGGTTGCTGTTGATGAGAGCGGGCAAAGGAAAGGAACGGGCCAGGGAGACACTGGCGAA | 4 |
| AG876-81240-81360 | 36280650 | ACGACCCCCTATATTTTTAAGACGCCTGCACAGGTTGCTGTTGATGAGAGCGGGCAAAGGAAAGGAACGGGCCAGGGAGACACTGGCGAAGGCCCCTAGGCGCACTTATGGCACACCTAG | 4 |
| AG876-81270-81390 | 36453415 | CAGGTTGCTGTTGATGAGAGCGGGCAAAGGAAAGGAACGGGCCAGGGAGACACTGGCGAAGGCCCCTAGGCGCACTTATGGCACACCTAGGCCGCCAGTTCAGAAACCAAGACCAGAGGT | 4 |
| AG876-81300-81420 | 36626180 | AAAGGAACGGGCCAGGGAGACACTGGCGAAGGCCCCTAGGCGCACTTATGGCACACCTAGGCCGCCAGTTCAGAAACCAAGACCAGAGGTCCCGCAAAGCTATGAGACAGCTACCAGTCA | 4 |
| AG876-81330-81450 | 36798945 | GGCCCCTAGGCGCACTTATGGCACACCTAGGCCGCCAGTTCAGAAACCAAGACCAGAGGTCCCGCAAAGCTATGAGACAGCTACCAGTCACGGGTCGGCGCAAGTCCCAGAACCCCCACC | 4 |
| AG876-81360-81480 | 36971710 | GCCGCCAGTTCAGAAACCAAGACCAGAGGTCCCGCAAAGCTATGAGACAGCTACCAGTCACGGGTCGGCGCAAGTCCCAGAACCCCCACCCACCCACCCATTACATCAGCAACACAGCAT | 4 |
| AG876-81390-81510 | 37144475 | CCCGCAAAGCTATGAGACAGCTACCAGTCACGGGTCGGCGCAAGTCCCAGAACCCCCACCCACCCACCCATTACATCAGCAACACAGCATGGCCCCGTGTATGGTAGCTCAGAACCCACG | 4 |
| AG876-81420-81540 | 37317240 | CGGGTCGGCGCAAGTCCCAGAACCCCCACCCACCCACCCATTACATCAGCAACACAGCATGGCCCCGTGTATGGTAGCTCAGAACCCACGTGCACCCTTAGGGGACCAACTCCCAGGTGT | 4 |
| AG876-81450-81570 | 37490005 | CACCCACCCATTACATCAGCAACACAGCATGGCCCCGTGTATGGTAGCTCAGAACCCACGTGCACCCTTAGGGGACCAACTCCCAGGTGTTCCTAAAGATGGACGAGGGGCGTGTGCACC | 4 |
| AG876-81480-81600 | 37662770 | GGCCCCGTGTATGGTAGCTCAGAACCCACGTGCACCCTTAGGGGACCAACTCCCAGGTGTTCCTAAAGATGGACGAGGGGCGTGTGCACCGGTACCCGCCCTGGCCGGGCCTATTGTCCG | 4 |
| AG876-81510-81630 | 37835535 | TGCACCCTTAGGGGACCAACTCCCAGGTGTTCCTAAAGATGGACGAGGGGCGTGTGCACCGGTACCCGCCCTGGCCGGGCCTATTGTCCGGCCCTGGGAGTCATCCCTGTTACAGTCTCC | 4 |
| AG876-81540-81660 | 38008300 | TCCTAAAGATGGACGAGGGGCGTGTGCACCGGTACCCGCCCTGGCCGGGCCTATTGTCCGGCCCTGGGAGTCATCCCTGTTACAGTCTCCGGGAAGGGCCTTTGCACCCGTTAGCCCACA | 4 |
| AG876-81600-81720 | 38353830 | GCCCTGGGAGTCATCCCTGTTACAGTCTCCGGGAAGGGCCTTTGCACCCGTTAGCCCACAACCCATGCCAGTAGAACCCGTCCCTGTCCCTACTGTGGCACTTGAGCGACCAGTTTGTCC | 4 |
| AG876-81630-81750 | 38526595 | GGGAAGGGCCTTTGCACCCGTTAGCCCACAACCCATGCCAGTAGAACCCGTCCCTGTCCCTACTGTGGCACTTGAGCGACCAGTTTGTCCCGCACCTCCTGAGATTGCTATGCAGGGCCC | 4 |
| AG876-81660-81780 | 38699360 | ACCCATGCCAGTAGAACCCGTCCCTGTCCCTACTGTGGCACTTGAGCGACCAGTTTGTCCCGCACCTCCTGAGATTGCTATGCAGGGCCCGGGGGAACCTTCTGGCATTAAACGCACACG | 4 |
| AG876-81690-81810 | 38872125 | TACTGTGGCACTTGAGCGACCAGTTTGTCCCGCACCTCCTGAGATTGCTATGCAGGGCCCGGGGGAACCTTCTGGCATTAAACGCACACGGGAGCGTTGGAGGCCCGCACCGTGGACGCC | 4 |
| AG876-81720-81840 | 39044890 | CGCACCTCCTGAGATTGCTATGCAGGGCCCGGGGGAACCTTCTGGCATTAAACGCACACGGGAGCGTTGGAGGCCCGCACCGTGGACGCCAAACCCACCCCGCTCTCCCAGTCAGATGTC | 4 |
| AG876-81780-81900 | 39390420 | GGAGCGTTGGAGGCCCGCACCGTGGACGCCAAACCCACCCCGCTCTCCCAGTCAGATGTCCGTGCGTGACCGTCTGGCTCGTTTGCGTGCTGAGGCACAGGCCCGTCAGGCTAGTGTTGA | 8 |
| AG876-81810-81930 | 39563185 | AAACCCACCCCGCTCTCCCAGTCAGATGTCCGTGCGTGACCGTCTGGCTCGTTTGCGTGCTGAGGCACAGGCCCGTCAGGCTAGTGTTGAGGTGCAGCCCACCCAGTTGACCCAAGTATC | 4 |
| AG876-81960-82080 | 39735950 | ACCAGAGCAGCAGATGTTCCCTGGTTCCCCCTTTAGCCAGGTCGCTGATGTTGCCCGGGAATCTGGGGTACCTGCAATGCAGCCTCAGTACTTTGACCTCCCCTTAACTCAACCCATTAG | 4 |
| AG876-81990-82110 | 39908715 | CTTTAGCCAGGTCGCTGATGTTGCCCGGGAATCTGGGGTACCTGCAATGCAGCCTCAGTACTTTGACCTCCCCTTAACTCAACCCATTAGCCAGGGGGCACCCGCGGCCCCGTTGAGGGC | 4 |
| AG876-82020-82140 | 40081480 | ATCTGGGGTACCTGCAATGCAGCCTCAGTACTTTGACCTCCCCTTAACTCAACCCATTAGCCAGGGGGCACCCGCGGCCCCGTTGAGGGCTAGTATGGGCCCGGTACCTCCGGTACCGGC | 4 |
| AG876-82050-82170 | 40254245 | CTTTGACCTCCCCTTAACTCAACCCATTAGCCAGGGGGCACCCGCGGCCCCGTTGAGGGCTAGTATGGGCCCGGTACCTCCGGTACCGGCAACACAGCCACAGTATTTTGACATCCCCTT | 4 |
| AG876-83640-83760 | 40427010 | CAGAGGCGGTGGAAGCTCCTGTCACCTTACAGAACCTGGCGTATGGGTTACCGTACGCAAACCCTCAATGTAAACAGTTTTGAGACAGGGGGAGATAAAGTCCACCCACTCCTTGTGACT | 4 |
| AG876-83670-83790 | 40599775 | AGAACCTGGCGTATGGGTTACCGTACGCAAACCCTCAATGTAAACAGTTTTGAGACAGGGGGAGATAAAGTCCACCCACTCCTTGTGACTGCTACGCTAGGATGTGAAGAGGGCCTGCGG | 4 |
| AG876-83700-83820 | 40772540 | ACCCTCAATGTAAACAGTTTTGAGACAGGGGGAGATAAAGTCCACCCACTCCTTGTGACTGCTACGCTAGGATGTGAAGAGGGCCTGCGGCATGCAATAACTTACAGTGCTGGCATTGTA | 4 |
| AG876-83730-83850 | 40945305 | GGAGATAAAGTCCACCCACTCCTTGTGACTGCTACGCTAGGATGTGAAGAGGGCCTGCGGCATGCAATAACTTACAGTGCTGGCATTGTACAGCTACCACGAATGTCAGACCAAAACCAA | 4 |
| AG876-83760-83880 | 41118070 | GCTACGCTAGGATGTGAAGAGGGCCTGCGGCATGCAATAACTTACAGTGCTGGCATTGTACAGCTACCACGAATGTCAGACCAAAACCAAAAGATAGAAACAGCCTTCTTGATGGCACGT | 4 |
| AG876-83790-83910 | 41290835 | CATGCAATAACTTACAGTGCTGGCATTGTACAGCTACCACGAATGTCAGACCAAAACCAAAAGATAGAAACAGCCTTCTTGATGGCACGTCGTGCTAGGTCACTTTCGGCAGAAAGATAT | 4 |
| AG876-83820-83940 | 41463600 | CAGCTACCACGAATGTCAGACCAAAACCAAAAGATAGAAACAGCCTTCTTGATGGCACGTCGTGCTAGGTCACTTTCGGCAGAAAGATATACTTTGTTCTTTGATTTAGTATCCTCTGGA | 4 |
| AG876-84000-84120 | 41636365 | TTTGTAGGATGGCTATGTAAGAAGGACCACACTCATATACGTGAATGGTTCCGCCAGTGCACCGGGAGACCCTCACCATCCAAGCCATGGATGAGAGCGCATCCCGTCGCCGTTCCTTAT | 4 |
| AG876-84030-84150 | 41809130 | ACTCATATACGTGAATGGTTCCGCCAGTGCACCGGGAGACCCTCACCATCCAAGCCATGGATGAGAGCGCATCCCGTCGCCGTTCCTTATGACGATCCATTAACAAGTGAGGAGACTGAC | 4 |

| | | | | |
|---|---|---|---|---|
| AG876-84060-84180 | 41981895 | ACCGGGAGACCCTCACCATCCAAGCCATGGATGAGAGCGCATCCCGTCGCCGTTCCTTATGACGATCCATTAACAAGTGAGGAGACTGACCTGGCCTATGCCCGTGGACTGGCCATGAGT | | 4 |
| AG876-84090-84210 | 42154660 | ATGAGAGCGCATCCCGTCGCCGTTCCTTATGACGATCCATTAACAAGTGAGGAGACTGACCTGGCCTATGCCCGTGGACTGGCCATGAGTATCGAGGCTGCTAGACTGCCAGATGATCCA | | 4 |
| AG876-84120-84240 | 42327425 | GACGATCCATTAACAAGTGAGGAGACTGACCTGGCCTATGCCCGTGGACTGGCCATGAGTATCGAGGCTGCTAGACTGCCAGATGATCCAATAATTGTTGAGGATGACGATGAAAGTGAG | | 4 |
| AG876-84150-84270 | 42500190 | CTGGCCTATGCCCGTGGACTGGCCATGAGTATCGAGGCTGCTAGACTGCCAGATGATCCAATAATTGTTGAGGATGACGATGAAAGTGAGGAAATTGAAGATAAATGTGATAAGGATGAA | | 4 |
| AG876-84180-84300 | 42672955 | ATCGAGGCTGCTAGACTGCCAGATGATCCAATAATTGTTGAGGATGACGATGAAAGTGAGGAAATTGAAGATAAATGTGATAAGGATGAAGAGGAAAGTGGAACGGAAGATGTTACAAGC | | 4 |
| AG876-84210-84330 | 42845720 | ATAATTGTTGAGGATGACGATGAAAGTGAGGAAATTGAAGATAAATGTGATAAGGATGAAGAGGAAAGTGGAACGGAAGATGTTACAAGCATACCGCAAACACTGCCGCACAGTCCAACA | | 4 |
| AG876-84240-84360 | 43018485 | GAAATTGAAGATAAATGTGATAAGGATGAAGAGGAAAGTGGAACGGAAGATGTTACAAGCATACCGCAAACACTGCCGCACAGTCCAACAGTATACGGCAGGCCCTCGGTGTTTTACCGA | | 4 |
| AG876-84270-84390 | 43191250 | GAGGAAAGTGGAACGGAAGATGTTACAAGCATACCGCAAACACTGCCGCACAGTCCAACAGTATACGGCAGGCCCTCGGTGTTTTACCGAAAGCCAGATACTAAATCAACCAAAAAATGC | | 4 |
| AG876-84300-84420 | 43364015 | ATACCGCAAACACTGCCGCACAGTCCAACAGTATACGGCAGGCCCTCGGTGTTTTACCGAAAGCCAGATACTAAATCAACCAAAAAATGCAGGGCCATAGTGACTGACCTTAGTATAATC | | 4 |
| AG876-84330-84450 | 43536780 | GTATACGGCAGGCCCTCGGTGTTTTACCGAAAGCCAGATACTAAATCAACCAAAAAATGCAGGGCCATAGTGACTGACCTTAGTATAATCAAGGTCATTGAAGATGAACACAGAAAGAAG | | 4 |
| AG876-84360-84480 | 43709545 | AAGCCAGATACTAAATCAACCAAAAAATGCAGGGCCATAGTGACTGACCTTAGTATAATCAAGGTCATTGAAGATGAACACAGAAAGAAGAAGACAGCCAGAACAGAGCAACCAAGAGCC | | 4 |
| AG876-84390-84510 | 43882310 | AGGGCCATAGTGACTGACCTTAGTATAATCAAGGTCATTGAAGATGAACACAGAAAGAAGAAGACAGCCAGAACAGAGCAACCAAGAGCCAAGCCTGATTCCCCTGCCCCCACAGTGGTC | | 4 |
| AG876-84420-84540 | 44055075 | AAGGTCATTGAAGATGAACACAGAAAGAAGAAGACAGCCAGAACAGAGCAACCAAGAGCCAAGCCTGATTCCCCTGCCCCCACAGTGGTCCTTCGGCGACCACCCACGCAAAAGGTGACT | | 4 |
| AG876-84450-84570 | 44227840 | AAGACAGCCAGAACAGAGCAACCAAGAGCCAAGCCTGATTCCCCTGCCCCCACAGTGGTCCTTCGGCGACCACCCACGCAAAAGGTGACTGGCCCTGCCGGTTCACTGAGTGTCCAGGCT | | 4 |
| AG876-84480-84600 | 44400605 | AAGCCTGATTCCCCTGCCCCCACAGTGGTCCTTCGGCGACCACCCACGCAAAAGGTGACTGGCCCTGCCGGTTCACTGAGTGTCCAGGCTCAGCTGGAGCCATGGCAACCTTTGTCCTGG | | 4 |
| AG876-84510-84630 | 44573370 | CTTCGGCGACCACCCACGCAAAAGGTGACTGGCCCTGCCGGTTCACTGAGTGTCCAGGCTCAGCTGGAGCCATGGCAACCTTTGTCCTGGCCACATGAGACAAGAGTTATACTTCACGGA | | 4 |
| AG876-84540-84660 | 44746135 | GGCCCTGCCGGTTCACTGAGTGTCCAGGCTCAGCTGGAGCCATGGCAACCTTTGTCCTGGCCACATGAGACAAGAGTTATACTTCACGGACCACCCACGCAGGGTGACCAAGCACACGGT | | 4 |
| AG876-84570-84690 | 44918900 | CAGCTGGAGCCATGGCAACCTTTGTCCTGGCCACATGAGACAAGAGTTATACTTCACGGACCACCCACGCAGGGTGACCAAGCACACGGTTCCATGCTAGACCTTCTTGAAAAGGACGAC | | 4 |
| AG876-84600-84720 | 45091665 | CCACATGAGACAAGAGTTATACTTCACGGACCACCCACGCAGGGTGACCAAGCACACGGTTCCATGCTAGACCTTCTTGAAAAGGACGACCAGCACATGGAGCAGCAGGTTATGGCAACC | | 4 |
| AG876-84630-84750 | 45264430 | CCACCCACGCAGGGTGACCAAGCACACGGTTCCATGCTAGACCTTCTTGAAAAGGACGACCAGCACATGGAGCAGCAGGTTATGGCAACCCTACTGCCACCAGAACCACACCAGCCCCGG | | 4 |
| AG876-84660-84780 | 45437195 | TCCATGCTAGACCTTCTTGAAAAGGACGACCAGCACATGGAGCAGCAGGTTATGGCAACCCTACTGCCACCAGAACCACACCAGCCCCGGTCTGGGAGAAGAGCCCCTTGTGTCTACACC | | 4 |
| AG876-84690-84810 | 45609960 | CAGCACATGGAGCAGCAGGTTATGGCAACCCTACTGCCACCAGAACCACACCAGCCCCGGTCTGGGAGAAGAGCCCCTTGTGTCTACACCGCTGACCTAGACATAGAAAGTGATGAGCCC | | 4 |
| AG876-84720-84840 | 45782725 | CTACTGCCACCAGAACCACACCAGCCCCGGTCTGGGAGAAGAGCCCCTTGTGTCTACACCGCTGACCTAGACATAGAAAGTGATGAGCCCGCCACGTCAGAGCCGGTTCTTGATCAGCTA | | 4 |
| AG876-84750-84870 | 45955490 | TCTGGGAGAAGAGCCCCTTGTGTCTACACCGCTGACCTAGACATAGAAAGTGATGAGCCCGCCACGTCAGAGCCGGTTCTTGATCAGCTACTGCCCGCCCCAGGACTTGGACCTCTTGCA | | 4 |
| AG876-84780-84900 | 46128255 | GCTGACCTAGACATAGAAAGTGATGAGCCCGCCACGTCAGAGCCGGTTCTTGATCAGCTACTGCCCGCCCCAGGACTTGGACCTCTTGCAATTCAACCATTAACGTCCCCCACCACGTCT | | 4 |
| AG876-84810-84930 | 46301020 | GCCACGTCAGAGCCGGTTCTTGATCAGCTACTGCCCGCCCCAGGACTTGGACCTCTTGCAATTCAACCATTAACGTCCCCCACCACGTCTCAACTCCGCAGTTCAGCACCGAGCCACGCA | | 4 |
| AG876-84840-84960 | 46473785 | CTGCCCGCCCCAGGACTTGGACCTCTTGCAATTCAACCATTAACGTCCCCCACCACGTCTCAACTCCGCAGTTCAGCACCGAGCCACGCACAAACTCCATGGCCGGTTACCCACCCAAGT | | 4 |
| AG876-84870-84990 | 46646550 | ATTCAACCATTAACGTCCCCCACCACGTCTCAACTCCGCAGTTCAGCACCGAGCCACGCACAAACTCCATGGCCGGTTACCCACCCAAGTCAGACTCCAGGTGGCCCAACGACACAGTCC | | 4 |
| AG876-84900-85020 | 46819315 | CAACTCCGCAGTTCAGCACCGAGCCACGCACAAACTCCATGGCCGGTTACCCACCCAAGTCAGACTCCAGGTGGCCCAACGACACAGTCCCTGGCACCGGAAACAGAAGCCCCGCGCCAG | | 4 |
| AG876-84930-85050 | 46992080 | CAAACTCCATGGCCGGTTACCCACCCAAGTCAGACTCCAGGTGGCCCAACGACACAGTCCCTGGCACCGGAAACAGAAGCCCCGCGCCAGTGGCCAATGCCACTGCGACCTATCCCTCTG | | 4 |
| AG876-84960-85080 | 47164845 | CAGACTCCAGGTGGCCCAACGACACAGTCCCTGGCACCGGAAACAGAAGCCCCGCGCCAGTGGCCAATGCCACTGCGACCTATCCCTCTGCACCCCTTGCGGATGCAGCCAATATCATTT | | 4 |
| AG876-84990-85110 | 47337610 | CTGGCACCGGAAACAGAAGCCCCGCGCCAGTGGCCAATGCCACTGCGACCTATCCCTCTGCACCCCTTGCGGATGCAGCCAATATCATTTAATCCTGCAGTGAGACCCACTCCCCATCAG | | 4 |
| AG876-85020-85140 | 47510375 | TGGCCAATGCCACTGCGACCTATCCCTCTGCACCCCTTGCGGATGCAGCCAATATCATTTAATCCTGCAGTGAGACCCACTCCCCATCAGCCACCTCAGGTGGAGCCCACTTTCTATCAG | | 4 |
| AG876-85050-85170 | 47683140 | CACCCCTTGCGGATGCAGCCAATATCATTTAATCCTGCAGTGAGACCCACTCCCCATCAGCCACCTCAGGTGGAGCCCACTTTCTATCAGTCCACTTGGGTGAAACCCCCTCAACAATAC | | 4 |
| AG876-85080-85200 | 47855905 | AATCCTGCAGTGAGACCCACTCCCCATCAGCCACCTCAGGTGGAGCCCACTTTCTATCAGTCCACTTGGGTGAAACCCCCTCAACAATACCAGCCTCAGATGGGGCACATTCCATATCAG | | 4 |
| AG876-85110-85230 | 48028670 | CCACCTCAGGTGGAGCCCACTTTCTATCAGTCCACTTGGGTGAAACCCCCTCAACAATACCAGCCTCAGATGGGGCACATTCCATATCAGCCCCGACCAACGGGTCACTCTACTATGCTC | | 4 |
| AG876-85140-85260 | 48201435 | TCCACTTGGGTGAAACCCCCTCAACAATACCAGCCTCAGATGGGGCACATTCCATATCAGCCCCGACCAACGGGTCACTCTACTATGCTCCGGCCCCAGTGGGCACCCACCACCATGCAG | | 4 |
| AG876-85200-85320 | 48546965 | CCCCGACCAACGGGTCACTCTACTATGCTCCGGCCCCAGTGGGCACCCACCACCATGCAGCCACCACCAAGGGCGCCCACTCCCATGCCGCCACCTCAGGGGCCACCCACCGCTATGCAG | | 8 |
| AG876-85230-85350 | 48719730 | CGGCCCCAGTGGGCACCCACCACCATGCAGCCACCACCAAGGGCGCCCACTCCCATGCCGCCACCTCAGGGGCCACCCACCGCTATGCAGAGGCCTCAGGGGGCGCCCACTCCCATGCCG | | 8 |
| AG876-85260-85380 | 48892495 | CCACCACCAAGGGCGCCCACTCCCATGCCGCCACCTCAGGGGCCACCCACCGCTATGCAGAGGCCTCAGGGGGCGCCCACTCCCATGCCGCCACCTCAGGGGACACCCACCGCCATGCAG | | 8 |
| AG876-85290-85410 | 49065260 | CCACCTCAGGGGCCACCCACCGCTATGCAGAGGCCTCAGGGGGCGCCCACTCCCATGCCGCCACCTCAGGGGACACCCACCGCCATGCAGAGGCCTCAGGGTGCGCCCACTCCCATGCCG | | 8 |
| AG876-85320-85440 | 49238025 | AGGCCTCAGGGGGCGCCCACTCCCATGCCGCCACCTCAGGGGACACCCACCGCCATGCAGAGGCCTCAGGGTGCGCCCACTCCCATGCCGCCACCTCAGGGGACACCCACCGCCATGCAG | | 8 |
| AG876-85350-85470 | 49410790 | CCACCTCAGGGGACACCCACCGCCATGCAGAGGCCTCAGGGTGCGCCCACTCCCATGCCGCCACCTCAGGGGACACCCACCGCCATGCAGAGGCCTCAGGGTGCGCCCACTCCCATGCCG | | 8 |
| AG876-85380-85500 | 49583555 | AGGCCTCAGGGTGCGCCCACTCCCATGCCGCCACCTCAGGGGACACCCACCGCCATGCAGAGGCCTCAGGGTGCGCCCACTCCCATGCCGCCACCTCAGGGGACACCCACCGCCATACAG | | 8 |
| AG876-85410-85530 | 49756320 | CCACCTCAGGGGACACCCACCGCCATGCAGAGGCCTCAGGGTGCGCCCACTCCCATGCCGCCACCTCAGGGGACACCCACCGCCATACAGAGGCCTCAGGGTGCGCCCACTCCCATGCCG | | 8 |
| AG876-85440-85560 | 49929085 | AGGCCTCAGGGTGCGCCCACTCCCATGCCGCCACCTCAGGGGACACCCACCGCCATACAGAGGCCTCAGGGTGCGCCCACTCCCATGCCGCCACCTCAGGGGACACCCACCGCCATGCAG | | 8 |
| AG876-85470-85590 | 50101850 | CCACCTCAGGGGACACCCACCGCCATACAGAGGCCTCAGGGTGCGCCCACTCCCATGCCGCCACCTCAGGGGACACCCACCGCCATGCAGAGGCCTCAGGGGGCGCCCACTCCCATGCCG | | 8 |
| AG876-85500-85620 | 50274615 | AGGCCTCAGGGTGCGCCCACTCCCATGCCGCCACCTCAGGGGACACCCACCGCCATGCAGAGGCCTCAGGGGGCGCCCACTCCCATGCCGCCACCTCAGGGGACACCCACCGCCATGCAG | | 8 |
| AG876-85530-85650 | 50447380 | CCACCTCAGGGGACACCCACCGCCATGCAGAGGCCTCAGGGGGCGCCCACTCCCATGCCGCCACCTCAGGGGACACCCACCGCCATGCAGAGGCCTCAGGGGGCGCCCACTCCCATGCCG | | 8 |
| AG876-85560-85680 | 50620145 | AGGCCTCAGGGGGCGCCCACTCCCATGCCGCCACCTCAGGGGACACCCACCGCCATGCAGAGGCCTCAGGGGGCGCCCACTCCCATGCCGCCACCTCAGGGGCCACCCACCGCCATGCAG | | 8 |
| AG876-85590-85710 | 50792910 | CCACCTCAGGGGACACCCACCGCCATGCAGAGGCCTCAGGGGGCGCCCACTCCCATGCCGCCACCTCAGGGGCCACCCACCGCCATGCAGAGGCCTCGGGGGGCGCCCACTCCCATGCCG | | 8 |

| | | | |
|---|---|---|---|
| AG876-85620-85740 | 50965675 | AGGCCTCAGGGGGCGCCCACTCCCATGCCGCCACCTCAGGGGCCACCCACCGCCATGCAGAGGCCTCGGGGGGCGCCCACTCCCATGCCGCCACCTCAGGGGCCACCCACCGCCATGCAG | 12 |
| AG876-85650-85770 | 51138440 | CCACCTCAGGGGCCACCCACCGCCATGCAGAGGCCTCGGGGGGCGCCCACTCCCATGCCGCCACCTCAGGGGCCACCCACCGCCATGCAGAGGCCTCAGGGGGCGCCCACTCCCATGCCG | 12 |
| AG876-85680-85800 | 51311205 | AGGCCTCGGGGGGCGCCCACTCCCATGCCGCCACCTCAGGGGCCACCCACCGCCATGCAGAGGCCTCAGGGGGCGCCCACTCCCATGCCGCCACCTCAGGGGCCACCCACCGCCATGCAG | 12 |
| AG876-85710-85830 | 51483970 | CCACCTCAGGGGCCACCCACCGCCATGCAGAGGCCTCAGGGGGCGCCCACTCCCATGCCGCCACCTCAGGGGCCACCCACCGCCATGCAGAGGCCTCGGGGGGCGCCCACTCCCATGCCG | 12 |
| AG876-85740-85860 | 51656735 | AGGCCTCAGGGGGCGCCCACTCCCATGCCGCCACCTCAGGGGCCACCCACCGCCATGCAGAGGCCTCGGGGGGCGCCCACTCCCATGCCGCCACCTCAGGGGCCACCCACCGCCATGCAG | 12 |
| AG876-85770-85890 | 51829500 | CCACCTCAGGGGCCACCCACCGCCATGCAGAGGCCTCGGGGGGCGCCCACTCCCATGCCGCCACCTCAGGGGCCACCCACCGCCATGCAGAGGCCTCAGGGGGCGCCCACTCCCATGCCG | 12 |
| AG876-85800-85920 | 52002265 | AGGCCTCGGGGGGCGCCCACTCCCATGCCGCCACCTCAGGGGCCACCCACCGCCATGCAGAGGCCTCAGGGGGCGCCCACTCCCATGCCGCCACCTCAGGGGCCACCCACCGCCATGCAG | 12 |
| AG876-85830-85950 | 52175030 | CCACCTCAGGGGCCACCCACCGCCATGCAGAGGCCTCAGGGGGCGCCCACTCCCATGCCGCCACCTCAGGGGCCACCCACCGCCATGCAGAGGCCTCGGGGGGCGCCCACTCCCATGCCG | 12 |
| AG876-85860-85980 | 52347795 | AGGCCTCAGGGGGCGCCCACTCCCATGCCGCCACCTCAGGGGCCACCCACCGCCATGCAGAGGCCTCGGGGGGCGCCCACTCCCATGCCGCCACCTCAGGGGCCACCCACCGCCATGCAG | 12 |
| AG876-85890-86010 | 52520560 | CCACCTCAGGGGCCACCCACCGCCATGCAGAGGCCTCGGGGGGCGCCCACTCCCATGCCGCCACCTCAGGGGCCACCCACCGCCATGCAGCTGTCACCAAGGGCACTTACCGGCCAGAAG | 8 |
| AG876-85920-86040 | 52693325 | AGGCCTCGGGGGGCGCCCACTCCCATGCCGCCACCTCAGGGGCCACCCACCGCCATGCAGCTGTCACCAAGGGCACTTACCGGCCAGAAGGGGCCAGCAAAGCACATTCTGCGCCAGTTG | 8 |
| AG876-85950-86070 | 52866090 | CCACCTCAGGGGCCACCCACCGCCATGCAGCTGTCACCAAGGGCACTTACCGGCCAGAAGGGGCCAGCAAAGCACATTCTGCGCCAGTTGTTAACGGGGGGCGTCAAGAGTGGGAGACCA | 4 |
| AG876-85980-86100 | 53038855 | CTGTCACCAAGGGCACTTACCGGCCAGAAGGGGCCAGCAAAGCACATTCTGCGCCAGTTGTTAACGGGGGGCGTCAAGAGTGGGAGACCATCACTTAAGTTTAAGGCTGCCCTTGAGCGT | 4 |
| AG876-86010-86130 | 53211620 | GGGCCAGCAAAGCACATTCTGCGCCAGTTGTTAACGGGGGGCGTCAAGAGTGGGAGACCATCACTTAAGTTTAAGGCTGCCCTTGAGCGTCAAGCCGCTGCGGGCTTGCGACCTTCACCA | 4 |
| AG876-86040-86160 | 53384385 | TTAACGGGGGGCGTCAAGAGTGGGAGACCATCACTTAAGTTTAAGGCTGCCCTTGAGCGTCAAGCCGCTGCGGGCTTGCGACCTTCACCAGGGTCTGGAACGGGTGCCAAGATTGTGCAG | 4 |
| AG876-86070-86190 | 53557150 | TCACTTAAGTTTAAGGCTGCCCTTGAGCGTCAAGCCGCTGCGGGCTTGCGACCTTCACCAGGGTCTGGAACGGGTGCCAAGATTGTGCAGGCACCTGTTTTCTATCCACCCGTCCTACAG | 4 |
| AG876-86100-86220 | 53729915 | CAAGCCGCTGCGGGCTTGCGACCTTCACCAGGGTCTGGAACGGGTGCCAAGATTGTGCAGGCACCTGTTTTCTATCCACCCGTCCTACAGCCCATACAAGTTATGTGGCAAGTGGGTTCC | 4 |
| AG876-86130-86250 | 53902680 | GGGTCTGGAACGGGTGCCAAGATTGTGCAGGCACCTGTTTTCTATCCACCCGTCCTACAGCCCATACAAGTTATGTGGCAAGTGGGTTCCTCAAAGGCCGTGGCCGCCTCAACGGTGACA | 4 |
| AG876-86160-86280 | 54075445 | GCACCTGTTTTCTATCCACCCGTCCTACAGCCCATACAAGTTATGTGGCAAGTGGGTTCCTCAAAGGCCGTGGCCGCCTCAACGGTGACACAGGCACCCACGGAATATACCGGGGAAAGG | 4 |
| AG876-86190-86310 | 54248210 | CCCATACAAGTTATGTGGCAAGTGGGTTCCTCAAAGGCCGTGGCCGCCTCAACGGTGACACAGGCACCCACGGAATATACCGGGGAAAGGAGGTTAGGGGGGCCTATGTCTCCCACTGAT | 4 |
| AG876-86220-86340 | 54420975 | TCAAAGGCCGTGGCCGCCTCAACGGTGACACAGGCACCCACGGAATATACCGGGGAAAGGAGGTTAGGGGGGCCTATGTCTCCCACTGATATTCCGCCGTCTAAACGGGTGAAGAAAAAG | 4 |
| AG876-86250-86370 | 54593740 | CAGGCACCCACGGAATATACCGGGGAAAGGAGGTTAGGGGGGCCTATGTCTCCCACTGATATTCCGCCGTCTAAACGGGTGAAGAAAAAGGCCTATCCAGAGCGCAAGACGCCGCATGGG | 4 |
| AG876-86280-86400 | 54766505 | AGGTTAGGGGGGCCTATGTCTCCCACTGATATTCCGCCGTCTAAACGGGTGAAGAAAAAGGCCTATCCAGAGCGCAAGACGCCGCATGGGGGGCCCTCACACTCTTCCACCGTTATGTGG | 4 |
| AG876-86310-86430 | 54939270 | ATTCCGCCGTCTAAACGGGTGAAGAAAAAGGCCTATCCAGAGCGCAAGACGCCGCATGGGGGGCCCTCACACTCTTCCACCGTTATGTGGGAGAATGTCAGCCAGGGACAACAGCAGACT | 4 |
| AG876-86340-86460 | 55112035 | GCCTATCCAGAGCGCAAGACGCCGCATGGGGGGCCCTCACACTCTTCCACCGTTATGTGGGAGAATGTCAGCCAGGGACAACAGCAGACTCTGGAGTGCGGAGGAACTGATAAACAGGAA | 4 |
| AG876-86370-86490 | 55284800 | GGGCCCTCACACTCTTCCACCGTTATGTGGGAGAATGTCAGCCAGGGACAACAGCAGACTCTGGAGTGCGGAGGAACTGATAAACAGGAAAGGAACATGTTGGGGATGGGGGACATTGCA | 4 |
| AG876-86400-86520 | 55457565 | GAGAATGTCAGCCAGGGACAACAGCAGACTCTGGAGTGCGGAGGAACTGATAAACAGGAAAGGAACATGTTGGGGATGGGGGACATTGCAGTTTCTTCCCCTTCCTCTTCTGAAACATCG | 4 |
| AG876-86430-86550 | 55630330 | CTGGAGTGCGGAGGAACTGATAAACAGGAAAGGAACATGTTGGGGATGGGGGACATTGCAGTTTCTTCCCCTTCCTCTTCTGAAACATCGAATGATGAGTGATTTCAACCACGTAACAAG | 4 |
| AG876-86580-86700 | 55803095 | CTGCGGGGAGGGGCAGTGATAAGTCATGACAATTTTAGATGAGGTAGACATTTTGCATATTTTCAGACCCACCATGGAATCATTTGAAGGAGAGGGGGACTCTATACAGTCACCTGACAA | 4 |
| AG876-86610-86730 | 55975860 | AATTTTAGATGAGGTAGACATTTTGCATATTTTCAGACCCACCATGGAATCATTTGAAGGAGAGGGGGACTCTATACAGTCACCTGACAATGCGCGGGGAGATGATGTACAGAATACTGG | 4 |
| AG876-86640-86760 | 56148625 | TTTCAGACCCACCATGGAATCATTTGAAGGAGAGGGGGACTCTATACAGTCACCTGACAATGCGCGGGGAGATGATGTACAGAATACTGGTGAGCATATTCAGGACCCCGGGCCGGGGCC | 4 |
| AG876-86670-86790 | 56321390 | AGAGGGGGACTCTATACAGTCACCTGACAATGCGCGGGGAGATGATGTACAGAATACTGGTGAGCATATTCAGGACCCCGGGCCGGGGCCTTCAACCGGCGGGGCTTCTGAGGGATTGGT | 4 |
| AG876-86700-86820 | 56494155 | TGCGCGGGGAGATGATGTACAGAATACTGGTGAGCATATTCAGGACCCCGGGCCGGGGCCTTCAACCGGCGGGGCTTCTGAGGGATTGGTGCAGAACGAGCCGGACTCAAGAGATCAACA | 4 |
| AG876-86730-86850 | 56666920 | TGAGCATATTCAGGACCCCGGGCCGGGGCCTTCAACCGGCGGGGCTTCTGAGGGATTGGTGCAGAACGAGCCGGACTCAAGAGATCAACAGTCCCGGGGGCAGAGAAGGGGTGATGAAAA | 4 |
| AG876-86760-86880 | 56839685 | TTCAACCGGCGGGGCTTCTGAGGGATTGGTGCAGAACGAGCCGGACTCAAGAGATCAACAGTCCCGGGGGCAGAGAAGGGGTGATGAAAACAGAGGCTGGATGCAGCGCATCAGGCGAAG | 4 |
| AG876-86790-86910 | 57012450 | GCAGAACGAGCCGGACTCAAGAGATCAACAGTCCCGGGGGCAGAGAAGGGGTGATGAAAACAGAGGCTGGATGCAGCGCATCAGGCGAAGGCGGAGAAGACGGGCCGCATTGTCCGGCCA | 4 |
| AG876-86820-86940 | 57185215 | GTCCCGGGGGCAGAGAAGGGGTGATGAAAACAGAGGCTGGATGCAGCGCATCAGGCGAAGGCGGAGAAGACGGGCCGCATTGTCCGGCCATCTTTTAGACATGGAAGACAATGTGCCGCC | 4 |
| AG876-86850-86970 | 57357980 | CAGAGGCTGGATGCAGCGCATCAGGCGAAGGCGGAGAAGACGGGCCGCATTGTCCGGCCATCTTTTAGACATGGAAGACAATGTGCCGCCGTGGTTTCCTCCACACGATATCACACCATA | 4 |
| AG876-86880-87000 | 57530745 | GCGGAGAAGACGGGCCGCATTGTCCGGCCATCTTTTAGACATGGAAGACAATGTGCCGCCGTGGTTTCCTCCACACGATATCACACCATATGTCGCAAGGAATATCAGGGACGCTGCCTG | 4 |
| AG876-86910-87030 | 57703510 | TCTTTTAGACATGGAAGACAATGTGCCGCCGTGGTTTCCTCCACACGATATCACACCATATGTCGCAAGGAATATCAGGGACGCTGCCTGCCAGGCTGTCAAGGTGAGCATGCCTCTAAC | 4 |
| AG876-86940-87060 | 57876275 | GTGGTTTCCTCCACACGATATCACACCATATGTCGCAAGGAATATCAGGGACGCTGCCTGCCAGGCTGTCAAGGTGAGCATGCCTCTAACTGGGTTCATGGGGGCCATCTAAGGCCCACG | 4 |
| AG876-87390-87510 | 58049040 | TTCTCCGCCACCGTGTGGATACCACCCCCTGCTGGACCAAGAGAGCAAGAAAGATACGCTCGGGAAGCCGAGGTGCGCTTCCTTCGTGGTAAATGGCAGAGGCGGTTCCGAAGAATCTTT | 4 |
| AG876-87420-87540 | 58221805 | GCTGGACCAAGAGAGCAAGAAAGATACGCTCGGGAAGCCGAGGTGCGCTTCCTTCGTGGTAAATGGCAGAGGCGGTTCCGAAGAATCTTTGATTTGATAGAACTGTGTGGCTCTCTGCAC | 4 |
| AG876-87990-88110 | 58394570 | CCTACACCCAAGACGCACCCAGTGAAGCGCACAATTGTTAAGACCTCCTATAGATCTGATGAGGCAGAAGAAGCACAGAGCACCCCTGAAAGGCCGGGCCCTTCCAAACAACCATCAGAG | 4 |
| AG876-88020-88140 | 58567335 | ACAATTGTTAAGACCTCCTATAGATCTGATGAGGCAGAAGAAGCACAGAGCACCCCTGAAAGGCCGGGCCCTTCCAAACAACCATCAGAGCCCGTGGAGCCCGCCCACACAACCCCAGCG | 4 |
| AG876-88050-88170 | 58740100 | GAGGCAGAAGAAGCACAGAGCACCCCTGAAAGGCCGGGCCCTTCCAAACAACCATCAGAGCCCGTGGAGCCCGCCCACACAACCCCAGCGGGGAGGTCAACGGTGATTCTCCACGAACCA | 4 |
| AG876-88080-88200 | 58912865 | AGGCCGGGCCCTTCCAAACAACCATCAGAGCCCGTGGAGCCCGCCCACACAACCCCAGCGGGGAGGTCAACGGTGATTCTCCACGAACCACCTCGAGAGCCCGAAGCTGTTTCCTTCAAG | 4 |
| AG876-88140-88260 | 59258395 | GGGAGGTCAACGGTGATTCTCCACGAACCACCTCGAGAGCCCGAAGCTGTTTCCTTCAAGCCTCCGCCACCACCTTCCCGGAGGAGAAGGGGAGCGTGTGTTGTATATGACGATGATATC | 4 |
| AG876-88170-88290 | 59431160 | CCTCGAGAGCCCGAAGCTGTTTCCTTCAAGCCTCCGCCACCACCTTCCCGGAGGAGAAGGGGAGCGTGTGTTGTATATGACGATGATATCATAGAGGTGATTGATGTTGAAACCACCGAA | 4 |
| AG876-88200-88320 | 59603925 | CCTCCGCCACCACCTTCCCGGAGGAGAAGGGGAGCGTGTGTTGTATATGACGATGATATCATAGAGGTGATTGATGTTGAAACCACCGAAGAGGAGACGACGTCAATGCAAAGACAGCCT | 4 |
| AG876-88230-88350 | 59776690 | GGAGCGTGTGTTGTATATGACGATGATATCATAGAGGTGATTGATGTTGAAACCACCGAAGAGGAGACGACGTCAATGCAAAGACAGCCTCCGCTCGGGCAACAACCGCCCCCCCCCGTG | 4 |

| | | | |
|---|---|---|---|
| AG876-88260-88380 | 59949455 | ATAGAGGTGATTGATGTTGAAACCACCGAAGAGGAGACGACGTCAATGCAAAGACAGCCTCCGCTCGGGCAACAACCGCCCCCCCCCGTGATTTCTACCGGGAGTGCCATGTCTTCCAGT | 4 |
| AG876-88290-88410 | 60122220 | GAGGAGACGACGTCAATGCAAAGACAGCCTCCGCTCGGGCAACAACCGCCCCCCCCCGTGATTTCTACCGGGAGTGCCATGTCTTCCAGTCACACGGATCCATCAGTAACACAGCCAAGT | 4 |
| AG876-88320-88440 | 60294985 | CCGCTCGGGCAACAACCGCCCCCCCCCGTGATTTCTACCGGGAGTGCCATGTCTTCCAGTCACACGGATCCATCAGTAACACAGCCAAGTAAGCCACATCGGAAACCTCAAGACGGATTT | 4 |
| AG876-88350-88470 | 60467750 | ATTTCTACCGGGAGTGCCATGTCTTCCAGTCACACGGATCCATCAGTAACACAGCCAAGTAAGCCACATCGGAAACCTCAAGACGGATTTCAACGTTCAGGCCGACGTCAAAAACGAGCC | 4 |
| AG876-88380-88500 | 60640515 | CACACGGATCCATCAGTAACACAGCCAAGTAAGCCACATCGGAAACCTCAAGACGGATTTCAACGTTCAGGCCGACGTCAAAAACGAGCCATGCCTCCCCCAGTGAGTCCTTCTGACGCT | 4 |
| AG876-88410-88530 | 60813280 | AAGCCACATCGGAAACCTCAAGACGGATTTCAACGTTCAGGCCGACGTCAAAAACGAGCCATGCCTCCCCCAGTGAGTCCTTCTGACGCTGGGCCTCCTTCCACCAGGCCTCGTGTCATG | 4 |
| AG876-88440-88560 | 60986045 | CAACGTTCAGGCCGACGTCAAAAACGAGCCATGCCTCCCCCAGTGAGTCCTTCTGACGCTGGGCCTCCTTCCACCAGGCCTCGTGTCATGGCGCCTCCTTCCACCGGGCCTCGTGTCATG | 4 |
| AG876-88470-88590 | 61158810 | ATGCCTCCCCCAGTGAGTCCTTCTGACGCTGGGCCTCCTTCCACCAGGCCTCGTGTCATGGCGCCTCCTTCCACCGGGCCTCGTGTCATGGCGACTCCTTCCACCGGGCCTCGTGACATG | 8 |
| AG876-88500-88620 | 61331575 | GGGCCTCCTTCCACCAGGCCTCGTGTCATGGCGCCTCCTTCCACCGGGCCTCGTGTCATGGCGACTCCTTCCACCGGGCCTCGTGACATGGCGCCTCCTTCCACCGGGCCTCGTGACATG | 8 |
| AG876-88530-88650 | 61504340 | GCGCCTCCTTCCACCGGGCCTCGTGTCATGGCGACTCCTTCCACCGGGCCTCGTGACATGGCGCCTCCTTCCACCGGGCCTCGTGACATGGCGCCTCCTTCCACCGGGCCTCGTGACATG | 8 |
| AG876-88560-88680 | 61677105 | GCGACTCCTTCCACCGGGCCTCGTGACATGGCGCCTCCTTCCACCGGGCCTCGTGACATGGCGCCTCCTTCCACCGGGCCTCGTGACATGGCGCCTCCTTCCACCGGGCCTCGTGACATG | 8 |
| AG876-88590-88710 | 61849870 | GCGCCTCCTTCCACCGGGCCTCGTGACATGGCGCCTCCTTCCACCGGGCCTCGTGACATGGCGCCTCCTTCCACCGGGCCTCGTGACATGGCGCCTACCGTCGTACATATGTTTACGAGG | 8 |
| AG876-88620-88740 | 62022635 | GCGCCTCCTTCCACCGGGCCTCGTGACATGGCGCCTCCTTCCACCGGGCCTCGTGACATGGCGCCTACCGTCGTACATATGTTTACGAGGGAGCGCCTGCTCACCCAGTCCACCGGCCCT | 8 |
| AG876-88650-88770 | 62195400 | GCGCCTCCTTCCACCGGGCCTCGTGACATGGCGCCTACCGTCGTACATATGTTTACGAGGGAGCGCCTGCTCACCCAGTCCACCGGCCCTGCACCTCGGAGCTTCTGGGAAATGCGGGCC | 8 |
| AG876-88680-88800 | 62368165 | GCGCCTACCGTCGTACATATGTTTACGAGGGAGCGCCTGCTCACCCAGTCCACCGGCCCTGCACCTCGGAGCTTCTGGGAAATGCGGGCCGGCCGTGATGCTCCCAAAATTCAACAAGAA | 4 |
| AG876-88710-88830 | 62540930 | GAGCGCCTGCTCACCCAGTCCACCGGCCCTGCACCTCGGAGCTTCTGGGAAATGCGGGCCGGCCGTGATGCTCCCAAAATTCAACAAGAACCAAGTTCACAACAGCAGCCAGCCACTCAG | 4 |
| AG876-88740-88860 | 62713695 | GCACCTCGGAGCTTCTGGGAAATGCGGGCCGGCCGTGATGCTCCCAAAATTCAACAAGAACCAAGTTCACAACAGCAGCCAGCCACTCAGTCTACACCGCCTTGCCAATCATGGGTCCCA | 4 |
| AG876-88770-88890 | 62886460 | GGCCGTGATGCTCCCAAAATTCAACAAGAACCAAGTTCACAACAGCAGCCAGCCACTCAGTCTACACCGCCTTGCCAATCATGGGTCCCATCTGTCTATGTCCTCCCGGCAGTGGATGCT | 4 |
| AG876-88800-88920 | 63059225 | CCAAGTTCACAACAGCAGCCAGCCACTCAGTCTACACCGCCTTGCCAATCATGGGTCCCATCTGTCTATGTCCTCCCGGCAGTGGATGCTGGTAATGCCCAGCCCCTACAAATATCACAC | 4 |
| AG876-88830-88950 | 63231990 | TCTACACCGCCTTGCCAATCATGGGTCCCATCTGTCTATGTCCTCCCGGCAGTGGATGCTGGTAATGCCCAGCCCCTACAAATATCACACTTGAGCTCCATGTCGCCCACGCAGCCGATA | 4 |
| AG876-88860-88980 | 63404755 | TCTGTCTATGTCCTCCCGGCAGTGGATGCTGGTAATGCCCAGCCCCTACAAATATCACACTTGAGCTCCATGTCGCCCACGCAGCCGATATCGCATGAAGAACAACCCCGGTATGAGGAT | 4 |
| AG876-88890-89010 | 63577520 | GGTAATGCCCAGCCCCTACAAATATCACACTTGAGCTCCATGTCGCCCACGCAGCCGATATCGCATGAAGAACAACCCCGGTATGAGGATCCTGACACTCCTCTGGATTTGAGTTTACAT | 4 |
| AG876-88920-89040 | 63750285 | TTGAGCTCCATGTCGCCCACGCAGCCGATATCGCATGAAGAACAACCCCGGTATGAGGATCCTGACACTCCTCTGGATTTGAGTTTACATCCAGATACTGCAACTCTGCCGCCCACCCAG | 4 |
| AG876-88950-89070 | 63923050 | TCGCATGAAGAACAACCCCGGTATGAGGATCCTGACACTCCTCTGGATTTGAGTTTACATCCAGATACTGCAACTCTGCCGCCCACCCAGGATTTATACCCTGGACGCGAGGATCTGCAG | 4 |
| AG876-88980-89100 | 64095815 | CCTGACACTCCTCTGGATTTGAGTTTACATCCAGATACTGCAACTCTGCCGCCCACCCAGGATTTATACCCTGGACGCGAGGATCTGCAGGCCACCCAGGCTCCATACCCGGGATACGAG | 4 |
| AG876-89010-89130 | 64268580 | CCAGATACTGCAACTCTGCCGCCCACCCAGGATTTATACCCTGGACGCGAGGATCTGCAGGCCACCCAGGCTCCATACCCGGGATACGAGGAGCCACGGCCCCCCCAGGCTCCTTTTGTA | 4 |
| AG876-89040-89160 | 64441345 | GATTTATACCCTGGACGCGAGGATCTGCAGGCCACCCAGGCTCCATACCCGGGATACGAGGAGCCACGGCCCCCCCAGGCTCCTTTTGTAGGTGACTATGGTTTTGTTCAAATTCCCTCG | 4 |
| AG876-89070-89190 | 64614110 | GCCACCCAGGCTCCATACCCGGGATACGAGGAGCCACGGCCCCCCCAGGCTCCTTTTGTAGGTGACTATGGTTTTGTTCAAATTCCCTCGGCTCAATGGGAGCCACACCCCTCTCAGGGC | 4 |
| AG876-89100-89220 | 64786875 | GAGCCACGGCCCCCCCAGGCTCCTTTTGTAGGTGACTATGGTTTTGTTCAAATTCCCTCGGCTCAATGGGAGCCACACCCCTCTCAGGGCACATACCAGGGCCACATTGATCCCCAGCTT | 4 |
| AG876-89130-89250 | 64959640 | GGTGACTATGGTTTTGTTCAAATTCCCTCGGCTCAATGGGAGCCACACCCCTCTCAGGGCACATACCAGGGCCACATTGATCCCCAGCTTCCGGCTGCTCTAGATTTAGGGCCAGAGCAG | 4 |
| AG876-89160-89280 | 65132405 | GCTCAATGGGAGCCACACCCCTCTCAGGGCACATACCAGGGCCACATTGATCCCCAGCTTCCGGCTGCTCTAGATTTAGGGCCAGAGCAGCCCCGGTTCCCACAAGATCCATATGTGTAT | 4 |
| AG876-89190-89310 | 65305170 | ACATACCAGGGCCACATTGATCCCCAGCTTCCGGCTGCTCTAGATTTAGGGCCAGAGCAGCCCCGGTTCCCACAAGATCCATATGTGTATTCCGGGGGCCAATTGTCTTCATGTCCAGGT | 4 |
| AG876-89220-89340 | 65477935 | CCGGCTGCTCTAGATTTAGGGCCAGAGCAGCCCCGGTTCCCACAAGATCCATATGTGTATTCCGGGGGCCAATTGTCTTCATGTCCAGGTTATGCAGGTCCCTGGCCCTCGAGGCCTCAA | 4 |
| AG876-89250-89370 | 65650700 | CCCCGGTTCCCACAAGATCCATATGTGTATTCCGGGGGCCAATTGTCTTCATGTCCAGGTTATGCAGGTCCCTGGCCCTCGAGGCCTCAACATCCAAGATATAGGCACACCTTGGCATTG | 4 |
| AG876-89280-89400 | 65823465 | TCCGGGGGCCAATTGTCTTCATGTCCAGGTTATGCAGGTCCCTGGCCCTCGAGGCCTCAACATCCAAGATATAGGCACACCTTGGCATTGTGGCCTCGAGAACCCAGGCATGGGCACTCT | 4 |
| AG876-89310-89430 | 65996230 | TATGCAGGTCCCTGGCCCTCGAGGCCTCAACATCCAAGATATAGGCACACCTTGGCATTGTGGCCTCGAGAACCCAGGCATGGGCACTCTCAGGGCCCATGGAAACCCTGGTCAGCACAT | 4 |
| AG876-89340-89460 | 66168995 | CATCCAAGATATAGGCACACCTTGGCATTGTGGCCTCGAGAACCCAGGCATGGGCACTCTCAGGGCCCATGGAAACCCTGGTCAGCACATCTCCCACCTCAGTGGGATGGATCTGCAGGA | 4 |
| AG876-89370-89490 | 66341760 | TGGCCTCGAGAACCCAGGCATGGGCACTCTCAGGGCCCATGGAAACCCTGGTCAGCACATCTCCCACCTCAGTGGGATGGATCTGCAGGACATGGCCAGGATCAGGTCTCCCAGTTCCCA | 4 |
| AG876-89400-89520 | 66514525 | CAGGGCCCATGGAAACCCTGGTCAGCACATCTCCCACCTCAGTGGGATGGATCTGCAGGACATGGCCAGGATCAGGTCTCCCAGTTCCCACATCTGCATTCGGAGACAGGCCCACCACGT | 4 |
| AG876-89430-89550 | 66687290 | CTCCCACCTCAGTGGGATGGATCTGCAGGACATGGCCAGGATCAGGTCTCCCAGTTCCCACATCTGCATTCGGAGACAGGCCCACCACGTCTTCAACTTTCATCGGTGCCACAGGTGCTG | 4 |
| AG876-89550-89670 | 66860055 | TACCCGCAGCCACTGGTCTCATCCTCTGCACCATCGTGGTCATCTCCCCAGCCCCGAGCCCCCATACGCCCCATTCCAACAAGATTCCCCCCTCCCCCTATGCCGTTACAAGATAGCATG | 4 |
| AG876-89580-89700 | 67032820 | CCATCGTGGTCATCTCCCCAGCCCCGAGCCCCCATACGCCCCATTCCAACAAGATTCCCCCCTCCCCCTATGCCGTTACAAGATAGCATGGCCGTGGGGTGCGACTCATCAGGTACAGCA | 4 |
| AG876-91110-91230 | 67205585 | CTGCCGCCCTCCTTGAGCCCCACCATCCGCTGCCACCCTCTTTGAGCCCCACCGTCCGCTGCCACCCTCTTTGAGCCCCTTCTTACCGATTCTGGCTGTAGTGGTTTCCTTGTACGTCGT | 4 |
| AG876-91140-91260 | 67378350 | TGCCACCCTCTTTGAGCCCCACCGTCCGCTGCCACCCTCTTTGAGCCCCTTCTTACCGATTCTGGCTGTAGTGGTTTCCTTGTACGTCGTGCCGGGGCAGCCGCTGGTGCAGGCTGTGGA | 4 |
| AG876-91170-91290 | 67551115 | GCCACCCTCTTTGAGCCCCTTCTTACCGATTCTGGCTGTAGTGGTTTCCTTGTACGTCGTGCCGGGGCAGCCGCTGGTGCAGGCTGTGGAGCACCAATGTCTGCTAGCTGTTGTCCTTGG | 4 |
| AG876-91200-91320 | 67723880 | TCTGGCTGTAGTGGTTTCCTTGTACGTCGTGCCGGGGCAGCCGCTGGTGCAGGCTGTGGAGCACCAATGTCTGCTAGCTGTTGTCCTTGGTTAGCCCCGGGACAAGCAAACACCACTGCT | 4 |
| AG876-96750-96870 | 67896645 | GTGGAACAGGAGCAGGAGGAGGGGCAGGAGCAGGAGGGGCAGGAGCAGGAGGGGCAGGAGCAGGAGGGGCAGGAGCAGGAGGGGCAGGAGCAGGAGGGGCAGGAGCAGGAGGGGCAGGAG | 8 |
| AG876-96780-96900 | 68069410 | CAGGAGGGGCAGGAGCAGGAGGGGCAGGAGCAGGAGGGGCAGGAGCAGGAGGGGCAGGAGCAGGAGGGGCAGGAGCAGGAGGGGCAGGAGCAGGAGGGGCAGGAGCAGGAGGGGCAGGAG | 8 |
| AG876-96810-96930 | 68242175 | CAGGAGGGGCAGGAGCAGGAGGGGCAGGAGCAGGAGGGGCAGGAGCAGGAGGGGCAGGAGCAGGAGGGGCAGGAGCAGGAGGGGCAGGAGCAGGAGGGGCAGGAGCAGGAGGAGGGGCAG | 8 |
| AG876-96840-96960 | 68414940 | CAGGAGGGGCAGGAGCAGGAGGGGCAGGAGCAGGAGGGGCAGGAGCAGGAGGGGCAGGAGCAGGAGGGGCAGGAGCAGGAGGAGGGGCAGGAGCAGGAGGGGCAGGAGCAGGAGGGGCAG | 8 |
| AG876-96870-96990 | 68587705 | CAGGAGGGGCAGGAGCAGGAGGGGCAGGAGCAGGAGGGGCAGGAGCAGGAGGAGGGGCAGGAGCAGGAGGGGCAGGAGCAGGAGGGGCAGGAGCAGGAGGAGGGGCAGGAGCAGGAGGAG | 8 |

| | | | |
|---|---|---|---|
| AG876-96900-97020 | 68760470 | CAGGAGGGGCAGGAGCAGGAGGAGGGGCAGGAGCAGGAGGGGCAGGAGCAGGAGGGGCAGGAGCAGGAGGAGGGGCAGGAGCAGGAGGAGGGGCAGGAGCAGGAGGAGGGGCAGGAGCAG | 8 |
| AG876-97050-97170 | 68933235 | GAGCAGGAGGAGGGGCAGGAGCAGGAGGAGGGGCAGGAGCAGGAGGAGGGGCAGGAGCAGGAGGAGGGGCAGGAGCAGGCGGGGCAGGAGCAGGCGGGGCAGGAGCAGGAGGAGGGGCAG | 8 |
| AG876-97080-97200 | 69106000 | GGGCAGGAGCAGGAGGAGGGGCAGGAGCAGGAGGAGGGGCAGGAGCAGGCGGGGCAGGAGCAGGCGGGGCAGGAGCAGGAGGAGGGGCAGGAGCAGGAGGAGGGGCAGGAGCAGGAGGAG | 8 |
| AG876-97110-97230 | 69278765 | GAGGAGGGGCAGGAGCAGGCGGGGCAGGAGCAGGCGGGGCAGGAGCAGGAGGAGGGGCAGGAGCAGGAGGAGGGGCAGGAGCAGGAGGAGGGGCAGGAGCAGGAGGAGGGGCAGGAGCAG | 8 |
| AG876-97140-97260 | 69451530 | CAGGCGGGGCAGGAGCAGGAGGAGGGGCAGGAGCAGGAGGAGGGGCAGGAGCAGGAGGAGGGGCAGGAGCAGGAGGAGGGGCAGGAGCAGGAGGAGGGGCAGGAGCAGGAGGAGGGGCAG | 8 |
| AG876-97170-97290 | 69624295 | GAGCAGGAGGAGGGGCAGGAGCAGGAGGAGGGGCAGGAGCAGGAGGAGGGGCAGGAGCAGGAGGAGGGGCAGGAGCAGGAGGAGGGGCAGGAGCAGGAGGAGGGGCAGGAGCAGGAGGAG | 8 |
| AG876-97200-97320 | 69797060 | GGGCAGGAGCAGGAGGAGGGGCAGGAGCAGGAGGAGGGGCAGGAGCAGGAGGAGGGGCAGGAGCAGGAGGAGGGGCAGGAGCAGGAGGAGGGGCAGGAGCAGGAGGAGGGGCAGGAGCAG | 8 |
| AG876-97230-97350 | 69969825 | GAGGAGGGGCAGGAGCAGGAGGAGGGGCAGGAGCAGGAGGAGGGGCAGGAGCAGGAGGAGGGGCAGGAGCAGGAGGAGGGGCAGGAGCAGGAGGAGGGGCAGGAGCAGGAGGAGGGGCAG | 8 |
| AG876-97260-97380 | 70142590 | GAGCAGGAGGAGGGGCAGGAGCAGGAGGAGGGGCAGGAGCAGGAGGAGGGGCAGGAGCAGGAGGAGGGGCAGGAGCAGGAGGAGGGGCAGGAGCAGGAGGAGGGGCAGGAGCAGGAGGAG | 8 |
| AG876-97290-97410 | 70315355 | GGGCAGGAGCAGGAGGAGGGGCAGGAGCAGGAGGAGGGGCAGGAGCAGGAGGAGGGGCAGGAGCAGGAGGAGGGGCAGGAGCAGGAGGAGGGGCAGGAGCAGGAGGAGGGGCAGGAGCAG | 8 |
| AG876-97320-97440 | 70488120 | GAGGAGGGGCAGGAGCAGGAGGAGGGGCAGGAGCAGGAGGAGGGGCAGGAGCAGGAGGAGGGGCAGGAGCAGGAGGAGGGGCAGGAGCAGGAGGAGGGGCAGGAGCAGGAGGAGGGGCAG | 8 |
| AG876-97350-97470 | 70660885 | GAGCAGGAGGAGGGGCAGGAGCAGGAGGAGGGGCAGGAGCAGGAGGAGGGGCAGGAGCAGGAGGAGGGGCAGGAGCAGGAGGAGGGGCAGGAGCAGGAGGAGGGGCAGGAGCAGGAGGTG | 8 |
| AG876-97380-97500 | 70833650 | GGGCAGGAGCAGGAGGAGGGGCAGGAGCAGGAGGAGGGGCAGGAGCAGGAGGAGGGGCAGGAGCAGGAGGAGGGGCAGGAGCAGGAGGTGGAGGCCGGGGTCGAGGAGGCAGTGGAGGCC | 8 |
| AG876-97410-97530 | 71006415 | GAGGAGGGGCAGGAGCAGGAGGAGGGGCAGGAGCAGGAGGAGGGGCAGGAGCAGGAGGTGGAGGCCGGGGTCGAGGAGGCAGTGGAGGCCGGGGTCGAGGAGGTAGTGGAGGCCGGGGTC | 8 |
| AG876-97440-97560 | 71179180 | GAGCAGGAGGAGGGGCAGGAGCAGGAGGTGGAGGCCGGGGTCGAGGAGGCAGTGGAGGCCGGGGTCGAGGAGGTAGTGGAGGCCGGGGTCGAGGAGGTAGTGGAGGCCGCCGGGGTAGAG | 8 |
| AG876-141690-141810 | 71351945 | CGGCCCTGGAGCTCGGGGGGCGGCCGGGTGGCCCACCGGGTCCGCTGGGTCCGCTGCCCCGCTCCCGCGGGGGGGTGGCCGGCTGCAGCCGGGTCCGGGGTTCCGGCCCTGGAGCTCGGG | 12 |
| AG876-141720-141840 | 71524710 | GCCCACCGGGTCCGCTGGGTCCGCTGCCCCGCTCCCGCGGGGGGGTGGCCGGCTGCAGCCGGGTCCGGGGTTCCGGCCCTGGAGCTCGGGGGGCGGCCGGGTGGCCCACCGGGTCCGCTG | 12 |
| AG876-141750-141870 | 71697475 | GCTCCCGCGGGGGGGTGGCCGGCTGCAGCCGGGTCCGGGGTTCCGGCCCTGGAGCTCGGGGGGCGGCCGGGTGGCCCACCGGGTCCGCTGGGTCCGCTGCCCCGCTCCGGCGGGGGGTGG | 12 |
| AG876-141780-141900 | 71870240 | GGGTCCGGGGTTCCGGCCCTGGAGCTCGGGGGGCGGCCGGGTGGCCCACCGGGTCCGCTGGGTCCGCTGCCCCGCTCCGGCGGGGGGTGGCCGGCTGCAGCCGGGTCCGGGGTTCCGGCC | 12 |
| AG876-141960-142080 | 72043005 | GGCGGGGGGGTGGCCGGCTGCAGCCGGGTCCGGGGTTCCGGCCCTGGAGCTCGGGGGGCGGCCGGGTGGCCCACCGGGTCCGCTGGGTCCGCTGCCCCGCTCCGGCGGGGGGGTGGCCGG | 16 |
| AG876-141990-142110 | 72215770 | CGGGGTTCCGGCCCTGGAGCTCGGGGGGCGGCCGGGTGGCCCACCGGGTCCGCTGGGTCCGCTGCCCCGCTCCGGCGGGGGGGTGGCCGGCTGCAGCCGGGTCCGGGGTTCCGGCCCTGG | 12 |
| AG876-142020-142140 | 72388535 | GCCGGGTGGCCCACCGGGTCCGCTGGGTCCGCTGCCCCGCTCCGGCGGGGGGGTGGCCGGCTGCAGCCGGGTCCGGGGTTCCGGCCCTGGAGCTCGGGGGGCGGCCGGGTGGCCCACCGG | 12 |
| AG876-142050-142170 | 72561300 | GCTGCCCCGCTCCGGCGGGGGGGTGGCCGGCTGCAGCCGGGTCCGGGGTTCCGGCCCTGGAGCTCGGGGGGCGGCCGGGTGGCCCACCGGGTCCGCTGGGTCCGCTGCCCCGCTCCGGCG | 12 |
| AG876-142080-142200 | 72734065 | CTGCAGCCGGGTCCGGGGTTCCGGCCCTGGAGCTCGGGGGGCGGCCGGGTGGCCCACCGGGTCCGCTGGGTCCGCTGCCCCGCTCCGGCGGGGGGGTGGCCGGCTGCAGCCGGGTCCGGG | 12 |
| AG876-142110-142230 | 72906830 | AGCTCGGGGGGCGGCCGGGTGGCCCACCGGGTCCGCTGGGTCCGCTGCCCCGCTCCGGCGGGGGGGTGGCCGGCTGCAGCCGGGTCCGGGGTTCCGGCCCTGGAGCTCGGGGGGCGGCCG | 12 |
| AG876-142140-142260 | 73079595 | GTCCGCTGGGTCCGCTGCCCCGCTCCGGCGGGGGGGTGGCCGGCTGCAGCCGGGTCCGGGGTTCCGGCCCTGGAGCTCGGGGGGCGGCCGGGTGGCCCACCGGGTCCGCTGGGTCCGCTG | 12 |
| AG876-142170-142290 | 73252360 | GGGGGGTGGCCGGCTGCAGCCGGGTCCGGGGTTCCGGCCCTGGAGCTCGGGGGGCGGCCGGGTGGCCCACCGGGTCCGCTGGGTCCGCTGCCCCGCTCCGGCGGGGGGGTGGCCGGCTGC | 12 |
| AG876-142200-142320 | 73425125 | GTTCCGGCCCTGGAGCTCGGGGGGCGGCCGGGTGGCCCACCGGGTCCGCTGGGTCCGCTGCCCCGCTCCGGCGGGGGGGTGGCCGGCTGCAGCCGGGTCCGGGGTTCCGGCCCTGGAGCT | 12 |
| AG876-142230-142350 | 73597890 | GGTGGCCCACCGGGTCCGCTGGGTCCGCTGCCCCGCTCCGGCGGGGGGGTGGCCGGCTGCAGCCGGGTCCGGGGTTCCGGCCCTGGAGCTCGGGGGGCGGCCGGGTGGCCCACCGGGTCC | 12 |
| AG876-142260-142380 | 73770655 | CCCCGCTCCGGCGGGGGGGTGGCCGGCTGCAGCCGGGTCCGGGGTTCCGGCCCTGGAGCTCGGGGGGCGGCCGGGTGGCCCACCGGGTCCGCTGGGTCCGCTGCCCCGCTCCGGCGGGGG | 16 |
| AG876-142290-142410 | 73943420 | AGCCGGGTCCGGGGTTCCGGCCCTGGAGCTCGGGGGGCGGCCGGGTGGCCCACCGGGTCCGCTGGGTCCGCTGCCCCGCTCCGGCGGGGGGGTGGCCGGCTGCAGCCGGGTCCGGGGTTC | 12 |
| AG876-142320-142440 | 74116185 | CGGGGGGCGGCCGGGTGGCCCACCGGGTCCGCTGGGTCCGCTGCCCCGCTCCGGCGGGGGGGTGGCCGGCTGCAGCCGGGTCCGGGGTTCCGGCCCTGGAGCTCGGGGGGCGGCCGGGTG | 16 |
| AG876-142350-142470 | 74288950 | GCTGGGTCCGCTGCCCCGCTCCGGCGGGGGGGTGGCCGGCTGCAGCCGGGTCCGGGGTTCCGGCCCTGGAGCTCGGGGGGCGGCCGGGTGGCCCACCGGGTCCGCTGGGTCCGCTGCCCC | 12 |
| AG876-142380-142500 | 74461715 | GGTGGCCGGCTGCAGCCGGGTCCGGGGTTCCGGCCCTGGAGCTCGGGGGGCGGCCGGGTGGCCCACCGGGTCCGCTGGGTCCGCTGCCCCGCTCCGGCGGGGGGGTGGCCGGCTGCAGCC | 12 |
| AG876-142410-142530 | 74634480 | CGGCCCTGGAGCTCGGGGGGCGGCCGGGTGGCCCACCGGGTCCGCTGGGTCCGCTGCCCCGCTCCGGCGGGGGGGTGGCCGGCTGCAGCCGGGTCCGGGGTTCCGGCCCTGGAGCTCGGG | 12 |
| AG876-142440-142560 | 74807245 | GCCCACCGGGTCCGCTGGGTCCGCTGCCCCGCTCCGGCGGGGGGGTGGCCGGCTGCAGCCGGGTCCGGGGTTCCGGCCCTGGAGCTCGGGGGGCGGCCGGGTGGCCCACCGGGTCCGCTG | 12 |
| AG876-142470-142590 | 74980010 | GCTCCGGCGGGGGGGTGGCCGGCTGCAGCCGGGTCCGGGGTTCCGGCCCTGGAGCTCGGGGGGCGGCCGGGTGGCCCACCGGGTCCGCTGGGTCCGCTGCCCCGCTCCGGCGGGGGGGTG | 12 |
| AG876-142500-142620 | 75152775 | GGGTCCGGGGTTCCGGCCCTGGAGCTCGGGGGGCGGCCGGGTGGCCCACCGGGTCCGCTGGGTCCGCTGCCCCGCTCCGGCGGGGGGGTGGCCGGCTGCAGCCGGGTCCGGGGTTCCGGC | 12 |
| AG876-142530-142650 | 75325540 | GGGCGGCCGGGTGGCCCACCGGGTCCGCTGGGTCCGCTGCCCCGCTCCGGCGGGGGGGTGGCCGGCTGCAGCCGGGTCCGGGGTTCCGGCCCTGGAGCTCGGGGGGCGGCCGGGTGGCCC | 16 |
| AG876-142560-142680 | 75498305 | GGTCCGCTGCCCCGCTCCGGCGGGGGGGTGGCCGGCTGCAGCCGGGTCCGGGGTTCCGGCCCTGGAGCTCGGGGGGCGGCCGGGTGGCCCACCGGGTCCGCTGGGTCCGCTGCCCCGCTC | 12 |
| AG876-142590-142710 | 75671070 | GCCGGCTGCAGCCGGGTCCGGGGTTCCGGCCCTGGAGCTCGGGGGGCGGCCGGGTGGCCCACCGGGTCCGCTGGGTCCGCTGCCCCGCTCCGGCGGGGGGGTGGCCGGCTGCAGCCGGGT | 12 |
| AG876-142620-142740 | 75843835 | CCTGGAGCTCGGGGGGCGGCCGGGTGGCCCACCGGGTCCGCTGGGTCCGCTGCCCCGCTCCGGCGGGGGGGTGGCCGGCTGCAGCCGGGTCCGGGGTTCCGGCCCTGGAGCTCGGGGGGC | 12 |
| AG876-142650-142770 | 76016600 | ACCGGGTCCGCTGGGTCCGCTGCCCCGCTCCGGCGGGGGGGTGGCCGGCTGCAGCCGGGTCCGGGGTTCCGGCCCTGGAGCTCGGGGGGCGGCCGGGTGGCCCACCGGGTCCGCTGGGTC | 12 |
| AG876-142680-142800 | 76189365 | CGGCGGGGGGGTGGCCGGCTGCAGCCGGGTCCGGGGTTCCGGCCCTGGAGCTCGGGGGGCGGCCGGGTGGCCCACCGGGTCCGCTGGGTCCGCTGCCCCGCTCCGGCGGGGGGGTGGCCG | 16 |
| AG876-142710-142830 | 76362130 | CCGGGGTTCCGGCCCTGGAGCTCGGGGGGCGGCCGGGTGGCCCACCGGGTCCGCTGGGTCCGCTGCCCCGCTCCGGCGGGGGGGTGGCCGGCTGCAGCCGGGTCCGGGGTTCCGGCCCTG | 12 |
| AG876-142740-142860 | 76534895 | GGCCGGGTGGCCCACCGGGTCCGCTGGGTCCGCTGCCCCGCTCCGGCGGGGGGGTGGCCGGCTGCAGCCGGGTCCGGGGTTCCGGCCCTGGAGCTCGGGGGGCGGCCGGGTGGCCCACCG | 12 |
| AG876-142770-142890 | 76707660 | CGCTGCCCCGCTCCGGCGGGGGGGTGGCCGGCTGCAGCCGGGTCCGGGGTTCCGGCCCTGGAGCTCGGGGGGCGGCCGGGTGGCCCACCGGGTCCGCTGGGTCCGCTGCCCCGCTCCGGC | 12 |
| AG876-142800-142920 | 76880425 | GCTGCAGCCGGGTCCGGGGTTCCGGCCCTGGAGCTCGGGGGGCGGCCGGGTGGCCCACCGGGTCCGCTGGGTCCGCTGCCCCGCTCCGGCGGGGGGGTGGCCGGCTGCAGCCGGGTCCGG | 12 |
| AG876-142830-142950 | 77053190 | GAGCTCGGGGGGCGGCCGGGTGGCCCACCGGGTCCGCTGGGTCCGCTGCCCCGCTCCGGCGGGGGGGTGGCCGGCTGCAGCCGGGTCCGGGGTTCCGGCCCTGGAGCTCGGGGGGCGGCC | 12 |
| AG876-142860-142980 | 77225955 | GGTCCGCTGGGTCCGCTGCCCCGCTCCGGCGGGGGGGTGGCCGGCTGCAGCCGGGTCCGGGGTTCCGGCCCTGGAGCTCGGGGGGCGGCCGGGTGGCCCACCGGGTCCGCTGGGTCCGCT | 12 |
| AG876-142890-143010 | 77398720 | GGGGGGGTGGCCGGCTGCAGCCGGGTCCGGGGTTCCGGCCCTGGAGCTCGGGGGGCGGCCGGGTGGCCCACCGGGTCCGCTGGGTCCGCTGCCCCGCTCCGGCGGGGGGGTGGCCGGCTG | 12 |

| | | | |
|---|---|---|---|
| AG876-142920-143040 | 77571485 | GGTTCCGGCCCTGGAGCTCGGGGGGCGGCCGGGTGGCCCACCGGGTCCGCTGGGTCCGCTGCCCCGCTCCGGCGGGGGGGTGGCCGGCTGCAGCCGGGTCCGGGGTTCCGGCCCTGGAGC | 12 |
| AG876-142950-143070 | 77744250 | GGGTGGCCCACCGGGTCCGCTGGGTCCGCTGCCCCGCTCCGGCGGGGGGGTGGCCGGCTGCAGCCGGGTCCGGGGTTCCGGCCCTGGAGCTCGGGGGGCGGCCGGGTGGCCCACCGGGTC | 12 |
| AG876-142980-143100 | 77917015 | GCCCCGCTCCGGCGGGGGGGTGGCCGGCTGCAGCCGGGTCCGGGGTTCCGGCCCTGGAGCTCGGGGGGCGGCCGGGTGGCCCACCGGGTCCGCTGGGTCCGCTGCCCCGCTCCGGCGGGG | 16 |
| AG876-143010-143130 | 78089780 | CAGCCGGGTCCGGGGTTCCGGCCCTGGAGCTCGGGGGGCGGCCGGGTGGCCCACCGGGTCCGCTGGGTCCGCTGCCCCGCTCCGGCGGGGGGGTGGCCGGCTGCAGCCGGGTCCGGGGTT | 12 |
| AG876-143040-143160 | 78262545 | TCGGGGGGCGGCCGGGTGGCCCACCGGGTCCGCTGGGTCCGCTGCCCCGCTCCGGCGGGGGGGTGGCCGGCTGCAGCCGGGTCCGGGGTTCCGGCCCTGGAGCTCGGGGGGCGGCCGGGT | 12 |
| AG876-143070-143190 | 78435310 | CGCTGGGTCCGCTGCCCCGCTCCGGCGGGGGGGTGGCCGGCTGCAGCCGGGTCCGGGGTTCCGGCCCTGGAGCTCGGGGGGCGGCCGGGTGGCCCACCGGGTCCGCTGGGTCCGCTGCCC | 12 |
| AG876-143100-143220 | 78608075 | GGGTGGCCGGCTGCAGCCGGGTCCGGGGTTCCGGCCCTGGAGCTCGGGGGGCGGCCGGGTGGCCCACCGGGTCCGCTGGGTCCGCTGCCCCGCTCCGGCGGGGGGGTGGCCGGCTGCAGC | 12 |
| AG876-143130-143250 | 78780840 | CCGGCCCTGGAGCTCGGGGGGCGGCCGGGTGGCCCACCGGGTCCGCTGGGTCCGCTGCCCCGCTCCGGCGGGGGGGTGGCCGGCTGCAGCCGGGTCCGGGGTTCCGGCCCTGGAGCTCGG | 12 |
| AG876-143160-143280 | 78953605 | GGCCCACCGGGTCCGCTGGGTCCGCTGCCCCGCTCCGGCGGGGGGGTGGCCGGCTGCAGCCGGGTCCGGGGTTCCGGCCCTGGAGCTCGGGGGGCGGCCGGGTGGCCCACCGGGTCCGCT | 12 |
| AG876-143190-143310 | 79126370 | CGCTCCGGCGGGGGGGTGGCCGGCTGCAGCCGGGTCCGGGGTTCCGGCCCTGGAGCTCGGGGGGCGGCCGGGTGGCCCACCGGGTCCGCTGGGTCCGCTGCCCCGCTCCGGCGGGGGGGT | 12 |
| AG876-143220-143340 | 79299135 | CGGGTCCGGGGTTCCGGCCCTGGAGCTCGGGGGGCGGCCGGGTGGCCCACCGGGTCCGCTGGGTCCGCTGCCCCGCTCCGGCGGGGGGGTGGCCGGCTGCAGCCGGGTCCGGGGTTCCGG | 12 |
| AG876-143250-143370 | 79471900 | GGGGCGGCCGGGTGGCCCACCGGGTCCGCTGGGTCCGCTGCCCCGCTCCGGCGGGGGGGTGGCCGGCTGCAGCCGGGTCCGGGGTTCCGGCCCTGGAGCTCGGGGGGCGGCCGGGTGGCC | 16 |
| AG876-143280-143400 | 79644665 | GGGTCCGCTGCCCCGCTCCGGCGGGGGGGTGGCCGGCTGCAGCCGGGTCCGGGGTTCCGGCCCTGGAGCTCGGGGGGCGGCCGGGTGGCCCACCGGGTCCGCTGGGTCCGCTGCCCCGCT | 12 |
| AG876-143310-143430 | 79817430 | GGCCGGCTGCAGCCGGGTCCGGGGTTCCGGCCCTGGAGCTCGGGGGGCGGCCGGGTGGCCCACCGGGTCCGCTGGGTCCGCTGCCCCGCTCCGGCGGGGGGGTGGCCGGCTGCAGCCGGG | 12 |
| AG876-143340-143460 | 79990195 | CCCTGGAGCTCGGGGGGCGGCCGGGTGGCCCACCGGGTCCGCTGGGTCCGCTGCCCCGCTCCGGCGGGGGGGTGGCCGGCTGCAGCCGGGTCCGGGGTTCCGGCCCTGGAGCTCGGGGGG | 12 |
| AG876-143370-143490 | 80162960 | CACCGGGTCCGCTGGGTCCGCTGCCCCGCTCCGGCGGGGGGGTGGCCGGCTGCAGCCGGGTCCGGGGTTCCGGCCCTGGAGCTCGGGGGGCGGCCGGGTGGCCCACCGGGTCCGCTGGGT | 12 |
| AG876-143400-143520 | 80335725 | CCGGCGGGGGGGTGGCCGGCTGCAGCCGGGTCCGGGGTTCCGGCCCTGGAGCTCGGGGGGCGGCCGGGTGGCCCACCGGGTCCGCTGGGTCCGCTGCCCCGCTCCGGCGGGGGGGTGGCC | 16 |
| AG876-143430-143550 | 80508490 | TCCGGGGTTCCGGCCCTGGAGCTCGGGGGGCGGCCGGGTGGCCCACCGGGTCCGCTGGGTCCGCTGCCCCGCTCCGGCGGGGGGGTGGCCGGCTGCAGCCGGGTCCGGGGTTCCGGCCCT | 12 |
| AG876-143460-143580 | 80681255 | CGGCCGGGTGGCCCACCGGGTCCGCTGGGTCCGCTGCCCCGCTCCGGCGGGGGGGTGGCCGGCTGCAGCCGGGTCCGGGGTTCCGGCCCTGGAGCTCGGGGGGCGGCCGGGTGGCCCACC | 12 |
| AG876-143490-143610 | 80854020 | CCGCTGCCCCGCTCCGGCGGGGGGGTGGCCGGCTGCAGCCGGGTCCGGGGTTCCGGCCCTGGAGCTCGGGGGGCGGCCGGGTGGCCCACCGGGTCCGCTGGGTCCGCTGCCCCGCTCCGG | 12 |
| AG876-143520-143640 | 81026785 | GGCTGCAGCCGGGTCCGGGGTTCCGGCCCTGGAGCTCGGGGGGCGGCCGGGTGGCCCACCGGGTCCGCTGGGTCCGCTGCCCCGCTCCGGCGGGGGGGTGGCCGGCTGCAGCCGGGTCCG | 12 |
| AG876-143550-143670 | 81199550 | GGAGCTCGGGGGGCGGCCGGGTGGCCCACCGGGTCCGCTGGGTCCGCTGCCCCGCTCCGGCGGGGGGGTGGCCGGCTGCAGCCGGGTCCGGGGTTCCGGCCCTGGAGCTCGGGGGGCGGC | 12 |
| AG876-143580-143700 | 81372315 | GGGTCCGCTGGGTCCGCTGCCCCGCTCCGGCGGGGGGGTGGCCGGCTGCAGCCGGGTCCGGGGTTCCGGCCCTGGAGCTCGGGGGGCGGCCGGGTGGCCCACCGGGTCCGCTGGGTCCGC | 12 |
| AG876-143610-143730 | 81545080 | CGGGGGGGTGGCCGGCTGCAGCCGGGTCCGGGGTTCCGGCCCTGGAGCTCGGGGGGCGGCCGGGTGGCCCACCGGGTCCGCTGGGTCCGCTGCCCCGCTCCGGCGGGGGGGTGGCCGGCT | 12 |
| AG876-143640-143760 | 81717845 | GGGTTCCGGCCCTGGAGCTCGGGGGGCGGCCGGGTGGCCCACCGGGTCCGCTGGGTCCGCTGCCCCGCTCCGGCGGGGGGGTGGCCGGCTGCAGCCGGGTCCGGGGTTCCGGCCCTGGAG | 12 |
| AG876-143670-143790 | 81890610 | CGGGTGGCCCACCGGGTCCGCTGGGTCCGCTGCCCCGCTCCGGCGGGGGGGTGGCCGGCTGCAGCCGGGTCCGGGGTTCCGGCCCTGGAGCTCGGGGGGCGGCCGGGTGGCCCACCGGGT | 12 |
| AG876-143700-143820 | 82063375 | TGCCCCGCTCCGGCGGGGGGGTGGCCGGCTGCAGCCGGGTCCGGGGTTCCGGCCCTGGAGCTCGGGGGGCGGCCGGGTGGCCCACCGGGTCCGCTGGGTCCGCTGCCCCGCTCCGGCGGG | 12 |
| AG876-143730-143850 | 82236140 | GCAGCCGGGTCCGGGGTTCCGGCCCTGGAGCTCGGGGGGCGGCCGGGTGGCCCACCGGGTCCGCTGGGTCCGCTGCCCCGCTCCGGCGGGGGGGTGGCCGGCTGCAGCCGGGTCCGGGGT | 12 |
| AG876-143760-143880 | 82408905 | CTCGGGGGGCGGCCGGGTGGCCCACCGGGTCCGCTGGGTCCGCTGCCCCGCTCCGGCGGGGGGGTGGCCGGCTGCAGCCGGGTCCGGGGTTCCGGCCCTGGAGCTCGGGGGGCGGCCGGG | 16 |
| AG876-143790-143910 | 82581670 | CCGCTGGGTCCGCTGCCCCGCTCCGGCGGGGGGGTGGCCGGCTGCAGCCGGGTCCGGGGTTCCGGCCCTGGAGCTCGGGGGGCGGCCGGGTGGCCCACCGGGTCCGCTGGGTCCGCTGCC | 12 |
| AG876-143820-143940 | 82754435 | GGGGTGGCCGGCTGCAGCCGGGTCCGGGGTTCCGGCCCTGGAGCTCGGGGGGCGGCCGGGTGGCCCACCGGGTCCGCTGGGTCCGCTGCCCCGCTCCGGCGGGGGGGTGGCCGGCTGCAG | 12 |
| AG876-143850-143970 | 82927200 | TCCGGCCCTGGAGCTCGGGGGGCGGCCGGGTGGCCCACCGGGTCCGCTGGGTCCGCTGCCCCGCTCCGGCGGGGGGGTGGCCGGCTGCAGCCGGGTCCGGGGTTCCGGCCCTGGAGCTCG | 12 |
| AG876-143880-144000 | 83099965 | TGGCCCACCGGGTCCGCTGGGTCCGCTGCCCCGCTCCGGCGGGGGGGTGGCCGGCTGCAGCCGGGTCCGGGGTTCCGGCCCTGGAGCTCGGGGGGCGGCCGGGTGGCCCACCGGGTCCGC | 12 |
| AG876-143910-144030 | 83272730 | CCGCTCCGGCGGGGGGGTGGCCGGCTGCAGCCGGGTCCGGGGTTCCGGCCCTGGAGCTCGGGGGGCGGCCGGGTGGCCCACCGGGTCCGCTGGGTCCGCTGCCCCGCTCCGGCGGGGGGG | 16 |
| AG876-143940-144060 | 83445495 | CCGGGTCCGGGGTTCCGGCCCTGGAGCTCGGGGGGCGGCCGGGTGGCCCACCGGGTCCGCTGGGTCCGCTGCCCCGCTCCGGCGGGGGGGTGGCCGGCTGCAGCCGGGTCCGGGGTTCCG | 12 |
| AG876-143970-144090 | 83618260 | GGGGGCGGCCGGGTGGCCCACCGGGTCCGCTGGGTCCGCTGCCCCGCTCCGGCGGGGGGGTGGCCGGCTGCAGCCGGGTCCGGGGTTCCGGCCCTGGAGCTCGGGGGGCGGCCGGGTGGC | 16 |
| AG876-144000-144120 | 83791025 | TGGGTCCGCTGCCCCGCTCCGGCGGGGGGGTGGCCGGCTGCAGCCGGGTCCGGGGTTCCGGCCCTGGAGCTCGGGGGGCGGCCGGGTGGCCCACCGGGTCCGCTGGGTCCGCTGCCCCGC | 12 |
| AG876-144030-144150 | 83963790 | TGGCCGGCTGCAGCCGGGTCCGGGGTTCCGGCCCTGGAGCTCGGGGGGCGGCCGGGTGGCCCACCGGGTCCGCTGGGTCCGCTGCCCCGCTCCGGCGGGGGGGTGGCCGGCTGCAGCCGG | 12 |
| AG876-144060-144180 | 84136555 | GCCCTGGAGCTCGGGGGGCGGCCGGGTGGCCCACCGGGTCCGCTGGGTCCGCTGCCCCGCTCCGGCGGGGGGGTGGCCGGCTGCAGCCGGGTCCGGGGTTCCGGCCCTGGAGCTCGGGGG | 12 |
| AG876-144090-144210 | 84309320 | CCACCGGGTCCGCTGGGTCCGCTGCCCCGCTCCGGCGGGGGGGTGGCCGGCTGCAGCCGGGTCCGGGGTTCCGGCCCTGGAGCTCGGGGGGCGGCCGGGTGGCCCACCGGGTCCGCTGGG | 12 |
| AG876-144120-144240 | 84482085 | TCCGGCGGGGGGGTGGCCGGCTGCAGCCGGGTCCGGGGTTCCGGCCCTGGAGCTCGGGGGGCGGCCGGGTGGCCCACCGGGTCCGCTGGGTCCGCTGCCCCGCTCCGGCGGGGGGGTGGC | 12 |
| AG876-144150-144270 | 84654850 | GTCCGGGGTTCCGGCCCTGGAGCTCGGGGGGCGGCCGGGTGGCCCACCGGGTCCGCTGGGTCCGCTGCCCCGCTCCGGCGGGGGGGTGGCCGGCTGCAGCCGGGTCCGGGGTTCCGGCCC | 12 |
| AG876-144180-144300 | 84827615 | GCGGCCGGGTGGCCCACCGGGTCCGCTGGGTCCGCTGCCCCGCTCCGGCGGGGGGGTGGCCGGCTGCAGCCGGGTCCGGGGTTCCGGCCCTGGAGCTCGGGGGGCGGCCGGGTGGCCCAC | 12 |
| AG876-144210-144330 | 85000380 | TCCGCTGCCCCGCTCCGGCGGGGGGGTGGCCGGCTGCAGCCGGGTCCGGGGTTCCGGCCCTGGAGCTCGGGGGGCGGCCGGGTGGCCCACCGGGTCCGCTGGGTCCGCTGCCCCGCTCCG | 12 |
| AG876-144240-144360 | 85173145 | CGGCTGCAGCCGGGTCCGGGGTTCCGGCCCTGGAGCTCGGGGGGCGGCCGGGTGGCCCACCGGGTCCGCTGGGTCCGCTGCCCCGCTCCGGCGGGGGGGTGGCCGGCTGCAGCCGGGTCC | 12 |
| AG876-144270-144390 | 85345910 | TGGAGCTCGGGGGGCGGCCGGGTGGCCCACCGGGTCCGCTGGGTCCGCTGCCCCGCTCCGGCGGGGGGGTGGCCGGCTGCAGCCGGGTCCGGGGTTCCGGCCCTGGAGCTCGGGGGGCGG | 12 |
| AG876-144300-144420 | 85518675 | CGGGTCCGCTGGGTCCGCTGCCCCGCTCCGGCGGGGGGGTGGCCGGCTGCAGCCGGGTCCGGGGTTCCGGCCCTGGAGCTCGGGGGGCGGCCGGGTGGCCCACCGGGTCCGCTGGGTCCG | 12 |
| AG876-144330-144450 | 85691440 | GCGGGGGGGTGGCCGGCTGCAGCCGGGTCCGGGGTTCCGGCCCTGGAGCTCGGGGGGCGGCCGGGTGGCCCACCGGGTCCGCTGGGTCCGCTGCCCCGCTCCGGCGGGGGGGTGGCCGGC | 16 |
| AG876-144360-144480 | 85864205 | GGGGTTCCGGCCCTGGAGCTCGGGGGGCGGCCGGGTGGCCCACCGGGTCCGCTGGGTCCGCTGCCCCGCTCCGGCGGGGGGGTGGCCGGCTGCAGCCGGGTCCGGGGTTCCGGCCCTGGA | 12 |
| AG876-144390-144510 | 86036970 | CCGGGTGGCCCACCGGGTCCGCTGGGTCCGCTGCCCCGCTCCGGCGGGGGGGTGGCCGGCTGCAGCCGGGTCCGGGGTTCCGGCCCTGGAGCTCGGGGGGCGGCCGGGTGGCCCACCGGG | 12 |
| AG876-144420-144540 | 86209735 | CTGCCCCGCTCCGGCGGGGGGGTGGCCGGCTGCAGCCGGGTCCGGGGTTCCGGCCCTGGAGCTCGGGGGGCGGCCGGGTGGCCCACCGGGTCCGCTGGGTCCGCTGCCCCGCTCCGGCGG | 12 |

| | | | |
|---|---|---|---|
| AG876-144450-144570 | 86382500 | TGCAGCCGGGTCCGGGGTTCCGGCCCTGGAGCTCGGGGGGCGGCCGGGTGGCCCACCGGGTCCGCTGGGTCCGCTGCCCCGCTCCGGCGGGGGGGTGGCCGGCTGCAGCCGGGTCCGGGG | 12 |
| AG876-144480-144600 | 86555265 | GCTCGGGGGGCGGCCGGGTGGCCCACCGGGTCCGCTGGGTCCGCTGCCCCGCTCCGGCGGGGGGGTGGCCGGCTGCAGCCGGGTCCGGGGTTCCGGCCCTGGAGCTCGGGGGGCGGCCGG | 16 |
| AG876-144510-144630 | 86728030 | TCCGCTGGGTCCGCTGCCCCGCTCCGGCGGGGGGGTGGCCGGCTGCAGCCGGGTCCGGGGTTCCGGCCCTGGAGCTCGGGGGGCGGCCGGGTGGCCCACCGGGTCCGCTGGGTCCGCTGC | 12 |
| AG876-144540-144660 | 86900795 | GGGGGTGGCCGGCTGCAGCCGGGTCCGGGGTTCCGGCCCTGGAGCTCGGGGGGCGGCCGGGTGGCCCACCGGGTCCGCTGGGTCCGCTGCCCCGCTCCGGCGGGGGGGTGGCCGGCTGCA | 12 |
| AG876-144570-144690 | 87073560 | TTCCGGCCCTGGAGCTCGGGGGGCGGCCGGGTGGCCCACCGGGTCCGCTGGGTCCGCTGCCCCGCTCCGGCGGGGGGGTGGCCGGCTGCAGCCGGGTCCGGGGTTCCGGCCCTGGAGCTC | 12 |
| AG876-144600-144720 | 87246325 | GTGGCCCACCGGGTCCGCTGGGTCCGCTGCCCCGCTCCGGCGGGGGGGTGGCCGGCTGCAGCCGGGTCCGGGGTTCCGGCCCTGGAGCTCGGGGGGGGGGGCGCTCCCAGGCCGGACCCT | 12 |
| AG876-144630-144750 | 87419090 | CCCGCTCCGGCGGGGGGGTGGCCGGCTGCAGCCGGGTCCGGGGTTCCGGCCCTGGAGCTCGGGGGGGGGGGCGCTCCCAGGCCGGACCCTGGTGCCAGGCAGGGACCCCGCGCCACCCGC | 12 |
| AG876-144660-144780 | 87591855 | GCCGGGTCCGGGGTTCCGGCCCTGGAGCTCGGGGGGGGGGGCGCTCCCAGGCCGGACCCTGGTGCCAGGCAGGGACCCCGCGCCACCCGCTTCATGGGGGGGGAGGCCGCCGCAAGGACG | 12 |
| AG876-144690-144810 | 87764620 | GGGGGGGGGGGCGCTCCCAGGCCGGACCCTGGTGCCAGGCAGGGACCCCGCGCCACCCGCTTCATGGGGGGGGAGGCCGCCGCAAGGACGCCGGGCCGGCTGGGAGGTGTGCACCCCCCG | 12 |
| AG876-144720-144840 | 87937385 | GGTGCCAGGCAGGGACCCCGCGCCACCCGCTTCATGGGGGGGGAGGCCGCCGCAAGGACGCCGGGCCGGCTGGGAGGTGTGCACCCCCCGAGCGTCTGGACGACGCTGGCGAGCCGGGCC | 12 |
| AG876-145170-145290 | 88110150 | TGGAGGTATCCTAAGCTCCGCCCCTATATACCAGGTGGGTGGAGCTAGGTAGGATTCAGCTAGGTTCCTACTGGGGTACCCCCCTACCCTACCTTAAGGTGCGCCACCCTTCCTCCTTCC | 4 |
| AG876-147060-147180 | 88282915 | GTATCAGAGTGGGGGATGCTTCTGGTTCCTGGTCTTCTCTGTGCACAAAAAGGGGCCAATGGCCACGGCCCCGCGGCTTTTTGTGCCGGTGCGGAGCCAATTTAGCTTCCCCTCCCCTTA | 4 |
| AG876-147090-147210 | 88455680 | GGTCTTCTCTGTGCACAAAAAGGGGCCAATGGCCACGGCCCCGCGGCTTTTTGTGCCGGTGCGGAGCCAATTTAGCTTCCCCTCCCCTTAGCGGGGGGTCTCGCGGGGTGCCAATTGTCG | 4 |
| AG876-147150-147270 | 88801210 | GCGGAGCCAATTTAGCTTCCCCTCCCCTTAGCGGGGGGTCTCGCGGGGTGCCAATTGTCGCCTGCCTTCCCCTGCTTCCCCTTGTTAACTTATAGCATGATAGGTAGGTCACCTAACGTG | 4 |
| AG876-147180-147300 | 88973975 | GCGGGGGGTCTCGCGGGGTGCCAATTGTCGCCTGCCTTCCCCTGCTTCCCCTTGTTAACTTATAGCATGATAGGTAGGTCACCTAACGTGGAAGCCTGGTGGGTGATCCTTCCTCGGTAG | 4 |
| AG876-169290-169410 | 89146740 | CCACCGTCTGTCATCGAAGGCGGGCCCCGGTCACCTCCTTTGTTTTCAACCTCTTCCGTCAAATTTGGTGGGCCTCCATCATTTCCAGCAGAGTCGCTAGGGTTATGAGGCAGCGGGTCA | 4 |
| AG876-169320-169440 | 89319505 | TCACCTCCTTTGTTTTCAACCTCTTCCGTCAAATTTGGTGGGCCTCCATCATTTCCAGCAGAGTCGCTAGGGTTATGAGGCAGCGGGTCATGTGGGCCATTGTCAGCAGTGTTGTCAGGG | 4 |
| AG876-171240-171360 | 89492270 | AGGGGAACGGGGGGGTCGGGGGGCGCAGCGGCCGCGCAGCCATGCGTGACCGCGAGGAGGGGACAGGGTCGCAGGGGGCGTGTCTGGTGGGGGCGGGAGCGGGGGGCGGCGCGGGAGCCT | 12 |
| AG876-171270-171390 | 89665035 | GCCGCGCAGCCATGCGTGACCGCGAGGAGGGGACAGGGTCGCAGGGGGCGTGTCTGGTGGGGGCGGGAGCGGGGGGCGGCGCGGGAGCCTGCACGCCGTTGGAGGGTAGAATGACAGGGG | 12 |
| AG876-171630-171750 | 89837800 | GGGGTGGGGCCATGGGCCGCGCATTCCTGGAAAAAGTGGAGGGGGCGTGGCCTGCCCCCCGCGGCCCCCTAGCCCCCCCGCACAGAGCGGCGCAACGGCGGGCGGGCGGCGGGGGGTCGG | 12 |
| AG876-171660-171780 | 90010565 | AAAAAGTGGAGGGGGCGTGGCCTGCCCCCCGCGGCCCCCTAGCCCCCCCGCACAGAGCGGCGCAACGGCGGGCGGGCGGCGGGGGGTCGGGGTCCGCGGGCTCCGGGGGCTGCGGGCGGT | 12 |
| AG876-171690-171810 | 90183330 | GCGGCCCCCTAGCCCCCCCGCACAGAGCGGCGCAACGGCGGGCGGGCGGCGGGGGGTCGGGGTCCGCGGGCTCCGGGGGCTGCGGGCGGTGGATGGCGGCTGGCGTTCAGGGGAACGGGG | 12 |
| AG876-171810-171930 | 90356095 | GGGTCGGGGGGCGCAGCGGCCGCGCAGCCATGCGTGACCGCGAGGAGGGGACAGGGTCGCAGGGGGCGTGTCTGGTGGGGGCGGGAGCGGGGGGCGGCGCGGGAGCCTGCACGCCGTTGG | 12 |
| AG876-171840-171960 | 90528860 | TGCGTGACCGCGAGGAGGGGACAGGGTCGCAGGGGGCGTGTCTGGTGGGGGCGGGAGCGGGGGGCGGCGCGGGAGCCTGCACGCCGTTGGAGGGTAGAATGACAGGGGGGCGGGGACAGA | 8 |
| AG876-172170-172290 | 90701625 | GGTGTTGGTGGGGCCATGGGCCGCGCATTCCTGGAAAAAGTGGAGGGGGCGTGGCCTGCCCCCCGCGGCCCCCTAGCCCCCCCGCACAGAGCGGCGCAACGGCGGGCGGGCGGCGGGGGG | 12 |
| AG876-172200-172320 | 90874390 | CTGGAAAAAGTGGAGGGGGCGTGGCCTGCCCCCCGCGGCCCCCTAGCCCCCCCGCACAGAGCGGCGCAACGGCGGGCGGGCGGCGGGGGGTCGGGGTCCGCGGGCTCCGGGGGCTGCGGG | 12 |
| AG876-172230-172350 | 91047155 | CCCCGCGGCCCCCTAGCCCCCCCGCACAGAGCGGCGCAACGGCGGGCGGGCGGCGGGGGGTCGGGGTCCGCGGGCTCCGGGGGCTGCGGGCGGTGGATGGCGGCTGGCGTTCAGGGGAAC | 12 |
| AG876-172380-172500 | 91219920 | GCCATGCGTGACCGCGAGGAGGGGACAGGGTCGCAGGGGGCGTGTCTGGTGGGGGCGGGAGCGGGGGGCGGCGCGGGAGCCTGCACGCCGTTGGAGGGTAGAATGACAGGGGGGCGGGGA | 12 |
| AG876-172620-172740 | 91392685 | GGGTGACCCAGCCAAGCGTGACCAAGGGGCCGCTGGGTGACACAGACAACCCTGACAAAGGCCCCCCAGGAAAGACCCCCGGGGGGCATCGGGGGGTGTTGGTGGGGCCATGGGGGGTCG | 8 |
| AG876-172644-172764 | 91565450 | AGGGGCCGCTGGGTGACACAGACAACCCTGACAAAGGCCCCCCAGGAAAGACCCCCGGGGGGCATCGGGGGGTGTTGGTGGGGCCATGGGGGGTCGGATTTCGCCCTTATTGCCCTGTTT | 8 |
| AG876-172644-172764 | 91738215 | AGGGGCCGCTGGGTGACACAGACAACCCTGACAAAGGCCCCCCAGGAAAGACCCCCGGGGGGCATCGGGGGGTGTTGGTGGGGCCATGGGGGGTCGGATTTCGCCCTTATTGCCCTGTTT | 8 |
| AG876-172644-172764 | 91910980 | AGGGGCCGCTGGGTGACACAGACAACCCTGACAAAGGCCCCCCAGGAAAGACCCCCGGGGGGCATCGGGGGGTGTTGGTGGGGCCATGGGGGGTCGGATTTCGCCCTTATTGCCCTGTTT | 8 |
| AG876-172644-172764 | 92083745 | AGGGGCCGCTGGGTGACACAGACAACCCTGACAAAGGCCCCCCAGGAAAGACCCCCGGGGGGCATCGGGGGGTGTTGGTGGGGCCATGGGGGGTCGGATTTCGCCCTTATTGCCCTGTTT | 8 |
| Mutu-7410-7530 | 171688 | CTGTATATGCATGAGGATAGCATATGCTACCCGGATACAGATTAGGATAGCATATACTACCCAGATATAGATTAGGATAGCATATGCTACCCAGATATAGATTAGGATAGCCTATGCTAC | 4 |
| Mutu-10140-10260 | 343376 | CATGACAATTAAAGCCAGGTGATAATTCCATACCCTGGAAGCAGGAGATGAGTGCATTTCACCCCAACCCCCCCTTTCGACTGTCATTTACAATAAAATGAAACCTTTTATTCTTGATTG | 4 |
| Mutu-10770-10890 | 515064 | CCCCGGCCGTGTCCACTCACATATTCCACTCTTATTTTTAAATTAATGTGTCCCAATTAGAAACCCAAGCGCAGAAATTAGTTGAGAGGCTAGTGTTTTAAACCTGCACCCTAGGCCAGC | 4 |
| Mutu-12900-13020 | 686752 | CCAAAGAATGTCTGAGGGGAGTCACCCTCGGGGCCCAGGCCCCAGAGTCCAGAGGTCAGGGGCACCTCAGGGTGCCTCCCCGGGTCCCAGGCCAGCCGGAGGGACCCCGGCAGCCCGGGC | 8 |
| Mutu-12930-13050 | 858440 | GGGCCCAGGCCCCAGAGTCCAGAGGTCAGGGGCACCTCAGGGTGCCTCCCCGGGTCCCAGGCCAGCCGGAGGGACCCCGGCAGCCCGGGCGGCTCCAGAGGCCGGTTCCTCGCCCCTTCC | 12 |
| Mutu-13200-13320 | 1030128 | GCGGCCCTTCTCTCTGTCCCCCTGCTCCTCTCCAACCTTCGCTCCACCCTAGACCCCAGCTTCTGGCCTCCCCGGGTCCACCAGGCCAGCCGGAGGGACCCCGGCAGCCCGGGCGAGTCG | 8 |
| Mutu-13740-13860 | 1201816 | CCCCTGGCGGCCCAGCCCGACCCCCGGGCGCCCCCAAACTTTGTCCAGATGTCCAGGGGTCCCCGAGGGCGAGGCCCAGCCCCCTCCCGCCCCTGTCCACTGCCCCAGTCCCCCCAGAAG | 12 |
| Mutu-13770-13890 | 1373504 | CCCCCAAACTTTGTCCAGATGTCCAGGGGTCCCCGAGGGCGAGGCCCAGCCCCCTCCCGCCCCTGTCCACTGCCCCAGTCCCCCCAGAAGCCCCCAAAAGTAGAGGCTCAGGCCATGCGC | 8 |
| Mutu-13950-14070 | 1545192 | GGGTGCAGTAACAGGTAATCTCTGGTAGTGATTTGGACCCGAAATCTGACACTTTAGAGCTCTGGAGGACTTTAAAACTCTAAAAATCAAAACTTTAGAGGCGAATGGGCGCCATTTTGT | 4 |
| Mutu-13980-14100 | 1716880 | ATTTGGACCCGAAATCTGACACTTTAGAGCTCTGGAGGACTTTAAAACTCTAAAAATCAAAACTTTAGAGGCGAATGGGCGCCATTTTGTCCCCACGCGCGCATAATGGCGGACCTAGGC | 4 |
| Mutu-14010-14130 | 1888568 | TCTGGAGGACTTTAAAACTCTAAAAATCAAAACTTTAGAGGCGAATGGGCGCCATTTTGTCCCCACGCGCGCATAATGGCGGACCTAGGCCTAAAACCCCCAGGAAGCGGGTCTATGGTT | 4 |
| Mutu-14040-14160 | 2060256 | AACTTTAGAGGCGAATGGGCGCCATTTTGTCCCCACGCGCGCATAATGGCGGACCTAGGCCTAAAACCCCCAGGAAGCGGGTCTATGGTTGGCTGCGCTGCTGCTATCTTTAGAGGGGAA | 4 |
| Mutu-14370-14490 | 2231944 | GGAGTCCACACAAATGTAAGAGGGGGTCTTCTACCTCTCCCTAGCCCTCCGCCCCCTCCAAGGACTCGGGCCCAGTTTCTAACTTTTTCCCCTTCCCTCCCTCGTCTTGCCCTGCGCCCG | 4 |
| Mutu-15810-15930 | 2403632 | GACCGGCCCCGCGCCGGGCGCCTCCTCGGGGCCAGCCGCCGGGGTTGGTTCTGCCCCTCTCTCTGTCCTTCAGAGGAACCCGGGACCTCGGGCACCCCAGAGCCCCCCGGGCCCGCCTCC | 12 |
| Mutu-15840-15960 | 2575320 | GCCAGCCGCCGGGGTTGGTTCTGCCCCTCTCTCTGTCCTTCAGAGGAACCCGGGACCTCGGGCACCCCAGAGCCCCCCGGGCCCGCCTCCAGGCGCCCTCCTGGTCTCCGCTCCCCTCTG | 8 |
| Mutu-15870-15990 | 2747008 | CTCTGTCCTTCAGAGGAACCCGGGACCTCGGGCACCCCAGAGCCCCCCGGGCCCGCCTCCAGGCGCCCTCCTGGTCTCCGCTCCCCTCTGAGCCCCGTTAAACCCAAAGAATGTCTGAGG | 8 |
| Mutu-16170-16290 | 2918696 | CCCTCTCCTCCCCTCCAGGCCCGAGCCTCTCCCTCGCGGAGAGGGGCCTCTTTGGGCCCTCACGTCCAGCCCCACCGAGACCCGAGTGGCCCGGATCCCCCCAGCGGCCCTTCTCTCTGT | 8 |
| Mutu-17790-17910 | 3090384 | TCCTACCAGAGGGGGCCAAGAACCCAGACGAGTCCGTAGAAGGGTCCTCGTCCAGCAAGAAGAGGAGGTGGTAAGCGGTTCACCTTCAGGGGTAAGTAACCTGACCTCTCCAGGGCTCAC | 4 |

| | | | | |
|---|---|---|---|---|
| Mutu-17820-17940 | 3262072 | AGTCCGTAGAAGGGTCCTCGTCCAGCAAGAAGAGGAGGTGGTAAGCGGTTCACCTTCAGGGGTAAGTAACCTGACCTCTCCAGGGCTCACATAAAGGGAGGCTTAGTATACATGCTTCTT | | 4 |
| Mutu-19080-19200 | 3433760 | CCAGGCCCCAGAGTCCAGAGGTCAGGGGCACCTCAGGGTGCCTCCCCGGGTCCCAGGCCAGCCGGAGGGACCCCGGCAGCCCGGGCGGCTCCAGAGGCCGGTTCCTCGCCCCTTCCCCGG | | 12 |
| Mutu-19110-19230 | 3605448 | CCTCAGGGTGCCTCCCCGGGTCCCAGGCCAGCCGGAGGGACCCCGGCAGCCCGGGCGGCTCCAGAGGCCGGTTCCTCGCCCCTTCCCCGGGCTTCAGAGCCCAGGATGTCCCCCAGAAGG | | 12 |
| Mutu-19140-19260 | 3777136 | GCCGGAGGGACCCCGGCAGCCCGGGCGGCTCCAGAGGCCGGTTCCTCGCCCCTTCCCCGGGCTTCAGAGCCCAGGATGTCCCCCAGAAGGGACCCTAGGCGTCCCCTCTCCTCCCCTCCA | | 8 |
| Mutu-19410-19530 | 3948824 | GGCCTCCCCGGGTCCACCAGGCCAGCCGGAGGGACCCCGGCAGCCCGGGCGAGTCGCCTTCCCTCTCCCCTGGCCTCTCCTTCCCGCCTCCCACCCGAGCCCCCTCAGCTTGCCTCCCCA | | 12 |
| Mutu-19440-19560 | 4120512 | GGGACCCCGGCAGCCCGGGCGAGTCGCCTTCCCTCTCCCCTGGCCTCTCCTTCCCGCCTCCCACCCGAGCCCCCTCAGCTTGCCTCCCCACCGGGTCCATCAGGCCGGCCGGAGGGACCC | | 12 |
| Mutu-19470-19590 | 4292200 | CCCTCTCCCCTGGCCTCTCCTTCCCGCCTCCCACCCGAGCCCCCTCAGCTTGCCTCCCCACCGGGTCCATCAGGCCGGCCGGAGGGACCCCGGCGGCCCGGTGTCAGTCCCCCCTGCAGC | | 12 |
| Mutu-19860-19980 | 4463888 | TAGGTGCCTCCAGAGCCCCTTTTGCCCCCCTGGCGGCCCAGCCCGACCCCCGGGCGCCCCCAAACTTTGTCCAGATGTCCAGGGGTCCCCGAGGGCGAGGCCCAGCCCCCTCCCGCCCCT | | 8 |
| Mutu-21480-21600 | 4635576 | AGGCCCGCGGCCATTGGGCCCAGATTGAGAGACCAGTCCAGGGGCCCGAGGTTGGAGCCAGCGGGCACCCGAGGTCCCAGCACCCGGTCCCTCTGGTGGGCAGAGACAGGCAGGGCCCCC | | 8 |
| Mutu-22020-22140 | 4807264 | GTCCTTCAGAGGAACCCGGGACCTCGGGCACCCCAGAGCCCCCCGGGCCCGCCTCCAGGCGCCCTCCTGGTCTCCGCTCCCCTCTGAGCCCCGTTAAACCCAAAGAATGTCTGAGGGGAG | | 8 |
| Mutu-22050-22170 | 4978952 | CCCCAGAGCCCCCCGGGCCCGCCTCCAGGCGCCCTCCTGGTCTCCGCTCCCCTCTGAGCCCCGTTAAACCCAAAGAATGTCTGAGGGGAGTCACCCTCGGGGCCCAGGCCCCAGAGTCCA | | 8 |
| Mutu-22080-22200 | 5150640 | GCCCTCCTGGTCTCCGCTCCCCTCTGAGCCCCGTTAAACCCAAAGAATGTCTGAGGGGAGTCACCCTCGGGGCCCAGGCCCCAGAGTCCAGAGGTCAGGGGCACCTCAGGGTGCCTCCCC | | 8 |
| Mutu-23280-23400 | 5322328 | CCATTTTGTCCCCACGCGCGCATAATGGCGGACCTAGGCCTAAAACCCCCAGGAAGCGGGTCTATGGTTGGCTGCGCTGCTGCTATCTTTAGAGGGGAAAAGAGGAATAAGCCCCCAGAC | | 4 |
| Mutu-23310-23430 | 5494016 | GACCTAGGCCTAAAACCCCCAGGAAGCGGGTCTATGGTTGGCTGCGCTGCTGCTATCTTTAGAGGGGAAAAGAGGAATAAGCCCCCAGACAGGGGAGTGGGCTTGTTTGTGACTTCACCA | | 4 |
| Mutu-24600-24720 | 5665704 | GAGGTTGGAGCCAGCGGGCACCCGAGGTCCCAGCACCCGGTCCCTCTGGTGGGCAGAGACAGGCAGGGCCCCCCGGCAGCTGGCCCCGAGGAGGCGCCCGGAGTGGGGCCGGTCGGCTGG | | 12 |
| Mutu-27030-27150 | 5837392 | ACCCAGACGAGTCCGTAGAAGGGTCCTCGTCCAGCAAGAAGAGGAGGTGGTAAGCGGTTCACCTTCAGGGGTAAGTAACCTGACCTCTCCAGGGCTCACATAAAGGGAGGCTTAGTATAC | | 4 |
| Mutu-27540-27660 | 6009080 | CGGGCCAGGCCACCTTAGACCCGGCCAAGCCCCATCCCTGAAGACCCAGCGGCCATTCTCTCTGGTAACGAGCAGAGAAGAAGTAGAGGCCCGCGGCCATTGGGCCCAGATTGAGAGACC | | 4 |
| Mutu-27810-27930 | 6180768 | CTGGGAGGTCTGGGGTGGCGAGCCTGCTGTCTCAGGAGGGGCCTGGCTCCGCCGGGTGGCCCTGGGGTACGTCTGGGAGGCAGAGGGTCGGCCTAGGCCCGGGGAAGTGGAGGGGGATCG | | 8 |
| Mutu-29820-29940 | 6352456 | TTTCCCCTTCCCTCCCTCGTCTTGCCCTGCGCCCGGGGCCACCTTCATCACCGTCGCTGACTCCGCCATCCAAGCCTAGGGGAGACCGAAGTGAAGTCCCTGGACCAGCCCGGCCCGGGC | | 8 |
| Mutu-29850-29970 | 6524144 | GCCCGGGGCCACCTTCATCACCGTCGCTGACTCCGCCATCCAAGCCTAGGGGAGACCGAAGTGAAGTCCCTGGACCAGCCCGGCCCGGGCCCCCGGTATCGGGCCAGAGGTAAGTGGAC | | 8 |
| Mutu-30720-30840 | 6695832 | AGATTGAGAGACCAGTCCAGGGGCCCGAGGTTGGAGCCAGCGGGCACCCGAGGTCCCAGCACCCGGTCCCTCTGGTGGGCAGAGACAGGCAGGGCCCCCCGGCAGCTGGCCCCGAGGAGG | | 8 |
| Mutu-30900-31020 | 6867520 | GCGAGCCTGCTGTCTCAGGAGGGGCCTGGCTCCGCCGGGTGGCCCTGGGGTACGTCTGGGAGGCAGAGGGTCGGCCTAGGCCCGGGGAAGTGGAGGGGGATCGCCCGGGTCTCTGTTGGC | | 8 |
| Mutu-31230-31350 | 7039208 | CCTCTCTCTGTCCTTCAGAGGAACCCGGGACCTCGGGCACCCCAGAGCCCCCCGGGCCCGCCTCCAGGCGCCCTCCTGGTCTCCGCTCCCCTCTGAGCCCCGTTAAACCCAAAGAATGTC | | 8 |
| Mutu-32250-32370 | 7210896 | GGCCCAGCCCCCTCCCGCCCCTGTCCACTGCCCCAGTCCCCCCAGAAGCCCCCAAAAGTAGAGGCTCAGGCCATGCGCGCCCTGTCACCAGGCCTGCCAAAGAGCCAGATCTAAGGCCAG | | 8 |
| Mutu-32280-32400 | 7382584 | CCCCAGTCCCCCCAGAAGCCCCCAAAAGTAGAGGCTCAGGCCATGCGCGCCCTGTCACCAGGCCTGCCAAAGAGCCAGATCTAAGGCCAGGAGAGGCAGCCCCAAAGCGGGTGCAGTAAC | | 4 |
| Mutu-32310-32430 | 7554272 | GAGGCTCAGGCCATGCGCGCCCTGTCACCAGGCCTGCCAAAGAGCCAGATCTAAGGCCAGGAGAGGCAGCCCCAAAGCGGGTGCAGTAACAGGTAATCTCTGGTAGTGATTTGGACCCGA | | 4 |
| Mutu-32340-32460 | 7725960 | GGCCTGCCAAAGAGCCAGATCTAAGGCCAGGAGAGGCAGCCCCAAAGCGGGTGCAGTAACAGGTAATCTCTGGTAGTGATTTGGACCCGAAATCTGACACTTTAGAGCTCTGGAGGACTT | | 4 |
| Mutu-32640-32760 | 7897648 | ACTTCACCAAAGGTCAGGGCCCAAGGGGGTTCGCGTTGCTAGGCCACCTTCTCAGTCCAGCGCGTTTACGTAAGCCAGACAGCAGCCAATTGTCAGTTCTAGGGAGGGGGACCACTGCCC | | 4 |
| Mutu-32670-32790 | 8069336 | TCGCGTTGCTAGGCCACCTTCTCAGTCCAGCGCGTTTACGTAAGCCAGACAGCAGCCAATTGTCAGTTCTAGGGAGGGGGACCACTGCCCCTGGTATAAAGTGGTCCTGCAGCTATTTCT | | 4 |
| Mutu-34110-34230 | 8241024 | CTGAGACCCTCCGGGCCCGGACGGCCTCCCTCAGCCCCCCAGACAGACCCCAGGGTCTCCAGGCAGGGTCCGGCATCTCCAGGGGCAGCAGGCTCACCACCACAGGCCCACCAGACCCGG | | 8 |
| Mutu-34140-34260 | 8412712 | TCAGCCCCCCAGACAGACCCCAGGGTCTCCAGGCAGGGTCCGGCATCTCCAGGGGCAGCAGGCTCACCACCACAGGCCCACCAGACCCGGGTCTCGGCCAGCCGAGCCGACCGGCCCCGC | | 8 |
| Mutu-34170-34290 | 8584400 | AGGCAGGGTCCGGCATCTCCAGGGGCAGCAGGCTCACCACCACAGGCCCACCAGACCCGGGTCTCGGCCAGCCGAGCCGACCGGCCCCGCGCCGGGCGCCTCCTCGGGGCCAGCCGCCGG | | 12 |
| Mutu-34200-34320 | 8756088 | GGCTCACCACCACAGGCCCACCAGACCCGGGTCTCGGCCAGCCGAGCCGACCGGCCCCGCGCCGGGCGCCTCCTCGGGGCCAGCCGCCGGGGTTGGTTCTGCCCCTCTCTCTGTCCTTCA | | 12 |
| Mutu-36390-36510 | 8927776 | CCCCACCACCTCCACCACCTCCACCACCTCCACCACCTCCACCACCTTCACCACCTTCACCACCTTCACCACCCCCCCGCCCCCACCACCCCCACCACCTCAGCGCAGGGATGCCTGGA | | 8 |
| Mutu-36420-36540 | 9099464 | CACCACCTCCACCACCTTCACCACCTTCACCACCTTCACCACCCCCCCGCCCCCACCACCCCCACCACCTCAGCGCAGGGATGCCTGGACACAAGAGCCATCACCTCTTGATAGGGATC | | 4 |
| Mutu-36450-36570 | 9271152 | CACCTTCACCACCCCCCCGCCCCCACCACCCCCACCACCTCAGCGCAGGGATGCCTGGACACAAGAGCCATCACCTCTTGATAGGGATCCGCTAGGATATGACGTCGGGCATGGACCTC | | 4 |
| Mutu-36480-36600 | 9442840 | CCCCACCACCTCAGCGCAGGGATGCCTGGACACAAGAGCCATCACCTCTTGATAGGGATCCGCTAGGATATGACGTCGGGCATGGACCTCTAGCATCTGCTATGCGAATGCTTTGGATGG | | 4 |
| Mutu-36510-36630 | 9614528 | CACAAGAGCCATCACCTCTTGATAGGGATCCGCTAGGATATGACGTCGGGCATGGACCTCTAGCATCTGCTATGCGAATGCTTTGGATGGCTAATTATATTGTAAGACAATCACGGGGTG | | 4 |
| Mutu-37260-37380 | 9786216 | GCAAGGGCAAGTCCAGGGACAAGCAACGCAAGCCCGGTGGACCATGGAGACCAGAGCCAAACACCTCCAGTCCTAGCATGCCTGAACTAAGTCCAGTCCTCGGTCTTCATCAGGGACAAG | | 4 |
| Mutu-39090-39210 | 9957904 | GCCGCTGCCCCGCTCCGGGTGGGGGGTGGCCCCGCTGGGCACCGCTGCGCCGCCGCCAGGTCCTGGGGCAGCCGGGGTTCCTGGCGCTCCGGGGGCAGCCGGGCGGCCGCCGGTGGGTCC | | 12 |
| Mutu-40770-40890 | 10129592 | TAGTGGGTGGAGCCTAGAGGTAGGTATCCATAGGGTTCCATTATCCTGGAGGTATCCTAAGCCTCCGCCCCTATATACCAGGTGGGTGGAGCTAGGTAGGATTCAGCTAGGTTCCTACTG | | 4 |
| Mutu-40800-40920 | 10301280 | TAGGGTTCCATTATCCTGGAGGTATCCTAAGCCTCCGCCCCTATATACCAGGTGGGTGGAGCTAGGTAGGATTCAGCTAGGTTCCTACTGGGGTACCCCCCTACCCTACCTTAAGGTGCG | | 4 |
| Mutu-40830-40950 | 10472968 | GCCTCCGCCCCTATATACCAGGTGGGTGGAGCTAGGTAGGATTCAGCTAGGTTCCTACTGGGGTACCCCCCTACCCTACCTTAAGGTGCGCCACCCTTCCTCCTTCCGTTTTAATGGTAG | | 4 |
| Mutu-40860-40980 | 10644656 | GCTAGGTAGGATTCAGCTAGGTTCCTACTGGGGTACCCCCCTACCCTACCTTAAGGTGCGCCACCCTTCCTCCTTCCGTTTTAATGGTAGAATAACCTATAGGTTATTAACCTAGTGGTG | | 4 |
| Mutu-40890-41010 | 10816344 | GGGTACCCCCCTACCCTACCTTAAGGTGCGCCACCCTTCCTCCTTCCGTTTTAATGGTAGAATAACCTATAGGTTATTAACCTAGTGGTGGAATAGGGTATTGCAGCTGGGTATATACCT | | 4 |
| Mutu-41280-41400 | 10988032 | CCACTAAGCCCCCGTTGCTCGCCTTGCCTGCCTCACCATGACACACTAAGCCCCTGCTAATCCATGAGCCCCGCCTTTAGGAAGCACCACGTCCCGGGGACGGAAGGGGACTTGGGGTGA | | 4 |
| Mutu-41310-41430 | 11159720 | CCTCACCATGACACACTAAGCCCCTGCTAATCCATGAGCCCCGCCTTTAGGAAGCACCACGTCCCGGGGACGGAAGGGGACTTGGGGTGATTTTCTATGTGGGGGTGGAAATATGAGCAA | | 4 |
| Mutu-59490-59610 | 11331408 | CCCCTCCTCCCGGCACGCCCCTGAGATCTGCCTCCCCTCCCTCTCCCTCGTCCTGGTCGGAAAAGTCTGAGGAGGAAAAGGAGAATGGGGAGGAGTCCAAAACGGCACGCCACCTGCCGT | | 4 |
| Mutu-95040-95160 | 11503096 | ATCCTTGAACTCAGCCCAGTCTGAGGGGACCTCCCCCGCCGTGTCCTTAGTATGTCCCATTGCCTCCCCGGTGGCCTCCAGGGCCAAGCAGCACCGCGGGGGCACTGGATCCTCACAGGC | | 8 |
| Mutu-95070-95190 | 11674784 | CTCCCCCGCCGTGTCCTTAGTATGTCCCATTGCCTCCCCGGTGGCCTCCAGGGCCAAGCAGCACCGCGGGGGCACTGGATCCTCACAGGCCGCACCCAGCTCTTCTTCCGTTGCCCCAGT | | 8 |
| Mutu-95100-95220 | 11846472 | TGCCTCCCCGGTGGCCTCCAGGGCCAAGCAGCACCGCGGGGGCACTGGATCCTCACAGGCCGCACCCAGCTCTTCTTCCGTTGCCCCAGTAGCATCTCCGTCTGGTGACCTTGAAGAGGA | | 8 |

| | | | | |
|---|---|---|---|---|
| Mutu-95130-95250 | 12018160 | GCACCGCGGGGGCACTGGATCCTCACAGGCCGCACCCAGCTCTTCTTCCGTTGCCCCAGTAGCATCTCCGTCTGGTGACCTTGAAGAGGAAGAGGAGGGGTCCCGAGAATCCCCATCCCT | | 4 |
| Mutu-96330-96450 | 12189848 | GAGGGGCAGGAGCAGGAGGAGGGGCAGGAGGGGCAGGAGCAGGAGGAGGGGCAGGAGGGGCAGGAGCAGGAGGGGCAGGAGGGGCAGGAGCAGGAGGGGCAGGAGGGGCAGGAGCAGGAG | | 8 |
| Mutu-140880-141000 | 12361536 | GCTGCCCCGCTCCGGCGGGGGGTGGCCGGCTGCAGCCGGGTCCGGGGTTCCGGCCCTGGAGCTCGGGGGGCGGCCGGGTGGCCCACCGGGTCCGCTGGGTCCGCTGCCCCGCTCCGGCGG | | 12 |
| Mutu-140910-141030 | 12533224 | TGCAGCCGGGTCCGGGGTTCCGGCCCTGGAGCTCGGGGGGCGGCCGGGTGGCCCACCGGGTCCGCTGGGTCCGCTGCCCCGCTCCGGCGGGGGGTGGCCGGCTGCAGCCGGGTCCGGGGT | | 12 |
| Mutu-140940-141060 | 12704912 | GCTCGGGGGGCGGCCGGGTGGCCCACCGGGTCCGCTGGGTCCGCTGCCCCGCTCCGGCGGGGGGTGGCCGGCTGCAGCCGGGTCCGGGGTTCCGGCCCTGGAGCTCGGGGGGCGGCCGGG | | 16 |
| Mutu-140970-141090 | 12876600 | TCCGCTGGGTCCGCTGCCCCGCTCCGGCGGGGGGTGGCCGGCTGCAGCCGGGTCCGGGGTTCCGGCCCTGGAGCTCGGGGGGCGGCCGGGTGGCCCACCGGGTCCGCTGGGTCCGCTGCC | | 12 |
| Mutu-141120-141240 | 13048288 | CGGGTCCGGGGTTCCGGCCCTGGAGCTCGGGGGGCGGCCGGGTGGCCCACCGGGTCCGCTGGGTCCGCTGCCCCGCTCCGGCGGGGGGTGGCCGGCTGCAGCCGGGTCCGGGGTTCCGGC | | 12 |
| Mutu-141150-141270 | 13219976 | GGGGCGGCCGGGTGGCCCACCGGGTCCGCTGGGTCCGCTGCCCCGCTCCGGCGGGGGGTGGCCGGCTGCAGCCGGGTCCGGGGTTCCGGCCCTGGAGCTCGGGGGGCGGCCGGGTGGCCC | | 16 |
| Mutu-141180-141300 | 13391664 | GGGTCCGCTGCCCCGCTCCGGCGGGGGGTGGCCGGCTGCAGCCGGGTCCGGGGTTCCGGCCCTGGAGCTCGGGGGGCGGCCGGGTGGCCCACCGGGTCCGCTGGGTCCGCTGCCCCGCTC | | 12 |
| Mutu-141210-141330 | 13563352 | GCCGGCTGCAGCCGGGTCCGGGGTTCCGGCCCTGGAGCTCGGGGGGCGGCCGGGTGGCCCACCGGGTCCGCTGGGTCCGCTGCCCCGCTCCGGCGGGGGGTGGCCGGCTGCAGCCGGGTC | | 12 |
| Mutu-141240-141360 | 13735040 | CCTGGAGCTCGGGGGGCGGCCGGGTGGCCCACCGGGTCCGCTGGGTCCGCTGCCCCGCTCCGGCGGGGGGTGGCCGGCTGCAGCCGGGTCCGGGGTTCCGGCCCTGGAGCTCGGGGGGCG | | 12 |
| Mutu-141270-141390 | 13906728 | ACCGGGTCCGCTGGGTCCGCTGCCCCGCTCCGGCGGGGGGTGGCCGGCTGCAGCCGGGTCCGGGGTTCCGGCCCTGGAGCTCGGGGGGCGGCCGGGTGGCCCACCGGGTCCGCTGGGTCC | | 12 |
| Mutu-141450-141570 | 14078416 | GCTCGGGGGGCGGCCGGGTGGCCCACCGGGTCCGCTGGGTCCGCTGCCCCGCTCCGGCGGGGGGTGGCCGGCTGCAGCCGGGTCCGGGGTTCCGGCCCTGGAGCTCGGGGGGCGGCCGGG | | 16 |
| Mutu-141540-141660 | 14250104 | TCCGGCCCTGGAGCTCGGGGGGCGGCCGGGTGGCCCACCGGGTCCGCTGGGTCCGCTGCCCCGCTCCGGCGGGGGGTGGCCGGCTGCAGCCGGGTCCGGGGTTCCGGCCCTGGAGCTCGG | | 12 |
| Mutu-141630-141750 | 14421792 | CGGGTCCGGGGTTCCGGCCCTGGAGCTCGGGGGGCGGCCGGGTGGCCCACCGGGTCCGCTGGGTCCGCTGCCCCGCTCCGGCGGGGGGTGGCCGGCTGCAGCCGGGTCCGGGGTTCCGGC | | 12 |
| Mutu-141660-141780 | 14593480 | GGGGCGGCCGGGTGGCCCACCGGGTCCGCTGGGTCCGCTGCCCCGCTCCGGCGGGGGGTGGCCGGCTGCAGCCGGGTCCGGGGTTCCGGCCCTGGAGCTCGGGGGGCGGCCGGGTGGCCC | | 16 |
| Mutu-141810-141930 | 14765168 | CGGCGGGGGGTGGCCGGCTGCAGCCGGGTCCGGGGTTCCGGCCCTGGAGCTCGGGGGGCGGCCGGGTGGCCCACCGGGTCCGCTGGGTCCGCTGCCCCGCTCCGGCGGGGGGTGGCCGGC | | 16 |
| Mutu-141840-141960 | 14936856 | CGGGGTTCCGGCCCTGGAGCTCGGGGGGCGGCCGGGTGGCCCACCGGGTCCGCTGGGTCCGCTGCCCCGCTCCGGCGGGGGGTGGCCGGCTGCAGCCGGGTCCGGGGTTCCGGCCCTGGA | | 12 |
| Mutu-141870-141990 | 15108544 | GCCGGGTGGCCCACCGGGTCCGCTGGGTCCGCTGCCCCGCTCCGGCGGGGGGTGGCCGGCTGCAGCCGGGTCCGGGGTTCCGGCCCTGGAGCTCGGGGGGCGGCCGGGTGGCCCACCGGG | | 12 |
| Mutu-141900-142020 | 15280232 | GCTGCCCCGCTCCGGCGGGGGGTGGCCGGCTGCAGCCGGGTCCGGGGTTCCGGCCCTGGAGCTCGGGGGGCGGCCGGGTGGCCCACCGGGTCCGCTGGGTCCGCTGCCCCGCTCCGGCGG | | 12 |
| Mutu-142230-142350 | 15451920 | GCCGGCTGCAGCCGGGTCCGGGGTTCCGGCCCTGGAGCTCGGGGGGCGGCCGGGTGGCCCACCGGGTCCGCTGGGTCCGCTGCCCCGCTCCGGCGGGGGGTGGCCGGCTGCAGCCGGGTC | | 12 |
| Mutu-142260-142380 | 15623608 | CCTGGAGCTCGGGGGGCGGCCGGGTGGCCCACCGGGTCCGCTGGGTCCGCTGCCCCGCTCCGGCGGGGGGTGGCCGGCTGCAGCCGGGTCCGGGGTTCCGGCCCTGGAGCTCGGGGGGCG | | 12 |
| Mutu-142290-142410 | 15795296 | ACCGGGTCCGCTGGGTCCGCTGCCCCGCTCCGGCGGGGGGTGGCCGGCTGCAGCCGGGTCCGGGGTTCCGGCCCTGGAGCTCGGGGGGCGGCCGGGTGGCCCACCGGGTCCGCTGGGTCC | | 12 |
| Mutu-142320-142440 | 15966984 | CGGCGGGGGGTGGCCGGCTGCAGCCGGGTCCGGGGTTCCGGCCCTGGAGCTCGGGGGGCGGCCGGGTGGCCCACCGGGTCCGCTGGGTCCGCTGCCCCGCTCCGGCGGGGGGTGGCCGGC | | 16 |
| Mutu-142440-142560 | 16138672 | TGCAGCCGGGTCCGGGGTTCCGGCCCTGGAGCTCGGGGGGCGGCCGGGTGGCCCACCGGGTCCGCTGGGTCCGCTGCCCCGCTCCGGCGGGGGGTGGCCGGCTGCAGCCGGGTCCGGGGT | | 12 |
| Mutu-142470-142590 | 16310360 | GCTCGGGGGGCGGCCGGGTGGCCCACCGGGTCCGCTGGGTCCGCTGCCCCGCTCCGGCGGGGGGTGGCCGGCTGCAGCCGGGTCCGGGGTTCCGGCCCTGGAGCTCGGGGGGCGGCCGGG | | 16 |
| Mutu-142770-142890 | 16482048 | CCTGGAGCTCGGGGGGCGGCCGGGTGGCCCACCGGGTCCGCTGGGTCCGCTGCCCCGCTCCGGCGGGGGGTGGCCGGCTGCAGCCGGGTCCGGGGTTCCGGCCCTGGAGCTCGGGGGGCG | | 12 |
| Mutu-142830-142950 | 16653736 | CGGCGGGGGGTGGCCGGCTGCAGCCGGGTCCGGGGTTCCGGCCCTGGAGCTCGGGGGGCGGCCGGGTGGCCCACCGGGTCCGCTGGGTCCGCTGCCCCGCTCCGGCGGGGGGTGGCCGGC | | 16 |
| Mutu-142860-142980 | 16825424 | CGGGGTTCCGGCCCTGGAGCTCGGGGGGCGGCCGGGTGGCCCACCGGGTCCGCTGGGTCCGCTGCCCCGCTCCGGCGGGGGGTGGCCGGCTGCAGCCGGGTCCGGGGTTCCGGCCCTGGA | | 12 |
| Mutu-169530-169650 | 16997112 | GGGAGGGGGCGTGGCCTTCCCCCGCGGCCCCCCCGCCCCCCCGCACAGAGCGGCGCAACGGCGGGCGGGCGGCGGGGGGTCGGGGTCCGCGGGCTCCGGGGGCTGCGGGCGGTGGATGGC | | 16 |
| Mutu-169770-169890 | 17168800 | CGCGGGAGCCTGCACGCCGTTGGAGGGTAGAATGACAGGGGGGCGGGGATAGAGAGGCGGTCGCGCCCCCGGCCGCGGCAGCCAAGCCAAGCCCCCGAGGGCAACGGAGCGTGACGAAGG | | 8 |
| Mutu-169800-169920 | 17340488 | AATGACAGGGGGGCGGGGATAGAGAGGCGGTCGCGCCCCCGGCCGCGGCAGCCAAGCCAAGCCCCCGAGGGCAACGGAGCGTGACGAAGGGCCCCAGGGCTGACCCCGGCAAACGTGACC | | 8 |
| Mutu-169830-169950 | 17512176 | TCGCGCCCCCGGCCGCGGCAGCCAAGCCAAGCCCCCGAGGGCAACGGAGCGTGACGAAGGGCCCCAGGGCTGACCCCGGCAAACGTGACCCGGGGCTCCGGGGAGACCCAGCCAAGCGTG | | 12 |
| Mutu-169860-169980 | 17683864 | GCCCCCGAGGGCAACGGAGCGTGACGAAGGGCCCCAGGGCTGACCCCGGCAAACGTGACCCGGGGCTCCGGGGAGACCCAGCCAAGCGTGACCAAGGGGCCCCTGGGTGACACAGGCAAC | | 8 |
| Mutu-169890-170010 | 17855552 | GCCCCAGGGCTGACCCCGGCAAACGTGACCCGGGGCTCCGGGGAGACCCAGCCAAGCGTGACCAAGGGGCCCCTGGGTGACACAGGCAACCCTGACAAAGGCCCCCCAGGAAAGACCCCC | | 8 |
| Mutu-169920-170040 | 18027240 | CGGGGCTCCGGGGAGACCCAGCCAAGCGTGACCAAGGGGCCCCTGGGTGACACAGGCAACCCTGACAAAGGCCCCCCAGGAAAGACCCCCGGGGGGCATCGGGGGGGGGGTCATGGGGGG | | 8 |
| Mutu-170250-170370 | 18198928 | GGGGGCAGGGTCGCGGGGGGGCGTGTCTGGTGGGGGCGGGAGCGGGGGCGGCGCGGGAGCCTGCACGCCGTTGGAGGGTAGAATGACAGGGGGGCGGGGATAGAGAGGCGGTCGCGCCCC | | 12 |
| Mutu-170310-170430 | 18370616 | CTGCACGCCGTTGGAGGGTAGAATGACAGGGGGGCGGGGATAGAGAGGCGGTCGCGCCCCCGGCCGCGGCAGCCAAGCCAAGCCCCCGAGGGCAACGGAGCGTGACGAAGGGCCCCAGGG | | 8 |
| Mutu-170340-170460 | 18542304 | GGGGCGGGGATAGAGAGGCGGTCGCGCCCCCGGCCGCGGCAGCCAAGCCAAGCCCCCGAGGGCAACGGAGCGTGACGAAGGGCCCCAGGGCTGACCCCGGCAAACGTGACCCGGGGCTCC | | 12 |
| Mutu-170370-170490 | 18713992 | CGGCCGCGGCAGCCAAGCCAAGCCCCCGAGGGCAACGGAGCGTGACGAAGGGCCCCAGGGCTGACCCCGGCAAACGTGACCCGGGGCTCCGGGGAGACCCAGCCAAGCGTGACCAAGGGG | | 8 |
| Mutu-170400-170520 | 18885680 | GGCAACGGAGCGTGACGAAGGGCCCCAGGGCTGACCCCGGCAAACGTGACCCGGGGCTCCGGGGAGACCCAGCCAAGCGTGACCAAGGGGCCCCTGGGTGACACAGGCAACCCTGACAAA | | 8 |
| Mutu-170910-171030 | 19057368 | CAGCCAAGCCAAGCCCCCGAGGGCAACGGAGCGTGACGAAGGGCCCCAGGGCTGACCCCGGCAAACGTGACCCGGGGCTCCGGGGAGACCCAGCCAAGCGTGACCAAGGGGCCCCTGGGT | | 8 |
| Mutu-170940-171060 | 19229056 | GCGTGACGAAGGGCCCCAGGGCTGACCCCGGCAAACGTGACCCGGGGCTCCGGGGAGACCCAGCCAAGCGTGACCAAGGGGCCCCTGGGTGACACAGGCAACCCTGACAAAGGCCCCCCA | | 8 |
| Mutu-171090-171210 | 19400744 | TGGTGGGAGGAGCATGGGGGGTCATGGGGGGCCGCGCATTCCTGGAAAAAGTGGGAGGGGGCGTGGCCTTCCCCCGCGGCCCCCCCGCCCCCCCGCACAGAGCGGCGCAACGGCGGGCGG | | 12 |
| Mutu-171120-171240 | 19572432 | GCCGCGCATTCCTGGAAAAAGTGGGAGGGGGCGTGGCCTTCCCCCGCGGCCCCCCCGCCCCCCCGCACAGAGCGGCGCAACGGCGGGCGGGCGGCGGGGGGTCGGGGTCCGCGGGCTCCG | | 12 |
| Mutu-171450-171570 | 19744120 | CCGGCCGCGGCAGCCAAGCCAAGCCCCCGAGGGCAACGGAGCGTGACGAAGGGCCCCAGGGCTGACCCCGGCAAACGTGACCCGGGGCTCCGGGGAGACCCAGCCAAGCGTGACCAAGGG | | 8 |
| Mutu-171480-171600 | 19915808 | GGGCAACGGAGCGTGACGAAGGGCCCCAGGGCTGACCCCGGCAAACGTGACCCGGGGCTCCGGGGAGACCCAGCCAAGCGTGACCAAGGGGCCCCTGGGTGACACAGGCAACCCTGACAA | | 8 |
| Mutu-171567-171687 | 20087496 | GGGGCCCCTGGGTGACACAGGCAACCCTGACAAAGGCCCCCCAGGAAAGACCCCCGGGGGGCATCGGGGGGGTGTTGGGAGGAGCATGGGGGGTCGGATTTCGCCCTTATTGCCCTGTTT | | 8 |
| Mutu-171567-171687 | 20259184 | GGGGCCCCTGGGTGACACAGGCAACCCTGACAAAGGCCCCCCAGGAAAGACCCCCGGGGGGCATCGGGGGGGTGTTGGGAGGAGCATGGGGGGTCGGATTTCGCCCTTATTGCCCTGTTT | | 8 |
| Mutu-171567-171687 | 20430872 | GGGGCCCCTGGGTGACACAGGCAACCCTGACAAAGGCCCCCCAGGAAAGACCCCCGGGGGGCATCGGGGGGGTGTTGGGAGGAGCATGGGGGGTCGGATTTCGCCCTTATTGCCCTGTTT | | 8 |
| Mutu-171567-171687 | 20602560 | GGGGCCCCTGGGTGACACAGGCAACCCTGACAAAGGCCCCCCAGGAAAGACCCCCGGGGGGCATCGGGGGGGTGTTGGGAGGAGCATGGGGGGTCGGATTTCGCCCTTATTGCCCTGTTT | | 8 |

| | | | |
|---|---|---|---|
| Akata-7530-7650 | 171324 | CTACCCAAAGAGAGATTAGGGTAGTATATGCTATCCGAATCTCTATCTGGATAGCATATGCTACCCAATCTATATGCGGGTAGTACATGCTATCCTAAATCTATATTCGGGTAGTATATG | 4 |
| Akata-7560-7680 | 342648 | CTATCCGAATCTCTATCTGGATAGCATATGCTACCCAATCTATATGCGGGTAGTACATGCTATCCTAAATCTATATTCGGGTAGTATATGCTATCCTAATCTCTATGTGGGTAGTACATG | 4 |
| Akata-7590-7710 | 513972 | CTACCCAATCTATATGCGGGTAGTACATGCTATCCTAAATCTATATTCGGGTAGTATATGCTATCCTAATCTCTATGTGGGTAGTACATGCTATCCTAATCTCTATGTGGGTAGCATATG | 4 |
| Akata-7620-7740 | 685296 | TATCCTAAATCTATATTCGGGTAGTATATGCTATCCTAATCTCTATGTGGGTAGTACATGCTATCCTAATCTCTATGTGGGTAGCATATGCTATCCTAATCTATATCTGGGTAGCATATG | 4 |
| Akata-7650-7770 | 856620 | CTATCCTAATCTCTATGTGGGTAGTACATGCTATCCTAATCTCTATGTGGGTAGCATATGCTATCCTAATCTATATCTGGGTAGCATATGCTACCCAAATAGAGATTAGGGTAGCATATG | 4 |
| Akata-7680-7800 | 1027944 | CTATCCTAATCTCTATGTGGGTAGCATATGCTATCCTAATCTATATCTGGGTAGCATATGCTACCCAAATAGAGATTAGGGTAGCATATGCTATCCTAATCTCTATCTGGATAGCATATG | 4 |
| Akata-7710-7830 | 1199268 | CTATCCTAATCTATATCTGGGTAGCATATGCTACCCAAATAGAGATTAGGGTAGCATATGCTATCCTAATCTCTATCTGGATAGCATATGCTATCCAATCTATATGCGGGTAGTACATGC | 4 |
| Akata-7740-7860 | 1370592 | CTACCCAAATAGAGATTAGGGTAGCATATGCTATCCTAATCTCTATCTGGATAGCATATGCTATCCAATCTATATGCGGGTAGTACATGCTATCCTAAATCTATATTCGGGTAGTATATG | 4 |
| Akata-7770-7890 | 1541916 | CTATCCTAATCTCTATCTGGATAGCATATGCTATCCAATCTATATGCGGGTAGTACATGCTATCCTAAATCTATATTCGGGTAGTATATGCTATCCTAATCTCTATGTGGGTAGTACATG | 4 |
| Akata-7800-7920 | 1713240 | CTATCCAATCTATATGCGGGTAGTACATGCTATCCTAAATCTATATTCGGGTAGTATATGCTATCCTAATCTCTATGTGGGTAGTACATGCTATCCTAATCTCTATGTGGGTAGCATATG | 4 |
| Akata-7830-7950 | 1884564 | TATCCTAAATCTATATTCGGGTAGTATATGCTATCCTAATCTCTATGTGGGTAGTACATGCTATCCTAATCTCTATGTGGGTAGCATATGCTATCCAACGCAACATTATTCCACCGTGCT | 4 |
| Akata-7860-7980 | 2055888 | CTATCCTAATCTCTATGTGGGTAGTACATGCTATCCTAATCTCTATGTGGGTAGCATATGCTATCCAACGCAACATTATTCCACCGTGCTCTCAGCGACCTCGTGAATATGAGGACCAAC | 4 |
| Akata-7890-8010 | 2227212 | CTATCCTAATCTCTATGTGGGTAGCATATGCTATCCAACGCAACATTATTCCACCGTGCTCTCAGCGACCTCGTGAATATGAGGACCAACAACCCTGTGCTTGGCGCTCAGGCGCAAGTG | 4 |
| Akata-7920-8040 | 2398536 | CTATCCAACGCAACATTATTCCACCGTGCTCTCAGCGACCTCGTGAATATGAGGACCAACAACCCTGTGCTTGGCGCTCAGGCGCAAGTGTGTGTAACTTGTCCTCCAGATCGCAGCAAT | 4 |
| Akata-10050-10170 | 2569860 | ATAGAAGCATACATGACAATTAAAGCCAGGTGATAATTCCATACCCTGGAAGCAGGAGATGGGTGCATTTCACCCCAACCCCCCCTTTCGACTGTCATTTACAATAAAATGAAACCTTTT | 4 |
| Akata-12630-12750 | 2741184 | CCAGACCCGGGTCTCGGCCAGCCGAGCCGACCGGCCCCGCGCCGGGCGCCTCCTCGGGGCCAGCCGCCGGGGTTGGTTCTGCCCCTCTCTCTGTCCTTCAGAGGAACCAGGGACCTCGGG | 12 |
| Akata-13800-13920 | 2912508 | AGGCCATGCGCGCCCTGTCACCAGGCCTGCCAAAGAGCCAGATCTAAGGCCAGGAGAGGCAGCCCCAAAGCGGGTGCAGTAACAGGTAATCTCTGGTAGTGATTTGGACCCGAAATCTGA | 4 |
| Akata-13860-13980 | 3083832 | AGCCCCAAAGCGGGTGCAGTAACAGGTAATCTCTGGTAGTGATTTGGACCCGAAATCTGACACTTTAGAGCTCTGGAGGACTTTAAAACTCTAAAAATCAAAACTTTAGAGGCGAATGGG | 4 |
| Akata-13890-14010 | 3255156 | CTCTGGTAGTGATTTGGACCCGAAATCTGACACTTTAGAGCTCTGGAGGACTTTAAAACTCTAAAAATCAAAACTTTAGAGGCGAATGGGCGCCATTTTGTCCCCACGCGCGCATAATGG | 4 |
| Akata-13920-14040 | 3426480 | CACTTTAGAGCTCTGGAGGACTTTAAAACTCTAAAAATCAAAACTTTAGAGGCGAATGGGCGCCATTTTGTCCCCACGCGCGCATAATGGCGGACCTAGGCCTAAAACCCCCAGGAAGCG | 4 |
| Akata-13950-14070 | 3597804 | CTAAAAATCAAAACTTTAGAGGCGAATGGGCGCCATTTTGTCCCCACGCGCGCATAATGGCGGACCTAGGCCTAAAACCCCCAGGAAGCGGGTCTATGGTTGGCTGCGCTGCTGCTATCT | 4 |
| Akata-15990-16110 | 3769128 | CGGAGGGACCCCGGCAGCCCGGGCGGCCCCAGAGGCCGGTTCCTCGCCCCTTCCCCGGGCTTCAGAGCCCAGGATGTCCCCCAGAAGGGACCCTAGGCGTCCCCTCTCCTCCCCTCCAGG | 8 |
| Akata-16320-16440 | 3940452 | CTCTCCCCTGGCCTCTCCTTCCCGCCTCCCACCCGAGCCCCCTCAGCTTGCCTCCCCACCGGGTCCATCAGGCCGGCCGGAGGGACCCCGGCGGCCCGGTGTCAGTCCCCCCTGCAGCCG | 12 |
| Akata-16680-16800 | 4111776 | ATAGACTCCCATGTAAGCCTGCCTCGAGTAGGTGCCTCCAGAGCCCCTTTTGCCCCCCTGGCGGCCCAGCCCGACCCCCGGGCGCCCCCAAACTTTGTCCAGATGTCCAGGGGTCCCCGA | 8 |
| Akata-16740-16860 | 4283100 | GCGGCCCAGCCCGACCCCCGGGCGCCCCCAAACTTTGTCCAGATGTCCAGGGGTCCCCGAGGGCGAGGCCCAGCCCCCTCCCACCCCTGTCCACTGCCCCGGTCCCCCCAGAAGCCCCCA | 12 |
| Akata-17700-17820 | 4454424 | CGGTCGCCCAGTCCTACCAGAGGGGGCCAAGAACCCAGACGAGTCCGTAGAAGGGTCCTCGTCCAGCAAGAAGAGGAGGTGGTAAGCGGTTCACCTTCAGGGGTAAGTAACCTGACCTCT | 4 |
| Akata-17730-17850 | 4625748 | GAACCCAGACGAGTCCGTAGAAGGGTCCTCGTCCAGCAAGAAGAGGAGGTGGTAAGCGGTTCACCTTCAGGGGTAAGTAACCTGACCTCTCCAGGGCTCACATAAAGGGAGGCTTAGTAT | 4 |
| Akata-18300-18420 | 4797072 | TCTCTGGTAACGAGCAGAGAAGAAGTAGAGGCCCGCGGCCATTGGGCCCAGATTGAGAGACCAGTCCAGGGGCCCGAGGTTGGAGCCAGCGGGCACCCGAGGTCCCAGCACCCGGTCCCT | 8 |
| Akata-18330-18450 | 4968396 | GCCCGCGGCCATTGGGCCCAGATTGAGAGACCAGTCCAGGGGCCCGAGGTTGGAGCCAGCGGGCACCCGAGGTCCCAGCACCCGGTCCCTCCGGGGGGCAGAGACAGGCAGGGCCCCCCG | 8 |
| Akata-18360-18480 | 5139720 | CCAGTCCAGGGGCCCGAGGTTGGAGCCAGCGGGCACCCGAGGTCCCAGCACCCGGTCCCTCCGGGGGGCAGAGACAGGCAGGGCCCCCCGGCAGCTGGCCCCGAGGAGGCGCCCGGAGTG | 12 |
| Akata-18960-19080 | 5311044 | GTTAAACCCAAAGAATGTCTGAGGGGAGCCACCCTCGGGGCCCAGGCCCCAGAGTCCAGAGGTCAGGGGCACCTCAGGGTGCCTCCCCGGGTCCCAGGCCAGCCGGAGGGACCCCGGCAG | 8 |
| Akata-18990-19110 | 5482368 | ACCCTCGGGGCCCAGGCCCCAGAGTCCAGAGGTCAGGGGCACCTCAGGGTGCCTCCCCGGGTCCCAGGCCAGCCGGAGGGACCCCGGCAGCCCGGGCGGCCCCAGAGGCCGGTTCCTCGC | 12 |
| Akata-19020-19140 | 5653692 | GGTCAGGGGCACCTCAGGGTGCCTCCCCGGGTCCCAGGCCAGCCGGAGGGACCCCGGCAGCCCGGGCGGCCCCAGAGGCCGGTTCCTCGCCCCTTCCCCGGGCTTCAGAGCCCAGGATGT | 12 |
| Akata-19740-19860 | 5825016 | GGCCTGGCTCCCCATAGACTCCCATGTAAGCCTGCCTCGAGTAGGTGCCTCCAGAGCCCCTTTTGCCCCCCTGGCGGCCCAGCCCGACCCCCGGGCGCCCCCAAACTTTGTCCAGATGTC | 8 |
| Akata-19830-19950 | 5996340 | CCGGGCGCCCCCAAACTTTGTCCAGATGTCCAGGGGTCCCCGAGGGCGAGGCCCAGCCCCCTCCCACCCCTGTCCACTGCCCCGGTCCCCCCAGAAGCCCCCAAAAGTAGAGGCTCAGGC | 8 |
| Akata-19860-19980 | 6167664 | CAGGGGTCCCCGAGGGCGAGGCCCAGCCCCCTCCCACCCCTGTCCACTGCCCCGGTCCCCCCAGAAGCCCCCAAAAGTAGAGGCTCAGGCCATGCGCGCCCTGTCACCAGGCCTGCCAAA | 8 |
| Akata-20520-20640 | 6338988 | TTTTTCCCCTTCCCTCCCTCGTCTTGCCCTGCGCCCGGGGCCACCTTCATCACCGTCGCTGACTCCGCCATCCAAGCCTAGGGGAGACCGAAGTGAAGTCCCTGGACCAGCCCGGCCCGG | 8 |
| Akata-21660-21780 | 6510312 | GGAGGCAGAGGGTCGGCCTAGGCCCGGGGAAGTGGAGGGGGATCGCCCGGGTCTCTGTTGGCAGAGTCCGGGCGATCCTCTGAGACCCTCCGGGCCCGGACGGCCGCCCTCAGCCCCCCA | 8 |
| Akata-23010-23130 | 6681636 | TAGAGGCTCAGGCCATGCGCGCCCTGTCACCAGGCCTGCCAAAGAGCCAGATCTAAGGCCAGGAGAGGCAGCCCCAAAGCGGGTGCAGTAACAGGTAATCTCTGGTAGTGATTTGGACCC | 4 |
| Akata-23040-23160 | 6852960 | CAGGCCTGCCAAAGAGCCAGATCTAAGGCCAGGAGAGGCAGCCCCAAAGCGGGTGCAGTAACAGGTAATCTCTGGTAGTGATTTGGACCCGAAATCTGACACTTTAGAGCTCTGGAGGAC | 4 |
| Akata-23070-23190 | 7024284 | AGGAGAGGCAGCCCCAAAGCGGGTGCAGTAACAGGTAATCTCTGGTAGTGATTTGGACCCGAAATCTGACACTTTAGAGCTCTGGAGGACTTTAAAACTCTAAAAATCAAAACTTTAGAG | 4 |
| Akata-23100-23220 | 7195608 | ACAGGTAATCTCTGGTAGTGATTTGGACCCGAAATCTGACACTTTAGAGCTCTGGAGGACTTTAAAACTCTAAAAATCAAAACTTTAGAGGCGAATGGGCGCCATTTTGTCCCCACGCGC | 4 |
| Akata-23640-23760 | 7366932 | CATCACCGTCGCTGACTCCGCCATCCAAGCCTAGGGGAGACCGAAGTGAAGTCCCTGGACCAGCCCGGCCCGGGCCCCCCGGTATCGGGCCAGAGGTAAGTGGACTTTAATTTTTTCTGC | 4 |
| Akata-23670-23790 | 7538256 | CTAGGGGAGACCGAAGTGAAGTCCCTGGACCAGCCCGGCCCGGGCCCCCCGGTATCGGGCCAGAGGTAAGTGGACTTTAATTTTTTCTGCTAAGCCCAACACTCCACCACACCCAGGCAC | 4 |
| Akata-25980-26100 | 7709580 | GCGCCCCCAAACTTTGTCCAGATGTCCAGGGGTCCCCGAGGGCGAGGCCCAGCCCCCTCCCACCCCTGTCCACTGCCCCGGTCCCCCCAGAAGCCCCCAAAAGTAGAGGCTCAGGCCATG | 8 |
| Akata-27930-28050 | 7880904 | GACCCCAGGGTCTCCAGGCAGGGTCCGGCATCTCCAGGGGCAGCAGGCTCACCACCACAGGCCCCCCAGACCCGGGTCTCGGCCAGCCGAGCCGACCGGCCCCGCGCCGGGCGCCTCCTC | 12 |
| Akata-29100-29220 | 8052228 | CCCAGCCCCCTCCCACCCCTGTCCACTGCCCCGGTCCCCCCAGAAGCCCCCAAAAGTAGAGGCTCAGGCCATGCGCGCCCTGTCACCAGGCCTGCCAAAGAGCCAGATCTAAGGCCAGGA | 8 |
| Akata-29760-29880 | 8223552 | TCTTGCCCTGCGCCCGGGGCCACCTTCATCACCGTCGCTGACTCCGCCATCCAAGCCTAGGGGAGACCGAAGTGAAGTCCCTGGACCAGCCCGGCCCGGGCCCCCCGGTATCGGGCCAGA | 8 |
| Akata-30540-30660 | 8394876 | AGGCCACCTTAGACCCGGCCAAGCCCCATCCCTGAAGACCCAGCGGCCATTCTCTCTGGTAACGAGCAGAGAAGAAGTAGAGGCCCGCGGCCATTGGGCCCAGATTGAGAGACCAGTCCA | 4 |
| Akata-30570-30690 | 8566200 | CCTGAAGACCCAGCGGCCATTCTCTCTGGTAACGAGCAGAGAAGAAGTAGAGGCCCGCGGCCATTGGGCCCAGATTGAGAGACCAGTCCAGGGGCCCGAGGTTGGAGCCAGCGGGCACCC | 4 |
| Akata-31170-31290 | 8737524 | GGAACCAGGGACCTCGGGCACCCCAGAGCCCCTCGGGCCCGCCTCCAGGCGCCCTCCTGGTCTCCGCTCCCCTCTTAGCCCCGTTAAACCCAAAGAATGTCTGAGGGGAGCCACCCTCGG | 8 |

| | | | |
|---|---|---|---|
| Akata-31200-31320 | 8908848 | CCTCGGGCCCGCCTCCAGGCGCCCTCCTGGTCTCCGCTCCCCTCTTAGCCCCGTTAAACCCAAAGAATGTCTGAGGGGAGCCACCCTCGGGGCCCAGGCCCCAGAGTCCAGAGGTCAGGG | 8 |
| Akata-32580-32700 | 9080172 | CCCAAGGGGGTTCGCGTTGCTAGGCCACCTTCTCAGTCCAGCGCGTTTACGTAAGCCAGACAGCAGCCAATTGTCAGTTCTAGGGAGGGGGACCACTGCCCCTGGTATAAAGTGGTCCTG | 4 |
| Akata-33420-33540 | 9251496 | GCCAGACGCCTGGGCCTCTAAGGCCCTCGGGTCCCCCTGGACCCCGGCCTCAGCAACCCTGCTGCTCCCCTCCTGCCACCCCAGCCTCCCCCCCTCCCCGTCCCCCTTCGCTCCTGATCC | 8 |
| Akata-33450-33570 | 9422820 | GTCCCCCTGGACCCCGGCCTCAGCAACCCTGCTGCTCCCCTCCTGCCACCCCAGCCTCCCCCCCTCCCCGTCCCCCTTCGCTCCTGATCCTCCCCCGGTCCCCAGTAGGGCCGCCTGCCC | 12 |
| Akata-34080-34200 | 9594144 | CCCAGGGTCTCCAGGCAGGGTCCGGCATCTCCAGGGGCAGCAGGCTCACCACCACAGGCCCCCCAGACCCGGGTCTCGGCCAGCCGAGCCGACCGGCCCCGCGCCGGGCGCCTCCTCGGG | 12 |
| Akata-34110-34230 | 9765468 | CCAGGGGCAGCAGGCTCACCACCACAGGCCCCCCAGACCCGGGTCTCGGCCAGCCGAGCCGACCGGCCCCGCGCCGGGCGCCTCCTCGGGGCCAGCCGCCGGGGTTGGTTCTGCCCCTCT | 12 |
| Akata-34140-34260 | 9936792 | CCCCAGACCCGGGTCTCGGCCAGCCGAGCCGACCGGCCCCGCGCCGGGCGCCTCCTCGGGGCCAGCCGCCGGGGTTGGTTCTGCCCCTCTCTCTGTCCTTCAGAGGAACCAGGGACCTCG | 12 |
| Akata-37170-37290 | 10108116 | GCAGGGGCAGGGGCAAGGGCAAGTCCAGGGACAAGCAACGCAAGCCCGGTGGACCATGGAGACCAGAGCCAAACACCTCCAGTCCTAGCATGCCTGAACTAAGTCCAGTCCTCGGTCTTC | 4 |
| Akata-39600-39720 | 10279440 | CCGGGGGCAGCAGGGCGGCCGCCGGTGGGTCCGCTGGGCCGCTGCCCCGCTCCGGGTGGGGGGTGGCCCCGCTGGACACCGCTGCGCCGCCGCCAGGTCCTGGGGCAGCCGGGGTTCCTG | 12 |
| Akata-39630-39750 | 10450764 | CCGCTGGGCCGCTGCCCCGCTCCGGGTGGGGGGTGGCCCCGCTGGACACCGCTGCGCCGCCGCCAGGTCCTGGGGCAGCCGGGGTTCCTGGCGCTCCACTGCACCTGGAATGCAGGGTGG | 12 |
| Akata-39660-39780 | 10622088 | GGGTGGCCCCGCTGGACACCGCTGCGCCGCCGCCAGGTCCTGGGGCAGCCGGGGTTCCTGGCGCTCCACTGCACCTGGAATGCAGGGTGGGGGCGTGGTCCCCTGGACCCCAGCCCCGCC | 12 |
| Akata-39690-39810 | 10793412 | CGCCAGGTCCTGGGGCAGCCGGGGTTCCTGGCGCTCCACTGCACCTGGAATGCAGGGTGGGGGCGTGGTCCCCTGGACCCCAGCCCCGCCGATCCCTCCCCCAGGGCGTACCCGGCTTGC | 8 |
| Akata-40590-40710 | 10964736 | GGGGCTAGAGGTCCGCGAGATTTGGGGTAGGCGGAGCCTCAAGAGGGTCCCCTCCATAGGGTTTAACCATGAGGGGGAGGATTGGGCTCCGCCCTGATATACCTAGTGGGTGGAGCCTAG | 4 |
| Akata-40620-40740 | 11136060 | GCGGAGCCTCAAGAGGGTCCCCTCCATAGGGTTTAACCATGAGGGGGAGGATTGGGCTCCGCCCTGATATACCTAGTGGGTGGAGCCTAGAGGTAGGTATCCATAGGGTTCCATTATCCT | 4 |
| Akata-40650-40770 | 11307384 | GTTTAACCATGAGGGGGAGGATTGGGCTCCGCCCTGATATACCTAGTGGGTGGAGCCTAGAGGTAGGTATCCATAGGGTTCCATTATCCTGGAGGTATCCTAAGCTCCGCCCCTATATAC | 4 |
| Akata-40680-40800 | 11478708 | GCCCTGATATACCTAGTGGGTGGAGCCTAGAGGTAGGTATCCATAGGGTTCCATTATCCTGGAGGTATCCTAAGCTCCGCCCCTATATACCAGGTGGGTGGAGCTAGGTAGGATTCAGCT | 4 |
| Akata-40800-40920 | 11650032 | AGGTTCCTACTGGGGTACCCCCCTACCCTACCTTAAGGTGCGCCACCCTTCCTCCTTCCGTTTTAATGGTAGAATAACCTATAGGTTATTAACCTAGTGGTGGAATAGGGTATTGCAGCT | 4 |
| Akata-40830-40950 | 11821356 | CCTTAAGGTGCGCCACCCTTCCTCCTTCCGTTTTAATGGTAGAATAACCTATAGGTTATTAACCTAGTGGTGGAATAGGGTATTGCAGCTGGGTATATACCTATAGGTATATAGAACCTA | 4 |
| Akata-40860-40980 | 11992680 | TTTTAATGGTAGAATAACCTATAGGTTATTAACCTAGTGGTGGAATAGGGTATTGCAGCTGGGTATATACCTATAGGTATATAGAACCTAGAGGAAGGGAACCCTATAGTGTAATCCCTC | 4 |
| Akata-59310-59430 | 12164004 | GTATGCTCCTTCTCCTCCTCCTCCTCCTCCCGGCACGCCCCTGAGATCTGCCTCCCCTCCCTCTCCCTCGTCCTGGTCGGAAAAGTCTGAGGAGGAAAAGGAGAATGGGGAGGAGTCCAA | 4 |
| Akata-77640-77760 | 12335328 | TGGGATTGAACTGGGTCTCAGTGACATGGAAGAGGTTGAACTTGAAGTTCTGTTATGTTGGCCTGTGGTAACATCACTGGTTGCATTTTTTGGTGGGCTGGTAACTACTGGGGTGGGACT | 4 |
| Akata-77670-77790 | 12506652 | AGAGGTTGAACTTGAAGTTCTGTTATGTTGGCCTGTGGTAACATCACTGGTTGCATTTTTTGGTGGGCTGGTAACTACTGGGGTGGGACTTGTTCCTCCTAAGGTGTGGTTGGTGGCATT | 4 |
| Akata-87960-88080 | 12677976 | GCCCCCCGGGCTCCATACCAGGGATACCAGGAGCCGCCGGCCCCCCGGGCTCCATACCAGGGATACCAGGAGCCGCCGGCCCCCCAGGCTCCATACCAGGGATACCAGGAGCCGCCGGCC | 8 |
| Akata-87990-88110 | 12849300 | GAGCCGCCGGCCCCCCGGGCTCCATACCAGGGATACCAGGAGCCGCCGGCCCCCCAGGCTCCATACCAGGGATACCAGGAGCCGCCGGCCCCCCAGGCTCCATACCAGGGATACCAGGAG | 8 |
| Akata-88020-88140 | 13020624 | GGATACCAGGAGCCGCCGGCCCCCCAGGCTCCATACCAGGGATACCAGGAGCCGCCGGCCCCCCAGGCTCCATACCAGGGATACCAGGAGCCGCCGGCCCCCCAGGCTCCATACCAGGGA | 8 |
| Akata-88050-88170 | 13191948 | CCATACCAGGGATACCAGGAGCCGCCGGCCCCCCAGGCTCCATACCAGGGATACCAGGAGCCGCCGGCCCCCCAGGCTCCATACCAGGGATACCAGGAGCCGCCGGCCCCCCGGGCTCCAA | 8 |
| Akata-96000-96120 | 13363272 | GAGCAGGAGGGGCAGGAGCAGGAGGGGCAGGAGGGGCAGGAGCAGGAGGGGCGGGAGCAGGAGGGGCAGGAGGGGCAGGAGCAGGAGGGGCGGGAGCAGGAGGGGCAGGAGGGGCGGGAG | 12 |
| Akata-96030-96150 | 13534596 | GAGGGGCAGGAGCAGGAGGGGCGGGAGCAGGAGGGGCAGGAGGGGCAGGAGCAGGAGGGGCGGGAGCAGGAGGGGCAGGAGGGGCGGGAGCAGGAGGGGCAGGAGGGGCAGGAGCAGGAG | 12 |
| Akata-96060-96180 | 13705920 | GAGGGGCAGGAGGGGCAGGAGCAGGAGGGGCGGGAGCAGGAGGGGCAGGAGGGGCGGGAGCAGGAGGGGCAGGAGGGGCAGGAGCAGGAGGGGCAGGAGGGGCAGGAGGTGGAGGGGCAG | 12 |
| Akata-96090-96210 | 13877244 | CGGGAGCAGGAGGGGCAGGAGGGGCGGGAGCAGGAGGGGCAGGAGGGGCAGGAGCAGGAGGGGCAGGAGGGGCAGGAGGTGGAGGGGCAGGAGGGGCAGGAGGTGGAGGCCGGGGTCGAG | 12 |
| Akata-96120-96240 | 14048568 | CAGGAGGGGCAGGAGGGGCAGGAGCAGGAGGGGCAGGAGGGGCAGGAGGTGGAGGGGCAGGAGGGGCAGGAGGTGGAGGCCGGGGTCGAGGAGGCAGTGGAGGCCGGGGTCGAGGAGGCA | 8 |
| Akata-96150-96270 | 14219892 | GGGCAGGAGGGGCAGGAGGTGGAGGGGCAGGAGGGGCAGGAGGTGGAGGCCGGGGTCGAGGAGGCAGTGGAGGCCGGGGTCGAGGAGGCAGTGGAGGCCGGGGTCGAGGAGGTAGTGGAG | 8 |
| Akata-125700-125820 | 14391216 | GGCACCAGGAGCTGTCTCCGCTCATGTCCAGAAGCCATGGAGGGCAGACGGATGTCAGGTCGGGCCCGGACCCGGCCCGGAGGCTGGTGGCCCTCCTGCGAAGGGAAGATGGGGCACCTA | 8 |
| Akata-125730-125850 | 14562540 | GAAGCCATGGAGGGCAGACGGATGTCAGGTCGGGCCCGGACCCGGCCCGGAGGCTGGTGGCCCTCCTGCGAAGGGAAGATGGGGCACCTAAGGACCCCCCTCTGGGACCGTTTGGACACC | 8 |
| Akata-140520-140640 | 14733864 | GAGCTCGGGGGGCGGCCGGGTGGCCCACCGGGTCCGCTGGGTCCGCTGCCCCGCTCCGGCGGGGGGTGGCCGGCTGCAGCCGGGTCCGGGGTTCCGGCCCTGGAGCTCGGGGGGCGGCCG | 12 |
| Akata-140550-140670 | 14905188 | GGTCCGCTGGGTCCGCTGCCCCGCTCCGGCGGGGGGTGGCCGGCTGCAGCCGGGTCCGGGGTTCCGGCCCTGGAGCTCGGGGGGCGGCCGGGTGGCCCACCGGGTCCGCTGGGTCCGCTG | 12 |
| Akata-140580-140700 | 15076512 | GGGGGGTGGCCGGCTGCAGCCGGGTCCGGGGTTCCGGCCCTGGAGCTCGGGGGGCGGCCGGGTGGCCCACCGGGTCCGCTGGGTCCGCTGCCCCGCTCCGGCGGGGGGTGGCCGGCTGCA | 12 |
| Akata-140610-140730 | 15247836 | GTTCCGGCCCTGGAGCTCGGGGGGCGGCCGGGTGGCCCACCGGGTCCGCTGGGTCCGCTGCCCCGCTCCGGCGGGGGGTGGCCGGCTGCAGCCGGGTCCGGGGTTCCGGCCCTGGAGCTC | 12 |
| Akata-140640-140760 | 15419160 | GGTGGCCCACCGGGTCCGCTGGGTCCGCTGCCCCGCTCCGGCGGGGGGTGGCCGGCTGCAGCCGGGTCCGGGGTTCCGGCCCTGGAGCTCGGGGGGCGGCCGGGTGGCCCACCGGGTCCG | 12 |
| Akata-140670-140790 | 15590484 | CCCCGCTCCGGCGGGGGGTGGCCGGCTGCAGCCGGGTCCGGGGTTCCGGCCCTGGAGCTCGGGGGGCGGCCGGGTGGCCCACCGGGTCCGCTGGGTCCGCTGCCCCGCTCCGGCGGGGGG | 16 |
| Akata-140700-140820 | 15761808 | GCCGGGTCCGGGGTTCCGGCCCTGGAGCTCGGGGGGCGGCCGGGTGGCCCACCGGGTCCGCTGGGTCCGCTGCCCCGCTCCGGCGGGGGGTGGCCGGCTGCAGCCGGGTCCGGGGTTCCG | 12 |
| Akata-140730-140850 | 15933132 | GGGGGGCGGCCGGGTGGCCCACCGGGTCCGCTGGGTCCGCTGCCCCGCTCCGGCGGGGGGTGGCCGGCTGCAGCCGGGTCCGGGGTTCCGGCCCTGGAGCTCGGGGGGCGGCCGGGTGGC | 16 |
| Akata-140760-140880 | 16104456 | CTGGGTCCGCTGCCCCGCTCCGGCGGGGGGTGGCCGGCTGCAGCCGGGTCCGGGGTTCCGGCCCTGGAGCTCGGGGGGCGGCCGGGTGGCCCACCGGGTCCGCTGGGTCCGCTGCCCCGC | 12 |
| Akata-140790-140910 | 16275780 | TGGCCGGCTGCAGCCGGGTCCGGGGTTCCGGCCCTGGAGCTCGGGGGGCGGCCGGGTGGCCCACCGGGTCCGCTGGGTCCGCTGCCCCGCTCCGGCGGGGGGTGGCCGGCTGCAGCCGGG | 12 |
| Akata-141060-141180 | 16447104 | GGTCCGCTGGGTCCGCTGCCCCGCTCCGGCGGGGGGTGGCCGGCTGCAGCCGGGTCCGGGGTTCCGGCCCTGGAGCTCGGGGGGCGGCCGGGTGGCCCACCGGGTCCGCTGGGTCCGCTG | 12 |
| Akata-141150-141270 | 16618428 | GGTGGCCCACCGGGTCCGCTGGGTCCGCTGCCCCGCTCCGGCGGGGGGTGGCCGGCTGCAGCCGGGTCCGGGGTTCCGGCCCTGGAGCTCGGGGGGCGGCCGGGTGGCCCACCGGGTCCG | 12 |
| Akata-141210-141330 | 16789752 | GCCGGGTCCGGGGTTCCGGCCCTGGAGCTCGGGGGGCGGCCGGGTGGCCCACCGGGTCCGCTGGGTCCGCTGCCCCGCTCCGGCGGGGGGTGGCCGGCTGCAGCCGGGTCCGGGGTTCCG | 12 |
| Akata-141240-141360 | 16961076 | GGGGGGCGGCCGGGTGGCCCACCGGGTCCGCTGGGTCCGCTGCCCCGCTCCGGCGGGGGGTGGCCGGCTGCAGCCGGGTCCGGGGTTCCGGCCCTGGAGCTCGGGGGGCGGCCGGGTGGC | 16 |
| Akata-141270-141390 | 17132400 | CTGGGTCCGCTGCCCCGCTCCGGCGGGGGGTGGCCGGCTGCAGCCGGGTCCGGGGTTCCGGCCCTGGAGCTCGGGGGGCGGCCGGGTGGCCCACCGGGTCCGCTGGGTCCGCTGCCCCGC | 12 |
| Akata-141300-141420 | 17303724 | TGGCCGGCTGCAGCCGGGTCCGGGGTTCCGGCCCTGGAGCTCGGGGGGCGGCCGGGTGGCCCACCGGGTCCGCTGGGTCCGCTGCCCCGCTCCGGCGGGGGGTGGCCGGCTGCAGCCGGG | 12 |
| Akata-141360-141480 | 17475048 | CCACCGGGTCCGCTGGGTCCGCTGCCCCGCTCCGGCGGGGGGTGGCCGGCTGCAGCCGGGTCCGGGGTTCCGGCCCTGGAGCTCGGGGGGCGGCCGGGTGGCCCACCGGGTCCGCTGGGT | 12 |

| | | | |
|---|---|---|---|
| Akata-141390-141510 | 17646372 | TCCGGCGGGGGGTGGCCGGCTGCAGCCGGGTCCGGGGTTCCGGCCCTGGAGCTCGGGGGGCGGCCGGGTGGCCCACCGGGTCCGCTGGGTCCGCTGCCCCGCTCCGGCGGGGGGTGGCCG | 12 |
| Akata-141570-141690 | 17817696 | GGTCCGCTGGGTCCGCTGCCCCGCTCCGGCGGGGGGTGGCCGGCTGCAGCCGGGTCCGGGGTTCCGGCCCTGGAGCTCGGGGGGCGGCCGGGTGGCCCACCGGGTCCGCTGGGTCCGCTG | 12 |
| Akata-141600-141720 | 17989020 | GGGGGGTGGCCGGCTGCAGCCGGGTCCGGGGTTCCGGCCCTGGAGCTCGGGGGGCGGCCGGGTGGCCCACCGGGTCCGCTGGGTCCGCTGCCCCGCTCCGGCGGGGGGTGGCCGGCTGCA | 12 |
| Akata-141630-141750 | 18160344 | GTTCCGGCCCTGGAGCTCGGGGGGCGGCCGGGTGGCCCACCGGGTCCGCTGGGTCCGCTGCCCCGCTCCGGCGGGGGGTGGCCGGCTGCAGCCGGGTCCGGGGTTCCGGCCCTGGAGCTC | 12 |
| Akata-141660-141780 | 18331668 | GGTGGCCCACCGGGTCCGCTGGGTCCGCTGCCCCGCTCCGGCGGGGGGTGGCCGGCTGCAGCCGGGTCCGGGGTTCCGGCCCTGGAGCTCGGGGGGCGGCCGGGTGGCCCACCGGGTCCG | 12 |
| Akata-141690-141810 | 18502992 | CCCCGCTCCGGCGGGGGGTGGCCGGCTGCAGCCGGGTCCGGGGTTCCGGCCCTGGAGCTCGGGGGGCGGCCGGGTGGCCCACCGGGTCCGCTGGGTCCGCTGCCCCGCTCCGGCGGGGGG | 16 |
| Akata-141720-141840 | 18674316 | GCCGGGTCCGGGGTTCCGGCCCTGGAGCTCGGGGGGCGGCCGGGTGGCCCACCGGGTCCGCTGGGTCCGCTGCCCCGCTCCGGCGGGGGGTGGCCGGCTGCAGCCGGGTCCGGGGTTCCG | 12 |
| Akata-141750-141870 | 18845640 | GGGGGGCGGCCGGGTGGCCCACCGGGTCCGCTGGGTCCGCTGCCCCGCTCCGGCGGGGGGTGGCCGGCTGCAGCCGGGTCCGGGGTTCCGGCCCTGGAGCTCGGGGGGCGGCCGGGTGGC | 16 |
| Akata-141780-141900 | 19016964 | CTGGGTCCGCTGCCCCGCTCCGGCGGGGGGTGGCCGGCTGCAGCCGGGTCCGGGGTTCCGGCCCTGGAGCTCGGGGGGCGGCCGGGTGGCCCACCGGGTCCGCTGGGTCCGCTGCCCCGC | 12 |
| Akata-141810-141930 | 19188288 | TGGCCGGCTGCAGCCGGGTCCGGGGTTCCGGCCCTGGAGCTCGGGGGGCGGCCGGGTGGCCCACCGGGTCCGCTGGGTCCGCTGCCCCGCTCCGGCGGGGGGTGGCCGGCTGCAGCCGGG | 12 |
| Akata-141870-141990 | 19359612 | CCACCGGGTCCGCTGGGTCCGCTGCCCCGCTCCGGCGGGGGGTGGCCGGCTGCAGCCGGGTCCGGGGTTCCGGCCCTGGAGCTCGGGGGGCGGCCGGGTGGCCCACCGGGTCCGCTGGGT | 12 |
| Akata-141960-142080 | 19530936 | CGGCCGGGTGGCCCACCGGGTCCGCTGGGTCCGCTGCCCCGCTCCGGCGGGGGGTGGCCGGCTGCAGCCGGGTCCGGGGTTCCGGCCCTGGAGCTCGGGGGGCGGCCGGGTGGCCCACCG | 12 |
| Akata-141990-142110 | 19702260 | CCGCTGCCCCGCTCCGGCGGGGGGTGGCCGGCTGCAGCCGGGTCCGGGGTTCCGGCCCTGGAGCTCGGGGGGCGGCCGGGTGGCCCACCGGGTCCGCTGGGTCCGCTGCCCCGCTCCGGC | 12 |
| Akata-142020-142140 | 19873584 | GCTGCAGCCGGGTCCGGGGTTCCGGCCCTGGAGCTCGGGGGGCGGCCGGGTGGCCCACCGGGTCCGCTGGGTCCGCTGCCCCGCTCCGGCGGGGGGTGGCCGGCTGCAGCCGGGTCCGGG | 12 |
| Akata-142050-142170 | 20044908 | GAGCTCGGGGGGCGGCCGGGTGGCCCACCGGGTCCGCTGGGTCCGCTGCCCCGCTCCGGCGGGGGGTGGCCGGCTGCAGCCGGGTCCGGGGTTCCGGCCCTGGAGCTCGGGGGGCGGCCG | 12 |
| Akata-142200-142320 | 20216232 | CCCCGCTCCGGCGGGGGGTGGCCGGCTGCAGCCGGGTCCGGGGTTCCGGCCCTGGAGCTCGGGGGGCGGCCGGGTGGCCCACCGGGTCCGCTGGGTCCGCTGCCCCGCTCCGGCGGGGGG | 16 |
| Akata-142230-142350 | 20387556 | GCCGGGTCCGGGGTTCCGGCCCTGGAGCTCGGGGGGCGGCCGGGTGGCCCACCGGGTCCGCTGGGTCCGCTGCCCCGCTCCGGCGGGGGGTGGCCGGCTGCAGCCGGGTCCGGGGTTCCG | 12 |
| Akata-142290-142410 | 20558880 | CTGGGTCCGCTGCCCCGCTCCGGCGGGGGGTGGCCGGCTGCAGCCGGGTCCGGGGTTCCGGCCCTGGAGCTCGGGGGGCGGCCGGGTGGCCCACCGGGTCCGCTGGGTCCGCTGCCCCGC | 12 |
| Akata-142320-142440 | 20730204 | TGGCCGGCTGCAGCCGGGTCCGGGGTTCCGGCCCTGGAGCTCGGGGGGCGGCCGGGTGGCCCACCGGGTCCGCTGGGTCCGCTGCCCCGCTCCGGCGGGGGGTGGCCGGCTGCAGCCGGG | 12 |
| Akata-142470-142590 | 20901528 | CGGCCGGGTGGCCCACCGGGTCCGCTGGGTCCGCTGCCCCGCTCCGGCGGGGGGTGGCCGGCTGCAGCCGGGTCCGGGGTTCCGGCCCTGGAGCTCGGGGGGCGGCCGGGTGGCCCACCG | 12 |
| Akata-142500-142620 | 21072852 | CCGCTGCCCCGCTCCGGCGGGGGGTGGCCGGCTGCAGCCGGGTCCGGGGTTCCGGCCCTGGAGCTCGGGGGGCGGCCGGGTGGCCCACCGGGTCCGCTGGGTCCGCTGCCCCGCTCCGGC | 12 |
| Akata-142530-142650 | 21244176 | GCTGCAGCCGGGTCCGGGGTTCCGGCCCTGGAGCTCGGGGGGCGGCCGGGTGGCCCACCGGGTCCGCTGGGTCCGCTGCCCCGCTCCGGCGGGGGGTGGCCGGCTGCAGCCGGGTCCGGG | 12 |
| Akata-142560-142680 | 21415500 | GAGCTCGGGGGGCGGCCGGGTGGCCCACCGGGTCCGCTGGGTCCGCTGCCCCGCTCCGGCGGGGGGTGGCCGGCTGCAGCCGGGTCCGGGGTTCCGGCCCTGGAGCTCGGGGGGCGGCCG | 12 |
| Akata-142590-142710 | 21586824 | GGTCCGCTGGGTCCGCTGCCCCGCTCCGGCGGGGGGTGGCCGGCTGCAGCCGGGTCCGGGGTTCCGGCCCTGGAGCTCGGGGGGCGGCCGGGTGGCCCACCGGGTCCGCTGGGTCCGCTG | 12 |
| Akata-142620-142740 | 21758148 | GGGGGGTGGCCGGCTGCAGCCGGGTCCGGGGTTCCGGCCCTGGAGCTCGGGGGGCGGCCGGGTGGCCCACCGGGTCCGCTGGGTCCGCTGCCCCGCTCCGGCGGGGGGTGGCCGGCTGCA | 12 |
| Akata-142680-142800 | 21929472 | GGTGGCCCACCGGGTCCGCTGGGTCCGCTGCCCCGCTCCGGCGGGGGGTGGCCGGCTGCAGCCGGGTCCGGGGTTCCGGCCCTGGAGCTCGGGGGGCGGCCGGGTGGCCCACCGGGTCCG | 12 |
| Akata-143370-143490 | 22100796 | TTCCTACTGGGGCACCCCCCTACCCTACCTTAAGGTGCGCCACCCTTCCTCCTTCCGTTTTAATGGTAGAATAACCTATAGGTTATTAACCTAGTGGTGGAATAGGGTATTGCAGCTGGG | 4 |
| Akata-143400-143520 | 22272120 | TAAGGTGCGCCACCCTTCCTCCTTCCGTTTTAATGGTAGAATAACCTATAGGTTATTAACCTAGTGGTGGAATAGGGTATTGCAGCTGGGTATATACCTATAGGTATATAGAACCTAGAG | 4 |
| Akata-143430-143550 | 22443444 | TAATGGTAGAATAACCTATAGGTTATTAACCTAGTGGTGGAATAGGGTATTGCAGCTGGGTATATACCTATAGGTATATAGAACCTAGAGGAAGGGAACCCTATAGTGTAATCCCTCCCC | 4 |
| Akata-167310-167430 | 22614768 | CCAAAAGCAGCGTAGGAAGGTGTGGATCACCGGCGCCCCCGTCTGTCATCGAAGGCGGGCCCCGGTCACCTCCTTTGTTTTCAAGCTCATCCGTCAAATTTGGAGGGCCTCCATCATTTC | 4 |
| Akata-167340-167460 | 22786092 | CGGCGCCCCCGTCTGTCATCGAAGGCGGGCCCCGGTCACCTCCTTTGTTTTCAAGCTCATCCGTCAAATTTGGAGGGCCTCCATCATTTCCAGCAGAGTCGCTAGGGTTATGAGGCAGCG | 4 |
| Akata-168000-168120 | 22957416 | AGACTGCGTTGAAAAAAAGAGTGTTAGGGCTGGAAAGGTGAGGGTGTGGTAAATAAGTCCTAGGGAATGTTAGATCTTACCAAGTAAGCAGCCAAAGATGAACAGCACAATTCCAAGGTA | 4 |
| Akata-168300-168420 | 23128740 | GATAAAGATGATGAGAATGATAATAAGAAGCATGAGAGCAAAGGAATAGAGGACAAGGAGCGCTCCTCCAGTCCAGTTACTCATAACGATATACAGCCAGAACAGTAGCGCCAAGAGAAG | 4 |
| Akata-168330-168450 | 23300064 | CATGAGAGCAAAGGAATAGAGGACAAGGAGCGCTCCTCCAGTCCAGTTACTCATAACGATATACAGCCAGAACAGTAGCGCCAAGAGAAGGAGAAGGAGAGCAAGGCCTATGGAAGAGGA | 4 |
| Akata-168360-168480 | 23471388 | CGCTCCTCCAGTCCAGTTACTCATAACGATATACAGCCAGAACAGTAGCGCCAAGAGAAGGAGAAGGAGAGCAAGGCCTATGGAAGAGGAGAGGGGGGGTCCTAGAGGGGGCCGTGGCGG | 4 |
| Akata-168390-168510 | 23642712 | ATACAGCCAGAACAGTAGCGCCAAGAGAAGGAGAAGGAGAGCAAGGCCTATGGAAGAGGAGAGGGGGGGTCCTAGAGGGGGCCGTGGCGGGCCCGGTGGGCCCCTCTCAAGGTCGCGTTC | 4 |
| Akata-168420-168540 | 23814036 | GAGAAGGAGAGCAAGGCCTATGGAAGAGGAGAGGGGGGGTCCTAGAGGGGGCCGTGGCGGGCCCGGTGGGCCCCTCTCAAGGTCGCGTTCCATGGTCAGGGTAGTGTGGCAGGAGCAAGG | 8 |
| Akata-168450-168570 | 23985360 | GAGGGGGGGTCCTAGAGGGGGCCGTGGCGGGCCCGGTGGGCCCCTCTCAAGGTCGCGTTCCATGGTCAGGGTAGTGTGGCAGGAGCAAGGCAACTGAGGAAAGAAGCTGACAGAGCAGTG | 8 |
| Akata-168480-168600 | 24156684 | GCCCGGTGGGCCCCTCTCAAGGTCGCGTTCCATGGTCAGGGTAGTGTGGCAGGAGCAAGGCAACTGAGGAAAGAAGCTGACAGAGCAGTGACAGAGGCTTATGTAGGCCGCCTACGAGAG | 4 |
| Akata-168510-168630 | 24328008 | CATGGTCAGGGTAGTGTGGCAGGAGCAAGGCAACTGAGGAAAGAAGCTGACAGAGCAGTGACAGAGGCTTATGTAGGCCGCCTACGAGAGAGTAACGCGTGTTTCTGGGGATGGAGGCCC | 4 |
| Akata-168540-168660 | 24499332 | CAACTGAGGAAAGAAGCTGACAGAGCAGTGACAGAGGCTTATGTAGGCCGCCTACGAGAGAGTAACGCGTGTTTCTGGGGATGGAGGCCCGGGGGGATTTGCGGGGTCTGCCGGAGGCAG | 4 |
| Akata-168570-168690 | 24670656 | ACAGAGGCTTATGTAGGCCGCCTACGAGAGAGTAACGCGTGTTTCTGGGGATGGAGGCCCGGGGGGATTTGCGGGGTCTGCCGGAGGCAGTACGGGCGCCGCTTTCCCGAAAGCGGCGTA | 4 |
| Akata-168600-168720 | 24841980 | AGTAACGCGTGTTTCTGGGGATGGAGGCCCGGGGGGATTTGCGGGGTCTGCCGGAGGCAGTACGGGCGCCGCTTTCCCGAAAGCGGCGTAGTGTGTGTGCATGTAAGCGTTAAAAGAGGA | 4 |
| Akata-168630-168750 | 25013304 | GGGGGGATTTGCGGGGTCTGCCGGAGGCAGTACGGGCGCCGCTTTCCCGAAAGCGGCGTAGTGTGTGTGCATGTAAGCGTTAAAAGAGGAAGTAGAAAGCGTGTGTCTGTGTTAGAAAAG | 4 |
| Akata-168660-168780 | 25184628 | TACGGGCGCCGCTTTCCCGAAAGCGGCGTAGTGTGTGTGCATGTAAGCGTTAAAAGAGGAAGTAGAAAGCGTGTGTCTGTGTTAGAAAAGCGAGTCCCCGAGGGGCAAGCTGTGGGAATG | 4 |
| Akata-168690-168810 | 25355952 | GTGTGTGTGCATGTAAGCGTTAAAAGAGGAAGTAGAAAGCGTGTGTCTGTGTTAGAAAAGCGAGTCCCCGAGGGGCAAGCTGTGGGAATGCGGTGGTAAAGCGTAGCAGGAAATGGAAAG | 4 |
| Akata-168720-168840 | 25527276 | AGTAGAAAGCGTGTGTCTGTGTTAGAAAAGCGAGTCCCCGAGGGGCAAGCTGTGGGAATGCGGTGGTAAAGCGTAGCAGGAAATGGAAAGGCAGTGCGGCAATCACAAGGGGGAGTGAGC | 4 |
| Akata-168750-168870 | 25698600 | CGAGTCCCCGAGGGGCAAGCTGTGGGAATGCGGTGGTAAAGCGTAGCAGGAAATGGAAAGGCAGTGCGGCAATCACAAGGGGGAGTGAGCAGTGTTGTGGGAAGCGGCAGTGTAATCCAC | 4 |
| Akata-168780-168900 | 25869924 | CGGTGGTAAAGCGTAGCAGGAAATGGAAAGGCAGTGCGGCAATCACAAGGGGGAGTGAGCAGTGTTGTGGGAAGCGGCAGTGTAATCCACACAAAGAGGCGTAGGGCGCGCAACGTTGGG | 4 |
| Akata-168810-168930 | 26041248 | GCAGTGCGGCAATCACAAGGGGGAGTGAGCAGTGTTGTGGGAAGCGGCAGTGTAATCCACACAAAGAGGCGTAGGGCGCGCAACGTTGGGAGGTCGTTGTCGGCAGGCGGGAGGCTGTGC | 4 |
| Akata-168840-168960 | 26212572 | AGTGTTGTGGGAAGCGGCAGTGTAATCCACACAAAGAGGCGTAGGGCGCGCAACGTTGGGAGGTCGTTGTCGGCAGGCGGGAGGCTGTGCTTTAGGGGGGTTCAGGTGAGGCAAGGCTGT | 4 |

| | | | |
|---|---|---|---|
| Akata-168870-168990 | 26383896 | ACAAAGAGGCGTAGGGCGCGCAACGTTGGGAGGTCGTTGTCGGCAGGCGGGAGGCTGTGCTTTAGGGGGGTTCAGGTGAGGCAAGGCTGTGGGGTAACCGTAGGGGAGGCGGGTGAGGCG | 8 |
| Akata-169110-169230 | 26555220 | GCGGAAAGTCGCTGCGGCTTGCTGGCGTGGGGCATGGGGGGCGCATTCCTGGAAAAAGTGGAGGGGGCGTGGCCTGCCCCCCGCGGCCCCCCTAGCCCCCCCGCACAGCGCGGCGCAACG | 8 |
| Akata-169140-169260 | 26726544 | GGCATGGGGGGCGCATTCCTGGAAAAAGTGGAGGGGGCGTGGCCTGCCCCCCGCGGCCCCCCTAGCCCCCCCGCACAGCGCGGCGCAACGGCGGGCGGGCGGGGGGGGTCGGGGTCCGCG | 12 |
| Akata-169170-169290 | 26897868 | GAGGGGGCGTGGCCTGCCCCCCGCGGCCCCCCTAGCCCCCCCGCACAGCGCGGCGCAACGGCGGGCGGGCGGGGGGGGTCGGGGTCCGCGGGCTCCGGGGGCTGCGGGCGGTGGATGGCG | 16 |
| Akata-169320-169440 | 27069192 | GGCGCGGCGCCCGCCCAGCCATGCGTGACCGCGAGGAGGGGACAGGGTCGCAGGGGGCGTGTCTGGTGGGGGCGGGAGCGGGGGGCGGCGCGGGAGCCTGCACGCCGTTGGAGGGGAGGA | 12 |
| Akata-169350-169470 | 27240516 | GCGAGGAGGGGACAGGGTCGCAGGGGGCGTGTCTGGTGGGGGCGGGAGCGGGGGGCGGCGCGGGAGCCTGCACGCCGTTGGAGGGGAGGATGACAGGGGGGCGGGGACAGAGAGGCGGTC | 12 |
| Akata-169380-169500 | 27411840 | GTCTGGTGGGGGCGGGAGCGGGGGGCGGCGCGGGAGCCTGCACGCCGTTGGAGGGGAGGATGACAGGGGGGCGGGGACAGAGAGGCGGTCGCGCCCCCGGCCGCGGCAGGCAAGCCCCCG | 12 |
| Akata-169440-169560 | 27583164 | TGACAGGGGGGCGGGGACAGAGAGGCGGTCGCGCCCCCGGCCGCGGCAGGCAAGCCCCCGGGGGGCGGGGCGCGGGCAGCGGAGCGTGACGAAGGGCCCCAGGGCTGACCCCGGCAAACG | 12 |
| Akata-169830-169950 | 27754488 | GTTCAGGGGAACGGGGGGGTCGGGGGCGCGGCGCCCGCCCAGCCATGCGTGACCGCGAGGAGGGGACAGGGTCGCAGGGGGCGTGTCTGGTGGGGGCGGGAGCGGGGGGCGGCGCGGGAG | 12 |
| Akata-169860-169980 | 27925812 | GCGCCCGCCCAGCCATGCGTGACCGCGAGGAGGGGACAGGGTCGCAGGGGGCGTGTCTGGTGGGGGCGGGAGCGGGGGGCGGCGCGGGAGCCTGCACGCCGTTGGAGGGGAGGATGACAG | 12 |
| Akata-169890-170010 | 28097136 | AGGGGACAGGGTCGCAGGGGGCGTGTCTGGTGGGGGCGGGAGCGGGGGGCGGCGCGGGAGCCTGCACGCCGTTGGAGGGGAGGATGACAGGGGGGCGGGGACAGAGAGGCGGTCGCGCCC | 12 |
| Akata-170220-170340 | 28268460 | GCATTCCTGGAAAAAGTGGAGGGGGCGTGGCCTGCCCCCCGCGGCCCCCCTAGCCCCCCCGCACAGCGCGGCGCAACGGCGGGCGGGCGGGGGGGGTCGGGGTCCGCGGGCTCCGGGGGC | 12 |
| Akata-170370-170490 | 28439784 | GGGAACGGGGGGGTCGGGGGCGCGGCGCCCGCCCAGCCATGCGTGACCGCGAGGAGGGGACAGGGTCGCAGGGGGCGTGTCTGGTGGGGGCGGGAGCGGGGGGCGGCGCGGGAGCCTGCA | 12 |
| Akata-170400-170520 | 28611108 | GCCCAGCCATGCGTGACCGCGAGGAGGGGACAGGGTCGCAGGGGGCGTGTCTGGTGGGGGCGGGAGCGGGGGGCGGCGCGGGAGCCTGCACGCCGTTGGAGGGGAGGATGACAGGGGGGC | 12 |
| Akata-170580-170700 | 28782432 | CGGGCAGCGGAGCGTGACGAAGGGCCCCAGGGCTGACCCCGGCAAACGTGACCCGGGGCTCCGGGGTGACCCAGCCAAGCGTGACCAAGGGGCCGCTGGGTGACACAGGCAACCCTGACA | 8 |
| Akata-170880-171000 | 28953756 | CGGTGGATGGCGGCTGGCGTTCAGGGGAACGGGGGGGTCGGGGGCGCGGCGCCCGCCCAGCCATGCGTGACCGCGAGGAGGGGACAGGGTCGCAGGGGGCGTGTCTGGTGGGGGCGGGAG | 12 |
| Akata-170910-171030 | 29125080 | GGGGGGGTCGGGGGCGCGGCGCCCGCCCAGCCATGCGTGACCGCGAGGAGGGGACAGGGTCGCAGGGGGCGTGTCTGGTGGGGGCGGGAGCGGGGGGCGGCGCGGGAGCCTGCACGCCGT | 12 |
| Akata-170940-171060 | 29296404 | CCATGCGTGACCGCGAGGAGGGGACAGGGTCGCAGGGGGCGTGTCTGGTGGGGGCGGGAGCGGGGGGCGGCGCGGGAGCCTGCACGCCGTTGGAGGGGAGGATGACAGGGGGGCGGGGAC | 12 |
| Akata-171203-171323 | 29467728 | GGGCCGCTGGGTGACACAGGCAACCCTGACAAAGGCCCCCCAGGAAAGACCCCCGGGGGGTATCGGGGGGGTGTTGATGGGGGCATGGGGGGGTCGGATTTCGCCCTTATTGCCCTGTTT | 8 |
| Akata-171203-171323 | 29639052 | GGGCCGCTGGGTGACACAGGCAACCCTGACAAAGGCCCCCCAGGAAAGACCCCCGGGGGGTATCGGGGGGGTGTTGATGGGGGCATGGGGGGGTCGGATTTCGCCCTTATTGCCCTGTTT | 8 |
| Akata-171203-171323 | 29810376 | GGGCCGCTGGGTGACACAGGCAACCCTGACAAAGGCCCCCCAGGAAAGACCCCCGGGGGGTATCGGGGGGGTGTTGATGGGGGCATGGGGGGGTCGGATTTCGCCCTTATTGCCCTGTTT | 8 |
| Akata-171203-171323 | 29981700 | GGGCCGCTGGGTGACACAGGCAACCCTGACAAAGGCCCCCCAGGAAAGACCCCCGGGGGGTATCGGGGGGGTGTTGATGGGGGCATGGGGGGGTCGGATTTCGCCCTTATTGCCCTGTTT | 8 |
| GD1-7980-8100 | 171658 | GACAGAGATTAGGATAGCATATGCTACCCAGATATAGATTAGGATAGCATATGCTACCCACATATTTGGGTAGTATATGCTATCCAAGGCAACATTATTCCACCGTGCTCTCAGCGACCT | 4 |
| GD1-8010-8130 | 343316 | GATATAGATTAGGATAGCATATGCTACCCACATATTTGGGTAGTATATGCTATCCAAGGCAACATTATTCCACCGTGCTCTCAGCGACCTCGTGAATATGAGGACCAACAACCCTGTGCT | 4 |
| GD1-12840-12960 | 514974 | GTTGGTTCTGCCCCTCTCTCTGTCCTTCAGAGGAACCAGGGACCTCGGGCACCCCAGAGCCCCTCGGGCCCGCCTCCAGGCGCCCTCCTGGTCTCCGCTCCCCTCTTAGCCCCGTTAAAC | 8 |
| GD1-12870-12990 | 686632 | AGGAACCAGGGACCTCGGGCACCCCAGAGCCCCTCGGGCCCGCCTCCAGGCGCCCTCCTGGTCTCCGCTCCCCTCTTAGCCCCGTTAAACCCAAAGAATGTCTGAGGGGAGCCACCCTCG | 8 |
| GD1-12900-13020 | 858290 | CCCTCGGGCCCGCCTCCAGGCGCCCTCCTGGTCTCCGCTCCCCTCTTAGCCCCGTTAAACCCAAAGAATGTCTGAGGGGAGCCACCCTCGGGGCCCAGGCCCCAGAGTCCAGAGGTCAGG | 8 |
| GD1-14280-14400 | 1029948 | GCCCAAGGGGGTTCGCGTTGCTAGGCCACCTTCTCAGTCCAGCGCGTTTACGTAAGCCAGACAGCAGCCAATTGTCAGTTCTAGGGAGGGGGACCACTGCCCCTGGTATAAAGTGGTCCT | 4 |
| GD1-15780-15900 | 1373264 | CCCAGGGTCTCCAGGCAGGGTCCGGCATCTCCAGGGGCAGCAGGCTCACCACCACAGGCCCCCCAGACCCGGGTCTCGGCCAGCCGAGCCGACCGGCCCCGCGCCGGGCGCCTCCTCGGG | 12 |
| GD1-17610-17730 | 1544922 | TGCCCTGCGCCCGGGGCCACCTTCATCACCGTCGCTGACTCCGCCATCCAAGCCTAGGGGAGACCGAAGTGAAGTCCCTGGACCAGCCCGGCCCGGGCCCCCCGGTATCGGGCCAGAGGT | 8 |
| GD1-18120-18240 | 1716580 | CGTCCTCCAGAGACCCGGGCTTCAGGCCCTGCCTCTCCTGTTACCCTTTTAGAACCACAGCCTGGACACATGTGCCAGACGCCTGGGCCTCTAAGGCCCTCGGGTCCCCCTGGACCCCGG | 8 |
| GD1-18150-18270 | 1888238 | GCCTCTCCTGTTACCCTTTTAGAACCACAGCCTGGACACATGTGCCAGACGCCTGGGCCTCTAAGGCCCTCGGGTCCCCCTGGACCCCGGCCTCAGCAACCCTGCTGCTCCCCTCCTGCC | 8 |
| GD1-18180-18300 | 2059896 | CCTGGACACATGTGCCAGACGCCTGGGCCTCTAAGGCCCTCGGGTCCCCCTGGACCCCGGCCTCAGCAACCCTGCTGCTCCCCTCCTGCCACCCCAGCCTCCCCCCCTCCCCGTCCCCCT | 8 |
| GD1-18210-18330 | 2231554 | CTAAGGCCCTCGGGTCCCCCTGGACCCCGGCCTCAGCAACCCTGCTGCTCCCCTCCTGCCACCCCAGCCTCCCCCCCTCCCCGTCCCCCTTCGCTCCTGATCCTCCCCCGGTCCCCAGTA | 8 |
| GD1-18240-18360 | 2403212 | CCTCAGCAACCCTGCTGCTCCCCTCCTGCCACCCCAGCCTCCCCCCCTCCCCGTCCCCCTTCGCTCCTGATCCTCCCCCGGTCCCCAGTAGGGCCGCCTGCCCCCCTGCACCCAGTACCT | 8 |
| GD1-21540-21660 | 2574870 | AGGCCCGCGGCCATTGGGCCCAGATTGAGAGACCAGTCCAGGGGCCCGAGGTTGGAGCCAGCGGGCACCCGAGGTCCCAGCACCCGGTCCCTCCGGGGGGCAGAGACAGGCAGGGCCCCC | 8 |
| GD1-21570-21690 | 2746528 | GACCAGTCCAGGGGCCCGAGGTTGGAGCCAGCGGGCACCCGAGGTCCCAGCACCCGGTCCCTCCGGGGGGCAGAGACAGGCAGGGCCCCCCGGCAGCTGGCCCCGAGGAGGCGCCCGGAG | 12 |
| GD1-21810-21930 | 2918186 | GTCGGCCTAGGCCCGGGGAAGTGGAGGGGGATCGCCCGGGTCTCTGTTGGCAGAGTCCGGGCGATCCTCTGAGACCCTCCGGGCCCGGACGGCCGCCCTCAGCCCCCCAGACAGACCCCA | 8 |
| GD1-23130-23250 | 3089844 | CCCCAGAAGCCCCCAAAAGTAGAGGCTCAGGCCATGCGCGCCCTGTCACCAGGCCTGCCAAAGAGCCAGATCTAAGGCCAGGAGAGGCAGCCCCAAAGCGGGTGCAGTAACAGGTAATCT | 4 |
| GD1-23160-23280 | 3261502 | GCCATGCGCGCCCTGTCACCAGGCCTGCCAAAGAGCCAGATCTAAGGCCAGGAGAGGCAGCCCCAAAGCGGGTGCAGTAACAGGTAATCTCTGGTAGTGATTTGGACCCGAAATCTGACA | 4 |
| GD1-23190-23310 | 3433160 | AAGAGCCAGATCTAAGGCCAGGAGAGGCAGCCCCAAAGCGGGTGCAGTAACAGGTAATCTCTGGTAGTGATTTGGACCCGAAATCTGACACTTTAGAGCTCTGGAGGACTTTAAAACTCT | 4 |
| GD1-23220-23340 | 3604818 | CCCCAAAGCGGGTGCAGTAACAGGTAATCTCTGGTAGTGATTTGGACCCGAAATCTGACACTTTAGAGCTCTGGAGGACTTTAAAACTCTAAAAATCAAAACTTTAGAGGCGAATGGGCG | 4 |
| GD1-23250-23370 | 3776476 | CTGGTAGTGATTTGGACCCGAAATCTGACACTTTAGAGCTCTGGAGGACTTTAAAACTCTAAAAATCAAAACTTTAGAGGCGAATGGGCGCCATTTTGTCCCCACGCGCGCATAATGGCG | 4 |
| GD1-23550-23670 | 3948134 | GTAAGCCAGACAGCAGCCAATTGTCAGTTCTAGGGAGGGGGACCACTGCCCCTGGTATAAAGTGGTCCTGCAGCTATTTCTGGTCGCATCAGAGCGCCAGGAGTCCACACAAATGTAAGA | 4 |
| GD1-23790-23910 | 4119792 | CTGACTCCGCCATCCAAGCCTAGGGGAGACCGAAGTGAAGTCCCTGGACCAGCCCGGCCCGGGCCCCCCGGTATCGGGCCAGAGGTAAGTGGACTTTAATTTTTTCTGCTAAGCCCAACA | 4 |
| GD1-24510-24630 | 4291450 | CTCTTGGCCACGCACCTCGGGCCAGGCCACCTTAGACCCGGCCAAGCCCCATCCCTGAAGACCCAGCGGCCATTCTCTCTGGTAACGAGCAGAGAAGAAGTAGAGGCCCGCGGCCATTGG | 4 |
| GD1-25260-25380 | 4463108 | TGTCTGAGGGGAGCCACCCTCGGGGCCCAGGCCCCAGAGTCCAGAGGTCAGGGGCACCTCAGGGTGCCTCCCCGGGTCCCAGGCCAGCCGGAGGGACCCCGGCAGCCCGGGCGGCCCCAG | 12 |
| GD1-25290-25410 | 4634766 | GCCCCAGAGTCCAGAGGTCAGGGGCACCTCAGGGTGCCTCCCCGGGTCCCAGGCCAGCCGGAGGGACCCCGGCAGCCCGGGCGGCCCCAGAGGCCGGTTCCTCGCCCCTTCCCCGGGCTT | 12 |
| GD1-25320-25440 | 4806424 | AGGGTGCCTCCCCGGGTCCCAGGCCAGCCGGAGGGACCCCGGCAGCCCGGGCGGCCCCAGAGGCCGGTTCCTCGCCCCTTCCCCGGGCTTCAGAGCCCAGGATGTCCCCCAGAAGGGACC | 12 |
| GD1-26130-26250 | 4978082 | CTTTGTCCAGATGTCCAGGGGTCCCCGAGGGCGAGGCCCAGCCCCCTCCCACCCCTGTCCACTGCCCCGGTCCCCCCAGAAGCCCCCAAAAGTAGAGGCTCAGGCCATGCGCGCCCTGTC | 8 |
| GD1-28740-28860 | 5149740 | AGTCGCCTTCCCTCTCCCCTGGCCTCTCCTTCCCGCCTCCCACCCGAGCCCCCTCAGCTTGCCTCCCCACCGGGTCCATCAGGCCGGCCGGAGGGACCCCGGCGGCCCGGTGTCAGTCCC | 8 |

| | | | |
|---|---|---|---|
| GD1-29160-29280 | 5321398 | TTGCCCCCCTGGCGGCCCAGCCCGACCCCCGGGCGCCCCCAAACTTTGTCCAGATGTCCAGGGGTCCCCGAGGGCGAGGCCCAGCCCCCTCCCACCCCTGTCCACTGCCCCGGTCCCCCC | 12 |
| GD1-30120-30240 | 5493056 | TAAGAAGGCACCGGTCGCCCAGTCCTACCAGAGGGGGCCAAGAACCCAGACGAGTCCGTAGAAGGGTCCTCGTCCAGCAAGAAGAGGAGGTGGTAAGCGGTTCACCTTCAGGGGTAAGTA | 4 |
| GD1-30150-30270 | 5664714 | GAGGGGGCCAAGAACCCAGACGAGTCCGTAGAAGGGTCCTCGTCCAGCAAGAAGAGGAGGTGGTAAGCGGTTCACCTTCAGGGGTAAGTAACCTGACCTCTCCAGGGCTCACATAAAGGG | 4 |
| GD1-32940-33060 | 5836372 | CAGTTTCTAACTTTTTCCCCTTCCCTCCCTCGTCTTGCCCTGCGCCCGGGGCCACCTTCATCACCGTCGCTGACTCCGCCATCCAAGCCTAGGGGAGACCGAAGTGAAGTCCCTGGACCA | 4 |
| GD1-34200-34320 | 6008030 | TCAGCCCCCCAGACAGACCCCAGGGTCTCCAGGCAGGGTCCGGCATCTCCAGGGGCAGCAGGCTCACCACCACAGGCCCCCCAGACCCGGGTCTCGGCCAGCCGAGCCGACCGGCCCCGC | 8 |
| GD1-34230-34350 | 6179688 | AGGCAGGGTCCGGCATCTCCAGGGGCAGCAGGCTCACCACCACAGGCCCCCCAGACCCGGGTCTCGGCCAGCCGAGCCGACCGGCCCCGCGCCGGGCGCCTCCTCGGGGCCAGCCGCCGG | 12 |
| GD1-34260-34380 | 6351346 | GGCTCACCACCACAGGCCCCCCAGACCCGGGTCTCGGCCAGCCGAGCCGACCGGCCCCGCGCCGGGCGCCTCCTCGGGGCCAGCCGCCGGGGTTGGTTCTGCCCCTCTCTCTGTCCTTCA | 12 |
| GD1-34290-34410 | 6523004 | GTCTCGGCCAGCCGAGCCGACCGGCCCCGCGCCGGGCGCCTCCTCGGGGCCAGCCGCCGGGGTTGGTTCTGCCCCTCTCTCTGTCCTTCAGAGGAACCAGGGACCTCGGGCACCCCAGAG | 8 |
| GD1-37200-37320 | 6694662 | GGGGCAGGGGCAGGGGCAGGGGCAAGTCCAGGGACAAGCAACGCAAGCCCGGTGGACCATGGAGACCAGAGCCAAACACCTCCAGTCCTAGCATGCCTGAACTAAGTCCAGTCCTCGGTC | 4 |
| GD1-38340-38460 | 6866320 | TGCGCCGCCGCCAGGTCCTGGGGCAGCCGGCGTTCCTGGCGCTCCGGGGGCAGCCGGGCGGCCGCCGGTGGGTCCGCTGGGCCGCTGCCCCGCTCCGGGTGGGGGTGGCCCCGCTGGGC | 12 |
| GD1-38370-38490 | 7037978 | CGTTCCTGGCGCTCCGGGGGCAGCCGGGCGGCCGCCGGTGGGTCCGCTGGGCCGCTGCCCCGCTCCGGGTGGGGGTGGCCCCGCTGGGCACCGCTGCGCCGCCGCCAGGTCCTGGGGCA | 12 |
| GD1-38460-38580 | 7209636 | ACCGCTGCGCCGCCGCCAGGTCCTGGGGCAGCCGGGGTTCCTGGCGCTCCGGGGGCAGCAGGGCGGCCGCCGGTGGGTCCGCTGGGCCGCTGCCCCGCTCCGGGTGGGGGTGGCCCCGC | 12 |
| GD1-38490-38610 | 7381294 | GCCGGGGTTCCTGGCGCTCCGGGGGCAGCAGGGCGGCCGCCGGTGGGTCCGCTGGGCCGCTGCCCCGCTCCGGGTGGGGGTGGCCCCGCTGGGCACCGCTGCGCCGCCGCCAGGTCCTG | 12 |
| GD1-38760-38880 | 7552952 | GGGGGCAGCAGGGCGGCCGCCGGTGGGTCCGCTGGGCCGCTGCCCCGCTCCGGGTGGGGGTGGCCCCGCTGGGCACCGCTGCGCCGCCGCCAGGTCCTGGGGCAGCCGGGGTTCCTGGC | 12 |
| GD1-38790-38910 | 7724610 | GCTGGGCCGCTGCCCCGCTCCGGGTGGGGGTGGCCCCGCTGGGCACCGCTGCGCCGCCGCCAGGTCCTGGGGCAGCCGGGGTTCCTGGCGCTCCGGGGGCAGCAGGGCGGCCGCCGGTG | 12 |
| GD1-38820-38940 | 7896268 | GTGGCCCCGCTGGGCACCGCTGCGCCGCCGCCAGGTCCTGGGGCAGCCGGGGTTCCTGGCGCTCCGGGGGCAGCAGGGCGGCCGCCGGTGGGTCCGCTGGGCCGCTGCCCCGCTCCGGGT | 12 |
| GD1-39000-39120 | 8067926 | CTGGCGCTCCGGGGGCAGCAGGGCGGCCGCCGGTGGGTCCGCTGGGCCGCTGCCCCGCTCCGGGTGGGGGTGGCCCCGCTGGGCACCGCTGCGCCGCCGCCAGGTCCTGGGGCAGCCGG | 12 |
| GD1-39600-39720 | 8239584 | CCAGGTCCTGGGGCAGCCGGGGTTCCTGGCGCTCCGGGGGCAGCAGGCCGGCCGCCGGTGGGTCCGCAGGACCGATGCACCGCTCCGGGTGGGGGTGGCCCCGCTGGGCACCGCTGCGC | 12 |
| GD1-39630-39750 | 8411242 | GCTCCGGGGGCAGCAGGCCGGCCGCCGGTGGGTCCGCAGGACCGATGCACCGCTCCGGGTGGGGGTGGCCCCGCTGGGCACCGCTGCGCCGCCGCCAGGTCCTGGGGCAGCCGGGGTTC | 12 |
| GD1-40800-40920 | 8582900 | ATACCAGGTGGGTGGAGCTAGGTAGGATTCAGCTAGGTTCCTACTGGGGTACCCCCCTACCCTACCTTAAGGTGCGCCACCCTTCCTCCTTCCGTTTTAATGGTAGAATAACCTATAGGT | 4 |
| GD1-40830-40950 | 8754558 | AGCTAGGTTCCTACTGGGGTACCCCCCTACCCTACCTTAAGGTGCGCCACCCTTCCTCCTTCCGTTTTAATGGTAGAATAACCTATAGGTTATTAACCTAGTGGTGGAATAGGGTATTGC | 4 |
| GD1-40860-40980 | 8926216 | CCTACCTTAAGGTGCGCCACCCTTCCTCCTTCCGTTTTAATGGTAGAATAACCTATAGGTTATTAACCTAGTGGTGGAATAGGGTATTGCAGCTGGGTATATACCTATAGGTATATAGAA | 4 |
| GD1-40890-41010 | 9097874 | TCCGTTTTAATGGTAGAATAACCTATAGGTTATTAACCTAGTGGTGGAATAGGGTATTGCAGCTGGGTATATACCTATAGGTATATAGAACCTAGAGGAAGGGAACCCTATAGTGTAATC | 4 |
| GD1-40920-41040 | 9269532 | TATTAACCTAGTGGTGGAATAGGGTATTGCAGCTGGGTATATACCTATAGGTATATAGAACCTAGAGGAAGGGAACCCTATAGTGTAATCCCTCCCCCCCCTACCCCCCCCTCCCTTACG | 4 |
| GD1-40950-41070 | 9441190 | AGCTGGGTATATACCTATAGGTATATAGAACCTAGAGGAAGGGAACCCTATAGTGTAATCCCTCCCCCCCCTACCCCCCCCTCCCTTACGGTTGCCTGAGCCCATCCCCCACCCCAGCAC | 4 |
| GD1-40980-41100 | 9612848 | CCTAGAGGAAGGGAACCCTATAGTGTAATCCCTCCCCCCCCTACCCCCCCCTCCCTTACGGTTGCCTGAGCCCATCCCCCACCCCAGCACCCCGGGGTGACGTGGCACCCCGCGTGCCTT | 8 |
| GD1-57540-57660 | 9784506 | GGCTGCTGCTGAGTTGGAGGAGGAGAAGGAGCCGGGGCGGCTGATTGCAGGGGCTGCTGCTGAGTTGGAGGAGGAGAAAGAGTCGTGGCGGTGGGGGCTGCTGCTGCAGTCGGGGAAGGG | 8 |
| GD1-57570-57690 | 9956164 | GCCGGGGCGGCTGATTGCAGGGGCTGCTGCTGAGTTGGAGGAGGAGAAAGAGTCGTGGCGGTGGGGGCTGCTGCTGCAGTCGGGGAAGGGGATGGGGTGGTCAGAGGGATTTCTGGGTTC | 8 |
| GD1-58110-58230 | 10127822 | GAGTCCGGAGCCGCAGAGGCCGGAGCCGCAGAGGCCGGAGCCACAGAGGCCGGTGCCACAGAGGCCTGAGCTGCAGAGGCCGGAGCCGCAGAGGCCGGAGCCGCAGAGGCCGGAGCCGCA | 12 |
| GD1-58140-58260 | 10299480 | GAGGCCGGAGCCACAGAGGCCGGTGCCACAGAGGCCTGAGCTGCAGAGGCCGGAGCCGCAGAGGCCGGAGCCGCAGAGGCCGGAGCCGCAGAGGCCGGAGCCGCAGAGGCCGGAGACGGC | 12 |
| GD1-58170-58290 | 10471138 | GAGGCCTGAGCTGCAGAGGCCGGAGCCGCAGAGGCCGGAGCCGCAGAGGCCGGAGCCGCAGAGGCCGGAGCCGCAGAGGCCGGAGACGGCGGCGGGGAGTTGGTCTTTGCAGGACTATAC | 8 |
| GD1-58200-58320 | 10642796 | GAGGCCGGAGCCGCAGAGGCCGGAGCCGCAGAGGCCGGAGCCGCAGAGGCCGGAGACGGCGGCGGGGAGTTGGTCTTTGCAGGACTATACCTGGCGGCAGGGAATGAGTCGGATGTGAAA | 8 |
| GD1-61500-61620 | 10814454 | CGCCACCGCCCCCACACCCACAGCCGTCGCGGCGGGCGTAGAGGGGCCGCTGGTGCGCCGGGCTCGTCTGTCCTTGGCCTCGGCCTCCGCCCGCAGGTAGGCCGCCCGGGCCACACGGGC | 12 |
| GD1-61530-61650 | 10986112 | GGCGGGCGTAGAGGGGCCGCTGGTGCGCCGGGCTCGTCTGTCCTTGGCCTCGGCCTCCGCCCGCAGGTAGGCCGCCCGGGCCACACGGGCGAAGCGGCTCGTGGGGCTCGCGGTGGGCAG | 12 |
| GD1-61560-61680 | 11157770 | GGCTCGTCTGTCCTTGGCCTCGGCCTCCGCCCGCAGGTAGGCCGCCCGGGCCACACGGGCGAAGCGGCTCGTGGGGCTCGCGGTGGGCAGCAGTCGGAAAAAGTGCAGGGCAAAGCCCGA | 8 |
| GD1-77820-77940 | 11329428 | ATATGTGACGTTGTCTCTGAGATTGAACTGGGTCTCAGTGACATGGAAGAGGTTGAACTTGAAGTTATGTTATGTTGGCCTGTGGTAACAGCACTGGTTGCATTTTTTGGTTGGCTGGTA | 4 |
| GD1-81780-81900 | 11501086 | TATCCCCTCAGCAACCAATGGAGGGGCCTGGTGCCCCCTTTAGCCAGGTTGCTGATGTGGTCCACACACCTGGGGTACCGGCGATGCAGCCACAGTACTTTGACCTCCCCTTAATTCAAC | 4 |
| GD1-81810-81930 | 11672744 | GTGCCCCCTTTAGCCAGGTTGCTGATGTGGTCCACACACCTGGGGTACCGGCGATGCAGCCACAGTACTTTGACCTCCCCTTAATTCAACCCATTAGCCAGGGGGCACCCGTGGCCCCGT | 4 |
| GD1-81840-81960 | 11844402 | TCCACACACCTGGGGTACCGGCGATGCAGCCACAGTACTTTGACCTCCCCTTAATTCAACCCATTAGCCAGGGGGCACCCGTGGCCCCGTTGAGGGCTAGTATGGGCCCGGTACCTCCGG | 4 |
| GD1-87690-87810 | 12016060 | CTCCTGCCGCGGGGCCTCCTGCCGCGGGGCCTCACATACTGACGCCTCCTTCCGCCCGGCCTCGTATAATGGCGCCTCCCGTCGTACGTATGTTTATGAGGGAGCGACAGCTCCCCCAGT | 8 |
| GD1-87720-87840 | 12187718 | CTCACATACTGACGCCTCCTTCCGCCCGGCCTCGTATAATGGCGCCTCCCGTCGTACGTATGTTTATGAGGGAGCGACAGCTCCCCCAGTCCACCGGCCGTAAACCTCAGTGCTTCTGGG | 4 |
| GD1-96360-96480 | 12359376 | CAGGAGCAGGAGGGGCAGGAGGGGCAGGAGCAGGGGAGGAGGCAGGAGGGGCAGGAGCAGGAGGAGGGGCAGGAGGGGCAGGAGCAGGAGGGGCAGGAGGGGCAGGAGCAGGAGGGGCAG | 8 |
| GD1-96390-96510 | 12531034 | CAGGGGAGGAGGCAGGAGGGGCAGGAGCAGGAGGAGGGGCAGGAGGGGCAGGAGCAGGAGGGGCAGGAGGGGCAGGAGCAGGAGGGGCAGGAGGGGCAGGAGCAGGAGGGGCAGGAGGGG | 8 |
| GD1-140790-140910 | 12702692 | CTCGGGGGGCGGCCGGGTGGCCCACCGGGTCCGCTGGGTCCGCTGCCCCGCTCCGGCGGGGGGTGGCCGGCTGCAGCCGGGTCCGGGGTTCCGGCCCTGGAGCTCGGGGGGCGGCCGGGT | 12 |
| GD1-140850-140970 | 12874350 | GGGTGGCCGGCTGCAGCCGGGTCCGGGGTTCCGGCCCTGGAGCTCGGGGGGCGGCCGGGTGGCCCACCGGGTCCGCTGGGTCCGCTGCCCCGCTCCGGCGGGGGGTGGCCGGCTGCAGCC | 12 |
| GD1-140880-141000 | 13046008 | CCGGCCCTGGAGCTCGGGGGGCGGCCGGGTGGCCCACCGGGTCCGCTGGGTCCGCTGCCCCGCTCCGGCGGGGGGTGGCCGGCTGCAGCCGGGTCCGGGGTTCCGGCCCTGGAGCTCGGG | 12 |
| GD1-140910-141030 | 13217666 | GGCCCACCGGGTCCGCTGGGTCCGCTGCCCCGCTCCGGCGGGGGGTGGCCGGCTGCAGCCGGGTCCGGGGTTCCGGCCCTGGAGCTCGGGGGGCGGCCGGGTGGCCCACCGGGTCCGCTG | 12 |
| GD1-140940-141060 | 13389324 | CGCTCCGGCGGGGGGTGGCCGGCTGCAGCCGGGTCCGGGGTTCCGGCCCTGGAGCTCGGGGGGCGGCCGGGTGGCCCACCGGGTCCGCTGGGTCCGCTGCCCCGCTCCGGCGGGGGGTGG | 12 |
| GD1-140970-141090 | 13560982 | GGGTCCGGGGTTCCGGCCCTGGAGCTCGGGGGGCGGCCGGGTGGCCCACCGGGTCCGCTGGGTCCGCTGCCCCGCTCCGGCGGGGGGTGGCCGGCTGCAGCCGGGTCCGGGGTTCCGGCC | 12 |
| GD1-141000-141120 | 13732640 | GGGCGGCCGGGTGGCCCACCGGGTCCGCTGGGTCCGCTGCCCCGCTCCGGCGGGGGGTGGCCGGCTGCAGCCGGGTCCGGGGTTCCGGCCCTGGAGCTCGGGGGGCGGCCGGGTGGCCCA | 12 |
| GD1-141030-141150 | 13904298 | GGTCCGCTGCCCCGCTCCGGCGGGGGGTGGCCGGCTGCAGCCGGGTCCGGGGTTCCGGCCCTGGAGCTCGGGGGGCGGCCGGGTGGCCCACCGGGTCCGCTGGGTCCGCTGCCCCGCTCC | 12 |

| | | | |
|---|---|---|---|
| GD1-141060-141180 | 14075956 | CCGGCTGCAGCCGGGTCCGGGGTTCCGGCCCTGGAGCTCGGGGGGCGGCCGGGTGGCCCACCGGGTCCGCTGGGTCCGCTGCCCCGCTCCGGCGGGGGGTGGCCGGCTGCAGCCGGGTCC | 12 |
| GD1-141090-141210 | 14247614 | CTGGAGCTCGGGGGGCGGCCGGGTGGCCCACCGGGTCCGCTGGGTCCGCTGCCCCGCTCCGGCGGGGGGTGGCCGGCTGCAGCCGGGTCCGGGGTTCCGGCCCTGGAGCTCGGGGGGCGG | 12 |
| GD1-141120-141240 | 14419272 | CCGGGTCCGCTGGGTCCGCTGCCCCGCTCCGGCGGGGGGTGGCCGGCTGCAGCCGGGTCCGGGGTTCCGGCCCTGGAGCTCGGGGGGCGGCCGGGTGGCCCACCGGGTCCGCTGGGTCCG | 12 |
| GD1-141150-141270 | 14590930 | GGCGGGGGGTGGCCGGCTGCAGCCGGGTCCGGGGTTCCGGCCCTGGAGCTCGGGGGGCGGCCGGGTGGCCCACCGGGTCCGCTGGGTCCGCTGCCCCGCTCCGGCGGGGGGTGGCCGGCT | 12 |
| GD1-141180-141300 | 14762588 | GGGGTTCCGGCCCTGGAGCTCGGGGGGCGGCCGGGTGGCCCACCGGGTCCGCTGGGTCCGCTGCCCCGCTCCGGCGGGGGGTGGCCGGCTGCAGCCGGGTCCGGGGTTCCGGCCCTGGAG | 12 |
| GD1-141210-141330 | 14934246 | CCGGGTGGCCCACCGGGTCCGCTGGGTCCGCTGCCCCGCTCCGGCGGGGGGTGGCCGGCTGCAGCCGGGTCCGGGGTTCCGGCCCTGGAGCTCGGGGGGCGGCCGGGTGGCCCACCGGGT | 12 |
| GD1-141300-141420 | 15105904 | CTCGGGGGGCGGCCGGGTGGCCCACCGGGTCCGCTGGGTCCGCTGCCCCGCTCCGGCGGGGGGTGGCCGGCTGCAGCCGGGTCCGGGGTTCCGGCCCTGGAGCTCGGGGGGCGGCCGGGT | 12 |
| GD1-141330-141450 | 15277562 | CCGCTGGGTCCGCTGCCCCGCTCCGGCGGGGGGTGGCCGGCTGCAGCCGGGTCCGGGGTTCCGGCCCTGGAGCTCGGGGGGCGGCCGGGTGGCCCACCGGGTCCGCTGGGTCCGCTGCCC | 12 |
| GD1-141420-141540 | 15449220 | GGCCCACCGGGTCCGCTGGGTCCGCTGCCCCGCTCCGGCGGGGGGTGGCCGGCTGCAGCCGGGTCCGGGGTTCCGGCCCTGGAGCTCGGGGGGCGGCCGGGTGGCCCACCGGGTCCGCTG | 12 |
| GD1-141450-141570 | 15620878 | CGCTCCGGCGGGGGGTGGCCGGCTGCAGCCGGGTCCGGGGTTCCGGCCCTGGAGCTCGGGGGGCGGCCGGGTGGCCCACCGGGTCCGCTGGGTCCGCTGCCCCGCTCCGGCGGGGGGTGG | 12 |
| GD1-141480-141600 | 15792536 | GGGTCCGGGGTTCCGGCCCTGGAGCTCGGGGGGCGGCCGGGTGGCCCACCGGGTCCGCTGGGTCCGCTGCCCCGCTCCGGCGGGGGGTGGCCGGCTGCAGCCGGGTCCGGGGTTCCGGCC | 12 |
| GD1-141510-141630 | 15964194 | GGGCGGCCGGGTGGCCCACCGGGTCCGCTGGGTCCGCTGCCCCGCTCCGGCGGGGGGTGGCCGGCTGCAGCCGGGTCCGGGGTTCCGGCCCTGGAGCTCGGGGGGCGGCCGGGTGGCCCA | 12 |
| GD1-141540-141660 | 16135852 | GGTCCGCTGCCCCGCTCCGGCGGGGGGTGGCCGGCTGCAGCCGGGTCCGGGGTTCCGGCCCTGGAGCTCGGGGGGCGGCCGGGTGGCCCACCGGGTCCGCTGGGTCCGCTGCCCCGCTCC | 12 |
| GD1-141600-141720 | 16307510 | CTGGAGCTCGGGGGGCGGCCGGGTGGCCCACCGGGTCCGCTGGGTCCGCTGCCCCGCTCCGGCGGGGGGTGGCCGGCTGCAGCCGGGTCCGGGGTTCCGGCCCTGGAGCTCGGGGGGCGG | 12 |
| GD1-141630-141750 | 16479168 | CCGGGTCCGCTGGGTCCGCTGCCCCGCTCCGGCGGGGGGTGGCCGGCTGCAGCCGGGTCCGGGGTTCCGGCCCTGGAGCTCGGGGGGCGGCCGGGTGGCCCACCGGGTCCGCTGGGTCCG | 12 |
| GD1-141660-141780 | 16650826 | GGCGGGGGGTGGCCGGCTGCAGCCGGGTCCGGGGTTCCGGCCCTGGAGCTCGGGGGGCGGCCGGGTGGCCCACCGGGTCCGCTGGGTCCGCTGCCCCGCTCCGGCGGGGGGTGGCCGGCT | 12 |
| GD1-141690-141810 | 16822484 | GGGGTTCCGGCCCTGGAGCTCGGGGGGCGGCCGGGTGGCCCACCGGGTCCGCTGGGTCCGCTGCCCCGCTCCGGCGGGGGGTGGCCGGCTGCAGCCGGGTCCGGGGTTCCGGCCCTGGAG | 12 |
| GD1-141720-141840 | 16994142 | CCGGGTGGCCCACCGGGTCCGCTGGGTCCGCTGCCCCGCTCCGGCGGGGGGTGGCCGGCTGCAGCCGGGTCCGGGGTTCCGGCCCTGGAGCTCGGGGGGCGGCCGGGTGGCCCACCGGGT | 12 |
| GD1-141750-141870 | 17165800 | CTGCCCCGCTCCGGCGGGGGGTGGCCGGCTGCAGCCGGGTCCGGGGTTCCGGCCCTGGAGCTCGGGGGGCGGCCGGGTGGCCCACCGGGTCCGCTGGGTCCGCTGCCCCGCTCCGGCGGG | 12 |
| GD1-141840-141960 | 17337458 | CCGCTGGGTCCGCTGCCCCGCTCCGGCGGGGGGTGGCCGGCTGCAGCCGGGTCCGGGGTTCCGGCCCTGGAGCTCGGGGGGCGGCCGGGTGGCCCACCGGGTCCGCTGGGTCCGCTGCCC | 12 |
| GD1-141870-141990 | 17509116 | GGGTGGCCGGCTGCAGCCGGGTCCGGGGTTCCGGCCCTGGAGCTCGGGGGGCGGCCGGGTGGCCCACCGGGTCCGCTGGGTCCGCTGCCCCGCTCCGGCGGGGGGTGGCCGGCTGCAGCC | 12 |
| GD1-141900-142020 | 17680774 | CCGGCCCTGGAGCTCGGGGGGCGGCCGGGTGGCCCACCGGGTCCGCTGGGTCCGCTGCCCCGCTCCGGCGGGGGGTGGCCGGCTGCAGCCGGGTCCGGGGTTCCGGCCCTGGAGCTCGGG | 12 |
| GD1-141930-142050 | 17852432 | GGCCCACCGGGTCCGCTGGGTCCGCTGCCCCGCTCCGGCGGGGGGTGGCCGGCTGCAGCCGGGTCCGGGGTTCCGGCCCTGGAGCTCGGGGGGCGGCCGGGTGGCCCACCGGGTCCGCTG | 12 |
| GD1-141960-142080 | 18024090 | CGCTCCGGCGGGGGGTGGCCGGCTGCAGCCGGGTCCGGGGTTCCGGCCCTGGAGCTCGGGGGGCGGCCGGGTGGCCCACCGGGTCCGCTGGGTCCGCTGCCCCGCTCCGGCGGGGGGTGG | 12 |
| GD1-142110-142230 | 18195748 | CTGGAGCTCGGGGGGCGGCCGGGTGGCCCACCGGGTCCGCTGGGTCCGCTGCCCCGCTCCGGCGGGGGGTGGCCGGCTGCAGCCGGGTCCGGGGTTCCGGCCCTGGAGCTCGGGGGGCGG | 12 |
| GD1-142140-142260 | 18367406 | CCGGGTCCGCTGGGTCCGCTGCCCCGCTCCGGCGGGGGGTGGCCGGCTGCAGCCGGGTCCGGGGTTCCGGCCCTGGAGCTCGGGGGGCGGCCGGGTGGCCCACCGGGTCCGCTGGGTCCG | 12 |
| GD1-142170-142290 | 18539064 | GGCGGGGGGTGGCCGGCTGCAGCCGGGTCCGGGGTTCCGGCCCTGGAGCTCGGGGGGCGGCCGGGTGGCCCACCGGGTCCGCTGGGTCCGCTGCCCCGCTCCGGCGGGGGGTGGCCGGCT | 12 |
| GD1-142200-142320 | 18710722 | GGGGTTCCGGCCCTGGAGCTCGGGGGGCGGCCGGGTGGCCCACCGGGTCCGCTGGGTCCGCTGCCCCGCTCCGGCGGGGGGTGGCCGGCTGCAGCCGGGTCCGGGGTTCCGGCCCTGGAG | 12 |
| GD1-142230-142350 | 18882380 | CCGGGTGGCCCACCGGGTCCGCTGGGTCCGCTGCCCCGCTCCGGCGGGGGGTGGCCGGCTGCAGCCGGGTCCGGGGTTCCGGCCCTGGAGCTCGGGGGGCGGCCGGGTGGCCCACCGGGT | 12 |
| GD1-142260-142380 | 19054038 | CTGCCCCGCTCCGGCGGGGGGTGGCCGGCTGCAGCCGGGTCCGGGGTTCCGGCCCTGGAGCTCGGGGGGCGGCCGGGTGGCCCACCGGGTCCGCTGGGTCCGCTGCCCCGCTCCGGCGGG | 12 |
| GD1-142290-142410 | 19225696 | GCAGCCGGGTCCGGGGTTCCGGCCCTGGAGCTCGGGGGGCGGCCGGGTGGCCCACCGGGTCCGCTGGGTCCGCTGCCCCGCTCCGGCGGGGGGTGGCCGGCTGCAGCCGGGTCCGGGGTT | 12 |
| GD1-142320-142440 | 19397354 | CTCGGGGGGCGGCCGGGTGGCCCACCGGGTCCGCTGGGTCCGCTGCCCCGCTCCGGCGGGGGGTGGCCGGCTGCAGCCGGGTCCGGGGTTCCGGCCCTGGAGCTCGGGGGGCGGCCGGGT | 12 |
| GD1-142350-142470 | 19569012 | CCGCTGGGTCCGCTGCCCCGCTCCGGCGGGGGGTGGCCGGCTGCAGCCGGGTCCGGGGTTCCGGCCCTGGAGCTCGGGGGGCGGCCGGGTGGCCCACCGGGTCCGCTGGGTCCGCTGCCC | 12 |
| GD1-142380-142500 | 19740670 | GGGTGGCCGGCTGCAGCCGGGTCCGGGGTTCCGGCCCTGGAGCTCGGGGGGCGGCCGGGTGGCCCACCGGGTCCGCTGGGTCCGCTGCCCCGCTCCGGCGGGGGGTGGCCGGCTGCAGCC | 12 |
| GD1-142410-142530 | 19912328 | CCGGCCCTGGAGCTCGGGGGGCGGCCGGGTGGCCCACCGGGTCCGCTGGGTCCGCTGCCCCGCTCCGGCGGGGGGTGGCCGGCTGCAGCCGGGTCCGGGGTTCCGGCCCTGGAGCTCGGG | 12 |
| GD1-142440-142560 | 20083986 | GGCCCACCGGGTCCGCTGGGTCCGCTGCCCCGCTCCGGCGGGGGGTGGCCGGCTGCAGCCGGGTCCGGGGTTCCGGCCCTGGAGCTCGGGGGGCGGCCGGGTGGCCCACCGGGTCCGCTG | 12 |
| GD1-142470-142590 | 20255644 | CGCTCCGGCGGGGGGTGGCCGGCTGCAGCCGGGTCCGGGGTTCCGGCCCTGGAGCTCGGGGGGCGGCCGGGTGGCCCACCGGGTCCGCTGGGTCCGCTGCCCCGCTCCGGCGGGGGGTGG | 12 |
| GD1-142500-142620 | 20427302 | GGGTCCGGGGTTCCGGCCCTGGAGCTCGGGGGGCGGCCGGGTGGCCCACCGGGTCCGCTGGGTCCGCTGCCCCGCTCCGGCGGGGGGTGGCCGGCTGCAGCCGGGTCCGGGGTTCCGGCC | 12 |
| GD1-142530-142650 | 20598960 | GGGCGGCCGGGTGGCCCACCGGGTCCGCTGGGTCCGCTGCCCCGCTCCGGCGGGGGGTGGCCGGCTGCAGCCGGGTCCGGGGTTCCGGCCCTGGAGCTCGGGGGGCGGCCGGGTGGCCCA | 12 |
| GD1-142560-142680 | 20770618 | GGTCCGCTGCCCCGCTCCGGCGGGGGGTGGCCGGCTGCAGCCGGGTCCGGGGTTCCGGCCCTGGAGCTCGGGGGGCGGCCGGGTGGCCCACCGGGTCCGCTGGGTCCGCTGCCCCGCTCC | 12 |
| GD1-142740-142860 | 20942276 | CCGGGTGGCCCACCGGGTCCGCTGGGTCCGCTGCCCCGCTCCGGCGGGGGGTGGCCGGCTGCAGCCGGGTCCGGGGTTCCGGCCCTGGAGCTCGGGGGGCGGCCGGGTGGCCCACCGGGT | 12 |
| GD1-142770-142890 | 21113934 | CTGCCCCGCTCCGGCGGGGGGTGGCCGGCTGCAGCCGGGTCCGGGGTTCCGGCCCTGGAGCTCGGGGGGCGGCCGGGTGGCCCACCGGGTCCGCTGGGTCCGCTGCCCCGCTCCGGCGGG | 12 |
| GD1-142800-142920 | 21285592 | GCAGCCGGGTCCGGGGTTCCGGCCCTGGAGCTCGGGGGGCGGCCGGGTGGCCCACCGGGTCCGCTGGGTCCGCTGCCCCGCTCCGGCGGGGGGTGGCCGGCTGCAGCCGGGTCCGGGGTT | 12 |
| GD1-142830-142950 | 21457250 | CTCGGGGGGCGGCCGGGTGGCCCACCGGGTCCGCTGGGTCCGCTGCCCCGCTCCGGCGGGGGGTGGCCGGCTGCAGCCGGGTCCGGGGTTCCGGCCCTGGAGCTCGGGGGGCGGCCGGGT | 12 |
| GD1-142860-142980 | 21628908 | CCGCTGGGTCCGCTGCCCCGCTCCGGCGGGGGGTGGCCGGCTGCAGCCGGGTCCGGGGTTCCGGCCCTGGAGCTCGGGGGGCGGCCGGGTGGCCCACCGGGTCCGCTGGGTCCGCTGCCC | 12 |
| GD1-142890-143010 | 21800566 | GGGTGGCCGGCTGCAGCCGGGTCCGGGGTTCCGGCCCTGGAGCTCGGGGGGCGGCCGGGTGGCCCACCGGGTCCGCTGGGTCCGCTGCCCCGCTCCGGCGGGGGGTGGCCGGCTGCAGCC | 12 |
| GD1-142920-143040 | 21972224 | CCGGCCCTGGAGCTCGGGGGGCGGCCGGGTGGCCCACCGGGTCCGCTGGGTCCGCTGCCCCGCTCCGGCGGGGGGTGGCCGGCTGCAGCCGGGTCCGGGGTTCCGGCCCTGGAGCTCGGG | 12 |
| GD1-142950-143070 | 22143882 | GGCCCACCGGGTCCGCTGGGTCCGCTGCCCCGCTCCGGCGGGGGGTGGCCGGCTGCAGCCGGGTCCGGGGTTCCGGCCCTGGAGCTCGGGGGGCGGCCGGGTGGCCCACCGGGTCCGCTG | 12 |
| GD1-142980-143100 | 22315540 | CGCTCCGGCGGGGGGTGGCCGGCTGCAGCCGGGTCCGGGGTTCCGGCCCTGGAGCTCGGGGGGCGGCCGGGTGGCCCACCGGGTCCGCTGGGTCCGCTGCCCCGCTCCGGCGGGGGGTGG | 12 |
| GD1-143010-143130 | 22487198 | GGGTCCGGGGTTCCGGCCCTGGAGCTCGGGGGGCGGCCGGGTGGCCCACCGGGTCCGCTGGGTCCGCTGCCCCGCTCCGGCGGGGGGTGGCCGGCTGCAGCCGGGTCCGGGGTTCCGGCC | 12 |
| GD1-143040-143160 | 22658856 | GGGCGGCCGGGTGGCCCACCGGGTCCGCTGGGTCCGCTGCCCCGCTCCGGCGGGGGGTGGCCGGCTGCAGCCGGGTCCGGGGTTCCGGCCCTGGAGCTCGGGGGGCGGCCGGGTGGCCCA | 12 |

| | | | |
|---|---|---|---|
| GD1-143070-143190 | 22830514 | GGTCCGCTGCCCCGCTCCGGCGGGGGGTGGCCGGCTGCAGCCGGGTCCGGGGTTCCGGCCCTGGAGCTCGGGGGGCGGCCGGGTGGCCCACCGGGTCCGCTGGGTCCGCTGCCCCGCTCC | 12 |
| GD1-143100-143220 | 23002172 | CCGGCTGCAGCCGGGTCCGGGGTTCCGGCCCTGGAGCTCGGGGGGCGGCCGGGTGGCCCACCGGGTCCGCTGGGTCCGCTGCCCCGCTCCGGCGGGGATGGGGGTGCGCTCCCAGGCCGG | 12 |
| GD1-143760-143880 | 23173830 | ACCCTACCTTAAGGTGCGCACCCTTCCTCCTTCCGTTTTAATGAAAGAATAACCTATAGGTTATTAACCTAGTGGTGGAATAGGGTATTGCAGCTGGGTATATACCTATAGGTATATAGA | 4 |
| GD1-143790-143910 | 23345488 | TTCCGTTTTAATGAAAGAATAACCTATAGGTTATTAACCTAGTGGTGGAATAGGGTATTGCAGCTGGGTATATACCTATAGGTATATAGAACCTAGAGGAAGGGAACCCTATAGTGTAAT | 4 |
| GD1-143820-143940 | 23517146 | TTATTAACCTAGTGGTGGAATAGGGTATTGCAGCTGGGTATATACCTATAGGTATATAGAACCTAGAGGAAGGGAACCCTATAGTGTAATCCCTCCCCCCCCTACCCCCCCCTCCCTTAC | 4 |
| GD1-143850-143970 | 23688804 | CAGCTGGGTATATACCTATAGGTATATAGAACCTAGAGGAAGGGAACCCTATAGTGTAATCCCTCCCCCCCCTACCCCCCCCTCCCTTACGGTTGCTTGAGCCCATCCCCCACCCCAGCA | 4 |
| GD1-143880-144000 | 23860462 | ACCTAGAGGAAGGGAACCCTATAGTGTAATCCCTCCCCCCCCTACCCCCCCCTCCCTTACGGTTGCTTGAGCCCATCCCCCACCCCAGCACCCCGGGGTGACGTGGCACCCCGCGTGCCT | 8 |
| GD1-143910-144030 | 24032120 | CCCTCCCCCCCCTACCCCCCCCTCCCTTACGGTTGCTTGAGCCCATCCCCCACCCCAGCACCCCGGGGTGACGTGGCACCCCGCGTGCCTTACTGACTTGTCACCTTTGCACATTTGGTC | 8 |
| GD1-143940-144060 | 24203778 | GGTTGCTTGAGCCCATCCCCCACCCCAGCACCCCGGGGTGACGTGGCACCCCGCGTGCCTTACTGACTTGTCACCTTTGCACATTTGGTCAGCTGACCGATGCTCGCCACTTCCTGGGTC | 4 |
| GD1-143970-144090 | 24375436 | CCCCGGGGTGACGTGGCACCCCGCGTGCCTTACTGACTTGTCACCTTTGCACATTTGGTCAGCTGACCGATGCTCGCCACTTCCTGGGTCATGACCTGGCCTGTGCCTTGTCCCGTGGAC | 4 |
| GD1-144000-144120 | 24547094 | TACTGACTTGTCACCTTTGCACATTTGGTCAGCTGACCGATGCTCGCCACTTCCTGGGTCATGACCTGGCCTGTGCCTTGTCCCGTGGACAATGTCCCTCCAGCGTGGTGGCTGCCTTTG | 4 |
| GD1-144030-144150 | 24718752 | AGCTGACCGATGCTCGCCACTTCCTGGGTCATGACCTGGCCTGTGCCTTGTCCCGTGGACAATGTCCCTCCAGCGTGGTGGCTGCCTTTGGGATGCATCACTTTGAGCCACTAAGCCCCC | 4 |
| GD1-167910-168030 | 24890410 | TCAGGGTCCTGTGGGCCATTGTCATCAGTGTTGTCAGGGTCCTGTGGGCCATTGTCAGGACCACCTCCAGGTGCGCCTAGGTTTTGAGAGCAGAGTGGGGGTCCGTCGCCGGCCCCACTC | 4 |
| GD1-169500-169620 | 25062068 | GGGGGGCGCATTCCTGGAAAAAGTGGAGGGGGCGTGGCCTGCCCCCCGCGGCCCCCCTAGCCCCCCCGCACAGCGCGGCGCAACGGCGGGCGGGCGGGGGGGGTCGGGGTCCGCGGGCTC | 12 |
| GD1-169710-169830 | 25233726 | GAGGGGACAGGGTCGCAGGGGGCGTGTCTGGTGGGGGCGGGAGCGGGGGGCGGCGCGGGAGCCTGCACGCCGTTGGAGGGGAGAATGACAGGGGGGCGGGGACAGAGAGGCGGTCGCGCC | 12 |
| GD1-169740-169860 | 25405384 | GTGGGGGCGGGAGCGGGGGGCGGCGCGGGAGCCTGCACGCCGTTGGAGGGGAGAATGACAGGGGGGCGGGGACAGAGAGGCGGTCGCGCCCCCGGCCGCGGCAGGCAAGCCCCCGGGGGG | 12 |
| GD1-169770-169890 | 25577042 | GCCTGCACGCCGTTGGAGGGGAGAATGACAGGGGGGCGGGGACAGAGAGGCGGTCGCGCCCCCGGCCGCGGCAGGCAAGCCCCCGGGGGGCGGGGCGCGGGCCGCGGAGCGTGACGAAGG | 12 |
| GD1-169800-169920 | 25748700 | GGGGGGCGGGGACAGAGAGGCGGTCGCGCCCCCGGCCGCGGCAGGCAAGCCCCCGGGGGGCGGGGCGCGGGCCGCGGAGCGTGACGAAGGGCCCCAGGGCTGACCCCGGCAAACGTGACC | 12 |
| GD1-170010-170130 | 25920358 | GGGGGGGGGGGCGTGGGGCATGGGGGGCGCATTCCTGGAAAAAGTGGAGGGGGCGTGGCCTGCCCCCCGCGGCCCCCCTAGCCCCCCCGCACAGCGCGGCGCAACGGCGGGCGGGCGGGG | 12 |
| GD1-170190-170310 | 26092016 | GAAGGGGGGGGTCGGGGGCGCGGCGCCCGCCCAGCCATGCGTGACCGCGATGAGGGGACAGGGTCGCAGGGGGCGTGTCTGGTGGGGGCGGGAGCGGGGGGCGGCGCGGGAGCCTGCACG | 12 |
| GD1-170220-170340 | 26263674 | CCAGCCATGCGTGACCGCGATGAGGGGACAGGGTCGCAGGGGGCGTGTCTGGTGGGGGCGGGAGCGGGGGGCGGCGCGGGAGCCTGCACGCCGTTGGAGGGGAGAATGACAGGGGGGCGG | 12 |
| GD1-170250-170370 | 26435332 | GGGTCGCAGGGGGCGTGTCTGGTGGGGGCGGGAGCGGGGGGCGGCGCGGGAGCCTGCACGCCGTTGGAGGGGAGAATGACAGGGGGGCGGGGACAGAGAGGCGGTCGCGCCCCCGGCCGC | 12 |
| GD1-170520-170640 | 26606990 | GGCCCCCCAGGAAAGACCCCCGGGGGCATGGGGGGCGCATTCCTGGAAAAAGTGGAGGGGGCGTGGCCTGCCCCCCGCGGCCCCCCTAGCCCCCCCGCACAGCGCGGCGCAACGGCGGGC | 12 |
| GD1-170550-170670 | 26778648 | GGGGGCGCATTCCTGGAAAAAGTGGAGGGGGCGTGGCCTGCCCCCCGCGGCCCCCCTAGCCCCCCCGCACAGCGCGGCGCAACGGCGGGCGGGCGGGGGGGGTCGGGGTCCGCGGGCTCC | 12 |
| GD1-170700-170820 | 26950306 | GTTCAGGGGAAGGGGGGGGTCGGGGGCGCGGCGCCCGCCCAGCCATGCGTGACCGTGATGAGGGGACAGGGTCGCAGGGGGCGTGTCTGGTGGGGGCGGGAGCGGGGGGCGGCGCGGGAG | 12 |
| GD1-170730-170850 | 27121964 | GCGCCCGCCCAGCCATGCGTGACCGTGATGAGGGGACAGGGTCGCAGGGGGCGTGTCTGGTGGGGGCGGGAGCGGGGGGCGGCGCGGGAGCCTGCACGCCGTTGGAGGGGAGAATGACAG | 8 |
| GD1-170760-170880 | 27293622 | AGGGGACAGGGTCGCAGGGGGCGTGTCTGGTGGGGGCGGGAGCGGGGGGCGGCGCGGGAGCCTGCACGCCGTTGGAGGGGAGAATGACAGGGGGGCGGGGACAGAGAGGCGGTCGCGCCC | 12 |
| GD1-170790-170910 | 27465280 | TGGGGGCGGGAGCGGGGGGCGGCGCGGGAGCCTGCACGCCGTTGGAGGGGAGAATGACAGGGGGGCGGGGACAGAGAGGCGGTCGCGCCCCCGGCCGCGGCAGGCAAGCCCCCGGGGGGC | 12 |
| GD1-170820-170940 | 27636938 | CCTGCACGCCGTTGGAGGGGAGAATGACAGGGGGGCGGGGACAGAGAGGCGGTCGCGCCCCCGGCCGCGGCAGGCAAGCCCCCGGGGGGCGGGGCGCGGGCCGCGGAGCGTGACGAAGGG | 12 |
| GD1-170850-170970 | 27808596 | GGGGGCGGGGACAGAGAGGCGGTCGCGCCCCCGGCCGCGGCAGGCAAGCCCCCGGGGGGCGGGGCGCGGGCCGCGGAGCGTGACGAAGGGCCCCAGGGCTGACCCCGGCAAACGTGACCC | 12 |
| GD1-170880-171000 | 27980254 | CCGGCCGCGGCAGGCAAGCCCCCGGGGGGCGGGGCGCGGGCCGCGGAGCGTGACGAAGGGCCCCAGGGCTGACCCCGGCAAACGTGACCCGGGGCTCCGGGGTGACCCAGCCAAGCGTGA | 12 |
| GD1-170910-171030 | 28151912 | GGGGCGCGGGCCGCGGAGCGTGACGAAGGGCCCCAGGGCTGACCCCGGCAAACGTGACCCGGGGCTCCGGGGTGACCCAGCCAAGCGTGACCAAGGGGCCGCTGGGTGACACAGGCAACC | 8 |
| GD1-171060-171180 | 28323570 | GGGGGGGGGGGCGTGGGGCATGGGGGGCGCATTCCTGGAAAAAGTGGAGGGGGCGTGGCCTGCCCCCCGCGGCCCCCCTAGCCCCCCCGCACAGCGCGGCGCAACGGCGGGCGGGCGGGGG | 12 |
| GD1-171090-171210 | 28495228 | TTCCTGGAAAAAGTGGAGGGGGCGTGGCCTGCCCCCCGCGGCCCCCCTAGCCCCCCCGCACAGCGCGGCGCAACGGCGGGCGGGCGGGGGGGGTCGGGGTCCGCGGGCTCCGGGGGCTGC | 12 |
| GD1-171240-171360 | 28666886 | AAGGGGGGGGTCGGGGGCGCGGCGCCCGCCCAGCCATGCGTGACCGTGATGAGGGGGCAGGGTCGCAGGGGGTGTGTCTGGTGGGGCCGGGAGCGGGGGGCGGCGCGGGAGCCTGCACGC | 12 |
| GD1-171420-171540 | 28838544 | GCAGGCAAGCCCCCGGGGGGCGGGGCGCGGGCCGCGGAGCGTGACGAAGGGCCCCAGGGCTGACCCCGGCAAACGTGACCCGGGGCTCCGGGGTGACCCAGCCAAGCGTGACCAAGGGGC | 12 |
| GD1-171450-171570 | 29010202 | GCCGCGGAGCGTGACGAAGGGCCCCAGGGCTGACCCCGGCAAACGTGACCCGGGGCTCCGGGGTGACCCAGCCAAGCGTGACCAAGGGGCCGCTGGGTGACACAGGCAACCCTGACAAAG | 8 |
| GD1-171537-171657 | 29181860 | GGCCGCTGGGTGACACAGGCAACCCTGACAAAGGCCCCCCAGGAAAGACCCCCGGGGGGGATCCGGGGGGGTGTTGGCGGGGGCATGGGGGGGTCGGATTTCGCCCTTATTGCCCTGTTT | 8 |
| GD1-171537-171657 | 29353518 | GGCCGCTGGGTGACACAGGCAACCCTGACAAAGGCCCCCCAGGAAAGACCCCCGGGGGGGATCCGGGGGGGTGTTGGCGGGGGCATGGGGGGGTCGGATTTCGCCCTTATTGCCCTGTTT | 8 |
